# Diversity in DNA sequence, structure, and heterozygosity of nuclear rRNA gene region in *Neopyropia yezoensis*

**DOI:** 10.1101/2022.03.03.482932

**Authors:** Yukino Mizutani, Yukio Nagano, Kei Kimura, Genta Kobayashi, Yoshio Kawamura

**Affiliations:** Analytical Research Center for Experimental Sciences, Saga University, 1 Honjo-machi, Saga 840-8502, Japan; Faculty of Agriculture, Saga University, 1 Honjo-machi, Saga 840-8502, Japan

**Author notes:** These authors contributed equally to this work (Yukino Mizutani, Yukio Nagano). Corresponding author: Dr. Yukio Nagano.

## Abstract

DNA sequence reads of *Neopyropia yezoensis* (susabi-nori), its relative *N. tenera* (asakusa-nori), and their hybrids available in the sequence read archive produced assembled sequences of the nuclear rRNA gene region for 82 samples. Analysis of the assembled sequences revealed structural differences in the region of nuclear rRNA genes, with 17 forms depending on the presence or absence of introns and their lengths. The samples were divided into three groups based on differences in DNA sequences: Japanese *N. yezoensis*, Chinese *N. yezoensis*, and *N. tenera*/hybrids of *N. tenera* and *N. yezoensis* that exist in both countries. Despite genetic differentiation, the Japanese and Chinese *N. yezoensis* exhibit common structural forms. One sample of Chinese *N. yezoensis* presented almost 1:1 heterozygosity, whereas five other samples of Chinese *N. yezoensis* showed non-1:1 heterozygosity. In the latter case, neither the ratio of alleles nor the ratio of the number of introns was 1:1, suggesting the existence of an ongoing mechanism to eliminate the nuclear rRNA gene region on one of the homologous chromosomes in *N. yezoensis*.

## Introduction

*Neopyropia yezoensis* (susabi-nori) is a red alga belonging to the Bangiaceae family^1^. This species is well known worldwide by its Japanese name, Nori, and is cultivated in shallow waters in East Asia. *N. tenera* (asakusa-nori), which is closely related to *N. yezoensis*, was consumed in Japan before the beginning of *N. yezoensis* cultivation. The hybrids of *N. yezoensis* and *N. tenera* are allotetraploid in conchocelis cells and allodiploid in blade cells^2–4^. In these hybrids, the nuclear rRNA genes are from *N. tenera* whereas the chloroplast and mitochondrial genomes are derived from *N. yezoensis* or *N. tenera*. The genetic diversity of the chloroplast and mitochondrial genomes of *N. yezoensis, N. tenera*, and their hybrids has been studied in detail^4,5^. However, it would be pertinent to study the genetic diversity and structure of their nuclear rRNA genes.

In eukaryotes, the region of nuclear rRNA genes is tandemly repeated in the nucleolus organizer regions of one or more chromosomes. This region sequentially consists of the intergenic spacer (IGS), small subunit (SSU) rRNA gene, internal transcribed spacer 1 (ITS1), 5.8S rRNA gene, internal transcribed spacer 2 (ITS2), and large subunit (LSU) rRNA gene^6,7^. Many eukaryotes have hundreds of copies of this region, and each region does not evolve independently, but in a concerted fashion. It is speculated that this is caused by DNA recombination, repair, and replication, including gene conversion and unequal crossover^8–10^. Therefore, nuclear rRNA genes are useful molecular markers for classifying various organisms. These rRNA genes contain rich taxonomic information because they evolve at different rates, depending on the region^11^. The SSU and LSU rRNA genes, which are functional RNAs and represent the most conserved regions of nuclear ribosomal nuclear RNA genes, are useful for classification at the species level and above^12,13^, whereas the ITS and IGS, which are not functional RNAs, are useful for classification at the species level and below^7^.

Due to the presence of introns, rRNA genes have a variety of forms. The rRNA genes of various organisms occasionally contain group I introns, which are self-splicing ribozymes^14^. The introns of rRNA genes have been proposed to be remnants of the sequential linking of functional RNAs during the evolution of rRNA^15^. In the red algal order Bangiales, ribosomal introns have been well studied in the genera *Bangia* and *Porphyra* (formerly, *Neopyropia* was classified as *Porphyra*)^16–19^. Among the group I introns of red algae, the introns located in the SSU rRNA gene have been well studied. *Porphyra spiralis* var. amplifolia contains a His-Cys box homing endonuclease pseudogene located on the antisense strand in a group I intron^19^. These introns can transfer to other DNA regions. *P. spiralis* and other species such as *N. yezoensis, N. tenera*, and *Neoporphyra haitaensis*, which are used as food in East Asia, have been reported to have introns in their nuclear rRNA gene region^20–23^.

Introns in rRNA genes have been analyzed in a variety of red algae, but these sequences were obtained using specific primers to amplify the region^13,16,18,21,24,25^. However, introns in nuclear rRNA genes are heterogeneous, that is, some introns are absent in one copy of an rRNA gene region but present in another copy within the same individual organism. Therefore, short templates lacking introns are preferentially amplified in PCR-based methods, resulting in an underestimation of the ratio of introns present^25^. In addition, because the study of the group I introns of red algae was mainly focused on SSU rRNA gene, the knowledge of introns in the LSU rRNA gene is less than that in SSU rRNA gene.

The reads obtained from high-throughput DNA sequencing can be used to assemble rRNA gene regions^26^. This technique, based on high-throughput sequencing, can also be applied to determine the amount of each region in heterogeneous DNA regions. Therefore, we obtained DNA sequences of the nuclear rRNA gene region of *N. yezoensis, N. tenera*, and their hybrids collected in China and Japan, using publicly available sequence reads^4,5^ and analyzed diversity in DNA sequence, structure, and heterozygosity of their nuclear rRNA gene regions.

## Results

### Assembly of nuclear rRNA gene regions

DNA sequence reads already available in the sequence read archive were used in this study, including 1 sample of *N. tenera* from Japan (Pyr_19)^4^, 1 sample of a hybrid between *N. tenera* and *N. yezoensis* from Japan (Pyr_27)^4^, 34 samples of *N. yezoensis* from Japan (samples whose names begin with Pyr_, except for Pyr_19 and Pyr_27)^4^, and 53 samples of possible *N. yezoensis* from China (samples whose names start with SRR; names in the sequence read archive)^5^. The sequences of the nuclear rRNA gene region were assembled using these data. The assembly program^26^ failed to use the data from the two Japanese samples (Pyr_3 and Pyr_42) and five Chinese samples (SRR9587948, SRR9587928, SRR9587956, SRR9587969, and SRR9587920) because multiple assembled sequences were generated. The program successfully assembled the sequences of the nuclear rRNA gene region using data of the other 82 samples.

We compared 82 assembled sequences and extracted the conserved core sequences (Supplementary Fig. 1). We verified the validity of each core sequence by mapping the reads to the core sequence, and found no mapping results that would break the continuity of the DNA sequence (Supplementary Fig. 2).

### Classification of samples based on structural differences of the nuclear rRNA gene region

Comparison of the sequences divided these 82 sequences into 17 forms according to their structural differences (Fig. 1, Supplementary Table 1). Both the SSU rRNA and LSU rRNA genes had two maximal introns. Secondary structure inference predicted that these four introns are group I introns (Supplementary Fig. 3). Differences in introns are due to their presence or absence as well as their lengths. In addition, four forms (N, O, P, and Q) have a 157 bp DNA sequence insertion due to a duplication in the 5’ upstream region of the SSU rRNA gene.

**Figure 1.**
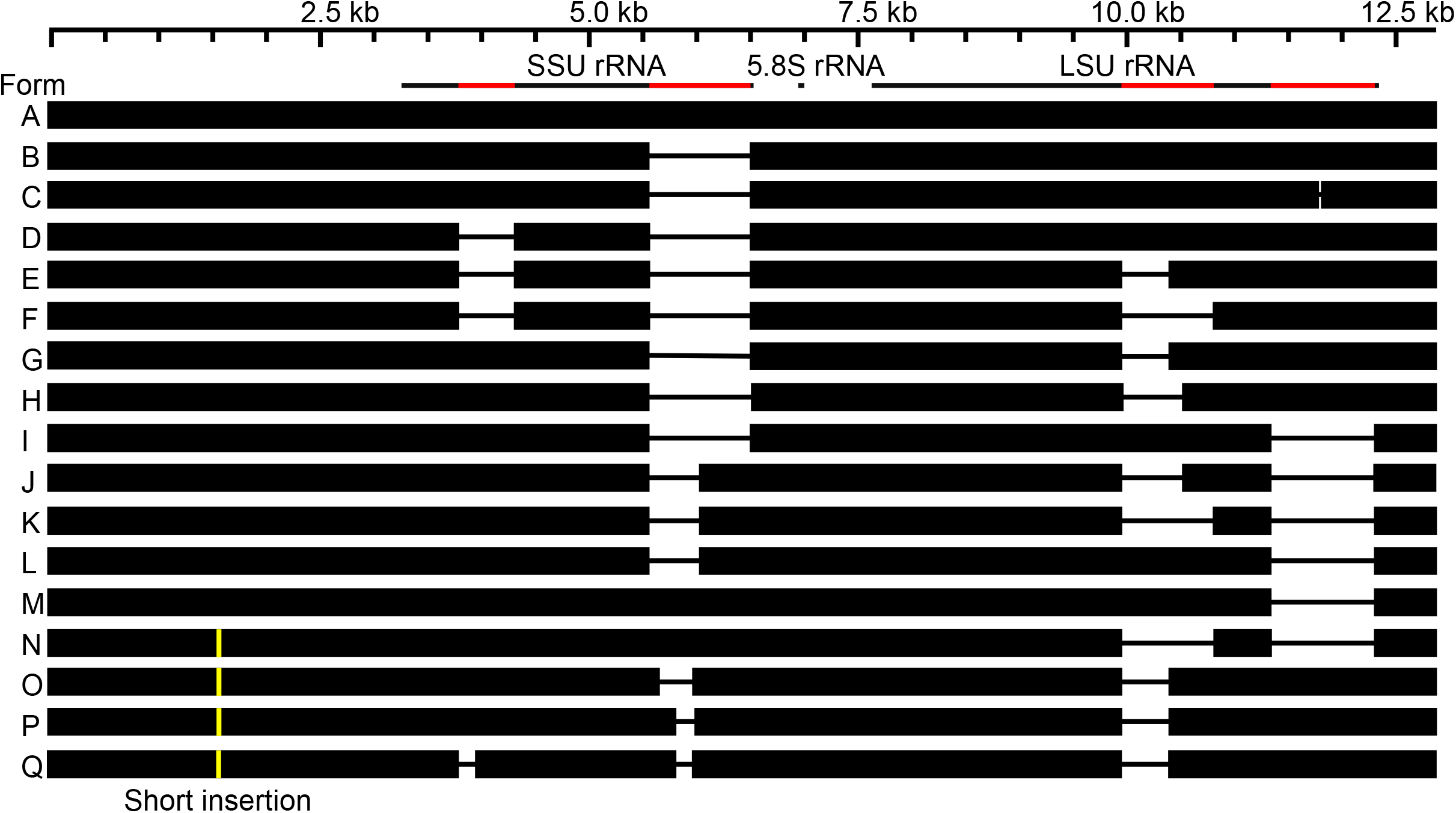
Structural differences involving the region of nuclear rRNA genes. The top line indicates the length of the DNA. The second line from the top illustrates gene structures, with exons as black lines and introns in red. Genes are oriented to the right. For each form, regions where the DNA sequence is present are indicated by a box, and regions where it is absent are indicated by a line. The yellow vertical line indicates 157 bp DNA sequence insertion. The figure was generated manually using BLAST results.

### Classification of samples based on nuclear rRNA gene region sequences

Multiple alignments of these 82 DNA sequences (Supplementary Fig. 4) were used to perform principal component analysis (PCA) and to classify the samples based on their DNA sequence differences (Fig. 2). Principal component 1, with a contribution of 77.2%, separated the four forms from the other forms based on sequence differences. The four forms were found in one *N. tenera* sample (form P), one hybrid sample between *N. tenera* and *N. yezoensis* (form N), and six samples of the possible *N. yezoensis* (five samples in form O and one sample in form Q), all of which had a 157 bp DNA sequence insertion, as described above. These results indicated that these six possible *N. yezoensis* were not *N. yezoensis*, but are hybrids between *N. tenera* and *N. yezoensis*^2–4^ because they have *N. yezoensis* chloroplast and mitochondrial genomes^4,5^.

**Figure 2.**
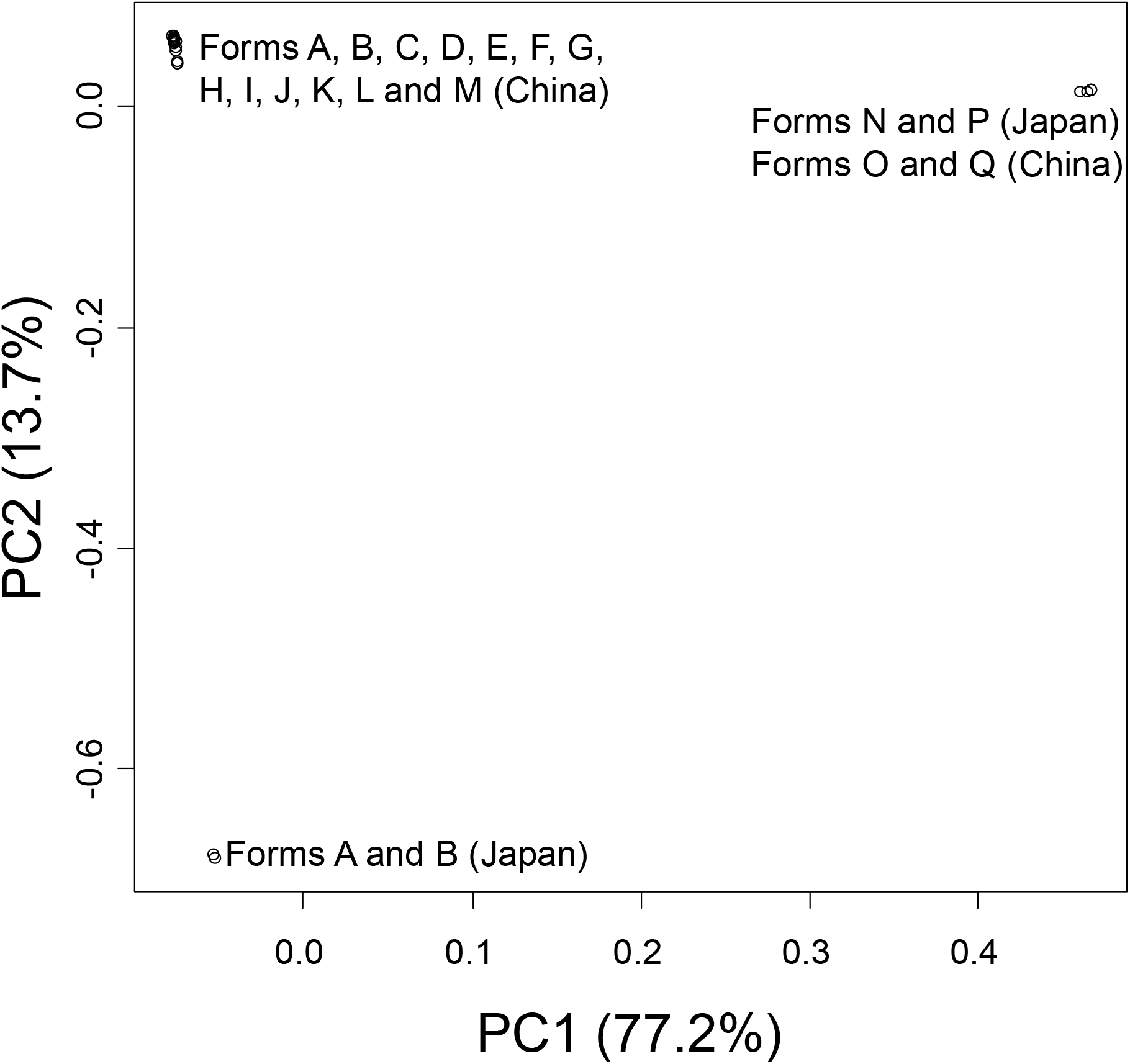
Principal component analysis (PCA) of samples based on their DNA sequences. The first two major component data are shown. The contribution rate of each principal component is shown in parentheses. The figure was generated using R software (v.4.1.2)^37^.

Principal component 2, with a contribution of 13.7%, separated the two forms of Japanese *N. yezoensis* from the other forms based on sequence differences. This low contribution rate indicated that the remaining samples of possible *N. yezoensis* from China were closely related to *N. yezoensis* from Japan. Therefore, these possible *N. yezoensis* from China could have been *N. yezoensis*.

Thus, PCA divided the 82 sequences into three groups based on sequence differences. Phylogenetic trees were created based on neighbor-joining (NJ) and maximum-likelihood (ML) methods (Supplementary Fig. 5). These trees also divided the 82 sequences into three groups.

Comparing Figs. 1 and 2 reveals that forms A and B are present in both Chinese and Japanese *N. yezoensis*. In other words, unlike PCA, structural differences do not clearly distinguish between Japanese and Chinese samples.

### Heterozygous sequences in nuclear rRNA gene region

Inspection of Supplementary Fig. 2 shows that only one sample (SRR9587922) carried multiple near-1:1 heterozygous sites (Fig. 3a). We mapped sequence reads of the Chinese samples to the SRR9587944 sequence (form A sequence carrying four introns) (Supplementary Fig. 6). This result showed that the number of reads of SRR9587922 mapped to the first intron of the *28S* rRNA gene was almost half the number of reads mapped to other locations, indicating that SRR9587922 has forms D and E in a ratio of approximately 1:1 (Fig. 3b). We mapped short reads of the Japanese samples to the Pyr_16 sequence (form A sequence carrying four introns) (Supplementary Fig. 7) and did not detect any samples with multiple heterozygous sites.

**Figure 3.**
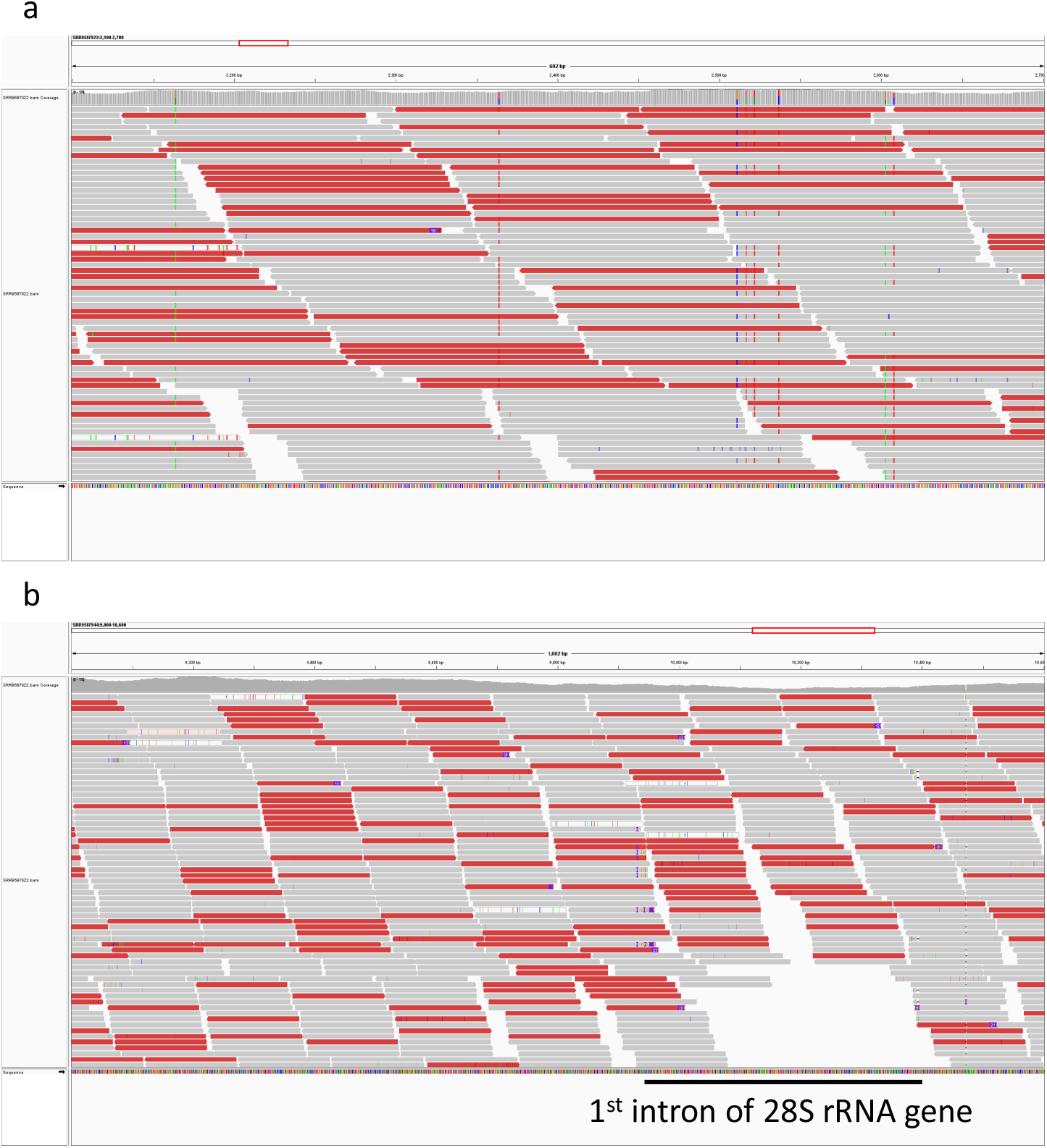
Near-1:1 heterozygous sequence in nuclear rRNA genes of sample SRR9587922. Genotypes were visualized using the integrative genomics viewer^40^. (**a**) Region carrying multiple near-1:1 heterozygous variants. (**b**) Intronic region with one-half reads.

Inspection of Supplementary Fig. 2 reveals non-1:1 heterozygosity for SRR9587923, SRR9587933, SRR9587936, SRR9587966, and SRR9587967 (SRR9587967 results are shown in Fig. 4a). Among these, four samples, except for SRR9587936, showed non-1:1 heterozygosity, even in the intronic region (SRR9587967 results are shown in Fig. 4b). Contamination of reads from other samples may have contributed to these results. However, when we mapped the reads to their respective mitochondrial genomes, we did not detect any heteroplasmy, indicating no evidence of contamination (Supplementary Fig. 8).

**Figure 4.**
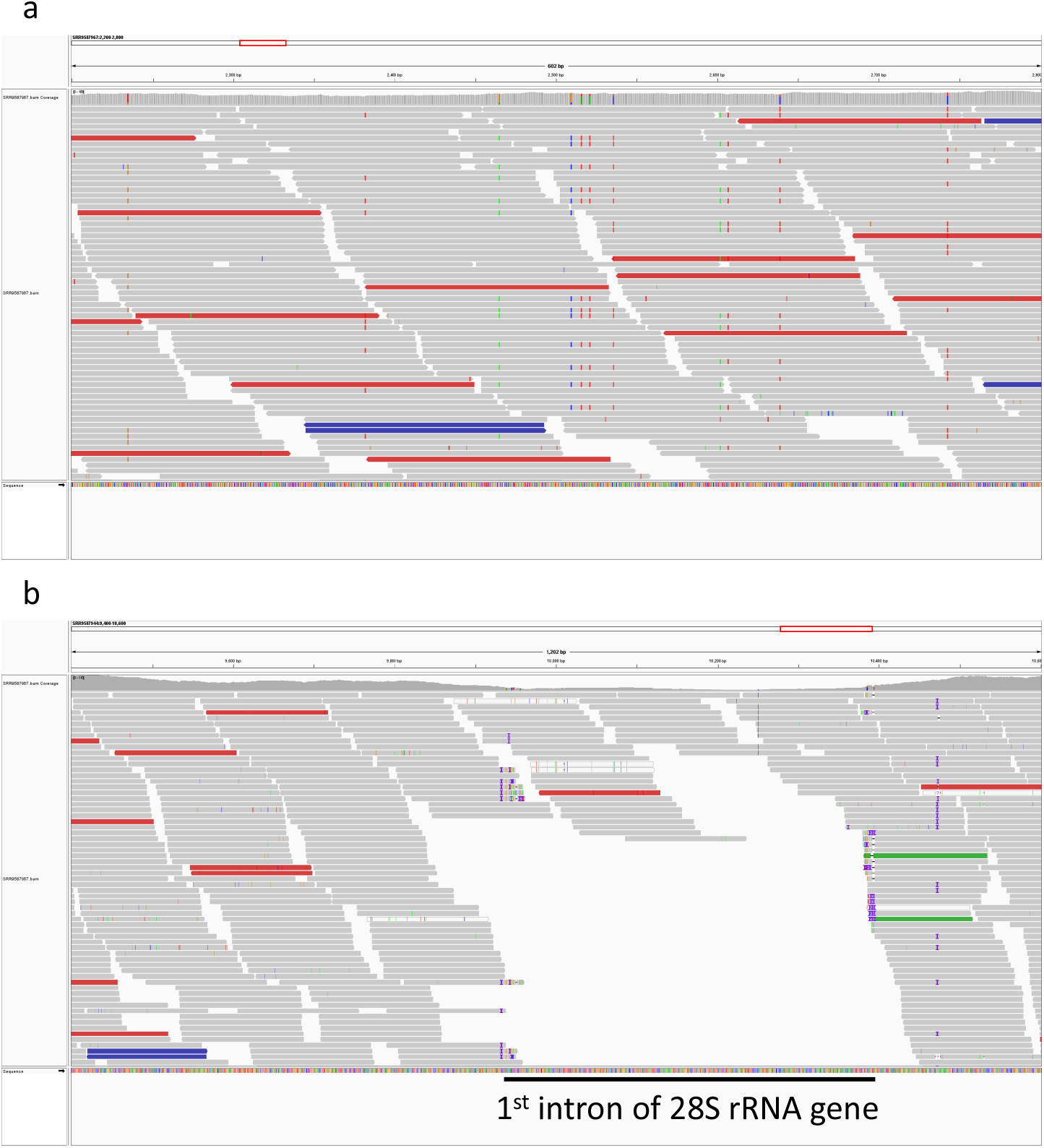
Non-1:1 heterozygous sequence in nuclear rRNA genes of sample SRR9587967. Genotypes were visualized using the integrative genomics viewer^40^. (**a**) Region carrying multiple non-1:1 heterozygous variants. (**b**) Intronic region with a low number of reads.

## Discussion

Various forms of the nuclear rRNA gene region were detected in the 82 samples. The presence or absence of group I introns and their different lengths have been reported for the nuclear rRNA genes of red algae^13,16,18,21,24,25^. The presence of various forms identified in this study is consistent with these previous reports. In the present study, we identified 13 forms in *N. yezoensis*. Because we examined a region with lengths of up to approximately 13 kb, we detected many forms.

In the Japanese *N. yezoensis*, two forms of the nuclear rRNA gene region (forms A and B) are present. These same forms are also present in Chinese *N. yezoensis*, which differs genetically from Japanese *N. yezoensis*. One possible mechanism for the presence of forms A and B in both groups is that the common ancestor of the two groups had various forms, which continue to exist today. It should be noted that these four introns are group I introns, which may exhibit mobility. If the second intron of SSU rRNA gene is excised from form A, it becomes form B. Therefore, another possible mechanism for the presence of forms A and B in both the groups is that such excision events occur independently in Japanese and Chinese *N. yezoensis*.

The samples used in this study were divided into three groups: Japanese *N. yezoensis*, Chinese *N. yezoensis*, and *N. tenera*/hybrids between the two species. Notably, *N. yezoensis* was genetically divided into Japanese and Chinese groups. A previous analysis based on chloroplast genome sequences also indicated clear genetic separation of the Japanese and Chinese groups,^4^ corroborating our results. It is likely that little or no genetic exchange occurred between the two groups after they were separated from a common ancestor. However, the Japanese samples used in this study were presumed to be specimens used for cultivation or wild specimens that escaped from cultivation farms^4^. Therefore, we cannot rule out the possibility that some of the non-cultivated *N. yezoensis* from Japan continue to undergo genetic exchanges with *N. yezoensis* from China.

While the previous analysis based on chloroplast genome sequences and the current PCA based on nuclear rRNA genes divided *N. yezoensis* into two groups (Japanese and Chinese), the previous analysis of mitochondrial genome sequences divided *N. yezoensis* into three Chinese groups and one Japanese cohort^4^. In our previous study^4^, we explained that this discrepancy arose because the evolutionary rate of the mitochondrial genome was more rapid than that of the chloroplast genome. However, certain hidden mechanisms may be responsible for the unusual evolution of the mitochondrial genome. The group containing *N. tenera* or hybrids of the two species was clearly distinct from the other groups. Interestingly, this group is not divided into two groups (Japanese and Chinese). Therefore, this result suggested that genetic exchange occurs between the Japanese and Chinese groups within this group. However, we are not elaborating the differences in the ease of migration between *N. yezoensis* and *N. tenera*, which is difficult because human activities may have played a role in the migration of species belonging to the Bangiaceae family ^24,27^. Of the Chinese samples that were previously morphologically identified as *N. yezoensis* ^5^, six had nuclear rRNA sequences of *N. tenera*, and these are likely to be hybrids between *N. yezoensis* and *N. tenera*^2–4^. Although hybrids between the two species exist in Japan, it is not easy to distinguish them morphologically from *N. yezoensis*. Therefore, when studying *N. yezoensis* candidates, it is important to examine DNA sequences of both the nucleus and organelles.

Three groups were present in both principal component and phylogenetic tree analyses. However, analyses using phylogenetic trees with nuclear rRNA genes may not always be relevant in populations where nuclear rRNA gene recombination occurs. In Supplementary Fig. 5, which compares the sequences of Chinese *N. yezoensis*, certain samples exhibited multiple variations in the intermediate region (ITS1) between the SSU rRNA and 5.8S rRNA genes. In contrast, in other samples, there were multiple variations in the 5’ upstream region of the SSUrRNA gene (part of the IGS sequence). This variation bias is likely due to the recombination of rRNA genes between homologous chromosomes.

In Japanese *N. yezoensis*, no heterozygosity in nuclear rRNA sequences was identified. This is because most of these samples were studied using DNA extracted from diploidized haploid samples. In contrast, there is heterozygosity in rRNA sequences of Chinese *N. yezoensis*. Only one of these samples showed near-1:1 heterozygosity. Five samples showed non-1:1 heterozygosity. Chinese samples were obtained from the wild. Nevertheless, the small number of heterozygous samples suggested that there is considerable self-fertilization or crossing involving genetically-related individuals. Red algae do not have flagella and form non-motile sperm. Because of this low motility, it is plausible that sperm are able to fertilize nearby oocytes. However, a mechanism that erases rRNA sequences of one of the homologous chromosomes may contribute to the low observed number of heterozygous samples.

High-throughput sequencing was used to determine the number of DNA sequences. Examination of the number of DNA sequences revealed that there were five samples exhibiting non-1:1 heterozygosity. Intron heterogeneity can be explained by intronic mobility. However, the simultaneous occurrence of base and intron heterogeneity suggested that an ongoing mechanism that eliminates the rRNA gene region of one of the homologous chromosomes might be responsible for this phenomenon. Complete disappearance of the rRNA gene region of one of the homologous chromosomes has been reported in studies of hybrids between *N. yezoensis* and *N. tenera*^2–4^. Similar mechanisms may be responsible for reductions of this gene region in non-hybrid *Neopyropia* species.

## Methods

### Assembly of nuclear rRNA gene regions

The DNA sequencing reads of *N. yezoensis, N. tenera*, and their hybrids, which have been published^4,5^, were used to assemble nuclear rRNA genes. Adapter sequences and low-quality bases were trimmed using fastp (v.0.20.1) with default parameters^28^. Nuclear rRNA gene assembly was performed using GetOrganelle (v.1.7.5) (-R 10)^26^. Sequences of nuclear rRNA gene region of *Pyropia nitida* (accession number: KR149268)^29^ were used as seed sequence. The assembled sequences were compared using BLAST^30^. Based on BLAST results, well-conserved core sequences were extracted by excluding non-conserved sequences in their upstream and downstream regions. Classification into structural forms was performed based on a comparison of well-conserved core sequences using BLAST. StructRNAfinder^31^ was used to predict intron structures.

### Classification of samples based on nuclear rRNA gene sequences

Multiple sequence alignments of well-conserved core sequences were performed using MAFFT (v.7.490) using default parameters^32^. An NJ tree was constructed using the MEGA 11 program (v. 11.0.10)^33^ (test of phylogeny = bootstrap method, number of bootstrap replications = 1000, model/method = Tamura–Nei method, substitutions to include = transitions + transversions, rate among sites = uniform rates, pattern among lineages = same (homogeneous), and gaps/missing data treatment = complete deletion). An ML tree was constructed using the MEGA 11 program^33^ (test of phylogeny = bootstrap method, number of bootstrap replications = 1000, model/method = Tamura–Nei method, rate among sites = uniform rates, gaps/missing date treatment = complete deletion, ML heuristic method = nearest-neighbour-interchange (NNI), initial tree for ML = make initial tree automatically, nd branch swap filter = none). For PCA, multiple alignments were trimmed using trimAL (v.1.4.1) (-gt 1)^34^. The trimmed sequences were converted to vcf files using the msa2vcf command of JVarkit (jvarkit-msa2vcf-201904251722)^35^. PCA was conducted based on the vcf file using SNPRelate (v.1.28.0)^36^ in R software (version 4.1.2)^37^.

### Mapping of reads against assembled sequences

For visual confirmation of the assembled results, filtered reads were aligned with the assembled genome using bowtie2 (v.2.4.4)^38^. Samtools (v.1.12) was used to process aligned data^39^. The aligned data were visually inspected using integrative genomics viewer (v.2.11.4)^40^. To examine differences within groups and confirm heterozygosity, sequencing reads of Chinese *N. yezoensis* were mapped to the sequence of SRR9587944, and reads of Japanese *N. yezoensis* were mapped to the Pyr_16 sequence using the same method. The mitochondrial sequences for mapping reads to their sequences were created using the GetOrganelle program.

## Acknowledgments

This work was supported by the “Projects for sophistication of production and utilization technology supporting local agriculture and marine industry” from Saga University. We would like to thank Editage for English language editing.

## Author contributions

Y. M., Y. N., K. K., and Y. K. designed this study. Y. M. and Y. N. wrote the original manuscript draft and performed bioinformatic analysis. All authors reviewed the drafts of the manuscript and approved the final version.

