## Supplementary information for "Diversity in DNA sequence, structure, and heterozygosity of nuclear rRNA gene region in *Neopyropia yezoensis*"

**Diversity in DNA sequence, structure and heterozygosity of the region of nuclear rRNA genes in *Neopyropia yezoensis* and its related species**

Yukino Mizutani[1,2,3], Yukio Nagano[1,3✉], Kei Kimura[2], Genta Kobayashi[2], Yoshio Kawamura[2]

[1]Analytical Research Center for Experimental Sciences, Saga University, 1 Honjo-machi, Saga 840-8502, Japan.

[2]Faculty of Agriculture, Saga University, 1 Honjo-machi, Saga 840-8502, Japan.

[3]These authors contributed equally: Yukino Mizutani and Yukio Nagano.

✉

**Supplementary Table 1.** Samples belonging to each of the Forms

| Form | Samples |
|---|---|
| A | Pyr_16, Pyr_18, Pyr_20, Pyr_21, Pyr_24, Pyr_30, Pyr_38, Pyr_40, SRR9587944, SRR9587962 |
| B | Pyr_1, Pyr_2, Pyr_4, Pyr_7, Pyr_8, Pyr_9, Pyr_10, Pyr_11, Pyr_12, Pyr_13, Pyr_14, Pyr_15, Pyr_17, Pyr_22, Pyr_23, Pyr_25, Pyr_26, Pyr_28, Pyr_29, Pyr_33, Pyr_34, Pyr_36, Pyr_39, Pyr_41, SRR9587931, SRR9587932, SRR9587936, SRR9587955 |
| C | SRR9587933 |
| D | SRR9587934 |
| E | SRR9587922, SRR9587923, SRR9587924, SRR9587930, SRR9587965, SRR9587968 |
| F | SRR9587966 |
| G | SRR9587967 |
| H | SRR9587949, SRR9587950, SRR9587952, SRR9587954, SRR9587960, SRR9587961, SRR9587964 |
| I | SRR9587927, SRR9587929, SRR9587942, SRR9587963 |
| J | SRR9587941, SRR9587946, SRR9587959 |
| K | SRR9587921, SRR9587937, SRR9587947 |
| L | SRR9587917, SRR9587918, SRR9587938, SRR9587939, SRR9587943, SRR9587958 |
| M | SRR9587919, SRR9587951, SRR9587953 |
| N | Pyr_27 |
| O | SRR9587925, SRR9587935, SRR9587940, SRR9587945, SRR9587957 |
| P | Pyr_19 |
| Q | SRR9587926 |

The Japanese sample name starts with Pyr_, while the Chinese sample name starts with SRR.

**Supplementary Figure 1.** The conserved core sequences of 82 samples.

>Pyr_1_Pyr_2_Pyr_4_Pyr_7_Pyr_8_Pyr_9_Pyr_10_Pyr_11_Pyr_12_Pyr_13_Pyr_14_Pyr_15_Pyr_17_Pyr_22_Pyr_23_Pyr_25_Pyr_26_Pyr_28_Pyr_29_Pyr_33_Pyr_34_Pyr_36_Pyr_39_Pyr_41

GCATGTTTTGGTATGTGTTTCGACAGTCATGCAAGTCATAGCCGTAGGATGGGGAGTGAG
CTTCATAGCCCGACCCCTTCTCAGGAGCATGTTTTCATAGCATATTGAGCATACTGGGGT
GGTCTCCCATGTAGCTCGCTCTCTCGGTGCAGGACCTCCCTTTTGCGGGCCACTTCGATG
GCCTCCAATAGGGAGAGCATGTTTTGGCATATGAAGCGTTGGTGGTGAGGCTCGTAGCCT
GTATATACATACCAACGCAGTCGTCATATGTCTCTCTGGCGGGGGTCGGTCGTATGACCT
TGTCCCTACCTGGAGCATGTATTTCGTATGATATACTGGACGGCAGCTGCGCTGTTCAGT
CGTCAGCCGTACCTCATGTGCGCGTTTCCAGGTCCGAACGTTTGGCATGCGAGGGCATCA
TACCAGAAGCCTGTTTTCTCGTATGAGCATGTAGGTAGGCAGGCTCTTCAGTCGTCTACC
CATAAATGCACTTCGAAGGAACTTTTTGGTATTGAGTGGCTTTGTGCCCAGACGTTCTTG
TGAGCTTGTTCTCGCTCTACGATACGCTGGATGCTGTGGTATTACCTACAGCAGACGCAT
TCCTAGGGCGGGCGGCCTCTCTGCAGATGAGCACATCTGTAGAAAGAAGCTCCCCTTGAT
GACGCAATGCAATGGCTGGGTAAACGGAGTGCTATACAGAAGGGTCAAACCTCTGTGGAA
GTACACCCATACCAGCTTTGTTTGTTGAGAGGGGTCGGCTCTACGAGTGAGGCCCTATAC
TGCCATCTCATTCAGATGCGTAGACGGTCTTATCTCAGACTGCCTTTCTGGCTGATTGTC
CTCGGTCCCTCGTGCACACGCGTCGCTGGCTGAATGGGCTTGTCCTTATGGAGTGCCGCG
TTCTTCAAAGAATCCGTCTTCATACAGCGACCTCGTTTACATATTGCCGGCTGAATGGGC
TTGTCCTTATGGAGCACTGCATTCTTCTAGGAATTGGTATCTATACAGCGACCTCATTTA
CATACAAGGCACCTGTTTTCTACATAGACTTGCCATTTGGCTGATTGTTCTGACAGTCGC
TGCCTCTTCGGAGTTCTGTGTGGACGAGTGGCCTTCTGAGTGGGCCGGTCCTTACGGAGT
CCCCATATCTGTTATGACTGGGTTCTCCATACAGCGACCTCACTCAGAATTGTATGGCAC
CGTGCCGATTGAATGGACAAGCAGTCCTTAGGGAATCGCATGAAAGCATGACGGCACATG
GTGTATGGACCCGTTGGGCCTTGCCTGTGCTATCTTCTCGAGTCGACCGGTCCTCGTCTC
CTCCTTTCGCGACGGGAAGCTCTTGAGTTTACCGTGGGTCAGGACGCTGTGGTATCTGTT
TGTCTGTAGTGTGGGCTCTCTATCAGAACAGCGTATCAGCACACGTCGTCTGGGTCTATG
GTATTTTTGCTTAACGGCCCTGGGTTTGGTAGTATGGTTCCATGTCGGTCCTCTCACCTC
CCATCCGTCATGTCTCTGATATCATCTGCACAGTACGTCTTGTCAGAATATCATTCTGTA
TGGCAGTCGATAGACTAGGCCCTATGCCGCAGCTGTCTCTGACCACTATGCGAGCGCATA
CCTACCTCAATGTGTTCATAGCTGTTATGGATCTTGGGTGTGGTGTGCCAGAACGATACG
CCACCAGATTGTGGAATGGTGCTTGTAGTATTAGAAGAGGACATGTATGAAGAGCACGTT
CTTTTCTCTTATGATGGTGCCGCGGGTGGGGGCCGTTCTCATTCTGCTTGTCGGGTGCTC
AAGGTGCATGTAGGATTGCTGAACCCACTGAATGGTCATGAGAATGGTCATATGGTGAGG

TTCACTTCTCCATCGCTTTGTGTGTGTTGGATGTTTGGGTGGCTTCGGCCGCATGGAAAA

TCTCGGTCTGTAACTCATCAGGGAGGGTTTCCATGTGAGCAATCTCTCCAGTCGAACACA

CACCCATGGGCGTCCCGCTAGGCAAAATGGAACTCTCGATCCGACACAGCCTGTGGAGTG

TCTGTTCCGTGTGGCGTGAAACGGCTTATACTGGATGTCTTCCCTTTGGACACCAGTCTG

TTGGTATGTACGCATACCATCTAGAGGGAGCAAGAAAAAGGCATGCAAGTGCCACAAAGG

AAGGGAGGCCTTCCTGCTACCGTGTGTCCTCTTGACGATCACGGGGGAGTTTATAGAGAC

ATTTGGACGAGATCCTTTTTTCACGAGGCGGTGCCCTCGCGCAAAGGTGTTGCTCTGCAC

GCATCTTTATGATGATAGTTGCTCGCTCCTATGCGTCCACTCGTGAAAATACCTGCTCTC

TCTGTCTGTTTCCATGACACCCCCATGCGTTGTCATAGAGCACGCTCAGGTACAGAAAGT

GCTGTGACTTTCTTGGCCTGCTGTCACTGAGTCAGCCTGATACTTTGGAGGAATCTTTTC

TGGGCGCTGCCCAGCTGAGAGTTCTTATCAGGCTCTGGAGTGTGCATATGGCTATAATCA

TAATAGTGCATGTGGCCACGCAGAAAAAGCATCGACGTGGAATGGAGAAGAACACCATTA

TGAATGGAACGGTTCTTACTCTGCTTCCGTTGCTTGACGCTGTATGGGAATCGCTTTGCC

CTGGGGGTACACCTTCCATTCCATAACAAAGCATTTGATCTATGCCCCGAGGGTCATTCT

CTATGAAGAGCCCGAAACTTGTCTGAGTGAGAGGCGAAGGTAGCCCTTCATTTATGCGGG

GTCCGCCCTCACACTGTAGTCGGCACAGTCCCTTGTTTATGGACTGCGCAGCTCAATCTT

CGACTGGAGAGGAAAAACAAAACACAAGTATGACACGCACATACGTGTTTTCTAGGACTG

TGACCTGTCGGCTGCCTCGCGGTGGTTGGCATGATGCCCCAAGTGGGGGGGGGGAAACAA

TCAGTATGATTGATAAAAACCAGTGGGGAAAAAGGCGCGCTGTCGCACACCCTTGGTCAC

GGCCTCAAGTCGACGGCCCTCACTGGCCCACCCTGCGACTCCTGTTCGCTATTGACAAGA

CACACCTGCAATACCGGAGGAAATGCCTGAGCAGCTGTGCCGCAGCAACTGCGGACGCGT

AGCGATATGCATCCCTATTGTTTGCGACTTGGTACAAAGAACGTGCGCCGGGAGACGGAT

TGCCCTTTGACTTTTCCACCCAGCCCGCGGCTCTATACCCTTGACGTACCTATTTACCTG

GTTGATCCTGCCAGTAGTCATATGCTTGTCTCAAAGACTAAGCCATGCATGTCTAAGTAT

AAACACTTTTATACTCGTGAAACTGCGAATGGCTCATTAAAACAGTTATAGTTCCTTTGG

GAACAACACTACTTGGATAACCGTAGTAATTCTAGAGCTAATACATGCCTCAACGCCCGA

CTCACGAAGGGTGGTATTTATTGGATAAAAAACCATCGTGCTCTCCAGAGCGCTTTGAGA

TGATTCACAATAACTTGTCGGATCGCATGGCCTCGTGCTGGCGACTCCTCATTCAAATTT

CTGCCCTATCAACTTTCGATGGTAGAGTATTGGTCTACCATGGTGTCGACGGGTGACGGG

GAATTAGGGTTCGATTCCGGAGAGGGAGCCTGAGAAACGGCTACCACATCCAAGGAAGGC

AGCAGGCGCGCAAATTACCCAATCCCGACTCGGGGAGGTAGTGACAAAAAATAACAATAG

GGGGCCCTTTGGGTCTTCTAATTGGAATGAGAACAATTTAAATCCCTTATCGAGGATCCA

TTGGAGGGCAAGTCTGGTGAACGTATGCATCAAAGCCCTCCGTAATAAAAAAAAACAGCG

CCAGATGTCTCTCTCTGGAAGCACACCACGCACGGGGTTCGTTCCTCTCTGCATGGCCGT

GTGAAAAAGGCGAGTAGGCAAACGATGGGAAACCCGGCTTGGAATAGTAACCAGCCACAG

AGCCCCCATCATACACTTCTAAATTGCCGGGGAGTGCCCGCGAGGCCTTGACTACCGCGG

TCCGCCGAGAGGTGAGAGCGAGCAATCGCTCAGGACGCAGCAGTGTACGTAGTGGTACAT
GGATGGTAAAAACGTCAAGGATAGGGTTGATCCGCAGGGAAGCGCGAGGGCCAACCTGCC
TTTGGGCAACAGTTGGCAAAGGCGAACCTTCAGAGACTCTAATGGAGTGGGCGCAAGCTT
AAGGGAGAGTCCAATCCACTGGGAAACCAGGTCCACAGCAGTAACGGGGTGATGCTCGCC
CCAGAGGACTGTGGAAGGCAACCACCAAAGCGGTGTGTGTTGTCGGGTAGCAATGGCCAG
CAGCCGCGGTAATTCCAGCTCCAATAGCGTATATTAAAGTTGTTGCAGTTAAAACGCTCG
TAGTCGGACCTCGGAGCGCCCCCACGGGCCGGGCTTTTGCCTTGTTGCCAGTGGGAGTTT
GTCTCCTTTTGTCGGGACGCGGTGTGCAGGGCCTTTACTGTCTCTGCGCGTTTACCTGCG
GGCCGACCGTTTACTGTGAAGAAAGTAAAGTGTTCAAAGCAGGCCAATGCCTTGAATATG
TGAGCATGGAATAATAGAATAGGACTTGGGCTCTATTTTGTTGGTTTCCAGTGACCAAGT
AATGATTAATAGGGACGGTTGGGGGCATTCGTATTTCATTGTCAGAGGTGAAATTCTTGG
ATTGATGGAAGACGTACAACTGCGAAAGCATCTGCCATGGATGTTTTCATTGATCAAGAA
CGAAAGTTAGGGGATCGAAGACGATCAGATACCGTCGTAGTCTTAACCATAAACGATGCC
GACTGGGGATTGGCGGGGGCACTTTTATATGACTCTGTCAGCACCCTGAGGGAAACCAAA
GTCTTTGGGTTCTGGGGGGAGTATGGTCGCAAGGCTGAAACTTAAAGGAATTGACGGAAG
GGCACCACCAGGAGTGGAGCCTGCGGCTTAATTTGACTCAACACGGGAAAACTTACCAGG
TCCGGACAGAAGAATGATTGACAGACTGAAAGCTCTTTCTTGATTTTTTGGTTGGTGGTG
CATGGCCGTTCTTAGTTGGTGGAGTGATTTGTCTGGTTAATTCCGTTAACGAACGAGACC
TCGTCCTGCTAAATAGGTGCGCGCATGCCAGCACTGCGTTCTTACCTTCTTAGAGGGACT
ATGCGCGTCTAGCGTATGGAAGATTGAGGCAATAACAGGTCTGTGATGCCCTTAGATGTT
CTGGGCCGCACGCGCGCTACACTGATGCATTCAACGAGTTTCACTTGATAGTCCTGGGTC
GGAAGGCTCGGGTAATCTTTTGAAAGTGCATCGTGCTGGGGATAGATCATTGCAATTATT
GATCTTCAACGAGGAATTCCTTGTAGGCGCAGGTCATCAGCCTGCGCCGAATACGTCCCT
GCCCTTTGTACACACCGCCCGTCGCTCCTACCGATTGAATGGTCCGATGAAATGTCCGGA
TCGCGGCTAGTGTGATGGACTCGTCCAGAGCGGTAGCCGTGAGAAGCTCATTAAATCTTA
TCATTTAGAGGAAGGAGAAGTCGTAACAAGGTTTCCGTAGGTGAACCTGCGGAAGGATCA
TTCACAAACTATTGACACAAACACACGCGAACCAAATCGTCCGCACAGGTGGTGGCAATG
AAAGAGAGAATCTGCATGTCGCCTTTCGGGGTATAGCAAGCAGCACTCTTTTGCCATCGC
CTCTGTGCCGGGCGTAAATTCTCATTGAGAGGATGTGAGGGCACCACAGGAAGCTTTTCC
ACAGGAAGTCGCCATCCTTCTCCCTCCACGGCGGCGCTTCTGTGCTTGGCAGTTTTTTTT
TTTGCCTTCCAGGGAGGATGCCGCCAATGGAGCCCCATATAATATATACATCATCATATG
CCCCTTTTTTTCTTAACCGCTTGCCACAGCTTCTTCTATGAGGAGCTTGTGGGAAGACTG
TCTCCATACAATAACAAAGATACAACTCTTAGCGGTGGATATCTTGGTTCTCGCAACGAT
GAAGAACGCAGCCAACTGCGATAACTAATGTGAATTGCAGGACTTCGTGAATCATTAAGT
CTTTGAACGCAAGTTGCGCTCATCTCCATGAGGAGGTGAGCATGCCTGTTTGAGTGTCGA
TGGGAAAAACAATACATAACAACAATCGGCTCTCTTATCTTCCTGCCTTCTTGGGGCAAA

AGGTAGTGCATGCCGGAGATGGGCTTTGTCGCCCCTTTCTCTTGAAAGAGGCGTCGTCCG
AAACACGTATCGTTGCCCGCTGGAGGTCATAGAGCATGAGTTTAAAGCACCACATCTCTC
AAGTCAGAGTGAGCGTCTTGAGTAGGTCCGCATTGATTCAGTTTCAACCCAGGTGGCGTT
GGTTACTTGCCATAAGCCTCTCTGGGGTTTCTTTGCTTGGACCTCTCTTGTCAGTTTCTC
TGACGTAGAGACAGGTGCCGTCTCATGGGCGTTGCCCTCTGGAACGTGCTAAAAAAAAGA
AAAAACCGTACCACATGTCAAAAGAGGCAGATTGTGCTTGACGGCGGGAGATCTTGCATT
TCTTGCTTGTCCTGAGCGTCTGCCAGTCTCTGACGACGAGTGTGTTCCTTCCATAAACCC
GATCTCAAATCAGGTAGGACCACCCGCTGAATTTAAGCATATAATTAAGCGGAGGAAAAG
AAACTAACAAGGATTCCCTCAGTAGCGGCGAGCGAAGCGGGAAGAGCTCAACATGAAAAC
CGATGGGGGTTCGCGCTTTCCTTCGGGTTGTATAGTAAAGTGGTGTCGCCTGCGTCGCGC
CGGGTATAAATCCCGTCTGAAATGGGGTGTCGTAGAGGGTTATAACCCCGTCTGTGACCC
GGTGTGCGATGCGTTACGGTGACCTTACGATGAGTCGGGTTGCTTGGGCATGCAGCCCTA
AGTGGGTGGTAAACTTCATCTAAAGCTAAATACTAGCGAGAGACCGATAGCGTACAAGTA
CCGTAAGGGAAAGATGCAAAGAACTTTGAAAAGAGAGTTAAACAGGGCTTGAAACCGTTA
AGAGTGAAGCGAAGGAGGTCAGTGATCATGGGGCCAGATCCCCAAGTGCCTGTCATAGGG
CACCGTAGCGTTCTGGTTCCCGGGGCAGCGCCGGCCGGGGGCGGGTGTCAACGGGGAGGT
GGTTGAGGGATTTATTTCCCGCCTGTACTCCTTTCTAGCTTGCACCCTGGCCGAGGGATC
CGCGACTTGTCGTGTGAGGGCGCTGTCAAAATGGCCTTCTCCACCCCGTCTTGAAACACG
GACCAAGGAGTCTAACATACTCGCGAGTGTTTGGGTGGTTAAAACCCGAACGCACAATGA
AAGTAAACGGTGGCTCTGCGCCGCTGCTTGGTGGGAACCCTGCTCTTATAATGGGGGGCA
CCATCACCCGACCCCGATCCCTTCGGGGTGGTTGGTTTTGAGGCAGAGCGAGTATGATGG
GACCCGAAAGATGGTGAACTATTCCTGAGTAGGGCGAAGCCAGAGGAAACTCTGGTGGAG
GCTCGTAGCGATGCTGACGTGCAAATCGCTCGTCGAACTTGGGTATAGGGGCGAAAGACT
AATCGAACCATCTAGTAGCTGGTTCCCTCCGAAGTTTCCCTCAGGATAGCGGGAGCAAAT
GCAAGTCAGTATCATCCGGTAAAGCGAATGATTAGAGGCCTCGGGGGAGTAACTCCTTCG
ACCTATTCTCAAACTTTAAATAGGTGAGGAGCCGGCAACGTTGCTTAACACAATGAACGC
CGTGCGGACAATGATTGCTCTCAGTGGGCCGATTTTGGTAAGCAGAACTGGCGATGCGGG
ATGAACCGAAAGCGACGCTAAGGTGCCAAACTCCCAGCTCATTCAGACACCAGAAAAGGT
GTTGATTCATACAGACAGCAGGACGGTGGCCATGGAAGTCGGAATCCGCTAAGGAGTGTG
TAACAACTCACCTGCCGAATGGATCAGCCCTGAAAATGGATGGCGCTTAAGCGGGGGACC
GAGGCGTCGCCGTTGGGGTAAATACGAAGCCCCAATGAGTAGGAGGGCGTGGGGGTTTTG
AAACACGGTACACTCGCATGGAAGCCTTGGGCGTAAGCTCGGGTGGAGCGACCCCTAGTG
TAGATCTTGGTGGTAGTAGCAAATATTCAAGTGAGAACCTTGAAGACTGAAGTGGGGAAG
GGTTCCACGCGAACAGTGCTTGGACGTGGGTTAGTCGGTCCTAAGAGATGGGGAAACTCC
GTATTCAAGTGCGCTTTTGTAGCGCAGTCCATCGAAAGGGAAGCGGGTTAACATTCCCGC
ACCAGGATGTGGAAACTGTGGTTTACCGCAAACTGCGGCAACGCAACTGAACTTGGGGAC

GCCCGTAGGAGCTCCGAGCAGGGATCTCTTTCCTGTTTAACGCGCCCAACCCCCTGGAAT
CGGTTTGTCCGGAGATAGGGGGCGAAAGGCCGGAAAAGCACCGCACGTCTTGTGGTGTTC
GGAACGCTTGTAATGGCCCTTGAAAACCCAGGGGAAAGCAGCCGTCTTACACTTGTGTGA
GACGAGGAATTCTCACACCTGGCCGTACCCATAACCGCATCAGGTCCCCAAGGTGAACAG
CCTCTAGTCAATAGAATAATGTAGATAAGGGAAGTCGGCAAAATAGATCCGTAACTTTGG
GAAAAGGATTGGCTCTAATGTTCGGGCTTGTCGGGACGCGCGGGCGTAGACGCCAATGAG
CGGCAGACGTCTAGCCTCTTTGGAGGCTGGATTGACGCTGCTTGTGGGAAGAAACCGTGT
CGATGGAGGCAGGAGCGCTTATCGTTCTTGTCTCTTCCCGCTTGCGATCAACGGACGGAT
TAGAACTGGTACGGACAAGGGGAATCCGACTGTTTAATTAAAACAAAGCATTGCGACGGC
CGGAAACGGTGTTGGCGCAATGTGATTTCTGCCCAGTGCTCTGAATGTCAAAGTGAAGAA
ATTCAACCAAGCGCGGGTAAACGGCGGGAGTAACTATGACTCTTAAACAAAAAATACTAC
CGACGGGAGTCAAAAGTGGGGGCGGCGTCTGGTGTAGAAATGCACGTATGACACGCTTTT
TTCCTGCTAGTCGATCTGACCTTTGTTCTTTGGACTGTGGGCGGTCAGCGGCAACATAAC
CTGGTACGGGGAAGGCCTCGACGTTGAGTGCTGCATCTTGGAAGTATCTTGACACAATAA
GAGAAAAGAGATTGCGCTTGATAGGGGGTTGATCTCGTGGCGAGTCGACGCCGGTGTGAA
CTGGTGGCAATGGCCGTCGTAGAGCACGCAAAGGTATGGGCTGACTTTTTCTCCTCAAAA
GTTGGCTTAAGGGACGTGCCATCCCACACGATAATAAAGTGTGCAGTCTGTAAGAGCTCC
CACAGGGCAAAGACAGACTGGGATGCAGACGGAGACGGCAAAAGCAACATCCGTTCTGCT
TCTGGGCGCAAATGCTTAAGGTAGCCAAATGCCTCGTCATCTAAACTAATTGGATGAGAT
AGTCTGAGTGAAAAGCTTATGCAGGCTAGTACCGGAAAAAAAAAAAAAAAAGAGGGGCTC
TGCCACTCTGACGGTGCGACACTCCTTGTACAGGGAAACCCTCCGCATGGAAAGGCACGT
GGTACCACCCTCGCGGGGAAACTCGCGAGCGGGATCCCGGGTAATGACCGGAGCCGATGG
TAAAAACTCACGTGACCGATGGGGTGACCCTGTGGCGAGACGCTAAGTTGTGGGACATCA
CCCTCTCACAATACGCATCCGTCGCAGAGACTGAACGGGAGTGGGCTCGGGTCGTCGATG
GCTAACCAGCCTTCGTCGTGGGGCTTAAGATACAGTCCGGGCCTCAGGGATGGAAAAAAA
AACCTGGGGAACTACCAGAATTAGTGACGCGCATGAATGGATTAACGAGATTCCCACTGT
CCCTATCTACTATCTAGCGAAACCACAGCCAAGGGAACGGGCTTGGAAAAATCAGCGGGG
AAAGAAGACCCTGTTGAGCTTGACTCTAGTCCGACTTTGTGAGATGAGTTGGGAGGTGTA
GTCTAAGAGGGAGGGGCAACTCGACCGTGAAATACCTCTACTCCCAAGCTTGTTTTACTT
ACTCCGGCGAGCAGGGATGGGTGGTAACACCCTTTTTCTAGGATTAACACCTCTTCGGGG
GTGGACCTGCGCGGAGGACATTGTCAGGTGGGGAGTTTGGCTGGGGCGGCACATCTGTTA
AAAGATAACGCAGGTGTCCTAAGATGAGCTCAGTGAGAACAGAAATCTCACGTAGAACAA
AAGGGTAAAAGCTCATTTGATTTTGATTTTCAGTACGAATACAAACTGTGAAAGCATGGC
CTATCGATCCTTTTGACCTTCAAGGTTTGAAGCAAGAGGTGTCAGAAAAGTTAACACGCG
TTGTGGTAGCTTTTAAGAGTGGACTCAAAGTGTTCCGCTAGTCCCCCCTCGGACCGCCAC
TTACAAAAGGCCTGGGGATGGGGGCGACACTGTCAAATTGCGGGGACTCCCTAAAGCCTG

TCTCTACCAAGCCAGCGCGGCGAAAGCCCCTGGTGGCCGGGGTAATGACCTAGGGTATGG
TAAAAGCGAGACAGGATGGAACAATGGGCAATCCGCAGCCAAGTCCTAAGGCTTTTTGAA
AAGCGGCTATGGATGCAGTTCAACGACTAGATGGCAGTGGGCTGAGAGAGCCCACCGCGG
CACCCTTTCGGCTTAAGATATAGTCTTGCCCACGTGAAAGCGTGCCTGCAGATGAACGCC
ACCATGTATGGCCGGAGCTGCAGGAGGCCGCTAAACATACCAGCGGCTGGGAACATAACA
GTACCACAGGGATGAACTATCGTCCCAGGCGGGTGTCCCTCTCTCACCCGTCTAGTCCGG
CCTTGGTGCGCAAAGACTTGCGTGGATGCACCTTGGTTAAGGAAAATGCTCCTAGTAGTT
TTATAAGGGCAATTCTTGTAGGGCAACACATCTGGATGCGAAGAACTCCCGGAAAGACAA
CGGTACCAAGTGCTTGGGTTAGCACCCCATTCATGAAGGCAAATAATCGAAAGAGGTGCC
GGGTAATAATCCGTTGTATAGGGACAATCCGCAGCTGACTTGGCTCCCTGCCCATCCATG
GGCACAGCTTAGCGGTTCAACGCTCGGTAAGATGTGGGTTGGCGCGCTGTCCGACAGCGC
ACTGGCTTAAGGTACGGGCTACTCCCACCGGAGACGGTGTCTTTGTCTACCTCGGGTAAC
ACGCCTCGGGAGAAGCAAAGAGGGTGCAATGGAAAAAACTGGTTGCGGGGAAATCCCTAC
CAGAGTCACCCGGTACAAAGGAACTGGCTTGTGGCAGCCAAGCGTTCATAGCGACGTTGC
TTTTTGATCCTTCGATGTCGGCTCTTCCTATCATTGCGAAGCAGAATTCGCCAAGTGTTG
GATTGTTCACCCACTAATAGGGAACGTGAGCTGGGTTTTTTTTTATTGCCCAAAAGCTT
GAAGCTGGGTCCTGTGTGTGACGGGTTGCATTTCTCACGGCTCCCCGTCCGTCGTTTAAT
AGGCGCCACAGTCCTAGATGGATCTTCTGGCTCGACTAGTAGAGGATAGATAGAGATCGT
CAAATTGCGGGAAAGCCCTAAGAGCCCGGAGCTACCAAGTGCGTGTCGAAAGGCATAGAC
CCGGCCAGTGATGGCCGGCGCATGGCTTGATTTCCATCAAGAGGTAAAAAGGCTCTGGGA
TTGGGTAATCCGCAGCCAAGCCCCTAAGGTAGAACATCCGTTTCTGCTATGGGGAAGGTT
CAACGACTGTAAGGCGTACCGTGCAAGCGGAAGAGACAGTCTAGGCCCACTGGAAAGCAG
TGCCAAAGTAAGTAGGCAGGTCGTGGTTCTATCCTC
>Pyr_16_Pyr_18_Pyr_20_Pyr_21_Pyr_24_Pyr_30_Pyr_38_Pyr_40
GCATGTTTTGGTATGTGTTTCGACAGTCATGCAAGTCATAGCCGTAGGATGGGGAGTGAG
CTTCATAGCCCGACCCCTTCTCAGGAGCATGTTTTCATAGCATATTGAGCATACTGGGGT
GGTCTCCCATGTAGCTCGCTCTCTCGGTGCAGGACCTCCCTTTTGCGGGCCACTTCGATG
GCCTCCAATAGGGAGAGCATGTTTTGGCATATGAAGCGTTGGTGGTGAGGCTCGTAGCCT
GTATATACATACCAACGCAGTCGTCATATGTCTCTCTGGCGGGGGTCGGTCGTATGACCT
TGTCCCTACCTGGAGCATGTATTTCGTATGATATACTGGACGGCAGCTGCGCTGTTCAGT
CGTCAGCCGTACCTCATGTGCGCGTTTCCAGGTCCGAACGTTTGGCATGCGAGGGCATCA
TACCAGAAGCCTGTTTTCTCGTATGAGCATGTAGGTAGGCAGGCTCTTCAGTCGTCTACC
CATAAATGCACTTCGAAGGAACTTTTTGGTATTGAGTGGCTTTGTGCCCAGACGTTCTTG
TGAGCTTGTTCTCGCTCTACGATACGCTGGATGCTGTGGTATTACCTACAGCAGACGCAT
TCCTAGGGCGGGCGGCCTCTCTGCAGATGAGCACATCTGTAGAAAGAAGCTCCCCTTGAT
GACGCAATGCAATGGCTGGGTAAACGGAGTGCTATACAGAAGGGTCAAACCTCTGTGGAA

GTACACCCATACCAGCTTTGTTTGTTGAGAGGGGTCGGCTCTACGAGTGAGGCCCTATAC
TGCCATCTCATTCAGATGCGTAGACGGTCTTATCTCAGACTGCCTTTCTGGCTGATTGTC
CTCGGTCCCTCGTGCACACGCGTCGCTGGCTGAATGGGCTTGTCCTTATGGAGTGCCGCG
TTCTTCAAAGAATCCGTCTTCATACAGCGACCTCGTTTACATATTGCCGGCTGAATGGGC
TTGTCCTTATGGAGCACTGCATTCTTCTAGGAATTGGTATCTATACAGCGACCTCATTTA
CATACAAGGCACCTGTTTTCTACATAGACTTGCCATTTGGCTGATTGTTCTGACAGTCGC
TGCCTCTTCGGAGTTCTGTGTGGACGAGTGGCCTTCTGAGTGGGCCGGTCCTTACGGAGT
CCCCATATCTGTTATGACTGGGTTCTCCATACAGCGACCTCACTCAGAATTGTATGGCAC
CGTGCCGATTGAATGGACAAGCAGTCCTTAGGGAATCGCATGAAAGCATGACGGCACATG
GTGTATGGACCCGTTGGGCCTTGCCTGTGCTATCTTCTCGAGTCGACCGGTCCTCGTCTC
CTCCTTTCGCGACGGGAAGCTCTTGAGTTTACCGTGGGTCAGGACGCTGTGGTATCTGTT
TGTCTGTAGTGTGGGCTCTCTATCAGAACAGCGTATCAGCACACGTCGTCTGGGTCTATG
GTATTTTTGCTTAACGGCCCTGGGTTTGGTAGTATGGTTCCATGTCGGTCCTCTCACCTC
CCATCCGTCATGTCTCTGATATCATCTGCACAGTACGTCTTGTCAGAATATCATTCTGTA
TGGCAGTCGATAGACTAGGCCCTATGCCGCAGCTGTCTCTGACCACTATGCGAGCGCATA
CCTACCTCAATGTGTTCATAGCTGTTATGGATCTTGGGTGTGGTGTGCCAGAACGATACG
CCACCAGATTGTGGAATGGTGCTTGTAGTATTAGAAGAGGACATGTATGAAGAGCACGTT
CTTTTCTCTTATGATGGTGCCGCGGGTGGGGGCCGTTCTCATTCTGCTTGTCGGGTGCTC
AAGGTGCATGTAGGATTGCTGAACCCACTGAATGGTCATGAGAATGGTCATATGGTGAGG
TTCACTTCTCCATCGCTTTGTGTGTGTTGGATGTTTGGGTGGCTTCGGCCGCATGGAAAA
TCTCGGTCTGTAACTCATCAGGGAGGGTTTCCATGTGAGCAATCTCTCCAGTCGAACACA
CACCCATGGGCGTCCCGCTAGGCAAAATGGAACTCTCGATCCGACACAGCCTGTGGAGTG
TCTGTTCCGTGTGGCGTGAAACGGCTTATACTGGATGTCTTCCCTTTGGACACCAGTCTG
TTGGTATGTACGCATACCATCTAGAGGGAGCAAGAAAAAGGCATGCAAGTGCCACAAAGG
AAGGGAGGCCTTCCTGCTACCGTGTGTCCTCTTGACGATCACGGGGGAGTTTATAGAGAC
ATTTGGACGAGATCCTTTTTTCACGAGGCGGTGCCCTCGCGCAAAGGTGTTGCTCTGCAC
GCATCTTTATGATGATAGTTGCTCGCTCCTATGCGTCCACTCGTGAAAATACCTGCTCTC
TCTGTCTGTTTCCATGACACCCCCATGCGTTGTCATAGAGCACGCTCAGGTACAGAAAGT
GCTGTGACTTTCTTGGCCTGCTGTCACTGAGTCAGCCTGATACTTTGGAGGAATCTTTTC
TGGGCGCTGCCCAGCTGAGAGTTCTTATCAGGCTCTGGAGTGTGCATATGGCTATAATCA
TAATAGTGCATGTGGCCACGCAGAAAAAGCATCGACGTGGAATGGAGAAGAACACCATTA
TGAATGGAACGGTTCTTACTCTGCTTCCGTTGCTTGACGCTGTATGGGAATCGCTTTGCC
CTGGGGGTACACCTTCCATTCCATAACAAAGCATTTGATCTATGCCCCGAGGGTCATTCT
CTATGAAGAGCCCGAAACTTGTCTGAGTGAGAGGCGAAGGTAGCCCTTCATTTATGCGGG
GTCCGCCCTCACACTGTAGTCGGCACAGTCCCTTGTTTATGGACTGCGCAGCTCAATCTT
CGACTGGAGAGGAAAAACAAAACACAAGTATGACACGCACATACGTGTTTTCTAGGACTG

TGACCTGTCGGCTGCCTCGCGGTGGTTGGCATGATGCCCCAAGTGGGGGGGGGGAAACAA

TCAGTATGATTGATAAAAACCAGTGGGGAAAAAGGCGCGCTGTCGCACACCCTTGGTCAC

GGCCTCAAGTCGACGGCCCTCACTGGCCCACCCTGCGACTCCTGTTCGCTATTGACAAGA

CACACCTGCAATACCGGAGGAAATGCCTGAGCAGCTGTGCCGCAGCAACTGCGGACGCGT

AGCGATATGCATCCCTATTGTTTGCGACTTGGTACAAAGAACGTGCGCCGGGAGACGGAT

TGCCCTTTGACTTTTCCACCCAGCCCGCGGCTCTATACCCTTGACGTACCTATTTACCTG

GTTGATCCTGCCAGTAGTCATATGCTTGTCTCAAAGACTAAGCCATGCATGTCTAAGTAT

AAACACTTTTATACTCGTGAAACTGCGAATGGCTCATTAAAACAGTTATAGTTCCTTTGG

GAACAACACTACTTGGATAACCGTAGTAATTCTAGAGCTAATACATGCCTCAACGCCCGA

CTCACGAAGGGTGGTATTTATTGGATAAAAAACCATCGTGCTCTCCAGAGCGCTTTGAGA

TGATTCACAATAACTTGTCGGATCGCATGGCCTCGTGCTGGCGACTCCTCATTCAAATTT

CTGCCCTATCAACTTTCGATGGTAGAGTATTGGTCTACCATGGTGTCGACGGGTGACGGG

GAATTAGGGTTCGATTCCGGAGAGGGAGCCTGAGAAACGGCTACCACATCCAAGGAAGGC

AGCAGGCGCGCAAATTACCCAATCCCGACTCGGGGAGGTAGTGACAAAAAATAACAATAG

GGGGCCCTTTGGGTCTTCTAATTGGAATGAGAACAATTTAAATCCCTTATCGAGGATCCA

TTGGAGGGCAAGTCTGGTGAACGTATGCATCAAAGCCCTCCGTAATAAAAAAAAACAGCG

CCAGATGTCTCTCTCTGGAAGCACACCACGCACGGGGTTCGTTCCTCTCTGCATGGCCGT

GTGAAAAAGGCGAGTAGGCAAACGATGGGAAACCCGGCTTGGAATAGTAACCAGCCACAG

AGCCCCCATCATACACTTCTAAATTGCCGGGGAGTGCCCGCGAGGCCTTGACTACCGCGG

TCCGCCGAGAGGTGAGAGCGAGCAATCGCTCAGGACGCAGCAGTGTACGTAGTGGTACAT

GGATGGTAAAAACGTCAAGGATAGGGTTGATCCGCAGGGAAGCGCGAGGGCCAACCTGCC

TTTGGGCAACAGTTGGCAAAGGCGAACCTTCAGAGACTCTAATGGAGTGGGCGCAAGCTT

AAGGGAGAGTCCAATCCACTGGGAAACCAGGTCCACAGCAGTAACGGGGTGATGCTCGCC

CCAGAGGACTGTGGAAGGCAACCACCAAAGCGGTGTGTGTTGTCGGGTAGCAATGGCCAG

CAGCCGCGGTAATTCCAGCTCCAATAGCGTATATTAAAGTTGTTGCAGTTAAAACGCTCG

TAGTCGGACCTCGGAGCGCCCCCACGGGCCGGGCTTTTGCCTTGTTGCCAGTGGGAGTTT

GTCTCCTTTTGTCGGGACGCGGTGTGCAGGGCCTTTACTGTCTCTGCGCGTTTACCTGCG

GGCCGACCGTTTACTGTGAAGAAAGTAAAGTGTTCAAAGCAGGCCAATGCCTTGAATATG

TGAGCATGGAATAATAGAATAGGACTTGGGCTCTATTTTGTTGGTTTCCAGTGACCAAGT

AATGATTAATAGGGACGGTTGGGGGCATTCGTATTTCATTGTCAGAGGTGAAATTCTTGG

ATTGATGGAAGACGTACAACTGCGAAAGCATCTGCCATGGATGTTTTCATTGATCAAGAA

CGAAAGTTAGGGGATCGAAGACGATCAGATACCGTCGTAGTCTTAACCATAAACGATGCC

GACTGGGGATTGGCGGGGGCACTTTTATATGACTCTGTCAGCACCCTGAGGGAAACCAAA

GTCTTTGGGTTCTGGGGGGAGTATGGTCGCAAGGCTGAAACTTAAAGGAATTGACGGAAG

GGCACCACCAGGAGTGGAGCCTGCGGCTTAATTTGACTCAACACGGGAAAACTTACCAGG

TCCGGACAGAAGAATGATTGACAGACTGAAAGCTCTTTCTTGATTTTTTGGTTGGTGGTG

CATGGCCGTTCTTAGTTGGTGGAGTGATTTGTCTGGTTAATTCCGTTAACGAACGAGACC
TCGTCCTGCTAAATAGGTGCGCGCATGCCAGCACTGCGTTCTTACCTTCTTAGAGGGACT
ATGCGCGTCTAGCGTATGGAAGATTGAGGCAATAACAGGTCTGTGATGCCCTTAGATGTT
CTGGGCCGCACGCGCGCTACACTGATGCATTCAACGAGTTTCACTTGATAGTCCTGGGTC
GGAAGGCTCGGGTAATCTTTTGAAAGTGCATCGTGCTGGGGATAGATCATTGCAATTATT
GATCTTCAACGAGGAATTCCTTGTAGGTGCAGGTCATCAGCCTGCGCCGAATACGTCCCT
GCCCTTTGTACACACCGCCCGTCGCTCCTACCGATTGAATGGTCCGATGAAATGTCCGGA
TCGCGGCTAGTGTGATGGACTCGTCCAGAGCGGTAGCCGTGAGAAGCTCATTAAATCTTA
TCATTTAGAGGAAGGAGAAGTCGTAACAAGGTTTCCGTATTTTGTGGGTGGATACCGATT
CAATAACATCCGCCGTTTTTCTCAATGTACTATATCCCTAGTCTGCTTGCCTAGTTCTTG
GAGTCTTGTGCGATGTGTGCTATTACAGGCATGCCGGGTCATGAAGACAGACAAGGTTGG
GATCTTCACGGAAGTACGCGCAATAGGAGCGCGTCTTGTTCAACGCACCTGACTCAAGAG
TCATATGCGAGGGACGGACGCAGCGCTTGTCGCCGCACAGATGGGAAGCTTCCTCTCCAG
GCATAGGATGGCGCTGCATTTGCATCACCGCTGCTAGTAAGTGGCAATAATACTTGACTC
CCCGCCAGCGCACTTGTACGTAGCCGTTGGCGGCGGGAACGTAGTTGGTGATGTGACAGC
CTTGAGCGTTCACCTCTGTGTGTCCGCCAATCTGCACAAAGATGGACGCCATATCGGAGA
CAGGGGGAGGTGCCATTGTGATGATGACGATGTTGTTGGGTGGTCCTTCCGTAGGAAGAC
GTACACGAAGCCTTTGCGGCCCGAAAGGGTGCACAGTGCGCGACTATAAACACTGATTCT
GTGGTCAACGCAAGCAGCGGTTCTGTGGAAGAACAGCTGCGACTTTCTCAAATTGCGGGA
ACACCCTAAAGTCAACAAGCACCGCGGCTTGACGTGAAAAGCGTCAGGCGCAGCAGTCGG
GGTAGCGCCCTGGCGGATGGTAATAGCCTTGTTGAATGGCACAATGGGCAATCCGCAGCC
AAGCCCCTAAGGTCCCCCGCTGGGCTATGGGGAAGGTTCACAGACTGTAAGGGAGGGGGT
TTCTTCCATGAAAGAAGCTTAAGAGACAGTCGGTCCCCTGCGAGAGCAGTGTCGCATGTG
AGGAAGGCACCCATGAAAAGTGGGTGCTGGAGAGCCATGTGAGCCCTCTATGAGGGTGGG
AAGTTTTCCGATACAATGAGAAACAAACACGTTCCGTAGGTGAACCTGCGGAAGGATCAT
TCACAAACTATTGACACAAACACACGCGAACCAAATCGTCCGCACAGGTGGTGGCAATGA
AAGAGAGAATCTGCATGTCGCCTTTCGGGGTATAGCAAGCAGCACTCTTTTGCCATCGCC
TCTGTGCCGGGCGTAAATTCTCATTGAGAGGATGTGAGGGCACCACAGGAAGCTTTTCCA
CAGGAAGTCGCCATCCTTCTCCCTCCACGGCGGCGCTTCTGTGCTTGGCAGTTTTTTTTT
TTGCCTTCCAGGGAGGATGCCGCCAATGGAGCCCCATATAATATATACATCATCATATGC
CCCTTTTTTTCTTAACCGCTTGCCACAGCTTCTTCTATGAGGAGCTTGTGGGAAGACTGT
CTCCATACAATAACAAAGATACAACTCTTAGCGGTGGATATCTTGGTTCTCGCAACGATG
AAGAACGCAGCCAACTGCGATAACTAATGTGAATTGCAGGACTTCGTGAATCATTAAGTC
TTTGAACGCAAGTTGCGCTCATCTCCATGAGGAGGTGAGCATGCCTGTTTGAGTGTCGAT
GGGAAAAACAATACATAACAACAATCGGCTCTCTTATCTTCCTGCCTTCTTGGGGCAAAA
GGTAGTGCATGCCGGAGATGGGCTTTGTCGCCCCTTTCTCTTGAAAGAGGCGTCGTCCGA

AACACGTATCGTTGCCCGCTGGAGGTCATAGAGCATGAGTTTAAAGCACCACATCTCTCA
AGTCAGAGTGAGCGTCTTGAGTAGGTCCGCATTGATTCAGTTTCAACCCAGGTGGCGTTG
GTTACTTGCCATAAGCCTCTCTGGGGTTTCTTTGCTTGGACCTCTCTTGTCAGTTTCTCT
GACGTAGAGACAGGTGCCGTCTCATGGGCGTTGCCCTCTGGAACGTGCTAAAAAAAAGAA
AAAACCGTACCACATGTCAAAAGAGGCAGATTGTGCTTGACGGCGGGAGATCTTGCATTT
CTTGCTTGTCCTGAGCGTCTGCCAGTCTCTGACGACGAGTGTGTTCCTTCCATAAACCCG
ATCTCAAATCAGGTAGGACCACCCGCTGAATTTAAGCATATAATTAAGCGGAGGAAAAGA
AACTAACAAGGATTCCCTCAGTAGCGGCGAGCGAAGCGGGAAGAGCTCAACATGAAAACC
GATGGGGGTTCGCGCTTTCCTTCGGGTTGTATAGTAAAGTGGTGTCGCCTGCGTCGCGCC
GGGTATAAATCCCGTCTGAAATGGGGTGTCGTAGAGGGTTATAACCCCGTCTGTGACCCG
GTGTGCGATGCGTTACGGTGACCTTACGATGAGTCGGGTTGCTTGGGCATGCAGCCCTAA
GTGGGTGGTAAACTTCATCTAAAGCTAAATACTAGCGAGAGACCGATAGCGTACAAGTAC
CGTAAGGGAAAGATGCAAAGAACTTTGAAAAGAGAGTTAAACAGGGCTTGAAACCGTTAA
GAGTGAAGCGAAGGAGGTCAGTGATCATGGGGCCAGATCCCCAAGTGCCTGTCATAGGGC
ACCGTAGCGTTCTGGTTCCCGGGGCAGCGCCGGCCGGGGGCGGGTGTCAACGGGGAGGTG
GTTGAGGGATTTATTTCCCGCCTGTACTCCTTTCTAGCTTGCACCCTGGCCGAGGGATCC
GCGACTTGTCGTGTGAGGGCGCTGTCAAAATGGCCTTCTCCACCCCGTCTTGAAACACGG
ACCAAGGAGTCTAACATACTCGCGAGTGTTTGGGTGGTTAAAACCCGAACGCACAATGAA
AGTAAACGGTGGCTCTGCGCCGCTGCTTGGTGGGAACCCTGCTCTTATAATGGGGGGCAC
CATCACCCGACCCCGATCCCTTCGGGGTGGTTGGTTTTGAGGCAGAGCGAGTATGATGGG
ACCCGAAAGATGGTGAACTATTCCTGAGTAGGGCGAAGCCAGAGGAAACTCTGGTGGAGG
CTCGTAGCGATGCTGACGTGCAAATCGCTCGTCGAACTTGGGTATAGGGGCGAAAGACTA
ATCGAACCATCTAGTAGCTGGTTCCCTCCGAAGTTTCCCTCAGGATAGCGGGAGCAAATG
CAAGTCAGTATCATCCGGTAAAGCGAATGATTAGAGGCCTCGGGGGAGTAACTCCTTCGA
CCTATTCTCAAACTTTAAATAGGTGAGGAGCCGGCAACGTTGCTTAACACAATGAACGCC
GTGCGGACAATGATTGCTCTCAGTGGGCCGATTTTGGTAAGCAGAACTGGCGATGCGGGA
TGAACCGAAAGCGACGCTAAGGTGCCAAACTCCCAGCTCATTCAGACACCAGAAAAGGTG
TTGATTCATACAGACAGCAGGACGGTGGCCATGGAAGTCGGAATCCGCTAAGGAGTGTGT
AACAACTCACCTGCCGAATGGATCAGCCCTGAAAATGGATGGCGCTTAAGCGGGGGACCG
AGGCGTCGCCGTTGGGGTAAATACGAAGCCCCAATGAGTAGGAGGGCGTGGGGGTTTTGA
AACACGGTACACTCGCATGGAAGCCTTGGGCGTAAGCTCGGGTGGAGCGACCCCTAGTGT
AGATCTTGGTGGTAGTAGCAAATATTCAAGTGAGAACCTTGAAGACTGAAGTGGGGAAGG
GTTCCACGCGAACAGTGCTTGGACGTGGGTTAGTCGGTCCTAAGAGATGGGGAAACTCCG
TATTCAAGTGCGCTTTTGTAGCGCAGTCCATCGAAAGGGAAGCGGGTTAACATTCCCGCA
CCAGGATGTGGAAACTGTGGTTTACCGCAAACTGCGGCAACGCAACTGAACTTGGGGACG
CCCGTAGGAGCTCCGAGCAGGGATCTCTTTCCTGTTTAACGCGCCCAACCCCCTGGAATC

GGTTTGTCCGGAGATAGGGGGCGAAAGGCCGGAAAAGCACCGCACGTCTTGTGGTGTTCG
GAACGCTTGTAATGGCCCTTGAAAACCCAGGGGAAAGCAGCCGTCTTACACTTGTGTGAG
ACGAGGAATTCTCACACCTGGCCGTACCCATAACCGCATCAGGTCCCCAAGGTGAACAGC
CTCTAGTCAATAGAATAATGTAGATAAGGGAAGTCGGCAAAATAGATCCGTAACTTTGGG
AAAAGGATTGGCTCTAATGTTCGGGCTTGTCGGGACGCGCGGGCGTAGACGCCAATGAGC
GGCAGACGTCTAGCCTCTTTGGAGGCTGGATTGACGCTGCTTGTGGGAAGAAACCGTGTC
GATGGAGGCAGGAGCGCTTATCGTTCTTGTCTCTTCCCGCTTGCGATCAACGGACGGATT
AGAACTGGTACGGACAAGGGGAATCCGACTGTTTAATTAAAACAAAGCATTGCGACGGCC
GGAAACGGTGTTGGCGCAATGTGATTTCTGCCCAGTGCTCTGAATGTCAAAGTGAAGAAA
TTCAACCAAGCGCGGGTAAACGGCGGGAGTAACTATGACTCTTAAACAAAAAATACTACC
GACGGGAGTCAAAAGTGGGGGCGGCGTCTGGTGTAGAAATGCACGTATGACACGCTTTTT
TCCTGCTAGTCGATCTGACCTTTGTTCTTTGGACTGTGGGCGGTCAGCGGCAACATAACC
TGGTACGGGGAAGGCCTCGACGTTGAGTGCTGCATCTTGGAAGTATCTTGACACAATAAG
AGAAAAGAGATTGCGCTTGATAGGGGGTTGATCTCGTGGCGAGTCGACGCCGGTGTGAAC
TGGTGGCAATGGCCGTCGTAGAGCACGCAAAGGTATGGGCTGACTTTTTCTCCTCAAAAG
TTGGCTTAAGGGACGTGCCATCCCACACGATAATAAAGTGTGCAGTCTGTAAGAGCTCCC
ACAGGGCAAAGACAGACTGGGATGCAGACGGAGACGGCAAAAGCAACATCCGTTCTGCTT
CTGGGCGCAAATGCTTAAGGTAGCCAAATGCCTCGTCATCTAAACTAATTGGATGAGATA
GTCTGAGTGAAAAGCTTATGCAGGCTAGTACCGGAAAAAAAAAAAAAAAAGAGGGGCTCT
GCCACTCTGACGGTGCGACACTCCTTGTACAGGGAAACCCTCCGCATGGAAAGGCACGTG
GTACCACCCTCGCGGGGAAACTCGCGAGCGGGATCCCGGGTAATGACCGGAGCCGATGGT
AAAAACTCACGTGACCGATGGGGTGACCCTGTGGCGAGACGCTAAGTTGTGGGACATCAC
CCTCTCACAATACGCATCCGTCGCAGAGACTGAACGGGAGTGGGCTCGGGTCGTCGATGG
CTAACCAGCCTTCGTCGTGGGGCTTAAGATACAGTCCGGGCCTCAGGGATGGAAAAAAAA
ACCTGGGGAACTACCAGAATTAGTGACGCGCATGAATGGATTAACGAGATTCCCACTGTC
CCTATCTACTATCTAGCGAAACCACAGCCAAGGGAACGGGCTTGGAAAAATCAGCGGGGA
AAGAAGACCCTGTTGAGCTTGACTCTAGTCCGACTTTGTGAGATGAGTTGGGAGGTGTAG
TCTAAGAGGGAGGGGCAACTCGACCGTGAAATACCTCTACTCCCAAGCTTGTTTTACTTA
CTCCGGCGAGCAGGGATGGGTGGTAACACCCTTTTTCTAGGATTAACACCTCTTCGGGGG
TGGACCTGCGCGGAGGACATTGTCAGGTGGGGAGTTTGGCTGGGGCGGCACATCTGTTAA
AAGATAACGCAGGTGTCCTAAGATGAGCTCAGTGAGAACAGAAATCTCACGTAGAACAAA
AGGGTAAAAGCTCATTTGATTTTGATTTTCAGTACGAATACAAACTGTGAAAGCATGGCC
TATCGATCCTTTTGACCTTCAAGGTTTGAAGCAAGAGGTGTCAGAAAAGTTAACACGCGT
TGTGGTAGCTTTTAAGAGTGGACTCAAAGTGTTCCGCTAGTCCCCCCTCGGACCGCCACT
TACAAAAGGCCTGGGGATGGGGGCGACACTGTCAAATTGCGGGGACTCCCTAAAGCCTGT
CTCTACCAAGCCAGCGCGGCGAAAGCCCCTGGTGGCCGGGGTAATGACCTAGGGTATGGT

AAAAGCGAGACAGGATGGAACAATGGGCAATCCGCAGCCAAGTCCTAAGGCTTTTTGAAA
AGCGGCTATGGATGCAGTTCAACGACTAGATGGCAGTGGGCTGAGAGAGCCCACCGCGGC
ACCCTTTCGGCTTAAGATATAGTCTTGCCCACGTGAAAGCGTGCCTGCAGATGAACGCCA
CCATGTATGGCCGGAGCTGCAGGAGGCCGCTAAACATACCAGCGGCTGGGAACATAACAG
TACCACAGGGATGAACTATCGTCCCAGGCGGGTGTCCCTCTCTCACCCGTCTAGTCCGGC
CTTGGTGCGCAAAGACTTGCGTGGATGCACCTTGGTTAAGGAAAATGCTCCTAGTAGTTT
TATAAGGGCAATTCTTGTAGGGCAACACATCTGGATGCGAAGAACTCCCGGAAAGACAAC
GGTACCAAGTGCTTGGGTTAGCACCCCATTCATGAAGGCAAATAATCGAAAGAGGTGCCG
GGTAATAATCCGTTGTATAGGGACAATCCGCAGCTGACTTGGCTCCCTGCCCATCCATGG
GCACAGCTTAGCGGTTCAACGCTCGGTAAGATGTGGGTTGGCGCGCTGTCCGACAGCGCA
CTGGCTTAAGGTACGGGCTACTCCCACCGGAGACGGTGTCTTTGTCTACCTCGGGTAACA
CGCCTCGGGAGAAGCAAAGAGGGTGCAATGGAAAAAACTGGTTGCGGGGAAATCCCTACC
AGAGTCACCCGGTACAAAGGAACTGGCTTGTGGCAGCCAAGCGTTCATAGCGACGTTGCT
TTTTGATCCTTCGATGTCGGCTCTTCCTATCATTGCGAAGCAGAATTCGCCAAGTGTTGG
ATTGTTCACCCACTAATAGGGAACGTGAGCTGGGTTTTTTTTTATTGCCCAAAAGCTTG
AAGCTGGGTCCTGTGTGTGACGGGTTGCATTTCTCACGGCTCCCCGTCCGTCGTTTAATA
GGCGCCACAGTCCTAGATGGATCTTCTGGCTCGACTAGTAGAGGATAGATAGAGATCGTC
AAATTGCGGGAAAGCCCTAAGAGCCCGGAGCTACCAAGTGCGTGTCGAAAGGCATAGACC
CGGCCAGTGATGGCCGGCGCATGGCTTGATTTCCATCAAGAGGTAAAAAGGCTCTGGGAT
TGGGTAATCCGCAGCCAAGCCCCTAAGGTAGAACATCCGTTTCTGCTATGGGGAAGGTTC
AACGACTGTAAGGCGTACCGTGCAAGCGGAAGAGACAGTCTAGGCCCACTGGAAAGCAGT
GCCAAAGTAAGTAGGCAGGTCGTGGTTCTATCCTC
>Pyr_19
GCCTGTTTTGATATGTGTTTTGACAGTCATACGGGTCATGGCCGAGGGATGGTGGGTGAG
CTTCATAGCGCAATCCCCCGTCAGGAGCATGTTTTCATAGCATTTTGAGCATACTAGGGT
GGTCTCCCATGTAGCTCGCTCTCTCGGCTCAGGACCTTCCGTATGGGGCACTTTTTGACG
AGCTCCTTTGGGGAGGAGCATGTTCTGGCATATGAAGCGTTGGTCACGAGCTTCGTAGCC
CGACACATACCAAATCAGTCTTCATATGTCTTTCTGGCGGGGGTCGGTCGTATGACCACG
CCCTTACCTTGAGCATGTTTTTCGTATGATACCATACCTGACGGGCAGCGCCGCTGTTCA
GTCGTCGGCCGTACCTCCTATGCGCGTTTTCTGATTCGAACGTGTGACATGCAAGCGCAT
CATATCAGAAGCATGTTTTTTCATATGAGCATGTTGGTAGGCAGGCTCTTCAGTCGTCTA
CCCATAAATGCACTTTGAAAGACGTTTTTTGGTACTGAACGGCTTTGTGGTCAGACGTTC
TCGTGAGCCTATTCTCGCCCTCTGATACGCTGGATGCTGCGGTATTACCCACAGCAGACA
TCCAGGAGCTCCCTCTGCGGATAAGCATCTGTAGAAGGGGGTTTCTCTCGGTGACGCACT
GGCTGGGAAAACGGAGTGCTATCCAAAAGGGTGAAACCTCTTGGGGATGTACTCCCATAC
CAGCATAATATGCTGAGAGGAATTGACTCAATGAGTGAGGTCCTATACTGCAACCTCTTT

TGGATATGTAGGCGGTCTGTACCCAGACTGTCCTACTGGCTGATTTCCCTTGGTCCCACG
CGTCGCTGGCTGAATGGGCTAGTCCTTATGAAGTGCCGCGTTCATCAAAAGAACCGGTCT
TCATACAGCGACCTCATTCTATACTGCCAGCTGGATGGGCTGGTCCTTATGGAGCGCCGC
ATTCTTCGAAGAATCGGTTTCTATAGAGCGACCCCATTTACATATAAGGAACCGTTACAT
AGACTTGCCATTTGGCTGATGGCTCTGACAGTCGCTGCCTCTTCGGAGTTCTGTGTAAAC
GTGGCGCCATTTTTGAGTGGGCCAGTCCTTATGGAGTGTCCGTGTCTTCTATGACAGGGT
CCCCCATACAGCGACCTCACTCAGAACCGTATGGGCACGATGGTGAAAGACCATGCAGAT
AGTCCGCCTTCCAGAGGGGGAAGTAGTATATGGGGAATGCTGGCGAAGCATGAGGTACAT
GGTTCATGGACCCGTTGGGCCATGCCTGTGCCATCTTCTCGAGTCAAACCGGTCCGTGTC
GCTTCCTTATCTCTTCCTTTCGCGACGACAAGCTCTTGAGCTGGTCGTGGGTCAGGACGC
TGTGGTATCTGTTTGTCTGTAGTGTGGGCTTCCACCAGAACAGCGTATCAGCACACGTCG
TCTGGGCATATGATGTTTTTGCTTTGTGGTCCCGGGTTTGGTAGTATGGTTCCATGTCGG
TCCGCTCACCTCCCAACCATAATGTCTTTCACATCATCTTCACAGTACGTCTTGTCAGCA
TACCATTCTGTATAGAACAGTGTATCAGCACACGTCGTCTGAGCATAGGGTATTTTTGCT
TTGTGGTCCCGGGTTTGGTAGTATGGTTCCATGTCGGTCCGCTCACCTCCCAACCATAAT
GTCATTTCTATCATCTGCACAGTACGTCTTGTCAGAATACCATTCTCTATGGCAGTCGAT
AGACTAGGCCTTATGCCGCAGCAGTCTCTGATCACTGTGCGAGTGCATATATCTCTCAAT
GTGTTCATGGCTAGCATGGGTCTTGAGTGAGATGTGCCAGAGAGATACGCCACCAGTCTA
TGGAAAGGTGCTTTTTGTATGAGAAGAGTACATGTATGAAGAGCATGTTCTCTCCTCTTA
TGATGGTACCGCGGGTACGGGCTGCTCGCATTATGCTTGGCAGGTGTCCACTGTGCATGC
AGGATTGCTGAACCCACTTGTATGGCCATGAGAATGGTCATTGGGTGAGGTTCACTTCTC
CATCGCTTTGTGTGTGTTAGATGTTTGGGTGGCTTCGGCCGCATGGAAATTCTCGGTCTG
TAACTCATCAGGGAGGGTTTCCATGTGAGTATTCTCCAGTCGACCACACACCCTTGGGCG
TCCCGCTAGACAAAATGTAACTCTCGATCCGACCTAGCCTGTGGAGTGTCTTTCCGTATG
GCGTGAAAGGGCCCAGACTGGATGTTTTCCCTTTTGGACACCAGTCTGTTATATAGATCA
TCTACAGGGAGCAAAGGACAGGCATGCAAGTGCCACAAAGGCAGGGAGGCCTTCCTGCTA
CTGTGTGTCCTTCTTGACGATCACGGGGGAGTATATAGAAACATTTGAAGGAGATCCTTT
TCCACAAGGCGGTGCCCTCGCGCAAAGGTGTTGCTCTGCACGCATCTGTATGATGATAGT
TGCTCGCTCCTATGCGTCCACTTGTGAAAATACCTTCTCTCTTTGTCTGTTTCTATGACG
CCCCTTCTGCTTTGTCATAGAGCATACTCAGGTACAGAAAGTGCCGTGACTTCCTTGGCC
TGCTGTCAAAGAAATAGCCTGGTGCTATAGAGGACTCTTCCTGGGCGCCGCCTGGTGAAG
AGGTCCTATCAGGCTCTGGAGTGTGCGTATGCCTATGTTCATCATAGCTCATGTGCACGC
GCGGAAAAGCATCGACGTGGAATGGGGAAGAACACCATTCTGAATGAGCCAGTTCTTCCT
CTGCTCCCCTTGCTTGACGCTGTATGGGAATCGCTTTGCCTTGGGGGTACACCTTCCATT
CCATGACAAAGCATTCGATCTATGTCCCGAGGGGTGATTCTACATGAAGAACCCGAAACT
TGTCTGAGTGAGAGGCGAAGGTAGTCCTTCATTTTATGCGGGATCCGCCCTCACACTGTA

GTCGGCACAGTCCCTTGTTTAGGGACTGCGCAGCTCAATCTTCGACTGGAGAGGAAAAAA
ACAAAACACAAGTATGACACGCACATACGTGTTTTCTAGGCCATTGACCTGTTGGCTGCC
TCGCGGTGGTTGGCATGATGCCCCAAGTGGGGGGGGGGGAAATCAATCAGTATGGTTGAC
AAAATCCAGTGGGGAAAAAAGGCGCGCTGTCGCACACCCTTGGTCACGGCCTCAAAAAGT
CGACGGCCCTCACTGGCCCACCCCTGCGACTCCTGTTCGCTTTTGACAAGACACACCTGC
AATATGGAGGAAATGCCTGAGCAGCTGTGCCGCAGCAGCTGTGGACGCGTAGCGATATGC
ATCCCTATTGTTTGCGACTTGGTACAAAGAACGCGTGCCGGGAGACGGATTGCCCTTTGA
CTTTTCCACCCGGCCCGCGGCTCTGCCCTTTACGTACCCTTATTTACCTGGTTGATCCTG
CCAGTAGTCATATGCTTGTCTCAAAGACTAAGCCATGCATGTCTAAGTATAAACACTTTT
ATACTCGTGAAACTGCGAATGGCTCATTAAAACAGTTATAGTTCCTTTGGGAACAACACT
ACTTGGATAACCGTAGTAATTCTAGAGCTAATACATGCCTCAACGCCCGACTCACGAAGG
GTGGTATTTATTGGATAAAAAACCATCGTGCTCTCTCCAGAGCGCTTTGAGATGATTCAC
AATAACTTGTCGGATCGCATGGCCTCGTGCTGGCGACTCCTCATTCAAATTTCTGCCCTA
TCAACTTTCGATGGTAGAGTATTGGTCTACCATGGTGTCGACGGGTGACGGGGAATTAGG
GTTCGATTCCGGAGAGGGAGCCTGAGAAACGGCTACCACATCCAAGGAAGGCAGCAGGCG
CGCAAATTACCCAATCCCGACTCGGGGAGGTAGTGACAAAAAATAACAATAGGGGGCCCT
TTGGGTCTTCTAATTGGAATGAGAACAATTTAAATCCCTTATCGAGGATCCATTGGAGGG
CAAGTCTGGTGAACGTATGCATCAAAGCCCTCGATAAAAAAAAACAGCGCCAGATGTCTC
TCTCTGGAAGCGCACCATGCATGGCTTTGTTCCTCTCTGCATGGCCGTGTGGAAGGCGAG
TAGGCAAACGATGGGAAACCAGGCTTGGAATAGTAACCAACCACAGAGCCCCCATCATAC
ACTTCTAAATTGCCGGGGAGTGCCCGCGAGGCCTTGACTACCGCGGTCCGCCGAGAGGTG
AGAGCGAGCAATCGTTCAGGACGCAGCAGTGTACGTAGTGGTACATGGATGGTAAAAACG
TCAAGGATAGGGTTGATCCGCAGGGAAGCGCGAGTGCCGCCATGGTCGGTGGGCAAAGGC
GAACCTTCAGAGACTCTAATGGAGTGGGCGCAAGCTTAAGGGAGAGTCCAATCCACTGGG
AAACCAGGTCCACAGCAGTAACGGGGTGATGCTCGCCCCAGAGGACTGTGGAAGGCACCA
CCACCCCAAGAGGGGGTGGGTGTTGTCGGGTAGCAAATGGCCAGCAGCCGCGGTAATTCC
AGCTCCAATAGCGTATATTAAAGTTGTTGCAGTTAAAACGCTCGTAGTCGGACCTCGGAG
CGCCCCCACGGGCCGGGCTTTTGCCTTGTTGCCAGTGGGAGTTTGTCTCCTTTTGTCGGG
ACGCGGTGTGCAGGGCCTTTACTGTCTCTGCGCGTTTACCTGCGGGCCGACCGTTTACTG
TGAAGAAAGTAAAGTGTTCAAAGCAGGCCAATGCCTTGAATATGTGAGCATGGAATAATA
GAATAGGACTTGGGCTCTATTTTGTTGGTTTCCAGTGACCAAGTAATGATTAATAGGGAC
GGTTGGGGGCATTCGTATTTCATTGTCAGAGGTGAAATTCTTGGATTGATGGAAGACGTA
CAACTGCGAAAGCATCTGCCATGGATGTTTTCATTGATCAAGAACGAAAGTTAGGGGATC
GAAGACGATCAGATACCGTCGTAGTCTTAACCATAAACGATGCCGACTGGGGATTGGCGG
GGGCACTATTATATGACTCTGTCAGCACCCTGAGGGAAACCAAAGTCTTTGGGTTCTGGG
GGGAGTATGGTCGCAAGGCTGAAACTTAAAGGAATTGACGGAAGGGCACCACCAGGAGTG

GAGCCTGCGGCTTAATTTGACTCAACACGGGAAAACTTACCAGGTCCGGACAGAAGAATG
ATTGACAGACTGAAAGCTCTTTCTTGATTTTTTGGTTGGTGGTGCATGGCCGTTCTTAGT
TGGTGGAGTGATTTGTCTGGTTAATTCCGTTAACGAACGAGACCTCGTCCTGCTAAATAG
GTGCGCGCATGCCAGCACTGCGTTCTTACCTTCTTAGAGGGACTATGCGCGTCTAGCGTA
TGGAAGATTGAGGCAATAACAGGTCTGTGATGCCCTTAGATGTTCTGGGCCGCACGCGCG
CTACACTGATGCATTCAACGAGTTTCACTTGATAGTCCTGGGTCGGAAGGCTCGGGTAAT
CTTTTGAAAGTGCATCGTGCTGGGGATAGATCATTGCAATTATTGATCTTCAACGAGGAA
TTCCTTGTAGGTGCAGGTCATCAGCCTGCGCCGAATACGTCCCTGCCCTTTGTACACACC
GCCCGTCGCTCCTACCGATTGAATGGTCCGATGAAATGTCCGGATCGCGGCTAGTGTGAT
GGACTCGTCCAGAGCGGTAGCCGTGAGAAGCTCATTAAATCTTATCATTTAGAGGAAGGA
GAAGTCGTAACAAGGTTTCCGTATTTTGTGGGTGGATACCGATTCAATAACATCCGCCGT
TTTTCTCAATGTACTATATCCCTAGTCTGCTTGCCTAGTTCTTGGAGTCTTGTGCGATGT
GTGCTATTACAGGCATGCCGGGTCATGAAGACAGACAAGGTTGGGATCTTCACGGAAGTA
CGCGCAATAGGAGCGCGTCTTGTTCAACGCACCCGACTCAAGAGTCATATGCGAGGGACG
GACGCAGCGCTTGTCGCCGCACAGATGGGACGCGCCATATCGGAGACAGGGGGACGTGCC
ATTGTGATGATGATGATGATGTTGTTGGGTGGTCCTTCCGTAGGAAAACGTACACGAAGC
CTTTGCGGCCCGAAAGGGTGCACAGTGCGCGACTATAAACACTGATTCTGTGGTCAACGC
AAGCAGCGGTTCTGTGGAAGAACAGCTGCGACTTTCTCAAATTGCGGGAACACCCTAAAG
TCAACAAACACCGCGGCCGGACGTGAAAGCGTCAGGCGCAGCAGTCGGGGTAGCGCCCTG
GTGGAAGGTAATAGCTTTGTTGAATGGCACAATGGGCAATCCGCAGCCAAGCCCCTAAGG
TCCCCCGCTGGGCTATGGGGAAGGTTCACAGACTGTAAGGGAGGGGGTTTCTTCCATGAA
AGAAGCTTAAGAGACAGTCGGTCCCCTGCGAGAGCAGTGTCGCATGTGAGGAAGGCACCC
ATGAAAAGTGGGTGCTGGAGAGCCATGTGAGCCCTCTATGAGGGTGGGAAGTTTTCCGAT
ACAATGAGAAACAAACACGTTCCGTAGGTGAACCTGCGGAAGGATCATTCACAAACTATT
GACACAAACACACGCGAACCAAATCGTCCACACAGGTGCCGATGAAAGAGAGAATCTGTT
TGTCGCCTTTTGGGGTATACAAGCACTCTTTTCCATCGCCTCTGTGCAGGGCGTAACTTC
CCATTGAGAGGATGTGAGGGCACCACAGGAAGCTTTTCCACAGGAAGTCGCCATCCTTCT
CCCTCCACGACGGCGGCGCTTCACAGGCTTGGCAGTTTTTTTTTTTTTTTATTGCCTTCC
AGGGAGGATGCCGCCAATGGAGCCACATAGACATTTACATCATCATATGCCCGTTTTTCT
TTCTTAACCGCTTGCCACAGTTTCTCCTGTGAGGAGCTTGTGGGAAGACCGTCTCCATAC
AATAACAAAAGATACAACTCTTAGCGGTGGATATCTTGGTTCTCGCAACGATGAAGAACG
CAGCCAACTGCGATACCTAGTGTGAATTGCAGGACTCCGTGAATCATTAAGTCTTTGAAC
GCAAGTTGCGCTCATCTCCATGAGGAGGTGAGCATGCCTGTTTGAGTGTCGATGGGAAAA
ACAATACATAACAACAATCGGCTCTCTTCCCTTCCTGCCTTCCCCGGCAAAAAAGGTAGT
GCATGCCGGAGATGGGCCTGGTCGCCCCTTCATGAAAGAGGCGTCGTCCGAAACATGTGT
CGTTGCCCGCTGGAGGTCATACAGCATGAGTTTCAAAAACACCATGGCTCTCAAGGCAGA

GTAAATCGTCTTGAGACGGTCTGCATTGATTCACTTTCAACCCAGGTGGCGTTGGTTACT
TGCCATAAGCCTCTCTGGGGTTTCTTTGCTTGGACCTCTCTTGTCAGTTCTTCTGACGTA
GAGACAGGTGCGCGGTCTCATGGGCGTTGCCCTCTGGAACGTGCTAAAAAAAACAAAACA
AACAACAACAACAACCGTATCACACGTCAAGAGGGTCGGATCGTACTTGACGGCGGGAGA
TCCTTTGTTTCTCGCTTGTCCTGAGCGTCTGCCAAGCCTTCTTTGACGAGTGTGTTTGTT
CCTTCCATAAACCCGATCTCACATCAGGTAGGACCACCCGCTGAATTTAAGCATATAATT
AAGCGGAGGAAAAGAAACTAACAAGGATTCCCTCAGTAGCGGCGAGCGAAGCGGGAAGAG
CTCAACATGAAAACCGTAGGGGGTTCGCGCTCACCTTCGGGTTGTATAGTAAAGTGGTGT
CGCCTGCGTCGCGCCGGGTATAAATCCCGTTTGAAATGGGGTGTCGTAGAGGGTTATAAC
CCCGTCTGTGACCCGGTGTGCGATGCGTTACGGTGACCTTACGATGAGTCGGGTTGCTTG
GGCATGCAGCCCTAAGTGGGTGGTAAACTTCATCTAAAGCTAAATACTAGCGAGAGACCG
ATAGCGTACAAGTACCGTAAGGGAAAGATGCAAAGAACTTTGAAAAGAGAGTTAAACAGG
GCTTGAAACCGTTAAGAGTGAAGCGAAGGAGGTCAGTGATCATGGGGCCAGATCCCCACG
TGCCCGTCATAGGGCACCGTAGCGTTCTGGTTCCCGGGGCAGCGCCGGCCGGGGGCGGGT
GTCAACAGGGAGGTGGTTGAGGGATTAACTTTCCGCCTGTACTCCTTTCTAGCTTGCACC
CCGGCCGAGGGATCCGCGACTTGTCGTGTGAGGGCGCTGTCAAAATGGCCTTCTCCACCC
CGTCTTGAAACACGGACCAAGGAGTCTAACATACTCGCGAGTGTTTGGGTGGTTAAAACC
CGAACGCACAATGAAAGTAAACGGTGGCTCTGCGCTGCTGCTTGGTGGGAACCCTGCTCT
TATAACGGGGGGCACCATCACCCGACCCCGATCCCTTTGGGGTGGTTGGTTTTGAGGCAG
AGCGAGTATGATGGGACCCGAAAGATGGTGAACTATTCCTGAGTAGGGCGAAGCCAGAGG
AAACTCTGGTGGAGGCTCGTAGCGATGCTGACGTGCAAATCGCTCGTCGAACTTGGGTAT
AGGGGCGAAAGACTAATCGAACCATCTAGTAGCTGGTTCCCTCCGAAGTTTCCCTCAGGA
TAGCGGGAGCAAATGCAAGTCAGTATCATCCGGTAAAGCGAATGATTAGAGGCCTCGGGG
GAGTAACTCCTTCGACCTATTCTCAAACTTTAAATAGGTGAGGAGCCGGCAACGTTGCTT
AAAACAATGAACGCCGTGCGGACAATGATTGCTCTCAGTGGGCCGATTTTGGTAAGCAGA
ACTGGCGATGCGGGATGAACCGAAAGCGACGCTAAGGTGCCAAACTCCCAGCTCATTCAG
ACACCAGAAAAGGTGTTGATTCATACAGACAGCAGGACGGTGGCCATGGAAGTCGGAATC
CGCTAAGGAGTGTGTAACAACTCACCTGCCGAATGGATCAGCCCTGAAAATGGATGGCGC
TTAAGCGGGGGACCGAGGCGTCGCCGTTGGGGTAAATACGAAGCCTCAATGAGTAGGAGG
GCGTGGGGGTTTTGAAACACGGTACACACGCATGGAAGCCTTGGGCGTAAGCTCGGGTGG
AGCGACCCCTAGTGTAGATCTTGGTGGTAGTAGCAAATATTCAAGTGAGAACCTTGAAGA
CTGAAGTGGGGAAGGGTTCCACGCGAACAGTGCTTGGACGTGGGTTAGTCGGTCCTAAGA
GATGGGGAAACTCCGTATTCAAGTGCGCTCTTCCAGCGCAGTCCATCGAAAGGGAAGCGG
GTTAACATTCCCGCACCAGGATGTGGAAACTGTGGTTACCGCAAACTGCGGCAACGCAAC
TGAACTTGGGGACGCCCGTAGGAGCTCCGAGCAGGGATCTCTTTCCTGTTTAACGCGCCC
AACCCCCTGGAATCGGTTTGTCCGGAGATAGGGGGCGAAAGGCCGGAAAAGCACCGCACG

TCTTGTGGTGTTCGGAACGCTTGTAATGGCCCTTGAAAACCCAGGGGAAAGCAGCCGTCT

TACACTTTTTTGTGGGACGAGGAATTCTCACACCTGGCCGTACCCATCACCGCATCAGGT

CCCCAAGGTGAACAGCCTCTAGTCAATAGAATAATGTAGATAAGGGAAGTCGGCAAAATA

GATCCGTAACTTTGGGAAAAGGATTGGCTCTAATGTTCGGGCTTGCTGGGACGCGCGGGC

GTAGACGCCAATGAGCGGCAGACGTCTAGCCTCCTCGGAGGCTGGATTGACGCTGCTTGT

GGGAAGAAACCGTGTCGATGGAGGCAGGAGCGCTTGCCGTTCTTGTCTCTTCCTGCTTGC

GATCAACGAACGGATTAGAACTGGTACGGACAAGGGGAATCCGACTGTTTAATTAAAACA

AAGCATTGCGACGGCCGGAAACGGTGTTGGCGCAATGTGATTTCTGCCCAGTGCTCTGAA

TGTCAAAGTGAAGAAATTCAACCAAGCGCGGGTAAACGGCGGGAGTAACTATGACTCTCT

TAACTACATGGACATCTTTTGAGAGAGATAGGCGGGCTGAAAAGGTTATAGCCGTCTAGT

CTCGCTCTTTCATGAGCGGGGCGACACACCTGGTTGCGGGAACCCTGCCCACCTCTCTAT

CAGAAGTGGGAACCAGTCCATTCCTACTCCTGCTTGTGAGGCATATAGTAAAAAGTAGCA

GCAGGAGGATCTCGGCGAGAGGCCGAACCCGAAAGGAATAACGTGGGCTGGGAGGCAATC

CGCAGGCGAGTCCGTAACTATATGATTATCGACGGGCCGTCTCAACGACTGGTAAGGTGT

GGGTTTGTCCTCACGCGGGACAGGCTTAAGGTACAGTCTAATCCCACCCGCCAAGGGTGC

TTGGGTGTAATCCCTCTTACATAGAGGGCACTAGCCCAGGGGCTGGGACGCTGGGAGGAA

ATGCCCCGTCCCTATAGCCGGTAGAAATGAAGGTAGCCAAATGCCTCGTCATCTAAACTA

ATTGGATGAGATAGTCTGAGTGAAAAGCTTATGCAGGCTAGTACCGGTAAAAAAACGAGA

GGCTCTGCCACTCTGACGGTGCGACACTCCTTGTACAGGGAAACCCTCCGCATGGAAAGG

CACGTGGTACCACCCTCGCGAGGAAACTCGTGAGCGGGATCCCGGGTAATGACCGGAGCC

GATGGTAAAAACCCACGTGACCGATGGGGTGACCCTGTGGCGAGAAGCTACGTTGTGACT

TGTGTGCACAATATGCTTCCGTCGCAGAGACTGAACGGGAGTGGGCTCCGTCGACGGCCG

GCCAGCCGTCTTCCAGGAGCTTAAGATACAGTCCGGCCCACAGGGATGGCTTAAACCTGT

GGAACAAGCAGAATTAGTGACGCGCATGAATGGATTAACGAGATTCCCACTGTCCCTATC

TACTATCTAGCGAAACCACAGCCAAGGGAACGGGCTTGGAAGAATCAGCGGGGAAAGAAG

ACCCTGTTGAGCTTGACTCTAGTCCGACTTTGTGAGATGAGTTGGGAGGTGTAGTCTAAG

AGGGAGGGGCAACTCGACCGTGAAATACCTCTACTCCCAAGCTTGTTTTACTTACTCCGG

CGAGCAGGAATGGGTGGCAACACCCTTTTTCTAGGATTAACACCTCTTCGGGGGTGGACC

TGCGCGGAGGACATTGTCAGGTGGGGAGTTTGGCTGGGGCGGCACATCTGTTAAAAGATA

ACGCAGGTGTCCTAAGATGAGCTCAGTGAGAACAGAAATCTCACGTAGAACAAAAGGGTA

AAAGCTCATTTGATTTTGATTTTCAGTACGAATACAAACTGTGAAAGCATGGCCTATCGA

TCCTTTTGACCTTCAAGGTTTGAAGCAAGAGGTGTCAGAAAAGTTAGCACGCGTTGTGGT

AGCTTTTAAGAGTGGACTCAAAGTGTTCCGCTAGTCCCCCCTCGGACCGCCACTTACAAA

AAGGACAGAGGAGGGGGCGACACTGTCAAATTGCGGGGACTCCCTAAAGCCTGTCTCTAC

CAAGCCAGGCGGCGAAAGCTCCTGGTGGCCGGGGTAATGACCTAGGGTATGGTAAAAGCG

AGACAGGATGAGAACAATGGGCAATCCGCAGCCAAGTCCTACGGCTCTTTTCTTGGAGAA

GAGCTATGGATGCAGTTCAACGACTAGATGGCAGTGGGCTGAGGGAGCCTACGGCGGCAC
CCTTTCGGCTTAAGATATAGTCTTGCCCACGTGAAAGCGTGCCTGCAGATGAACGCCACC
ATGTATGGCAGGAGCTGCAGGAGGCCGCTAAACATACCAGCGGCTGGGAACATAACAGTA
CCACAGGGATGAACAAAAACACACAATTGTTGTGGCAATCATACAGTCGTATGATATTAT
CGTCCCAGGCGGGTGCGCCTCTCTCACCCGTCTAGTCCGGCCTTGGTGCGCAAAAAATGC
TTGCGTGGATGCGCCTTGGTTAAGGAAAATGCTCCTCGTAGTTTTACAAGGACATTTCTT
GTAGGGCAACACATCTGGATGCGAAGAACTCCCGGAAAGACAACGGTACCAAGTGACTGG
GGTAACGCCCCATTCATAAAGGCAATGAATAAAGAGGTGCCGGGTAATAATCTGTTGTAT
AGGGACAATCCGCAGCTGACTTGGCTCCCTGCCCCCATCGGTGGGCATAGCTTAGCGGTT
CAACGCTCGGTAAGATGTGGGTTGGCGCGCTGTCCAACAGCGCACTGGCTTAAGGTACGG
GCTACTCCCACCGGAGACGGTGTCTTTGTCTACCTCGGGTAACACGCCTCGGGAGAAGCA
AAGAGGGTGCAAGTAAAAAAACTGGTAGTGGGGAAATCCCTACCAGAGTCACCTGGTACA
AAGGAACTGGCTTGTGGCAGCCAAGCGTTCATAGCGACGTTGCTTTTTGATCCTTCGATG
TCGGCTCTTCCTATCATTGCGAAGCAGAATTCGCCAAGTGTTGGATTGTTCACCCACTAA
TAGGGAACGTGAGCTGGGTAGGCATTTTATGCCCTATGCCCAACAGCTGGATAGCTGGGC
TTCTGTCCTAGGCCGGGACTTCTCACGACCCGTGCCTTGGTCACCCAAAAAAGGGCGACC
ACAGAGCTCAGGATGGATCTTCTGGCTCAACTAGTAGAGGACAGGTAGAGATCGTCAAAT
TGCGGGAAAACCCTTAGAGCCTCGAGCTACCAAGTGCGTGTCGAAAGGCATAGACACAGC
CAGCAATGGCTGTCGCATGGCTTGATTCCATCAAGATAGGTAAAAATGCTCGAGGATTGG
GCAATCCGCAGCCAAGCCCCTAAGGTCCCCCCGTGGGGCTATGGGGAAGGTTCAACGACT
GTAAGGCGTACCGTGCAAGCGGAAGATACAGTCTAGCCCCGCTGGAAAGCAGCGCCAAAG
TAAGTAGGTTGCAGTGGATCTGTCCTC
>Pyr_27
GCCTGTTTTGATATGTGTTTTGACAGTCATACGGGTCATGGCCGAGGGATGGTGGGTGAG
CTTCATAGCGCAACCCCCCGTCAGGAGCATGTTTTCATAGCATTTTGAGCATACTAGGGT
GGTCTCCCATGTAGCTCGCTCTCTCGGCTCAGGACCTTCCGTATGGGGCACTTTTTGACG
AGCTCCTTTGGGGAGGAGCATGTTCTGGCATATGAAGCGTTGGTCACGAGCTTCGTAGCC
CGACACATACCAAATCAGTCTTCATATGTCTTTCTGGCGGGGGTCGGTCGTATGACCACG
CCCTTACCTTGAGCATGTTTTTCGTATGATACCTGACGGGCAGCGCCGCTGTTCAGTCGT
CGGCCGTACCTCCTATGCGCGTTTTCTGATTCGAACGTGTGACATTCAAGCGCATCATAT
CAGAAGCATGTTTTTTCATATGAGCATGTTGGTAGGCAGGCTCTTCAGTCGTCTACCCAT
AAATGCACTTTGAAAGACGTTTTTTGGTACTGAACGGCTTTGTGGTCAGACGTTCTCGTG
AGCTTATTCTCGCCCTCTGATACGCTGGATGCTGCGGTATTACCCACAGCAGACATCCAG
GAGCTCCCTCTGCGGATAAGCATCTGTAGAAGGGGGTTTCTCTCGGTGACGCAATGGCTG
GGAAAACGGAGTGCTATCCAGAAGGGTGAAACCTCTTGGGGATGTACTCCCATACCAGCA
TAATATGCTGAGAGGAATTGACTCAATGAGTGAGGTCCTATACTGCAACCTCTTTTGGAT

ATGTAGGCGGTCTGTACCCAGACTGTCCTACTGGCTGATTTCCCTTGGTCCCACGCGTCG
CTGGCTGAATGGGCTAGTCCTTATGAAGTGCCGCGTTCATCAAAAGAACCGGTCTTCATA
CAGCGACCTCATTCTATACTGCCAGCTGGATGGGCTGGTCCTTATGGAGCGCCGCATTCT
TCGAAGAATCGGTTTCTATAGAGCGACCCCATTTACATATAAGGAACCGTTACATAGACT
TGCCATTTGGCTGATTGCTCTGACAGTCGCTGCCTCTTCGGAGTTCTGTGTAAACGTGGC
GCCATTTTTGAGTGGGCCAGTCCTTATGGAGTGTCCGTGTCTTCTATGACAGGGTCCCCC
ATACAGCGACCTCACTCAGAACCGTATGGGCACGATGGTGAAAGACCATGCAGATAGTCC
GCCTTCCAAAGGGGGAAGCAGTATATGGGGAATGCTGGCGAAGCATGAGGTACATGGTTC
ATGGACCCGTTGGGCCATGCCTGTGCCATCTTCTCGAGTCAAACCGGTCCGTGTCGCTTC
CTTATCTCTTCCTTTCGCGACGACAAGCTCTTGAGCTGGTCGTGGGTCAGGACGCTGTGG
TATCTGTTTGTCTGTAGTGTGGGCTTCCACCAGAACAGCGTATCAGCACACGTCGTCTGG
GCATATGATGTTTTTGCTTTGTGGTCCCGGGTTTGGTAGTATGGTTCCATGTCGGTCCGC
TCACCTCCCAACCATAATGTCTTTCACATCATCTTCACAGTACGTCTTGTCAGCATACCA
TTCTGTATAGAACAGTGTATCAGCACACGTCGTCTGAGCATAGGGTATTTTTGCTTTGTG
GTCCCGGGTTTGGTAGTATGGTTCCATGTCGGTCCGCTCACCTCCCAACCATAATGTCAT
TTCTATCATCTGCACAGTACGTCTTGTCAGAATACCATTCTCTATGGCAGTCGATAGACT
AGGCCTTATGCCGCAGCAGTCTCTGATCACTGTGCGAGTGCATATATCTCTCAATGTGTT
CATGGCTAGCATGGGTCTTGAGTGAGTTGTGCCAGAGAGATACGCCACCAGTCTATGGAA
AGGTGCTTTTTGTATGAGAAGAGTACATGTATGAAGAGCATGTTCTCTCCTCTTATGATG
GTACCGCGGGTACGGGCTGCTCGCATTATGCTTGGCAGGTGTCCACTGTGCATGCAGGAT
TGCTGAACCCACTTGTATGGCCATGAGAATGGTCATTGGGTGAGGTTCACTTCTCCATCG
CTTTGTGTGTGTTAGATGTTTGGGTGGCTTCGGCCGCATGGAAATTCTCGGTCTGTAACT
CATCAGGGAGGGTTTCCATGTGAGTATTCTCCAGTCGACCACACACCCTTGGGCGTCCCG
CTAGGCAAAATGTAACTCTCGATCCGACCTAGCCTGTGGAGTGTCTTTCCGTATGGCGTG
AAAGGGCCCACACTGGATGTTTTCCCTTTTGGACACCAGTCTGGTAGTATGTATATATAG
ATCATCTACAGGGAGCAAAGGACAGGCATGCAAGTGCCACAAAGGCAGGGAGGCCTTCCT
GCTACTGTGTGTCCTTCTTGACGATCACGGGGGAGTATATAGAAACATTTGAAGGAGATC
CTTTTCCACAAGGCGGTGCCCTCGCGCAAAGGTGTTGCTCTGCACGCATCTGTATGATGA
TAGTTGCTCGCTCCTATGCGTCCACTTGTGAAAATACCTTCTCTCTTTGTCTGTTTCTAT
GACGCCCCTTCTGCTTTGTCATAGAGCATACTCAGGTACAGAAAGTGCCGTGACTTCCTT
GGCCTGCTGTCAAAGAAATAGCCTGGTGCTATAGAGGACTCTTCCTGGGCGCCGCCTGGT
GAAGAGGTCCTATCAGGCTCTGGAGTGTGCGTATGCCTATGTTCATCATAGCTCATTTGC
ACGCGCGGAAAAGCATCGACGTGGAATGGGGAAGAACACCATTCTGAATGAGCCAGTTCT
TCCTCTGCTCCCCTTGCTTGACGCTGTATGGGAATCGTTTTGCCTTGGGGGTACACCTTC
CATTCCATGACAAAGCATTCGATCTATGTCCCGAGGGGTGATTCTACATGAAGAACCCGA
AACTTGTCTGAGTGAGAGGCGAAGGTAGTCCTTCATTTTAATGCGGGATCCGCCCTCACA

CTGTAGTCGGCACAGTCCCTTGTTTAGGGACTGCGCAGCTCAATCTTCGACTGGAGAGGA
AAAAAACAAAACACAAGTATGACACGCACATACGTGTTTTCTAGGCCATGGACCTGTTGG
CTGCCTCGCGGTGGTTGGCATGATGCCCCAAGTGGGGGGGGGGGAAATCAATCAGTATGG
TTGACAAAATCCAGTGGGGAAAAAGGCGCGCTGTCGCACACCCTTGGTCACGGCCTCAAA
AAGTCGACGGCCCTCACTGGCCCACCCCTGCGACTCCTGTTCGCTTTTGACAAGACACAC
CTGCAATATGGAGGAAATGCCTGAGCAGCTGTGCCGCAGCAGCTGTGGACGCGTAGCGAT
ATGCATCCCTATTGTTTGCGACTTGGTACAAAGAACGCGTGCCGGGAGACGGATTGCCCT
TTGACTTTTCCACCCGGCCCGCGGCTCTGCCCTTTACGTACCCTTATTTACCTGGTTGAT
CCTGCCAGTAGTCATATGCTTGTCTCAAAGACTAAGCCATGCATGTCTAAGTATAAACAC
TTTTATACTCGTGAAACTGCGAATGGCTCATTAAAACAGTTATAGTTCCTTTGGGAACAA
CACTACTTGGATAACCGTAGTAATTCTAGAGCTAATACATGCCTCAACGCCCGACTCACG
AAGGGTGGTATTTATTGGATAAAAAACCATCGTGCTCTCTCCAGAGCGCTTTGAGATGAT
TCACAATAACTTGTCGGATCGCATGGCCTCGTGCTGGCGACTCCTCATTCAAATTTCTGC
CCTATCAACTTTCGATGGTAGAGTATTGGTCTACCATGGTGTCGACGGGTGACGGGGAAT
TAGGGTTCGATTCCGGAGAGGGAGCCTGAGAAACGGCTACCACATCCAAGGAAGGCAGCA
GGCGCGCAAATTACCCAATCCCGACTCGGGGAGGTAGTGACAAAAAATAACAATAGGGGG
CCCTTTGGGTCTTCTAATTGGAATGAGAACAATTTAAATCCCTTATCGAGGATCCATTGG
AGGGCAAGTCTGGTGAACGTATGCATCAAAGCCCTCGATAAAAAAAAACAGCGCCAGATG
TCTCTCTCTGGAAGCGCACCATGCATGGCTTTGTTCCTCTCTGCATGGCCGTGTGGAAGG
CGAGTAGGCAAACGATGGGAAACCAGGCTTGGAATAGTAACCAACCACAGAGCCCCCATC
ATACACTTCTAAATTGCCGGGGAGTGCCCGTGAGGCCTTGACTACCGCGGTCCGCCGAGA
GGTGAGAGCGAGTAATCGTTCAGGACGCAGCAGTGTACGTAGTGGTACATGGATGGTAAA
AACGTCAAGGATAGGGTTGATCCGCAGGGAAGCGCGAGTGCCGCCATCGTCGGTGGGCAA
AGGCGAACCTTCAGAGACTCTAATGGAGTGGGCGCAAGCTTAAGGGAGAGTCCAATCCAC
TGGGAAACCAGGTCCACAGCAGTAACGGGGTGATGCTCGCCCCAGAGGACTGTGGAAGGC
ACCACCACCCCAAGAGGGGGTGGGTGTTGTCGGGTAGCAAATGGCCAGCAGCCGCGGTAA
TTCCAGCTCCAATAGCGTATATTAAAGTTGTTGCAGTTAAAACGCTCGTAGTCGGACCTC
GGAGCGCCCCCACGGGCCGGGCTTTTGCCTTGTTGCCAGTGGGAGTTTGTCTCCTTTTGT
CGGGACGCGGTGTGCAGGGCCTTTACTGTCTCTGCGCGTTTACCTGCGGGCCGACCGTTT
ACTGTGAAGAAAGTAAAGTGTTCAAAGCAGGCCAATGCCTTGAATATGTGAGCATGGAAT
AATAGAATAGGACTTGGGCTCTATTTTGTTGGTTTCCAGTGACCAAGTAATGATTAATAG
GGACGGTTGGGGGCATTCGTATTTCATTGTCAGAGGTGAAATTCTTGGATTGATGGAAGA
CGTACAACTGCGAAAGCATCTGCCATGGATGTTTTCATTGATCAAGAACGAAAGTTAGGG
GATCGAAGACGATCAGATACCGTCGTAGTCTTAACCATAAACGATGCCGACTGGGGATTG
GCGGGGGCACTATTATATGACTCTGTCAGCACCCTGAGGGAAACCAAAGTCTTTGGGTTC
TGGGGGGAGTATGGTCGCAAGGCTGAAACTTAAAGGAATTGACGGAAGGGCACCACCAGG

AGTGGAGCCTGCGGCTTAATTTGACTCAACACGGGAAAACTTACCAGGTCCGGACAGAAG
AATGATTGACAGACTGAAAGCTCTTTCTTGATTTTTTGGTTGGTGGTGCATGGCCGTTCT
TAGTTGGTGGAGTGATTTGTCTGGTTAATTCCGTTAACGAACGAGACCTCGTCCTGCTAA
ATAGGTGCGCGCATGCCAGCACTGCGTTCTTACCTTCTTAGAGGGACTATGCGCGTCTAG
CGTATGGAAGATTGAGGCAATAACAGGTCTGTGATGCCCTTAGATGTTCTGGGCCGCACG
CGCGCTACACTGATGCATTCAACGAGTTTCACTTGATAGTCCTGGGTCGGAAGGCTCGGG
TAATCTTTTGAAAGTGCATCGTGCTGGGGATAGATCATTGCAATTATTGATCTTCAACGA
GGAATTCCTTGTAGGTGCAGGTCATCAGCCTGCGCCGAATACGTCCCTGCCCTTTGTACA
CACCGCCCGTCGCTCCTACCGATTGAATGGTCCGATGAAATGTCCGGATCGCGGCTAGTG
TGATGGACTCGTCCAGAGCGGTAGCCGTGAGAAGCTCATTAAATCTTATCATTTAGAGGA
AGGAGAAGTCGTAACAAGGTTTCCGTATTTTGTGGGTGGATACCGATTCAATAACATCCG
CCGTTTTTCTCAATGTACTATATCCCTAGTCTGCTTGCCTAGTTCTTGGAGTCTTGTGCG
ATGTGTGCTATTACAGGCATGCCGGGTCATGAAGACAGACAAGGTTGGGATCTTCACGGA
AGTACGCGCAATAGGAGCGCGTCTTGTTCAACGCACCTGACTCAAGAGTCATATGCGAGG
GACGGACGCAGCGCTTGTCGCCGCACAGATGGGAAGCTTCCTCTCCAGGCATAGGATGGC
GCTGCATTTGCATCACCGCTGCTAGTAAGTGGCAATAATACTTGACTCCCCGCCAGCGCA
CTTGTACGTAGCCGTTGGCGGCGGGAACGTAGTTGGTGATGTGACAGCCTTGAGCGTTCA
CCTCTGTGTGTCCGCCAATCTGCACAAAGATGGACGCCATATCGGAGACAGGGGGAGGTG
CCATTGTGATGATGACGATGTTGTTGGGTGGTCCTTCCGTAGGAAGACGTACACGAAGCC
TTTGCGGCCCGAAAGGGTGCACAGTGCGCGACTATAAACACTGATTCTGTGGTCAACGCA
AGCAGCGGTTCTGTGGAAGAACAGCTGCGACTTTCTCAAATTGCGGGAACACCCTAAAGT
CAACAAGCACCGCGGCTTGACGTGAAAAGCGTCAGGCGCAGCAGTCGGGGTAGCGCCCTG
GCGGATGGTAATAGCCTTGTTGAATGGCACAATGGGCAATCCGCAGCCAAGCCCCTAAGG
TCCCCCGCTGGGCTATGGGGAAGGTTCACAGACTGTAAGGGAGGGGGTTTCTTCCATGAA
AGAAGCTTAAGAGACAGTCGGTCCCCTGCGAGAGCAGTGTCGCATGTGAGGAAGGCACCC
ATGAAAAGTGGGTGCTGGAGAGCCATGTGAGCCCTCTATGAGGGTGGGAAGTTTTCCGAT
ACAATGAGAAACAAACACGTTCCGTAGGTGAACCTGCGGAAGGATCATTCACAAACTATT
GACACAAACACACGCGAACCAAATCGTCCACACAGGTGCCGATGAAAGAGAGAATCTGTT
TGTCGCCTTTTGGGGTATACAAGCACTCTTTTCCATCGCCTCTGTGCAGGGCGTAACTTC
CCATTGAGAGGATGTGAGGGCACCACAGGAAGCTTTTCCACAGGAAGTCGCCATCCTTCT
CCCTCCACGACGGCGGCGCTTCACAGGCTTGGCAGTTTTTTTTTTTTTTATTGCCTTCCA
GGGAGGATGCCGCCAATGGAGCCACATAGACATTTACATCACCATATGCCCGTTTTTCTT
TCTTAACCGCTTGCCACAGTTTCTCCTGTGAGGAGCTTGTGGGAAGACCGTCTCCATACA
ATAACAAAAGATACAACTCTTAGCGGTGGATATCTTGGTTCTCGCAACGATGAAGAACGC
AGCCAACTGCGATACCTAGTGTGAATTGCAGGACTCCGTGAATCATTAAGTCTTTGAACG
CAAGTTGCGCTCATCTCCATGAGGAGGTGAGCATGCCTGTTTGAGTGTCGATGGGAAAAA

CAATACATAACAACAATCGGCTCTCTTCCCTTCCTGCCTTCCCCGGCAAAAAAGGTAGTG

CATGCCGGAGATGGGCCTGGTCGCCCCTTCATGAAAGAGGCGTCGTCCGAAACACGTGTC

GTTGCCCGCTGGAGGTCATACAGCATGAGTTTCAAAAACACCATGGCTCTCAAGGCAGCG

TAAATCGTCTTGAGACGGTCTGGCATTGATTCACTTTCAACCCAGGTGGCGTTGGTTACT

TGCCATAAGCCTCTCTGGGGTTTCTTTGCTTGGACCTCTCTTGTCAGTTCTTCTGACGTA

GAGACAGGTGCGCGGTCTCATGGGCGTTGCCCTCTGGAACGTGCTAAAAAAACAAAACAA

ACAACAACAACAACAACCGTATCACACGTCAAGAGGGTCGGATCGTACTTGACGGCGGGA

GATCCTTTGTTTCTCGCTTGTCCTGAGCGTCTGCCAAGCCTTCTTTGACGAGTGTGTTTG

TTCCTTCCATAAACCCGATCTCACATCAGGTAGGACCACCCGCTGAATTTAAGCATATAA

TTAAGCGGAGGAAAAGAAACTAACAAGGATTCCCTCAGTAGCGGCGAGCGAAGCGGGAAG

AGCTCAACATGAAAACCGTAGGGGGTTCGCGCTCACCTTCGGGTTGTATAGTAAAGTGGT

GTCGCCTGCGTCGCGCCGGGTATAAATCCCGTTTGAAATGGGGTGTCGTAGAGGGTTATA

ACCCCGTCTGTGACCCGGTGTGCGATGCGTTACGGTGACCTTACGATGAGTCGGGTTGCT

TGGGCATGCAGCCCTAAGTGGGTGGTAAACTTCATCTAAAGCTAAATACTAGCGAGAGAC

CGATAGCGTACAAGTACCGTAAGGGAAAGATGCAAAGAACTTTGAAAAGAGAGTTAAACA

GGGCTTGAAACCGTTAAGAGTGAAGCGAAGGAGGTCAGTGATCATGGGGCCAGATCCCCA

CGTGCCCGTCATAGGGCACCGTAGCGTTCTGGTTCCCGGGGCAGCGCCGGCCGGGGGCGG

GTGTCAATAGGGAGGTGGTTGAGGGATTAACTTTCCGCCTGTACTCCTTTCTAGCTTGCA

CCCCGGCCGAGGGATCCGCGACTCGTCGTGTGAGGGCGCTGTCAAAATGGCCTTCTCCAC

CCCGTCTTGAAACACGGACCAAGGAGTCTAACATACTCGCGAGTGTTTGGGTGGTTAAAA

CCCGAACGCACAATGAAAGTAAACGGTGGCTCTGCGCTGCTGCTTGGTGGGAACCCTGCT

CTTATAACGGGGGGCACCATCACCCGACCCCGATCCCTTTGGGGTGGTTGGTTTTGAGGC

AGAGCGAGTATGATGGGACCCGAAAGATGGTGAACTATTCCTGAGTAGGGCGAAGCCAGA

GGAAACTCTGGTGGAGGCTCGTAGCGATGCTGACGTGCAAATCGCTCGTCGAACTTGGGT

ATAGGGGCGAAAGACTAATCGAACCATCTAGTAGCTGGTTCCCTCCGAAGTTTCCCTCAG

GATAGCGGGAGCAAATGCAAGTCAGTATCATCCGGTAAAGCGAATGATTAGAGGCCTCGG

GGGAGTAACTCCTTCGACCTATTCTCAAACTTTAAATAGGTGAGGAGCCGGCAACGTTGC

TTAAAACAATGAACGCCGTGCGGACAATGATTGCTCTCAGTGGGCCGATTTTGGTAAGCA

GAACTGGCGATGCGGGATGAACCGAAAGCGACGCTAAGGTGCCAAACTCCCAGCTCATTC

AGACACCAGAAAAGGTGTTGATTCATACAGACAGCAGGACGGTGGCCATGGAAGTCGGAA

TCCGCTAAGGAGTGTGTAACAACTCACCTGCCGAATGGATCAGCCCTGAAAATGGATGGC

GCTTAAGCGGGGGACCGAGGCGTCGCCGTTGGGGTAAATACGAAGCCTCAATGAGTAGGA

GGGCGTGGGGGTTTTGAAACACGGTACACACGCATGGAAGCCTTGGGCGTAAGCTCGGGT

GGAGCGACCCCTAGTGTAGATCTTGGTGGTAGTAGCAAATATTCAAGTGAGAACCTTGAA

GACTGAAGTGGGGAAGGGTTCCACGCGAACAGTGCTTGGACGTGGGTTAGTCGGTCCTAA

GAGATGGGGAAACTCCGTATTCAAGTGCGCTCTTCCAGCGCAGTCCATCGAAAGGGAAGC

GGGTTAACATTCCCGCACCAGGATGTGGAAACTGTGGTTACCGCAAACTGCGGCAACGCA
ACTGAACTTGGGGACGCCCGTAGGAGCTCCGAGCAGGGATCTCTTTCCTGTTTAACGCGC
CCAACCCCCTGGAATCGGTTTGTCCGGAGATAGGGGGCGAAAGGCCGGAAAAGCACCGCA
CGTCTTGTGGTGTTCGGAACGCTTGTAATGGCCCTTGAAAACCCAGGGGAAAGCAGCCGT
CTTACACTTTTTTGTGGGACGAGGAATTCTCACACCTGGCCGTACCCATCACCGCATCAG
GTCCCCAAGGTGAACAGCCTCTAGTCAATAGAATAATGTAGATAAGGGAAGTCGGCAAAA
TAGATCCGTAACTTTGGGAAAAGGATTGGCTCTAATGTTCGGGCTTGCTGGGACGCGCGG
GCGTAGACGCCAATGAGCGGCAGACGTCTAGCCTCCTCGGAGGCTGGATTGACGCTGCTT
GTGGGAAGAAACCGTGTCGATGGAGGCAGGAGCGCTTGCCGTTCTTGTCTCTTCCTGCTT
GCGATCAACGAACGGATTAGAACTGGTACGGACAAGGGGAATCCGACTGTTTAATTAAAA
CAAAGCATTGCGACGGCCGGAAACGGTGTTGGCGCAATGTGATTTCTGCCCAGTGCTCTG
AATGTCAAAGTGAAGAAATTCAACCAAGCGCGGGTAAACGGCGGGAGTAACTATGACTCT
CTTAAGGTAGCCAAATGCCTCGTCATCTAATTAGTGACGCGCATGAATGGATTAACGAGA
TTCCCACTGTCCCTATCTACTATCTAGCGAAACCACAGCCAAGGGAACGGGCTTGGAAGA
ATCAGCGGGGAAAGAAGACCCTGTTGAGCTTGACTCTAGTCCGACTTTGTGAGATGAGTT
GGGAGGTGTAGTCTAAGAGGGAGGGGCAACTCGACCGTGAAATACCTCTACTCCCAAGCT
TGTTTTACTTACTCCGGCGAGCAGGAATGGGTGGCAACACCCTTTTTCTAGGATTAACAC
CTCTTCGGGGGTGGACCTGCGCGGAGGACATTGTCAGGTGGGGAGTTTGGCTGGGGCGGC
ACATCTGTTAAAAGATAACGCAGGTGTCCTAAGATGAGCTCAGTGAGAACAGAAATCTCA
CGTAGAACAAAAGGGTAAAAGCTCATTTGATTTTGATTTTCAGTACGAATACAAACTGTG
AAAGCATGGCCTATCGATCCTTTTGACCTTCAAGGTTTGAAGCAAGAGGTGTCAGAAAAG
TTACCACAGGGATAACTGGCTTGTGGCAGCCAAGCGTTCATAGCGACGTTGCTTTTTGAT
CCTTCGATGTCGGCTCTTCCTATCATTGCGAAGCAGAATTCGCCAAGTGTTGGATTGTTC
ACCCACTAATAGGGAACGTGAGCTGGGTAGGCATTTTATGCCCTATGCCCAACAGCTGGA
TAGCTGGGCTTCTGTCCTAGGCCGGGACTTCTCACGACCCGTGCCTTGGTCACCCAAAAA
AAGGGCGACCACAGAGCTCAGGATGGATCTTCTGGCTCAACTAGTAGAGGACAGGTAGAG
ATCGTCAAATTGCGGGAAAACCCTTAGAGCCTCGAGCTACCAAGTGCGTGTCGAAAGGCA
TAGACACAGCCAGCAATGGCTGTCGCATGGCTTGATTCCATCAAGATAGGTAAAAATGCT
CGAGGATTGGGCAATCCGCAGCCAAGCCCCTAAGGTCCCCCCGTGGGGCTATGGGGAAGG
TTCAACGACTGTAAGGCGTACCGTGCAAGCGGAAGATACAGTCTAGCCCCGCTGGAAAGC
AGCGCCAAAGTAAGTAGGTTGCAGTGGATCTGTCCTC
>SRR9587917_SRR9587918_SRR9587938_SRR9587939_SRR9587943_SRR9587958
GCATGTTTTGGTATGTGTTTCGACAGTCATGCATGTCATAGCCGTAGGATGGGGAGTGAG
CTTCATAGCCCGACCCCTTCTCAGGAGCATGTTTTCATAGCATATTGAGCATACTGGGGT
GGTCTCCCATGTAGCTCGCTCTCTCGGTGCAGGACCTCCCTTTTGCGGGCCACTTCGATG
GCCTCCATTATGGAGAGCATGTTTTGGCATATGAAGCGTTGGTGGTGAGGCTCGTAGCCT

GTATATACATACCAACGCAGTCGTCATATGTCTCTCTGGCGGGGGTCGGTCGTATGACCT
TGTCCCTACCTGGAGCATGTATTTTGTATGATATACTGGACGGCAGCTGCGCTGTTCAGT
CGTCAGCCGTACCTCATGTGCGCGTTTCCAGGTCCGAACGTTTGACATGCAAGGGCATCA
TACCAGAAGCCTGTTTTCTCGTATGAGCATGTAGGTAGGCAGGCTCTTCAGTCGTCTACC
CATAAATGCACTTCGAAGGAACTTTTTGGTATTGAATGGCTTTGTGCCCAGACGTTCTTG
TGAGCTTGTTCTCGCTCTGCGATACGCTGGAAGCTGTGGTATTACCTACAGCAGACGCAT
TCCTAGGGCGGACGGCCTCTCTGCAGATGGGCACATCTGTAGAAAGAGGCTCCCCTTGAT
GACGCAATGCAATGGCTGGGTAAACGGAGTGCTATACAGAAGGGTCAAACCTCTGTGGAA
GTACACTCATACCAGCATTGTTTGTTGAGAGGGGTCGGCTCTACGAGTGAGGCCCTATAC
TGCCATCTCATTCAGATGCGTAGACGGTCTTTACTCAGACTGCCTTTCTGGCTGATTGTC
CTCGGTCCCTCGTGCGCACGCGCCGCTGGCTGAATGGGCTTGTCCTTATGGAGTGCCGCG
TTCTTCAAAGAATCCGTCTTCATACAGCGACCTCATTTACATATTGCCGGCTGAATGGGC
TTGTCCTTATGGAGCACTGCATTCTTCTAGGAATTGCTATCTATACAGCGACCTCATTTA
CATACAAGGCACCTTTTTCTACATAGACTTGCCATTTGGCTGATTGTTCTGACAGTCGCT
GCCTCTTCGGAGTTCTGTGTGGACGAGTGGCCTCCTGAGTGGGCCGGTCCTTACGGAGTG
CCCAAGTCTGTTATGACTGGGTTCTCCATACAGCGACCTCACTCAGAATTGTATGGCACC
GTGCCGATTGAATGGACAAGCAGTCCTTAGGGAATCGCATGAAAGCATGACGGCAACATG
GTGTATGGACCCGTTGGGCCTTGCCTGTGCTATCTTCTCGAGTCGACCGGTCCTCGTCTC
CTCCTTTCGCGACGGGAAGCTCTTGAGTTTACCGTGGGTCAGGACGCTGTGGTATCTGTT
TGTCTGTAGTGTGGGCTCTCTATCAGAACAGCGTATCAGCACACGTCGTCTGGGTCTATG
GTATTTTTGCTTAACGGCCCTGGGTTTGGTAGTATGGTTCCATGTCGGTCCTCTCACCTC
CCAACCGTCATGTCTCTGATATCATCTGCACAGTACGTCTTGTCAGAATATCATTCTGTA
TGGCAGTCGATAGACTAGGCCCTATGCCGCAGCTGTCTCTGACCACTATGCGAGCGCATA
CCTACCTCAATGTGTTCATAGCTGTTATGGATCTTGGGTGTGGTGTGCCAGAACGATACG
CCACCAGATTGTGGAATGGTGCTTGTAGTATTAGAAGAGGACATGTATGAAGAGCACGTT
CTTTTCTCTTATGATGGTGCCGCGGGTGCGGGCCGTTCTCATTCTGCTTGTCGGGTGCTC
AAGGTGCATGTAGGATTGCTGAACCCACTGAATGGTCATGAGAATGGTCATATGGTGAGG
TTCACTTCTCCATCGCTTTGTGTGTGTTGGATGTTTGGGTGGCTTCGGCCGCATGGAAAA
TCTCGGTCTGTAACTCATCAGGGAGGGTTTCCATGTGAGCAATCTCTCCAGTCGAACACA
CACCCATGGGCGTCCCGCTAGGCAAAATGGAACTCTCGATCCGACACAGCCTGTGGAGTG
TTTGTTCCGTGTGGCGTGAAACGGCTTATACTGGATGTCTTCCCTTTGGACACCAGTCTG
TTGGTATGTACGCATACCATCTAGAGGGAGCAAGAAAAAGGCATGCAAGTGCCACAAAGG
AAGGGAGGCCTTCCTGCTACCGTGTGTCCTCTTGACGATCACGGGGGAGTATATAGAGAC
ATTTGGACGAGATCCGTTTTTCACGAGGCGGTGCCCTCGCGCAAAGGTGTTGCTCTGCAC
GCATCTTTATGATGATAGTTGCTCGCTCCTATGCGTCCACTCGTGAAAATACCTGCTCTC
TCTGTCTGTTTCCATGACACCCCCATGCGTTGTCATAGAGCATGCTCAGGTACAGAAAGT

GCTGTGACTTTCTTGGCCTGCTGTCACTGAGTCAGCCTGATACTTTGGAGGAATCTTTTC
TGGGCACTGCCCAGCTGAGAGTTCTTATCAGGCTCTGGAGTGTGCATATGCCTATATTCA
TTATAGTGCATGTGGCTACGCAGAAAAAGCATCGACGTGGAATGGAGAAGAACACCATTC
TGAATGGAACGGTTCTTACTCTACTTCTGTTGCTTGACGCTGTATGGGAATCGCTTTGCT
CTGGGGGTACACCTTCCATTCCATGACAAAGCATTTGATCTATGCCCCGAGGGTCATTCT
CTATGAAGAGCCCGAAACTTGTCTGAGTGAGAGGCGAAGGTAGTCCTTCATTTATGCGGG
GTCCGCCCTCACACTGTAGTCGGCACAGTCCCTTGTTTATGGACTGCGCAGCTCAATCTT
CGACTGGAGAGGAAAAACAAAACACAAGTATGACACGCACATACGTGTTTTCTAGGACTG
TGACCTGTCGGCTGCCTCGCGGTGGTTGGCATGATGCCCCAAGTGGGGGGGGGAGAAAC
AATCAGTATGATTGATAAAAACCAGTGGGGAAAAAGGCGCGCTGTCGCACACCCTTGGTC
ACGGCCTCAAGTCGACGGCCCTCACTGGCCCACCCTGCGACTCCTGTTCGCTTTTGACAA
GACACACCTGCAATACGGAGGAAATGCCTGAGCAGCTGTGCCGCAGCAACTGCGGACGCG
TAGCGATATGCATCCCTGTTGTTTGCGACTTGGTACAAAGAACGTGCGCCGGGAGACGGA
TTGCCCTTTGACTTTTCCACCCAGCCCGCGGCTCTATACCCTTGACGTACCTATTTACCT
GGTTGATCCTGCCAGTAGTCATATGCTTGTCTCAAAGACTAAGCCATGCATGTCTAAGTA
TAAACACTTTTATACTCGTGAAACTGCGAATGGCTCATTAAAACAGTTATAGTTCCTTTG
GGAACAACACTACTTGGATAACCGTAGTAATTCTAGAGCTAATACATGCCTCAACGCCCG
ACTCACGAAGGGTGGTATTTATTGGATAAAAAACCATCGTGCTCTGTCCAGAGCGCTTTG
AGATGATTCACAATAACTTGTCGGATCGCATGGCCTCGTGCTGGCGACTCCTCATTCAAA
TTTCTGCCCTATCAACTTTCGATGGTAGAGTATTGGTCTACCATGGTGTCGACGGGTGAC
GGGGAATTAGGGTTCGATTCCGGAGAGGGAGCCTGAGAAACGGCTACCACATCCAAGGAA
GGCAGCAGGCGCGCAAATTACCCAATCCCGACTCGGGGAGGTAGTGACAAAAAATAACAA
TAGGGGGCCCTTTGGGTCTTCTAATTGGAATGAGAACAATTTAAATCCCTTATCGAGGAT
CCATTGGAGGGCAAGTCTGGTGAACGTATGCATTAAAGCCCTCCGTAATAAAAAAAAACA
GCGCCAGATGTCTCTCTCTGGAAGCACACCATGCATGGGGTTTGTTCCTCTCTGCATGGC
CGTGTGAAAAAGGCGAGTAGGCAAACGATGGGAAACCCGGCTTGGAATAGTAACCAGCCA
CAGAGCCCCCATCATACACTTCTAAATTGCCGGGGAGTGCCCGCGAGGCCTTGACTACCG
CGGTCCGCCGAGAGGTGAGAGCGAGCAATCGTTCAGGACGCAGCAGTGTACGTAGTGGTA
CATGGATGGTAAAAACGTCAAGGATAGGGTTGATCCGCAGGGAAGCGCGAGTGCCGCCAT
GGTCGGTGGGCAAAGGCGAACCTTCAGAGACTCTAATGGAGTGGGCGCAAGCTTAAGGGA
GAGTCCAATCCACTGGGAAACCAGGTCCACAGCAGTAACGGGGTGATGCTCGCCCCAGAG
GACTGTGGAAGGCAACCCCACCAAAAGGGTGTGTGTTGTCGGGTAGCAATGGCCAGCAGC
CGCGGTAATTCCAGCTCCAATAGCGTATATTAAAGTTGTTGCAGTTAAAACGCTCGTAGT
CGGACCTCGGAGCGCCCCCACGGGCCGGGCTTTTGCCTTGTTGCCAGTGGGAGTTTGTCT
CCTTTTGTCGGGACGCGGTGTGCAGGGCCTTTACTGTCTCTGCGCGTTTACCTGCGGGCC
GACCGTTTACTGTGAAGAAAGTAAAGTGTTCAAAGCAGGCCAATGCCTTGAATATGTGAG

CATGGAATAATAGAATAGGACTTGGGCTCTATTTTGTTGGTTTCCAGTGACCAAGTAATG

ATTAATAGGGACGGTTGGGGGCATTCGTATTTCATTGTCAGAGGTGAAATTCTTGGATTG

ATGGAAGACGTACAACTGCGAAAGCATCTGCCATGGATGTTTTCATTGATCAAGAACGAA

AGTTAGGGGATCGAAGACGATCAGATACCGTCGTAGTCTTAACCATAAACGATGCCGACT

GGGGATTGGCGGGGGCACTTTTATATGACTCTGTCAGCACCCTGAGGGAAACCAAAGTCT

TTGGGTTCTGGGGGGAGTATGGTCGCAAGGCTGAAACTTAAAGGAATTGACGGAAGGGCA

CCACCAGGAGTGGAGCCTGCGGCTTAATTTGACTCAACACGGGAAAACTTACCAGGTCCG

GACAGAAGAATGATTGACAGACTGAAAGCTCTTTCTTGATTTTTTGGTTGGTGGTGCATG

GCCGTTCTTAGTTGGTGGAGTGATTTGTCTGGTTAATTCCGTTAACGAACGAGACCTCGT

CCTGCTAAATAGGTGCGCGCATGCCAGCACTGCGTTCTTACCTTCTTAGAGGGACTATGC

GCGTCTAGCGTATGGAAGATTGAGGCAATAACAGGTCTGTGATGCCCTTAGATGTTCTGG

GCCGCACGCGCGCTACACTGATGCATTCAACGAGTTTCACTTGATAGTCCTGGGTCGGAA

GGCTCGGGTAATCTTTTGAAAGTGCATCGTGCTGGGGATAGATCATTGCAATTATTGATC

TTCAACGAGGAATTCCTTGTAGGCGCAGGTCATCAGCCTGCGCCGAATACGTCCCTGCCC

TTTGTACACACCGCCCGTCGCTCCTACCGATTGAATGGTCCGATGAAATGTCCGGATCGC

GGCTAGTGTGATGGACTCGTCCAGAGCGGTAGCCGTGAGAAGCTCATTAAATCTTATCAT

TTAGAGGAAGGAGAAGTCGTAACAAGGTTTCCGTATTTTGTGGATGGGCACTCGTCGTGG

CGTTCTTTGCGCCACGCGGATGTCCCCCTTCCGTAGGAAAACGTACACGAAGCCTTTGCG

GCCCGAAAGGGTGCACAGTGCGCGACTATAAACACTGATTCTGTGGTCAACGCAAGCAGC

GGTTCTGTGACAGAACAGCTGCGACTTTCTCAAATTGCGGGAACACCCTAAAGTCAACAA

ACACCGCGGCCGGACGTGAAAGCGTCAGGCGCAGCAGTCGGGGTAGCGCCCTGGCGGATG

GTAAGAGCTTTGTTGAATGGCACAATGGGCAATCCGCAGCCAAGCCCCTAAGGTCCTTCG

CGGCTATGGGGAAGGTTCACAGACTGTAAGGGAGGGGGTTTCTTCCATGAAGAAGAAGCT

TAAGAGACAGTCGGTCCCCTGCGAGAGCAGTGTCTCATGTGAGGAAGGTGCCCATGAAAA

GTGGGCGCTGGAGAGCCATGGGAGCCCTCTATGAGGGTGGGGAGTTTTCCTAGATAATAA

TGAGAAACAAACACGTTCCGTAGGTGAACCTGCGGAAGGATCATTCACAAACTATTGACA

CAAACACACGCGAACCAAATCGTCCGCACAGCAGGTGGCAATGAAAGAGAGACTCTGCAT

GTCGCCTTTCGGGGTAGCAAGCAGCACTCTTCTGCCATCGCCTCTGTGCAGGGCGTAAAT

TCCCATTGAGAGGATGTGAGGGCACCACAGGAAGCTTTTCCACAGGAAGTCGCCATCCTT

CTCCCTCCACGGCGGCGCTTCTGTGCTTGGCAGTTTTTTTTGCCTTCTAGGGAGGATGCC

GCCAATGGAGCCCCTTATATATATACATCATCATATGCCCTTTTTTTCTTAACCGCTTGC

CACAGCTTCTTCTATTAGGAGCTTGTGGGAAGACTGTCTCCATACAATAACAAAGATACA

ACTCTTAGCGGTGGATATCTTGGTTCTCGCAACGATGAAGAACGCAGCCAACTGCGATAA

CTAATGTGAATTGCAGGACTTCGTGAATCATTAAGTCTTTGAACGCAAGTTGCGCTCATC

TCCATGAGGAGGTGAGCATGCCTGTTTGAGTGTCGATGGGAAAAACAATACATAACAACA

ATCGGCTCTCTTATCTTCCTGCCTTCTTGGGGGCAAAAGGTAGTGCATGCCGGAGATGGG

CTTTGTCGCCCCTTTCTCTTGAAAGAGGCGTCGTCCGAAACACGTATCGTTGCCCGCTGG
AGGTCATAGAGCATGAGTTTAAAGCACCACATCTCTCAAGTCAGAGTGAGCGTCTTGAGT
AGGTCCGCATTGATTCAGTTTCAACCCAGGTGGCGTTGGTTACTTGCCATAAGCCTCTCT
GGGGTTTCTTTGCTTGGACCTCTCTTGTCAGTTTCTCTGACGTAGAGACAGGTGCCGTCT
CATGGACGTTGCCCTCTGGAACGTGCTAAAAAAAAAAAAGAAAAAACCGTACCACATGTC
AAAAGAGGCAGATCGTGCTTGACGGCGGGAGGTGTTGCATTTCTTGCTTGTCCTGAGCGT
CTGCCAGTCTCTGACGAGTGTGTTCCTTCCATAAACCCGATCTCAAATCAGGTAGGACCA
CCCGCTGAATTTAAGCATATAATTAAGCGGAGGAAAAGAAACTAACAAGGATTCCCTCAG
TAGCGGCGAGCGAAGCGGGAAGAGCTCAACATGAAAACCGTTGGGGGTTCGCGCTTTCCT
TCGGGTTGTATAGTAAAGTGGTGTCGCCTGCGTCGCGCCGGGTATAAATCCCGTCTGAAA
TGGGGTGTCGTAGAGGGTTATATCCCCGTCTGTGACCCGGTGTGCGATGCGTTACGGTGA
CCTTACGATGAGTCGGGTTGCTTGGGCATGCAGCCCTAAGTGGGTGGTAAACTTCATCTA
AAGCTAAATACTAGCGAGAGACCGATAGCGTACAAGTACCGTAAGGGAAAGATGCAAAGA
ACTTTGAAAAGAGAGTTAAACAGGGCTTGAAACCGTTAAGAGTGAAGCGAAGGAGGTCAG
TGATCATGGGGCCAGATCCCCAAGTGCCTGTCATAGGGCACCGTAGCGTTCTGGTTTCCC
GGGGCAGCGCCGGCCGGGGGCGGGGGTTAACAGGGAGGTGGTTGAGGGATTCGTTTTCCG
CCTGTACTCCTTTATAGCTTGCACCCCGGCCGAGGGATCCGCGACTTGTCGTGTGAGGGC
GCTGTCAAAATGGCCTTCTCCACCCCGTCTTGAAACACGGACCAAGGAGTCTAACATACT
CGCGAGTGTTTGGGTGGTTAAAACCCGAACGCACAATGAAAGTAAACGGTGGCTCTGCGC
CGCTGCTTGGTGGGAACCCTGCTTTTATAATGGGGGGCACTATCACCCGACCCCGATCCC
TTTGGGGTGGTTGGTTTTGAGGTAGAGCGAGTATGATGGGACCCGAAAGATGGTGAACTA
TTCCTGAGTAGGGCGAAGCCAGAGGAAACTCTGGTGGAGGCTCGTAGCGATGCTGACGTG
CAAATCGCTCGTCGAACTTGGGTATAGGGGCGAAAGACTAATCGAACCATCTAGTAGCTG
GTTCCCTCCGAAGTTTCCCTCAGGATAGCGGGAGCAAATGCAAGTCAGTATCATCCGGTA
AAGCGAATGATTAGAGGCCTCGGGGGAGTAACTCCTTCGACCTATTCTCAAACTTTAAAT
AGGTGAGGAGCCGGCAACGTTGCTTAAAACAATGAACGCCGTGCGGACAATGATTGCTCT
CAGTGGGCCGATTTTGGTAAGCAGAACTGGCGATGCGGGATGAACCGAAAGCGACGCTAA
GGTGCCAAACTCCCAGCTCATTCAGACACCAGAAAAGGTGTTGATTCATACAGACAGCAG
GACGGTGGCCATGGAAGTCGGAATCCGCTAAGGAGTGTGTAACAACTCACCTGCCGAATG
GATCAGCCCTGAAAATGGATGGCGCTTAAGCGGGGGACCGAGGCGTCGCCGTTGGGGTAA
ATACGAAGCCCCAATGAGTAGGAGGGCGTGGGGGTTTTGAAACACGGTACACTCGCATGG
AAGCCTTGGGCGTAAGCTCGGGTGGAGCGACCCCTAGTGTAGATCTTGGTGGTAGTAGCA
AATATTCAAGTGAGAACCTTGAAGACTGAAGTGGGGAAGGGTTCCACGCGAACAGTGCTT
GGACGTGGGTTAGTCGGTCCTAAGAGATGGGGAAACTCCGTATTCAAGTGCGCTTTTGTA
GCGCAGTCCATCGAAAGGGAAGCGGGTTAACATTCCCGCACCAGGATGTGGAAACTGTGG
TTTACCGCAAACTGCGGCAACGCAACTGAACCTGGGGACGCCCGTAGGAGCTCCGAGCAG

GGATCTCTTTCCTGTTTAACGCGCCCAACCCCCTGGAATCGGTTTGTCCGGAGATAGGGG
GCGAAAGGCCGGAAAAGCACCGCACGTCTTGTGGTGTTCGGAACGCTTGTAATGGCCCTT
GAAAACCCAGGGGAAAGCAGCCGTCTTACACTTGTGTGAGACGAGGAATTCTCACACCTG
GCCGTACCCATCACCGCATCAGGTCCCCAAGGTGAACAGCCTCTAGTCAATAGAATAATG
TAGATAAGGGAAGTCGGCAAAATAGATCCGTAACTTTGGGAAAAGGATTGGCTCTAATGT
TCGGGCTTGTCGGGACGCGCGGGCGTAGATGCCAATGAGCGGCAGACGTCTAGCCTCTTT
GGAGGCTGGATTAACGCTGCTTGTGGGAAGAAACCGTGTCGATGGAGGCAGGAACGCTTA
TCGATCTTGTCTCTTCCCGCTTGCGATCAACGAACGGATTAGAACTGGTACGGACAAGGG
GAATCCGACTGTTTAATTAAAACAAAGCATTGCGACGGCCGGAAACGGTGTTGGCGCAAT
GTGATTTCTGCCCAGTGCTCTGAATGTCAAAGTGAAGAAATTCAACCAAGCGCGGGTAAA
CGGCGGGAGTAACTATGACTCTTAAACAAACCATACTACCTACGGGAGTCAAAAGTGGGG
GCGGCGTCTGGTGTAGAAATACACGTACGACACGCTTTTTTTCCTGCTAGTCGATCTGAC
CTTTGTTCTTTGGACTGTGGGCGGTCAGCGGCAACATAACCTGGTACGGGGAAGGCCTCG
ACGTTGAGTGCTGCATCTTGGAAGTATCTTGACACAATAAGAGAAAAGAGATTGCGCTTA
ATAGGGGGTTGATCTCGTGGCGAGCCGACGCCGGTGTGAACTGGTGGCAATGGCCGTCGT
AGAGCACGCAAAGGTATGGGCTGACTTTTTCTCCTCAAAAGTTGGCTTAAGGGACGTGCC
ATCCCACACGATAATAAAGTGTGCAGTCTGTAAGAGCTCCCACAGGGCAAAGACAGACTG
GGATGCAGACGGAGACGGCAAAAACAACAACATCCGTTCTGCTTCTGGGCGCAAATGCTT
AAGGTAGCCAAATGCCTCGTCATCTAAACTAATTGGATGAGATAGTCTGAGTGAAAAGCT
TATGCAGGCTAGTACCGGAAAAAAAAAAAAGAGGGGCTCTGCCACTCTGACGGTGCGACA
CTCCTTGTACAGGGAAACCCTCCGCATGGAAAGGCACGTGGTACCACCCTCGCGGGGAAA
CTCGCGAGCGGGATCCCGGGTAATGACCGGAGCCGATGGTAAAAACTCACGTGACCGATG
GGGTGACCCTGTGGCGAGACGCTAAGTTGTGGGACATCACCCTCTCACAATACGCATCCG
TCGCAGAGACTGAACGGGAGTGGGCTCCGTTGACGGCCGGCCGGCCAGCCAGCCGTCTTC
CAGGGGCTTAAGATACAGTCCGGGCCTCAGGGATGGAAAAACCTGGGGAACTACCAGAAT
TAGTGACGCGCATGAATGGATTAACGAGATTCCCACTGTCCCTATCTACTATCTAGCGAA
ACCACAGCCAAGGGAACGGGCTTGGAAGAATCAGCGGGGAAAGAAGACCCTGTTGAGCTT
GACTCTAGTCCGACTTTGTGAGATGAGTTGGGAGGTGTAGTCTAAGAGGGAGGGGCAACT
CGACCGTGAAATACCTCTACTCCCAAGCTTGTTTTACTTACTCCGGCGAGCAGGGATGGG
TGGTAACACCCTTTTTCTAGGATTAACACCTCTTCGGGGGTGGACCTGCGCGGAGGACAT
TGTCAGGTGGGGAGTTTGGCTGGGGCGGCACATCTGTTAAAAGATAACGCAGGTGTCCTA
AGATGAGCTCAGTGAGAACAGAAATCTCACGTAGAACAAAAGGGTAAAAGCTCATTTGAT
TTTGATTTTCAGTACGAATACAAACTGTGAAAGCATGGCCTATCGATCCTTTTGACCTTC
AAGGTTTGAAGCAAGAGGTGTCAGAAAAGTTACCACAGGGATAACTGGCTTGTGGCAGCC
AAGCGTTCATAGCGACGTTGCTTTTTGATCCTTCGATGTCGGCTCTTCCTATCATTGCGA
AGCAGAATTCGCCAAGTGTTGGATTGTTCACCCACTAATAGGGAACGTGAGCTGGGTAGC

CACTTTATGCCCAACAGCTGGATAGCTGGGCTTCTGTCTTAGGCCGGGACTTCTCACGAC
CCGTGCCTTGGTCACCCAAAAAAGGGCGACCGCAGAGCTCAGGATGGAGCTTCTGGCTTA
ACTAGTAGAGGACAGATAGAGATCGTCAAATTGCGGGAAACCCCTTAGAGCCTCGAGCTA
CCAAGTGCGTGTCGAAAGGCATAGACGCAGCCAGCGATGGCTGTCGCATGGCTTGATTCC
ATCAAGATAGGTAAAAATGCTCGAGGATTGGGCAATCCGCAGCCAAGCCTCTAAGGTCCC
CCCGCGGGCTATGGGGAAGGTTCAACGACTGTAAGGCGTACCGTGCAAGCGGAAGATACA
GTCTAGCCCCGCTGGAAAGCAGCGCCAAAGTAAGTAGGTTGCAGTGGGTCTGTCCTC
>SRR9587919_SRR9587951_SRR9587953
GCATGTTTTGGTATGTGTTTCGACAGTCATGCATGTCATAGCCGTAGGATGGGGAGTGAG
CTTCATAGCCCGACCCCTTCTCAGGAGCATGTTTTCATAGCATATTGAGCATACTGGGGT
GGTCTCCCATGTAGCTCGCTCTCTCGGTGCAGGACCTCCCTTTTGCGGGCCACTTCGATG
GCCTCCATTATGGAGAGCATGTTTTGGCATATGAAGCGTTGGTGGTGAGGCTCGTAGCCT
GTATATACATACCAACGCAGTCGTCATATGTCTCTCTGGCGGGGGTCGGTCGTATGACCT
TGTCCCTACCTGGAGCATGTATTTTGTATGATATACTGGACGGCAGCTGCGCTGTTCAGT
CGTCAGCCGTACCTCATGTGCGCGTTTCCAGGTCCGAACGTTTGACATGCAAGGGCATCA
TACCAGAAGCCTGTTTTCTCGTATGAGCATGTAGGTAGGCAGGCTCTTCAGTCGTCTACC
CATAAATGCACTTCGAAGGAACTTTTTGGTATTGAATGGCTTTGTGCCCAGACGTTCTTG
TGAGCTTGTTCTCGCTCTGCGATACGCTGGAAGCTGTGGTATTACCTACAGCAGACGCAT
TCCTAGGGCGGACGGCCTCTCTGCAGATGGGCACATCTGTAGAAAGAGGCTCCCCTTGAT
GACGCAATGCAATGGCTGGGTAAACGGAGTGCTATACAGAAGGGTCAAACCTCTGTGGAA
GTACACTCATACCAGCATTGTTTGTTGAGAGGGGTCGGCTCTACGAGTGAGGACCTATAC
TGCCATCTCATTCAGATGCGTAGACGGTCTTTACTCAGACTGCCTTTCTGGCTGATTGTC
CTCGGTCCCTCGTGCGCACGCGCCGCTGGCTGAATGGGCTTGTCCTTATGGAGTGCCGCG
TTCTTCAAAGAATCCGTCTTCATACAGCGACCTCATTTACATATTGCCGGCTGAATGGGC
TTGTCCTTATGGAGCACTGCATTCTTCTAGGAATTGCTATCTATACAGCGACCTCATTTA
CATACAAGGCACCTTTTTCTACATAGACTTGCCATTTGGCTGATTGTTCTGACAGTCGCT
GCCTCTTCGGAGTTCTGTGTGGACGAGTGGCCTCCTGAGTGGGCCGGTCCTTACGGAGTG
CCCACGTCTGTTATGACTGGGTTCTCCATACAGCGACCTCACTCAGAATTGTATGGCACC
GTGCCGATTGAATGGACAAGCAGTCCTTAGGGAATCGCATGAAAGCATGACGGCACATGG
TGTATGGACCCGTTGGGCCTTGCCTGTGCTATCTTCTCGAGTCGACCGGTCCTCGTCTCC
TCCTTTCGCGACGGGAAGCTCTTGAGTTTACCGTGGGTCAGGACGCTGTGGTATCTGTTT
GTCTGTAGTGTGGGCTCTCTATCAGAACAGCGTATCAGCACACGTCGTCTGGGTCTATGG
TATTTTTGCTTAACGGCCCTGGGTTTGGTAGTATGGTTCCATGTCGGTCCTCTCACCTCC
CAACCGTCATGTCTCTGATATCATCTGCACAGTACGTCTTGTCAGAATATCATTCTGTAT
GGCAGTCGATAGACTAGGCCCTATGCCGCAGCTGTCTCTGACCACTATGCGAGCGCATAC
CTACCTCAATGTGTTCATAGCTGTTATGGATCTTGGGTGTGGTGTGCCAGAACGATACGC

CACCAGATTGTGGAATGGTGCTTGTAGTATTAGAAGAGGACATGTATGAAGAGCACGTTC
TTTTCTCTTATGATGGTGCCGCGGGTGCGGGCCGTTCTCATTCTGCTTGTCGGGCGCTCA
AGGTGCATGTAGGATTGCTGAACCCACTGAATGGTCATGAGAATGGTCATATGGTGAGGT
TCACTTCTCCATCGCTTTGTGTGTGTTGGATGTTTGGGTGGCTTCGGCCGCATGGAAAAT
CTCGGTCTGTAACTCATCAGGGAGGGTTTCCATGTGAGCAATCTCTCCAGTCGAACACAC
ACCCATGGGCGTCCCGCTAGGCAAAATGGAACTCTCGATCCGACACAGCCTGTGGAGTGT
TTGTTCCGTGTGGCGTGAAACGGCTTATACTGGATGTCTTCCCTTTGGACACCAGTCTGT
TGGTATGTACGCATACCATCTAGAGGGAGCAAGAAAAAGGCATGCAAGTGCCACAAAGGA
AGGGAGGCCTTCCTGCTACCGTGTGTCCTCTTGACGATCACGGGGGAGTATATAGAGACA
TTTGGACGAGATCCGTTTTTCACGAGGCGGTGCCCTCGCGCAAAGGTGTTGCTCTGCACG
CATCTTTATGATGATAGTTGCTCGCTCCTATGCGTCCACTCGTGAAAATACCTGCTCTCT
CTGTCTGTTTCCATGACACCCCCATGCGTTGTCATAGAGCATGCTCAGGTACAGAAAGTG
CTGTGACTTTCTTGGCCTGCTGTCACTGAGTCAGCCTGATACTTTGGAGGAATCTTTTCT
GGGCACTGCCCAGCTGAGAGTTCTTATCAGGCTCTGGAGTGTGCATATGCCTATATTCAT
TATAGTGCATGTGGCTACGCAGAAAAAGCATCGACGTGGAATGGAGAAGAACACCATTCT
GAATGGAACGGTTCTTACTCTACTTCTGTTGCTTGACGCTGTATGGGAATCGCTTTGCTC
TGGGGGTACACCTTCCATTCCATGACAAAGCATTTGATCTATGCCCCGAGGGTCATTCTC
TATGAAGAGCCCGAAACTTGTCTGAGTGAGAGGCGAAGGTAGTCCTTCATTTATGCGGGG
TCCGCCCTCACACTGTAGTCGGCACAGTCCCTTGTTTATGGACTGCGCAGCTCAATCTTC
GACTGGAGAGGAAAAACAAAACACAAGTATGACACGCACATACGTGTTTTCTAGGACTGT
GACCTGTCGGCTGCCTCGCGGTGGTTGGCATGATGCCCCAAGTGGGGGGGGGAGAAACAA
TCAGTATGATTGATAAAAACCAGTGGGGAAAAAGGCGCGCTGTCGCACACCCTTGGTCAC
GGCCTCAAGTCGACGGCCCTCACTGGCCCACCCTGCGACTCCTGTTCGCTTTTGACAAGA
CACACCTGCAATACGGAGGAAATGCCTGAGCAGCTGTGCCGCAGCAACTGCGGACGCGTA
GCGATATGCATCCCTGTTGTTTGCGACTTGGTACAAAGAACGTGCGCCGGGAGACGGATT
GCCCTTTGACTTTTCCACCCAGCCCGCGGCTCTATACCCTTGACGTACCTATTTACCTGG
TTGATCCTGCCAGTAGTCATATGCTTGTCTCAAAGACTAAGCCATGCATGTCTAAGTATA
AACACTTTTATACTCGTGAAACTGCGAATGGCTCATTAAAACAGTTATAGTTCCTTTGGG
AACAACACTACTTGGATAACCGTAGTAATTCTAGAGCTAATACATGCCTCAACGCCCGAC
TCACGAAGGGTGGTATTTATTGGATAAAAAACCATCGTGCTCTGTCCAGAGCGCTTTGAG
ATGATTCACAATAACTTGTCGGATCGCATGGCCTCGTGCTGGCGACTCCTCATTCAAATT
TCTGCCCTATCAACTTTCGATGGTAGAGTATTGGTCTACCATGGTGTCGACGGGTGACGG
GGAATTAGGGTTCGATTCCGGAGAGGGAGCCTGAGAAACGGCTACCACATCCAAGGAAGG
CAGCAGGCGCGCAAATTACCCAATCCCGACTCGGGGAGGTAGTGACAAAAAATAACAATA
GGGGGCCCTTTGGGTCTTCTAATTGGAATGAGAACAATTTAAATCCCTTATCGAGGATCC
ATTGGAGGGCAAGTCTGGTGAACGTATGCATTAAAGCCCTCCGTAATAAAAAAAAACAGC

GCCAGATGTCTCTCTCTGGAAGCACACCATGCATGGGGTTTGTTCCTCTCTGCATGGCCG
TGTGAAAAAGGCGAGTAGGCAAACGATGGGAAACCCGGCTTGGAATAGTAACCAGCCACA
GAGCCCCCATCATACACTTCTAAATTGCCGGGGAGTGCCCGCGAGGCCTTGACTACCGCG
GTCCGCCGAGAGGTGAGAGCGAGCAATCGTTCAGGACGCAGCAGTGTACGTAGTGGTACA
TGGATGGTAAAAACGTCAAGGATAGGGTTGATCCGCAGGGAAGCGCGAGTGCCGCCATGG
TCGGTGGGCAAAGGCGAACCTTCAGAGACTCTAATGGAGTGGGCGCAAGCTTAAGGGAGA
GTCCAATCCACTGGGAAACCAGGTCCACAGCAGTAACGGGGTGATGCTCGCCACAGAGGA
CTGTGGAAGGCAACCCCACCAAAAGGGTGTGTGTTGTCGGGTAGCAATGGCCAGCAGCCG
CGGTAATTCCAGCTCCAATAGCGTATATTAAAGTTGTTGCAGTTAAAACGCTCGTAGTCG
GACCTCGGAGCGCCCCCACGGGCCGGGCTTTTGCCTTGTTGCCAGTGGGAGTTTGTCTCC
TTTTGTCGGGACGCGGTGTGCAGGGCCTTTACTGTCTCTGCGCGTTTACCTGCGGGCCGA
CCGTTTACTGTGAAGAAAGTAAAGTGTTCAAAGCAGGCCAATGCCTTGAATATGTGAGCA
TGGAATAATAGAATAGGACTTGGGCTCTATTTTGTTGGTTTCCAGTGACCAAGTAATGAT
TAATAGGGACGGTTGGGGGCATTCGTATTTCATTGTCAGAGGTGAAATTCTTGGATTGAT
GGAAGACGTACAACTGCGAAAGCATCTGCCATGGATGTTTTCATTGATCAAGAACGAAAG
TTAGGGGATCGAAGACGATCAGATACCGTCGTAGTCTTAACCATAAACGATGCCGACTGG
GGATTGGCGGGGGCACTTTTATATGACTCTGTCAGCACCCTGAGGGAAACCAAAGTCTTT
GGGTTCTGGGGGGAGTATGGTCGCAAGGCTGAAACTTAAAGGAATTGACGGAAGGGCACC
ACCAGGAGTGGAGCCTGCGGCTTAATTTGACTCAACACGGGAAAACTTACCAGGTCCGGA
CAGAAGAATGATTGACAGACTGAAAGCTCTTTCTTGATTTTTTGGTTGGTGGTGCATGGC
CGTTCTTAGTTGGTGGAGTGATTTGTCTGGTTAATTCCGTTAACGAACGAGACCTCGTCC
TGCTAAATAGGTGCGCGCATGCCAGCACTGCGTTCTTACCTTCTTAGAGGGACTATGCGC
GTCTAGCGTATGGAAGATTGAGGCAATAACAGGTCTGTGATGCCCTTAGATGTTCTGGGC
CGCACGCGCGCTACACTGATGCATTCAACGAGTTTCACTTGATAGTCCTGGGTCGGAAGG
CTCGGGTAATCTTTTGAAAGTGCATCGTGCTGGGGATAGATCATTGCAATTATTGATCTT
CAACGAGGAATTCCTTGTAGGCGCAGGTCATCAGCCTGCGCCGAATACGTCCCTGCCCTT
TGTACACACCGCCCGTCGCTCCTACCGATTGAATGGTCCGATGAAATGTCCGGATCGCGG
CTAGTGTGATGGACTCGTCCAGAGCGGTAGCCGTGAGAAGCTCATTAAATCTTATCATTT
AGAGGAAGGAGAAGTCGTAACAAGGTTTCCGTATTTTGTGGGTGGATACCGATTCAATAA
CATCTGCCGTTTTTCTCAATGTACTATATCCCTAGTCTGCTTGCCCAGTTCTTGGAGTCT
CGTGCGATGTGTGCTATTACAGGCACGCTGGGTCATGAAGACAGACAAGGTTAGGATCTT
CACGGAAGTACGCGCAATAGGAGCGCGTCTTGTTCAACGCACCTGACTCAAGAGTCATAT
GCGAGGGACGGACGCAGCGCTTGTCGCCGCACAGATGGGAAGCTTCCTCTCCAGGCATAG
GATGGCGCTGCATTTGCATCATCGCTGCTAGTAAGTGGCAATAATACTTGATTCCCCGCC
AGCGCACTTGTACGTAGCCGTTGGCGGCGGGAACGTAGTTGGTGATGTGACAGCCTCGAG
CGTTCACCTCTGTGTGTCCGCCAATCTGCACAAAGATGAACGCCATATCGGAGACAGGGG

GAGGTGCCATTGTGATGATGACGATGTTGTTGGGTGGTCCTTCCGTAGGAAAACGTACAC
GAAGCCTTTGCGGCCCGAAAGGGTGCACAGTGCGCGACTATAAACACTGATTCTGTGGTC
AACGCAAGCAGCGGTTCTGTGGAAGAACAGCTGCGACTTTCTCAAATTGCGGGAACACCC
TAAAGTCAACAAGCACCGCGGCTTGACGTGAAAAGCGTCAGGCGCAGCAGTCGGGGTAGC
GCCCTGGCGGATGGTAATAGCCTTGTTGAATGGCACAATGGGCAATCCGCAGCCAAGCCC
CTAAGGTCCCCCGCTGGGCTATGGGGAAGGTTCACAGACTGTAAGGGAGGGGGTTTCTTC
CATGAAAGAAGCTTAAGAGACAGTCGGTCCCCTGCGAGAGCAGTGTCGCATGTGAGGAAG
GCACCCATGAAAAGTGGGTGCTGGAGAGCCATGTGAGCCCTCTATGAGGGCGGGAAGTTT
TCCGATACAATGAGAAACAAACACGTTCCGTAGGTGAACCTGCGGAAGGATCATTCACAA
ACTATTGACACAAACACACGCGAACCAAATCGTCCGCACAGCAGGTGGCAATGAAAGAGA
GACTCTGCATGTCGCCTTTCGGGGTAGCAAGCAGCACTCTTCTGCCATCGCCTCTGTGCA
GGGCGTAAATTCCCATTGAGAGGATGTGAGGGCACCACAGGAAGCTTTTCCACAGGAAGT
CGCCATCCTTCTCCCTCCACGGCGGCGCTTCTGTGCTTGGCAGTTTTTTTTGCCTTCTAG
GGAGGATGCCGCCAATGGAGCCCCTTATATATATACATCATCATATGCCCCTTTTTTTCT
TAACCGCTTGCCACAGCTTCTTCTATGAGGAGCTTGTGGGAAGACTGTCTCCATACAATA
ACAAAGATACAACTCTTAGCGGTGGATATCTTGGTTCTCGCAACGATGAAGAACGCAGCC
AACTGCGATAACTAATGTGAATTGCAGGACTTCGTGAATCATTAAGTCTTTGAACGCAAG
TTGCGCTCATCTCCATGAGGAGGTGAGCATGCCTGTTTGAGTGTCGATGGGAAAAACAAT
ACATAACAACAATCGGCTCTCTTATCTTCCTGCCTTCTTGGGGGCAAAAGGTAGTGCATG
CCGGAGATGGGCTTTGTCGCCCCTTTCTCTTGAAAGAGGCGTCGTCCGAAACACGTATCG
TTGCCCGCTGGAGGTCATAGAGCATGAGTTTAAAGCACCACATCTCTCAAGTCAGAGTGA
GCGTCTTGAGTAGGTCCGCATTGATTCAGTTTCAACCCAGGTGGCGTTGGTTACTTGCCA
TAAGCCTCTCTGGGGTTTCTTTGCTTGGACCTCTCTTGTCAGTTTCTCTGACGTAGAGAC
AGGTGCCGTCTCATGGACGTTGCCCTCTGGAACGTGCTAAAAAAAAAAAAAAGAAAAAAC
CGTACCACATGTCAAAAGAGGCAGATCGTGCTTGACGGCGGGAGGTGTTGCATTTCTTGC
TTGTCCTGAGCGTCTGCCAGTCTCTGACGAGTGTGTTCCTTCCATAAACCCGATCTCAAA
TCAGGTAGGACCACCCGCTGAATTTAAGCATATAATTAAGCGGAGGAAAAGAAACTAACA
AGGATTCCCTCAGTAGCGGCGAGCGAAGCGGGAAGAGCTCAACATGAAAACCGTTGGGGG
TTCGCGCTTTCCTTCGGGTTGTATAGTAAAGTGGTGTCGCCTGCGTCGCGCCGGGTATAA
ATCCCGTCTGAAATGGGGTGTCGTAGAGGGTTATATCCCCGTCTGTGACCCGGTGTGCGA
TGCGTTACGGTGACCTTACGATGAGTCGGGTTGCTTGGGCATGCAGCCCTAAGTGGGTGG
TAAACTTCATCTAAAGCTAAATACTAGCGAGAGACCGATAGCGTACAAGTACCGTAAGGG
AAAGATGCAAAGAACTTTGAAAAGAGAGTTAAACAGGGCTTGAAACCGTTAAGAGTGAAG
CGAAGGAGGTCAGTGATCATGGGGCCAGATCCCCAAGTGCCTGTCATAGGGCACCGTAGC
GTTCTGGTTTCCCGGGGCAGCGCCGGCCGGGGGCGGGGGTTAACAGGGAGGTGGTTGAGG
GATTCGTTTTCCGCCTGTACTCCTTTATAGCTTGCACCCCGGCCGAGGGATCCGCGACTT

GTCGTGTGAGGGCGCTGTCAAAATGGCCTTCTCCACCCCGTCTTGAAACACGGACCAAGG
AGTCTAACATACTCGCGAGTGTTTGGGTGGTTAAAACCCGAACGCACAATGAAAGTAAAC
GGTGGCTCTGCGCCGCTGCTTGGTGGGAACCCTGCTTTTATAATGGGGGGCACTATCACC
CGACCCCGATCCCTTTGGGGTGGTTGGTTTTGAGGTAGAGCGAGTATGATGGGACCCGAA
AGATGGTGAACTATTCCTGAGTAGGGCGAAGCCAGAGGAAACTCTGGTGGAGGCTCGTAG
CGATGCTGACGTGCAAATCGCTCGTCGAACTTGGGTATAGGGGCGAAAGACTAATCGAAC
CATCTAGTAGCTGGTTCCCTCCGAAGTTTCCCTCAGGATAGCGGGAGCAAATGCAAGTCA
GTATCATCCGGTAAAGCGAATGATTAGAGGCCTCGGGGGAGTAACTCCTTCGACCTATTC
TCAAACTTTAAATAGGTGAGGAGCCGGCAACGTTGCTTAAAACAATGAACGCCGTGCGGA
CAATGATTGCTCTCAGTGGGCCGATTTTGGTAAGCAGAACTGGCGATGCGGGATGAACCG
AAAGCGACGCTAAGGTGCCAAACTCCCAGCTCATTCAGACACCAGAAAAGGTGTTGATTC
ATACAGACAGCAGGACGGTGGCCATGGAAGTCGGAATCCGCTAAGGAGTGTGTAACAACT
CACCTGCCGAATGGATCAGCCCTGAAAATGGATGGCGCTTAAGCGGGGGACCGAGGCGTC
GCCGTTGGGGTAAATACGAAGCCCCAATGAGTAGGAGGGCGTGGGGGTTTTGAAACACGG
TACACTCGCATGGAAGCCTTGGGCGTAAGCTCGGGTGGAGCGACCCCTAGTGTAGATCTT
GGTGGTAGTAGCAAATATTCAAGTGAGAACCTTGAAGACTGAAGTGGGGAAGGGTTCCAC
GCGAACAGTGCTTGGACGTGGGTTAGTCGGTCCTAAGAGATGGGGAAACTCCGTATTCAA
GTGCGCTTTTGTAGCGCAGTCCATCGAAAGGGAAGCGGGTTAACATTCCCGCACCAGGAT
GTGGAAACTGTGGTTTACCGCAAACTGCGGCAACGCAACTGAACCTGGGGACGCCCGTAG
GAGCTCCGAGCAGGGATCTCTTTCCTGTTTAACGCGCCCAACCCCCTGGAATCGGTTTGT
CCGGAGATAGGGGGCGAAAGGCCGGAAAAGCACCGCACGTCTTGTGGTGTTCGGAACGCT
TGTAATGGCCCTTGAAAACCCAGGGGAAAGCAGCCGTCTTACACTTGTGTGAGACGAGGA
ATTCTCACACCTGGCCGTACCCATCACCGCATCAGGTCCCCAAGGTGAACAGCCTCTAGT
CAATAGAATAATGTAGATAAGGGAAGTCGGCAAAATAGATCCGTAACTTTGGGAAAAGGA
TTGGCTCTAATGTTCGGGCTTGTCGGGACGCGCGGGCGTAGATGCCAATGAGCGGCAGAC
GTCTAGCCTCTTTGGAGGCTGGATTAACGCTGCTTGTGGGAAGAAACCGTGTCGATGGAG
GCAGGAACGCTTATCGATCTTGTCTCTTCCCGCTTGCGATCAACGAACGGATTAGAACTG
GTACGGACAAGGGGAATCCGACTGTTTAATTAAAACAAAGCATTGCGACGGCCGGAAACG
GTGTTGGCGCAATGTGATTTCTGCCCAGTGCTCTGAATGTCAAAGTGAAGAAATTCAACC
AAGCGCGGGTAAACGGCGGGAGTAACTATGACTCTTAAACAAACCATACTACCTACGGGA
GTCAAAAGTGGGGGCGGCGTCTGGTGTAGAAATACACGTACGACACGCTTTTTTTCCTGC
TAGTCGATCTGACCTTTGTTCTTTGGACTGTGGGCGGTCAGCGGCAACATAACCTGGTAC
GGGGAAGGCCTCGACGTTGAGTGCTGCATCTTGGAAGTATCTTGACACAATAAGAGAAAA
GAGATTGCGCTTAATAGGGGGTTGATCTCGTGGCGAGCCGACGCCGGTGTGAACTGGTGG
CAATGGCCGTCGTAGAGCACGCAAAGGTATGGGCTGACTTTTTCTCCTCAAAAGTTGGCT
TAAGGGACGTGCCATCCCACACGATAATAAAGTGTGCAGTCTGTAAGAGCTCCCACAGGG

CAAAGACAGACTGGGATGCAGACGGAGACGGCAAAAACAACAACATCCGTTCTGCTTCTG
GGCGCAAATGCTTAAGGTAGCCAAATGCCTCGTCATCTAAACTAATTGGATGAGATAGTC
TGAGTGAAAAGCTTATGCAGGCTAGTACCGGAAAAAAAAAAAAAGAGGGGCTCTGCCACT
CTGACGGTGCGACACTCCTTGTACAGGGAAACCCTCCGCATGGAAAGGCACGTGGTACCA
CCCTCGCGGGGAAACTCGCGAGCGGGATCCCGGGTAATGACCGGAGCCGATGGTAAAAAC
TCACGTGACCGATGGGGTGACCCTGTGGCGAGACGCTAAGTTGTGGGACATCACCCTCTC
ACAATACGCATCCGTCGCAGAGACTGAACGGGAGTGGGCTCCGTTGACGGCCGGCCGGCC
AGCCAGCCGTCTTCCAGGGGCTTAAGATACAGTCCGGGCCTCAGGGATGGAAAAACCTGG
GGAACTACCAGAATTAGTGACGCGCATGAATGGATTAACGAGATTCCCACTGTCCCTATC
TACTATCTAGCGAAACCACAGCCAAGGGAACGGGCTTGGAAGAATCAGCGGGGAAAGAAG
ACCCTGTTGAGCTTGACTCTAGTCCGACTTTGTGAGATGAGTTGGGAGGTGTAGTCTAAG
AGGGAGGGGCAACTCGACCGTGAAATACCTCTACTCCCAAGCTTGTTTTACTTACTCCGG
CGAGCAGGGATGGGTGGTAACACCCTTTTTCTAGGATTAACACCTCTTCGGGGGTGGACC
TGCGCGGAGGACATTGTCAGGTGGGGAGTTTGGCTGGGGCGGCACATCTGTTAAAAGATA
ACGCAGGTGTCCTAAGATGAGCTCAGTGAGAACAGAAATCTCACGTAGAACAAAAGGGTA
AAAGCTCATTTGATTTTGATTTTCAGTACGAATACAAACTGTGAAAGCATGGCCTATCGA
TCCTTTTGACCTTCAAGGTTTGAAGCAAGAGGTGTCAGAAAAGTTACCACAGGGATAACT
GGCTTGTGGCAGCCAAGCGTTCATAGCGACGTTGCTTTTTGATCCTTCGATGTCGGCTCT
TCCTATCATTGCGAAGCAGAATTCGCCAAGTGTTGGATTGTTCACCCACTAATAGGGAAC
GTGAGCTGGGTAGCCACTTTATGCCCAACAGCTGGATAGCTGGGCTTCTGTCTTAGGCCG
GGACTTCTCACGACCCGTGCCTTGGTCACCCAAAAAAGGGCGACCGCAGAGCTCAGGATG
GAGCTTCTGGCTTAACTAGTAGAGGACAGATAGAGATCGTCAAATTGCGGGAAACCCCTT
AGAGCCTCGAGCTACCAAGTGCGTGTCGAAAGGCATAGACGCAGCCAGCGATGGCTGTCG
CATGGCTTGATTCCATCAAGATAGGTAAAAATGCTCGAGGATTGGGCAATCCGCAGCCAA
GCCCCTAAGGTCCCCCCGCGGGCTATGGGGAAGGTTCAACGACTGTAAGGCGTACCGTGC
AAGCGGAAGATACAGTCTAGCCCCGCTGGAAAGCAGCGCCAAAGTAAGTAGGTTGCAGTG
GGTCTGTCCTC
>SRR9587921_SRR9587937_SRR9587947
GCATGTTTTGGTATGTGTTTCGACAGTCATGCATGTCATAGCCGTAGGATGGGGAGTGAG
CTTCATAGCCCGACCCCTTCTCAGGAGCATGTTTTCATAGCATATTGAGCATACTGGGGT
GGTCTCCCATGTAGCTCGCTCTCTCGGTGCAGGACCTCCCTTTTGCGGGCCACTTCGATG
GCCTCCATTATGGAGAGCATGTTTTGGCATATGAAGCGTTGGTGGTGAGGCTCGTAGCCT
GTATATACATACCAACGCAGTCGTCATATGTCTCTCTGGCGGGGGTCGGTCGTATGACCT
TGTCCCTACCTGGAGCATGTATTTTGTATGATATACTGGACGGCAGCTGCGCTGTTCAGT
CGTCAGCCGTACCTCATGTGCGCGTTTCCAGGTCCGAACGTTTGACATGCAAGGGCATCA
TACCAGAAGCCTGTTTTCTCGTATGAGCATGTAGGTAGGCAGGCTCTTCAGTCGTCTACC

CATAAATGCACTTCGAAGGAACTTTTTGGTATTGAATGGCTTTGTGCCCAGACGTTCTTG
TGAGCTTGTTCTCGCTCTGCGATACGCTGGAAGCTGTGGTATTACCTACAGCAGACGCAT
TCCTAGGGCGGACGGCCTCTCTGCAGATGGGCACATCTGTAGAAAGAGGCTCCCCTTGAT
GACGCAATGCAATGGCTGGGTAAACGGAGTGCTATACAGAAGGGTCAAACCTCTGTGGAA
GTACACTCATACCAGCATTGTTTGTTGAGAGGGGTCGGCTCTACGAGTGAGGACCTATAC
TGCCATCTCATTCAGATGCGTAGACGGTCTTTACTCAGACTGCCTTTCTGGCTGATTGTC
CTCGGTCCCTCGTGCGCACGCGCCGCTGGCTGAATGGGCTTGTCCTTATGGAGTGCCGCG
TTCTTCAAAGAATCCGTCTTCATACAGCGACCTCATTTACATATTGCCGGCTGAATGGGC
TTGTCCTTATGGAGCACTGCATTCTTCTAGGAATTGCTATCTATACAGCGACCTCATTTA
CATACAAGGCACCTTTTTCTACATAGACTTGCCATTTGGCTGATTGTTCTGACAGTCGCT
GCCTCTTCGGAGTTCTGTGTGGACGAGTGGCCTCCTGAGTGGGCCGGTCCTTACGGAGTG
CCCACGTCTGTTATGACTGGGTTCTCCATACAGCGACCTCACTCAGAATTGTATGGCACC
GTGCCGATTGAATGGACAAGCAGTCCTTAGGGAATCGCATGAAAGCATGACGGCACATGG
TGTATGGACCCGTTGGGCCTTGCCTGTGCTATCTTCTCGAGTCGACCGGTCCTCGTCTCC
TCCTTTCGCGACGGGAAGCTCTTGAGTTTACCGTGGGTCAGGACGCTGTGGTATCTGTTT
GTCTGTAGTGTGGGCTCTCTATCAGAACAGCGTATCAGCACACGTCGTCTGGGTCTATGG
TATTTTTGCTTAACGGCCCTGGGTTTGGTAGTATGGTTCCATGTCGGTCCTCTCACCTCC
CAACCGTCATGTCTCTGATATCATCTGCACAGTACGTCTTGTCAGAATATCATTCTGTAT
GGCAGTCGATAGACTAGGCCCTATGCCGCAGCTGTCTCTGACCACTATGCGAGCGCATAC
CTACCTCAATGTGTTCATAGCTGTTATGGATCTTGGGTGTGGTGTGCCAGAACGATACGC
CACCAGATTGTGGAATGGTGCTTGTAGTATTAGAAGAGGACATGTATGAAGAGCACGTTC
TTTTCTCTTATGATGGTGCCGCGGGTGCGGGCCGTTCTCATTCTGCTTGTCGGGTGCTCA
AGGTGCATGTAGGATTGCTGAACCCACTGAATGGTCATGAGAATGGTCATATGGTGAGGT
TCACTTCTCCATCGCTTTGTGTGTGTTGGATGTTTGGGTGGCTTCGGCCGCATGGAAAAT
CTCGGTCTGTAACTCATCAGGGAGGGTTTCCATGTGAGCAATCTCTCCAGTCGAACACAC
ACCCATGGGCGTCCCGCTAGGCAAAATGGAACTCTCGATCCGACACAGCCTGTGGAGTGT
TTGTTCCGTGTGGCGTGAAACGGCTTATACTGGATGTCTTCCCTTTGGACACCAGTCTGT
TGGTATGTACGCATACCATCTAGAGGGAGCAAGAAAAAGGCATGCAAGTGCCACAAAGGA
AGGGAGGCCTTCCTGCTACCGTGTGTCCTCTTGACGATCACGGGGGAGTATATAGAGACA
TTTGGACGAGATCCGTTTTTCACGAGGCGGTGCCCTCGCGCAAAGGTGTTGCTCTGCACG
CATCTTTATGATGATAGTTGCTCGCTCCTATGCGTCCACTCGTGAAAATACCTGCTCTCT
CTGTCTGTTTCCATGACACCCCCATGCGTTGTCATAGAGCATGCTCAGGTACAGAAAGTG
CTGTGACTTTCTTGGCCTGCTGTCACTGAGTCAGCCTGATACTTTGGAGGAATCTTTTCT
GGGCACTGCCCAGCTGAGAGTTCTTATCAGGCTCTGGAGTGTGCATATGCCTATATTCAT
TATAGTGCATGTGGCTACGCAGAAAAAGCATCGACGTGGAATGGAGAAGAACACCATTCT
GAATGGAACGGTTCTTACTCTACTTCTGTTGCTTGACGCTGTATGGGAATCGCTTTGCTC

TGGGGGTACACCTTCCATTCCATGACAAAGCATTTGATCTATGCCCCGAGGGTCATTCTC
TATGAAGAGCCCGAAACTTGTCTGAGTGAGAGGCGAAGGTAGTCCTTCATTTATGCGGGG
TCCGCCCTCACACTGTAGTCGGCACAGTCCCTTGTTTATGGACTGCGCAGCTCAATCTTC
GACTGGAGAGGAAAAACAAAACACAAGTATGACACGCACATACGTGTTTTCTAGGACTGT
GACCTGTCGGCTGCCTCGCGGTGGTTGGCATGATGCCCCAAGTGGGGGGGGGGAGAAACA
ATCAGTATGATTGATAAAAACCAGTGGGGAAAAAGGCGCGCTGTCGCACACCCTTGGTCA
CGGCCTCAAGTCGACGGCCCTCACTGGCCCACCCTGCGACTCCTGTTCGCTTTTGACAAG
ACACACCTGCAATACGGAGGAAATGCCTGAGCAGCTGTGCCGCAGCAACTGCGGACGCGT
AGCGATATGCATCCCTATTGTTTGCGACTTGGTACAAAGAACGTGCGCCGGGAGACGGAT
TGCCCTTTGACTTTTCCACCCAGCCCGCGGCTCTATACCCTTGACGTACCTATTTACCTG
GTTGATCCTGCCAGTAGTCATATGCTTGTCTCAAAGACTAAGCCATGCATGTCTAAGTAT
AAACACTTTTATACTCGTGAAACTGCGAATGGCTCATTAAAACAGTTATAGTTCCTTTGG
GAACAACACTACTTGGATAACCGTAGTAATTCTAGAGCTAATACATGCCTCAACGCCCGA
CTCACGAAGGGTGGTATTTATTGGATAAAAAACCATCGTGCTCTGTCCAGAGCGCTTTGA
GATGATTCACAATAACTTGTCGGATCGCATGGCCTCGTGCTGGCGACTCCTCATTCAAAT
TTCTGCCCTATCAACTTTCGATGGTAGAGTATTGGTCTACCATGGTGTCGACGGGTGACG
GGGAATTAGGGTTCGATTCCGGAGAGGGAGCCTGAGAAACGGCTACCACATCCAAGGAAG
GCAGCAGGCGCGCAAATTACCCAATCCCGACTCGGGGAGGTAGTGACAAAAAATAACAAT
AGGGGGCCCTTTGGGTCTTCTAATTGGAATGAGAACAATTTAAATCCCTTATCGAGGATC
CATTGGAGGGCAAGTCTGGTGAACGTATGCATTAAAGCCCTCCGTAATAAAAAAAAACAG
CGCCAGATGTCTCTCTCTGGAAGCACACCATGCATGGGGTTTGTTCCTCTCTGCATGGCC
GTGTGAAAAAGGCGAGTAGGCAAACGATGGGAAACCCGGCTTGGAATAGTAACCAGCCAC
AGAGCCCCCATCATACACTTCTAAATTGCCGGGGAGTGCCCGCGAGGCCTTGACTACCGC
GGTCCGCCGAGAGGTGAGAGCGAGCAATCGTTCAGGACGCAGCAGTGTACGTAGTGGTAC
ATGGATGGTAAAAACGTCAAGGATAGGGTTGATCCGCAGGGAAGCGCGAGTGCCGCCATG
GTCGGTGGGCAAAGGCGAACCTTCAGAGACTCTAATGGAGTGGGCGCAAGCTTAAGGGAG
AGTCCAATCCACTGGGAAACCAGGTCCACAGCAGTAACGGGGTGATGCTCGCCCCAGAGG
ACTGTGGAAGGCAACCCCACCAAAAGGGTGTGTGTTGTCGGGTAGCAATGGCCAGCAGCC
GCGGTAATTCCAGCTCCAATAGCGTATATTAAAGTTGTTGCAGTTAAAACGCTCGTAGTC
GGACCTCGGAGCGCCCCCACGGGCCGGGCTTTTGCCTTGTTGCCAGTGGGAGTTTGTCTC
CTTTTGTCGGGACGCGGTGTGCAGGGCCTTTACTGTCTCTGCGCGTTTACCTGCGGGCCG
ACCGTTTACTGTGAAGAAAGTAAAGTGTTCAAAGCAGGCCAATGCCTTGAATATGTGAGC
ATGGAATAATAGAATAGGACTTGGGCTCTATTTTGTTGGTTTCCAGTGACCAAGTAATGA
TTAATAGGGACGGTTGGGGGCATTCGTATTTCATTGTCAGAGGTGAAATTCTTGGATTGA
TGGAAGACGTACAACTGCGAAAGCATCTGCCATGGATGTTTTCATTGATCAAGAACGAAA
GTTAGGGGATCGAAGACGATCAGATACCGTCGTAGTCTTAACCATAAACGATGCCGACTG

GGGATTGGCGGGGGCACTTTTATATGACTCTGTCAGCACCCTGAGGGAAACCAAAGTCTT
TGGGTTCTGGGGGGAGTATGGTCGCAAGGCTGAAACTTAAAGGAATTGACGGAAGGGCAC
CACCAGGAGTGGAGCCTGCGGCTTAATTTGACTCAACACGGGAAAACTTACCAGGTCCGG
ACAGAAGAATGATTGACAGACTGAAAGCTCTTTCTTGATTTTTTGGTTGGTGGTGCATGG
CCGTTCTTAGTTGGTGGAGTGATTTGTCTGGTTAATTCCGTTAACGAACGAGACCTCGTC
CTGCTAAATAGGTGCGCGCATGCCAGCACTGCGTTCTTACCTTCTTAGAGGGACTATGCG
CGTCTAGCGTATGGAAGATTGAGGCAATAACAGGTCTGTGATGCCCTTAGATGTTCTGGG
CCGCACGCGCGCTACACTGATGCATTCAACGAGTTTCACTTGATAGTCCTGGGTCGGAAG
GCTCGGGTAATCTTTTGAAAGTGCATCGTGCTGGGGATAGATCATTGCAATTATTGATCT
TCAACGAGGAATTCCTTGTAGGCGCAGGTCATCAGCCTGCGCCGAATACGTCCCTGCCCT
TTGTACACACCGCCCGTCGCTCCTACCGATTGAATGGTCCGATGAAATGTCCGGATCGCG
GCTAGTGTGATGGACTCGTCCAGAGCGGTAGCCGTGAGAAGCTCATTAAATCTTATCATT
TAGAGGAAGGAGAAGTCGTAACAAGGTTTCCGTATTTTGTGGATGGGCACTCGTCGTGGC
GTTCTTTGCGCCACGCGGATGTCCCCCTTCCGTAGGAAAACGTACACGAAGCCTTTGCGG
CCCGAAAGGGTGCACAGTGCGCGACTATAAACACTGATTCTGTGGTCAACGCAAGCAGCG
GTTCTGTGACAGAACAGCTGCGACTTTCTCAAATTGCGGGAACACCCTAAAGTCAACAAA
CACCGCGGCCGGACGTGAAAGCGTCAGGCGCAGCAGTCGGGGTAGCGCCCTGGCGGATGG
TAAGAGCTTTGTTGAATGGCACAATGGGCAATCCGCAGCCAAGCCCCTAAGGTCCTTCGC
GGCTATGGGGAAGGTTCACAGACTGTAAGGGAGGGGGTTTCTTCCATGAAGAAGCTTAAG
AGACAGTCGGTCCCCTGCGAGAGCAGTGTCTCATGTGAGGAAGGTGCCCATGAAAAGTGG
GCGCTGGAGAGCCATGGGAGCCCTCTATGAGGGTGGGGAGTTTTCCTAGATAATAATGAG
AAACAAACACGTTCCGTAGGTGAACCTGCGGAAGGATCATTCACAAACTATTGACACAAA
CACACGCGAACCAAATCGTCCGCACAGCAGGTGGCAATGAAAGAGAGACTCTGCATGTCG
CCTTTCGGGGTAGCAAGCAGCACTCTTCTGCCATCGCCTCTGTGCAGGGCGTAAATTCCC
ATTGAGAGGATGTGAGGGCACCACAGGAAGCTTTTCCACAGGAAGTCGCCATCCTTCTCC
CTCCACGGCGGCGCTTCTGTGCTTGGCAGTTTTTTTTGCCTTCTAGGGAGGATGCCGCCA
ATGGAGCCCCTTATATATATACATCATCATATGCCCCTTTTTTTCTTAACCGCTTGCCAC
AGCTTCTTCTATGAGGAGCTTGTGGGAAGACTGTCTCCATACAATAACAAAGATACAACT
CTTAGCGGTGGATATCTTGGTTCTCGCAACGATGAAGAACGCAGCCAACTGCGATAACTA
ATGTGAATTGCAGGACTTCGTGAATCATTAAGTCTTTGAACGCAAGTTGCGCTCATCTCC
ATGAGGAGGTGAGCATGCCTGTTTGAGTGTCGATGGGAAAAACAATACATAACAACAATC
GGCTCTCTTATCTTCCTGCCTTCTTGGGGGCAAAAGGTAGTGCATGCCGGAGATGGGCTT
TGTCGCCCCTTTCTCTTGAAAGAGGCGTCGTCCGAAACACGTATCGTTGCCCGCTGGAGG
TCATAGAGCATGAGTTTAAAGCACCACATCTCTCAAGTCAGAGTGAGCGTCTTGAGTAGG
TCCGCATTGATTCAGTTTCAACCCAGGTGGCGTTGGTTACTTGCCATAAGCCTCTCTGGG
GTTTCTTTGCTTGGACCTCTCTTGTCAGTTTCTCTGACGTAGAGACAGGTGCCGTCTCAT

GGACGTTGCCCTCTGGAACGTGCTAAAAAAAAAAAAAAGAAAAAACCGTACCACATGTCAA
AAGAGGCAGATCGTGCTTGACGGCGGGAGGTGTTGCATTTCTTGCTTGTCCTGAGCGTCT
GCCAGTCTCTGACGAGTGTGTTCCTTCCATAAACCCGATCTCAAATCAGGTAGGACCACC
CGCTGAATTTAAGCATATAATTAAGCGGAGGAAAAGAAACTAACAAGGATTCCCTCAGTA
GCGGCGAGCGAAGCGGGAAGAGCTCAACATGAAAACCGTTGGGGGTTCGCGCTTTCCTTC
GGGTTGTATAGTAAAGTGGTGTCGCCTGCGTCGCGCCGGGTATAAATCCCGTCTGAAATG
GGGTGTCGTAGAGGGTTATATCCCCGTCTGTGACCCGGTGTGCGATGCGTTACGGTGACC
TTACGATGAGTCGGGTTGCTTGGGCATGCAGCCCTAAGTGGGTGGTAAACTTCATCTAAA
GCTAAATACTAGCGAGAGACCGATAGCGTACAAGTACCGTAAGGGAAAGATGCAAAGAAC
TTTGAAAAGAGAGTTAAACAGGGCTTGAAACCGTTAAGAGTGAAGCGAAGGAGGTCAGTG
ATCATGGGGCCAGATCCCCAAGTGCCTGTCATAGGGCACCGTAGCGTTCTGGTTTCCCGG
GGCAGCGCCGGCCGGGGGCGGGGGTTAACAGGGAGGTGGTTGAGGGATTCGTTTTCCGCC
TGTACTCCTTTATAGCTTGCACCCCGGCCGAGGGATCCGCGACTTGTCGTGTGAGGGCGC
TGTCAAAATGGCCTTCTCCACCCCGTCTTGAAACACGGACCAAGGAGTCTAACATACTCG
CGAGTGTTTGGGTGGTTAAAACCCGAACGCACAATGAAAGTAAACGGTGGCTCTGCGCCG
CTGCTTGGTGGGAACCCTGCTTTTATAATGGGGGGCACTATCACCCGACCCCGATCCCTT
TGGGGTGGTTGGTTTTGAGGTAGAGCGAGTATGATGGGACCCGAAAGATGGTGAACTATT
CCTGAGTAGGGCGAAGCCAGAGGAAACTCTGGTGGAGGCTCGTAGCGATGCTGACGTGCA
AATCGCTCGTCGAACTTGGGTATAGGGGCGAAAGACTAATCGAACCATCTAGTAGCTGGT
TCCCTCCGAAGTTTCCCTCAGGATAGCGGGAGCAAATGCAAGTCAGTATCATCCGGTAAA
GCGAATGATTAGAGGCCTCGGGGGAGTAACTCCTTCGACCTATTCTCAAACTTTAAATAG
GTGAGGAGCCGGCAACGTTGCTTAAAACAATGAACGCCGTGCGGACAATGATTGCTCTCA
GTGGGCCGATTTTGGTAAGCAGAACTGGCGATGCGGGATGAACCGAAAGCGACGCTAAGG
TGCCAAACTCCCAGCTCATTCAGACACCAGAAAAGGTGTTGATTCATACAGACAGCAGGA
CGGTGGCCATGGAAGTCGGAATCCGCTAAGGAGTGTGTAACAACTCACCTGCCGAATGGA
TCAGCCCTGAAAATGGATGGCGCTTAAGCGGGGGACCGAGGCGTCGCCGTTGGGGTAAAT
ACGAAGCCCCAATGAGTAGGAGGGCGTGGGGGTTTTGAAACACGGTACACTCGCATGGAA
GCCTTGGGCGTAAGCTCGGGTGGAGCGACCCCTAGTGTAGATCTTGGTGGTAGTAGCAAA
TATTCAAGTGAGAACCTTGAAGACTGAAGTGGGGAAGGGTTCCACGCGAACAGTGCTTGG
ACGTGGGTTAGTCGGTCCTAAGAGATGGGGAAACTCCGTATTCAAGTGCGCTTTTGTAGC
GCAGTCCATCGAAAGGGAAGCGGGTTAACATTCCCGCACCAGGATGTGGAAACTGTGGTT
TACCGCAAACTGCGGCAACGCAACTGAACCTGGGGACGCCCGTAGGAGCTCCGAGCAGGG
ATCTCTTTCCTGTTTAACGCGCCCAACCCCCTGGAATCGGTTTGTCCGGAGATAGGGGGC
GAAAGGCCGGAAAAGCACCGCACGTCTTGTGGTGTTCGGAACGCTTGTAATGGCCCTTGA
AAACCCAGGGGAAAGCAGCCGTCTTACACTTGTGTGAGACGAGGAATTCTCACACCTGGC
CGTACCCATCACCGCATCAGGTCCCCAAGGTGAACAGCCTCTAGTCAATAGAATAATGTA

GATAAGGGAAGTCGGCAAAATAGATCCGTAACTTTGGGAAAAGGATTGGCTCTAATGTTC
GGGCTTGTCGGGACGCGCGGGCGTAGATGCCAATGAGCGGCAGACGTCTAGCCTCTTTGG
AGGCTGGATTAACGCTGCTTGTGGGAAGAAACCGTGTCGATGGAGGCAGGAACGCTTATC
GATCTTGTCTCTTCCCGCTTGCGATCAACGAACGGATTAGAACTGGTACGGACAAGGGGA
ATCCGACTGTTTAATTAAAACAAAGCATTGCGACGGCCGGAAACGGTGTTGGCGCAATGT
GATTTCTGCCCAGTGCTCTGAATGTCAAAGTGAAGAAATTCAACCAAGCGCGGGTAAACG
GCGGGAGTAACTATGACTCTCTTAAGGTAGCCAAATGCCTCGTCATCTAATTAGTGACGC
GCATGAATGGATTAACGAGATTCCCACTGTCCCTATCTACTATCTAGCGAAACCACAGCC
AAGGGAACGGGCTTGGAAGAATCAGCGGGGAAAGAAGACCCTGTTGAGCTTGACTCTAGT
CCGACTTTGTGAGATGAGTTGGGAGGTGTAGTCTAAGAGGGAGGGGCAACTCGACCGTGA
AATACCTCTACTCCCAAGCTTGTTTTACTTACTCCGGCGAGCAGGGATGGGTGGTAACAC
CCTTTTTCTAGGATTAACACCTCTTCGGGGGTGGACCTGCGCGGAGGACATTGTCAGGTG
GGGAGTTTGGCTGGGGCGGCACATCTGTTAAAAGATAACGCAGGTGTCCTAAGATGAGCT
CAGTGAGAACAGAAATCTCACGTAGAACAAAAGGGTAAAAGCTCATTTGATTTTGATTTT
CAGTACGAATACAAACTGTGAAAGCATGGCCTATCGATCCTTTTGACCTTCAAGGTTTGA
AGCAAGAGGTGTCAGAAAAGTTACCACAGGGATAACTGGCTTGTGGCAGCCAAGCGTTCA
TAGCGACGTTGCTTTTTGATCCTTCGATGTCGGCTCTTCCTATCATTGCGAAGCAGAATT
CGCCAAGTGTTGGATTGTTCACCCACTAATAGGGAACGTGAGCTGGGTAGCCACTTTATG
CCCAACAGCTGGATAGCTGGGCTTCTGTCTTAGGCCGGGACTTCTCACGACCCGTGCCTT
GGTCACCCAAAAAAGGGCGACCGCAGAGCTCAGGATGGAGCTTCTGGCTTAACTAGTAGA
GGACAGATAGAGATCGTCAAATTGCGGGAAACCCCTTAGAGCCTCGAGCTACCAAGTGCG
TGTCGAAAGGCATAGACGCAGCCAGCGATGGCTGTCGCATGGCTTGATTCCATCAAGATA
GGTAAAAATGCTCGAGGATTGGGCAATCCGCAGCCAAGCCCCTAAGGTCCCCCCGCGGGC
TATGGGGAAGGTTCAACGACTGTAAGGCGTACCGTGCAAGCGGAAGATACAGTCTAGCCC
CGCTGGAAAGCAGCGCCAAAGTAAGTAGGTTGCAGTGGGTCTGTCCTC
>SRR9587922
GCATGTTTTGGTATGTGTTTCGACAGTCATGCATGTCATAGCCGTAGGATGGGGAGTGAG
CTTCATAGCCCGACCCCTTCTCAGGAGCATGTTTTCATAGCATATTGAGCATACTGGGGT
GGTCTCCCATGTAGCTCGCTCTCTCGGTGCAGGACCTCCCTTTTGCGGGCCACTTCGATG
GCCTCCATTATGGAGAGCATGTTTTGGCATATGAAGCGTTGGTGGTGAGGCTCGTAGCCT
GTATATACATACCAACGCAGTCGTCATATGTCTCTCTGGCGGGGGTCGGTCGTATGACCT
TGTCCCTACCTGGAGCATGTATTTTGTATGATATACTGGACGGCAGCTGCGCTGTTCAGT
CGTCGGCCGTACCTCATGTGCGCGTTTCCAGGTCCGAACGTTTGACATGCAAGGGCATCA
TACCAGAAGCCTGTTTTCTTGTATGAGCATGTAGGTAGGCAGGCTCTTCAGTCGTCTACC
CATAAATGCACTTCGAAGGAACTTTTTGGTATTGAATGGCTTTGTGCCCAGACGTTCTTG
TGAGCTTGTTCTCGCTCTGCGATACGCTGGAAGCTGTGGTATTACCTACAGCAGACGCAT

TCCTAGGGCGGACGGCCTCTCTGCAGATGGGCACATCTGTAGAAAGAGGCTCCCCTTGAT
GACGCAATGCAATGGCTGGGTAAACGGAGTGCTATACAGAAGGGTCAAACCTCTGTGGAA
GTACACTCATACCAGCATTGTTTGTTGAGAGGGGTCGGCTCTACGAGTGAGGCCCTATAC
TGCCATCTCATTCAGATGCGTAGACGGTCTTTACTCAGACTGCCTTTCTGGCTGATTGTC
CTCGGTCCCTCGTGCGCACGCGCCGCTGGCTGAATGGGCTTGTCCTTATGGAGTGCCGCG
TTCTTCAAAGAATCCGTCTTCATACAGCGACCTCATTTACATATTGCCGGCTGAATGGGC
TTGTCCTTATGGAGCACTGCATTCTTCTAGGAATTGCTATCTATACAGCGACCTCATTTA
CATACAAGGCACCTTTTTCTACATAGACTTGCCATTTGGCTGATTGTTCTGACAGTCGCT
GCCTCTTCGGAGTTCTGTGTGGACGAGTGGCCTCCTGAGTGGGCCGGTCCTTACGGAGTG
CCCAAGTCTGTTATGACTGGGTTCTCCATACAGCGACCTCACTCAGAATTGTATGGCACC
GTGCCGATTGAATGGACAAGCAGTCCTTAGGGAATCGCATGAAAGCATGACGGCAACATG
GTGTATGGACCCGTTGGGCCTTGCCTGTGCTATCTTCTCGAGTCGACCGGTCCTCGTCTC
CTCCTTTCGCGACGGGAAGCTCTTGAGTTTACCGTGGGTCAGGACGCTGTGGTATCTGTT
TGTCTGTAGTGTGGGCTCTCTATCAGAACAGCGTATCAGCACACGTCGTCTGGGTCTATG
GTATTTTTGCTTAACGGCCCTGGGTTTGGTAGTATGGTTCCATGTCGGTCCTCTCACCTC
CCAACCGTCATGTCTCTGATATCATCTGCACAGTACGTCTTGTCAGAATATCATTCTGTA
TGGCAGTCGATAGACTAGGCCCTATGCCGCAGCTGTCTCTGACCACTATGCGAGCGCATA
CCTACCTCAATGTGTTCATAGCTGTTATGGATCTTGGGTGTGGTGTGCCAGAACGATACG
CCACCAGATTGTGGAATGGTGCTTGTAGTATTAGAAGAGGACATGTATGAAGAGCACGTT
CTTTTCTCTTATGATGGTGCCGCGGGTGCGGGCCGTTCTCATTCTGCTTGTCGGGTGCTC
AAGGTGCATGTAGGATTGCTGAACCCACTGAATGGTCATGAGAATGGTCATATGGTGAGG
TTCACTTCTCCATCGCTTTGTGTGTGTTGGATGTTTGGGTGGCTTCGGCCGCATGGAAAA
TCTCGGTCTGTAACTCATCAGGGAGGGTTTCCATGTGAGCAATCTCTCCAGTCGAACACA
CACCCATGGGCGTCCCGCTAGGCAAAATGGAACTCTCGATCCGACACAGCCTGTGGAGTG
TTTGTTCCGTGTGGCGTGAAACGGCTTATACTGGATGTCTTCCCTTTGGACACCAGTCTG
TTGGTATGTACGCATACCATCTAGAGGGAGCAAGAAAAAGGCATGCAAGTGCCACAAAGG
AAGGGAGGCCTTCCTGCTACCGTGTGTCCTCTTGACGATCACGGGGGAGTATATAGAGAC
ATTTGGACGAGATCCGTTTTTCACGAGGCGGTGCCCTCGCGCAAAGGTGTTGCTCTGCAC
GCATCTTTATGATGATAGTTGCTCGCTCCTATGCGTCCACTCGTGAAAATACCTGCTCTC
TCTGTCTGTTTCCATGACACCCCCATGCGTTGTCATAGAGCATGCTCAGGTACAGAAAGT
GCTGTGACTTTCTTGGCCTGCTGTCACTGAGTCAGCCTGATACTTTGGAGGAATCTTTTC
TGGGCACTGCCCAGCTGAGAGTTCTTATCAGGCTCTGGAGTGTGCATATGGCTATAATCA
TAATAGTGCATGTGGCCACGCAGAAAAAGCATCGACGTGGAATGGAGAAGAACACCATTC
TGAATGGAACGGTTCTTACTCTGCTTCCGTTGCTTGACGCTGTATGGGAATCGCTTTGCT
CTGGGGGTACACCTTCCATTCCATGACAAAGCATTTGATCTATGCCCCGAGGGTCATTCT
CTATGAAGAGCCCGAAACTTGTCTGAGTGAGAGGCGAAGGTAGTCCTTCATTTATGCGGG

GTCCGCCCTCACACTGTAGTCGGCACAGTCCCTTGTTTATGGACTGCGCAGCTCAATCTT

CGACTGGAGAGGAAAAACAAAACACAAGTATGACACGCACATACGTGTTTTCTAGGACTG

TGACCTGTCGGCTGCCTCGCGGTGGTTGGCATGATGCCCCAAGTGGGGGGGGGAGAAACA

ATCAGTATGATTGATAAAAACCAGTGGGGAAAAAGGCGCGCTGTCGCACACCCTTGGTCA

CGGCCTCAAGTCGACGGCCCTCACTGGCCCACCCTGCGACTCCTGTTCGCTTTTGACAAG

ACACACCTGCAATACGGAGGAAATGCCTGAGCAGCTGTGCCGCAGCAACTGCGGACGCGT

AGCGATATGCATCCCTGTTGTTTGCGACTTGGTACAAAGAACGTGCGCCGGGAGACGGAT

TGCCCTTTGACTTTTCCACCCAGCCCGCGGCTCTATACCCTTGACGTACCTATTTACCTG

GTTGATCCTGCCAGTAGTCATATGCTTGTCTCAAAGACTAAGCCATGCATGTCTAAGTAT

AAACACTTTTATACTCGTGAAACTGCGAATGGCTCATTAAAACAGTTATAGTTCCTTTGG

GAACAACACTACTTGGATAACCGTAGTAATTCTAGAGCTAATACATGCCTCAACGCCCGA

CTCACGAAGGGTGGTATTTATTGGATAAAAAACCATCGTGCTCTGTCCAGAGCGCTTTGA

GATGATTCACAATAACTTGTCGGATCGCATGGCCTCGTGCTGGCGACTCCTCATTCAAAT

TTCTGCCCTATCAACTTTCGATGGTAGAGTATTGGTCTACCATGGTGTCGACGGGTGACG

GGGAATTAGGGTTCGATTCCGGAGAGGGAGCCTGAGAAACGGCTACCACATCCAAGGAAG

GCAGCAGGCGCGCAAATTACCCAATCCCGACTCGGGGAGGTAGTGACAAAAAATAACAAT

AGGGGGCCCTTTGGGTCTTCTAATTGGAATGAGAACAATTTAAATCCCTTATCGAGGATC

CATTGGAGGGCAAGTCTGGTGCCAGCAGCCGCGGTAATTCCAGCTCCAATAGCGTATATT

AAAGTTGTTGCAGTTAAAACGCTCGTAGTCGGACCTCGGAGCGCCCCCACGGGCCGGGCT

TTTGCCTTGTTGCCAGTGGGAGTTTGTCTCCTTTTGTCGGGACGCGGTGTGCAGGGCCTT

TACTGTCTCTGCGCGTTTACCTGCGGGCCGACCGTTTACTGTGAAGAAAGTAAAGTGTTC

AAAGCAGGCCAATGCCTTGAATATGTGAGCATGGAATAATAGAATAGGACTTGGGCTCTA

TTTTGTTGGTTTTCAGTGACCAAGTAATGATTAATAGGGACGGTTGGGGGCATTCGTATT

TCATTGTCAGAGGTGAAATTCTTGGATTGATGGAAGACGTACAACTGCGAAAGCATCTGC

CATGGATGTTTTCATTGATCAAGAACGAAAGTTAGGGGATCGAAGACGATCAGATACCGT

CGTAGTCTTAACCATAAACGATGCCGACTGGGGATTGGCGGGGGCACTTTTATATGACTC

TGTCAGCACCCTGAGGGAAACCAAAGTCTTTGGGTTCTGGGGGGAGTATGGTCGCAAGGC

TGAAACTTAAAGGAATTGACGGAAGGGCACCACCAGGAGTGGAGCCTGCGGCTTAATTTG

ACTCAACACGGGAAAACTTACCAGGTCCGGACAGAAGAATGATTGACAGACTGAAAGCTC

TTTCTTGATTTTTTGGTTGGTGGTGCATGGCCGTTCTTAGTTGGTGGAGTGATTTGTCTG

GTTAATTCCGTTAACGAACGAGACCTCGTCCTGCTAAATAGGTGCGCGCATGCCAGCACT

GCGTTCTTACCTTCTTAGAGGGACTATGCGCGTCTAGCGTATGGAAGATTGAGGCAATAA

CAGGTCTGTGATGCCCTTAGATGTTCTGGGCCGCACGCGCGCTACACTGATGCATTCAAC

GAGTTTCACTTGATAGTCCTGGGTCGGAAGGCTCGGGTAATCTTTTGAAAGTGCATCGTG

CTGGGGATAGATCATTGCAATTATTGATCTTCAACGAGGAATTCCTTGTAGGCGCAGGTC

ATCAGCCTGCGCCGAATACGTCCCTGCCCTTTGTACACACCGCCCGTCGCTCCTACCGAT

TGAATGGTCCGATGAAATGTCCGGATCGCGGCTAGTGTGATGGACTCGTCCAGAGCGGTA
GCCGTGAGAAGCTCATTAAATCTTATCATTTAGAGGAAGGAGAAGTCGTAACAAGGTTTC
CGTAGGTGAACCTGCGGAAGGATCATTCACAAACTATTGACACAAACACACGCGAACCAA
ATCGTCCGCACAGCAGGTGGCAATGAAAGAGAGACTCTGCATGTCGCCTTTCGGGGTAGC
AAGCAGCACTCTTCTGCCATCGCCTCTGTGCAGGGCGTAAATTCCCATTGAGAGGATGTG
AGGGCACCACAGGAAGCTTTTCCACAGGAAGTCGCCATCCTTCTCCCTCCACGGCGGCGC
TTCTGTGCTTGGCAGTTTTTTTTTGCCTTCTAGGGAGGATGCCGCCAATGGAGCCCCTTA
TATATATACATCATCATATGCCCCTTTTTTTCTTAACCGCTTGCCACAGCTTCTTCTATG
AGGAGCTTGTGGGAAGACTGTCTCCATACAATAACAAAGATACAACTCTTAGCGGTGGAT
ATCTTGGTTCTCGCAACGATGAAGAACGCAGCCAACTGCGATAACTAATGTGAATTGCAG
GACTTCGTGAATCATTAAGTCTTTGAACGCAAGTTGCGCTCATCTCCATGAGGAGGTGAG
CATGCCTGTTTGAGTGTCGATGGAAAAACAATACATAACAACAATCGGCTCTCTTATCT
TCCTGCCTTCTTGGGGGCAAAAGGTAGTGCATGCCGGAGATGGGCTTTGTCGCCCCTTTC
TCTTGAAAGAGGCGTCGTCCGAAACACGTATCGTTGCCCGCTGGAGGTCATAGAGCATGA
GTTTAAAGCACCACATCTCTCAAGTCAGAGTGAGCGTCTTGAGTAGGTCCGCATTGATTC
AGTTTCAACCCAGGTGGCGTTGGTTACTTGCCATAAGCCTCTCTGGGGTTTCTTTGCTTG
GACCTCTCTTGTCAGTTTCTCTGACGTAGAGACAGGTGCCGTCTCATGGACGTTGCCCTC
TGGAACGTGCTAAAAAAAAAAAAAAAGAAAAAACCGTACCACATGTCAAAAGAGACAGATC
GTGCTTGACGGCGGGAGGTGTTGCATTTCTTGCTTGTCCTGAGCGTCTGCCAGTCTCTGA
CGAGTGTGTTCCTTCCATAAACCCGATCTCAAATCAGGTAGGACCACCCGCTGAATTTAA
GCATATAATTAAGCGGAGGAAAAGAAACTAACAAGGATTCCCTCAGTAGCGGCGAGCGAA
GCGGGAAGAGCTCAACATGAAAACCGTTGGGGGTTCGCGCTTTCCTTCGGGTTGTATAGT
AAAGTGGTGTCGCCTGCGTCGCGCCGGGTATAAATCCCGTCTGAAATGGGGTGTCGTAGA
GGGTTATATCCCCGTCTGTGACCCGGTGTGCGATGCGTTACGGTGACCTTACGATGAGTC
GGGTTGCTTGGGCATGCAGCCCTAAGTGGGTGGTAAACTTCATCTAAAGCTAAATACTAG
CGAGAGACCGATAGCGTACAAGTACCGTAAGGGAAAGATGCAAAGAACTTTGAAAAGAGA
GTTAAACAGGGCTTGAAACCGTTAAGAGTGAAGCGAAGGAGGTCAGTGATCATGGGGCCA
GATCCCCAAGTGCCTGTCATAGGGCACCGTAGCGTTCTGGTTTCCCGGGGCAGCGCCGGC
CGGGGGCGGGGGTTAACAGGGAGGTGGTTGAGGGATTCGTTTTCCGCCTGTACTCCTTTA
TAGCTTGCACCCCGGCCGAGGGATCCGCGACTTGTCGTGTGAGGGCGCTGTCAAAATGGC
CTTCTCCACCCCGTCTTGAAACACGGACCAAGGAGTCTAACATACTCGCGAGTGTTTGGG
TGGTTAAAACCCGAACGCACAATGAAAGTAAACGGTGGCTCTGCGCCGCTGCTTGGTGGG
AACCCTGCTTTTATAATGGGGGGCACTATCACCCGACCCCGATCCCTTTGGGGTGGTTGG
TTTTGAGGTAGAGCGAGTATGATGGGACCCGAAAGATGGTGAACTATTCCTGAGTAGGGC
GAAGCCAGAGGAAACTCTGGTGGAGGCTCGTAGCGATGCTGACGTGCAAATCGCTCGTCG
AACTTGGGTATAGGGGCGAAAGACTAATCGAACCATCTAGTAGCTGGTTCCCTCCGAAGT

TTCCCTCAGGATAGCGGGAGCAAATGCAAGTCAGTATCATCCGGTAAAGCGAATGATTAG
AGGCCTCGGGGGAGTAACTCCTTCGACCTATTCTCAAACTTTAAATAGGTGAGGAGCCGG
CAACGTTGCTTAAAACAATGAACGCCGTGCGGACAATGATTGCTCTCAGTGGGCCGATTT
TGGTAAGCAGAACTGGCGATGCGGGATGAACCGAAAGCGACGCTAAGGTGCCAAACTCCC
AGCTCATTCAGACACCAGAAAAGGTGTTGATTCATACAGACAGCAGGACGGTGGCCATGG
AAGTCGGAATCCGCTAAGGAGTGTGTAACAACTCACCTGCCGAATGGATCAGCCCTGAAA
ATGGATGGCGCTTAAGCGGGGGACCGAGGCGTCGCCGTTGGGGTAAATACGAAGCCCCAA
TGAGTAGGAGGGCGTGGGGGTTTTGAAACACGGTACACTCGCATGGAAGCCTTGGGCGTA
AGCTCGGGTGGAGCGACCCCTAGTGTAGATCTTGGTGGTAGTAGCAAATATTCAAGTGAG
AACCTTGAAGACTGAAGTGGGGAAGGGTTCCACGCGAACAGTGCTTGGACGTGGGTTAGT
CGGTCCTAAGAGATGGGGAAACTCCGTATTCAAGTGCGCTTTTGTAGCGCAGTCCATCGA
AAGGGAAGCGGGTTAACATTCCCGCACCAGGATGTGGAAACTGTGGTTTACCGCAAACTG
CGGCAACGCAACTGAACCTGGGGACGCCCGTAGGAGCTCCGAGCAGGGATCTCTTTCCTG
TTTAACGCGCCCAACCCCCTGGAATCGGTTTGTCCGGAGATAGGGGGCGAAAGGCCGGAA
AAGCACCGCACGTCTTGTGGTGTTCGGAACGCTTGTAATGGCCCTTGAAAACCCAGGGGA
AAGCAGCCGTCTTACACTTGTGTGAGACGAGGAATTCTCACACCTGGCCGTACCCATCAC
CGCATCAGGTCCCCAAGGTGAACAGCCTCTAGTCAATAGAATAATGTAGATAAGGGAAGT
CGGCAAAATAGATCCGTAACTTTGGGAAAAGGATTGGCTCTAATGTTCGGGCTTGTCGGG
ACGCGCGGGCGTAGATGCCAATGAGCGGCAGACGTCTAGCCTCTTTGGAGGCTGGATTAA
CGCTGCTTGTGGGAAGAAACCGTGTCGATGGAGGCAGGAACGCTTATCGATCTTGTCTCT
TCCCGCTTGCGATCAACGAACGGATTAGAACTGGTACGGACAAGGGGAATCCGACTGTTT
AATTAAAACAAAGCATTGCGACGGCCGGAAACGGTGTTGGCGCAATGTGATTTCTGCCCA
GTGCTCTGAATGTCAAAGTGAAGAAATTCAACCAAGCGCGGGTAAACGGCGGGAGTAACT
ATGACTCTCTTAGACAGATCAAGACACCTTCAATAAATTGGTTCATTATGTCGGGTGGTC
CGTCGTCCGGGAAGCGACCAAAGCAAAACCCCGTAGACTGTGATGTGCATCTGTCAGACG
AAACTGACGTCCGTGTTCATCAAGCCTTAGCTATCTTTCAACGCCGGCTTCGAGACGTGG
CGTCTCTTAAGGTGTTTGAAGTGGCTCTCTCTGTGGGAACTACTGATGCACTGGTAGAGA
TGGGCACTATCACTTGCTGGAGTATTGATGCTGTTGGGAAAGGTTTCTCAAGGAACAGCA
GCGGTCGCTGGCAATTTGACCTCACTGGCAAGATTAATGGCAAGCCTCTAATATCCACCA
CAAAGGATCGTTGGAGGCCGGTTGTGACGGCTGTGCCCTACTACCTTTCGTGCGCCATCG
ATGGGGACAAGACGACGCTGGAGGAAAAGCAGTGGACTTCATCACACATGTGTCACCATG
AATGGTGTGTGAACGTGGAGCACATTGTTTATGAAAGCCTCGACGTCAACAAGTCTCGGA
ACTGGTGCCAGGGAAAAAACGCATGCCTCCATCTGCCTGTTTGCATTCGAACCGGCCGCC
GTGTGCAGGAGCTCAGGAAGAGCAATGGGGTTCGTGCAATGGACCTCCTCCTGCAGAAAT
TGGCTGGATTGGAGATTGAGGTTGAGGAGCCCGCTGCAGCTGTACTCCGAGAGTCTGTCG
AACGGGAGGACGCCTAGTGAGTGACTCGTGACACGGGTTGGCTGCGCATGATGCGTCAAC

TACATGGACATCTTTTGAGAGAGATAGGCGGGCTGAAAAGGTTATAGCCGTCTAGTCTCG
CTCTCTTTCATTGAGCGGGGCGACACACCTGGTTGCGGGAACCCTGCCCACCTTTCTATC
AAGAAGTGGGAACCAGTCCATTCCTACTCCTGCTTGTGAGGCATATAGTAACGTAGCAGC
AGGAGGATCTCGGCGAGAGGCCGAACCCGAAAGGAATAACGTGGGCTGGGAGGCGATCCG
CAGGCGAGTCCGTAACTATATGATTATCGACGGGCCGTCTCAACGACTGGTAAGGTGTGG
GTTTGTCCTCCTCCTCACGTGGGGACAGGCTTAAGGTACAGTCTAATCCCACCCGCCAAG
GGTGCTTGGGTGTAATCCCTCTTCCATAGAGGGCACTAGCCCAGGGGCTGGGACGCTGGG
AGGAAATGCCCCGTCCCTATAGCCGGTAGAAATGAAGGTAGCCAAATGCCTCGTCATCTA
AACTAATTGGATGAGATAGTCTGAGTGAAAAGCTTATGCAGGCTAGTACCGGAAAAAAAA
AAAAAGAGGGGCTCTGCCACTCTGACGGTGCGACACTCCTTGTACAGGGAAACCCTCCGC
ATGGAAAGGCACGTGGTACCACCCTCGCGGGGAAACTCGCGAGCGGGATCCCGGGTAATG
ACCGGAGCCGATGGTAAAAACTCACGTGACCGATGGGGTGACCCTGTGGCGAGACGCTAA
GTTGTGGGACATCACCCTCTCACAATACGCATCCGTCGCAGAGACTGAACGGGAGTGGGC
TCCGTTGACGGCCGGCCGGCCAGCCAGCCGTCTTCCAGGGGCTTAAGATACAGTCCGGGC
CTCAGGGATGGAAAAACCTGGGGAACTACCAGAATTAGTGACGCGCATGAATGGATTAAC
GAGATTCCCACTGTCCCTATCTACTATCTAGCGAAACCACAGCCAAGGGAACGGGCTTGG
AAGAATCAGCGGGGAAAGAAGACCCTGTTGAGCTTGACTCTAGTCCGACTTTGTGAGATG
AGTTGGGAGGTGTAGTCTAAGAGGGAGGGGCAACTCGACCGTGAAATACCTCTACTCCCA
AGCTTGTTTTACTTACTCCGGCGAGCAGGGATGGGTGGTAACACCCTTTTTCTAGGATTA
ACACCTCTTCGGGGGTGGACCTGCGCGGAGGACATTGTCAGGTGGGGAGTTTGGCTGGGG
CGGCACATCTGTTAAAAGATAACGCAGGTGTCCTAAGATGAGCTCAGTGAGAACAGAAAT
CTCACGTAGAACAAAAGGGTAAAAGCTCATTTGATTTTGATTTTCAGTACGAATACAAAC
TGTGAAAGCATGGCCTATCGATCCTTTTGACCTTCAAGGTTTGAAGCAAGAGGTGTCAGA
AAAGTTAACACGCGTTGTGGTAGCTTTTAAGAGTGGACTCAAAGTGTTCCGCTAGTCCCC
CCTCGGACCGCCACTTACAAAAGGCCTGGGGATGGGGGCGACACTGTCAAATTGCGGGGA
CTCCCTAAAGCCTGTCTCTACCAAGCCAGCGCGGCGAAAGCCCCTGGTGGCCGGGGTAAT
GACCTAGGGTATGGTAAAAGCGAGACAGGATGGAACAATGGGCAATCCGCAGCCAAGTCC
TAAGGCTTTTTGAAAAGCGGCTATGGATGCAGTTCAACGACTAGATGGCAGTGGGCTGAG
AGAGCCCACCGCGGCACCCTTTCGGCTTAAGATATAGTCTTGCCCACGTGAAAGCGTGCC
TGCAGATGAACGCCACCATGTATGGCCGGAGCTGCAGGAGACCGCACTTGATGTGCAGTC
GGGAATCGACAGTACCACAGGGATGAACAAAAACACACAATTAATTTGTGGCCATCATAC
AGTCGTATGATATTATCGTCCCAGGCGGGTGCGCCTCTCTCACCCGTCTAGTCCGGCCTT
GGTGCGCAAAAAATGCTTGCGTGGATGCGCCTTGGTTAAGGAAAATGCTCCTAGTAGTTT
TACAAGGACACTTCTTGTAGGGCAACACATCTGGATGCGAAGAACTCCCGGAAAAGACAA
CGGTACCAAGTGACTGGGGTAACGCCCCATTCATGAAGGCAAATAATCAAAAAGAGGTGC
CGGGTAAGAACCCGTTGTATAGGGACAATCCGCAGCTGACTTGGCTCCCTGCCCATCCCC

ATGGGCATAGCTTAGCGGTTCAACGCTCGGTAAGATGTGGGTTGGCGCGCTGTCCGACAG
CGCACTGGCTTAAGGTACGGGCTACTCCCACCGGAGACGGTGTCTTTGTCTACCTCGGGT
AACACGCCTCGGGAGAAGCAAAGAGGGTGCAATGGAAAAAAACTGGTTGCGGGGAAATCC
CTACCAGAGTCACCTGGTACAAAGGAACTGGCTTGTGGCAGCCAAGCGTTCATAGCGACG
TTGCTTTTTGATCCTTCGATGTCGGCTCTTCCTATCATTGCGAAGCAGAATTCGCCAAGT
GTTGGATTGTTCACCCACTAATAGGGAACGTGAGCTGGGTAGCCACTTTATGCCCAACAG
CTGGATAGCTGGGCTTCTGTCTTAGGCCGGGACTTCTCACGACCCGTGCCTTGGTCACCC
AAAAAAGGGCGACCGCAGAGCTCAGGATGGAGCTTCTGGCTTAACTAGTAGAGGACAGAT
AGAGATCGTCAAATTGCGGGAAACCCCTTAGAGCCTCGAGCTACCAAGTGCGTGTCGAAA
GGCATAGACGCAGCCAGCGATGGCTGTCGCATGGCTTGATTCCATCAAGATAGGTAAAAA
TGCTCGAGGATTGGGCAATCCGCAGCCAAGCCCCTAAGGTCCCCCCGCGGGCTATGGGGA
AGGTTCAACGACTGTAAGGCGTACCGTGCAAGCGGAAGATACAGTCTAGCCCCGCTGGAA
AGCAGCGCCAAAGTAAGTAGGTTGCAGTGGGTCTGTCCTC
>SRR9587923
GCATGTTTTGGTATGTGTTTCGACAGTCATGCATGTCATAGCCGTAGGATGGGGAGTGAG
CTTCATAGCCCGACCCCTTCTCAGGAGCATGTTTTCATAGCATATTGAGCATACTGGGGT
GGTCTCCCATGTAGCTCGCTCTCTCGGTGCAGGACCTCCCTTTTGCGGGCCACTTCGATG
GCCTCCATTATGGAGAGCATGTTTTGGCATATGAAGCGTTGGTGGTGAGGCTCGTAGCCT
GTATATACATACCAACGCAGTCGTCATATGTCTCTCTGGCGGGGGTCGGTCGTATGACCT
TGTCCCTACCTGGAGCATGTATTTTGTATGATATACTGGACGGCAGCTGCGCTGTTCAGT
CGTCGGCCGTACCTCATGTGCGCGTTTCCAGGTCCGAACGTTTGACATGCAAGGGCATCA
TACCAGAAGCCTGTTTTCTCGTATGAGCATGTAGGTAGGCAGGCTCTTCAGTCGTCTACC
CATAAATGCACTTCGAAGGAACTTTTTGGTATTGAATGGCTTTGTGCCCAGACGTTCTTG
TGAGCTTGTTCTCGCTCTGCGATACGCTGGAAGCTGTGGTATTACCTACAGCAGACGCAT
TCCTAGGGCGGACGGCCTCTCTGCAGATGGGCACATCTGTAGAAAGAGGCTCCCCTTGAT
GACGCAATGCAATGGCTGGGTAAACGGAGTGCTATACAGAAGGGTCAAACCTCTGTGGAA
GTACACTCATACCAGCATTGTTTGTTGAGAGGGGTCGGCTCTACGAGTGAGGCCCTATAC
TGCCATCTCATTCAGATGCGTAGACGGTCTTTACTCAGACTGCCTTTCTGGCTGATTGTC
CTCGGTCCCTCGTGCGCACGCGCCGCTGGCTGAATGGGCTTGTCCTTATGGAGTGCCGCG
TTCTTCAAAGAATCCGTCTTCATACAGCGACCTCATTTACATATTGCCGGCTGAATGGGC
TTGTCCTTATGGAGCACTGCATTCTTCTAGGAATTGCTATCTATACAGCGACCTCATTTA
CATACAAGGCACCTTTTTCTACATAGACTTGCCATTTGGCTGATTGTTCTGACAGTCGCT
GCCTCTTCGGAGTTCTGTGTGGACGAGTGGCCTCCTGAGTGGGCCGGTCCTTACGGAGTG
CCCAAGTCTGTTATGACTGGGTTCTCCATACAGCGACCTCACTCAGAATTGTATGGCACC
GTGCCGATTGAATGGACAAGCAGTCCTTAGGGAATCGCATGAAAGCATGACGGCAACATG
GTGTATGGACCCGTTGGGCCTTGCCTGTGCTATCTTCTCGAGTCGACCGGTCCTCGTCTC

CTCCTTTCGCGACGGGAAGCTCTTGAGTTTACCGTGGGTCAGGACGCTGTGGTATCTGTT
TGTCTGTAGTGTGGGCTCTCTATCAGAACAGCGTATCAGCACACGTCGTCTGGGTCTATG
GTATTTTTGCTTAACGGCCCTGGGTTTGGTAGTATGGTTCCATGTCGGTCCTCTCACCTC
CCAACCGTCATGTCTCTGATATCATCTGCACAGTACGTCTTGTCAGAATATCATTCTGTA
TGGCAGTCGATAGACTAGGCCCTATGCCGCAGCTGTCTCTGACCACTATGCGAGCGCATA
CCTACCTCAATGTGTTCATAGCTGTTATGGATCTTGGGTGTGGTGTGCCAGAACGATACG
CCACCAGATTGTGGAATGGTGCTTGTAGTATTAGAAGAGGACATGTATGAAGAGCACGTT
CTTTTCTCTTATGATGGTGCCGCGGGTGCGGGCCGTTCTCATTCTGCTTGTCGGGTGCTC
AAGGTGCATGTAGGATTGCTGAACCCACTGAATGGTCATGAGAATGGTCATATGGTGAGG
TTCACTTCTCCATCGCTTTGTGTGTGTTGGATGTTTGGGTGGCTTCGGCCGCATGGAAAA
TCTCGGTCTGTAACTCATCAGGGAGGGTTTCCATGTGAGCAATCTCTCCAGTCGAACACA
CACCCATGGGCGTCCCGCTAGGCAAAATGGAACTCTCGATCCGACACAGCCTGTGGAGTG
TTTGTTCCGTGTGGCGTGAAACGGCTTATACTGGATGTCTTCCCTTTGGACACCAGTCTG
TTGGTATGTACGCATACCATCTAGAGGGAGCAAGAAAAAGGCATGCAAGTGCCACAAAGG
AAGGGAGGCCTTCCTGCTACCGTGTGTCCTCTTGACGATCACGGGGGAGTATATAGAGAC
ATTTGGACGAGATCCGTTTTTCACGAGGCGGTGCCCTCGCGCAAAGGTGTTGCTCTGCAC
GCATCTTTATGATGATAGTTGCTCGCTCCTATGCGTCCACTCGTGAAAATACCTGCTCTC
TCTGTCTGTTTCCATGACACCCCCATGCGTTGTCATAGAGCATGCTCAGGTACAGAAAGT
GCTGTGACTTTCTTGGCCTGCTGTCACTGAGTCAGCCTGATACTTTGGAGGAATCTTTTC
TGGGCACTGCCCAGCTGAGAGTTCTTATCAGGCTCTGGAGTGTGCATATGCCTATATTCA
TTATAGTGCATGTGGCTACGCAGAAAAAGCATCGACGTGGAATGGAGAAGAACACCATTC
TGAATGGAACGGTTCTTACTCTACTTCTGTTGCTTGACGCTGTATGGGAATCGCTTTGCT
CTGGGGGTACACCTTCCATTCCATGACAAAGCATTTGATCTATGCCCCGAGGGTCATTCT
CTATGAAGAGCCCGAAACTTGTCTGAGTGAGAGGCGAAGGTAGTCCTTCATTTATGCGGG
GTCCGCCCTCACACTGTAGTCGGCACAGTCCCTTGTTTATGGACTGCGCAGCTCAATCTT
CGACTGGAGAGGAAAAACAAAACACAAGTATGACACGCACATACGTGTTTTCTAGGACTG
TGACCTGTCGGCTGCCTCGCGGTGGTTGGCATGATGCCCCAAGTGGGGGGGGGGAGAAAC
AATCAGTATGATTGATAAAAACCAGTGGGGAAAAAGGCGCGCTGTCGCACACCCTTGGTC
ACGGCCTCAAGTCGACGGCCCTCACTGGCCCACCCTGCGACTCCTGTTCGCTTTTGACAA
GACACACCTGCAATACGGAGGAAATGCCTGAGCAGCTGTGCCGCAGCAACTGCGGACGCG
TAGCGATATGCATCCCTGTTGTTTGCGACTTGGTACAAAGAACGTGCGCCGGGAGACGGA
TTGCCCTTTGACTTTTCCACCCAGCCCGCGGCTCTATACCCTTGACGTACCTATTTACCT
GGTTGATCCTGCCAGTAGTCATATGCTTGTCTCAAAGACTAAGCCATGCATGTCTAAGTA
TAAACACTTTTATACTCGTGAAACTGCGAATGGCTCATTAAAACAGTTATAGTTCCTTTG
GGAACAACACTACTTGGATAACCGTAGTAATTCTAGAGCTAATACATGCCTCAACGCCCG
ACTCACGAAGGGTGGTATTTATTGGATAAAAAACCATCGTGCTCTGTCCAGAGCGCTTTG

AGATGATTCACAATAACTTGTCGGATCGCATGGCCTCGTGCTGGCGACTCCTCATTCAAA
TTTCTGCCCTATCAACTTTCGATGGTAGAGTATTGGTCTACCATGGTGTCGACGGGTGAC
GGGGAATTAGGGTTCGATTCCGGAGAGGGAGCCTGAGAAACGGCTACCACATCCAAGGAA
GGCAGCAGGCGCGCAAATTACCCAATCCCGACTCGGGGAGGTAGTGACAAAAAATAACAA
TAGGGGGCCCTTTGGGTCTTCTAATTGGAATGAGAACAATTTAAATCCCTTATCGAGGAT
CCATTGGAGGGCAAGTCTGGTGCCAGCAGCCGCGGTAATTCCAGCTCCAATAGCGTATAT
TAAAGTTGTTGCAGTTAAAACGCTCGTAGTCGGACCTCGGAGCGCCCCCACGGGCCGGGC
TTTTGCCTTGTTGCCAGTGGGAGTTTGTCTCCTTTTGTCGGGACGCGGTGTGCAGGGCCT
TTACTGTCTCTGCGCGTTTACCTGCGGGCCGACCGTTTACTGTGAAGAAAGTAAAGTGTT
CAAAGCAGGCCAATGCCTTGAATATGTGAGCATGGAATAATAGAATAGGACTTGGGCTCT
ATTTTGTTGGTTTTCAGTGACCAAGTAATGATTAATAGGGACGGTTGGGGGCATTCGTAT
TTCATTGTCAGAGGTGAAATTCTTGGATTGATGGAAGACGTACAACTGCGAAAGCATCTG
CCATGGATGTTTTCATTGATCAAGAACGAAAGTTAGGGGATCGAAGACGATCAGATACCG
TCGTAGTCTTAACCATAAACGATGCCGACTGGGGATTGGCGGGGGCACTTTTATATGACT
CTGTCAGCACCCTGAGGGAAACCAAAGTCTTTGGGTTCTGGGGGGAGTATGGTCGCAAGG
CTGAAACTTAAAGGAATTGACGGAAGGGCACCACCAGGAGTGGAGCCTGCGGCTTAATTT
GACTCAACACGGGAAAACTTACCAGGTCCGGACAGAAGAATGATTGACAGACTGAAAGCT
CTTTCTTGATTTTTTGGTTGGTGGTGCATGGCCGTTCTTAGTTGGTGGAGTGATTTGTCT
GGTTAATTCCGTTAACGAACGAGACCTCGTCCTGCTAAATAGGTGCGCGCATGCCAGCAC
TGCGTTCTTACCTTCTTAGAGGGACTATGCGCGTCTAGCGTATGGAAGATTGAGGCAATA
ACAGGTCTGTGATGCCCTTAGATGTTCTGGGCCGCACGCGCGCTACACTGATGCATTCAA
CGAGTTTCACTTGATAGTCCTGGGTCGGAAGGCTCGGGTAATCTTTTGAAAGTGCATCGT
GCTGGGGATAGATCATTGCAATTATTGATCTTCAACGAGGAATTCCTTGTAGGCGCAGGT
CATCAGCCTGCGCCGAATACGTCCCTGCCCTTTGTACACACCGCCCGTCGCTCCTACCGA
TTGAATGGTCCGATGAAATGTCCGGATCGCGGCTAGTGTGATGGACTCGTCCAGAGCGGT
AGCCGTGAGAAGCTCATTAAATCTTATCATTTAGAGGAAGGAGAAGTCGTAACAAGGTTT
CCGTAGGTGAACCTGCGGAAGGATCATTCACAAACTATTGACACAAACACACGCGAACCA
AATCGTCCGCACAGCAGGTGGCAATGAAAGAGAGACTCTGCATGTCGCCTTTCGGGGTAG
CAAGCAGCACTCTTCTGCCATCGCCTCTGTGCAGGGCGTAAATTCCCATTGAGAGGATGT
GAGGGCACCACAGGAAGCTTTTCCACAGGAAGTCGCCATCCTTCTCCCTCCACGGCGGCG
CTTCTGTGCTTGGCAGTTTTTTTTTGCCTTCTAGGGAGGATGCCGCCAATGGAGCCCCTT
ATATATATACATCATCATATGCCCCTTTTTTTCTTAACCGCTTGCCACAGCTTCTTCTAT
GAGGAGCTTGTGGGAAGACTGTCTCCATACAATAACAAAGATACAACTCTTAGCGGTGGA
TATCTTGGTTCTCGCAACGATGAAGAACGCAGCCAACTGCGATAACTAATGTGAATTGCA
GGACTTCGTGAATCATTAAGTCTTTGAACGCAAGTTGCGCTCATCTCCATGAGGAGGTGA
GCATGCCTGTTTGAGTGTCGATGGGAAAAACAATACATAACAACAATCGGCTCTCTTATC

TTCCTGCCTTCTTGGGGGCAAAAGGTAGTGCATGCCGGAGATGGGCTTTGTCGCCCCTTT
CTCTTGAAAGAGGCGTCGTCCGAAACACGTATCGTTGCCCGCTGGAGGTCATAGAGCATG
AGTTTAAAGCACCACATCTCTCAAGTCAGAGTGAGCGTCTTGAGTAGGTCCGCATTGATT
CAGTTTCAACCCAGGTGGCGTTGGTTACTTGCCATAAGCCTCTCTGGGGTTTCTTTGCTT
GGACCTCTCTTGTCAGTTTCTCTGACGTAGAGACAGGTGCCGTCTCATGGACGTTGCCCT
CTGGAACGTGCTAAAAAAAAAAAAAAAGAAAAAACCGTACCACATGTCAAAAGAGACAGAT
CGTGCTTGACGGCGGGAGGTGTTGCATTTCTTGCTTGTCCTGAGCGTCTGCCAGTCTCTG
ACGAGTGTGTTCCTTCCATAAACCCGATCTCAAATCAGGTAGGACCACCCGCTGAATTTA
AGCATATAATTAAGCGGAGGAAAAGAAACTAACAAGGATTCCCTCAGTAGCGGCGAGCGA
AGCGGGAAGAGCTCAACATGAAAACCGTTGGGGGTTCGCGCTTTCCTTCGGGTTGTATAG
TAAAGTGGTGTCGCCTGCGTCGCGCCGGGTATAAATCCCGTCTGAAATGGGGTGTCGTAG
AGGGTTATATCCCCGTCTGTGACCCGGTGTGCGATGCGTTACGGTGACCTTACGATGAGT
CGGGTTGCTTGGGCATGCAGCCCTAAGTGGGTGGTAAACTTCATCTAAAGCTAAATACTA
GCGAGAGACCGATAGCGTACAAGTACCGTAAGGGAAAGATGCAAAGAACTTTGAAAAGAG
AGTTAAACAGGGCTTGAAACCGTTAAGAGTGAAGCGAAGGAGGTCAGTGATCATGGGGCC
AGATCCCCAAGTGCCTGTCATAGGGCACCGTAGCGTTCTGGTTTCCCGGGGCAGCGCCGG
CCGGGGGCGGGGGTTAACAGGGAGGTGGTTGAGGGATTCGTTTTCCGCCTGTACTCCTTT
ATAGCTTGCACCCCGGCCGAGGGATCCGCGACTTGTCGTGTGAGGGCGCTGTCAAAATGG
CCTTCTCCACCCCGTCTTGAAACACGGACCAAGGAGTCTAACATACTCGCGAGTGTTTGG
GTGGTTAAAACCCGAACGCACAATGAAAGTAAACGGTGGCTCTGCGCCGCTGCTTGGTGG
GAACCCTGCTTTTATAATGGGGGGCACTATCACCCGACCCCGATCCCTTTGGGGTGGTTG
GTTTTGAGGTAGAGCGAGTATGATGGGACCCGAAAGATGGTGAACTATTCCTGAGTAGGG
CGAAGCCAGAGGAAACTCTGGTGGAGGCTCGTAGCGATGCTGACGTGCAAATCGCTCGTC
GAACTTGGGTATAGGGGCGAAAGACTAATCGAACCATCTAGTAGCTGGTTCCCTCCGAAG
TTTCCCTCAGGATAGCGGGAGCAAATGCAAGTCAGTATCATCCGGTAAAGCGAATGATTA
GAGGCCTCGGGGGAGTAACTCCTTCGACCTATTCTCAAACTTTAAATAGGTGAGGAGCCG
GCAACGTTGCTTAAAACAATGAACGCCGTGCGGACAATGATTGCTCTCAGTGGGCCGATT
TTGGTAAGCAGAACTGGCGATGCGGGATGAACCGAAAGCGACGCTAAGGTGCCAAACTCC
CAGCTCATTCAGACACCAGAAAGGTGTTGATTCATACAGACAGCAGGACGGTGGCCATG
GAAGTCGGAATCCGCTAAGGAGTGTGTAACAACTCACCTGCCGAATGGATCAGCCCTGAA
AATGGATGGCGCTTAAGCGGGGGACCGAGGCGTCGCCGTTGGGGTAAATACGAAGCCCCA
ATGAGTAGGAGGGCGTGGGGGTTTTGAAACACGGTACACTCGCATGGAAGCCTTGGGCGT
AAGCTCGGGTGGAGCGACCCCTAGTGTAGATCTTGGTGGTAGTAGCAAATATTCAAGTGA
GAACCTTGAAGACTGAAGTGGGGAAGGGTTCCACGCGAACAGTGCTTGGACGTGGGTTAG
TCGGTCCTAAGAGATGGGGAAACTCCGTATTCAAGTGCGCTTTTGTAGCGCAGTCCATCG
AAAGGGAAGCGGGTTAACATTCCCGCACCAGGATGTGGAAACTGTGGTTTACCGCAAACT

GCGGCAACGCAACTGAACCTGGGGACGCCCGTAGGAGCTCCGAGCAGGGATCTCTTTCCT
GTTTAACGCGCCCAACCCCCTGGAATCGGTTTGTCCGGAGATAGGGGGCGAAAGGCCGGA
AAAGCACCGCACGTCTTGTGGTGTTCGGAACGCTTGTAATGGCCCTTGAAAACCCAGGGG
AAAGCAGCCGTCTTACACTTGTGTGAGACGAGGAATTCTCACACCTGGCCGTACCCATCA
CCGCATCAGGTCCCCAAGGTGAACAGCCTCTAGTCAATAGAATAATGTAGATAAGGGAAG
TCGGCAAAATAGATCCGTAACTTTGGGAAAAGGATTGGCTCTAATGTTCGGGCTTGTCGG
GACGCGCGGGCGTAGATGCCAATGAGCGGCAGACGTCTAGCCTCTTTGGAGGCTGGATTA
ACGCTGCTTGTGGGAAGAAACCGTGTCGATGGAGGCAGGAACGCTTATCGATCTTGTCTC
TTCCCGCTTGCGATCAACGAACGGATTAGAACTGGTACGGACAAGGGGAATCCGACTGTT
TAATTAAAACAAAGCATTGCGACGGCCGGAAACGGTGTTGGCGCAATGTGATTTCTGCCC
AGTGCTCTGAATGTCAAAGTGAAGAAATTCAACCAAGCGCGGGTAAACGGCGGGAGTAAC
TATGACTCTCTTAAGGTAGCCAAATGCCTCGTCATCTAAACTAATTGGATGAGATAGTCT
GAGTGAAAAGCTTATGCAGGCTAGTACCGGAAAAAAAAAAAAAGAGGGGCTCTGCCACTC
TGACGGTGCGACACTCCTTGTACAGGGAAACCCTCCGCATGGAAAGGCACGTGGTACCAC
CCTCGCGGGGAAACTCGCGAGCGGGATCCCGGGTAATGACCGGAGCCGATGGTAAAAACT
CACGTGACCGATGGGGTGACCCTGTGGCGAGACGCTAAGTTGTGGGACATCACCCTCTCA
CAATACGCATCCGTCGCAGAGACTGAACGGGAGTGGGCTCCGTTGACGGCCGGCCGGCCA
GCCAGCCGTCTTCCAGGGGCTTAAGATACAGTCCGGGCCTCAGGGATGGAAAAACCTGGG
GAACTACCAGAATTAGTGACGCGCATGAATGGATTAACGAGATTCCCACTGTCCCTATCT
ACTATCTAGCGAAACCACAGCCAAGGGAACGGGCTTGGAAGAATCAGCGGGGAAAGAAGA
CCCTGTTGAGCTTGACTCTAGTCCGACTTTGTGAGATGAGTTGGGAGGTGTAGTCTAAGA
GGGAGGGGCAACTCGACCGTGAAATACCTCTACTCCCAAGCTTGTTTTACTTACTCCGGC
GAGCAGGGATGGGTGGTAACACCCTTTTTCTAGGATTAACACCTCTTCGGGGGTGGACCT
GCGCGGAGGACATTGTCAGGTGGGGAGTTTGGCTGGGGCGGCACATCTGTTAAAAGATAA
CGCAGGTGTCCTAAGATGAGCTCAGTGAGAACAGAAATCTCACGTAGAACAAAAGGGTAA
AAGCTCATTTGATTTTGATTTTCAGTACGAATACAAACTGTGAAAGCATGGCCTATCGAT
CCTTTTGACCTTCAAGGTTTGAAGCAAGAGGTGTCAGAAAAGTTAACACGCGTTGTGGTA
GCTTTTAAGAGTGGACTCAAAGTGTTCCGCTAGTCCCCCCTCGGACCGCCACTTACAAAA
GGCCTGGGGATGGGGGCGACACTGTCAAATTGCGGGGACTCCCTAAAGCCTGTCTCTACC
AAGCCAGCGCGGCGAAAGCCCCTGGTGGCCGGGGTAATGACCTAGGGTATGGTAAAAGCG
AGACAGGATGGAACAATGGGCAATCCGCAGCCAAGTCCTAAGGCTTTTTGAAAAGCGGCT
ATGGATGCAGTTCAACGACTAGATGGCAGTGGGCTGAGAGAGCCCACCGCGGCACCCTTT
CGGCTTAAGATATAGTCTTGCCCACGTGAAAGCGTGCCTGCAGATGAACGCCACCATGTA
TGGCCGGAGCTGCAGGAGACCGCACTTGATGTGCAGTCGGGAATCGACAGTACCACAGGG
ATGAACAAAAACACACAATTGTGGCCATCATACAGTCGTATGATATTATCGTCCCAGGCG
GGTGCGCCTCTCTCACCCGTCTAGTCCGGCCTTGGTGCGCAAAAAATGCTTGCGTGGATG

CGCCTTGGTTAAGGAAAATGCTCCTAGTAGTTTTACAAGGACACTTCTTGTAGGGCAACA
CATCTGGATGCGAAGAACTCCCGGAAAAGACAACGGTACCAAGTGACTGGGGTAACGCCC
CATTCATGAAGGCAAATAATCAAAAAGAGGTGCCGGGTAAGAACCCGTTGTATAGGGACA
ATCCGCAGCTGACTTGGCTCCCTGCCCATCCATGGGCATAGCTTAGCGGTTCAACGCTCG
GTAAGATGTGGGTTGGCGCGCTGTCCGACAGCGCACTGGCTTAAGGTACGGGCTACTCCC
ACCGGAGACGGTGTCTTTGTCTACCTCGGGTAACACGCCTCGGGAGAAGCAAAGAGGGTG
CAATGGAAAAAAACTGGTTGCGGGGAAATCCCTACCAGAGTCACCTGGTACAAAGGAACT
GGCTTGTGGCAGCCAAGCGTTCATAGCGACGTTGCTTTTTGATCCTTCGATGTCGGCTCT
TCCTATCATTGCGAAGCAGAATTCGCCAAGTGTTGGATTGTTCACCCACTAATAGGGAAC
GTGAGCTGGGTAGCCACTTTATGCCCAACAGCTGGATAGCTGGGCTTCTGTCTTAGGCCG
GGACTTCTCACGACCCGTGCCTTGGTCACCCAAAAAAGGGCGACCGCAGAGCTCAGGATG
GAGCTTCTGGCTTAACTAGTAGAGGACAGATAGAGATCGTCAAATTGCGGGAAACCCCTT
AGAGCCTCGAGCTACCAAGTGCGTGTCGAAAGGCATAGACGCAGCCAGCGATGGCTGTCG
CATGGCTTGATTCCATCAAGATAGGTAAAAATGCTCGAGGATTGGGCAATCCGCAGCCAA
GCCCCTAAGGTCCCCCCGCGGGCTATGGGGAAGGTTCAACGACTGTAAGGCGTACCGTGC
AAGCGGAAGATACAGTCTAGCCCCGCTGGAAAGCAGCGCCAAAGTAAGTAGGTTGCAGTG
GGTCTGTCCTC
>SRR9587924_SRR9587965
GCATGTTTTGGTATGTGTTTCGACAGTCATGCATGTCATAGCCGTAGGATGGGGAGTGAG
CTTCATAGCCCGACCCCTTCTCAGGAGCATGTTTTCATAGCATATTGAGCATACTGGGGT
GGTCTCCCATGTAGCTCGCTCTCTCGGTGCAGGACCTCCCTTTTGCGGGCCACTTCGATG
GCCTCCATTATGGAGAGCATGTTTTGGCATATGAAGCGTTGGTGGTGAGGCTCGTAGCCT
GTATATACATACCAACGCAGTCGTCATATGTCTCTCTGGCGGGGGTCGGTCGTATGACCT
TGTCCCTACCTGGAGCATGTATTTTGTATGATATACTGGACGGCAGCTGCGCTGTTCAGT
CGTCGGCCGTACCTCATGTGCGCGTTTCCAGGTCCGAACGTTTGACATGCAAGGGCATCA
TACCAGAAGCCTGTTTTCTCGTATGAGCATGTAGGTAGGCAGGCTCTTCAGTCGTCTACC
CATAAATGCACTTCGAAGGAACTTTTTGGTATTGAATGGCTTTGTGCCCAGACGTTCTTG
TGAGCTTGTTCTCGCTCTGCGATACGCTGGAAGCTGTGGTATTACCTACAGCAGACGCAT
TCCTAGGGCGGACGGCCTCTCTGCAGATGGGCACATCTGTAGAAAGAGGCTCCCCTTGAT
GACGCAATGCAATGGCTGGGTAAACGGAGTGCTATACAGAAGGGTCAAACCTCTGTGGAA
GTACACTCATACCAGCATTGTTTGTTGAGAGGGGTCGGCTCTACGAGTGAGGCCCTATAC
TGCCATCTCATTCAGATGCGTAGACGGTCTTTACTCAGACTGCCTTTCTGGCTGATTGTC
CTCGGTCCCTCGTGCGCACGCGCCGCTGGCTGAATGGGCTTGTCCTTATGGAGTGCCGCG
TTCTTCAAAGAATCCGTCTTCATACAGCGACCTCATTTACATATTGCCGGCTGAATGGGC
TTGTCCTTATGGAGCACTGCATTCTTCTAGGAATTGCTATCTATACAGCGACCTCATTTA
CATACAAGGCACCTTTTTCTACATAGACTTGCCATTTGGCTGATTGTTCTGACAGTCGCT

GCCTCTTCGGAGTTCTGTGTGGACGAGTGGCCTCCTGAGTGGGCCGGTCCTTACGGAGTG
CCCAAGTCTGTTATGACTGGGTTCTCCATACAGCGACCTCACTCAGAATTGTATGGCACC
GTGCCGATTGAATGGACAAGCAGTCCTTAGGGAATCGCATGAAAGCATGACGGCAACATG
GTGTATGGACCCGTTGGGCCTTGCCTGTGCTATCTTCTCGAGTCGACCGGTCCTCGTCTC
CTCCTTTCGCGACGGGAAGCTCTTGAGTTTACCGTGGGTCAGGACGCTGTGGTATCTGTT
TGTCTGTAGTGTGGGCTCTCTATCAGAACAGCGTATCAGCACACGTCGTCTGGGTCTATG
GTATTTTTGCTTAACGGCCCTGGGTTTGGTAGTATGGTTCCATGTCGGTCCTCTCACCTC
CCAACCGTCATGTCTCTGATATCATCTGCACAGTACGTCTTGTCAGAATATCATTCTGTA
TGGCAGTCGATAGACTAGGCCCTATGCCGCAGCTGTCTCTGACCACTATGCGAGCGCATA
CCTACCTCAATGTGTTCATAGCTGTTATGGATCTTGGGTGTGGTGTGCCAGAACGATACG
CCACCAGATTGTGGAATGGTGCTTGTAGTATTAGAAGAGGACATGTATGAAGAGCACGTT
CTTTTCTCTTATGATGGTGCCGCGGGTGCGGGCCGTTCTCATTCTGCTTGTCGGGTGCTC
AAGGTGCATGTAGGATTGCTGAACCCACTGAATGGTCATGAGAATGGTCATATGGTGAGG
TTCACTTCTCCATCGCTTTGTGTGTGTTGGATGTTTGGGTGGCTTCGGCCGCATGGAAAA
TCTCGGTCTGTAACTCATCAGGGAGGGTTTCCATGTGAGCAATCTCTCCAGTCGAACACA
CACCCATGGGCGTCCCGCTAGGCAAAATGGAACTCTCGATCCGACACAGCCTGTGGAGTG
TTTGTTCCGTGTGGCGTGAAACGGCTTATACTGGATGTCTTCCCTTTGGACACCAGTCTG
TTGGTATGTACGCATACCATCTAGAGGGAGCAAGAAAAAGGCATGCAAGTGCCACAAAGG
AAGGGAGGCCTTCCTGCTACCGTGTGTCCTCTTGACGATCACGGGGGAGTATATAGAGAC
ATTTGGACGAGATCCGTTTTTCACGAGGCGGTGCCCTCGCGCAAAGGTGTTGCTCTGCAC
GCATCTTTATGATGATAGTTGCTCGCTCCTATGCGTCCACTCGTGAAAATACCTGCTCTC
TCTGTCTGTTTCCATGACACCCCCATGCGTTGTCATAGAGCATGCTCAGGTACAGAAAGT
GCTGTGACTTTCTTGGCCTGCTGTCACTGAGTCAGCCTGATACTTTGGAGGAATCTTTTC
TGGGCACTGCCCAGCTGAGAGTTCTTATCAGGCTCTGGAGTGTGCATATGCCTATATTCA
TTATAGTGCATGTGGCTACGCAGAAAAAGCATCGACGTGGAATGGAGAAGAACACCATTC
TGAATGGAACGGTTCTTACTCTACTTCTGTTGCTTGACGCTGTATGGGAATCGCTTTGCT
CTGGGGGTACACCTTCCATTCCATGACAAAGCATTTGATCTATGCCCCGAGGGTCATTCT
CTATGAAGAGCCCGAAACTTGTCTGAGTGAGAGGCGAAGGTAGTCCTTCATTTATGCGGG
GTCCGCCCTCACACTGTAGTCGGCACAGTCCCTTGTTTATGGACTGCGCAGCTCAATCTT
CGACTGGAGAGGAAAAACAAAACACAAGTATGACACGCACATACGTGTTTTCTAGGACTG
TGACCTGTCGGCTGCCTCGCGGTGGTTGGCATGATGCCCCAAGTGGGGGGGGGGAGAAAC
AATCAGTATGATTGATAAAAACCAGTGGGGAAAAAGGCGCGCTGTCGCACACCCTTGGTC
ACGGCCTCAAGTCGACGGCCCTCACTGGCCCACCCTGCGACTCCTGTTCGCTTTTGACAA
GACACACCTGCAATACGGAGGAAATGCCTGAGCAGCTGTGCCGCAGCAACTGCGGACGCG
TAGCGATATGCATCCCTGTTGTTTGCGACTTGGTACAAAGAACGTGCGCCGGGAGACGGA
TTGCCCTTTGACTTTTCCACCCAGCCCGCGGCTCTATACCCTTGACGTACCTATTTACCT

GGTTGATCCTGCCAGTAGTCATATGCTTGTCTCAAAGACTAAGCCATGCATGTCTAAGTA

TAAACACTTTTATACTCGTGAAACTGCGAATGGCTCATTAAAACAGTTATAGTTCCTTTG

GGAACAACACTACTTGGATAACCGTAGTAATTCTAGAGCTAATACATGCCTCAACGCCCG

ACTCACGAAGGGTGGTATTTATTGGATAAAAAACCATCGTGCTCTGTCCAGAGCGCTTTG

AGATGATTCACAATAACTTGTCGGATCGCATGGCCTCGTGCTGGCGACTCCTCATTCAAA

TTTCTGCCCTATCAACTTTCGATGGTAGAGTATTGGTCTACCATGGTGTCGACGGGTGAC

GGGGAATTAGGGTTCGATTCCGGAGAGGGAGCCTGAGAAACGGCTACCACATCCAAGGAA

GGCAGCAGGCGCGCAAATTACCCAATCCCGACTCGGGGAGGTAGTGACAAAAAATAACAA

TAGGGGGCCCTTTGGGTCTTCTAATTGGAATGAGAACAATTTAAATCCCTTATCGAGGAT

CCATTGGAGGGCAAGTCTGGTGCCAGCAGCCGCGGTAATTCCAGCTCCAATAGCGTATAT

TAAAGTTGTTGCAGTTAAAACGCTCGTAGTCGGACCTCGGAGCGCCCCCACGGGCCGGGC

TTTTGCCTTGTTGCCAGTGGGAGTTTGTCTCCTTTTGTCGGGACGCGGTGTGCAGGGCCT

TTACTGTCTCTGCGCGTTTACCTGCGGGCCGACCGTTTACTGTGAAGAAAGTAAAGTGTT

CAAAGCAGGCCAATGCCTTGAATATGTGAGCATGGAATAATAGAATAGGACTTGGGCTCT

ATTTTGTTGGTTTTCAGTGACCAAGTAATGATTAATAGGGACGGTTGGGGGCATTCGTAT

TTCATTGTCAGAGGTGAAATTCTTGGATTGATGGAAGACGTACAACTGCGAAAGCATCTG

CCATGGATGTTTTCATTGATCAAGAACGAAAGTTAGGGGATCGAAGACGATCAGATACCG

TCGTAGTCTTAACCATAAACGATGCCGACTGGGGATTGGCGGGGGCACTTTTATATGACT

CTGTCAGCACCCTGAGGGAAACCAAAGTCTTTGGGTTCTGGGGGGAGTATGGTCGCAAGG

CTGAAACTTAAAGGAATTGACGGAAGGGCACCACCAGGAGTGGAGCCTGCGGCTTAATTT

GACTCAACACGGGAAAACTTACCAGGTCCGGACAGAAGAATGATTGACAGACTGAAAGCT

CTTTCTTGATTTTTTGGTTGGTGGTGCATGGCCGTTCTTAGTTGGTGGAGTGATTTGTCT

GGTTAATTCCGTTAACGAACGAGACCTCGTCCTGCTAAATAGGTGCGCGCATGCCAGCAC

TGCGTTCTTACCTTCTTAGAGGGACTATGCGCGTCTAGCGTATGGAAGATTGAGGCAATA

ACAGGTCTGTGATGCCCTTAGATGTTCTGGGCCGCACGCGCGCTACACTGATGCATTCAA

CGAGTTTCACTTGATAGTCCTGGGTCGGAAGGCTCGGGTAATCTTTTGAAAGTGCATCGT

GCTGGGGATAGATCATTGCAATTATTGATCTTCAACGAGGAATTCCTTGTAGGCGCAGGT

CATCAGCCTGCGCCGAATACGTCCCTGCCCTTTGTACACACCGCCCGTCGCTCCTACCGA

TTGAATGGTCCGATGAAATGTCCGGATCGCGGCTAGTGTGATGGACTCGTCCAGAGCGGT

AGCCGTGAGAAGCTCATTAAATCTTATCATTTAGAGGAAGGAGAAGTCGTAACAAGGTTT

CCGTAGGTGAACCTGCGGAAGGATCATTCACAAACTATTGACACAAACACACGCGAACCA

AATCGTCCGCACAGCAGGTGGCAATGAAAGAGAGACTCTGCATGTCGCCTTTCGGGGTAG

CAAGCAGCACTCTTCTGCCATCGCCTCTGTGCAGGGCGTAAATTCCCATTGAGAGGATGT

GAGGGCACCACAGGAAGCTTTTCCACAGGAAGTCGCCATCCTTCTCCCTCCACGGCGGCG

CTTCTGTGCTTGGCAGTTTTTTTTTGCCTTCTAGGGAGGATGCCGCCAATGGAGCCCCTT

ATATATATACATCATCATATGCCCCTTTTTTTCTTAACCGCTTGCCACAGCTTCTTCTAT

GAGGAGCTTGTGGGAAGACTGTCTCCATACAATAACAAAGATACAACTCTTAGCGGTGGA
TATCTTGGTTCTCGCAACGATGAAGAACGCAGCCAACTGCGATAACTAATGTGAATTGCA
GGACTTCGTGAATCATTAAGTCTTTGAACGCAAGTTGCGCTCATCTCCATGAGGAGGTGA
GCATGCCTGTTTGAGTGTCGATGGGAAAAACAATACATAACAACAATCGGCTCTCTTATC
TTCCTGCCTTCTTGGGGGCAAAAGGTAGTGCATGCCGGAGATGGGCTTTGTCGCCCCTTT
CTCTTGAAAGAGGCGTCGTCCGAAACACGTATCGTTGCCCGCTGGAGGTCATAGAGCATG
AGTTTAAAGCACCACATCTCTCAAGTCAGAGTGAGCGTCTTGAGTAGGTCCGCATTGATT
CAGTTTCAACCCAGGTGGCGTTGGTTACTTGCCATAAGCCTCTCTGGGGTTTCTTTGCTT
GGACCTCTCTTGTCAGTTTCTCTGACGTAGAGACAGGTGCCGTCTCATGGACGTTGCCCT
CTGGAACGTGCTAAAAAAAAAAAAAAAGAAAAAACCGTACCACATGTCAAAAGAGACAGAT
CGTGCTTGACGGCGGGAGGTGTTGCATTTCTTGCTTGTCCTGAGCGTCTGCCAGTCTCTG
ACGAGTGTGTTCCTTCCATAAACCCGATCTCAAATCAGGTAGGACCACCCGCTGAATTTA
AGCATATAATTAAGCGGAGGAAAAGAAACTAACAAGGATTCCCTCAGTAGCGGCGAGCGA
AGCGGGAAGAGCTCAACATGAAAACCGTTGGGGGTTCGCGCTTTCCTTCGGGTTGTATAG
TAAAGTGGTGTCGCCTGCGTCGCGCCGGGTATAAATCCCGTCTGAAATGGGGTGTCGTAG
AGGGTTATATCCCCGTCTGTGACCCGGTGTGCGATGCGTTACGGTGACCTTACGATGAGT
CGGGTTGCTTGGGCATGCAGCCCTAAGTGGGTGGTAAACTTCATCTAAAGCTAAATACTA
GCGAGAGACCGATAGCGTACAAGTACCGTAAGGGAAAGATGCAAAGAACTTTGAAAAGAG
AGTTAAACAGGGCTTGAAACCGTTAAGAGTGAAGCGAAGGAGGTCAGTGATCATGGGGCC
AGATCCCCAAGTGCCTGTCATAGGGCACCGTAGCGTTCTGGTTTCCCGGGGCAGCGCCGG
CCGGGGGCGGGGGTTAACAGGGAGGTGGTTGAGGGATTCGTTTTCCGCCTGTACTCCTTT
ATAGCTTGCACCCCGGCCGAGGGATCCGCGACTTGTCGTGTGAGGGCGCTGTCAAAATGG
CCTTCTCCACCCCGTCTTGAAACACGGACCAAGGAGTCTAACATACTCGCGAGTGTTTGG
GTGGTTAAAACCCGAACGCACAATGAAAGTAAACGGTGGCTCTGCGCCGCTGCTTGGTGG
GAACCCTGCTTTTATAATGGGGGGCACTATCACCCGACCCCGATCCCTTTGGGGTGGTTG
GTTTTGAGGTAGAGCGAGTATGATGGGACCCGAAAGATGGTGAACTATTCCTGAGTAGGG
CGAAGCCAGAGGAAACTCTGGTGGAGGCTCGTAGCGATGCTGACGTGCAAATCGCTCGTC
GAACTTGGGTATAGGGGCGAAAGACTAATCGAACCATCTAGTAGCTGGTTCCCTCCGAAG
TTTCCCTCAGGATAGCGGGAGCAAATGCAAGTCAGTATCATCCGGTAAAGCGAATGATTA
GAGGCCTCGGGGGAGTAACTCCTTCGACCTATTCTCAAACTTTAAATAGGTGAGGAGCCG
GCAACGTTGCTTAAAACAATGAACGCCGTGCGGACAATGATTGCTCTCAGTGGGCCGATT
TTGGTAAGCAGAACTGGCGATGCGGGATGAACCGAAAGCGACGCTAAGGTGCCAAACTCC
CAGCTCATTCAGACACCAGAAAAGGTGTTGATTCATACAGACAGCAGGACGGTGGCCATG
GAAGTCGGAATCCGCTAAGGAGTGTGTAACAACTCACCTGCCGAATGGATCAGCCCTGAA
AATGGATGGCGCTTAAGCGGGGGACCGAGGCGTCGCCGTTGGGGTAAATACGAAGCCCCA
ATGAGTAGGAGGGCGTGGGGGTTTTGAAACACGGTACACTCGCATGGAAGCCTTGGGCGT

AAGCTCGGGTGGAGCGACCCCTAGTGTAGATCTTGGTGGTAGTAGCAAATATTCAAGTGA
GAACCTTGAAGACTGAAGTGGGGAAGGGTTCCACGCGAACAGTGCTTGGACGTGGGTTAG
TCGGTCCTAAGAGATGGGGAAACTCCGTATTCAAGTGCGCTTTTGTAGCGCAGTCCATCG
AAAGGGAAGCGGGTTAACATTCCCGCACCAGGATGTGGAAACTGTGGTTTACCGCAAACT
GCGGCAACGCAACTGAACCTGGGGACGCCCGTAGGAGCTCCGAGCAGGGATCTCTTTCCT
GTTTAACGCGCCCAACCCCCTGGAATCGGTTTGTCCGGAGATAGGGGGCGAAAGGCCGGA
AAAGCACCGCACGTCTTGTGGTGTTCGGAACGCTTGTAATGGCCCTTGAAAACCCAGGGG
AAAGCAGCCGTCTTACACTTGTGTGAGACGAGGAATTCTCACACCTGGCCGTACCCATCA
CCGCATCAGGTCCCCAAGGTGAACAGCCTCTAGTCAATAGAATAATGTAGATAAGGGAAG
TCGGCAAAATAGATCCGTAACTTTGGGAAAAGGATTGGCTCTAATGTTCGGGCTTGTCGG
GACGCGCGGGCGTAGATGCCAATGAGCGGCAGACGTCTAGCCTCTTTGGAGGCTGGATTA
ACGCTGCTTGTGGGAAGAAACCGTGTCGATGGAGGCAGGAACGCTTATCGATCTTGTCTC
TTCCCGCTTGCGATCAACGAACGGATTAGAACTGGTACGGACAAGGGGAATCCGACTGTT
TAATTAAAACAAAGCATTGCGACGGCCGGAAACGGTGTTGGCGCAATGTGATTTCTGCCC
AGTGCTCTGAATGTCAAAGTGAAGAAATTCAACCAAGCGCGGGTAAACGGCGGGAGTAAC
TATGACTCTCTTAGACAGATCAAGACACCTTCAATAAATTGGTTCATTATGTCGGGTGGT
CCGTCGTCCGGGAAGCGACCAAAGCAAAACCCCGTAGACTGTGATGTGCATCTGTCAGAC
GAAACTGACGTCCGTGTTCATCAAGCCTTAGCTATCTTTCAACGCCGGCTTCGAGACGTG
GCGTCTCTTAAGGTGTTTGAAGTGGCTCTCTCTGTGGGAACTACTGATGCACTGGTAGAG
ATGGGCACTATCACTTGCTGGAGTATTGATGCTGTTGGGAAAGGTTTCTCAAGGAACAGC
AGCGGTCGCTGGCAATTTGACCTCACTGGCAAGATTAATGGCAAGCCTCTAATATCCACC
ACAAAGGATCGTTGGAGGCCGGTTGTGACGGCTGTGCCCTACTACCTTTCGTGCGCCATC
GATGGGGACAAGACGACGCTGGAGGAAAAGCAGTGGACTTCATCACACATGTGTCACCAT
GAATGGTGTGTGAACGTGGAGCACATTGTTTATGAAAGCCTCGACGTCAACAAGTCTCGG
AACTGGTGCCAGGGAAAAAACGCATGCCTCCATCTGCCTGTTTGCATTCGAACCGGCCGC
CGTGTGCAGGAGCTCAGGAAGAGCAATGGGGTTCGTGCAATGGACCTCCTCCTGCAGAAA
TTGGCTGGATTGGAGATTGAGGTTGAGGAGCCCGCTGCAGCTGTACTCCGAGAGTCTGTC
GAACGGGAGGACGCCTAGTGAGTGACTCGTGACACGGGTTGGCTGCGCATGATGCGTCAA
CTACATGGACATCTTTTGAGAGAGATAGGCGGGCTGAAAAGGTTATAGCCGTCTAGTCTC
GCTCTCTTTCATTGAGCGGGGCGACACACCTGGTTGCGGGAACCCTGCCCACCTTTCTAT
CAAGAAGTGGGAACCAGTCCATTCCTACTCCTGCTTGTGAGGCATATAGTAACGTAGCAG
CAGGAGGATCTCGGCGAGAGGCCGAACCCGAAAGGAATAACGTGGGCTGGGAGGCGATCC
GCAGGCGAGTCCGTAACTATATGATTATCGACGGGCCGTCTCAACGACTGGTAAGGTGTG
GGTTTGTCCTCCTCCTCACGTGGGGACAGGCTTAAGGTACAGTCTAATCCCACCCGCCAA
GGGTGCTTGGGTGTAATCCCTCTTCCATAGAGGGCACTAGCCCAGGGGCTGGGACGCTGG
GAGGAAATGCCCCGTCCCTATAGCCGGTAGAAATGAAGGTAGCCAAATGCCTCGTCATCT

AAACTAATTGGATGAGATAGTCTGAGTGAAAAGCTTATGCAGGCTAGTACCGGAAAAAAA
AAAAAAGAGGGGCTCTGCCACTCTGACGGTGCGACACTCCTTGTACAGGGAAACCCTCCG
CATGGAAAGGCACGTGGTACCACCCTCGCGGGGAAACTCGCGAGCGGGATCCCGGGTAAT
GACCGGAGCCGATGGTAAAAACTCACGTGACCGATGGGGTGACCCTGTGGCGAGACGCTA
AGTTGTGGGACATCACCCTCTCACAATACGCATCCGTCGCAGAGACTGAACGGGAGTGGG
CTCCGTTGACGGCCGGCCGGCCAGCCAGCCGTCTTCCAGGGGCTTAAGATACAGTCCGGG
CCTCAGGGATGGAAAAACCTGGGGAACTACCAGAATTAGTGACGCGCATGAATGGATTAA
CGAGATTCCCACTGTCCCTATCTACTATCTAGCGAAACCACAGCCAAGGGAACGGGCTTG
GAAGAATCAGCGGGGAAAGAAGACCCTGTTGAGCTTGACTCTAGTCCGACTTTGTGAGAT
GAGTTGGGAGGTGTAGTCTAAGAGGGAGGGGCAACTCGACCGTGAAATACCTCTACTCCC
AAGCTTGTTTTACTTACTCCGGCGAGCAGGGATGGGTGGTAACACCCTTTTTCTAGGATT
AACACCTCTTCGGGGGTGGACCTGCGCGGAGGACATTGTCAGGTGGGGAGTTTGGCTGGG
GCGGCACATCTGTTAAAAGATAACGCAGGTGTCCTAAGATGAGCTCAGTGAGAACAGAAA
TCTCACGTAGAACAAAAGGGTAAAAGCTCATTTGATTTTGATTTTCAGTACGAATACAAA
CTGTGAAAGCATGGCCTATCGATCCTTTTGACCTTCAAGGTTTGAAGCAAGAGGTGTCAG
AAAAGTTAACACGCGTTGTGGTAGCTTTTAAGAGTGGACTCAAAGTGTTCCGCTAGTCCC
CCCTCGGACCGCCACTTACAAAAGGCCTGGGGATGGGGGCGACACTGTCAAATTGCGGGG
ACTCCCTAAAGCCTGTCTCTACCAAGCCAGCGCGGCGAAAGCCCCTGGTGGCCGGGGTAA
TGACCTAGGGTATGGTAAAAGCGAGACAGGATGGAACAATGGGCAATCCGCAGCCAAGTC
CTAAGGCTTTTTGAAAAGCGGCTATGGATGCAGTTCAACGACTAGATGGCAGTGGGCTGA
GAGAGCCCACCGCGGCACCCTTTCGGCTTAAGATATAGTCTTGCCCACGTGAAAGCGTGC
CTGCAGATGAACGCCACCATGTATGGCCGGAGCTGCAGGAGACCGCACTTGATGTGCAGT
CGGGAATCGACAGTACCACAGGGATGAACAAAAACACACAATTGTGGCCATCATACAGTC
GTATGATATTATCGTCCCAGGCGGGTGCGCCTCTCTCACCCGTCTAGTCCGGCCTTGGTG
CGCAAAAAATGCTTGCGTGGATGCGCCTTGGTTAAGGAAAATGCTCCTAGTAGTTTTACA
AGGACACTTCTTGTAGGGCAACACATCTGGATGCGAAGAACTCCCGGAAAAGACAACGGT
ACCAAGTGACTGGGGTAACGCCCCATTCATGAAGGCAAATAATCAAAAAGAGGTGCCGGG
TAAGAACCCGTTGTATAGGGACAATCCGCAGCTGACTTGGCTCCCTGCCCATCCATGGGC
ATAGCTTAGCGGTTCAACGCTCGGTAAGATGTGGGTTGGCGCGCTGTCCGACAGCGCACT
GGCTTAAGGTACGGGCTACTCCCACCGGAGACGGTGTCTTTGTCTACCTCGGGTAACACG
CCTCGGGAGAAGCAAAGAGGGTGCAATGGAAAAAAACTGGTTGCGGGGAAATCCCTACCA
GAGTCACCTGGTACAAAGGAACTGGCTTGTGGCAGCCAAGCGTTCATAGCGACGTTGCTT
TTTGATCCTTCGATGTCGGCTCTTCCTATCATTGCGAAGCAGAATTCGCCAAGTGTTGGA
TTGTTCACCCACTAATAGGGAACGTGAGCTGGGTAGCCACTTTATGCCCAACAGCTGGAT
AGCTGGGCTTCTGTCTTAGGCCGGGACTTCTCACGACCCGTGCCTTGGTCACCCAAAAAA
GGGCGACCGCAGAGCTCAGGATGGAGCTTCTGGCTTAACTAGTAGAGGACAGATAGAGAT

CGTCAAATTGCGGGAAACCCCTTAGAGCCTCGAGCTACCAAGTGCGTGTCGAAAGGCATA
GACGCAGCCAGCGATGGCTGTCGCATGGCTTGATTCCATCAAGATAGGTAAAAATGCTCG
AGGATTGGGCAATCCGCAGCCAAGCCCCTAAGGTCCCCCCGCGGGCTATGGGGAAGGTTC
AACGACTGTAAGGCGTACCGTGCAAGCGGAAGATACAGTCTAGCCCCGCTGGAAAGCAGC
GCCAAAGTAAGTAGGTTGCAGTGGGTCTGTCCTC
>SRR9587925_SRR9587935_SRR9587940_SRR9587945_SRR9587957
GCCTGTTTTGATATGTGTTTTGACAGTCATACGGGTCATGGCCGAGGGATGGTGGGTGAG
CTTCATAGCGCAACCCCCCGTCAGGAGCATGTTTTCATAGCATTTTGAGCATACTAGGGT
GGTCTCCCATGTAGCTCGCTCTCTCGGCTCAGGACCTTCCGTATGGGGCACTTTTTGACG
AGCTCCTTTGGGGAGGAGCATGTTCTGGCATATGAAGCGTTGGTCACGAGCTTCGTAGCC
CGACACATACCAAATCAGTCTTCATATGTCTTTCTGGCGGGGGTCGGTCGTATGACCACG
CCCTTACCTTGAGCATGTTTTTCGTATGATACCTGACGGGCAGCGCCGCTCTTCAGTCGT
CGGCCGTACCTCGTATGCGCGTTTTCTGATTCGAACGTGTGACATGCAAGCGCATCATAT
CAGAAGCATGTTTTTTCATATGAGCATGTTGGTAGGCAGGCTCTTCAGTCGTCTACCCAT
AAATGCACTTTGAAAGACGTTTTTTGGTACTGAACGGCTTTGTGGTCAGACGTTCTCGTG
AGCTTATTCTCGCCCTCTGATACGCTGGATGCTGCGGTATTACCCACAGCAGACATCCAG
GAGCTCCCTCTGCGGATAAGCATCTGTAGAAGGGGGTTTCTCTCGGTGACGCAATGGCTG
GGAAAACGGAGTGCTATCCAGAAGGGTGAAACCTCTTGGGGATGTACTCCCATACCAGCA
TAATATGCTGAGAGGAATTGACTCAATGAGTGAGGTCCTATACTGCAACCTCTTTTGGAT
ATGTAGGCGGTCTGTACCCAGACTGTCCTACTGGCTGATTTCCCTTGGTCCCACGCGTCG
CTGGCTGAATGGGCTAGTCCTTATGAAGTGCCGCGTTCATCAAAAGAACCGGTCTTCATA
CAGCGACCTCATTCTATACTGCCAGCTGGATGGGCTGGTCCTTATGGAGCGCCGCATTCT
TCGAAGAATCGGTTTCTATAGAGCGACCCCATTTACATATAAGGAACCGTTACATAGACT
TGCCATTTGGCTGATTGCTCTGACAGTCGCTGCCTCTTCGGAGTTCTGTGTAAACGTGGC
GCCATTTTTGAGTGGGCCAGTCCTTATGGAGTGTCCGTGTCTTCTATGACAGGGTCCCCC
ATACAGCGACCTCACTCAGAACCGTATGGGCACGATGGTGAAAGACCATGCAGATAGTCC
GCCTTCCAAAGGGGGAAGTAGTATATGGGGAATGCTGGCGAAGCATGAGGTACATGGTTC
ATGGACCCGTTGGGCCATGCCTGTGCCATCTTCTCGAGTCAAACCGGTCCGTGTCGCTTC
CTTATCTCTTCCTTTCGCGACGACAAGCTCTTGAGCTGGTCGTGGGTCAGGACGCTGTGG
TATCTGTTTGTCTGTAGTGTGGGCTTCCACCAGAACAGCGTATCAGCACACGTCGTCTGG
GCATATGATGTTTTTGCTTTGTGGTCCCGGGTTTGGTAGTATGGTTCCATGTCGGTCCGC
TCACCTCCCAACCATAATGTCTTTCACATCATCTTCACAGAACGTCTTGTCAGCATACCA
TTCTGTATAGAACAGTGTATCAGCACACGTCGTCTGAGCATAGGGTATTTTTGCTTTGTG
GTCCCGGGTTTGGTAGTATGGTTCCATGTCGGTCCGCTCACCTCCCAACCATAATGTCAT
TTCTATCATCTGCACAGTACGTCTTGTCAGAATACCATTCTCTATGGCAGTCGATAGACT
AGGCCTTATGCCGCAGCAGTCTCTGATCACTGTGCGAGTGCATATATCTCTCAATGTGTT

CATGGCTAGCATGGGTCTTGAGTGAGATGTGCCAGAGAGATGCGCCACCAGTCTATGGAA
AGGTGCTTTTTGTATGAGAAGAGTACATGTATGAAGAGCATGTTCTCTCCTCTTATGATG
GTACCGCGGGTACGGGCTGCTCGCATTATGCTTGGCAGGTGTCCACTGTGCATGCAGGAT
TGCTGAACCCACTTGTATGGCCATGAGAATGGTCATTGGGTGAGGTTCACTTCTCCATCG
CTTTGTGTGTGTTAGATGTTTGGGTGGCTTCGGCCGCATGGAAATTCTCGGTCTGTAACT
CATCAGGGAGGGTTTCCATGTGAGTATTCTCCAGTCGACCACACACCCTTGGGCGTCCCG
CTAGACAAAATGTAACTCTCGATCCGACCTAGCCTGTGGAGTGTCTTTCCGTATGGCGTG
AAAGGGCCCACACTGGATGTTTTCCCTTTTGGACACCAGTCTGTTAGTATGTATATATAG
ATCATCTACAGGGAGCAAAGGACAGGCATGCAAGTGCCACAAAGGCAGGGAGGCCTTCCT
GCTACTGTGTGTCCTTCTTGACGATCACGGGGGAGTATATAGAAACATTTGAAGGAGATC
CTTTTCCACAAGGCGGTGCCCTCGCGCAAAGGTGTTGCTCTGCACGCATCTGTATGATGA
TAGTTGCTCGCTCCTATGCGTCCACTTGTGAAAATACCTTCTCTCTTTGTCTGTTTCTAT
GACGCCCCTTCTGCTTTGTCATAGAGCATACTCAGGTACAGAAAGTGCCGTGACTTCCTT
GGCCTGCTGTCAAAGAAATAGCCTGGTGCTATAGAGGACTCTTCCTGGGCGCCGCCTGGT
GAAGAGGTCCTATCAGGCTCTGGAGTGTGCGTATGCCTATGTTCATCATAGCTCATGTGC
ACGTGCGGAAAAGCATCGACGTGGAATGGGGAAGAACACCATTCTGAATGAGCCAGTTCT
TCCTCTGCTCCCCTTGCTTGACGCTGTATGGGAATCGCTTTGCCTTGGGGGTACACCTTC
CATTCCATGACAAAGCATTCGATCTATGTCCCGAGGGGTGATTCTACATGAAGAACCCGA
AACTTGTCTGAGTGAGAGGCGAAGGTAGTCCTTCTGGATCCGCCCTCACACTGTAGTCGG
CACAGTCCCTTGTTTAGGGACTGCGCAGCTCAATCTTCGACTGGAGAGGAAAAAAACAAA
ACACAAGTATGACACGCACATACGTGTTTTCTAGGCCATTGACCTGTTGGCTGCCTCGCG
GTGGTTGGCATGATGCCCCAAGTGGGGGGGGGGAAATCAATCAGTATGGTTGACAAAAT
CCAGTGGGGAAAAAGGCGCGCTGTCGCACACCCTTGGTCACGGCCTCAAAAAGTCGACGG
CCCTCACTGGCCCACCCCTGCGACTCCTGTTCGCTTTTGACAAGACACACCTGCAATATG
GAGGAAATGCCTGAGCAGCTGTGCCGCAGCAGCTGTGGACGCGTAGCGATATGCATCCCT
ATTGTTTGCGACTTGGTACAAAGAACGCGTGCCGGGAGACGGATTGCCCTTTGACTTTTC
CACCCGGCCCGCGGCTCTGCCCTTTACGTACCCTTATTTACCTGGTTGATCCTGCCAGTA
GTCATATGCTTGTCTCAAAGACTAAGCCATGCATGTCTAAGTATAAACACTTTTATACTC
GTGAAACTGCGAATGGCTCATTAAAACAGTTATAGTTCCTTTGGGAACAACACTACTTGG
ATAACCGTAGTAATTCTAGAGCTAATACATGCCTCAACGCCCGACTCACGAAGGGTGGTA
TTTATTGGATAAAAAACCATCGTGCTCTCTCCAGAGCGCTTTGAGATGATTCACAATAAC
TTGTCGGATCGCATGGCCTCGTGCTGGCGACTCCTCATTCAAATTTCTGCCCTATCAACT
TTCGATGGTAGAGTATTGGTCTACCATGGTGTCGACGGGTGACGGGGAATTAGGGTTCGA
TTCCGGAGAGGGAGCCTGAGAAACGGCTACCACATCCAAGGAAGGCAGCAGGCGCGCAAA
TTACCCAATCCCGACTCGGGGAGGTAGTGACAAAAAATAACAATAGGGGGCCCTTTGGGT
CTTCTAATTGGAATGAGAACAATTTAAATCCCTTATCGAGGATCCATTGGAGGGCAAGTC

TGGTGAACGTATGCATCAAAGCCCTCGATAAAAAAAAACAGCGCCAGATGTCTCTCTCTG
GAAGCGCACCATGCATGGCTTTGTTCCTCTCTGCATGGCCGTGTGGAAGGCGAGTAGGCA
AACGATGGGAAACCAGGCTTGGAATAGTAACCAACCACAGAGCCCCCATCATACACTTCT
AAATTGCCGGGGAGTGCCCGCGAGGCCTTGACTACCGCGGTCCGCCGAGAGGTGAGAGCG
AGCAATCGTTCAGGACGCAGCAGTGTACGTAGTGGTACATGGATGGTAAAAACGTCAAGG
ATAGGGTTGATCCGCAGGGAAGCGCGAGTGCCGCCATGGTCGGTGGGCAAAGGCGAACCT
TCAGAGACTCTAATGGAGTGGGCGCAAGCTTAAGGGAGAGTCCAATCCACTGGGAAACCA
GGTCCACAGCAGTAACGGGGTGATGCTCGCCCCAGAGGACTGTGGAAGGCACCACCACCC
CAAGAGGGGGTGGGTGTTGTCGGGTAGCAAATGGCCAGCAGCCGCGGTAATTCCAGCTCC
AATAGCGTATATTAAAGTTGTTGCAGTTAAAACGCTCGTAGTCGGACCTCGGAGCGCCCC
CACGGGCCGGGCTTTTGCCTTGTTGCCAGTGGGAGTTTGTCTCCTTTTGTCGGGACGCGG
TGTGCAGGGCCTTTACTGTCTCTGCGCGTTTACCTGCGGGCCGACCGTTTACTGTGAAGA
AAGTAAAGTGTTCAAAGCAGGCCAATGCCTTGAATATGTGAGCATGGAATAATAGAATAG
GACTTGGGCTCTATTTTGTTGGTTTCCAGTGACCAAGTAATGATTAATAGGGACGGTTGG
GGGCATTCGTATTTCATTGTCAGAGGTGAAATTCTTGGATTGATGGAAGACGTACAACTG
CGAAAGCATCTGCCATGGATGTTTTCATTGATCAAGAACGAAAGTTAGGGGATCGAAGAC
GATCAGATACCGTCGTAGTCTTAACCATAAACGATGCCGACTGGGGATTGGCGGGGGCAC
TATTATATGACTCTGTCAGCACCCTGAGGGAAACCAAAGTCTTTGGGTTCTGGGGGGAGT
ATGGTCGCAAGGCTGAAACTTAAAGGAATTGACGGAAGGGCACCACCAGGAGTGGAGCCT
GCGGCTTAATTTGACTCAACACGGGAAAACTTACCAGGTCCGGACAGAAGAATGATTGAC
AGACTGAAAGCTCTTTCTTGATTTTTTGGTTGGTGGTGCATGGCCGTTCTTAGTTGGTGG
AGTGATTTGTCTGGTTAATTCCGTTAACGAACGAGACCTCGTCCTGCTAAATAGGTGCGC
GCATGCCAGCACTGCGTTCTTACCTTCTTAGAGGGACTATGCGCGTCTAGCGTATGGAAG
ATTGAGGCAATAACAGGTCTGTGATGCCCTTAGATGTTCTGGGCCGCACGCGCGCTACAC
TGATGCATTCAACGAGTTTCACTTGATAGTCCTGGGTCGGAAGGCTCGGGTAATCTTTTG
AAAGTGCATCGTGCTGGGGATAGATCATTGCAATTATTGATCTTCAACGAGGAATTCCTT
GTAGGTGCAGGTCATCAGCCTGCGCCGAATACGTCCCTGCCCTTTGTACACACCGCCCGT
CGCTCCTACCGATTGAATGGTCCGATGAAATGTCCGGATCGCGGCTAGTGTGATGGACTC
GTCCAGAGCGGTAGCCGTGAGAAGCTCATTAAATCTTATCATTTAGAGGAAGGAGAAGTC
GTAACAAGGTTTCCGTATTTTGTGGGTGGATACCGATTCAATAACATCCGCCGTTTTTCT
CAATGTACTATATCCCTAGTCTGCTTGCCTAGTTCTTGGAGTCTCGTGCGATGTGTGCAA
AGATGGACGCCATATCGGAGACAGGGGGAGGTGCCATTGTGATGATGACGATGTTGTTGG
GTGGTCCTTCCGTAGGAAGACGTACACGAAGCCTTTGCGGCCCGAAAGGGTGCACAGTGC
GCGACTATAAACACTGATTCTGTGGTCAACGCAAGCAGCGGTTCTGTGGAAGAACAGCTG
CGACTTTCTCAAATTGCGGGAACACCCTAAAGTCAACAAGCACCGCGGCTTGACGTGAAA
AGCGTCAGGCGCAGCAGTCGGGGTAGCGCCCTGGCGGATGGTAATAGCCTTGTTGAATGG

CACAATGGGCAATCCGCAGCCAAGCCCCTAAGGTCCCCCGCTGGGCTATGGGGAAGGTTC
ACAGACTGTAAGGGAGGGGGTTTCTTCCATGAAAGAAGCTTAAGAGACAGTCGGTCCCCT
GCGAGAGCAGTGTCGCATGTGAGGAAGGCACCCATGAAAAGTGGGTGCTGGAGAGCCATG
TGAGCCCTCTATGAGGGTGGGAAGTTTTCCGATACAATGAGAAACAAACACGTTCCGTAG
GTGAACCTGCGGAAGGATCATTCACAAACTATTGACACAAACACACGCGAACCAAATCGT
CCACACAGGTGCCGATGAAAGAGAGAATCTGATTGTCGCCTTTTGGGGTATACAAGCACT
CTTTTCCATCGCCTCTGTGCAGGGCGTAACTTCCCATTGAGAGGATGTGAGGGCACCACA
GGAAGCTTTTCCACAGGAAGTCGCCATCCTTCTCCCTCCACGACGGCGGCGCTTCACAGG
CTTGGCAGTTTTTTTTTTTTTTATTGCCTTCCAGGGAGGATGCCGCCAATGGAGCCACAT
AGACATTTACATCATCATATGCCCGTTTTTCTTTCTTAACCGCTTGCCACAGTTTCTCCT
GTGAGGAGCTTGTGGGAAGACCGTCTCCATACAATAACAAAAGATACAACTCTTAGCGGT
GGATATCTTGGTTCTCGCAACGATGAAGAACGCAGCCAACTGCGATACCTAGTGTGAATT
GCAGGACTCCGTGAATCATTAAGTCTTTGAACGCAAGTTGCGCTCATCTCCATGAGGAGG
TGAGCATGCCTGTTTGAGTGTCGATGGGAAAAACAATACAATAACAACAATCGGCTCTCT
TCCCTTCCTGCCTTCCCCGGCAAAAAGGTAGTGCATGCCGGAGATGGGCCTGGTCGCCCC
TTCATGAAAGAGGCGTCGTCCGAAACACGTGTCGTTGCCCGCTGGAGGTCATACAGCATG
AGTTTCAAAACACCATGGCTCTCAAGGCAGAGTAAATCGTCTTGAGACGGTCTGCATTGA
TTCACTTTCAACCCAGGTGGCGTTGGTTACTTGCCATAAGCCTCTCTGGGGTTTCTTTGC
TTGGACCTCTCTTGTCAGTTCTTCTGACGTAGAGACAGGTGCGGTCTCATGGGCGTTGCC
CTCTGGAACGTGCTAAAAAAAACAAAACAAACAACAACAACAACAACCGTATCACACGTC
AAGAGGGTCGGATCGTACTTGACGGCGGGAGATCCTTTGTTTCTCGCTTGTCCTGAGCGT
CTGCCAAGCCTTCTTTGACGCGTGTGTGTTTGTTCCTTCCATAAACCCGATCTCACATCA
GGTAGGACCACCCGCTGAATTTAAGCATATAATTAAGCGGAGGAAAAGAAACTAACAAGG
ATTCCCTCAGTAGCGGCGAGCGAAGCGGGAAGAGCTCAACATGAAAACCGTAGGGGGTTC
GCGCTCACCTTCGGGTTGTATAGTAAAGTGGTGTCGCCTGCGTCGCGCCGGGTATAAATC
CCGTTTGAAATGGGGTGTCGTAGAGGGTTATAACCCCGTCTGTGACCCGGTGTGCGATGC
GTTACGGTGACCTTACGATGAGTCGGGTTGCTTGGGCATGCAGCCCTAAGTGGGTGGTAA
ACTTCATCTAAAGCTAAATACTAGCGAGAGACCGATAGCGTACAAGTACCGTAAGGGAAA
GATGCAAAGAACTTTGAAAAGAGAGTTAAACAGGGCTTGAAACCGTTAAGAGTGAAGCGA
AGGAGGTCAGTGATCATGGGGCCAGATCCCCACGTGCCCGTCATAGGGGCACCGTAGCGT
TCTGGTTCCCGGGGCAGCGCCGGCCGGGGGCGGGTGTCAATAGGGAGGTGGTTGAGGGAT
TAACTTTCCGCCTGTACTCCTTTCTAGCTTGCACCCCGGCCGAGGGATCCGCGACTTGTC
GTGTGAGGGCGCTGTCAAAATGGCCTTCTCCACCCCGTCTTGAAACACGGACCAAGGAGT
CTAACATACTCGCGAGTGTTTGGGTGGTTAAAACCCGAACGCACAATGAAAGTAAACGGT
GGCTCTGCGCTGCTGCTTGGTGGGAACCCTGCTCTTATAACGGGGGGCACCATCACCCGA
CCCCGATCCCTTTGGGGTGGTTGGTTTTGAGGCAGAGCGAGTATGATGGGACCCGAAAGA

TGGTGAACTATTCCTGAGTAGGGCGAAGCCAGAGGAAACTCTGGTGGAGGCTCGTAGCGA

TGCTGACGTGCAAATCGCTCGTCGAACTTGGGTATAGGGGCGAAAGACTAATCGAACCAT

CTAGTAGCTGGTTCCCTCCGAAGTTTCCCTCAGGATAGCGGGAGCAAATGCAAGTCAGTA

TCATCCGGTAAAGCGAATGATTAGAGGCCTCGGGGGAGTAACTCCTTCGACCTATTCTCA

AACTTTAAATAGGTGAGGAGCCGGCAACGTTGCTTAAAACAATGAACGCCGTGCGGACAA

TGATTGCTCTCAGTGGGCCGATTTTGGTAAGCAGAACTGGCGATGCGGGATGAACCGAAA

GCGACGCTAAGGTGCCAAACTCCCAGCTCATTCAGACACCAGAAAAGGTGTTGATTCATA

CAGACAGCAGGACGGTGGCCATGGAAGTCGGAATCCGCTAAGGAGTGTGTAACAACTCAC

CTGCCGAATGGATCAGCCCTGAAAATGGATGGCGCTTAAGCGGGGGACCGAGGCGTCGCC

GTTGGGGTAAATACGAAGCCTCAATGAGTAGGAGGGCGTGGGGGTTTTGAAACACGGTAC

ACACGCATGGAAGCCTTGGGCGTAAGCTCGGGTGGAGCGACCCCTAGTGTAGATCTTGGT

GGTAGTAGCAAATATTCAAGTGAGAACCTTGAAGACTGAAGTGGGGAAGGGTTCCACGCG

AACAGTGCTTGGACGTGGGTTAGTCGGTCCTAAGAGATGGGGAAACTCCGTATTCAAGTG

CGCTCTTCCAGCGCAGTCCATCGAAAGGGAAGCGGGTTAACATTCCCGCACCAGGATGTG

GAAACTGTGGTTACCGCAAACTGCGGCAACGCAACTGAACTTGGGGACGCCCGTAGGAGC

TCCGAGCAGGGATCTCTTTCCTGTTTAACGCGCCCAACCCCCTGGAATCGGTTTGTCCGG

AGATAGGGGGCGAAAGGCCGGAAAAGCACCGCACGTCTTGTGGTGTTCGGAACGCTTGTA

ATGGCCCTTGAAAACCCAGGGGAAAGCAGCCGTCTTACACTTTTTTGTGGGACGAGGAAT

TCTCACACCTGGCCGTACCCATCACCGCATCAGGTCCCCAAGGTGAACAGCCTCTAGTCA

ATAGAATAATGTAGATAAGGGAAGTCGGCAAAATAGATCCGTAACTTTGGGAAAAGGATT

GGCTCTAATGTTCGGGCTTGCTGGGACGCGCGGGCGTAGACGCCAATGAGCGGCAGACGT

CTAGCCTCCTCGGAGGCTGGATTGACGCTGCTTGTGGGAAGAAACCGTGTCGATGGAGGC

AGGAGCGCTTGCCGTTCTTGTCTCTTCCTGCTTGCGATCAACGAACGGATTAGAACTGGT

ACGGACAAGGGGAATCCGACTGTTTAATTAAAACAAAGCATTGCGACGGCCGGAAACGGT

GTTGGCGCAATGTGATTTCTGCCCAGTGCTCTGAATGTCAAAGTGAAGAAATTCAACCAA

GCGCGGGTAAACGGCGGGAGTAACTATGACTCTCTTAGACATGGACATCTTTTGAGAGAG

ATAGGCGGGCTGAAAAGGTTATAGCCGTCTAGTCTCGCTCTTTCATGAGCGGGGCGACAC

ACCTGGTTGCGGGAACCCTGCCCACCTCTCTATCAGAAGTGGGAACCAGTCCATTCCTAC

TCCTGCTTGTGAGGCATATAGTAAAAGTAGCAGCAGGAGGATCTCGGCGAGAGGCCGAAC

CCGAAAGGAATAACGTGGGCTGGGAGGCAATCCGCAGGCGAGTCCGTAACTATATGATTA

TCGACGGGCCGTCTCAACGACTGGTAAGGTGTGGGTTTGTCCTCACGCGGGACAGGCTTA

AGGTACAGTCTAATCCCACCCGCCAAGGGTGCTTGGGTGTAATCCCTCTTACATAGAGGG

CACTAGCCCAGGGGCTGGGACGCTGGGAGGAAATGCCCCGTCCCTATAGCCGGTAGAAAT

GAAGGTAGCCAAATGCCTCGTCATCTAAACTAATTGGATGAGATAGTCTGAGTGAAAAGC

TTATGCAGGCTAGTACCGGGAAAAAAAAGAGGGGCTCTGCCACTCTGACTGACGGTGCGA

CACTCCTTGTACAGGGAAACCCTCCGCATGGAAAGGCACGTGGTACCACCCTCGCGGGGA

AACTCGCGAGCGGGATCCCGGGTAATGACCGGAGCCGATGGTAAAAACCCACGTGACCGA
TGGGGTGACCCTGTGGCGAGGCGCTAAGTTGTGGGACATCACCCTCTCACAATACGCATC
CGTCGCAGAGACTGAACGGGAGTGGGCTCCGTCGACGGCCGGCCAGCCAGCCGTCTTCCA
GGGGCTTAAGATACAGTCCGGGCCTCAGGGATGGAAAAACCTGGGGAACTACCAGAATTA
GTGACGCGCATGAATGGATTAACGAGATTCCCACTGTCCCTATCTACTATCTAGCGAAAC
CACAGCCAAGGGAACGGGCTTGGAAGAATCAGCGGGGAAAGAAGACCCTGTTGAGCTTGA
CTCTAGTCCGACTTTGTGAGATGAGTTGGGAGGTGTAGTCTAAGAGGGAGGGGCAACTCG
ACCGTGAAATACCTCTACTCCCAAGCTTGTTTTACTTACTCCGGCGAGCAGGGATGGGTG
GTAACACCCTTTTTCTAGGATTAACACCTCTTTGGGGGTGGACCTGCGCGGAGGACATTG
TCAGGTGGGGAGTTTGGCTGGGGCGGCACATCTGTTAAAAGATAACGCAGGTGTCCTAAG
ATGAGCTCAGTGAGAACAGAAATCTCACGTAGAACAAAAGGGTAAAAGCTCATTTGATTT
TGATTTTCAGTACGAATACAAACTGTGAAAGCATGGCCTATCGATCCTTTTGACCTTCAA
GGTTTGAAGCAAGAGGTGTCAGAAAAGTTAGCACGCGTTGTGGTAGCTTTTAAGAGTGGA
CTCAAAGTGTTCCGCTAGTCCCCCCTCGGACCGCCACTTACAAAAAGGACAGAGGAGGGG
GCGACACTGTCAAATTGCGGGGACTCCCTAAAGCCTGTCTCTACCAAGCCAGGCGGCGAA
AGCTCCTGGTGGCCGGGGTAATGACCTAGGGTATGGTAAAAGCGAGACAGGATGAGAACA
ATGGGCAATCCGCAGCCAAGTCCTACGGCTCTTTTCTTGGAGAAGAGCTATGGATGCAGT
TCAACGACTAGATGGCAGTGGGCTGAGGGAGCCTACGGCGGCACCCTTTCGGCTTAAGAT
ATAGTCTTGCCCACGTGAAAGCGTGCCTGCAGATGAACGCCACCATGTATGGCAGGAGCT
GCAGGAGGCCGCTAAACATACCAGCGGCTGGGAACATAACAGTACCACAGGGATGAACAA
AAACACACAATTGTTGTGGCAATCATCATACAGTCGTATGATAGTATCGTCCCAGGCGGG
TGCGCCTCTCTCACCCGTCTAGTCCGGCCTTGGTGCGCAAAAAATGCTTGCGTGGATGCG
CCTTGGTTAAGGAAAATGCTCCTCGTAGTTTTACAAGGACATTTCTTGTAGGGCAACACA
TCTGGATGCGAAGAACTCCCGGAAAGACAACGGTACCAAGTGCTTGGGGTAACGCCCCAT
TCATGAAGGCAATGAATAAAGAGGTGCCGGGTAATAATCTGTTGTATAGGGACAATCCGC
AGCTGACTTGGCTCCCTGCCCATCGGTGGGCATAGCTTAGCGGTTCAACGCTCGGTAAGA
TGTGGGTTGGCGCGCTGTCTGACAGCGCACTGGCTTAAGGTACGGGCTACTCCCACCGGA
GACGGTGTCTTTGTCTACCTCGGGTAACACGCCTCGGGAGAAGCAAAGAGGGTGCAAATA
AAAACTGGTAGCGGGGAAATCCCTACCAGAGTCACCTGGTACAAAGGAACTGGCTTGTGG
CAGCCAAGCGTTCATAGCGACGTTGCTTTTTGATCCTTCGATGTCGGCTCTTCCTATCAT
TGCGAAGCAGAATTCGCCAAGTGTTGGATTGTTCACCCACTAATAGGGAACGTGAGCTGG
GTAGGCATTTTATGCCCTATGCCCAACAGCTGGATAGCTGGGCTTCTGTCCTAGGCCGGG
ACTTCTCACGACCCGTGCCTTGGTCACCCAAAAAAGGGCGACCACAGAGCTCAGGATGGA
TCTTCTGGCTCAACTAGTAGAGGACAGGTAGAGATCGTCAAATTGCGGGAAACCCCTTAG
AGCCTCGAGCTACCAAGTGCGTGTCGAAAGGCATAGACACAGCCAGCAATGGCTGTCGCA
TGGCTTGATTCCATCAAGATAGGTAAAAATGCTCGAGGATTGGGCAATCCGCAGCCAAGC

CCCTAAGGTCCCCCCGTGGGGCTATGGGGAAGGTTCAACGACTGTAAGGCGTACCGTGCA
AGCGGAAGATACAGTCTAGCCCCGCTGGAAAGCAGCGCCAAAGTAAGTAGGTTGCAGTGG
ATCTGTCCTC
>SRR9587926
GCCTGTTTTGATATGTGTTTTGACAGTCATACGGGTCATGGCCGAGGGATGGTGGGTGAG
CTTCATAGCGCAATCCCCCGTCAGGAGCATGTTTTCATAGCATTTTGAGCATACTAGGGT
GGTCTCCCATGTAGCTCGCTCTCTCGGCTCAGGACCTTCCGTATGGGGCACTTTTTGACG
AGCTCCTTTGGGGAGGAGCATGTTCTGGCATATGAAGCGTTGGTCACGAGCTTCGTAGCC
CGACACATACCAAATCAGTCTTCATATGTCTTTCTGGCGGGGGTCGGTCGTATGACCACG
CCCTTACCTTGAGCATGTTTTTCGTATGATACCATACCTGACGGGCAGCGCCGCTGTTCA
GTCGTCGGCCGTACCTCCTATGCGCGTTTTCTGATTCGAACGTGTGACATGCAAGCGCAT
CATATCAGAAGCATGTTTTTTCATATGAGCATGTTGGTAGGCAGGCTCTTCAGTCGTCTA
CCCATAAATGCACTTTGAAAGACGTTTTTTGGTACTGAACGGCTTTGTGGTCAGACGTTC
TCGTGAGCCTATTCTCGCCCTCTGATACGCTGGATGCTGCGGTATTACCCACAGCAGACA
TCCAGGAGCTCCCTCTGCGGATAAGCATCTGTAGAAGGGGGTTTCTCTCGGTGACGCACT
GGCTGGGAAAACGGAGTGCTATCCAAAAGGGTGAAACCTCTTGGGGATGTACTCCCATAC
CAGCATAATATGCTGAGAGGAATTGACTCAATGAGTGAGGTCCTATACTGCAACCTCTTT
TGGATATGTAGGCGGTCTGTACCCAGACTGTCCTACTGGCTGATTTCCCTTGGTCCCACG
CGTCGCTGGCTGAATGGGCTAGTCCTTATGAAGTGCCGCGTTCATCAAAAGAACCGGTCT
TCATACAGCGACCTCATTCTATACTGCCAGCTGGATGGGCTGGTCCTTATGGAGCGCCGC
ATTCTTCGAAGAATCGGTTTCTATAGAGCGACCCCATTTACATATAAGGAACCGTTACAT
AGACTTGCCATTTGGCTGATGGCTCTGACAGTCGCTGCCTCTTCGGAGTTCTGTGTAAAC
GTGGCGCCATTTTTGAGTGGGCCAGTCCTTATGGAGTGTCCGTGTCTTCTATGACAGGGT
CCCCCATACAGCGACCTCACTCAGAACCGTATGGGCACGATGGTGAAAGACCATGCAGAT
AGTCCGCCTTCCAGAGGGGGAAGTAGTATATGGGGAATGCTGGCGAAGCATGAGGTACAT
GGTTCATGGACCCGTTGGGCCATGCCTGTGCCATCTTCTCGAGTCAAACCGGTCCGTGTC
GCTTCCTTATCTCTTCCTTTCGCGACGACAAGCTCTTGAGCTGGTCGTGGGTCAGGACGC
TGTGGTATCTGTTTGTCTGTAGTGTGGGCTTCCACCAGAACAGCGTATCAGCACACGTCG
TCTGGGCATATGATGTTTTTGCTTTGTGGTCCCGGGTTTGGTAGTATGGTTCCATGTCGG
TCCGCTCACCTCCCAACCATAATGTCTTTCACATCATCTTCACAGTACGTCTTGTCAGCA
TACCATTCTGTATAGAACAGTGTATCAGCACACGTCGTCTGAGCATAGGGTATTTTTGCT
TTGTGGTCCCGGGTTTGGTAGTATGGTTCCATGTCGGTCCGCTCACCTCCCAACCATAAT
GTCATTTCTATCATCTGCACAGTACGTCTTGTCAGAATACCATTCTCTATGGCAGTCGAT
AGACTAGGCCTTATGCCGCAGCAGTCTCTGATCACTGTGCGAGTGCATATATCTCTCAAT
GTGTTCATGGCTAGCATGGGTCTTGAGTGAGATGTGCCAGAGAGATACGCCACCAGTCTA
TGGAAAGGTGCTTTTTGTATGAGAAGAGTACATGTATGAAGAGCATGTTCTCTCCTCTTA

TGATGGTACCGCGGGTACGGGCTGCTCGCATTATGCTTGGCAGGTGTCCACTGTGCATGC
AGGATTGCTGAACCCACTTGTATGGCCATGAGAATGGTCATTGGGTGAGGTTCACTTCTC
CATCGCTTTGTGTGTGTTAGATGTTTGGGTGGCTTCGGCCGCATGGAAATTCTCGGTCTG
TAACTCATCAGGGAGGGTTTCCATGTGAGTATTCTCCAGTCGACCACACACCCTTGGGCG
TCCCGCTAGACAAAATGTAACTCTCGATCCGACCTAGCCTGTGGAGTGTCTTTCCGTATG
GCGTGAAAGGGCCCAGACTGGATGTTTTCCCTTTTGGACACCAGTCTGTTATATAGATCA
TCTACAGGGAGCAAAGGACAGGCATGCAAGTGCCACAAAGGCAGGGAGGCCTTCCTGCTA
CTGTGTGTCCTTCTTGACGATCACGGGGGAGTATATAGAAACATTTGAAGGAGATCCTTT
TCCACAAGGCGGTGCCCTCGCGCAAAGGTGTTGCTCTGCACGCATCTGTATGATGATAGT
TGCTCGCTCCTATGCGTCCACTTGTGAAAATACCTTCTCTCTTTGTCTGTTTCTATGACG
CCCCTTCTGCTTTGTCATAGAGCATACTCAGGTACAGAAAGTGCCGTGACTTCCTTGGCC
TGCTGTCAAAGAAATAGCCTGGTGCTATAGAGGACTCTTCCTGGGCGCCGCCTGGTGAAG
AGGTCCTATCAGGCTCTGGAGTGTGCGTATGCCTATGTTCATCATAGCTCATGTGCACGC
GCGGAAAAGCATCGACGTGGAATGGGGAAGAACACCATTCTGAATGAGCCAGTTCTTCCT
CTGCTCCCCTTGCTTGACGCTGTATGGGAATCGCTTTGCCTTGGGGGTACACCTTCCATT
CCATGACAAAGCATTCGATCTATGTCCCGAGGGGTGATTCTACATGAAGAACCCGAAACT
TGTCTGAGTGAGAGGCGAAGGTAGTCCTTCATTTTATGCGGGATCCGCCCTCACACTGTA
GTCGGCACAGTCCCTTGTTTAGGGACTGCGCAGCTCAATCTTCGACTGGAGAGGAAAAAA
ACAAAACACAAGTATGACACGCACATACGTGTTTTCTAGGCCATTGACCTGTTGGCTGCC
TCGCGGTGGTTGGCATGATGCCCCAAGTGGGGGGGGGGGAAATCAATCAGTATGGTTGAC
AAAATCCAGTGGGGAAAAAAGGCGCGCTGTCGCACACCCTTGGTCACGGCCTCAAAAAGT
CGACGGCCCTCACTGGCCCACCCCTGCGACTCCTGTTCGCTTTTGACAAGACACACCTGC
AATATGGAGGAAATGCCTGAGCAGCTGTGCCGCAGCAGCTGTGGACGCGTAGCGATATGC
ATCCCTATTGTTTGCGACTTGGTACAAAGAACGCGTGCCGGGAGACGGATTGCCCTTTGA
CTTTTCCACCCGGCCCGCGGCTCTGCCCTTTACGTACCCTTATTTACCTGGTTGATCCTG
CCAGTAGTCATATGCTTGTCTCAAAGACTAAGCCATGCATGTCTAAGTATAAACACTTTT
ATACTCGTGAAACTGCGAATGGCTCATTAAAACAGTTATAGTTCCTTTGGGAACAACACT
ACTTGGATAACCGTAGTAATTCTAGAGCTAATACATGCCTCAACGCCCGACTCACGAAGG
GTGGTATTTATTGGATAAAAAACCATCGTGCTCTCTCCAGAGCGCTTTGAGATGATTCAC
AATAACTTGTCGGATCGCATGGCCTCGTGCTGGCGACTCCTCATTCAAATTTCTGCCCTA
TCAACTTTCGATGGTAGAGTATTGGTCTACCATGGTGTCGACGGGTGACGGGGAATTAGG
GTTCGATTCCGGAGAGGGAGCCTGAGAAACGGCTACCACATCCAAGGAAGGCAGCAGGCG
CGCAAATTACCCAATCCCGACTCGGGGAGGTAGTGACAAAAAATAACAATAGGGGGCCCT
TTGGGTCTTCTAATTGGAATGAGAACAATTTAAATCCCTTATCGAGGATCCATTGGAGGG
CAAGTCTGGTGAACCTAGTTCCACCCGCTCCTCGTGTTGATCGCACGCAGGCGCACCGTG
TGAAGCATCAAGCCCTCGTTAAAATAAAACAGCGCCAGATGTCTCGTCCGGAAGGAGGTC

CCTCAGGTAGAGCACACAACAAAAGCGGCCAGTCAACTCTGGGCGTGTGTGATGCTGCGG
CTGAAGTGCACATAGATCGGTGCTTGCATGCTGTGACAAAAAGTGGGGTCGAGGTGCTGA
AATTGTCCAGAACGATGTGTCTCCTACAAGAAACACCTCGCGATGGACGCTACACGCACG
GTCAAAGAAAGCACTTCTATGTTTATCAGGCGGTGGCTTATGCCAGGTTTGGTAGACAGG
AAATGCTTAAAGTGACCACCTCAAAGATCGGCACAGACCTGACCATATCGCATGTGTGTG
GCACCCATCACTGCCTAGCAGCGCATCACTTAATGTTAGAACCAAAGCATGTGAATGATG
ATCGCGTGCACTGCCATGTTTTTCTCGGGCGCGTTTCTGACGCTTCAGAGAGGCATGCGT
TGGCAGAGCGTCATTGCCCACATTTACCGCGGTGCTATTCTGTTTGACTGCTAACGGACA
AGAGGCAAAAGATGGGAATCGTTTATAAAACCATTGAGCCCCCATCATACACTTCTAAAT
TGCCGGGGAGTGCCCGCGAGGCCTTGACTACCGCGGTCCGCCGAGAGGTGAGAGCGAGCA
ATCGTTCAGGACGCAGCAGTGTACGTAGTGGTACATGGATGGTAAAAACGTCAAGGATAG
GGTTGATCCGCAGGGAAGCGCGAGTGCCGCCATGGTCGGTGGGCAAAGGCGAACCTTCAG
AGACTCTAATGGAGTGGGCGCAAGCTTAAGGGAGAGTCCAATCCACTGGGAAACCAGGTC
CACAGCAGTAACGGGGTGATGCTCGCCCCAGAGGACTGTGGAAGGCACCACCACCCCAAG
AGGGGGTGGGTGTTGTCGGGTAGCAAATGGCCAGCAGCCGCGGTAATTCCAGCTCCAATA
GCGTATATTAAAGTTGTTGCAGTTAAAACGCTCGTAGTCGGACCTCGGAGCGCCCCCACG
GGCCGGGCTTTTGCCTTGTTGCCAGTGGGAGTTTGTCTCCTTTTGTCGGGACGCGGTGTG
CAGGGCCTTTACTGTCTCTGCGCGTTTACCTGCGGGCCGACCGTTTACTGTGAAGAAAGT
AAAGTGTTCAAAGCAGGCCAATGCCTTGAATATGTGAGCATGGAATAATAGAATAGGACT
TGGGCTCTATTTTGTTGGTTTCCAGTGACCAAGTAATGATTAATAGGGACGGTTGGGGGC
ATTCGTATTTCATTGTCAGAGGTGAAATTCTTGGATTGATGGAAGACGTACAACTGCGAA
AGCATCTGCCATGGATGTTTTCATTGATCAAGAACGAAAGTTAGGGGATCGAAGACGATC
AGATACCGTCGTAGTCTTAACCATAAACGATGCCGACTGGGGATTGGCGGGGGCACTATT
ATATGACTCTGTCAGCACCCTGAGGGAAACCAAAGTCTTTGGGTTCTGGGGGGAGTATGG
TCGCAAGGCTGAAACTTAAAGGAATTGACGGAAGGGCACCACCAGGAGTGGAGCCTGCGG
CTTAATTTGACTCAACACGGGAAAACTTACCAGGTCCGGACAGAAGAATGATTGACAGAC
TGAAAGCTCTTTCTTGATTTTTTGGTTGGTGGTGCATGGCCGTTCTTAGTTGGTGGAGTG
ATTTGTCTGGTTAATTCCGTTAACGAACGAGACCTCGTCCTGCTAAATAGGTGCGCGCAT
GCCAGCACTGCGTTCTTACCTTCTTAGAGGGACTATGCGCGTCTAGCGTATGGAAGATTG
AGGCAATAACAGGTCTGTGATGCCCTTAGATGTTCTGGGCCGCACGCGCGCTACACTGAT
GCATTCAACGAGTTTCACTTGATAGTCCTGGGTCGGAAGGCTCGGGTAATCTTTTGAAAG
TGCATCGTGCTGGGGATAGATCATTGCAATTATTGATCTTCAACGAGGAATTCCTTGTAG
GTGCAGGTCATCAGCCTGCGCCGAATACGTCCCTGCCCTTTGTACACACCGCCCGTCGCT
CCTACCGATTGAATGGTCCGATGAAATGTCCGGATCGCGGCTAGTGTGATGGACTCGTCC
AGAGCGGTAGCCGTGAGAAGCTCATTAAATCTTATCATTTAGAGGAAGGAGAAGTCGTAA
CAAGGTTTCCGTATTTTGTGGGTGGATACCGATTCAATAACATCCGCCGTTTTTCTCAAT

GTACTATATCCCTAGTCTGCTTGCCTAGTTCTTGGAGTCTTGTGCGATGTGTGCTATTAC
AGGCATGCCGGGTCATGAAGACAGACAAGGTTGGGATCTTCACGGAAGTACGCGCAATAG
GAGCGCGTCTTGTTCAACGCACCCGACTCAAGAGTCATATGCGAGGGACGGACGCAGCGC
TTGTCGCCGCACAGATGGGACGCGCCATATCGGAGACAGGGGGACGTGCCATTGTGATGA
TGATGATGATGTTGTTGGGTGGTCCTTCCGTAGGAAAACGTACACGAAGCCTTTGCGGCC
CGAAAGGGTGCACAGTGCGCGACTATAAACACTGATTCTGTGGTCAACGCAAGCAGCGGT
TCTGTGGAAGAACAGCTGCGACTTTCTCAAATTGCGGGAACACCCTAAAGTCAACAAACA
CCGCGGCCGGACGTGAAAGCGTCAGGCGCAGCAGTCGGGGTAGCGCCCTGGTGGAAGGTA
ATAGCTTTGTTGAATGGCACAATGGGCAATCCGCAGCCAAGCCCCTAAGGTCCCCCGCTG
GGCTATGGGGAAGGTTCACAGACTGTAAGGGAGGGGGTTTCTTCCATGAAAGAAGCTTAA
GAGACAGTCGGTCCCCTGCGAGAGCAGTGTCGCATGTGAGGAAGGCACCCATGAAAAGTG
GGTGCTGGAGAGCCATGTGAGCCCTCTATGAGGGTGGGAAGTTTTCCGATACAATGAGAA
ACAAACACGTTCCGTAGGTGAACCTGCGGAAGGATCATTCACAAACTATTGACACAAACA
CACGCGAACCAAATCGTCCACACAGGTGCCGATGAAAGAGAGAATCTGTTTGTCGCCTTT
TGGGGTATACAAGCACTCTTTTCCATCGCCTCTGTGCAGGGCGTAACTTCCCATTGAGAG
GATGTGAGGGCACCACAGGAAGCTTTTCCACAGGAAGTCGCCATCCTTCTCCCTCCACGA
CGGCGGCGCTTCACAGGCTTGGCAGTTTTTTTTTTTTTTTATTGCCTTCCAGGGAGGATG
CCGCCAATGGAGCCACATAGACATTTACATCATCATATGCCCGTTTTTCTTTCTTAACCG
CTTGCCACAGTTTCTCCTGTGAGGAGCTTGTGGGAAGACCGTCTCCATACAATAACAAAA
GATACAACTCTTAGCGGTGGATATCTTGGTTCTCGCAACGATGAAGAACGCAGCCAACTG
CGATACCTAGTGTGAATTGCAGGACTCCGTGAATCATTAAGTCTTTGAACGCAAGTTGCG
CTCATCTCCATGAGGAGGTGAGCATGCCTGTTTGAGTGTCGATGGGAAAAACAATACATA
ACAACAATCGGCTCTCTTCCCTTCCTGCCTTCCCCGGCAAAAAAGGTAGTGCATGCCGGA
GATGGGCCTGGTCGCCCCTTCATGAAAGAGGCGTCGTCCGAAACATGTGTCGTTGCCCGC
TGGAGGTCATACAGCATGAGTTTCAAAAACACCATGGCTCTCAAGGCAGAGTAAATCGTC
TTGAGACGGTCTGCATTGATTCACTTTCAACCCAGGTGGCGTTGGTTACTTGCCATAAGC
CTCTCTGGGGTTTCTTTGCTTGGACCTCTCTTGTCAGTTCTTCTGACGTAGAGACAGGTG
CGCGGTCTCATGGGCGTTGCCCTCTGGAACGTGCTAAAAAAAACAAAACAAACAACAACA
ACAACCGTATCACACGTCAAGAGGGTCGGATCGTACTTGACGGCGGGAGATCCTTTGTTT
CTCGCTTGTCCTGAGCGTCTGCCAAGCCTTCTTTGACGAGTGTGTTTGTTCCTTCCATAA
ACCCGATCTCACATCAGGTAGGACCACCCGCTGAATTTAAGCATATAATTAAGCGGAGGA
AAAGAAACTAACAAGGATTCCCTCAGTAGCGGCGAGCGAAGCGGGAAGAGCTCAACATGA
AAACCGTAGGGGGTTCGCGCTCACCTTCGGGTTGTATAGTAAAGTGGTGTCGCCTGCGTC
GCGCCGGGTATAAATCCCGTTTGAAATGGGGTGTCGTAGAGGGTTATAACCCCGTCTGTG
ACCCGGTGTGCGATGCGTTACGGTGACCTTACGATGAGTCGGGTTGCTTGGGCATGCAGC
CCTAAGTGGGTGGTAAACTTCATCTAAAGCTAAATACTAGCGAGAGACCGATAGCGTACA

AGTACCGTAAGGGAAAGATGCAAAGAACTTTGAAAAGAGAGTTAAACAGGGCTTGAAACC
GTTAAGAGTGAAGCGAAGGAGGTCAGTGATCATGGGGCCAGATCCCCACGTGCCCGTCAT
AGGGCACCGTAGCGTTCTGGTTCCCGGGGCAGCGCCGGCCGGGGGCGGGTGTCAACAGGG
AGGTGGTTGAGGGATTAACTTTCCGCCTGTACTCCTTTCTAGCTTGCACCCCGGCCGAGG
GATCCGCGACTTGTCGTGTGAGGGCGCTGTCAAAATGGCCTTCTCCACCCCGTCTTGAAA
CACGGACCAAGGAGTCTAACATACTCGCGAGTGTTTGGGTGGTTAAAACCCGAACGCACA
ATGAAAGTAAACGGTGGCTCTGCGCTGCTGCTTGGTGGGAACCCTGCTCTTATAACGGGG
GGCACCATCACCCGACCCCGATCCCTTTGGGGTGGTTGGTTTTGAGGCAGAGCGAGTATG
ATGGGACCCGAAAGATGGTGAACTATTCCTGAGTAGGGCGAAGCCAGAGGAAACTCTGGT
GGAGGCTCGTAGCGATGCTGACGTGCAAATCGCTCGTCGAACTTGGGTATAGGGGCGAAA
GACTAATCGAACCATCTAGTAGCTGGTTCCCTCCGAAGTTTCCCTCAGGATAGCGGGAGC
AAATGCAAGTCAGTATCATCCGGTAAAGCGAATGATTAGAGGCCTCGGGGGAGTAACTCC
TTCGACCTATTCTCAAACTTTAAATAGGTGAGGAGCCGGCAACGTTGCTTAAAACAATGA
ACGCCGTGCGGACAATGATTGCTCTCAGTGGGCCGATTTTGGTAAGCAGAACTGGCGATG
CGGGATGAACCGAAAGCGACGCTAAGGTGCCAAACTCCCAGCTCATTCAGACACCAGAAA
AGGTGTTGATTCATACAGACAGCAGGACGGTGGCCATGGAAGTCGGAATCCGCTAAGGAG
TGTGTAACAACTCACCTGCCGAATGGATCAGCCCTGAAAATGGATGGCGCTTAAGCGGGG
GACCGAGGCGTCGCCGTTGGGGTAAATACGAAGCCTCAATGAGTAGGAGGGCGTGGGGGT
TTTGAAACACGGTACACACGCATGGAAGCCTTGGGCGTAAGCTCGGGTGGAGCGACCCCT
AGTGTAGATCTTGGTGGTAGTAGCAAATATTCAAGTGAGAACCTTGAAGACTGAAGTGGG
GAAGGGTTCCACGCGAACAGTGCTTGGACGTGGGTTAGTCGGTCCTAAGAGATGGGGAAA
CTCCGTATTCAAGTGCGCTCTTCCAGCGCAGTCCATCGAAAGGGAAGCGGGTTAACATTC
CCGCACCAGGATGTGGAAACTGTGGTTACCGCAAACTGCGGCAACGCAACTGAACTTGGG
GACGCCCGTAGGAGCTCCGAGCAGGGATCTCTTTCCTGTTTAACGCGCCCAACCCCCTGG
AATCGGTTTGTCCGGAGATAGGGGGCGAAAGGCCGGAAAAGCACCGCACGTCTTGTGGTG
TTCGGAACGCTTGTAATGGCCCTTGAAAACCCAGGGGAAAGCAGCCGTCTTACACTTTTT
TGTGGGACGAGGAATTCTCACACCTGGCCGTACCCATCACCGCATCAGGTCCCCAAGGTG
AACAGCCTCTAGTCAATAGAATAATGTAGATAAGGGAAGTCGGCAAAATAGATCCGTAAC
TTTGGGAAAAGGATTGGCTCTAATGTTCGGGCTTGCTGGGACGCGCGGGCGTAGACGCCA
ATGAGCGGCAGACGTCTAGCCTCCTCGGAGGCTGGATTGACGCTGCTTGTGGGAAGAAAC
CGTGTCGATGGAGGCAGGAGCGCTTGCCGTTCTTGTCTCTTCCTGCTTGCGATCAACGAA
CGGATTAGAACTGGTACGGACAAGGGGAATCCGACTGTTTAATTAAAACAAAGCATTGCG
ACGGCCGGAAACGGTGTTGGCGCAATGTGATTTCTGCCCAGTGCTCTGAATGTCAAAGTG
AAGAAATTCAACCAAGCGCGGGTAAACGGCGGGAGTAACTATGACTCTCTTAACTACATG
GACATCTTTTGAGAGAGATAGGCGGGCTGAAAAGGTTATAGCCGTCTAGTCTCGCTCTTT
CATGAGCGGGGCGACACACCTGGTTGCGGGAACCCTGCCCACCTCTCTATCAGAAGTGGG

AACCAGTCCATTCCTACTCCTGCTTGTGAGGCATATAGTAAAAAGTAGCAGCAGGAGGAT
CTCGGCGAGAGGCCGAACCCGAAAGGAATAACGTGGGCTGGGAGGCAATCCGCAGGCGAG
TCCGTAACTATATGATTATCGACGGGCCGTCTCAACGACTGGTAAGGTGTGGGTTTGTCC
TCACGCGGGACAGGCTTAAGGTACAGTCTAATCCCACCCGCCAAGGGTGCTTGGGTGTAA
TCCCTCTTACATAGAGGGCACTAGCCCAGGGGCTGGGACGCTGGGAGGAAATGCCCCGTC
CCTATAGCCGGTAGAAATGAAGGTAGCCAAATGCCTCGTCATCTAAACTAATTGGATGAG
ATAGTCTGAGTGAAAAGCTTATGCAGGCTAGTACCGGTAAAAAAACGAGAGGCTCTGCCA
CTCTGACGGTGCGACACTCCTTGTACAGGGAAACCCTCCGCATGGAAAGGCACGTGGTAC
CACCCTCGCGAGGAAACTCGTGAGCGGGATCCCGGGTAATGACCGGAGCCGATGGTAAAA
ACCCACGTGACCGATGGGGTGACCCTGTGGCGAGAAGCTACGTTGTGACTTGTGTGCACA
ATATGCTTCCGTCGCAGAGACTGAACGGGAGTGGGCTCCGTCGACGGCCGGCCAGCCGTC
TTCCAGGAGCTTAAGATACAGTCCGGCCCACAGGGATGGCTTAAACCTGTGGAACAAGCA
GAATTAGTGACGCGCATGAATGGATTAACGAGATTCCCACTGTCCCTATCTACTATCTAG
CGAAACCACAGCCAAGGGAACGGGCTTGGAAGAATCAGCGGGGAAAGAAGACCCTGTTGA
GCTTGACTCTAGTCCGACTTTGTGAGATGAGTTGGGAGGTGTAGTCTAAGAGGGAGGGGC
AACTCGACCGTGAAATACCTCTACTCCCAAGCTTGTTTTACTTACTCCGGCGAGCAGGAA
TGGGTGGCAACACCCTTTTTCTAGGATTAACACCTCTTCGGGGGTGGACCTGCGCGGAGG
ACATTGTCAGGTGGGGAGTTTGGCTGGGGCGGCACATCTGTTAAAAGATAACGCAGGTGT
CCTAAGATGAGCTCAGTGAGAACAGAAATCTCACGTAGAACAAAAGGGTAAAAGCTCATT
TGATTTTGATTTTCAGTACGAATACAAACTGTGAAAGCATGGCCTATCGATCCTTTTGAC
CTTCAAGGTTTGAAGCAAGAGGTGTCAGAAAAGTTAGCACGCGTTGTGGTAGCTTTTAAG
AGTGGACTCAAAGTGTTCCGCTAGTCCCCCCTCGGACCGCCACTTACAAAAAGGACAGAG
GAGGGGGCGACACTGTCAAATTGCGGGGACTCCCTAAAGCCTGTCTCTACCAAGCCAGGC
GGCGAAAGCTCCTGGTGGCCGGGGTAATGACCTAGGGTATGGTAAAAGCGAGACAGGATG
AGAACAATGGGCAATCCGCAGCCAAGTCCTACGGCTCTTTTCTTGGAGAAGAGCTATGGA
TGCAGTTCAACGACTAGATGGCAGTGGGCTGAGGGAGCCTACGGCGGCACCCTTTCGGCT
TAAGATATAGTCTTGCCCACGTGAAAGCGTGCCTGCAGATGAACGCCACCATGTATGGCA
GGAGCTGCAGGAGGCCGCTAAACATACCAGCGGCTGGGAACATAACAGTACCACAGGGAT
GAACAAAAACACACAATTGTTGTGGCAATCATACAGTCGTATGATATTATCGTCCCAGGC
GGGTGCGCCTCTCTCACCCGTCTAGTCCGGCCTTGGTGCGCAAAAAATGCTTGCGTGGAT
GCGCCTTGGTTAAGGAAAATGCTCCTCGTAGTTTTACAAGGACATTTCTTGTAGGGCAAC
ACATCTGGATGCGAAGAACTCCCGGAAAGACAACGGTACCAAGTGACTGGGGTAACGCCC
CATTCATAAAGGCAATGAATAAAGAGGTGCCGGGTAATAATCTGTTGTATAGGGACAATC
CGCAGCTGACTTGGCTCCCTGCCCCCATCGGTGGGCATAGCTTAGCGGTTCAACGCTCGG
TAAGATGTGGGTTGGCGCGCTGTCCAACAGCGCACTGGCTTAAGGTACGGGCTACTCCCA
CCGGAGACGGTGTCTTTGTCTACCTCGGGTAACACGCCTCGGGAGAAGCAAAGAGGGTGC

AAGTAAAAAAACTGGTAGTGGGGAAATCCCTACCAGAGTCACCTGGTACAAAGGAACTGG
CTTGTGGCAGCCAAGCGTTCATAGCGACGTTGCTTTTTGATCCTTCGATGTCGGCTCTTC
CTATCATTGCGAAGCAGAATTCGCCAAGTGTTGGATTGTTCACCCACTAATAGGGAACGT
GAGCTGGGTAGGCATTTTATGCCCTATGCCCAACAGCTGGATAGCTGGGCTTCTGTCCTA
GGCCGGGACTTCTCACGACCCGTGCCTTGGTCACCCAAAAAAGGGCGACCACAGAGCTCA
GGATGGATCTTCTGGCTCAACTAGTAGAGGACAGGTAGAGATCGTCAAATTGCGGGAAAA
CCCTTAGAGCCTCGAGCTACCAAGTGCGTGTCGAAAGGCATAGACACAGCCAGCAATGGC
TGTCGCATGGCTTGATTCCATCAAGATAGGTAAAAATGCTCGAGGATTGGGCAATCCGCA
GCCAAGCCCCTAAGGTCCCCCCGTGGGGCTATGGGGAAGGTTCAACGACTGTAAGGCGTA
CCGTGCAAGCGGAAGATACAGTCTAGCCCCGCTGGAAAGCAGCGCCAAAGTAAGTAGGTT
GCAGTGGATCTGTCCTC
>SRR9587927
GCATGTTTTGGTATGTGTTTCGACAGTCATGCATGTCATAGCCGTAGGATGGGGAGTGAG
CTTCATAGCCCGACCCCTTCTCAGGAGCATGTTTTCATAGCATATTGAGCATACTGGGGT
GGTCTCCCATGTAGCTCGCTCTCTCGGTGCAGGACCTCCCTTTTGCGGGCCACTTCGATG
GCCTCCATTATGGAGAGCATGTTTTGGCATATGAAGCGTTGGTGGTGAGGCTCGTAGCCT
GTATATACATACCAACGCAGTCGTCATATGTCTCTCTGGCGGGGGTCGGTCGTATGACCT
TGTCCCTACCTGGAGCATGTATTTTGTATGATATACTGGACGGCAGCTGCGCTGTTCAGT
CGTCAGCCGTACCTCATGTGCGCGTTTCCAGGTCCGAACGTTTGACATGCAAGGGCATCA
TACCAGAAGCCTGTTTTCTCGTATGAGCATGTAGGTAGGCAGGCTCTTCAGTCGTCTACC
CATAAATGCACTTCGAAGGAACTTTTTGGTATTGAATGGCTTTGTGCCCAGACGTTCTTG
TGAGCTTGTTCTCGCTCTGCGATACGCTGGAAGCTGTGGTATTACCTACAGCAGACGCAT
TCCTAGGGCGGACGGCCTCTCTGCAGATGGGCACATCTGTAGAAAGAGGCTCCCCTTGAT
GACGCAATGCAATGGCTGGGTAAACGGAGTGCTATACAGAAGGGTCAAACCTCTGTGGAA
GTACACTCATACCAGCATTGTTTGTTGAGAGGGGTCGGCTCTACGAGTGAGGACCTATAC
TGCCATCTCATTCAGATGCGTAGACGGTCTTTACTCAGACTGCCTTTCTGGCTGATTGTC
CTCGGTCCCTCGTGCGCACGCGCCGCTGGCTGAATGGGCTTGTCCTTATGGAGTGCCGCG
TTCTTCAAAGAATCCGTCTTCATACAGCGACCTCATTTACATATTGCCGGCTGAATGGGC
TTGTCCTTATGGAGCACTGCATTCTTCTAGGAATTGCTATCTATACAGCGACCTCATTTA
CATACAAGGCACCTTTTTCTACATAGACTTGCCATTTGGCTGATTGTTCTGACAGTCGCT
GCCTCTTCGGAGTTCTGTGTGGACGAGTGGCCTCCTGAGTGGGCCGGTCCTTACGGAGTG
CCCAAGTCTGCTATGACTGGGTTCTCCATACAGCGACCTCACTCAGAATTGTATGGCACC
GTGCCGATTGAATGGACAAGCAGTCCTTAGGGAATCGCATGAAAGCATGACGGCAACATG
GTGTATGGACCCGTTGGGCCTTGCCTGTGCTATCTTCTCGAGTCGACCGGTCCTCGTCTC
CTCCTTTCGCGACGGGAAGCTCTTGAGTTTACCGTGGGTCAGGACGCTGTGGTATCTGTT
TGTCTGTAGTGTGGGCTCTCTATCAGAACAGCGTATCAGCACACGTCGTCTGGGTCTATG

GTATTTTTGCTTAACGGCCCTGGGTTTGGTAGTATGGTTCCATGTCGGTCCTCTCACCTC
CCAACCGTCATGTCTCTGATATCATCTGCACAGTACGTCTTGTCAGAATATCATTCTGTA
TGGCAGTCGATAGACTAGGCCCTATGCCGCAGCTGTCTCTGACCACTATGCGAGCGCATA
CCTACCTCAATGTGTTCATAGCTGTTATGGATCTTGGGTGTGGTGTGCCAGAACGATACG
CCACCAGATTGTGGAATGGTGCTTGTAGTATTAGAAGAGGACATGTATGAAGAGCACGTT
CTTTTCTCTTATGATGGTGCCGCGGGTGCGGGCCGTTCTCATTCTGCTTGTCGGGTGCTC
AAGGTGCATGTAGGATTGCTGAACCCACTGAATGGTCATGAGAATGGTCATATGGTGAGG
TTCACTTCTCCATCGCTTTGTGTGTGTTGGATGTTTGGGTGGCTTCGGCCGCATGGAAAA
TCTCGGTCTGTAACTCATCAGGGAGGGTTTCCATGTGAGCAATCTCTCCAGTCGAACACA
CACCCATGGGCGTCCCGCTAGGCAAAATGGAACTCTCGATCCGACACAGCCTGTGGAGTG
TTTGTTCCGTGTGGCGTGAAACGGCTTATACTGGATGTCTTCCCTTTGGACACCAGTCTG
TTGGTATGTACGCATACCATCTAGAGGGAGCAAGAAAAAGGCATGCAAGTGCCACAAAGG
AAGGGAGGCCTTCCTGCTACCGTGTGTCCTCTTGACGATCACGGGGGAGTATATAGAGAC
ATTTGGACGAGATCCGTTTTTCACGAGGCGGTGCCCTCGCGCAAAGGTGTTGCTCTGCAC
GCATCTTTATGATGATAGTTGCTCGCTCCTATGCGTCCACTCGTGAAAATACTTGCTCTC
TCTGTCTGTTTCCATGACACCCCCATGCGTTGTCATAGAGCATGCTCAGGTACAGAAAGT
GCTGTGACTTTCTTGGCCTGCTGTCACTGAGTCAGCCTGATACTTTGGAGGAATCTTTTC
TGGGCACTGCCCAGCTGAGAGTTCTTATCAGGCTCTGGAGTGTGCATATGCCTATATTCA
TTATAGTGCATGTGGCTACGCAGAAAAAGCATCGACGTGGAATGGAGAAGAACACCATTC
TGAATGGAACGGTTCTTACTCTACTTCTGTTGCTTGACGCTGTATGGGAATCGCTTTGCT
CTGGGGGTACACCTTCCATTCCATGACAAAGCATTTGATCTATGCCCCGAGGGTCATTCT
CTATGAAGAGCCCGAAACTTGTCTGAGTGAGAGGCGAAGGTAGTCCTTCATTTATGCGGG
GTCCGCCCTCACACTGTAGTCGGCACAGTCCCTTGTTTATGGACTGCGCAGCTCAATCTT
CGACTGGAGAGGAAAAACAAAACACAAGTATGACACGCACATACGTGTTTTCTAGGACTG
TGACCTGTCGGCTGCCTCGCGGTGGTTGGCATGATGCCCCAAGTGGGGGGGGGAGAAACA
ATCAGTATGATTGATAAAAACCAGTGGGGAAAAAGGCGCGCTGTCGCACACCCTTGGTCA
CGGCCTCAAGTCGACGGCCCTCACTGGCCCACCCTGCGACTCCTGTTCGCTTTTGACAAG
ACACACCTGCAATACGGAGGAAATGCCTGAGCAGCTGTGCCGCAGCAACTGCGGACGCGT
AGCGATATGCATCCCTGTTGTTTGCGACTTGGTACAAAGAACGTGCGCCGGGAGACGGAT
TGCCCTTTGACTTTTCCACCCAGCCCGCGGCTCTATACCCTTGACGTACCTATTTACCTG
GTTGATCCTGCCAGTAGTCATATGCTTGTCTCAAAGACTAAGCCATGCATGTCTAAGTAT
AAACACTTTTATACTCGTGAAACTGCGAATGGCTCATTAAAACAGTTATAGTTCCTTTGG
GAACAACACTACTTGGATAACCGTAGTAATTCTAGAGCTAATACATGCCTCAACGCCCGA
CTCACGAAGGGTGGTATTTATTGGATAAAAAACCATCGTGCTCTGTCCAGAGCGCTTTGA
GATGATTCACAATAACTTGTCGGATCGCATGGCCTCGTGCTGGCGACTCCTCATTCAAAT
TTCTGCCCTATCAACTTTCGATGGTAGAGTATTGGTCTACCATGGTGTCGACGGGTGACG

GGGAATTAGGGTTCGATTCCGGAGAGGGAGCCTGAGAAACGGCTACCACATCCAAGGAAG
GCAGCAGGCGCGCAAATTACCCAATCCCGACTCGGGGAGGTAGTGACAAAAAATAACAAT
AGGGGGCCCTTTGGGTCTTCTAATTGGAATGAGAACAATTTAAATCCCTTATCGAGGATC
CATTGGAGGGCAAGTCTGGTGAACGTATGCATTAAAGCCCTCCGTAATAAAAAAAAACAG
CGCCAGATGTCTCTCTCTGGAAGCACACCATGCATGGGGTTTGTTCCTCTCTGCATGGCC
GTGTGAAAAAGGCGAGTAGGCAAACGATGGGAAACCCGGCTTGGAATAGTAACCAGCCAC
AGAGCCCCCATCATACACTTCTAAATTGCCGGGGAGTGCCCGCGAGGCCTTGACTACCGC
GGTCCGCCGAGAGGTGAGAGCGAGCAATCGTTCAGGACGCAGCAGTGTACGTAGTGGTAC
ATGGATGGTAAAAACGTCAAGGATAGGGTTGATCCGCAGGGAAGCGCGAGTGCCGCCATG
GTCGGTGGGCAAAGGCGAACCTTCAGAGACTCTAATGGAGTGGGCGCAAGCTTAAGGGAG
AGTCCAATCCACTGGGAAACCAGGTCCACAGCAGTAACGGGGTGATGCTCGCCCCAGAGG
ACTGTGGAAGGCAACCCCACCAAAAGGGTGTGTGTTGTCGGGTAGCAATGGCCAGCAGCC
GCGGTAATTCCAGCTCCAATAGCGTATATTAAAGTTGTTGCAGTTAAAACGCTCGTAGTC
GGACCTCGGAGCGCCCCCACGGGCCGGGCTTTTGCCTTGTTGCCAGTGGGAGTTTGTCTC
CTTTTGTCGGGACGCGGTGTGCAGGGCCTTTACTGTCTCTGCGCGTTTACCTGCGGGCCG
ACCGTTTACTGTGAAGAAAGTAAAGTGTTCAAAGCAGGCCAATGCCTTGAATATGTGAGC
ATGGAATAATAGAATAGGACTTGGGCTCTATTTTGTTGGTTTCCAGTGACCAAGTAATGA
TTAATAGGGACGGTTGGGGGCATTCGTATTTCATTGTCAGAGGTGAAATTCTTGGATTGA
TGGAAGACGTACAACTGCGAAAGCATCTGCCATGGATGTTTTCATTGATCAAGAACGAAA
GTTAGGGGATCGAAGACGATCAGATACCGTCGTAGTCTTAACCATAAACGATGCCGACTG
GGGATTGGCGGGGGCACTTTTATATGACTCTGTCAGCACCCTGAGGGAAACCAAAGTCTT
TGGGTTCTGGGGGGAGTATGGTCGCAAGGCTGAAACTTAAAGGAATTGACGGAAGGGCAC
CACCAGGAGTGGAGCCTGCGGCTTAATTTGACTCAACACGGGAAAACTTACCAGGTCCGG
ACAGAAGAATGATTGACAGACTGAAAGCTCTTTCTTGATTTTTTGGTTGGTGGTGCATGG
CCGTTCTTAGTTGGTGGAGTGATTTGTCTGGTTAATTCCGTTAACGAACGAGACCTCGTC
CTGCTAAATAGGTGCGCGCATGCCAGCACTGCGTTCTTACCTTCTTAGAGGGACTATGCG
CGTCTAGCGTATGGAAGATTGAGGCAATAACAGGTCTGTGATGCCCTTAGATGTTCTGGG
CCGCACGCGCGCTACACTGATGCATTCAACGAGTTTCACTTGATAGTCCTGGGTCGGAAG
GCTCGGGTAATCTTTTGAAAGTGCATCGTGCTGGGGATAGATCATTGCAATTATTGATCT
TCAACGAGGAATTCCTTGTAGGCGCAGGTCATCAGCCTGCGCCGAATACGTCCCTGCCCT
TTGTACACACCGCCCGTCGCTCCTACCGATTGAATGGTCCGATGAAATGTCCGGATCGCG
GCTAGTGTGATGGACTCGTCCAGAGCGGTAGCCGTGAGAAGCTCATTAAATCTTATCATT
TAGAGGAAGGAGAAGTCGTAACAAGGTTTCCGTAGGTGAACCTGCGGAAGGATCATTCAC
AAACTATTGACACAAACACACGCGAACCAAATCGTCCGCACAGCAGGTGGCAATGAAAGA
GAGACTCTGCATGTCGCCTTTCGGGGTAGCAAGCAGCACTCTTCTGCCATCGCCTCTGTG
CAGGGCGTAAATTCCCATTGAGAGGATGTGAGGGCACCACAGGAAGCTTTTCCACAGGAA

GTCGCCATCCTTCTCCCTCCACGGCGGCGCTTCTGTGCTTGGCAGTTTTTTTTTGCCTTC

TAGGGAGGATGCCGCCAATGGAGCCCCTTATATATATACATCATCATATGCCCCTTTTTT

TCTTAACCGCTTGCCACAGCTTCTTCTATGAGGAGCTTGTGGGAAGACTGTCTCCATACA

ATAACAAAGATACAACTCTTAGCGGTGGATATCTTGGTTCTCGCAACGATGAAGAACGCA

GCCAACTGCGATAACTAATGTGAATTGCAGGACTTCGTGAATCATTAAGTCTTTGAACGC

AAGTTGCGCTCATCTCCATGAGGAGGTGAGCATGCCTGTTTGAGTGTCGATGGGAAAAAC

AATACATAACAACAATCGGCTCTCTTATCTTCCTGCCTTCTTGGGGGCAAAAGGTAGTGC

ATGCCGGAGATGGGCTTTGTCGCCCCTTTCTCTTGAAAGAGGCGTCGTCCGAAACACGTA

TCGTTGCCCGCTGGAGGTCATAGAGCATGAGTTTAAAGCACCACATCTCTCAAGTCAGAG

TGAGCGTCTTGAGTAGGTCCGCATTGATTCAGTTTCAACCCAGGTGGCGTTGGTTACTTG

CCATAAGCCTCTCTGGGGTTTCTTTGCTTGGACCTCTCTTGTCAGTTTCTCTGACGTAGA

GACAGGTGCCGTCTCATGGACGTTGCCCTCTGGAACGTGCTAAAAAAAAAAAAAGAAAAA

ACCGTACCACATGTCAAAAGAGGCAGATCGTGCTTGACGGCGGGAGGTGTTGCATTTCTT

GCTTGTCCTGAGCGTCTGCCAGTCTCTGACGAGTGTGTTCCTTCCATAAACCCGATCTCA

AATCAGGTAGGACCACCCGCTGAATTTAAGCATATAATTAAGCGGAGGAAAAGAAACTAA

CAAGGATTCCCTCAGTAGCGGCGAGCGAAGCGGGAAGAGCTCAACATGAAAACCGTTGGG

GGTTCGCGCTTTCCTTCGGGTTGTATAGTAAAGTGGTGTCGCCTGCGTCGCGCCGGGTAT

AAATCCCGTCTGAAATGGGGTGTCGTAGAGGGTTATATCCCCGTCTGTGACCCGGTGTGC

GATGCGTTACGGTGACCTTACGATGAGTCGGGTTGCTTGGGCATGCAGCCCTAAGTGGGT

GGTAAACTTCATCTAAAGCTAAATACTAGCGAGAGACCGATAGCGTACAAGTACCGTAAG

GGAAAGATGCAAAGAACTTTGAAAAGAGAGTTAAACAGGGCTTGAAACCGTTAAGAGTGA

AGCGAAGGAGGTCAGTGATCATGGGGCCAGATCCCCAAGTGCCTGTCATAGGGCACCGTA

GCGTTCTGGTTTCCCGGGGCAGCGCCGGCCGGGGGCGGGGGTTAACAGGGAGGTGGTTGA

GGGATTCGTTTTCCGCCTGTACTCCTTTATAGCTTGCACCCCGGCCGAGGGATCCGCGAC

TTGTCGTGTGAGGGCGCTGTCAAAATGGCCTTCTCCACCCCGTCTTGAAACACGGACCAA

GGAGTCTAACATACTCGCGAGTGTTTGGGTGGTTAAAACCCGAACGCACAATGAAAGTAA

ACGGTGGCTCTGCGCCGCTGCTTGGTGGGAACCCTGCTTTTATAATGGGGGGCACTATCA

CCCGACCCCGATCCCTTTGGGGTGGTTGGTTTTGAGGTAGAGCGAGTATGATGGGACCCG

AAAGATGGTGAACTATTCCTGAGTAGGGCGAAGCCAGAGGAAACTCTGGTGGAGGCTCGT

AGCGATGCTGACGTGCAAATCGCTCGTCGAACTTGGGTATAGGGGCGAAAGACTAATCGA

ACCATCTAGTAGCTGGTTCCCTCCGAAGTTTCCCTCAGGATAGCGGGAGCAAATGCAAGT

CAGTATCATCCGGTAAAGCGAATGATTAGAGGCCTCGGGGGAGTAACTCCTTCGACCTAT

TCTCAAACTTTAAATAGGTGAGGAGCCGGCAACGTTGCTTAAAACAATGAACGCCGTGCG

GACAATGATTGCTCTCAGTGGGCCGATTTTGGTAAGCAGAACTGGCGATGCGGGATGAAC

CGAAAGCGACGCTAAGGTGCCAAACTCCCAGCTCATTCAGACACCAGAAAAGGTGTTGAT

TCATACAGACAGCAGGACGGTGGCCATGGAAGTCGGAATCCGCTAAGGAGTGTGTAACAA

CTCACCTGCCGAATGGATCAGCCCTGAAAATGGATGGCGCTTAAGCGGGGGACCGAGGCG

TCGCCGTTGGGGTAAATACGAAGCCCCAATGAGTAGGAGGGCGTGGGGGTTTTGAAACAC

GGTACACTCGCATGGAAGCCTTGGGCGTAAGCTCGGGTGGAGCGACCCCTAGTGTAGATC

TTGGTGGTAGTAGCAAATATTCAAGTGAGAACCTTGAAGACTGAAGTGGGGAAGGGTTCC

ACGCGAACAGTGCTTGGACGTGGGTTAGTCGGTCCTAAGAGATGGGGAAACTCCGTATTC

AAGTGCGCTTTTGTAGCGCAGTCCATCGAAAGGGAAGCGGGTTAACATTCCCGCACCAGG

ATGTGGAAACTGTGGTTTACCGCAAACTGCGGCAACGCAACTGAACCTGGGGACGCCCGT

AGGAGCTCCGAGCAGGGATCTCTTTCCTGTTTAACGCGCCCAACCCCCTGGAATCGGTTT

GTCCGGAGATAGGGGGCGAAAGGCCGGAAAAGCACCGCACGTCTTGTGGTGTTCGGAACG

CTTGTAATGGCCCTTGAAAACCCAGGGGAAAGCAGCCGTCTTACACTTGTGTGAGACGAG

GAATTCTCACACCTGGCCGTACCCATCACCGCATCAGGTCCCCAAGGTGAACAGCCTCTA

GTCAATAGAATAATGTAGATAAGGGAAGTCGGCAAAATAGATCCGTAACTTTGGGAAAAG

GATTGGCTCTAATGTTCGGGCTTGTCGGGACGCGCGGGCGTAGATGCCAATGAGCGGCAG

ACGTCTAGCCTCTTTGGAGGCTGGATTAACGCTGCTTGTGGGAAGAAACCGTGTCGATGG

AGGCAGGAACGCTTATCGATCTTGTCTCTTCCCGCTTGCGATCAACGAACGGATTAGAAC

TGGTACGGACAAGGGGAATCCGACTGTTTAATTAAAACAAAGCATTGCGACGGCCGGAAA

CGGTGTTGGCGCAATGTGATTTCTGCCCAGTGCTCTGAATGTCAAAGTGAAGAAATTCAA

CCAAGCGCGGGTAAACGGCGGGAGTAACTATGACTCTTAAACAAACCATACTACCTACGG

GAGTCAAAAGTGGGGGCGGCGTCTGGTGTAGAAATACACGTACGACACGCTTTTTTTCCT

GCTAGTCGATCTGACCTTTGTTCTTTGGACTGTGGGCGGTCAGCGGCAACATAACCTGGT

ACGGGGAAGGCCTCGACGTTGAGTGCTGCATCTTGGAAGTATCTTGACACAATAAGAGAA

AAGAGATTGCGCTTAATAGGGGGTTGATCTCGTGGCGAGCCGACGCCGGTGTGAACTGGT

GGCAATGGCCGTCGTAGAGCACGCAAAGGTATGGGCTGACTTTTTCTCCTCAAAAGTTGG

CTTAAGGGACGTGCCATCCCACACGATAATAAAGTGTGCAGTCTGTAAGAGCTCCCACAG

GGCAAAGACAGACTGGGATGCAGACGGAGACGGCAAAAACAACAACATCCGTTCTGCTTC

TGGGCGCAAATGCTTAAGGTAGCCAAATGCCTCGTCATCTAAACTAATTGGATGAGATAG

TCTGAGTGAAAAGCTTATGCAGGCTAGTACCGGAAAAAAAAAAAAGAGGGGCTCTGCCAC

TCTGACGGTGCGACACTCCTTGTACAGGGAAACCCTCCGCATGGAAAGGCACGTGGTACC

ACCCTCGCGGGGAAACTCGCGAGCGGGATCCCGGGTAATGACCGGAGCCGATGGTAAAAA

CTCACGTGACCGATGGGGTGACCCTGTGGCGAGACGCTAAGTTGTGGGACATCACCCTCT

CACAATACGCATCCGTCGCAGAGACTGAACGGGAGTGGGCTCCGTTGACGGCCGGCCGGC

CAGCCAGCCGTCTTCCAGGGGCTTAAGATACAGTCCGGGCCTCAGGGATGGAAAAACCTG

GGGAACTACCAGAATTAGTGACGCGCATGAATGGATTAACGAGATTCCCACTGTCCCTAT

CTACTATCTAGCGAAACCACAGCCAAGGGAACGGGCTTGGAAGAATCAGCGGGGAAAGAA

GACCCTGTTGAGCTTGACTCTAGTCCGACTTTGTGAGATGAGTTGGGAGGTGTAGTCTAA

GAGGGAGGGGCAACTCGACCGTGAAATACCTCTACTCCCAAGCTTGTTTTACTTACTCCG

GCGAGCAGGGATGGGTGGTAACACCCTTTTTCTAGGATTAACACCTCTTCGGGGGTGGAC
CTGCGCGGAGGACATTGTCAGGTGGGGAGTTTGGCTGGGGCGGCACATCTGTTAAAAGAT
AACGCAGGTGTCCTAAGATGAGCTCAGTGAGAACAGAAATCTCACGTAGAACAAAAGGGT
AAAAGCTCATTTGATTTTGATTTTCAGTACGAATACAAACTGTGAAAGCATGGCCTATCG
ATCCTTTTGACCTTCAAGGTTTGAAGCAAGAGGTGTCAGAAAAGTTACCACAGGGATAAC
TGGCTTGTGGCAGCCAAGCGTTCATAGCGACGTTGCTTTTTGATCCTTCGATGTCGGCTC
TTCCTATCATTGCGAAGCAGAATTCGCCAAGTGTTGGATTGTTCACCCACTAATAGGGAA
CGTGAGCTGGGTAGCCACTTTATGCCCAACAGCTGGATAGCTGGGCTTCTGTCTTAGGCC
GGGACTTCTCACGACCCGTGCCTTGGTCACCCAAAAAAGGGCGACCGCAGAGCTCAGGAT
GGAGCTTCTGGCTTAACTAGTAGAGGACAGATAGAGATCGTCAAATTGCGGGAAACCCCT
TAGAGCCTCGAGCTACCAAGTGCGTGTCGAAAGGCATAGACGCAGCCAGCGATGGCTGTC
GCATGGCTTGATTCCATCAAGATAGGTAAAAATGCTCGAGGATTGGGCAATCCGCAGCCA
AGCCCCTAAGGTCCCCCCGCGGGCTATGGGGAAGGTTCAACGACTGTAAGGCGTACCGTG
CAAGCGGAAGATACAGTCTAGCCCCGCTGGAAAGCAGCGCCAAAGTAAGTAGGTTGCAGT
GGGTCTGTCCTC
>SRR9587929_SRR9587963
GCATGTTTTGGTATGTGTTTCGACAGTCATGCATGTCATAGCCGTAGGATGGGGAGTGAG
CTTCATAGCCCGACCCCTTCTCAGGAGCATGTTTTCATAGCATATTGAGCATACTGGGGT
GGTCTCCCATGTAGCTCGCTCTCTCGGTGCAGGACCTCCCTTTTGCGGGCCACTTCGATG
GCCTCCATTATGGAGAGCATGTTTTGGCATATGAAGCGTTGGTGGTGAGGCTCGTAGCCT
GTATATACATACCAACGCAGTCGTCATATGTCTCTCTGGCGGGGGTCGGTCGTATGACCT
TGTCCCTACCTGGAGCATGTATTTTGTATGATATACTGGACGGCAGCTGCGCTGTTCAGT
CGTCGGCCGTACCTCATGTGCGCGTTTCCAGGTCCGAACGTTTGACATGCAAGGGCATCA
TACCAGAAGCCTGTTTTCTCGTATGAGCATGTAGGTAGGCAGGCTCTTCAGTCGTCTACC
CATAAATGCACTTCGAAGGAACTTTTTGGTATTGAATGGCTTTGTGCCCAGACGTTCTTG
TGAGCTTGTTCTCGCTCTGCGATACGCTGGAAGCTGTGGTATTACCTACAGCAGACGCAT
TCCTAGGGCGGACGGCCTCTCTGCAGATGGGCACATCTGTAGAAAGAGGCTCCCCTTGAT
GACGCAATGCAATGGCTGGGTAAACGGAGTGCTATACAGAAGGGTCAAACCTCTGTGGAA
GTACACCCATACCAGCATTGTTTGTTGAGAGGGGTCGGCTCTACGAGTGAGGACCTATAC
TGCCATCTCATTCAGATGCGTAGACGGTCTTTACTCAGACTGCCTTTCTGGCTGATTGTC
CTCGGTCCCTCGTGCGCACGCGCCGCTGGCTGAATGGGCTTGTCCTTATGGAGTGCCGCG
TTCTTCAAAGAATCCGTCTTCATACAGCGACCTCATTTACATATTGCCGGCTGAATGGGC
TTGTCCTTATGGAGCACTGCATTCTTCTAGGAATTGCTATCTATACAGCGACCTCATTTA
CATACAAGGCACCTTTTTCTACATAGACTTGCCATTTGGCTGATTGTTCTGACAGTCGCT
GCCTCTTCGGAGTTCTGTGTGGACGAGTGGCCTCCTGAGTGGGCCGGTCCTTACGGAGTG
CCCACGTCTGTTATGACTGGGTTCTCCATACAGCGACCTCACTCAGAATTGTATGGCACC

GTGCCGATTGAATGGACAAGCAGTCCTTAGGGAATCGCATGAAAGCATGACGGCACATGG
TGTATGGACCCGTTGGGCCTTGCCTGTGCTATCTTCTCGAGTCGACCGGTCCTCGTCTCC
TCCTTTCGCGACGGGAAGCTCTTGAGTTTACCGTGGGTCAGGACGCTGTGGTATCTGTTT
GTCTGTAGTGTGGGCTCTCTATCAGAACAGCGTATCAGCACACGTCGTCTGGGTCTATGG
TATTTTTGCTTAACGGCCCTGGGTTTGGTAGTATGGTTCCATGTCGGTCCTCTCACCTCC
CAACCGTCATGTCTCTGATATCATCTGCACAGTACGTCTTGTCAGAATATCATTCTGTAT
GGCAGTCGATAGACTAGGCCCTATGCCGCAGCTGTCTCTGACCACTATGCGAGCGCATAC
CTACCTCAATGTGTTCATAGCTGTTATGGATCTTGGGTGTGGTGTGCCAGAACGATACGC
CACCAGATTGTGGAATGGTGCTTGTAGTATTAGAAGAGGACATGTATGAAGAGCACGTTC
TTTTCTCTTATGATGGTGCCGCGGGTGCGGGCCGTTTTCATTCTGCTTGTCGGGTGCTCA
AGGTGCATGTAGGATTGCTGAACCCACTGAATGGTCATGAGAATGGTCATATGGTGAGGT
TCACTTCTCCATCGCTTTGTGTGTGTTGGATGTTTGGGTGGCTTCGGCCGCATGGAAAAT
CTCGGTCTGTAACTCATCAGGGAGGGTTTCCATGTGAGCAATCTCTCCAGTCGAACACAC
ACCCATGGGCGTCCCGCTAGGCAAAATGGAACTCTCGATCCGACACAGCCTGTGGAGTGT
TTGTTCCGTGTGGCGTGAAACGGCTTATACTGGATGTCTTCCCTTTGGACACCAGTCTGT
TGGTATGTACGCATACCATCTAGAGGGAGCAAGAAAAAGGCATGCAAGTGCCACAAAGGA
AGGGAGGCCTTCCTGCTACCGTGTGTCCTCTTGACGATCACGGGGGAGTATATAGAGACA
TTTGGACGAGATCCGTTTTTCACGAGGCGGTGCCCTCGCGCAAAGGTGTTGCTCTGCACG
CATCTTTATGATGATAGTTGCTCGCTCCTATGCGTCCACTCGTGAAAATACCTGCTCTCT
CTGTCTGTTTCCATGACACCCCCATGCGTTGTCATAGAGCATGCTCAGGTACAGAAAGTG
CTGTGACTTTCTTGGCCTGCTGTCACTGAGTCAGCCTGATACTTTGGAGGAATCTTTTCT
GGGCACTGCCCAGCTGAGAGTTCTTATCAGGCTCTGGAGTGTGCATATGCCTATATTCAT
TATAGTGCATGTGGCTACGCAGAAAAAGCATCGACGTGGAATGGAGAAGAACACCATTCT
GAATGGAACGGTTCTTACTCTACTTCTGTTGCTTGACGCTGTATGGGAATCGCTTTGCTC
TGGGGGTACACCTTCCATTCCATGACAAAGCATTTGATCTATGCCCCGAGGGTCATTCTC
TATGAAGAGCCCGAAACTTGTCTGAGTGAGAGGCGAAGGTAGTCCTTCATTTATGCGGGG
TCCGCCCTCACACTGTAGTCGGCACAGTCCCTTGTTTATGGACTGCGCAGCTCAATCTTC
GACTGGAGAGGAAAAACAAAACACAAGTATGACACGCACATACGTGTTTTCTAGGACTGT
GACCTGTCGGCTGCCTCGCGGTGGTTGGCATGATGCCCCAAGTGGGGGGGGGAGAAACAA
TCAGTATGATTGATAAAAACCAGTGGGGAAAAAGGCGCGCTGTCGCACACCCTTGGTCAC
GGCCTCAAGTCGACGGCCCTCACTGGCCCACCCTGCGACTCCTGTTCGCTTTTGACAAGA
CACACCTGCAATACGGAGGAAATGCCTGAGCAGCTGTGCCGCAGCAACTGCGGACGCGTA
GCGATATGCATCCCTGTTGTTTGCGACTTGGTACAAAGAACGTGCGCCGGGAGACGGATT
GCCCTTTGACTTTTCCACCCAGCCCGCGGCTCTATACCCTTGACGTACCTATTTACCTGG
TTGATCCTGCCAGTAGTCATATGCTTGTCTCAAAGACTAAGCCATGCATGTCTAAGTATA
AACACTTTTATACTCGTGAAACTGCGAATGGCTCATTAAAACAGTTATAGTTCCTTTGGG

AACAACACTACTTGGATAACCGTAGTAATTCTAGAGCTAATACATGCCTCAACGCCCGAC
TCACGAAGGGTGGTATTTATTGGATAAAAAACCATCGTGCTCTGTCCAGAGCGCTTTGAG
ATGATTCACAATAACTTGTCGGATCGCATGGCCTCGTGCTGGCGACTCCTCATTCAAATT
TCTGCCCTATCAACTTTCGATGGTAGAGTATTGGTCTACCATGGTGTCGACGGGTGACGG
GGAATTAGGGTTCGATTCCGGAGAGGGAGCCTGAGAAACGGCTACCACATCCAAGGAAGG
CAGCAGGCGCGCAAATTACCCAATCCCGACTCGGGGAGGTAGTGACAAAAAATAACAATA
GGGGGCCCTTTGGGTCTTCTAATTGGAATGAGAACAATTTAAATCCCTTATCGAGGATCC
ATTGGAGGGCAAGTCTGGTGAACGTATGCATTAAAGCCCTCCGTAATAAAAAAAAACAGC
GCCAGATGTCTCTCTCTGGAAGCACACCATGCATGGGGTTTGTTCCTCTCTGCATGGCCG
TGTGAAAAAGGCGAGTAGGCAAACGATGGGAAACCCGGCTTGGAATAGTAACCAGCCACA
GAGCCCCCATCATACACTTCTAAATTGCCGGGGAGTGCCCGCGAGGCCTTGACTACCGCG
GTCCGCCGAGAGGTGAGAGCGAGCAATCGTTCAGGACGCAGCAGTGTACGTAGTGGTACA
TGGATGGTAAAAACGTCAAGGATAGGGTTGATCCGCAGGGAAGCGCGAGTGCCGCCATGG
TCGGTGGGCAAAGGCGAACCTTCAGAGACTCTAATGGAGTGGGCGCAAGCTTAAGGGAGA
GTCCAATCCACTGGGAAACCAGGTCCACAGCAGTAACGGGGTGATGCTCGCCCCAGAGGA
CTGTGGAAGGCAACCCCACCAAAAGGGTGTGTGTTGTCGGGTAGCAATGGCCAGCAGCCG
CGGTAATTCCAGCTCCAATAGCGTATATTAAAGTTGTTGCAGTTAAAACGCTCGTAGTCG
GACCTCGGAGCGCCCCCACGGGCCGGGCTTTTGCCTTGTTGCCAGTGGGAGTTTGTCTCC
TTTTGTCGGGACGCGGTGTGCGGGGCCTTTACTGTCTCTGCGCGTTTACCTGCGGGCCGA
CCGTTTACTGTGAAGAAAGTAAAGTGTTCAAAGCAGGCCAATGCCTTGAATATGTGAGCA
TGGAATAATAGAATAGGACTTGGGCTCTATTTTGTTGGTTTCCAGTGACCAAGTAATGAT
TAATAGGGACGGTTGGGGGCATTCGTATTTCATTGTCAGAGGTGAAATTCTTGGATTGAT
GGAAGACGTACAACTGCGAAAGCATCTGCCATGGATGTTTTCATTGATCAAGAACGAAAG
TTAGGGGATCGAAGACGATCAGATACCGTCGTAGTCTTAACCATAAACGATGCCGACTGG
GGATTGGCGGGGGCACTTTTATATGACTCTGTCAGCACCCTGAGGGAAACCAAAGTCTTT
GGGTTCTGGGGGGAGTATGGTCGCAAGGCTGAAACTTAAAGGAATTGACGGAAGGGCACC
ACCAGGAGTGGAGCCTGCGGCTTAATTTGACTCAACACGGGAAAACTTACCAGGTCCGGA
CAGAAGAATGATTGACAGACTGAAAGCTCTTTCTTGATTTTTTGGTTGGTGGTGCATGGC
CGTTCTTAGTTGGTGGAGTGATTTGTCTGGTTAATTCCGTTAACGAACGAGACCTCGTCC
TGCTAAATAGGTGCGCGCATGCCAGCACTGCGTTCTTACCTTCTTAGAGGGACTATGCGC
GTCTAGCGTATGGAAGATTGAGGCAATAACAGGTCTGTGATGCCCTTAGATGTTCTGGGC
CGCACGCGCGCTACACTGATGCATTCAACGAGTTTCACTTGATAGTCCTGGGTCGGAAGG
CTCGGGTAATCTTTTGAAAGTGCATCGTGCTGGGGATAGATCATTGCAATTATTGATCTT
CAACGAGGAATTCCTTGTAGGCGCAGGTCATCAGCCTGCGCCGAATACGTCCCTGCCCTT
TGTACACACCGCCCGTCGCTCCTACCGATTGAATGGTCCGATGAAATGTCCGGATCGCGG
CTAGTGTGATGGACTCGTCCAGAGCGGTAGCCGTGAGAAGCTCATTAAATCTTATCATTT

AGAGGAAGGAGAAGTCGTAACAAGGTTTCCGTAGGTGAACCTGCGGAAGGATCATTCACA
AACTATTGACACAAACACACGCGAACCAAATCGTCCGCACAGCAGGTGGCAATGAAAGAG
AGACTCTGCATGTCGCCTTTCGGGGTAGCAAGCAGCACTCTTCTGCCATCGCCTCTGTGC
AGGGCGTAAATTCCCATTGAGAGGATGTGAGGGCACCAAAGGAAGCTTTTCCACAGGAAG
TCGCCATCCTTCTCCCTCCACGGCGGCGCTTCTGTGCTTGGCAGTTTTTTTTGCCTTCTA
GGGAGGATGCCGCCAATGGAGCCCCTTATATATATACATCATCATATGCCCCTTTTTTTC
TTAACCGCTTGCCACAGCTTCTTCTATGAGGAGCTTGTGGGAAGACTGTCTCCATACAAT
AACAAAGATACAACTCTTAGCGGTGGATATCTTGGTTCTCGCAACGATGAAGAACGCAGC
CAACTGCGATAACTAATGTGAATTGCAGGACTTCGTGAATCATTAAGTCTTTGAACGCAA
GTTGCGCTCATCTCCATGAGGAGGTGAGCATGCCTGTTTGAGTGTCGATGGGAAAAACAA
TACATAACAACAATCGGCTCTCTTATCTTCCTGCCTTCTTGGGGGCAAAAGGTAGTGCAT
GCCGGAGATGGGCTTTGTCGCCCCTTTCTCTTGAAAGAGGCGTCGTCCGAAACACGTATC
GTTGCCCGCTGGAGGTCATAGAGCATGAGTTTAAAGCACCACATCTCTCAAGTCAGAGTG
AGCGTCTTGAGTAGGTCCGCATTGATTCAGTTTCAACCCAGGTGGCGTTGGTTACTTGCC
ATAAGCCTCTCTGGGGTTTCTTTGCTTGGACCTCTCTTGTCAGTTTCTCTGACGTAGAGA
CAGGTGCCGTCTCATGGACGTTGCCCTCTGGAACGTGCTAAAAAAAAAAAAAAGAAAAAA
CCGTACCACATGTCAAAAGAGGCAGATCGTGCTTGACGGCGGGAGGTGTTGCATTTCTTG
CTTGTCCTGAGCGTCTGCCAGTCTCTGACGAGTGTGTTCCTTCCATAAACCCGATCTCAA
ATCAGGTAGGACCACCCGCTGAATTTAAGCATATAATTAAGCGGAGGAAAAGAAACTAAC
AAGGATTCCCTCAGTAGCGGCGAGCGAAGCGGGAAGAGCTCAACATGAAAACCGTTGGGG
GTTCGCGCTTTCCTTCGGGTTGTATAGTAAAGTGGTGTCGCCTGCGTCGCGCCGGGTATA
AATCCCGTCTGAAATGGGGTGTCGTAGAGGGTTATATCCCCGTCTGTGACCCGGTGTGCG
ATGCGTTACGGTGACCTTACGATGAGTCGGGTTGCTTGGGCATGCAGCCCTAAGTGGGTG
GTAAACTTCATCTAAAGCTAAATACTAGCGAGAGACCGATAGCGTACAAGTACCGTAAGG
GAAAGATGCAAAGAACTTTGAAAAGAGAGTTAAACAGGGCTTGAAACCGTTAAGAGTGAA
GCGAAGGAGGTCAGTGATCATGGGGCCAGATCCCCAAGTGCCTGTCATAGGGCACCGTAG
CGTTCTGGTTTCCCGGGGCAGCGCCGGCCGGGGGCGGGGGTTAACAGGGAGGTGGTTGAG
GGATTCGTTTTCCGCCTGTACTCCTTTATAGCTTGCACCCCGGCCGAGGGATCCGCGACT
TGTCGTGTGAGGGCGCTGTCAAAATGGCCTTCTCCACCCCGTCTTGAAACACGGACCAAG
GAGTCTAACATACTCGCGAGTGTTTGGGTGGTTAAAACCCGAACGCACAATGAAAGTAAA
CGGTGGCTCTGCGCCGCTGCTTGGTGGGAACCCTGCTTTTATAATGGGGGGCACTATCAC
CCGACCCCGATCCCTTTGGGGTGGTTGGTTTTGAGGTAGAGCGAGTATGATGGGACCCGA
AAGATGGTGAACTATTCCTGAGTAGGGCGAAGCCAGAGGAAACTCTGGTGGAGGCTCGTA
GCGATGCTGACGTGCAAATCGCTCGTCGAACTTGGGTATAGGGGCGAAAGACTAATCGAA
CCATCTAGTAGCTGGTTCCCTCCGAAGTTTCCCTCAGGATAGCGGGAGCAAATGCAAGTC
AGTATCATCCGGTAAAGCGAATGATTAGAGGCCTCGGGGGAGTAACTCCTTCGACCTATT

CTCAAACTTTAAATAGGTGAGGAGCCGGCAACGTTGCTTAAAACAATGAACGCCGTGCGG
ACAATGATTGCTCTCAGTGGGCCGATTTTGGTAAGCAGAACTGGCGATGCGGGATGAACC
GAAAGCGACGCTAAGGTGCCAAACTCCCAGCTCATTCAGACACCAGAAAAGGTGTTGATT
CATACAGACAGCAGGACGGTGGCCATGGAAGTCGGAATCCGCTAAGGAGTGTGTAACAAC
TCACCTGCCGAATGGATCAGCCCTGAAAATGGATGGCGCTTAAGCGGGGGACCGAGGCGT
CGCCGTTGGGGTAAATACGAAGCCCCAATGAGTAGGAGGGCGTGGGGGTTTTGAAACACG
GTACACTCGCATGGAAGCCTTGGGCGTAAGCTCGGGTGGAGCGACCCCTAGTGTAGATCT
TGGTGGTAGTAGCAAATATTCAAGTGAGAACCTTGAAGACTGAAGTGGGGAAGGGTTCCA
CGCGAACAGTGCTTGGACGTGGGTTAGTCGGTCCTAAGAGATGGGGAAACTCCGTATTCA
AGTGCGCTTTTGTAGCGCAGTCCATCGAAAGGGAAGCGGGTTAACATTCCCGCACCAGGA
TGTGGAAACTGTGGTTTACCGCAAACTGCGGCAACGCAACTGAACCTGGGGACGCCCGTA
GGAGCTCCGAGCAGGGATCTCTTTCCTGTTTAACGCGCCCAACCCCCTGGAATCGGTTTG
TCCGGAGATAGGGGGCGAAAGGCCGGAAAAGCACCGCACGTCTTGTGGTGTTCGGAACGC
TTGTAATGGCCCTTGAAAACCCAGGGGAAAGCAGCCGTCTTACACTTGTGTGAGACGAGG
AATTCTCACACCTGGCCGTACCCATCACCGCATCAGGTCCCCAAGGTGAACAGCCTCTAG
TCAATAGAATAATGTAGATAAGGGAAGTCGGCAAAATAGATCCGTAACTTTGGGAAAAGG
ATTGGCTCTAATGTTCGGGCTTGTCGGGACGCGCGGGCGTAGATGCCAATGAGCGGCAGA
CGTCTAGCCTCTTTGGAGGCTGGATTAACGCTGCTTGTGGGAAGAAACCGTGTCGATGGA
GGCAGGAACGCTTATCGATCTTGTCTCTTCCCGCTTGCGATCAACGAACGGATTAGAACT
GGTACGGACAAGGGGAATCCGACTGTTTAATTAAAACAAAGCATTGCGACGGCCGGAAAC
GGTGTTGGCGCAATGTGATTTCTGCCCAGTGCTCTGAATGTCAAAGTGAAGAAATTCAAC
CAAGCGCGGGTAAACGGCGGGAGTAACTATGACTCTTAAACAAACCATACTACCTACGGG
AGTCAAAAGTGGGGGCGGCGTCTGGTGTAGAAATACACGTACGACACGCTTTTTTTCCTG
CTAGTCGATCTGACCTTTGTTCTTTGGACTGTGGGCGGTCAGCGGCAACATAACCTGGTA
CGGGGAAGGCCTCGACGTTGAGTGCTGCATCTTGGAAGTATCTTGACACAATAAGAGAAA
AGAGATTGCGCTTAATAGGGGGTTGATCTCGTGGCGAGCCGACGCCGGTGTGAACTGGTG
GCAATGGCCGTCGTAGAGCACGCAAAGGTATGGGCTGACTTTTTCTCCTCAAAAGTTGGC
TTAAGGGACGTGCCATCCCACACGATAATAAAGTGTGCAGTCTGTAAGAGCTCCCACAGG
GCAAAGACAGACTGGGATGCAGACGGAGACGGCAAAAACAACAACATCCGTTCTGCTTCT
GGGCGCAAATGCTTAAGGTAGCCAAATGCCTCGTCATCTAAACTAATTGGATGAGATAGT
CTGAGTGAAAAGCTTATGCAGGCTAGTACCGGAAAAAAAAAAAAAAAGAGGGGCTCTGCC
ACTCTGACGGTGCGACACTCCTTGTACAGGGAAACCCTCCGCATGGAAAGGCACGTGGTA
CCACCCTCGCGGGGAAACTCGCGAGCGGGATCCCGGGTAATGACCGGAGCCGATGGTAAA
AACTCACGTGACCGATGGGGTGACCCTGTGGCGAGACGCTAAGTTGTGGGACATCACCCT
CTCACAATACGCATCCGTCGCAGAGACTGAACGGGAGTGGGCTCCGTTGACGGCCGGCCG
GCCAGCCAGCCGTCTTCCAGGGGCTTAAGATACAGTCCGGGCCTCAGGGATGGAAAAACC

TGGGGAACTACCAGAATTAGTGACGCGCATGAATGGATTAACGAGATTCCCACTGTCCCT
ATCTACTATCTAGCGAAACCACAGCCAAGGGAACGGGCTTGGAAGAATCAGCGGGGAAAG
AAGACCCTGTTGAGCTTGACTCTAGTCCGACTTTGTGAGATGAGTTGGGAGGTGTAGTCT
AAGAGGGAGGGGCAACTCGACCGTGAAATACCTCTACTCCCAAGCTTGTTTTACTTACTC
CGGCGAGCAGGGATGGGTGGTAACACCCTTTTTCTAGGATTAACACCTCTTCGGGGGTGG
ACCTGCGCGGAGGACATTGTCAGGTGGGGAGTTTGGCTGGGGCGGCACATCTGTTAAAAG
ATAACGCAGGTGTCCTAAGATGAGCTCAGTGAGAACAGAAATCTCACGTAGAACAAAAGG
GTAAAAGCTCATTTGATTTTGATTTTCAGTACGAATACAAACTGTGAAAGCATGGCCTAT
CGATCCTTTTGACCTTCAAGGTTTGAAGCAAGAGGTGTCAGAAAAGTTACCACAGGGATA
ACTGGCTTGTGGCAGCCAAGCGTTCATAGCGACGTTGCTTTTTGATCCTTCGATGTCGGC
TCTTCCTATCATTGCGAAGCAGAATTCGCCAAGTGTTGGATTGTTCACCCACTAATAGGG
AACGTGAGCTGGGTAGCCACTTTATGCCCAACAGCTGGATAGCTGGGCTTCTGTCTTAGG
CCGGGACTTCTCACGACCCGTGCCTTGGTCACCCAAAAAAGGGCGACCGCAGAGCTCAGG
ATGGAGCTTCTGGCTTAACTAGTAGAGGACAGATAGAGATCGTCAAATTGCGGGAAACCC
CTTAGAGCCTCGAGCTACCAAGTGCGTGTCGAAAGGCATAGACGCAGCCAGCGATGGCTG
TCGCATGGCTTGATTCCATCAAGATAGGTAAAAATGCTCGAGGATTGGGCAATCCGCAGC
CAAGCCCCTAAGGTCCCCCCGCGGGCTATGGGGAAGGTTCAACGACTGTAAGGCGTACCG
TGCAAGCGGAAGATACAGTCTAGCCCCGCTGGAAAGCAGCGCCAAAGTAAGTAGGTTGCA
GTGGGTCTGTCCTC
>SRR9587930
GCATGTTTTGGTATGTGTTTCGACAGTCATGCATGTCATAGCCGTAGGATGGGGAGTGAG
CTTCATAGCCCGACCCCTTCTCAGGAGCATGTTTTCATAGCATATTGAGCATACTGGGGT
GGTCTCCCATGTAGCTCGCTCTCTCGGTGCAGGACCTCCCTTTTGCGGGCCACTTCGATG
GCCTCCATTATGGAGAGCATGTTTTGGCATATGAAGCGTTGGTGGTGAGGCTCGTAGCCT
GTATATACATACCAACGCAGTCGTCATATGTCTCTCTGGCGGGGGTCGGTCGTATGACCT
TGTCCCTACCTGGAGCATGTATTTCGTATGATATACTGGACGGCAGCTGCGCTGTTCAGT
CGTCGGCCGTACCTCATGTGCGCGTTTCCAGGTCCGAACGTTTGACATGCAAGGGCATCA
TACCAGAAGCCTGTTTTCTCGTATGAGCATGTAGGTAGGCAGGCTCTTCAGTCGTCTACC
CATAAATGCACTTCGAAGGAACTTTTTGGTATTGAATGGCTTTGTGCCCAGACGTTCTTG
TGAGCTTGTTCTCGCTCTGCGATACGCTGGAAGCTGTGGTATTACCTACAGCAGACGCAT
TCCTAGGGCGGACGGCCTCTCTGCAGATGGGCACATCTGTAGAAAGAGGCTCCCCTTGAT
GACGCAATGCAATGGCTGGGTAAACGGAGTGCTATACAGAAGGGTCAAACCTCTGTGGAA
GTACACTCATACCAGCATTGTTTGTTGAGAGGGGTCGGCTCTACGAGTGAGGACCTATAC
TGCCATCTCATTCAGATGCGTAGACGGTCTTTACTCAGACTGCCTTTCTGGCTGATTGTC
CTCGGTCCCTCGTGCGCACGCGCCGCTGGCTGAATGGGCTTGTCCTTATGGAGTGCCGCG
TTCTTCAAAGAATCCGTCTTCATACAGCGACCTCATTTACATATTGCCGGCTGAATGGGC

TTGTCCTTATGGAGCACTGCATTCTTCTAGGAATTGCTATCTATACAGCGACCTCATTTA
CATACAAGGCACCTTTTTCTACATAGACTTGCCATTTGGCTGATTGTTCTGACAGTCGCT
GCCTCTTCGGAGTTCTGTGTGGACGAGTGGCCTCCTGAGTGGGCCGGTCCTTACGGAGTG
CCCACGTCTGTTATGACTGGGTTCTCCATACAGCGACCTCACTCAGAATTGTATGGCACC
GTGCCGATTGAATGGACAAGCAGTCCTTAGGGAATCGCATGAAAGCATGATGGCACATGG
TGTATGGACCCGTTGGGCCTTGCCTGTGCTATCTTCTCGAGTCGACCGGTCCTCGTCTCC
TCCTTTCGCGACGGGAAGCTCTTGAGTTTACCGTGGGTCAGGACGCTGTGGTATCTGTTT
GTCTGTAGTGTGGGCTCTCTATCAGAACAGCGTATCAGCACACGTCGTCTGGGTCTATGG
TATTTTTGCTTAACGGCCCTGGGTTTGGTAGTATGGTTCCATGTCGGTCCTCTCACCTCC
CATCCGTCATGTCTCTGATATCATCTGCACAGTACGTCTTGTCAGAATATCATTCTGTAT
GGCAGTCGATAGACTAGGCCCTATGCCGCAGCTGTCTCTGACCACTATGCGAGCGCATAC
CTACCTCAATGTGTTCATAGCTGTTATGGATCTTGGGTGTGGTGTGCCAGAACGATACGC
CACCAGATTGTGGAATGGTGCTTGTAGTATTAGAAGAGGACATGTATGAAGAGCACGTTC
TTTTCTCTTATGATGGTGCCGCGGGTGGGGGCCGTTCTCATTCTGCTTGTCGGGTGCTCA
AGGTGCATGTAGGATTGCTGAACCCACTGAATGGTCATGAGAATGGTCATATGGTGAGGT
TCACTTCTCCATCGCTTTGTGTGTGTTGGATGTTTGGGTGGCTTCGGCCGCATGGAAAAT
CTCGGTCTGTAACTCATCAGGGAGGGTTTCCATGTGAGCAATCTCTCCAGTCGAACACAC
ACCCATGGGCGTCCCGCTAGGCAAAATGGAACTCTCGATCCGACACAGCCTGTGGAGTGT
CTGTTCCGTGTGGCGTGAAACGGCTTATACTGGATGTCTTCCCTTTGGACACCAGTCTGT
TGGTATGTACGCATACCATCTAGAGGGAGCAAGAAAAAGGCATGCAAGTGCCACAAAGGA
AGGGAGGCCTCCCTGCTACCGTGTGTCCTCTTGACGATCACGGGGGAGTATATAGAGACA
TTTGGACGAGATCCTTTTTTCACGAGGCGGTGCCCTCGCGCAAAGGTGTTGCTCTGCACG
CATCTTTATGATGATAGTTGCTCGCTCCTATGCGTCCACTCGTGAAAATACCTGCTCTCT
CTGTCTGTTTCCATGACACCCCCATGCGTTGTCATAGAGCACGCTCAGGTACAGAAAGTG
CTGTGACTTTCTTGGCCTGCTGTCACTGAGTCAGCCTGATACTTTGGAGGAATCTTTTCT
GGGCGCTGCCCAGCTGAGAGTTCTTATCAGGCTCTGGAGTGTGCATATGGCTATAATCAT
AATAGTGCATGTGGCCACGCAGAAAAAGCATCGACGTGGAATGGAGAAGAACACCATTCT
GAATGGAACGGTTCTTACTCTGCTTCCGTTGCTTGACGCTGTATGGGAATCGCTTTGCCC
TGGGGGTACACCTTCCATTCCATGACAAAGCATTTGATCTATGCCCCGAGGGTCATTCTC
TATGAAGAGCCCGAAACTTGTCTGAGTGAGAGGCGAAGGTAGCCCTTCATTTATGCGGGG
TCCGCCCTCACACTGTAGTCGGCACAGTCCCTTGTTTATGGACTGCGCAGCTCAATCTTC
GACTGGAGAGGAAAAACAAAACACAAGTATGACACGCACATACGTGTTTTCTAGGACTGT
GACCTGTCGGCTGCCTCGCGGTGGTTGGCATGATGCCCCAAGTGGGGGGGGGGAGAAACA
ATCAGTATGATTGATAAAAACCAGTGGGGAAAAAGGCGCGCTGTCGCACACCCTTGGTCA
CGGCCTCAAGTCGACGGCCCTCACTGGCCCACCCTGCGACTCCTGTTCGCTTTTGACAAG
ACACACCTGCAATACGGAGGAAATGCCTGAGCAGCTGTGCCGCAGCAACTGCGGACGCGT

AGCGATATGCATCCCTGTTGTTTGCGACTTGGTACAAAGAACGTGCGCCGGGAGACGGAT
TGCCCTTTGACTTTTCCACCCAGCCCGCGGCTCTATACCCTTGACGTACCTATTTACCTG
GTTGATCCTGCCAGTAGTCATATGCTTGTCTCAAAGACTAAGCCATGCATGTCTAAGTAT
AAACACTTTTATACTCGTGAAACTGCGAATGGCTCATTAAAACAGTTATAGTTCCTTTGG
GAACAACACTACTTGGATAACCGTAGTAATTCTAGAGCTAATACATGCCTCAACGCCCGA
CTCACGAAGGGTGGTATTTATTGGATAAAAAACCATCGTGCTCTGTCCAGAGCGCTTTGA
GATGATTCACAATAACTTGTCGGATCGCATGGCCTCGTGCTGGCGACTCCTCATTCAAAT
TTCTGCCCTATCAACTTTCGATGGTAGAGTATTGGTCTACCATGGTGTCGACGGGTGACG
GGGAATTAGGGTTCGATTCCGGAGAGGGAGCCTGAGAAACGGCTACCACATCCAAGGAAG
GCAGCAGGCGCGCAAATTACCCAATCCCGACTCGGGGAGGTAGTGACAAAAAATAACAAT
AGGGGGCCCTTTGGGTCTTCTAATTGGAATGAGAACAATTTAAATCCCTTATCGAGGATC
CATTGGAGGGCAAGTCTGGTGCCAGCAGCCGCGGTAATTCCAGCTCCAATAGCGTATATT
AAAGTTGTTGCAGTTAAAACGCTCGTAGTCGGACCTCGGAGCGCCCCCACGGGCCGGGCT
TTTGCCTTGTTGCCAGTGGGAGTTTGTCTCCTTTTGTCGGGACGCGGTGTGCGGGGCCTT
TACTGTCTCTGCGCGTTTACCTGCGGGCCGACCGTTTACTGTGAAGAAAGTAAAGTGTTC
AAAGCAGGCCAATGCCTTGAATATGTGAGCATGGAATAATAGAATAGGACTTGGGCTCTA
TTTTGTTGGTTTCCAGTGACCAAGTAATGATTAATAGGGACGGTTGGGGGCATTCGTATT
TCATTGTCAGAGGTGAAATTCTTGGATTGATGGAAGACGTACAACTGCGAAAGCATCTGC
CATGGATGTTTTCATTGATCAAGAACGAAAGTTAGGGGATCGAAGACGATCAGATACCGT
CGTAGTCTTAACCATAAACGATGCCGACTGGGGATTGGCGGGGGCACTTTTATATGACTC
TGTCAGCACCCTGAGGGAAACCAAAGTCTTTGGGTTCTGGGGGGAGTATGGTCGCAAGGC
TGAAACTTAAAGGAATTGACGGAAGGGCACCACCAGGAGTGGAGCCTGCGGCTTAATTTG
ACTCAACACGGGAAAACTTACCAGGTCCGGACAGAAGAATGATTGACAGACTGAAAGCTC
TTTCTTGATTTTTTGGTTGGTGGTGCATGGCCGTTCTTAGTTGGTGGAGTGATTTGTCTG
GTTAATTCCGTTAACGAACGAGACCTCGTCCTGCTAAATAGGTGCGCGCATGCCAGCACT
GCGTTCTTACCTTCTTAGAGGGACTATGCGCGTCTAGCGTATGGAAGATTGAGGCAATAA
CAGGTCTGTGATGCCCTTAGATGTTCTGGGCCGCACGCGCGCTACACTGATGCATTCAAC
GAGTTTCACTTGATAGTCCTGGGTCGGAAGGCTCGGGTAATCTTTTGAAAGTGCATCGTG
CTGGGGATAGATCATTGCAATTATTGATCTTCAACGAGGAATTCCTTGTAGGCGCAGGTC
ATCAGCCTGCGCCGAATACGTCCCTGCCCTTTGTACACACCGCCCGTCGCTCCTACCGAT
TGAATGGTCCGATGAAATGTCCGGATCGCGGCTAGTGTGATGGACTCGTCCAGAGCGGTA
GCCGTGAGAAGCTCATTAAATCTTATCATTTAGAGGAAGGAGAAGTCGTAACAAGGTTTC
CGTAGGTGAACCTGCGGAAGGATCATTCACAAACTATTGACACAAACACACGCGAACCAA
ATCGTCCGCACAGCAGGTGGCAATGAAAGAGAGACTCTGCATGTCGCCTTTCGGGGTAGC
AAGCAGCACTCTTCTGCCATCGCCTCTGTGCAGGGCGTAAATTCCCATTGAGAGGATGTG
AGGGCACCACAGGAAGCTTTTCCACAGGAAGTCGCCATCCTTCTCCCTCCACGGCGGCGC

TTCTGTGCTTGGCAGTTTTTTTTTGCCTTCTAGGGAGGATGCCGCCAATGGAGCCCCTTA
TATATATACATCATCATATGCCCCTTTTTTTCTTAACCGCTTGCCACAGCTTCTTCTATG
AGGAGCTTGTGGGAAGACTGTCTCCATACAATAACAAAGATACAACTCTTAGCGGTGGAT
ATCTTGGTTCTCGCAACGATGAAGAACGCAGCCAACTGCGATAACTAATGTGAATTGCAG
GACTTCGTGAATCATTAAGTCTTTGAACGCAAGTTGCGCTCATCTCCATGAGGAGGTGAG
CATGCCTGTTTGAGTGTCGATGGGAAAAACAATACATAACAACAATCGGCTCTCTTATCT
TCCTGCCTTCTTGGGGGCAAAAGGTAGTGCATGCCGGAGATGGGCTTTGTCGCCCCTTTC
TCTTGAAAGAGGCGTCGTCCGAAACACGTATCGTTGCCCGCTGGAGGTCATAGAGCATGA
GTTTAAAGCACCACATCTCTCAAGTCAGAGTGAGCGTCTTGAGTAGGTCCGCATTGATTC
AGTTTCAACCCAGGTGGCGTTGGTTACTTGCCATAAGCCTCTCTGGGGTTTCTTTGCTTG
GACCTCTCTTGTCAGTTTCTCTGACGTAGAGACAGGTGCCGTCTCATGGACGTTGCCCTC
TGGAACGTGCTAAAAAAAAAAAAAAGAAAAAACCGTACCACATGTCAAAAGAGGCAGATC
GTGCTTGACGGCGGGAGGTGTTGCATTTCTTGCTTGTCCTGAGCGTCTGCCAGTCTCTGA
CGAGTGTGTTCCTTCCATAAACCCGATCTCAAATCAGGTAGGACCACCCGCTGAATTTAA
GCATATAATTAAGCGGAGGAAAAGAAACTAACAAGGATTCCCTCAGTAGCGGCGAGCGAA
GCGGGAAGAGCTCAACATGAAAACCGTTGGGGGTTCGCGCTTTCCTTCGGGTTGTATAGT
AAAGTGGTGTCGCCTGCGTCGCGCCGGGTATAAATCCCGTCTGAAATGGGGTGTCGTAGA
GGGTTATATCCCCGTCTGTGACCCGGTGTGCGATGCGTTACGGTGACCTTACGATGAGTC
GGGTTGCTTGGGCATGCAGCCCTAAGTGGGTGGTAAACTTCATCTAAAGCTAAATACTAG
CGAGAGACCGATAGCGTACAAGTACCGTAAGGGAAAGATGCAAAGAACTTTGAAAAGAGA
GTTAAACAGGGCTTGAAACCGTTAAGAGTGAAGCGAAGGAGGTCAGTGATCATGGGGCCA
GATCCCCAAGTGCCTGTCATAGGGCACCGTAGCGTTCTGGTTTCCCGGGGCAGCGCCGGC
CGGGGGCGGGGGTTAACAGGGAGGTGGTTGAGGGATTCGTTTTCCGCCTGTACTCCTTTA
TAGCTTGCACCCCGGCCGAGGGATCCGCGACTTGTCGTGTGAGGGCGCTGTCAAAATGGC
CTTCTCCACCCCGTCTTGAAACACGGACCAAGGAGTCTAACATACTCGCGAGTGTTTGGG
TGGTTAAAACCCGAACGCACAATGAAAGTAAACGGTGGCTCTGCGCCGCTGCTTGGTGGG
AACCCTGCTTTTATAATGGGGGGCACTATCACCCGACCCCGATCCCTTTGGGGTGGTTGG
TTTTGAGGTAGAGCGAGTATGATGGGACCCGAAAGATGGTGAACTATTCCTGAGTAGGGC
GAAGCCAGAGGAAACTCTGGTGGAGGCTCGTAGCGATGCTGACGTGCAAATCGCTCGTCG
AACTTGGGTATAGGGGCGAAAGACTAATCGAACCATCTAGTAGCTGGTTCCCTCCGAAGT
TTCCCTCAGGATAGCGGGAGCAAATGCAAGTCAGTATCATCCGGTAAAGCGAATGATTAG
AGGCCTCGGGGGAGTAACTCCTTCGACCTATTCTCAAACTTTAAATAGGTGAGGAGCCGG
CAACGTTGCTTAAAACAATGAACGCCGTGCGGACAATGATTGCTCTCAGTGGGCCGATTT
TGGTAAGCAGAACTGGCGATGCGGGATGAACCGAAAGCGACGCTAAGGTGCCAAACTCCC
AGCTCATTCAGACACCAGAAAAGGTGTTGATTCATACAGACAGCAGGACGGTGGCCATGG
AAGTCGGAATCCGCTAAGGAGTGTGTAACAACTCACCTGCCGAATGGATCAGCCCTGAAA

ATGGATGGCGCTTAAGCGGGGGACCGAGGCGTCGCCGTTGGGGTAAATACGAAGCCCCAA
TGAGTAGGAGGGCGTGGGGGTTTTGAAACACGGTACACTCGCATGGAAGCCTTGGGCGTA
AGCTCGGGTGGAGCGACCCCTAGTGTAGATCTTGGTGGTAGTAGCAAATATTCAAGTGAG
AACCTTGAAGACTGAAGTGGGGAAGGGTTCCACGCGAACAGTGCTTGGACGTGGGTTAGT
CGGTCCTAAGAGATGGGGAAACTCCGTATTCAAGTGCGCTTTTGTAGCGCAGTCCATCGA
AAGGGAAGCGGGTTAACATTCCCGCACCAGGATGTGGAAACTGTGGTTTACCGCAAACTG
CGGCAACGCAACTGAACCTGGGGACGCCCGTAGGAGCTCCGAGCAGGGATCTCTTTCCTG
TTTAACGCGCCCAACCCCCTGGAATCGGTTTGTCCGGAGATAGGGGGCGAAAGGCCGGAA
AAGCACCGCACGTCTTGTGGTGTTCGGAACGCTTGTAATGGCCCTTGAAAACCCAGGGGA
AAGCAGCCGTCTTACACTTGTGTGAGACGAGGAATTCTCACACCTGGCCGTACCCATCAC
CGCATCAGGTCCCCAAGGTGAACAGCCTCTAGTCAATAGAATAATGTAGATAAGGGAAGT
CGGCAAAATAGATCCGTAACTTTGGGAAAAGGATTGGCTCTAATGTTCGGGCTTGTCGGG
ACGCGCGGGCGTAGATGCCAATGAGCGGCAGACGTCTAGCCTCTTTGGAGGCTGGATTAA
CGCTGCTTGTGGGAAGAAACCGTGTCGATGGAGGCAGGAACGCTTATCGATCTTGTCTCT
TCCCGCTTGCGATCAACGAACGGATTAGAACTGGTACGGACAAGGGGAATCCGACTGTTT
AATTAAAACAAAGCATTGCGACGGCCGGAAACGGTGTTGGCGCAATGTGATTTCTGCCCA
GTGCTCTGAATGTCAAAGTGAAGAAATTCAACCAAGCGCGGGTAAACGGCGGGAGTAACT
ATGACTCTCTTAAGGTAGCCAAATGCCTCGTCATCTAAACTAATTGGATGAGATAGTCTG
AGTGAAAAGCTTATGCAGGCTAGTACCGGAAAAAAAAAAAAAAAGAGGGGCTCTGCCACT
CTGACGGTGCGACACTCCTTGTACAGGGAAACCCTCCGCATGGAAAGGCACGTGGTACCA
CCCTCGCGGGGAAACTCGCGAGCGGGATCCCGGGTAATGACCGGAGCCGATGGTAAAAAC
TCACGTGACCGATGGGGTGACCCTGTGGCGAGACGCTAAGTTGTGGGACATCACCCTCTC
ACAATACGCATCCGTCGCAGAGACTGAACGGGAGTGGGCTCCGTTGACGGCCGGCCGGCC
AGCCAGCCGTCTTCCAGGGGCTTAAGATACAGTCCGGGCCTCAGGGATGGAAAAACCTGG
GGAACTACCAGAATTAGTGACGCGCATGAATGGATTAACGAGATTCCCACTGTCCCTATC
TACTATCTAGCGAAACCACAGCCAAGGGAACGGGCTTGGAAGAATCAGCGGGGAAAGAAG
ACCCTGTTGAGCTTGACTCTAGTCCGACTTTGTGAGATGAGTTGGGAGGTGTAGTCTAAG
AGGGAGGGGCAACTCGACCGTGAAATACCTCTACTCCCAAGCTTGTTTTACTTACTCCGG
CGAGCAGGGATGGGTGGTAACACCCTTTTTCTAGGATTAACACCTCTTCGGGGGTGGACC
TGCGCGGAGGACATTGTCAGGTGGGGAGTTTGGCTGGGGCGGCACATCTGTTAAAAGATA
ACGCAGGTGTCCTAAGATGAGCTCAGTGAGAACAGAAATCTCACGTAGAACAAAAGGGTA
AAAGCTCATTTGATTTTGATTTTCAGTACGAATACAAACTGTGAAAGCATGGCCTATCGA
TCCTTTTGACCTTCAAGGTTTGAAGCAAGAGGTGTCAGAAAAGTTAACACGCGTTGTGGT
AGCTTTTAAGAGTGGACTCAAAGTGTTCCGCTAGTCCCCCCTCGGACCGCCACTTACAAA
AGGCCTGGGGATGGGGGCGACACTGTCAAATTGCGGGGACTCCCTAAAGCCTGTCTCTAC
CAAGCCAGCGCGGCGAAAGCCCCTGGTGGCCGGGGTAATGACCTAGGGTATGGTAAAAGC

GAGACAGGATGGAACAATGGGCAATCCGCAGCCAAGTCCTAAGGCTTTTTGAAAAGCGGC
TATGGATGCAGTTCAACGACTAGATGGCAGTGGGCTGAGAGAGCCCACCGCGGCACCCTT
TCGGCTTAAGATATAGTCTTGCCCACGTGAAAGCGTGCCTGCAGATGAACGCCACCATGT
ATGGCCGGAGCTGCAGGAGACCGCACTTGATGTGCAGTCGGGAATCGACAGTACCACAGG
GATGAACAAAAACACACAATTAATTTGTGGCCATCATACAGTCGTATGATATTATCGTCC
CAGGCGGGTGCGCCTCTCTCACCCGTCTAGTCCGGCCTTGGTGCGCAAAAAATGCTTGCG
TGGATGCGCCTTGGTTAAGGAAAATGCTCCTAGTAGTTTTACAAGGACACTTCTTGTAGG
GCAACACATCTGGATGCGAAGAACTCCCGGAAAAGACAACGGTACCAAGTGACTGGGGTA
ACGCCCCATTCATGAAGGCAAATAATCAAAAAGAGGTGCCGGGTAAGAACCCGTTGTATA
GGGACAATCCGCAGCTGACTTGGCTCCCTGCCCATCCATGGGCATAGCTTAGCGGTTCAA
CGCTCGGTAAGATGTGGGTTGGCGCGCTGTCCGACAGCGCACTGGCTTAAGGTACGGGCT
ACTCCCACCGGAGACGGTGTCTTTGTCTACCTCGGGTAACACGCCTCGGGAGAAGCAAAG
AGGGTGCAATGGAAAAAAAACTGGTTGCGGGGAAATCCCTACCAGAGTCACCTGGTACAA
AGGAACTGGCTTGTGGCAGCCAAGCGTTCATAGCGACGTTGCTTTTTGATCCTTCGATGT
CGGCTCTTCCTATCATTGCGAAGCAGAATTCGCCAAGTGTTGGATTGTTCACCCACTAAT
AGGGAACGTGAGCTGGGTAGCCACTTTATGCCCAACAGCTGGATAGCTGGGCTTCTGTCT
TAGGCCGGGACTTCTCACGACCCGTGCCTTGGTCACCCAAAAAAGGGCGACCGCAGAGCT
CAGGATGGAGCTTCTGGCTTAACTAGTAGAGGACAGATAGAGATCGTCAAATTGCGGGAA
ACCCCTTAGAGCCTCGAGCTACCAAGTGCGTGTCGAAAGGCATAGACGCAGCCAGCGATG
GCTGTCGCATGGCTTGATTCCATCAAGATAGGTAAAAATGCTCGAGGATTGGGCAATCCG
CAGCCAAGCCCCTAAGGTCCCCCCGCGGGCTATGGGGAAGGTTCAACGACTGTAAGGCGT
ACCGTGCAAGCGGAAGATACAGTCTAGCCCCGCTGGAAAGCAGCGCCAAAGTAAGTAGGT
TGCAGTGGGTCTGTCCTC
>SRR9587931
GCATGTTTTGGTATGTGTTTCGACAGTCATGCATGTCATAGCCGTAGGATGGGGAGTGAG
CTTCATAGCCCGACCCCTTCTCAGGAGCATGTTTTCATAGCATATTGAGCATACTGGGGT
GGTCTCCCATGTAGCTCGCTCTCTCGGTGCAGGACCTCCCTTTTGCGGGCCACTTCGATG
GCCTCCATTATGGAGAGCATGTTTTGGCATATGAAGCGTTGGTGGTGAGGCTCGTAGCCT
GTATATACATACCAACGCAGTCGTCATATGTCTCTCTGGCGGGGGTCGGTCGTATGACCT
TGTCCCTACCTGGAGCATGTATTTTGTATGATATACTGGACGGCAGCTGCGCTGTTCAGT
CGTCGGCCGTACCTCATGTGCGCGTTTCCAGGTCCGAACGTTTGACATGCAAGGGCATCA
TACCAGAAGCCTGTTTTCTCGTATGAGCATGTAGGTAGGCAGGCTCTTCAGTCGTCTACC
CATAAATGCACTTCGAAGGAACTTTTTGGTATTGAATGGCTTTGTGCCCAGACGTTCTTG
TGAGCTTGTTCTCGCTCTGCGATACGCTGGAAGCTGTGGTATTACCTACAGCAGACGCAT
TCCTAGGGCGGACGGCCTCTCTGCAGATGGGCACATCTGTAGAAAGAGGCTCCCCTTGAT
GACGCAATGCAATGGCTGGGTAAACGGAGTGCTATACAGAAGGGTCAAACCTCTGTGGAA

GTACACCCATACCAGCATTGTTTGTTGAGAGGGGTCGGCTCTACGAGTGAGGACCTATAC

TGCCATCTCATTCAGATGCGTAGACGGTCTTTACTCAGACTGCCTTTCTGGCTGATTGTC

CTCGGTCCCTCGTGCGCACGCGCCGCTGGCTGAATGGGCTTGTCCTTATGGAGTGCCGCG

TTCTTCAAAGAATCCGTCTTCATACAGCGACCTCATTTACATATTGCCGGCTGAATGGGC

TTGTCCTTATGGAGCACTGCATTCTTCTAGGAATTGCTATCTATACAGCGACCTCATTTA

CATACAAGGCACCTTTTTCTACATAGACTTGCCATTTGGCTGATTGTTCTGACAGTCGCT

GCCTCTTCGGAGTTCTGTGTGGACGAGTGGCCTCCTGAGTGGGCCGGTCCTTACGGAGTG

CCCACGTCTGTTATGACTGGGTTCTCCATACAGCGACCTCACTCAGAATTGTATGGCACC

GTGCCGATTGAATGGACAAGCAGTCCTTAGGGAATCGCATGAAAGCATGACGGCACATGG

TGTATGGACCCGTTGGGCCTTGCCTGTGCTATCTTCTCGAGTCGACCGGTCCTCGTCTCC

TCCTTTCGCGACGGGAAGCTCTTGAGTTTACCGTGGGTCAGGACGCTGTGGTATCTGTTT

GTCTGTAGTGTGGGCTCTCTATCAGAACAGCGTATCAGCACACGTCGTCTGGGTCTATGG

TATTTTTGCTTAACGGCCCTGGGTTTGGTAGTATGGTTCCATGTCGGTCCTCTCACCTCC

CAACCGTCATGTCTCTGATATCATCTGCACAGTACGTCTTGTCAGAATATCATTCTGTAT

GGCAGTCGATAGACTAGGCCCTATGCCGCAGCTGTCTCTGACCACTATGCGAGCGCATAC

CTACCTCAATGTGTTCATAGCTGTTATGGATCTTGGGTGTGGTGTGCCAGAACGATACGC

CACCAGATTGTGGAATGGTGCTTGTAGTATTAGAAGAGGACATGTATGAAGAGCACGTTC

TTTTCTCTTATGATGGTGCCGCGGGTGCGGGCCGTTTTCATTCTGCTTGTCGGGTGCTCA

AGGTGCATGTAGGATTGCTGAACCCACTGAATGGTCATGAGAATGGTCATATGGTGAGGT

TCACTTCTCCATCGCTTTGTGTGTGTTGGATGTTTGGGTGGCTTCGGCCGCATGGAAAAT

CTCGGTCTGTAACTCATCAGGGAGGGTTTCCATGTGAGCAATCTCTCCAGTCGAACACAC

ACCCATGGGCGTCCCGCTAGGCAAAATGGAACTCTCGATCCGACACAGCCTGTGGAGTGT

TTGTTCCGTGTGGCGTGAAACGGCTTATACTGGATGTCTTCCCTTTGGACACCAGTCTGT

TGGTATGTACGCATACCATCTAGAGGGAGCAAGAAAAAGGCATGCAAGTGCCACAAAGGA

AGGGAGGCCTTCCTGCTACCGTGTGTCCTCTTGACGATCACGGGGGAGTATATAGAGACA

TTTGGACGAGATCCGTTTTTCACGAGGCGGTGCCCTCGCGCAAAGGTGTTGCTCTGCACG

CATCTTTATGATGATAGTTGCTCGCTCCTATGCGTCCACTCGTGAAAATACCTGCTCTCT

CTGTCTGTTTCCATGACACCCCCATGCGTTGTCATAGAGCATGCTCAGGTACAGAAAGTG

CTGTGACTTTCTTGGCCTGCTGTCACTGAGTCAGCCTGATACTTTGGAGGAATCTTTTCT

GGGCACTGCCCAGCTGAGAGTTCTTATCAGGCTCTGGAGTGTGCATATGCCTATATTCAT

TATAGTGCATGTGGCTACGCAGAAAAAGCATCGACGTGGAATGGAGAAGAACACCATTCT

GAATGGAACGGTTCTTACTCTACTTCTGTTGCTTGACGCTGTATGGGAATCGCTTTGCTC

TGGGGGTACACCTTCCATTCCATGACAAAGCATTTGATCTATGCCCCGAGGGTCATTCTC

TATGAAGAGCCCGAAACTTGTCTGAGTGAGAGGCGAAGGTAGTCCTTCATTTATGCGGGG

TCCGCCCTCACACTGTAGTCGGCACAGTCCCTTGTTTATGGACTGCGCAGCTCAATCTTC

GACTGGAGAGGAAAAACAAAACACAAGTATGACACGCACATACGTGTTTTCTAGGACTGT

GACCTGTCGGCTGCCTCGCGGTGGTTGGCATGATGCCCCAAGTGGGGGGGGGAGAAACAA

TCAGTATGATTGATAAAAACCAGTGGGGAAAAAGGCGCGCTGTCGCACACCCTTGGTCAC

GGCCTCAAGTCGACGGCCCTCACTGGCCCACCCTGCGACTCCTGTTCGCTTTTGACAAGA

CACACCTGCAATACGGAGGAAATGCCTGAGCAGCTGTGCCGCAGCAACTGCGGACGCGTA

GCGATATGCATCCCTGTTGTTTGCGACTTGGTACAAAGAACGTGCGCCGGGAGACGGATT

GCCCTTTGACTTTTCCACCCAGCCCGCGGCTCTATACCCTTGACGTACCTATTTACCTGG

TTGATCCTGCCAGTAGTCATATGCTTGTCTCAAAGACTAAGCCATGCATGTCTAAGTATA

AACACTTTTATACTCGTGAAACTGCGAATGGCTCATTAAAACAGTTATAGTTCCTTTGGG

AACAACACTACTTGGATAACCGTAGTAATTCTAGAGCTAATACATGCCTCAACGCCCGAC

TCACGAAGGGTGGTATTTATTGGATAAAAAACCATCGTGCTCTGTCCAGAGCGCTTTGAG

ATGATTCACAATAACTTGTCGGATCGCATGGCCTCGTGCTGGCGACTCCTCATTCAAATT

TCTGCCCTATCAACTTTCGATGGTAGAGTATTGGTCTACCATGGTGTCGACGGGTGACGG

GGAATTAGGGTTCGATTCCGGAGAGGGAGCCTGAGAAACGGCTACCACATCCAAGGAAGG

CAGCAGGCGCGCAAATTACCCAATCCCGACTCGGGGAGGTAGTGACAAAAAATAACAATA

GGGGGCCCTTTGGGTCTTCTAATTGGAATGAGAACAATTTAAATCCCTTATCGAGGATCC

ATTGGAGGGCAAGTCTGGTGAACGTATGCATTAAAGCCCTCCGTAATAAAAAAAAACAGC

GCCAGATGTCTCTCTCTGGAAGCACACCATGCATGGGGTTTGTTCCTCTCTGCATGGCCG

TGTGAAAAAGGCGAGTAGGCAAACGATGGGAAACCCGGCTTGGAATAGTAACCAGCCACA

GAGCCCCCATCATACACTTCTAAATTGCCGGGGAGTGCCCGCGAGGCCTTGACTACCGCG

GTCCGCCGAGAGGTGAGAGCGAGCAATCGTTCAGGACGCAGCAGTGTACGTAGTGGTACA

TGGATGGTAAAAACGTCAAGGATAGGGTTGATCCGCAGGGAAGCGCGAGTGCCGCCATGG

TCGGTGGGCAAAGGCGAACCTTCAGAGACTCTAATGGAGTGGGCGCAAGCTTAAGGGAGA

GTCCAATCCACTGGGAAACCAGGTCCACAGCAGTAACGGGGTGATGCTCGCCCCAGAGGA

CTGTGGAAGGCAACCCCACCAAAAGGGTGTGTGTTGTCGGGTAGCAATGGCCAGCAGCCG

CGGTAATTCCAGCTCCAATAGCGTATATTAAAGTTGTTGCAGTTAAAACGCTCGTAGTCG

GACCTCGGAGCGCCCCCACGGGCCGGGCTTTTGCCTTGTTGCCAGTGGGAGTTTGTCTCC

TTTTGTCGGGACGCGGTGTGCGGGGCCTTTACTGTCTCTGCGCGTTTACCTGCGGGCCGA

CCGTTTACTGTGAAGAAAGTAAAGTGTTCAAAGCAGGCCAATGCCTTGAATATGTGAGCA

TGGAATAATAGAATAGGACTTGGGCTCTATTTTGTTGGTTTCCAGTGACCAAGTAATGAT

TAATAGGGACGGTTGGGGGCATTCGTATTTCATTGTCAGAGGTGAAATTCTTGGATTGAT

GGAAGACGTACAACTGCGAAAGCATCTGCCATGGATGTTTTCATTGATCAAGAACGAAAG

TTAGGGGATCGAAGACGATCAGATACCGTCGTAGTCTTAACCATAAACGATGCCGACTGG

GGATTGGCGGGGGCACTTTTATATGACTCTGTCAGCACCCTGAGGGAAACCAAAGTCTTT

GGGTTCTGGGGGGAGTATGGTCGCAAGGCTGAAACTTAAAGGAATTGACGGAAGGGCACC

ACCAGGAGTGGAGCCTGCGGCTTAATTTGACTCAACACGGGAAAACTTACCAGGTCCGGA

CAGAAGAATGATTGACAGACTGAAAGCTCTTTCTTGATTTTTTGGTTGGTGGTGCATGGC

CGTTCTTAGTTGGTGGAGTGATTTGTCTGGTTAATTCCGTTAACGAACGAGACCTCGTCC
TGCTAAATAGGTGCGCGCATGCCAGCACTGCGTTCTTACCTTCTTAGAGGGACTATGCGC
GTCTAGCGTATGGAAGATTGAGGCAATAACAGGTCTGTGATGCCCTTAGATGTTCTGGGC
CGCACGCGCGCTACACTGATGCATTCAACGAGTTTCACTTGATAGTCCTGGGTCGGAAGG
CTCGGGTAATCTTTTGAAAGTGCATCGTGCTGGGGATAGATCATTGCAATTATTGATCTT
CAACGAGGAATTCCTTGTAGGCGCAGGTCATCAGCCTGCGCCGAATACGTCCCTGCCCTT
TGTACACACCGCCCGTCGCTCCTACCGATTGAATGGTCCGATGAAATGTCCGGATCGCGG
CTAGTGTGATGGACTCGTCCAGAGCGGTAGCCGTGAGAAGCTCATTAAATCTTATCATTT
AGAGGAAGGAGAAGTCGTAACAAGGTTTCCGTAGGTGAACCTGCGGAAGGATCATTCACA
AACTATTGACACAAACACACGCGAACCAAATCGTCCGCACAGCAGGTGGCAATGAAAGAG
AGACTCTGCATGTCGCCTTTCGGGGTAGCAAGCAGCACTCTTCTGCCATCGCCTCTGTGC
AGGGCGTAAATTCCCATTGAGAGGATGTGAGGGCACCAAAGGAAGCTTTTCCACAGGAAG
TCGCCATCCTTCTCCCTCCACGGCGGCGCTTCTGTGCTTGGCAGTTTTTTTTGCCTTCTA
GGGAGGATGCCGCCAATGGAGCCCCTTATATATATACATCATCATATGCCCCTTTTTTTC
TTAACCGCTTGCCACAGCTTCTTCTATGAGGAGCTTGTGGGAAGACTGTCTCCATACAAT
AACAAAGATACAACTCTTAGCGGTGGATATCTTGGTTCTCGCAACGATGAAGAACGCAGC
CAACTGCGATAACTAATGTGAATTGCAGGACTTCGTGAATCATTAAGTCTTTGAACGCAA
GTTGCGCTCATCTCCATGAGGAGGTGAGCATGCCTGTTTGAGTGTCGATGGGAAAAACAA
TACATAACAACAATCGGCTCTCTTATCTTCCTGCCTTCTTGGGGGCAAAAGGTAGTGCAT
GCCGGAGATGGGCTTTGTCGCCCCTTTCTCTTGAAAGAGGCGTCGTCCGAAACACGTATC
GTTGCCCGCTGGAGGTCATAGAGCATGAGTTTAAAGCACCACATCTCTCAAGTCAGAGTG
AGCGTCTTGAGTAGGTCCGCATTGATTCAGTTTCAACCCAGGTGGCGTTGGTTACTTGCC
ATAAGCCTCTCTGGGGTTTCTTTGCTTGGACCTCTCTTGTCAGTTTCTCTGACGTAGAGA
CAGGTGCCGTCTCATGGACGTTGCCCTCTGGAACGTGCTAAAAAAAAAAAAAAAGAAAAAA
CCGTACCACATGTCAAAAGAGGCAGATCGTGCTTGACGGCGGGAGGTGTTGCATTTCTTG
CTTGTCCTGAGCGTCTGCCAGTCTCTGACGAGTGTGTTCCTTCCATAAACCCGATCTCAA
ATCAGGTAGGACCACCCGCTGAATTTAAGCATATAATTAAGCGGAGGAAAAGAAACTAAC
AAGGATTCCCTCAGTAGCGGCGAGCGAAGCGGGAAGAGCTCAACATGAAAACCGTTGGGG
GTTCGCGCTTTCCTTCGGGTTGTATAGTAAAGTGGTGTCGCCTGCGTCGCGCCGGGTATA
AATCCCGTCTGAAATGGGGTGTCGTAGAGGGTTATATCCCCGTCTGTGACCCGGTGTGCG
ATGCGTTACGGTGACCTTACGATGAGTCGGGTTGCTTGGGCATGCAGCCCTAAGTGGGTG
GTAAACTTCATCTAAAGCTAAATACTAGCGAGAGACCGATAGCGTACAAGTACCGTAAGG
GAAAGATGCAAAGAACTTTGAAAAGAGAGTTAAACAGGGCTTGAAACCGTTAAGAGTGAA
GCGAAGGAGGTCAGTGATCATGGGGCCAGATCCCCAAGTGCCTGTCATAGGGCACCGTAG
CGTTCTGGTTTCCCGGGGCAGCGCCGGCCGGGGGCGGGGGTTAACAGGGAGGTGGTTGAG
GGATTCGTTTTCCGCCTGTACTCCTTTATAGCTTGCACCCCGGCCGAGGGATCCGCGACT

TGTCGTGTGAGGGCGCTGTCAAAATGGCCTTCTCCACCCCGTCTTGAAACACGGACCAAG

GAGTCTAACATACTCGCGAGTGTTTGGGTGGTTAAAACCCGAACGCACAATGAAAGTAAA

CGGTGGCTCTGCGCCGCTGCTTGGTGGGAACCCTGCTTTTATAATGGGGGGCACTATCAC

CCGACCCCGATCCCTTTGGGGTGGTTGGTTTTGAGGTAGAGCGAGTATGATGGGACCCGA

AAGATGGTGAACTATTCCTGAGTAGGGCGAAGCCAGAGGAAACTCTGGTGGAGGCTCGTA

GCGATGCTGACGTGCAAATCGCTCGTCGAACTTGGGTATAGGGGCGAAAGACTAATCGAA

CCATCTAGTAGCTGGTTCCCTCCGAAGTTTCCCTCAGGATAGCGGGAGCAAATGCAAGTC

AGTATCATCCGGTAAAGCGAATGATTAGAGGCCTCGGGGGAGTAACTCCTTCGACCTATT

CTCAAACTTTAAATAGGTGAGGAGCCGGCAACGTTGCTTAAAACAATGAACGCCGTGCGG

ACAATGATTGCTCTCAGTGGGCCGATTTTGGTAAGCAGAACTGGCGATGCGGGATGAACC

GAAAGCGACGCTAAGGTGCCAAACTCCCAGCTCATTCAGACACCAGAAAAGGTGTTGATT

CATACAGACAGCAGGACGGTGGCCATGGAAGTCGGAATCCGCTAAGGAGTGTGTAACAAC

TCACCTGCCGAATGGATCAGCCCTGAAAATGGATGGCGCTTAAGCGGGGGACCGAGGCGT

CGCCGTTGGGGTAAATACGAAGCCCCAATGAGTAGGAGGGCGTGGGGGTTTTGAAACACG

GTACACTCGCATGGAAGCCTTGGGCGTAAGCTCGGGTGGAGCGACCCCTAGTGTAGATCT

TGGTGGTAGTAGCAAATATTCAAGTGAGAACCTTGAAGACTGAAGTGGGGAAGGGTTCCA

CGCGAACAGTGCTTGGACGTGGGTTAGTCGGTCCTAAGAGATGGGGAAACTCCGTATTCA

AGTGCGCTTTTGTAGCGCAGTCCATCGAAAGGGAAGCGGGTTAACATTCCCGCACCAGGA

TGTGGAAACTGTGGTTTACCGCAAACTGCGGCAACGCAACTGAACCTGGGGACGCCCGTA

GGAGCTCCGAGCAGGGATCTCTTTCCTGTTTAACGCGCCCAACCCCCTGGAATCGGTTTG

TCCGGAGATAGGGGGCGAAAGGCCGGAAAAGCACCGCACGTCTTGTGGTGTTCGGAACGC

TTGTAATGGCCCTTGAAAACCCAGGGGAAAGCAGCCGTCTTACACTTGTGTGAGACGAGG

AATTCTCACACCTGGCCGTACCCATCACCGCATCAGGTCCCCAAGGTGAACAGCCTCTAG

TCAATAGAATAATGTAGATAAGGGAAGTCGGCAAAATAGATCCGTAACTTTGGGAAAAGG

ATTGGCTCTAATGTTCGGGCTTGTCGGGACGCGCGGGCGTAGATGCCAATGAGCGGCAGA

CGTCTAGCCTCTTTGGAGGCTGGATTAACGCTGCTTGTGGGAAGAAACCGTGTCGATGGA

GGCAGGAACGCTTATCGATCTTGTCTCTTCCCGCTTGCGATCAACGAACGGATTAGAACT

GGTACGGACAAGGGGAATCCGACTGTTTAATTAAAACAAAGCATTGCGACGGCCGGAAAC

GGTGTTGGCGCAATGTGATTTCTGCCCAGTGCTCTGAATGTCAAAGTGAAGAAATTCAAC

CAAGCGCGGGTAAACGGCGGGAGTAACTATGACTCTTAAACAAACCATACTACCTACGGG

AGTCAAAAGTGGGGGCGGCGTCTGGTGTAGAAATACACGTACGACACGCTTTTTTTCCTG

CTAGTCGATCTGACCTTTGTTCTTTGGACTGTGGGCGGTCAGCGGCAACATAACCTGGTA

CGGGGAAGGCCTCGACGTTGAGTGCTGCATCTTGGAAGTATCTTGACACAATAAGAGAAA

AGAGATTGCGCTTAATAGGGGGTTGATCTCGTGGCGAGCCGACGCCGGTGTGAACTGGTG

GCAATGGCCGTCGTAGAGCACGCAAAGGTATGGGCTGACTTTTTCTCCTCAAAAGTTGGC

TTAAGGGACGTGCCATCCCACACGATAATAAAGTGTGCAGTCTGTAAGAGCTCCCACAGG

GCAAAGACAGACTGGGATGCAGACGGAGACGGCAAAAACAACAACATCCGTTCTGCTTCT
GGGCGCAAATGCTTAAGGTAGCCAAATGCCTCGTCATCTAAACTAATTGGATGAGATAGT
CTGAGTGAAAAGCTTATGCAGGCTAGTACCGGAAAAAAAAAAAAAAAAGAGGGGCTCTGCC
ACTCTGACGGTGCGACACTCCTTGTACAGGGAAACCCTCCGCATGGAAAGGCACGTGGTA
CCACCCTCGCGGGGAAACTCGCGAGCGGGATCCCGGGTAATGACCGGAGCCGATGGTAAA
AACTCACGTGACCGATGGGGTGACCCTGTGGCGAGACGCTAAGTTGTGGGACATCACCCT
CTCACAATACGCATCCGTCGCAGAGACTGAACGGGAGTGGGCTCCGTTGACGGCCGGCCG
GCCAGCCAGCCGTCTTCCAGGGGCTTAAGATACAGTCCGGGCCTCAGGGATGGAAAAACC
TGGGGAACTACCAGAATTAGTGACGCGCATGAATGGATTAACGAGATTCCCACTGTCCCT
ATCTACTATCTAGCGAAACCACAGCCAAGGGAACGGGCTTGGAAGAATCAGCGGGGAAAG
AAGACCCTGTTGAGCTTGACTCTAGTCCGACTTTGTGAGATGAGTTGGGAGGTGTAGTCT
AAGAGGGAGGGGCAACTCGACCGTGAAATACCTCTACTCCCAAGCTTGTTTTACTTACTC
CGGCGAGCAGGGATGGGTGGTAACACCCTTTTTCTAGGATTAACACCTCTTCGGGGGTGG
ACCTGCGCGGAGGACATTGTCAGGTGGGGAGTTTGGCTGGGGCGGCACATCTGTTAAAAG
ATAACGCAGGTGTCCTAAGATGAGCTCAGTGAGAACAGAAATCTCACGTAGAACAAAAGG
GTAAAAGCTCATTTGATTTTGATTTTCAGTACGAATACAAACTGTGAAAGCATGGCCTAT
CGATCCTTTTGACCTTCAAGGTTTGAAGCAAGAGGTGTCAGAAAAGTTAACACGCGTTGT
GGTAGCTTTTAAGAGTGGACTCAAAGTGTTCCGCTAGTCCCCCCTCGGACCGCCACTTAC
AAAAGGCCTGGGGATGGGGGCGACACTGTCAAATTGCGGGGACTCCCTAAAGCCTGTCTC
TACCAAGCCAGCGCGGCGAAAGCCCCTGGTGGCCGGGGTAATGACCTAGGGTATGGTAAA
AGCGAGACAGGATGGAACAATGGGCAATCCGCAGCCAAGTCCTAAGGCTTTTTGAAAAGC
GGCTATGGATGCAGTTCAACGACTAGATGGCAGTGGGCTGAGAGAGCCCACCGCGGCACC
CTTTCGGCTTAAGATATAGTCTTGCCCACGTGAAAGCGTGCCTGCAGATGAACGCCACCA
TGTATGGCCGGAGCTGCAGGAGACCGCACTTGATGTGCAGTCGGGAATCGACAGTACCAC
AGGGATGAACAAAAACACACAATTGTGGCCATCATACAGTCGTATGATATTATCGTCCCA
GGCGGGTGCGCCTCTCTCACCCGTCTAGTCCGGCCTTGGTGCGCAAAAAATGCTTGCGTG
GATGCGCCTTGGTTAAGGAAAATGCTCCTAGTAGTTTTACAAGGACACTTCTTGTAGGGC
AACACATCTGGATGCGAAGAACTCCCGGAAAAGACAACGGTACCAAGTGACTGGGGTAAC
GCCCCATTCATGAAGGCAAATAATCAAAAAGAGGTGCCGGGTAAGAACCCGTTGTATAGG
GACAATCCGCAGCTGACTTGGCTCCCTGCCCATCCATGGGCATAGCTTAGCGGTTCAACG
CTCGGTAAGATGTGGGTTGGCGCGCTGTCCGACAGCGCACTGGCTTAAGGTACGGGCTAC
TCCCACCGGAGACGGTGTCTTTGTCTACCTCGGGTAACACGCCTCGGGAGAAGCAAAGAG
GGTGCAATGGAAAAAAACTGGTTGCGGGGAAATCCCTACCAGAGTCACCTGGTACAAAGG
AACTGGCTTGTGGCAGCCAAGCGTTCATAGCGACGTTGCTTTTTGATCCTTCGATGTCGG
CTCTTCCTATCATTGCGAAGCAGAATTCGCCAAGTGTTGGATTGTTCACCCACTAATAGG
GAACGTGAGCTGGGTAGCCACTTTATGCCCAACAGCTGGATAGCTGGGCTTCTGTCTTAG

GCCGGGACTTCTCACGACCCGTGCCTTGGTCACCCAAAAAAGGGCGACCGCAGAGCTCAG
GATGGAGCTTCTGGCTTAACTAGTAGAGGACAGATAGAGATCGTCAAATTGCGGGAAACC
CCTTAGAGCCTCGAGCTACCAAGTGCGTGTCGAAAGGCATAGACGCAGCCAGCGATGGCT
GTCGCATGGCTTGATTCCATCAAGATAGGTAAAAATGCTCGAGGATTGGGCAATCCGCAG
CCAAGCCCCTAAGGTCCCCCCGCGGGCTATGGGGAAGGTTCAACGACTGTAAGGCGTACC
GTGCAAGCGGAAGATACAGTCTAGCCCCGCTGGAAAGCAGCGCCAAAGTAAGTAGGTTGC
AGTGGGTCTGTCCTC
>SRR9587932
GCATGTTTTGGTATGTGTTTCGACAGTCATGCATGTCATAGCCGTAGGATGGGGAGTGAG
CTTCATAGCCCGACCCCTTCTCAGGAGCATGTTTTCATAGCATATTGAGCATACTGGGGT
GGTCTCCCATGTAGCTCGCTCTCTCGGTGCAGGACCTCCCTTTTGCGGGCCACTTCGATG
GCCTCCATTATGGAGAGCATGTTTTGGCATATGAAGCGTTGGTGGTGAGGCTCGTAGCCT
GTATATACATACCAACGCAGTCGTCATATGTCTCTCTGGCGGGGGTCGGTCGTATGACCT
TGTCCCTACCTGGAGCATGTATTTTGTATGATATACTGGACGGCAGCTGCGCTGTTCAGT
CGTCGGCCGTACCTCATGTGCGCGTTTCCAGGTCCGAACGTTTGACATGCAAGGGCATCA
TACCAGAAGCCTGTTTTCTCGTATGAGCATGTAGGTAGGCAGGCTCTTCAGTCGTCTACC
CATAAATGCACTTCGAAGGAACTTTTTGGTATTGAATGGCTTTGTGCCCAGACGTTCTTG
TGAGCTTGTTCTCGCTCTGCGATACGCTGGAAGCTGTGGTATTACCTACAGCAGACGCAT
TCCTAGGGCGGACGGCCTCTCTGCAGATGGGCACATCTGTAGAAAGAGGCTCCCCTTGAT
GACGCAATGCAATGGCTGGGTAAACGGAGTGCTATACAGAAGGGTCAAACCTCTGTGGAA
GTACACCCATACCAGCATTGTTTGTTGAGAGGGGTCGGCTCTACGAGTGAGGACCTATAC
TGCCATCTCATTCAGATGCGTAGACGGTCTTTACTCAGACTGCCTTTCTGGCTGATTGTC
CTCGGTCCCTCGTGCGCACGCGCCGCTGGCTGAATGGGCTTGTCCTTATGGAGTGCCGCG
TTCTTCAAAGAATCCGTCTTCATACAGCGACCTCATTTACATATTGCCGGCTGAATGGGC
TTGTCCTTATGGAGCACTGCATTCTTCTAGGAATTGCTATCTATACAGCGACCTCATTTA
CATACAAGGCACCTTTTTCTACATAGACTTGCCATTTGGCTGATTGTTCTGACAGTCGCT
GCCTCTTCGGAGTTCTGTGTGGACGAGTGGCCTCCTGAGTGGGCCGGTCCTTACGGAGTG
CCCACGTCTGTTATGACTGGGTTCTCCATACAGCGACCTCACTCAGAATTGTATGGCACC
GTGCCGATTGAATGGACAAGCAGTCCTTAGGGAATCGCATGAAAGCATGACGGCACATGG
TGTATGGACCCGTTGGGCCTTGCCTGTGCTATCTTCTCGAGTCGACCGGTCCTCGTCTCC
TCCTTTCGCGACGGGAAGCTCTTGAGTTTACCGTGGGTCAGGACGCTGTGGTATCTGTTT
GTCTGTAGTGTGGGCTCTCTATCAGAACAGCGTATCAGCACACGTCGTCTGGGTCTATGG
TATTTTTGCTTAACGGCCCTGGGTTTGGTAGTATGGTTCCATGTCGGTCCTCTCACCTCC
CATCCGTCATGTCTCTGATATCATCTGCACAGTACGTCTTGTCAGAATATCATTCTGTAT
GGCAGTCGATAGACTAGGCCCTATGCCGCAGCTGTCTCTGACCACTATGCGAGCGCATAC
CTACCTCAATGTGTTCATAGCTGTTATGGATCTTGGGTGTGGTGTGCCAGAACGATACGC

CACCAGATTGTGGAATGGTGCTTGTAGTATTAGAAGAGGACATGTATGAAGAGCACGTTC
TTTTCTCTTATGATGGTGCCGCGGGTGGGGGCCGTTCTCATTCTGCTTGTCGGGTGCTCA
AGGTGCATGTAGGATTGCTGAACCCACTGAATGGTCATGAGAATGGTCATATGGTGAGGT
TCACTTCTCCATCGCTTTGTGTGTGTTGGATGTTTGGGTGGCTTCGGCCGCATGGAAAAT
CTCGGTCTGTAACTCATCAGGGAGGGTTTCCATGTGAGCAATCTCTCCAGTCGAACACAC
ACCCATGGGCGTCCCGCTAGGCAAAATGGAACTCTCGATCCGACACAGCCTGTGGAGTGT
CTGTTCCGTGTGGCGTGAAACGGCTTATACTGGATGTCTTCCCTTTGGACACCAGTCTGT
TGGTATGTACGCATACCATCTAGAGGGAGCAAGAAAAAGGCATGCAAGTGCCACAAAGGA
AGGGAGGCCTTCCTGCTACCGTGTGTCCTCTTGACGATCACGGGGGAGTATATAGAGACA
TTTGGACGAGATCCTTTTTTCACGAGGCGGTGCCCTCGCGCAAAGGTGTTGCTCTGCACG
CATCTTTATGATGATAGTTGCTCGCTCCTATGCGTCCACTCGTGAAAATACCTGCTCTCT
CTGTCTGTTTCCATGACACCCCCATGCGTTGTCATAGAGCACGCTCAGGTACAGAAAGTG
CTGTGACTTTCTTGGCCTGCTGTCACTGAGTCAGCCTGATACTTTGGAGGAATCTTTTCT
GGGCACTGCCCAGCTGAGAGTTCTTATCAGGCTCTGGAGTGTGCATATGCCTATATTCAT
TATAGTGCATGTGGCTACGCAGAAAAAGCATCGACGTGGAATGGAGAAGAACACCATTCT
GAATGGAACGGTTCTTACTCTACTTCTGTTGCTTGACGCTGTATGGGAATCGCTTTGCTC
TGGGGGTACACCTTCCATTCCATGACAAAGCATTTGATCTATGCCCCGAGGGTCATTCTC
TATGAAGAGCCCGAAACTTGTCTGAGTGAGAGGCGAAGGTAGTCCTTCATTTATGCGGGG
TCCGCCCTCACACTGTAGTCGGCACAGTCCCTTGTTTATGGACTGCGCAGCTCAATCTTC
GACTGGAGAGGAAAAACAAAACACAAGTATGACACGCACATACGTGTTTTCTAGGACTGT
GACCTGTCGGCTGCCTCGCGGTGGTTGGCATGATGCCCCAAGTGGGGGGGGGAGAAACAA
TCAGTATGATTGATAAAAACCAGTGGGGAAAAAGGCGCGCTGTCGCACACCCTTGGTCAC
GGCCTCAAGTCGACGGCCCTCACTGGCCCACCCTGCGACTCCTGTTCGCTTTTGACAAGA
CACACCTGCAATACGGAGGAAATGCCTGAGCAGCTGTGCCGCAGCAACTGCGGACGCGTA
GCGATATGCATCCCTGTTGTTTGCGACTTGGTACAAAGAACGTGCGCCGGGAGACGGATT
GCCCTTTGACTTTTCCACCCAGCCCGCGGCTCTATACCCTTGACGTACCTATTTACCTGG
TTGATCCTGCCAGTAGTCATATGCTTGTCTCAAAGACTAAGCCATGCATGTCTAAGTATA
AACACTTTTATACTCGTGAAACTGCGAATGGCTCATTAAAACAGTTATAGTTCCTTTGGG
AACAACACTACTTGGATAACCGTAGTAATTCTAGAGCTAATACATGCCTCAACGCCCGAC
TCACGAAGGGTGGTATTTATTGGATAAAAAACCATCGTGCTCTGTCCAGAGCGCTTTGAG
ATGATTCACAATAACTTGTCGGATCGCATGGCCTCGTGCTGGCGACTCCTCATTCAAATT
TCTGCCCTATCAACTTTCGATGGTAGAGTATTGGTCTACCATGGTGTCGACGGGTGACGG
GGAATTAGGGTTCGATTCCGGAGAGGGAGCCTGAGAAACGGCTACCACATCCAAGGAAGG
CAGCAGGCGCGCAAATTACCCAATCCCGACTCGGGGAGGTAGTGACAAAAAATAACAATA
GGGGGCCCTTTGGGTCTTCTAATTGGAATGAGAACAATTTAAATCCCTTATCGAGGATCC
ATTGGAGGGCAAGTCTGGTGAACGTATGCATTAAAGCCCTCCGTAATAAAAAAAAACAGC

GCCAGATGTCTCTCTCTGGAAGCACACCATGCATGGGGTTTGTTCCTCTCTGCATGGCCG
TGTGAAAAAGGCGAGTAGGCAAACGATGGGAAACCCGGCTTGGAATAGTAACCAGCCACA
GAGCCCCCATCATACACTTCTAAATTGCCGGGGAGTGCCCGCGAGGCCTTGACTACCGCG
GTCCGCCGAGAGGTGAGAGCGAGCAATCGTTCAGGACGCAGCAGTGTACGTAGTGGTACA
TGGATGGTAAAAACGTCAAGGATAGGGTTGATCCGCAGGGAAGCGCGAGTGCCGCCATGG
TCGGTGGGCAAAGGCGAACCTTCAGAGACTCTAATGGAGTGGGCGCAAGCTTAAGGGAGA
GTCCAATCCACTGGGAAACCAGGTCCACAGCAGTAACGGGGTGATGCTCGCCCCAGAGGA
CTGTGGAAGGCAACCCCACCAAAAGGGTGTGTGTTGTCGGGTAGCAATGGCCAGCAGCCG
CGGTAATTCCAGCTCCAATAGCGTATATTAAAGTTGTTGCAGTTAAAACGCTCGTAGTCG
GACCTCGGAGCGCCCCCACGGGCCGGGCTTTTGCCTTGTTGCCAGTGGGAGTTTGTCTCC
TTTTGTCGGGACGCGGTGTGCAGGGCCTTTACTGTCTCTGCGCGTTTACCTGCGGGCCGA
CCGTTTACTGTGAAGAAAGTAAAGTGTTCAAAGCAGGCCAATGCCTTGAATATGTGAGCA
TGGAATAATAGAATAGGACTTGGGCTCTATTTTGTTGGTTTCCAGTGACCAAGTAATGAT
TAATAGGGACGGTTGGGGGCATTCGTATTTCATTGTCAGAGGTGAAATTCTTGGATTGAT
GGAAGACGTACAACTGCGAAAGCATCTGCCATGGATGTTTTCATTGATCAAGAACGAAAG
TTAGGGGATCGAAGACGATCAGATACCGTCGTAGTCTTAACCATAAACGATGCCGACTGG
GGATTGGCGGGGGCACTTTTATATGACTCTGTCAGCACCCTGAGGGAAACCAAAGTCTTT
GGGTTCTGGGGGGAGTATGGTCGCAAGGCTGAAACTTAAAGGAATTGACGGAAGGGCACC
ACCAGGAGTGGAGCCTGCGGCTTAATTTGACTCAACACGGGAAAACTTACCAGGTCCGGA
CAGAAGAATGATTGACAGACTGAAAGCTCTTTCTTGATTTTTTGGTTGGTGGTGCATGGC
CGTTCTTAGTTGGTGGAGTGATTTGTCTGGTTAATTCCGTTAACGAACGAGACCTCGTCC
TGCTAAATAGGTGCGCGCATGCCAGCACTGCGTTCTTACCTTCTTAGAGGGACTATGCGC
GTCTAGCGTATGGAAGATTGAGGCAATAACAGGTCTGTGATGCCCTTAGATGTTCTGGGC
CGCACGCGCGCTACACTGATGCATTCAACGAGTTTCACTTGATAGTCCTGGGTCGGAAGG
CTCGGGTAATCTTTTGAAAGTGCATCGTGCTGGGGATAGATCATTGCAATTATTGATCTT
CAACGAGGAATTCCTTGTAGGCGCAGGTCATCAGCCTGCGCCGAATACGTCCCTGCCCTT
TGTACACACCGCCCGTCGCTCCTACCGATTGAATGGTCCGATGAAATGTCCGGATCGCGG
CTAGTGTGATGGACTCGTCCAGAGCGGTAGCCGTGAGAAGCTCATTAAATCTTATCATTT
AGAGGAAGGAGAAGTCGTAACAAGGTTTCCGTAGGTGAACCTGCGGAAGGATCATTCACA
AACTATTGACACAAACACACGCGAACCAAATCGTCCGCACAGCAGGTGGCAATGAAAGAG
AGACTCTGCATGTCGCCTTTCGGGGTAGCAAGCAGCACTCTTCTGCCATCGCCTCTGTGC
AGGGCGTAAATTCCCATTGAGAGGATGTGAGGGCACCACAGGAAGCTTTTCCACAGGAAG
TCGCCATCCTTCTCCCTCCACGGCGGCGCTTCTGTGCTTGGCAGTTTTTTTTGCCTTCTA
GGGAGGATGCCGCCAATGGAGCCCCTTATATATATACATCATCATATGCCCCTTTTTTTC
TTAACCGCTTGCCACAGCTTCTTCTATGAGGAGCTTGTGGGAAGACTGTCTCCATACAAT
AACAAAGATACAACTCTTAGCGGTGGATATCTTGGTTCTCGCAACGATGAAGAACGCAGC

CAACTGCGATAACTAATGTGAATTGCAGGACTTCGTGAATCATTAAGTCTTTGAACGCAA
GTTGCGCTCATCTCCATGAGGAGGTGAGCATGCCTGTTTGAGTGTCGATGGGAAAAACAA
TACATAACAACAATCGGCTCTCTTATCTTCCTGCCTTCTTGGGGGCAAAAGGTAGTGCAT
GCCGGAGATGGGCTTTGTCGCCCCTTTCTCTTGAAAGAGGCGTCGTCCGAAACACGTATC
GTTGCCCGCTGGAGGTCATAGAGCATGAGTTTAAAGCACCACATCTCTCAAGTCAGAGTG
AGCGTCTTGAGTAGGTCCGCATTGATTCAGTTTCAACCCAGGTGGCGTTGGTTACTTGCC
ATAAGCCTCTCTGGGGTTTCTTTGCTTGGACCTCTCTTGTCAGTTTCTCTGACGTAGAGA
CAGGTGCCGTCTCATGGACGTTGCCCTCTGGAACGTGCTAAAAAAAAAAAAAAAGAAAAAA
CCGTACCACATGTCAAAAGAGGCAGATCGTGCTTGACGGCGGGAGGTGTTGCATTTCTTG
CTTGTCCTGAGCGTCTGCCAGTCTCTGACGAGTGTGTTCCTTCCATAAACCCGATCTCAA
ATCAGGTAGGACCACCCGCTGAATTTAAGCATATAATTAAGCGGAGGAAAAGAAACTAAC
AAGGATTCCCTCAGTAGCGGCGAGCGAAGCGGGAAGAGCTCAACATGAAAACCGTTGGGG
GTTCGCGCTTTCCTTCGGGTTGTATAGTAAAGTGGTGTCGCCTGCGTCGCGCCGGGTATA
AATCCCGTCTGAAATGGGGTGTCGTAGAGGGTTATATCCCCGTCTGTGACCCGGTGTGCG
ATGCGTTACGGTGACCTTACGATGAGTCGGGTTGCTTGGGCATGCAGCCCTAAGTGGGTG
GTAAACTTCATCTAAAGCTAAATACTAGCGAGAGACCGATAGCGTACAAGTACCGTAAGG
GAAAGATGCAAAGAACTTTGAAAAGAGAGTTAAACAGGGCTTGAAACCGTTAAGAGTGAA
GCGAAGGAGGTCAGTGATCATGGGGCCAGATCCCCAAGTGCCTGTCATAGGGCACCGTAG
CGTTCTGGTTTCCCGGGGCAGCGCCGGCCGGGGGCGGGGGTTAACAGGGAGGTGGTTGAG
GGATTCGTTTTCCGCCTGTACTCCTTTATAGCTTGCACCCCGGCCGAGGGATCCGCGACT
TGTCGTGTGAGGGCGCTGTCAAAATGGCCTTCTCCACCCCGTCTTGAAACACGGACCAAG
GAGTCTAACATACTCGCGAGTGTTTGGGTGGTTAAAACCCGAACGCACAATGAAAGTAAA
CGGTGGCTCTGCGCCGCTGCTTGGTGGGAACCCTGCTTTTATAATGGGGGGCACTATCAC
CCGACCCCGATCCCTTTGGGGTGGTTGGTTTTGAGGTAGAGCGAGTATGATGGGACCCGA
AAGATGGTGAACTATTCCTGAGTAGGGCGAAGCCAGAGGAAACTCTGGTGGAGGCTCGTA
GCGATGCTGACGTGCAAATCGCTCGTCGAACTTGGGTATAGGGGCGAAAGACTAATCGAA
CCATCTAGTAGCTGGTTCCCTCCGAAGTTTCCCTCAGGATAGCGGGAGCAAATGCAAGTC
AGTATCATCCGGTAAAGCGAATGATTAGAGGCCTCGGGGGAGTAACTCCTTCGACCTATT
CTCAAACTTTAAATAGGTGAGGAGCCGGCAACGTTGCTTAAAACAATGAACGCCGTGCGG
ACAATGATTGCTCTCAGTGGGCCGATTTTGGTAAGCAGAACTGGCGATGCGGGATGAACC
GAAAGCGACGCTAAGGTGCCAAACTCCCAGCTCATTCAGACACCAGAAAAGGTGTTGATT
CATACAGACAGCAGGACGGTGGCCATGGAAGTCGGAATCCGCTAAGGAGTGTGTAACAAC
TCACCTGCCGAATGGATCAGCCCTGAAAATGGATGGCGCTTAAGCGGGGGACCGAGGCGT
CGCCGTTGGGGTAAATACGAAGCCCCAATGAGTAGGAGGGCGTGGGGGTTTTGAAACACG
GTACACTCGCATGGAAGCCTTGGGCGTAAGCTCGGGTGGAGCGACCCCTAGTGTAGATCT
TGGTGGTAGTAGCAAATATTCAAGTGAGAACCTTGAAGACTGAAGTGGGGAAGGGTTCCA

CGCGAACAGTGCTTGGACGTGGGTTAGTCGGTCCTAAGAGATGGGGAAACTCCGTATTCA
AGTGCGCTTTTGTAGCGCAGTCCATCGAAAGGGAAGCGGGTTAACATTCCCGCACCAGGA
TGTGGAAACTGTGGTTTACCGCAAACTGCGGCAACGCAACTGAACCTGGGGACGCCCGTA
GGAGCTCCGAGCAGGGATCTCTTTCCTGTTTAACGCGCCCAACCCCCTGGAATCGGTTTG
TCCGGAGATAGGGGGCGAAAGGCCGGAAAAGCACCGCACGTCTTGTGGTGTTCGGAACGC
TTGTAATGGCCCTTGAAAACCCAGGGGAAAGCAGCCGTCTTACACTTGTGTGAGACGAGG
AATTCTCACACCTGGCCGTACCCATCACCGCATCAGGTCCCCAAGGTGAACAGCCTCTAG
TCAATAGAATAATGTAGATAAGGGAAGTCGGCAAAATAGATCCGTAACTTTGGGAAAAGG
ATTGGCTCTAATGTTCGGGCTTGTCGGGACGCGCGGGCGTAGATGCCAATGAGCGGCAGA
CGTCTAGCCTCTTTGGAGGCTGGATTAACGCTGCTTGTGGGAAGAAACCGTGTCGATGGA
GGCAGGAACGCTTATCGATCTTGTCTCTTCCCGCTTGCGATCAACGAACGGATTAGAACT
GGTACGGACAAGGGGAATCCGACTGTTTAATTAAAACAAAGCATTGCGACGGCCGGAAAC
GGTGTTGGCGCAATGTGATTTCTGCCCAGTGCTCTGAATGTCAAAGTGAAGAAATTCAAC
CAAGCGCGGGTAAACGGCGGGAGTAACTATGACTCTTAAACAAACCATACTACCTACGGG
AGTCAAAAGTGGGGGCGGCGTCTGGTGTAGAAATACACGTACGACACGCTTTTTTTCCTG
CTAGTCGATCTGACCTTTGTTCTTTGGACTGTGGGCGGTCAGCGGCAACATAACCTGGTA
CGGGGAAGGCCTCGACGTTGAGTGCTGCATCTTGGAAGTATCTTGACACAATAAGAGAAA
AGAGATTGCGCTTAATAGGGGGTTGATCTCGTGGCGAGCCGACGCCGGTGTGAACTGGTG
GCAATGGCCGTCGTAGAGCACGCAAAGGTATGGGCTGACTTTTTCTCCTCAAAAGTTGGC
TTAAGGGACGTGCCATCCCACACGATAATAAAGTGTGCAGTCTGTAAGAGCTCCCACAGG
GCAAAGACAGACTGGGATGCAGACGGAGACGGCAAAAACAACAACATCCGTTCTGCTTCT
GGGCGCAAATGCTTAAGGTAGCCAAATGCCTCGTCATCTAAACTAATTGGATGAGATAGT
CTGAGTGAAAAGCTTATGCAGGCTAGTACCGGAAAAAAAAAAAAAAAGAGGGGCTCTGCC
ACTCTGACGGTGCGACACTCCTTGTACAGGGAAACCCTCCGCATGGAAAGGCACGTGGTA
CCACCCTCGCGGGGAAACTCGCGAGCGGGATCCCGGGTAATGACCGGAGCCGATGGTAAA
AACTCACGTGACCGATGGGGTGACCCTGTGGCGAGACGCTAAGTTGTGGGACATCACCCT
CTCACAATACGCATCCGTCGCAGAGACTGAACGGGAGTGGGCTCCGTTGACGGCCGGCCG
GCCAGCCAGCCGTCTTCCAGGGGCTTAAGATACAGTCCGGGCCTCAGGGATGGAAAAACC
TGGGGAACTACCAGAATTAGTGACGCGCATGAATGGATTAACGAGATTCCCACTGTCCCT
ATCTACTATCTAGCGAAACCACAGCCAAGGGAACGGGCTTGGAAGAATCAGCGGGGAAAG
AAGACCCTGTTGAGCTTGACTCTAGTCCGACTTTGTGAGATGAGTTGGGAGGTGTAGTCT
AAGAGGGAGGGGCAACTCGACCGTGAAATACCTCTACTCCCAAGCTTGTTTTACTTACTC
CGGCGAGCAGGGATGGGTGGTAACACCCTTTTTCTAGGATTAACACCTCTTCGGGGGTGG
ACCTGCGCGGAGGACATTGTCAGGTGGGGAGTTTGGCTGGGGCGGCACATCTGTTAAAAG
ATAACGCAGGTGTCCTAAGATGAGCTCAGTGAGAACAGAAATCTCACGTAGAACAAAAGG
GTAAAAGCTCATTTGATTTTGATTTTCAGTACGAATACAAACTGTGAAAGCATGGCCTAT

CGATCCTTTTGACCTTCAAGGTTTGAAGCAAGAGGTGTCAGAAAAGTTAACACGCGTTGT
GGTAGCTTTTAAGAGTGGACTCAAAGTGTTCCGCTAGTCCCCCCTCGGACCGCCACTTAC
AAAAGGCCTGGGGATGGGGGCGACACTGTCAAATTGCGGGGACTCCCTAAAGCCTGTCTC
TACCAAGCCAGCGCGGCGAAAGCCCCTGGTGGCCGGGGTAATGACCTAGGGTATGGTAAA
AGCGAGACAGGATGGAACAATGGGCAATCCGCAGCCAAGTCCTAAGGCTTTTTGAAAAGC
GGCTATGGATGCAGTTCAACGACTAGATGGCAGTGGGCTGAGAGAGCCCACCGCGGCACC
CTTTCGGCTTAAGATATAGTCTTGCCCACGTGAAAGCGTGCCTGCAGATGAACGCCACCA
TGTATGGCCGGAGCTGCAGGAGACCGCACTTGATGTGCAGTCGGGAATCGACAGTACCAC
AGGGATGAACAAAAACACACAATTAATTTGTGGCCATCATACAGTCGTATGATATTATCG
TCCCAGGCGGGTGCGCCTCTCTCACCCGTCTAGTCCGGCCTTGGTGCGCAAAAAATGCTT
GCGTGGATGCGCCTTGGTTAAGGAAAATGCTCCTAGTAGTTTTACAAGGACACTTCTTGT
AGGGCAACACATCTGGATGCGAAGAACTCCCGGAAAAGACAACGGTACCAAGTGACTGGG
GTAACGCCCCATTCATGAAGGCAAATAATCAAAAAGAGGTGCCGGGTAAGAACCCGTTGT
ATAGGGACAATCCGCAGCTGACTTGGCTCCCTGCCCATCCATGGGCATAGCTTAGCGGTT
CAACGCTCGGTAAGATGTGGGTTGGCGCGCTGTCCGACAGCGCACTGGCTTAAGGTACGG
GCTACTCCCACCGGAGACGGTGTCTTTGTCTACCTCGGGTAACACGCCTCGGGAGAAGCA
AAGAGGGTGCAATGGAAAAAAACTGGTTGCGGGGAAATCCCTACCAGAGTCACCTGGTAC
AAAGGAACTGGCTTGTGGCAGCCAAGCGTTCATAGCGACGTTGCTTTTTGATCCTTCGAT
GTCGGCTCTTCCTATCATTGCGAAGCAGAATTCGCCAAGTGTTGGATTGTTCACCCACTA
ATAGGGAACGTGAGCTGGGTAGCCACTTTATGCCCAACAGCTGGATAGCTGGGCTTCTGT
CTTAGGCCGGGACTTCTCACGACCCGTGCCTTGGTCACCCAAAAAAGGGCGACCGCAGAG
CTCAGGATGGAGCTTCTGGCTTAACTAGTAGAGGACAGATAGAGATCGTCAAATTGCGGG
AAACCCCTTAGAGCCTCGAGCTACCAAGTGCGTGTCGAAAGGCATAGACGCAGCCAGCGA
TGGCTGTCGCATGGCTTGATTCCATCAAGATAGGTAAAAATGCTCGAGGATTGGGCAATC
CGCAGCCAAGCCCCTAAGGTCCCCCCGCGGGCTATGGGGAAGGTTCAACGACTGTAAGGC
GTACCGTGCAAGCGGAAGATACAGTCTAGCCCCGCTGGAAAGCAGCGCCAAAGTAAGTAG
GTTGCAGTGGGTCTGTCCTC
>SRR9587933
GCATGTTTTGGTATGTGTTTCGACAGTCATGCATGTCATAGCCGTAGGATGGGGAGTGAG
CTTCATAGCCCGACCCCTTCTCAGGAGCATGTTTTCATAGCATATTGAGCATACTGGGGT
GGTCTCCCATGTAGCTCGCTCTCTCGGTGCAGGACCTCCCTTTTGCGGGCCACTTCGATG
GCCTCCATTATGGAGAGCATGTTTTGGCATATGAAGCGTTGGTGGTGAGGCTCGTAGCCT
GTATATACATACCAACGCAGTCGTCATATGTCTCTCTGGCGGGGGTCGGTCGTATGACCT
TGTCCCTACCTGGAGCATGTATTTTGTATGATATACTGGACGGCAGCTGCGCTGTTCAGT
CGTCGGCCGTACCTCATGTGCGCGTTTCCAGGTCCGAACGTTTGACATGCAAGGGCATCA
TACCAGAAGCCCGTTTTCTCGTATGAGCATGTAGGTAGGCAGGCTCTTCAGTCGTCTACC

CATAAATGCACTTCGAAGGAACTTTTTGGTATTGAATGGCTTTGTGCCCAGACGTTCTTG
TGAGCTTGTTCTCGCTCTGCGATACGCTGGAAGCTGTGGTATTACCTACAGCAGACGCAT
TCCTAGGGCGGACGGCCTCTCTGCAGATGGGCACATCTGTAGAAAGAGGCTCCCCTTGAT
GACGCAATGCAATGGCTGGGTAAACGGAGTGCTATACAGAAGGGTCAAACCTCTGTGGAA
GTACACCCATACCAGCATTGTTTGTTGAGAGGGGTCGGCTCTACGAGTGAGGACCTATAC
TGCCATCTCATTCAGATGCGTAGACGGTCTTTACTCAGACTGCCTTTCTGGCTGATTGTC
CTCGGTCCCTCGTGCGCACGCGCCGCTGGCTGAATGGGCTTGTCCTTATGGAGTGCCGCG
TTCTTCAAAGAATCCGTCTTCATACAGCGACCTCATTTACATATTGCCGGCTGAATGGGC
TTGTCCTTATGGAGCACTGCATTCTTCTAGGAATTGCTATCTATACAGCGACCTCATTTA
CATACAAGGCACCTTTTTCTACATAGACTTGCCATTTGGCTGATTGTTCTGACAGTCGCT
GCCTCTTCGGAGTTCTGTGTGGACGAGTGGCCTCCTGAGTGGGCCGGTCCTTACGGAGTG
CCCACGTCTGTTATGACTGGGTTCTCCATACAGCGACCTCACTCAGAATTGTATGGCACC
GTGCCGATTGAATGGACAAGCAGTCCTTAGGGAATCGCATGAAAGCATGACGGCACATGG
TGTATGGACCCGTTGGGCCTTGCCTGTGCTATCTTCTCGAGTCGACCGGTCCTCGTCTCC
TCCTTTCGCGACGGGAAGCTCTTGAGTTTACCGTGGGTCAGGACGCTGTGGTATCTGTTT
GTCTGTAGTGTGGGCTCTCTATCAGAACAGCGTATCAGCACACGTCGTCTGGGTCTATGG
TATTTTTGCTTAACGGCCCTGGGTTTGGTAGTATGGTTCCATGTCGGTCCTCTCACCTCC
CAACCGTCATGTCTCTGATATCATCTGCACAGTACGTCTTGTCAGAATATCATTCTGTAT
GGCAGTCGATAGACTAGGCCCTATGCCGCAGCTGTCTCTGACCACTATGCGAGCGCATAC
CTACCTCAATGTGTTCATAGCTGTTATGGATCTTGGGTGTGGTGTGCCAGAACGATACGC
CACCAGATTGTGGAATGGTGCTTGTAGTATTAGAAGAGGACATGTATGAAGAGCACGTTC
TTTTCTCTTATGATGGTGCCGCGGGTGCGGGCCGTTCTCATTCTGCTTGTCGGGTGCTCA
AGGTGCATGTAGGATTGCTGAACCCACTGAATGGTCATGAGAATGGTCATATGGTGAGGT
TCACTTCTCCATCGCTTTGTGTGTGTTGGATGTTTGGGTGGCTTCGGCCGCATGGAAAAT
CTCGGTCTGTAACTCATCAGGGAGGGTTTCCATGTGAGCAATCTCTCCAGTCGAACACAC
ACCCATGGGCGTCCCGCTAGGCAAAATGGAACTCTCGATCCGACACAGCCTGTGGAGTGT
TTGTTCCGTGTGGCGTGAAACGGCTTATACTGGATGTCTTCCCTTTGGACACCAGTCTGT
TGGTATGTACGCATACCATCTAGAGGGAGCAAGAAAAAGGCATGCAAGTGCCACAAAGGA
AGGGAGGCCTTCCTGCTACCGTGTGTCCTCTTGACGATCACGGGGGAGTATATAGAGACA
TTTGGACGAGATCCGTTTTTCACGAGGCGGTGCCCTCGCGCAAAGGTGTTGCTCTGCACG
CATCTTTATGATGATAGTTGCTCGCTCCTATGCGTCCACTCGTGAAAATACCTGCTCTCT
CTGTCTGTTTCCATGACACCCCCATGCGTTGTCATAGAGCATGCTCAGGTACAGAAAGTG
CTGTGACTTTCTTGGCCTGCTGTCACTGAGTCAGCCTGATACTTTGGAGGAATCTTTTCT
GGGCACTGCCCAGCTGAGAGTTCTTATCAGGCTCTGGAGTGTGCATATGCCTATATTCAT
TATAGTGCATGTGGCTACGCAGAAAAAGCATCGACGTGGAATGGAGAAGAACACCATTCT
GAATGGAACGGTTCTTACTCTACTTCTGTTGCTTGACGCTGTATGGGAATCGCTTTGCTC

TGGGGGTACACCTTCCATTCCATGACAAAGCATTTGATCTATGCCCCGAGGGTCATTCTC
TATGAAGAGCCCGAAACTTGTCTGAGTGAGAGGCGAAGGTAGTCCTTCATTTATGCGGGG
TCCGCCCTCACACTGTAGTCGGCACAGTCCCTTGTTTATGGACTGCGCAGCTCAATCTTC
GACTGGAGAGGAAAAACAAAACACAAGTATGACACGCACATACGTGTTTTCTAGGACTGT
GACCTGTCGGCTGCCTCGCGGTGGTTGGCATGATGCCCCAAGTGGGGGGGGGAGAAACAA
TCAGTATGATTGATAAAAACCAGTGGGGAAAAAGGCGCGCTGTCGCACACCCTTGGTCAC
GGCCTCAAGTCGACGGCCCTCACTGGCCCACCCTGCGACTCCTGTTCGCTTTTGACAAGA
CACACCTGCAATACGGAGGAAATGCCTGAGCAGCTGTGCCGCAGCAACTGCGGACGCGTA
GCGATATGCATCCCTGTTGTTTGCGACTTGGTACAAAGAACGTGCGCCGGGAGACGGATT
GCCCTTTGACTTTTCCACCCAGCCCGCGGCTCTATACCCTTGACGTACCTATTTACCTGG
TTGATCCTGCCAGTAGTCATATGCTTGTCTCAAAGACTAAGCCATGCATGTCTAAGTATA
AACACTTTTATACTCGTGAAACTGCGAATGGCTCATTAAAACAGTTATAGTTCCTTTGGG
AACAACACTACTTGGATAACCGTAGTAATTCTAGAGCTAATACATGCCTCAACGCCCAAC
TCACGAAGGGTGGTATTTATTGGATAAAAAACCATCGTGCTCTGTCCAGAGCGCTTTGAG
ATGATTCACAATAACTTGTCGGATCGCATGGCCTCGTGCTGGCGACTCCTCATTCAAATT
TCTGCCCTATCAACTTTCGATGGTAGAGTATTGGTCTACCATGGTGTCGACGGGTGACGG
GGAATTAGGGTTCGATTCCGGAGAGGGAGCCTGAGAAACGGCTACCACATCCAAGGAAGG
CAGCAGGCGCGCAAATTACCCAATCCCGACTCGGGGAGGTAGTGACAAAAAATAACAATA
GGGGGCCCTTTGGGTCTTCTAATTGGAATGAGAACAATTTAAATCCCTTATCGAGGATCC
ATTGGAGGGCAAGTCTGGTGAACGTATGCATTAAAGCCCTCCGTAATAAAAAAAAACAGC
GCCAGATGTCTCTCTCTGGAAGCACACCATGCATGGGGTTTGTTCCTCTCTGCATGGCCG
TGTGAAAAAGGCGAGTAGGCAAACGATGGGAAACCCGGCTTGGAATAGTAACCAGCCACA
GAGCCCCCATCATACACTTCTAAATTGCCGGGGAGTGCCCGCGAGGCCTTGACTACCGCG
GTCCGCCGAGAGGTGAGAGCGAGCAATCGTTCAGGACGCAGCAGTGTACGTAGTGGTACA
TGGATGGTAAAAACGTCAAGGATAGGGTTGATCCGCAGGGAAGCGCGAGTGCCGCCATGG
TCGGTGGGCAAAGGCGAACCTTCAGAGACTCTAATGGAGTGGGCGCAAGCTTAAGGGAGA
GTCCAATCCACTGGGAAACCAGGTCCACAGCAGTAACGGGGTGATGCTCGCCCCAGAGGA
CTGTGGAAGGCAACCCCACCAAAAGGGTGTGTGTTGTCGGGTAGCAATGGCCAGCAGCCG
CGGTAATTCCAGCTCCAATAGCGTATATTAAAGTTGTTGCAGTTAAAACGCTCGTAGTCG
GACCTCGGAGCGCCCCCACGGGCCGGGCTTTTGCCTTGTTGCCAGTGGGAGTTTGTCTCC
TTTTGTCGGGACGCGGTGTGCAGGGCCTTTACTGTCTCTGCGCGTTTACCTGCGGGCCGA
CCGTTTACTGTGAAGAAAGTAAAGTGTTCAAAGCAGGCCAATGCCTTGAATATGTGAGCA
TGGAATAATAGAATAGGACTTGGGCTCTATTTTGTTGGTTTCCAGTGACCAAGTAATGAT
TAATAGGGACGGTTGGGGGCATTCGTATTTCATTGTCAGAGGTGAAATTCTTGGATTGAT
GGAAGACGTACAACTGCGAAAGCATCTGCCATGGATGTTTTCATTGATCAAGAACGAAAG
TTAGGGGATCGAAGACGATCAGATACCGTCGTAGTCTTAACCATAAACGATGCCGACTGG

GGATTGGCGGGGGCACTTTTATATGACTCTGTCAGCACCCTGAGGGAAACCAAAGTCTTT

GGGTTCTGGGGGGAGTATGGTCGCAAGGCTGAAACTTAAAGGAATTGACGGAAGGGCACC

ACCAGGAGTGGAGCCTGCGGCTTAATTTGACTCAACACGGGAAAACTTACCAGGTCCGGA

CAGAAGAATGATTGACAGACTGAAAGCTCTTTCTTGATTTTTTGGTTGGTGGTGCATGGC

CGTTCTTAGTTGGTGGAGTGATTTGTCTGGTTAATTCCGTTAACGAACGAGACCTCGTCC

TGCTAAATAGGTGCGCGCATGCCAGCACTGCGTTCTTACCTTCTTAGAGGGACTATGCGC

GTCTAGCGTATGGAAGATTGAGGCAATAACAGGTCTGTGATGCCCTTAGATGTTCTGGGC

CGCACGCGCGCTACACTGATGCATTCAACGAGTTTCACTTGATAGTCCTGGGTCGGAAGG

CTCGGGTAATCTTTTGAAAGTGCATCGTGCTGGGGATAGATCATTGCAATTATTGATCTT

CAACGAGGAATTCCTTGTAGGCGCAGGTCATCAGCCTGCGCCGAATACGTCCCTGCCCTT

TGTACACACCGCCCGTCGCTCCTACCGATTGAATGGTCCGATGAAATGTCCGGATCGCGG

CTAGTGTGATGGACTCGTCCAGAGCGGTAGCCGTGAGAAGCTCATTAAATCTTATCATTT

AGAGGAAGGAGAAGTCGTAACAAGGTTTCCGTAGGTGAACCTGCGGAAGGATCATTCACA

AACTATTGACACAAACACACGCGAACCAAATCGTCCGCACAGCAGGTGGCAATGAAAGAG

AGACTCTGCATGTCGCCTTTCGGGGTAGCAAGCAGCACTCTTCTGCCATCGCCTCTGTGC

AGGGCGTAAATTCCCATTGAGAGGATGTGAGGGCACCACAGGAAGCTTTTCCACAGGAAG

TCGCCATCCTTCTCCCTCCACGGCGGCGCTTCTGTGCTTGGCAGTTTTTTTGCCTTCTA

GGGAGGATGCCGCCAATGGAGCCCCTTATATATATACATCATCATATGCCCCTTTTTTTC

TTAACCGCTTGCCACAGCTTCTTCTATGAGGAGCTTGTGGGAAGACTGTCTCCATACAAT

AACAAAGATACAACTCTTAGCGGTGGATATCTTGGTTCTCGCAACGATGAAGAACGCAGC

CAACTGCGATAACTAATGTGAATTGCAGGACTTCGTGAATCATTAAGTCTTTGAACGCAA

GTTGCGCTCATCTCCATGAGGAGGTGAGCATGCCTGTTTGAGTGTCGATGGGAAAAACAA

TACATAACAACAATCGGCTCTCTTATCTTCCTGCCTTCTTGGGGGCAAAAGGTAGTGCAT

GCCGGAGATGGGCTTTGTCGCCCCTTTCTCTTGAAAGAGGCGTCGTCCGAAACACGTATC

GTTGCCCGCTGGAGGTCATAGAGCATGAGTTTAAAGCACCACATCTCTCAAGTCAGAGTG

AGCGTCTTGAGTAGGTCCGCATTGATTCAGTTTCAACCCAGGTGGCGTTGGTTACTTGCC

ATAAGCCTCTCTGGGGTTTCTTTGCTTGGACCTCTCTTGTCAGTTTCTCTGACGTAGAGA

CAGGTGCCGTCTCATGGACGTTGCCCTCTGGAACGTGCTAAAAAAAAAAAAAAGAAAAAA

CCGTACCACATGTCAAAAGAGGCAGATCGTGCTTGACGGCGGGAGGTGTTGCATTTCTTG

CTTGTCCTGAGCGTCTGCCAGTCTCTGACGAGTGTGTTCCTTCCATAAACCCGATCTCAA

ATCAGGTAGGACCACCCGCTGAATTTAAGCATATAATTAAGCGGAGGAAAAGAAACTAAC

AAGGATTCCCTCAGTAGCGGCGAGCGAAGCGGGAAGAGCTCAACATGAAAACCGTTGGGG

GTTCGCGCTTTCCTTCGGGTTGTATAGTAAAGTGGTGTCGCCTGCGTCGCGCCGGGTATA

AATCCCGTCTGAAATGGGGTGTCGTAGAGGGTTATATCCCCGTCTGTGACCCGGTGTGCG

ATGCGTTACGGTGACCTTACGATGAGTCGGGTTGCTTGGGCATGCAGCCCTAAGTGGGTG

GTAAACTTCATCTAAAGCTAAATACTAGCGAGAGACCGATAGCGTACAAGTACCGTAAGG

GAAAGATGCAAAGAACTTTGAAAAGAGAGTTAAACAGGGCTTGAAACCGTTAAGAGTGAA
GCGAAGGAGGTCAGTGATCATGGGGCCAGATCCCCAAGTGCCTGTCATAGGGCACCGTAG
CGTTCTGGTTTCCCGGGGCAGCGCCGGCCGGGGGCGGGGGTTAACAGGGAGGTGGTTGAG
GGATTCGTTTTCCGCCTGTACTCCTTTATAGCTTGCACCCCGGCCGAGGGATCCGCGACT
TGTCGTGTGAGGGCGCTGTCAAAATGGCCTTCTCCACCCCGTCTTGAAACACGGACCAAG
GAGTCTAACATACTCGCGAGTGTTTGGGTGGTTAAAACCCGAACGCACAATGAAAGTAAA
CGGTGGCTCTGCGCCGCTGCTTGGTGGGAACCCTGCTTTTATAATGGGGGGCACTATCAC
CCGACCCCGATCCCTTTGGGGTGGTTGGTTTTGAGGTAGAGCGAGTATGATGGGACCCGA
AAGATGGTGAACTATTCCTGAGTAGGGCGAAGCCAGAGGAAACTCTGGTGGAGGCTCGTA
GCGATGCTGACGTGCAAATCGCTCGTCGAACTTGGGTATAGGGGCGAAAGACTAATCGAA
CCATCTAGTAGCTGGTTCCCTCCGAAGTTTCCCTCAGGATAGCGGGAGCAAATGCAAGTC
AGTATCATCCGGTAAAGCGAATGATTAGAGGCCTCGGGGGAGTAACTCCTTCGACCTATT
CTCAAACTTTAAATAGGTGAGGAGCCGGCAACGTTGCTTAAAACAATGAACGCCGTGCGG
ACAATGATTGCTCTCAGTGGGCCGATTTTGGTAAGCAGAACTGGCGATGCGGGATGAACC
GAAAGCGACGCTAAGGTGCCAAACTCCCAGCTCATTCAGACACCAGAAAAGGTGTTGATT
CATACAGACAGCAGGACGGTGGCCATGGAAGTCGGAATCCGCTAAGGAGTGTGTAACAAC
TCACCTGCCGAATGGATCAGCCCTGAAAATGGATGGCGCTTAAGCGGGGGACCGAGGCGT
CGCCGTTGGGGTAAATACGAAGCCCCAATGAGTAGGAGGGCGTGGGGGTTTTGAAACACG
GTACACTCGCATGGAAGCCTTGGGCGTAAGCTCGGGTGGAGCGACCCCTAGTGTAGATCT
TGGTGGTAGTAGCAAATATTCAAGTGAGAACCTTGAAGACTGAAGTGGGGAAGGGTTCCA
CGCGAACAGTGCTTGGACGTGGGTTAGTCGGTCCTAAGAGATGGGGAAACTCCGTATTCA
AGTGCGCTTTTGTAGCGCAGTCCATCGAAAGGGAAGCGGGTTAACATTCCCGCACCAGGA
TGTGGAAACTGTGGTTTACCGCAAACTGCGGCAACGCAACTGAACCTGGGGACGCCCGTA
GGAGCTCCGAGCAGGGATCTCTTTCCTGTTTAACGCGCCCAACCCCCTGGAATCGGTTTG
TCCGGAGATAGGGGGCGAAAGGCCGGAAAAGCACCGCACGTCTTGTGGTGTTCGGAACGC
TTGTAATGGCCCTTGAAAACCCAGGGGAAAGCAGCCGTCTTACACTTGTGTGAGACGAGG
AATTCTCACACCTGGCCGTACCCATCACCGCATCAGGTCCCCAAGGTGAACAGCCTCTAG
TCAATAGAATAATGTAGATAAGGGAAGTCGGCAAAATAGATCCGTAACTTTGGGAAAAGG
ATTGGCTCTAATGTTCGGGCTTGTCGGGACGCGCGGGCGTAGATGCCAATGAGCGGCAGA
CGTCTAGCCTCTTTGGAGGCTGGATTAACGCTGCTTGTGGGAAGAAACCGTGTCGATGGA
GGCAGGAACGCTTATCGATCTTGTCTCTTCCCGCTTGCGATCAACGAACGGATTAGAACT
GGTACGGACAAGGGGAATCCGACTGTTTAATTAAAACAAAGCATTGCGACGGCCGGAAAC
GGTGTTGGCGCAATGTGATTTCTGCCCAGTGCTCTGAATGTCAAAGTGAAGAAATTCAAC
CAAGCGCGGGTAAACGGCGGGAGTAACTATGACTCTTAAACAAACCATACTACCTACGGG
AGTCAAAAGTGGGGGCGGCGTCTGGTGTAGAAATACACGTACGACACGCTTTTTTTCCTG
CTAGTCGATCTGACCTTTGTTCTTTGGACTGTGGGCGGTCAGCGGCAACATAACCTGGTA

CGGGGAAGGCCTCGACGTTGAGTGCTGCATCTTGGAAGTATCTTGACACAATAAGAGAAA
AGAGATTGCGCTTAATAGGGGGTTGATCTCGTGGCGAGCCGACGCCGGTGTGAACTGGTG
GCAATGGCCGTCGTAGAGCACGCAAAGGTATGGGCTGACTTTTTCTCCTCAAAAGTTGGC
TTAAGGGACGTGCCATCCCACACGATAATAAAGTGTGCAGTCTGTAAGAGCTCCCACAGG
GCAAAGACAGACTGGGATGCAGACGGAGACGGCAAAAACAACAACATCCGTTCTGCTTCT
GGGCGCAAATGCTTAAGGTAGCCAAATGCCTCGTCATCTAAACTAATTGGATGAGATAGT
CTGAGTGAAAAGCTTATGCAGGCTAGTACCGGAAAAAAAAAAAAAAAAGAGGGGCTCTGCC
ACTCTGACGGTGCGACACTCCTTGTACAGGGAAACCCTCCGCATGGAAAGGCACGTGGTA
CCACCCTCGCGGGGAAACTCGCGAGCGGGATCCCGGGTAATGACCGGAGCCGATGGTAAA
AACTCACGTGACCGATGGGGTGACCCTGTGGCGAGACGCTAAGTTGTGGGACATCACCCT
CTCACAATACGCATCCGTCGCAGAGACTGAACGGGAGTGGGCTCCGTTGACGGCCGGCCG
GCCAGCCAGCCGTCTTCCAGGGGCTTAAGATACAGTCCGGGCCTCAGGGATGGAAAAACC
TGGGGAACTACCAGAATTAGTGACGCGCATGAATGGATTAACGAGATTCCCACTGTCCCT
ATCTACTATCTAGCGAAACCACAGCCAAGGGAACGGGCTTGGAAGAATCAGCGGGGAAAG
AAGACCCTGTTGAGCTTGACTCTAGTCCGACTTTGTGAGATGAGTTGGGAGGTGTAGTCT
AAGAGGGAGGGGCAACTCGACCGTGAAATACCTCTACTCCCAAGCTTGTTTTACTTACTC
CGGCGAGCAGGGATGGGTGGTAACACCCTTTTTCTAGGATTAACACCTCTTCGGGGGTGG
ACCTGCGCGGAGGACATTGTCAGGTGGGGAGTTTGGCTGGGGCGGCACATCTGTTAAAAG
ATAACGCAGGTGTCCTAAGATGAGCTCAGTGAGAACAGAAATCTCACGTAGAACAAAAGG
GTAAAAGCTCATTTGATTTTGATTTTCAGTACGAATACAAACTGTGAAAGCATGGCCTAT
CGATCCTTTTGACCTTCAAGGTTTGAAGCAAGAGGTGTCAGAAAAGTTAACACGCGTTGT
GGTAGCTTTTAAGAGTGGACTCAAAGTGTTCCGCTAGTCCCCCCTCGGACCGCCACTTAC
AAAAGGCCTGGGGATGGGGGCGACACTGTCAAATTGCGGGGACTCCCTAAAGCCTGTCTC
TACCAAGCCAGCGCGGCGAAAGCCCCTGGTGGCCGGGGTAATGACCTAGGGTATGGTAAA
AGCGAGACAGGATGGAACAATGGGCAATCCGCAGCCAAGTCCTAAGGCTTTTTGAAAAGC
GGCTATGGATGCAGTTCAACGACTAGATGGCAGTGGGCTGAGAGAGCCCACCGCGGCACC
CTTTCGGCTTAAGATATAGTCTTGCCCACGTGAAAGCGTGCCTGCAGATGAACGCCACCA
TGTATGGCAGGAGCTGCAGGAGGCCGCTAAACATACCAGCGGCTGGGAACATAACAGTAC
CACAGGGATGAACCATACATAGAATTAGCCAGTTTGGTGAGACAATAATTGTGCACGAAC
GCGGTATTGTCCACTTTGTGCGGTGTCCTGCTCGGGGCCGTCCGTGTGCGTCATGGCTGA
CAGAGAACGCCCTCCTAAGCGAGTGGCGTTCGCCGTTGCGTCGGCAACCAACAGTGTGCA
TGTGCAGATACATTCAGGTGGAAGCTCACAGGAGTCCACGGACTCTTCGTTTACTCCTCA
CAGCGATCACCCGGATGCGGGAATGAGTGGTACTGAAGACAGCGGCGATGAAGACGGCGA
AGAGAGTGTTGAAGAGGCCCCTTTGGCGCGCACCACGGTATTCACTCAAATCTTGGAAAA
CACCGAGGTTGCCAAACAAGAACTCTTCAATGTTACTGAGAATCGCTCACACACGGAGGG
CTGGTGCTGCATTTCTGATCTTGCGCCCTCTAAGAAGAATGGATACGTCCAAGTGTCTAT

TCAGGGCCATAATCACTTAATGGTTATGCAGAACCTAGTCCTGGCTGCGAGCGGCTACTC
AAGCGCTGTGTACGATGTGAACGTTCAGGCGAGCCACCTTTGCGACAATCGTCGTTGTTG
CACGCTAGGACACATTGTTCCAGAGTCGGCCAAACTCAACAATAGCCGTAAAGGCTGCCG
TGGGTGGGTACCTTGCCCACACACGGCGTGCACCACTCCTATCTTCATTTGTCCGCACTC
TCCGCCTTGTATAAGGTACCACTCGGCGTACGCCGACATGCAGGCCTTGATCGACGCGCA
CACAGACGGTCATAAGTAAGGCACAGTAGTTTCGTGGCCTATCGCTGGAGTGCTGTGTGT
TTCATCTTCTTTTTCTGGAACTATCGTCCCAGGCGGGTGTCCCTCTCTCACCCGTCTAGT
CCGGCCTTGGTGCGCAAAGACGTGCGTGGATGCACCTTGGTTAAGGAAAATGCTCCTAGT
AGTTTTACAAGGACACTTCTTGTAGGGCAACACATCTGGATGCGAAGAACTCCCGGAAAA
GACAACGGTACCAAGTGACTGGGGTAACGCCCCATTCATGAAGGCAAATAATCAAAAAGA
GGTGCCGGGTAAGAACCCGTTGTATAGGGACAATCCGCAGCTGACTTGGCTCCCTGCCCA
TCCATGGGCATAGCTTAGCGGTTCAACGCTCGGTAAGATGTGGGTTGGCGCGCTGTCCGA
CAGCGCACTGGCTTAAGGTACGGGCTACTCCCACCGGAGACGGTGTCTTTGTCTACCTCG
GGTAACACGCCTCGGGAGAAGCAAAGAGGGTGCAATGGAAAAAAAACTGGTTGCGGGGAA
ATCCCTACCAGAGTCACCTGGTACAAAGGAACTGGCTTGTGGCAGCCAAGCGTTCATAGC
GACGTTGCTTTTTGATCCTTCGATGTCGGCTCTTCCTATCATTGCGAAGCAGAATTCGCC
AAGTGTTGGATTGTTCACCCACTAATAGGGAACGTGAGCTGGGTAGCCACTTTATGCCCA
ACAGCTGGATAGCTGGGCTTCTGTCTTAGGCCGGGACTTCTCACGACCCGTGCCTTGGTC
ACCCAAAAAAGGGCGACCGCAGAGCTCAGGATGGAGCTTCTGGCTTAACTAGTAGAGGAC
AGATAGAGATCGTCAAATTGCGGGAAACCCCTTAGAGCCTCGAGCTACCAAGTGCGTGTC
GAAAGGCATAGACGCAGCCAGCGATGGCTGTCGCATGGCTTGATTCCATCAAGATAGGTA
AAAATGCTCGAGGATTGGGCAATCCGCAGCCAAGCCCCTAAGGTCCCCCCGCGGGCTATG
GGGAAGGTTCAACGACTGTAAGGCGTACCGTGCAAGCGGAAGATACAGTCTAGCCCCGCT
GGAAAGCAGCGCCAAAGTAAGTAGGTTGCAGTGGGTCTGTCCTC
>SRR9587934
GCATGTTTTGGTATGTGTTTCGACAGTCATGCATGTCATAGCCGTAGGATGGGGAGTGAG
CTTCATAGCCCGACCCCTTCTCAGGAGCATGTTTTCATAGCATATTGAGCATACTGGGGT
GGTCTCCCATGTAGCTCGCTCTCTCGGTGCAGGACCTCCCTTTTGCGGGCCACTTCGATG
GCCTCCATTATGGAGAGCATGTTTTGGCATATGAAGCGTTGGTGGTGAGGCTCGTAGCCT
GTATATACATACCAACGCAGTCGTCATATGTCTCTCTGGCGGGGGTCGGTCGTATGACCT
TGTCCCTACCTGGAGCATGTATTTCGTATGATATACTGGACGGCAGCTGCGCTGTTCAGT
CGTCGGCCGTACCTCATGTGCGCGTTTCCAGGTCCGAACGTTTGACATGCAAGGGCATCA
TACCAGAAGCCTGTTTTCTCGTATGAGCATGTAGGTAGGCAGGCTCTTCAGTCGTCTACC
CATAAATGCACTTCGAAGGAACTTTTTGGTATTGAATGGCTTTGTGCCCAGACGTTCTTG
TGAGCTTGTTCTCGCTCTGCGATACGCTGGAAGCTGTGGTATTACCTACAGCAGACGCAT
TCCTAGGGCGGACGGCCTCTCTGCAGATGGGCACATCTGTAGAAAGAGGCTCCCCTTGAT

GACGCAATGCAATGGCTGGGTAAACGGAGTGCTATACAGAAGGGTCAAACCTCTGTGGAA
GTACACTCATACCAGCATTGTTTGTTGAGAGGGGTCGGCTCTACGAGTGAGGACCTATAC
TGCCATCTCATTCAGATGCGTAGACGGTCTTTACTCAGACTGCCTTTCTGGCTGATTGTC
CTCGGTCCCTCGTGCGCACGCGCCGCTGGCTGAATGGGCTTGTCCTTATGGAGTGCCGCG
TTCTTCAAAGAATCCGTCTTCATACAGCGACCTCATTTACATATTGCCGGCTGAATGGGC
TTGTCCTTATGGAGCACTGCATTCTTCTAGGAATTGCTATCTATACAGCGACCTCATTTA
CATACAAGGCACCTTTTTCTACATAGACTTGCCATTTGGCTGATTGTTCTGACAGTCGCT
GCCTCTTCGGAGTTCTGTGTGGACGAGTGGCCTCCTGAGTGGGCCGGTCCTTACGGAGTG
CCCACGTCTGTTATGACTGGGTTCTCCATACAGCGACCTCACTCAGAATTGTATGGCACC
GTGCCGATTGAATGGACAAGCAGTCCTTAGGGAATCGCATGAAAGCATGATGGCACATGG
TGTATGGACCCGTTGGGCCTTGCCTGTGCTATCTTCTCGAGTCGACCGGTCCTCGTCTCC
TCCTTTCGCGACGGGAAGCTCTTGAGTTTACCGTGGGTCAGGACGCTGTGGTATCTGTTT
GTCTGTAGTGTGGGCTCTCTATCAGAACAGCGTATCAGCACACGTCGTCTGGGTCTATGG
TATTTTTGCTTAACGGCCCTGGGTTTGGTAGTATGGTTCCATGTCGGTCCTCTCACCTCC
CATCCGTCATGTCTCTGATATCATCTGCACAGTACGTCTTGTCAGAATATCATTCTGTAT
GGCAGTCGATAGACTAGGCCCTATGCCGCAGCTGTCTCTGACCACTATGCGAGCGCATAC
CTACCTCAATGTGTTCATAGCTGTTATGGATCTTGGGTGTGGTGTGCCAGAACGATACGC
CACCAGATTGTGGAATGGTGCTTGTAGTATTAGAAGAGGACATGTATGAAGAGCACGTTC
TTTTCTCTTATGATGGTGCCGCGGGTGGGGGCCGTTCTCATTCTGCTTGTCGGGTGCTCA
AGGTGCATGTAGGATTGCTGAACCCACTGAATGGTCATGAGAATGGTCATATGGTGAGGT
TCACTTCTCCATCGCTTTGTGTGTGTTGGATGTTTGGGTGGCTTCGGCCGCATGGAAAAT
CTCGGTCTGTAACTCATCAGGGAGGGTTTCCATGTGAGCAATCTCTCCAGTCGAACACAC
ACCCATGGGCGTCCCGCTAGGCAAAATGGAACTCTCGATCCGACACAGCCTGTGGAGTGT
CTGTTCCGTGTGGCGTGAAACGGCTTATACTGGATGTCTTCCCTTTGGACACCAGTCTGT
TGGTATGTACGCATACCATCTAGAGGGAGCAAGAAAAAGGCATGCAAGTGCCACAAAGGA
AGGGAGGCCTCCCTGCTACCGTGTGTCCTCTTGACGATCACGGGGGAGTATATAGAGACA
TTTGGACGAGATCCTTTTTTCACGAGGCGGTGCCCTCGCGCAAAGGTGTTGCTCTGCACG
CATCTTTATGATGATAGTTGCTCGCTCCTATGCGTCCACTCGTGAAAATACCTGCTCTCT
CTGTCTGTTTCCATGACACCCCCATGCGTTGTCATAGAGCACGCTCAGGTACAGAAAGTG
CTGTGACTTTCTTGGCCTGCTGTCACTGAGTCAGCCTGATACTTTGGAGGAATCTTTTCT
GGGCGCTGCCCAGCTGAGAGTTCTTATCAGGCTCTGGAGTGTGCATATGGCTATAATCAT
AATAGTGCATGTGGCCACGCAGAAAAAGCATCGACGTGGAATGGAGAAGAACACCATTCT
GAATGGAACGGTTCTTACTCTGCTTCCGTTGCTTGACGCTGTATGGGAATCGCTTTGCCC
TGGGGGTACACCTTCCATTCCATGACAAAGCATTTGATCTATGCCCCGAGGGTCATTCTC
TATGAAGAGCCCGAAACTTGTCTGAGTGAGAGGCGAAGGTAGCCCTTCATTTATGCGGGG
TCCGCCCTCACACTGTAGTCGGCACAGTCCCTTGTTTATGGACTGCGCAGCTCAATCTTC

GACTGGAGAGGAAAAACAAAACACAAGTATGACACGCACATACGTGTTTTCTAGGACTGT
GACCTGTCGGCTGCCTCGCGGTGGTTGGCATGATGCCCCAAGTGGGGGGGGGGAGAAACA
ATCAGTATGATTGATAAAAACCAGTGGGGAAAAAGGCGCGCTGTCGCACACCCTTGGTCA
CGGCCTCAAGTCGACGGCCCTCACTGGCCCACCCTGCGACTCCTGTTCGCTTTTGACAAG
ACACACCTGCAATACGGAGGAAATGCCTGAGCAGCTGTGCCGCAGCAACTGCGGACGCGT
AGCGATATGCATCCCTGTTGTTTGCGACTTGGTACAAAGAACGTGCGCCGGGAGACGGAT
TGCCCTTTGACTTTTCCACCCAGCCCGCGGCTCTATACCCTTGACGTACCTATTTACCTG
GTTGATCCTGCCAGTAGTCATATGCTTGTCTCAAAGACTAAGCCATGCATGTCTAAGTAT
AAACACTTTTATACTCGTGAAACTGCGAATGGCTCATTAAAACAGTTATAGTTCCTTTGG
GAACAACACTACTTGGATAACCGTAGTAATTCTAGAGCTAATACATGCCTCAACGCCCGA
CTCACGAAGGGTGGTATTTATTGGATAAAAAACCATCGTGCTCTGTCCAGAGCGCTTTGA
GATGATTCACAATAACTTGTCGGATCGCATGGCCTCGTGCTGGCGACTCCTCATTCAAAT
TTCTGCCCTATCAACTTTCGATGGTAGAGTATTGGTCTACCATGGTGTCGACGGGTGACG
GGGAATTAGGGTTCGATTCCGGAGAGGGAGCCTGAGAAACGGCTACCACATCCAAGGAAG
GCAGCAGGCGCGCAAATTACCCAATCCCGACTCGGGGAGGTAGTGACAAAAAATAACAAT
AGGGGGCCCTTTGGGTCTTCTAATTGGAATGAGAACAATTTAAATCCCTTATCGAGGATC
CATTGGAGGGCAAGTCTGGTGCCAGCAGCCGCGGTAATTCCAGCTCCAATAGCGTATATT
AAAGTTGTTGCAGTTAAAACGCTCGTAGTCGGACCTCGGAGCGCCCCCACGGGCCGGGCT
TTTGCCTTGTTGCCAGTGGGAGTTTGTCTCCTTTTGTCGGGACGCGGTGTGCAGGGCCTT
TACTGTCTCTGCGCGTTTACCTGCGGGCCGACCGTTTACTGTGAAGAAAGTAAAGTGTTC
AAAGCAGGCCAATGCCTTGAATATGTGAGCATGGAATAATAGAATAGGACTTGGGCTCTA
TTTTGTTGGTTTCCAGTGACCAAGTAATGATTAATAGGGACGGTTGGGGGCATTCGTATT
TCATTGTCAGAGGTGAAATTCTTGGATTGATGGAAGACGTACAACTGCGAAAGCATCTGC
CATGGATGTTTTCATTGATCAAGAACGAAAGTTAGGGGATCGAAGACGATCAGATACCGT
CGTAGTCTTAACCATAAACGATGCCGACTGGGGATTGGCGGGGGCACTTTTATATGACTC
TGTCAGCACCCTGAGGGAAACCAAAGTCTTTGGGTTCTGGGGGGAGTATGGTCGCAAGGC
TGAAACTTAAAGGAATTGACGGAAGGGCACCACCAGGAGTGGAGCCTGCGGCTTAATTTG
ACTCAACACGGGAAAACTTACCAGGTCCGGACAGAAGAATGATTGACAGACTGAAAGCTC
TTTCTTGATTTTTTGGTTGGTGGTGCATGGCCGTTCTTAGTTGGTGGAGTGATTTGTCTG
GTTAATTCCGTTAACGAACGAGACCTCGTCCTGCTAAATAGGTGCGCGCATGCCAGCACT
GCGTTCTTACCTTCTTAGAGGGACTATGCGCGTCTAGCGTATGGAAGATTGAGGCAATAA
CAGGTCTGTGATGCCCTTAGATGTTCTGGGCCGCACGCGCGCTACACTGATGCATTCAAC
GAGTTTCACTTGATAGTCCTGGGTCGGAAGGCTCGGGTAATCTTTTGAAAGTGCATCGTG
CTGGGGATAGATCATTGCAATTATTGATCTTCAACGAGGAATTCCTTGTAGGCGCAGGTC
ATCAGCCTGCGCCGAATACGTCCCTGCCCTTTGTACACACCGCCCGTCGCTCCTACCGAT
TGAATGGTCCGATGAAATGTCCGGATCGCGGCTAGTGTGATGGACTCGTCCAGAGCGGTA

GCCGTGAGAAGCTCATTAAATCTTATCATTTAGAGGAAGGAGAAGTCGTAACAAGGTTTC
CGTAGGTGAACCTGCGGAAGGATCATTCACAAACTATTGACACAAACACACGCGAACCAA
ATCGTCCGCACAGCAGGTGGCAATGAAAGAGAGACTCTGCATGTCGCCTTTCGGGGTAGC
AAGCAGCACTCTTCTGCCATCGCCTCTGTGCAGGGCGTAAATTCCCATTGAGAGGATGTG
AGGGCACCACAGGAAGCTTTTCCACAGGAAGTCGCCATCCTTCTCCCTCCACGGCGGCGC
TTCTGTGCTTGGCAGTTTTTTTTTGCCTTCTAGGGAGGATGCCGCCAATGGAGCCCCTTA
TATATATACATCATCATATGCCCCTTTTTTTCTTAACCGCTTGCCACAGCTTCTTCTATG
AGGAGCTTGTGGGAAGACTGTCTCCATACAATAACAAAGATACAACTCTTAGCGGTGGAT
ATCTTGGTTCTCGCAACGATGAAGAACGCAGCCAACTGCGATAACTAATGTGAATTGCAG
GACTTCGTGAATCATTAAGTCTTTGAACGCAAGTTGCGCTCATCTCCATGAGGAGGTGAG
CATGCCTGTTTGAGTGTCGATGGGAAAAACAATACATAACAACAATCGGCTCTCTTATCT
TCCTGCCTTCTTGGGGGCAAAAGGTAGTGCATGCCGGAGATGGGCTTTGTCGCCCCTTTC
TCTTGAAAGAGGCGTCGTCCGAAACACGTATCGTTGCCCGCTGGAGGTCATAGAGCATGA
GTTTAAAGCACCACATCTCTCAAGTCAGAGTGAGCGTCTTGAGTAGGTCCGCATTGATTC
AGTTTCAACCCAGGTGGCGTTGGTTACTTGCCATAAGCCTCTCTGGGGTTTCTTTGCTTG
GACCTCTCTTGTCAGTTTCTCTGACGTAGAGACAGGTGCCGTCTCATGGACGTTGCCCTC
TGGAACGTGCTAAAAAAAAAAAAAAGAAAAAACCGTACCACATGTCAAAAGAGGCAGATC
GTGCTTGACGGCGGGAGGTGTTGCATTTCTTGCTTGTCCTGAGCGTCTGCCAGTCTCTGA
CGAGTGTGTTCCTTCCATAAACCCGATCTCAAATCAGGTAGGACCACCCGCTGAATTTAA
GCATATAATTAAGCGGAGGAAAAGAAACTAACAAGGATTCCCTCAGTAGCGGCGAGCGAA
GCGGGAAGAGCTCAACATGAAAACCGTTGGGGGTTCGCGCTTTCCTTCGGGTTGTATAGT
AAAGTGGTGTCGCCTGCGTCGCGCCGGGTATAAATCCCGTCTGAAATGGGGTGTCGTAGA
GGGTTATATCCCCGTCTGTGACCCGGTGTGCGATGCGTTACGGTGACCTTACGATGAGTC
GGGTTGCTTGGGCATGCAGCCCTAAGTGGGTGGTAAACTTCATCTAAAGCTAAATACTAG
CGAGAGACCGATAGCGTACAAGTACCGTAAGGGAAAGATGCAAAGAACTTTGAAAAGAGA
GTTAAACAGGGCTTGAAACCGTTAAGAGTGAAGCGAAGGAGGTCAGTGATCATGGGGCCA
GATCCCCAAGTGCCTGTCATAGGGCACCGTAGCGTTCTGGTTTCCCGGGGCAGCGCCGGC
CGGGGGCGGGGGTTAACAGGGAGGTGGTTGAGGGATTCGTTTTCCGCCTGTACTCCTTTA
TAGCTTGCACCCCGGCCGAGGGATCCGCGACTTGTCGTGTGAGGGCGCTGTCAAAATGGC
CTTCTCCACCCCGTCTTGAAACACGGACCAAGGAGTCTAACATACTCGCGAGTGTTTGGG
TGGTTAAAACCCGAACGCACAATGAAAGTAAACGGTGGCTCTGCGCCGCTGCTTGGTGGG
AACCCTGCTTTTATAATGGGGGGCACTATCACCCGACCCCGATCCCTTTGGGGTGGTTGG
TTTTGAGGTAGAGCGAGTATGATGGGACCCGAAAGATGGTGAACTATTCCTGAGTAGGGC
GAAGCCAGAGGAAACTCTGGTGGAGGCTCGTAGCGATGCTGACGTGCAAATCGCTCGTCG
AACTTGGGTATAGGGGCGAAAGACTAATCGAACCATCTAGTAGCTGGTTCCCTCCGAAGT
TTCCCTCAGGATAGCGGGAGCAAATGCAAGTCAGTATCATCCGGTAAAGCGAATGATTAG

AGGCCTCGGGGGAGTAACTCCTTCGACCTATTCTCAAACTTTAAATAGGTGAGGAGCCGG
CAACGTTGCTTAAAACAATGAACGCCGTGCGGACAATGATTGCTCTCAGTGGGCCGATTT
TGGTAAGCAGAACTGGCGATGCGGGATGAACCGAAAGCGACGCTAAGGTGCCAAACTCCC
AGCTCATTCAGACACCAGAAAAGGTGTTGATTCATACAGACAGCAGGACGGTGGCCATGG
AAGTCGGAATCCGCTAAGGAGTGTGTAACAACTCACCTGCCGAATGGATCAGCCCTGAAA
ATGGATGGCGCTTAAGCGGGGGACCGAGGCGTCGCCGTTGGGGTAAATACGAAGCCCCAA
TGAGTAGGAGGGCGTGGGGGTTTTGAAACACGGTACACTCGCATGGAAGCCTTGGGCGTA
AGCTCGGGTGGAGCGACCCCTAGTGTAGATCTTGGTGGTAGTAGCAAATATTCAAGTGAG
AACCTTGAAGACTGAAGTGGGGAAGGGTTCCACGCGAACAGTGCTTGGACGTGGGTTAGT
CGGTCCTAAGAGATGGGGAAACTCCGTATTCAAGTGCGCTTTTGTAGCGCAGTCCATCGA
AAGGGAAGCGGGTTAACATTCCCGCACCAGGATGTGGAAACTGTGGTTTACCGCAAACTG
CGGCAACGCAACTGAACCTGGGGACGCCCGTAGGAGCTCCGAGCAGGGATCTCTTTCCTG
TTTAACGCGCCCAACCCCCTGGAATCGGTTTGTCCGGAGATAGGGGGCGAAAGGCCGGAA
AAGCACCGCACGTCTTGTGGTGTTCGGAACGCTTGTAATGGCCCTTGAAAACCCAGGGGA
AAGCAGCCGTCTTACACTTGTGTGAGACGAGGAATTCTCACACCTGGCCGTACCCATCAC
CGCATCAGGTCCCCAAGGTGAACAGCCTCTAGTCAATAGAATAATGTAGATAAGGGAAGT
CGGCAAAATAGATCCGTAACTTTGGGAAAAGGATTGGCTCTAATGTTCGGGCTTGTCGGG
ACGCGCGGGCGTAGATGCCAATGAGCGGCAGACGTCTAGCCTCTTTGGAGGCTGGATTAA
CGCTGCTTGTGGGAAGAAACCGTGTCGATGGAGGCAGGAACGCTTATCGATCTTGTCTCT
TCCCGCTTGCGATCAACGAACGGATTAGAACTGGTACGGACAAGGGGAATCCGACTGTTT
AATTAAAACAAAGCATTGCGACGGCCGGAAACGGTGTTGGCGCAATGTGATTTCTGCCCA
GTGCTCTGAATGTCAAAGTGAAGAAATTCAACCAAGCGCGGGTAAACGGCGGGAGTAACT
ATGACTCTTAAACAAAACCATACTACCTACGGGAGTCAAAAGTGGGGGCGGCGTCTGGTG
TAGAAATACACGTACGACACGCTTTTTTTCCTGCTAGTCGATCTGACCTTTGTTCTTTGG
ACTGTGGGCGGTCAGCGGCAACATAACCTGGTACGGGGAAGGCCTCGACGTTGAGTGCTG
CATCTTGGAAGTATCTTGACACAATAAGAGAAAAGAGATTGCGCTTAATAGGGGGTTGAT
CTCGTGGCGAGCCGACGCCGGTGTGAACTGGTGGCAATGGCCGTCGTAGAGCACGCAAAG
GTATGGGCTGACTTTTTCTCCTCAAAAGTTGGCTTAAGGGACGTGCCATCCCACACGATA
ATAAAGTGTGCAGTCTGTAAGAGCTCCCACAGGGCAAAGACAGACTGGGATGCAGACGGA
GACGGCAAAAACAACAACATCCGTTCTGCTTCTGGGCGCAAATGCTTAAGGTAGCCAAAT
GCCTCGTCATCTAAACTAATTGGATGAGATAGTCTGAGTGAAAAGCTTATGCAGGCTAGT
ACCGGAAAAAAAAAAAAAAAGAGGGGCTCTGCCACTCTGACGGTGCGACACTCCTTGTAC
AGGGAAACCCTCCGCATGGAAAGGCACGTGGTACCACCCTCGCGGGGAAACTCGCGAGCG
GGATCCCGGGTAATGACCGGAGCCGATGGTAAAAACTCACGTGACCGATGGGGTGACCCT
GTGGCGAGACGCTAAGTTGTGGGACATCACCCTCTCACAATACGCATCCGTCGCAGAGAC
TGAACGGGAGTGGGCTCCGTTGACGGCCGGCCGGCCGGCCAGCCAGCCGTCTTCCAGGGG

CTTAAGATACAGTCCGGGCCTCAGGGATGGAAAAACCTGGGGAACTACCAGAATTAGTGA
CGCGCATGAATGGATTAACGAGATTCCCACTGTCCCTATCTACTATCTAGCGAAACCACA
GCCAAGGGAACGGGCTTGGAAGAATCAGCGGGGAAAGAAGACCCTGTTGAGCTTGACTCT
AGTCCGACTTTGTGAGATGAGTTGGGAGGTGTAGTCTAAGAGGGAGGGGCAACTCGACCG
TGAAATACCTCTACTCCCAAGCTTGTTTTACTTACTCCGGCGAGCAGGGATGGGTGGTAA
CACCCTTTTTCTAGGATTAACACCTCTTCGGGGGTGGACCTGCGCGGAGGACATTGTCAG
GTGGGGAGTTTGGCTGGGGCGGCACATCTGTTAAAAGATAACGCAGGTGTCCTAAGATGA
GCTCAGTGAGAACAGAAATCTCACGTAGAACAAAAGGGTAAAAGCTCATTTGATTTTGAT
TTTCAGTACGAATACAAACTGTGAAAGCATGGCCTATCGATCCTTTTGACCTTCAAGGTT
TGAAGCAAGAGGTGTCAGAAAAGTTAACACGCGTTGTGGTAGCTTTTAAGAGTGGACTCA
AAGTGTTCCGCTAGTCCCCCCTCGGACCGCCACTTACAAAAGGCCTGGGGATGGGGGCGA
CACTGTCAAATTGCGGGGACTCCCTAAAGCCTGTCTCTACCAAGCCAGCGCGGCGAAAGC
CCCTGGTGGCCGGGGTAATGACCTAGGGTATGGTAAAAGCGAGACAGGATGGAACAATGG
GCAATCCGCAGCCAAGTCCTAAGGCTTTTTGAAAAGCGGCTATGGATGCAGTTCAACGAC
TAGATGGCAGTGGGCTGAGAGAGCCCACCGCGGCACCCTTTCGGCTTAAGATATAGTCTT
GCCCACGTGAAAGCGTGCCTGCAGATGAACGCCACCATGTATGGCCGGAGCTGCAGGAGA
CCGCACTTGATGTGCAGTCGGGAATCGACAGTACCACAGGGATGAAAAAAAAACACACAA
TTAATTTGTGGCCATCATACAGTCGTATGATATTATCGTCCCAGGCGGGTGCGCCTCTCT
CACCCGTCTAGTCCGGCCTTGGTGCGCAAAAAATGCTTGCGTGGATGCGCCTTGGTTAAG
GAAAATGCTCCTAGTAGTTTTACAAGGACACTTCTTGTAGGGCAACACATCTGGATGCGA
AGAACTCCCGGAAAAGACAACGGTACCAAGTGACTGGGGTAACGCCCCATTCATGAAGGC
AAATAATCAAAAAGAGGTGCCGGGTAAGAACCCGTTGTATAGGGACAATCCGCAGCTGAC
TTGGCTCCCTGCCCATCCATGGGCATAGCTTAGCGGTTCAACGCTCGGTAAGATGTGGGT
TGGCGCGCTGTCCGACAGCGCACTGGCTTAAGGTACGGGCTACTCCCACCGGAGACGGTG
TCTTTGTCTACCTCGGGTAACACGCCTCGGGAGAAGCAAAGAGGGTGCAATGGAAAAAAA
ACTGGTTGCGGGGAAATCCCTACCAGAGTCACCTGGTACAAAGGAACTGGCTTGTGGCAG
CCAAGCGTTCATAGCGACGTTGCTTTTTGATCCTTCGATGTCGGCTCTTCCTATCATTGC
GAAGCAGAATTCGCCAAGTGTTGGATTGTTCACCCACTAATAGGGAACGTGAGCTGGGTA
GCCACTTTATGCCCAACAGCTGGATAGCTGGGCTTCTGTCTTAGGCCGGGACTTCTCACG
ACCCGTGCCTTGGTCACCCAAAAAAGGGCGACCGCAGAGCTCAGGATGGAGCTTCTGGCT
TAACTAGTAGAGGACAGATAGAGATCGTCAAATTGCGGGAAACCCCTTAGAGCCTCGAGC
TACCAAGTGCGTGTCGAAAGGCATAGACGCAGCCAGCGATGGCTGTCGCATGGCTTGATT
CCATCAAGATAGGTAAAAATGCTCGAGGATTGGGCAATCCGCAGCCAAGCCCCTAAGGTC
CCCCCGCGGGCTATGGGGAAGGTTCAACGACTGTAAGGCGTACCGTGCAAGCGGAAGATA
CAGTCTAGCCCCGCTGGAAAGCAGCGCCAAAGTAAGTAGGTTGCAGTGGGTCTGTCCTC
>SRR9587936

GCATGTTTTGGTATGTGTTTCGACAGTCATGCATGTCATAGCCGTAGGATGGGGAGTGAG
CTTCATAGCCCGACCCCTTCTCAGGAGCATGTTTTCATAGCATATTGAGCATACTGGGGT
GGTCTCCCATGTAGCTCGCTCTCTCGGTGCAGGACCTCCCTTTTGCGGGCCACTTCGATG
GCCTCCATTATGGAGAGCATGTTTTGGCATATGAAGCGTTGGTGGTGAGGCTCGTAGCCT
GTATATACATACCAACGCAGTCGTCATATGTCTCTCTGGCGGGGGTCGGTCGTATGACCT
TGTCCCTACCTGGAGCATGTATTTTGTATGATATACTGGACGGCAGCTGCGCTGTTCAGT
CGTCGGCCGTACCTCATGTGCGCGTTTCCAGGTCCGAACGTTTGACATGCAAGGGCATCA
TACCAGAAGCCTGTTTTCTCGTATGAGCATGTAGGTAGGCAGGCTCTTCAGTCGTCTACC
CATAAATGCACTTCGAAGGAACTTTTTGGTATTGAATGGCTTTGTGCCCAGACGTTCTTG
TGAGCTTGTTCTCGCTCTGCGATACGCTGGAAGCTGTGGTATTACCTACAGCAGACGCAT
TCCTAGGGCGGACGGCCTCTCTGCAGATGGGCACATCTGTAGAAAGAGGCTCCCCTTGAT
GACGCAATGCAATGGCTGGGTAAACGGAGTGCTATACAGAAGGGTCAAACCTCTGTGGAA
GTACACCCATACCAGCATTGTTTGTTGAGAGGGGTCGGCTCTACGAGTGAGGACCTATAC
TGCCATCTCATTCAGATGCGTAGACGGTCTTTACTCAGACTGCCTTTCTGGCTGATTGTC
CTCGGTCCCTCGTGCGCACGCGCCGCTGGCTGAATGGGCTTGTCCTTATGGAGTGCCGCG
TTCTTCAAAGAATCCGTCTTCATACAGCGACCTCATTTACATATTGCCGGCTGAATGGGC
TTGTCCTTATGGAGCACTGCATTCTTCTAGGAATTGCTATCTATACAGCGACCTCATTTA
CATACAAGGCACCTTTTTCTACATAGACTTGCCATTTGGCTGATTGTTCTGACAGTCGCT
GCCTCTTCGGAGTTCTGTGTGGACGAGTGGCCTCCTGAGTGGGCCGGTCCTTACGGAGTG
CCCACGTCTGTTATGACTGGGTTCTCCATACAGCGACCTCACTCAGAATTGTATGGCACC
GTGCCGATTGAATGGACAAGCAGTCCTTAGGGAATCGCATGAAAGCATGACGGCACATGG
TGTATGGACCCGTTGGGCCTTGCCTGTGCTATCTTCTCGAGTCGACCGGTCCTCGTCTCC
TCCTTTCGCGACGGGAAGCTCTTGAGTTTACCGTGGGTCAGGACGCTGTGGTATCTGTTT
GTCTGTAGTGTGGGCTCTCTATCAGAACAGCGTATCAGCACACGTCGTCTGGGTCTATGG
TATTTTTGCTTAACGGCCCTGGGTTTGGTAGTATGGTTCCATGTCGGTCCTCTCACCTCC
CAACCGTCATGTCTCTGATATCATCTGCACAGTACGTCTTGTCAGAATATCATTCTGTAT
GGCAGTCGATAGACTAGGCCCTATGCCGCAGCTGTCTCTGACCACTATGCGAGCGCATAC
CTACCTCAATGTGTTCATAGCTGTTATGGATCTTGGGTGTGGTGTGCCAGAACGATACGC
CACCAGATTGTGGAATGGTGCTTGTAGTATTAGAAGAGGACATGTATGAAGAGCACGTTC
TTTTCTCTTATGATGGTGCCGCGGGTGCGGGCCGTTTTCATTCTGCTTGTCGGGTGCTCA
AGGTGCATGTAGGATTGCTGAACCCACTGAATGGTCATGAGAATGGTCATATGGTGAGGT
TCACTTCTCCATCGCTTTGTGTGTGTTGGATGTTTGGGTGGCTTCGGCCGCATGGAAAAT
CTCGGTCTGTAACTCATCAGGGAGGGTTTCCATGTGAGCAATCTCTCCAGTCGAACACAC
ACCCATGGGCGTCCCGCTAGGCAAAATGGAACTCTCGATCCGACACAGCCTGTGGAGTGT
TTGTTCCGTGTGGCGTGAAACGGCTTATACTGGATGTCTTCCCTTTGGACACCAGTCTGT
TGGTATGTACGCATACCATCTAGAGGGAGCAAGAAAAAGGCATGCAAGTGCCACAAAGGA

AGGGAGGCCTTCCTGCTACCGTGTGTCCTCTTGACGATCACGGGGGAGTATATAGAGACA

TTTGGACGAGATCCGTTTTTCACGAGGCGGTGCCCTCGCGCAAAGGTGTTGCTCTGCACG

CATCTTTATGATGATAGTTGCTCGCTCCTATGCGTCCACTCGTGAAAATACCTGCTCTCT

CTGTCTGTTTCCATGACACCCCCATGCGTTGTCATAGAGCATGCTCAGGTACAGAAAGTG

CTGTGACTTTCTTGGCCTGCTGTCACTGAGTCAGCCTGATACTTTGGAGGAATCTTTTCT

GGGCACTGCCCAGCTGAGAGTTCTTATCAGGCTCTGGAGTGTGCATATGCTTATATTCAT

TATAGTGCATGTGGCTACGCAGAAAAAGCATCGACGTGGAATGGAGAAGAACACCATTCT

GAATGGAACGGTTCTTACTCTACTTCTGTTGCTTGACGCTGTATGGGAATCGCTTTGCTC

TGGGGGTACACCTTCCATTCCATGACAAAGCATTTGATCTATGCCCCGAGGGTCATTCTC

TATGAAGAGCCCGAAACTTGTCTGAGTGAGAGGCGAAGGTAGTCCTTCATTTATGCGGGG

TCCGCCCTCACACTGTAGTCGGCACAGTCCCTTGTTTATGGACTGCGCAGCTCAATCTTC

GACTGGAGAGGAAAAACAAAACACAAGTATGACACGCACATACGTGTTTTCTAGGACTGT

GACCTGTCGGCTGCCTCGCGGTGGTTGGCATGATGCCCCAAGTGGGGGGGGGAGAAACAA

TCAGTATGATTGATAAAAACCAGTGGGGAAAAAGGCGCGCTGTCGCACACCCTTGGTCAC

GGCCTCAAGTCGACGGCCCTCACTGGCCCACCCTGCGACTCCTGTTCGCTTTTGACAAGA

CACACCTGCAATACGGAGGAAATGCCTGAGCAGCTGTGCCGCAGCAACTGCGGACGCGTA

GCGATATGCATCCCTGTTGTTTGCGACTTGGTACAAAGAACGTGCGCCGGGAGACGGATT

GCCCTTTGACTTTTCCACCCAGCCCGCGGCTCTATACCCTTGACGTACCTATTTACCTGG

TTGATCCTGCCAGTAGTCATATGCTTGTCTCAAAGACTAAGCCATGCATGTCTAAGTATA

AACACTTTTATACTCGTGAAACTGCGAATGGCTCATTAAAACAGTTATAGTTCCTTTGGG

AACAACACTACTTGGATAACCGTAGTAATTCTAGAGCTAATACATGCCTCAACGCCCGAC

TCACGAAGGGTGGTATTTATTGGATAAAAAACCATCGTGCTCTGTCCAGAGCGCTTTGAG

ATGATTCACAATAACTTGTCGGATCGCATGGCCTCGTGCTGGCGACTCCTCATTCAAATT

TCTGCCCTATCAACTTTCGATGGTAGAGTATTGGTCTACCATGGTGTCGACGGGTGACGG

GGAATTAGGGTTCGATTCCGGAGAGGGAGCCTGAGAAACGGCTACCACATCCAAGGAAGG

CAGCAGGCGCGCAAATTACCCAATCCCGACTCGGGGAGGTAGTGACAAAAAATAACAATA

GGGGGCCCTTTGGGTCTTCTAATTGGAATGAGAACAATTTAAATCCCTTATCGAGGATCC

ATTGGAGGGCAAGTCTGGTGAACGTATGCATCAAAGCCCTCCGTAATAAAAAAAAACAGC

GCCAGATGTCTCTCTCTGGAAGCACACCACGCACGGGGTTCGTTCCTCTCTGCATGGCCG

TGTGAAAAAGGCGAGTAGGCAAACGATGGGAAACCCGGCTTGGAATAGTAACCAGCCACA

GAGCCCCCATCATACACTTCTAAATTGCCGGGGAGTGCCCGCGAGGCCTTGACTACCGCG

GTCCGCCGAGAGGTGAGAGCGAGCAATCGCTCAGGACGCAGCAGTGTACGTAGTGGTACA

TGGATGGTAAAAACGTCAAGGATAGGGTTGATCCGCAGGGAAGCGCGAGTGCCGCCATGG

TCGGTGGGCAAAGGCGAACCTTCAGAGACTCTAATGGAGTGGGCGCAAGCTTAAGGGAGA

GTCCAATCCACTGGGAAACCAGGTCCACAGCAGTAACGGGGTGATGCTCGCCCCCAGAGGA

CTGTGGAAGGCAACCCCACCAAAAGGGTGTGTGTTGTCGGGTAGCAATGGCCAGCAGCCG

CGGTAATTCCAGCTCCAATAGCGTATATTAAAGTTGTTGCAGTTAAAACGCTCGTAGTCG
GACCTCGGAGCGCCCCCACGGGCCGGGCTTTTGCCTTGTTGCCAGTGGGAGTTTGTCTCC
TTTTGTCGGGACGCGGTGTGCAGGGCCTTTACTGTCTCTGCGCGTTTACCTGCGGGCCGA
CCGTTTACTGTGAAGAAAGTAAAGTGTTCAAAGCAGGCCAATGCCTTGAATATGTGAGCA
TGGAATAATAGAATAGGACTTGGGCTCTATTTTGTTGGTTTTCAGTGACCAAGTAATGAT
TAATAGGGACGGTTGGGGGCATTCGTATTTCATTGTCAGAGGTGAAATTCTTGGATTGAT
GGAAGACGTACAACTGCGAAAGCATCTGCCATGGATGTTTTCATTGATCAAGAACGAAAG
TTAGGGGATCGAAGACGATCAGATACCGTCGTAGTCTTAACCATAAACGATGCCGACTGG
GGATTGGCGGGGGCACTTTTATATGACTCTGTCAGCACCCTGAGGGAAACCAAAGTCTTT
GGGTTCTGGGGGGAGTATGGTCGCAAGGCTGAAACTTAAAGGAATTGACGGAAGGGCACC
ACCAGGAGTGGAGCCTGCGGCTTAATTTGACTCAACACGGGAAAACTTACCAGGTCCGGA
CAGAAGAATGATTGACAGACTGAAAGCTCTTTCTTGATTTTTTGGTTGGTGGTGCATGGC
CGTTCTTAGTTGGTGGAGTGATTTGTCTGGTTAATTCCGTTAACGAACGAGACCTCGTCC
TGCTAAATAGGTGCGCGCATGCCAGCACTGCGTTCTTACCTTCTTAGAGGGACTATGCGC
GTCTAGCGTATGGAAGATTGAGGCAATAACAGGTCTGTGATGCCCTTAGATGTTCTGGGC
CGCACGCGCGCTACACTGATGCATTCAACGAGTTTCACTTGATAGTCCTGGGTCGGAAGG
CTCGGGTAATCTTTTGAAAGTGCATCGTGCTGGGGATAGATCATTGCAATTATTGATCTT
CAACGAGGAATTCCTTGTAGGCGCAGGTCATCAGCCTGCGCCGAATACGTCCCTGCCCTT
TGTACACACCGCCCGTCGCTCCTACCGATTGAATGGTCCGATGAAATGTCCGGATCGCGG
CTAGTGTGATGGACTCGTCCAGAGCGGTAGCCGTGAGAAGCTCATTAAATCTTATCATTT
AGAGGAAGGAGAAGTCGTAACAAGGTTTCCGTAGGTGAACCTGCGGAAGGATCATTCACA
AACTATTGACACAAACACACGCGAACCAAATCGTCCGCACAGCAGGTGGCAATGAAAGAG
AGACTCTGCATGTCGCCTTTCGGGGTAGCAAGCAGCACTCTTCTGCCATCGCCTCTGTGC
AGGGCGTAAATTCCCATTGAGAGGATGTGAGGGCACCACAGGAAGCTTTTCCACAGGAAG
TCGCCATCCTTCTCCCTCCACGGCGGCGCTTCTGTGCTTGGCAGTTTTTTTTTGCCTTCT
AGGGAGGATGCCGCCAATGGAGCCCCTTATATATATACATCATCATATGCCCCTTTTTTT
CTTAACCGCTTGCCACAGCTTCTTCTATGAGGAGCTTGTGGGAAGACTGTCTCCATACAA
TAACAAAGATACAACTCTTAGCGGTGGATATCTTGGTTCTCGCAACGATGAAGAACGCAG
CCAACTGCGATAACTAATGTGAATTGCAGGACTTCGTGAATCATTAAGTCTTTGAACGCA
AGTTGCGCTCATCTCCATGAGGAGGTGAGCATGCCTGTTTGAGTGTCGATGGGAAAAACA
ATACATAACAACAATCGGCTCTCTTATCTTCCTGCCTTCTTGGGGGCAAAAGGTAGTGCA
TGCCGGAGATGGGCTTTGTCGCCCCTTTCTCTTGAAAGAGGCGTCGTCCGAAACACGTAT
CGTTGCCCGCTGGAGGTCATAGAGCATGAGTTTAAAGCACCACATCTCTCAAGTCAGAGT
GAGCGTCTTGAGTAGGTCCGCATTGATTCAGTTTCAACCCAGGTGGCGTTGGTTACTTGC
CATAAGCCTCTCTGGGGTTTCTTTGCTTGGACCTCTCTTGTCAGTTTCTCTGACGTAGAG
ACAGGTGCCGTCTCATGGACGTTGCCCTCTGGAACGTGCTAAAAAAAAAAAAAAAGAAAAA

ACCGTACCACATGTCAAAAGAGGCAGATCGTGCTTGACGGCGGGAGGTGTTGCATTTCTT
GCTTGTCCTGAGCGTCTGCCAGTCTCTGACGAGTGTGTTCCTTCCATAAACCCGATCTCA
AATCAGGTAGGACCACCCGCTGAATTTAAGCATATAATTAAGCGGAGGAAAAGAAACTAA
CAAGGATTCCCTCAGTAGCGGCGAGCGAAGCGGGAAGAGCTCAACATGAAAACCGTTGGG
GGTTCGCGCTTTCCTTCGGGTTGTATAGTAAAGTGGTGTCGCCTGCGTCGCGCCGGGTAT
AAATCCCGTCTGAAATGGGGTGTCGTAGAGGGTTATATCCCCGTCTGTGACCCGGTGTGC
GATGCGTTACGGTGACCTTACGATGAGTCGGGTTGCTTGGGCATGCAGCCCTAAGTGGGT
GGTAAACTTCATCTAAAGCTAAATACTAGCGAGAGACCGATAGCGTACAAGTACCGTAAG
GGAAAGATGCAAAGAACTTTGAAAAGAGAGTTAAACAGGGCTTGAAACCGTTAAGAGTGA
AGCGAAGGAGGTCAGTGATCATGGGGCCAGATCCCCAAGTGCCTGTCATAGGGCACCGTA
GCGTTCTGGTTTCCCGGGGCAGCGCCGGCCGGGGGCGGGGGTTAACAGGGAGGTGGTTGA
GGGATTCGTTTTCCGCCTGTACTCCTTTATAGCTTGCACCCCGGCCGAGGGATCCGCGAC
TTGTCGTGTGAGGGCGCTGTCAAAATGGCCTTCTCCACCCCGTCTTGAAACACGGACCAA
GGAGTCTAACATACTCGCGAGTGTTTGGGTGGTTAAAACCCGAACGCACAATGAAAGTAA
ACGGTGGCTCTGCGCCGCTGCTTGGTGGGAACCCTGCTTTTATAATGGGGGGCACTATCA
CCCGACCCCGATCCCTTTGGGGTGGTTGGTTTTGAGGTAGAGCGAGTATGATGGGACCCG
AAAGATGGTGAACTATTCCTGAGTAGGGCGAAGCCAGAGGAAACTCTGGTGGAGGCTCGT
AGCGATGCTGACGTGCAAATCGCTCGTCGAACTTGGGTATAGGGGCGAAAGACTAATCGA
ACCATCTAGTAGCTGGTTCCCTCCGAAGTTTCCCTCAGGATAGCGGGAGCAAATGCAAGT
CAGTATCATCCGGTAAAGCGAATGATTAGAGGCCTCGGGGGAGTAACTCCTTCGACCTAT
TCTCAAACTTTAAATAGGTGAGGAGCCGGCAACGTTGCTTAAAACAATGAACGCCGTGCG
GACAATGATTGCTCTCAGTGGGCCGATTTTGGTAAGCAGAACTGGCGATGCGGGATGAAC
CGAAAGCGACGCTAAGGTGCCAAACTCCCAGCTCATTCAGACACCAGAAAAGGTGTTGAT
TCATACAGACAGCAGGACGGTGGCCATGGAAGTCGGAATCCGCTAAGGAGTGTGTAACAA
CTCACCTGCCGAATGGATCAGCCCTGAAAATGGATGGCGCTTAAGCGGGGGACCGAGGCG
TCGCCGTTGGGGTAAATACGAAGCCCCAATGAGTAGGAGGGCGTGGGGGTTTTGAAACAC
GGTACACTCGCATGGAAGCCTTGGGCGTAAGCTCGGGTGGAGCGACCCCTAGTGTAGATC
TTGGTGGTAGTAGCAAATATTCAAGTGAGAACCTTGAAGACTGAAGTGGGGAAGGGTTCC
ACGCGAACAGTGCTTGGACGTGGGTTAGTCGGTCCTAAGAGATGGGGAAACTCCGTATTC
AAGTGCGCTTTTGTAGCGCAGTCCATCGAAAGGGAAGCGGGTTAACATTCCCGCACCAGG
ATGTGGAAACTGTGGTTTACCGCAAACTGCGGCAACGCAACTGAACCTGGGGACGCCCGT
AGGAGCTCCGAGCAGGGATCTCTTTCCTGTTTAACGCGCCCAACCCCCTGGAATCGGTTT
GTCCGGAGATAGGGGGCGAAAGGCCGGAAAAGCACCGCACGTCTTGTGGTGTTCGGAACG
CTTGTAATGGCCCTTGAAAACCCAGGGGAAAGCAGCCGTCTTACACTTGTGTGAGACGAG
GAATTCTCACACCTGGCCGTACCCATCACCGCATCAGGTCCCCAAGGTGAACAGCCTCTA
GTCAATAGAATAATGTAGATAAGGGAAGTCGGCAAAATAGATCCGTAACTTTGGGAAAAG

GATTGGCTCTAATGTTCGGGCTTGTCGGGACGCGCGGGCGTAGATGCCAATGAGCGGCAG

ACGTCTAGCCTCTTTGGAGGCTGGATTAACGCTGCTTGTGGGAAGAAACCGTGTCGATGG

AGGCAGGAACGCTTATCGATCTTGTCTCTTCCCGCTTGCGATCAACGAACGGATTAGAAC

TGGTACGGACAAGGGGAATCCGACTGTTTAATTAAAACAAAGCATTGCGACGGCCGGAAA

CGGTGTTGGCGCAATGTGATTTCTGCCCAGTGCTCTGAATGTCAAAGTGAAGAAATTCAA

CCAAGCGCGGGTAAACGGCGGGAGTAACTATGACTCTTAAACAAAACCATACTACCTACG

GGAGTCAAAAGTGGGGGCGGCGTCTGGTGTAGAAATACACGTACGACACGCTTTTTTTCC

TGCTAGTCGATCTGACCTTTGTTCTTTGGACTGTGGGCGGTCAGCGGCAACATAACCTGG

TACGGGGAAGGCCTCGACGTTGAGTGCTGCATCTTGGAAGTATCTTGACACAATAAGAGA

AAAGAGATTGCGCTTAATAGGGGGTTGATCTCGTGGCGAGCCGACGCCGGTGTGAACTGG

TGGCAATGGCCGTCGTAGAGCACGCAAAGGTATGGGCTGACTTTTTCTCCTCAAAAGTTG

GCTTAAGGGACGTGCCATCCCACACGATAATAAAGTGTGCAGTCTGTAAGAGCTCCCACA

GGGCAAAGACAGACTGGGATGCAGACGGAGACGGCAAAAACAACAACATCCGTTCTGCTT

CTGGGCGCAAATGCTTAAGGTAGCCAAATGCCTCGTCATCTAAACTAATTGGATGAGATA

GTCTGAGTGAAAAGCTTATGCAGGCTAGTACCGGAAAAAAAAAAAAAAAGAGGGGCTCTG

CCACTCTGACGGTGCGACACTCCTTGTACAGGGAAACCCTCCGCATGGAAAGGCACGTGG

TACCACCCTCGCGGGGAAACTCGCGAGCGGGATCCCGGGTAATGACCGGAGCCGATGGTA

AAAACTCACGTGACCGATGGGGTGACCCTGTGGCGAGACGCTAAGTTGTGGGACATCACC

CTCTCACAATACGCATCCGTCGCAGAGACTGAACGGGAGTGGGCTCCGTTGACGGCCGGC

CGGCCGGCCAGCCAGCCGTCTTCCAGGGGCTTAAGATACAGTCCGGGCCTCAGGGATGGA

AAAACCTGGGGAACTACCAGAATTAGTGACGCGCATGAATGGATTAACGAGATTCCCACT

GTCCCTATCTACTATCTAGCGAAACCACAGCCAAGGGAACGGGCTTGGAAGAATCAGCGG

GGAAAGAAGACCCTGTTGAGCTTGACTCTAGTCCGACTTTGTGAGATGAGTTGGGAGGTG

TAGTCTAAGAGGGAGGGGCAACTCGACCGTGAAATACCTCTACTCCCAAGCTTGTTTTAC

TTACTCCGGCGAGCAGGGATGGGTGGTAACACCCTTTTTCTAGGATTAACACCTCTTCGG

GGGTGGACCTGCGCGGAGGACATTGTCAGGTGGGGAGTTTGGCTGGGGCGGCACATCTGT

TAAAAGATAACGCAGGTGTCCTAAGATGAGCTCAGTGAGAACAGAAATCTCACGTAGAAC

AAAAGGGTAAAAGCTCATTTGATTTTGATTTTCAGTACGAATACAAACTGTGAAAGCATG

GCCTATCGATCCTTTTGACCTTCAAGGTTTGAAGCAAGAGGTGTCAGAAAAGTTAACACG

CGTTGTGGTAGCTTTTAAGAGTGGACTCAAAGTGTTCCGCTAGTCCCCCCTCGGACCGCC

ACTTACAAAAGGCCTGGGGATGGGGGCGACACTGTCAAATTGCGGGGACTCCCTAAAGCC

TGTCTCTACCAAGCCAGCGCGGCGAAAGCCCCTGGTGGCCGGGGTAATGACCTAGGGTAT

GGTAAAAGCGAGACAGGATGGAACAATGGGCAATCCGCAGCCAAGTCCTAAGGCTTTTTG

AAAAGCGGCTATGGATGCAGTTCAACGACTAGATGGCAGTGGGCTGAGAGAGCCCACCGC

GGCACCCTTTCGGCTTAAGATATAGTCTTGCCCACGTGAAAGCGTGCCTGCAGATGAACG

CCACCATGTATGGCCGGAGCTGCAGGAGACCGCACTTGATGTGCAGTCGGGAATCGACAG

TACCACAGGGATGAAAAAAAAACACACAATTAATTTGTGGCCATCATACAGTCGTATGAT
ATTATCGTCCCAGGCGGGTGCGCCTCTCTCACCCGTCTAGTCCGGCCTTGGTGCGCAAAA
AATGCTTGCGTGGATGCGCCTTGGTTAAGGAAAATGCTCCTAGTAGTTTTACAAGGACAC
TTCTTGTAGGGCAACACATCTGGATGCGAAGAACTCCCGGAAAAGACAACGGTACCAAGT
GACTGGGGTAACGCCCCATTCATGAAGGCAAATAATCAAAAAGAGGTGCCGGGTAAGAAC
CCGTTGTATAGGGACAATCCGCAGCTGACTTGGCTCCCTGCCCATCCATGGGCATAGCTT
AGCGGTTCAACGCTCGGTAAGATGTGGGTTGGCGCGCTGTCCGACAGCGCACTGGCTTAA
GGTACGGGCTACTCCCACCGGAGACGGTGTCTTTGTCTACCTCGGGTAACACGCCTCGGG
AGAAGCAAAGAGGGTGCAATGGAAAAAAAACTGGTTGCGGGGAAATCCCTACCAGAGTCA
CCTGGTACAAAGGAACTGGCTTGTGGCAGCCAAGCGTTCATAGCGACGTTGCTTTTTGAT
CCTTCGATGTCGGCTCTTCCTATCATTGCGAAGCAGAATTCGCCAAGTGTTGGATTGTTC
ACCCACTAATAGGGAACGTGAGCTGGGTAGCCACTTTATGCCCAACAGCTGGATAGCTGG
GCTTCTGTCTTAGGCCGGGACTTCTCACGACCCGTGCCTTGGTCACCCAAAAAAGGGCGA
CCGCAGAGCTCAGGATGGAGCTTCTGGCTTAACTAGTAGAGGACAGATAGAGATCGTCAA
ATTGCGGGAAACCCCTTAGAGCCTCGAGCTACCAAGTGCGTGTCGAAAGGCATAGACGCA
GCCAGCGATGGCTGTCGCATGGCTTGATTCCATCAAGATAGGTAAAAATGCTCGAGGATT
GGGCAATCCGCAGCCAAGCCCCTAAGGTCCCCCCGCGGGCTATGGGGAAGGTTCAACGAC
TGTAAGGCGTACCGTGCAAGCGGAAGATACAGTCTAGCCCCGCTGGAAAGCAGCGCCAAA
GTAAGTAGGTTGCAGTGGGTCTGTCCTC
>SRR9587941_SRR9587946_SRR9587959
GCATGTTTTGGTATGTGTTTCGACAGTCATGCATGTCATAGCCGTAGGATGGGGAGTGAG
CTTCATAGCCCGACCCCTTCTCAGGAGCATGTTTTCATAGCATATTGAGCATACTGGGGT
GGTCTCCCATGTAGCTCGCTCTCTCGGTGCAGGACCTCCCTTTTGCGGGCCACTTCGATG
GCCTCCATTATGGAGTGCATGTTTTGGCATATGAAGCGTTGGTGGTGAGGCTCGTAGCCT
GTATATACATACCAACGCAGTCGTCATATGTCTCTCTGGCGGGGGTCGGTCGTATGACCT
TGTCCCTACCTGGAGCATGTATTTTGTATGATATACTGGACGGCAGCTGCGCTGTTCAGT
CGTCAGCCGTACCTCATGTGCGCGTTTCCAGGTCCGAACGTTTGACATGCAAGGGCATCA
TACCAGAAGCCTGTTTTCTCGTATGAGCATGTAGGTAGGCAGGCTCTTCAGTCGTCTACC
CATAAATGCACTTCGAAGGAACTTTTTGGTATTGAATGGCTTTGTGCCCAGACGTTCTTG
TGAGCTTGTTCTCGCTCTGCGATACGCTGGAAGCTGTGGTATTACCTACAGCAGACGCAT
TCCTAGGGCGGACGGCCTCTCTGCAGATGGGCACATCTGTAGAAAGAGGCTCCCCTTGAT
GACGCAATGCAATGGCTGGGTAAACGGAGTGCTATACAGAAGGGTCAAACCTCTGTGGAA
GTACACTCATACCAGCATTGTTTGTTGAGAGGGGTCGGCTCTACGAGTGAGGCCCTATAC
TGCCATCTCATTCAGATGCGTAGACGGTCTTTACTCAGACTGCCTTTCTGGCTGATTGTC
CTCGGTCCCTCGTGCGCACGCGCCGCTGGCTGAATGGGCTTGTCCTTATGGAGTGCCGCG
TTCTTCAAAGAATCCGTCTTCATACAGCGACCTCATTTACATATTGCCGGCTGAATGGGC

TTGTCCTTATGGAGCACTGCATTCTTCTAGGAATTGCTATCTATACAGCGACCTCATTTA
CATACAAGGCACCTTTTTCTACATAGACTTGCCATTTGGCTGATTGTTCTGACAGTCGCT
GCCTCTTCGGAGTTCTGTGTGGACGAGTGGCCTCCTGAGTGGGCCGGTCCTTACGGAGTG
CCCAAGTCTGTTATGACTGGGTTCTCCATACAGCGACCTCACTCAGAATTGTATGGCACC
GTGCCGATTGAATGGACAAGCAGTCCTTAGGGAATCGCATGAAAGCATGACGGCAACATG
GTGTATGGACCCGTTGGGCCTTGCCTGTGCTATCTTCTCGAGTCGACCGGTCCTCGTCTC
CTCCTTTCGCGACGGGAAGCTCTTGAGTTTACCGTGGGTCAGGACGCTGTGGTATCTGTT
TGTCTGTAGTGTGGGCTCTCTATCAGAACAGCGTATCAGCACACGTCGTCTGGGTCTATG
GTATTTTTGCTTAACGGCCCTGGGTTTGGTAGTATGGTTCCATGTCGGTCCTCTCACCTC
CCAACCGTCATGTCTCTATCATCTGCACAGTACGTCTTGTCAGAATATCATTCTGTATGG
CAGTCGATAGACTAGGCCCTATGCCGCAGCTGTCTCTGACCACTATGCGAGCGCATACCT
ACCTCAATGTGTTCATAGCTGTTATGGATCTTGGGTGTGGTGTGCCAGAACGATACGCCA
CCAGATTGTGGAATGGTGCTTGTAGTATTAGAAGAGGACATGTATGAAGAGCACGTTCTT
TTCTCTTATGATGGTGCCGCGGGTGCGGGCCGTTCTCATTCTGCTTGTCGGGTGCTCAAG
GTGCATGTAGGATTGCTGAACCCACTGAATGGTCATGAGAATGGTCATATGGTGAGGTTC
ACTTCTCCATCGCTTTGTGTGTGTTGGATGTTTGGGTGGCTTCGGCCGCATGGAAAATCT
CGGTCTGTAACTCATCAGGGAGGGTTTCCATGTGAGCAATCTCTCCAGTCGAACACACAC
CCATGGGCGTCCCGCTAGGCAAAATGGAATTCTCGATCCGACACAGCCTGTGGAGTGTTT
GTTCCGTGTGGCGTGAAACGGCTTATACTGGATGTCTTCCCTTTGGACACCAGTCTGTTG
GTATGTACGCATACCATCTAGAGGGAGCAAGAAAAAGGCATGCAAGTGCCACAAAGGAAG
GGAGGCCTTCCTGCTACCGTGTGTCCTCTTGACGATCACGGGGGAGTATATAGAGACATT
TGGACGAGATCCTTTTTTCACGAGGCGGTGCCCTCGCGCAAAGGTGTTGCTCTGCACGCA
TCTTTATGATGATAGTTGCTCGCTCCTATGCGTCCACTCGTGAAAATACCTGCTCTCTCT
GTCTGTTTCCATGACACCCCCATGCGTTGTCATAGAGCATGCTCAGGTACAGAAAGTGCT
GTGACTTTCTTGGCCTGCTGTCACTGAGTCAGCCTGATACTTTGGAGGAATCTTTTCTGG
GCACTGCCCAGCTGAGAGTTCTTATCAGGCTCTGGAGTGTGCATATGCCTATATTCATTA
TAGTGCATGTGGCTACGCAGAAAAAGCATCGACGTGGAATGGAGAAGAACACCATTCTGA
ATGGAACGGTTCTTACTCTACTTCTGTTGCTTGACGCTGTATGGGAATCGCTTTGCTCTG
GGGGTACACCTTCCATTCCATGACAAAGCATTTGATCTATGCCCCGAGGGTCATTCTCTA
TGAAGAGCCCGAAACTTGTCTGAGTGAGAGGCGAAGGTAGTCCTTCATTTATGCGGGGTC
CGCCCTCACACTGTAGTCGGCACAGTCCCTTGTTTATGGACTGCGCAGCTCAATCTTCGA
CTGGAGAGGAAAAACAAAACACAAGTATGACACGCACATACGTGTTTTCTAGGACTGTGA
CCTGTCGGCTGCCTCGCGGTGGTTGGCATGATGCCCCAAGTGGGGGGGGGAGAAACAATC
AGTATGATTGATAAAAACCAGTGGGGAAAAAGGCGCGCTGTCGCACACCCTTGGTCACGG
CCTCAAGTCGACGGCCCTCACTGGCCCACCCTGCGACTCCTGTTCGCTTTTGACAAGACA
CACCTGCAATACGGAGGAAATGCCTGAGCAGCTGTGCCGCAGCAACTGCGGACGCGTAGC

GATATGCATCCCTGTTGTTTGCGACTTGGTACAAAGAACGTGCGCCGGGAGACGGATTGC
CCTTTGACTTTTCCACCCAGCCCGCGGCTCTATACCCTTGACGTACCTATTTACCTGGTT
GATCCTGCCAGTAGTCATATGCTTGTCTCAAAGACTAAGCCATGCATGTCTAAGTATAAA
CACTTTTATACTCGTGAAACTGCGAATGGCTCATTAAAACAGTTATAGTTCCTTTGGGAA
CAACACTACTTGGATAACCGTAGTAATTCTAGAGCTAATACATGCCTCAACGCCCGACTC
ACGAAGGGTGGTATTTATTGGATAAAAAACCATCGTGCTCTGTCCAGAGCGCTTTGAGAT
GATTCACAATAACTTGTCGGATCGCATGGCCTCGTGCTGGCGACTCCTCATTCAAATTTC
TGCCCTATCAACTTTCGATGGTAGAGTATTGGTCTACCATGGTGTCGACGGGTGACGGGG
AATTAGGGTTCGATTCCGGAGAGGGAGCCTGAGAAACGGCTACCACATCCAAGGAAGGCA
GCAGGCGCGCAAATTACCCAATCCCGACTCGGGGAGGTAGTGACAAAAAATAACAATAGG
GGGCCCTTTGGGTCTTCTAATTGGAATGAGAACAATTTAAATCCCTTATCGAGGATCCAT
TGGAGGGCAAGTCTGGTGAACGTATGCATTAAAGCCCTCCGTAATAAAAAAAAAACAGCG
CCAGATGTCTCTCTCTGGAAGCACACCATGCATGGGGTTTGTTCCTCTCTGCATGGCCGT
GTGAAAAAGGCGAGTAGGCAAACGATGGGAAACCCGGCTTGGAATAGTAACCAGCCACAG
AGCCCCCATCATACACTTCTAAATTGCCGGGGAGTGCCCGCGAGGCCTTGACTACCGCGG
TCCGCCGAGAGGTGAGAGCGAGCAATCGTTCAGGACGCAGCAGTGTACGTAGTGGTACAT
GGATGGTAAAAACGTCAAGGATAGGGTTGATCCGCAGGGAAGCGCGAGTGCCGCCATGGT
CGGTGGGCAAAGGCGAACCTTCAGAGACTCTAATGGAGTGGGCGCAAGCTTAAGGGAGAG
TCCAATCCACTGGGAAACCAGGTCCACAGCAGTAACGGGGTGATGCTCGCCCCAGAGGAC
TGTGGAAGGCAACCCCACCAAAAGGGTGTGTGTTGTCGGGTAGCAATGGCCAGCAGCCGC
GGTAATTCCAGCTCCAATAGCGTATATTAAAGTTGTTGCAGTTAAAACGCTCGTAGTCGG
ACCTCGGAGCGCCCCCACGGGCCGGGCTTTTGCCTTGTTGCCAGTGGGAGTTTGTCTCCT
TTTGTCGGGACGCGGTGTGCAGGGCCTTTACTGTCTCTGCGCGTTTACCTGCGGGCCGAC
CGTTTACTGTGAAGAAAGTAAAGTGTTCAAAGCAGGCCAATGCCTTGAATATGTGAGCAT
GGAATAATAGAATAGGACTTGGGCTCTATTTTGTTGGTTTCCAGTGACCAAGTAATGATT
AATAGGGACGGTTGGGGGCATTCGTATTTCATTGTCAGAGGTGAAATTCTTGGATTGATG
GAAGACGTACAACTGCGAAAGCATCTGCCATGGATGTTTTCATTGATCAAGAACGAAAGT
TAGGGGATCGAAGACGATCAGATACCGTCGTAGTCTTAACCATAAACGATGCCGACTGGG
GATTGGCGGGGGCACTTTTATATGACTCTGTCAGCACCCTGAGGGAAACCAAAGTCTTTG
GGTTCTGGGGGGAGTATGGTCGCAAGGCTGAAACTTAAAGGAATTGACGGAAGGGCACCA
CCAGGAGTGGAGCCTGCGGCTTAATTTGACTCAACACGGGAAAACTTACCAGGTCCGGAC
AGAAGAATGATTGACAGACTGAAAGCTCTTTCTTGATTTTTTGGTTGGTGGTGCATGGCC
GTTCTTAGTTGGTGGAGTGATTTGTCTGGTTAATTCCGTTAACGAACGAGACCTCGTCCT
GCTAAATAGGTGCGCGCATGCCAGCACTGCGTTCTTACCTTCTTAGAGGGACTATGCGCG
TCTAGCGTATGGAAGATTGAGGCAATAACAGGTCTGTGATGCCCTTAGATGTTCTGGGCC
GCACGCGCGCTACACTGATGCATTCAACGAGTTTCACTTGATAGTCCTGGGTCGGAAGGC

TCGGGTAATCTTTTGAAAGTGCATCGTGCTGGGGATAGATCATTGCAATTATTGATCTTC
AACGAGGAATTCCTTGTAGGCGCAGGTCATCAGCCTGCGCCGAATACGTCCCTGCCCTTT
GTACACACCGCCCGTCGCTCCTACCGATTGAATGGTCCGATGAAATGTCCGGATCGCGGC
TAGTGTGATGGACTCGTCCAGAGCGGTAGCCGTGAGAAGCTCATTAAATCTTATCATTTA
GAGGAAGGAGAAGTCGTAACAAGGTTTCCGTATTTTGTGGATGGGCACTCGTCGTGGCGT
TCTTTGCGCCACGCGGATGTCCCCCTTCCGTAGGAAAACGTACACGAAGCCTTTGCGGCC
CGAAAGGGTGCACAGTGCGCGACTATAAACACTGATTCTGTGGTCAACGCAAGCAGCGGT
TCTGTGACAGAACAGCTGCGACTTTCTCAAATTGCGGGAACACCCTAAAGTCAACAAACA
CCGCGGCCGGACGTGAAAGCGTCAGGCGCAGCAGTCGGGGTAGCGCCCTGGCGGATGGTA
AGAGCTTTGTTGAATGGCACAATGGGCAATCCGCAGCCAAGCCCCTAAGGTCCTTCGCGG
CTATGGGGAAGGTTCACAGACTGTAAGGGAGGGGGTTTCTTCCATGAAGAAGAAGCTTAA
GAGACAGTCGGTCCCCTGCGAGAGCAGTGTCTCATGTGAGGAAGGTGCCCATGAAAAGTG
GGCGCTGGAGAGCCATGGGAGCCCTCTATGAGGGTGGGGAGTTTTCCTAGATAATAATGA
GAAACAAACACGTTCCGTAGGTGAACCTGCGGAAGGATCATTCACAAACTATTGACACAA
ACACACGCGAACCAAATCGTCCGCACAGCAGGTGGCAATGAAAGAGAGACTCTGCATGTC
GCCTTTCGGGGTAGCAAGCAGCACTCTTCTGCCATCGCCTCTGTACAGGGCGTAAATTCC
CATTGAGAGGATGTGAGGGCACCACAGGAAGCTTTTCCACAGGAAGTCGCCATCCTTCTC
CCTCCACGGCGGCGCTTCTGTGCTTGGCAGTTTTTTTTGCCTTCTAGGGAGGATGCCGCC
AATGGAGCCCCTTATATATATACATCATCATATGCCCCTTTTTTTCTTAACCGCTTGCCA
CAGCTTCTTCTATGAGGAGCTTGTGGGAAGACTGTCTCCATACAATAACAAAGATACAAC
TCTTAGCGGTGGATATCTTGGTTCTCGCAACGATGAAGAACGCAGCCAACTGCGATAACT
AATGTGAATTGCAGGACTTCGTGAATCATTAAGTCTTTGAACGCAAGTTGCGCTCATCTC
CATGAGGAGGTGAGCATGCCTGTTTGAGTGTCGATGGGAAAAACAATACATAACAACAAT
CGGCTCTCTTATCTTCCTGCCTTCTTGGGGGCAAAAGGTAGTGCATGCCGGAGATGGGCT
TTGTCGCCCCTTTCTCTTGAAAGAGGCGTCGTCCGAAACACGTATCGTTGCCCGCTGGAG
GTCATAGAGCATGAGTTTAAAGCACCACATCTCTCAAGTCAGAGTGAGCGTCTTGAGTAG
GTCCGCATTGATTCAGTTTCAACCCAGGTGGCGTTGGTTACTTGCCATAAAGCCTCTCTG
GGGTTTCTTTGCTTGGACCTCTCTTGTCAGTTTCTCTGACGTAGAGACAGGTGCCGTCTC
ATGGACGTTGCCCTCTGGAACGTGCTAAAAAAAAAAAAAGAAAAAACCGTACCACATGTC
AAAAGAGGCAGATCGTGCTTGACGGCGGGAGGTGTTGCATTTCTTGCTTGTCCTGAGCGT
CTGCCAGTCTCTGACGAGTGTGTTCCTTCCATAAACCCGATCTCAAATCAGGTAGGACCA
CCCGCTGAATTTAAGCATATAATTAAGCGGAGGAAAAGAAACTAACAAGGATTCCCTCAG
TAGCGGCGAGCGAAGCGGGAAGAGCTCAACATGAAAACCGTTGGGGGTTCGCGCTTTCCT
TCGGGTTGTATAGTAAAGTGGTGTCGCCTGCGTCGCGCCGGGTATAAATCCCGTCTGAAA
TGGGGTGTCGTAGAGGGTTATATCCCCGTCTGTGACCCGGTGTGCGATGCGTTACGGTGA
CCTTACGATGAGTCGGGTTGCTTGGGCATGCAGCCCTAAGTGGGTGGTAAACTTCATCTA

AAGCTAAATACTAGCGAGAGACCGATAGCGTACAAGTACCGTAAGGGAAAGATGCAAAGA
ACTTTGAAAAGAGAGTTAAACAGGGCTTGAAACCGTTAAGAGTGAAGCGAAGGAGGTCAG
TGATCATGGGGCCAGATCCCCAAGTGCCTGTCATAGGGCACCGTAGCGTTCTGGTTTCCC
GGGGCAGCGCCGGCCGGGGGCGGGGGTTAACAGGGAGGTGGTTGAGGGATTCGTTTTCCG
CCTGTACTCCTTTATAGCTTGCACCCCGGCCGAGGGATCCGCGACTTGTCGTGTGAGGGC
GCTGTCAAAATGGCCTTCTCCACCCCGTCTTGAAACACGGACCAAGGAGTCTAACATACT
CGCGAGTGTTTGGGTGGTTAAAACCCGAACGCACAATGAAAGTAAACGGTGGCTCTGCGC
CGCTGCTTGGTGGGAACCCTGCTTTTATAATGGGGGGCACTATCACCCGACCCCGATCCC
TTTGGGGTGGTTGGTTTTGAGGTAGAGCGAGTATGATGGGACCCGAAAGATGGTGAACTA
TTCCTGAGTAGGGCGAAGCCAGAGGAAACTCTGGTGGAGGCTCGTAGCGATGCTGACGTG
CAAATCGCTCGTCGAACTTGGGTATAGGGGCGAAAGACTAATCGAACCATCTAGTAGCTG
GTTCCCTCCGAAGTTTCCCTCAGGATAGCGGGAGCAAATGCAAGTCAGTATCATCCGGTA
AAGCGAATGATTAGAGGCCTCGGGGGAGTAACTCCTTCGACCTATTCTCAAACTTTAAAT
AGGTGAGGAGCCGGCAACGTTGCTTAAAACAATGAACGCCGTGCGGACAATGATTGCTCT
CAGTGGGCCGATTTTGGTAAGCAGAACTGGCGATGCGGGATGAACCGAAAGCGACGCTAA
GGTGCCAAACTCCCAGCTCATTCAGACACCAGAAAAGGTGTTGATTCATACAGACAGCAG
GACGGTGGCCATGGAAGTCGGAATCCGCTAAGGAGTGTGTAACAACTCACCTGCCGAATG
GATCAGCCCTGAAAATGGATGGCGCTTAAGCGGGGGACCGAGGCGTCGCCGTTGGGGTAA
ATACGAAGCCCCAATGAGTAGGAGGGCGTGGGGGTTTTGAAACACGGTACACTCGCATGG
AAGCCTTGGGCGTAAGCTCGGGTGGAGCGACCCCTAGTGTAGATCTTGGTGGTAGTAGCA
AATATTCAAGTGAGAACCTTGAAGACTGAAGTGGGGAAGGGTTCCACGCGAACAGTGCTT
GGACGTGGGTTAGTCGGTCCTAAGAGATGGGGAAACTCCGTATTCAAGTGCGCTTTTGTA
GCGCAGTCCATCGAAAGGGAAGCGGGTTAACATTCCCGCACCAGGATGTGGAAACTGTGG
TTTACCGCAAACTGCGGCAACGCAACTGAACCTGGGGACGCCCGTAGGAGCTCCGAGCAG
GGATCTCTTTCCTGTTTAACGCGCCCAACCCCCTGGAATCGGTTTGTCCGGAGATAGGGG
GCGAAAGGCCGGAAAAGCACCGCACGTCTTGTGGTGTTCGGAACGCTTGTAATGGCCCTT
GAAAACCTAGGGGAAAGCAGCCGTCTTACACTTGTGTGAGACGAGGAATTCTCACACCTG
GCCGTACCCATCACCGCATCAGGTCCCCAAGGTGAACAGCCTCTAGTCAATAGAATAATG
TAGATAAGGGAAGTCGGCAAAATAGATCCGTAACTTTGGGAAAAGGATTGGCTCTAATGT
TCGGGCTTGTCGGGACGCGCGGGCGTAGATGCCAATGAGCGGCAGACGTCTAGCCTCTTT
GGAGGCTGGATTAACGCTGCTTGTGGGAAGAAACCGTGTCGATGGAGGCAGGAACGCTTA
TCGATCTTGTCTCTTCCCGCTTGCGATCAACGAACGGATTAGAACTGGTACGGACAAGGG
GAATCCGACTGTTTAATTAAAACAAAGCATTGCGACGGCCGGAAACGGTGTTGGCGCAAT
GTGATTTCTGCCCAGTGCTCTGAATGTCAAAGTGAAGAAATTCAACCAAGCGCGGGTAAA
CGGCGGGAGTAACTATGACTCTCTTAAGGTAGCCAAATGCCTCGTCATCTAAACTAATTG
GATGAGATAGTCTGAGTGAAAAGCTTATGCAGGCTAGTACCTGTCTAATCGTCCACAGTG

CAATTCTATTCGTCAGCGACAGGCTCACTCACGTTAACACCACGAGAAGCAATGGAGCAG
CGCGGGTGGTGCTTGCACACCAACACCCATTGCTGCCCATCGTCGATGAAAACAACTCCC
GGACATCCGCGCCGCGATATGTTGATAAATCGACTCTCGTAGCACAGGTGGGCCGGGTTG
CAGCACCAGGTGTTGCCGCACAGGTGCGATGTGTCGGAATTGACCGGCTTAACACCGCAG
CGTACCTCGGCAACTAGCACGTGCACCATGGCATGTCTGCCCATAAAACTGGTGCGATAA
TAGCCAGACGGCTGGCGAAATACACCTGGTCCAGTCGGCTGGCGGCACTTGAAGATATTC
TTGTAACCACCAACTGACATGACACGGTCGGCCAACAAGTCGTTGACGAACGCATCTAGT
GTAGTCAACGAGGTGTACTGGTTGCGGTGGCGAGACATGGTGTAGAAACGTGTGTGTGTT
GACGAAGTGAAATGTATTGTGTAGATGGGTGCGACACTCCTTGTACAGGGAAACCCTCCG
CATGGAAAGGCACGTGGTACCACCCTCGCGGGGAAACTCGCGAGCGGGATCCCGGGTAAT
GACCGGAGCCGATGGTAAAAACCCACGTGACCGATGGGGTGACCCTGTGGCGAGACGCTA
AGTTGTGGGACATCACCCTCTCACAATACGCATCCGTCGCAGAGACTGAACGGGAGTGGG
CTCCGTCGACGGCCAGCCAGCCGTCTTCCAGGGGGGCTTAAGATACAGTCCGGGCCTCAG
GGATGGAAAAACCTGGGGAACTACCAGAATTAGTGACGCGCATGAATGGATTAACGAGAT
TCCCACTGTCCCTATCTACTATCTAGCGAAACCACAGCCAAGGGAACGGGCTTGGAAGAA
TCAGCGGGGAAAGAAGACCCTGTTGAGCTTGACTCTAGTCCGACTTTGTGAGATGAGTTG
GGAGGTGTAGTCTAAGAGGGAGGGGCAACTCGACCGTGAAATACCTCTACTCCCAAGCTT
GTTTTACTTACTCCGGCGAGCAGGGATGGGTGGTAACACCCTTTTTCTAGGATTAACACC
TCTTCGGGGGTGGACCTGCGCGGAGGACATTGTCAGGTGGGGAGTTTGGCTGGGGCGGCA
CATCTGTTAAAAGATAACGCAGGTGTCCTAAGATGAGCTCAGTGAGAACAGAAATCTCAC
GTAGAACAAAAGGGTAAAAGCTCATTTGATTTTGATTTTCAGTACGAATACAAACTGTGA
AAGCATGGCCTATCGATCCTTTTGACCTTCAAGGTTTGAAGCAAGAGGTGTCAGAAAAGT
TACCACAGGGATAACTGGCTTGTGGCAGCCAAGCGTTCATAGCGACGTTGCTTTTTGATC
CTTCGATGTCGGCTCTTCCTATCATTGCGAAGCAGAATTCGCCAAGTGTTGGATTGTTCA
CCCACTAATAGGGAACGTGAGCTGGGTAGCCACTTTATGCCCAACAGCTGGATAGCTGGG
CTTCTGTCTTAGGCCGGGACTTCTCACGACCCGTGCCTTGGTCACCCAAAAAAGGGCGAC
CGCAGAGCTCAGGATGGAGCTTCTGGCTTAACTAGTAGAGGACAGATAGAGATCGTCAAA
TTGCGGGAAACCCCTTAGAGCCTCGAGCTACCAAGTGCGTGTCGAAAGGCATAGACGCAG
CCAGCGATGGCTGTCGCATGGCTTGATTCCATCAAGATAGGTAAAAATGCTCGAGGATTG
GGCAATCCGCAGCCAAGCCTCTAAGGTCCCCCCGCGGGCTATGGGGAAGGTTCAACGACT
GTAAGGCGTACCGTGCAAGCGGAAGATACAGTCTAGCCCCGCTGGAAAGCAGCGCCAAAG
TAAGTAGGTTGCAGTGGGTCTGTCCTC
>SRR9587942
GCATGTTTTGGTATGTGTTTCGACAGTCATGCATGTCATAGCCGTAGGATGGGGAGTGAG
CTTCATAGCCCGACCCCTTCTCAGGAGCATGTTTTCATAGCATATTGAGCATACTGGGGT
GGTCTCCCATGTAGCTCGCTCTCTCGGTGCAGGACCTCCCTTTTGCGGGCCACTTCGATG

GCCTCCATTATGGAGAGCATGTTTTGGCATATGAAGCGTTGGTGGTGAGGCTCGTAGCCT
GTATATACATACCAACGCAGTCGTCATATGTCTCTCTGGCGGGGGTCGGTCGTATGACCT
TGTCCCTACCTGGAGCATGTATTTTGTATGATATACTGGACGGCAGCTGCGCTGTTCAGT
CGTCAGCCGTACCTCATGTGCGCGTTTCCAGGTCCGAACGTTTGACATGCAAGGGCATCA
TACCAGAAGCCTGTTTTCTCGTATGAGCATGTAGGTAGGCAGGCTCTTCAGTCGTCTACC
CATAAATGCACTTCGAAGGAACTTTTTGGTATTGAATGGCTTTGTGCCCAGACGTTCTTG
TGAGCTTGTTCTCGCTCTGCGATACGCTGGAAGCTGTGGTATTACCTACAGCAGACGCAT
TCCTAGGGCGGACGGCCTCTCTGCAGATGGGCACATCTGTAGAAAGAGGCTCCCCTTGAT
GACGCAATGCAATGGCTGGGTAAACGGAGTGCTATACAGAAGGGTCAAACCTCTGTGGAA
GTACACTCATACCAGCATTGTTTGTTGAGAGGGGTCGGCTCTACGAGTGAGGCCCTATAC
TGCCATCTCATTCAGATGCGTAGACGGTCTTTACTCAGACTGCCTTTCTGGCTGATTGTC
CTCGGTCCCTCGTGCGCACGCGCCGCTGGCTGAATGGGCTTGTCCTTATGGAGTGCCGCG
TTCTTCAAAGAATCCGTCTTCATACAGCGACCTCATTTACATATTGCCGGCTGAATGGGC
TTGTCCTTATGGAGCACTGCATTCTTCTAGGAATTGCTATCTATACAGCGACCTCATTTA
CATACAAGGCACCTTTTTCTACATAGACTTGCCATTTGGCTGATTGTTCTGACAGTCGCT
GCCTCTTCGGAGTTCTGTGTGGACGAGTGGCCTCCTGAGTGGGCCGGTCCTTACGGAGTG
CCCAAGTCTGTTATGACTGGGTTCTCCATACAGCGACCTCACTCAGAATTGTATGGCACC
GTGCCGATTGAATGGACAAGCAGTCCTTAGGGAATCGCATGAAAGCATGACGGCAACATG
GTGTATGGACCCGTTGGGCCTTGCCTGTGCTATCTTCTCGAGTCGACCGGTCCTCGTCTC
CTCCTTTCGCGACGGGAAGCTCTTGAGTTTACCGTGGGTCAGGACGCTGTGGTATCTGTT
TGTCTGTAGTGTGGGCTCTCTATCAGAACAGCGTATCAGCACACGTCGTCTGGGTCTATG
GTATTTTTGCTTAACGGCCCTGGGTTTGGTAGTATGGTTCCATGTCGGTCCTCTCACCTC
CCAACCGTCATGTCTCTGATATCATCTGCACAGTACGTCTTGTCAGAATATCATTCTGTA
TGGCAGTCGATAGACTAGGCCCTATGCCGCAGCTGTCTCTGACCACTATGCGAGCGCATA
CCTACCTCAATGTGTTCATAGCTGTTATGGATCTTGGGTGTGGTGTGCCAGAACGATACG
CCACCAGATTGTGGAATGGTGCTTGTAGTATTAGAAGAGGACATGTATGAAGAGCACGTT
CTTTTCTCTTATGATGGTGCCGCGGGTGCGGGCCGTTCTCATTCTGCTTGTCGGGTGCTC
AAGGTGCATGTAGGATTGCTGAACCCACTGAATGGTCATGAGAATGGTCATATGGTGAGG
TTCACTTCTCCATCGCTTTGTGTGTGTTGGATGTTTGGGTGGCTTCGGCCGCATGGAAAA
TCTCGGTCTGTAACTCATCAGGGAGGGTTTCCATGTGAGCAATCTCTCCAGTCGAACACA
CACCCATGGGCGTCCCGCTAGGCAAAATGGAACTCTCGATCCGACACAGCCTGTGGAGTG
TTTGTTCCGTGTGGCGTGAAACGGCTTATACTGGATGTCTTCCCTTTGGACACCAGTCTG
TTGGTATGTACGCATACCATCTAGAGGGAGCAAGAAAAAGGCATGCAAGTGCCACAAAGG
AAGGGAGGCCTTCCTGCTACCGTGTGTCCTCTTGACGATCACGGGGGAGTATATAGAGAC
ATTTGGACGAGATCCGTTTTTCACGAGGCGGTGCCCTCGCGCAAAGGTGTTGCTCTGCAC
GCATCTTTATGATGATAGTTGCTCGCTCCTATGCGTCCACTCGTGAAAATACCTGCTCTC

TCTGTCTGTTTCCATGACACCCCCATGCGTTGTCATAGAGCATGCTCAGGTACAGAAAGT
GCTGTGACTTTCTTGGCCTGCTGTCACTGAGTCAGCCTGATACTTTGGAGGAATCTTTTC
TGGGCACTGCCCAGCTGAGAGTTCTTATCAGGCTCTGGAGTGTGCATATGCCTATATTCA
TTATAGTGCATGTGGCTACGCAGAAAAAGCATCGACGTGGAATGGAGAAGAACACCATTC
TGAATGGAACGGTTCTTACTCTACTTCTGTTGCTTGACGCTGTATGGGAATCGCTTTGCT
CTGGGGGTACACCTTCCATTCCATGACAAAGCATTTGATCTATGCCCCGAGGGTCATTCT
CTATGAAGAGCCCGAAACTTGTCTGAGTGAGAGGCGAAGGTAGTCCTTCATTTATGCGGG
GTCCGCCCTCACACTGTAGTCGGCACAGTCCCTTGTTTATGGACTGCGCAGCTCAATCTT
CGACTGGAGAGGAAAAACAAAACACAAGTATGACACGCACATACGTGTTTTCTAGGACTG
TGACCTGTCGGCTGCCTCGCGGTGGTTGGCATGATGCCCCAAGTGGGGGGGGGGAGAAAC
AATCAGTATGATTGATAAAAACCAGTGGGGAAAAAGGCGCGCTGTCGCACACCCTTGGTC
ACGGCCTCAAGTCGACGGCCCTCACTGGCCCACCCTGCGACTCCTGTTCGCTTTTGACAA
GACACACCTGCAATACGGAGGAAATGCCTGAGCAGCTGTGCCGCAGCAACTGCGGACGCG
TAGCGATATGCATCCCTGTTGTTTGCGACTTGGTACAAAGAACGTGCGCCGGGAGACGGA
TTGCCCTTTGACTTTTCCACCCAGCCCGCGGCTCTATACCCTTGACGTACCTATTTACCT
GGTTGATCCTGCCAGTAGTCATATGCTTGTCTCAAAGACTAAGCCATGCATGTCTAAGTA
TAAACACTTTTATACTCGTGAAACTGCGAATGGCTCATTAAAACAGTTATAGTTCCTTTG
GGAACAACACTACTTGGATAACCGTAGTAATTCTAGAGCTAATACATGCCTCAACGCCCG
ACTCACGAAGGGTGGTATTTATTGGATAAAAAACCATCGTGCTCTGTCCAGAGCGCTTTG
AGATGATTCACAATAACTTGTCGGATCGCATGGCCTCGTGCTGGCGACTCCTCATTCAAA
TTTCTGCCCTATCAACTTTCGATGGTAGAGTATTGGTCTACCATGGTGTCGACGGGTGAC
GGGGAATTAGGGTTCGATTCCGGAGAGGGAGCCTGAGAAACGGCTACCACATCCAAGGAA
GGCAGCAGGCGCGCAAATTACCCAATCCCGACTCGGGGAGGTAGTGACAAAAAATAACAA
TAGGGGGCCCTTTGGGTCTTCTAATTGGAATGAGAACAATTTAAATCCCTTATCGAGGAT
CCATTGGAGGGCAAGTCTGGTGAACGTATGCATTAAAGCCCTCCGTAATAAAAAAAAACA
GCGCCAGATGTCTCTCTCTGGAAGCACACCATGCATGGGGTTTGTTCCTCTCTGCATGGC
CGTGTGAAAAAGGCGAGTAGGCAAACGATGGGAAACCCGGCTTGGAATAGTAACCAGCCA
CAGAGCCCCCATCATACACTTCTAAATTGCCGGGGAGTGCCCGCGAGGCCTTGACTACCG
CGGTCCGCCGAGAGGTGAGAGCGAGCAATCGTTCAGGACGCAGCAGTGTACGTAGTGGTA
CATGGATGGTAAAAACGTCAAGGATAGGGTTGATCCGCAGGGAAGCGCGAGTGCCGCCAT
GGTCGGTGGGCAAAGGCGAACCTTCAGAGACTCTAATGGAGTGGGCGCAAGCTTAAGGGA
GAGTCCAATCCACTGGGAAACCAGGTCCACAGCAGTAACGGGGTGATGCTCGCCCCAGAG
GACTGTGGAAGGCAACCCCACCAAAAGGGTGTGTGTTGTCGGGTAGCAATGGCCAGCAGC
CGCGGTAATTCCAGCTCCAATAGCGTATATTAAAGTTGTTGCAGTTAAAACGCTCGTAGT
CGGACCTCGGAGCGCCCCCACGGGCCGGGCTTTTGCCTTGTTGCCAGTGGGAGTTTGTCT
CCTTTTGTCGGGACGCGGTGTGCAGGGCCTTTACTGTCTCTGCGCGTTTACCTGCGGGCC

GACCGTTTACTGTGAAGAAAGTAAAGTGTTCAAAGCAGGCCAATGCCTTGAATATGTGAG
CATGGAATAATAGAATAGGACTTGGGCTCTATTTTGTTGGTTTCCAGTGACCAAGTAATG
ATTAATAGGGACGGTTGGGGGCATTCGTATTTCATTGTCAGAGGTGAAATTCTTGGATTG
ATGGAAGACGTACAACTGCGAAAGCATCTGCCATGGATGTTTTCATTGATCAAGAACGAA
AGTTAGGGGATCGAAGACGATCAGATACCGTCGTAGTCTTAACCATAAACGATGCCGACT
GGGGATTGGCGGGGGCACTTTTATATGACTCTGTCAGCACCCTGAGGGAAACCAAAGTCT
TTGGGTTCTGGGGGGAGTATGGTCGCAAGGCTGAAACTTAAAGGAATTGACGGAAGGGCA
CCACCAGGAGTGGAGCCTGCGGCTTAATTTGACTCAACACGGGAAAACTTACCAGGTCCG
GACAGAAGAATGATTGACAGACTGAAAGCTCTTTCTTGATTTTTTGGTTGGTGGTGCATG
GCCGTTCTTAGTTGGTGGAGTGATTTGTCTGGTTAATTCCGTTAACGAACGAGACCTCGT
CCTGCTAAATAGGTGCGCGCATGCCAGCACTGCGTTCTTACCTTCTTAGAGGGACTATGC
GCGTCTAGCGTATGGAAGATTGAGGCAATAACAGGTCTGTGATGCCCTTAGATGTTCTGG
GCCGCACGCGCGCTACACTGATGCATTCAACGAGTTTCACTTGATAGTCCTGGGTCGGAA
GGCTCGGGTAATCTTTTGAAAGTGCATCGTGCTGGGGATAGATCATTGCAATTATTGATC
TTCAACGAGGAATTCCTTGTAGGCGCAGGTCATCAGCCTGCGCCGAATACGTCCCTGCCC
TTTGTACACACCGCCCGTCGCTCCTACCGATTGAATGGTCCGATGAAATGTCCGGATCGC
GGCTAGTGTGATGGACTCGTCCAGAGCGGTAGCCGTGAGAAGCTCATTAAATCTTATCAT
TTAGAGGAAGGAGAAGTCGTAACAAGGTTTCCGTAGGTGAACCTGCGGAAGGATCATTCA
CAAACTATTGACACAAACACACGCGAACCAAATCGTCCGCACAGCAGGTGGCAATGAAAG
AGAGACTCTGCATGTCGCCTTTCGGGGTAGCAAGCAGCACTCTTCTGCCATCGCCTCTGT
GCAGGGCGTAAATTCCCATTGAGAGGATGTGAGGGCACCACAGGAAGCTTTTCCACAGGA
AGTCGCCATCCTTCTCCCTCCACGGCGGCGCTTCTGTGCTTGGCAGTTTTTTTGCCTTC
TAGGGAGGATGCCGCCAATGGAGCCCCTTATATATATACATCATCATATGCCCCTTTTTT
TCTTAACCGCTTGCCACAGCTTCTTCTATGAGGAGCTTGTGGGAAGACTGTCTCCATACA
ATAACAAAGATACAACTCTTAGCGGTGGATATCTTGGTTCTCGCAACGATGAAGAACGCA
GCCAACTGCGATAACTAATGTGAATTGCAGGACTTCGTGAATCATTAAGTCTTTGAACGC
AAGTTGCGCTCATCTCCATGAGGAGGTGAGCATGCCTGTTTGAGTGTCGATGGGAAAAAC
AATACATAACAACAATCGGCTCTCTTATCTTCCTGCCTTCTTGGGGGCAAAAGGTAGTGC
ATGCCGGAGATGGGCTTTGTCGCCCCTTTCTCTTGAAAGAGGCGTCGTCCGAAACACGTA
TCGTTGCCCGCTGGAGGTCATAGAGCATGAGTTTAAAGCACCACATCTCTCAAGTCAGAG
TGAGCGTCTTGAGTAGGTCCGCATTGATTCAGTTTCAACCCAGGTGGCGTTGGTTACTTG
CCATAAAGCCTCTCTGGGGTTTCTTTGCTTGGACCTCTCTTGTCAGTTTCTCTGACGTAG
AGACAGGTGCCGTCTCATGGACGTTGCCCTCTGGAACGTGCTAAAAAAAAAAAAGAAAAA
ACCGTACCACATGTCAAAAGAGGCAGATCGTGCTTGACGGCGGGAGGTGTTGCATTTCTT
GCTTGTCCTGAGCGTCTGCCAGTCTCTGACGAGTGTGTTCCTTCCATAAACCCGATCTCA
AATCAGGTAGGACCACCCGCTGAATTTAAGCATATAATTAAGCGGAGGAAAAGAAACTAA

CAAGGATTCCCTCAGTAGCGGCGAGCGAAGCGGGAAGAGCTCAACATGAAAACCGTTGGG
GGTTCGCGCTTTCCTTCGGGTTGTATAGTAAAGTGGTGTCGCCTGCGTCGCGCCGGGTAT
AAATCCCGTCTGAAATGGGGTGTCGTAGAGGGTTATATCCCCGTCTGTGACCCGGTGTGC
GATGCGTTACGGTGACCTTACGATGAGTCGGGTTGCTTGGGCATGCAGCCCTAAGTGGGT
GGTAAACTTCATCTAAAGCTAAATACTAGCGAGAGACCGATAGCGTACAAGTACCGTAAG
GGAAAGATGCAAAGAACTTTGAAAAGAGAGTTAAACAGGGCTTGAAACCGTTAAGAGTGA
AGCGAAGGAGGTCAGTGATCATGGGGCCAGATCCCCAAGTGCCTGTCATAGGGCACCGTA
GCGTTCTGGTTTCCCGGGGCAGCGCCGGCCGGGGGCGGGGGTTAACAGGGAGGTGGTTGA
GGGATTCGTTTTCCGCCTGTACTCCTTTATAGCTTGCACCCCGGCCGAGGGATCCGCGAC
TTGTCGTGTGAGGGCGCTGTCAAAATGGCCTTCTCCACCCCGTCTTGAAACACGGACCAA
GGAGTCTAACATACTCGCGAGTGTTTGGGTGGTTAAAACCCGAACGCACAATGAAAGTAA
ACGGTGGCTCTGCGCCGCTGCTTGGTGGGAACCCTGCTTTTATAATGGGGGGCACTATCA
CCCGACCCCGATCCCTTTGGGGTGGTTGGTTTTGAGGTAGAGCGAGTATGATGGGACCCG
AAAGATGGTGAACTATTCCTGAGTAGGGCGAAGCCAGAGGAAACTCTGGTGGAGGCTCGT
AGCGATGCTGACGTGCAAATCGCTCGTCGAACTTGGGTATAGGGGCGAAAGACTAATCGA
ACCATCTAGTAGCTGGTTCCCTCCGAAGTTTCCCTCAGGATAGCGGGAGCAAATGCAAGT
CAGTATCATCCGGTAAAGCGAATGATTAGAGGCCTCGGGGGAGTAACTCCTTCGACCTAT
TCTCAAACTTTAAATAGGTGAGGAGCCGGCAACGTTGCTTAAAACAATGAACGCCGTGCG
GACAATGATTGCTCTCAGTGGGCCGATTTTGGTAAGCAGAACTGGCGATGCGGGATGAAC
CGAAAGCGACGCTAAGGTGCCAAACTCCCAGCTCATTCAGACACCAGAAAAGGTGTTGAT
TCATACAGACAGCAGGACGGTGGCCATGGAAGTCGGAATCCGCTAAGGAGTGTGTAACAA
CTCACCTGCCGAATGGATCAGCCCTGAAAATGGATGGCGCTTAAGCGGGGGACCGAGGCG
TCGCCGTTGGGGTAAATACGAAGCCCCAATGAGTAGGAGGGCGTGGGGGTTTTGAAACAC
GGTACACTCGCATGGAAGCCTTGGGCGTAAGCTCGGGTGGAGCGACCCCTAGTGTAGATC
TTGGTGGTAGTAGCAAATATTCAAGTGAGAACCTTGAAGACTGAAGTGGGGAAGGGTTCC
ACGCGAACAGTGCTTGGACGTGGGTTAGTCGGTCCTAAGAGATGGGGAAACTCCGTATTC
AAGTGCGCTTTTGTAGCGCAGTCCATCGAAAGGGAAGCGGGTTAACATTCCCGCACCAGG
ATGTGGAAACTGTGGTTTACCGCAAACTGCGGCAACGCAACTGAACCTGGGGACGCCCGT
AGGAGCTCCGAGCAGGGATCTCTTTCCTGTTTAACGCGCCCAACCCCCTGGAATCGGTTT
GTCCGGAGATAGGGGGCGAAAGGCCGGAAAAGCACCGCACGTCTTGTGGTGTTCGGAACG
CTTGTAATGGCCCTTGAAAACCCAGGGGAAAGCAGCCGTCTTACACTTGTGTGAGACGAG
GAATTCTCACACCTGGCCGTACCCATCACCGCATCAGGTCCCCAAGGTGAACAGCCTCTA
GTCAATAGAATAATGTAGATAAGGGAAGTCGGCAAAATAGATCCGTAACTTTGGGAAAAG
GATTGGCTCTAATGTTCGGGCTTGTCGGGACGCGCGGGCGTAGATGCCAATGAGCGGCAG
ACGTCTAGCCTCTTTGGAGGCTGGATTAACGCTGCTTGTGGGAAGAAACCGTGTCGATGG
AGGCAGGAACGCTTATCGATCTTGTCTCTTCCCGCTTGCGATCAACGAACGGATTAGAAC

TGGTACGGACAAGGGGAATCCGACTGTTTAATTAAAACAAAGCATTGCGACGGCCGGAAA

CGGTGTTGGCGCAATGTGATTTCTGCCCAGTGCTCTGAATGTCAAAGTGAAGAAATTCAA

CCAAGCGCGGGTAAACGGCGGGAGTAACTATGACTCTTAAACAAACCATACTACCTACGG

GAGTCAAAAGTGGGGGCGGCGTCTGGTGTAGAAATACACGTACGACACGCTTTTTTTCCT

GCTAGTCGATCTGACCTTTGTTCTTTGGACTGTGGGCGGTCAGCGGCAACATAACCTGGT

ACGGGGAAGGCCTCGACGTTGAGTGCTGCATCTTGGAAGTATCTTGACACAATAAGAGAA

AAGAGATTGCGCTTAATAGGGGGTTGATCTCGTGGCGAGCCGACGCCGGTGTGAACTGGT

GGCAATGGCCGTCGTAGAGCACGCAAAGGTATGGGCTGACTTTTTCTCCTCAAAAGTTGG

CTTAAGGGACGTGCCATCCCACACGATAATAAAGTGTGCAGTCTGTAAGAGCTCCCACAG

GGCAAAGACAGACTGGGATGCAGACGGAGACGGCAAAAACAACAACATCCGTTCTGCTTC

TGGGCGCAAATGCTTAAGGTAGCCAAATGCCTCGTCATCTAAACTAATTGGATGAGATAG

TCTGAGTGAAAAGCTTATGCAGGCTAGTACCGGAAAAAAAAAAAAGAGGGGCTCTGCCAC

TCTGACGGTGCGACACTCCTTGTACAGGGAAACCCTCCGCATGGAAAGGCACGTGGTACC

ACCCTCGCGGGGAAACTCGCGAGCGGGATCCCGGGTAATGACCGGAGCCGATGGTAAAAA

CTCACGTGACCGATGGGGTGACCCTGTGGCGAGACGCTAAGTTGTGGGACATCACCCTCT

CACAATACGCATCCGTCGCAGAGACTGAACGGGAGTGGGCTCCGTTGACGGCCGGCCGGC

CAGTTAGCCGTCTTCCAGGGGCTTAAGATACAGTCCGGGCCTCAGGGATGGAAAAACCTG

GGGAACTACCAGAATTAGTGACGCGCATGAATGGATTAACGAGATTCCCACTGTCCCTAT

CTACTATCTAGCGAAACCACAGCCAAGGGAACGGGCTTGGAAGAATCAGCGGGGAAAGAA

GACCCTGTTGAGCTTGACTCTAGTCCGACTTTGTGAGATGAGTTGGGAGGTGTAGTCTAA

GAGGGAGGGGCAACTCGACCGTGAAATACCTCTACTCCCAAGCTTGTTTTACTTACTCCG

GCGAGCAGGGATGGGTGGTAACACCCTTTTTCTAGGATTAACACCTCTTCGGGGGTGGAC

CTGCGCGGAGGACATTGTCAGGTGGGGAGTTTGGCTGGGGCGGCACATCTGTTAAAAGAT

AACGCAGGTGTCCTAAGATGAGCTCAGTGAGAACAGAAATCTCACGTAGAACAAAAGGGT

AAAAGCTCATTTGATTTTGATTTTCAGTACGAATACAAACTGTGAAAGCATGGCCTATCG

ATCCTTTTGACCTTCAAGGTTTGAAGCAAGAGGTGTCAGAAAAGTTACCACAGGGATAAC

TGGCTTGTGGCAGCCAAGCGTTCATAGCGACGTTGCTTTTTGATCCTTCGATGTCGGCTC

TTCCTATCATTGCGAAGCAGAATTCGCCAAGTGTTGGATTGTTCACCCACTAATAGGGAA

CGTGAGCTGGGTAGCCACTTTATGCCCAACAGCTGGATAGCTGGGCTTCTGTCTTAGGCC

GGGACTTCTCACGACCCGTGCCTTGGTCACCCAAAAAAGGGCGACCGCAGAGCTCAGGAT

GGAGCTTCTGGCTTAACTAGTAGAGGACAGATAGAGATCGTCAAATTGCGGGAAACCCCT

TAGAGCCTCGAGCTACCAAGTGCGTGTCGAAAGGCATAGACGCAGCCAGCGATGGCTGTC

GCATGGCTTGATTCCATCAAGATAGGTAAAAATGCTCGAGGATTGGGCAATCCGCAGCCA

AGCCTCTAAGGTCCCCCCGCGGGCTATGGGGAAGGTTCAACGACTGTAAGGCGTACCGTG

CAAGCGGAAGATACAGTCTAGCCCCGCTGGAAAGCAGCGCCAAAGTAAGTAGGTTGCAGT

GGGTCTGTCCTC

>SRR9587944

GCATGTTTTGGTATGTGTTTCGACAGTCATGCATGTCATAGCCGTAGGATGGGGAGTGAG
CTTCATAGCCCGACCCCTTCTCAGGAGCATGTTTTCATAGCATATTGAGCATACTGGGGT
GGTCTCCCATGTAGCTCGCTCTCTCGGTGCAGGACCTCCCTTTTGCGGGCCACTTCGATG
GCCTCCATTATGGAGAGCATGTTTTGGCATATGAAGCGTTGGTGGTGAGGCTCGTAGCCT
GTATATACATACCAACGCAGTCGTCATATGTCTCTCTGGCGGGGGTCGGTCGTATGACCT
TGTCCCTACCTGGAGCATGTATTTTGTATGATATACTGGACGGCAGCTGCGCTGTTCAGT
CGTCAGCCGTACCTCATGTGCGCGTTTCCAGGTCCGAACGTTTGACATGCAAGGGCATCA
TACCAGAAGCCTGTTTTCTCGTATGAGCATGTAGGTAGGCAGGCTCTTCAGTCGTCTACC
CATAAATGCACTTCGAAGGAACTTTTTGGTATTGAATGGCTTTGTGCCCAGACGTTCTTG
TGAGCTTGTTCTCGCTCTGCGATACGCTGGAAGCTGTGGTATTACCTACAGCAGACGCAT
TCCTAGGGCGGACGGCCTCTCTGCAGATGGGCACATCTGTAGAAAGAGGCTCCCCTTGAT
GACGCAATGCAATGGCTGGGTAAACGGAGTGCTATACAGAAGGGTCAAACCTCTGTGGAA
GTACACTCATACCAGCATTGTTTGTTGAGAGGGGTCGGCTCTACGAGTGAGGACCTATAC
TGCCATCTCATTCAGATGCGTAGACGGTCTTTACTCAGACTGCCTTTCTGGCTGATTGTC
CTCGGTCCCTCGTGCGCACGCGCCGCTGGCTGAATGGGCTTGTCCTTATGGAGTGCCGCG
TTCTTCAAAGAATCCGTCTTCATACAGCGACCTCATTTACATATTGCTGGCTGAATGGGC
TTGTCCTTATGGAGCACTGCATTCTTCTAGGAATTGCTATCTATACAGCGACCTCATTTA
CATACAAGGCACCTTTTTCTACATAGACTTGCCATTTGGCTGATTGTTCTGACAGTCGCT
GCCTCTTCGGAGTTCTGTGTGGACGAGTGGCCTCCTGAGTGGGCCGGTCCTTACGGAGTG
CCCACGTCTGTTATGACTGGGTTCTCCATACAGCGACCTCACTCAGAATTGTATGGCACC
GTGCCGATTGAATGGACAAGCAGTCCTTAGGGAATCGCATGAAAGCATGACGGCACATGG
TGTATGGACCCGTTGGGCCTTGCCTGTGCTATCTTCTCGAGTCGACCGGTCCTCGTCTCC
TCCTTTCGCGACGGGAAGCTCTTGAGTTTACCGTGGGTCAGGACGCTGTGGTATCTGTTT
GTCTGTAGTGTGGGCTCTCTATCAGAACAGCGTATCAGCACACGTCGTCTGGGTCTATGG
TATTTTTGCTTAACGGCCCTGGGTTTGGTAGTATGGTTCCATGTCGGTCCTCTCACCTCC
CAACCGTCATGTCTCTGATATCATCTGCACAGTACGTCTTGTCAGAATATCATTCTGTAT
GGCAGTCGATAGACTAGGCCCTATGCCGCAGCTGTCTCTGACCACTATGCGAGCGCATAC
CTACCTCAATGTGTTCATAGCTGTTATGGATCTTGGGTGTGGTGTGCCAGAACGATACGC
CACCAGATTGTGGAATGGTGCTTGTAGTATTAGAAGAGGACATGTATGAAGAGCACGTTC
TTTTCTCTTATGATGGTGCCGCGGGTGCGGGCCGTTCTCATTCTGCTTGTCGGGTGCTCA
AGGTGCATGTAGGATTGCTGAACCCACTGAATGGTCATGAGAATGGTCATATGGTGAGGT
TCACTTCTCCATCGCTTTGTGTGTGTTGGATGTTTGGGTGGCTTCGGCCGCATGGAAAAT
CTCGGTCTGTAACTCATCAGGGAGGGTTTCCATGTGAGCAATCTCTCCAGTCGAACACAC
ACCCATGGGCGTCCCGCTAGGCAAAATGGAACTCTCGATCCGACACAGCCTGTGGAGTGT
TTGTTCCGTGTGGCGTGAAACGGCTTATACTGGATGTCTTCCCTTTGGACACCAGTCTGT

TGGTATGTACGCATACCATCTAGAGGGAGCAAGAAAAAGGCATGCAAGTGCCACAAAGGA
AGGGAGGCCTTCCTGCTACCGTGTGTCCTCTTGACGATCACGGGGGAGTATATAGAGACA
TTTGGACGAGATCCGTTTTTCACGAGGCGGTGCCCTCGCGCAAAGGTGTTGCTCTGCACG
CATCTTTATGATGATAGTTGCTCGCTCCTATGCGTCCACTCGTGAAAATACCTGCTCTCT
CTGTCTGTTTCCATGACACCCCCATGCGTTGTCATAGAGCATGCTCAGGTACAGAAAGTG
CTGTGACTTTCTTGGCCTGCTGTCACTGAGTCAGCCTGATACTTTGGAGGAATCTTTTCT
GGGCACTGCCCAGCTGAGAGTTCTTATCAGGCTCTGGAGTGTGCATATGCCTATATTCAT
TATAGTGCATGTGGCTACGCAGAAAAAGCATCGACGTGGAATGGAGAAGAACACCATTCT
GAATGGAACGGTTCTTACTCTACTTCTGTTGCTTGACGCTGTATGGGAATCGCTTTGCTC
TGGGGGTACACCTTCCATTCCATGACAAAGCATTTGATCTATGCCCCGAGGGTCATTCTC
TATGAAGAGCCCGAAACTTGTCTGAGTGAGAGGCGAAGGTAGTCCTTCATTTATGCGGGG
TCCGCCCTCACACTGTAGTCGGCACAGTCCCTTGTTTATGGACTGCGCAGCTCAATCTTC
GACTGGAGAGGAAAAACAAAACACAAGTATGACACGCACATACGTGTTTTCTAGGACTGT
GACCTGTCGGCTGCCTCGCGGTGGTTGGCATGATGCCCCAAGTGGGGGGGGGAGAAACAA
TCAGTATGATTGATAAAAACCAGTGGGGAAAAAGGCGCGCTGTCGCACACCCTTGGTCAC
GGCCTCAAGTCGACGGCCCTCACTGGCCCACCCTGCGACTCCTGTTCGCTTTTGACAAGA
CACACCTGCAATACGGAGGAAATGCCTGAGCAGCTGTGCCGCAGCAACTGCGGACGCGTA
GCGATATGCATCCCTGTTGTTTGCGACTTGGTACAAAGAACGTGCGCCGGGAGACGGATT
GCCCTTTGACTTTTCCACCCAGCCCGCGGCTCTATACCCTTGACGTACCTATTTACCTGG
TTGATCCTGCCAGTAGTCATATGCTTGTCTCAAAGACTAAGCCATGCATGTCTAAGTATA
AACACTTTTATACTCGTGAAACTGCGAATGGCTCATTAAAACAGTTATAGTTCCTTTGGG
AACAACACTACTTGGATAACCGTAGTAATTCTAGAGCTAATACATGCCTCAACGCCCGAC
TCACGAAGGGTGGTATTTATTGGATAAAAAACCATCGTGCTCTGTCCAGAGCGCTTTGAG
ATGATTCACAATAACTTGTCGGATCGCATGGCCTCGTGCTGGCGACTCCTCATTCAAATT
TCTGCCCTATCAACTTTCGATGGTAGAGTATTGGTCTACCATGGTGTCGACGGGTGACGG
GGAATTAGGGTTCGATTCCGGAGAGGGAGCCTGAGAAACGGCTACCACATCCAAGGAAGG
CAGCAGGCGCGCAAATTACCCAATCCCGACTCGGGGAGGTAGTGACAAAAAATAACAATA
GGGGGCCCTTTGGGTCTTCTAATTGGAATGAGAACAATTTAAATCCCTTATCGAGGATCC
ATTGGAGGGCAAGTCTGGTGAACGTATGCATTAAAGCCCTCCGTAATAAAAAAAAACAGC
GCCAGATGTCTCTCTCTGGAAGCACACCATGCATGGGGTTTGTTCCTCTCTGCATGGCCG
TGTGAAAAAGGCGAGTAGGCAAACGATGGGAAACCCGGCTTGGAATAGTAACCAGCCACA
GAGCCCCCATCATACACTTCTAAATTGCCGGGGAGTGCCCGCGAGGCCTTGACTACCGCG
GTCCGCCGAGAGGTGAGAGCGAGCAATCGTTCAGGACGCAGCAGTGTACGTAGTGGTACA
TGGATGGTAAAAACGTCAAGGATAGGGTTGATCCGCAGGGAAGCGCGAGTGCCGCCATGG
TCGGTGGGCAAAGGCGAACCTTCAGAGACTCTAATGGAGTGGGCGCAAGCTTAAGGGAGA
GTCCAATCCACTGGGAAACCAGGTCCACAGCAGTAACGGGGTGATGCTCGCCCCAGAGGA

CTGTGGAAGGCAACCCCACCAAAAGGGTGTGTGTTGTCGGGTAGCAATGGCCAGCAGCCG

CGGTAATTCCAGCTCCAATAGCGTATATTAAAGTTGTTGCAGTTAAAACGCTCGTAGTCG

GACCTCGGAGCGCCCCCACGGGCCGGGCTTTTGCCTTGTTGCCAGTGGGAGTTTGTCTCC

TTTTGTCGGGACGCGGTGTGCAGGGCCTTTACTGTCTCTGCGCGTTTACCTGCGGGCCGA

CCGTTTACTGTGAAGAAAGTAAAGTGTTCAAAGCAGGCCAATGCCTTGAATATGTGAGCA

TGGAATAATAGAATAGGACTTGGGCTCTATTTTGTTGGTTTCCAGTGACCAAGTAATGAT

TAATAGGGACGGTTGGGGGCATTCGTATTTCATTGTCAGAGGTGAAATTCTTGGATTGAT

GGAAGACGTACAACTGCGAAAGCATCTGCCATGGATGTTTTCATTGATCAAGAACGAAAG

TTAGGGGATCGAAGACGATCAGATACCGTCGTAGTCTTAACCATAAACGATGCCGACTGG

GGATTGGCGGGGGCACTTTTATATGACTCTGTCAGCACCCTGAGGGAAACCAAAGTCTTT

GGGTTCTGGGGGGAGTATGGTCGCAAGGCTGAAACTTAAAGGAATTGACGGAAGGGCACC

ACCAGGAGTGGAGCCTGCGGCTTAATTTGACTCAACACGGGAAAACTTACCAGGTCCGGA

CAGAAGAATGATTGACAGACTGAAAGCTCTTTCTTGATTTTTTGGTTGGTGGTGCATGGC

CGTTCTTAGTTGGTGGAGTGATTTGTCTGGTTAATTCCGTTAACGAACGAGACCTCGTCC

TGCTAAATAGGTGCGCGCATGCCAGCACTGCGTTCTTACCTTCTTAGAGGGACTATGCGC

GTCTAGCGTATGGAAGATTGAGGCAATAACAGGTCTGTGATGCCCTTAGATGTTCTGGGC

CGCACGCGCGCTACACTGATGCATTCAACGAGTTTCACTTGATAGTCCTGGGTCGGAAGG

CTCGGGTAATCTTTTGAAAGTGCATCGTGCTGGGGATAGATCATTGCAATTATTGATCTT

CAACGAGGAATTCCTTGTAGGCGCAGGTCATCAGCCTGCGCCGAATACGTCCCTGCCCTT

TGTACACACCGCCCGTCGCTCCTACCGATTGAATGGTCCGATGAAATGTCCGGATCGCGG

CTAGTGTGATGGACTCGTCCAGAGCGGTAGCCGTGAGAAGCTCATTAAATCTTATCATTT

AGAGGAAGGAGAAGTCGTAACAAGGTTTCCGTATTTTGTGGGTGGATACCGATTCAATAA

CATCTGCCGTTTTTCTCAATGTACTATATCCCTAGTCTGCTTGCCCAGTTCTTGGAGTCT

CGTGCGATGTGTGCTATTACAGGCACGCTGGGTCATGAAGACAGACAAGGTTAGGATCTT

CACGGAAGTACGCGCAATAGGAGCGCGTCTTGTTCAACGCACCTGACTCAAGAGTCATAT

GCGAGGGACGGACGCAGCGCTTGTCGCCGCACAGATGGGAAGCTTCCTCTCCAGGCATAG

GATGGCGCTGCATTTGCATCATCGCTGCTAGTAAGTGGCAATAATACTTGATTCCCCGCC

AGCGCACTTGTACGTAGCCGTTGGCGGCGGGAACGTAGTTGGTGATGTGACAGCCTCGAG

CGTTCACCTCTGTGTGTCCGCCAATCTGCACAAAGATGAACGCCATATCGGAGACAGGGG

GAGGTGCCATTGTGATGATGACGATGTTGTTGGGTGGTCCTTCCGTAGGAAAACGTACAC

GAAGCCTTTGCGGCCCGAAAGGGTGCACAGTGCGCGACTATAAACACTGATTCTGTGGTC

AACGCAAGCAGCGGTTCTGTGGAAGAACAGCTGCGACTTTCTCAAATTGCGGGAACACCC

TAAAGTCAACAAGCACCGCGGCTTGACGTGAAAAGCGTCAGGCGCAGCAGTCGGGGTAGC

GCCCTGGCGGATGGTAATAGCCTTGTTGAATGGCACAATGGGCAATCCGCAGCCAAGCCC

CTAAGGTCCCCCGCTGGGCTATGGGGAAGGTTCACAGACTGTAAGGGAGGGGGTTTCTTC

CATGAAAGAAGCTTAAGAGACAGTCGGTCCCCTGCGAGAGCAGTGTCGCATGTGAGGAAG

GCACCCATGAAAAGTGGGTGCTGGAGAGCCATGTGAGCCCTCTATGAGGGCGGGAAGTTT
TCCGATACAATGAGAAACAAACACGTTCCGTAGGTGAACCTGCGGAAGGATCATTCACAA
ACTATTGACACAAACACACGCGAACCAAATCGTCCGCACAGCAGGTGGCAATGAAAGAGA
GACTCTGCATGTCGCCTTTCGGGGTAGCAAGCAGCACTCTTCTGCCATCGCCTCTGTGCA
GGGCGTAAATTCCCATTGAGAGGATGTGAGGGCACCACAGGAAGCTTTTCCACAGGAAGT
CGCCATCCTTCTCCCTCCACGGCGGCGCTTCTGTGCTTGGCAGTTTTTTTTGCCTTCTAG
GGAGGATGCCGCCAATGGAGCCCCTTATATATATACATCATCATATGCCCCTTTTTTTCT
TAACCGCTTGCCACAGCTTCTTCTATGAGGAGCTTGTGGGAAGACTGTCTCCATACAATA
ACAAAGATACAACTCTTAGCGGTGGATATCTTGGTTCTCGCAACGATGAAGAACGCAGCC
AACTGCGATAACTAATGTGAATTGCAGGACTTCGTGAATCATTAAGTCTTTGAACGCAAG
TTGCGCTCATCTCCATGAGGAGGTGAGCATGCCTGTTTGAGTGTCGATGGGAAAAACAAT
ACATAACAACAATCGGCTCTCTTATCTTCCTGCCTTCTTGGGGGCAAAAGGTAGTGCATG
CCGGAGATGGGCTTTGTCGCCCCTTTCTCTTGAAAGAGGCGTCGTCCGAAACACGTATCG
TTGCCCGCTGGAGGTCATAGAGCATGAGTTTAAAGCACCACATCTCTCAAGTCAGAGTGA
GCGTCTTGAGTAGGTCCGCATTGATTCAGTTTCAACCCAGGTGGCGTTGGTTACTTGCCA
TAAGCCTCTCTGGGGTTTCTTTGCTTGGACCTCTCTTGTCAGTTTCTCTGACGTAGAGAC
AGGTGCCGTCTCATGGACGTTGCCCTCTGGAACGTGCTAAAAAAAAAAAAAAAGAAAAAA
CCGTACCACATGTCAAAAGAGGCAGATCGTGCTTGACGGCGGGAGGTGTTGCATTTCTTG
CTTGTCCTGAGCGTCTGCCAGTCTCTGACGAGTGTGTTCCTTCCATAAACCCGATCTCAA
ATCAGGTAGGACCACCCGCTGAATTTAAGCATATAATTAAGCGGAGGAAAAGAAACTAAC
AAGGATTCCCTCAGTAGCGGCGAGCGAAGCGGGAAGAGCTCAACATGAAAACCGTTGGGG
GTTCGCGCTTTCCTTCGGGTTGTATAGTAAAGTGGTGTCGCCTGCGTCGCGCCGGGTATA
AATCCCGTCTGAAATGGGGTGTCGTAGAGGGTTATATCCCCGTCTGTGACCCGGTGTGCG
ATGCGTTACGGTGACCTTACGATGAGTCGGGTTGCTTGGGCATGCAGCCCTAAGTGGGTG
GTAAACTTCATCTAAAGCTAAATACTAGCGAGAGACCGATAGCGTACAAGTACCGTAAGG
GAAAGATGCAAAGAACTTTGAAAAGAGAGTTAAACAGGGCTTGAAACCGTTAAGAGTGAA
GCGAAGGAGGTCAGTGATCATGGGGCCAGATCCCCAAGTGCCTGTCATAGGGCACCGTAG
CGTTCTGGTTTCCCGGGGCAGCGCCGGCCGGGGGCGGGGGTTAACAGGGAGGTGGTTGAG
GGATTCGTTTTCCGCCTGTACTCCTTTATAGCTTGCACCCCGGCCGAGGGATCCGCGACT
TGTCGTGTGAGGGCGCTGTCAAAATGGCCTTCTCCACCCCGTCTTGAAACACGGACCAAG
GAGTCTAACATACTCGCGAGTGTTTGGGTGGTTAAAACCCGAACGCACAATGAAAGTAAA
CGGTGGCTCTGCGCCGCTGCTTGGTGGGAACCCTGCTTTTATAATGGGGGGCACTATCAC
CCGACCCCGATCCCTTTGGGGTGGTTGGTTTTGAGGTAGAGCGAGTATGATGGGACCCGA
AAGATGGTGAACTATTCCTGAGTAGGGCGAAGCCAGAGGAAACTCTGGTGGAGGCTCGTA
GCGATGCTGACGTGCAAATCGCTCGTCGAACTTGGGTATAGGGGCGAAAGACTAATCGAA
CCATCTAGTAGCTGGTTCCCTCCGAAGTTTCCCTCAGGATAGCGGGAGCAAATGCAAGTC

AGTATCATCCGGTAAAGCGAATGATTAGAGGCCTCGGGGGAGTAACTCCTTCGACCTATT
CTCAAACTTTAAATAGGTGAGGAGCCGGCAACGTTGCTTAAAACAATGAACGCCGTGCGG
ACAATGATTGCTCTCAGTGGGCCGATTTTGGTAAGCAGAACTGGCGATGCGGGATGAACC
GAAAGCGACGCTAAGGTGCCAAACTCCCAGCTCATTCAGACACCAGAAAAGGTGTTGATT
CATACAGACAGCAGGACGGTGGCCATGGAAGTCGGAATCCGCTAAGGAGTGTGTAACAAC
TCACCTGCCGAATGGATCAGCCCTGAAAATGGATGGCGCTTAAGCGGGGGACCGAGGCGT
CGCCGTTGGGGTAAATACGAAGCCCCAATGAGTAGGAGGGCGTGGGGGTTTTGAAACACG
GTACACTCGCATGGAAGCCTTGGGCGTAAGCTCGGGTGGAGCGACCCCTAGTGTAGATCT
TGGTGGTAGTAGCAAATATTCAAGTGAGAACCTTGAAGACTGAAGTGGGGAAGGGTTCCA
CGCGAACAGTGCTTGGACGTGGGTTAGTCGGTCCTAAGAGATGGGGAAACTCCGTATTCA
AGTGCGCTTTTGTAGCGCAGTCCATCGAAAGGGAAGCGGGTTAACATTCCCGCACCAGGA
TGTGGAAACTGTGGTTTACCGCAAACTGCGGCAACGCAACTGAACCTGGGGACGCCCGTA
GGAGCTCCGAGCAGGGATCTCTTTCCTGTTTAACGCGCCCAACCCCCTGGAATCGGTTTG
TCCGGAGATAGGGGGCGAAAGGCCGGAAAAGCACCGCACGTCTTGTGGTGTTCGGAACGC
TTGTAATGGCCCTTGAAAACCCAGGGGAAAGCAGCCGTCTTACACTTGTGTGAGACGAGG
AATTCTCACACCTGGCCGTACCCATCACCGCATCAGGTCCCCAAGGTGAACAGCCTCTAG
TCAATAGAATAATGTAGATAAGGGAAGTCGGCAAAATAGATCCGTAACTTTGGGAAAAGG
ATTGGCTCTAATGTTCGGGCTTGTCGGGACGCGCGGGCGTAGATGCCAATGAGCGGCAGA
CGTCTAGCCTCTTTGGAGGCTGGATTAACGCTGCTTGTGGGAAGAAACCGTGTCGATGGA
GGCAGGAACGCTTATCGATCTTGTCTCTTCCCGCTTGCGATCAACGAACGGATTAGAACT
GGTACGGACAAGGGGAATCCGACTGTTTAATTAAAACAAAGCATTGCGACGGCCGGAAAC
GGTGTTGGCGCAATGTGATTTCTGCCCAGTGCTCTGAATGTCAAAGTGAAGAAATTCAAC
CAAGCGCGGGTAAACGGCGGGAGTAACTATGACTCTTAAACAAACCATACTACCTACGGG
AGTCAAAAGTGGGGGCGGCGTCTGGTGTAGAAATACACGTACGACACGCTTTTTTTCCTG
CTAGTCGATCTGACCTTTGTTCTTTGGACTGTGGGCGGTCAGCGGCAACATAACCTGGTA
CGGGGAAGGCCTCGACGTTGAGTGCTGCATCTTGGAAGTATCTTGACACAATAAGAGAAA
AGAGATTGCGCTTAATAGGGGGTTGATCTCGTGGCGAGCCGACGCCGGTGTGAACTGGTG
GCAATGGCCGTCGTAGAGCACGCAAAGGTATGGGCTGACTTTTTCTCCTCAAAAGTTGGC
TTAAGGGACGTGCCATCCCACACGATAATAAAGTGTGCAGTCTGTAAGAGCTCCCACAGG
GCAAAGACAGACTGGGATGCAGACGGAGACGGCAAAAACAACAACATCCGTTCTGCTTCT
GGGCGCAAATGCTTAAGGTAGCCAAATGCCTCGTCATCTAAACTAATTGGATGAGATAGT
CTGAGTGAAAAGCTTATGCAGGCTAGTACCGGAAAAAAAAAAAAAGAGGGGCTCTGCCAC
TCTGACGGTGCGACACTCCTTGTACAGGGAAACCCTCCGCATGGAAAGGCACGTGGTACC
ACCCTCGCGGGGAAACTCGCGAGCGGGATCCCGGGTAATGACCGGAGCCGATGGTAAAAA
CTCACGTGACCGATGGGGTGACCCTGTGGCGAGACGCTAAGTTGTGGGACATCACCCTCT
CACAATACGCATCCGTCGCAGAGACTGAACGGGAGTGGGCTCCGTCGACGGCTAACCAGC

CTTCGTCGTGGGGCTTAAGATACAGTCCGGGCCTCAGGGATGGAAAAACCTGGGGAACTA
CCAGAATTAGTGACGCGCATGAATGGATTAACGAGATTCCCACTGTCCCTATCTACTATC
TAGCGAAACCACAGCCAAGGGAACGGGCTTGGAAGAATCAGCGGGGAAAGAAGACCCTGT
TGAGCTTGACTCTAGTCCGACTTTGTGAGATGAGTTGGGAGGTGTAGTCTAAGAGGGAGG
GGCAACTCGACCGTGAAATACCTCTACTCCCAAGCTTGTTTTACTTACTCCGGCGAGCAG
GGATGGGTGGTAACACCCTTTTTCTAGGATTAACACCTCTTCGGGGGTGGACCTGCGCGG
AGGACATTGTCAGGTGGGGAGTTTGGCTGGGGCGGCACATCTGTTAAAAGATAACGCAGG
TGTCCTAAGATGAGCTCAGTGAGAACAGAAATCTCACGTAGAACAAAAGGGTAAAAGCTC
ATTTGATTTTGATTTTCAGTACGAATACAAACTGTGAAAGCATGGCCTATCGATCCTTTT
GACCTTCAAGGTTTGAAGCAAGAGGTGTCAGAAAAGTTAACACGCGTTGTGGTAGCTTTT
AAGAGTGGACTCAAAGTGTTCCGCTAGTCCCCCCTCGGACCGCCACTTACAAAAGGCCTG
GGGATGGGGGCGACACTGTCAAATTGCGGGGACTCCCTAAAGCCTGTCTCTACCAAGCCA
GCGCGGCGAAAGCCCCTGGTGGCCGGGGTAATGACCTAGGGTATGGTAAAAGCGAGACAG
GATGGAACAATGGGCAATCCGCAGCCAAGTCCTAAGGCTCTTTTTGAAAAGCGGCTATGG
ATGCAGTTCAACGACTAGATGGCAGTGGGCTGAGAGAGCCCACCGCGGCACCCTTTCGGC
TTAAGATATAGTCTTGCCCACGTGAAAGCGTGCCTGCAGATGAACGCCACCATGTATGGC
CGGAGCTGCAGGAGACCGCACTTGATGTGCAGTCGGGAATCGACAGTACCACAGGGATGA
ACAAAAACACACAATTGTGGCCATCATACAGTCGTATGATATTATCGTCCCAGGCGGGTG
CGCCTCTCTCACCCGTCTAGTCCGGCCTTGGTGCGCAAAAAATGCTTGCGTGGATGCGCC
TTGGTTAAGGAAAATGCTCCTAGTAGTTTTACAAGGACACTTCTTGTAGGGCAACACATC
TGGATGCGAAGAACTCCCGGAAAAGACAACGGTACCAAGTGACTGGGGTAACGCCCCATT
CATGAAGGCAAATAATCAAAAAGAGGTGCCGGGTAAGAACCCGTTGTATAGGGACAATCC
GCAGCTGACTTGGCTCCCTGCCCATCCCCATGGGCATAGCTTAGCGGTTCAACGCTCGGT
AAGATGTGGGTTGGCGCGCTGTCCGACAGCGCACTGGCTTAAGGTACGGGCTACTCCCAC
CGGAGACGGTGTCTTTGTCTACCTCGGGTAACACGCCTCGGGAGAAGCAAAGAGGGTGCA
ATGGAAAAAAACTGGTTGCGGGGAAATCCCTACCAGAGTCACCTGGTACAAAGGAACTGG
CTTGTGGCAGCCAAGCGTTCATAGCGACGTTGCTTTTTGATCCTTCGATGTCGGCTCTTC
CTATCATTGCGAAGCAGAATTCGCCAAGTGTTGGATTGTTCACCCACTAATAGGGAACGT
GAGCTGGGTAGCCACTTTATGCCCAACAGCTGGATAGCTGGGCTTCTGTCTTAGGCCGGG
ACTTCTCACGACCCGTGCCTTGGTCACCCAAAAAAGGGCGACCGCAGAGCTCAGGATGGA
GCTTCTGGCTTAACTAGTAGAGGACAGATAGAGATCGTCAAATTGCGGGAAACCCCTTAG
AGCCTCGAGCTACCAAGTGCGTGTCGAAAGGCATAGACGCAGCCAGCGATGGCTGTCGCA
TGGCTTGATTCCATCAAGATAGGTAAAAATGCTCGAGGATTGGGCAATCCGCAGCCAAGC
CCCTAAGGTCCCCCCGCGGGCTATGGGGAAGGTTCAACGACTGTAAGGCGTACCGTGCAA
GCGGAAGATACAGTCTAGCCCCGCTGGAAAGCAGCGCCAAAGTAAGTAGGTTGCAGTGGG
TCTGTCCTC

>SRR9587949_SRR9587950_SRR9587952_SRR9587954_SRR9587960_SRR9587961_SRR9587964

GCATGTTTTGGTATGTGTTTCGACAGTCATGCATGTCATAGCCGTAGGATGGGGAGTGAG

CTTCATAGCCCGACCCCTTCTCAGGAGCATGTTTTCATAGCATATTGAGCATACTGGGGT

GGTCTCCCATGTAGCTCGCTCTCTCGGTGCAGGACCTCCCTTTTGCGGGCCACTTCGATG

GCCTCCATTATGGAGAGCATGTTTTGGCATATGAAGCGTTGGTGGTGAGGCTCGTAGCCT

GTATATACATACCAACGCAGTCGTCATATGTCTCTCTGGCGGGGGTCGGTCGTATGACCT

TGTCCCTACCTGGAGCATGTATTTTGTATGATATACTGGACGGCAGCTGCGCTGTTCAGT

CGTCGGCCGTACCTCATGTGCGCGTTTCCAGGTCCGAACGTTTGACATGCAAGGGCATCA

TACCAGAAGCCTGTTTTCTCGTATGAGCATGTAGGTAGGCAGGCTCTTCAGTCGTCTACC

CATAAATGCACTTCGAAGGAACTTTTTGGTATTGAATGGCTTTGTGCCCAGACGTTCTTG

TGAGCTTGTTCTCGCTCTGCGATACGCTGGAAGCTGTGGTATTACCTACAGCAGACGCAT

TCCTAGGGCGGACGGCCTCTCTGCAGATGGGCACATCTGTAGAAAGAGGCTCCCCTTGAT

GACGCAATGCAATGGCTGGGTAAACGGAGTGCTATACAGAAGGGTCAAACCTCTGTGGAA

GTACACTCATACCAGCATTGTTTGTTGAGAGGGGTCGGCTCTACGAGTGAGGCCCTATAC

TGCCATCTCATTCAGATGCGTAGACGGTCTTTACTCAGACTGCCTTTCTGGCTGATTGTC

CTCGGTCCCTCGTGCGCACGCGCCGCTGGCTGAATGGGCTTGTCCTTATGGAGTGCCGCG

TTCTTCAAAGAATCCGTCTTCATACAGCGACCTCATTTACATATTGCCGGCTGAATGGGC

TTGTCCTTATGGAGCACTGCATTCTTCTAGGAATTGCTATCTATACAGCGACCTCATTTA

CATACAAGGCACCTTTTTCTACATAGACTTGCCATTTGGCTGATTGTTCTGACAGTCGCT

GCCTCTTCGGAGTTCTGTGTGGACGAGTGGCCTCCTGAGTGGGCCGGTCCTTACGGAGTG

CCCAAGTCTGTTATGACTGGGTTCTCCATACAGCGACCTCACTCAGAATTGTATGGCACC

GTGCCGATTGAATGGACAAGCAGTCCTTAGGGAATCGCATGAAAGCATGACGGCAACATG

GTGTATGGACCCGTTGGGCCTTGCCTGTGCTATCTTCTCGAGTCGACCGGTCCTCGTCTC

CTCCTTTCGCGACGGGAAGCTCTTGAGTTTACCGTGGGTCAGGACGCTGTGGTATCTGTT

TGTCTGTAGTGTGGGCTCTCTATCAGAACAGCGTATCAGCACACGTCGTCTGGGTCTATG

GTATTTTTGCTTAACGGCCCTGGGTTTGGTAGTATGGTTCCATGTCGGTCCTCTCACCTC

CCAACCGTCATGTCTCTGATATCATCTGCACAGTACGTCTTGTCAGAATATCATTCTGTA

TGGCAGTCGATAGACTAGGCCCTATGCCGCAGCTGTCTCTGACCACTATGCGAGCGCATA

CCTACCTCAATGTGTTCATAGCTGTTATGGATCTTGGGTGTGGTGTGCCAGAACGATACG

CCACCAGATTGTGGAATGGTGCTTGTAGTATTAGAAGAGGACATGTATGAAGAGCACGTT

CTTTTCTCTTATGATGGTGCCGCGGGTGCGGGCCGTTCTCATTCTGCTTGTCGGGTGCTC

AAGGTGCATGTAGGATTGCTGAACCCACTGAATGGTCATGAGAATGGTCATATGGTGAGG

TTCACTTCTCCATCGCTTTGTGTGTGTTGGATGTTTGGGTGGCTTCGGCCGCATGGAAAA

TCTCGGTCTGTAACTCATCAGGGAGGGTTTCCATGTGAGCAATCTCTCCAGTCGAACACA

CACCCATGGGCGTCCCGCTAGGCAAAATGGAACTCTCGATCCGACACAGCCTGTGGAGTG

TTTGTTCCGTGTGGCGTGAAACGGCTTATACTGGATGTCTTCCCTTTGGACACCAGTCTG

TTGGTATGTACGCATACCATCTAGAGGGAGCAAGAAAAAGGCATGCAAGTGCCACAAAGG
AAGGGAGGCCTTCCTGCTACCGTGTGTCCTCTTGACGATCACGGGGGAGTATATAGAGAC
ATTTGGACGAGATCCGTTTTTCACGAGGCGGTGCCCTCGCGCAAAGGTGTTGCTCTGCAC
GCATCTTTATGATGATAGTTGCTCGCTCCTATGCGTCCACTCGTGAAAATACCTGCTCTC
TCTGTCTGTTTCCATGACACCCCCATGCGTTGTCATAGAGCATGCTCAGGTACAGAAAGT
GCTGTGACTTTCTTGGCCTGCTGTCACTGAGTCAGCCTGATACTTTGGAGGAATCTTTTC
TGGGCACTGCCCAGCTGAGAGTTCTTATCAGGCTCTGGAGTGTGCATATGCCTATATTCA
TTATAGTGCATGTGGCTACGCAGAAAAAGCATCGACGTGGAATGGAGAAGAACACCATTC
TGAATGGAACGGTTCTTACTCTACTTCTGTTGCTTGACGCTGTATGGGAATCGCTTTGCT
CTGGGGGTACACCTTCCATTCCATGACAAAGCATTTGATCTATGCCCCGAGGGTCATTCT
CTATGAAGAGCCCGAAACTTGTCTGAGTGAGAGGCGAAGGTAGTCCTTCATTTATGCGGG
GTCCGCCCTCACACTGTAGTCGGCACAGTCCCTTGTTTATGGACTGCGCAGCTCAATCTT
CGACTGGAGAGGAAAAACAAAACACAAGTATGACACGCACATACGTGTTTTCTAGGACTG
TGACCTGTCGGCTGCCTCGCGGTGGTTGGCATGATGCCCCAAGTGGGGGGGGGAGAAACA
ATCAGTATGATTGATAAAAACCAGTGGGGAAAAAGGCGCGCTGTCGCACACCCTTGGTCA
CGGCCTCAAGTCGACGGCCCTCACTGGCCCACCCTGCGACTCCTGTTCGCTTTTGACAAG
ACACACCTGCAATACGGAGGAAATGCCTGAGCAGCTGTGCCGCAGCAACTGCGGACGCGT
AGCGATATGCATCCCTGTTGTTTGCGACTTGGTACAAAGAACGTGCGCCGGGAGACGGAT
TGCCCTTTGACTTTTCCACCCAGCCCGCGGCTCTATACCCTTGACGTACCTATTTACCTG
GTTGATCCTGCCAGTAGTCATATGCTTGTCTCAAAGACTAAGCCATGCATGTCTAAGTAT
AAACACTTTTATACTCGTGAAACTGCGAATGGCTCATTAAAACAGTTATAGTTCCTTTGG
GAACAACACTACTTGGATAACCGTAGTAATTCTAGAGCTAATACATGCCTCAACGCCCGA
CTCACGAAGGGTGGTATTTATTGGATAAAAAACCATCGTGCTCTGTCCAGAGCGCTTTGA
GATGATTCACAATAACTTGTCGGATCGCATGGCCTCGTGCTGGCGACTCCTCATTCAAAT
TTCTGCCCTATCAACTTTCGATGGTAGAGTATTGGTCTACCATGGTGTCGACGGGTGACG
GGGAATTAGGGTTCGATTCCGGAGAGGGAGCCTGAGAAACGGCTACCACATCCAAGGAAG
GCAGCAGGCGCGCAAATTACCCAATCCCGACTCGGGGAGGTAGTGACAAAAAATAACAAT
AGGGGGCCCTTTGGGTCTTCTAATTGGAATGAGAACAATTTAAATCCCTTATCGAGGATC
CATTGGAGGGCAAGTCTGGTGAACGTATGCATTAAAGCCCTCCGTAATAAAAAAAAACAG
CGCCAGATGTCTCTCTCTGGAAGCACACCATGCATGGGGTTTGTTCCTCTCTGCATGGCC
GTGTGAAAAAGGCGAGTAGGCAAACGATGGGAAACCCGGCTTGGAATAGTAACCAGCCAC
AGAGCCCCCATCATACACTTCTAAATTGCCGGGGAGTGCCCGCGAGGCCTTGACTACCGC
GGTCCGCCGAGAGGTGAGAGCGAGCAATCGTTCAGGACGCAGCAGTGTACGTAGTGGTAC
ATGGATGGTAAAAACGTCAAGGATAGGGTTGATCCGCAGGGAAGCGCGAGTGCCGCCATG
GTCGGTGGGCAAAGGCGAACCTTCAGAGACTCTAATGGAGTGGGCGCAAGCTTAAGGGAG
AGTCCAATCCACTGGGAAACCAGGTCCACAGCAGTAACGGGGTGATGCTCGCCCCAGAGG

ACTGTGGAAGGCAACCCCACCAAAAGGGTGTGTGTTGTCGGGTAGCAATGGCCAGCAGCC
GCGGTAATTCCAGCTCCAATAGCGTATATTAAAGTTGTTGCAGTTAAAACGCTCGTAGTC
GGACCTCGGAGCGCCCCCACGGGCCGGGCTTTTGCCTTGTTGCCAGTGGGAGTTTGTCTC
CTTTTGTCGGGACGCGGTGTGCAGGGCCTTTACTGTCTCTGCGCGTTTACCTGCGGGCCG
ACCGTTTACTGTGAAGAAAGTAAAGTGTTCAAAGCAGGCCAATGCCTTGAATATGTGAGC
ATGGAATAATAGAATAGGACTTGGGCTCTATTTTGTTGGTTTCCAGTGACCAAGTAATGA
TTAATAGGGACGGTTGGGGGCATTCGTATTTCATTGTCAGAGGTGAAATTCTTGGATTGA
TGGAAGACGAACAACTGCGAAAGCATCTGCCATGGATGTTTTCATTGATCAAGAACGAAA
GTTAGGGGATCGAAGACGATCAGATACCGTCGTAGTCTTAACCATAAACGATGCCGACTG
GGGATTGGCGGGGGCACTTTTATATGACTCTGTCAGCACCCTGAGGGAAACCAAAGTCTT
TGGGTTCTGGGGGGAGTATGGTCGCAAGGCTGAAACTTAAAGGAATTGACGGAAGGGCAC
CACCAGGAGTGGAGCCTGCGGCTTAATTTGACTCAACACGGGAAAACTTACCAGGTCCGG
ACAGAAGAATGATTGACAGACTGAAAGCTCTTTCTTGATTTTTTGGTTGGTGGTGCATGG
CCGTTCTTAGTTGGTGGAGTGATTTGTCTGGTTAATTCCGTTAACGAACGAGACCTCGTC
CTGCTAAATAGGTGCGCGCATGCCAGCACTGCGTTCTTACCTTCTTAGAGGGACTATGCG
CGTCTAGCGTATGGAAGATTGAGGCAATAACAGGTCTGTGATGCCCTTAGATGTTCTGGG
CCGCACGCGCGCTACACTGATGCATTCAACGAGTTTCACTTGATAGTCCTGGGTCGGAAG
GCTCGGGTAATCTTTTGAAAGTGCATCGTGCTGGGGATAGATCATTGCAATTATTGATCT
TCAACGAGGAATTCCTTGTAGGCGCAGGTCATCAGCCTGCGCCGAATACGTCCCTGCCCT
TTGTACACACCGCCCGTCGCTCCTACCGATTGAATGGTCCGATGAAATGTCCGGATCGCG
GCTAGTGTGATGGACTCGTCCAGAGCGGTAGCCGTGAGAAGCTCATTAAATCTTATCATT
TAGAGGAAGGAGAAGTCGTAACAAGGTTTCCGTAGGTGAACCTGCGGAAGGATCATTCAC
AAACTATTGACACAAACACACGCGAACCAAATCGTCCGCACAGCAGGTGGCAATGAAAGA
GAGACTCTGCATGTCGCCTTTCGGGGTAGCAAGCAGCACTCTTCTGCCATCGCCTCTGTG
CAGGGCGTAAATTCCCATTGAGAGGATGTGAGGGCACCACAGGAAGCTTTTCCACAGGAA
GTCGCCATCCTTCTCCCTCCACGGCGGCGCTTCTGTGCTTGGCAGTTTTTTTTGCCTTCT
AGGGAGGATGCCGCCAATGGAGCCCCTTATATATATACATCATCATATGCCCCTTTTTTT
CTTAACCGCTTGCCACAGCTTCTTCTATGAGGAGCTTGTGGGAAGACTGTCTCCATACAA
TAACAAAGATACAACTCTTAGCGGTGGATATCTTGGTTCTCGCAACGATGAAGAACGCAG
CCAACTGCGATAACTAATGTGAATTGCAGGACTTCGTGAATCATTAAGTCTTTGAACGCA
AGTTGCGCTCATCTCCATGAGGAGGTGAGCATGCCTGTTTGAGTGTCGATGGGAAAAACA
ATACAGAACAACAATCGGCTCTCTTATCTTCCTGCCTTCTTGGGGGCAAAAGGTAGTGCA
TGCCGGAGATGGGCTTTGTCGCCCCTTTCTCTTGAAAGAGGCGTCGTCCGAAACACGTAT
CGTTGCCCGCTGGAGGTCATAGAGCATGAGTTTAAAGCACCACATCTCTCAAGTCAGAGT
GAGCGTCTTGAGTAGGTCCGCATTGATTCAGTTTCAACCCAGGTGGCGTTGGTTACTTGC
CATAAGCCTCTCTGGGGTTTCTTTGCTTGGACCTCTCTTGTCAGTTTCTCTGACGTAGAG

ACAGGTGCCGTCTCATGGACGTTGCCCTCTGGAACGTGCTAAAAAAAAAAAAAAAGAAAAA
ACCGTACCACATGTCAAAAGAGGCAGATCGTGCTTGACGGCGGGAGGTGTTGCATTTCTT
GCTTGTCCTGAGCGTCTGCCAGTCTCTGACGAGTGTGTTCCTTCCATAAACCCGATCTCA
AATCAGGTAGGACCACCCGCTGAATTTAAGCATATAATTAAGCGGAGGAAAAGAAACTAA
CAAGGATTCCCTCAGTAGCGGCGAGCGAAGCGGGAAGAGCTCAACATGAAAACCGTTGGG
GGTTCGCGCTTTCCTTCGGGTTGTATAGTAAAGTGGTGTCGCCTGCGTCGCGCCGGGTAT
AAATCCCGTCTGAAATGGGGTGTCGTAGAGGGTTATATCCCCGTCTGTGACCCGGTGTGC
GATGCGTTACGGTGACCTTACGATGAGTCGGGTTGCTTGGGCATGCAGCCCTAAGTGGGT
GGTAAACTTCATCTAAAGCTAAATACTAGCGAGAGACCGATAGCGTACAAGTACCGTAAG
GGAAAGATGCAAAGAACTTTGAAAAGAGAGTTAAACAGGGCTTGAAACCGTTAAGAGTGA
AGCGAAGGAGGTCAGTGATCATGGGGCCAGATCCCCAAGTGCCTGTCATAGGGCACTGTA
GCGTTCTGGTTTCCCGGGGCAGCGCCGGCCGGGGGCGGGGGTTAACAGGGAGGTGGTTGA
GGGATTCGTTTTCCGCCTGTACTCCTTTATAGCTTGCACCCCGGCCGAGGGATCCGCGAC
TTGTCGTGTGAGGGCGCTGTCAAAATGGCCTTCTCCACCCCGTCTTGAAACACGGACCAA
GGAGTCTAACATACTCGCGAGTGTTTGGGTGGTTAAAACCCGAACGCACAATGAAAGTAA
ACGGTGGCTCTGCGCCGCTGCTTGGTGGGAACCCTGCTTTTATAATGGGGGGCACTATCA
CCCGACCCCGATCCCTTTGGGGTGGTTGGTTTTGAGGTAGAGCGAGTATGATGGGACCCG
AAAGATGGTGAACTATTCCTGAGTAGGGCGAAGCCAGAGGAAACTCTGGTGGAGGCTCGT
AGCGATGCTGACGTGCAAATCGCTCGTCGAACTTGGGTATAGGGGCGAAAGACTAATCGA
ACCATCTAGTAGCTGGTTCCCTCCGAAGTTTCCCTCAGGATAGCGGGAGCAAATGCAAGT
CAGTATCATCCGGTAAAGCGAATGATTAGAGGCCTCGGGGGAGTAACTCCTTCGACCTAT
TCTCAAACTTTAAATAGGTGAGGAGCCGGCAACGTTGCTTAAAACAATGAACGCCGTGCG
GACAATGATTGCTCTCAGTGGGCCGATTTTGGTAAGCAGAACTGGCGATGCGGGATGAAC
CGAAAGCGACGCTAAGGTGCCAAACTCCCAGCTCATTCAGACACCAGAAAAGGTGTTGAT
TCATACAGACAGCAGGACGGTGGCCATGGAAGTCGGAATCCGCTAAGGAGTGTGTAACAA
CTCACCTGCCGAATGGATCAGCCCTGAAAATGGATGGCGCTTAAGCGGGGGACCGAGGCG
TCGCCGTTGGGGTAAATACGAAGCCCCAATGAGTAGGAGGGCGTGGGGGTTTTGAAACAC
GGTACACTCGCATGGAAGCCTTGGGCGTAAGCTCGGGTGGAGCGACCCCTAGTGTAGATC
TTGGTGGTAGTAGCAAATATTCAAGTGAGAACCTTGAAGACTGAAGTGGGGAAGGGTTCC
ACGCGAACAGTGCTTGGACGTGGGTTAGTCGGTCCTAAGAGATGGGGAAACTCCGTATTC
AAGTGCGCTTTTGTAGCGCAGTCCATCGAAAGGGAAGCGGGTTAACATTCCCGCACCAGG
ATGTGGAAACTGTGGTTTACCGCAAACTGCGGCAACGCAACTGAACCTGGGGACGCCCGT
AGGAGCTCCGAGCAGGGATCTCTTTCCTGTTTAACGCGCCCAACCCCCTGGAATCGGTTT
GTCCGGAGATAGGGGGCGAAAGGCCGGAAAAGCACCGCACGTCTTGTGGTGTTCGGAACG
CTTGTAATGGCCCTTGAAAACCCAGGGGAAAGCAGCCGTCTTACACTTGTGTGAGACGAG
GAATTCTCACACCTGGCCGTACCCATCACCGCATCAGGTCCCCAAGGTGAACAGCCTCTA

GTCAATAGAATAATGTAGATAAGGGAAGTCGGCAAAATAGATCCGTAACTTTGGGAAAAG

GATTGGCTCTAATGTTCGGGCTTGTCGGGACGCGCGGGCGTAGATGCCAATGAGCGGCAG

ACGTCTAGCCTCTTTGGAGGCTGGATTAACGCTGCTTGTGGGAAGAAACCGTGTCGATGG

AGGCAGGAACGCTTATCGATCTTGTCTCTTCCCGCTTGCGATCAACGAACGGATTAGAAC

TGGTACGGACAAGGGGAATCCGACTGTTTAATTAAAACAAAGCATTGCGACGGCCGGAAA

CGGTGTTGGCGCAATGTGATTTCTGCCCAGTGCTCTGAATGTCAAAGTGAAGAAATTCAA

CCAAGCGCGGGTAAACGGCGGGAGTAACTATGACTCTCTTAAGGTAGCCAAATGCCTCGT

CATCTAGACACTACTTGGATGAGATAGTCTGGGTTAAAAGCTTATGCAGGCTAGTACCCT

CTGGGTAAAGAACAAATATACGACTCCACAGGCGCATCACATTTCACACCAAGCGAAACA

ATCTTGCAATGCGGCGTGTGCTGGCACACGCGGACCCACTGCTGCTGCTCCTCGATGAAG

ACAGTTCCCGCACATCCGCGGCGGGAGATGTTTTTGTCGCGGCTTTCGAAGCATAAGTGG

TCCGGGTTGCAGCACCACGCGTTGCCGCAGAGGTGAGAGGTGTCATCGGCATCACACTCC

TTTGTGACGAGCGTCTCGTGGATGAACTTGTGCAGGGTGACCTTGCGTCCTTTGTAGTTC

GTGCGGAAGTAGCCGTTGGCGGCTTGGTCAACGCCAATTCCCATCGGCATTTTGCACTTG

TAGATGTTCTTGACGCCTCCGATCGAGTAGACGCGGTTCGCCATGAATTCGTCGACGAAC

GCCTCCGGCGTGACGTCCGGCGTGTATTGGTTGCTCTTCCGCGACATCGTGTTGGCGTGC

AGCGGGTGCGACACTCCTTGTACAGGGGAACCCTCCAAACAGAAAAAGGCGCGTGGTACC

ACCCCTGCGAGGAAACGCGCGGGCGGGATCCCGGGTAATGACCGGAGCCGAAGGTAAAAA

CCCACGCGACCGATGGGGCAACCCTGTGGCGAGAGGCTAAGTTGCGAGCGTCTTCGCGCG

CGATATGCCTCCGTCGCAGAGACTGAACGGGAGTGGGCTCGAGATGCTACCACTGGGGAC

ACGTGATGTTCCTTGTGCCAAGTACCGGGCTTAAGATACAGTCCAGGCCCCAGGTATGGA

AAACCTGGGGAACTACCCTGAATTAGTGACGCGCATGAATGGATTAACGAGATTCCCACT

GTCCCTATCTACTATCTAGCGAAACCACAGCCAAGGGAACGGGCTTGGAAGAATCAGCGG

GGAAAGAAGACCCTGTTGAGCTTGACTCTAGTCCGACTTTGTGAGATGAGTTGGGAGGTG

TAGTCTAAGAGGGAGGGGCAACTCGACCGTGAAATACCTCTACTCCCAAGCTTGTTTTAC

TTACTCCGGCGAGCAGGGATGGGTGGTAACACCCTTTTTCTAGGATTAACACCTCTTCGG

GGGTGGACCTGCGCGGAGGACATTGTCAGGTGGGGAGTTTGGCTGGGGCGGCACATCTGT

TAAAAGATAACGCAGGTGTCCTAAGATGAGCTCAGTGAGAACAGAAATCTCACGTAGAAC

AAAAGGGTAAAAGCTCATTTGATTTTGATTTTCAGTACGAATACAAACTGTGAAAGCATG

GCCTATCGATCCTTTTGACCTTCAAGGTTTGAAGCAAGAGGTGTCAGAAAAGTTAACACG

CGTTGTGGTAGCTTTTAAGAGTGGACTCAAAGTGTTCCGCTAGTCCCCCCTCGGACCGCC

ACTTACAAAAGGCCTGGGGATGGGGGCGACACTGTCAAATTGCGGGGACTCCCTAAAGCC

TGTCTCTACCAAGCCAGCGCGGCGAAAGCCCCTGGTGGCCGGGGTAATGACCTAGGGTAT

GGTAAAAGCGAGACAGGATGGAACAATGGGCAATCCGCAGCCAAGTCCTAAGGCTTTTTG

AAAAGCGGCTATGGATGCAGTTCAACGACTAGATGGCAGTGGGCTGAGAGAGCCCACCGC

GGCACCCTTTCGGCTTAAGATATAGTCTTGCCCACGTGAAAGCGTGCCTGCAGATGAACG

CCACCATGTATGGCCGGAGCTGCAGGAGACCGCACTTGATGTGCAGTCGGGAATCGACAG
TACCACAGGGATGAACAAAAACACACAATTAATTTGTGGCCATCATACAGTCGTATGATA
TTATCGTCCCAGGCGGGTGCGCCTCTCTCACCCGTCTAGTCCGGCCTTGGTGCGCAAAAA
ATGCTTGCGTGGATGCGCCTTGGTTAAAGAAAATGCTCCTAGTAGTTTTACAAGGACACT
TCTTGTAGGGCAACACATCTGGATGCGAAGAACTCCCGGAAAAGACAACGGTACCAAGTG
ACTGGGGTAACGCCCCATTCATGAAGGCAAATAATCAAAAAGAGGTGCCGGGTAAGAACC
CGTTGTATAGGGACAATCCGCAGCTGACTTGGCTCCCTGCCCATCCCCATGGGCATAGCT
TAGCGGTTCAACGCTCGGTAAGATGTGGGTTGGCGCGCTGTCCGACAGCGCACTGGCTTA
AGGTACGGGCTACTCCCACCGGAGACGGTGTCTTTGTCTACCTCGGGTAACACGCCTCGG
GAGAAGCAAAGAGGGTGCAATGGAAAAAAACTGGTTGCGGGGAAATCCCTACCAGAGTCA
CCTGGTACAAAGGAACTGGCTTGTGGCAGCCAAGCGTTCATAGCGACGTTGCTTTTTGAT
CCTTCGATGTCGGCTCTTCCTATCATTGCGAAGCAGAATTCGCCAAGTGTTGGATTGTTC
ACCCACTAATAGGGAACGTGAGCTGGGTAGCCACTTTATGCCCAACAGCTGGATAGCTGG
GCTTCTGTCTTAGGCCGGGACTTCTCACGACCCGTGCCTTGGTCACCCAAAAAAGGGCGA
CCGCAGAGCTCAGGATGGAGCTTCTGGCTTAACTAGTAGAGGACAGATAGAGATCGTCAA
ATTGCGGGAAACCCCTTAGAGCCTCGAGCTACCAAGTGCGTGTCGAAAGGCATAGACGCA
GCCAGCGATGGCTGTCGCATGGCTTGATTCCATCAAGATAGGTAAAAATGCTCGAGGATT
GGGCAATCCGCAGCCAAGCCCCTAAGGTCCCCCCGCGGGCTATGGGGAAGGTTCAACGAC
TGTAAGGCGTACCGTGCAAGCGGAAGATACAGTCTAGCCCCGCTGGAAAGCAGCGCCAAA
GTAAGTAGGTTGCAGTGGGTCTGTCCTC
>SRR9587955
GCATGTTTTGGTATGTGTTTCGACAGTCATGCATGTCATAGCCGTAGGATGGGGAGTGAG
CTTCATAGCCCGACCCCTTCTCAGGAGCATGTTTTCATAGCATATTGAGCATACTGGGGT
GGTCTCCCATGTAGCTCGCTCTCTCGGTGCAGGACCTCCCTTTTGCGGGCCACTTCGATG
GCCTCCATTATGGAGAGCATGTTTTGGCATATGAAGCGTTGGTGGTGAGGCTCGTAGCCT
GTATATACATACCAACGCAGTCGTCATATGTCTCTCTGGCGGGGGTCGGTCGTATGACCT
TGTCCCTACCTGGAGCATGTATTTTGTATGATATACTGGACGGCAGCTGCGCTGTTCAGT
CGTCGGCCGTACCTCATGTGCGCGTTTCCAGGTCCGAACGTTTGACATGCAAGGGCATCA
TACCAGAAGCCTGTTTTCTTGTATGAGCATGTAGGTAGGCAGGCTCTTCAGTCGTCTACC
CATAAATGCACTTCGAAGGAACTTTTTGGTATTGAATGGCTTTGTGCCCAGACGTTCTTG
TGAGCTTGTTCTCGCTCTGCGATACGCTGGAAGCTGTGGTATTACCTACAGCAGACGCAT
TCCTAGGGCGGACGGCCTCTCTGCAGATGCGCACATCTGTAGAAAGAGGCTCCCCTTGAT
GACGCAATGCAATGGCTGGGTAAACGGAGTGCTATACAGAAGGGTCAAACCTCTGTGGAA
GTACACTCATACCAGCATTGTTTGTTGAGAGGGGTCGGCTCTACGAGTGAGGACCTATAC
TGCCATCTCATTCAGATGCGTAGACGGTCTTTACTCAGACTGCCTTTCTGGCTGATTGTC
CTCGGTCCCTCGTGCGCACGCGCCGCTGGCTGAATGGGCTTGTCCTTATGGAGTGCCGCG

TTCTTCAAAGAATCCGTCTTCATACAGCGACCTCATTTACATATTGCCGGCTGAATGGGC
TTGTCCTTATGGAGCACTGCATTCTTCTAGGAATTGCTATCTATACAGCGACCTCATTTA
CATACAAGGCACCTTTTTCTACATAGACTTGCCATTTGGCTGATTGTTCTGACAGTCGCT
GCCTCTTCGGAGTTCTGTGTGGACGAGTGGCCTCCTGAGTGGGCCGGTCCTTACGGAGTG
CCCACGTCTGTTATGACTGGGTTCTCCATACAGCGACCTCACTCAGAATTGTATGGCACC
GTGCCGATTGAATGGACAAGCAGTCCTTAGGGAATCGCATGAAAGCATGACGGCACATGG
TGTATGGACCCGTTGGGCCTTGCCTGTGCTATCTTCTCGAGTCGACCGGTCCTCGTCTCC
TCCTTTCGCGACGGGAAGCTCTTGAGTTTACCGTGGGTCAGGACGCTGTGGTATCTGTTT
GTCTGTAGTGTGGGCTCTCTATCAGAACAGCGTATCAGCACACGTCGTCTGGGTCTATGG
TATTTTTGCTTAACGGCCCTGGGTTTGGTAGTATGGTTCCATGTCGGTCCTCTCACCTCC
CAACCGTCATGTCTCTGATATCATCTGCACAGTACGTCTTGTCAGAATATCATTCTGTAT
GGCAGTCGATAGACTAGGCCCTATGCCGCAGCTGTCTCTGACCACTATGCGAGCGCATAC
CTACCTCAATGTGTTCATAGCTGTTATGGATCTTGGGTGTGGTGTGCCAGAACGATACGC
CACCAGATTGTGGAATGGTGCTTGTAGTATTAGAAGAGGACATGTATGAAGAGCACGTTC
TTTTCTCTTATGATGGTGCCGCGGGTGCGGGCCGTTCTCATTCTGCTTGTCGGGTGCTCA
AGGTGCATGTAGGATTGCTGAACCCACTGAATGGTCATGAGAATGGTCATATGGTGAGGT
TCACTTCTCCATCGCTTTGTGTGTGTTGGATGTTTGGGTGGCTTCGGCCGCATGGAAAAT
CTCGGTCTGTAACTCATCAGGGAGGGTTTCCATGTGAGCAATCTCTCCAGTCGAACACAC
ACCCATGGGCGTCCCGCTAGGCAAAATGGAACTCTCGATCCGACACAGCCTGTGGAGTGT
TTGTTCCGTGTGGCGTGAAACGGCTTATACTGGATGTCTTCCCTTTGGACACCAGTCTGT
TGGTATGTACGCATACCATCTAGAGGGAGCAAGAAAAAGGCATGCAAGTGCCACAAAGGA
AGGGAGGCCTTCCTGCTACCGTGTGTCCTCTTGACGATCACGGGGGAGTATATAGAGACA
TTTGGACGAGATCCGTTTTTCACGAGGCGGTGCCCTCGCGCAAAGGTGTTGCTCTGCACG
CATCTTTATGATGATAGTTGCTCGCTCCTATGCGTCCACTCGTGAAAATACCTGCTCTCT
CTGTCTGTTTCCATGACACCCCCATGCGTTGTCATAGAGCATGCTCAGGTACAGAAAGTG
CTGTGACTTTCTTGGCCTGCTGTCACTGAGTCAGCCTGATACTTTGGAGGAATCTTTTCT
GGGCACTGCCCAGCTGAGAGTTCTTATCAGGCTCTGGAGTGTGCATATGCCTATATTCAT
TATAGTGCATGTGGCTACGCAGAAAAAGCATCGACGTGGAATGGAGAAGAACACCATTCT
GAATGGAACGGTTCTTACTCTACTTCTGTTGCTTGACGCTGTATGGGAATCGCTTTGCTC
TGGGGGTACACCTTCCATTCCATGACAAAGCATTTGATCTATGCCCCGAGGGTCATTCTC
TATGAAGAGCCCGAAACTTGTCTGAGTGAGAGGCGAAGGTAGTCCTTCATTTATGCGGGG
TCCGCCCTCACACTGTAGTCGGCACAGTCCCTTGTTTACGGACTGCGCAGCTCAATCTTC
GACTGGAGAGGAAAAACAAAACACAAGTATGACACGCACATACGTGTTTTCTAGGACTGT
GACCTGTCGGCTGCCTCGCGGTGGTTGGCATGATGCCCCAAGTGGGGGGGGGAGAAACAA
TCAGTATGATTGATAAAAACCAGTGGGGAAAAAGGCGCGCTGTCGCACACCCTTGGTCAC
GGCCTCAAGTCGACGGCCCTCACTGGCCCACCCTGCGACTCCTGTTCGCTTTTGACAAGA

CACACCTGCAATACGGAGGAAATGCCTGAGCAGCTGTGCCGCAGCAACTGCGGACGCGTA
GCGATATGCATCCCTGTTGTTTGCGACTTGGTACAAAGAACGTGCGCCGGGAGACGGATT
GCCCTTTGACTTTTCCACCCAGCCCGCGGCTCTATACCCTTGACGTACCTATTTACCTGG
TTGATCCTGCCAGTAGTCATATGCTTGTCTCAAAGACTAAGCCATGCATGTCTAAGTATA
AACACTTTTATACTCGTGAAACTGCGAATGGCTCATTAAAACAGTTATAGTTCCTTTGGG
AACAACACTACTTGGATAACCGTAGTAATTCTAGAGCTAATACATGCCTCAACGCCCGAC
TCACGAAGGGTGGTATTTATTGGATAAAAAACCATCGTGCTCTGTCCAGAGCGCTTTGAG
ATGATTCACAATAACTTGTCGGATCGCATGGCCTCGTGCTGGCGACTCCTCATTCAAATT
TCTGCCCTATCAACTTTCGATGGTAGAGTATTGGTCTACCATGGTGTCGACGGGTGACGG
GGAATTAGGGTTCGATTCCGGAGAGGGAGCCTGAGAAACGGCTACCACATCCAAGGAAGG
CAGCAGGCGCGCAAATTACCCAATCCCGACTCGGGGAGGTAGTGACAAAAAATAACAATA
GGGGGCCCTTTGGGTCTTCTAATTGGAATGAGAACAATTTAAATCCCTTATCGAGGATCC
ATTGGAGGGCAAGTCTGGTGAACGTATGCATTAAAGCCCTCCGTAATAAAAAAAAACAGC
GCCAGATGTCTCTCTCTGGAAGCACACCATGCATGGGGTTTGTTCCTCTCTGCATGGCCG
TGTGAAAAAGGCGAGTAGGCAAACGATGGGAAACCCGGCTTGGAATAGTAACCAGCCACA
GAGCCCCCATCATACACTTCTAAATTGCCGGGGAGTGCCCGCGAGGCCTTGACTACCGCG
GTCCGCCGAGAGGTGAGAGCGAGCAATCGTTCAGGACGCAGCAGTGTACGTAGTGGTACA
TGGATGGTAAAAACGTCAAGTATAGGGTTGATCCGCAGGGAAGCGCGAGTGCCGCCATGG
TCGGTGGGCAAAGGCGAACCTTCAGAGACTCTAATGGAGTGGGCGCAAGCTTAAGGGAGA
GTCCAATCCACTGGGAAACCAGGTCCACAGCAGTAACGGGGTGATGCTCGCCCCAGAGGA
CTGTGGAAGGCAACCCCACCAAAAGGGTGTGTGTTGTCGGGTAGCAATGGCCAGCAGCCG
CGGTAATTCCAGCTCCAATAGCGTATATTAAAGTTGTTGCAGTTAAAACGCTCGTAGTCG
GACCTCGGAGCGCCCCCACGGGCCGGGCTTTTGCCTTGTTGCCAGTGGGAGTTTGTCTCC
TTTTGTCGGGACGCGGTGTGCAGGGCCTTTACTGTCTCTGCGCGTTTACCTGCGGGCCGA
CCGTTTACTGTGAAGAAAGTAAAGTGTTCAAAGCAGGCCAATGCCTTGAATATGTGAGCA
TGGAATAATAGAATAGGACTTGGGCTCTATTTTGTTGGTTTCCAGTGACCAAGTAATGAT
TAATAGGGACGGTTGGGGGCATTCGTATTTCATTGTCAGAGGTGAAATTCTTGGATTGAT
GGAAGACGAACAACTGCGAAAGCATCTGCCATGGATGTTTTCATTGATCAAGAACGAAAG
TTAGGGGATCGAAGACGATCAGATACCGTCGTAGTCTTAACCATAAACGATGCCGACTGG
GGATTGGCGGGGGCACTTTTATATGACTCTGTCAGCACCCTGAGGGAAACCAAAGTCTTT
GGGTTCTGGGGGGAGTATGGTCGCAAGGCTGAAACTTAAAGGAATTGACGGAAGGGCACC
ACCAGGAGTGGAGCCTGCGGCTTAATTTGACTCAACACGGGAAAACTTACCAGGTCCGGA
CAGAAGAATGATTGACAGACTGAAAGCTCTTTCTTGATTTTTTGGTTGGTGGTGCATGGC
CGTTCTTAGTTGGTGGAGTGATTTGTCTGGTTAATTCCGTTAACGAACGAGACCTCGTCC
TGCTAAATAGGTGCGCGCATGCCAGCACTGCGTTCTTACCTTCTTAGAGGGACTATGCGC
GTCTAGCGTATGGAAGATTGAGGCAATAACAGGTCTGTGATGCCCTTAGATGTTCTGGGC

CGCACGCGCGCTACACTGATGCATTCAACGAGTTTCACTTGATAGTCCTGGGTCGGAAGG
CTCGGGTAATCTTTTGAAAGTGCATCGTGCTGGGGATAGATCATTGCAATTATTGATCTT
CAACGAGGAATTCCTTGTAGGCGCAGGTCATCAGCCTGCGCCGAATACGTCCCTGCCCTT
TGTACACACCGCCCGTCGCTCCTACCGATTGAATGGTCCGATGAAATGTCCGGATCGCGG
CTAGTGTGATGGACTCGTCCAGAGCGGTAGCCGTGAGAAGCTCATTAAATCTTATCATTT
AGAGGAAGGAGAAGTCGTAACAAGGTTTCCGTAGGTGAACCTGCGGAAGGATCATTCACA
AACTATTGACACAAACACACGCGAACCAAATCGTCCGCACAGCAGGTGGCAATGAAAGAG
AGACTCTGCATGTCGCCTTTCGGGGTAGCAAGCAGCACTCTTCTGCCATCGCCTCTGTGC
AGGGCGTAAATTCCCATTGAGAGGATGTGAGGGCACCACAGGAAGCTTTTCCACAGGAAG
TCGCCATCCTTCTCCCTCCACGGCGGCGCTTCTGTGCTTGGCAGTTTTTTTTGCCTTCTA
GGGAGGATGCCGCCAATGGAGCCCCTTATATATATACATCATCATATGCCCCTTTTTTTC
TTAACCGCTTGCCACAGCTTCTTCTATGAGGAGCTTGTGGGAAGACTGTCTCCATACAAT
AACAAAGATACAACTCTTAGCGGTGGATATCTTGGTTCTCGCAACGATGAAGAACGCAGC
CAACTGCGATAACTAATGTGAATTGCAGGACTTCGTGAATCATTCAGTCTTTGAACGCAA
GTTGCGCTCATCTCCATGAGGAGGTGAGCATGCCTGTTTGAGTGTCGATGGGAAAAACAA
TACATAACAACAATCGGCTCTCTTATCTTCCTGCCTTCTTGGGGGCAAAAGGTAGTGCAT
GCCGGAGATGGGCTTTGTCGCCCCTTTCTCTTGAAAGAGGCGTCGTCCGAAACACGTATC
GTTGCCCGCTGGAGGTCATAGAGCATGAGTTTAAAGCACCACATCTCTCAAGTCAGAGTG
AGCGTCTTGAGTAGGTCCGCATTGATTCAGTTTCAACCCAGGTGGCGTTGGTTACTTGCC
ATAAGCCTCTCTGGGGTTTCTTTGCTTGGACCTCTCTTGTCAGTTTCTCTGACGTAGAGA
CAGGTGCCGTCTCATGGACGTTGCCCTCTGGAACGTGCTAAAAAAAAAAAAAAAGAAAAAA
CCGTACCACATGTCAAAAGAGGCAGATCGTGCTTGACGGCGGGAGGTGTTGCATTTCTTG
CTTGTCCTGAGCGTCTGCCAGTCTCTGACGAGTGTGTTCCTTCCATAAACCCGATCTCAA
ATCAGGTAGGACCACCCGCTGAATTTAAGCATATAATTAAGCGGAGGAAAAGAAACTAAC
AAGGATTCCCTCAGTAGCGGCGAGCGAAGCGGGAAGAGCTCAACATGAAAACCGTTGGGG
GTTCGCGCTTTCCTTCGGGTTGTATAGTAAAGTGGTGTCGCCTGCGTCGCGCCGGGTATA
AATCCCGTCTGAAATGGGGTGTCGTAGAGGGTTATATCCCCGTCTGTGACCCGGTGTGCG
ATGCGTTACGGTGACCTTACGATGAGTCGGGTTGCTTGGGCATGCAGCCCTAAGTGGGTG
GTAAACTTCATCTAAAGCTAAATACTAGCGAGAGACCGATAGCGTACAAGTACCGTAAGG
GAAAGATGCAAAGAACTTTGAAAAGAGAGTTAAACAGGGCTTGAAACCGTTAAGAGTGAA
GCGAAGGAGGTCAGTGATCATGGGGCCAGATCCCCAAGTGCCTGTCATAGGGCACCGTAG
CGTTCTGGTTTCCCGGGGCAGCGCCGGCCGGGGGCGGGGGTTAACAGGGAGGTGGTTGAG
GGATTCGTTTTCCGCCTGTACTCCTTTATAGCTTGCACCCCGGCCGAGGGATCCGCGACT
TGTCGTGTGAGGGCGCTGTCAAAATGGCCTTCTCCACCCCGTCTTGAAACACGGACCAAG
GAGTCTAACATACTCGCGAGTGTTTGGGTGGTTAAAACCCGAACGCACAATGAAAGTAAA
CGGTGGCTCTGCGCCGCTGCTTGGTGGGAACCCTGCTTTTATAATGGGGGGCACTATCAC

CCGACCCCGATCCCTTTGGGGTGGTTGGTTTTGAGGTAGAGCGAGTATGATGGGACCCGA
AAGATGGTGAACTATTCCTGAGTAGGGCGAAGCCAGAGGAAACTCTGGTGGAGGCTCGTA
GCGATGCTGACGTGCAAATCGCTCGTCGAACTTGGGTATAGGGGCGAAAGACTAATCGAA
CCATCTAGTAGCTGGTTCCCTCCGAAGTTTCCCTCAGGATAGCGGGAGCAAATGCAAGTC
AGTATCATCCGGTAAAGCGAATGATTAGAGGCCTCGGGGGAGTAACTCCTTCGACCTATT
CTCAAACTTTAAATAGGTGAGGAGCCGGCAACGTTGCTTAAAACAATGAACGCCGTGCGG
ACAATGATTGCTCTCAGTGGGCCGATTTTGGTAAGCAGAACTGGCGATGCGGGATGAACC
GAAAGCGACGCTAAGGTGCCAAACTCCCAGCTCATTCAGACACCAGAAAAGGTGTTGATT
CATACAGACAGCAGGACGGTGGCCATGGAAGTCGGAATCCGCTAAGGAGTGTGTAACAAC
TCACCTGCCGAATGGATCAGCCCTGAAAATGGATGGCGCTTAAGCGGGGGACCGAGGCGT
CGCCGTTGGGGTAAATACGAAGCCCCAATGAGTAGGAGGGCGTGGGGGTTTTGAAACACG
GTACACTCGCATGGAAGCCTTGGGCGTAAGCTCGGGTGGAGCGACCCCTAGTGTAGATCT
TGGTGGTAGTAGCAAATATTCAAGTGAGAACCTTGAAGACTGAAGTGGGGAAGGGTTCCA
CGCGAACAGTGCTTGGACGTGGGTTAGTCGGTCCTAAGAGATGGGGAAACTCCGTATTCA
AGTGCGCTTTTGTAGCGCAGTCCATCGAAAGGGAAGCGGGTTAACATTCCCGCACCAGGA
TGTGGAAACTGTGGTTTACCGCAAACTGCGGCAACGCAACTGAACCTGGGGACGCCCGTA
GGAGCTCCGAGCAGGGATCTCTTTCCTGTTTAACGCGCCCAACCCCCTGGAATCGGTTTG
TCCGGAGATAGGGGGCGAAAGGCCGGAAAAGCACCGCACGTCTTGTGGTGTTCGGAACGC
TTGTAATGGCCCTTGAAAACCCAGGGGAAAGCAGCCGTCTTACACTTGTGTGAGACGAGG
AATTCTCACACCTGGCCGTACCCATCACCGCATCAGGTCCCCAAGGTGAACAGCCTCTAG
TCAATAGAATAATGTAGATAAGGGAAGTCGGCAAAATAGATCCGTAACTTTGGGAAAAGG
ATTGGCTCTAATGTTCGGGCTTGTCGGGACGCGCGGGCGTAGATGCCAATGAGCGGCAGA
CGTCTAGCCTCTTTGGAGGCTGGATTAACGCTGCTTGTGGGAAGAAACCGTGTCGATGGA
GGCAGGAACGCTTATCGATCTTGTCTCTTCCCGCTTGCGATCAACGAACGGATTAGAACT
GGTACGGACAAGGGGAATCCGACTGTTTAATTAAAACAAAGCATTGCGACGGCCGGAAAC
GGTGTTGGCGCAATGTGATTTCTGCCCAGTGCTCTGAATGTCAAAGTGAAGAAATTCAAC
CAAGCGCGGGTAAACGGCGGGAGTAACTATGACTCTTAAACAAACCATACTACCTACGGG
AGTCAAAAGTGGGGGCGGCGTCTGGTGTAGAAATACACGTACGACACGCTTTTTTTCCTG
CTAGTCGATCTGACCTTTGTTCTTTGGACTGTGGGCGGTCAGCGGCAACATAACCTGGTA
CGGGGAAGGCCTCGACGTTGAGTGCTGCATCTTGGAAGTATCTTGACACAATAAGAGAAA
AGAGATTGCGCTTAATAGGGGGTTGATCTCGTGGCGAGCCGACGCCGGTGTGAACTGGTG
GCAATGGCCGTCGTAGAGCACGCAAAGGTATGGGCTGACTTTTTCTCCTCAAAAGTTGGC
TTAAGGGACGTGCCATCCCACACGATAATAAAGTGTGCAGTCTGTAAGAGCTCCCACAGG
GCAAAGACAGACTGGGATGCAGACGGAGACGGCAAAAACAACAACATCCGTTCTGCTTCT
GGGCGCAAATGCTTAAGGTAGCCAAATGCCTCGTCATCTAAACTAATTGGATGAGATAGT
CTGAGTGAAAAGCTTATGCAGGCTAGTACCGGAAAAAAAAAAAAAAGAGGGGCTCTGCCAC

TCTGACGGTGCGACACTCCTTGTACAGGGAAACCCTCCGCATGGAAAGGCACGTGGTACC
ACCCTCGCGGGGAAACTCGCGAGCGGGATCCCGGGTAATGACCGGAGCCGATGGTAAAAA
CTCACGTGACCGATGGGGTGACCCTGTGGCGAGACGCTAAGTTGTGGGACATCACCCTCT
CACAATACGCATCCGTCGCAGAGACTGAACGGGAGTGGGCTCCGTTGACGGCCGGCCGGC
CAGCCAGCCGTCTTCCAGGGGCTTAAGATACAGTCCGGGCCTCAGGGATGGAAAAACCTG
GGGAACTACCAGAATTAGTGACGCGCATGAATGGATTAACGAGATTCCCACTGTCCCTAT
CTACTATCTAGCGAAACCACAGCCAAGGGAACGGGCTTGGAAGAATCAGCGGGGAAAGAA
GACCCTGTTGAGCTTGACTCTAGTCCGACTTTGTGAGATGAGTTGGGAGGTGTAGTCTAA
GAGGGAGGGGCAACTCGACCGTGAAATACCTCTACTCCCAAGCTTGTTTTACTTACTCCG
GCGAGCAGGGATGGGTGGTAACACCCTTTTTCTAGGATTAACACCTCTTCGGGGGTGGAC
CTGCGCGGAGGACATTGTCAGGTGGGGAGTTTGGCTGGGGCGGCACATCTGTTAAAAGAT
AACGCAGGTGTCCTAAGATGAGCTCAGTGAGAACAGAAATCTCACGTAGAACAAAAGGGT
AAAAGCTCATTTGATTTTGATTTTCAGTACGAATACAAACTGTGAAAGCATGGCCTATCG
ATCCTTTTGACCTTCAAGGTTTGAAGCAAGAGGTGTCAGAAAAGTTAACACGCGTTGTGG
TAGCTTTTAAGAGTGGACTCAAAGTGTTCCGCTAGTCCCCCCTCGGACCGCCACTTACAA
AAGGCCTGGGGATGGGGGCGACACTGTCAAATTGCGGGGACTCCCTAAAGCCTGTCTCTA
CCAAGCCAGCGCGGCGAAAGCCCCTGGTGGCCGGGGTAATGACCTAGGGTATGGTAAAAG
CGAGACAGGATGGAACAATGGGCAATCCGCAGCCAAGTCCTAAGGCTTTTTGAAAAGCGG
CTATGGATGCAGTTCAACGACTAGATGGCAGTGGGCTGAGAGAGCCCACCGCGGCACCCT
TTCGGCTTAAGATATAGTCTTGCCCACGTGAAAGCGTGCCTGCAGATGAACGCCACCATG
TATGGCCGGAGCTGCAGGAGACCGCACTTGATGTGCAGTCGGGAATCGACAGTACCACAG
GGATGAACAAAAACACACAATTGTGGCCATCATACAGTCGTATGATATTATCGTCCCAGG
CGGGTGCGCCTCTCTCACCCGTCTAGTCCGGCCTTGGTGCGCAAAAAATGCTTGCGTGGA
TGCGCCTTGGTTAAGGAAAATGCTCCTAGTAGTTTTACAAGGACACTTCTTGTAGGGCAA
CACATCTGGATGCGAAGAACTCCCGGAAAAGACAACGGTACCAAGTGACTGGGGTAACGC
CCCATTCATGAAGGCAAATAATCAAAAAGAGGTGCCGGGTAAGAACCCGTTGTATAGGGA
CAATCCGCAGCTGACTTGGCTCCCTGCCCATCCCCATGGGCATAGCTTAGCGGTTCAACG
CTCGGTAAGATGTGGGTTGGCGCGCTGTCCGACAGCGCACTGGCTTAAGGTACGGGCTAC
TCCCACCGGAGACGGTGTCTTTGTCTACCTCGGGTAACACGCCTCGGGAGAAGCAAAGAG
GGTGCAATGGAAAAAAACTGGTTGCGGGGAAATCCCTACCAGAGTCACCTGGTACAAAGG
AACTGGCTTGTGGCAGCCAAGCGTTCATAGCGACGTTGCTTTTTGATCCTTCGATGTCGG
CTCTTCCTATCATTGCGAAGCAGAATTCGCCAAGTGTTGGATTGTTCACCCACTAATAGG
GAACGTGAGCTGGGTAGCCACTTTATGCCCAACAGCTGGATAGCTGGGCTTCTGTCTTAG
GCCGGGACTTCTCACGACCCGTGCCTTGGTCACCCAAAAAAGGGCGACCGCAGAGCTCAG
GATGGAGCTTCTGGCTTAACTAGTAGAGGACAGATAGAGATCGTCAAATTGCGGGAAACC
CCTTAGAGCCTCGAGCTACCAAGTGCGTGTCGAAAGGCATAGACGCAGCCAGCGATGGCT

GTCGCATGGCTTGATTCCATCAAGATAGGTAAAAATGCTCGAGGATTGGGCAATCCGCAG
CCAAGCCCCTAAGGTCCCCCCGCGGGCTATGGGGAAGGTTCAACGACTGTAAGGCGTACC
GTGCAAGCGGAAGATACAGTCTAGCCCCGCTGGAAAGCAGCGCCAAAGTAAGTAGGTTGC
AGTGGGTCTGTCCTC
>SRR9587962
GCATGTTTTGGTATGTGTTTCGACAGTCATGCATGTCATAGCCGTAGGATGGGGAGTGAG
CTTCATAGCCCGACCCCTTCTCAGGAGCATGTTTTCATAGCATATTGAGCATACTGGGGT
GGTCTCCCATGTAGCTCGCTCTCTCGGTGCAGGACCTCCCTTTTGCGGGCCACTTCGATG
GCCTCCATTATGGAGAGCATGTTTTGGCATATGAAGCGTTGGTGGTGAGGCTCGTAGCCT
GTATATACATACCAACGCAGTCGTCATATGTCTCTCTGGCGGGGGTCGGTCGTATGACCT
TGTCCCTACCTGGAGCATGTATTTTGTATGATATACTGGACGGCAGCTGCGCTGTTCAGT
CGTCAGCCGTACCTCATGTGCGCGTTTCCAGGTCCGAACGTTTGACATGCAAGGGCATCA
TACCAGAAGCCTGTTTTCTCGTATGAGCATGTAGGTAGGCAGGCTCTTCAGTCGTCTACC
CATAAATGCACTTCGAAGGAACTTTTTGGTATTGAATGGCTTTGTGCCCAGACGTTCTTG
TGAGCTTGTTCTCGCTCTGCGATACGCTGGAAGCTGTGGTATTACCTACAGCAGACGCAT
TCCTAGGGCGGACGGCCTCTCTGCAGATGGGCACATCTGTAGAAAGAGGCTCCCCTTGAT
GACGCAATGCAATGGCTGGGTAAACGGAGTGCTATACAGAAGGGTCAAACCTCTGTGGAA
GTACACTCATACCAGCATTGTTTGTTGAGAGGGGTCGGCTCTACGAGTGAGGACCTATAC
TGCCATCTCATTCAGATGCGTAGACGGTCTTTACTCAGACTGCCTTTCTGGCTGATTGTC
CTCGGTCCCTCGTGCGCACGCGCCGCTGGCTGAATGGGCTTGTCCTTATGGAGTGCCGCG
TTCTTCAAAGAATCCGTCTTCATACAGCGACCTCATTTACATATTGCCGGCTGAATGGGC
TTGTCCTTATGGAGCACTGCATTCTTCTAGGAATTGCTATCTATACAGCGACCTCATTTA
CATACAAGGCACCTTTTTCTACATAGACTTGCCATTTGGCTGATTGTTCTGACAGTCGCT
GCCTCTTCGGAGTTCTGTGTGGACGAGTGGCCTCCTGAGTGGGCCGGTCCTTACGGAGTG
CCCACGTCTGTTATGACTGGGTTCTCCATACAGCGACCTCACTCAGAATTGTATGGCACC
GTGCCGATTGAATGGACAAGCAGTCCTTAGGGAATCGCATGAAAGCATGACGGCACATGG
TGTATGGACCCGTTGGGCCTTGCCTGTGCTATCTTCTCGAGTCGACCGGTCCTCGTCTCC
TCCTTTCGCGACGGGAAGCTCTTGAGTTTACCGTGGGTCAGGACGCTGTGGTATCTGTTT
GTCTGTAGTGTGGGCTCTCTATCAGAACAGCGTATCAGCACACGTCGTCTGGGTCTATGG
TATTTTTGCTTAACGGCCCTGGGTTTGGTAGTATGGTTCCATGTCGGTCCTCTCACCTCC
CAACCGTCATGTCTCTGATATCATCTGCACAGTACGTCTTGTCAGAATATCATTCTGTAT
GGCAGTCGATAGACTAGGCCCTATGCCGCAGCTGTCTCTGACCACTATGCGAGCGCATAC
CTACCTCAATGTGTTCATAGCTGTTATGGATCTTGGGTGTGGTGTGCCAGAACGATACGC
CACCAGATTGTGGAATGGTGCTTGTAGTATTAGAAGAGGACATGTATGAAGAGCACGTTC
TTTTCTCTTATGATGGTGCCGCGGGTGCGGGCCGTTCTCATTCTGCTTGTCGGGCGCTCA
AGGTGCATGTAGGATTGCTGAACCCACTGAATGGTCATGAGAATGGTCATATGGTGAGGT

TCACTTCTCCATCGCTTTGTGTGTGTTGGATGTTTGGGTGGCTTCGGCCGCATGGAAAAT
CTCGGTCTGTAACTCATCAGGGAGGGTTTCCATGTGAGCAATCTCTCCAGTCGAACACAC
ACCCATGGGCGTCCCGCTAGGCAAAATGGAACTCTCGATCCGACACAGCCTGTGGAGTGT
TTGTTCCGTGTGGCGTGAAACGGCTTATACTGGATGTCTTCCCTTTGGACACCAGTCTGT
TGGTATGTACGCATACCATCTAGAGGGAGCAAGAAAAAGGCATGCAAGTGCCACAAAGGA
AGGGAGGCCTTCCTGCTACCGTGTGTCCTCTTGACGATCACGGGGGAGTATATAGAGACA
TTTGGACGAGATCCGTTTTTCACGAGGCGGTGCCCTCGCGCAAAGGTGTTGCTCTGCACG
CATCTTTATGATGATAGTTGCTCGCTCCTATGCGTCCACTCGTGAAAATACCTGCTCTCT
CTGTCTGTTTCCATGACACCCCCATGCGTTGTCATAGAGCATGCTCAGGTACAGAAAGTG
CTGTGACTTTCTTGGCCTGCTGTCACTGAGTCAGCCTGATACTTTGGAGGAATCTTTTCT
GGGCACTGCCCAGCTGAGAGTTCTTATCAGGCTCTGGAGTGTGCATATGCCTATATTCAT
TATAGTGCATGTGGCTACGCAGAAAAAGCATCGACGTGGAATGGAGAAGAACACCATTCT
GAATGGAACGGTTCTTACTCTACTTCTGTTGCTTGACGCTGTATGGGAATCGCTTTGCTC
TGGGGGTACACCTTCCATTCCATGACAAAGCATTTGATCTATGCCCCGAGGGTCATTCTC
TATGAAGAGCCCGAAACTTGTCTGAGTGAGAGGCGAAGGTAGTCCTTCATTTATGCGGGG
TCCGCCCTCACACTGTAGTCGGCACAGTCCCTTGTTTATGGACTGCGCAGCTCAATCTTC
GACTGGAGAGGAAAAACAAAACACAAGTATGACACGCACATACGTGTTTTCTAGGACTGT
GACCTGTCGGCTGCCTCGCGGTGGTTGGCATGATGCCCCAAGTGGGGGGGGGAGAAACAA
TCAGTATGATTGATAAAAACCAGTGGGGAAAAAGGCGCGCTGTCGCACACCCTTGGTCAC
GGCCTCAAGTCGACGGCCCTCACTGGCCCACCCTGCGACTCCTGTTCGCTTTTGACAAGA
CACACCTGCAATACGGAGGAAATGCCTGAGCAGCTGTGCCGCAGCAACTGCGGACGCGTA
GCGATATGCATCCCTGTTGTTTGCGACTTGGTACAAAGAACGTGCGCCGGGAGACGGATT
GCCCTTTGACTTTTCCACCCAGCCCGCGGCTCTATACCCTTGACGTACCTATTTACCTGG
TTGATCCTGCCAGTAGTCATATGCTTGTCTCAAAGACTAAGCCATGCATGTCTAAGTATA
AACACTTTTATACTCGTGAAACTGCGAATGGCTCATTAAAACAGTTATAGTTCCTTTGGG
AACAACACTACTTGGATAACCGTAGTAATTCTAGAGCTAATACATGCCTCAACGCCCGAC
TCACGAAGGGTGGTATTTATTGGATAAAAAACCATCGTGCTCTGTCCAGAGCGCTTTGAG
ATGATTCACAATAACTTGTCGGATCGCATGGCCTCGTGCTGGCGACTCCTCATTCAAATT
TCTGCCCTATCAACTTTCGATGGTAGAGTATTGGTCTACCATGGTGTCGACGGGTGACGG
GGAATTAGGGTTCGATTCCGGAGAGGGAGCCTGAGAAACGGCTACCACATCCAAGGAAGG
CAGCAGGCGCGCAAATTACCCAATCCCGACTCGGGGAGGTAGTGACAAAAAATAACAATA
GGGGGCCCTTTGGGTCTTCTAATTGGAATGAGAACAATTTAAATCCCTTATCGAGGATCC
ATTGGAGGGCAAGTCTGGTGAACGTATGCATTAAAGCCCTCCGTAATAAAAAAAAACAGC
GCCAGATGTCTCTCTCTGGAAGCACACCATGCATGGGGTTTGTTCCTCTCTGCATGGCCG
TGTGAAAAAGGCGAGTAGGCAAACGATGGGAAACCCGGCTTGGAATAGTAACCAGCCACA
GAGCCCCCATCATACACTTCTAAATTGCCGGGGAGTGCCCGCGAGGCCTTGACTACCGCG

GTCCGCCGAGAGGTGAGAGCGAGCAATCGTTCAGGACGCAGCAGTGTACGTAGTGGTACA

TGGATGGTAAAAACGTCAAGGATAGGGTTGATCCGCAGGGAAGCGCGAGTGCCGCCATGG

TCGGTGGGCAAAGGCGAACCTTCAGAGACTCTAATGGAGTGGGCGCAAGCTTAAGGGAGA

GTCCAATCCACTGGGAAACCAGGTCCACAGCAGTAACGGGGTGATGCTCGCCACAGAGGA

CTGTGGAAGGCAACCCCACCAAAAGGGTGTGTGTTGTCGGGTAGCAATGGCCAGCAGCCG

CGGTAATTCCAGCTCCAATAGCGTATATTAAAGTTGTTGCAGTTAAAACGCTCGTAGTCG

GACCTCGGAGCGCCCCCACGGGCCGGGCTTTTGCCTTGTTGCCAGTGGGAGTTTGTCTCC

TTTTGTCGGGACGCGGTGTGCAGGGCCTTTACTGTCTCTGCGCGTTTACCTGCGGGCCGA

CCGTTTACTGTGAAGAAAGTAAAGTGTTCAAAGCAGGCCAATGCCTTGAATATGTGAGCA

TGGAATAATAGAATAGGACTTGGGCTCTATTTTGTTGGTTTCCAGTGACCAAGTAATGAT

TAATAGGGACGGTTGGGGGCATTCGTATTTCATTGTCAGAGGTGAAATTCTTGGATTGAT

GGAAGACGTACAACTGCGAAAGCATCTGCCATGGATGTTTTCATTGATCAAGAACGAAAG

TTAGGGGATCGAAGACGATCAGATACCGTCGTAGTCTTAACCATAAACGATGCCGACTGG

GGATTGGCGGGGGCACTTTTATATGACTCTGTCAGCACCCTGAGGGAAACCAAAGTCTTT

GGGTTCTGGGGGGAGTATGGTCGCAAGGCTGAAACTTAAAGGAATTGACGGAAGGGCACC

ACCAGGAGTGGAGCCTGCGGCTTAATTTGACTCAACACGGGAAAACTTACCAGGTCCGGA

CAGAAGAATGATTGACAGACTGAAAGCTCTTTCTTGATTTTTTGGTTGGTGGTGCATGGC

CGTTCTTAGTTGGTGGAGTGATTTGTCTGGTTAATTCCGTTAACGAACGAGACCTCGTCC

TGCTAAATAGGTGCGCGCATGCCAGCACTGCGTTCTTACCTTCTTAGAGGGACTATGCGC

GTCTAGCGTATGGAAGATTGAGGCAATAACAGGTCTGTGATGCCCTTAGATGTTCTGGGC

CGCACGCGCGCTACACTGATGCATTCAACGAGTTTCACTTGATAGTCCTGGGTCGGAAGG

CTCGGGTAATCTTTTGAAAGTGCATCGTGCTGGGGATAGATCATTGCAATTATTGATCTT

CAACGAGGAATTCCTTGTAGGCGCAGGTCATCAGCCTGCGCCGAATACGTCCCTGCCCTT

TGTACACACCGCCCGTCGCTCCTACCGATTGAATGGTCCGATGAAATGTCCGGATCGCGG

CTAGTGTGATGGACTCGTCCAGAGCGGTAGCCGTGAGAAGCTCATTAAATCTTATCATTT

AGAGGAAGGAGAAGTCGTAACAAGGTTTCCGTATTTTGTGGGTGGATACCGATTCAATAA

CATCTGCCGTTTTTCTCAATGTACTATATCCCTAGTCTGCTTGCCCAGTTCTTGGAGTCT

CGTGCGATGTGTGCTATTACAGGCACGCTGGGTCATGAAGACAGACAAGGTTAGGATCTT

CACGGAAGTACGCGCAATAGGAGCGCGTCTTGTTCAACGCACCTGACTCAAGAGTCATAT

GCGAGGGACGGACGCAGCGCTTGTCGCCGCACAGATGGGAAGCTTCCTCTCCAGGCATAG

GATGGCGCTGCATTTGCATCATCGCTGCTAGTAAGTGGCAATAATACTTGATTCCCCGCC

AGCGCACTTGTACGTAGCCGTTGGCGGCGGGAACGTAGTTGGTGATGTGACAGCCTCGAG

CGTTCACCTCTGTGTGTCCGCCAATCTGCACAAAGATGAACGCCATATCGGAGACAGGGG

GAGGTGCCATTGTGATGATGACGATGTTGTTGGGTGGTCCTTCCGTAGGAAAACGTACAC

GAAGCCTTTGCGGCCCGAAAGGGTGCACAGTGCGCGACTATAAACACTGATTCTGTGGTC

AACGCAAGCAGCGGTTCTGTGGAAGAACAGCTGCGACTTTCTCAAATTGCGGGAACACCC

TAAAGTCAACAAGCACCGCGGCTTGACGTGAAAAGCGTCAGGCGCAGCAGTCGGGGTAGC
GCCCTGGCGGATGGTAATAGCCTTGTTGAATGGCACAATGGGCAATCCGCAGCCAAGCCC
CTAAGGTCCCCCGCTGGGCTATGGGGAAGGTTCACAGACTGTAAGGGAGGGGGTTTCTTC
CATGAAAGAAGCTTAAGAGACAGTCGGTCCCCTGCGAGAGCAGTGTCGCATGTGAGGAAG
GCACCCATGAAAAGTGGGTGCTGGAGAGCCATGTGAGCCCTCTATGAGGGCGGGAAGTTT
TCCGATACAATGAGAAACAAACACGTTCCGTAGGTGAACCTGCGGAAGGATCATTCACAA
ACTATTGACACAAACACACGCGAACCAAATCGTCCGCACAGCAGGTGGCAATGAAAGAGA
GACTCTGCATGTCGCCTTTCGGGGTAGCAAGCAGCACTCTTCTGCCATCGCCTCTGTGCA
GGGCGTAAATTCCCATTGAGAGGATGTGAGGGCACCACAGGAAGCTTTTCCACAGGAAGT
CGCCATCCTTCTCCCTCCACGGCGGCGCTTCTGTGCTTGGCAGTTTTTTTTGCCTTCTAG
GGAGGATGCCGCCAATGGAGCCCCTTATATATATACATCATCATATGCCCCTTTTTTTCT
TAACCGCTTGCCACAGCTTCTTCTATGAGGAGCTTGTGGGAAGACTGTCTCCATACAATA
ACAAAGATACAACTCTTAGCGGTGGATATCTTGGTTCTCGCAACGATGAAGAACGCAGCC
AACTGCGATAACTAATGTGAATTGCAGGACTTCGTGAATCATTAAGTCTTTGAACGCAAG
TTGCGCTCATCTCCATGAGGAGGTGAGCATGCCTGTTTGAGTGTCGATGGGAAAAACAAT
ACATAACAACAATCGGCTCTCTTATCTTCCTGCCTTCTTGGGGGCAAAAGGTAGTGCATG
CCGGAGATGGGCTTTGTCGCCCCTTTCTCTTGAAAGAGGCGTCGTCCGAAACACGTATCG
TTGCCCGCTGGAGGTCATAGAGCATGAGTTTAAAGCACCACATCTCTCAAGTCAGAGTGA
GCGTCTTGAGTAGGTCCGCATTGATTCAGTTTCAACCCAGGTGGCGTTGGTTACTTGCCA
TAAAGCCTCTCTGGGGTTTCTTTGCTTGGACCTCTCTTGTCAGTTTCTCTGACGTAGAGA
CAGGTGCCGTCTCATGGACGTTGCCCTCTGGAACGTGCTAAAAAAAAAAAAGAAAAAACC
GTACCACATGTCAAAAGAGGCAGATCGTGCTTGACGGCGGGAGGTGTTGCATTTCTTGCT
TGTCCTGAGCGTCTGCCAGTCTCTGACGAGTGTGTTCCTTCCATAAACCCGATCTCAAAT
CAGGTAGGACCACCCGCTGAATTTAAGCATATAATTAAGCGGAGGAAAAGAAACTAACAA
GGATTCCCTCAGTAGCGGCGAGCGAAGCGGGAAGAGCTCAACATGAAAACCGTTGGGGGT
TCGCGCTTTCCTTCGGGTTGTATAGTAAAGTGGTGTCGCCTGCGTCGCGCCGGGTATAAA
TCCCGTCTGAAATGGGGTGTCGTAGAGGGTTATATCCCCGTCTGTGACCCGGTGTGCGAT
GCGTTACGGTGACCTTACGATGAGTCGGGTTGCTTGGGCATGCAGCCCTAAGTGGGTGGT
AAACTTCATCTAAAGCTAAATACTAGCGAGAGACCGATAGCGTACAAGTACCGTAAGGGA
AAGATGCAAAGAACTTTGAAAAGAGAGTTAAACAGGGCTTGAAACCGTTAAGAGTGAAGC
GAAGGAGGTCAGTGATCATGGGGCCAGATCCCCAAGTGCCTGTCATAGGGCACCGTAGCG
TTCTGGTTTCCCGGGGCAGCGCCGGCCGGGGGCGGGGGTTAACAGGGAGGTGGTTGAGGG
ATTCGTTTTCCGCCTGTACTCCTTTATAGCTTGCACCCCGGCCGAGGGATCCGCGACTTG
TCGTGTGAGGGCGCTGTCAAAATGGCCTTCTCCACCCCGTCTTGAAACACGGACCAAGGA
GTCTAACATACTCGCGAGTGTTTGGGTGGTTAAAACCCGAACGCACAATGAAAGTAAACG
GTGGCTCTGCGCCGCTGCTTGGTGGGAACCCTGCTTTTATAATGGGGGGCACTATCACCC

GACCCCGATCCCTTTGGGGTGGTTGGTTTTGAGGTAGAGCGAGTATGATGGGACCCGAAA
GATGGTGAACTATTCCTGAGTAGGGCGAAGCCAGAGGAAACTCTGGTGGAGGCTCGTAGC
GATGCTGACGTGCAAATCGCTCGTCGAACTTGGGTATAGGGGCGAAAGACTAATCGAACC
ATCTAGTAGCTGGTTCCCTCCGAAGTTTCCCTCAGGATAGCGGGAGCAAATGCAAGTCAG
TATCATCCGGTAAAGCGAATGATTAGAGGCCTCGGGGGAGTAACTCCTTCGACCTATTCT
CAAACTTTAAATAGGTGAGGAGCCGGCAACGTTGCTTAAAACAATGAACGCCGTGCGGAC
AATGATTGCTCTCAGTGGGCCGATTTTGGTAAGCAGAACTGGCGATGCGGGATGAACCGA
AAGCGACGCTAAGGTGCCAAACTCCCAGCTCATTCAGACACCAGAAAAGGTGTTGATTCA
TACAGACAGCAGGACGGTGGCCATGGAAGTCGGAATCCGCTAAGGAGTGTGTAACAACTC
ACCTGCCGAATGGATCAGCCCTGAAAATGGATGGCGCTTAAGCGGGGGACCGAGGCGTCG
CCGTTGGGGTAAATACGAAGCCCCAATGAGTAGGAGGGCGTGGGGGTTTTGAAACACGGT
ACACTCGCATGGAAGCCTTGGGCGTAAGCTCGGGTGGAGCGACCCCTAGTGTAGATCTTG
GTGGTAGTAGCAAATATTCAAGTGAGAACCTTGAAGACTGAAGTGGGGAAGGGTTCCACG
CGAACAGTGCTTGGACGTGGGTTAGTCGGTCCTAAGAGATGGGGAAACTCCGTATTCAAG
TGCGCTTTTGTAGCGCAGTCCATCGAAAGGGAAGCGGGTTAACATTCCCGCACCAGGATG
TGGAAACTGTGGTTTACCGCAAACTGCGGCAACGCAACTGAACCTGGGGACGCCCGTAGG
AGCTCCGAGCAGGGATCTCTTTCCTGTTTAACGCGCCCAACCCCCTGGAATCGGTTTGTC
CGGAGATAGGGGGCGAAAGGCCGGAAAAGCACCGCACGTCTTGTGGTGTTCGGAACGCTT
GTAATGGCCCTTGAAAACCCAGGGGAAAGCAGCCGTCTTACACTTGTGTGAGACGAGGAA
TTCTCACACCTGGCCGTACCCATCACCGCATCAGGTCCCCAAGGTGAACAGCCTCTAGTC
AATAGAATAATGTAGATAAGGGAAGTCGGCAAAATAGATCCGTAACTTTGGGAAAAGGAT
TGGCTCTAATGTTCGGGCTTGTCGGGACGCGCGGGCGTAGATGCCAATGAGCGGCAGACG
TCTAGCCTCTTTGGAGGCTGGATTAACGCTGCTTGTGGGAAGAAACCGTGTCGATGGAGG
CAGGAACGCTTATCGATCTTGTCTCTTCCCGCTTGCGATCAACGAACGGATTAGAACTGG
TACGGACAAGGGGAATCCGACTGTTTAATTAAAACAAAGCATTGCGACGGCCGGAAACGG
TGTTGGCGCAATGTGATTTCTGCCCAGTGCTCTGAATGTCAAAGTGAAGAAATTCAACCA
AGCGCGGGTAAACGGCGGGAGTAACTATGACTCTTAAACAAACCATACTACCTACGGGAG
TCAAAAGTGGGGGCGGCGTCTGGTGTAGAAATACACGTACGACACGCTTTTTTTCCTGCT
AGTCGATCTGACCTTTGTTCTTTGGACTGTGGGCGGTCAGCGGCAACATAACCTGGTACG
GGGAAGGCCTCGACGTTGAGTGCTGCATCTTGGAAGTATCTTGACACAATGAGAGAAAAG
AGATTGCGCTTAATAGGGGGTTGATCTCGTGGCGAGCCGACGCCGGTGTGAACTGGTGGC
AATGGCCGTCGTAGAGCACGCAAAGGTATGGGCTGACTTTTTCTCCTCAAAAGTTGGCTT
AAGGGACGTGCCATCCCACACGATAATAAAGTGTGCAGTCTGTAAGAGCTCCCACAGGGC
AAAGACAGACTGGGATGCAGACGGAGACGGCAAAAACAACAACATCCGTTCTGCTTCTGG
GCGCAAATGCTTAAGGTAGCCAAATGCCTCGTCATCTAAACTAATTGGATGAGATAGTCT
GAGTGAAAAGCTTATGCAGGCTAGTACCGGAAAAAAAAAAAAAGAGGGGCTCTGCCACTCT

GACGGTGCGACACTCCTTGTACAGGGAAACCCTCCGCATGGAAAGGCACGTGGTACCACC

CTCGCGGGGAAACTCGCGAGCGGGATCCCGGGTAATGACCGGAGCCGATGGTAAAAACTC

ACGTGACCGATGGGGTGACCCTGTGGCGAGACGCTAAGTTGTGGGACATCACCCTCTCAC

AATACGCATCCGTCGCAGAGACTGAACGGGAGTGGGCTCCGTTGACGGCCGGCCGGCCAG

TTAGCCGTCTTCCAGGGGCTTAAGATACAGTCCGGGCCTCAGGGATGGAAAAACCTGGGG

AACTACCAGAATTAGTGACGCGCATGAATGGATTAACGAGATTCCCACTGTCCCTATCTA

CTATCTAGCGAAACCACAGCCAAGGGAACGGGCTTGGAAGAATCAGCGGGGAAAGAAGAC

CCTGTTGAGCTTGACTCTAGTCCGACTTTGTGAGATGAGTTGGGAGGTGTAGTCTAAGAG

GGAGGGGCAACTCGACCGTGAAATACCTCTACTCCCAAGCTTGTTTTACTTACTCCGGCG

AGCAGGGATGGGTGGTAACACCCTTTTTCTAGGATTAACACCTCTTCGGGGGTGGACCTG

CGCGGAGGACATTGTCAGGTGGGGAGTTTGGCTGGGGCGGCACATCTGTTAAAAGATAAC

GCAGGTGTCCTAAGATGAGCTCAGTGAGAACAGAAATCTCACGTAGAACAAAAGGGTAAA

AGCTCATTTGATTTTGATTTTCAGTACGAATACAAACTGTGAAAGCATGGCCTATCGATC

CTTTTGACCTTCAAGGTTTGAAGCAAGAGGTGTCAGAAAAGTTAACACGCGTTGTGGTAG

CTTTTAAGAGTGGACTCAAAGTGTTCCGCTAGTCCCCCCTCGGACCGCCACTTACAAAAG

GCCTGGGGATGGGGGCGACACTGTCAAATTGCGGGGACTCCCTAAAGCCTGTCTCTACCA

AGCCAGCGCGGCGAAAGCCCCTGGTGGCCGGGGTAATGACCTAGGGTATGGTAAAAGCGA

GACAGGATGGAACAATGGGCAATCCGCAGCCAAGTCCTAAGGCTTTTTGAAAAGCGGCTA

TGGATGCAGTTCAACGACTAGATGGCAGTGGGCTGAGAGAGCCCACCGCGGCACCCTTTC

GGCTTAAGATATAGTCTTGCCCACGTGAAAGCGTGCCTGCAGATGAACGCCACCATGTAT

GGCCGGAGCTGCAGGAGACCGCACTTGATGTGCAGTCGGGAATCGACAGTACCACAGGGA

TGAACAAAAACACACAATTAATTTGTGGCCATCATACAGTCGTATGATATTATCGTCCCA

GGCGGGTGCGCCTCTCTCACCCGTCTAGTCCGGCCTTGGTGCGCAAAAAATGCTTGCGTG

GATGCGCCTTGGTTAAGGAAAATGCTCCTAGTAGTTTTACAAGGACACTTCTTGTAGGGC

AACACATCTGGATGCGAAGAACTCCCGGAAAAGACAACGGTACCAAGTGACTGGGGTAAC

GCCCCATTCATGAAGGCAAATAATCAAAAAGAGGTGCCGGGTAAGAACCCGTTGTATAGG

GACAATCCGCAGCTGACTTGGCTCCCTGCCCATCCATGGGCATAGCTTAGCGGTTCAACG

CTCGGTAAGATGTGGGTTGGCGCGCTGTCCGACAGCGCACTGGCTTAAGGTACGGGCTAC

TCCCACCGGAGACGGTGTCTTTGTCTACCTCGGGTAACACGCCTCGGGAGAAGCAAAGAG

GGTGCAATGGAAAAAAAACTGGTTGCGGGGAAATCCCTACCAGAGTCACCTGGTACAAAG

GAACTGGCTTGTGGCAGCCAAGCGTTCATAGCGACGTTGCTTTTTGATCCTTCGATGTCG

GCTCTTCCTATCATTGCGAAGCAGAATTCGCCAAGTGTTGGATTGTTCACCCACTAATAG

GGAACGTGAGCTGGGTAGCCACTTTATGCCCAACAGCTGGATAGCTGGGCTTCTGTCTTA

GGCCGGGACTTCTCACGACCCGTGCCTTGGTCACCCAAAAAAGGGCGACCGCAGAGCTCA

GGATGGAGCTTCTGGCTTAACTAGTAGAGGACAGATAGAGATCGTCAAATTGCGGGAAAC

CCCTTAGAGCCTCGAGCTACCAAGTGCGTGTCGAAAGGCATAGACGCAGCCAGCGATGGC

TGTCGCATGGCTTGATTCCATCAAGATAGGTAAAAATGCTCGAGGATTGGGCAATCCGCA
GCCAAGCCCCTAAGGTCCCCCCGCGGGCTATGGGGAAGGTTCAACGACTGTAAGGCGTAC
CGTGCAAGCGGAAGATACAGTCTAGCCCCGCTGGAAAGCAGCGCCAAAGTAAGTAGGTTG
CAGTGGGTCTGTCCTC
>SRR9587966
GCATGTTTTGGTATGTGTTTCGACAGTCATGCATGTCATAGCCGTAGGATGGGGAGTGAG
CTTCATAGCCCGACCCCTTCTCAGGAGCATGTTTTCATAGCATATTGAGCATACTGGGGT
GGTCTCCCATGTAGCTCGCTCTCTCGGTGCAGGACCTCCCTTTTGCGGGCCACTTCGATG
GCCTCCATTATGGAGAGCATGTTTTGGCATATGAAGCGTTGGTGGTGAGGCTCGTAGCCT
GTATATACATACCAACGCAGTCGTCATATGTCTCTCTGGCGGGGGTCGGTCGTATGACCT
TGTCCCTACCTGGAGCATGTATTTCGTATGATATACTGGACGGCAGCTGCGCTGTTCAGT
CGTCGGCCGTACCTCATGTGCGCGTTTCCAGGTCCGAACGTTTGACATGCAAGGGCATCA
TACCAGAAGCCTGTTTTCTCGTATGAGCATGTAGGTAGGCAGGCTCTTCAGTCGTCTACC
CATAAATGCACTTCGAAGGAACTTTTTGGTATTGAATGGCTTTGTGCCCAGACGTTCTTG
TGAGCTTGTTCTCGCTCTGCGATACGCTGGAAGCTGTGGTATTACCTACAGCAGACGCAT
TCCTAGGGCGGACGGCCTCTCTGCAGATGGGCACATCTGTAGAAAGAGGCTCCCCTTGAT
GACGCAATGCAATGGCTGGGTAAACGGAGTGCTATACAGAAGGGTCAAACCTCTGTGGAA
GTACACTCATACCAGCATTGTTTGTTGAGAGGGGTCGGCTCTACGAGTGAGGACCTATAC
TGCCATCTCATTCAGATGCGTAGACGGTCTTTACTCAGACTGCCTTTCTGGCTGATTGTC
CTCGGTCCCTCGTGCGCACGCGCCGCTGGCTGAATGGGCTTGTCCTTATGGAGTGCCGCG
TTCTTCAAAGAATCCGTCTTCATACAGCGACCTCATTTACATATTGCCGGCTGAATGGGC
TTGTCCTTATGGAGCACTGCATTCTTCTAGGAATTGCTATCTATACAGCGACCTCATTTA
CATACAAGGCACCTTTTTCTACATAGACTTGCCATTTGGCTGATTGTTCTGACAGTCGCT
GCCTCTTCGGAGTTCTGTGTGGACGAGTGGCCTCCTGAGTGGGCCGGTCCTTACGGAGTG
CCCACGTCTGTTATGACTGGGTTCTCCATACAGCGACCTCACTCAGAATTGTATGGCACC
GTGCCGATTGAATGGACAAGCAGTCCTTAGGGAATCGCATGAAAGCATGATGGCACATGG
TGTATGGACCCGTTGGGCCTTGCCTGTGCTATCTTCTCGAGTCGACCGGTCCTCGTCTCC
TCCTTTCGCGACGGGAAGCTCTTGAGTTTACCGTGGGTCAGGACGCTGTGGTATCTGTTT
GTCTGTAGTGTGGGCTCTCTATCAGAACAGCGTATCAGCACACGTCGTCTGGGTCTATGG
TATTTTTGCTTAACGGCCCTGGGTTTGGTAGTATGGTTCCATGTCGGTCCTCTCACCTCC
CATCCGTCATGTCTCTGATATCATCTGCACAGTACGTCTTGTCAGAATATCATTCTGTAT
GGCAGTCGATAGACTAGGCCCTATGCCGCAGCTGTCTCTGACCACTATGCGAGCGCATAC
CTACCTCAATGTGTTCATAGCTGTTATGGATCTTGGGTGTGGTGTGCCAGAACGATACGC
CACCAGATTGTGGAATGGTGCTTGTAGTATTAGAAGAGGACATGTATGAAGAGCACGTTC
TTTTCTCTTATGATGGTGCCGCGGGTGGGGGCCGTTCTCATTCTGCTTGTCGGGTGCTCA
AGGTGCATGTAGGATTGCTGAACCCACTGAATGGTCATGAGAATGGTCATATGGTGAGGT

TCACTTCTCCATCGCTTTGTGTGTGTTGGATGTTTGGGTGGCTTCGGCCGCATGGAAAAT
CTCGGTCTGTAACTCATCAGGGAGGGTTTCCATGTGAGCAATCTCTCCAGTCGAACACAC
ACCCATGGGCGTCCCGCTAGGCAAAATGGAACTCTCGATCCGACACAGCCTGTGGAGTGT
CTGTTCCGTGTGGCGTGAAACGGCTTATACTGGATGTCTTCCCTTTGGACACCAGTCTGT
TGGTATGTACGCATACCATCTAGAGGGAGCAAGAAAAAGGCATGCAAGTGCCACAAAGGA
AGGGAGGCCTCCCTGCTACCGTGTGTCCTCTTGACGATCACGGGGGAGTATATAGAGACA
TTTGGACGAGATCCTTTTTTCACGAGGCGGTGCCCTCGCGCAAAGGTGTTGCTCTGCACG
CATCTTTATGATGATAGTTGCTCGCTCCTATGCGTCCACTCGTGAAAATACCTGCTCTCT
CTGTCTGTTTCCATGACACCCCCATGCGTTGTCATAGAGCACGCTCAGGTACAGAAAGTG
CTGTGACTTTCTTGGCCTGCTGTCACTGAGTCAGCCTGATACTTTGGAGGAATCTTTTCT
GGGCGCTGCCCAGCTGAGAGTTCTTATCAGGCTCTGGAGTGTGCATATGGCTATAATCAT
AATAGTGCATGTGGCCACGCAGAAAAAGCATCGACGTGGAATGGAGAAGAACACCATTCT
GAATGGAACGGTTCTTACTCTGCTTCCGTTGCTTGACGCTGTATGGGAATCGCTTTGCCC
TGGGGGTACACCTTCCATTCCATGACAAAGCATTTGATCTATGCCCCGAGGGTCATTCTC
TATGAAGAGCCCGAAACTTGTCTGAGTGAGAGGCGAAGGTAGCCCTTCATTTATGCGGGG
TCCGCCCTCACACTGTAGTCGGCACAGTCCCTTGTTTATGGACTGCGCAGCTCAATCTTC
GACTGGAGAGGAAAAACAAAACACAAGTATGACACGCACATACGTGTTTTCTAGGACTGT
GACCTGTCGGCTGCCTCGCGGTGGTTGGCATGATGCCCCAAGTGGGGGGGGGGAGAAACA
ATCAGTATGATTGATAAAAACCAGTGGGGAAAAAGGCGCGCTGTCGCACACCCTTGGTCA
CGGCCTCAAGTCGACGGCCCTCACTGGCCCACCCTGCGACTCCTGTTCGCTTTTGACAAG
ACACACCTGCAATACGGAGGAAATGCCTGAGCAGCTGTGCCGCAGCAACTGCGGACGCGT
AGCGATATGCATCCCTGTTGTTTGCGACTTGGTACAAAGAACGTGCGCCGGGAGACGGAT
TGCCCTTTGACTTTTCCACCCAGCCCGCGGCTCTATACCCTTGACGTACCTATTTACCTG
GTTGATCCTGCCAGTAGTCATATGCTTGTCTCAAAGACTAAGCCATGCATGTCTAAGTAT
AAACACTTTTATACTCGTGAAACTGCGAATGGCTCATTAAAACAGTTATAGTTCCTTTGG
GAACAACACTACTTGGATAACCGTAGTAATTCTAGAGCTAATACATGCCTCAACGCCCGA
CTCACGAAGGGTGGTATTTATTGGATAAAAAACCATCGTGCTCTGTCCAGAGCGCTTTGA
GATGATTCACAATAACTTGTCGGATCGCATGGCCTCGTGCTGGCGACTCCTCATTCAAAT
TTCTGCCCTATCAACTTTCGATGGTAGAGTATTGGTCTACCATGGTGTCGACGGGTGACG
GGGAATTAGGGTTCGATTCCGGAGAGGGAGCCTGAGAAACGGCTACCACATCCAAGGAAG
GCAGCAGGCGCGCAAATTACCCAATCCCGACTCGGGGAGGTAGTGACAAAAAATAACAAT
AGGGGGCCCTTTGGGTCTTCTAATTGGAATGAGAACAATTTAAATCCCTTATCGAGGATC
CATTGGAGGGCAAGTCTGGTGCCAGCAGCCGCGGTAATTCCAGCTCCAATAGCGTATATT
AAAGTTGTTGCAGTTAAAACGCTCGTAGTCGGACCTCGGAGCGCCCCCACGGGCCGGGCT
TTTGCCTTGTTGCCAGTGGGAGTTTGTCTCCTTTTGTCGGGACGCGGTGTGCAGGGCCTT
TACTGTCTCTGCGCGTTTACCTGCGGGCCGACCGTTTACTGTGAAGAAAGTAAAGTGTTC

AAAGCAGGCCAATGCCTTGAATATGTGAGCATGGAATAATAGAATAGGACTTGGGCTCTA

TTTTGTTGGTTTCCAGTGACCAAGTAATGATTAATAGGGACGGTTGGGGGCATTCGTATT

TCATTGTCAGAGGTGAAATTCTTGGATTGATGGAAGACGTACAACTGCGAAAGCATCTGC

CATGGATGTTTTCATTGATCAAGAACGAAAGTTAGGGGATCGAAGACGATCAGATACCGT

CGTAGTCTTAACCATAAACGATGCCGACTGGGGATTGGCGGGGGCACTTTTATATGACTC

TGTCAGCACCCTGAGGGAAACCAAAGTCTTTGGGTTCTGGGGGGAGTATGGTCGCAAGGC

TGAAACTTAAAGGAATTGACGGAAGGGCACCACCAGGAGTGGAGCCTGCGGCTTAATTTG

ACTCAACACGGGAAAACTTACCAGGTCCGGACAGAAGAATGATTGACAGACTGAAAGCTC

TTTCTTGATTTTTTGGTTGGTGGTGCATGGCCGTTCTTAGTTGGTGGAGTGATTTGTCTG

GTTAATTCCGTTAACGAACGAGACCTCGTCCTGCTAAATAGGTGCGCGCATGCCAGCACT

GCGTTCTTACCTTCTTAGAGGGACTATGCGCGTCTAGCGTATGGAAGATTGAGGCAATAA

CAGGTCTGTGATGCCCTTAGATGTTCTGGGCCGCACGCGCGCTACACTGATGCATTCAAC

GAGTTTCACTTGATAGTCCTGGGTCGGAAGGCTCGGGTAATCTTTTGAAAGTGCATCGTG

CTGGGGATAGATCATTGCAATTATTGATCTTCAACGAGGAATTCCTTGTAGGCGCAGGTC

ATCAGCCTGCGCCGAATACGTCCCTGCCCTTTGTACACACCGCCCGTCGCTCCTACCGAT

TGAATGGTCCGATGAAATGTCCGGATCGCGGCTAGTGTGATGGACTCGTCCAGAGCGGTA

GCCGTGAGAAGCTCATTAAATCTTATCATTTAGAGGAAGGAGAAGTCGTAACAAGGTTTC

CGTAGGTGAACCTGCGGAAGGATCATTCACAAACTATTGACACAAACACACGCGAACCAA

ATCGTCCGCACAGCAGGTGGCAATGAAAGAGAGACTCTGCATGTCGCCTTTCGGGGTAGC

AAGCAGCACTCTTCTGCCATCGCCTCTGTGCAGGGCGTAAATTCCCATTGAGAGGATGTG

AGGGCACCACAGGAAGCTTTTCCACAGGAAGTCGCCATCCTTCTCCCTCCACGGCGGCGC

TTCTGTGCTTGGCAGTTTTTTTTTGCCTTCTAGGGAGGATGCCGCCAATGGAGCCCCTTA

TATATATACATCATCATATGCCCCTTTTTTTCTTAACCGCTTGCCACAGCTTCTTCTATG

AGGAGCTTGTGGGAAGACTGTCTCCATACAATAACAAAGATACAACTCTTAGCGGTGGAT

ATCTTGGTTCTCGCAACGATGAAGAACGCAGCCAACTGCGATAACTAATGTGAATTGCAG

GACTTCGTGAATCATTAAGTCTTTGAACGCAAGTTGCGCTCATCTCCATGAGGAGGTGAG

CATGCCTGTTTGAGTGTCGATGGGAAAAACAATACATAACAACAATCGGCTCTCTTATCT

TCCTGCCTTCTTGGGGGCAAAAGGTAGTGCATGCCGGAGATGGGCTTTGTCGCCCCTTTC

TCTTGAAAGAGGCGTCGTCCGAAACACGTATCGTTGCCCGCTGGAGGTCATAGAGCATGA

GTTTAAAGCACCACATCTCTCAAGTCAGAGTGAGCGTCTTGAGTAGGTCCGCATTGATTC

AGTTTCAACCCAGGTGGCGTTGGTTACTTGCCATAAGCCTCTCTGGGGTTTCTTTGCTTG

GACCTCTCTTGTCAGTTTCTCTGACGTAGAGACAGGTGCCGTCTCATGGACGTTGCCCTC

TGGAACGTGCTAAAAAAAAAAAAAAAGAAAAAACCGTACCACATGTCAAAAGAGGCAGATC

GTGCTTGACGGCGGGAGGTGTTGCATTTCTTGCTTGTCCTGAGCGTCTGCCAGTCTCTGA

CGAGTGTGTTCCTTCCATAAACCCGATCTCAAATCAGGTAGGACCACCCGCTGAATTTAA

GCATATAATTAAGCGGAGGAAAAGAAACTAACAAGGATTCCCTCAGTAGCGGCGAGCGAA

GCGGGAAGAGCTCAACATGAAAACCGTTGGGGGTTCGCGCTTTCCTTCGGGTTGTATAGT
AAAGTGGTGTCGCCTGCGTCGCGCCGGGTATAAATCCCGTCTGAAATGGGGTGTCGTAGA
GGGTTATATCCCCGTCTGTGACCCGGTGTGCGATGCGTTACGGTGACCTTACGATGAGTC
GGGTTGCTTGGGCATGCAGCCCTAAGTGGGTGGTAAACTTCATCTAAAGCTAAATACTAG
CGAGAGACCGATAGCGTACAAGTACCGTAAGGGAAAGATGCAAAGAACTTTGAAAAGAGA
GTTAAACAGGGCTTGAAACCGTTAAGAGTGAAGCGAAGGAGGTCAGTGATCATGGGGCCA
GATCCCCAAGTGCCTGTCATAGGGCACCGTAGCGTTCTGGTTTCCCGGGGCAGCGCCGGC
CGGGGGCGGGGGTTAACAGGGAGGTGGTTGAGGGATTCGTTTTCCGCCTGTACTCCTTTA
TAGCTTGCACCCCGGCCGAGGGATCCGCGACTTGTCGTGTGAGGGCGCTGTCAAAATGGC
CTTCTCCACCCCGTCTTGAAACACGGACCAAGGAGTCTAACATACTCGCGAGTGTTTGGG
TGGTTAAAACCCGAACGCACAATGAAAGTAAACGGTGGCTCTGCGCCGCTGCTTGGTGGG
AACCCTGCTTTTATAATGGGGGGCACTATCACCCGACCCCGATCCCTTTGGGGTGGTTGG
TTTTGAGGTAGAGCGAGTATGATGGGACCCGAAAGATGGTGAACTATTCCTGAGTAGGGC
GAAGCCAGAGGAAACTCTGGTGGAGGCTCGTAGCGATGCTGACGTGCAAATCGCTCGTCG
AACTTGGGTATAGGGGCGAAAGACTAATCGAACCATCTAGTAGCTGGTTCCCTCCGAAGT
TTCCCTCAGGATAGCGGGAGCAAATGCAAGTCAGTATCATCCGGTAAAGCGAATGATTAG
AGGCCTCGGGGGAGTAACTCCTTCGACCTATTCTCAAACTTTAAATAGGTGAGGAGCCGG
CAACGTTGCTTAAAACAATGAACGCCGTGCGGACAATGATTGCTCTCAGTGGGCCGATTT
TGGTAAGCAGAACTGGCGATGCGGGATGAACCGAAAGCGACGCTAAGGTGCCAAACTCCC
AGCTCATTCAGACACCAGAAAAGGTGTTGATTCATACAGACAGCAGGACGGTGGCCATGG
AAGTCGGAATCCGCTAAGGAGTGTGTAACAACTCACCTGCCGAATGGATCAGCCCTGAAA
ATGGATGGCGCTTAAGCGGGGGACCGAGGCGTCGCCGTTGGGGTAAATACGAAGCCCCAA
TGAGTAGGAGGGCGTGGGGGTTTTGAAACACGGTACACTCGCATGGAAGCCTTGGGCGTA
AGCTCGGGTGGAGCGACCCCTAGTGTAGATCTTGGTGGTAGTAGCAAATATTCAAGTGAG
AACCTTGAAGACTGAAGTGGGGAAGGGTTCCACGCGAACAGTGCTTGGACGTGGGTTAGT
CGGTCCTAAGAGATGGGGAAACTCCGTATTCAAGTGCGCTTTTGTAGCGCAGTCCATCGA
AAGGGAAGCGGGTTAACATTCCCGCACCAGGATGTGGAAACTGTGGTTTACCGCAAACTG
CGGCAACGCAACTGAACCTGGGGACGCCCGTAGGAGCTCCGAGCAGGGATCTCTTTCCTG
TTTAACGCGCCCAACCCCCTGGAATCGGTTTGTCCGGAGATAGGGGGCGAAAGGCCGGAA
AAGCACCGCACGTCTTGTGGTGTTCGGAACGCTTGTAATGGCCCTTGAAAACCCAGGGGA
AAGCAGCCGTCTTACACTTGTGTGAGACGAGGAATTCTCACACCTGGCCGTACCCATCAC
CGCATCAGGTCCCCAAGGTGAACAGCCTCTAGTCAATAGAATAATGTAGATAAGGGAAGT
CGGCAAAATAGATCCGTAACTTTGGGAAAAGGATTGGCTCTAATGTTCGGGCTTGTCGGG
ACGCGCGGGCGTAGATGCCAATGAGCGGCAGACGTCTAGCCTCTTTGGAGGCTGGATTAA
CGCTGCTTGTGGGAAGAAACCGTGTCGATGGAGGCAGGAACGCTTATCGATCTTGTCTCT
TCCCGCTTGCGATCAACGAACGGATTAGAACTGGTACGGACAAGGGGAATCCGACTGTTT

AATTAAAACAAAGCATTGCGACGGCCGGAAACGGTGTTGGCGCAATGTGATTTCTGCCCA
GTGCTCTGAATGTCAAAGTGAAGAAATTCAACCAAGCGCGGGTAAACGGCGGGAGTAACT
ATGACTCTCTTAGACAGATCAAGACACCTTCAATAAATTGGTTCATTATGTCGGGTGGTC
CGTCGTCCGGGAAGCGACCAAAGCAAAACCCCGTAGACTGTGATGTGCATCTGTCAGACG
AAACTGACGTCCGTGTTCATCAAGCCTTAGCTATCTTTCAACGCCGGCTTCGAGACGTGG
CGTCTCTTAAGGTGTTTGAAGTGGCTCTCTCTGTGGGAACTACTGATGCACTGGTAGAGA
TGGGCACTATCACTTGCTGGAGTATTGATGCTGTTGGGAAAGGTTTCTCAAGGAACAGCA
GCGGTCGCTGGCAATTTGACCTCACTGGCAAGATTAATGGCAAGCCTCTAATATCCACCA
CAAAGGATCGTTGGAGGCCGGTTGTGACGGCTGTGCCCTACTACCTTTCGTGCGCCATCG
ATGGGGACAAGACGACGCTGGAGGAAAAGCAGTGGACTTCATCACACATGTGTCACCATG
AATGGTGTGTGAACGTGGAGCACATTGTTTATGAAAGCCTCGACGTCAACAAGTCTCGGA
ACTGGTGCCAGGGAAAAAACGCATGCCTCCATCTGCCTGTTTGCATTCGAACCGGCCGCC
GTGTGCAGGAGCTCAGGAAGAGCAATGGGGTTCGTGCAATGGACCTCCTCCTGCAGAAAT
TGGCTGGATTGGAGATTGAGGTTGAGGAGCCCGCTGCAGCTGTACTCCGAGAGTCTGTCG
AACGGGAGGACGCCTAGTGAGTGACTCGTGACACGGGTTGGCTGCGCATGATGCGTCAAC
TACATGGACATCTTTTGAGAGAGATAGGCGGGCTGAAAAGGTTATAGCCGTCTAGTCTCG
CTCTCTTTCATTGAGCGGGGCGACACACCTGGTTGCGGGAACCCTGCCCACCTTTCTATC
AAGAAGTGGGAACCAGTCCATTCCTACTCCTGCTTGTGAGGCATATAGTAACGTAGCAGC
AGGAGGATCTCGGCGAGAGGCCGAACCCGAAAGGAATAACGTGGGCTGGGAGGCGATCCG
CAGGCGAGTCCGTAACTATATGATTATCGACGGGCCGTCTCAACGACTGGTAAGGTGTGG
GTTTGTCCTCCTCCTCACGTGGGGACAGGCTTAAGGTACAGTCTAATCCCACCCGCCAAG
GGTGCTTGGGTGTAATCCCTCTTCCATAGAGGGCACTAGCCCAGGGGCTGGGACGCTGGG
AGGAAATGCCCCGTCCCTATAGCCGGTAGAAATGAAGGTAGCCAAATGCCTCGTCATCTA
ATTAGTGACGCGCATGAATGGATTAACGAGATTCCCACTGTCCCTATCTACTATCTAGCG
AAACCACAGCCAAGGGAACGGGCTTGGAAGAATCAGCGGGGAAAGAAGACCCTGTTGAGC
TTGACTCTAGTCCGACTTTGTGAGATGAGTTGGGAGGTGTAGTCTAAGAGGGAGGGGCAA
CTCGACCGTGAAATACCTCTACTCCCAAGCTTGTTTTACTTACTCCGGCGAGCAGGGATG
GGTGGTAACACCCTTTTTCTAGGATTAACACCTCTTCGGGGGTGGACCTGCGCGGAGGAC
ATTGTCAGGTGGGGAGTTTGGCTGGGGCGGCACATCTGTTAAAAGATAACGCAGGTGTCC
TAAGATGAGCTCAGTGAGAACAGAAATCTCACGTAGAACAAAAGGGTAAAAGCTCATTTG
ATTTTGATTTTCAGTACGAATACAAACTGTGAAAGCATGGCCTATCGATCCTTTTGACCT
TCAAGGTTTGAAGCAAGAGGTGTCAGAAAAGTTAACACGCGTTGTGGTAGCTTTTAAGAG
TGGACTCAAAGTGTTCCGCTAGTCCCCCCTCGGACCGCCACTTACAAAAGGCCTGGGGAT
GGGGGCGACACTGTCAAATTGCGGGGACTCCCTAAAGCCTGTCTCTACCAAGCCAGCGCG
GCGAAAGCCCCTGGTGGCCGGGGTAATGACCTAGGGTATGGTAAAAGCGAGACAGGATGG
AACAATGGGCAATCCGCAGCCAAGTCCTAAGGCTTTTTGAAAAGCGGCTATGGATGCAGT

TCAACGACTAGATGGCAGTGGGCTGAGAGAGCCCACCGCGGCACCCTTTCGGCTTAAGAT
ATAGTCTTGCCCACGTGAAAGCGTGCCTGCAGATGAACGCCACCATGTATGGCCGGAGCT
GCAGGAGACCGCACTTGATGTGCAGTCGGGAATCGACAGTACCACAGGGATGAACAAAAA
CACACAATTAATTTGTGGCCATCATACAGTCGTATGATATTATCGTCCCAGGCGGGTGCG
CCTCTCTCACCCGTCTAGTCCGGCCTTGGTGCGCAAAAAATGCTTGCGTGGATGCGCCTT
GGTTAAGGAAAATGCTCCTAGTAGTTTTACAAGGACACTTCTTGTAGGGCAACACATCTG
GATGCGAAGAACTCCCGGAAAAGACAACGGTACCAAGTGACTGGGGTAACGCCCCATTCA
TGAAGGCAAATAATCAAAAAGAGGTGCCGGGTAAGAACCCGTTGTATAGGGACAATCCGC
AGCTGACTTGGCTCCCTGCCCATCCATGGGCATAGCTTAGCGGTTCAACGCTCGGTAAGA
TGTGGGTTGGCGCGCTGTCCGACAGCGCACTGGCTTAAGGTACGGGCTACTCCCACCGGA
GACGGTGTCTTTGTCTACCTCGGGTAACACGCCTCGGGAGAAGCAAAGAGGGTGCAATGG
AAAAAAAACTGGTTGCGGGGAAATCCCTACCAGAGTCACCTGGTACAAAGGAACTGGCTT
GTGGCAGCCAAGCGTTCATAGCGACGTTGCTTTTTGATCCTTCGATGTCGGCTCTTCCTA
TCATTGCGAAGCAGAATTCGCCAAGTGTTGGATTGTTCACCCACTAATAGGGAACGTGAG
CTGGGTAGCCACTTTATGCCCAACAGCTGGATAGCTGGGCTTCTGTCTTAGGCCGGGACT
TCTCACGACCCGTGCCTTGGTCACCCAAAAAAGGGCGACCGCAGAGCTCAGGATGGAGCT
TCTGGCTTAACTAGTAGAGGACAGATAGAGATCGTCAAATTGCGGGAAACCCCTTAGAGC
CTCGAGCTACCAAGTGCGTGTCGAAAGGCATAGACGCAGCCAGCGATGGCTGTCGCATGG
CTTGATTCCATCAAGATAGGTAAAAATGCTCGAGGATTGGGCAATCCGCAGCCAAGCCCC
TAAGGTCCCCCCGCGGGCTATGGGGAAGGTTCAACGACTGTAAGGCGTACCGTGCAAGCG
GAAGATACAGTCTAGCCCCGCTGGAAAGCAGCGCCAAAGTAAGTAGGTTGCAGTGGGTCT
GTCCTC
>SRR9587967
GCATGTTTTGGTATGTGTTTCGACAGTCATGCATGTCATAGCCGTAGGATGGGGAGTGAG
CTTCATAGCCCGACCCCTTCTCAGGAGCATGTTTTCATAGCATATTGAGCATACTGGGGT
GGTCTCCCATGTAGCTCGCTCTCTCGGTGCAGGACCTCCCTTTTGCGGGCCACTTCGATG
GCCTCCATTATGGAGAGCATGTTTTGGCATATGAAGCGTTGGTGGTGAGGCTCGTAGCCT
GTATATACATACCAACGCAGTCGTCATATGTCTCTCTGGCGGGGGTCGGTCGTATGACCT
TGTCCCTACCTGGAGCATGTATTTCGTATGATATACTGGACGGCAGCTGCGCTGTTCAGT
CGTCGGCCGTACCTCATGTGCGCGTTTCCAGGTCCGAACGTTTGACATGCAAGGGCATCA
TACCAGAAGCCTGTTTTCTCGTATGAGCATGTAGGTAGGCAGGCTCTTCAGTCGTCTACC
CATAAATGCACTTCGAAGGAACTTTTTGGTATTGAATGGCTTTGTGCCCAGACGTTCTTG
TGAGCTTGTTCTCGCTCTGCGATACGCTGGAAGCTGTGGTATTACCTACAGCAGACGCAT
TCCTAGGGCGGACGGCCTCTCTGCAGATGGGCACATCTGTAGAAAGAGGCTCCCCTTGAT
GACGCAATGCAATGGCTGGGTAAACGGAGTGCTATACAGAAGGGTCAAACCTCTGTGGAA
GTACACTCATACCAGCATTGTTTGTTGAGAGGGGTCGGCTCTACGAGTGAGGACCTATAC

TGCCATCTCATTCAGATGCGTAGACGGTCTTTACTCAGACTGCCTTTCTGGCTGATTGTC
CTCGGTCCCTCGTGCGCACGCGCCGCTGGCTGAATGGGCTTGTCCTTATGGAGTGCCGCG
TTCTTCAAAGAATCCGTCTTCATACAGCGACCTCATTTACATATTGCCGGCTGAATGGGC
TTGTCCTTATGGAGCACTGCATTCTTCTAGGAATTGCTATCTATACAGCGACCTCATTTA
CATACAAGGCACCTTTTTCTACATAGACTTGCCATTTGGCTGATTGTTCTGACAGTCGCT
GCCTCTTCGGAGTTCTGTGTGGACGAGTGGCCTCCTGAGTGGGCCGGTCCTTACGGAGTG
CCCACGTCTGTTATGACTGGGTTCTCCATACAGCGACCTCACTCAGAATTGTATGGCACC
GTGCCGATTGAATGGACAAGCAGTCCTTAGGGAATCGCATGAAAGCATGATGGCACATGG
TGTATGGACCCGTTGGGCCTTGCCTGTGCTATCTTCTCGAGTCGACCGGTCCTCGTCTCC
TCCTTTCGCGACGGGAAGCTCTTGAGTTTACCGTGGGTCAGGACGCTGTGGTATCTGTTT
GTCTGTAGTGTGGGCTCTCTATCAGAACAGCGTATCAGCACACGTCGTCTGGGTCTATGG
TATTTTTGCTTAACGGCCCTGGGTTTGGTAGTATGGTTCCATGTCGGTCCTCTCACCTCC
CATCCGTCATGTCTCTGATATCATCTGCACAGTACGTCTTGTCAGAATATCATTCTGTAT
GGCAGTCGATAGACTAGGCCCTATGCCGCAGCTGTCTCTGACCACTATGCGAGCGCATAC
CTACCTCAATGTGTTCATAGCTGTTATGGATCTTGGGTGTGGTGTGCCAGAACGATACGC
CACCAGATTGTGGAATGGTGCTTGTAGTATTAGAAGAGGACATGTATGAAGAGCACGTTC
TTTTCTCTTATGATGGTGCCGCGGGTGGGGGCCGTTCTCATTCTGCTTGTCGGGTGCTCA
AGGTGCATGTAGGATTGCTGAACCCACTGAATGGTCATGAGAATGGTCATATGGTGAGGT
TCACTTCTCCATCGCTTTGTGTGTGTTGGATGTTTGGGTGGCTTCGGCCGCATGGAAAAT
CTCGGTCTGTAACTCATCAGGGAGGGTTTCCATGTGAGCAATCTCTCCAGTCGAACACAC
ACCCATGGGCGTCCCGCTAGGCAAAATGGAACTCTCGATCCGACACAGCCTGTGGAGTGT
CTGTTCCGTGTGGCGTGAAACGGCTTATACTGGATGTCTTCCCTTTGGACACCAGTCTGT
TGGTATGTACGCATACCATCTAGAGGGAGCAAGAAAAAGGCATGCAAGTGCCACAAAGGA
AGGGAGGCCTCCCTGCTACCGTGTGTCCTCTTGACGATCACGGGGGAGTATATAGAGACA
TTTGGACGAGATCCTTTTTTCACGAGGCGGTGCCCTCGCGCAAAGGTGTTGCTCTGCACG
CATCTTTATGATGATAGTTGCTCGCTCCTATGCGTCCACTCGTGAAAATACCTGCTCTCT
CTGTCTGTTTCCATGACACCCCCATGCGTTGTCATAGAGCACGCTCAGGTACAGAAAGTG
CTGTGACTTTCTTGGCCTGCTGTCACTGAGTCAGCCTGATACTTTGGAGGAATCTTTTCT
GGGCGCTGCCCAGCTGAGAGTTCTTATCAGGCTCTGGAGTGTGCATATGGCTATAATCAT
AATAGTGCATGTGGCCACGCAGAAAAAGCATCGACGTGGAATGGAGAAGAACACCATTCT
GAATGGAACGGTTCTTACTCTGCTTCCGTTGCTTGACGCTGTATGGGAATCGCTTTGCCC
TGGGGGTACACCTTCCATTCCATGACAAAGCATTTGATCTATGCCCCGAGGGTCATTCTC
TATGAAGAGCCCGAAACTTGTCTGAGTGAGAGGCGAAGGTAGCCCTTCATTTATGCGGGG
TCCGCCCTCACACTGTAGTCGGCACAGTCCCTTGTTTATGGACTGCGCAGCTCAATCTTC
GACTGGAGAGGAAAAACAAAACACAAGTATGACACGCACATACGTGTTTTCTAGGACTGT
GACCTGTCGGCTGCCTCGCGGTGGTTGGCATGATGCCCCAAGTGGGGGGGGGGAGAAACA

ATCAGTATGATTGATAAAAACCAGTGGGGAAAAAGGCGCGCTGTCGCACACCCTTGGTCA
CGGCCTCAAGTCGACGGCCCTCACTGGCCCACCCTGCGACTCCTGTTCGCTTTTGACAAG
ACACACCTGCAATACGGAGGAAATGCCTGAGCAGCTGTGCCGCAGCAACTGCGGACGCGT
AGCGATATGCATCCCTGTTGTTTGCGACTTGGTACAAAGAACGTGCGCCGGGAGACGGAT
TGCCCTTTGACTTTTCCACCCAGCCCGCGGCTCTATACCCTTGACGTACCTATTTACCTG
GTTGATCCTGCCAGTAGTCATATGCTTGTCTCAAAGACTAAGCCATGCATGTCTAAGTAT
AAACACTTTTATACTCGTGAAACTGCGAATGGCTCATTAAAACAGTTATAGTTCCTTTGG
GAACAACACTACTTGGATAACCGTAGTAATTCTAGAGCTAATACATGCCTCAACGCCCGA
CTCACGAAGGGTGGTATTTATTGGATAAAAAACCATCGTGCTCTGTCCAGAGCGCTTTGA
GATGATTCACAATAACTTGTCGGATCGCATGGCCTCGTGCTGGCGACTCCTCATTCAAAT
TTCTGCCCTATCAACTTTCGATGGTAGAGTATTGGTCTACCATGGTGTCGACGGGTGACG
GGGAATTAGGGTTCGATTCCGGAGAGGGAGCCTGAGAAACGGCTACCACATCCAAGGAAG
GCAGCAGGCGCGCAAATTACCCAATCCCGACTCGGGGAGGTAGTGACAAAAAATAACAAT
AGGGGGCCCTTTGGGTCTTCTAATTGGAATGAGAACAATTTAAATCCCTTATCGAGGATC
CATTGGAGGGCAAGTCTGGTGAACGTATGCATCAAAGCCCTCCGTAATAAAAAAAAACAG
CGCCAGATGTCTCTCTCTGGAAGCACACCACGCACGGGGTTCGTTCCTCTCTGCATGGCC
GTGTGAAAAAGGCGAGTAGGCAAACGATGGGAAACCCGGCTTGGAATAGTAACCAGCCAC
AGAGCCCCCATCATACACTTCTAAATTGCCGGGGAGTGCCCGCGAGGCCTTGACTACCGC
GGTCCGCCGAGAGGTGAGAGCGAGCAATCGCTCAGGACGCAGCAGTGTACGTAGTGGTAC
ATGGATGGTAAAAACGTCAAGGATAGGGTTGATCCGCAGGGAAGCGCGAGTGCCGCCATG
GTCGGTGGGCAAAGGCGAACCTTCAGAGACTCTAATGGAGTGGGCGCAAGCTTAAGGGAG
AGTCCAATCCACTGGGAAACCAGGTCCACAGCAGTAACGGGGTGATGCTCGCCCCAGAGG
ACTGTGGAAGGCAACCCCACCAAAAGGGTGTGTGTTGTCGGGTAGCAATGGCCAGCAGCC
GCGGTAATTCCAGCTCCAATAGCGTATATTAAAGTTGTTGCAGTTAAAACGCTCGTAGTC
GGACCTCGGAGCGCCCCCACGGGCCGGGCTTTTGCCTTGTTGCCAGTGGGAGTTTGTCTC
CTTTTGTCGGGACGCGGTGTGCAGGGCCTTTACTGTCTCTGCGCGTTTACCTGCGGGCCG
ACCGTTTACTGTGAAGAAAGTAAAGTGTTCAAAGCAGGCCAATGCCTTGAATATGTGAGC
ATGGAATAATAGAATAGGACTTGGGCTCTATTTTGTTGGTTTCCAGTGACCAAGTAATGA
TTAATAGGGACGGTTGGGGGCATTCGTATTTCATTGTCAGAGGTGAAATTCTTGGATTGA
TGGAAGACGTACAACTGCGAAAGCATCTGCCATGGATGTTTTCATTGATCAAGAACGAAA
GTTAGGGGATCGAAGACGATCAGATACCGTCGTAGTCTTAACCATAAACGATGCCGACTG
GGGATTGGCGGGGGCACTTTTATATGACTCTGTCAGCACCCTGAGGGAAACCAAAGTCTT
TGGGTTCTGGGGGGAGTATGGTCGCAAGGCTGAAACTTAAAGGAATTGACGGAAGGGCAC
CACCAGGAGTGGAGCCTGCGGCTTAATTTGACTCAACACGGGAAAACTTACCAGGTCCGG
ACAGAAGAATGATTGACAGACTGAAAGCTCTTTCTTGATTTTTTGGTTGGTGGTGCATGG
CCGTTCTTAGTTGGTGGAGTGATTTGTCTGGTTAATTCCGTTAACGAACGAGACCTCGTC

CTGCTAAATAGGTGCGCGCATGCCAGCACTGCGTTCTTACCTTCTTAGAGGGACTATGCG
CGTCTAGCGTATGGAAGATTGAGGCAATAACAGGTCTGTGATGCCCTTAGATGTTCTGGG
CCGCACGCGCGCTACACTGATGCATTCAACGAGTTTCACTTGATAGTCCTGGGTCGGAAG
GCTCGGGTAATCTTTTGAAAGTGCATCGTGCTGGGGATAGATCATTGCAATTATTGATCT
TCAACGAGGAATTCCTTGTAGGCGCAGGTCATCAGCCTGCGCCGAATACGTCCCTGCCCT
TTGTACACACCGCCCGTCGCTCCTACCGATTGAATGGTCCGATGAAATGTCCGGATCGCG
GCTAGTGTGATGGACTCGTCCAGAGCGGTAGCCGTGAGAAGCTCATTAAATCTTATCATT
TAGAGGAAGGAGAAGTCGTAACAAGGTTTCCGTAGGTGAACCTGCGGAAGGATCATTCAC
AAACTATTGACACAAACACACGCGAACCAAATCGTCCGCACAGCAGGTGGCAATGAAAGA
GAGACTCTGCATGTCGCCTTTCGGGGTAGCAAGCAGCACTCTTCTGCCATCGCCTCTGTG
CAGGGCGTAAATTCCCATTGAGAGGATGTGAGGGCACCACAGGAAGCTTTTCCACAGGAA
GTCGCCATCCTTCTCCCTCCACGGCGGCGCTTCTGTGCTTGGCAGTTTTTTTTGCCTTCT
AGGGAGGATGCCGCCAATGGAGCCCCTTATATATATACATCATCATATGCCCCTTTTTTT
CTTAACCGCTTGCCACAGCTTCTTCTATGAGGAGCTTGTGGGAAGACTGTCTCCATACAA
TAACAAAGATACAACTCTTAGCGGTGGATATCTTGGTTCTCGCAACGATGAAGAACGCAG
CCAACTGCGATAACTAATGTGAATTGCAGGACTTCGTGAATCATTAAGTCTTTGAACGCA
AGTTGCGCTCATCTCCATGAGGAGGTGAGCATGCCTGTTTGAGTGTCGATGGGAAAAACA
ATACATAACAACAATCGGCTCTCTTATCTTCCTGCCTTCTTGGGGGCAAAAGGTAGTGCA
TGCCGGAGATGGGCTTTGTCGCCCCTTTCTCTTGAAAGAGGCGTCGTCCGAAACACGTAT
CGTTGCCCGCTGGAGGTCATAGAGCATGAGTTTAAAGCACCACATCTCTCAAGTCAGAGT
GAGCGTCTTGAGTAGGTCCGCATTGATTCAGTTTCAACCCAGGTGGCGTTGGTTACTTGC
CATAAGCCTCTCTGGGGTTTCTTTGCTTGGACCTCTCTTGTCAGTTTCTCTGACGTAGAG
ACAGGTGCCGTCTCATGGACGTTGCCCTCTGGAACGTGCTAAAAAAAAAAAAAAGAAAAA
ACCGTACCACATGTCAAAAGAGGCAGATCGTGCTTGACGGCGGGAGGTGTTGCATTTCTT
GCTTGTCCTGAGCGTCTGCCAGTCTCTGACGAGTGTGTTCCTTCCATAAACCCGATCTCA
AATCAGGTAGGACCACCCGCTGAATTTAAGCATATAATTAAGCGGAGGAAAAGAAACTAA
CAAGGATTCCCTCAGTAGCGGCGAGCGAAGCGGGAAGAGCTCAACATGAAAACCGTTGGG
GGTTCGCGCTTTCCTTCGGGTTGTATAGTAAAGTGGTGTCGCCTGCGTCGCGCCGGGTAT
AAATCCCGTCTGAAATGGGGTGTCGTAGAGGGTTATATCCCCGTCTGTGACCCGGTGTGC
GATGCGTTACGGTGACCTTACGATGAGTCGGGTTGCTTGGGCATGCAGCCCTAAGTGGGT
GGTAAACTTCATCTAAAGCTAAATACTAGCGAGAGACCGATAGCGTACAAGTACCGTAAG
GGAAAGATGCAAAGAACTTTGAAAAGAGAGTTAAACAGGGCTTGAAACCGTTAAGAGTGA
AGCGAAGGAGGTCAGTGATCATGGGGCCAGATCCCCAAGTGCCTGTCATAGGGCACCGTA
GCGTTCTGGTTTCCCGGGGCAGCGCCGGCCGGGGGCGGGGGTTAACAGGGAGGTGGTTGA
GGGATTCGTTTTCCGCCTGTACTCCTTTATAGCTTGCACCCCGGCCGAGGGATCCGCGAC
TTGTCGTGTGAGGGCGCTGTCAAAATGGCCTTCTCCACCCCGTCTTGAAACACGGACCAA

GGAGTCTAACATACTCGCGAGTGTTTGGGTGGTTAAAACCCGAACGCACAATGAAAGTAA
ACGGTGGCTCTGCGCCGCTGCTTGGTGGGAACCCTGCTTTTATAATGGGGGGCACTATCA
CCCGACCCCGATCCCTTTGGGGTGGTTGGTTTTGAGGTAGAGCGAGTATGATGGGACCCG
AAAGATGGTGAACTATTCCTGAGTAGGGCGAAGCCAGAGGAAACTCTGGTGGAGGCTCGT
AGCGATGCTGACGTGCAAATCGCTCGTCGAACTTGGGTATAGGGGCGAAAGACTAATCGA
ACCATCTAGTAGCTGGTTCCCTCCGAAGTTTCCCTCAGGATAGCGGGAGCAAATGCAAGT
CAGTATCATCCGGTAAAGCGAATGATTAGAGGCCTCGGGGGAGTAACTCCTTCGACCTAT
TCTCAAACTTTAAATAGGTGAGGAGCCGGCAACGTTGCTTAAAACAATGAACGCCGTGCG
GACAATGATTGCTCTCAGTGGGCCGATTTTGGTAAGCAGAACTGGCGATGCGGGATGAAC
CGAAAGCGACGCTAAGGTGCCAAACTCCCAGCTCATTCAGACACCAGAAAAGGTGTTGAT
TCATACAGACAGCAGGACGGTGGCCATGGAAGTCGGAATCCGCTAAGGAGTGTGTAACAA
CTCACCTGCCGAATGGATCAGCCCTGAAAATGGATGGCGCTTAAGCGGGGGACCGAGGCG
TCGCCGTTGGGGTAAATACGAAGCCCCAATGAGTAGGAGGGCGTGGGGGTTTTGAAACAC
GGTACACTCGCATGGAAGCCTTGGGCGTAAGCTCGGGTGGAGCGACCCCTAGTGTAGATC
TTGGTGGTAGTAGCAAATATTCAAGTGAGAACCTTGAAGACTGAAGTGGGGAAGGGTTCC
ACGCGAACAGTGCTTGGACGTGGGTTAGTCGGTCCTAAGAGATGGGGAAACTCCGTATTC
AAGTGCGCTTTTGTAGCGCAGTCCATCGAAAGGGAAGCGGGTTAACATTCCCGCACCAGG
ATGTGGAAACTGTGGTTTACCGCAAACTGCGGCAACGCAACTGAACCTGGGGACGCCCGT
AGGAGCTCCGAGCAGGGATCTCTTTCCTGTTTAACGCGCCCAACCCCCTGGAATCGGTTT
GTCCGGAGATAGGGGGCGAAAGGCCGGAAAAGCACCGCACGTCTTGTGGTGTTCGGAACG
CTTGTAATGGCCCTTGAAAACCCAGGGGAAAGCAGCCGTCTTACACTTGTGTGAGACGAG
GAATTCTCACACCTGGCCGTACCCATCACCGCATCAGGTCCCCAAGGTGAACAGCCTCTA
GTCAATAGAATAATGTAGATAAGGGAAGTCGGCAAAATAGATCCGTAACTTTGGGAAAAG
GATTGGCTCTAATGTTCGGGCTTGTCGGGACGCGCGGGCGTAGATGCCAATGAGCGGCAG
ACGTCTAGCCTCTTTGGAGGCTGGATTAACGCTGCTTGTGGGAAGAAACCGTGTCGATGG
AGGCAGGAACGCTTATCGATCTTGTCTCTTCCCGCTTGCGATCAACGAACGGATTAGAAC
TGGTACGGACAAGGGGAATCCGACTGTTTAATTAAAACAAAGCATTGCGACGGCCGGAAA
CGGTGTTGGCGCAATGTGATTTCTGCCCAGTGCTCTGAATGTCAAAGTGAAGAAATTCAA
CCAAGCGCGGGTAAACGGCGGGAGTAACTATGACTCTCTTAGACAGATCAAGACACCTTC
AATAAATTGGTTCATTATGTCGGGTGGTCCGTCGTCCGGGAAGCGACCAAAGCAAAACCC
CGTAGACTGTGATGTGCATCTGTCAGACGAAACTGACGTCCGTGTTCATCAAGCCTTAGC
TATCTTTCAACGCCGGCTTCGAGACGTGGCGTCTCTTAAGGTGTTTGAAGTGGCTCTCTC
TGTGGGAACTACTGATGCACTGGTAGAGATGGGCACTATCACTTGCTGGAGTATTGATGC
TGTTGGGAAAGGTTTCTCAAGGAACAGCAGCGGTCGCTGGCAATTTGACCTCACTGGCAA
GATTAATGGCAAGCCTCTAATATCCACCACAAAGGATCGTTGGAGGCCGGTTGTGACGGC
TGTGCCCTACTACCTTTCGTGCGCCATCGATGGGGACAAGACGACGCTGGAGGAAAAGCA

GTGGACTTCATCACACATGTGTCACCATGAATGGTGTGTGAACGTGGAGCACATTGTTTA
TGAAAGCCTCGACGTCAACAAGTCTCGGAACTGGTGCCAGGGAAAAAACGCATGCCTCCA
TCTGCCTGTTTGCATTCGAACCGGCCGCCGTGTGCAGGAGCTCAGGAAGAGCAATGGGGT
TCGTGCAATGGACCTCCTCCTGCAGAAATTGGCTGGATTGGAGATTGAGGTTGAGGAGCC
CGCTGCAGCTGTACTCCGAGAGTCTGTCGAACGGGAGGACGCCTAGTGAGTGACTCGTGA
CACGGGTTGGCTGCGCATGATGCGTCAACTACATGGACATCTTTTGAGAGAGATAGGCGG
GCTGAAAAGGTTATAGCCGTCTAGTCTCGCTCTCTTTCATTGAGCGGGGCGACACACCTG
GTTGCGGGAACCCTGCCCACCTTTCTATCAAGAAGTGGGAACCAGTCCATTCCTACTCCT
GCTTGTGAGGCATATAGTAACGTAGCAGCAGGAGGATCTCGGCGAGAGGCCGAACCCGAA
AGGAATAACGTGGGCTGGGAGGCGATCCGCAGGCGAGTCCGTAACTATATGATTATCGAC
GGGCCGTCTCAACGACTGGTAAGGTGTGGGTTTGTCCTCCTCCTCACGTGGGGACAGGCT
TAAGGTACAGTCTAATCCCACCCGCCAAGGGTGCTTGGGTGTAATCCCTCTTCCATAGAG
GGCACTAGCCCAGGGGCTGGGACGCTGGGAGGAAATGCCCCGTCCCTATAGCCGGTAGAA
ATGAAGGTAGCCAAATGCCTCGTCATCTAAACTAATTGGATGAGATAGTCTGAGTGAAAA
GCTTATGCAGGCTAGTACCGGAAAAAAAAAAAAAAGAGGGGCTCTGCCACTCTGACGGTGC
GACACTCCTTGTACAGGGAAACCCTCCGCATGGAAAGGCACGTGGTACCACCCTCGCGGG
GAAACTCGCGAGCGGGATCCCGGGTAATGACCGGAGCCGATGGTAAAAACTCACGTGACC
GATGGGGTGACCCTGTGGCGAGACGCTAAGTTGTGGGACATCACCCTCTCACAATACGCA
TCCGTCGCAGAGACTGAACGGGAGTGGGCTCCGTTGACGGCCGGCCGGCCAGCCAGCCGT
CTTCCAGGGGCTTAAGATACAGTCCGGGCCTCAGGGATGGAAAAACCTGGGGAACTACCA
GAATTAGTGACGCGCATGAATGGATTAACGAGATTCCCACTGTCCCTATCTACTATCTAG
CGAAACCACAGCCAAGGGAACGGGCTTGGAAGAATCAGCGGGGAAAGAAGACCCTGTTGA
GCTTGACTCTAGTCCGACTTTGTGAGATGAGTTGGGAGGTGTAGTCTAAGAGGGAGGGGC
AACTCGACCGTGAAATACCTCTACTCCCAAGCTTGTTTTACTTACTCCGGCGAGCAGGGA
TGGGTGGTAACACCCTTTTTCTAGGATTAACACCTCTTCGGGGGTGGACCTGCGCGGAGG
ACATTGTCAGGTGGGGAGTTTGGCTGGGGCGGCACATCTGTTAAAAGATAACGCAGGTGT
CCTAAGATGAGCTCAGTGAGAACAGAAATCTCACGTAGAACAAAAGGGTAAAAGCTCATT
TGATTTTGATTTTCAGTACGAATACAAACTGTGAAAGCATGGCCTATCGATCCTTTTGAC
CTTCAAGGTTTGAAGCAAGAGGTGTCAGAAAAGTTAACACGCGTTGTGGTAGCTTTTAAG
AGTGGACTCAAAGTGTTCCGCTAGTCCCCCCTCGGACCGCCACTTACAAAAGGCCTGGGG
ATGGGGGCGACACTGTCAAATTGCGGGGACTCCCTAAAGCCTGTCTCTACCAAGCCAGCG
CGGCGAAAGCCCCTGGTGGCCGGGGTAATGACCTAGGGTATGGTAAAAGCGAGACAGGAT
GGAACAATGGGCAATCCGCAGCCAAGTCCTAAGGCTTTTTGAAAAGCGGCTATGGATGCA
GTTCAACGACTAGATGGCAGTGGGCTGAGAGAGCCCACCGCGGCACCCTTTCGGCTTAAG
ATATAGTCTTGCCCACGTGAAAGCGTGCCTGCAGATGAACGCCACCATGTATGGCCGGAG
CTGCAGGAGACCGCACTTGATGTGCAGTCGGGAATCGACAGTACCACAGGGATGAACAAA

AACACACAATTAATTTGTGGCCATCATACAGTCGTATGATATTATCGTCCCAGGCGGGTG
CGCCTCTCTCACCCGTCTAGTCCGGCCTTGGTGCGCAAAAAATGCTTGCGTGGATGCGCC
TTGGTTAAGGAAAATGCTCCTAGTAGTTTTACAAGGACACTTCTTGTAGGGCAACACATC
TGGATGCGAAGAACTCCCGGAAAAGACAACGGTACCAAGTGACTGGGGTAACGCCCCATT
CATGAAGGCAAATAATCAAAAAGAGGTGCCGGGTAAGAACCCGTTGTATAGGGACAATCC
GCAGCTGACTTGGCTCCCTGCCCATCCATGGGCATAGCTTAGCGGTTCAACGCTCGGTAA
GATGTGGGTTGGCGCGCTGTCCGACAGCGCACTGGCTTAAGGTACGGGCTACTCCCACCG
GAGACGGTGTCTTTGTCTACCTCGGGTAACACGCCTCGGGAGAAGCAAAGAGGGTGCAAT
GGAAAAAAAACTGGTTGCGGGGAAATCCCTACCAGAGTCACCTGGTACAAAGGAACTGGC
TTGTGGCAGCCAAGCGTTCATAGCGACGTTGCTTTTTGATCCTTCGATGTCGGCTCTTCC
TATCATTGCGAAGCAGAATTCGCCAAGTGTTGGATTGTTCACCCACTAATAGGGAACGTG
AGCTGGGTAGCCACTTTATGCCCAACAGCTGGATAGCTGGGCTTCTGTCTTAGGCCGGGA
CTTCTCACGACCCGTGCCTTGGTCACCCAAAAAAGGGCGACCGCAGAGCTCAGGATGGAG
CTTCTGGCTTAACTAGTAGAGGACAGATAGAGATCGTCAAATTGCGGGAAACCCCTTAGA
GCCTCGAGCTACCAAGTGCGTGTCGAAAGGCATAGACGCAGCCAGCGATGGCTGTCGCAT
GGCTTGATTCCATCAAGATAGGTAAAAATGCTCGAGGATTGGGCAATCCGCAGCCAAGCC
CCTAAGGTCCCCCCGCGGGCTATGGGGAAGGTTCAACGACTGTAAGGCGTACCGTGCAAG
CGGAAGATACAGTCTAGCCCCGCTGGAAAGCAGCGCCAAAGTAAGTAGGTTGCAGTGGGT
CTGTCCTC
>SRR9587968
GCATGTTTTGGTATGTGTTTCGACAGTCATGCATGTCATAGCCGTAGGATGGGGAGTGAG
CTTCATAGCCCGACCCCTTCTCAGGAGCATGTTTTCATAGCATATTGAGCATACTGGGGT
GGTCTCCCATGTAGCTCGCTCTCTCGGTGCAGGACCTCCCTTTTGCGGGCCACTTCGATG
GCCTCCATTATGGAGAGCATGTTTTGGCATATGAAGCGTTGGTGGTGAGGCTCGTAGCCT
GTATATACATACCAACGCAGTCGTCATATGTCTCTCTGGCGGGGGTCGGTCGTATGACCT
TGTCCCTACCTGGAGCATGTATTTTGTATGATATACTGGACGGCAGCTGCGCTGTTCAGT
CGTCGGCCGTACCTCATGTGCGCGTTTCCAGGTCCGAACGTTTGACATGCAAGGGCATCA
TACCAGAAGCCTGTTTTCTCGTATGAGCATGTAGGTAGGCAGGCTCTTCAGTCGTCTACC
CATAAATGCACTTCGAAGGAACTTTTTGGTATTGAATGGCTTTGTGCCCAGACGTTCTTG
TGAGCTTGTTCTCGCTCTGCGATACGCTGGAAGCTGTGGTATTACCTACAGCAGACGCAT
TCCTAGGGCGGACGGCCTCTCTGCAGATGGGCACATCTGTAGAAAGAGGCTCCCCTTGAT
GACGCAATGCAATGGCTGGGTAAACGGAGTGCTATACAGAAGGGTCAAACCTCTGTGGAA
GTACACTCATACCAGCATTGTTTGTTGAGAGGGGTCGGCTCTACGAGTGAGGCCCTATAC
TGCCATCTCATTCAGATGCGTAGACGGTCTTTACTCAGACTGCCTTTCTGGCTGATTGTC
CTCGGTCCCTCGTGCGCACGCGCCGCTGGCTGAATGGGCTTGTCCTTATGGAGTGCCGCG
TTCTTCAAAGAATCCGTCTTCATACAGCGACCTCATTTACATATTGCCGGCTGAATGGGC

TTGTCCTTATGGAGCACTGCATTCTTCTAGGAATTGCTATCTATACAGCGACCTCATTTA
CATACAAGGCACCTTTTTCTACATAGACTTGCCATTTGGCTGATTGTTCTGACAGTCGCT
GCCTCTTCGGAGTTCTGTGTGGACGAGTGGCCTCCTGAGTGGGCCGGTCCTTACGGAGTG
CCCAAGTCTGTTATGACTGGGTTCTCCATACAGCGACCTCACTCAGAATTGTATGGCACC
GTGCCGATTGAATGGACAAGCAGTCCTTAGGGAATCGCATGAAAGCATGACGGCAACATG
GTGTATGGACCCGTTGGGCCTTGCCTGTGCTATCTTCTCGAGTCGACCGGTCCTCGTCTC
CTCCTTTCGCGACGGGAAGCTCTTGAGTTTACCGTGGGTCAGGACGCTGTGGTATCTGTT
TGTCTGTAGTGTGGGCTCTCTATCAGAACAGCGTATCAGCACACGTCGTCTGGGTCTATG
GTATTTTTGCTTAACGGCCCTGGGTTTGGTAGTATGGTTCCATGTCGGTCCTCTCACCTC
CCAACCGTCATGTCTCTGATATCATCTGCACAGTACGTCTTGTCAGAATATCATTCTGTA
TGGCAGTCGATAGACTAGGCCCTATGCCGCAGCTGTCTCTGACCACTATGCGAGCGCATA
CCTACCTCAATGTGTTCATAGCTGTTATGGATCTTGGGTGTGGTGTGCCAGAACGATACG
CCACCAGATTGTGGAATGGTGCTTGTAGTATTAGAAGAGGACATGTATGAAGAGCACGTT
CTTTTCTCTTATGATGGTGCCGCGGGTGCGGGCCGTTCTCATTCTGCTTGTCGGGTGCTC
AAGGTGCATGTAGGATTGCTGAACCCACTGAATGGTCATGAGAATGGTCATATGGTGAGG
TTCACTTCTCCATCGCTTTGTGTGTGTTGGATGTTTGGGTGGCTTCGGCCGCATGGAAAA
TCTCGGTCTGTAACTCATCAGGGAGGGTTTCCATGTGAGCAATCTCTCCAGTCGAACACA
CACCCATGGGCGTCCCGCTAGGCAAAATGGAACTCTCGATCCGACACAGCCTGTGGAGTG
TTTGTTCCGTGTGGCGTGAAACGGCTTATACTGGATGTCTTCCCTTTGGACACCAGTCTG
TTGGTATGTACGCATACCATCTAGAGGGAGCAAGAAAAAGGCATGCAAGTGCCACAAAGG
AAGGGAGGCCTTCCTGCTACCGTGTGTCCTCTTGACGATCACGGGGGAGTATATAGAGAC
ATTTGGACGAGATCCGTTTTTCACGAGGCGGTGCCCTCGCGCAAAGGTGTTGCTCTGCAC
GCATCTTTATGATGATAGTTGCTCGCTCCTATGCGTCCACTCGTGAAAATACCTGCTCTC
TCTGTCTGTTTCCATGACACCCCCATGCGTTGTCATAGAGCATGCTCAGGTACAGAAAGT
GCTGTGACTTTCTTGGCCTGCTGTCACTGAGTCAGCCTGATACTTTGGAGGAATCTTTTC
TGGGCACTGCCCAGCTGAGAGTTCTTATCAGGCTCTGGAGTGTGCATATGCCTATATTCA
TTATAGTGCATGTGGCTACGCAGAAAAAGCATCGACGTGGAATGGAGAAGAACACCATTC
TGAATGGAACGGTTCTTACTCTACTTCTGTTGCTTGACGCTGTATGGGAATCGCTTTGCT
CTGGGGGTACACCTTCCATTCCATGACAAAGCATTTGATCTATGCCCCGAGGGTCATTCT
CTATGAAGAGCCCGAAACTTGTCTGAGTGAGAGGCGAAGGTAGTCCTTCATTTATGCGGG
GTCCGCCCTCACACTGTAGTCGGCACAGTCCCTTGTTTATGGACTGCGCAGCTCAATCTT
CGACTGGAGAGGAAAAACAAAACACAAGTATGACACGCACATACGTGTTTTCTAGGACTG
TGACCTGTCGGCTGCCTCGCGGTGGTTGGCATGATGCCCCAAGTGGGGGGGGGGAGAAAC
AATCAGTATGATTGATAAAAACCAGTGGGGAAAAAGGCGCGCTGTCGCACACCCTTGGTC
ACGGCCTCAAGTCGACGGCCCTCACTGGCCCACCCTGCGACTCCTGTTCGCTTTTGACAA
GACACACCTGCAATACGGAGGAAATGCCTGAGCAGCTGTGCCGCAGCAACTGCGGACGCG

TAGCGATATGCATCCCTGTTGTTTGCGACTTGGTACAAAGAACGTGCGCCGGGAGACGGA
TTGCCCTTTGACTTTTCCACCCAGCCCGCGGCTCTATACCCTTGACGTACCTATTTACCT
GGTTGATCCTGCCAGTAGTCATATGCTTGTCTCAAAGACTAAGCCATGCATGTCTAAGTA
TAAACACTTTTATACTCGTGAAACTGCGAATGGCTCATTAAAACAGTTATAGTTCCTTTG
GGAACAACACTACTTGGATAACCGTAGTAATTCTAGAGCTAATACATGCCTCAACGCCCG
ACTCACGAAGGGTGGTATTTATTGGATAAAAAACCATCGTGCTCTGTCCAGAGCGCTTTG
AGATGATTCACAATAACTTGTCGGATCGCATGGCCTCGTGCTGGCGACTCCTCATTCAAA
TTTCTGCCCTATCAACTTTCGATGGTAGAGTATTGGTCTACCATGGTGTCGACGGGTGAC
GGGGAATTAGGGTTCGATTCCGGAGAGGGAGCCTGAGAAACGGCTACCACATCCAAGGAA
GGCAGCAGGCGCGCAAATTACCCAATCCCGACTCGGGGAGGTAGTGACAAAAAATAACAA
TAGGGGGCCCTTTGGGTCTTCTAATTGGAATGAGAACAATTTAAATCCCTTATCGAGGAT
CCATTGGAGGGCAAGTCTGGTGCCAGCAGCCGCGGTAATTCCAGCTCCAATAGCGTATAT
TAAAGTTGTTGCAGTTAAAACGCTCGTAGTCGGACCTCGGAGCGCCCCCACGGGCCGGGC
TTTTGCCTTGTTGCCAGTGGGAGTTTGTCTCCTTTTGTCGGGACGCGGTGTGCAGGGCCT
TTACTGTCTCTGCGCGTTTACCTGCGGGCCGACCGTTTACTGTGAAGAAAGTAAAGTGTT
CAAAGCAGGCCAATGCCTTGAATATGTGAGCATGGAATAATAGAATAGGACTTGGGCTCT
ATTTTGTTGGTTTCCAGTGACCAAGTAATGATTAATAGGGACGGTTGGGGGCATTCGTAT
TTCATTGTCAGAGGTGAAATTCTTGGATTGATGGAAGACGTACAACTGCGAAAGCATCTG
CCATGGATGTTTTCATTGATCAAGAACGAAAGTTAGGGGATCGAAGACGATCAGATACCG
TCGTAGTCTTAACCATAAACGATGCCGACTGGGGATTGGCGGGGGCACTTTTATATGACT
CTGTCAGCACCCTGAGGGAAACCAAAGTCTTTGGGTTCTGGGGGGAGTATGGTCGCAAGG
CTGAAACTTAAAGGAATTGACGGAAGGGCACCACCAGGAGTGGAGCCTGCGGCTTAATTT
GACTCAACACGGGAAAACTTACCAGGTCCGGACAGAAGAATGATTGACAGACTGAAAGCT
CTTTCTTGATTTTTTGGTTGGTGGTGCATGGCCGTTCTTAGTTGGTGGAGTGATTTGTCT
GGTTAATTCCGTTAACGAACGAGACCTCGTCCTGCTAAATAGGTGCGCGCATGCCAGCAC
TGCGTTCTTACCTTCTTAGAGGGACTATGCGCGTCTAGCGTATGGAAGATTGAGGCAATA
ACAGGTCTGTGATGCCCTTAGATGTTCTGGGCCGCACGCGCGCTACACTGATGCATTCAA
CGAGTTTCACTTGATAGTCCTGGGTCGGAAGGCTCGGGTAATCTTTTGAAAGTGCATCGT
GCTGGGGATAGATCATTGCAATTATTGATCTTCAACGAGGAATTCCTTGTAGGCGCAGGT
CATCAGCCTGCGCCGAATACGTCCCTGCCCTTTGTACACACCGCCCGTCGCTCCTACCGA
TTGAATGGTCCGATGAAATGTCCGGATCGCGGCTAGTGTGATGGACTCGTCCAGAGCGGT
AGCCGTGAGAAGCTCATTAAATCTTATCATTTAGAGGAAGGAGAAGTCGTAACAAGGTTT
CCGTAGGTGAACCTGCGGAAGGATCATTCACAAACTATTGACACAAACACACGCGAACCA
AATCGTCCGCACAGCAGGTGGCAATGAAAGAGAGACTCTGCATGTCGCCTTTCGGGGTAG
CAAGCAGCACTCTTCTGCCATCGCCTCTGTGCAGGGCGTAAATTCCCATTGAGAGGATGT
GAGGGCACCACAGGAAGCTTTTCCACAGGAAGTCGCCATCCTTCTCCCTCCACGGCGGCG

CTTCTGTGCTTGGCAGTTTTTTTTGCCTTCTAGGGAGGATGCCGCCAATGGAGCCCCTTA
TATATATACATCATCATATGCCCCTTTTTTTCTTAACCGCTTGCCACAGCTTCTTCTATG
AGGAGCTTGTGGGAAGACTGTCTCCATACAATAACAAAGATACAACTCTTAGCGGTGGAT
ATCTTGGTTCTCGCAACGATGAAGAACGCAGCCAACTGCGATAACTAATGTGAATTGCAG
GACTTCGTGAATCATTAAGTCTTTGAACGCAAGTTGCGCTCATCTCCATGAGGAGGTGAG
CATGCCTGTTTGAGTGTCGATGGGAAAAACAATACATAACAACAATCGGCTCTCTTATCT
TCCTGCCTTCTTGGGGGCAAAAGGTAGTGCATGCCGGAGATGGGCTTTGTCGCCCCTTTC
TCTTGAAAGAGGCGTCGTCCGAAACACGTATCGTTGCCCGCTGGAGGTCATAGAGCATGA
GTTTAAAGCACCACATCTCTCAAGTCAGAGTGAGCGTCTTGAGTAGGTCCGCATTGATTC
AGTTTCAACCCAGGTGGCGTTGGTTACTTGCCATAAGCCTCTCTGGGGTTTCTTTGCTTG
GACCTCTCTTGTCAGTTTCTCTGACGTAGAGACAGGTGCCGTCTCATGGACGTTGCCCTC
TGGAACGTGCTAAAAAAAAAAAAAAGAAAAAACCGTACCACATGTCAAAAGAGACAGATC
GTGCTTGACGGCGGGAGGTGTTGCATTTCTTGCTTGTCCTGAGCGTCTGCCAGTCTCTGA
CGAGTGTGTTCCTTCCATAAACCCGATCTCAAATCAGGTAGGACCACCCGCTGAATTTAA
GCATATAATTAAGCGGAGGAAAAGAAACTAACAAGGATTCCCTCAGTAGCGGCGAGCGAA
GCGGGAAGAGCTCAACATGAAAACCGTTGGGGGTTCGCGCTTTCCTTCGGGTTGTATAGT
AAAGTGGTGTCGCCTGCGTCGCGCCGGGTATAAATCCCGTCTGAAATGGGGTGTCGTAGA
GGGTTATATCCCCGTCTGTGACCCGGTGTGCGATGCGTTACGGTGACCTTACGATGAGTC
GGGTTGCTTGGGCATGCAGCCCTAAGTGGGTGGTAAACTTCATCTAAAGCTAAATACTAG
CGAGAGACCGATAGCGTACAAGTACCGTAAGGGAAAGATGCAAAGAACTTTGAAAAGAGA
GTTAAACAGGGCTTGAAACCGTTAAGAGTGAAGCGAAGGAGGTCAGTGATCATGGGGCCA
GATCCCCAAGTGCCTGTCATAGGGCACCGTAGCGTTCTGGTTTCCCGGGGCAGCGCCGGC
CGGGGGCGGGGGTTAACAGGGAGGTGGTTGAGGGATTCGTTTTCCGCCTGTACTCCTTTA
TAGCTTGCACCCCGGCCGAGGGATCCGCGACTTGTCGTGTGAGGGCGCTGTCAAAATGGC
CTTCTCCACCCCGTCTTGAAACACGGACCAAGGAGTCTAACATACTCGCGAGTGTTTGGG
TGGTTAAAACCCGAACGCACAATGAAAGTAAACGGTGGCTCTGCGCCGCTGCTTGGTGGG
AACCCTGCTTTTATAATGGGGGGCACTATCACCCGACCCCGATCCCTTTGGGGTGGTTGG
TTTTGAGGTAGAGCGAGTATGATGGGACCCGAAAGATGGTGAACTATTCCTGAGTAGGGC
GAAGCCAGAGGAAACTCTGGTGGAGGCTCGTAGCGATGCTGACGTGCAAATCGCTCGTCG
AACTTGGGTATAGGGGCGAAAGACTAATCGAACCATCTAGTAGCTGGTTCCCTCCGAAGT
TTCCCTCAGGATAGCGGGAGCAAATGCAAGTCAGTATCATCCGGTAAAGCGAATGATTAG
AGGCCTCGGGGGAGTAACTCCTTCGACCTATTCTCAAACTTTAAATAGGTGAGGAGCCGG
CAACGTTGCTTAAAACAATGAACGCCGTGCGGACAATGATTGCTCTCAGTGGGCCGATTT
TGGTAAGCAGAACTGGCGATGCGGGATGAACCGAAAGCGACGCTAAGGTGCCAAACTCCC
AGCTCATTCAGACACCAGAAAAGGTGTTGATTCATACAGACAGCAGGACGGTGGCCATGG
AAGTCGGAATCCGCTAAGGAGTGTGTAACAACTCACCTGCCGAATGGATCAGCCCTGAAA

ATGGATGGCGCTTAAGCGGGGGACCGAGGCGTCGCCGTTGGGGTAAATACGAAGCCCCAA
TGAGTAGGAGGGCGTGGGGGTTTTGAAACACGGTACACTCGCATGGAAGCCTTGGGCGTA
AGCTCGGGTGGAGCGACCCCTAGTGTAGATCTTGGTGGTAGTAGCAAATATTCAAGTGAG
AACCTTGAAGACTGAAGTGGGGAAGGGTTCCACGCGAACAGTGCTTGGACGTGGGTTAGT
CGGTCCTAAGAGATGGGGAAACTCCGTATTCAAGTGCGCTTTTGTAGCGCAGTCCATCGA
AAGGGAAGCGGGTTAACATTCCCGCACCAGGATGTGGAAACTGTGGTTTACCGCAAACTG
CGGCAACGCAACTGAACCTGGGGACGCCCGTAGGAGCTCCGAGCAGGGATCTCTTTCCTG
TTTAACGCGCCCAACCCCCTGGAATCGGTTTGTCCGGAGATAGGGGGCGAAAGGCCGGAA
AAGCACCGCACGTCTTGTGGTGTTCGGAACGCTTGTAATGGCCCTTGAAAACCCAGGGGA
AAGCAGCCGTCTTACACTTGTGTGAGACGAGGAATTCTCACACCTGGCCGTACCCATCAC
CGCATCAGGTCCCCAAGGTGAACAGCCTCTAGTCAATAGAATAATGTAGATAAGGGAAGT
CGGCAAAATAGATCCGTAACTTTGGGAAAAGGATTGGCTCTAATGTTCGGGCTTGTCGGG
ACGCGCGGGCGTAGATGCCAATGAGCGGCAGACGTCTAGCCTCTTTGGAGGCTGGATTAA
CGCTGCTTGTGGGAAGAAACCGTGTCGATGGAGGCAGGAACGCTTATCGATCTTGTCTCT
TCCCGCTTGCGATCAACGAACGGATTAGAACTGGTACGGACAAGGGGAATCCGACTGTTT
AATTAAAACAAAGCATTGCGACGGCCGGAAACGGTGTTGGCGCAATGTGATTTCTGCCCA
GTGCTCTGAATGTCAAAGTGAAGAAATTCAACCAAGCGCGGGTAAACGGCGGGAGTAACT
ATGACTCTCTTAGACAGATCAAGACACCTTCAATAAATTGGTTCATTATGTCGGGTGGTC
CGTCGTCCGGGAAGCGACCAAAGCAAAACCCCGTAGACTGTGATGTGCATCTGTCAGACG
AAACTGACGTCCGTGTTCATCAAGCCTTAGCTATCTTTCAACGCCGGCTTCGAGACGTGG
CGTCTCTTAAGGTGTTTGAAGTGGCTCTCTCTGTGGGAACTACTGATGCACTGGTAGAGA
TGGGCACTATCACTTGCTGGAGTATTGATGCTGTTGGGAAAGGTTTCTCAAGGAACAGCA
GCGGTCGCTGGCAATTTGACCTCACTGGCAAGATTAATGGCAAGCCTCTAATATCCACCA
CAAAGGATCGTTGGAGGCCGGTTGTGACGGCTGTGCCCTACTACCTTTCGTGCGCCATCG
ATGGGGACAAGACGACGCTGGAGGAAAAGCAGTGGACTTCATCACACATGTGTCACCATG
AATGGTGTGTGAACGTGGAGCACATTGTTTATGAAAGCCTCGACGTCAACAAGTCTCGGA
ACTGGTGCCAGGGAAAAAACGCATGCCTCCATCTGCCTGTTTGCATTCGAACCGGCCGCC
GTGTGCAGGAGCTCAGGAAGAGCAATGGGGTTCGTGCAATGGACCTCCTCCTGCAGAAAT
TGGCTGGATTGGAGATTGAGGTTGAGGAGCCCGCTGCAGCTGTACTCCGAGAGTCTGTCG
AACGGGAGGACGCCTAGTGAGTGACTCGTGACACGGGTTGGCTGCGCATGATGCGTCAAC
TACATGGACATCTTTTGAGAGAGATAGGCGGGCTGAAAAGGTTATAGCCGTCTAGTCTCG
CTCTCTTTCATTGAGCGGGGCGACACACCTGGTTGCGGGAACCCTGCCCACCTTTCTATC
AAGAAGTGGGAACCAGTCCATTCCTACTCCTGCTTGTGAGGCATATAGTAACGTAGCAGC
AGGAGGATCTCGGCGAGAGGCCGAACCCGAAAGGAATAACGTGGGCTGGGAGGCGATCCG
CAGGCGAGTCCGTAACTATATGATTATCGACGGGCCGTCTCAACGACTGGTAAGGTGTGG
GTTTGTCCTCCTCCTCACGTGGGGACAGGCTTAAGGTACAGTCTAATCCCACCCGCCAAG

GGTGCTTGGGTGTAATCCCTCTTCCATAGAGGGCACTAGCCCAGGGGCTGGGACGCTGGG
AGGAAATGCCCCGTCCCTATAGCCGGTAGAAATGAAGGTAGCCAAATGCCTCGTCATCTA
AACTAATTGGATGAGATAGTCTGAGTGAAAAGCTTATGCAGGCTAGTACCGGAAAAAAAA
AAAAAGAGGGGCTCTGCCACTCTGACGGTGCGACACTCCTTGTACAGGGAAACCCTCCGC
ATGGAAAGGCACGTGGTACCACCCTCGCGGGGAAACTCGCGAGCGGGATCCCGGGTAATG
ACCGGAGCCGATGGTAAAAACTCACGTGACCGATGGGGTGACCCTGTGGCGAGACGCTAA
GTTGTGGGACATCACCCTCTCACAATACGCATCCGTCGCAGAGACTGAACGGGAGTGGGC
TCCGTTGACGGCCGGCCGGCCAGCCAGCCGTCTTCCAGGGGCTTAAGATACAGTCCGGGC
CTCAGGGATGGAAAAACCTGGGGAACTACCAGAATTAGTGACGCGCATGAATGGATTAAC
GAGATTCCCACTGTCCCTATCTACTATCTAGCGAAACCACAGCCAAGGGAACGGGCTTGG
AAGAATCAGCGGGGAAAGAAGACCCTGTTGAGCTTGACTCTAGTCCGACTTTGTGAGATG
AGTTGGGAGGTGTAGTCTAAGAGGGAGGGGCAACTCGACCGTGAAATACCTCTACTCCCA
AGCTTGTTTTACTTACTCCGGCGAGCAGGGATGGGTGGTAACACCCTTTTTCTAGGATTA
ACACCTCTTCGGGGGTGGACCTGCGCGGAGGACATTGTCAGGTGGGGAGTTTGGCTGGGG
CGGCACATCTGTTAAAAGATAACGCAGGTGTCCTAAGATGAGCTCAGTGAGAACAGAAAT
CTCACGTAGAACAAAAGGGTAAAAGCTCATTTGATTTTGATTTTCAGTACGAATACAAAC
TGTGAAAGCATGGCCTATCGATCCTTTTGACCTTCAAGGTTTGAAGCAAGAGGTGTCAGA
AAAGTTAACACGCGTTGTGGTAGCTTTTAAGAGTGGACTCAAAGTGTTCCGCTAGTCCCC
CCTCGGACCGCCACTTACAAAAGGCCTGGGGATGGGGGCGACACTGTCAAATTGCGGGGA
CTCCCTAAAGCCTGTCTCTACCAAGCCAGCGCGGCGAAAGCCCCTGGTGGCCGGGGTAAT
GACCTAGGGTATGGTAAAAGCGAGACAGGATGGAACAATGGGCAATCCGCAGCCAAGTCC
TAAGGCTTTTTGAAAAGCGGCTATGGATGCAGTTCAACGACTAGATGGCAGTGGGCTGAG
AGAGCCCACCGCGGCACCCTTTCGGCTTAAGATATAGTCTTGCCCACGTGAAAGCGTGCC
TGCAGATGAACGCCACCATGTATGGCCGGAGCTGCAGGAGACCGCACTTGATGTGCAGTC
GGGAATCGACAGTACCACAGGGATGAACAAAAACACACAATTGTGGCCATCATACAGTCG
TATGATATTATCGTCCCAGGCGGGTGCGCCTCTCTCACCCGTCTAGTCCGGCCTTGGTGC
GCAAAAAAATGCTTGCGTGGATGCGCCTTGGTTAAGGAAAATGCTCCTAGTAGTTTTACAA
GGACACTTCTTGTAGGGCAACACATCTGGATGCGAAGAACTCCCGGAAAAGACAACGGTA
CCAAGTGACTGGGGTAACGCCCCATTCATGAAGGCAAATAATCAAAAAGAGGTGCCGGGT
AAGAACCCGTTGTATAGGGACAATCCGCAGCTGACTTGGCTCCCTGCCCATCCATGGGCA
TAGCTTAGCGGTTCAACGCTCGGTAAGATGTGGGTTGGCGCGCTGTCCGACAGCGCACTG
GCTTAAGGTACGGGCTACTCCCACCGGAGACGGTGTCTTTGTCTACCTCGGGTAACACGC
CTCGGGAGAAGCAAAGAGGGTGCAATGGAAAAAAACTGGTTGCGGGGAAATCCCTACCAG
AGTCACCTGGTACAAAGGAACTGGCTTGTGGCAGCCAAGCGTTCATAGCGACGTTGCTTT
TTGATCCTTCGATGTCGGCTCTTCCTATCATTGCGAAGCAGAATTCGCCAAGTGTTGGAT
TGTTCACCCACTAATAGGGAACGTGAGCTGGGTAGCCACTTTATGCCCAACAGCTGGATA

GCTGGGCTTCTGTCTTAGGCCGGGACTTCTCACGACCCGTGCCTTGGTCACCCAAAAAAG
GGCGACCGCAGAGCTCAGGATGGAGCTTCTGGCTTAACTAGTAGAGGACAGATAGAGATC
GTCAAATTGCGGGAAACCCCTTAGAGCCTCGAGCTACCAAGTGCGTGTCGAAAGGCATAG
ACGCAGCCAGCGATGGCTGTCGCATGGCTTGATTCCATCAAGATAGGTAAAAATGCTCGA
GGATTGGGCAATCCGCAGCCAAGCCCCTAAGGTCCCCCCGCGGGCTATGGGGAAGGTTCA
ACGACTGTAAGGCGTACCGTGCAAGCGGAAGATACAGTCTAGCCCCGCTGGAAAGCAGCG
CCAAAGTAAGTAGGTTGCAGTGGGTCTGTCCTC

**Supplementary Figure 2.** The result of mapping the reads of each sample to its own core sequence.

### Pyr_1

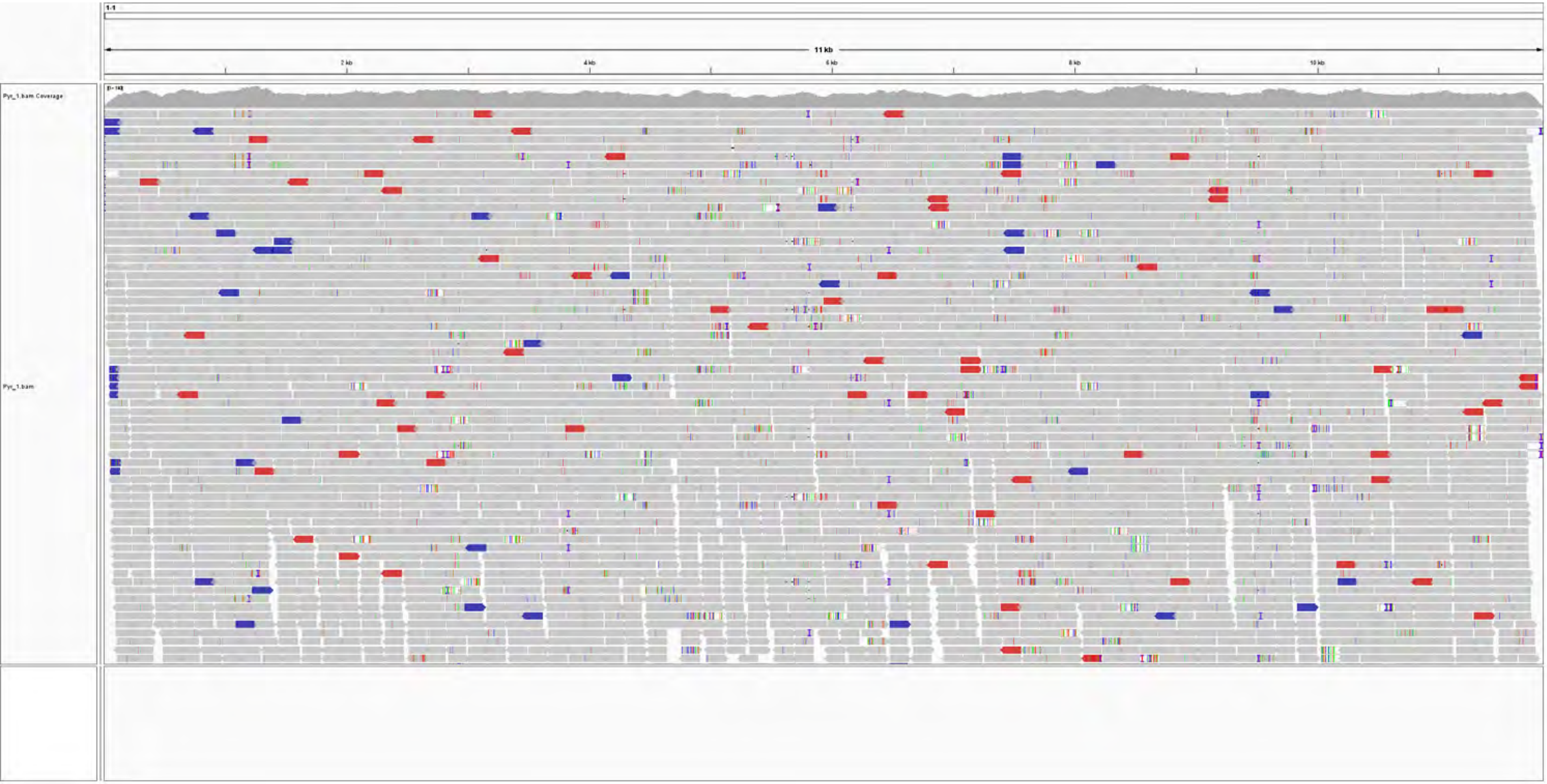

### Pyr_2

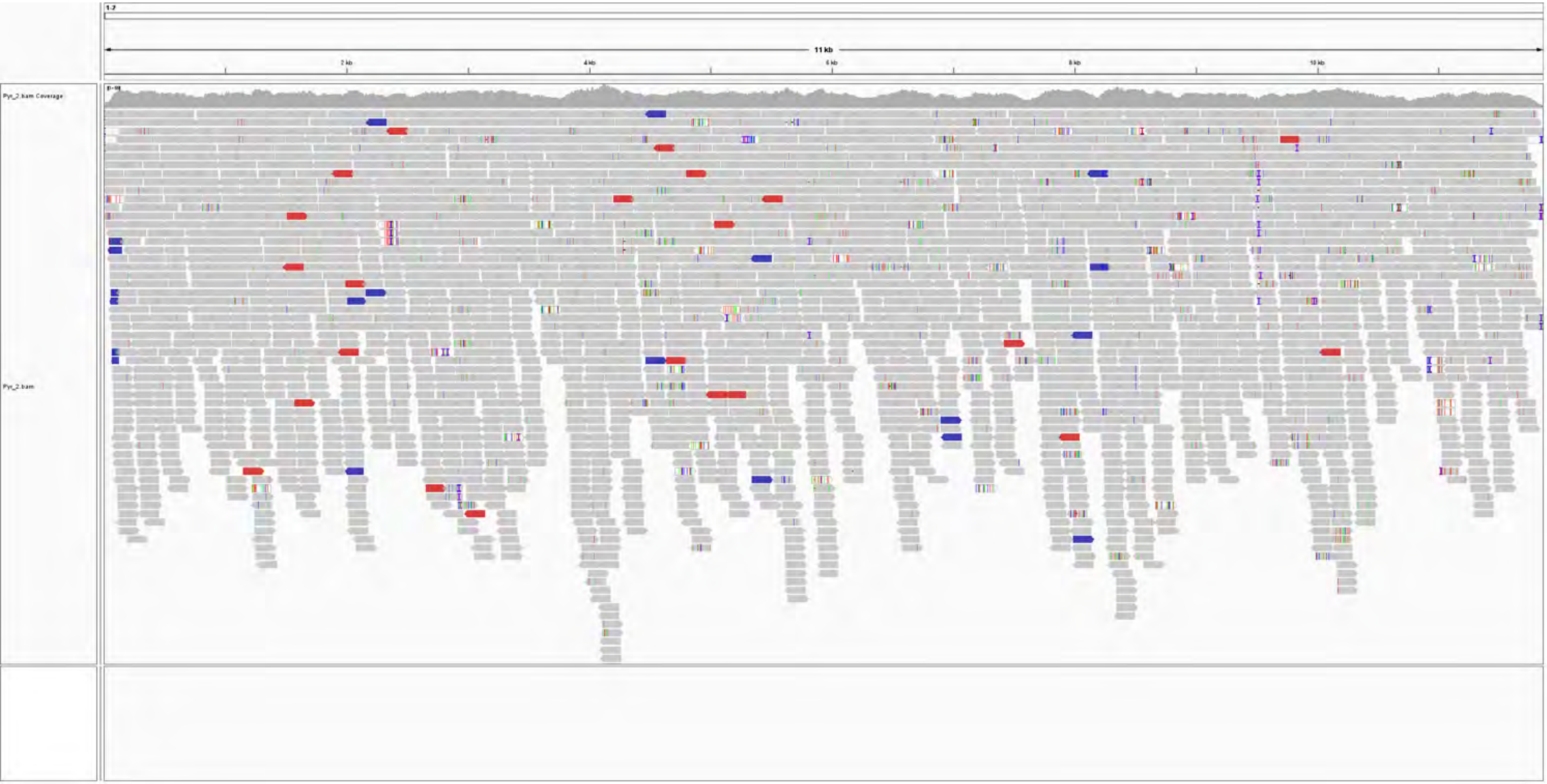

### Pyr_4

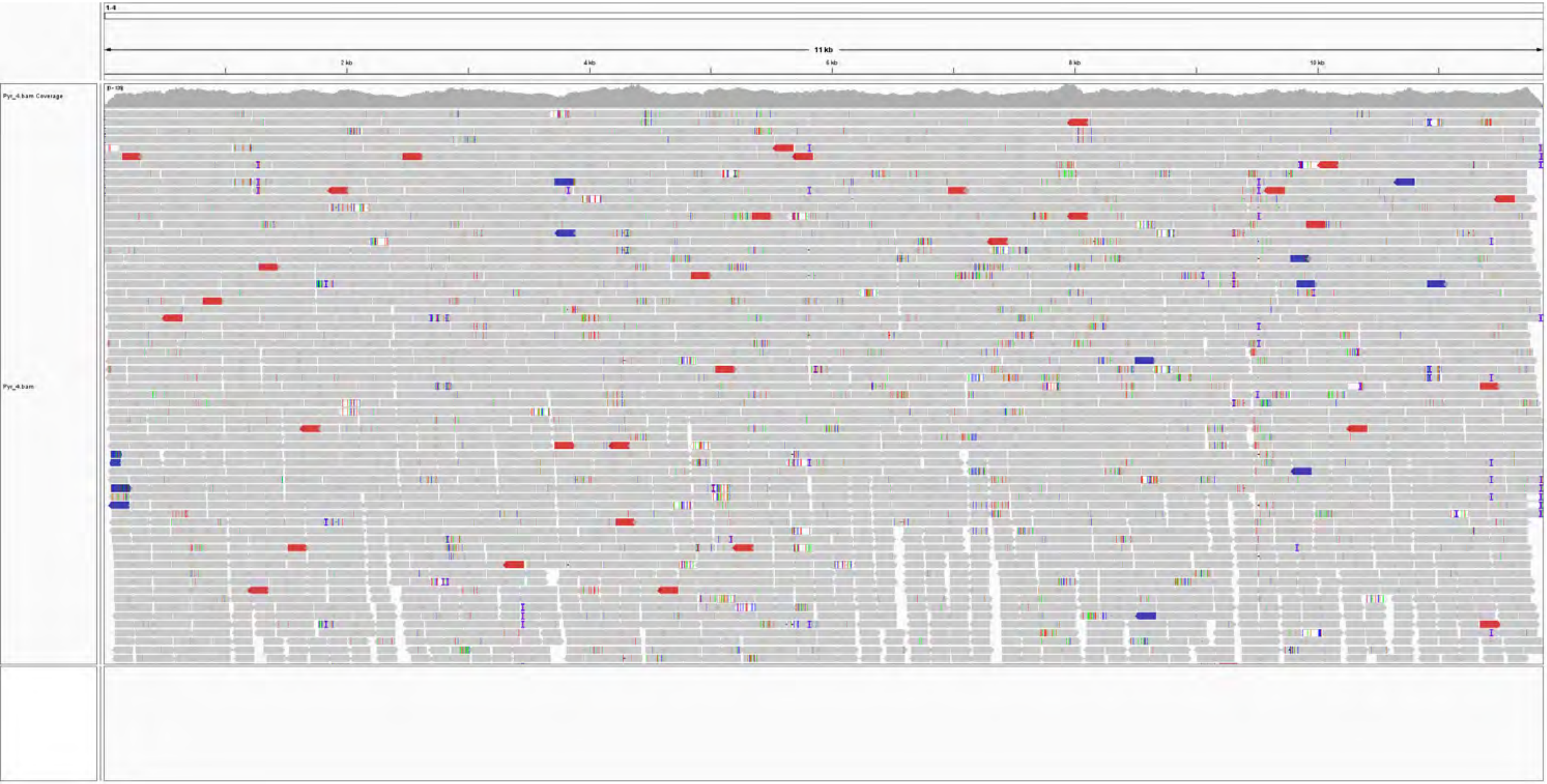

### Pyr_7

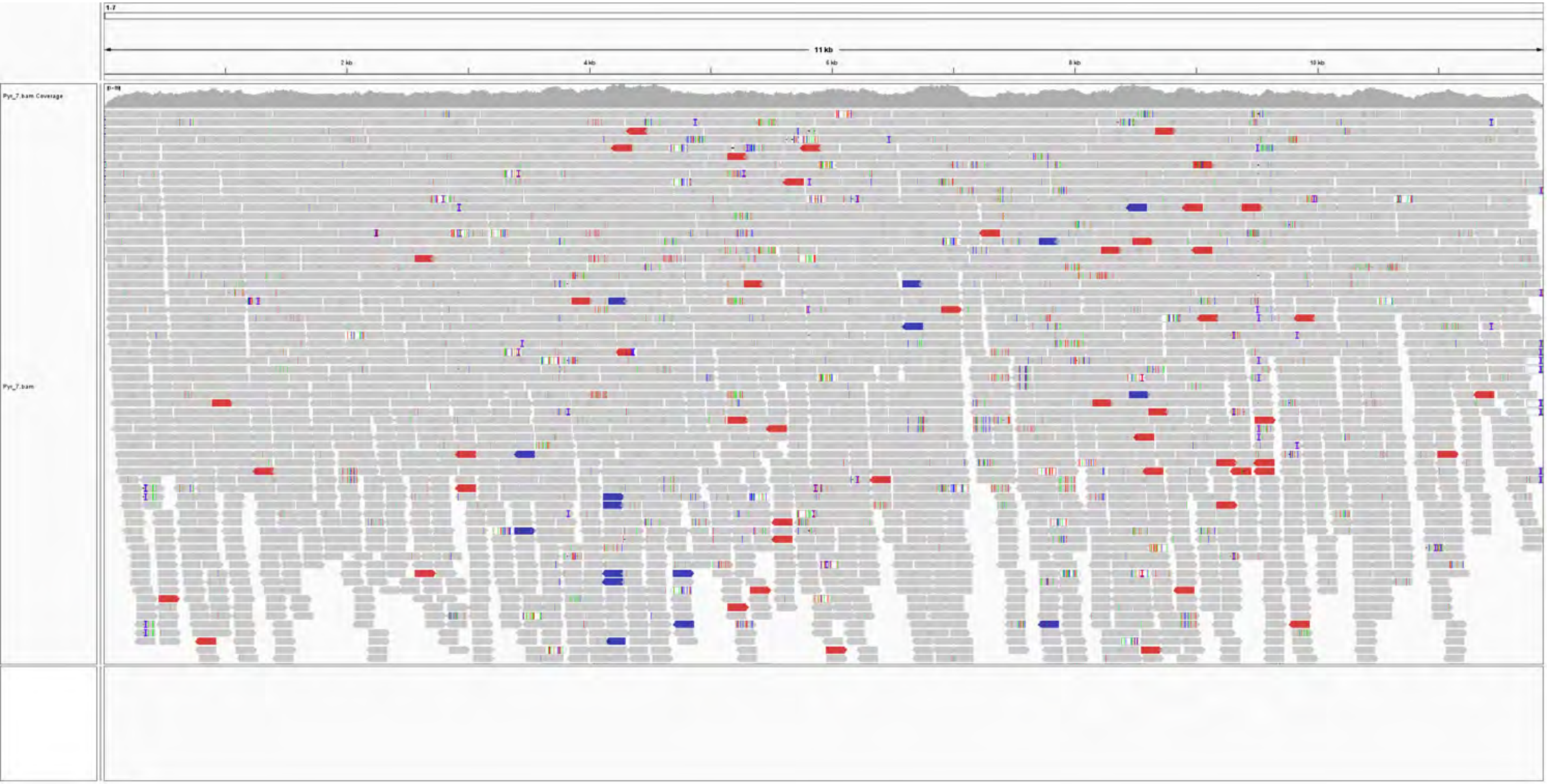

### Pyr_8

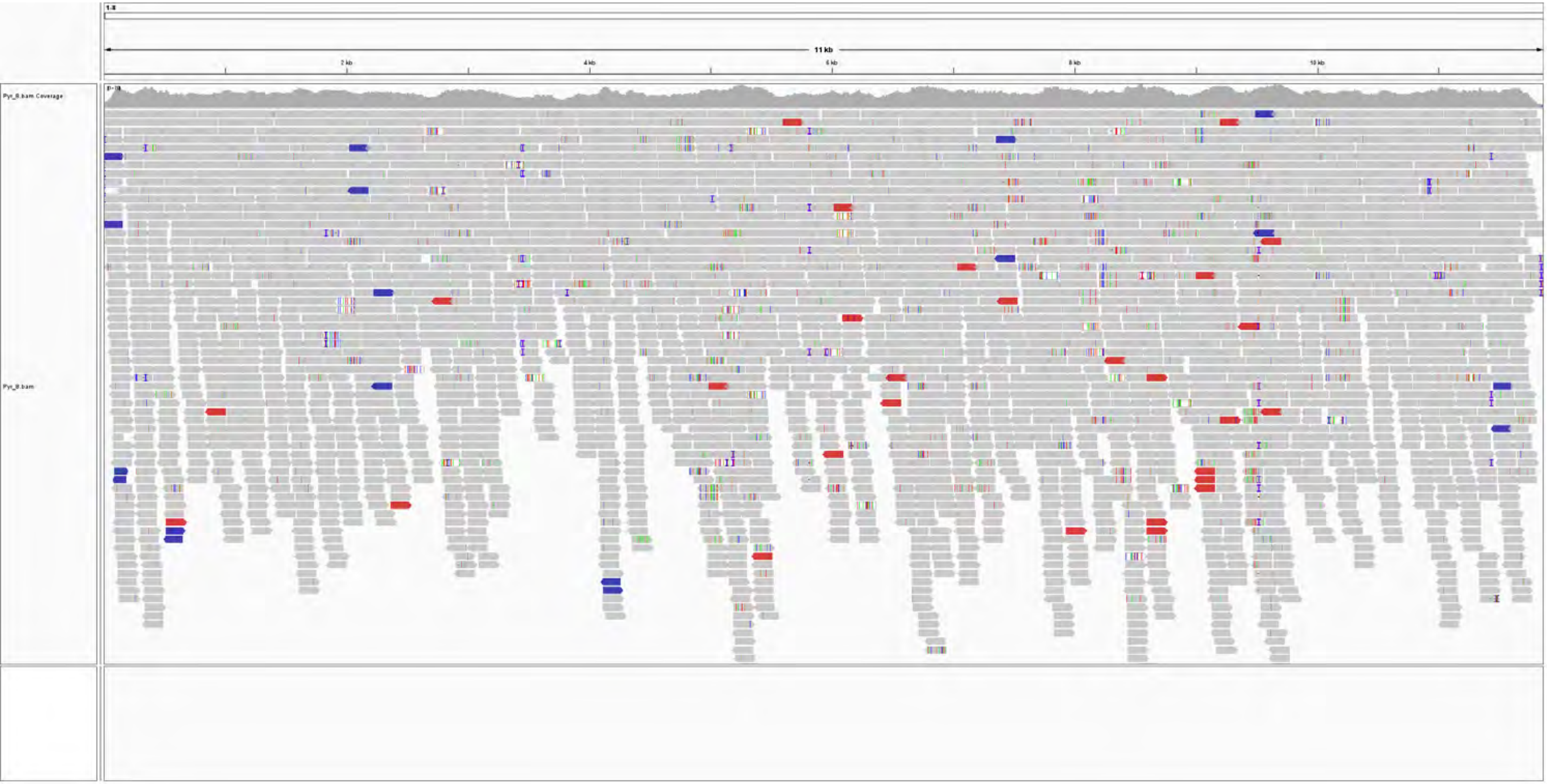

### Pyr_9

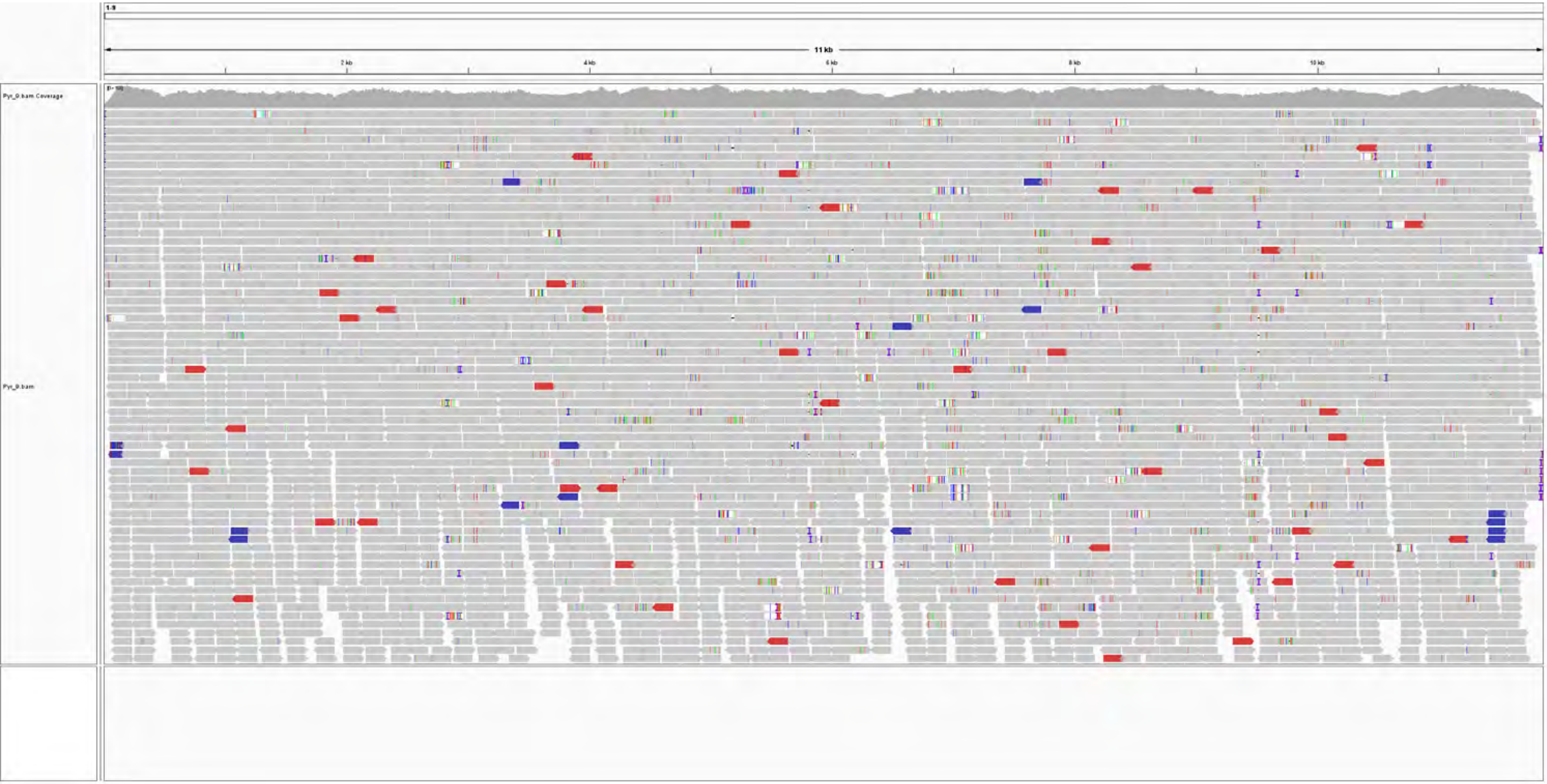

### Pyr_10

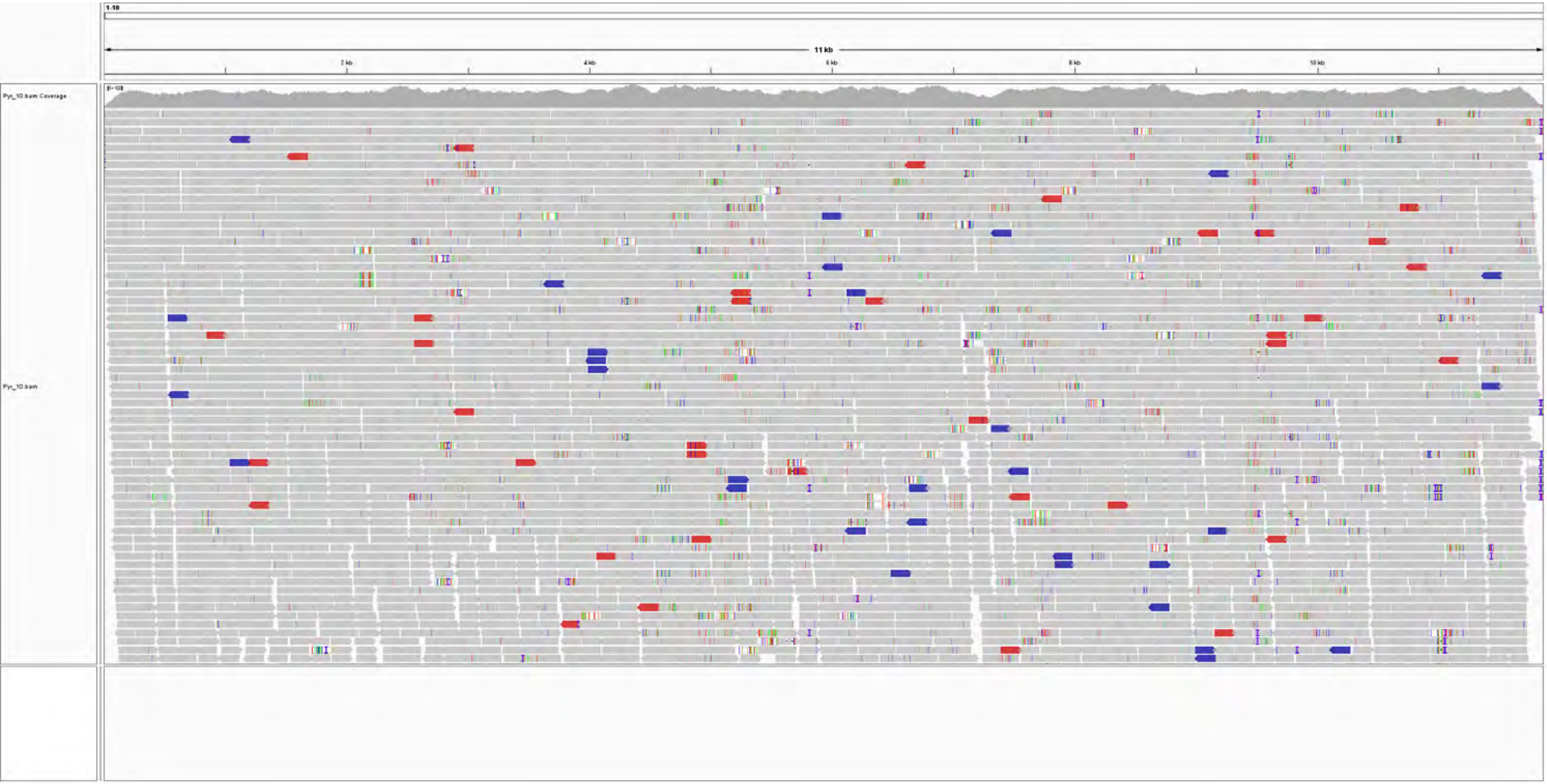

### Pyr_11

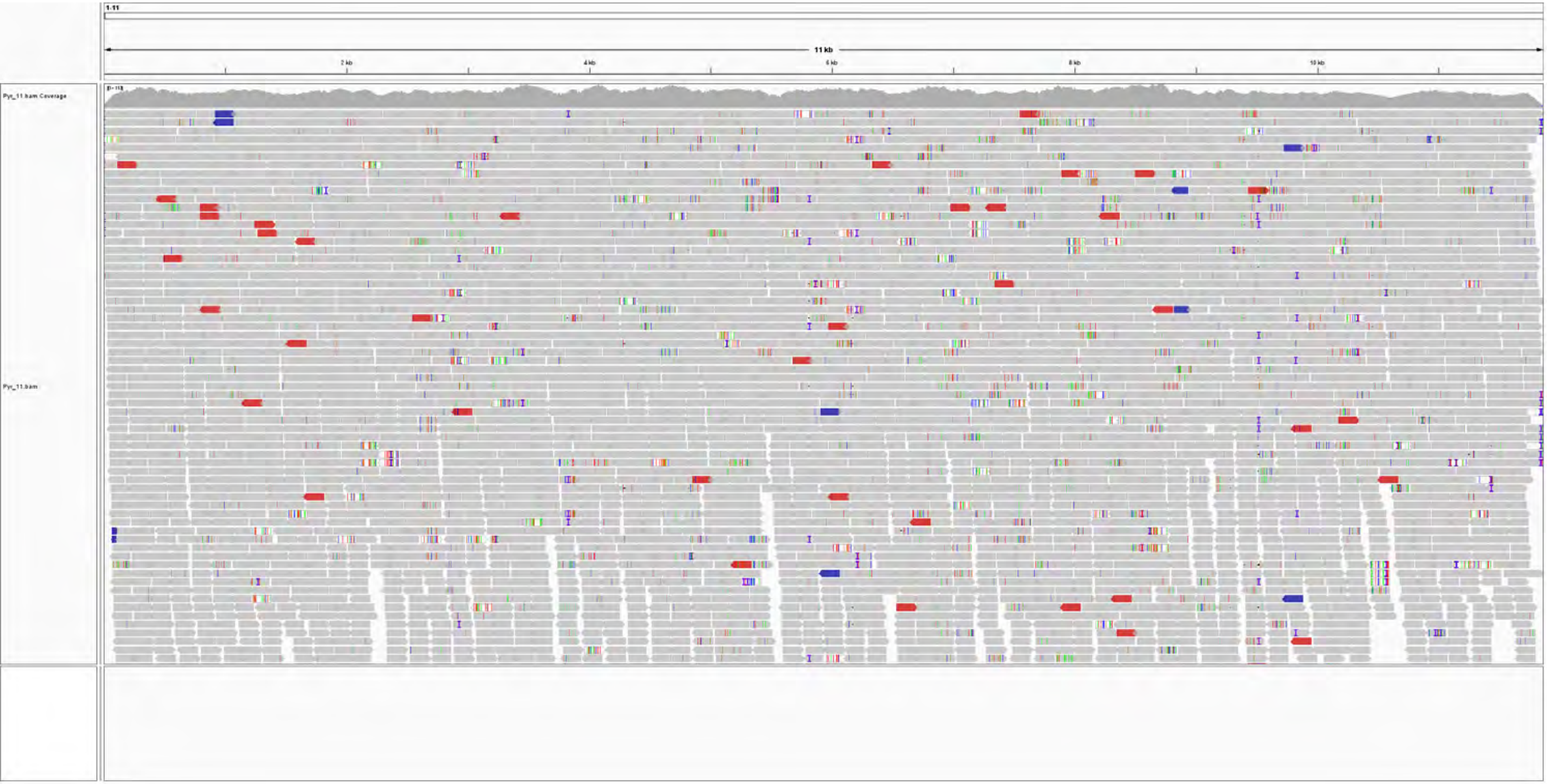

### Pyr_12

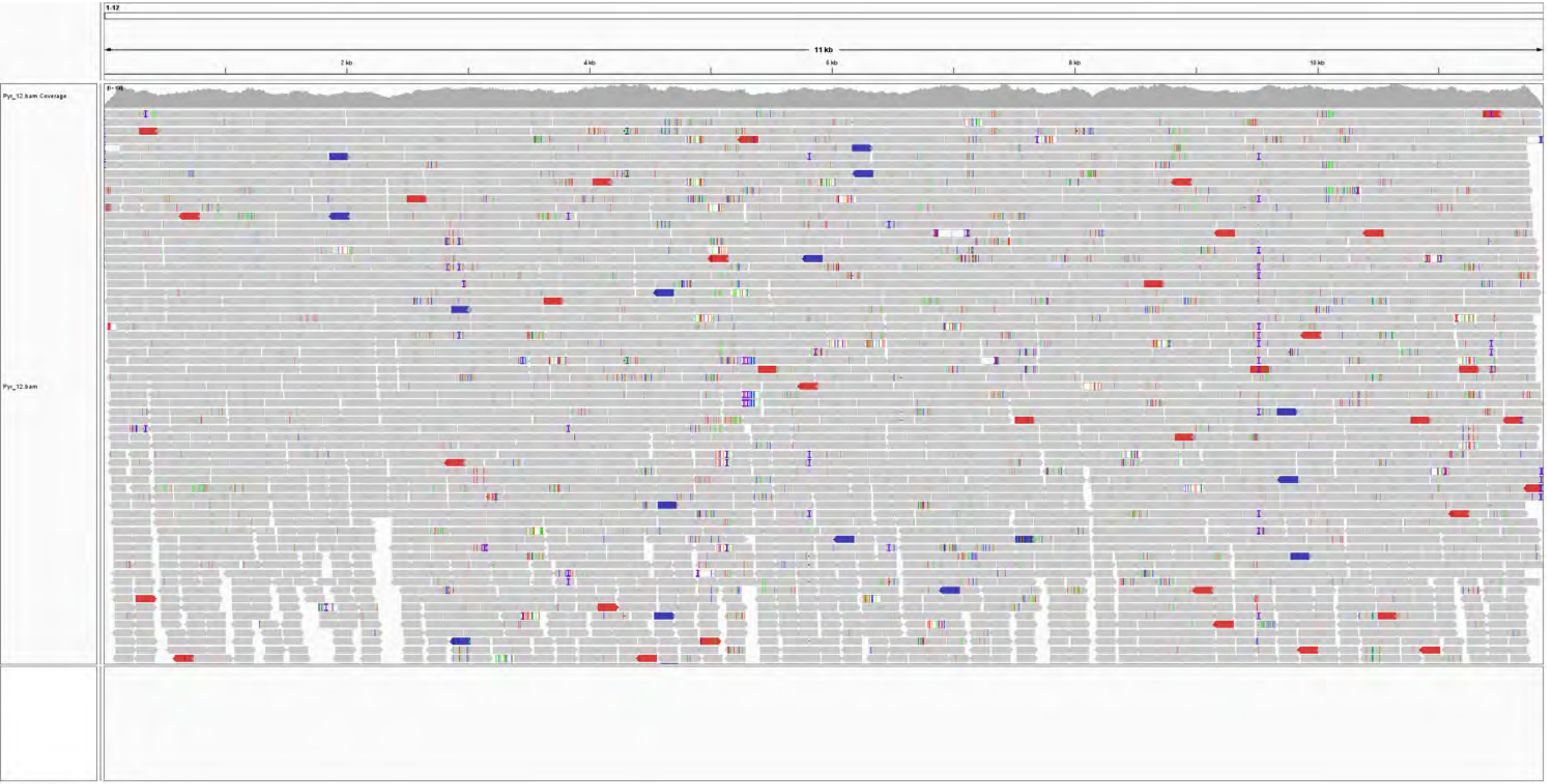

### Pyr_13

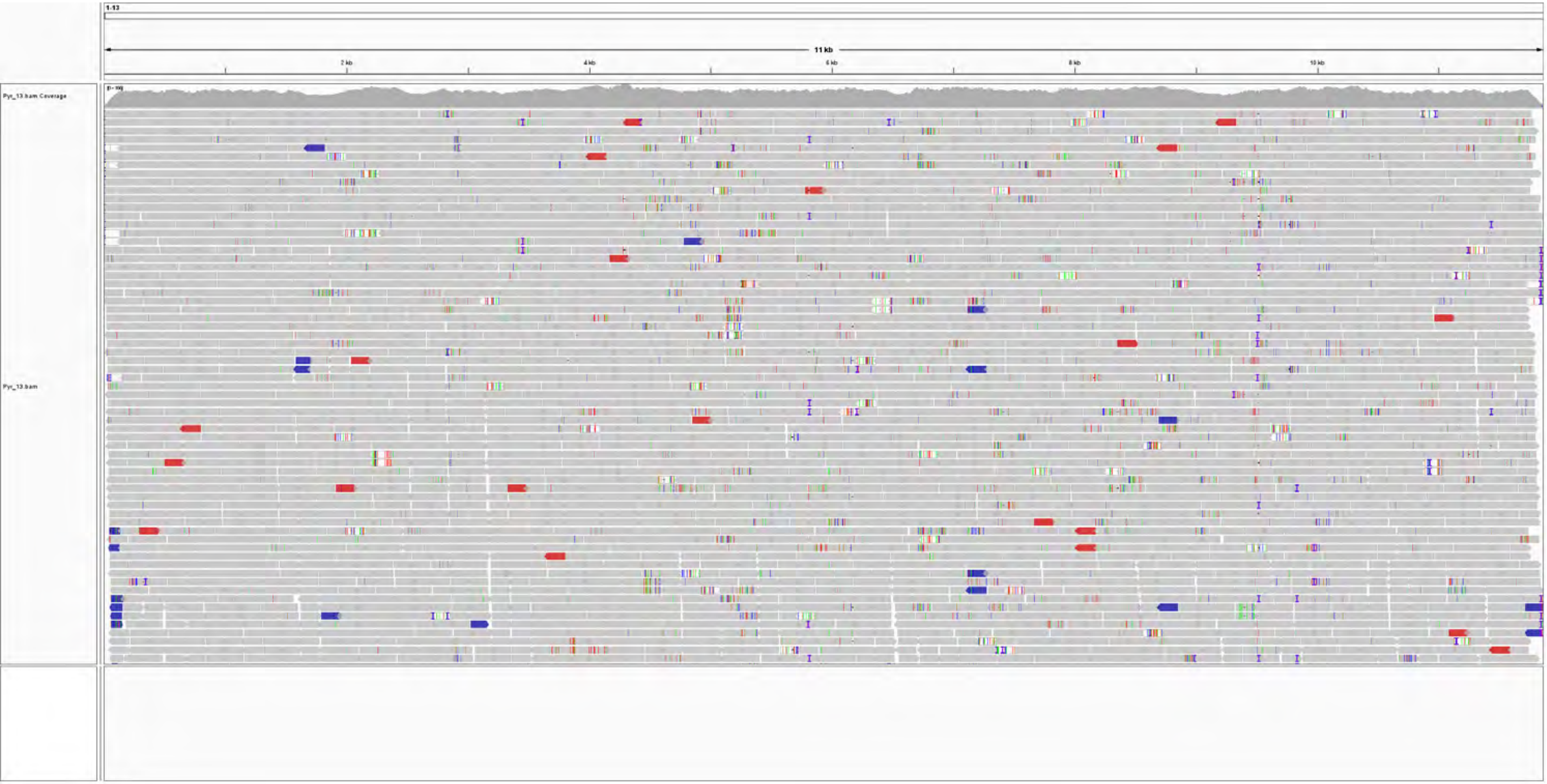

### Pyr_14

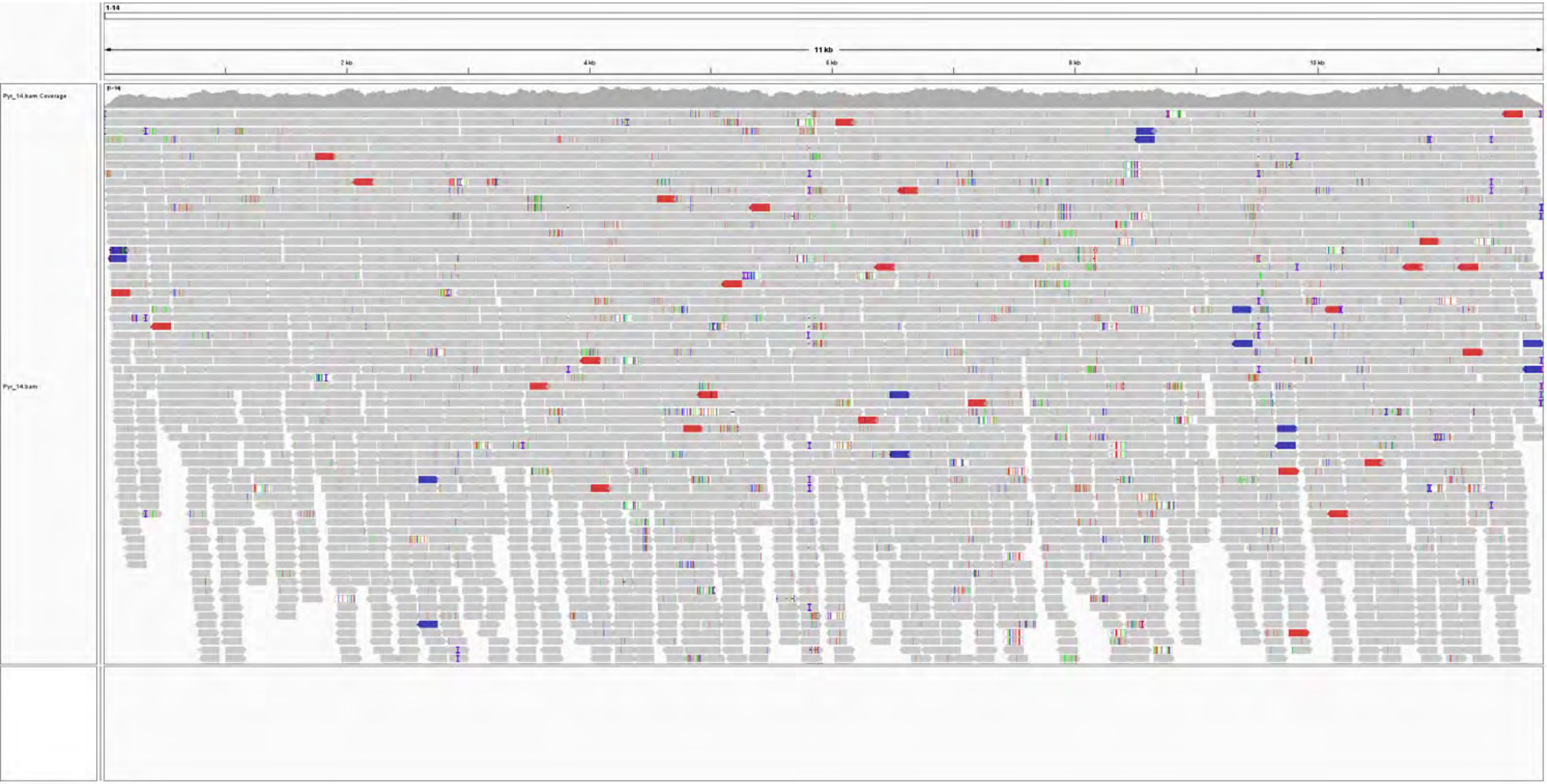

### Pyr_15

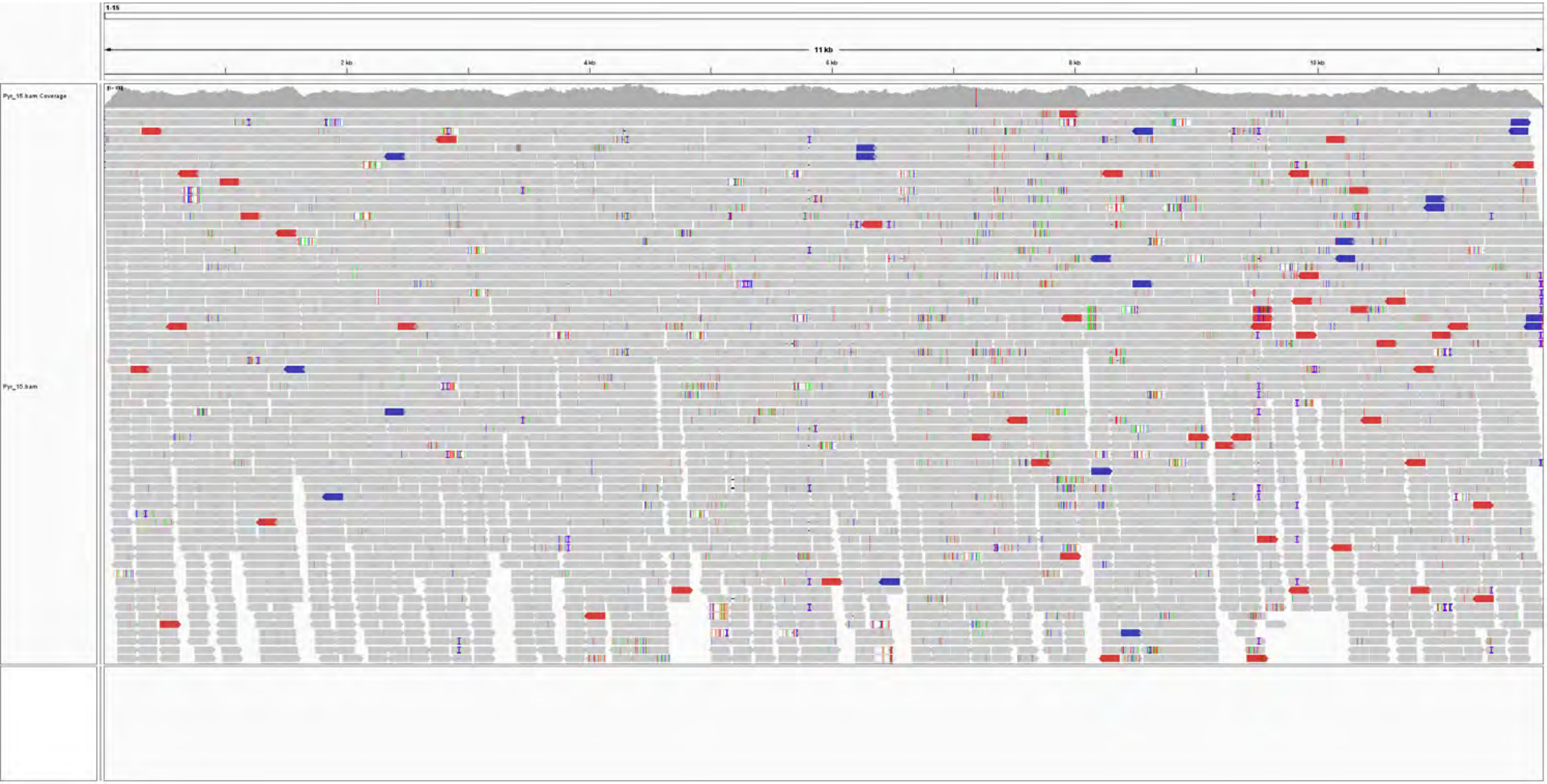

### Pyr_16

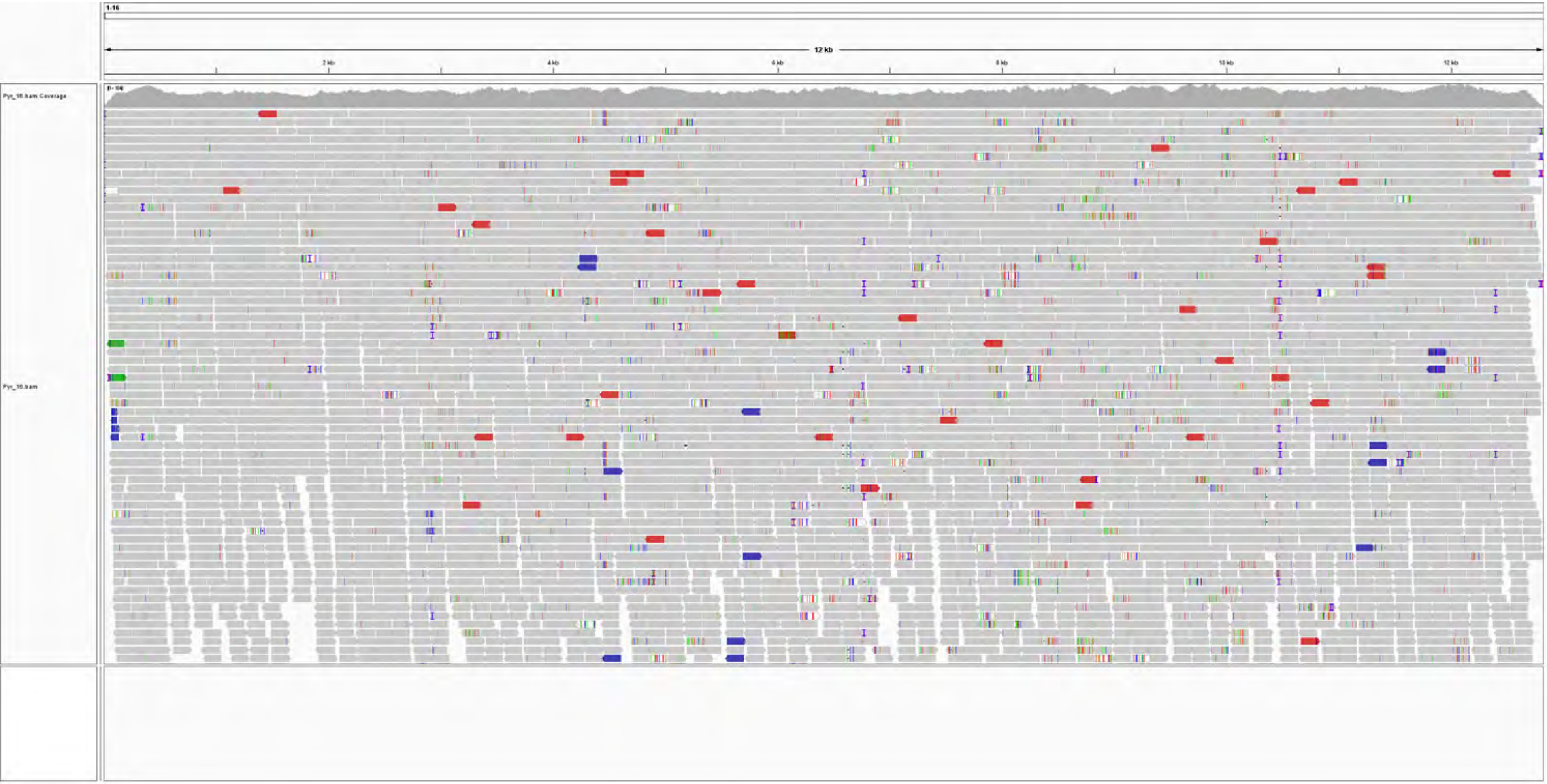

1-16
12 kb
2 kb
4 kb
6 kb
8 kb
10 kb
12 kb
Pyr_16.bam Coverage
Pyr_16.bam

### Pyr_17

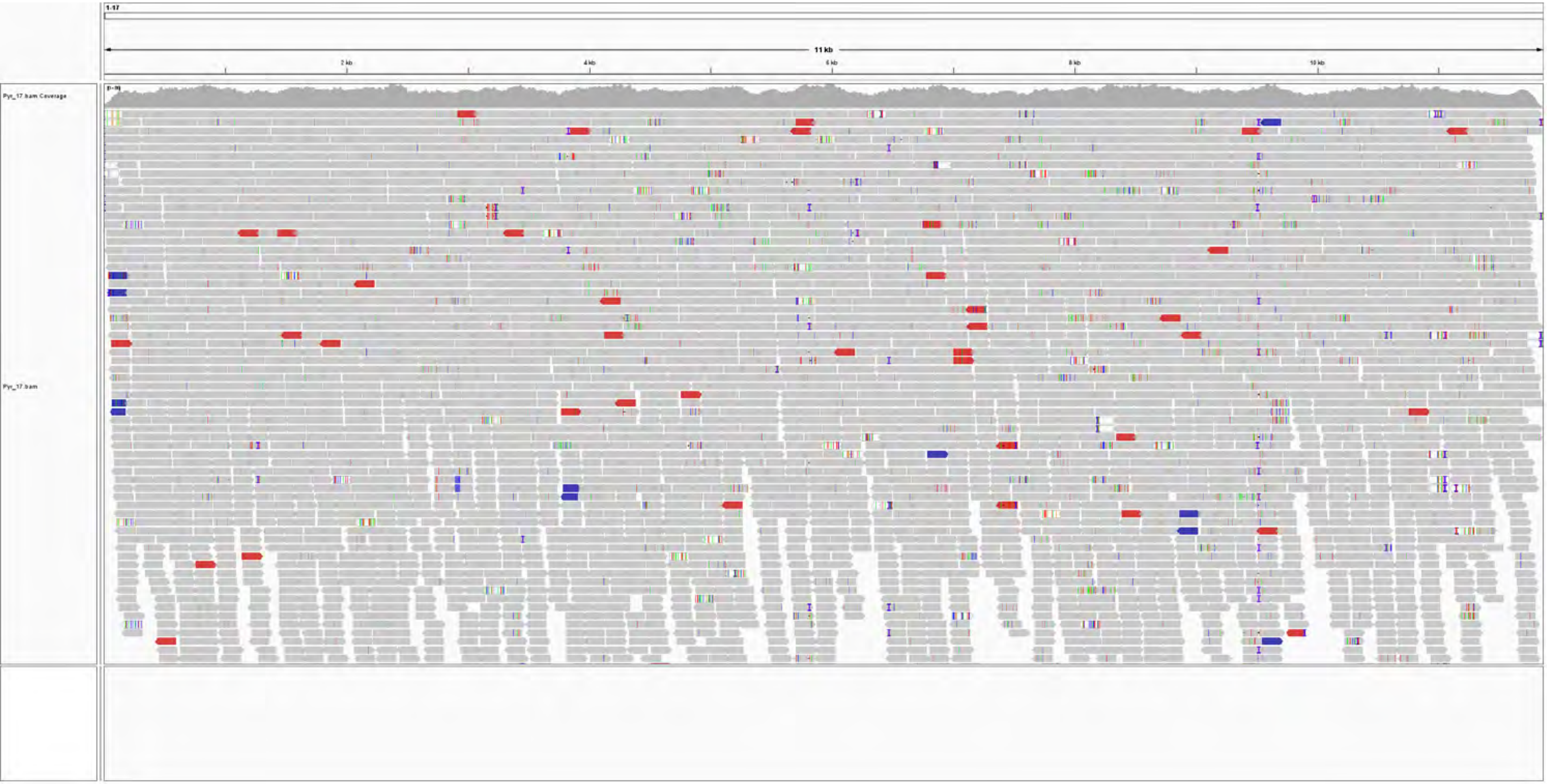

### Pyr_18

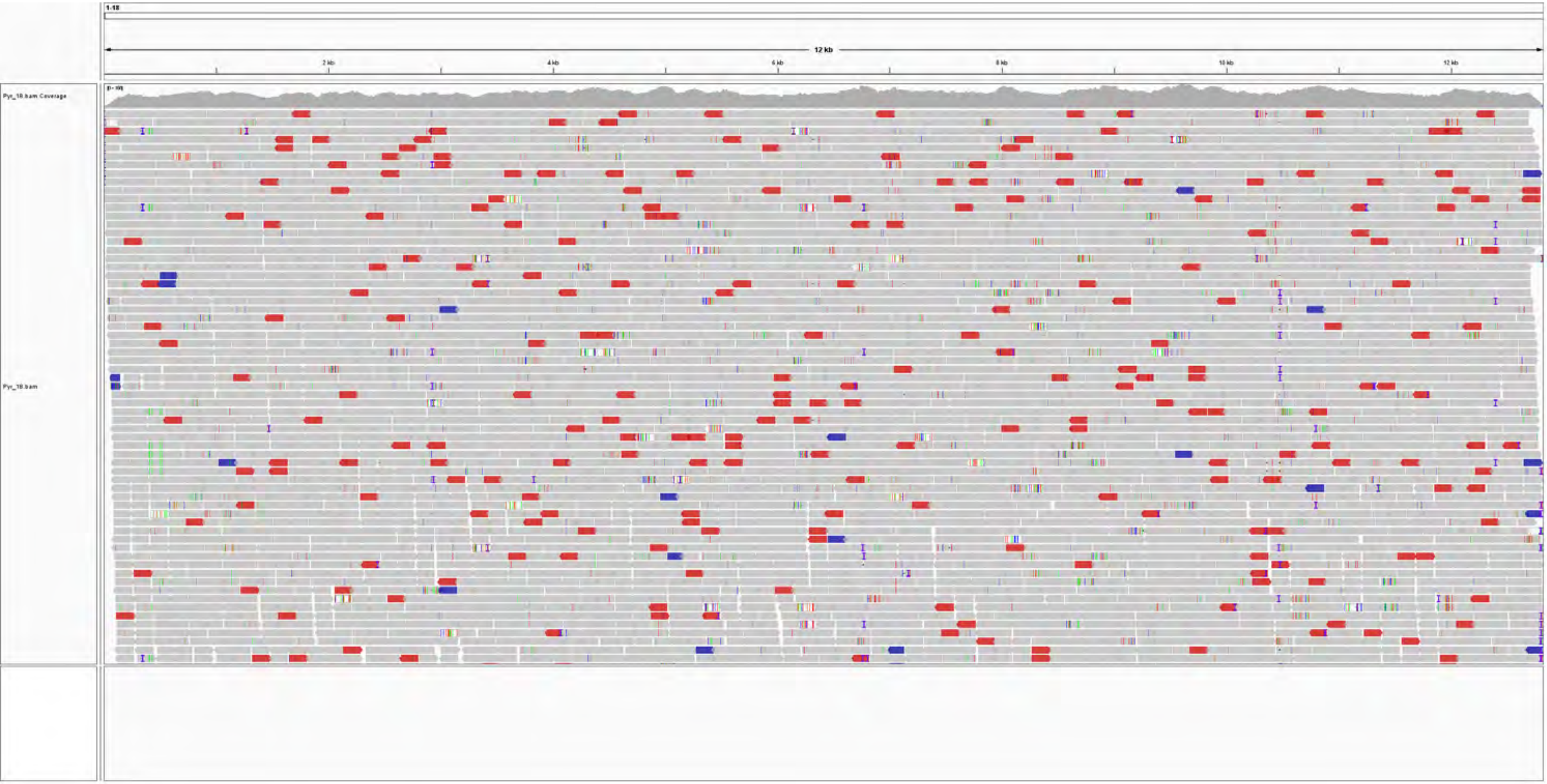

### Pyr_19

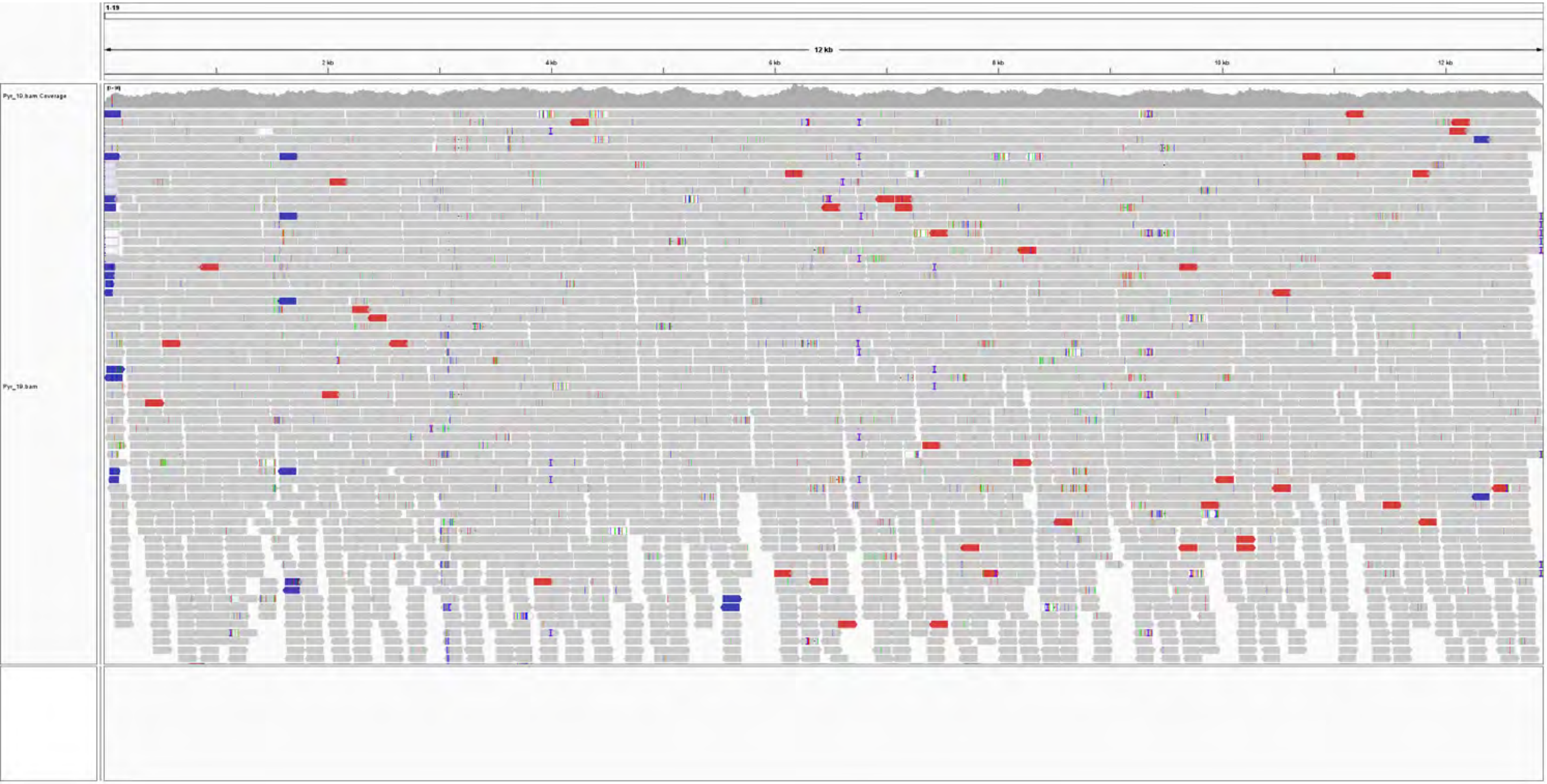

1-19
12 kb
2 kb
4 kb
6 kb
8 kb
10 kb
12 kb
Pyr_19.bam Coverage
Pyr_19.bam

### Pyr_20

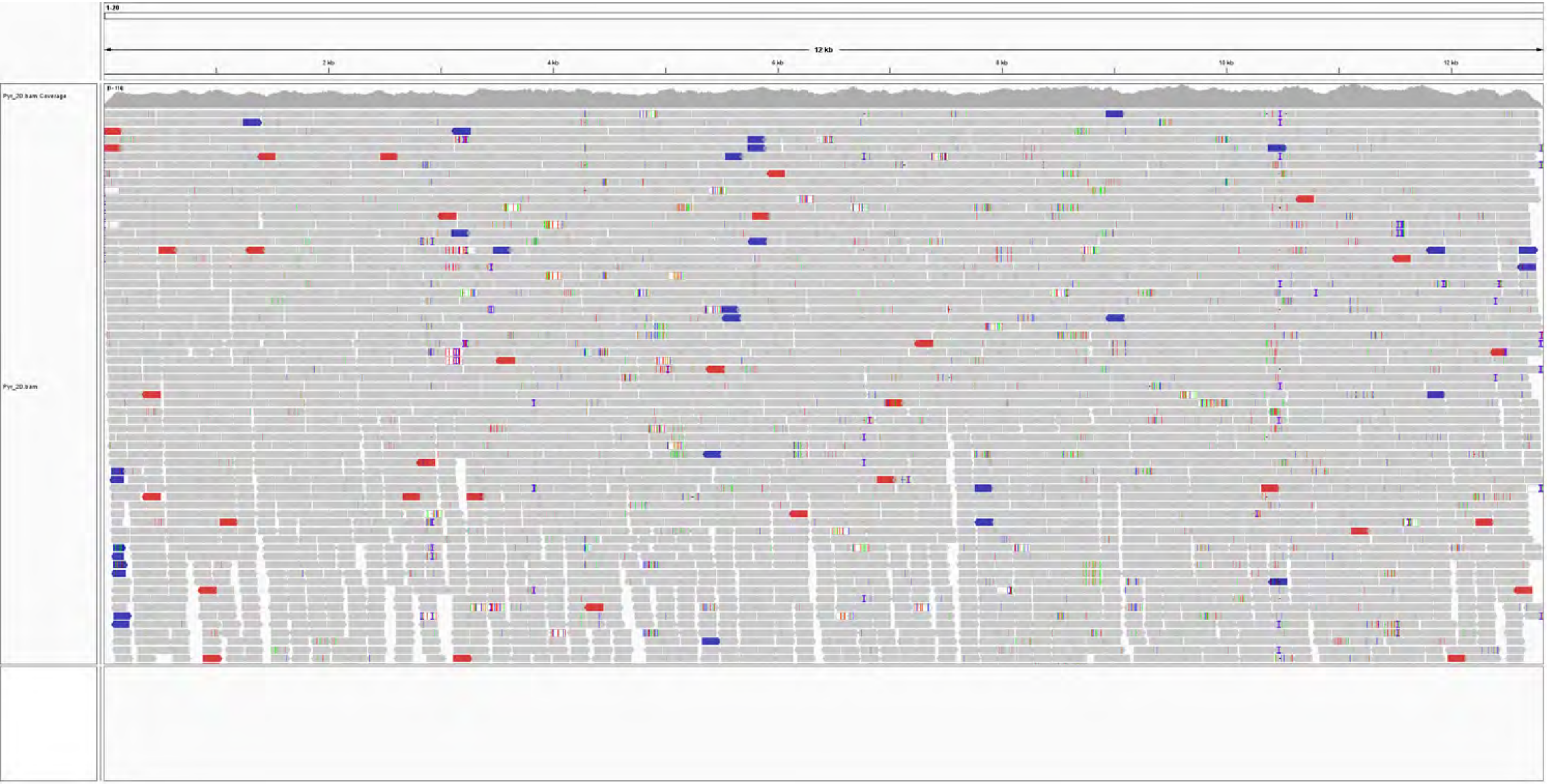

1-20
12 kb
2 kb
4 kb
6 kb
8 kb
10 kb
12 kb
Pyr_20.bam Coverage
Pyr_20.bam

### Pyr_21

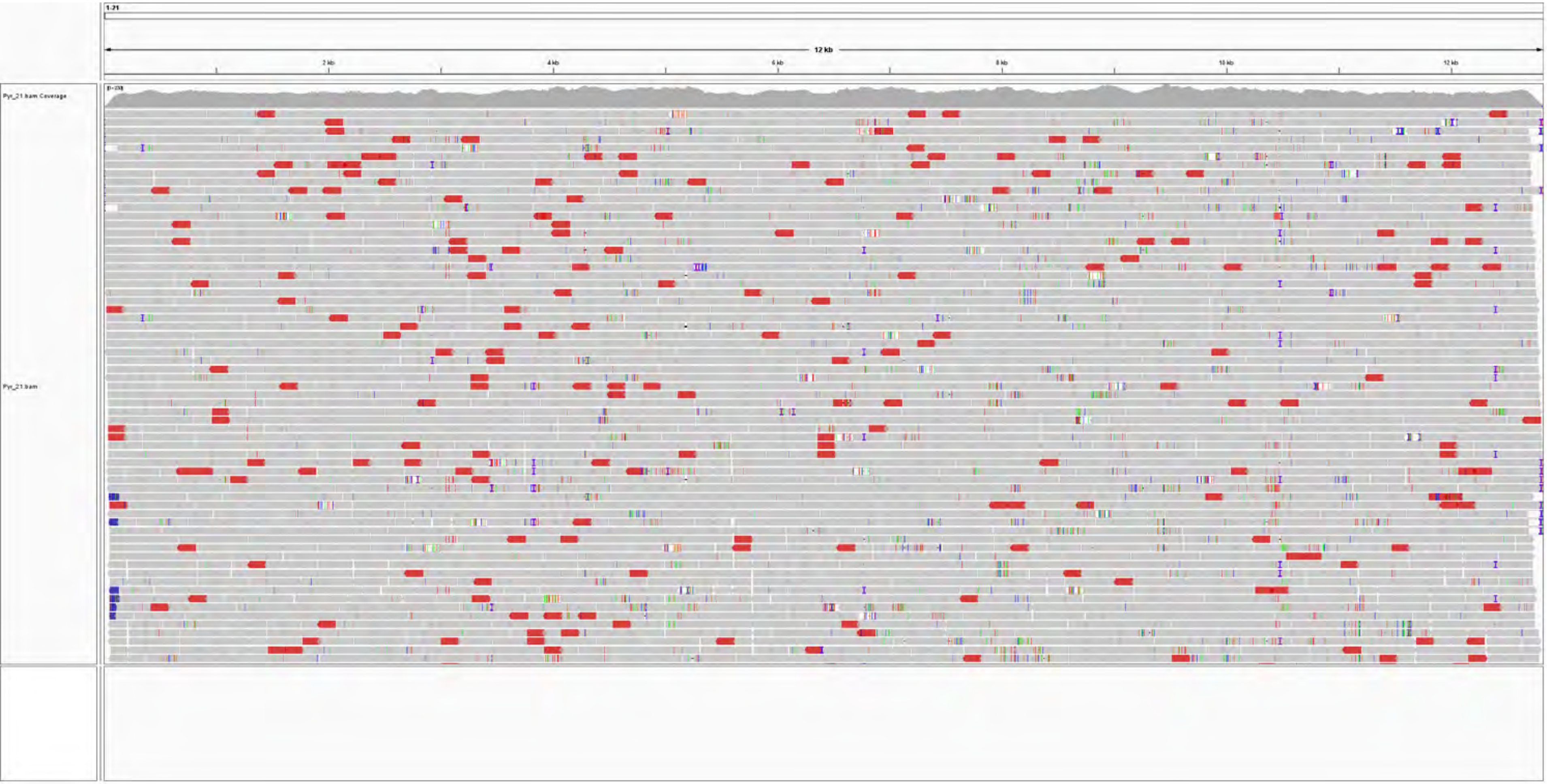

### Pyr_22

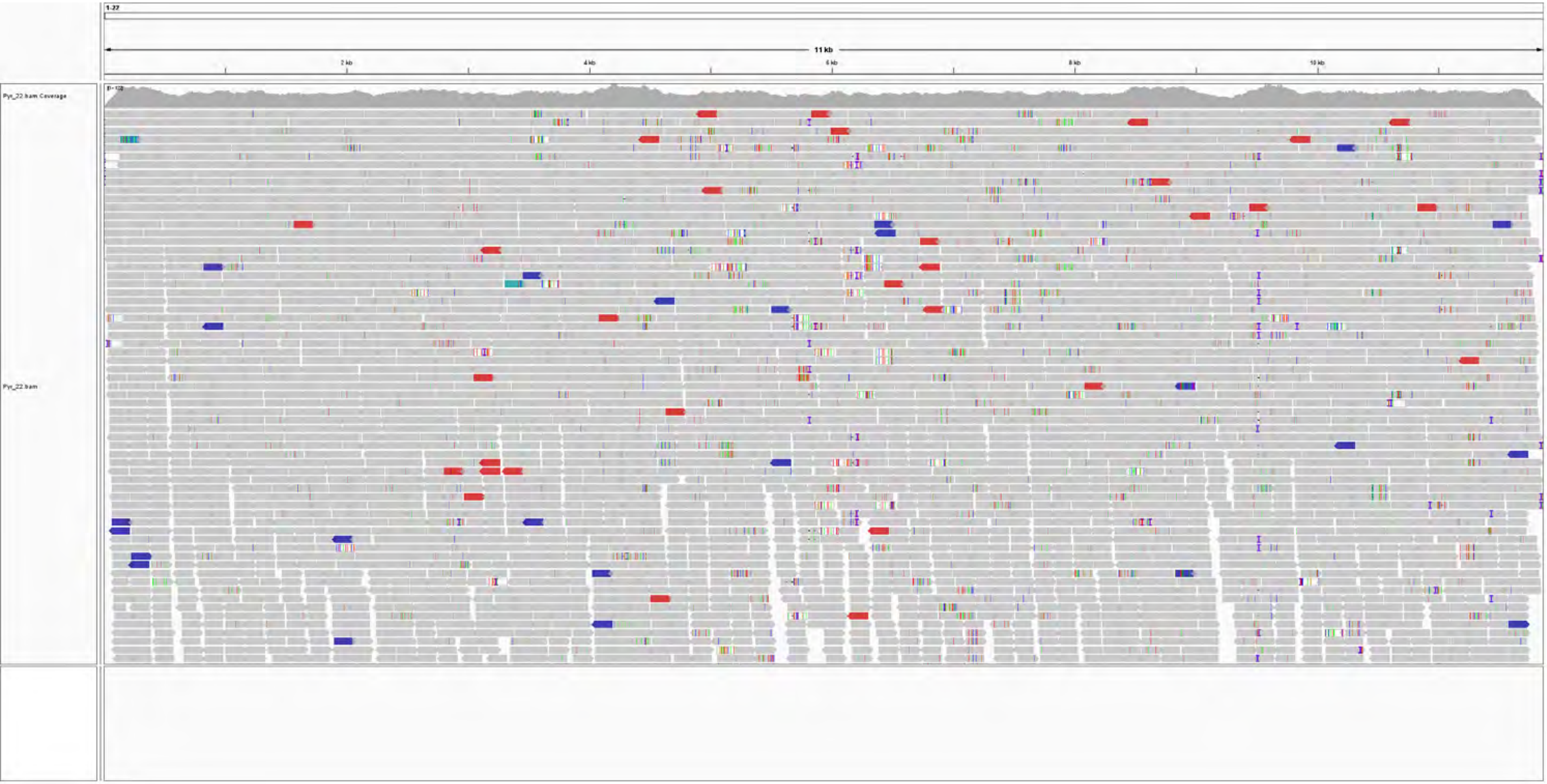

### Pyr_23

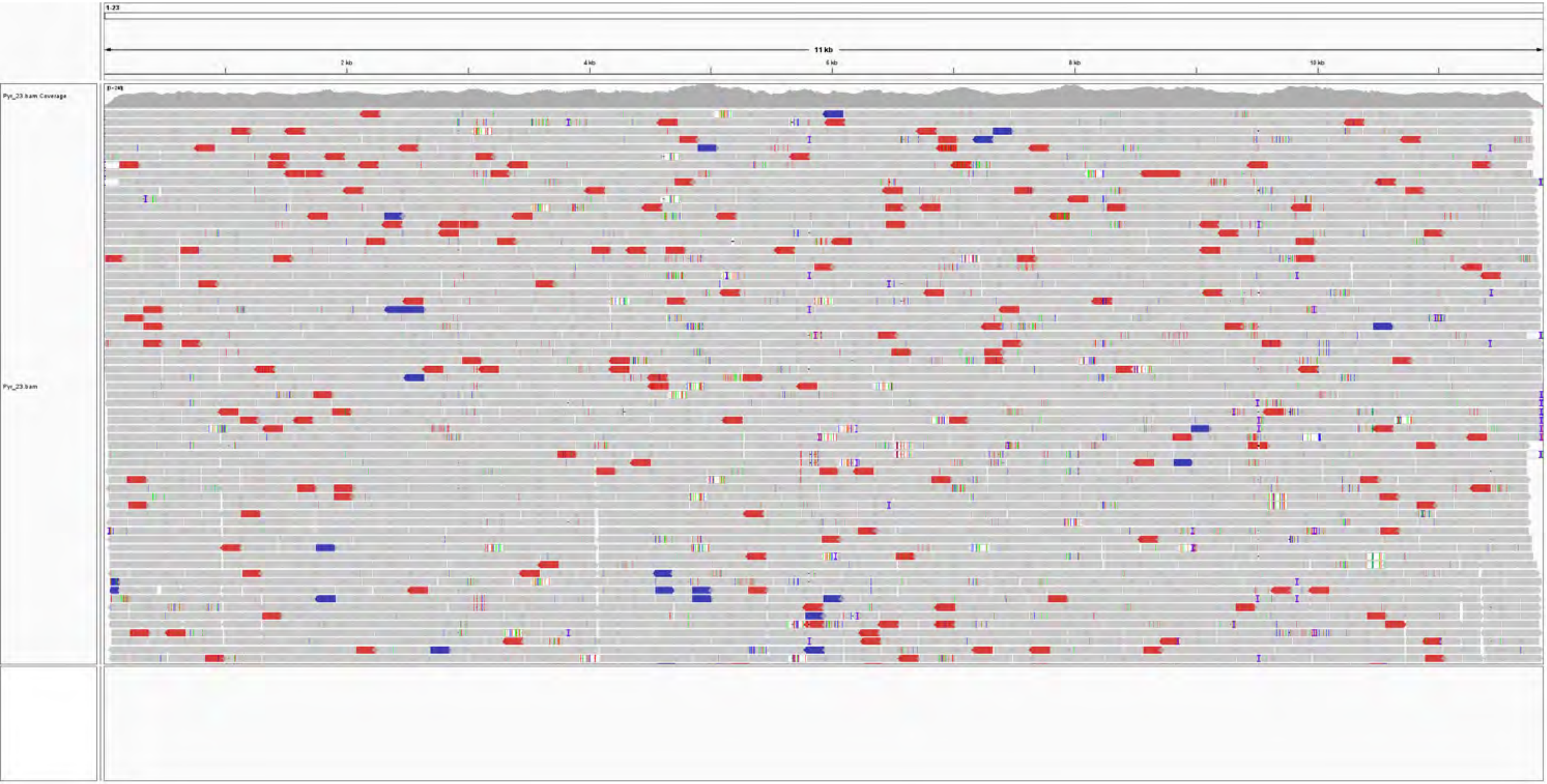

### Pyr_24

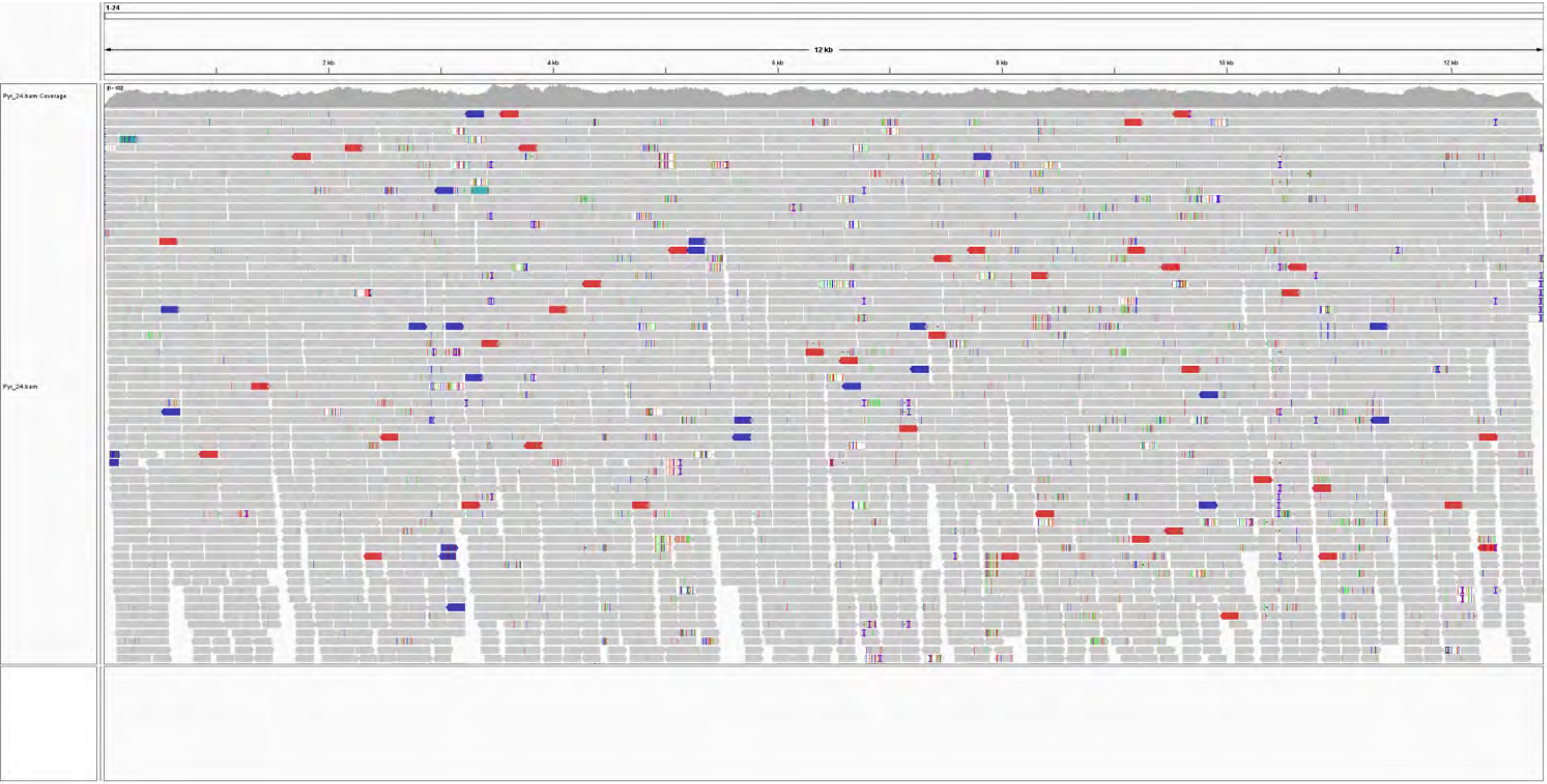
1-24
12 kb
2 kb
4 kb
6 kb
8 kb
10 kb
12 kb
Pyr_24.bam Coverage
Pyr_24.bam

### Pyr_25

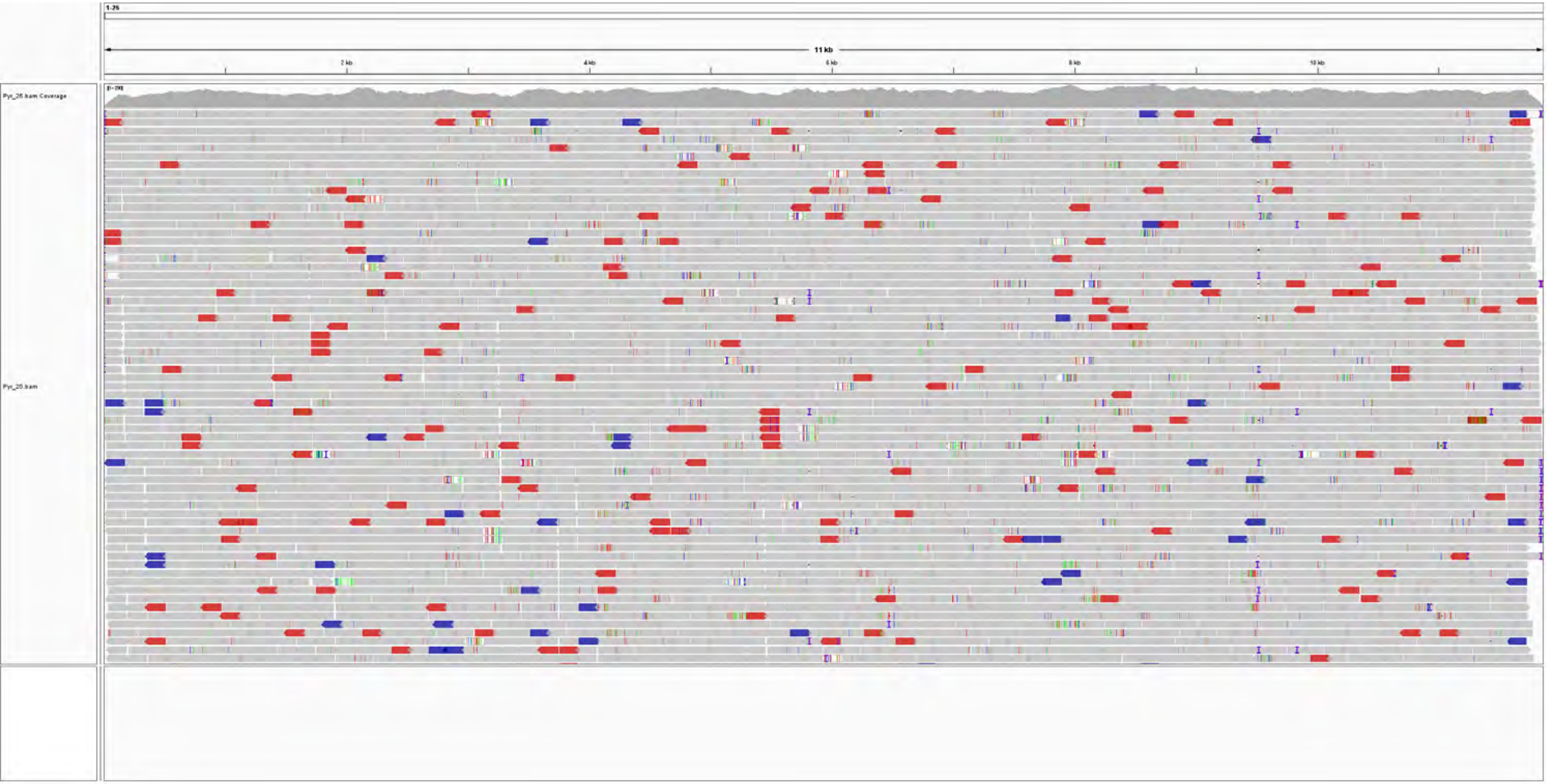

### Pyr_26

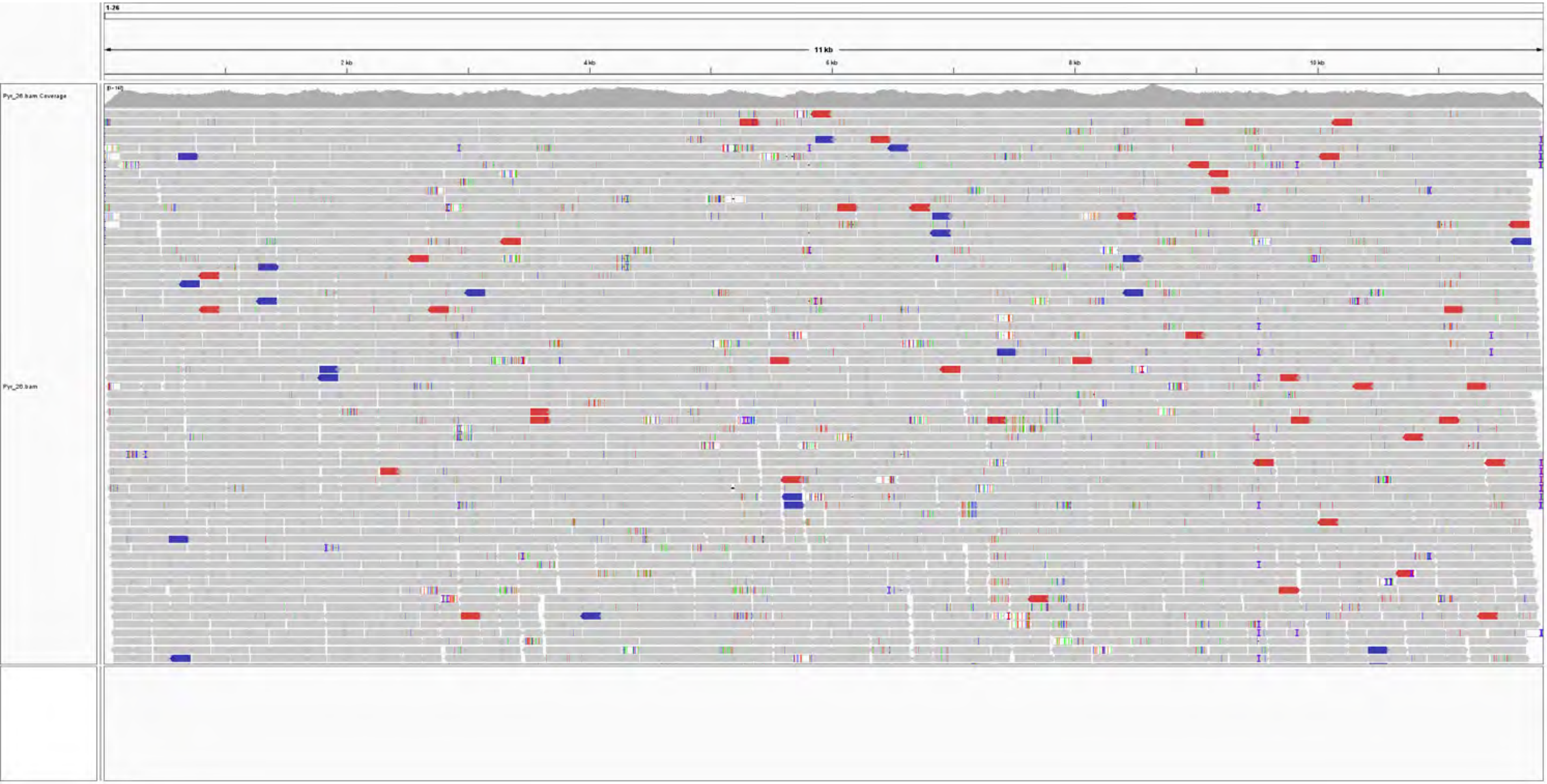

### Pyr_27

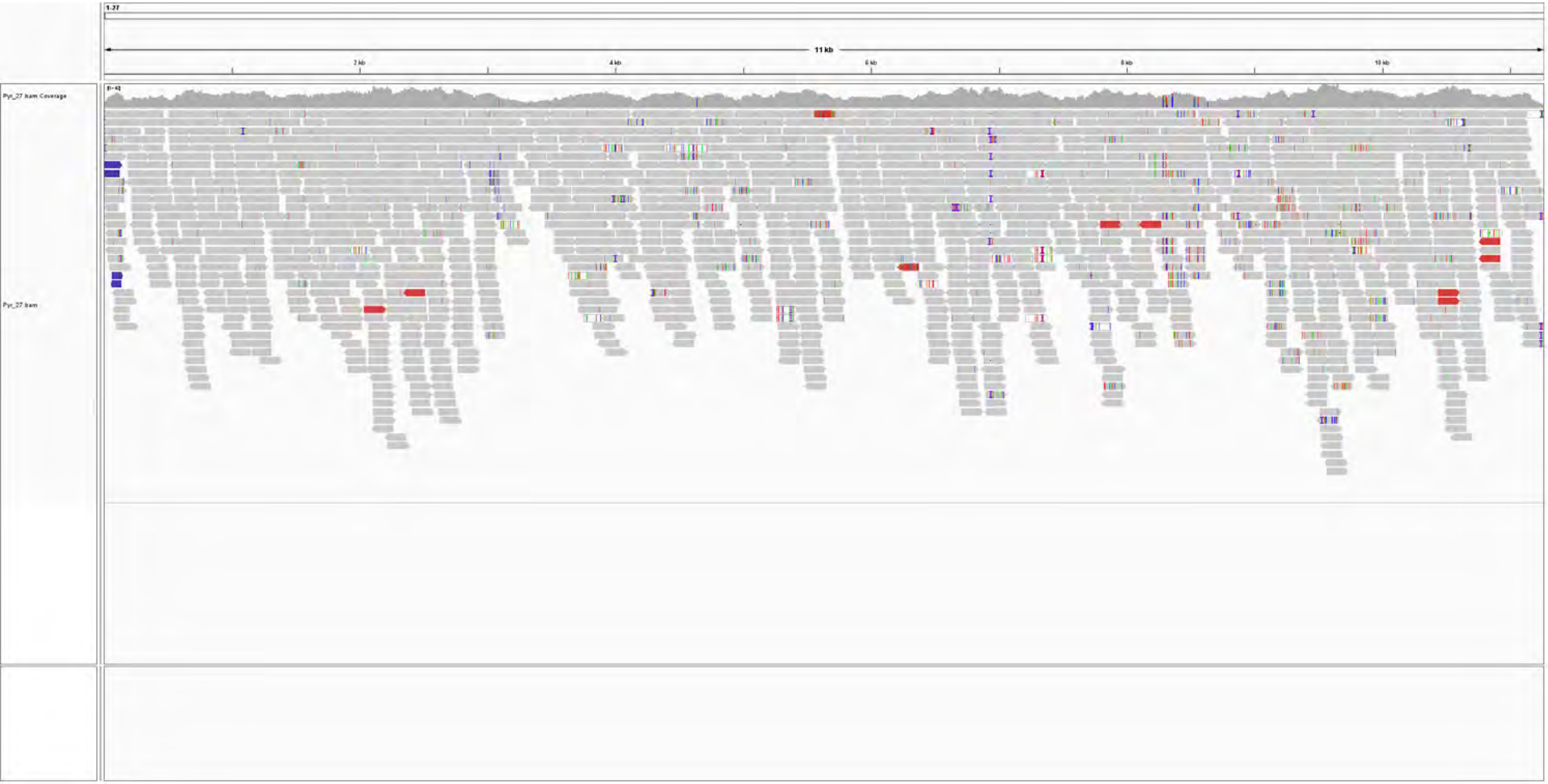

### Pyr_28

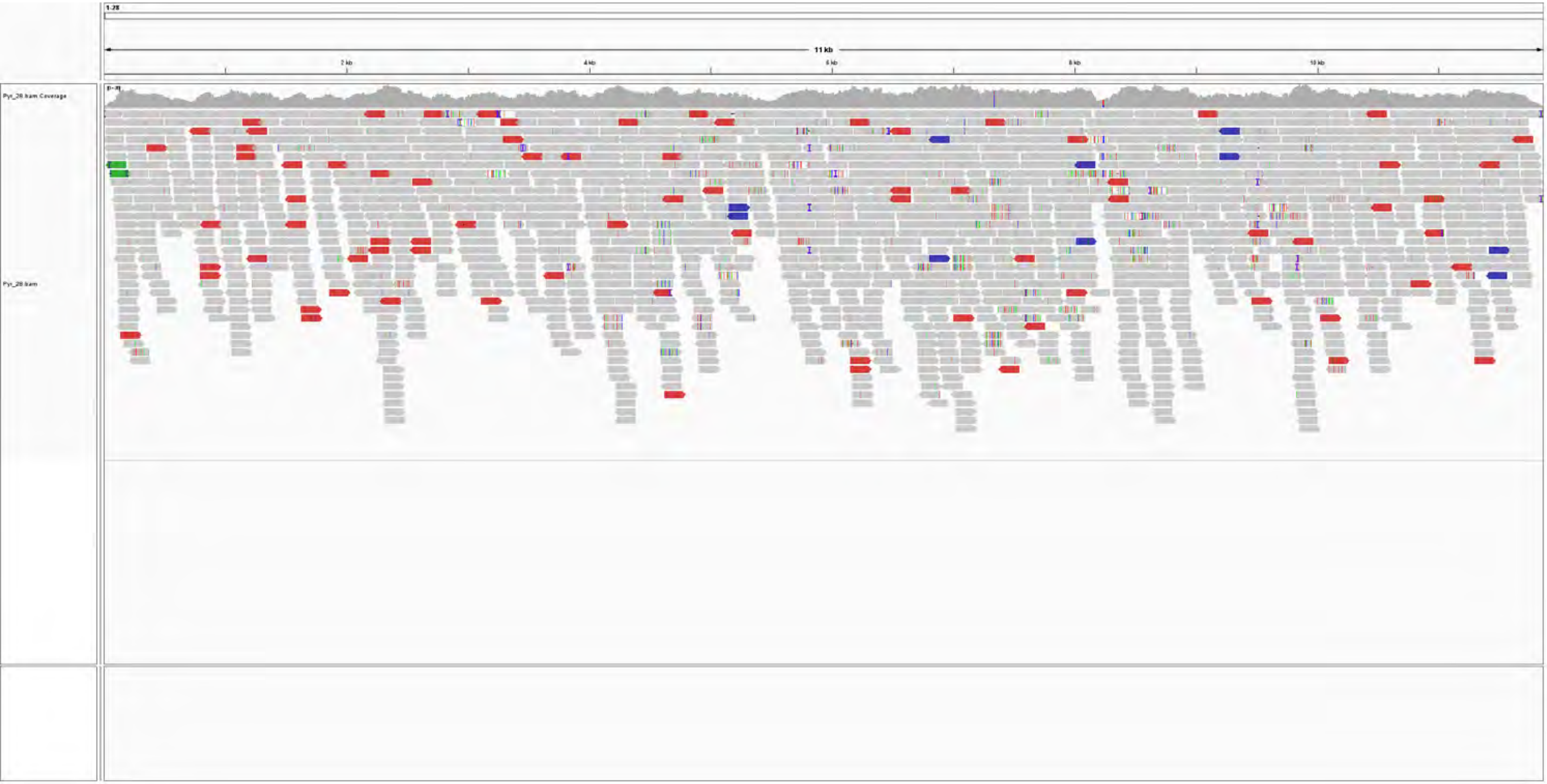

### Pyr_29

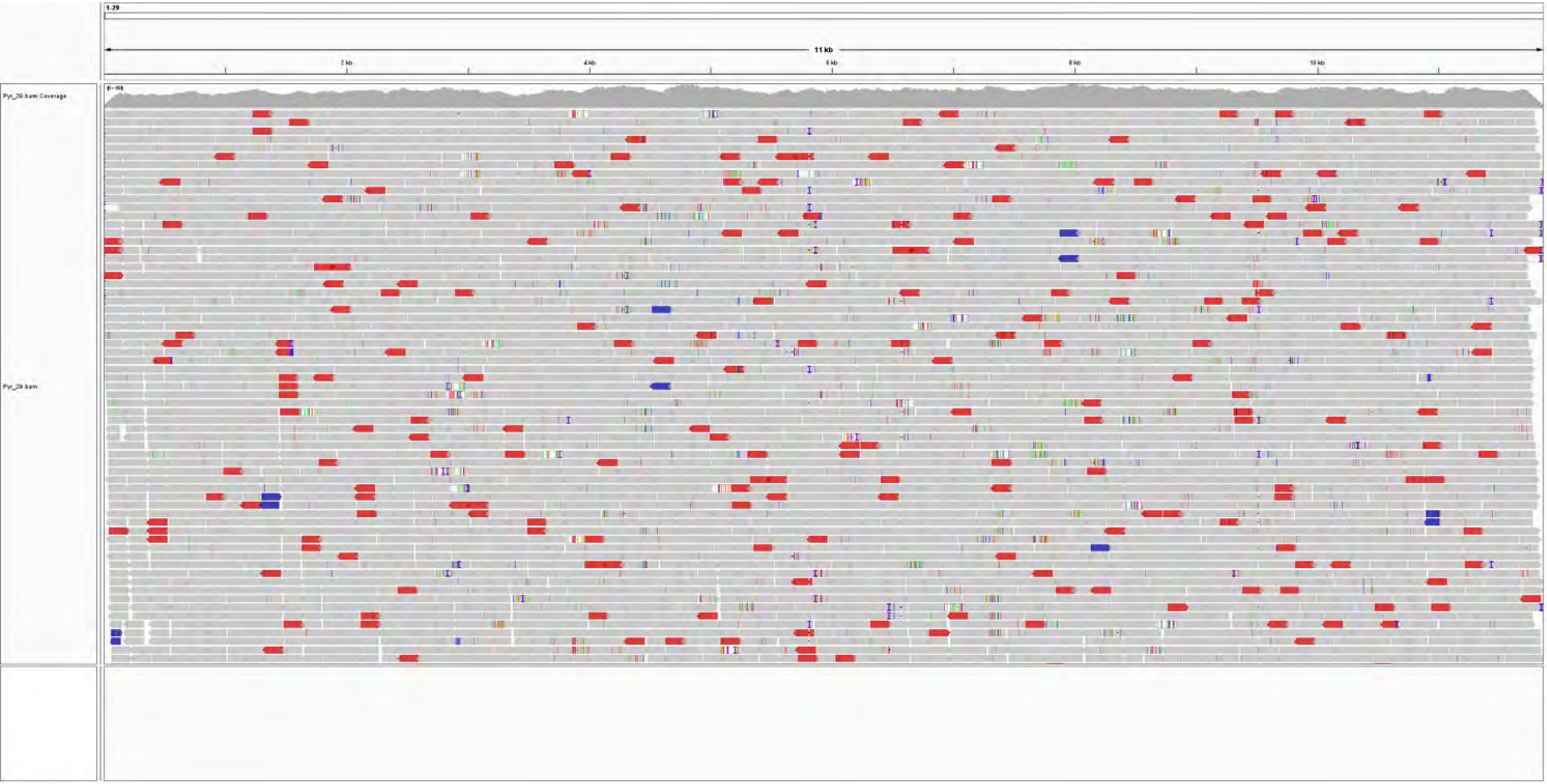

### Pyr_30

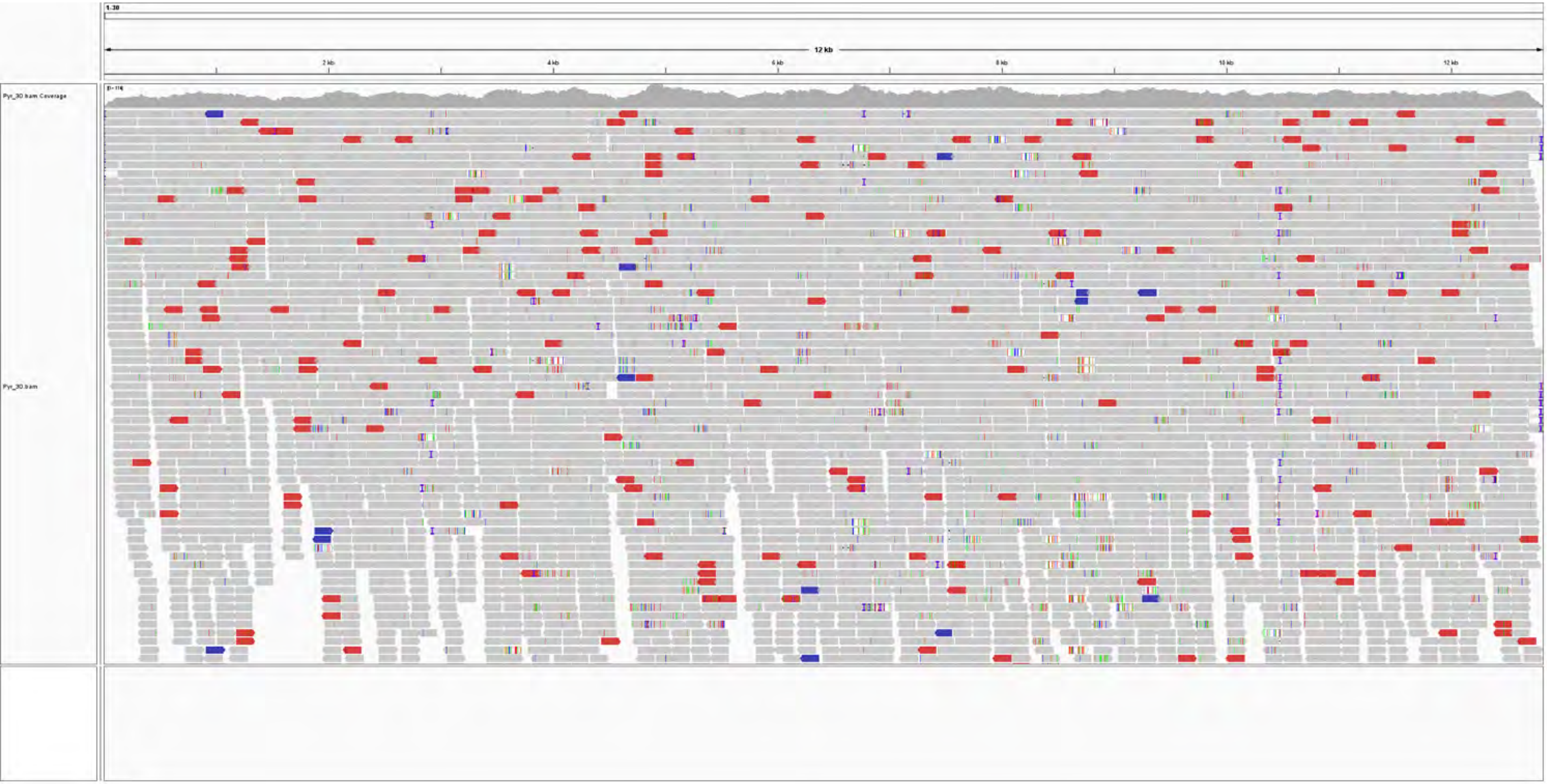

1:30
12 kb
2 kb
4 kb
6 kb
8 kb
10 kb
12 kb
Pyr_30.bam Coverage
[0-114]
Pyr_30.bam

### Pyr_33

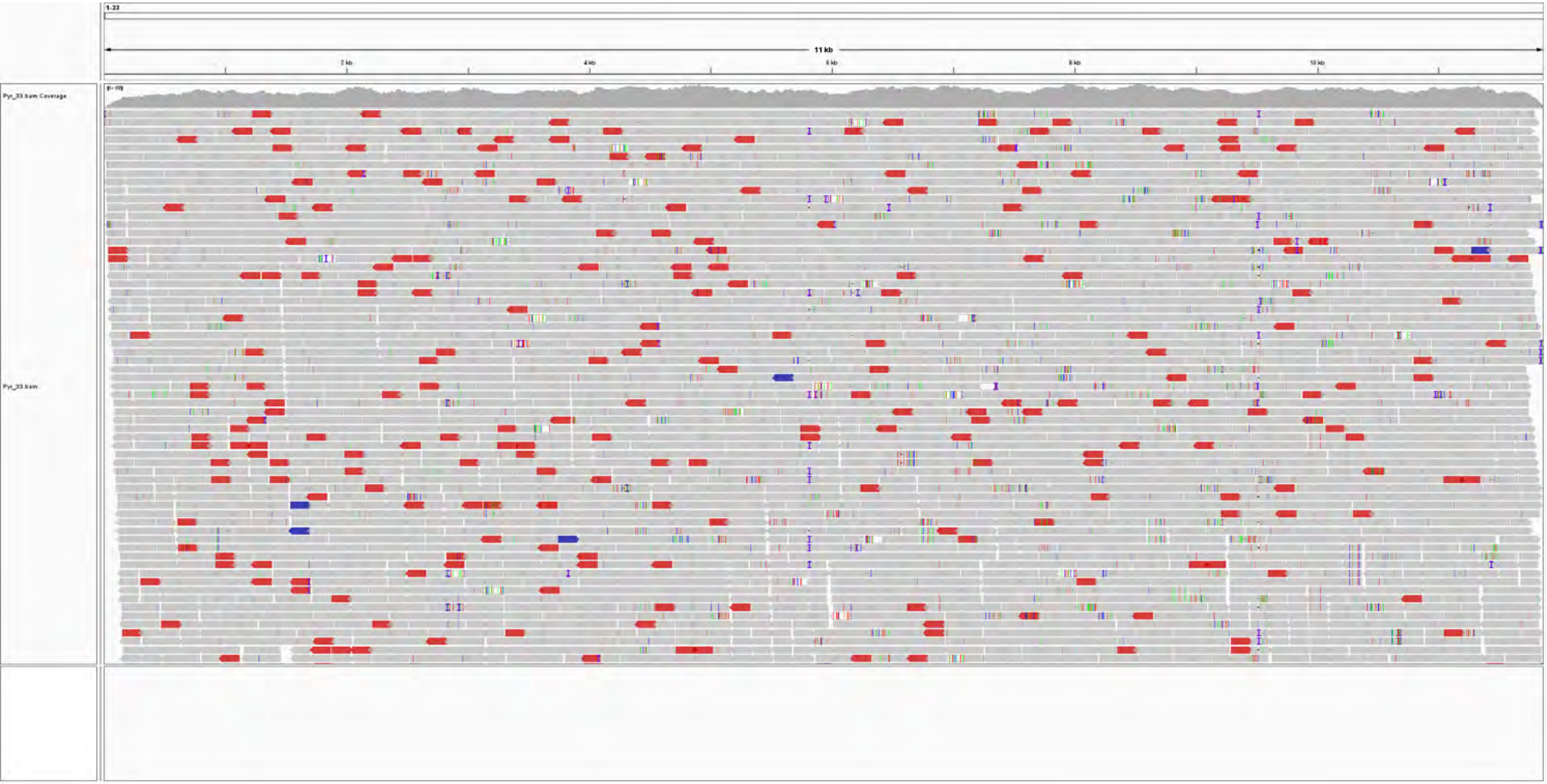

### Pyr_34

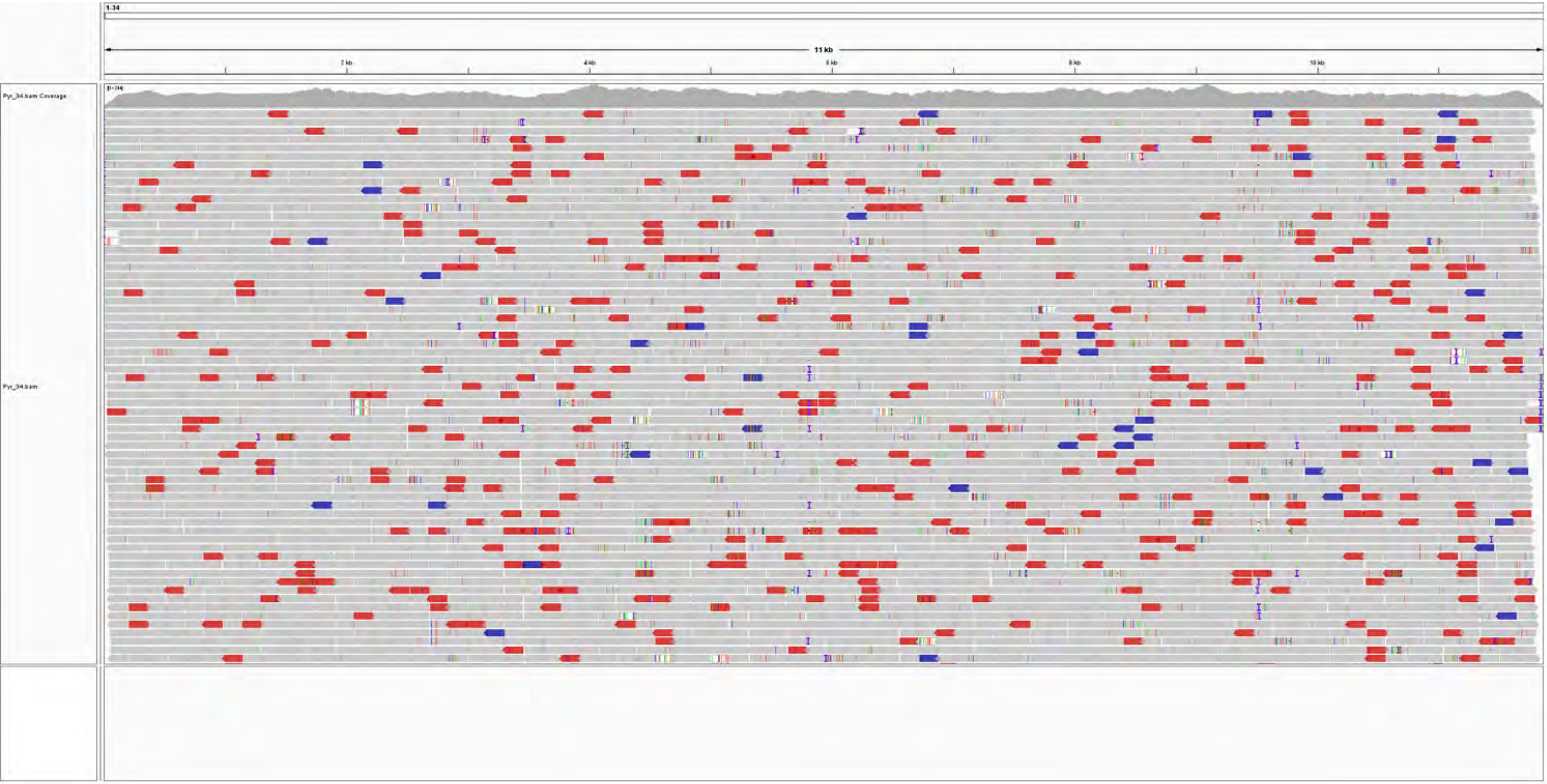

1-34
11 kb
2 kb
4 kb
6 kb
8 kb
10 kb
Pyr_34.bam Coverage
[0-24]
Pyr_34.bam

### Pyr_36

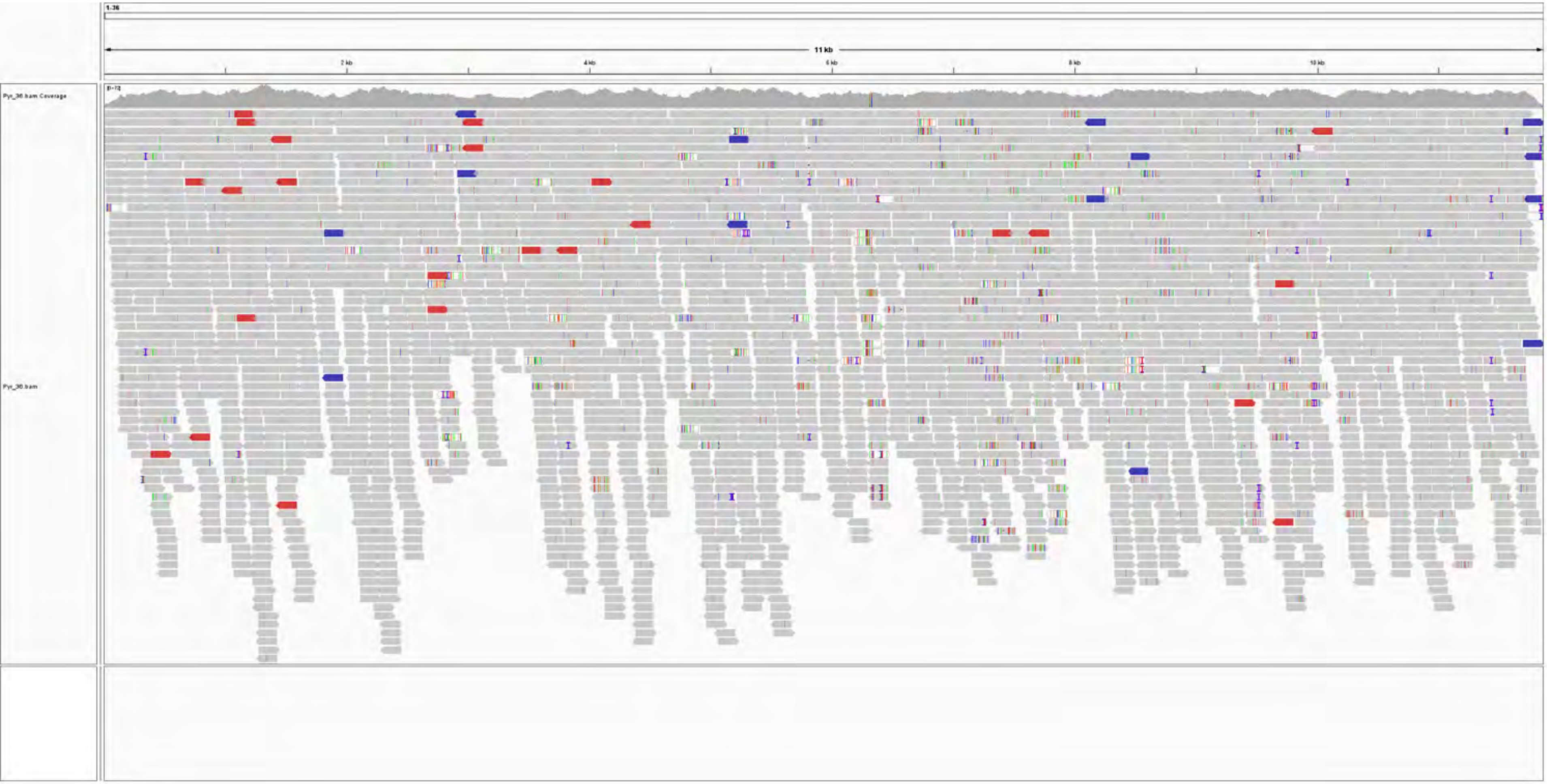

### Pyr_38

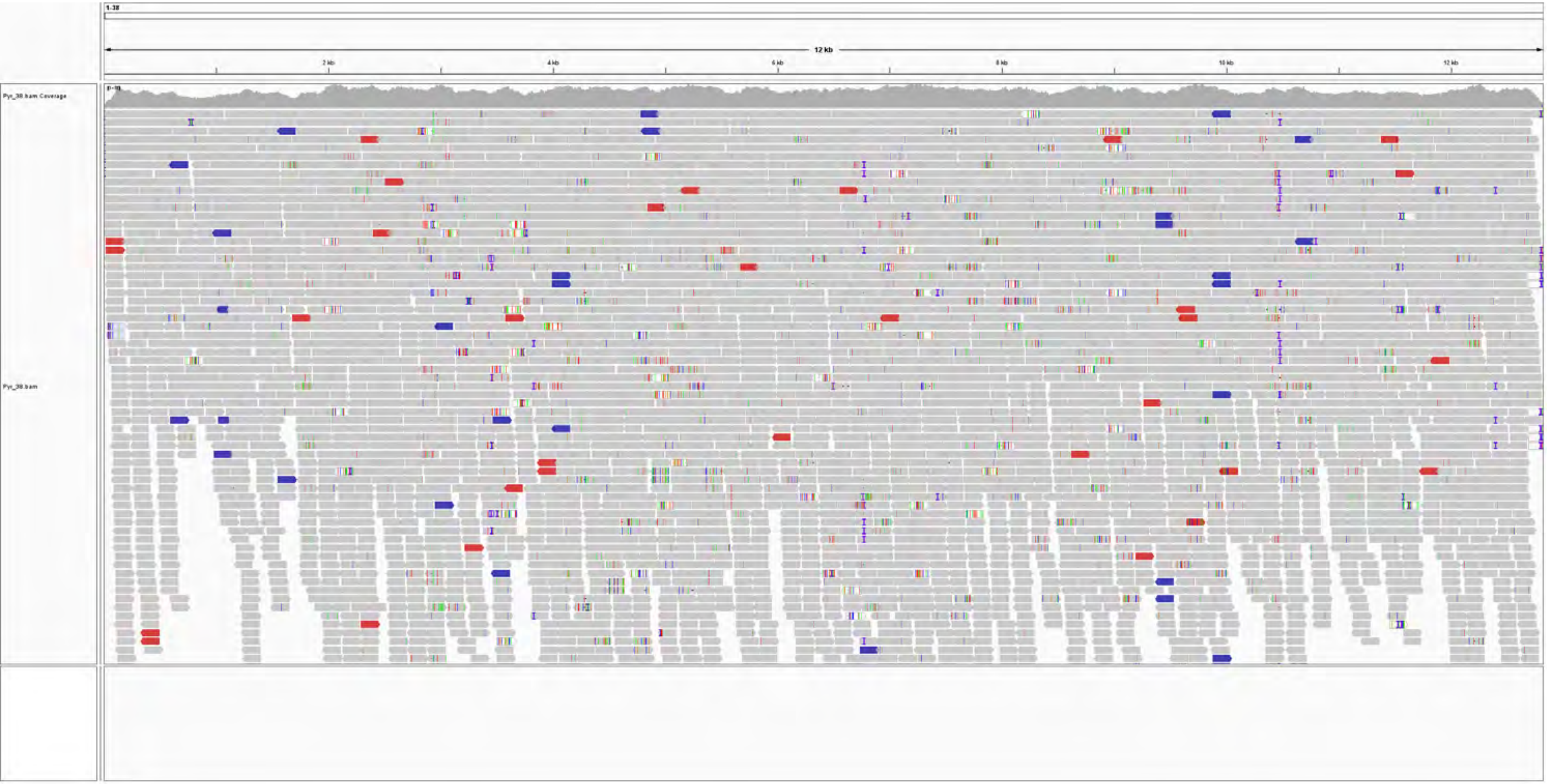

### Pyr_39

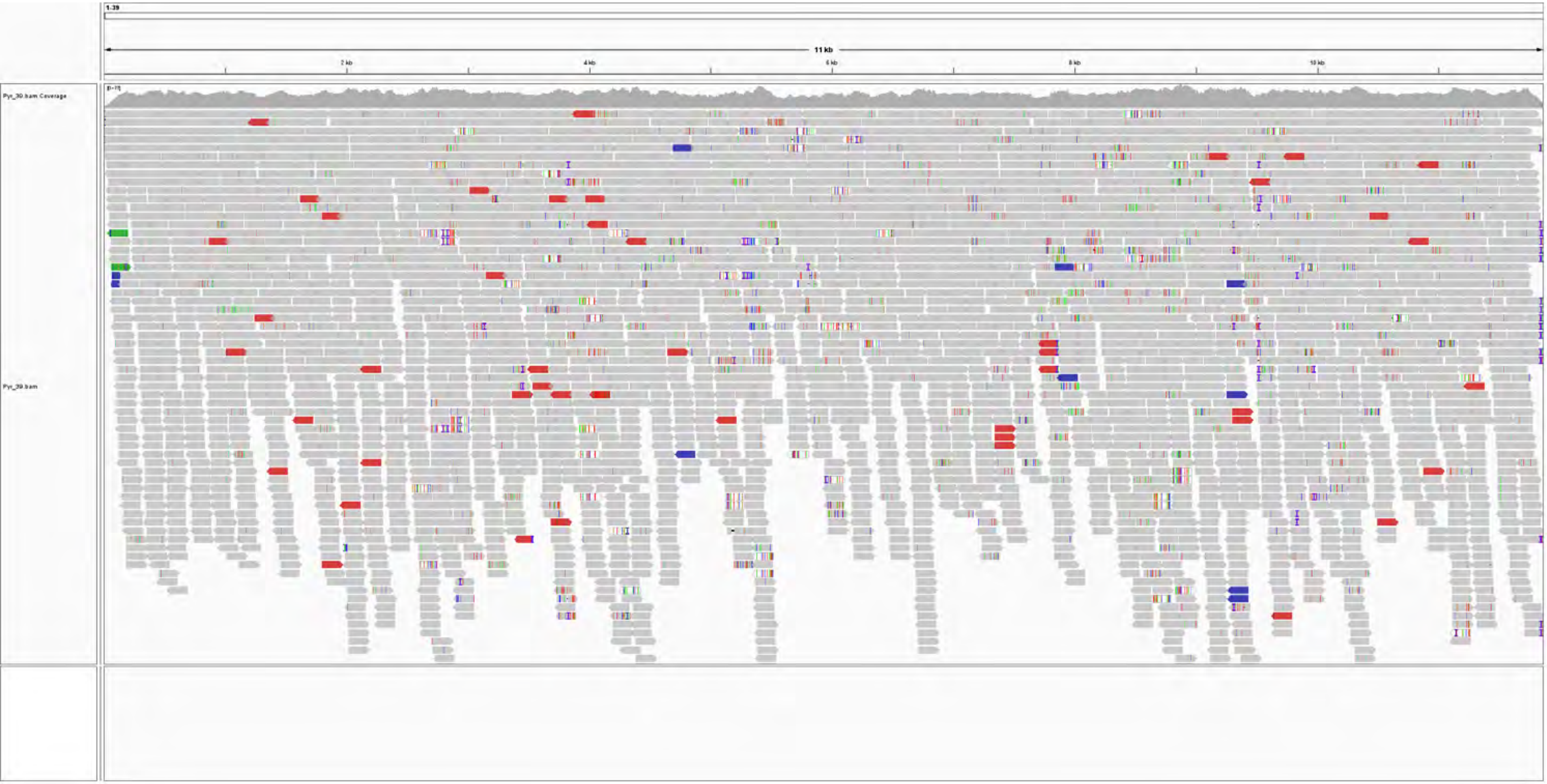

### Pyr_40

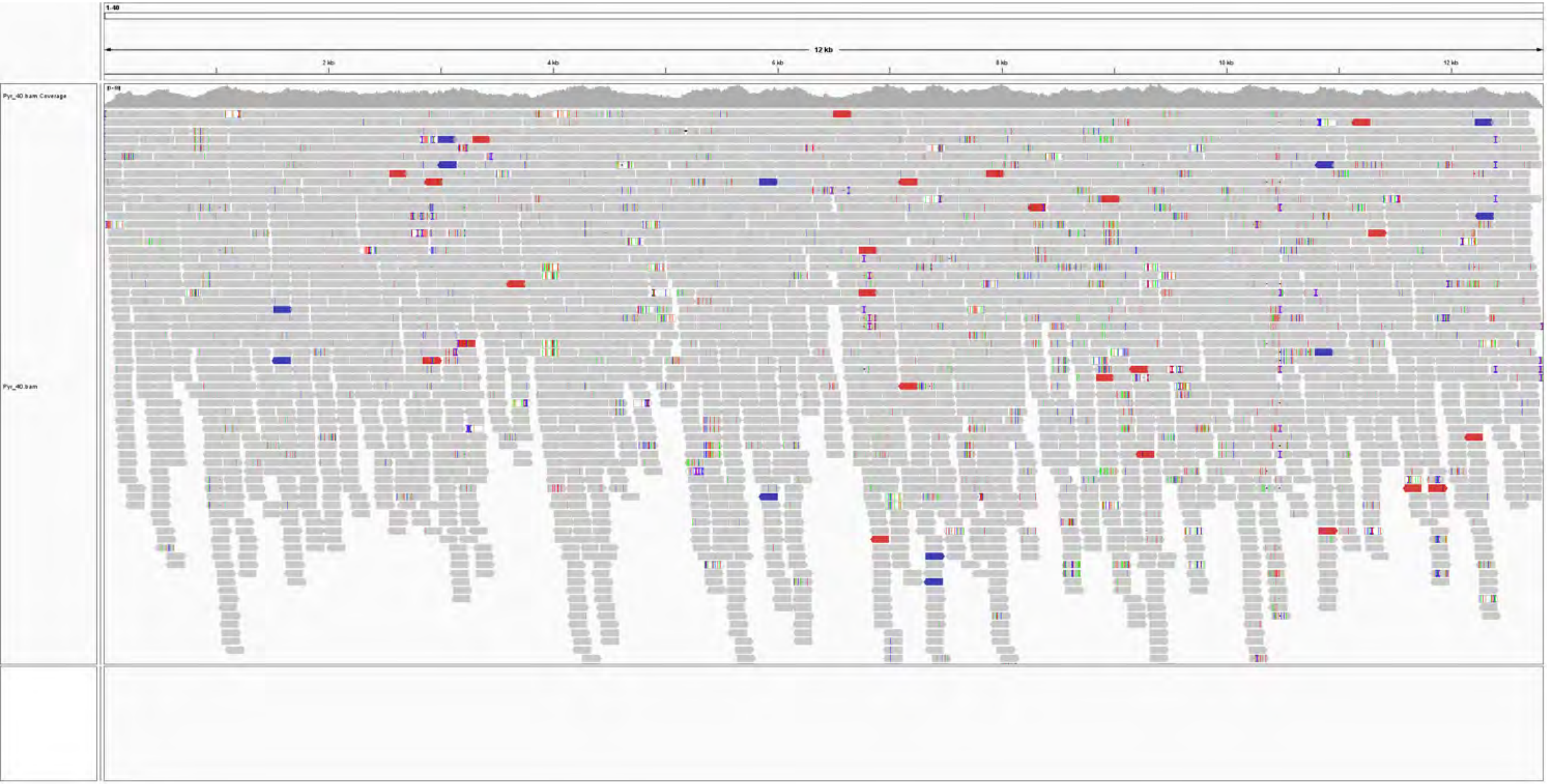

### Pyr_41

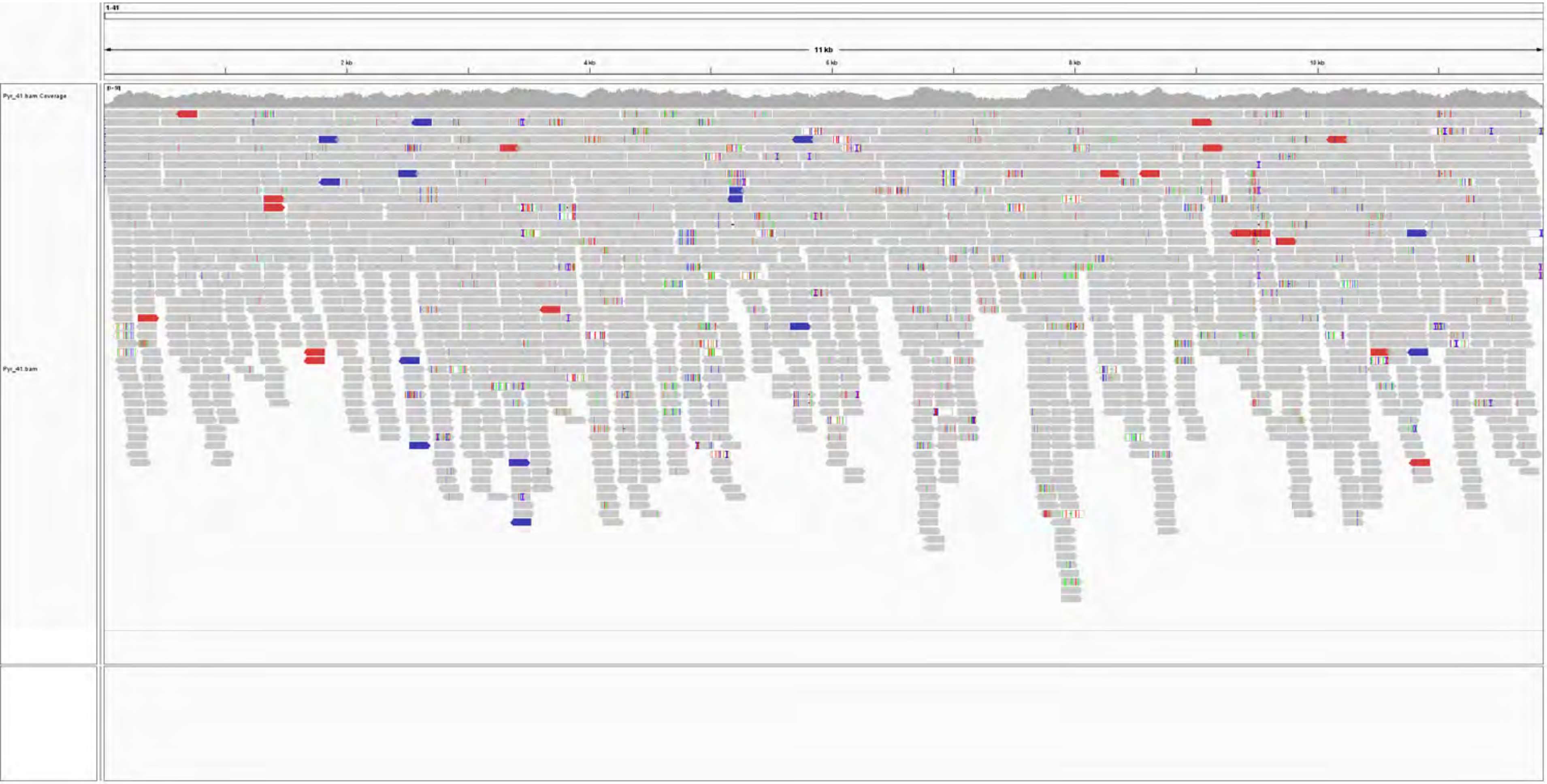

### SRR9587917

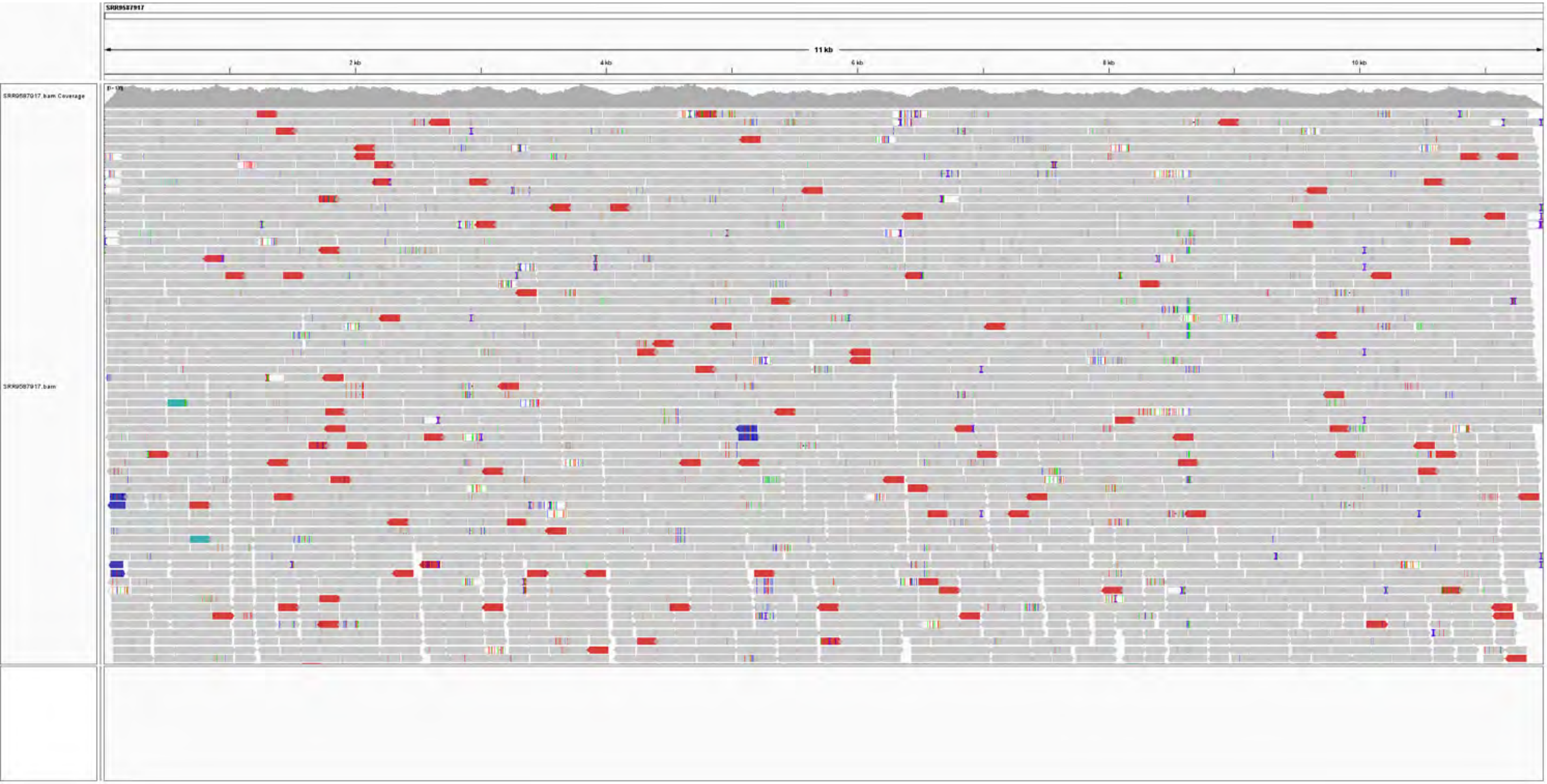

### SRR9587918

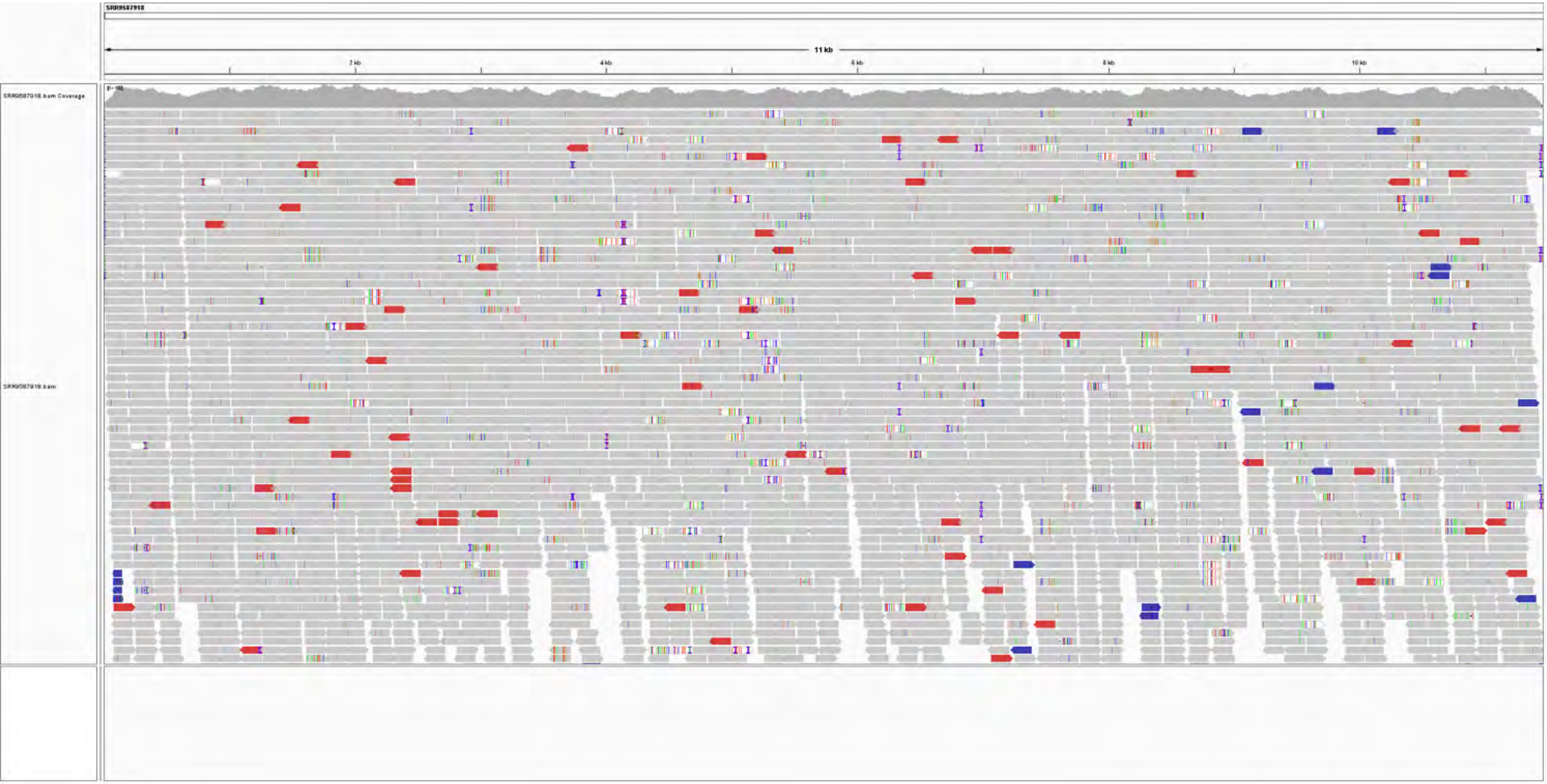
SRR9587918
11 kb
2 kb
4 kb
6 kb
8 kb
10 kb
SRR9587918.bam Coverage
SRR9587918.bam

### SRR9587919

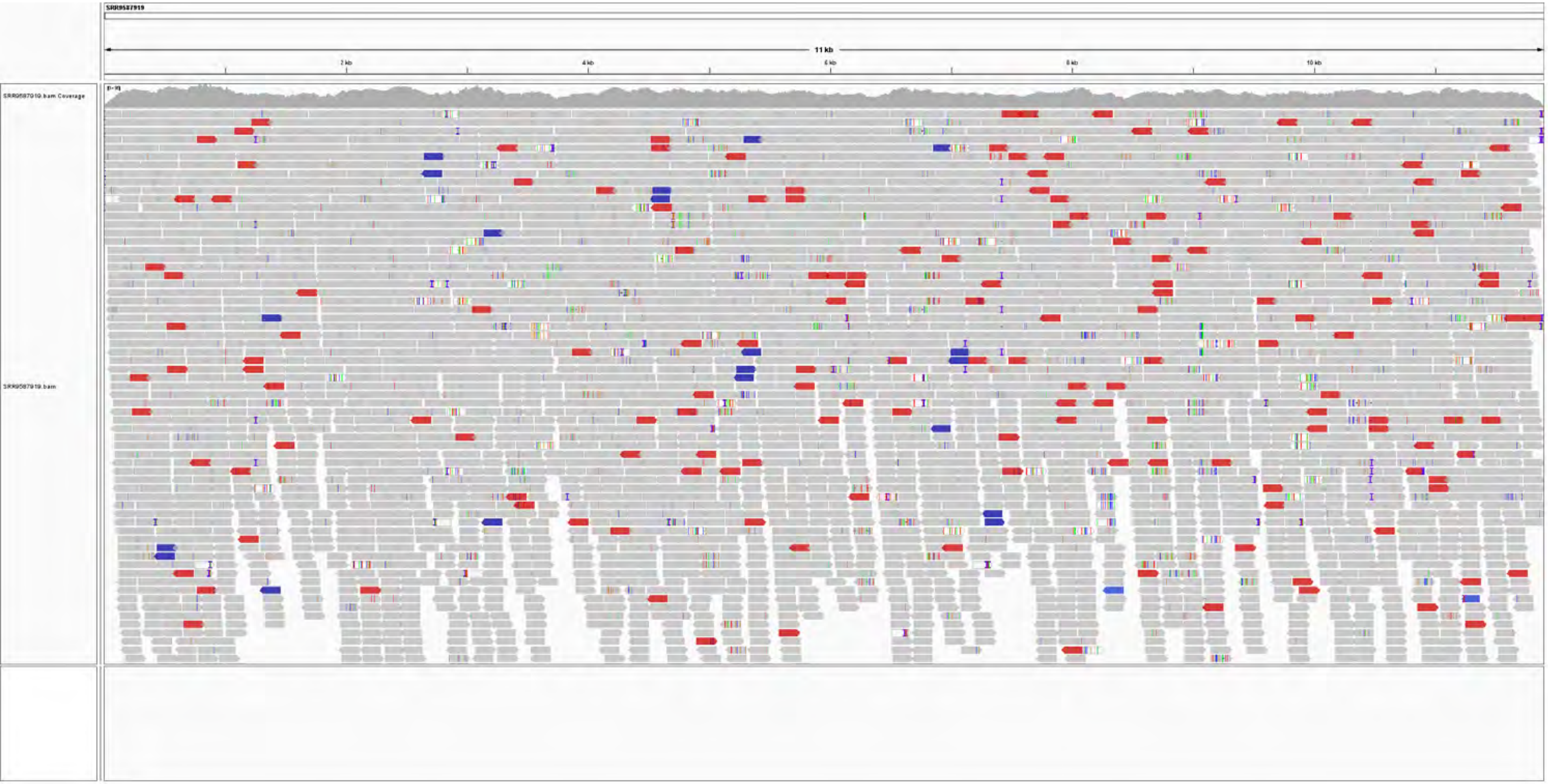

SRR9587919
11 kb
2 kb
4 kb
6 kb
8 kb
10 kb
SRR9587919.bam Coverage
SRR9587919.bam

### SRR9587921

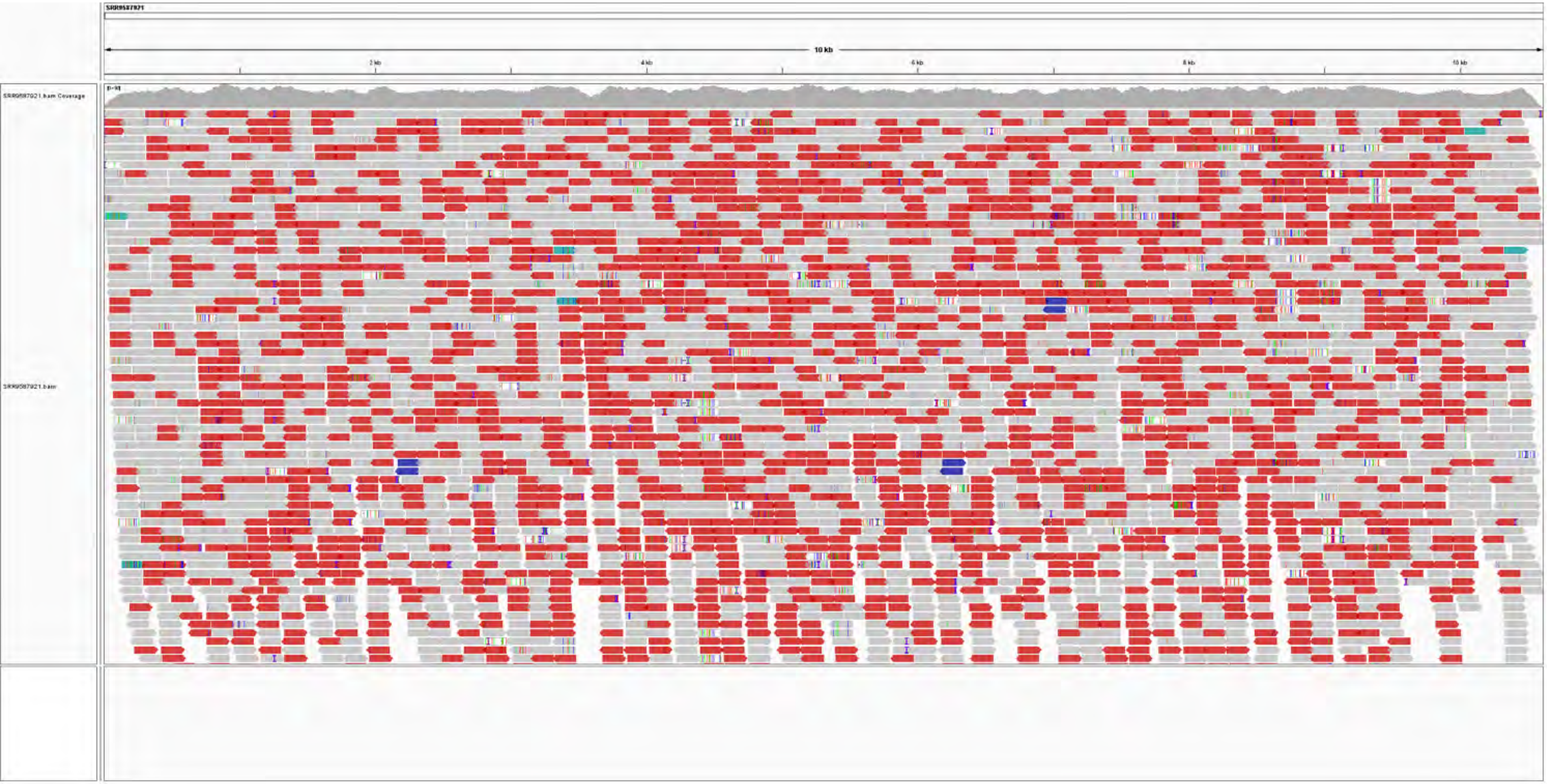

SRR9587921
10 kb
2 kb
4 kb
6 kb
8 kb
10 kb
SRR9587921.bam Coverage
SRR9587921.bam

### SRR9587922

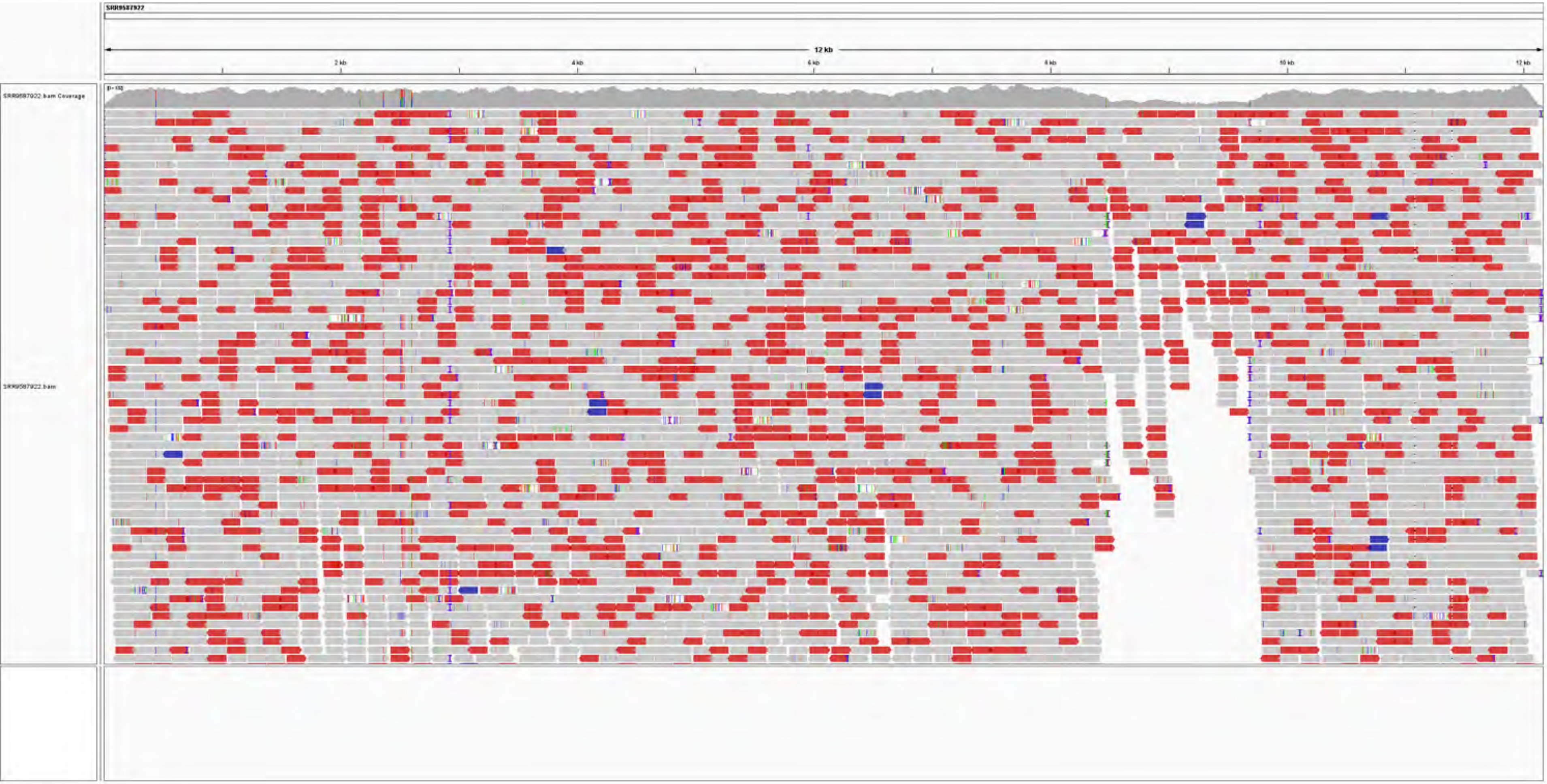

### SRR9587923

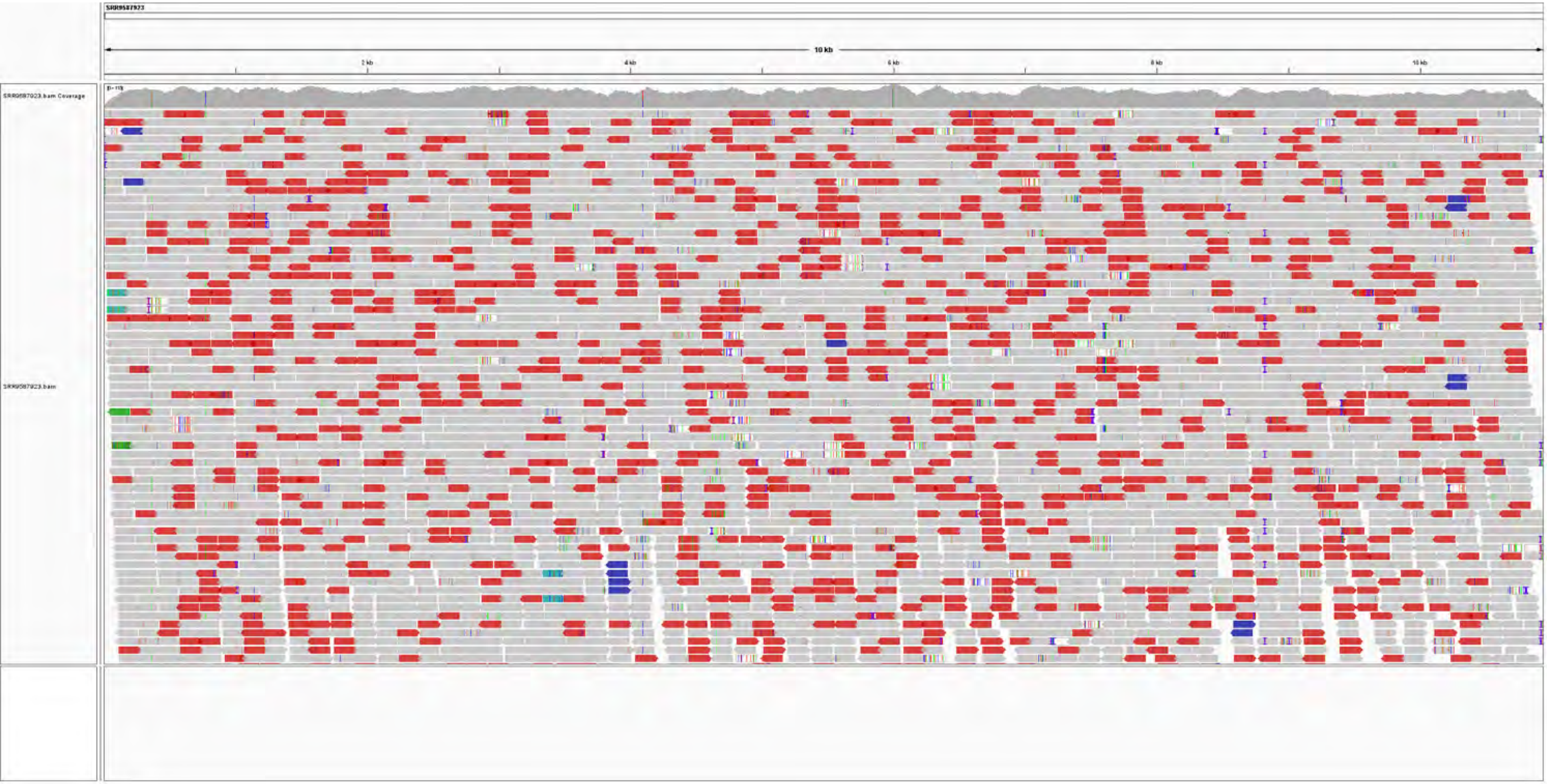
SRR9587923
10 kb
2 kb
4 kb
6 kb
8 kb
10 kb
SRR9587923.bam Coverage
SRR9587923.bam

### SRR9587924

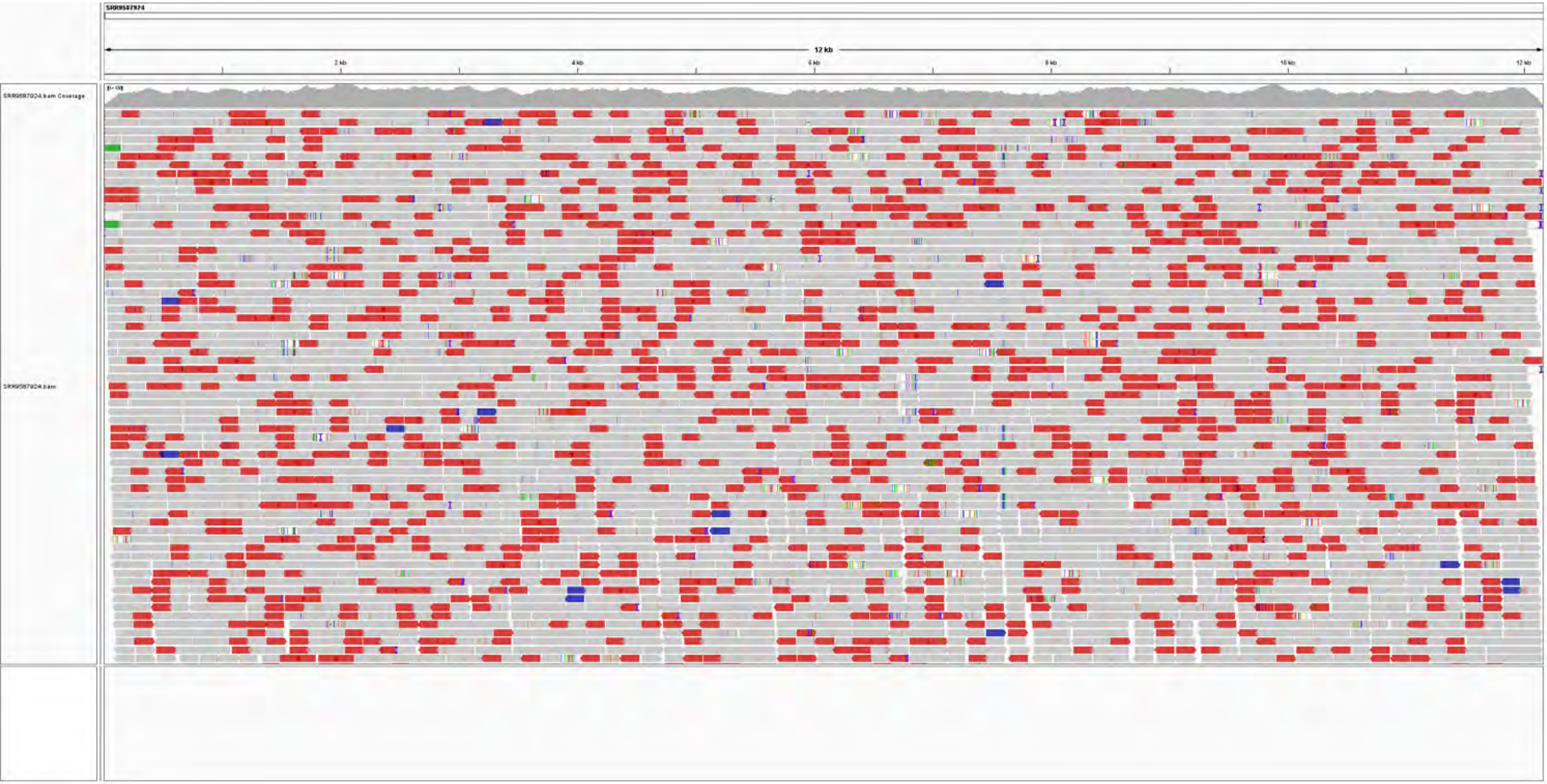
SRR9587924
12 kb
2 kb
4 kb
6 kb
8 kb
10 kb
12 kb
SRR9587924.bam Coverage
SRR9587924.bam

### SRR9587925

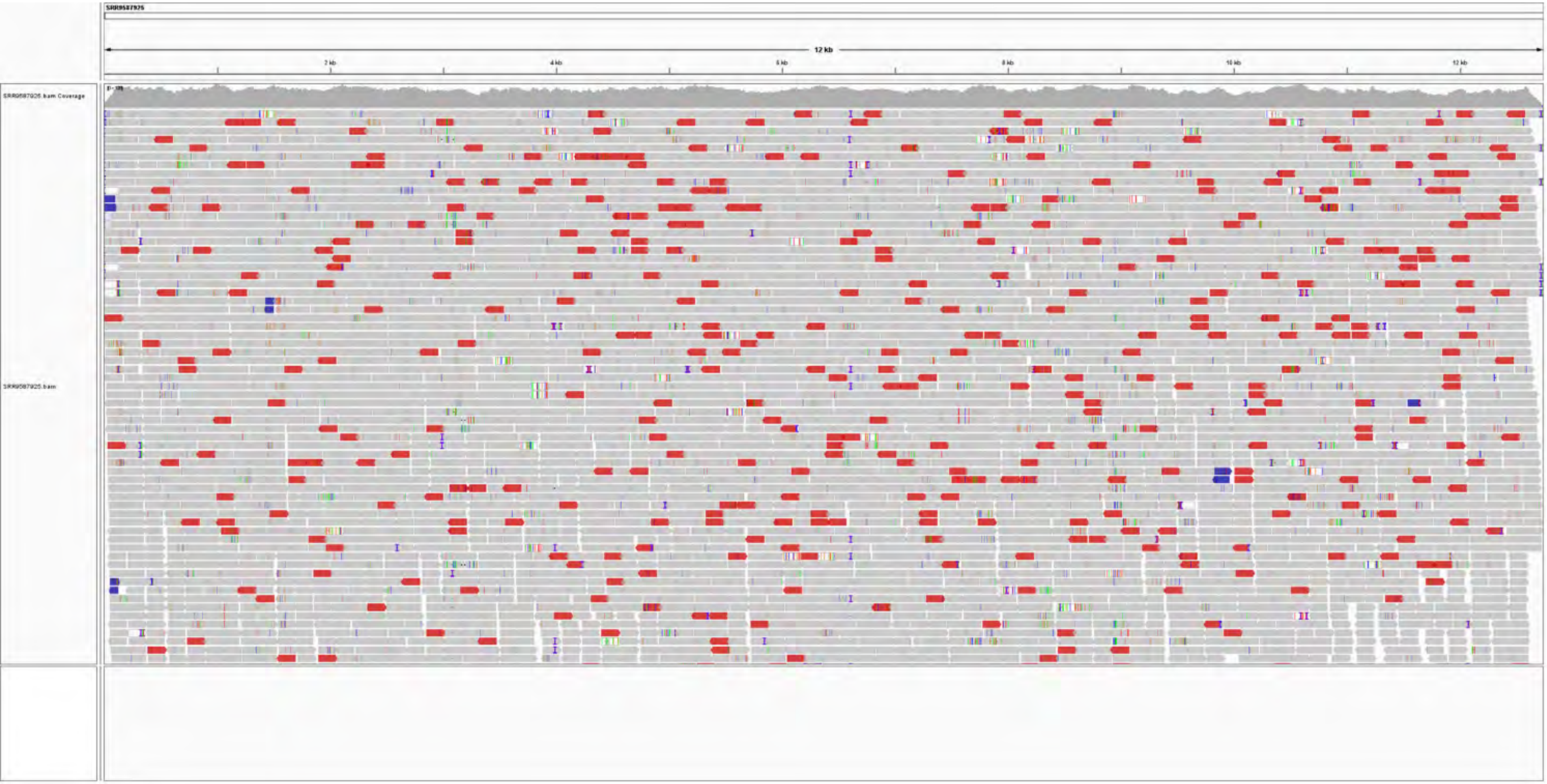
SRR9587925
12 kb
2 kb
4 kb
6 kb
8 kb
10 kb
12 kb
SRR9587925.bam Coverage
SRR9587925.bam

### SRR9587926

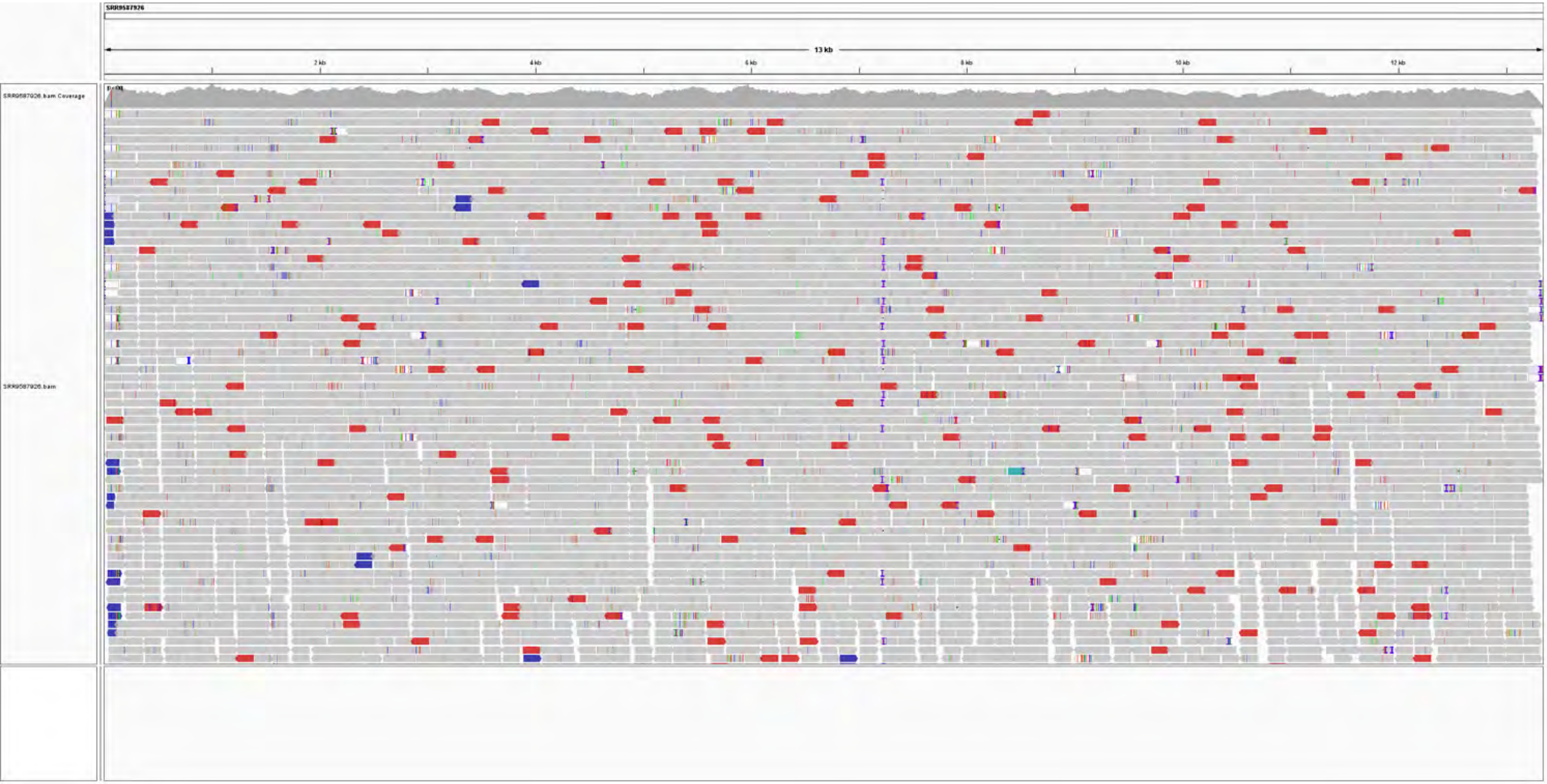

SRR9587926
13 kb
2 kb
4 kb
6 kb
8 kb
10 kb
12 kb
SRR9587926.bam Coverage
SRR9587926.bam

### SRR9587927

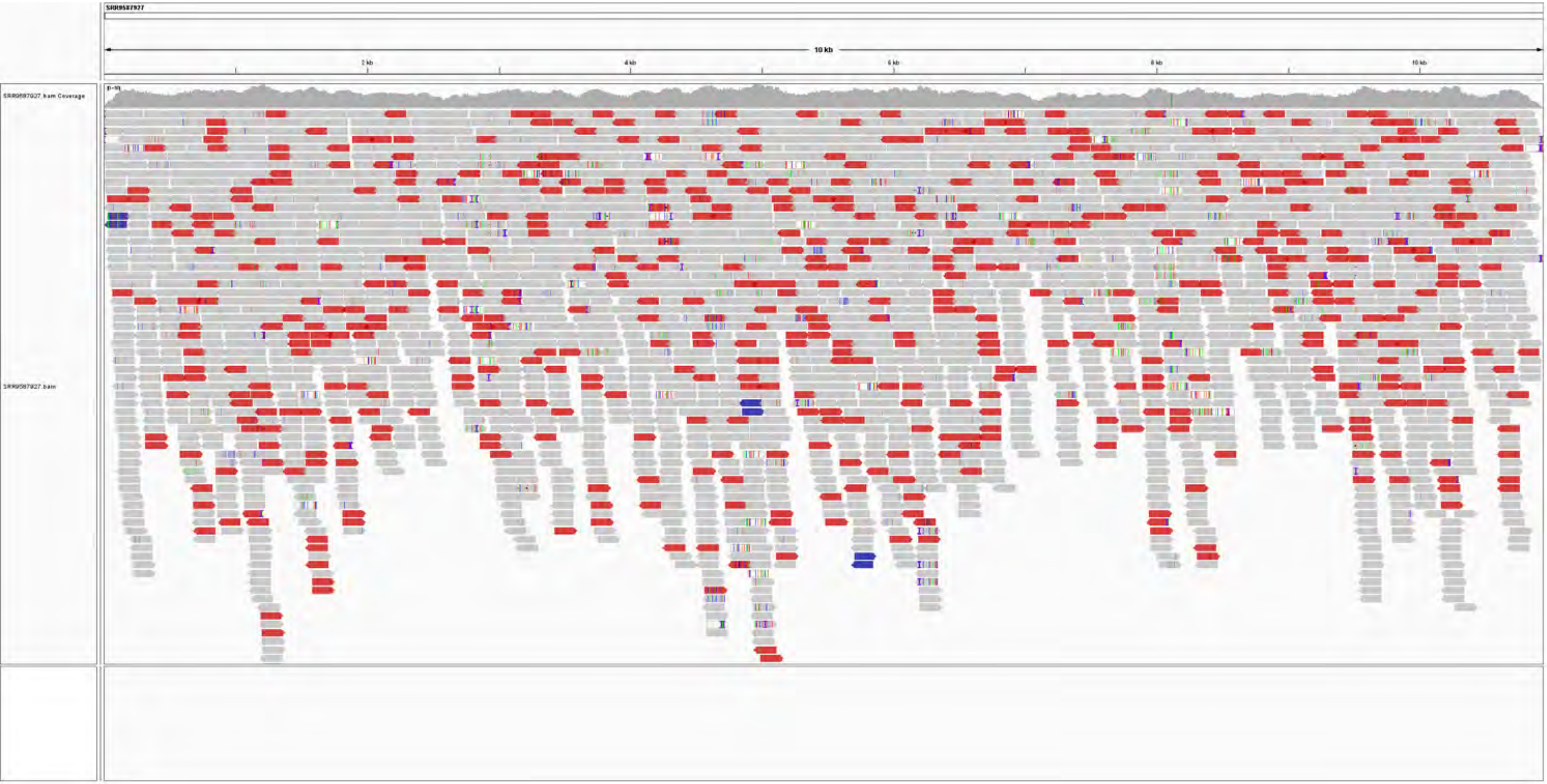

SRR9587927
10 kb
2 kb
4 kb
6 kb
8 kb
10 kb
SRR9587927.bam Coverage
SRR9587927.bam

### SRR9587929

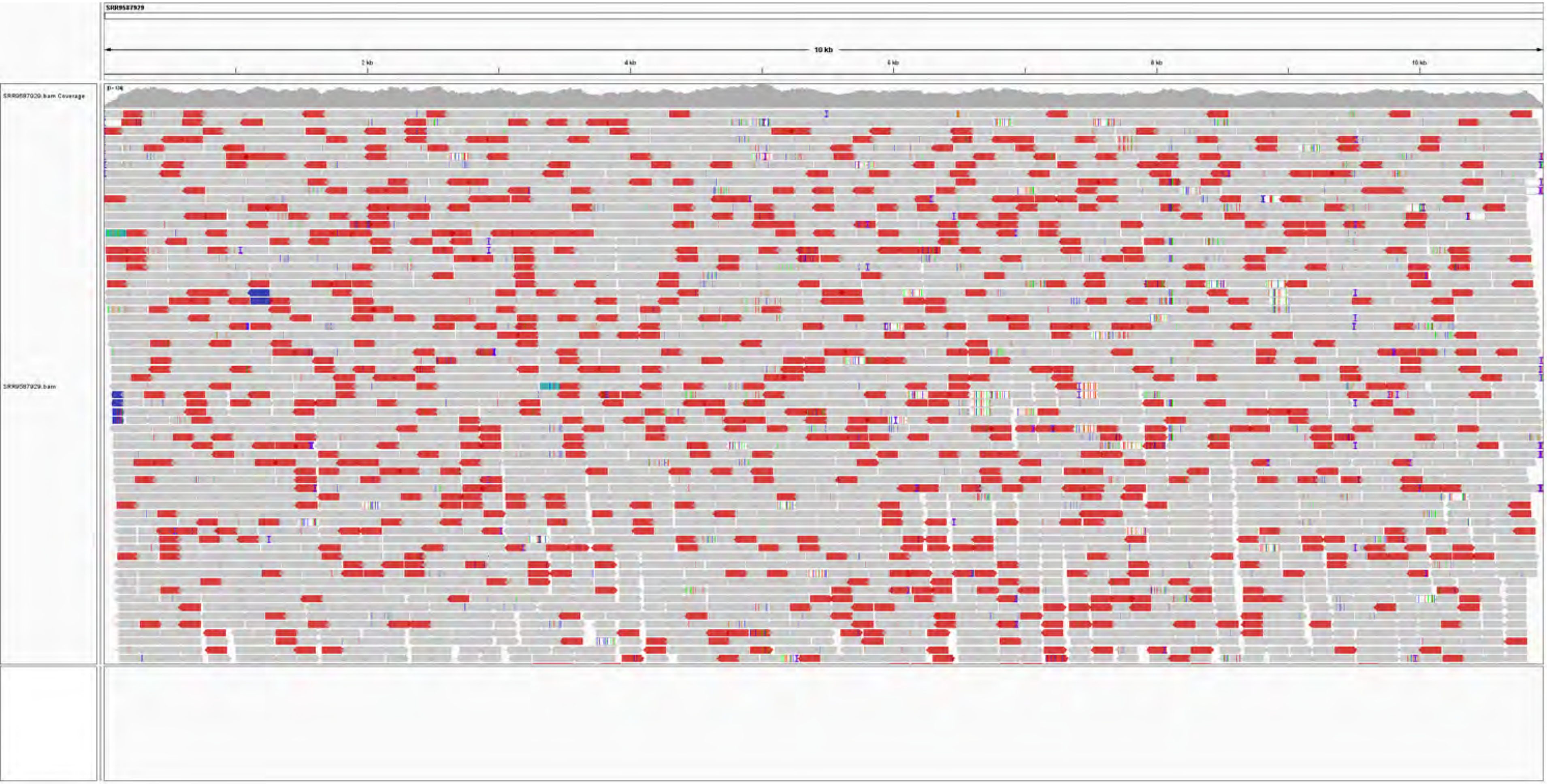
SRR9587929
10 kb
2 kb
4 kb
6 kb
8 kb
10 kb
SRR9587929.bam Coverage
SRR9587929.bam

### SRR9587930

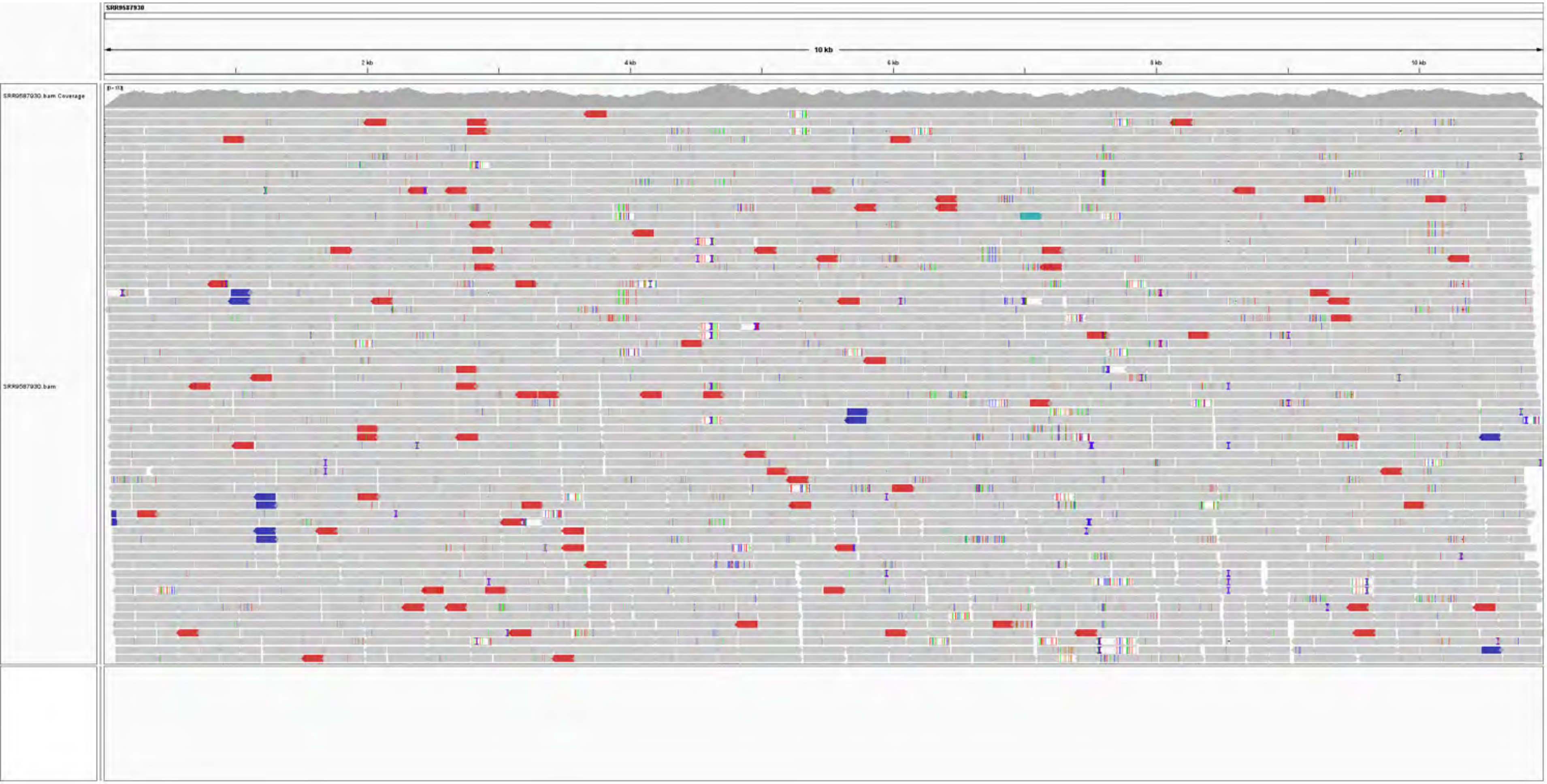

SRR9587930
10 kb
2 kb
4 kb
6 kb
8 kb
10 kb
SRR9587930.bam Coverage
SRR9587930.bam

### SRR9587931

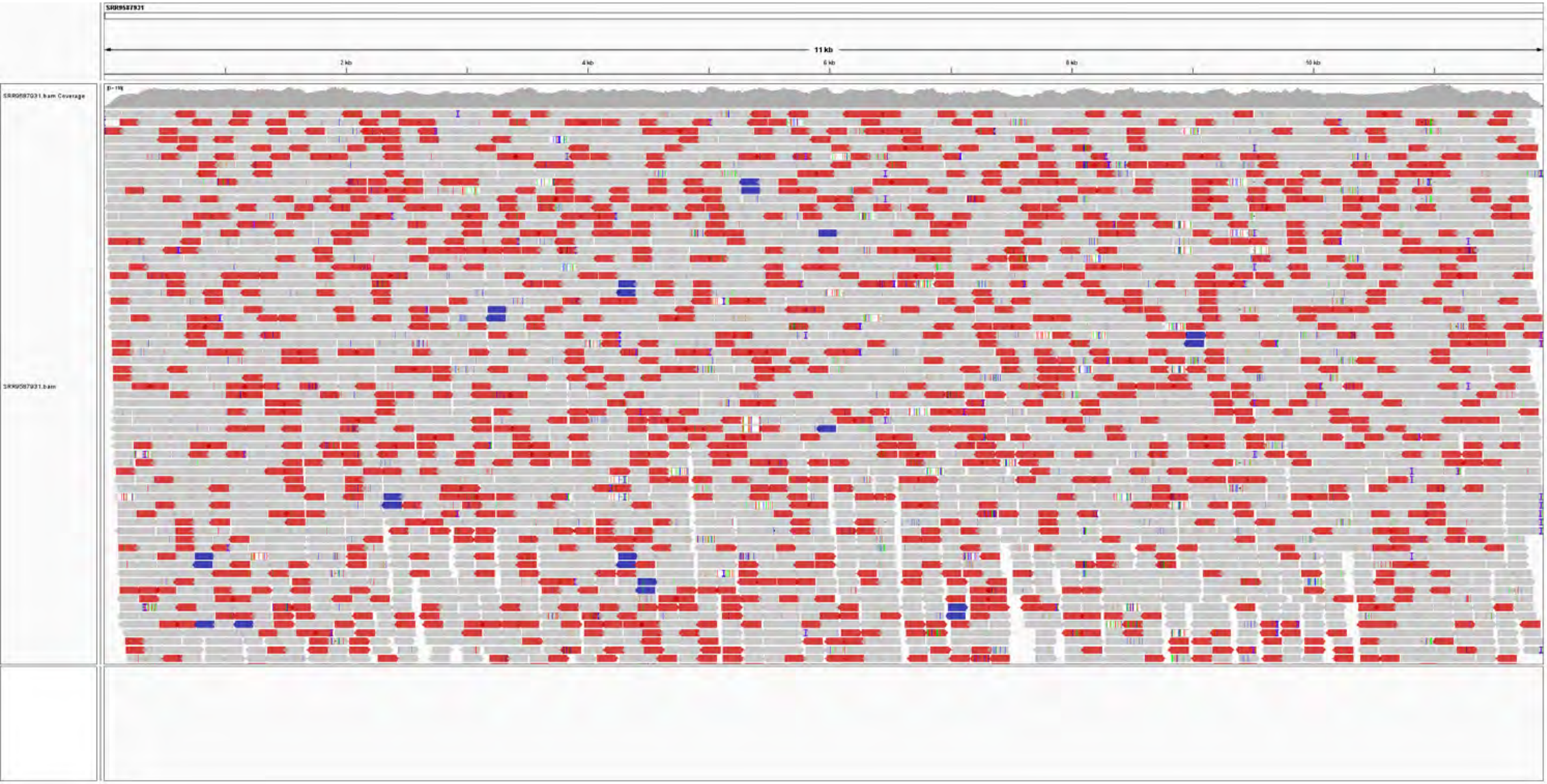

### SRR9587932

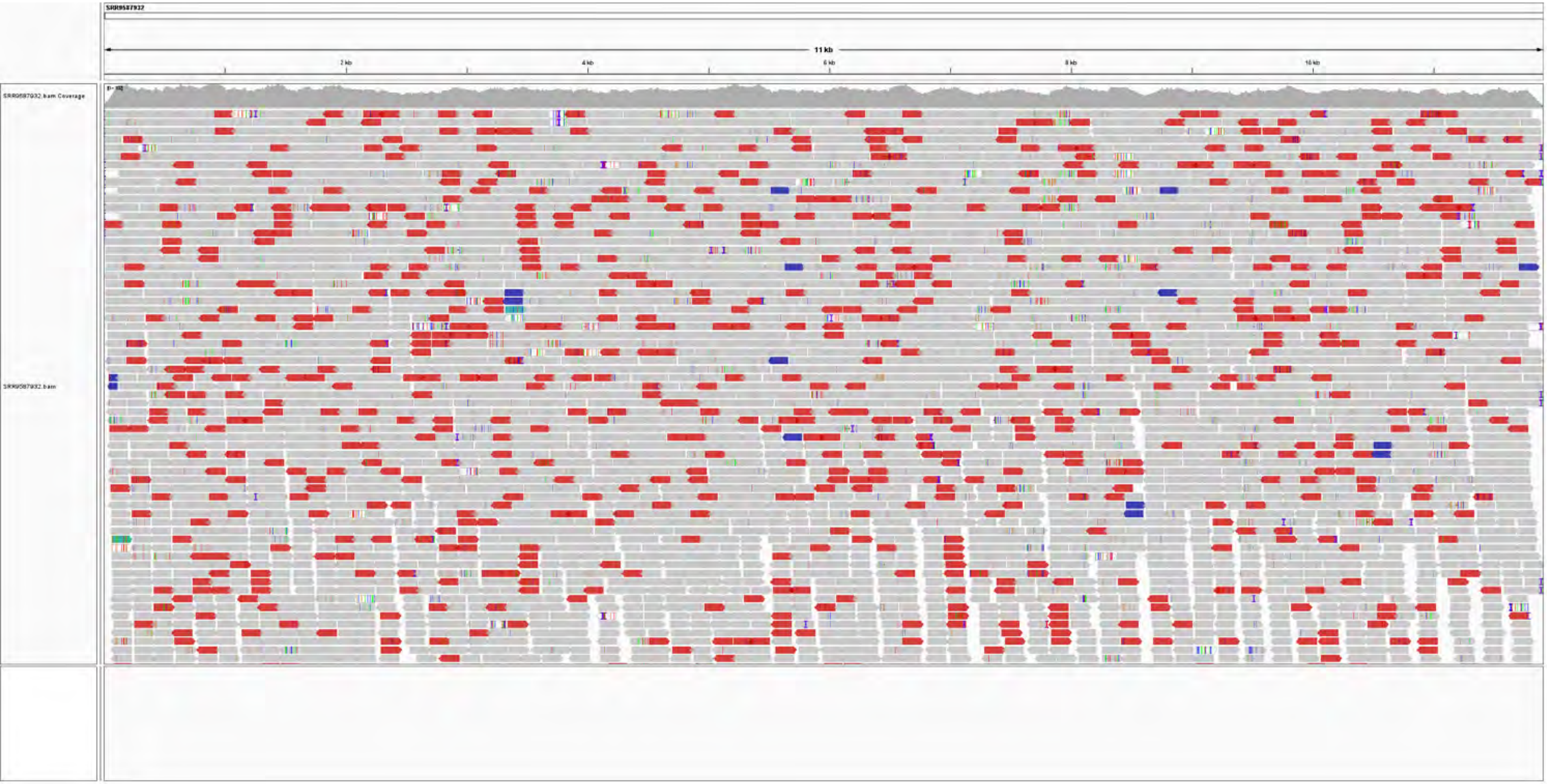

SRR9587932
11 kb
2 kb
4 kb
6 kb
8 kb
10 kb
SRR9587932.bam Coverage
SRR9587932.bam

### SRR9587933

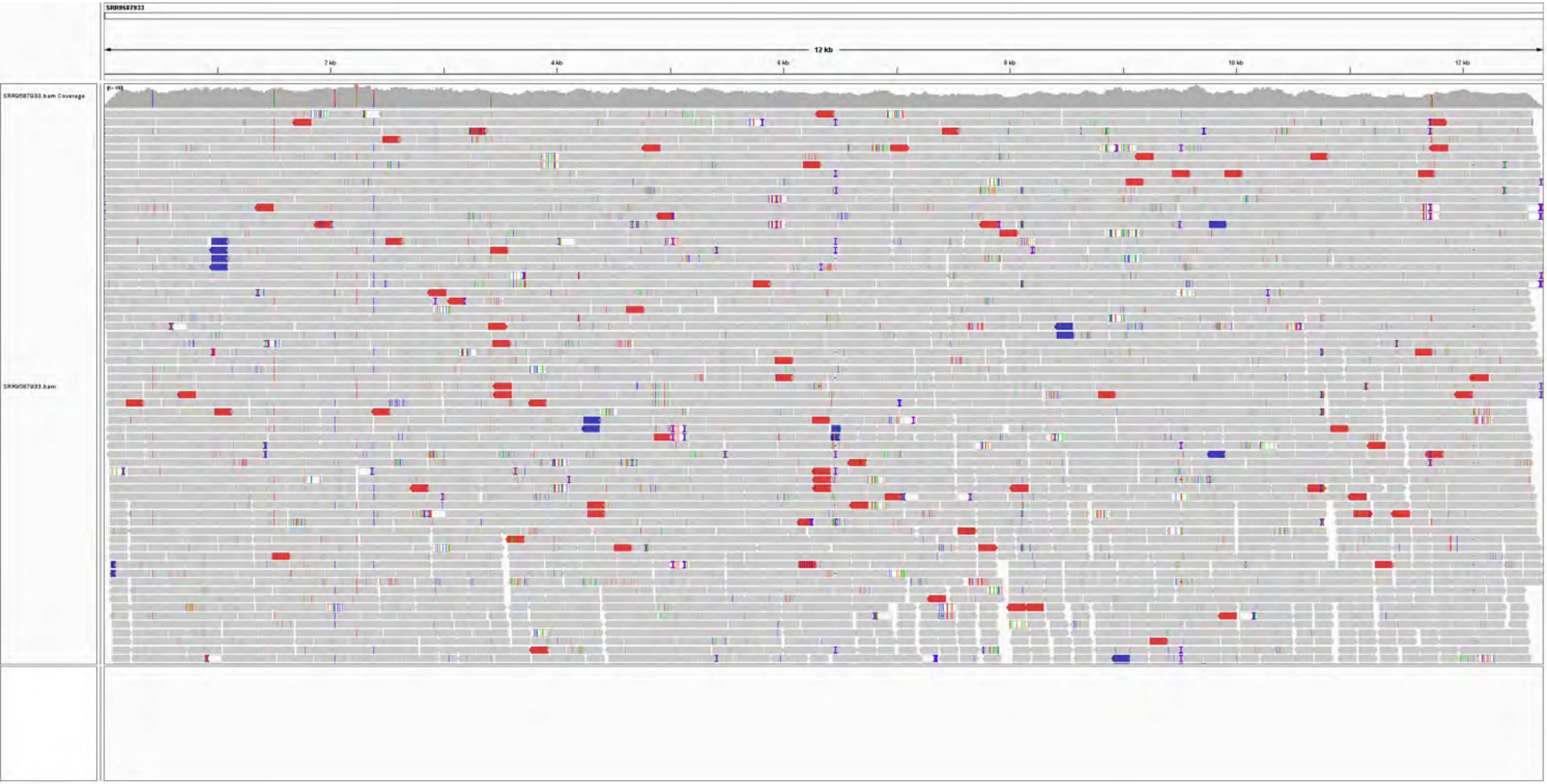
SRR9587933
12 kb
2 kb
4 kb
6 kb
8 kb
10 kb
12 kb
SRR9587933.bam Coverage
SRR9587933.bam

### SRR9587934

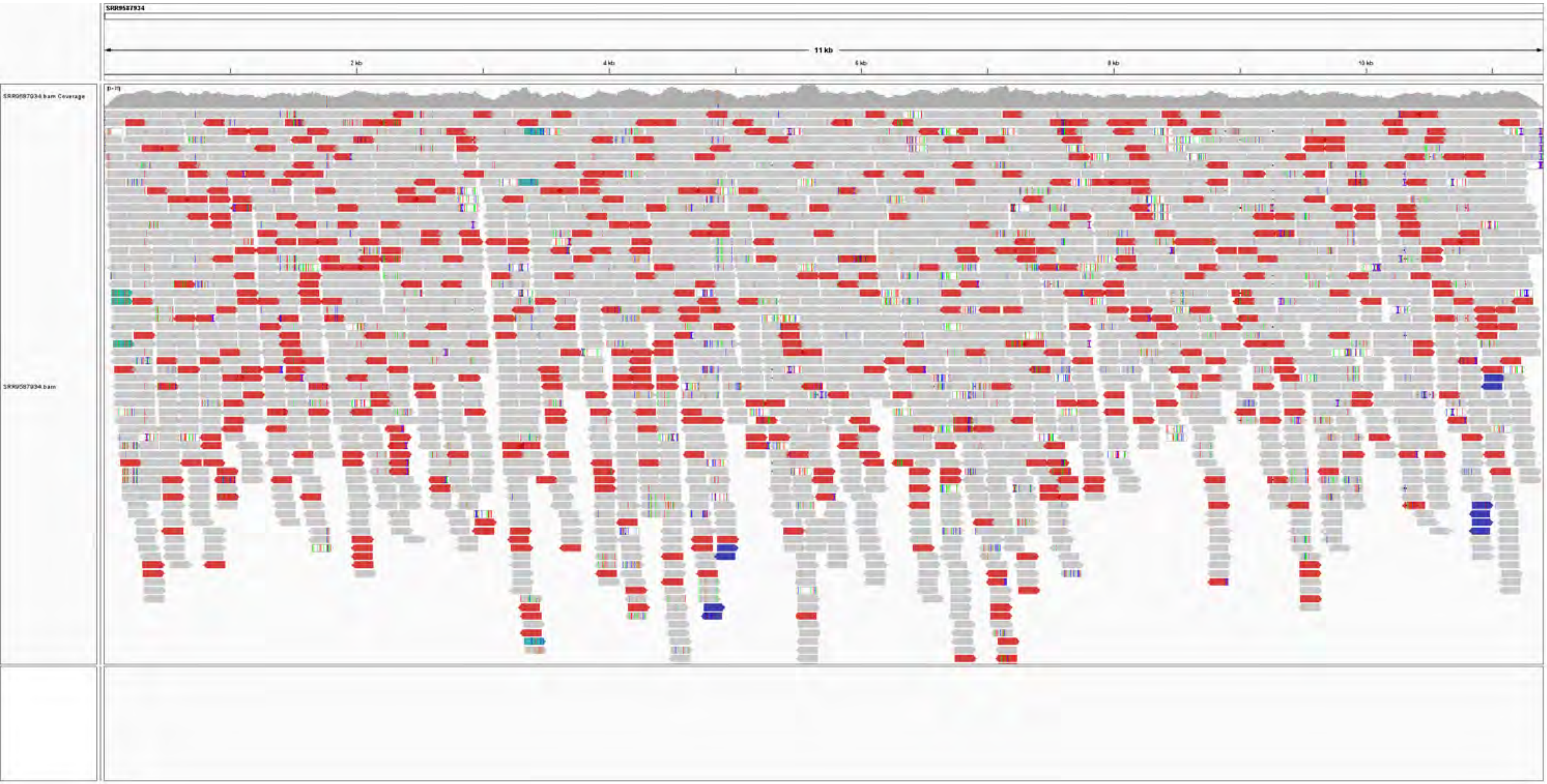

### SRR9587935

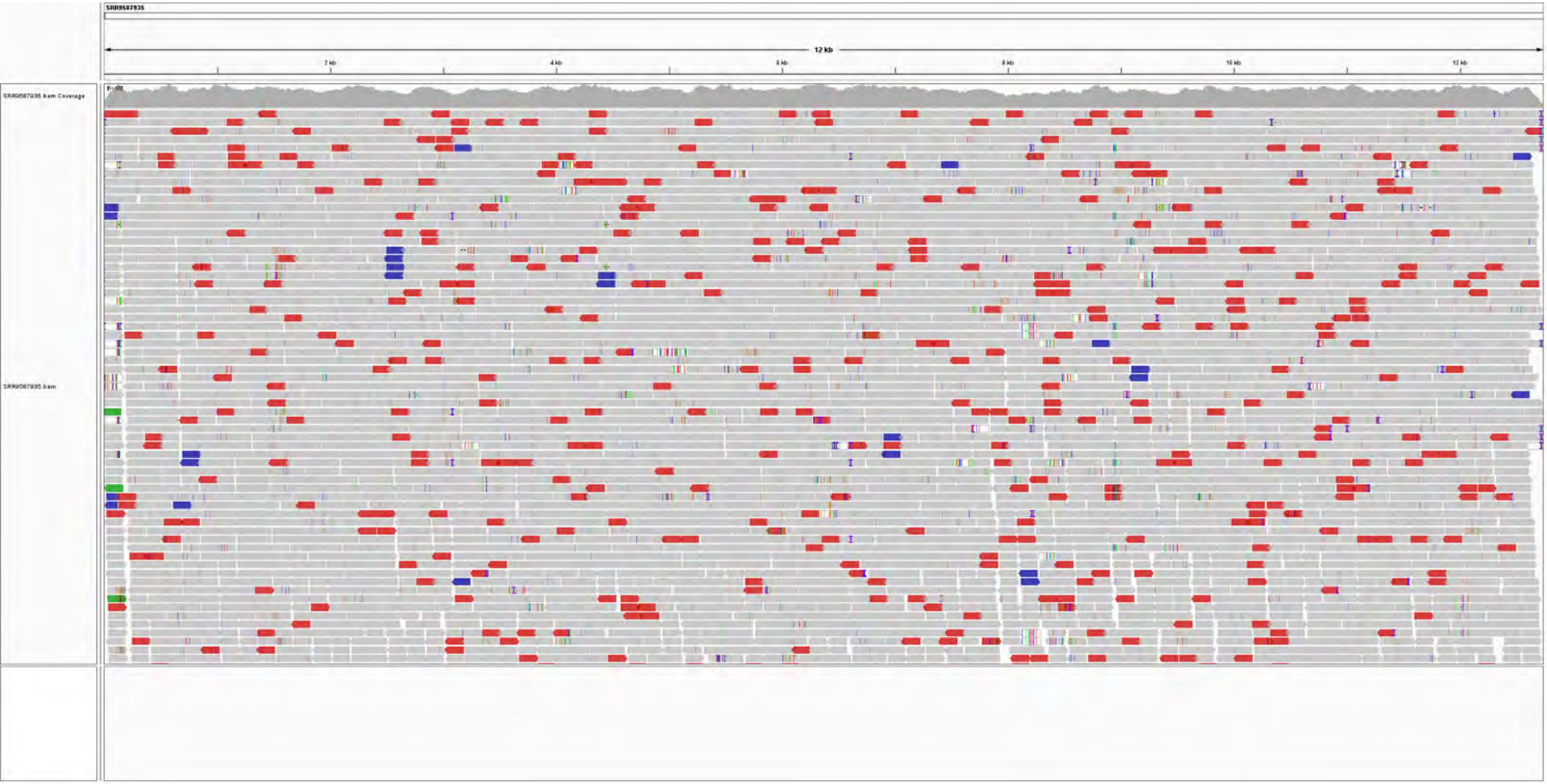

SRR9587935
12 kb
2 kb
4 kb
6 kb
8 kb
10 kb
12 kb
SRR9587935.bam Coverage
SRR9587935.bam

### SRR9587936

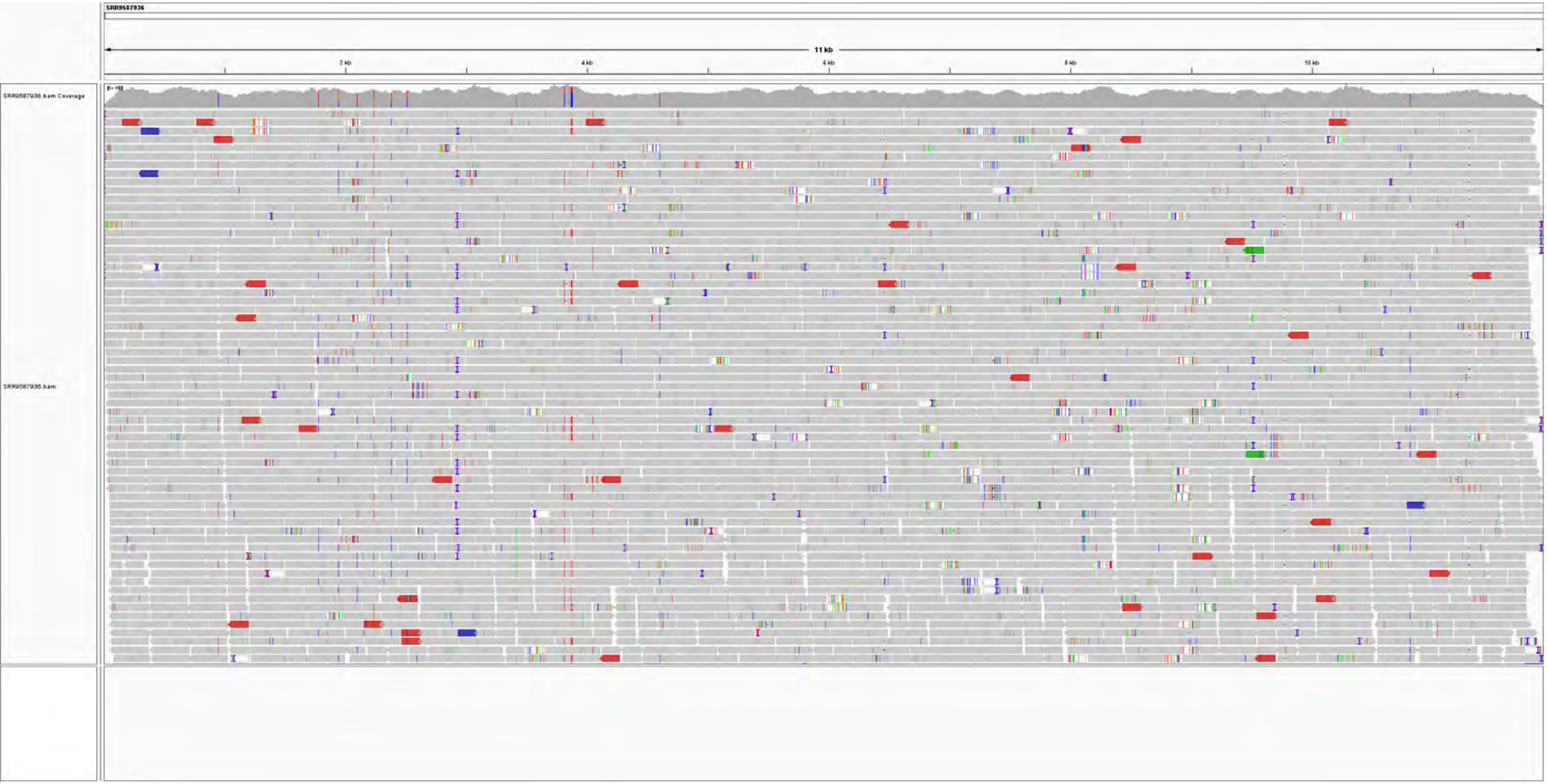

### SRR9587937

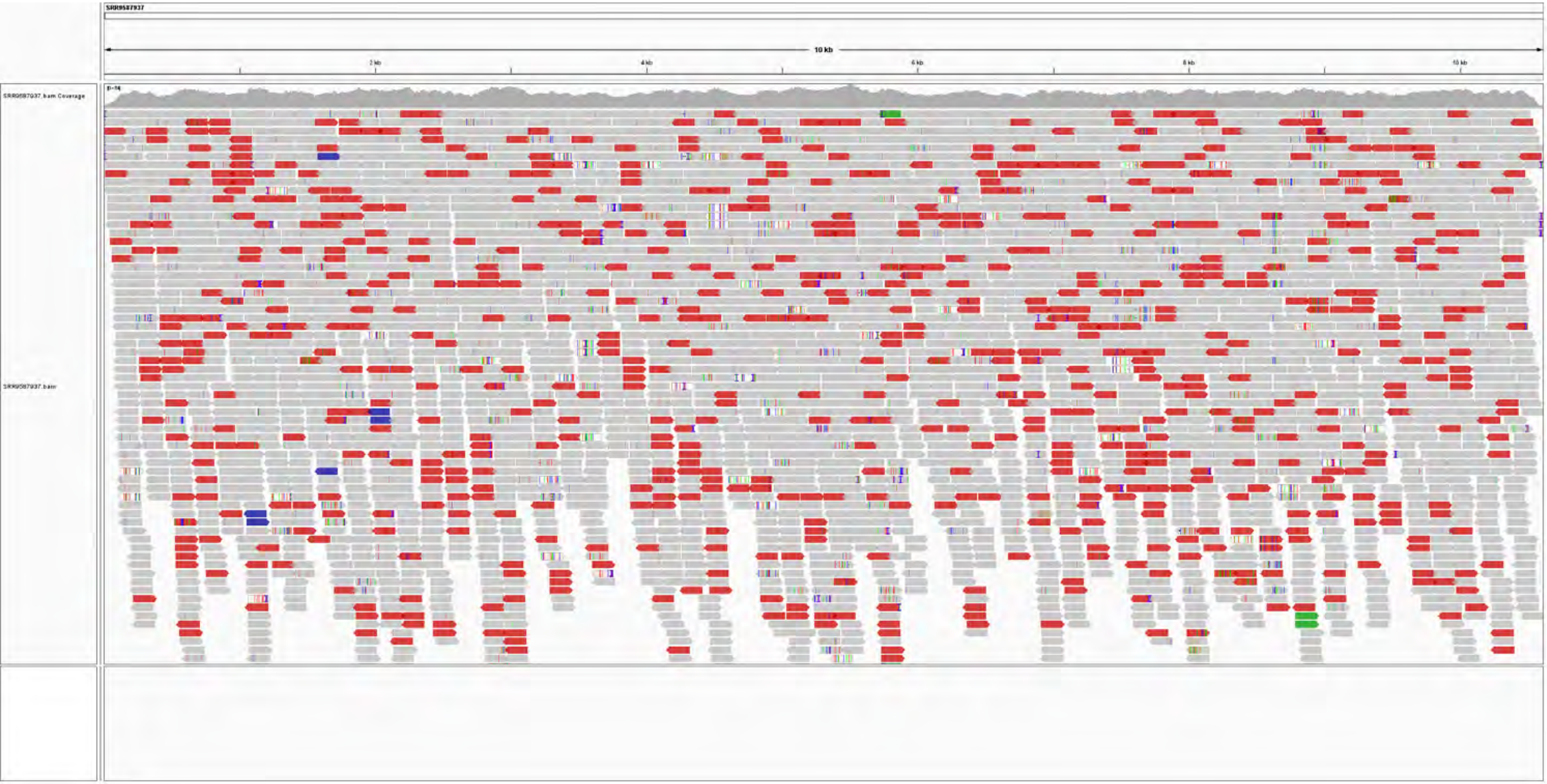
SRR9587937
10 kb
2 kb
4 kb
6 kb
8 kb
10 kb
SRR9587937.bam Coverage
SRR9587937.bam

### SRR9587938

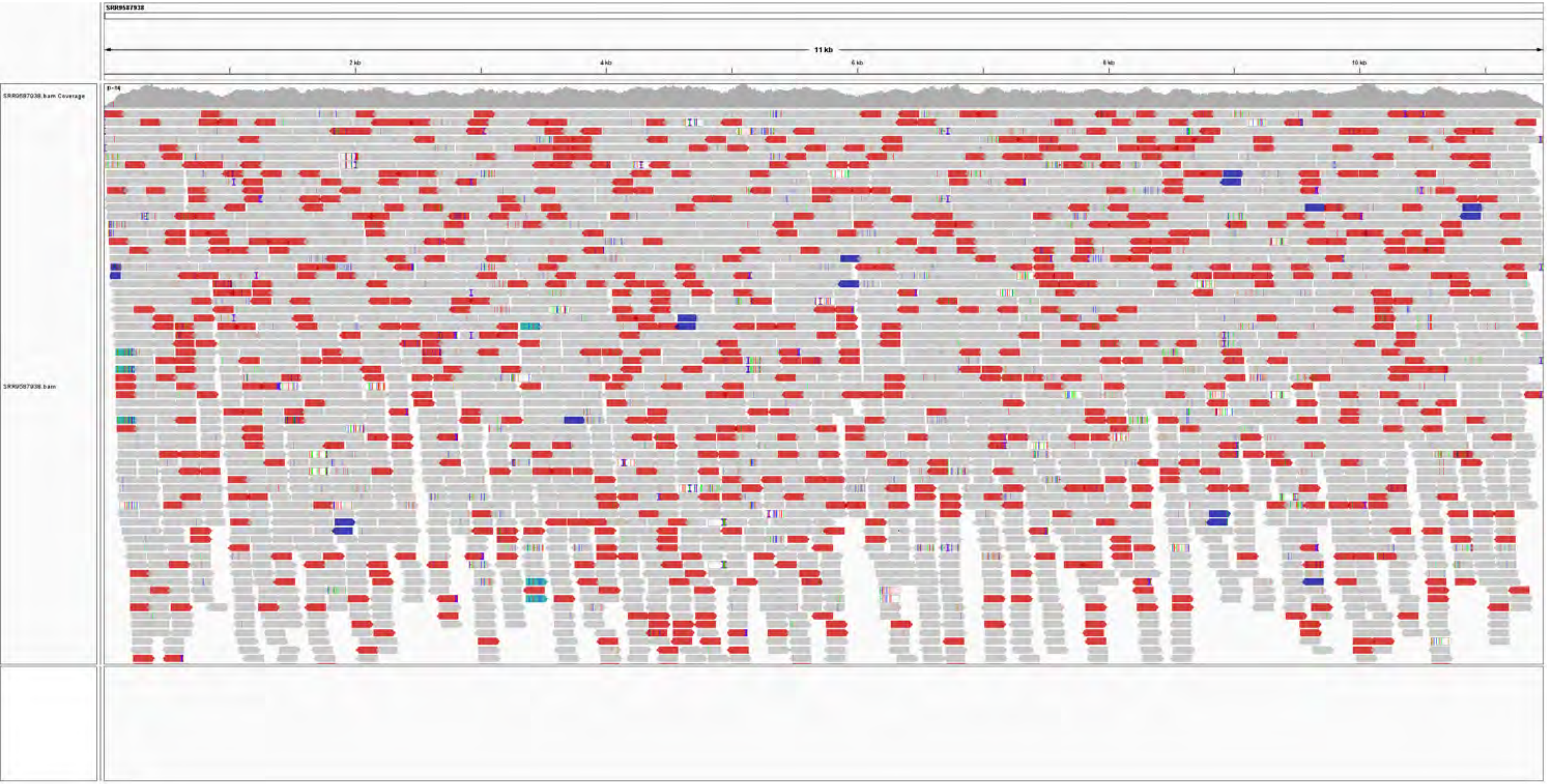
SRR9587938
11 kb
2 kb
4 kb
6 kb
8 kb
10 kb
[0-14]
SRR9587938.bam Coverage
SRR9587938.bam

### SRR9587939

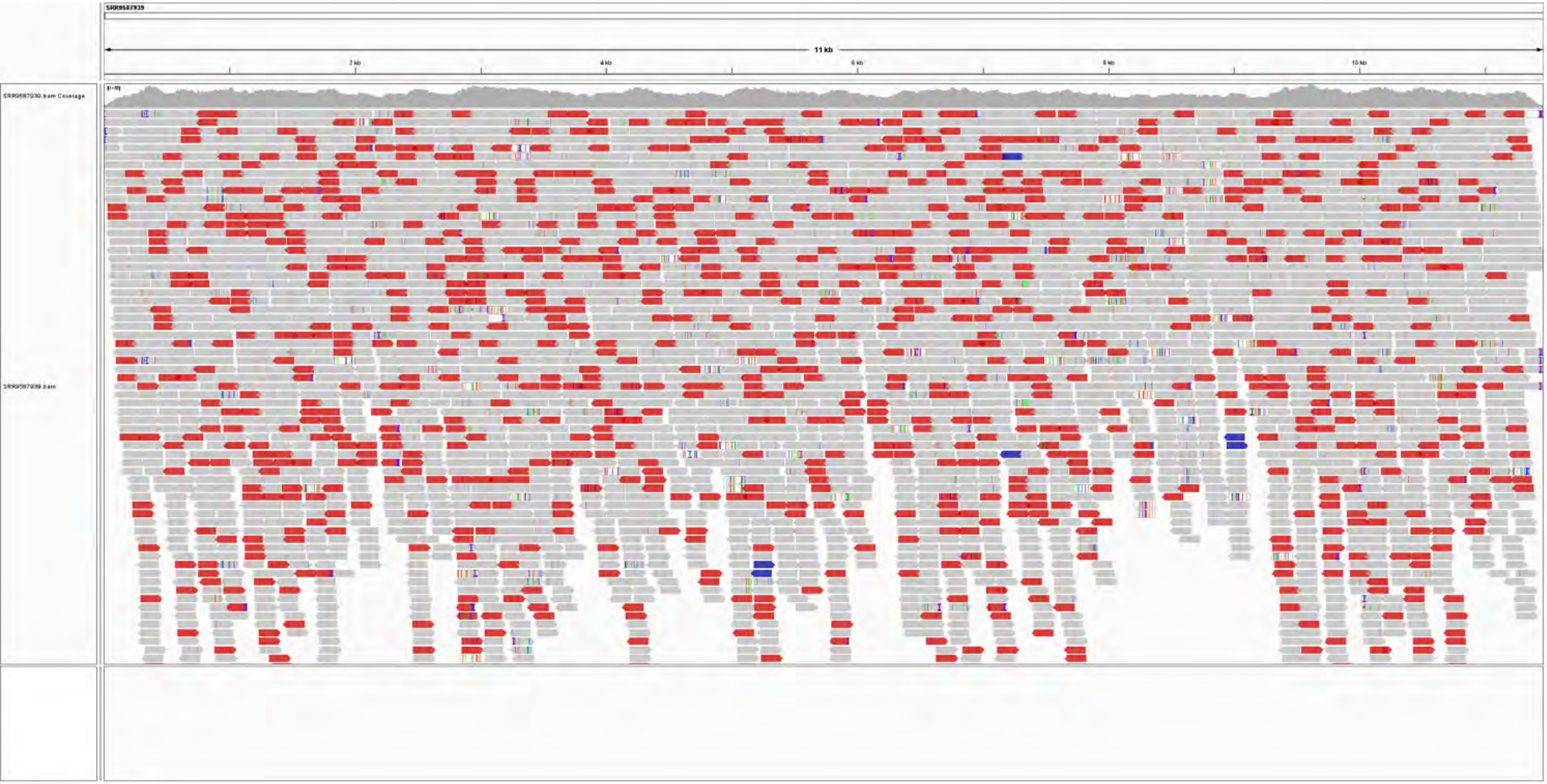

### SRR9587940

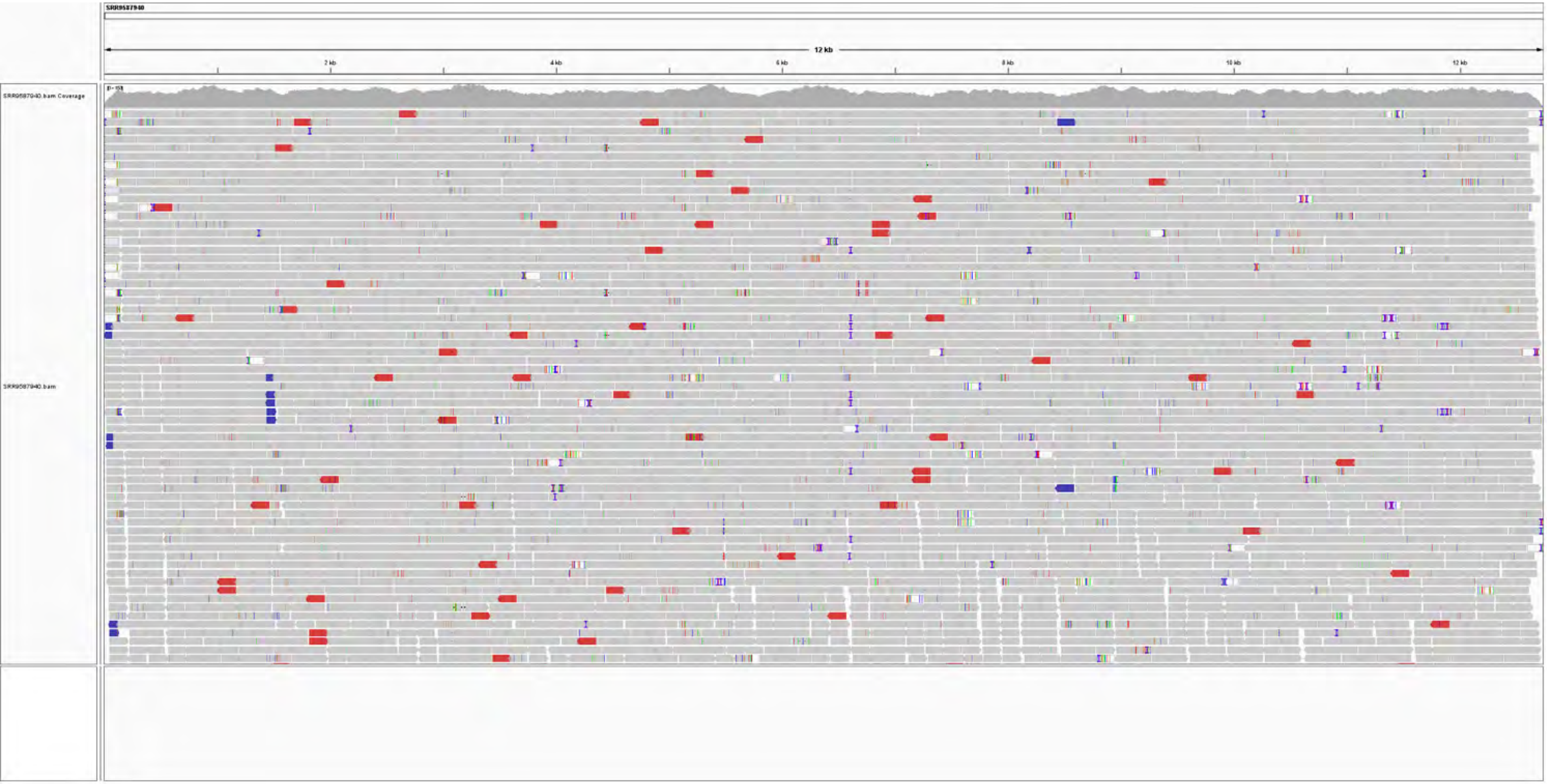
SRR9587940
12 kb
2 kb
4 kb
6 kb
8 kb
10 kb
12 kb
SRR9587940.bam Coverage
SRR9587940.bam

### SRR9587941

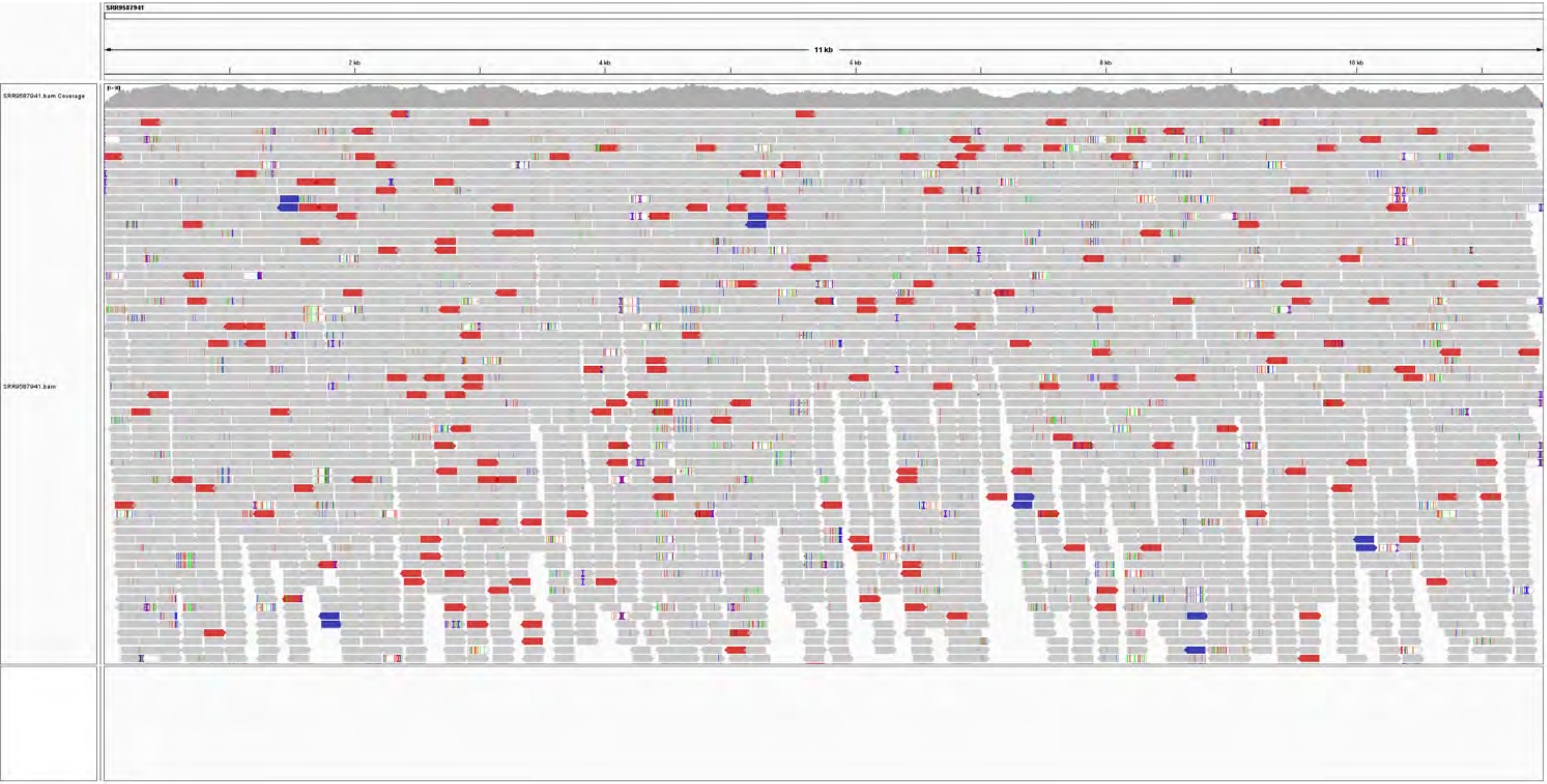
SRR9587941
11 kb
2 kb
4 kb
6 kb
8 kb
10 kb
SRR9587941.bam Coverage
SRR9587941.bam

### SRR9587942

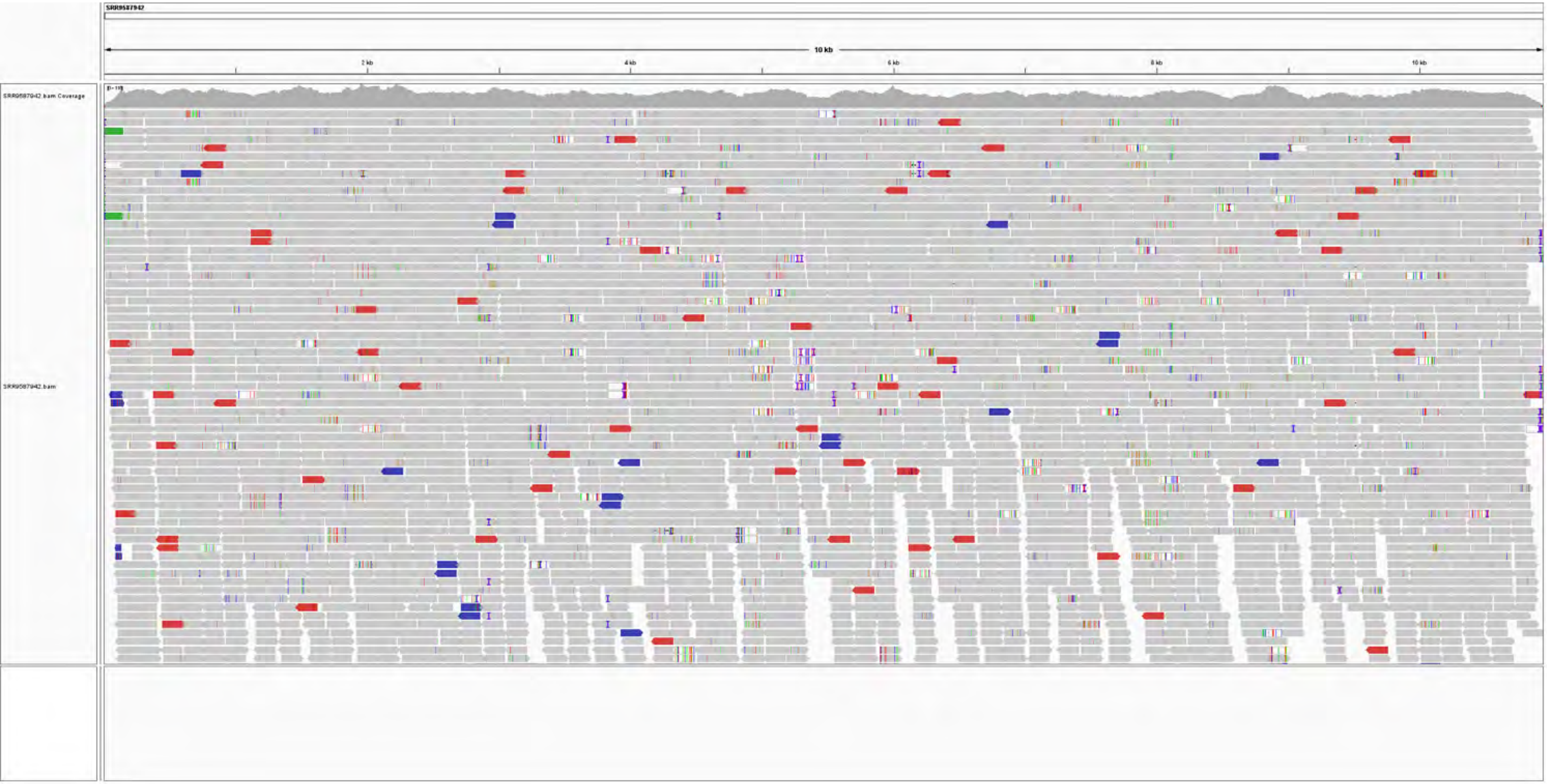

### SRR9587943

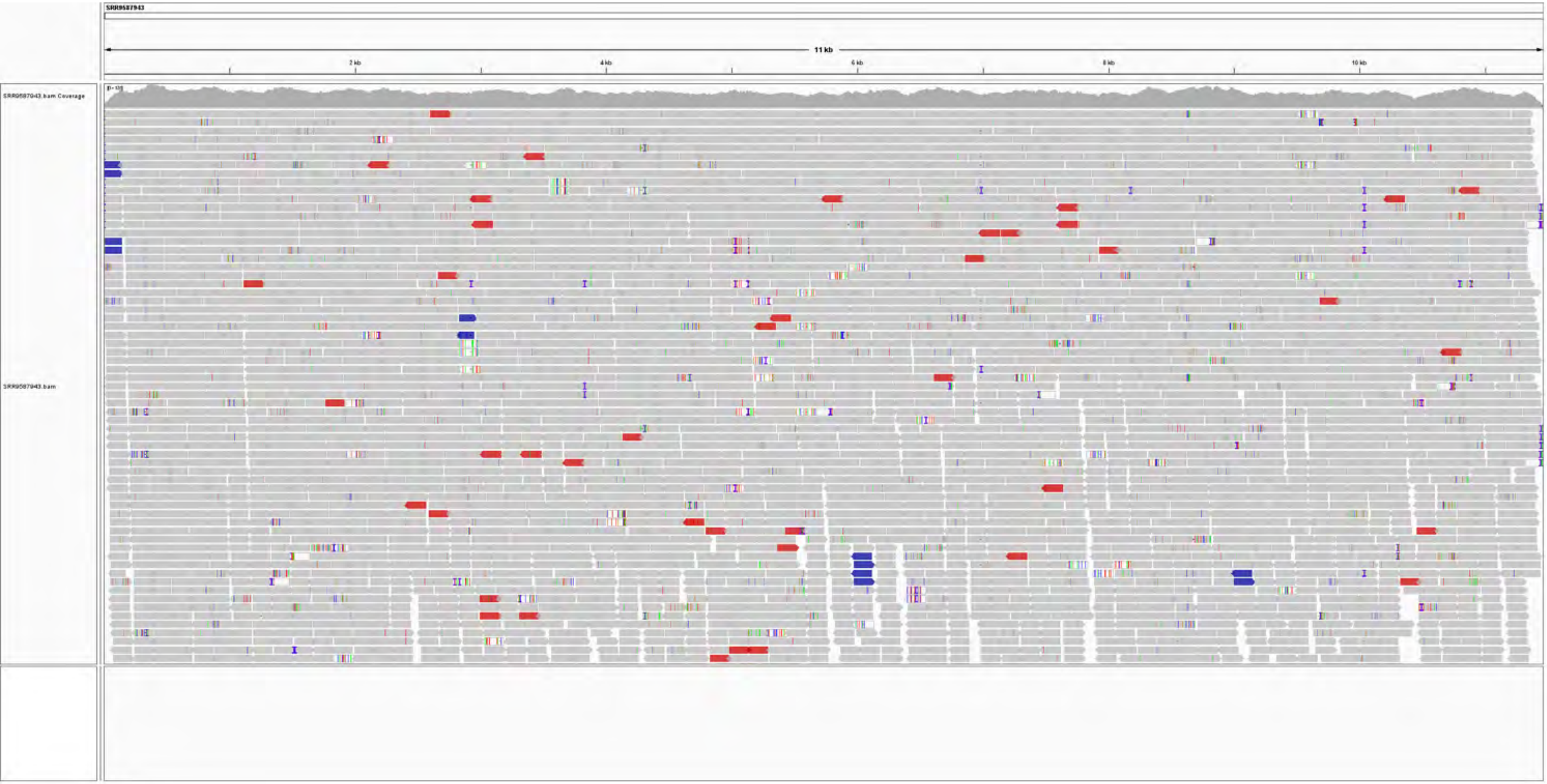
SRR9587943
11 kb
2 kb
4 kb
6 kb
8 kb
10 kb
SRR9587943.bam Coverage
SRR9587943.bam

### SRR9587944

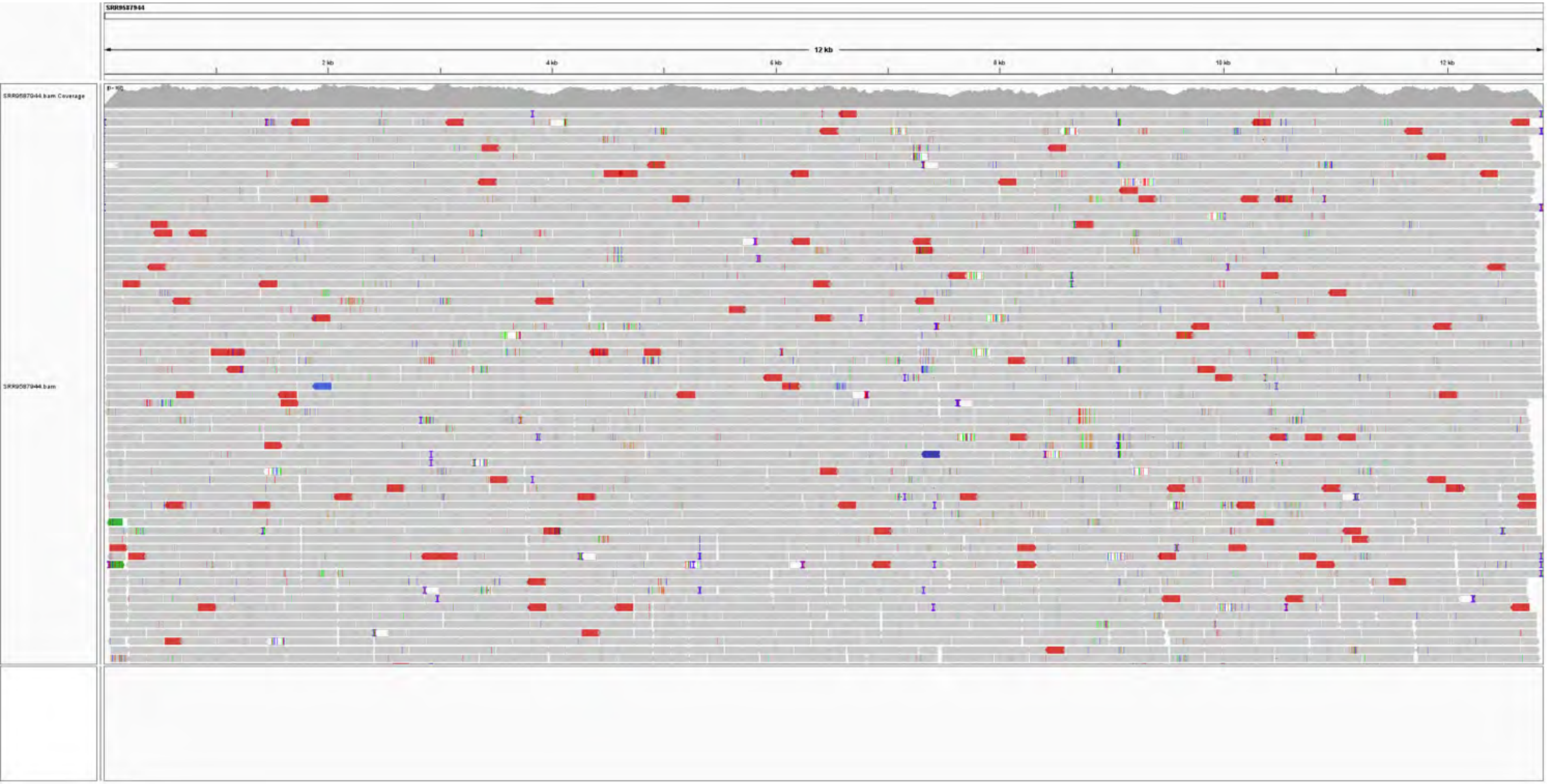
SRR9587944
12 kb
2 kb
4 kb
6 kb
8 kb
10 kb
12 kb
SRR9587944.bam Coverage
SRR9587944.bam

### SRR9587945

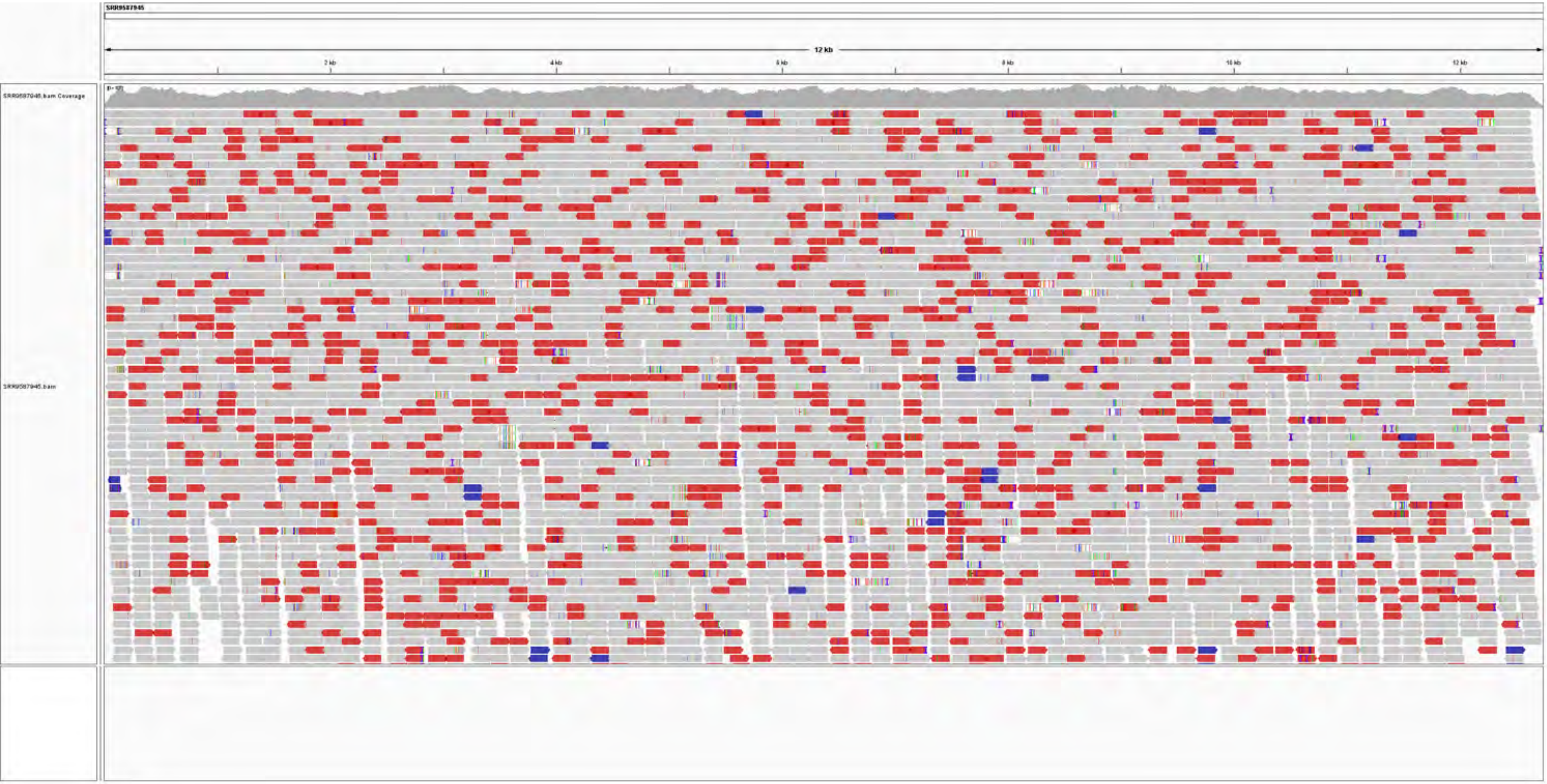

### SRR9587946

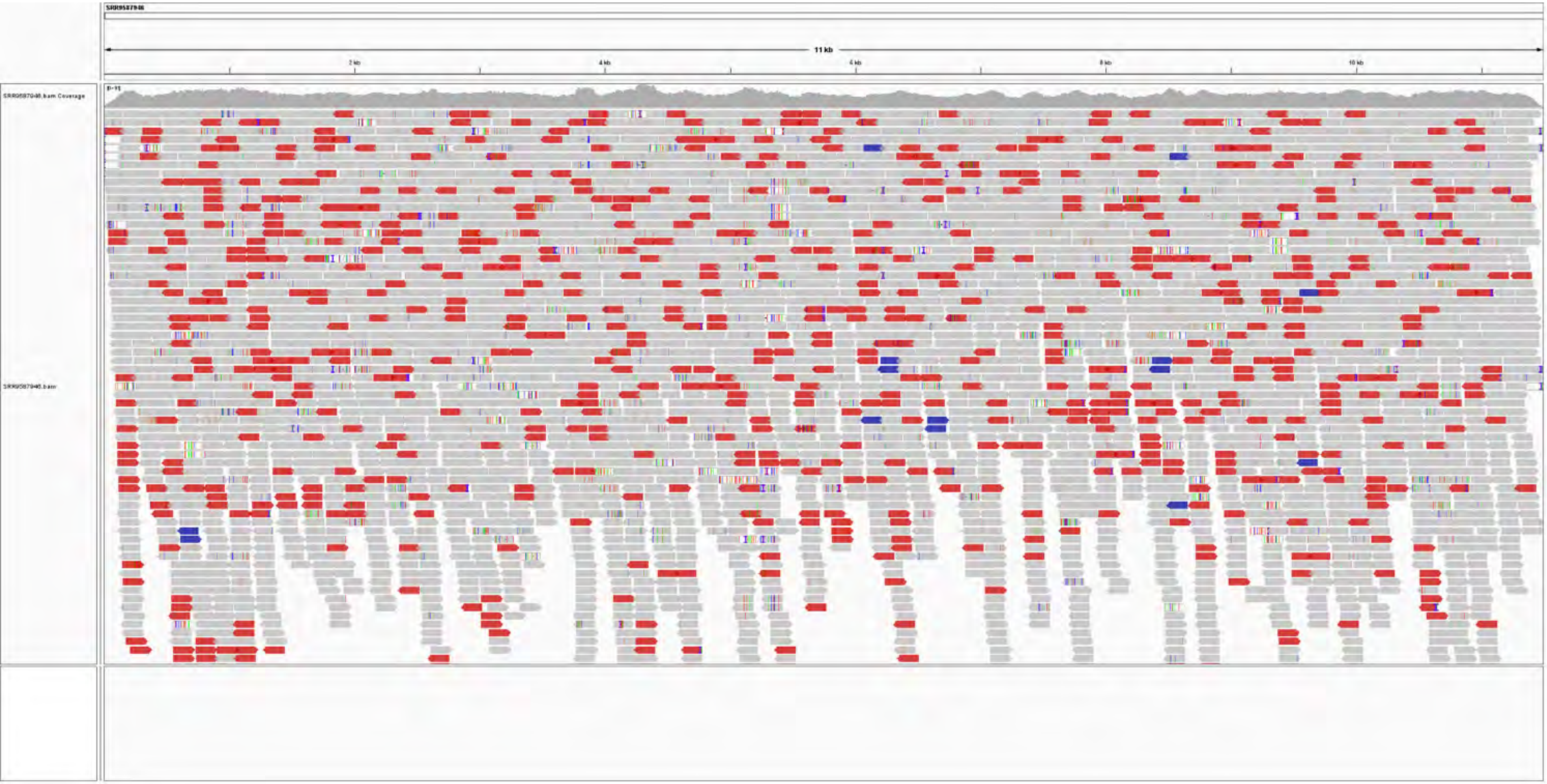

SRR9587946
11 kb
2 kb
4 kb
6 kb
8 kb
10 kb
SRR9587946.bam Coverage
SRR9587946.bam

### SRR9587947

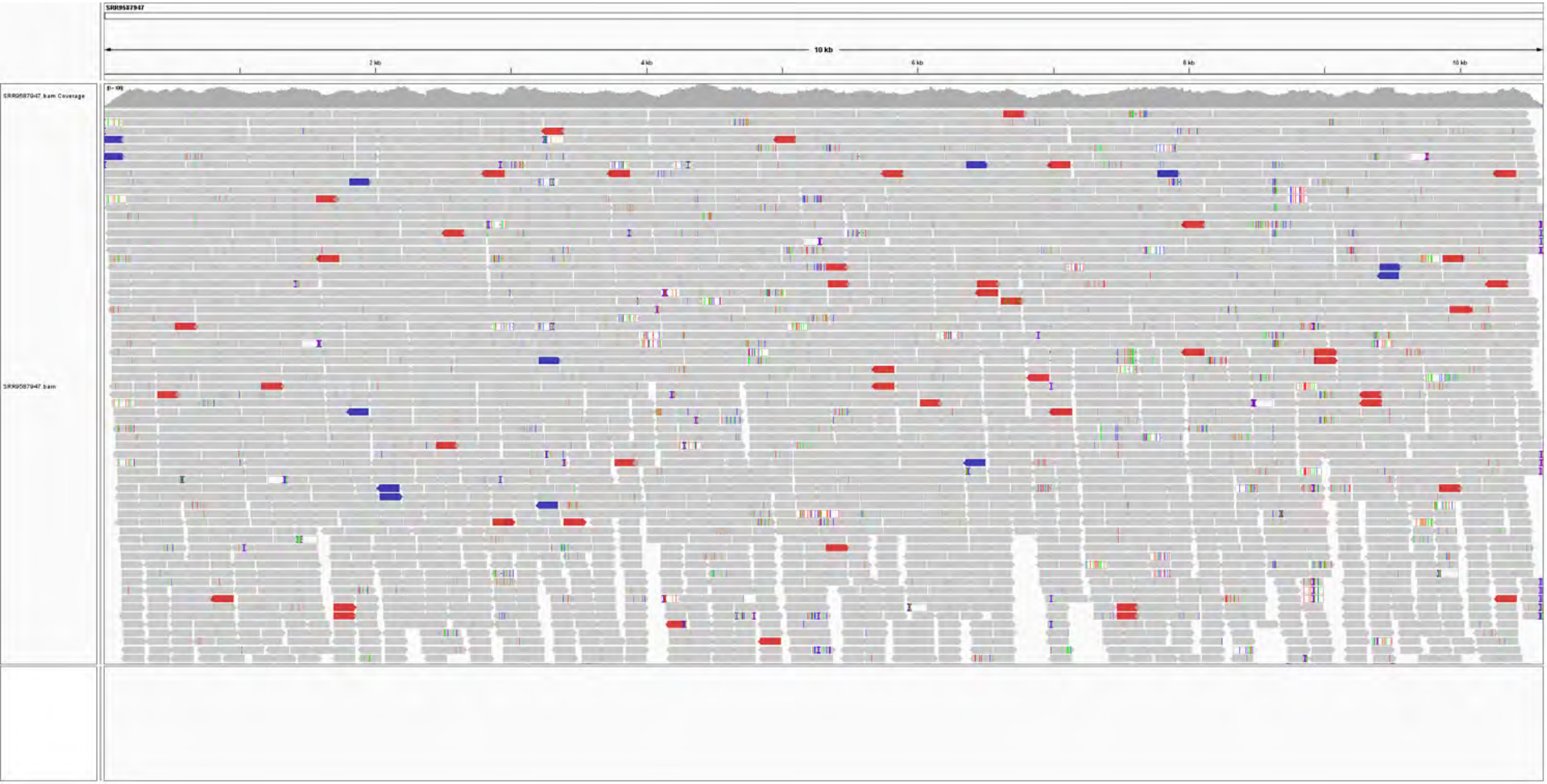
SRR9587947
10 kb
2 kb
4 kb
6 kb
8 kb
10 kb
SRR9587947.bam Coverage
SRR9587947.bam

### SRR9587949

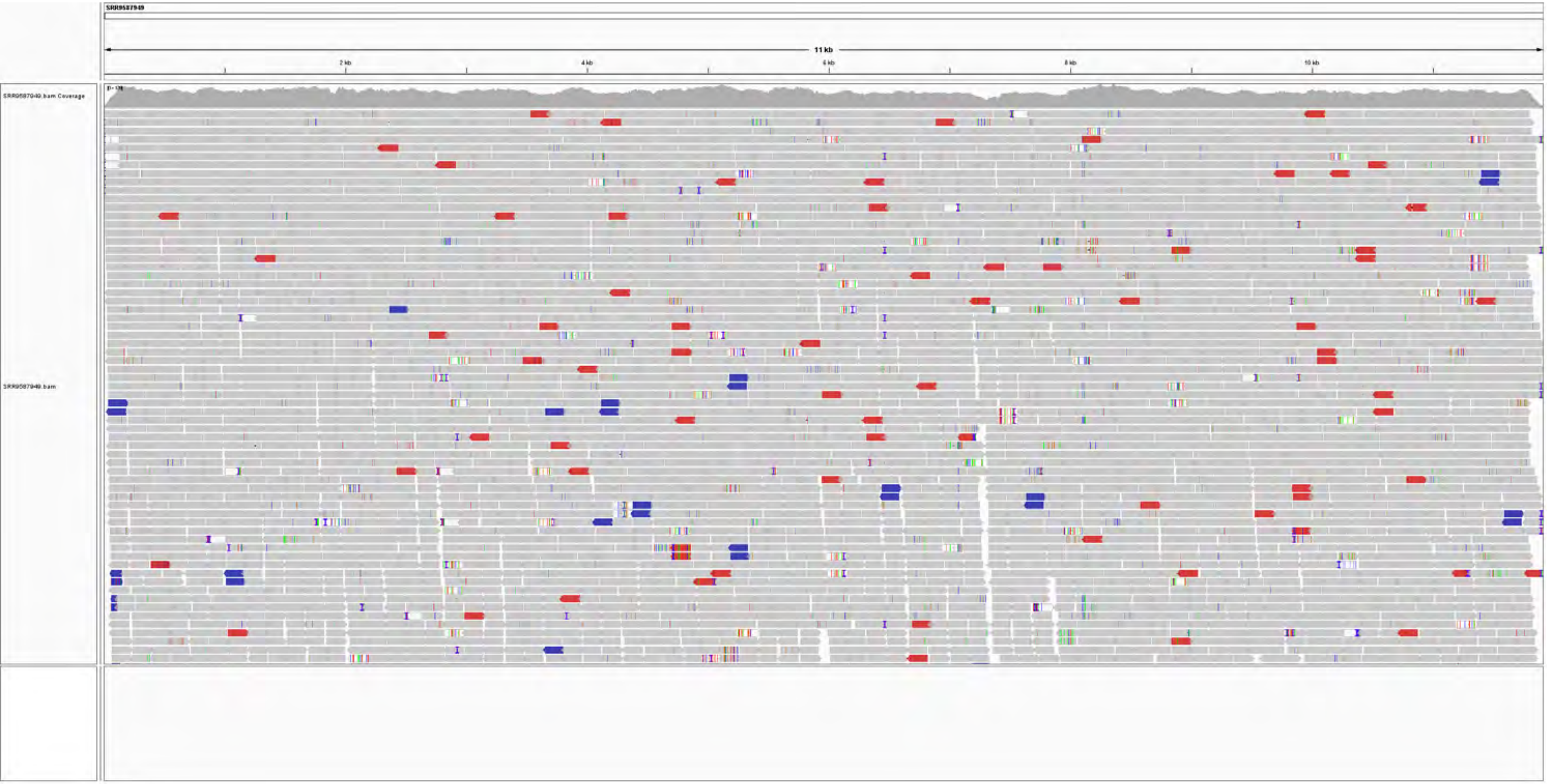

SRR9587949
11 kb
2 kb
4 kb
6 kb
8 kb
10 kb
SRR9587949.bam Coverage
SRR9587949.bam

### SRR9587950

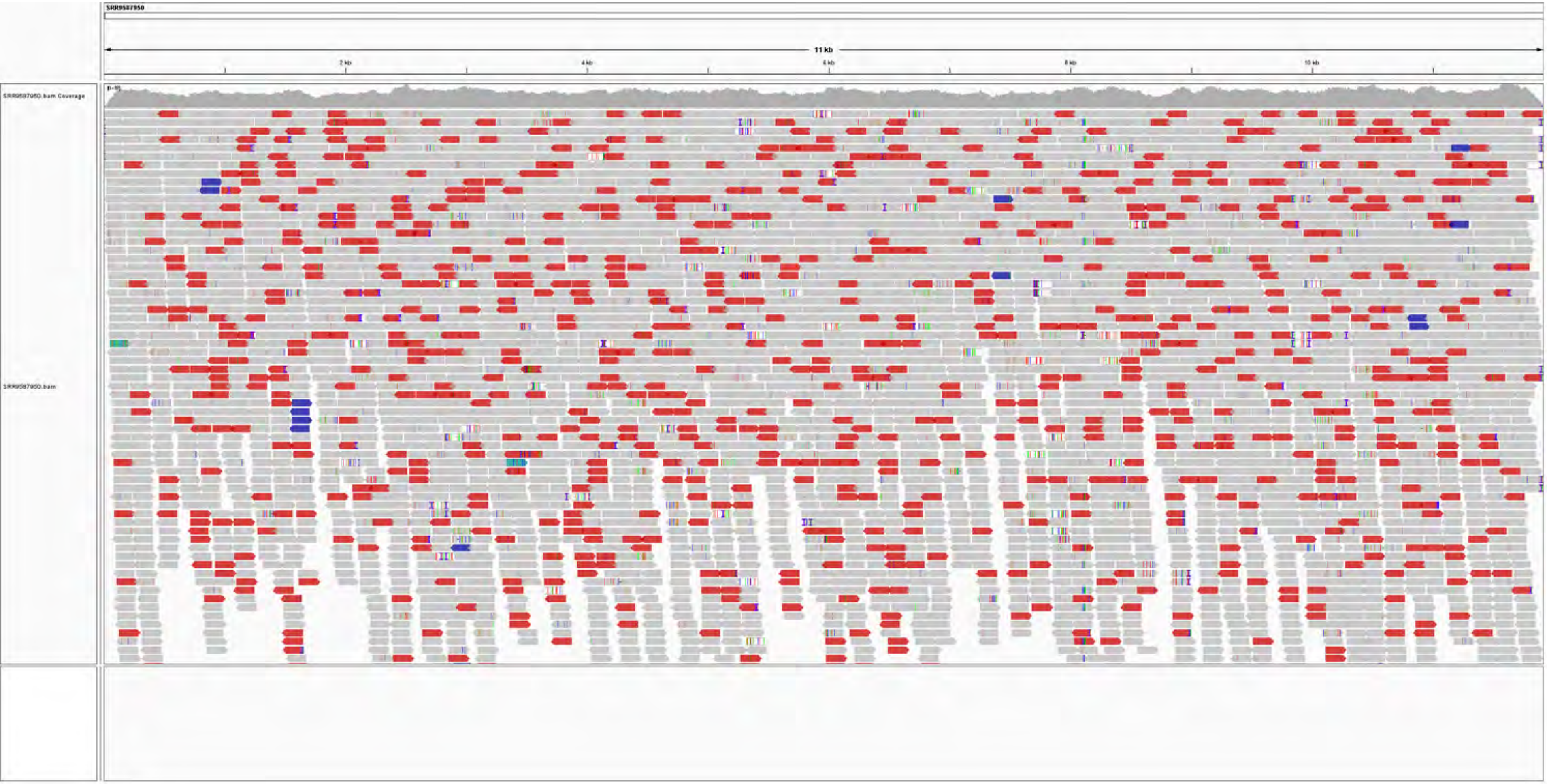
SRR9587950
11 kb
2 kb
4 kb
6 kb
8 kb
10 kb
SRR9587950.bam Coverage
SRR9587950.bam

### SRR9587951

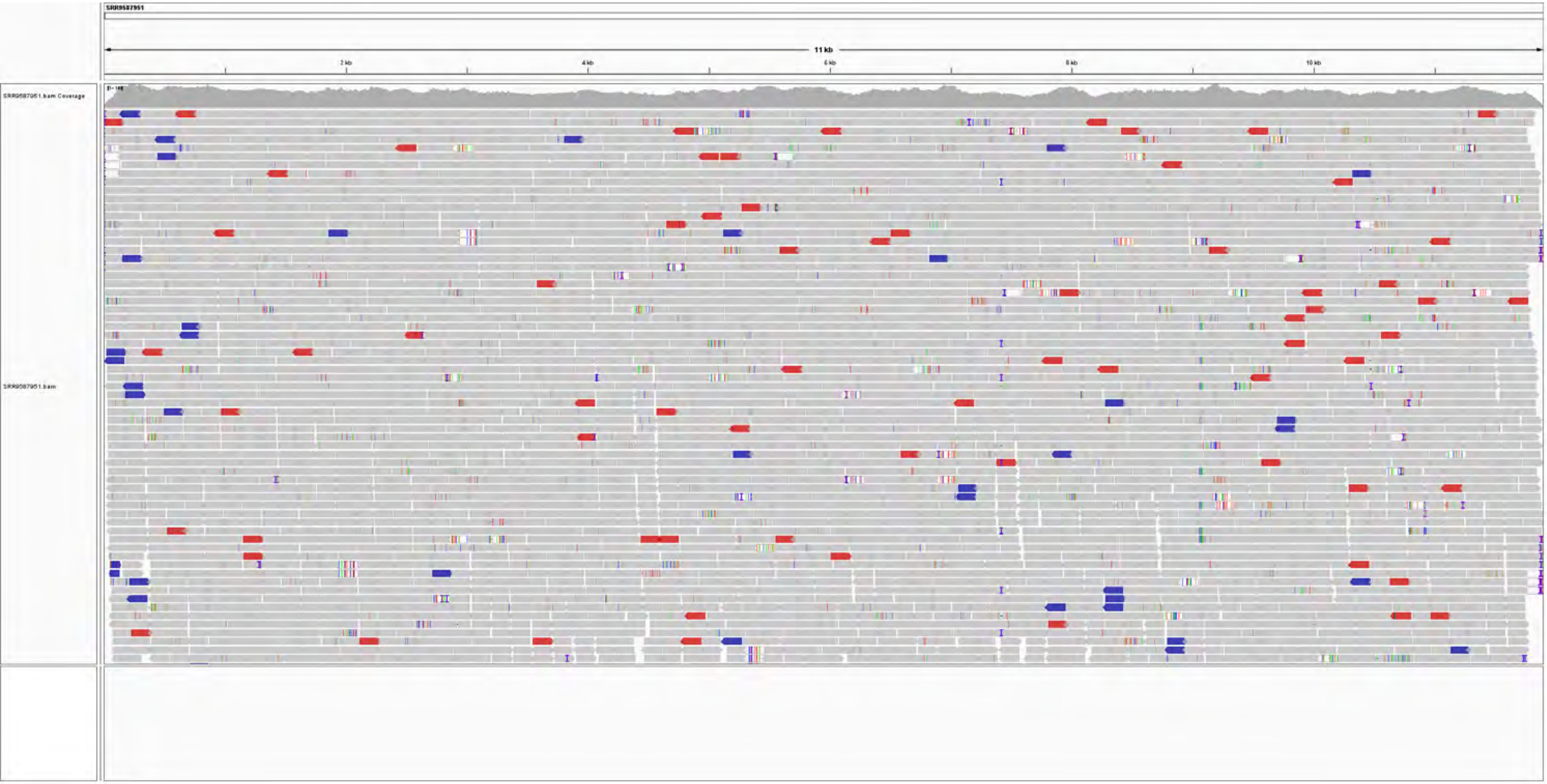

### SRR9587952

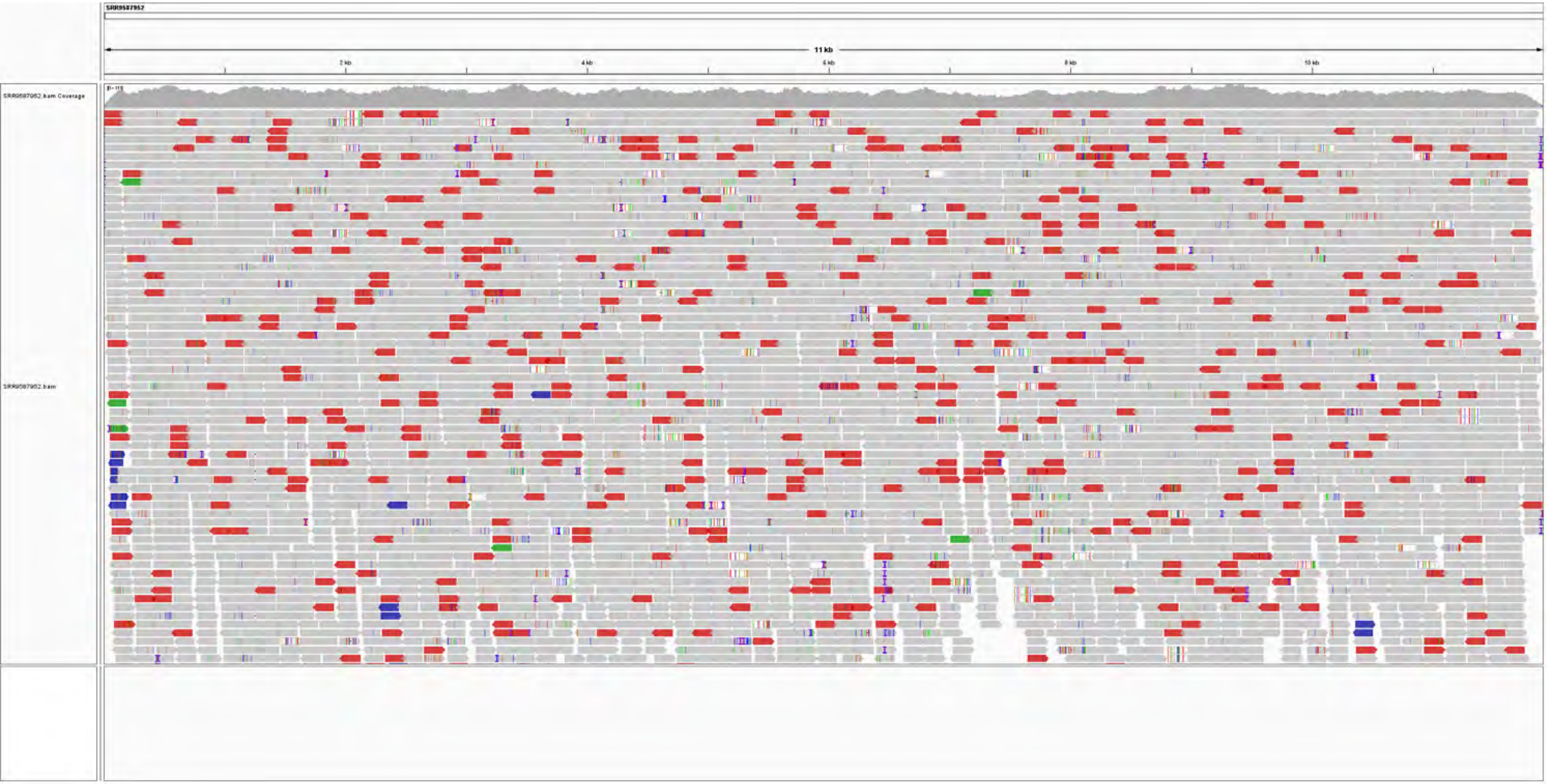
SRR9587952
11 kb
2 kb
4 kb
6 kb
8 kb
10 kb
SRR9587952.bam Coverage
SRR9587952.bam

### SRR9587953

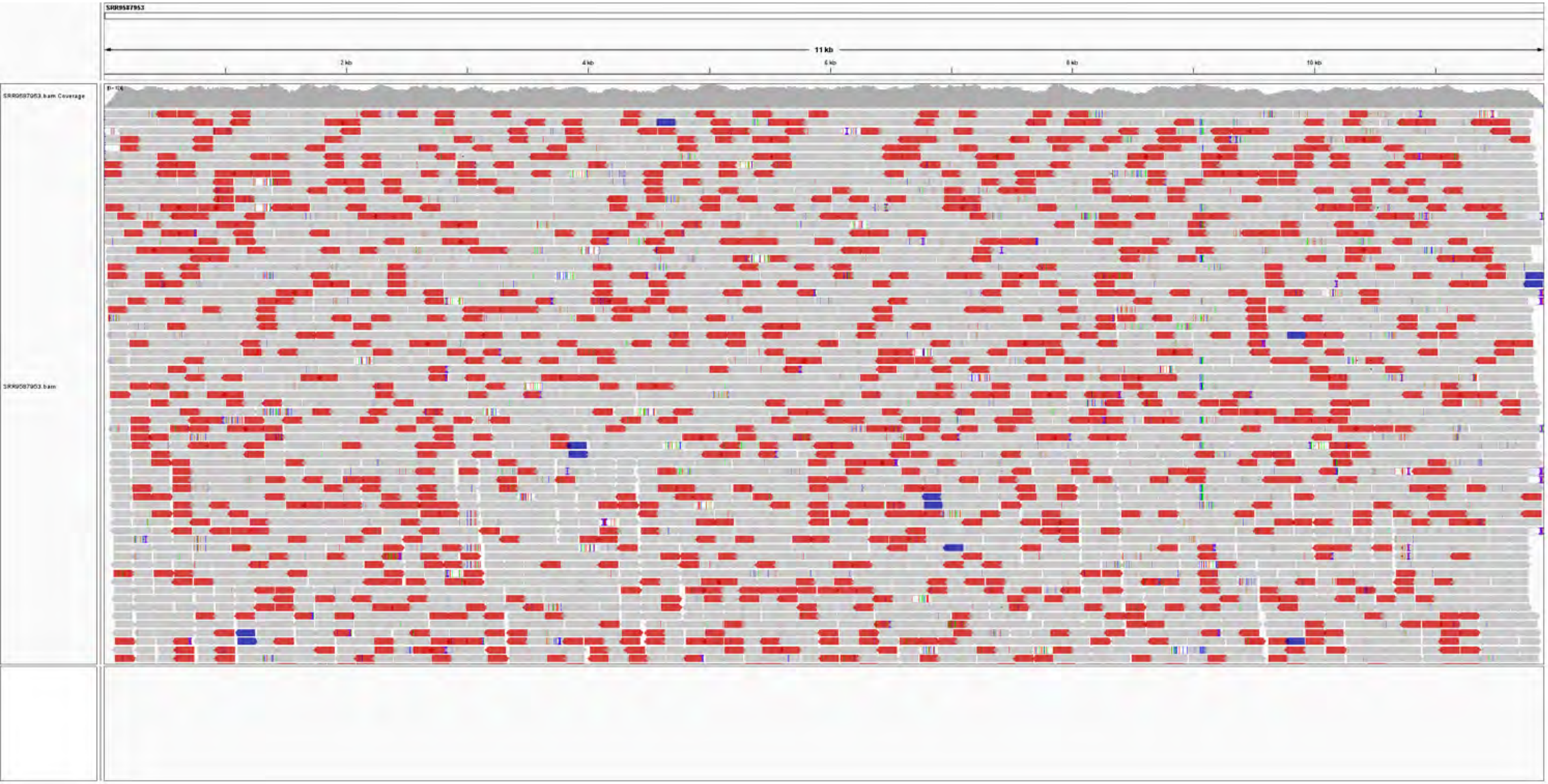
SRR9587953
11 kb
2 kb
4 kb
6 kb
8 kb
10 kb
SRR9587953.bam Coverage
SRR9587953.bam

### SRR9587954

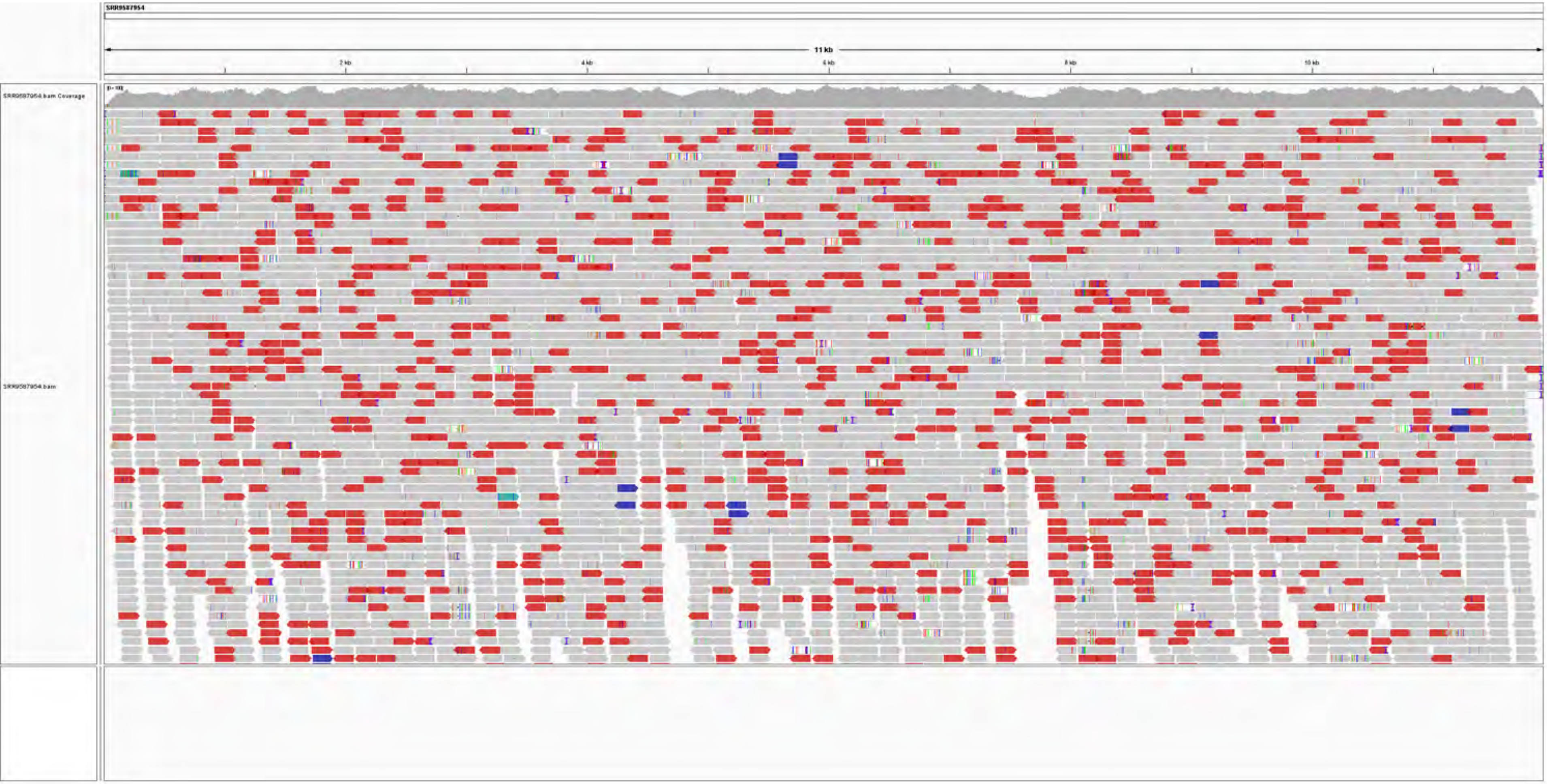

SRR9587954
11 kb
2 kb
4 kb
6 kb
8 kb
10 kb
SRR9587954.bam Coverage
SRR9587954.bam

### SRR9587955

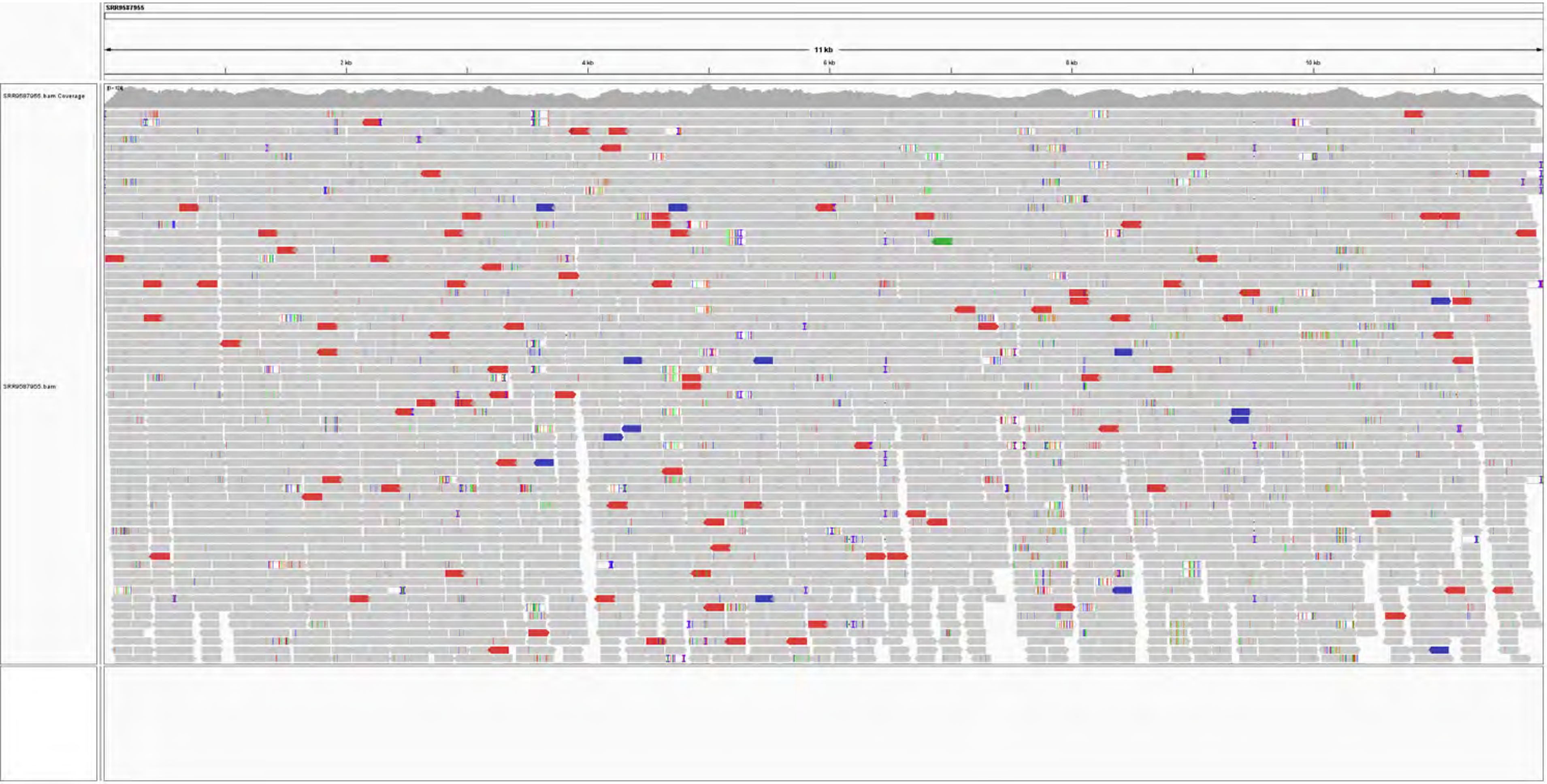

### SRR9587957

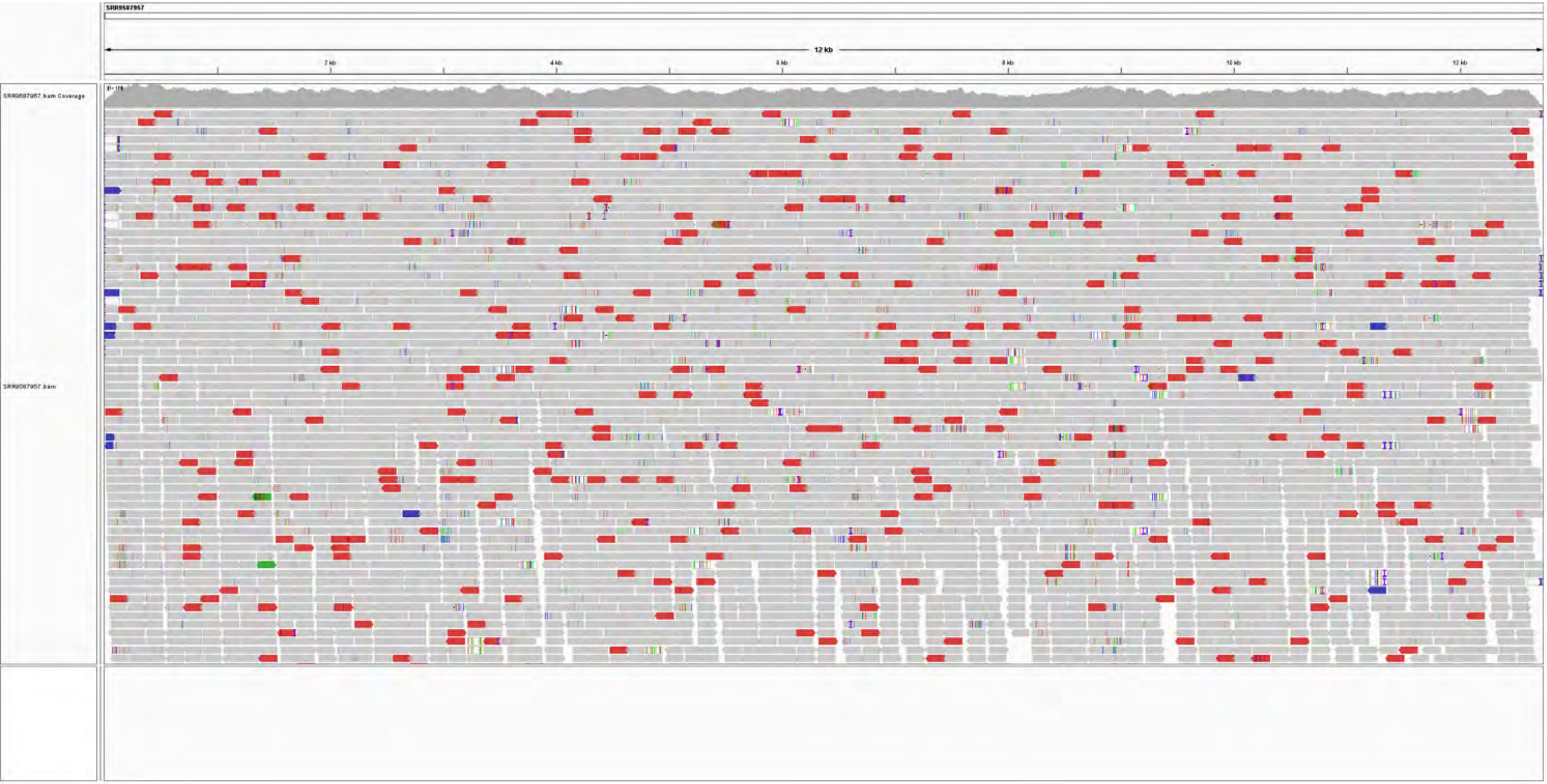

### SRR9587958

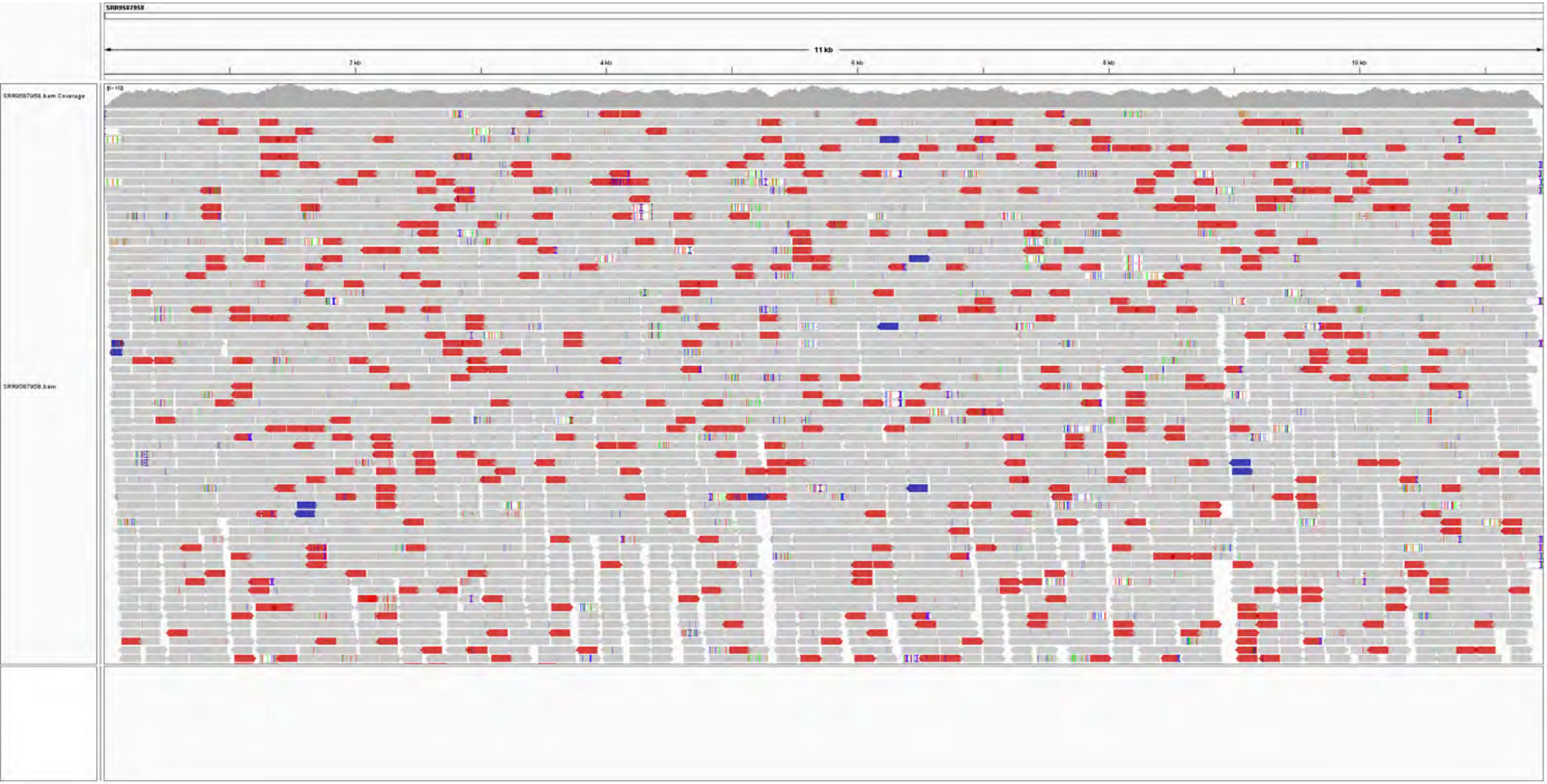

SRR9587958
11 kb
2 kb
4 kb
6 kb
8 kb
10 kb
SRR9587958.bam Coverage
[0 - 112]
SRR9587958.bam

### SRR9587959

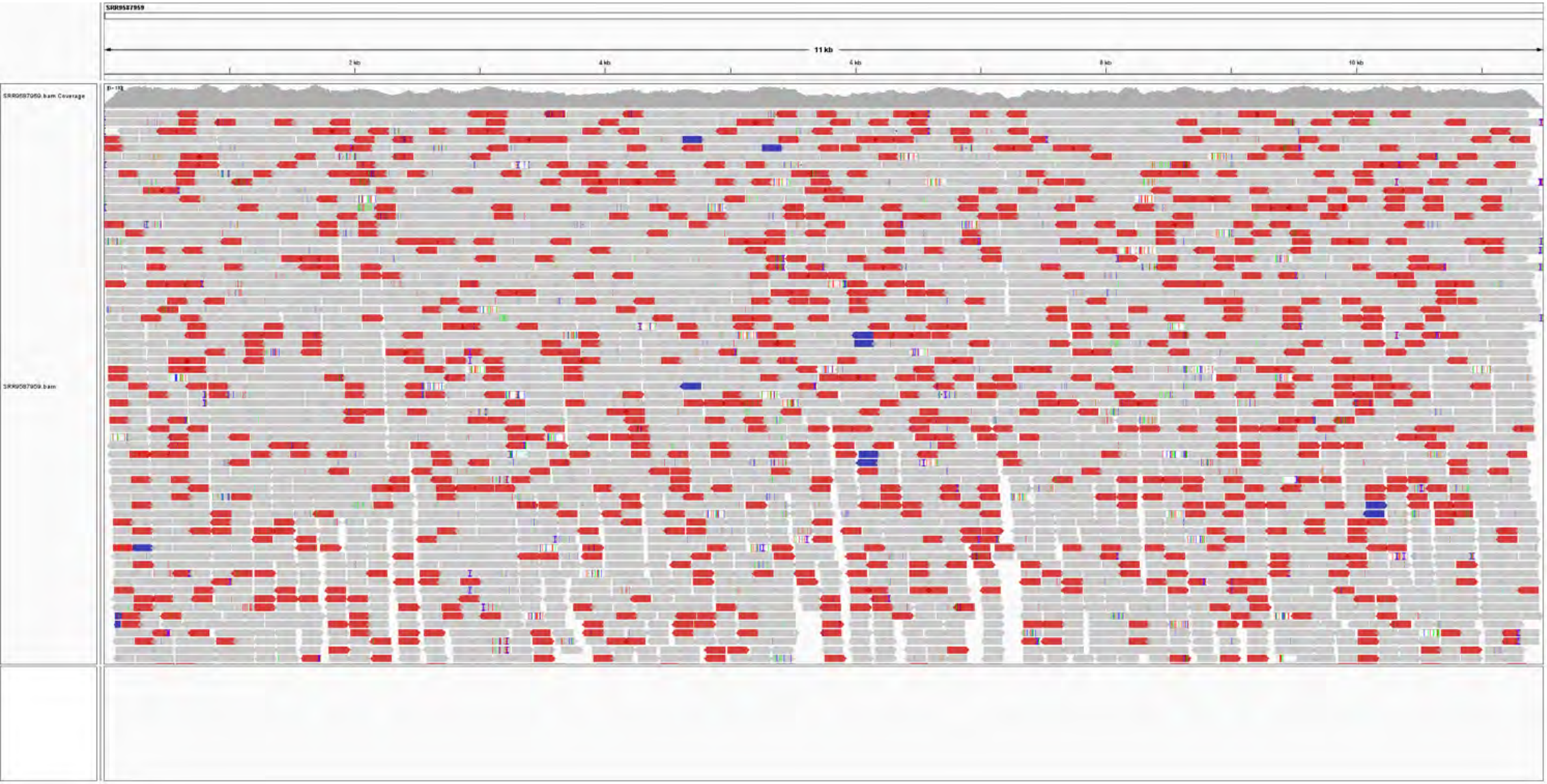

SRR9587959
11 kb
2 kb
4 kb
6 kb
8 kb
10 kb
SRR9587959.bam Coverage
SRR9587959.bam

### SRR9587960

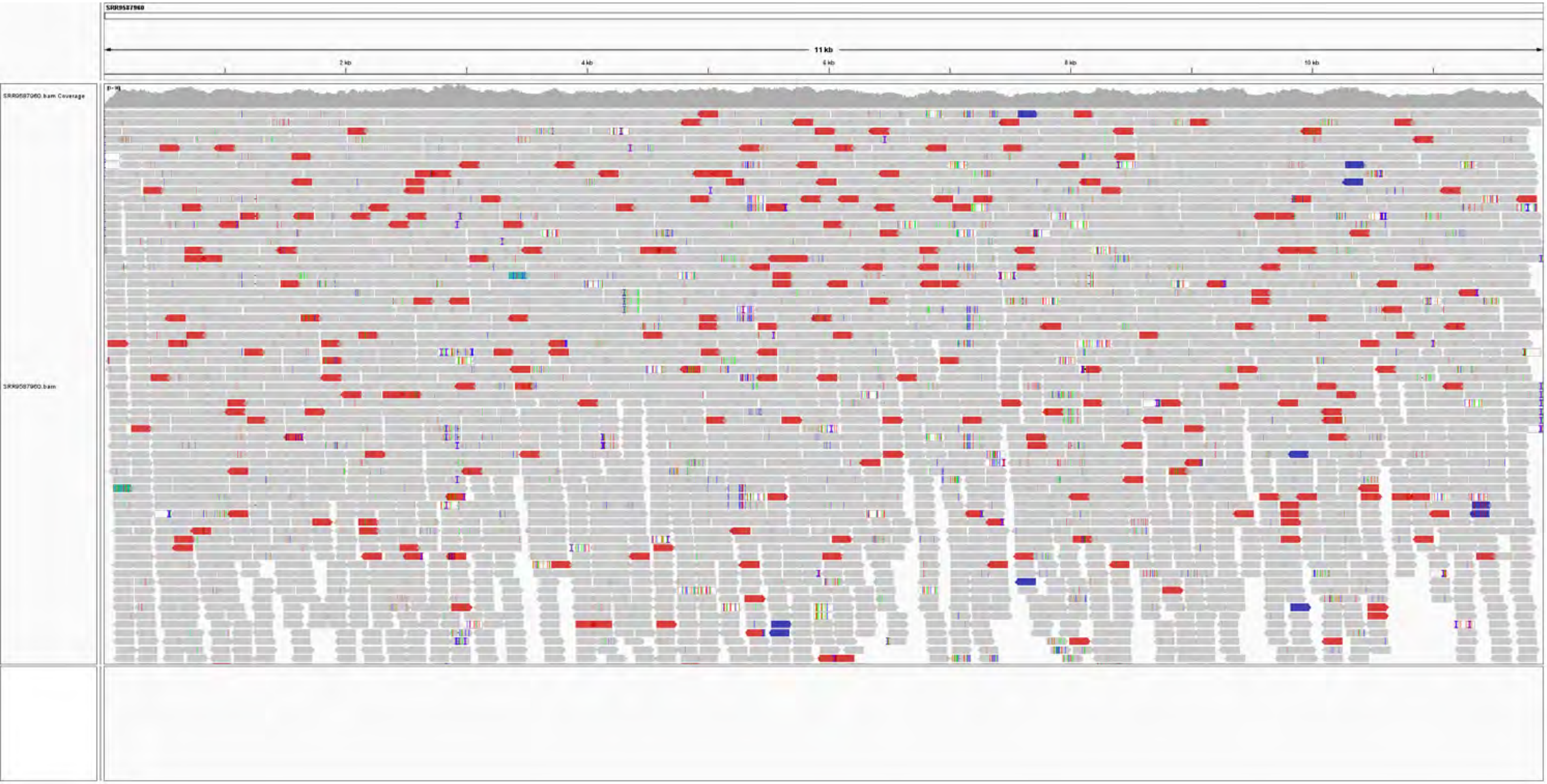
SRR9587960
11 kb
2 kb
4 kb
6 kb
8 kb
10 kb
SRR9587960.bam Coverage
SRR9587960.bam

### SRR9587961

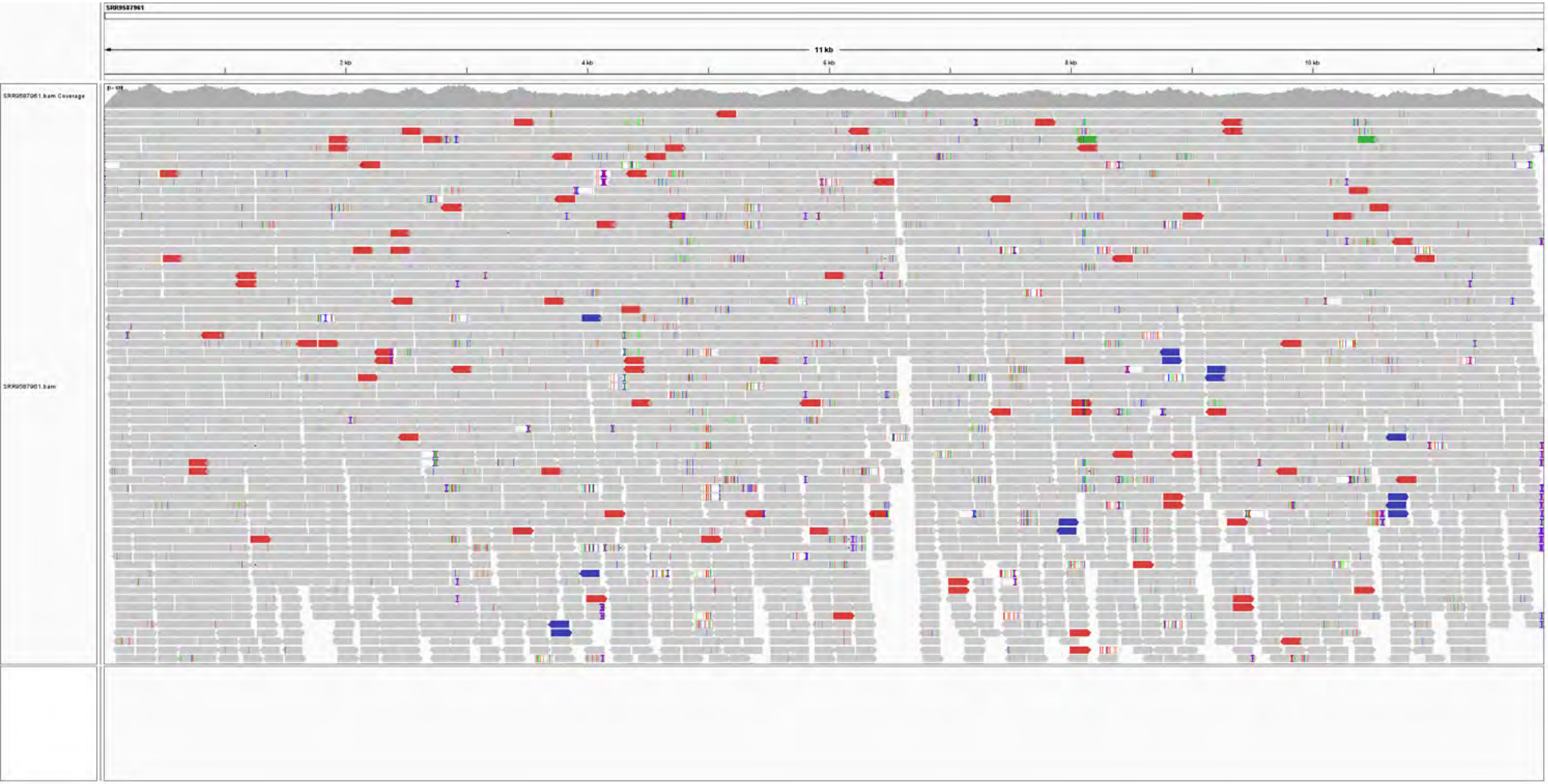
SRR9587961
11 kb
2 kb
4 kb
6 kb
8 kb
10 kb
SRR9587961.bam Coverage
SRR9587961.bam

### SRR9587962

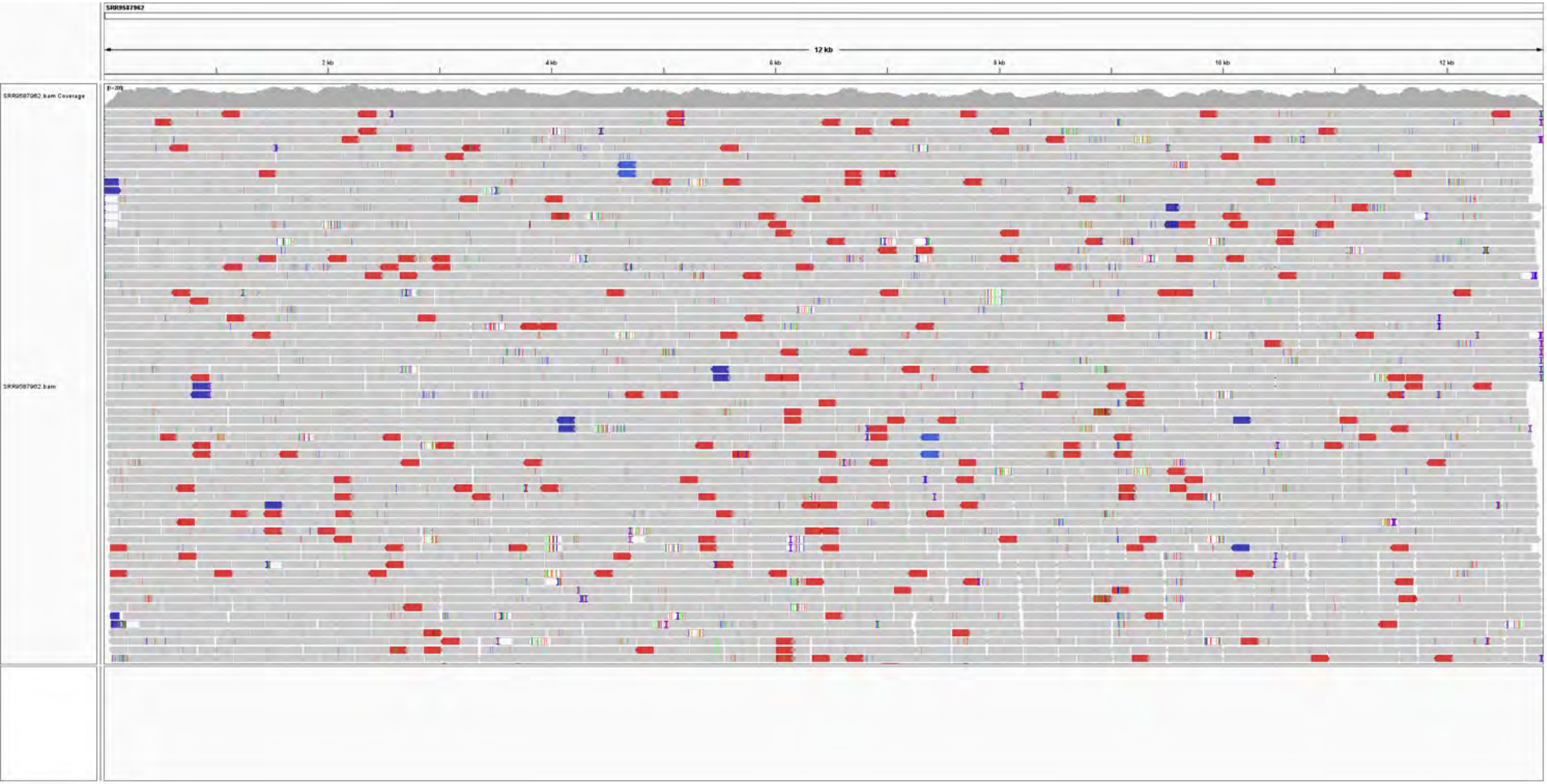
SRR9587962
12 kb
2 kb
4 kb
6 kb
8 kb
10 kb
12 kb
SRR9587962.bam Coverage
SRR9587962.bam

### SRR9587963

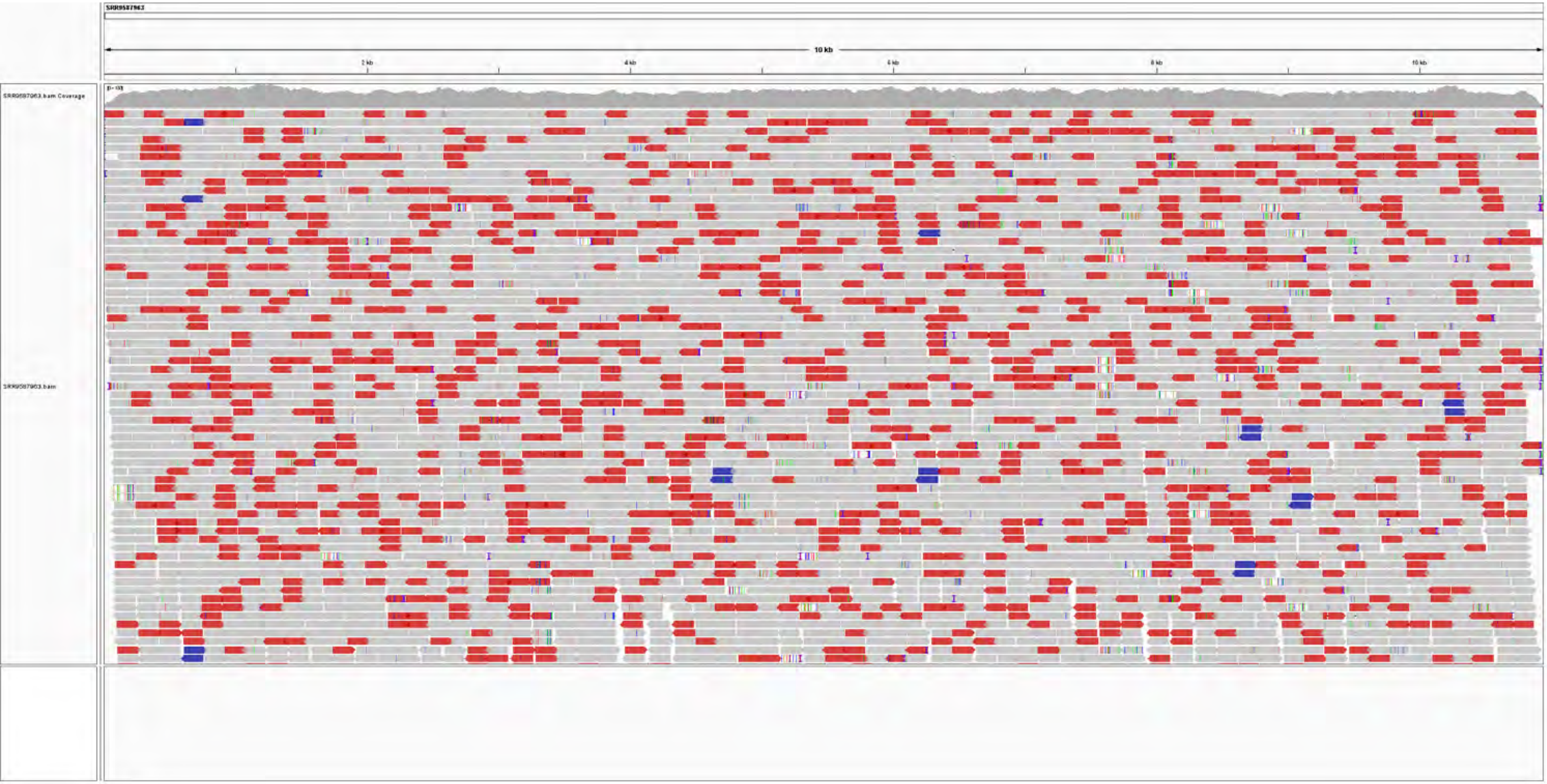

SRR9587963
10 kb
2 kb
4 kb
6 kb
8 kb
10 kb
SRR9587963.bam Coverage
SRR9587963.bam

### SRR9587964

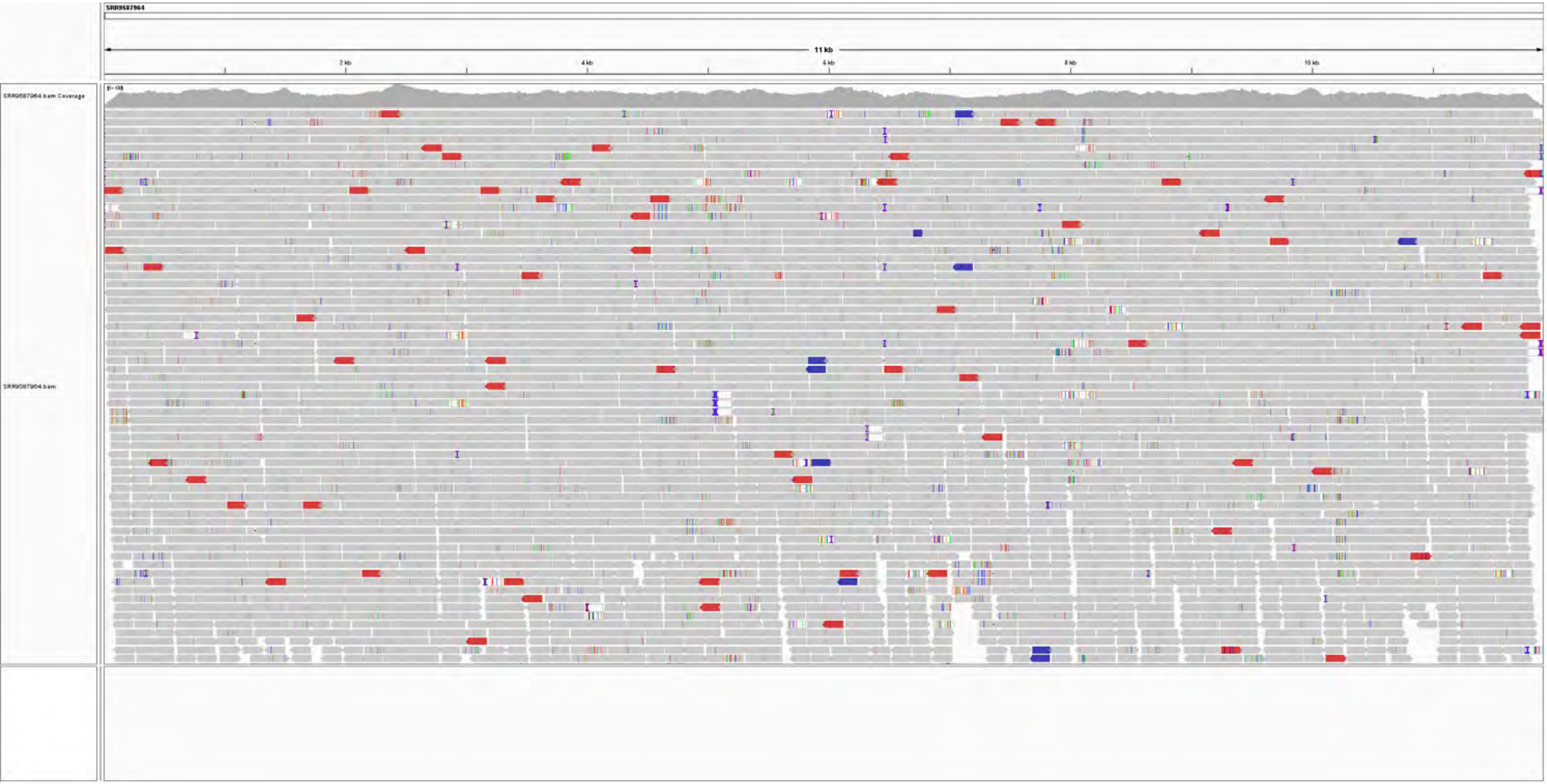

SRR9587964
11 kb
2 kb
4 kb
6 kb
8 kb
10 kb
SRR9587964.bam Coverage
SRR9587964.bam

### SRR9587965

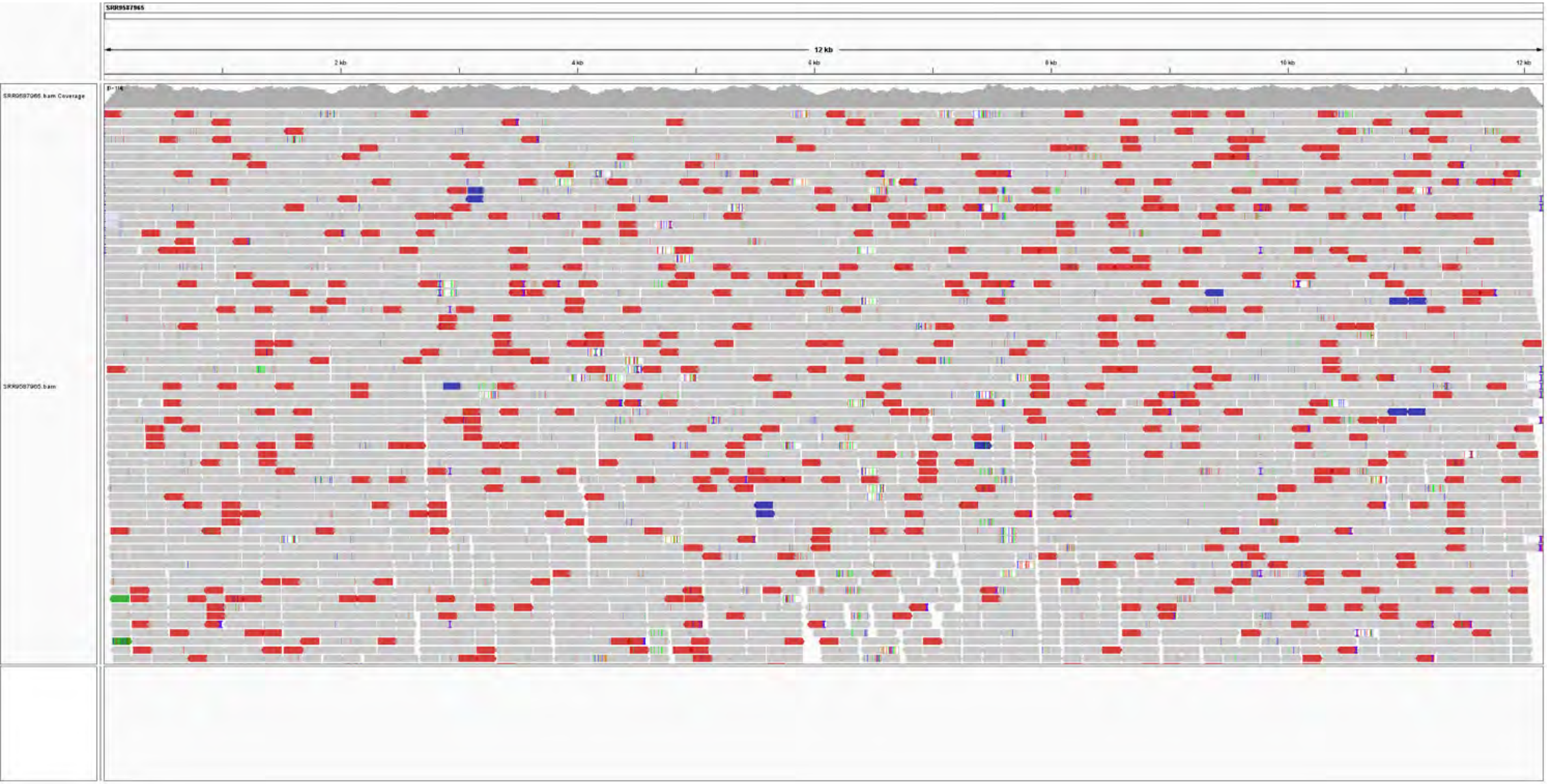

SRR9587965
12 kb
2 kb
4 kb
6 kb
8 kb
10 kb
12 kb
SRR9587965.bam Coverage
SRR9587965.bam

### SRR9587966

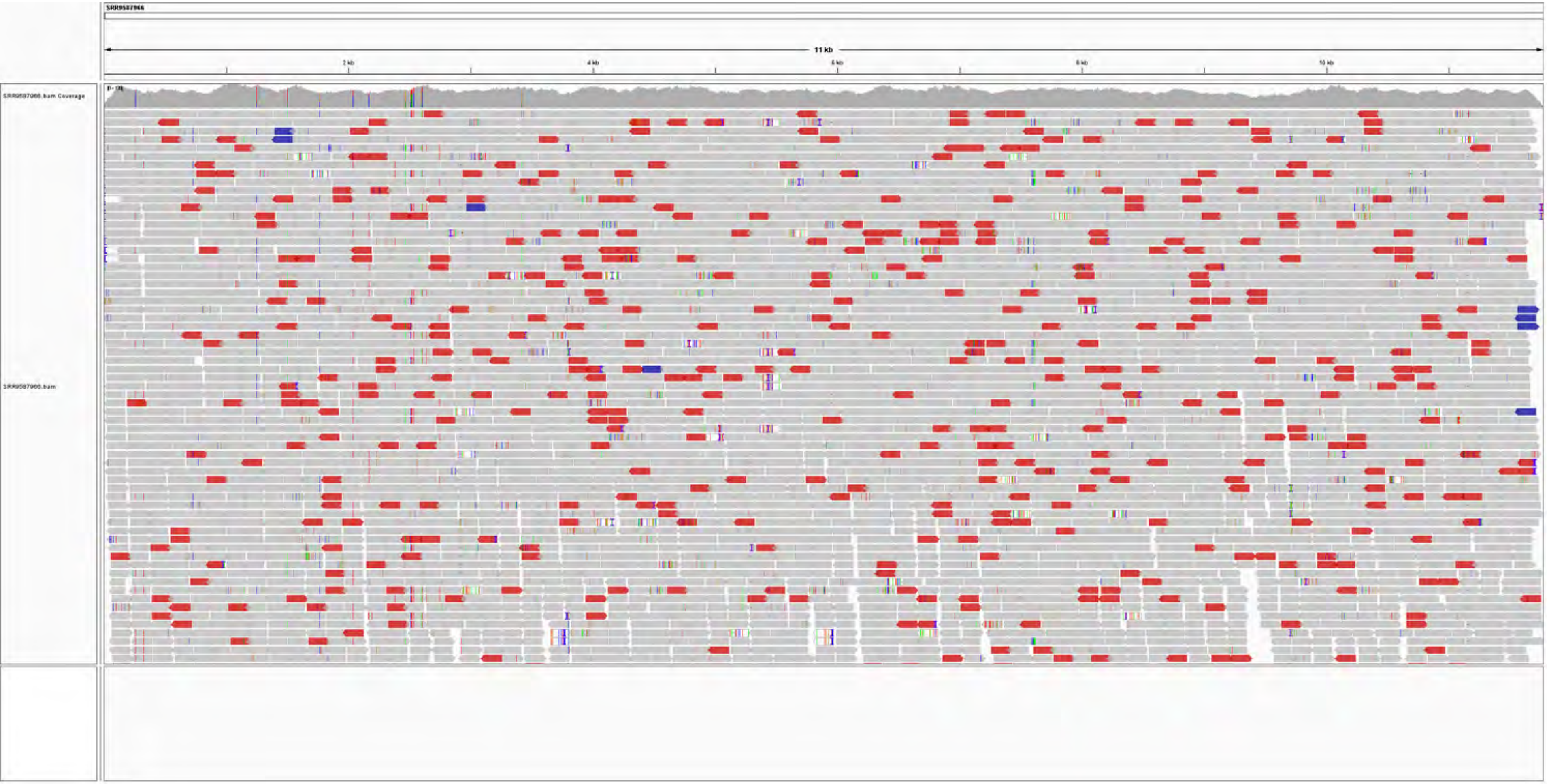

SRR9587966
11 kb
2 kb
4 kb
6 kb
8 kb
10 kb
SRR9587966.bam Coverage
SRR9587966.bam

### SRR9587967

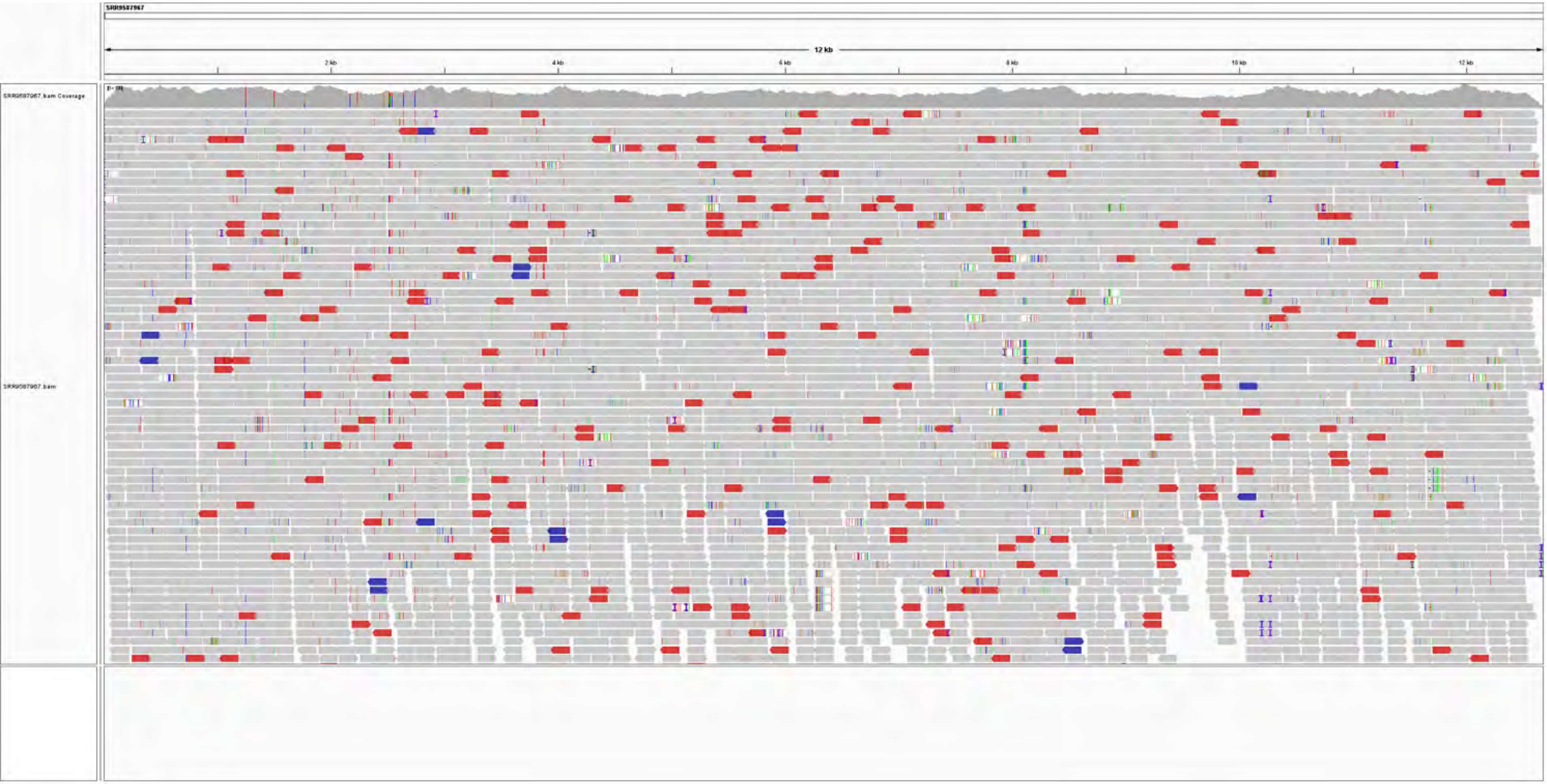
SRR9587967
12 kb
2 kb
4 kb
6 kb
8 kb
10 kb
12 kb
SRR9587967.bam Coverage
SRR9587967.bam

### SRR9587968

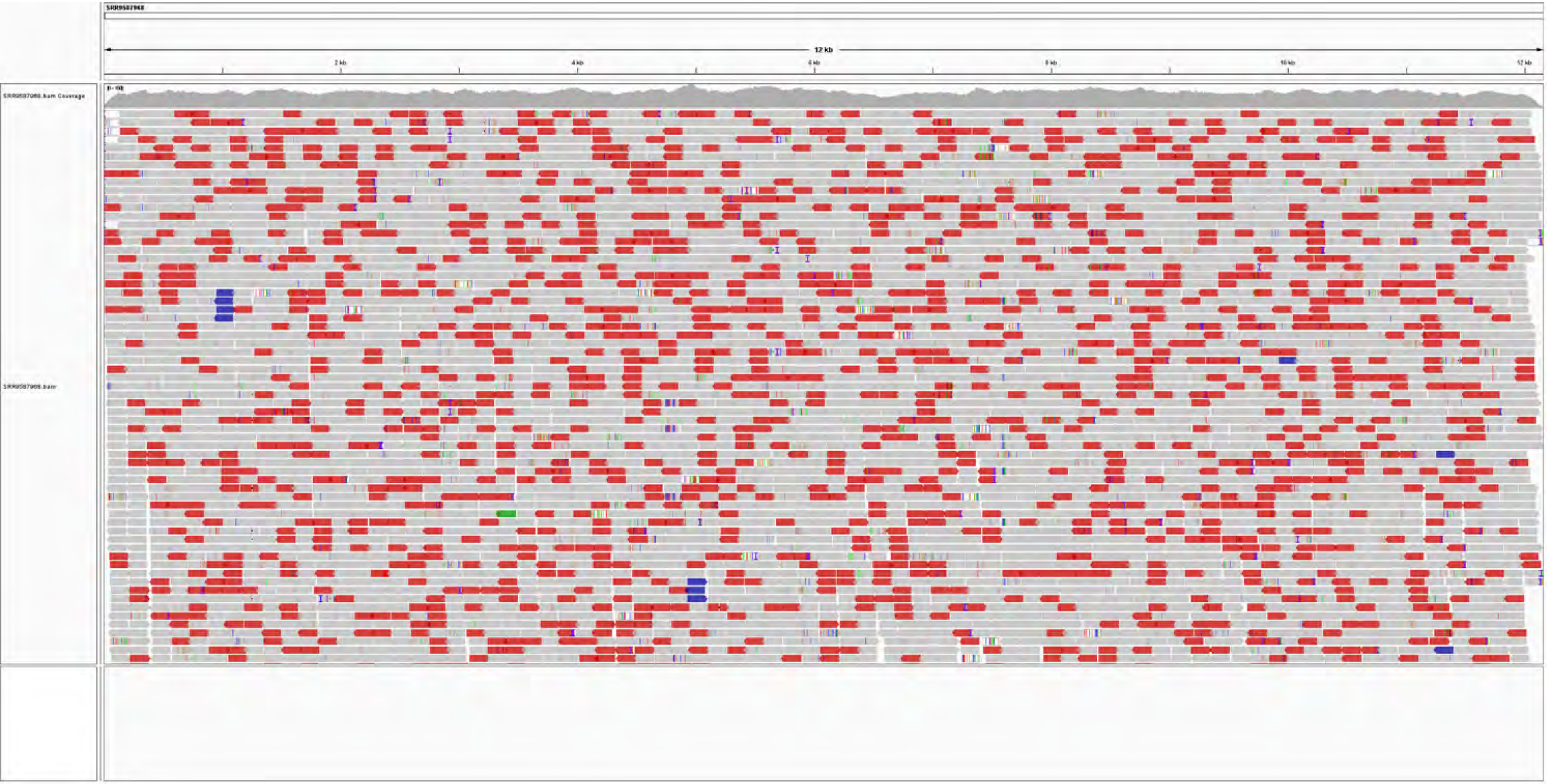

SRR9587968
12 kb
2 kb
4 kb
6 kb
8 kb
10 kb
12 kb
SRR9587968.bam Coverage
SRR9587968.bam

**Supplementary Figure 3.** Results of StructRNAfinder analysis using the sequence SRR9587944 (sequence with four introns) as query.

| | Infernal | | | | | | RNAfold | |
|---|---|---|---|---|---|---|---|---|
| Sequence | RNA family | Id | From_seq | To_seq | Score | Evalue | Score | Struct |
| **Intron** | | | | | | | | |
| FormA_SRR9587944_2 | Intron_gpI | RF00028 | 11309 | 11674 | 106.7 | 7.7e-30 | -132.70 | 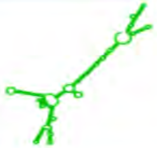 |
| FormA_SRR9587944_3 | Intron_gpI | RF00028 | 6100 | 6406 | 74.9 | 1.1e-20 | -109.80 | 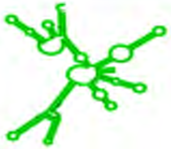 |
| FormA_SRR9587944_4 | Intron_gpI | RF00028 | 12498 | 12816 | 64.3 | 1.2e-17 | -115.80 | 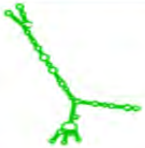 |
| FormA_SRR9587944_5 | Intron_gpI | RF00028 | 3892 | 4225 | 33.9 | 6.7e-09 | -108.60 | 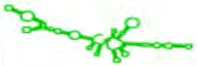 |
| FormA_SRR9587944_6 | Intron_gpI | RF00028 | 10410 | 10787 | 31.1 | 4.5e-08 | -145.90 | 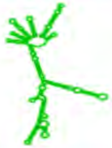 |

**Supplementary Figure 4.** Multiple alignments of the 82 DNA sequences.

>Form_A_Japan
gcatgttttggtatgtgtttcgacagtcatgcaagtcatagccgtaggatggggagtgag
cttcatagcccgacccccttctcaggagcatgttttcatagcatattgagcatactggggt
ggtctcccatgtagctcgctctctcggtgcaggacctcccttttgcgggccacttcgatg
gcctccaataggga-gagcatgttttggcatatgaagcgttggtggtgaggctcgtagcc
tgtatatacataccaacgcagtcgtcatatgtctctctggcgggggtcggtcgtatgacc
ttgtccctacctggagcatgtatttcgtatgat---atactggac-ggcagctgcgctgt
tcagtcgtcagccgtacctcatgtgcgcgtttccaggtccgaacgtttggcatgcgaggg
catcataccagaagcctgttttctcgtatgagcatgtaggtaggcaggctcttcagtcgt
ctacccataaatgcacttcgaagga-actttttggtattgagtggctttgtgcccagacg
ttcttgtgagcttgttctcgctctacgatacgctggatgctgtggtattacctacagcag
acgcattcctagggcgggcggcctctctgcagatgagcacatctgtagaaagaagctccc
cttgatgacgcaatgcaatggctgggtaaacggagtgctatacagaagggtcaaacctc-
tgtggaagtacacccataccagctttgtttgttgagaggggtcggctctacgagtgaggc
cctatactgccatctcattcagatgcgtagacggtcttatctcagactgcctttctggct
gattgtcctcggtccctcgtgcacacgcgtcgctggctgaatgggcttgtccttatggag
tgccgcgttcttc-aaagaatccgtcttcatacagcgacctcgtttacatattgccggct
gaatgggcttgtccttatggagcactgcattcttctaggaattggtatctatacagcgac
ctcatttacatacaaggcacctgttttctacatagacttgccatttggctgattgttctg
acagtcgctgcctcttcggagttctgtgtggacgagtggcc--ttctgagtgggccggtc
cttacggagtccccatatctgttatgactgggttctccatacagcgacctcactcagaat
tgtatgg--------------caccgtgccgat-----------tgaatggacaagcagt
ccttagggaatcgcatgaaagcatgacggc-acatggtgtatggacccgttgggccttgc
ctgtgctatcttctcgagtcgaccgg------------tcctcgtctcctcctttcgcga
cgggaagctcttgagtttaccgtgggtcaggacgctgtggtatctgtttgtctgtagtgt
gggctctctatcagaacagcgtatcagcacacgtcgtctgggtctatggtatttttgctt
aacggccctgggtttggtagtatggttccatgtcggtcctctcacctcccatccgtcatg
tctctga-----------------------------------------------------
------------------------------------------------------------
--------------------------------------------tatcatctgcacagta
cgtcttgtcagaatatcattctgtatggcagtcgatagactaggccctatgccgcagctg
tctctgaccactatgcgagcgcatacctacctcaatgtgttcatagctgttatggatctt
gggtgtggtgtgccagaacgatacgccaccagattgtggaatggtgcttgtagtattaga
agaggacatgtatgaagagcacgttcttttctcttatgatggtgccgcgggtgggggccg

ttctcattctgcttgtcgggtgctcaaggtgcatgtaggattgctgaacccac-tgaatg
gtcatgagaatggtcatatggtgaggttcacttctccatcgctttgtgtgtgttggatgt
ttgggtggcttcggccgcatggaaaatctcggtctgtaactcatcagggagggtttccat
gtgagcaatctctccagtcgaacacacacccatgggcgtcccgctaggcaaaatggaact
ctcgatccgacacagcctgtggagtgtctgttccgtgtggcgtgaaacggcttatactgg
atgtcttccc-tttggacaccagtctgttggtatg--tacgcataccatctagagggagc
aagaaaaaggcatgcaagtgccacaaaggaagggaggccttcctgctaccgtgtgtcc-t
cttgacgatcacgggggagtttatagagacatttggacgagatcctttttttcacgaggcg
gtgccctcgcgcaaaggtgttgctctgcacgcatctttatgatgatagttgctcgctcct
atgcgtccactcgtgaaaatacctgctctctctgtctgtttccatgacacccc-catgcg
ttgtcatagagcacgctcaggtacagaaagtgctgtgactttcttggcctgctgtcactg
agtcagcctgatactttggaggaatcttttctgggcgctgcccagctgagagttcttatc
aggctctggagtgtgcatatggctataatcataatagtgcatgtggccacgcagaaaaag
catcgacgtggaatggagaagaacaccattatgaatggaacggttcttactctgcttccg
ttgcttgacgctgtatgggaatcgctttgccctgggggtacaccttccattccataacaa
agcatttgatctatgccccga-gggtcattctctatgaagagcccgaaacttgtctgagt
gagaggcgaaggtagcccttcattt--atgcggggtccgccctcacactgtagtcggcac
agtcccttgtttatggactgcgcagctcaatcttcgactggagagg--aaaaacaaaaca
caagtatgacacgcacatacgtgttttctaggactgtgacctgtcggctgcctcgcggtg
gttggcatgatgccccaagt--gggggggggaaacaatcagtatgattgataaaacca
gtgggg-aaaaaggcgcgctgtcgcacacccttggtcacggcctc---aagtcgacggcc
ctcactggccca-ccctgcgactcctgttcgctattgacaagacacacctgcaataccgg
aggaaatgcctgagcagctgtgccgcagcaactgcggacgcgtagcgatatgcatcccta
ttgtttgcgacttggtacaaagaacgtgcgccgggagacggattgccctttgacttttcc
acccagcccgcggctctatacccttgacgtacctatttacctggttgatcctgccagtag
tcatatgcttgtctcaaagactaagccatgcatgtctaagtataaacacttttatactcg
tgaaactgcgaatggctcattaaaacagttatagttcctttgggaacaacactacttgga
taaccgtagtaattctagagctaatacatgcctcaacgcccgactcacgaagggtggtat
ttattggataaaaaaccatcgtgctc--tccagagcgctttgagatgattcacaataact
tgtcggatcgcatggcctcgtgctggcgactcctcattcaaatttctgccctatcaactt
tcgatggtagagtattggtctaccatggtgtcgacgggtgacggggaattagggttcgat
tccggagagggagcctgagaaacggctaccacatccaaggaaggcagcaggcgcgcaaat
tacccaatcccgactcggggaggtagtgacaaaaaataacaatagggggccctttgggtc
ttctaattggaatgagaacaatttaaatcccttatcgaggatccattggagggcaagtct
ggtgaacgta----------------------------------tgcatca---------
--aagccctccgtaat-aaaaaaaaacagcgccagatgtct-------------------

------------------------------------------------------------
------------------------------------------------------------
------------------------------------------------------------
------------------------------------------------------------
------------------------------------------------------------
-------ctctctggaagcacacca-----------------------------------
---cgcacggggttcgttcctctc-----------------------------tgcatgg
ccgtg-----------------------------------------tgaaaaaggcgagt
aggcaaacgatgggaaacccggcttggaatagtaaccagccacagagcccccatcataca
cttctaaattgccggggagtgcccgcgaggccttgactaccgcggtccgccgagaggtga
gagcgagcaatcgctcaggacgcagcagtgtacgtagtggtacatggatggtaaaaacgt
caaggatagggttgatccgcagggaagcgcgagggccaacctgcctttgggcaacagttg
gcaaaggcgaaccttcagagactctaatggagtgggcgcaagcttaagggagagtccaat
ccactgggaaaccaggtccacagcagtaacggggtgatgctcgccccagaggactgtgga
aggcaa--ccacc----aaagcggtgtgtgttgtcgggtagc-aatggccagcagccgcg
gtaattccagctccaatagcgtatattaaagttgttgcagttaaaacgctcgtagtcgga
cctcggagcgcccccacgggccgggcttttgccttgttgccagtgggagtttgtctcctt
ttgtcgggacgcggtgtgcagggcctttactgtctctgcgcgtttacctgcgggccgacc
gtttactgtgaagaaagtaaagtgttcaaagcaggccaatgccttgaatatgtgagcatg
gaataatagaataggacttgggctctattttgttggtttccagtgaccaagtaatgatta
atagggacggttgggggcattcgtatttcattgtcagaggtgaaattcttggattgatgg
aagacgtacaactgcgaaagcatctgccatggatgttttcattgatcaagaacgaaagtt
aggggatcgaagacgatcagataccgtcgtagtcttaaccataaacgatgccgactgggg
attggcgggggcacttttatatgactctgtcagcaccctgagggaaaccaaagtctttgg
gttctgggggggagtatggtcgcaaggctgaaacttaaaggaattgacggaagggcaccac
caggagtggagcctgcggcttaatttgactcaacacgggaaaacttaccaggtccggaca
gaagaatgattgacagactgaaagctctttcttgatttttttggttggtggtgcatggccg
ttcttagttggtggagtgatttgtctggttaattccgttaacgaacgagacctcgtcctg
ctaaataggtgcgcgcatgccagcactgcgttcttaccttcttagagggactatgcgcgt
ctagcgtatggaagattgaggcaataacaggtctgtgatgcccttagatgttctgggccg
cacgcgcgctacactgatgcattcaacgagtttcacttgatagtcctgggtcggaaggct
cgggtaatcttttgaaagtgcatcgtgctggggatagatcattgcaattattgatcttca
acgaggaattccttgtaggcgcaggtcatcagcctgcgccgaatacgtccctgccctttg
tacacaccgcccgtcgctcctaccgattgaatggtccgatgaaatgtccggatcgcggct
agtgtgatggactcgtccagagcggtagccgtgagaagctcattaaatcttatcatttag
aggaaggagaagtcgtaacaaggt------------------------------------

------------------------------------------------------------
------------------------------------------------------------
------------------------------------------------------------
------------------------------------------------------------
------------------------------------------------------------
------------------------------------------------------------
------------------------------------------------------------
------------------------------------------------------------
------------------------------------------------------------
------------------------------------------------------------
------------------------------------------------------------
------------------------------------------------------------
------------------------------------------------------------
------------------------------------------------------------
------------------------------------------------------------
-------------------------------ttccgtaggtgaacctgcggaaggatcat
tcacaaactattgacacaaacacacgcgaaccaaatcgtccgcacaggtggtggcaatga
aagagagaatctgcatgtcgcctttcggggtatagcaagcagcactcttttgccatcgcc
tctgtgccgggcgtaaattctcattgagaggatgtgagggcaccacaggaagcttttcca
caggaagtcgccatccttctccctccacggcggcg----cttctgtgcttggcag-----
--tttttttttttgccttccagggaggatgccgccaatggagccccatataatatataca
tcatcatatgccc--ctttttttcttaaccgcttgccacagcttcttctatgaggagctt
gtgggaagactgtctccatacaataac-aaagatacaactcttagcggtggatatcttgg
ttctcgcaacgatgaagaacgcagccaactgcgataactaatgtgaattgcaggacttcg
tgaatcattaagtctttgaacgcaagttgcgctcatctccatgaggaggtgagcatgcct
gtttgagtgtcgatgggaaaaacaatac-ataacaacaatcggctctcttatcttcctgc
cttctt-ggggcaaaaggtagtgcatgccggagatgggctttgtcgcccctttctcttga
aagaggcgtcgtccgaaacacgtatcgttgcccgctggaggtcatagagcatgagttt--
aaagcaccacatctctcaagtcagagtgag-cgtcttgagtaggtcc-gcattgattcag
tttcaacccaggtggcgttggttacttgccat-aagcctctctggggtttctttgcttgg
acctctcttgtcagtttctctgacgtagagacaggt--gccgtctcatgggcgttgccct
ctggaacgtgcta-------aaaaaaaga----------aaaaaccgtaccacatgtcaa
aagaggcagattgtgcttgacggcgggagatcttgcatttcttgcttgtcctgagcgtct
gccagtctctg-acga--cgagtg----tgttccttccataaacccgatctcaaatcagg
taggaccacccgctgaatttaagcatataattaagcggaggaaaagaaactaacaaggat
tccctcagtagcggcgagcgaagcgggaagagctcaacatgaaaaccgatgggggttcgc

gctttccttcgggttgtatagtaaagtggtgtcgcctgcgtcgcgccgggtataaatccc
gtctgaaatggggtgtcgtagagggttataaccccgtctgtgacccggtgtgcgatgcgt
tacggtgaccttacgatgagtcgggttgcttgggcatgcagccctaagtgggtggtaaac
ttcatctaaagctaaatactagcgagagaccgatagcgtacaagtaccgtaagggaaaga
tgcaaagaactttgaaaagagagttaaacagggcttgaaaccgttaagagtgaagcgaag
gaggtcagtgatcatggggccagatccccaagtgcctgtcata-gggcaccgtagcgttc
tgg-ttcccggggcagcgccggccgggggcgggtgtcaacggggaggtggttgagggatt
tatttcccgcctgtactcctttctagcttgcaccctggccgagggatccgcgacttgtcg
tgtgagggcgctgtcaaaatggccttctccaccccgtcttgaaacacggaccaaggagtc
taacatactcgcgagtgtttgggtggttaaaacccgaacgcacaatgaaagtaaacggtg
gctctgcgccgctgcttggtgggaaccctgctcttataatgggggcaccatcacccgac
cccgatcccttcggggtggttggttttgaggcagagcgagtatgatgggacccgaaagat
ggtgaactattcctgagtagggcgaagccagaggaaactctggtggaggctcgtagcgat
gctgacgtgcaaatcgctcgtcgaacttgggtataggggcgaaagactaatcgaaccatc
tagtagctggttccctccgaagtttccctcaggatagcgggagcaaatgcaagtcagtat
catccggtaaagcgaatgattagaggcctcgggggagtaactccttcgacctattctcaa
actttaaataggtgaggagccggcaacgttgcttaacacaatgaacgccgtgcggacaat
gattgctctcagtgggccgattttggtaagcagaactggcgatgcgggatgaaccgaaag
cgacgctaaggtgccaaactcccagctcattcagacaccagaaaaggtgttgattcatac
agacagcaggacggtggccatggaagtcggaatccgctaaggagtgtgtaacaactcacc
tgccgaatggatcagccctgaaaatggatggcgcttaagcgggggaccgaggcgtcgccg
ttggggtaaatacgaagccccaatgagtaggagggcgtgggggttttgaaacacggtaca
ctcgcatggaagccttgggcgtaagctcgggtggagcgacccctagtgtagatcttggtg
gtagtagcaaatattcaagtgagaaccttgaagactgaagtggggaagggttccacgcga
acagtgcttggacgtgggttagtcggtcctaagagatggggaaactccgtattcaagtgc
gcttttgtagcgcagtccatcgaaagggaagcgggttaacattcccgcaccaggatgtgg
aaactgtggtttaccgcaaactgcggcaacgcaactgaacttggggacgcccgtaggagc
tccgagcagggatctctttcctgtttaacgcgcccaaccccctggaatcggtttgtccgg
agatagggggcgaaaggccggaaaagcaccgcacgtcttgtggtgttcggaacgcttgta
atggcccttgaaaacccaggggaaagcagccgtcttacac--ttgtgtgagacgaggaat
tctcacacctggccgtacccataaccgcatcaggtccccaaggtgaacagcctctagtca
atagaataatgtagataagggaagtcggcaaaatagatccgtaactttgggaaaaggatt
ggctctaatgttcgggcttgtcgggacgcgcgggcgtagacgccaatgagcggcagacgt
ctagcctctttggaggctggattgacgctgcttgtgggaagaaaccgtgtcgatggaggc
aggagcgcttatcgttcttgtctcttcccgcttgcgatcaacggacggattagaactggt
acggacaaggggaatccgactgtttaattaaaacaaagcattgcgacggccggaaacggt

gttggcgcaatgtgatttctgcccagtgctctgaatgtcaaagtgaagaaattcaaccaa
gcgcgggtaaacggcgggagtaactatgactct--taaac-aaaaaatactaccgac---
-------------------gggagtcaaaagtgggggcggcgtctggtgtagaaatgcac
gtat--------------------------------------------------------
-------------------gacacgct-tttttcctgctagtcgatctgacctttgttct
ttggactgtgg-------------------------------------------------
--------------------------gcggtcagcggcaacataacc-------------
------------------------------------------------------------
------------------------------------------------------------
----------------------------tggtacggg---------------------ga
aggcctcgacgttga---------------------------------------------
--------------------------------gtgctgcatcttggaagtatc-------
-----------------------------ttgacacaataagaga---------------
------------------------------aaagagattgcgcttgatagg---------
----gggttgatctcgt-------------------------------------------
-------------------------------------------ggcgagtcgacgc----
---------------------------------cggtgtgaactggt-------------
------------------------------------------------------------
-----------------------ggcaat-------------------------------
----ggccgtcgtagagcacgcaaaggtatgggctgactttttctcctca----aaagtt
ggcttaagggac-gtgccatcccacacgataataaagtgtgcagtctg------------
--------------------taagagctcccacagggcaaagac---------agactgg
gatgcagacggagacggcaaa---agcaacatccgttctgcttctgggcgcaaatgctta
aggtagccaaatgcctcgtcatctaaactaattgg-at----------------------
-----------------------gagatagtc----------tgagtgaaaagcttatgc
ag-----------------gctagtaccggaaaaaaaaaaaaaaaagaggggctctgcca
ct----ctgac------------------------ggtgcgacactccttgtacagggaa
accctcc--gcatggaaaggcacgtggtaccaccctcgcggggaaactcgcgagcgggat
cccgggtaatgaccggagccgatggtaaaaactcacgtgaccgatggggtgaccctgtgg
cgagacgctaagttgtgggacatcaccctctcacaatacgcatccgtcgcagagactgaa
cgggagtgggctcgggtcgtcga---------tggctaaccagccttcgtcgt-------
---ggggcttaagatacagtccgggcctcagggatggaaaaaaaaacctggggaacta-c
cagaattagtgacgcgcatgaatggattaacgagattcccactgtccctatctactatct
agcgaaaccacagccaagggaacgggcttggaaaaatcagcggggaaagaagaccctgtt
gagcttgactctagtccgactttgtgagatgagttgggaggtgtagtctaagagggaggg
gcaactcgaccgtgaaatacctctactcccaagcttgttttacttactccggcgagcagg

gatgggtggtaacaccctttttctaggattaacacctcttcgggggtggacctgcgcgga
ggacattgtcaggtggggagtttggctggggcggcacatctgttaaaagataacgcaggt
gtcctaagatgagctcagtgagaacagaaatctcacgtagaacaaaagggtaaaagctca
tttgattttgattttcagtacgaatacaaactgtgaaagcatggcctatcgatcctttg
accttcaaggtttgaagcaagaggtgtcagaaaagttaacacgcgttgtggtagctttta
agagtggactcaaagtgttccgctagtccccccctcggaccgccacttacaaaaggcctgg
ggatgggggcgacactgtcaaattgcggggactccctaaagcctgtctctaccaagccag
cgcggcgaaagcccctggtggccggggtaatgacctagggtatggtaaaagcgagacagg
atg-gaacaatgggcaatccgcagccaagtcctaagg----ctttttgaaaagcggctat
ggatgcagttcaacgactagatggcagtgggctgagagagcccaccgcggcaccctttcg
gcttaagatatagtcttgcccacgtgaaagcgtgcctgcagatgaacgccaccatgtatg
gccggagctgcaggaggccgctaaacataccagcggctgggaacataacagtaccacagg
gatgaac-----------------------------------------------------
------------------------------------------------------------
------------------------------------------------------------
------------------------------------------------------------
------------------------------------------------------------
------------------------------------------------------------
------------------------------------------------------------
------------------------------------------------------------
------------------------------------------------------------
------------------------------------------------------------
------------------------------------------------------------
------------------------------------------------------------
------------------------------------------------------------
------------------------------------------------------------
------------------tatcgtcccaggcgggtgtccctctctcacccgtctagtccg
gccttggtgcgcaaaga---cttgcgtggatgcaccttggttaaggaaaatgctcctagt
agttttataagggcaattcttgtagggcaacacatctggatgcgaagaactcccgg-aaa
gacaacggtaccaagtgcttgggttagcaccccattcatgaaggcaaataatc-gaaaga
ggtgccgggtaataatccgttgtatagggacaatccgcagctgacttggctccctg--cc
cat--ccatgggcacagcttagcggttcaacgctcggtaagatgtgggttggcgcgctgt
ccgacagcgcactggcttaaggtacgggctactcccaccggagacggtgtctttgtctac
ctcgggtaacacgcctcgggagaagcaaagagggtgcaatgg--aaaaaactggttgcgg
ggaaatccctaccagagtcacccggtacaaaggaactggcttgtggcagccaagcgttca
tagcgacgttgctttttgatccttcgatgtcggctcttcctatcattgcgaagcagaatt

cgccaagtgttggattgttcacccactaatagggaacgtgagctgggttttttttttat-
-----tgcccaaaagcttgaagctgggtcctgtgtgtgacgggttgcatttctcacggct
c-cccgtccgtcgttt-aata------ggcgccacagtcctagatggatcttctggctcg
actagtagaggatagatagagatcgtcaaattgcgggaaagccctaagagcccggagcta
ccaagtgcgtgtcgaaaggcatagacccggccagtgatggccggcgcatggcttgatttc
catcaag--aggtaaaaaggctctgggattgggtaatccgcagccaagcccctaaggtag
aacatccgtttctgctatggggaaggttcaacgactgtaaggcgtaccgtgcaagcggaa
gagacagtctaggcccactggaaagcagtgccaaagtaagtaggcaggtcgtggttctat
cctc
>Form_B_Japan
gcatgttttggtatgtgtttcgacagtcatgcaagtcatagccgtaggatggggagtgag
cttcatagcccgaccccttctcaggagcatgttttcatagcatattgagcatactggggt
ggtctcccatgtagctcgctctctcggtgcaggacctcccttttgcgggccacttcgatg
gcctccaataggga-gagcatgttttggcatatgaagcgttggtggtgaggctcgtagcc
tgtatatacataccaacgcagtcgtcatatgtctctctggcgggggtcggtcgtatgacc
ttgtccctacctggagcatgtatttcgtatgat---atactggac-ggcagctgcgctgt
tcagtcgtcagccgtacctcatgtgcgcgtttccaggtccgaacgtttggcatgcgaggg
catcataccagaagcctgttttctcgtatgagcatgtaggtaggcaggctcttcagtcgt
ctacccataaatgcacttcgaagga-actttttggtattgagtggctttgtgcccagacg
ttcttgtgagcttgttctcgctctacgatacgctggatgctgtggtattacctacagcag
acgcattcctagggcgggcggcctctctgcagatgagcacatctgtagaaagaagctccc
cttgatgacgcaatgcaatggctgggtaaacggagtgctatacagaagggtcaaacctc-
tgtggaagtacacccataccagctttgtttgttgagaggggtcggctctacgagtgaggc
cctatactgccatctcattcagatgcgtagacggtcttatctcagactgcctttctggct
gattgtcctcggtccctcgtgcacacgcgtcgctggctgaatgggcttgtccttatggag
tgccgcgttcttc-aaagaatccgtcttcatacagcgacctcgtttacatattgccggct
gaatgggcttgtccttatggagcactgcattcttctaggaattggtatctatacagcgac
ctcatttacatacaaggcacctgttttctacatagacttgccatttggctgattgttctg
acagtcgctgcctcttcggagttctgtgtggacgagtggcc--ttctgagtgggccggtc
cttacggagtccccatatctgttatgactgggttctccatacagcgacctcactcagaat
tgtatgg--------------caccgtgccgat-----------tgaatggacaagcagt
ccttagggaatcgcatgaaagcatgacggc-acatggtgtatggacccgttgggccttgc
ctgtgctatcttctcgagtcgaccgg------------tcctcgtctcctcctttcgcga
cgggaagctcttgagtttaccgtgggtcaggacgctgtggtatctgtttgtctgtagtgt
gggctctctatcagaacagcgtatcagcacacgtcgtctgggtctatggtatttttgctt
aacggccctgggtttggtagtatggttccatgtcggtcctctcacctcccatccgtcatg

tctctga-----------------------------------------------------

------------------------------------------------------------

--------------------------------------------tatcatctgcacagta

cgtcttgtcagaatatcattctgtatggcagtcgatagactaggccctatgccgcagctg

tctctgaccactatgcgagcgcatacctacctcaatgtgttcatagctgttatggatctt

gggtgtggtgtgccagaacgatacgccaccagattgtggaatggtgcttgtagtattaga

agaggacatgtatgaagagcacgttcttttctcttatgatggtgccgcgggtgggggccg

ttctcattctgcttgtcgggtgctcaaggtgcatgtaggattgctgaacccac-tgaatg

gtcatgagaatggtcatatggtgaggttcacttctccatcgctttgtgtgtgttggatgt

ttgggtggcttcggccgcatggaaaatctcggtctgtaactcatcagggagggtttccat

gtgagcaatctctccagtcgaacacacacccatgggcgtcccgctaggcaaaatggaact

ctcgatccgacacagcctgtggagtgtctgttccgtgtggcgtgaaacggcttatactgg

atgtcttccc-tttggacaccagtctgttggtatg--tacgcataccatctagagggagc

aagaaaaaggcatgcaagtgccacaaaggaagggaggccttcctgctaccgtgtgtcc-t

cttgacgatcacgggggagtttatagagacatttggacgagatcctttttttcacgaggcg

gtgccctcgcgcaaaggtgttgctctgcacgcatctttatgatgatagttgctcgctcct

atgcgtccactcgtgaaaatacctgctctctctgtctgtttccatgacacccc-catgcg

ttgtcatagagcacgctcaggtacagaaagtgctgtgactttcttggcctgctgtcactg

agtcagcctgatactttggaggaatcttttctgggcgctgcccagctgagagttcttatc

aggctctggagtgtgcatatggctataatcataatagtgcatgtggccacgcagaaaaag

catcgacgtggaatggagaagaacaccattatgaatggaacggttcttactctgcttccg

ttgcttgacgctgtatgggaatcgctttgccctgggggtacaccttccattccataacaa

agcatttgatctatgccccga-gggtcattctctatgaagagcccgaaacttgtctgagt

gagaggcgaaggtagcccttcattt--atgcggggtccgccctcacactgtagtcggcac

agtcccttgtttatggactgcgcagctcaatcttcgactggagagg--aaaaacaaaaca

caagtatgacacgcacatacgtgttttctaggactgtgacctgtcggctgcctcgcggtg

gttggcatgatgccccaagt--gggggggggaaacaatcagtatgattgataaaaacca

gtgggg-aaaaaggcgcgctgtcgcacacccttggtcacggcctc---aagtcgacggcc

ctcactggccca-ccctgcgactcctgttcgctattgacaagacacacctgcaataccgg

aggaaatgcctgagcagctgtgccgcagcaactgcggacgcgtagcgatatgcatcccta

ttgtttgcgacttggtacaaagaacgtgcgccgggagacggattgccctttgacttttcc

acccagcccgcggctctatacccttgacgtacctatttacctggttgatcctgccagtag

tcatatgcttgtctcaaagactaagccatgcatgtctaagtataaacacttttatactcg

tgaaactgcgaatggctcattaaaacagttatagttcctttgggaacaacactacttgga

taaccgtagtaattctagagctaatacatgcctcaacgcccgactcacgaagggtggtat

ttattggataaaaaaccatcgtgctc--tccagagcgctttgagatgattcacaataact tgtcggatcgcatggcctcgtgctggcgactcctcattcaaatttctgccctatcaactt

tcgatggtagagtattggtctaccatggtgtcgacgggtgacggggaattagggttcgat

tccggagagggagcctgagaaacggctaccacatccaaggaaggcagcaggcgcgcaaat

tacccaatcccgactcggggaggtagtgacaaaaataacaatagggggccctttgggtc

ttctaattggaatgagaacaatttaaatcccttatcgaggatccattggagggcaagtct

ggtgaacgta----------------------------------tgcatca---------

--aagccctccgtaat-aaaaaaaaacagcgccagatgtct-------------------

------------------------------------------------------------

------------------------------------------------------------

------------------------------------------------------------

------------------------------------------------------------

------------------------------------------------------------

-------ctctctggaagcacacca-----------------------------------

---cgcacggggttcgttcctctc-----------------------------tgcatgg

ccgtg-----------------------------------------tgaaaaaggcgagt

aggcaaacgatgggaaacccggcttggaatagtaaccagccacagagcccccatcataca

cttctaaattgccggggagtgcccgcgaggccttgactaccgcggtccgccgagaggtga

gagcgagcaatcgctcaggacgcagcagtgtacgtagtggtacatggatggtaaaaacgt

caaggatagggttgatccgcagggaagcgcgagggccaacctgcctttgggcaacagttg

gcaaaggcgaaccttcagagactctaatggagtgggcgcaagcttaagggagagtccaat

ccactgggaaaccaggtccacagcagtaacggggtgatgctcgccccagaggactgtgga

aggcaa--ccacc----aaagcggtgtgtgttgtcgggtagc-aatggccagcagccgcg

gtaattccagctccaatagcgtatattaaagttgttgcagttaaaacgctcgtagtcgga

cctcggagcgcccccacgggccgggcttttgccttgttgccagtgggagtttgtctcctt

ttgtcgggacgcggtgtgcagggcctttactgtctctgcgcgtttacctgcgggccgacc

gtttactgtgaagaaagtaaagtgttcaaagcaggccaatgccttgaatatgtgagcatg

gaataatagaataggacttgggctctattttgttggtttccagtgaccaagtaatgatta

atagggacggttgggggcattcgtatttcattgtcagaggtgaaattcttggattgatgg

aagacgtacaactgcgaaagcatctgccatggatgttttcattgatcaagaacgaaagtt

aggggatcgaagacgatcagataccgtcgtagtcttaaccataaacgatgccgactgggg

attggcgggggcacttttatatgactctgtcagcaccctgagggaaaccaaagtctttgg

gttctgggggggagtatggtcgcaaggctgaaacttaaaggaattgacggaagggcaccac

caggagtggagcctgcggcttaatttgactcaacacgggaaaacttaccaggtccggaca

gaagaatgattgacagactgaaagctctttcttgatttttttggttggtggtgcatggccg

ttcttagttggtggagtgatttgtctggttaattccgttaacgaacgagacctcgtcctg

ctaaataggtgcgcgcatgccagcactgcgttcttaccttcttagagggactatgcgcgt

ctagcgtatggaagattgaggcaataacaggtctgtgatgcccttagatgttctgggccg
cacgcgcgctacactgatgcattcaacgagtttcacttgatagtcctgggtcggaaggct
cgggtaatcttttgaaagtgcatcgtgctggggatagatcattgcaattattgatcttca
acgaggaattccttgtaggtgcaggtcatcagcctgcgccgaatacgtccctgccctttg
tacacaccgcccgtcgctcctaccgattgaatggtccgatgaaatgtccggatcgcggct
agtgtgatggactcgtccagagcggtagccgtgagaagctcattaaatcttatcatttag
aggaaggagaagtcgtaacaaggtttccgtattttgtgggtggataccgattcaataaca
tccgccgtttttctcaatgtactatatccctagtctgcttgcctagttcttggagtcttg
tgcgatgtgtgctattacaggcatgccgggtcatgaagacagacaaggttgggatcttca
cggaagtacgcgcaataggagcgcgtcttgttcaacgcacctgactcaagagtcatatgc
gagggacggacgcagcgcttgtcgccgcacagatgggaagcttcctctccaggcatagga
tggcgctgcatttgcatcaccgctgctagtaagtggcaataatacttgactccccgccag
cgcacttgtacgtagccgttggcggcgggaacgtagttggtgatgtgacagccttgagcg
ttcacctctgtgtgtccgccaatctgcacaaagatggacgccatatcggagacaggggga
ggtgccattg---tgatgatgacgatgttgttgggtggtccttccgtaggaagacgtaca
cgaagcctttgcggcccgaaagggtgcacagtgcgcgactataaacactgattctgtggt
caacgcaagcagcggttctgtggaagaacagctgcgactttctcaaattgcgggaacacc
ctaaagtcaacaagcaccgcggcttgacgtgaaaagcgtcaggcgcagcagtcggggtag
cgccctggcggatggtaatagccttgttgaatggcacaatgggcaatccgcagccaagcc
cctaaggtcccccgctgggctatggggaaggttcacagactgtaagggagggggtttctt
ccatga--aagaagcttaagagacagtcggtcccctgcgagagcagtgtcgcatgtgagg
aaggcacccatgaaaagtgggtgctggagagccatgtgagccctctatgagggtgggaag
ttttccga---tacaatgagaaacaaacacgttccgtaggtgaacctgcggaaggatcat
tcacaaactattgacacaaacacacgcgaaccaaatcgtccgcacaggtggtggcaatga
aagagagaatctgcatgtcgcctttcggggtatagcaagcagcactcttttgccatcgcc
tctgtgccgggcgtaaattctcattgagaggatgtgagggcaccacaggaagcttttcca
caggaagtcgccatccttctccctccacggcggcg----cttctgtgcttggcag-----
--ttttttttttgccttccagggaggatgccgccaatggagccccatataatatataca
tcatcatatgccc--cttttttttcttaaccgcttgccacagcttcttctatgaggagctt
gtgggaagactgtctccatacaataac-aaagatacaactcttagcggtggatatcttgg
ttctcgcaacgatgaagaacgcagccaactgcgataactaatgtgaattgcaggacttcg
tgaatcattaagtctttgaacgcaagttgcgctcatctccatgaggaggtgagcatgcct
gtttgagtgtcgatgggaaaaacaatac-ataacaacaatcggctctcttatcttcctgc
cttctt-ggggcaaaaggtagtgcatgccggagatgggctttgtcgcccctttctcttga
aagaggcgtcgtccgaaacacgtatcgttgcccgctggaggtcatagagcatgagttt--
aaagcaccacatctctcaagtcagagtgag-cgtcttgagtaggtcc-gcattgattcag

tttcaacccaggtggcgttggttacttgccat-aagcctctctggggtttctttgcttgg
acctctcttgtcagtttctctgacgtagagacaggt--gccgtctcatgggcgttgccct
ctggaacgtgcta-------aaaaaaaga----------aaaaaccgtaccacatgtcaa
aagaggcagattgtgcttgacggcgggagatcttgcatttcttgcttgtcctgagcgtct
gccagtctctg-acga--cgagtg----tgttccttccataaacccgatctcaaatcagg
taggaccacccgctgaatttaagcatataattaagcggaggaaaagaaactaacaaggat
tccctcagtagcggcgagcgaagcgggaagagctcaacatgaaaaccgatgggggttcgc
gctttccttcgggttgtatagtaaagtggtgtcgcctgcgtcgcgccgggtataaatccc
gtctgaaatggggtgtcgtagagggttataaccccgtctgtgacccggtgtgcgatgcgt
tacggtgaccttacgatgagtcgggttgcttgggcatgcagccctaagtgggtggtaaac
ttcatctaaagctaaatactagcgagagaccgatagcgtacaagtaccgtaagggaaaga
tgcaaagaactttgaaaagagagttaaacagggcttgaaaccgttaagagtgaagcgaag
gaggtcagtgatcatggggccagatccccaagtgcctgtcata-gggcaccgtagcgttc
tgg-ttcccggggcagcgccggccgggggcgggtgtcaacggggaggtggttgagggatt
tatttcccgcctgtactcctttctagcttgcaccctggccgagggatccgcgacttgtcg
tgtgagggcgctgtcaaaatggccttctccaccccgtcttgaaacacggaccaaggagtc
taacatactcgcgagtgtttgggtggttaaaacccgaacgcacaatgaaagtaaacggtg
gctctgcgccgctgcttggtgggaaccctgctcttataatgggggcaccatcacccgac
cccgatcccttcggggtggttggttttgaggcagagcgagtatgatgggacccgaaagat
ggtgaactattcctgagtagggcgaagccagaggaaactctggtggaggctcgtagcgat
gctgacgtgcaaatcgctcgtcgaacttgggtataggggcgaaagactaatcgaaccatc
tagtagctggttccctccgaagtttccctcaggatagcgggagcaaatgcaagtcagtat
catccggtaaagcgaatgattagaggcctcgggggagtaactccttcgacctattctcaa
actttaaataggtgaggagccggcaacgttgcttaacacaatgaacgccgtgcggacaat
gattgctctcagtgggccgattttggtaagcagaactggcgatgcgggatgaaccgaaag
cgacgctaaggtgccaaactcccagctcattcagacaccagaaaaggtgttgattcatac
agacagcaggacggtggccatggaagtcggaatccgctaaggagtgtgtaacaactcacc
tgccgaatggatcagccctgaaaatggatggcgcttaagcgggggaccgaggcgtcgccg
ttggggtaaatacgaagccccaatgagtaggagggcgtgggggttttgaaacacggtaca
ctcgcatggaagccttgggcgtaagctcgggtggagcgacccctagtgtagatcttggtg
gtagtagcaaatattcaagtgagaaccttgaagactgaagtggggaagggttccacgcga
acagtgcttggacgtgggttagtcggtcctaagagatggggaaactccgtattcaagtgc
gcttttgtagcgcagtccatcgaaagggaagcgggttaacattcccgcaccaggatgtgg
aaactgtggtttaccgcaaactgcggcaacgcaactgaacttggggacgcccgtaggagc
tccgagcagggatctctttcctgtttaacgcgcccaacccccctggaatcggtttgtccgg
agatagggggcgaaaggccggaaaagcaccgcacgtcttgtggtgttcggaacgcttgta

atggcccttgaaaacccaggggaaagcagccgtcttacac--ttgtgtgagacgaggaat
tctcacacctggccgtacccataaccgcatcaggtccccaaggtgaacagcctctagtca
atagaataatgtagataagggaagtcggcaaaatagatccgtaactttgggaaaaggatt
ggctctaatgttcgggcttgtcgggacgcgcgggcgtagacgccaatgagcggcagacgt
ctagcctctttggaggctggattgacgctgcttgtgggaagaaaccgtgtcgatggaggc
aggagcgcttatcgttcttgtctcttcccgcttgcgatcaacggacggattagaactggt
acggacaaggggaatccgactgtttaattaaaacaaagcattgcgacggccggaaacggt
gttggcgcaatgtgatttctgcccagtgctctgaatgtcaaagtgaagaaattcaaccaa
gcgcgggtaaacggcgggagtaactatgactct--taaac-aaaaaatactaccgac---
-------------------gggagtcaaaagtgggggcggcgtctggtgtagaaatgcac
gtat--------------------------------------------------------
-------------------gacacgct-tttttcctgctagtcgatctgacctttgttct
ttggactgtgg-------------------------------------------------
--------------------------gcggtcagcggcaacataacc-------------
------------------------------------------------------------
------------------------------------------------------------
----------------------------tggtacggg---------------------ga
aggcctcgacgttga---------------------------------------------
--------------------------------gtgctgcatcttggaagtatc-------
-----------------------------ttgacacaataagaga---------------
------------------------------aaagagattgcgcttgatagg---------
----gggttgatctcgt-------------------------------------------
-------------------------------------------ggcgagtcgacgc----
---------------------------------cggtgtgaactggt-------------
------------------------------------------------------------
-----------------------ggcaat-------------------------------
----ggccgtcgtagagcacgcaaaggtatgggctgactttttctcctca----aaagtt
ggcttaagggac-gtgccatcccacacgataataaagtgtgcagtctg------------
--------------------taagagctcccacagggcaaagac---------agactgg
gatgcagacggagacggcaaa---agcaacatccgttctgcttctgggcgcaaatgctta
aggtagccaaatgcctcgtcatctaaactaattgg-at----------------------
-----------------------gagatagtc----------tgagtgaaaagcttatgc
ag-----------------gctagtaccggaaaaaaaaaaaaaaaagaggggctctgcca
ct----ctgac------------------------ggtgcgacactccttgtacagggaa
accctcc--gcatggaaaggcacgtggtaccaccctcgcggggaaactcgcgagcgggat
cccgggtaatgaccggagccgatggtaaaaactcacgtgaccgatggggtgaccctgtgg

cgagacgctaagttgtgggacatcaccctctcacaatacgcatccgtcgcagagactgaa
cgggagtgggctcgggtcgtcga---------tggctaaccagccttcgtcgt-------
---ggggcttaagatacagtccgggcctcagggatggaaaaaaaaacctggggaacta-c
cagaattagtgacgcgcatgaatggattaacgagattcccactgtccctatctactatct
agcgaaaccacagccaagggaacgggcttggaaaaatcagcggggaaagaagaccctgtt
gagcttgactctagtccgactttgtgagatgagttgggaggtgtagtctaagagggaggg
gcaactcgaccgtgaaatacctctactcccaagcttgttttacttactccggcgagcagg
gatgggtggtaacacccttttttctaggattaacacctcttcgggggtggacctgcgcgga
ggacattgtcaggtggggagtttggctggggcggcacatctgttaaaagataacgcaggt
gtcctaagatgagctcagtgagaacagaaatctcacgtagaacaaaagggtaaaagctca
tttgattttgattttcagtacgaatacaaactgtgaaagcatggcctatcgatccttttg
accttcaaggtttgaagcaagaggtgtcagaaaagttaacacgcgttgtggtagctttta
agagtggactcaaagtgttccgctagtccccccctcggaccgccacttacaaaaggcctgg
ggatgggggcgacactgtcaaattgcggggactccctaaagcctgtctctaccaagccag
cgcggcgaaagcccctggtggccggggtaatgacctagggtatggtaaaagcgagacagg
atg-gaacaatgggcaatccgcagccaagtcctaagg----ctttttgaaaagcggctat
ggatgcagttcaacgactagatggcagtgggctgagagagcccaccgcggcaccctttcg
gcttaagatatagtcttgcccacgtgaaagcgtgcctgcagatgaacgccaccatgtatg
gccggagctgcaggaggccgctaaacataccagcggctgggaacataacagtaccacagg
gatgaac-----------------------------------------------------
------------------------------------------------------------
------------------------------------------------------------
------------------------------------------------------------
------------------------------------------------------------
------------------------------------------------------------
------------------------------------------------------------
------------------------------------------------------------
------------------------------------------------------------
------------------------------------------------------------
------------------------------------------------------------
------------------------------------------------------------
------------------------------------------------------------
------------------------------------------------------------
------------------tatcgtcccaggcgggtgtccctctctcacccgtctagtccg
gccttggtgcgcaaaga---cttgcgtggatgcaccttggttaaggaaaatgctcctagt
agttttataagggcaattcttgtagggcaacacatctggatgcgaagaactcccgg-aaa

gacaacggtaccaagtgcttgggttagcaccccattcatgaaggcaaataatc-gaaaga
ggtgccgggtaataatccgttgtatagggacaatccgcagctgacttggctccctg--cc
cat--ccatgggcacagcttagcggttcaacgctcggtaagatgtgggttggcgcgctgt
ccgacagcgcactggcttaaggtacgggctactcccaccggagacggtgtctttgtctac
ctcgggtaacacgcctcgggagaagcaaagagggtgcaatgg--aaaaaactggttgcgg
ggaaatccctaccagagtcacccggtacaaaggaactggcttgtggcagccaagcgttca
tagcgacgttgctttttgatccttcgatgtcggctcttcctatcattgcgaagcagaatt
cgccaagtgttggattgttcacccactaatagggaacgtgagctgggtttttttttat-
-----tgcccaaaagcttgaagctgggtcctgtgtgtgacgggttgcatttctcacggct
c-cccgtccgtcgttt-aata------ggcgccacagtcctagatggatcttctggctcg
actagtagaggatagatagagatcgtcaaattgcgggaaagccctaagagcccggagcta
ccaagtgcgtgtcgaaaggcatagacccggccagtgatggccggcgcatggcttgatttc
catcaag--aggtaaaaaggctctgggattgggtaatccgcagccaagcccctaaggtag
aacatccgtttctgctatggggaaggttcaacgactgtaaggcgtaccgtgcaagcggaa
gagacagtctaggcccactggaaagcagtgccaaagtaagtaggcaggtcgtggttctat
cctc
>Form_L_China
gcatgttttggtatgtgtttcgacagtcatgcatgtcatagccgtaggatggggagtgag
cttcatagcccgaccccttctcaggagcatgttttcatagcatattgagcatactggggt
ggtctcccatgtagctcgctctctcggtgcaggacctcccttttgcgggccacttcgatg
gcctccattatgga-gagcatgttttggcatatgaagcgttggtggtgaggctcgtagcc
tgtatatacataccaacgcagtcgtcatatgtctctctggcgggggtcggtcgtatgacc
ttgtccctacctggagcatgtattttgtatgat---atactggac-ggcagctgcgctgt
tcagtcgtcagccgtacctcatgtgcgcgtttccaggtccgaacgtttgacatgcaaggg
catcataccagaagcctgttttctcgtatgagcatgtaggtaggcaggctcttcagtcgt
ctacccataaatgcacttcgaagga-actttttggtattgaatggctttgtgcccagacg
ttcttgtgagcttgttctcgctctgcgatacgctggaagctgtggtattacctacagcag
acgcattcctagggcggacggcctctctgcagatgggcacatctgtagaaagaggctccc
cttgatgacgcaatgcaatggctgggtaaacggagtgctatacagaagggtcaaacctc-
tgtggaagtacactcataccagcattgtttgttgagaggggtcggctctacgagtgaggc
cctatactgccatctcattcagatgcgtagacggtctttactcagactgcctttctggct
gattgtcctcggtccctcgtgcgcacgcgccgctggctgaatgggcttgtccttatggag
tgccgcgttcttc-aaagaatccgtcttcatacagcgacctcatttacatattgccggct
gaatgggcttgtccttatggagcactgcattcttctaggaattgctatctatacagcgac
ctcatttacatacaaggcacct-ttttctacatagacttgccatttggctgattgttctg
acagtcgctgcctcttcggagttctgtgtggacgagtggcc--tcctgagtgggccggtc

cttacggagtgcccaagtctgttatgactgggttctccatacagcgacctcactcagaat
tgtatgg--------------caccgtgccgat-----------tgaatggacaagcagt
ccttagggaatcgcatgaaagcatgacggcaacatggtgtatggacccgttgggccttgc
ctgtgctatcttctcgagtcgaccgg------------tcctcgtctcctcctttcgcga
cgggaagctcttgagtttaccgtgggtcaggacgctgtggtatctgtttgtctgtagtgt
gggctctctatcagaacagcgtatcagcacacgtcgtctgggtctatggtatttttgctt
aacggccctgggtttggtagtatggttccatgtcggtcctctcacctcccaaccgtcatg
tctctga-----------------------------------------------------
------------------------------------------------------------
--------------------------------------------tatcatctgcacagta
cgtcttgtcagaatatcattctgtatggcagtcgatagactaggccctatgccgcagctg
tctctgaccactatgcgagcgcatacctacctcaatgtgttcatagctgttatggatctt
gggtgtggtgtgccagaacgatacgccaccagattgtggaatggtgcttgtagtattaga
agaggacatgtatgaagagcacgttcttttctcttatgatggtgccgcgggtgcgggccg
ttctcattctgcttgtcgggtgctcaaggtgcatgtaggattgctgaacccac-tgaatg
gtcatgagaatggtcatatggtgaggttcacttctccatcgctttgtgtgtgttggatgt
ttgggtggcttcggccgcatggaaaatctcggtctgtaactcatcagggagggtttccat
gtgagcaatctctccagtcgaacacacacccatgggcgtcccgctaggcaaaatggaact
ctcgatccgacacagcctgtggagtgtttgttccgtgtggcgtgaaacggcttatactgg
atgtcttccc-tttggacaccagtctgttggtatg--tacgcataccatctagagggagc
aagaaaaaggcatgcaagtgccacaaaggaagggaggccttcctgctaccgtgtgtcc-t
cttgacgatcacgggggagtatatagagacatttggacgagatccgtttttcacgaggcg
gtgccctcgcgcaaaggtgttgctctgcacgcatctttatgatgatagttgctcgctcct
atgcgtccactcgtgaaaatacctgctctctctgtctgtttccatgacacccc-catgcg
ttgtcatagagcatgctcaggtacagaaagtgctgtgactttcttggcctgctgtcactg
agtcagcctgatactttggaggaatcttttctgggcactgcccagctgagagttcttatc
aggctctggagtgtgcatatgcctatattcattatagtgcatgtggctacgcagaaaaag
catcgacgtggaatggagaagaacaccattctgaatggaacggttcttactctacttctg
ttgcttgacgctgtatgggaatcgctttgctctgggggtacaccttccattccatgacaa
agcatttgatctatgccccga-gggtcattctctatgaagagcccgaaacttgtctgagt
gagaggcgaaggtagtccttcattt--atgcggggtccgccctcacactgtagtcggcac
agtcccttgtttatggactgcgcagctcaatcttcgactggagagg--aaaaacaaaaca
caagtatgacacgcacatacgtgttttctaggactgtgacctgtcggctgcctcgcggtg
gttggcatgatgccccaagtgggggggggagaaacaatcagtatgattgataaaaacca
gtgggg-aaaaaggcgcgctgtcgcacacccttggtcacggcctc---aagtcgacggcc
ctcactggccca-ccctgcgactcctgttcgcttttgacaagacacacctgcaata-cgg

aggaaatgcctgagcagctgtgccgcagcaactgcggacgcgtagcgatatgcatccctg
ttgtttgcgacttggtacaaagaacgtgcgccgggagacggattgccctttgacttttcc
acccagcccgcggctctatacccttgacgtacctatttacctggttgatcctgccagtag
tcatatgcttgtctcaaagactaagccatgcatgtctaagtataaacacttttatactcg
tgaaactgcgaatggctcattaaaacagttatagttcctttgggaacaacactacttgga
taaccgtagtaattctagagctaatacatgcctcaacgcccgactcacgaagggtggtat
ttattggataaaaaaccatcgtgctctgtccagagcgctttgagatgattcacaataact
tgtcggatcgcatggcctcgtgctggcgactcctcattcaaatttctgccctatcaactt
tcgatggtagagtattggtctaccatggtgtcgacgggtgacggggaattagggttcgat
tccggagagggagcctgagaaacggctaccacatccaaggaaggcagcaggcgcgcaaat
tacccaatcccgactcggggaggtagtgacaaaaaataacaataggggccctttgggtc
ttctaattggaatgagaacaatttaaatcccttatcgaggatccattggagggcaagtct
ggtgaacgta----------------------------------tgcatta---------
--aagccctccgtaat-aaaaaaaaacagcgccagatgtct-------------------
------------------------------------------------------------
------------------------------------------------------------
------------------------------------------------------------
------------------------------------------------------------
------------------------------------------------------------
-------ctctctggaagcacacca-----------------------------------
---tgcatggggtttgttcctctc-----------------------------tgcatgg
ccgtg-----------------------------------------tgaaaaaggcgagt
aggcaaacgatgggaaacccggcttggaatagtaaccagccacagagcccccatcataca
cttctaaattgccggggagtgcccgcgaggccttgactaccgcggtccgccgagaggtga
gagcgagcaatcgttcaggacgcagcagtgtacgtagtggtacatggatggtaaaaacgt
caaggatagggttgatccgcagggaagcgcgagtgccgccatg---------gtcggtgg
gcaaaggcgaaccttcagagactctaatggagtgggcgcaagcttaagggagagtccaat
ccactgggaaaccaggtccacagcagtaacggggtgatgctcgccccagaggactgtgga
aggcaaccccacc----aaaagggtgtgtgttgtcgggtagc-aatggccagcagccgcg
gtaattccagctccaatagcgtatattaaagttgttgcagttaaaacgctcgtagtcgga
cctcggagcgcccccacgggccgggcttttgccttgttgccagtgggagtttgtctcctt
ttgtcgggacgcggtgtgcagggcctttactgtctctgcgcgtttacctgcgggccgacc
gtttactgtgaagaaagtaaagtgttcaaagcaggccaatgccttgaatatgtgagcatg
gaataatagaataggacttgggctctattttgttggtttccagtgaccaagtaatgatta
atagggacggttgggggcattcgtatttcattgtcagaggtgaaattcttggattgatgg
aagacgtacaactgcgaaagcatctgccatggatgttttcattgatcaagaacgaaagtt

aggggatcgaagacgatcagataccgtcgtagtcttaaccataaacgatgccgactgggg
attggcgggggcacttttatatgactctgtcagcaccctgagggaaaccaaagtctttgg
gttctgggggggagtatggtcgcaaggctgaaacttaaaggaattgacggaagggcaccac
caggagtggagcctgcggcttaatttgactcaacacgggaaaacttaccaggtccggaca
gaagaatgattgacagactgaaagctctttcttgatttttttggttggtggtgcatggccg
ttcttagttggtggagtgatttgtctggttaattccgttaacgaacgagacctcgtcctg
ctaaataggtgcgcgcatgccagcactgcgttcttaccttcttagagggactatgcgcgt
ctagcgtatggaagattgaggcaataacaggtctgtgatgcccttagatgttctgggccg
cacgcgcgctacactgatgcattcaacgagtttcacttgatagtcctgggtcggaaggct
cgggtaatcttttgaaagtgcatcgtgctggggatagatcattgcaattattgatcttca
acgaggaattccttgtaggcgcaggtcatcagcctgcgccgaatacgtccctgccctttg
tacacaccgcccgtcgctcctaccgattgaatggtccgatgaaatgtccggatcgcggct
agtgtgatggactcgtccagagcggtagccgtgagaagctcattaaatcttatcatttag
aggaaggagaagtcgtaacaaggtttccgtattttgtggatgggcactcgtc--------
------------------------------------------------------------
------------------------------------------------------------
------------------------------------------------------------
---------gtggcgttctttgcgccacgcggat--------------------------
------------------------------------------------------------
------------------------------------------------------------
------------------------------------------------------------
-----------------------------------gtccccttccgtaggaaaacgtaca
cgaagcctttgcggcccgaaagggtgcacagtgcgcgactataaacactgattctgtggt
caacgcaagcagcggttctgtgacagaacagctgcgactttctcaaattgcgggaacacc
ctaaagtcaacaaacaccgcggccggacgtg-aaagcgtcaggcgcagcagtcggggtag
cgccctggcggatggtaagagctttgttgaatggcacaatgggcaatccgcagccaagcc
cctaaggtccttcgc--ggctatggggaaggttcacagactgtaagggagggggtttctt
ccatgaagaagaagcttaagagacagtcggtcccctgcgagagcagtgtctcatgtgagg
aaggtgcccatgaaaagtgggcgctggagagccatgggagccctctatgagggtggggag
ttttcctagataataatgagaaacaaacacgttccgtaggtgaacctgcggaaggatcat
tcacaaactattgacacaaacacacgcgaaccaaatcgtccgcacagcaggtggcaatga
aagagagactctgcatgtcgcctttcgggg--tagcaagcagcactcttctgccatcgcc
tctgtgcagggcgtaaattcccattgagaggatgtgagggcaccacaggaagcttttcca
caggaagtcgccatccttctccctccacggcggcg----cttctgtgcttggcag-----
-----ttttttttgccttctagggaggatgccgccaatggagccccttat-atatataca
tcatcatatg-cc--ctttttttcttaaccgcttgccacagcttcttctattaggagctt

gtgggaagactgtctccatacaataac-aaagatacaactcttagcggtggatatcttgg
ttctcgcaacgatgaagaacgcagccaactgcgataactaatgtgaattgcaggacttcg
tgaatcattaagtctttgaacgcaagttgcgctcatctccatgaggaggtgagcatgcct
gtttgagtgtcgatgggaaaaacaatac-ataacaacaatcggctctcttatcttcctgc
cttcttgggggcaaaaggtagtgcatgccggagatgggctttgtcgcccctttctcttga
aagaggcgtcgtccgaaacacgtatcgttgcccgctggaggtcatagagcatgagttt--
aaagcaccacatctctcaagtcagagtgag-cgtcttgagtaggtcc-gcattgattcag
tttcaacccaggtggcgttggttacttgccat-aagcctctctggggtttctttgcttgg
acctctcttgtcagtttctctgacgtagagacaggt--gccgtctcatggacgttgccct
ctggaacgtgct---aaaaaaaaaaaaga----------aaaaaccgtaccacatgtcaa
aagaggcagatcgtgcttgacggcgggaggtgttgcatttcttgcttgtcctgagcgtct
gccagtctct----ga--cgagtg----tgttccttccataaacccgatctcaaatcagg
taggaccacccgctgaatttaagcatataattaagcggaggaaaagaaactaacaaggat
tccctcagtagcggcgagcgaagcgggaagagctcaacatgaaaaccgttggggggttcgc
gctttccttcgggttgtatagtaaagtggtgtcgcctgcgtcgcgccgggtataaatccc
gtctgaaatggggtgtcgtagagggttatatccccgtctgtgacccggtgtgcgatgcgt
tacggtgaccttacgatgagtcgggttgcttgggcatgcagccctaagtgggtggtaaac
ttcatctaaagctaaatactagcgagagaccgatagcgtacaagtaccgtaagggaaaga
tgcaaagaactttgaaaagagagttaaacagggcttgaaaccgttaagagtgaagcgaag
gaggtcagtgatcatggggccagatccccaagtgcctgtcata-gggcaccgtagcgttc
tggtttcccggggcagcgccggccgggggcgggggttaacagggaggtggttgagggatt
cgttttccgcctgtactcctttatagcttgcaccccggccgagggatccgcgacttgtcg
tgtgagggcgctgtcaaaatggccttctccaccccgtcttgaaacacggaccaaggagtc
taacatactcgcgagtgtttgggtggttaaaacccgaacgcacaatgaaagtaaacggtg
gctctgcgccgctgcttggtgggaaccctgcttttataatggggggcactatcacccgac
cccgatccctttggggtggttggttttgaggtagagcgagtatgatgggacccgaaagat
ggtgaactattcctgagtagggcgaagccagaggaaactctggtggaggctcgtagcgat
gctgacgtgcaaatcgctcgtcgaacttgggtataggggcgaaagactaatcgaaccatc
tagtagctggttccctccgaagtttccctcaggatagcgggagcaaatgcaagtcagtat
catccggtaaagcgaatgattagaggcctcgggggagtaactccttcgacctattctcaa
actttaaataggtgaggagccggcaacgttgcttaaaacaatgaacgccgtgcggacaat
gattgctctcagtgggccgattttggtaagcagaactggcgatgcgggatgaaccgaaag
cgacgctaaggtgccaaactcccagctcattcagacaccagaaaaggtgttgattcatac
agacagcaggacggtggccatggaagtcggaatccgctaaggagtgtgtaacaactcacc
tgccgaatggatcagccctgaaaatggatggcgcttaagcgggggaccgaggcgtcgccg
ttggggtaaatacgaagccccaatgagtaggagggcgtgggggttttgaaacacggtaca

ctcgcatggaagccttgggcgtaagctcgggtggagcgacccctagtgtagatcttggtg

gtagtagcaaatattcaagtgagaaccttgaagactgaagtggggaagggttccacgcga

acagtgcttggacgtgggttagtcggtcctaagagatggggaaactccgtattcaagtgc

gcttttgtagcgcagtccatcgaaagggaagcgggttaacattcccgcaccaggatgtgg

aaactgtggtttaccgcaaactgcggcaacgcaactgaacctggggacgcccgtaggagc

tccgagcagggatctctttcctgtttaacgcgcccaacccccctggaatcggtttgtccgg

agatagggggcgaaaggccggaaaagcaccgcacgtcttgtggtgttcggaacgcttgta

atggcccttgaaaacccaggggaaagcagccgtcttacac--ttgtgtgagacgaggaat

tctcacacctggccgtacccatcaccgcatcaggtccccaaggtgaacagcctctagtca

atagaataatgtagataagggaagtcggcaaaatagatccgtaactttgggaaaaggatt

ggctctaatgttcgggcttgtcgggacgcgcgggcgtagatgccaatgagcggcagacgt

ctagcctctttggaggctggattaacgctgcttgtgggaagaaaccgtgtcgatggaggc

aggaacgcttatcgatcttgtctcttcccgcttgcgatcaacgaacggattagaactggt

acggacaaggggaatccgactgtttaattaaaacaaagcattgcgacggccggaaacggt

gttggcgcaatgtgatttctgcccagtgctctgaatgtcaaagtgaagaaattcaaccaa

gcgcgggtaaacggcgggagtaactatgactct--taaac-aaaccatactacctac---

-------------------gggagtcaaaagtgggggcggcgtctggtgtagaaatacac

gtac--------------------------------------------------------

-------------------gacacgctttttttcctgctagtcgatctgacctttgttct

ttggactgtgg-------------------------------------------------

--------------------------gcggtcagcggcaacataacc-------------

------------------------------------------------------------

------------------------------------------------------------

----------------------------tggtacggg---------------------ga

aggcctcgacgttga---------------------------------------------

--------------------------------gtgctgcatcttggaagtatc-------

-----------------------------ttgacacaataagaga---------------

------------------------------aaagagattgcgcttaatagg---------

----gggttgatctcgt-------------------------------------------

-------------------------------------------ggcgagccgacgc----

---------------------------------cggtgtgaactggt-------------

------------------------------------------------------------

-----------------------ggcaat-------------------------------

----ggccgtcgtagagcacgcaaaggtatgggctgactttttctcctca----aaagtt

ggcttaagggac-gtgccatcccacacgataataaagtgtgcagtctg------------

--------------------taagagctcccacagggcaaagac---------agactgg

gatgcagacggagacggcaaaaacaacaacatccgttctgcttctgggcgcaaatgctta
aggtagccaaatgcctcgtcatctaaactaattgg-at----------------------
-----------------------gagatagtc----------tgagtgaaaagcttatgc
ag-----------------gctagtaccgg----aaaaaaaaaaaagaggggctctgcca
ct----ctgac------------------------ggtgcgacactccttgtacagggaa
accctcc--gcatggaaaggcacgtggtaccaccctcgcggggaaactcgcgagcgggat
cccgggtaatgaccggagccgatggtaaaaactcacgtgaccgatggggtgaccctgtgg
cgagacgctaagttgtgggacatcaccctctcacaatacgcatccgtcgcagagactgaa
cgggagtgggctccgttga-----cggccggccggccagccagccgtcttcca-------
---ggggcttaagatacagtccgggcctcagggatgg----aaaaacctggggaacta-c
cagaattagtgacgcgcatgaatggattaacgagattcccactgtccctatctactatct
agcgaaaccacagccaagggaacgggcttggaagaatcagcggggaaagaagaccctgtt
gagcttgactctagtccgactttgtgagatgagttgggaggtgtagtctaagagggaggg
gcaactcgaccgtgaaatacctctactcccaagcttgttttacttactccggcgagcagg
gatgggtggtaacacccttttttctaggattaacacctcttcgggggtggacctgcgcgga
ggacattgtcaggtggggagtttggctggggcggcacatctgttaaaagataacgcaggt
gtcctaagatgagctcagtgagaacagaaatctcacgtagaacaaaagggtaaaagctca
tttgattttgattttcagtacgaatacaaactgtgaaagcatggcctatcgatccttttg
accttcaaggtttgaagcaagaggtgtcagaaaagt------------------------
------------------------------------------------------------
------------------------------------------------------------
------------------------------------------------------------
------------------------------------------------------------
------------------------------------------------------------
------------------------------------------------------------
---------------------------------------------------taccacagg
gat---------------------------------------------------------
------------------------------------------------------------
------------------------------------------------------------
------------------------------------------------------------
------------------------------------------------------------
------------------------------------------------------------
------------------------------------------------------------
------------------------------------------------------------
------------------------------------------------------------
------------------------------------------------------------

------------------------------------------------------------
------------------------------------------------------------
------------------------------------------------------------
------------------------------------------------------------
------------------------------------------------------------
------------------------------------------------------------
------------------------------------------------------------
------------------------------------------------------------
------------------------------------------------------------
------------------------------------------------------------
------------------------------------------------------------
------------------------------------------------------------
---------------------------------aactggcttgtggcagccaagcgttca
tagcgacgttgctttttgatccttcgatgtcggctcttcctatcattgcgaagcagaatt
cgccaagtgttggattgttcacccactaatagggaacgtgagctgggtagccactttta--
-----tgcccaacagctggatagctgggcttctgtcttа-ggccgggacttctcacgacc
cgtgccttggtcaccc-aaaaaagggcgaccgcagagctcaggatggagcttctggctta
actagtagaggacagatagagatcgtcaaattgcgggaaaccccttagagcctcgagcta
ccaagtgcgtgtcgaaaggcatagacgcagccagcgatggctgtcgcatggcttga-ttc
catcaagataggtaaaaatgctcgaggattgggcaatccgcagccaagcctctaaggt--
---ccccccgcgggctatggggaaggttcaacgactgtaaggcgtaccgtgcaagcggaa
gatacagtctagccccgctggaaagcagcgccaaagtaagtaggt-tgcagtgggtctgt
cctc
>Form_I_China_SRR9587942
gcatgttttggtatgtgtttcgacagtcatgcatgtcatagccgtaggatggggagtgag
cttcatagcccgaccccttctcaggagcatgttttcatagcatattgagcatactggggt
ggtctcccatgtagctcgctctctcggtgcaggacctcccttttgcgggccacttcgatg
gcctccattatgga-gagcatgttttggcatatgaagcgttggtggtgaggctcgtagcc
tgtatatacataccaacgcagtcgtcatatgtctctctggcgggggtcggtcgtatgacc
ttgtccctacctggagcatgtattttgtatgat---atactggac-ggcagctgcgctgt
tcagtcgtcagccgtacctcatgtgcgcgtttccaggtccgaacgtttgacatgcaaggg
catcataccagaagcctgttttctcgtatgagcatgtaggtaggcaggctcttcagtcgt
ctacccataaatgcacttcgaagga-actttttggtattgaatggctttgtgcccagacg
ttcttgtgagcttgttctcgctctgcgatacgctggaagctgtggtattacctacagcag
acgcattcctagggcggacggcctctctgcagatgggcacatctgtagaaagaggctccc
cttgatgacgcaatgcaatggctgggtaaacggagtgctatacagaagggtcaaacctc-

tgtggaagtacactcataccagcattgtttgttgagaggggtcggctctacgagtgaggc

cctatactgccatctcattcagatgcgtagacggtctttactcagactgcctttctggct

gattgtcctcggtccctcgtgcgcacgcgccgctggctgaatgggcttgtccttatggag

tgccgcgttcttc-aaagaatccgtcttcatacagcgacctcatttacatattgccggct

gaatgggcttgtccttatggagcactgcattcttctaggaattgctatctatacagcgac

ctcatttacatacaaggcacct-ttttctacatagacttgccatttggctgattgttctg

acagtcgctgcctcttcggagttctgtgtggacgagtggcc--tcctgagtgggccggtc

cttacggagtgcccaagtctgttatgactgggttctccatacagcgacctcactcagaat

tgtatgg--------------caccgtgccgat-----------tgaatggacaagcagt

ccttagggaatcgcatgaaagcatgacggcaacatggtgtatggacccgttgggccttgc

ctgtgctatcttctcgagtcgaccgg------------tcctcgtctcctcctttcgcga

cgggaagctcttgagtttaccgtgggtcaggacgctgtggtatctgtttgtctgtagtgt

gggctctctatcagaacagcgtatcagcacacgtcgtctgggtctatggtatttttgctt

aacggccctgggtttggtagtatggttccatgtcggtcctctcacctcccaaccgtcatg

tctctga-----------------------------------------------------

------------------------------------------------------------

--------------------------------------------tatcatctgcacagta

cgtcttgtcagaatatcattctgtatggcagtcgatagactaggccctatgccgcagctg

tctctgaccactatgcgagcgcatacctacctcaatgtgttcatagctgttatggatctt

gggtgtggtgtgccagaacgatacgccaccagattgtggaatggtgcttgtagtattaga

agaggacatgtatgaagagcacgttcttttctcttatgatggtgccgcgggtgcgggccg

ttctcattctgcttgtcgggtgctcaaggtgcatgtaggattgctgaacccac-tgaatg

gtcatgagaatggtcatatggtgaggttcacttctccatcgctttgtgtgtgttggatgt

ttgggtggcttcggccgcatggaaaatctcggtctgtaactcatcagggagggtttccat

gtgagcaatctctccagtcgaacacacacccatgggcgtcccgctaggcaaaatggaact

ctcgatccgacacagcctgtggagtgtttgttccgtgtggcgtgaaacggcttatactgg

atgtcttccc-tttggacaccagtctgttggtatg--tacgcataccatctagagggagc

aagaaaaaggcatgcaagtgccacaaaggaagggaggccttcctgctaccgtgtgtcc-t

cttgacgatcacgggggagtatatagagacatttggacgagatccgtttttcacgaggcg

gtgccctcgcgcaaaggtgttgctctgcacgcatctttatgatgatagttgctcgctcct

atgcgtccactcgtgaaaatacctgctctctctgtctgtttccatgacacccc-catgcg

ttgtcatagagcatgctcaggtacagaaagtgctgtgactttcttggcctgctgtcactg

agtcagcctgatactttggaggaatcttttctgggcactgcccagctgagagttcttatc

aggctctggagtgtgcatatgcctatattcattatagtgcatgtggctacgcagaaaaag

catcgacgtggaatggagaagaacaccattctgaatggaacggttcttactctacttctg

ttgcttgacgctgtatgggaatcgctttgctctgggggtacaccttccattccatgacaa

agcatttgatctatgccccga-gggtcattctctatgaagagcccgaaacttgtctgagt
gagaggcgaaggtagtccttcattt--atgcggggtccgccctcacactgtagtcggcac
agtcccttgtttatggactgcgcagctcaatcttcgactggagagg--aaaaacaaaaca
caagtatgacacgcacatacgtgttttctaggactgtgacctgtcggctgcctcgcggtg
gttggcatgatgccccaagtgggggggggagaaacaatcagtatgattgataaaaacca
gtgggg-aaaaaggcgcgctgtcgcacacccttggtcacggcctc---aagtcgacggcc
ctcactggccca-ccctgcgactcctgttcgcttttgacaagacacacctgcaata-cgg
aggaaatgcctgagcagctgtgccgcagcaactgcggacgcgtagcgatatgcatccctg
ttgtttgcgacttggtacaaagaacgtgcgccgggagacggattgccctttgacttttcc
acccagcccgcggctctatacccttgacgtacctatttacctggttgatcctgccagtag
tcatatgcttgtctcaaagactaagccatgcatgtctaagtataaacacttttatactcg
tgaaactgcgaatggctcattaaaacagttatagttcctttgggaacaacactacttgga
taaccgtagtaattctagagctaatacatgcctcaacgcccgactcacgaagggtggtat
ttattggataaaaaaccatcgtgctctgtccagagcgctttgagatgattcacaataact
tgtcggatcgcatggcctcgtgctggcgactcctcattcaaatttctgccctatcaactt
tcgatggtagagtattggtctaccatggtgtcgacgggtgacggggaattagggttcgat
tccggagagggagcctgagaaacggctaccacatccaaggaaggcagcaggcgcgcaaat
tacccaatcccgactcggggaggtagtgacaaaaaataacaataggggggcccctttgggtc
ttctaattggaatgagaacaatttaaatcccttatcgaggatccattggagggcaagtct
ggtgaacgta----------------------------------tgcatta---------
--aagccctccgtaat-aaaaaaaaacagcgccagatgtct-------------------
------------------------------------------------------------
------------------------------------------------------------
------------------------------------------------------------
------------------------------------------------------------
------------------------------------------------------------
-------ctctctggaagcacacca-----------------------------------
---tgcatggggtttgttcctctc-----------------------------tgcatgg
ccgtg-----------------------------------------tgaaaaaggcgagt
aggcaaacgatgggaaacccggcttggaatagtaaccagccacagagcccccatcataca
cttctaaattgccggggagtgcccgcgaggccttgactaccgcggtccgccgagaggtga
gagcgagcaatcgttcaggacgcagcagtgtacgtagtggtacatggatggtaaaaacgt
caaggatagggttgatccgcagggaagcgcgagtgccgccatg---------gtcggtgg
gcaaaggcgaaccttcagagactctaatggagtgggcgcaagcttaagggagagtccaat
ccactgggaaaccaggtccacagcagtaacggggtgatgctcgccccagaggactgtgga
aggcaaccccacc----aaaagggtgtgtgttgtcgggtagc-aatggccagcagccgcg

gtaattccagctccaatagcgtatattaaagttgttgcagttaaaacgctcgtagtcgga
cctcggagcgcccccacgggccgggcttttgccttgttgccagtgggagtttgtctcctt
ttgtcgggacgcggtgtgcagggcctttactgtctctgcgcgtttacctgcgggccgacc
gtttactgtgaagaaagtaaagtgttcaaagcaggccaatgccttgaatatgtgagcatg
gaataatagaataggacttgggctctattttgttggtttccagtgaccaagtaatgatta
atagggacggttgggggcattcgtatttcattgtcagaggtgaaattcttggattgatgg
aagacgtacaactgcgaaagcatctgccatggatgttttcattgatcaagaacgaaagtt
aggggatcgaagacgatcagataccgtcgtagtcttaaccataaacgatgccgactgggg
attggcgggggcacttttatatgactctgtcagcaccctgagggaaaccaaagtctttgg
gttctgggggggagtatggtcgcaaggctgaaacttaaaggaattgacggaagggcaccac
caggagtggagcctgcggcttaatttgactcaacacgggaaaacttaccaggtccggaca
gaagaatgattgacagactgaaagctctttcttgatttttttggttggtggtgcatggccg
ttcttagttggtggagtgatttgtctggttaattccgttaacgaacgagacctcgtcctg
ctaaataggtgcgcgcatgccagcactgcgttcttaccttcttagagggactatgcgcgt
ctagcgtatggaagattgaggcaataacaggtctgtgatgcccttagatgttctgggccg
cacgcgcgctacactgatgcattcaacgagtttcacttgatagtcctgggtcggaaggct
cgggtaatcttttgaaagtgcatcgtgctggggatagatcattgcaattattgatcttca
acgaggaattccttgtaggcgcaggtcatcagcctgcgccgaatacgtccctgccctttg
tacacaccgcccgtcgctcctaccgattgaatggtccgatgaaatgtccggatcgcggct
agtgtgatggactcgtccagagcggtagccgtgagaagctcattaaatcttatcatttag
aggaaggagaagtcgtaacaaggt------------------------------------

------------------------------------------------------------

------------------------------------------------------------

------------------------------------------------------------

------------------------------------------------------------

------------------------------------------------------------

------------------------------------------------------------

------------------------------------------------------------

------------------------------------------------------------

------------------------------------------------------------

------------------------------------------------------------

------------------------------------------------------------

------------------------------------------------------------

------------------------------------------------------------

------------------------------------------------------------

------------------------------------------------------------

-------------------------------ttccgtaggtgaacctgcggaaggatcat
tcacaaactattgacacaaacacacgcgaaccaaatcgtccgcacagcaggtggcaatga
aagagagactctgcatgtcgcctttcgggg--tagcaagcagcactcttctgccatcgcc
tctgtgcagggcgtaaattcccattgagaggatgtgagggcaccacaggaagcttttcca
caggaagtcgccatccttctccctccacggcggcg----cttctgtgcttggcag-----
-----ttttttttgccttctagggaggatgccgccaatggagccccttat-atatataca
tcatcatatgccc--ctttttttcttaaccgcttgccacagcttcttctatgaggagctt
gtgggaagactgtctccatacaataac-aaagatacaactcttagcggtggatatcttgg
ttctcgcaacgatgaagaacgcagccaactgcgataactaatgtgaattgcaggacttcg
tgaatcattaagtctttgaacgcaagttgcgctcatctccatgaggaggtgagcatgcct
gtttgagtgtcgatgggaaaaacaatac-ataacaacaatcggctctcttatcttcctgc
cttcttgggggcaaaaggtagtgcatgccggagatgggctttgtcgcccctttctcttga
aagaggcgtcgtccgaaacacgtatcgttgcccgctggaggtcatagagcatgagttt--
aaagcaccacatctctcaagtcagagtgag-cgtcttgagtaggtcc-gcattgattcag
tttcaacccaggtggcgttggttacttgccataaagcctctctggggtttctttgcttgg
acctctcttgtcagtttctctgacgtagagacaggt--gccgtctcatggacgttgccct
ctggaacgtgct---aaaaaaaaaaaaga----------aaaaaccgtaccacatgtcaa
aagaggcagatcgtgcttgacggcgggaggtgttgcatttcttgcttgtcctgagcgtct
gccagtctct----ga--cgagtg----tgttccttccataaacccgatctcaaatcagg
taggaccacccgctgaatttaagcatataattaagcggaggaaaagaaactaacaaggat
tccctcagtagcggcgagcgaagcgggaagagctcaacatgaaaaccgttggggggttcgc
gctttccttcgggttgtatagtaaagtggtgtcgcctgcgtcgcgccgggtataaatccc
gtctgaaatggggtgtcgtagagggttatatccccgtctgtgacccggtgtgcgatgcgt
tacggtgaccttacgatgagtcgggttgcttgggcatgcagccctaagtgggtggtaaac
ttcatctaaagctaaatactagcgagagaccgatagcgtacaagtaccgtaagggaaaga
tgcaaagaactttgaaaagagagttaaacagggcttgaaaccgttaagagtgaagcgaag
gaggtcagtgatcatggggccagatccccaagtgcctgtcata-gggcaccgtagcgttc
tggtttcccggggcagcgccggccgggggcgggggttaacagggaggtggttgagggatt
cgttttccgcctgtactcctttatagcttgcaccccggccgagggatccgcgacttgtcg
tgtgagggcgctgtcaaaatggccttctccaccccgtcttgaaacacggaccaaggagtc
taacatactcgcgagtgtttgggtggttaaaacccgaacgcacaatgaaagtaaacggtg
gctctgcgccgctgcttggtgggaaccctgcttttataatgggggggcactatcacccgac
cccgatccctttggggtggttggttttgaggtagagcgagtatgatgggacccgaaagat
ggtgaactattcctgagtagggcgaagccagaggaaactctggtggaggctcgtagcgat
gctgacgtgcaaatcgctcgtcgaacttgggtataggggcgaaagactaatcgaaccatc
tagtagctggttccctccgaagtttccctcaggatagcgggagcaaatgcaagtcagtat

catccggtaaagcgaatgattagaggcctcgggggagtaactccttcgacctattctcaa
actttaaataggtgaggagccggcaacgttgcttaaaacaatgaacgccgtgcggacaat
gattgctctcagtgggccgattttggtaagcagaactggcgatgcgggatgaaccgaaag
cgacgctaaggtgccaaactcccagctcattcagacaccagaaaaggtgttgattcatac
agacagcaggacggtggccatggaagtcggaatccgctaaggagtgtgtaacaactcacc
tgccgaatggatcagccctgaaaatggatggcgcttaagcgggggaccgaggcgtcgccg
ttggggtaaatacgaagccccaatgagtaggagggcgtgggggttttgaaacacggtaca
ctcgcatggaagccttgggcgtaagctcgggtggagcgacccctagtgtagatcttggtg
gtagtagcaaatattcaagtgagaaccttgaagactgaagtggggaagggttccacgcga
acagtgcttggacgtgggttagtcggtcctaagagatggggaaactccgtattcaagtgc
gcttttgtagcgcagtccatcgaaagggaagcgggttaacattcccgcaccaggatgtgg
aaactgtggtttaccgcaaactgcggcaacgcaactgaacctggggacgcccgtaggagc
tccgagcagggatctctttcctgtttaacgcgcccaacccccctggaatcggtttgtccgg
agatagggggcgaaaggccggaaaagcaccgcacgtcttgtggtgttcggaacgcttgta
atggcccttgaaaacccaggggaaagcagccgtcttacac--ttgtgtgagacgaggaat
tctcacacctggccgtacccatcaccgcatcaggtccccaaggtgaacagcctctagtca
atagaataatgtagataagggaagtcggcaaaatagatccgtaactttgggaaaaggatt
ggctctaatgttcgggcttgtcgggacgcgcgggcgtagatgccaatgagcggcagacgt
ctagcctctttggaggctggattaacgctgcttgtgggaagaaaccgtgtcgatggaggc
aggaacgcttatcgatcttgtctcttcccgcttgcgatcaacgaacggattagaactggt
acggacaaggggaatccgactgtttaattaaaacaaagcattgcgacggccggaaacggt
gttggcgcaatgtgatttctgcccagtgctctgaatgtcaaagtgaagaaattcaaccaa
gcgcgggtaaacggcgggagtaactatgactct--taaac-aaaccatactacctac---
-------------------gggagtcaaaagtgggggcggcgtctggtgtagaaatacac
gtac--------------------------------------------------------
-------------------gacacgctttttttcctgctagtcgatctgacctttgttct
ttggactgtgg-------------------------------------------------
--------------------------gcggtcagcggcaacataacc-------------
------------------------------------------------------------
------------------------------------------------------------
----------------------------tggtacggg--------------------ga
aggcctcgacgttga---------------------------------------------
---------------------------------gtgctgcatcttggaagtatc-------
------------------------------ttgacacaataagaga---------------
-------------------------------aaagagattgcgcttaatagg---------
----gggttgatctcgt-------------------------------------------

-------------------------------------------ggcgagccgacgc----
---------------------------------cggtgtgaactggt-------------
------------------------------------------------------------
-----------------------ggcaat-------------------------------
----ggccgtcgtagagcacgcaaaggtatgggctgactttttctcctca----aaagtt
ggcttaagggac-gtgccatcccacacgataataaagtgtgcagtctg------------
--------------------taagagctcccacagggcaaagac---------agactgg
gatgcagacggagacggcaaaaacaacaacatccgttctgcttctgggcgcaaatgctta
aggtagccaaatgcctcgtcatctaaactaattgg-at----------------------
-----------------------gagatagtc----------tgagtgaaaagcttatgc
ag-----------------gctagtaccgg----aaaaaaaaaaaagaggggctctgcca
ct----ctgac------------------------ggtgcgacactccttgtacagggaa
accctcc--gcatggaaaggcacgtggtaccaccctcgcggggaaactcgcgagcgggat
cccgggtaatgaccggagccgatggtaaaaactcacgtgaccgatggggtgaccctgtgg
cgagacgctaagttgtgggacatcaccctctcacaatacgcatccgtcgcagagactgaa
cgggagtgggctccgttga-----cggccggccggccagttagccgtcttcca-------
---ggggcttaagatacagtccgggcctcagggatgg----aaaaacctggggaacta-c
cagaattagtgacgcgcatgaatggattaacgagattcccactgtccctatctactatct
agcgaaaccacagccaagggaacgggcttggaagaatcagcggggaaagaagaccctgtt
gagcttgactctagtccgactttgtgagatgagttgggaggtgtagtctaagagggaggg
gcaactcgaccgtgaaatacctctactcccaagcttgttttacttactccggcgagcagg
gatgggtggtaacaccctttttctaggattaacacctcttcgggggtggacctgcgcgga
ggacattgtcaggtggggagtttggctggggcggcacatctgttaaaagataacgcaggt
gtcctaagatgagctcagtgagaacagaaatctcacgtagaacaaaagggtaaaagctca
tttgattttgattttcagtacgaatacaaactgtgaaagcatggcctatcgatccttttg
accttcaaggtttgaagcaagaggtgtcagaaaagt------------------------
------------------------------------------------------------
------------------------------------------------------------
------------------------------------------------------------
------------------------------------------------------------
------------------------------------------------------------
------------------------------------------------------------
---------------------------------------------------taccacagg
gat---------------------------------------------------------
------------------------------------------------------------
------------------------------------------------------------

------------------------------------------------------------
------------------------------------------------------------
------------------------------------------------------------
------------------------------------------------------------
------------------------------------------------------------
------------------------------------------------------------
------------------------------------------------------------
------------------------------------------------------------
------------------------------------------------------------
------------------------------------------------------------
------------------------------------------------------------
------------------------------------------------------------
------------------------------------------------------------
------------------------------------------------------------
------------------------------------------------------------
------------------------------------------------------------
------------------------------------------------------------
------------------------------------------------------------
------------------------------------------------------------
---------------------------------aactggcttgtggcagccaagcgttca
tagcgacgttgctttttgatccttcgatgtcggctcttcctatcattgcgaagcagaatt
cgccaagtgttggattgttcacccactaatagggaacgtgagctgggtagccacttta--
-----tgcccaacagctggatagctgggcttctgtctta-ggccgggacttctcacgacc
cgtgccttggtcaccc-aaaaaagggcgaccgcagagctcaggatggagcttctggctta
actagtagaggacagatagagatcgtcaaattgcgggaaaccccttagagcctcgagcta
ccaagtgcgtgtcgaaaggcatagacgcagccagcgatggctgtcgcatggcttga-ttc
catcaagataggtaaaaatgctcgaggattgggcaatccgcagccaagcctctaaggt--
---ccccccgcgggctatggggaaggttcaacgactgtaaggcgtaccgtgcaagcggaa
gatacagtctagccccgctggaaagcagcgccaaagtaagtaggt-tgcagtgggtctgt
cctc
>Form_I_China_SRR9587927
gcatgttttggtatgtgtttcgacagtcatgcatgtcatagccgtaggatggggagtgag
cttcatagcccgaccccttctcaggagcatgttttcatagcatattgagcatactggggt
ggtctcccatgtagctcgctctctcggtgcaggacctcccttttgcgggccacttcgatg
gcctccattatgga-gagcatgttttggcatatgaagcgttggtggtgaggctcgtagcc
tgtatatacataccaacgcagtcgtcatatgtctctctggcgggggtcggtcgtatgacc

ttgtccctacctggagcatgtattttgtatgat---atactggac-ggcagctgcgctgt
tcagtcgtcagccgtacctcatgtgcgcgtttccaggtccgaacgtttgacatgcaaggg
catcataccagaagcctgttttctcgtatgagcatgtaggtaggcaggctcttcagtcgt
ctacccataaatgcacttcgaagga-actttttggtattgaatggctttgtgcccagacg
ttcttgtgagcttgttctcgctctgcgatacgctggaagctgtggtattacctacagcag
acgcattcctagggcggacggcctctctgcagatgggcacatctgtagaaagaggctccc
cttgatgacgcaatgcaatggctgggtaaacggagtgctatacagaagggtcaaacctc-
tgtggaagtacactcataccagcattgtttgttgagaggggtcggctctacgagtgagga
cctatactgccatctcattcagatgcgtagacggtctttactcagactgcctttctggct
gattgtcctcggtccctcgtgcgcacgcgccgctggctgaatgggcttgtccttatggag
tgccgcgttcttc-aaagaatccgtcttcatacagcgacctcatttacatattgccggct
gaatgggcttgtccttatggagcactgcattcttctaggaattgctatctatacagcgac
ctcatttacatacaaggcacct-ttttctacatagacttgccatttggctgattgttctg
acagtcgctgcctcttcggagttctgtgtggacgagtggcc--tcctgagtgggccggtc
cttacggagtgcccaagtctgctatgactgggttctccatacagcgacctcactcagaat
tgtatgg--------------caccgtgccgat-----------tgaatggacaagcagt
ccttagggaatcgcatgaaagcatgacggcaacatggtgtatggacccgttgggccttgc
ctgtgctatcttctcgagtcgaccgg------------tcctcgtctcctcctttcgcga
cgggaagctcttgagtttaccgtgggtcaggacgctgtggtatctgtttgtctgtagtgt
gggctctctatcagaacagcgtatcagcacacgtcgtctgggtctatggtatttttgctt
aacggccctgggtttggtagtatggttccatgtcggtcctctcacctcccaaccgtcatg
tctctga-----------------------------------------------------
------------------------------------------------------------
--------------------------------------------tatcatctgcacagta
cgtcttgtcagaatatcattctgtatggcagtcgatagactaggccctatgccgcagctg
tctctgaccactatgcgagcgcatacctacctcaatgtgttcatagctgttatggatctt
gggtgtggtgtgccagaacgatacgccaccagattgtggaatggtgcttgtagtattaga
agaggacatgtatgaagagcacgttcttttctcttatgatggtgccgcgggtgcgggccg
ttctcattctgcttgtcgggtgctcaaggtgcatgtaggattgctgaacccac-tgaatg
gtcatgagaatggtcatatggtgaggttcacttctccatcgctttgtgtgtgttggatgt
ttgggtggcttcggccgcatggaaaatctcggtctgtaactcatcagggagggtttccat
gtgagcaatctctccagtcgaacacacacccatgggcgtcccgctaggcaaaatggaact
ctcgatccgacacagcctgtggagtgtttgttccgtgtggcgtgaaacggcttatactgg
atgtcttccc-tttggacaccagtctgttggtatg--tacgcataccatctagagggagc
aagaaaaaggcatgcaagtgccacaaaggaagggaggccttcctgctaccgtgtgtcc-t
cttgacgatcacgggggagtatatagagacatttggacgagatccgtttttcacgaggcg

gtgccctcgcgcaaaggtgttgctctgcacgcatctttatgatgatagttgctcgctcct
atgcgtccactcgtgaaaatacttgctctctctgtctgtttccatgacacccc-catgcg
ttgtcatagagcatgctcaggtacagaaagtgctgtgactttcttggcctgctgtcactg
agtcagcctgatactttggaggaatcttttctgggcactgcccagctgagagttcttatc
aggctctggagtgtgcatatgcctatattcattatagtgcatgtggctacgcagaaaaag
catcgacgtggaatggagaagaacaccattctgaatggaacggttcttactctacttctg
ttgcttgacgctgtatgggaatcgctttgctctgggggtacaccttccattccatgacaa
agcatttgatctatgccccga-gggtcattctctatgaagagcccgaaacttgtctgagt
gagaggcgaaggtagtccttcattt--atgcggggtccgccctcacactgtagtcggcac
agtcccttgtttatggactgcgcagctcaatcttcgactggagagg--aaaaacaaaaca
caagtatgacacgcacatacgtgttttctaggactgtgacctgtcggctgcctcgcggtg
gttggcatgatgccccaagt-gggggggggagaaacaatcagtatgattgataaaaacca
gtgggg-aaaaaggcgcgctgtcgcacacccttggtcacggcctc---aagtcgacggcc
ctcactggccca-ccctgcgactcctgttcgcttttgacaagacacacctgcaata-cgg
aggaaatgcctgagcagctgtgccgcagcaactgcggacgcgtagcgatatgcatccctg
ttgtttgcgacttggtacaaagaacgtgcgccgggagacggattgccctttgacttttcc
acccagcccgcggctctatacccttgacgtacctatttacctggttgatcctgccagtag
tcatatgcttgtctcaaagactaagccatgcatgtctaagtataaacacttttatactcg
tgaaactgcgaatggctcattaaaacagttatagttcctttgggaacaacactacttgga
taaccgtagtaattctagagctaatacatgcctcaacgcccgactcacgaagggtggtat
ttattggataaaaaaccatcgtgctctgtccagagcgctttgagatgattcacaataact
tgtcggatcgcatggcctcgtgctggcgactcctcattcaaatttctgccctatcaactt
tcgatggtagagtattggtctaccatggtgtcgacgggtgacggggaattagggttcgat
tccggagagggagcctgagaaacggctaccacatccaaggaaggcagcaggcgcgcaaat
tacccaatcccgactcggggaggtagtgacaaaaataacaatagggggccctttgggtc
ttctaattggaatgagaacaatttaaatcccttatcgaggatccattggagggcaagtct
ggtgaacgta----------------------------------tgcatta---------
--aagccctccgtaat-aaaaaaaaacagcgccagatgtct-------------------
------------------------------------------------------------
------------------------------------------------------------
------------------------------------------------------------
------------------------------------------------------------
------------------------------------------------------------
-------ctctctggaagcacacca-----------------------------------
---tgcatggggtttgttcctctc-----------------------------tgcatgg
ccgtg-----------------------------------------tgaaaaaggcgagt

aggcaaacgatgggaaacccggcttggaatagtaaccagccacagagcccccatcataca
cttctaaattgccggggagtgcccgcgaggccttgactaccgcggtccgccgagaggtga
gagcgagcaatcgttcaggacgcagcagtgtacgtagtggtacatggatggtaaaaacgt
caaggatagggttgatccgcagggaagcgcgagtgccgccatg---------gtcggtgg
gcaaaggcgaaccttcagagactctaatggagtgggcgcaagcttaagggagagtccaat
ccactgggaaaccaggtccacagcagtaacggggtgatgctcgccccagaggactgtgga
aggcaaccccacc----aaaagggtgtgtgttgtcgggtagc-aatggccagcagccgcg
gtaattccagctccaatagcgtatattaaagttgttgcagttaaaacgctcgtagtcgga
cctcggagcgcccccacgggccgggcttttgccttgttgccagtgggagtttgtctcctt
ttgtcgggacgcggtgtgcagggcctttactgtctctgcgcgtttacctgcgggccgacc
gtttactgtgaagaaagtaaagtgttcaaagcaggccaatgccttgaatatgtgagcatg
gaataatagaataggacttgggctctattttgttggtttccagtgaccaagtaatgatta
atagggacggttgggggcattcgtatttcattgtcagaggtgaaattcttggattgatgg
aagacgtacaactgcgaaagcatctgccatggatgttttcattgatcaagaacgaaagtt
aggggatcgaagacgatcagataccgtcgtagtcttaaccataaacgatgccgactgggg
attggcgggggcacttttatatgactctgtcagcaccctgagggaaaccaaagtctttgg
gttctgggggagtatggtcgcaaggctgaaacttaaaggaattgacggaagggcaccac
caggagtggagcctgcggcttaatttgactcaacacgggaaaacttaccaggtccggaca
gaagaatgattgacagactgaaagctctttcttgatttttggttggtggtgcatggccg
ttcttagttggtggagtgatttgtctggttaattccgttaacgaacgagacctcgtcctg
ctaaataggtgcgcgcatgccagcactgcgttcttaccttcttagagggactatgcgcgt
ctagcgtatggaagattgaggcaataacaggtctgtgatgcccttagatgttctgggccg
cacgcgcgctacactgatgcattcaacgagtttcacttgatagtcctgggtcggaaggct
cgggtaatcttttgaaagtgcatcgtgctggggatagatcattgcaattattgatcttca
acgaggaattccttgtaggcgcaggtcatcagcctgcgccgaatacgtccctgccctttg
tacacaccgcccgtcgctcctaccgattgaatggtccgatgaaatgtccggatcgcggct
agtgtgatggactcgtccagagcggtagccgtgagaagctcattaaatcttatcatttag
aggaaggagaagtcgtaacaaggt------------------------------------
------------------------------------------------------------
------------------------------------------------------------
------------------------------------------------------------
------------------------------------------------------------
------------------------------------------------------------
------------------------------------------------------------
------------------------------------------------------------
------------------------------------------------------------

------------------------------------------------------------
------------------------------------------------------------
------------------------------------------------------------
------------------------------------------------------------
------------------------------------------------------------
------------------------------------------------------------
------------------------------------------------------------
-------------------------------ttccgtaggtgaacctgcggaaggatcat
tcacaaactattgacacaaacacacgcgaaccaaatcgtccgcacagcaggtggcaatga
aagagagactctgcatgtcgcctttcgggg--tagcaagcagcactcttctgccatcgcc
tctgtgcagggcgtaaattcccattgagaggatgtgagggcaccacaggaagcttttcca
caggaagtcgccatccttctccctccacggcggcg----cttctgtgcttggcag-----
----tttttttttgccttctagggaggatgccgccaatggagccccttat-atatataca
tcatcatatgccc--ctttttttcttaaccgcttgccacagcttcttctatgaggagctt
gtgggaagactgtctccatacaataac-aaagatacaactcttagcggtggatatcttgg
ttctcgcaacgatgaagaacgcagccaactgcgataactaatgtgaattgcaggacttcg
tgaatcattaagtctttgaacgcaagttgcgctcatctccatgaggaggtgagcatgcct
gtttgagtgtcgatgggaaaaacaatac-ataacaacaatcggctctcttatcttcctgc
cttcttgggggcaaaaggtagtgcatgccggagatgggctttgtcgcccctttctcttga
aagaggcgtcgtccgaaacacgtatcgttgcccgctggaggtcatagagcatgagttt--
aaagcaccacatctctcaagtcagagtgag-cgtcttgagtaggtcc-gcattgattcag
tttcaacccaggtggcgttggttacttgccat-aagcctctctggggtttctttgcttgg
acctctcttgtcagtttctctgacgtagagacaggt--gccgtctcatggacgttgccct
ctggaacgtgct--aaaaaaaaaaaaaga----------aaaaaccgtaccacatgtcaa
aagaggcagatcgtgcttgacggcgggaggtgttgcatttcttgcttgtcctgagcgtct
gccagtctct----ga--cgagtg----tgttccttccataaacccgatctcaaatcagg
taggaccacccgctgaatttaagcatataattaagcggaggaaaagaaactaacaaggat
tccctcagtagcggcgagcgaagcgggaagagctcaacatgaaaaccgttgggggttcgc
gctttccttcgggttgtatagtaaagtggtgtcgcctgcgtcgcgccgggtataaatccc
gtctgaaatggggtgtcgtagagggttatatccccgtctgtgacccggtgtgcgatgcgt
tacggtgaccttacgatgagtcgggttgcttgggcatgcagccctaagtgggtggtaaac
ttcatctaaagctaaatactagcgagagaccgatagcgtacaagtaccgtaagggaaaga
tgcaaagaactttgaaaagagagttaaacagggcttgaaaccgttaagagtgaagcgaag
gaggtcagtgatcatggggccagatccccaagtgcctgtcata-gggcaccgtagcgttc
tggtttcccggggcagcgccggccgggggcgggggttaacagggaggtggttgagggatt
cgttttccgcctgtactcctttatagcttgcaccccggccgagggatccgcgacttgtcg

tgtgagggcgctgtcaaaatggccttctccaccccgtcttgaaacacggaccaaggagtc
taacatactcgcgagtgtttgggtggttaaaacccgaacgcacaatgaaagtaaacggtg
gctctgcgccgctgcttggtgggaaccctgcttttataatgggggcactatcacccgac
cccgatccctttggggtggttggttttgaggtagagcgagtatgatgggacccgaaagat
ggtgaactattcctgagtagggcgaagccagaggaaactctggtggaggctcgtagcgat
gctgacgtgcaaatcgctcgtcgaacttgggtataggggcgaaagactaatcgaaccatc
tagtagctggttccctccgaagtttccctcaggatagcgggagcaaatgcaagtcagtat
catccggtaaagcgaatgattagaggcctcgggggagtaactccttcgacctattctcaa
actttaaataggtgaggagccggcaacgttgcttaaaacaatgaacgccgtgcggacaat
gattgctctcagtgggccgattttggtaagcagaactggcgatgcgggatgaaccgaaag
cgacgctaaggtgccaaactcccagctcattcagacaccagaaaaggtgttgattcatac
agacagcaggacggtggccatggaagtcggaatccgctaaggagtgtgtaacaactcacc
tgccgaatggatcagccctgaaaatggatggcgcttaagcgggggaccgaggcgtcgccg
ttggggtaaatacgaagccccaatgagtaggagggcgtgggggttttgaaacacggtaca
ctcgcatggaagccttgggcgtaagctcgggtggagcgacccctagtgtagatcttggtg
gtagtagcaaatattcaagtgagaaccttgaagactgaagtggggaagggttccacgcga
acagtgcttggacgtgggttagtcggtcctaagagatggggaaactccgtattcaagtgc
gcttttgtagcgcagtccatcgaaagggaagcgggttaacattcccgcaccaggatgtgg
aaactgtggtttaccgcaaactgcggcaacgcaactgaacctggggacgcccgtaggagc
tccgagcagggatctctttcctgtttaacgcgcccaacccccctggaatcggtttgtccgg
agatagggggcgaaaggccggaaaagcaccgcacgtcttgtggtgttcggaacgcttgta
atggcccttgaaaacccaggggaaagcagccgtcttacac--ttgtgtgagacgaggaat
tctcacacctggccgtacccatcaccgcatcaggtccccaaggtgaacagcctctagtca
atagaataatgtagataagggaagtcggcaaaatagatccgtaactttgggaaaaggatt
ggctctaatgttcgggcttgtcgggacgcgcgggcgtagatgccaatgagcggcagacgt
ctagcctctttggaggctggattaacgctgcttgtgggaagaaaccgtgtcgatggaggc
aggaacgcttatcgatcttgtctcttcccgcttgcgatcaacgaacggattagaactggt
acggacaaggggaatccgactgtttaattaaaacaaagcattgcgacggccggaaacggt
gttggcgcaatgtgatttctgcccagtgctctgaatgtcaaagtgaagaaattcaaccaa
gcgcgggtaaacggcgggagtaactatgactct--taaac-aaaccatactacctac---
-------------------gggagtcaaaagtgggggcggcgtctggtgtagaaatacac
gtac--------------------------------------------------------
-------------------gacacgcttttttcctgctagtcgatctgacctttgttct
ttggactgtgg-------------------------------------------------
--------------------------gcggtcagcggcaacataacc-------------
------------------------------------------------------------

------------------------------------------------------------
----------------------------tggtacggg---------------------ga
aggcctcgacgttga---------------------------------------------
--------------------------------gtgctgcatcttggaagtatc-------
-----------------------------ttgacacaataagaga---------------
------------------------------aaagagattgcgcttaatagg---------
----gggttgatctcgt-------------------------------------------
-------------------------------------------ggcgagccgacgc----
---------------------------------cggtgtgaactggt-------------
------------------------------------------------------------
-----------------------ggcaat-------------------------------
----ggccgtcgtagagcacgcaaaggtatgggctgactttttctcctca----aaagtt
ggcttaagggac-gtgccatcccacacgataataaagtgtgcagtctg------------
--------------------taagagctcccacagggcaaagac---------agactgg
gatgcagacggagacggcaaaaacaacaacatccgttctgcttctgggcgcaaatgctta
aggtagccaaatgcctcgtcatctaaactaattgg-at----------------------
-----------------------gagatagtc----------tgagtgaaaagcttatgc
ag-----------------gctagtaccgg----aaaaaaaaaaaagaggggctctgcca
ct----ctgac------------------------ggtgcgacactccttgtacagggaa
accctcc--gcatggaaaggcacgtggtaccaccctcgcggggaaactcgcgagcgggat
cccgggtaatgaccggagccgatggtaaaaactcacgtgaccgatggggtgaccctgtgg
cgagacgctaagttgtgggacatcaccctctcacaatacgcatccgtcgcagagactgaa
cgggagtgggctccgttga-----cggccggccggccagccagccgtcttcca-------
---ggggcttaagatacagtccgggcctcagggatgg----aaaaacctggggaacta-c
cagaattagtgacgcgcatgaatggattaacgagattcccactgtccctatctactatct
agcgaaaccacagccaagggaacgggcttggaagaatcagcggggaaagaagaccctgtt
gagcttgactctagtccgactttgtgagatgagttgggaggtgtagtctaagagggaggg
gcaactcgaccgtgaaatacctctactcccaagcttgttttacttactccggcgagcagg
gatgggtggtaacacccttttctaggattaacacctcttcgggggtggacctgcgcgga
ggacattgtcaggtggggagtttggctggggcggcacatctgttaaaagataacgcaggt
gtcctaagatgagctcagtgagaacagaaatctcacgtagaacaaaagggtaaaagctca
tttgattttgattttcagtacgaatacaaactgtgaaagcatggcctatcgatcctttttg
accttcaaggtttgaagcaagaggtgtcagaaaagt------------------------
------------------------------------------------------------
------------------------------------------------------------
------------------------------------------------------------

------------------------------------------------------------
------------------------------------------------------------
------------------------------------------------------------
---------------------------------------------------taccacagg
gat---------------------------------------------------------
------------------------------------------------------------
------------------------------------------------------------
------------------------------------------------------------
------------------------------------------------------------
------------------------------------------------------------
------------------------------------------------------------
------------------------------------------------------------
------------------------------------------------------------
------------------------------------------------------------
------------------------------------------------------------
------------------------------------------------------------
------------------------------------------------------------
------------------------------------------------------------
------------------------------------------------------------
------------------------------------------------------------
------------------------------------------------------------
------------------------------------------------------------
------------------------------------------------------------
------------------------------------------------------------
------------------------------------------------------------
------------------------------------------------------------
---------------------------------aactggcttgtggcagccaagcgttca
tagcgacgttgctttttgatccttcgatgtcggctcttcctatcattgcgaagcagaatt
cgccaagtgttggattgttcacccactaatagggaacgtgagctgggtagccacttta--
-----tgcccaacagctggatagctgggcttctgtctta-ggccgggacttctcacgacc
cgtgccttggtcaccc-aaaaaagggcgaccgcagagctcaggatggagcttctggctta
actagtagaggacagatagagatcgtcaaattgcgggaaaccccttagagcctcgagcta
ccaagtgcgtgtcgaaaggcatagacgcagccagcgatggctgtcgcatggcttga-ttc
catcaagataggtaaaaatgctcgaggattgggcaatccgcagccaagcccctaaggt--
---ccccccgcgggctatggggaaggttcaacgactgtaaggcgtaccgtgcaagcggaa
gatacagtctagccccgctggaaagcagcgccaaagtaagtaggt-tgcagtgggtctgt

cctc

>Form_I_China_SRR9587929_SRR9587963

gcatgttttggtatgtgtttcgacagtcatgcatgtcatagccgtaggatggggagtgag
cttcatagcccgaccccttctcaggagcatgttttcatagcatattgagcatactggggt
ggtctcccatgtagctcgctctctcggtgcaggacctcccttttgcgggccacttcgatg
gcctccattatgga-gagcatgttttggcatatgaagcgttggtggtgaggctcgtagcc
tgtatatacataccaacgcagtcgtcatatgtctctctggcgggggtcggtcgtatgacc
ttgtccctacctggagcatgtattttgtatgat---atactggac-ggcagctgcgctgt
tcagtcgtcggccgtacctcatgtgcgcgtttccaggtccgaacgtttgacatgcaaggg
catcataccagaagcctgttttctcgtatgagcatgtaggtaggcaggctcttcagtcgt
ctacccataaatgcacttcgaagga-actttttggtattgaatggctttgtgcccagacg
ttcttgtgagcttgttctcgctctgcgatacgctggaagctgtggtattacctacagcag
acgcattcctagggcggacggcctctctgcagatgggcacatctgtagaaagaggctccc
cttgatgacgcaatgcaatggctgggtaaacggagtgctatacagaagggtcaaacctc-
tgtggaagtacacccataccagcattgtttgttgagaggggtcggctctacgagtgagga
cctatactgccatctcattcagatgcgtagacggtctttactcagactgcctttctggct
gattgtcctcggtccctcgtgcgcacgcgccgctggctgaatgggcttgtccttatggag
tgccgcgttcttc-aaagaatccgtcttcatacagcgacctcatttacatattgccggct
gaatgggcttgtccttatggagcactgcattcttctaggaattgctatctatacagcgac
ctcatttacatacaaggcacct-ttttctacatagacttgccatttggctgattgttctg
acagtcgctgcctcttcggagttctgtgtggacgagtggcc--tcctgagtgggccggtc
cttacggagtgcccacgtctgttatgactgggttctccatacagcgacctcactcagaat
tgtatgg--------------caccgtgccgat-----------tgaatggacaagcagt
ccttagggaatcgcatgaaagcatgacggc-acatggtgtatggacccgttgggccttgc
ctgtgctatcttctcgagtcgaccgg------------tcctcgtctcctcctttcgcga
cgggaagctcttgagtttaccgtgggtcaggacgctgtggtatctgtttgtctgtagtgt
gggctctctatcagaacagcgtatcagcacacgtcgtctgggtctatggtatttttgctt
aacggccctgggtttggtagtatggttccatgtcggtcctctcacctcccaaccgtcatg
tctctga-----------------------------------------------------
------------------------------------------------------------
--------------------------------------------tatcatctgcacagta
cgtcttgtcagaatatcattctgtatggcagtcgatagactaggccctatgccgcagctg
tctctgaccactatgcgagcgcatacctacctcaatgtgttcatagctgttatggatctt
gggtgtggtgtgccagaacgatacgccaccagattgtggaatggtgcttgtagtattaga
agaggacatgtatgaagagcacgttcttttctcttatgatggtgccgcgggtgcgggccg
ttttcattctgcttgtcgggtgctcaaggtgcatgtaggattgctgaacccac-tgaatg

gtcatgagaatggtcatatggtgaggttcacttctccatcgctttgtgtgtgttggatgt
ttgggtggcttcggccgcatggaaaatctcggtctgtaactcatcagggagggtttccat
gtgagcaatctctccagtcgaacacacacccatgggcgtcccgctaggcaaaatggaact
ctcgatccgacacagcctgtggagtgtttgttccgtgtggcgtgaaacggcttatactgg
atgtcttccc-tttggacaccagtctgttggtatg--tacgcataccatctagagggagc
aagaaaaaggcatgcaagtgccacaaaggaagggaggccttcctgctaccgtgtgtcc-t
cttgacgatcacgggggagtatatagagacatttggacgagatccgtttttcacgaggcg
gtgccctcgcgcaaaggtgttgctctgcacgcatctttatgatgatagttgctcgctcct
atgcgtccactcgtgaaaatacctgctctctctgtctgtttccatgacacccc-catgcg
ttgtcatagagcatgctcaggtacagaaagtgctgtgactttcttggcctgctgtcactg
agtcagcctgatactttggaggaatcttttctgggcactgcccagctgagagttcttatc
aggctctggagtgtgcatatgcctatattcattatagtgcatgtggctacgcagaaaaag
catcgacgtggaatggagaagaacaccattctgaatggaacggttcttactctacttctg
ttgcttgacgctgtatgggaatcgctttgctctgggggtacaccttccattccatgacaa
agcatttgatctatgccccga-gggtcattctctatgaagagcccgaaacttgtctgagt
gagaggcgaaggtagtccttcattt--atgcggggtccgccctcacactgtagtcggcac
agtcccttgtttatggactgcgcagctcaatcttcgactggagagg--aaaaacaaaaca
caagtatgacacgcacatacgtgttttctaggactgtgacctgtcggctgcctcgcggtg
gttggcatgatgccccaagt-gggggggggagaaacaatcagtatgattgataaaaacca
gtgggg-aaaaaggcgcgctgtcgcacacccttggtcacggcctc---aagtcgacggcc
ctcactggccca-ccctgcgactcctgttcgcttttgacaagacacacctgcaata-cgg
aggaaatgcctgagcagctgtgccgcagcaactgcggacgcgtagcgatatgcatccctg
ttgtttgcgacttggtacaaagaacgtgcgccgggagacggattgccctttgacttttcc
acccagcccgcggctctatacccttgacgtacctatttacctggttgatcctgccagtag
tcatatgcttgtctcaaagactaagccatgcatgtctaagtataaacacttttatactcg
tgaaactgcgaatggctcattaaaacagttatagttcctttgggaacaacactacttgga
taaccgtagtaattctagagctaatacatgcctcaacgcccgactcacgaagggtggtat
ttattggataaaaaaccatcgtgctctgtccagagcgctttgagatgattcacaataact
tgtcggatcgcatggcctcgtgctggcgactcctcattcaaatttctgccctatcaactt
tcgatggtagagtattggtctaccatggtgtcgacgggtgacggggaattagggttcgat
tccggagagggagcctgagaaacggctaccacatccaaggaaggcagcaggcgcgcaaat
tacccaatcccgactcggggaggtagtgacaaaaaataacaatagggggccctttgggtc
ttctaattggaatgagaacaatttaaatcccttatcgaggatccattggagggcaagtct
ggtgaacgta----------------------------------tgcatta---------
--aagccctccgtaat-aaaaaaaaacagcgccagatgtct-------------------
------------------------------------------------------------

------------------------------------------------------------

------------------------------------------------------------

------------------------------------------------------------

------------------------------------------------------------

-------ctctctggaagcacacca-----------------------------------

---tgcatggggtttgttcctctc-----------------------------tgcatgg

ccgtg-----------------------------------------tgaaaaaggcgagt

aggcaaacgatgggaaacccggcttggaatagtaaccagccacagagcccccatcataca

cttctaaattgccggggagtgcccgcgaggccttgactaccgcggtccgccgagaggtga

gagcgagcaatcgttcaggacgcagcagtgtacgtagtggtacatggatggtaaaaacgt

caaggatagggttgatccgcagggaagcgcgagtgccgccatg---------gtcggtgg

gcaaaggcgaaccttcagagactctaatggagtgggcgcaagcttaagggagagtccaat

ccactgggaaaccaggtccacagcagtaacggggtgatgctcgccccagaggactgtgga

aggcaaccccacc----aaaagggtgtgtgttgtcgggtagc-aatggccagcagccgcg

gtaattccagctccaatagcgtatattaaagttgttgcagttaaaacgctcgtagtcgga

cctcggagcgcccccacgggccgggcttttgccttgttgccagtgggagtttgtctcctt

ttgtcgggacgcggtgtgcggggcctttactgtctctgcgcgtttacctgcgggccgacc

gtttactgtgaagaaagtaaagtgttcaaagcaggccaatgccttgaatatgtgagcatg

gaataatagaataggacttgggctctattttgttggtttccagtgaccaagtaatgatta

atagggacggttgggggcattcgtatttcattgtcagaggtgaaattcttggattgatgg

aagacgtacaactgcgaaagcatctgccatggatgttttcattgatcaagaacgaaagtt

aggggatcgaagacgatcagataccgtcgtagtcttaaccataaacgatgccgactgggg

attggcgggggcacttttatatgactctgtcagcaccctgagggaaaccaaagtctttgg

gttctgggggggagtatggtcgcaaggctgaaacttaaaggaattgacggaagggcaccac

caggagtggagcctgcggcttaatttgactcaacacgggaaaacttaccaggtccggaca

gaagaatgattgacagactgaaagctctttcttgatttttggttggtggtgcatggccg

ttcttagttggtggagtgatttgtctggttaattccgttaacgaacgagacctcgtcctg

ctaaataggtgcgcgcatgccagcactgcgttcttaccttcttagagggactatgcgcgt

ctagcgtatggaagattgaggcaataacaggtctgtgatgcccttagatgttctgggccg

cacgcgcgctacactgatgcattcaacgagtttcacttgatagtcctgggtcggaaggct

cgggtaatcttttgaaagtgcatcgtgctggggatagatcattgcaattattgatcttca

acgaggaattccttgtaggcgcaggtcatcagcctgcgccgaatacgtccctgccctttg

tacacaccgcccgtcgctcctaccgattgaatggtccgatgaaatgtccggatcgcggct

agtgtgatggactcgtccagagcggtagccgtgagaagctcattaaatcttatcatttag

aggaaggagaagtcgtaacaaggt------------------------------------

------------------------------------------------------------

------------------------------------------------------------
------------------------------------------------------------
------------------------------------------------------------
------------------------------------------------------------
------------------------------------------------------------
------------------------------------------------------------
------------------------------------------------------------
------------------------------------------------------------
------------------------------------------------------------
------------------------------------------------------------
------------------------------------------------------------
------------------------------------------------------------
------------------------------------------------------------
------------------------------------------------------------
-------------------------------ttccgtaggtgaacctgcggaaggatcat
tcacaaactattgacacaaacacacgcgaaccaaatcgtccgcacagcaggtggcaatga
aagagagactctgcatgtcgcctttcgggg--tagcaagcagcactcttctgccatcgcc
tctgtgcagggcgtaaattcccattgagaggatgtgagggcaccaaaggaagcttttcca
caggaagtcgccatccttctccctccacggcggcg----cttctgtgcttggcag-----
-----ttttttttgccttctagggaggatgccgccaatggagccccttat-atatataca
tcatcatatgccc--ctttttttcttaaccgcttgccacagcttcttctatgaggagctt
gtgggaagactgtctccatacaataac-aaagatacaactcttagcggtggatatcttgg
ttctcgcaacgatgaagaacgcagccaactgcgataactaatgtgaattgcaggacttcg
tgaatcattaagtctttgaacgcaagttgcgctcatctccatgaggaggtgagcatgcct
gtttgagtgtcgatgggaaaaacaatac-ataacaacaatcggctctcttatcttcctgc
cttcttgggggcaaaaggtagtgcatgccggagatgggctttgtcgcccctttctcttga
aagaggcgtcgtccgaaacacgtatcgttgcccgctggaggtcatagagcatgagttt--
aaagcaccacatctctcaagtcagagtgag-cgtcttgagtaggtcc-gcattgattcag
tttcaacccaggtggcgttggttacttgccat-aagcctctctggggtttctttgcttgg
acctctcttgtcagtttctctgacgtagagacaggt--gccgtctcatggacgttgccct
ctggaacgtgcta-aaaaaaaaaaaaaga----------aaaaaccgtaccacatgtcaa
aagaggcagatcgtgcttgacggcgggaggtgttgcatttcttgcttgtcctgagcgtct
gccagtctct----ga--cgagtg----tgttccttccataaacccgatctcaaatcagg
taggaccacccgctgaatttaagcatataattaagcggaggaaaagaaactaacaaggat
tccctcagtagcggcgagcgaagcgggaagagctcaacatgaaaaccgttgggggttcgc
gctttccttcgggttgtatagtaaagtggtgtcgcctgcgtcgcgccgggtataaatccc

gtctgaaatggggtgtcgtagagggttatatccccgtctgtgacccggtgtgcgatgcgt
tacggtgaccttacgatgagtcgggttgcttgggcatgcagccctaagtgggtggtaaac
ttcatctaaagctaaatactagcgagagaccgatagcgtacaagtaccgtaagggaaaga
tgcaaagaactttgaaaagagagttaaacagggcttgaaaccgttaagagtgaagcgaag
gaggtcagtgatcatggggccagatccccaagtgcctgtcata-gggcaccgtagcgttc
tggtttcccggggcagcgccggccgggggcgggggttaacagggaggtggttgagggatt
cgttttccgcctgtactcctttatagcttgcaccccggccgagggatccgcgacttgtcg
tgtgagggcgctgtcaaaatggccttctccaccccgtcttgaaacacggaccaaggagtc
taacatactcgcgagtgtttgggtggttaaacccgaacgcacaatgaaagtaaacggtg
gctctgcgccgctgcttggtgggaaccctgcttttataatgggggcactatcacccgac
cccgatccctttggggtggttggttttgaggtagagcgagtatgatgggacccgaaagat
ggtgaactattcctgagtagggcgaagccagaggaaactctggtggaggctcgtagcgat
gctgacgtgcaaatcgctcgtcgaacttgggtataggggcgaaagactaatcgaaccatc
tagtagctggttccctccgaagtttccctcaggatagcgggagcaaatgcaagtcagtat
catccggtaaagcgaatgattagaggcctcgggggagtaactccttcgacctattctcaa
actttaaataggtgaggagccggcaacgttgcttaaaacaatgaacgccgtgcggacaat
gattgctctcagtgggccgattttggtaagcagaactggcgatgcgggatgaaccgaaag
cgacgctaaggtgccaaactcccagctcattcagacaccagaaaaggtgttgattcatac
agacagcaggacggtggccatggaagtcggaatccgctaaggagtgtgtaacaactcacc
tgccgaatggatcagccctgaaaatggatggcgcttaagcgggggaccgaggcgtcgccg
ttggggtaaatacgaagccccaatgagtaggagggcgtgggggttttgaaacacggtaca
ctcgcatggaagccttgggcgtaagctcgggtggagcgacccctagtgtagatcttggtg
gtagtagcaaatattcaagtgagaaccttgaagactgaagtggggaagggttccacgcga
acagtgcttggacgtgggttagtcggtcctaagagatggggaaactccgtattcaagtgc
gcttttgtagcgcagtccatcgaaagggaagcgggttaacattcccgcaccaggatgtgg
aaactgtggtttaccgcaaactgcggcaacgcaactgaacctggggacgcccgtaggagc
tccgagcagggatctctttcctgtttaacgcgcccaacccccctggaatcggtttgtccgg
agatagggggcgaaaggccggaaaagcaccgcacgtcttgtggtgttcggaacgcttgta
atggcccttgaaaacccaggggaaagcagccgtcttacac--ttgtgtgagacgaggaat
tctcacacctggccgtacccatcaccgcatcaggtccccaaggtgaacagcctctagtca
atagaataatgtagataagggaagtcggcaaaatagatccgtaactttgggaaaaggatt
ggctctaatgttcgggcttgtcgggacgcgcgggcgtagatgccaatgagcggcagacgt
ctagcctctttggaggctggattaacgctgcttgtgggaagaaaccgtgtcgatggaggc
aggaacgcttatcgatcttgtctcttcccgcttgcgatcaacgaacggattagaactggt
acggacaaggggaatccgactgtttaattaaaacaaagcattgcgacggccggaaacggt
gttggcgcaatgtgatttctgcccagtgctctgaatgtcaaagtgaagaaattcaaccaa

gcgcgggtaaacggcgggagtaactatgactct--taaac-aaaccatactacctac---
-------------------gggagtcaaaagtgggggcggcgtctggtgtagaaatacac
gtac--------------------------------------------------------
-------------------gacacgctttttttcctgctagtcgatctgacctttgttct
ttggactgtgg-------------------------------------------------
--------------------------gcggtcagcggcaacataacc-------------
------------------------------------------------------------
------------------------------------------------------------
----------------------------tggtacggg---------------------ga
aggcctcgacgttga---------------------------------------------
--------------------------------gtgctgcatcttggaagtatc-------
-----------------------------ttgacacaataagaga---------------
------------------------------aaagagattgcgcttaatagg---------
----gggttgatctcgt-------------------------------------------
-------------------------------------------ggcgagccgacgc----
---------------------------------cggtgtgaactggt-------------
------------------------------------------------------------
-----------------------ggcaat-------------------------------
----ggccgtcgtagagcacgcaaaggtatgggctgactttttctcctca----aaagtt
ggcttaagggac-gtgccatcccacacgataataaagtgtgcagtctg------------
--------------------taagagctcccacagggcaaagac---------agactgg
gatgcagacggagacggcaaaaacaacaacatccgttctgcttctgggcgcaaatgctta
aggtagccaaatgcctcgtcatctaaactaattgg-at----------------------
-----------------------gagatagtc----------tgagtgaaaagcttatgc
ag-----------------gctagtaccggaa-aaaaaaaaaaaaagaggggctctgcca
ct----ctgac------------------------ggtgcgacactccttgtacagggaa
accctcc--gcatggaaaggcacgtggtaccaccctcgcggggaaactcgcgagcgggat
cccgggtaatgaccggagccgatggtaaaaactcacgtgaccgatggggtgaccctgtgg
cgagacgctaagttgtgggacatcaccctctcacaatacgcatccgtcgcagagactgaa
cgggagtgggctccgttga-----cggccggccggccagccagccgtcttcca-------
---ggggcttaagatacagtccgggcctcagggatgg----aaaaacctggggaacta-c
cagaattagtgacgcgcatgaatggattaacgagattcccactgtccctatctactatct
agcgaaaccacagccaagggaacgggcttggaagaatcagcggggaaagaagaccctgtt
gagcttgactctagtccgactttgtgagatgagttgggaggtgtagtctaagagggaggg
gcaactcgaccgtgaaatacctctactcccaagcttgttttacttactccggcgagcagg
gatgggtggtaacaccctttttctaggattaacacctcttcgggggtggacctgcgcgga

ggacattgtcaggtggggagtttggctggggcggcacatctgttaaaagataacgcaggt
gtcctaagatgagctcagtgagaacagaaatctcacgtagaacaaaagggtaaaagctca
tttgattttgattttcagtacgaatacaaactgtgaaagcatggcctatcgatccttttg
accttcaaggtttgaagcaagaggtgtcagaaaagt------------------------
------------------------------------------------------------
------------------------------------------------------------
------------------------------------------------------------
------------------------------------------------------------
------------------------------------------------------------
------------------------------------------------------------
---------------------------------------------------taccacagg
gat---------------------------------------------------------
------------------------------------------------------------
------------------------------------------------------------
------------------------------------------------------------
------------------------------------------------------------
------------------------------------------------------------
------------------------------------------------------------
------------------------------------------------------------
------------------------------------------------------------
------------------------------------------------------------
------------------------------------------------------------
------------------------------------------------------------
------------------------------------------------------------
------------------------------------------------------------
------------------------------------------------------------
------------------------------------------------------------
------------------------------------------------------------
------------------------------------------------------------
------------------------------------------------------------
------------------------------------------------------------
------------------------------------------------------------
------------------------------------------------------------
---------------------------------aactggcttgtggcagccaagcgttca
tagcgacgttgctttttgatccttcgatgtcggctcttcctatcattgcgaagcagaatt
cgccaagtgttggattgttcacccactaatagggaacgtgagctgggtagccacttta--

-----tgcccaacagctggatagctgggcttctgtctta-ggccgggacttctcacgacc
cgtgccttggtcaccc-aaaaaagggcgaccgcagagctcaggatggagcttctggctta
actagtagaggacagatagagatcgtcaaattgcgggaaaccccttagagcctcgagcta
ccaagtgcgtgtcgaaaggcatagacgcagccagcgatggctgtcgcatggcttga-ttc
catcaagataggtaaaaatgctcgaggattgggcaatccgcagccaagcccctaaggt--
---ccccccgcgggctatggggaaggttcaacgactgtaaggcgtaccgtgcaagcggaa
gatacagtctagccccgctggaaagcagcgccaaagtaagtaggt-tgcagtgggtctgt
cctc
>Form_B_China_SRR9587931
gcatgttttggtatgtgtttcgacagtcatgcatgtcatagccgtaggatggggagtgag
cttcatagcccgaccccttctcaggagcatgttttcatagcatattgagcatactggggt
ggtctcccatgtagctcgctctctcggtgcaggacctcccttttgcgggccacttcgatg
gcctccattatgga-gagcatgttttggcatatgaagcgttggtggtgaggctcgtagcc
tgtatatacataccaacgcagtcgtcatatgtctctctggcgggggtcggtcgtatgacc
ttgtccctacctggagcatgtattttgtatgat---atactggac-ggcagctgcgctgt
tcagtcgtcggccgtacctcatgtgcgcgtttccaggtccgaacgtttgacatgcaaggg
catcataccagaagcctgttttctcgtatgagcatgtaggtaggcaggctcttcagtcgt
ctacccataaatgcacttcgaagga-actttttggtattgaatggctttgtgcccagacg
ttcttgtgagcttgttctcgctctgcgatacgctggaagctgtggtattacctacagcag
acgcattcctagggcggacggcctctctgcagatgggcacatctgtagaaagaggctccc
cttgatgacgcaatgcaatggctgggtaaacggagtgctatacagaagggtcaaacctc-
tgtggaagtacacccataccagcattgtttgttgagaggggtcggctctacgagtgagga
cctatactgccatctcattcagatgcgtagacggtctttactcagactgcctttctggct
gattgtcctcggtccctcgtgcgcacgcgccgctggctgaatgggcttgtccttatggag
tgccgcgttcttc-aaagaatccgtcttcatacagcgacctcatttacatattgccggct
gaatgggcttgtccttatggagcactgcattcttctaggaattgctatctatacagcgac
ctcatttacatacaaggcacct-ttttctacatagacttgccatttggctgattgttctg
acagtcgctgcctcttcggagttctgtgtggacgagtggcc--tcctgagtgggccggtc
cttacggagtgcccacgtctgttatgactgggttctccatacagcgacctcactcagaat
tgtatgg--------------caccgtgccgat-----------tgaatggacaagcagt
ccttagggaatcgcatgaaagcatgacggc-acatggtgtatggacccgttgggccttgc
ctgtgctatcttctcgagtcgaccgg------------tcctcgtctcctcctttcgcga
cgggaagctcttgagtttaccgtgggtcaggacgctgtggtatctgtttgtctgtagtgt
gggctctctatcagaacagcgtatcagcacacgtcgtctgggtctatggtatttttgctt
aacggccctgggtttggtagtatggttccatgtcggtcctctcacctcccaaccgtcatg
tctctga-----------------------------------------------------

------------------------------------------------------------
--------------------------------------------tatcatctgcacagta
cgtcttgtcagaatatcattctgtatggcagtcgatagactaggccctatgccgcagctg
tctctgaccactatgcgagcgcatacctacctcaatgtgttcatagctgttatggatctt
gggtgtggtgtgccagaacgatacgccaccagattgtggaatggtgcttgtagtattaga
agaggacatgtatgaagagcacgttcttttctcttatgatggtgccgcgggtgcgggccg
ttttcattctgcttgtcgggtgctcaaggtgcatgtaggattgctgaacccac-tgaatg
gtcatgagaatggtcatatggtgaggttcacttctccatcgctttgtgtgtgttggatgt
ttgggtggcttcggccgcatggaaaatctcggtctgtaactcatcagggagggtttccat
gtgagcaatctctccagtcgaacacacacccatgggcgtcccgctaggcaaaatggaact
ctcgatccgacacagcctgtggagtgtttgttccgtgtggcgtgaaacggcttatactgg
atgtcttccc-tttggacaccagtctgttggtatg--tacgcataccatctagagggagc
aagaaaaaggcatgcaagtgccacaaaggaagggaggccttcctgctaccgtgtgtcc-t
cttgacgatcacgggggagtatatagagacatttggacgagatccgtttttcacgaggcg
gtgccctcgcgcaaaggtgttgctctgcacgcatctttatgatgatagttgctcgctcct
atgcgtccactcgtgaaaatacctgctctctctgtctgtttccatgacacccc-catgcg
ttgtcatagagcatgctcaggtacagaaagtgctgtgactttcttggcctgctgtcactg
agtcagcctgatactttggaggaatcttttctgggcactgcccagctgagagttcttatc
aggctctggagtgtgcatatgcctatattcattatagtgcatgtggctacgcagaaaaag
catcgacgtggaatggagaagaacaccattctgaatggaacggttcttactctacttctg
ttgcttgacgctgtatgggaatcgctttgctctgggggtacaccttccattccatgacaa
agcatttgatctatgccccga-gggtcattctctatgaagagcccgaaacttgtctgagt
gagaggcgaaggtagtccttcattt--atgcggggtccgccctcacactgtagtcggcac
agtcccttgtttatggactgcgcagctcaatcttcgactggagagg--aaaaacaaaaca
caagtatgacacgcacatacgtgttttctaggactgtgacctgtcggctgcctcgcggtg
gttggcatgatgccccaagt-ggggggggagaaacaatcagtatgattgataaaaacca
gtgggg-aaaaaggcgcgctgtcgcacacccttggtcacggcctc---aagtcgacggcc
ctcactggccca-ccctgcgactcctgttcgcttttgacaagacacacctgcaata-cgg
aggaaatgcctgagcagctgtgccgcagcaactgcggacgcgtagcgatatgcatccctg
ttgtttgcgacttggtacaaagaacgtgcgccgggagacggattgccctttgacttttcc
acccagcccgcggctctatacccttgacgtacctatttacctggttgatcctgccagtag
tcatatgcttgtctcaaagactaagccatgcatgtctaagtataaacacttttatactcg
tgaaactgcgaatggctcattaaaacagttatagttcctttgggaacaacactacttgga
taaccgtagtaattctagagctaatacatgcctcaacgcccgactcacgaagggtggtat
ttattggataaaaaaccatcgtgctctgtccagagcgctttgagatgattcacaataact
tgtcggatcgcatggcctcgtgctggcgactcctcattcaaatttctgccctatcaactt

tcgatggtagagtattggtctaccatggtgtcgacgggtgacggggaattagggttcgat
tccggagagggagcctgagaaacggctaccacatccaaggaaggcagcaggcgcgcaaat
tacccaatcccgactcggggaggtagtgacaaaaaataacaatagggggccctttgggtc
ttctaattggaatgagaacaatttaaatcccttatcgaggatccattggagggcaagtct
ggtgaacgta----------------------------------tgcatta---------
--aagccctccgtaat-aaaaaaaaacagcgccagatgtct-------------------
------------------------------------------------------------
------------------------------------------------------------
------------------------------------------------------------
------------------------------------------------------------
------------------------------------------------------------
-------ctctctggaagcacacca-----------------------------------
---tgcatggggtttgttcctctc-----------------------------tgcatgg
ccgtg-----------------------------------------tgaaaaaggcgagt
aggcaaacgatgggaaacccggcttggaatagtaaccagccacagagcccccatcataca
cttctaaattgccggggagtgcccgcgaggccttgactaccgcggtccgccgagaggtga
gagcgagcaatcgttcaggacgcagcagtgtacgtagtggtacatggatggtaaaaacgt
caaggatagggttgatccgcagggaagcgcgagtgccgccatg---------gtcggtgg
gcaaaggcgaaccttcagagactctaatggagtgggcgcaagcttaagggagagtccaat
ccactgggaaaccaggtccacagcagtaacggggtgatgctcgccccagaggactgtgga
aggcaaccccacc----aaaagggtgtgtgttgtcgggtagc-aatggccagcagccgcg
gtaattccagctccaatagcgtatattaaagttgttgcagttaaaacgctcgtagtcgga
cctcggagcgcccccacgggccgggcttttgccttgttgccagtgggagtttgtctcctt
ttgtcgggacgcggtgtgcggggcctttactgtctctgcgcgtttacctgcgggccgacc
gtttactgtgaagaaagtaaagtgttcaaagcaggccaatgccttgaatatgtgagcatg
gaataatagaataggacttgggctctattttgttggtttccagtgaccaagtaatgatta
atagggacggttgggggcattcgtatttcattgtcagaggtgaaattcttggattgatgg
aagacgtacaactgcgaaagcatctgccatggatgttttcattgatcaagaacgaaagtt
aggggatcgaagacgatcagataccgtcgtagtcttaaccataaacgatgccgactgggg
attggcgggggcacttttatatgactctgtcagcaccctgagggaaaccaaagtctttgg
gttctgggggggagtatggtcgcaaggctgaaacttaaaggaattgacggaagggcaccac
caggagtggagcctgcggcttaatttgactcaacacgggaaaacttaccaggtccggaca
gaagaatgattgacagactgaaagctctttcttgatttttttggttggtggtgcatggccg
ttcttagttggtggagtgatttgtctggttaattccgttaacgaacgagacctcgtcctg
ctaaataggtgcgcgcatgccagcactgcgttcttaccttcttagagggactatgcgcgt
ctagcgtatggaagattgaggcaataacaggtctgtgatgcccttagatgttctgggccg

cacgcgcgctacactgatgcattcaacgagtttcacttgatagtcctgggtcggaaggct
cgggtaatcttttgaaagtgcatcgtgctggggatagatcattgcaattattgatcttca
acgaggaattccttgtaggcgcaggtcatcagcctgcgccgaatacgtccctgccctttg
tacacaccgcccgtcgctcctaccgattgaatggtccgatgaaatgtccggatcgcggct
agtgtgatggactcgtccagagcggtagccgtgagaagctcattaaatcttatcatttag
aggaaggagaagtcgtaacaaggt------------------------------------
------------------------------------------------------------
------------------------------------------------------------
------------------------------------------------------------
------------------------------------------------------------
------------------------------------------------------------
------------------------------------------------------------
------------------------------------------------------------
------------------------------------------------------------
------------------------------------------------------------
------------------------------------------------------------
------------------------------------------------------------
------------------------------------------------------------
------------------------------------------------------------
------------------------------------------------------------
------------------------------------------------------------
-------------------------------ttccgtaggtgaacctgcggaaggatcat
tcacaaactattgacacaaacacacgcgaaccaaatcgtccgcacagcaggtggcaatga
aagagagactctgcatgtcgcctttcgggg--tagcaagcagcactcttctgccatcgcc
tctgtgcagggcgtaaattcccattgagaggatgtgagggcaccaaaggaagcttttcca
caggaagtcgccatccttctccctccacggcggcg----cttctgtgcttggcag-----
-----ttttttttgccttctagggaggatgccgccaatggagccccttat-atatataca
tcatcatatgccc--ctttttttcttaaccgcttgccacagcttcttctatgaggagctt
gtgggaagactgtctccatacaataac-aaagatacaactcttagcggtggatatcttgg
ttctcgcaacgatgaagaacgcagccaactgcgataactaatgtgaattgcaggacttcg
tgaatcattaagtctttgaacgcaagttgcgctcatctccatgaggaggtgagcatgcct
gtttgagtgtcgatgggaaaaacaatac-ataacaacaatcggctctcttatcttcctgc
cttcttgggggcaaaaggtagtgcatgccggagatgggctttgtcgcccctttctcttga
aagaggcgtcgtccgaaacacgtatcgttgcccgctggaggtcatagagcatgagttt--
aaagcaccacatctctcaagtcagagtgag-cgtcttgagtaggtcc-gcattgattcag
tttcaacccaggtggcgttggttacttgccat-aagcctctctggggtttctttgcttgg

acctctcttgtcagtttctctgacgtagagacaggt--gccgtctcatggacgttgccct
ctggaacgtgcta-aaaaaaaaaaaaaga----------aaaaaccgtaccacatgtcaa
aagaggcagatcgtgcttgacggcgggaggtgttgcatttcttgcttgtcctgagcgtct
gccagtctct----ga--cgagtg----tgttccttccataaacccgatctcaaatcagg
taggaccacccgctgaatttaagcatataattaagcggaggaaaagaaactaacaaggat
tccctcagtagcggcgagcgaagcgggaagagctcaacatgaaaaccgttgggggttcgc
gctttccttcgggttgtatagtaaagtggtgtcgcctgcgtcgcgccgggtataaatccc
gtctgaaatggggtgtcgtagagggttatatccccgtctgtgacccggtgtgcgatgcgt
tacggtgaccttacgatgagtcgggttgcttgggcatgcagccctaagtgggtggtaaac
ttcatctaaagctaaatactagcgagagaccgatagcgtacaagtaccgtaagggaaaga
tgcaaagaactttgaaaagagagttaaacagggcttgaaaccgttaagagtgaagcgaag
gaggtcagtgatcatggggccagatccccaagtgcctgtcata-gggcaccgtagcgttc
tggtttcccggggcagcgccggccgggggcgggggttaacagggaggtggttgagggatt
cgttttccgcctgtactcctttatagcttgcaccccggccgagggatccgcgacttgtcg
tgtgagggcgctgtcaaaatggccttctccaccccgtcttgaaacacggaccaaggagtc
taacatactcgcgagtgtttgggtggttaaacccgaacgcacaatgaaagtaaacggtg
gctctgcgccgctgcttggtgggaaccctgcttttataatgggggcactatcacccgac
cccgatccctttggggtggttggttttgaggtagagcgagtatgatgggacccgaaagat
ggtgaactattcctgagtagggcgaagccagaggaaactctggtggaggctcgtagcgat
gctgacgtgcaaatcgctcgtcgaacttgggtataggggcgaaagactaatcgaaccatc
tagtagctggttccctccgaagtttccctcaggatagcgggagcaaatgcaagtcagtat
catccggtaaagcgaatgattagaggcctcgggggagtaactccttcgacctattctcaa
actttaaataggtgaggagccggcaacgttgcttaaaacaatgaacgccgtgcggacaat
gattgctctcagtgggccgattttggtaagcagaactggcgatgcgggatgaaccgaaag
cgacgctaaggtgccaaactcccagctcattcagacaccagaaaaggtgttgattcatac
agacagcaggacggtggccatggaagtcggaatccgctaaggagtgtgtaacaactcacc
tgccgaatggatcagccctgaaaatggatggcgcttaagcgggggaccgaggcgtcgccg
ttggggtaaatacgaagccccaatgagtaggagggcgtgggggttttgaaacacggtaca
ctcgcatggaagccttgggcgtaagctcgggtggagcgacccctagtgtagatcttggtg
gtagtagcaaatattcaagtgagaaccttgaagactgaagtggggaagggttccacgcga
acagtgcttggacgtgggttagtcggtcctaagagatggggaaactccgtattcaagtgc
gcttttgtagcgcagtccatcgaaagggaagcgggttaacattcccgcaccaggatgtgg
aaactgtggtttaccgcaaactgcggcaacgcaactgaacctggggacgcccgtaggagc
tccgagcagggatctctttcctgtttaacgcgcccaacccccctggaatcggtttgtccgg
agatagggggcgaaaggccggaaaagcaccgcacgtcttgtggtgttcggaacgcttgta
atggcccttgaaaacccaggggaaagcagccgtcttacac--ttgtgtgagacgaggaat

tctcacacctggccgtacccatcaccgcatcaggtccccaaggtgaacagcctctagtca

atagaataatgtagataagggaagtcggcaaaatagatccgtaactttgggaaaaggatt

ggctctaatgttcgggcttgtcgggacgcgcgggcgtagatgccaatgagcggcagacgt

ctagcctctttggaggctggattaacgctgcttgtgggaagaaaccgtgtcgatggaggc

aggaacgcttatcgatcttgtctcttcccgcttgcgatcaacgaacggattagaactggt

acggacaaggggaatccgactgtttaattaaaacaaagcattgcgacggccggaaacggt

gttggcgcaatgtgatttctgcccagtgctctgaatgtcaaagtgaagaaattcaaccaa

gcgcgggtaaacggcgggagtaactatgactct--taaac-aaaccatactacctac---

-------------------gggagtcaaaagtgggggcggcgtctggtgtagaaatacac

gtac--------------------------------------------------------

-------------------gacacgctttttttcctgctagtcgatctgacctttgttct

ttggactgtgg-------------------------------------------------

--------------------------gcggtcagcggcaacataacc-------------

------------------------------------------------------------

------------------------------------------------------------

----------------------------tggtacggg---------------------ga

aggcctcgacgttga---------------------------------------------

--------------------------------gtgctgcatcttggaagtatc-------

-----------------------------ttgacacaataagaga---------------

------------------------------aaagagattgcgcttaatagg---------

----gggttgatctcgt-------------------------------------------

-------------------------------------------ggcgagccgacgc----

---------------------------------cggtgtgaactggt-------------

------------------------------------------------------------

-----------------------ggcaat-------------------------------

----ggccgtcgtagagcacgcaaaggtatgggctgactttttctcctca----aaagtt

ggcttaagggac-gtgccatcccacacgataataaagtgtgcagtctg------------

--------------------taagagctcccacagggcaaagac---------agactgg

gatgcagacggagacggcaaaaacaacaacatccgttctgcttctgggcgcaaatgctta

aggtagccaaatgcctcgtcatctaaactaattgg-at----------------------

-----------------------gagatagtc----------tgagtgaaaagcttatgc

ag-----------------gctagtaccggaa-aaaaaaaaaaaaagaggggctctgcca

ct----ctgac------------------------ggtgcgacactccttgtacagggaa

accctcc--gcatggaaaggcacgtggtaccaccctcgcggggaaactcgcgagcgggat

cccgggtaatgaccggagccgatggtaaaaactcacgtgaccgatggggtgaccctgtgg

cgagacgctaagttgtgggacatcaccctctcacaatacgcatccgtcgcagagactgaa

cgggagtgggctccgttga-----cggccggccggccagccagccgtcttcca-------
---ggggcttaagatacagtccgggcctcagggatgg----aaaaacctggggaacta-c
cagaattagtgacgcgcatgaatggattaacgagattcccactgtccctatctactatct
agcgaaaccacagccaagggaacgggcttggaagaatcagcggggaaagaagaccctgtt
gagcttgactctagtccgactttgtgagatgagttgggaggtgtagtctaagagggaggg
gcaactcgaccgtgaaatacctctactcccaagcttgttttacttactccggcgagcagg
gatgggtggtaacacccttttttctaggattaacacctcttcgggggtggacctgcgcgga
ggacattgtcaggtggggagtttggctggggcggcacatctgttaaaagataacgcaggt
gtcctaagatgagctcagtgagaacagaaatctcacgtagaacaaaagggtaaaagctca
tttgattttgattttcagtacgaatacaaactgtgaaagcatggcctatcgatcctttttg
accttcaaggtttgaagcaagaggtgtcagaaaagttaacacgcgttgtggtagcttttta
agagtggactcaaagtgttccgctagtccccccctcggaccgccacttacaaaaggcctgg
ggatgggggcgacactgtcaaattgcggggactccctaaagcctgtctctaccaagccag
cgcggcgaaagcccctggtggccggggtaatgacctagggtatggtaaaagcgagacagg
atg-gaacaatgggcaatccgcagccaagtcctaagg----ctttttgaaaagcggctat
ggatgcagttcaacgactagatggcagtgggctgagagagcccaccgcggcaccctttcg
gcttaagatatagtcttgcccacgtgaaagcgtgcctgcagatgaacgccaccatgtatg
gccggagctgcaggagaccgc--acttgatgtgcagtcgggaat-cgacagtaccacagg
gatgaac-----------------------------------------------------
------------------------------------------------------------
------------------------------------------------------------
------------------------------------------------------------
------------------------------------------------------------
-------------------------------------------------aaaaacac---
------------------------------------------------------------
------------------------------------------------------------
------------------------------------------------------------
--------------------------------------acaattgtggcc---atcatac
----------------------------------------agtcgta-------------
------------------------------------------------------------
------------------------------------------------------------
------------------------------------------------------------
------------tgatattatcgtcccaggcgggtgcgcctctctcacccgtctagtccg
gccttggtgcgcaaaaaatgcttgcgtggatgcgccttggttaaggaaaatgctcctagt
agttttacaaggacacttcttgtagggcaacacatctggatgcgaagaactcccggaaaa
gacaacggtaccaagtgactggggtaacgccccattcatgaaggcaaataatcaaaaaga

ggtgccgggtaagaacccgttgtatagggacaatccgcagctgacttggctccctg--cc
cat--ccatgggcatagcttagcggttcaacgctcggtaagatgtgggttggcgcgctgt
ccgacagcgcactggcttaaggtacgggctactcccaccggagacggtgtctttgtctac
ctcgggtaacacgcctcgggagaagcaaagagggtgcaatgg-aaaaaaactggttgcgg
ggaaatccctaccagagtcacctggtacaaaggaactggcttgtggcagccaagcgttca
tagcgacgttgctttttgatccttcgatgtcggctcttcctatcattgcgaagcagaatt
cgccaagtgttggattgttcacccactaatagggaacgtgagctgggtagccacttta--
-----tgcccaacagctggatagctgggcttctgtctta-ggccgggacttctcacgacc
cgtgccttggtcaccc-aaaaaagggcgaccgcagagctcaggatggagcttctggctta
actagtagaggacagatagagatcgtcaaattgcgggaaaccccttagagcctcgagcta
ccaagtgcgtgtcgaaaggcatagacgcagccagcgatggctgtcgcatggcttga-ttc
catcaagataggtaaaaatgctcgaggattgggcaatccgcagccaagcccctaaggt--
---ccccccgcgggctatggggaaggttcaacgactgtaaggcgtaccgtgcaagcggaa
gatacagtctagccccgctggaaagcagcgccaaagtaagtaggt-tgcagtgggtctgt
cctc
>Form_C_China
gcatgttttggtatgtgtttcgacagtcatgcatgtcatagccgtaggatggggagtgag
cttcatagcccgaccccttctcaggagcatgttttcatagcatattgagcatactggggt
ggtctcccatgtagctcgctctctcggtgcaggacctcccttttgcgggccacttcgatg
gcctccattatgga-gagcatgttttggcatatgaagcgttggtggtgaggctcgtagcc
tgtatatacataccaacgcagtcgtcatatgtctctctggcgggggtcggtcgtatgacc
ttgtccctacctggagcatgtattttgtatgat---atactggac-ggcagctgcgctgt
tcagtcgtcggccgtacctcatgtgcgcgtttccaggtccgaacgtttgacatgcaaggg
catcataccagaagcccgttttctcgtatgagcatgtaggtaggcaggctcttcagtcgt
ctacccataaatgcacttcgaagga-actttttggtattgaatggctttgtgcccagacg
ttcttgtgagcttgttctcgctctgcgatacgctggaagctgtggtattacctacagcag
acgcattcctagggcggacggcctctctgcagatgggcacatctgtagaaagaggctccc
cttgatgacgcaatgcaatggctgggtaaacggagtgctatacagaagggtcaaacctc-
tgtggaagtacacccataccagcattgtttgttgagaggggtcggctctacgagtgagga
cctatactgccatctcattcagatgcgtagacggtctttactcagactgcctttctggct
gattgtcctcggtccctcgtgcgcacgcgccgctggctgaatgggcttgtccttatggag
tgccgcgttcttc-aaagaatccgtcttcatacagcgacctcatttacatattgccggct
gaatgggcttgtccttatggagcactgcattcttctaggaattgctatctatacagcgac
ctcatttacatacaaggcacct-ttttctacatagacttgccatttggctgattgttctg
acagtcgctgcctcttcggagttctgtgtggacgagtggcc--tcctgagtgggccggtc
cttacggagtgcccacgtctgttatgactgggttctccatacagcgacctcactcagaat

tgtatgg--------------caccgtgccgat-----------tgaatggacaagcagt
ccttagggaatcgcatgaaagcatgacggc-acatggtgtatggacccgttgggccttgc
ctgtgctatcttctcgagtcgaccgg------------tcctcgtctcctcctttcgcga
cgggaagctcttgagtttaccgtgggtcaggacgctgtggtatctgtttgtctgtagtgt
gggctctctatcagaacagcgtatcagcacacgtcgtctgggtctatggtatttttgctt
aacggccctgggtttggtagtatggttccatgtcggtcctctcacctcccaaccgtcatg
tctctga-----------------------------------------------------
------------------------------------------------------------
--------------------------------------------tatcatctgcacagta
cgtcttgtcagaatatcattctgtatggcagtcgatagactaggccctatgccgcagctg
tctctgaccactatgcgagcgcatacctacctcaatgtgttcatagctgttatggatctt
gggtgtggtgtgccagaacgatacgccaccagattgtggaatggtgcttgtagtattaga
agaggacatgtatgaagagcacgttcttttctcttatgatggtgccgcgggtgcgggccg
ttctcattctgcttgtcgggtgctcaaggtgcatgtaggattgctgaacccac-tgaatg
gtcatgagaatggtcatatggtgaggttcacttctccatcgctttgtgtgtgttggatgt
ttgggtggcttcggccgcatggaaaatctcggtctgtaactcatcagggagggtttccat
gtgagcaatctctccagtcgaacacacacccatgggcgtcccgctaggcaaaatggaact
ctcgatccgacacagcctgtggagtgtttgttccgtgtggcgtgaaacggcttatactgg
atgtcttccc-tttggacaccagtctgttggtatg--tacgcataccatctagagggagc
aagaaaaaggcatgcaagtgccacaaaggaagggaggccttcctgctaccgtgtgtcc-t
cttgacgatcacgggggagtatatagagacatttggacgagatccgtttttcacgaggcg
gtgccctcgcgcaaaggtgttgctctgcacgcatctttatgatgatagttgctcgctcct
atgcgtccactcgtgaaaatacctgctctctctgtctgtttccatgacacccc-catgcg
ttgtcatagagcatgctcaggtacagaaagtgctgtgactttcttggcctgctgtcactg
agtcagcctgatactttggaggaatcttttctgggcactgcccagctgagagttcttatc
aggctctggagtgtgcatatgcctatattcattatagtgcatgtggctacgcagaaaaag
catcgacgtggaatggagaagaacaccattctgaatggaacggttcttactctacttctg
ttgcttgacgctgtatgggaatcgctttgctctgggggtacaccttccattccatgacaa
agcatttgatctatgccccga-gggtcattctctatgaagagcccgaaacttgtctgagt
gagaggcgaaggtagtccttcattt--atgcggggtccgccctcacactgtagtcggcac
agtcccttgtttatggactgcgcagctcaatcttcgactggagagg--aaaaacaaaaca
caagtatgacacgcacatacgtgttttctaggactgtgacctgtcggctgcctcgcggtg
gttggcatgatgccccaagt-gggggggggagaaacaatcagtatgattgataaaaacca
gtgggg-aaaaaggcgcgctgtcgcacacccttggtcacggcctc---aagtcgacggcc
ctcactggccca-ccctgcgactcctgttcgcttttgacaagacacacctgcaata-cgg
aggaaatgcctgagcagctgtgccgcagcaactgcggacgcgtagcgatatgcatccctg

ttgtttgcgacttggtacaaagaacgtgcgccgggagacggattgccctttgacttttcc
acccagcccgcggctctataccсttgacgtacctatttacctggttgatcctgccagtag
tcatatgcttgtctcaaagactaagccatgcatgtctaagtataaacacttttatactcg
tgaaactgcgaatggctcattaaaacagttatagttcctttgggaacaacactacttgga
taaccgtagtaattctagagctaatacatgcctcaacgcccaactcacgaagggtggtat
ttattggataaaaaaccatcgtgctctgtccagagcgctttgagatgattcacaataact
tgtcggatcgcatggcctcgtgctggcgactcctcattcaaatttctgccctatcaactt
tcgatggtagagtattggtctaccatggtgtcgacgggtgacggggaattagggttcgat
tccggagagggagcctgagaaacggctaccacatccaaggaaggcagcaggcgcgcaaat
tacccaatcccgactcggggaggtagtgacaaaaaataacaatagggggccctttgggtc
ttctaattggaatgagaacaatttaaatcccttatcgaggatccattggagggcaagtct
ggtgaacgta----------------------------------tgcatta---------
--aagccctccgtaat-aaaaaaaaacagcgccagatgtct-------------------
------------------------------------------------------------
------------------------------------------------------------
------------------------------------------------------------
------------------------------------------------------------
------------------------------------------------------------
-------ctctctggaagcacacca-----------------------------------
---tgcatggggtttgttcctctc-----------------------------tgcatgg
ccgtg-----------------------------------------tgaaaaaggcgagt
aggcaaacgatgggaaacccggcttggaatagtaaccagccacagagcccccatcataca
cttctaaattgccggggagtgcccgcgaggccttgactaccgcggtccgccgagaggtga
gagcgagcaatcgttcaggacgcagcagtgtacgtagtggtacatggatggtaaaaacgt
caaggatagggttgatccgcagggaagcgcgagtgccgccatg---------gtcggtgg
gcaaaggcgaaccttcagagactctaatggagtgggcgcaagcttaagggagagtccaat
ccactgggaaaccaggtccacagcagtaacggggtgatgctcgccccagaggactgtgga
aggcaaccccacc----aaaagggtgtgtgttgtcgggtagc-aatggccagcagccgcg
gtaattccagctccaatagcgtatattaaagttgttgcagttaaaacgctcgtagtcgga
cctcggagcgcccccacgggccgggcttttgccttgttgccagtgggagtttgtctcctt
ttgtcgggacgcggtgtgcagggcctttactgtctctgcgcgtttacctgcgggccgacc
gtttactgtgaagaaagtaaagtgttcaaagcaggccaatgccttgaatatgtgagcatg
gaataatagaataggacttgggctctattttgttggtttccagtgaccaagtaatgatta
atagggacggttgggggcattcgtatttcattgtcagaggtgaaattcttggattgatgg
aagacgtacaactgcgaaagcatctgccatggatgttttcattgatcaagaacgaaagtt
aggggatcgaagacgatcagataccgtcgtagtcttaaccataaacgatgccgactgggg

attggcgggggcacttttatatgactctgtcagcaccctgagggaaaccaaagtctttgg
gttctgggggggagtatggtcgcaaggctgaaacttaaaggaattgacggaagggcaccac
caggagtggagcctgcggcttaatttgactcaacacgggaaaacttaccaggtccggaca
gaagaatgattgacagactgaaagctctttcttgatttttggttggtggtgcatggccg
ttcttagttggtggagtgatttgtctggttaattccgttaacgaacgagacctcgtcctg
ctaaataggtgcgcgcatgccagcactgcgttcttaccttcttagagggactatgcgcgt
ctagcgtatggaagattgaggcaataacaggtctgtgatgcccttagatgttctgggccg
cacgcgcgctacactgatgcattcaacgagtttcacttgatagtcctgggtcggaaggct
cgggtaatcttttgaaagtgcatcgtgctggggatagatcattgcaattattgatcttca
acgaggaattccttgtaggcgcaggtcatcagcctgcgccgaatacgtccctgccctttg
tacacaccgcccgtcgctcctaccgattgaatggtccgatgaaatgtccggatcgcggct
agtgtgatggactcgtccagagcggtagccgtgagaagctcattaaatcttatcatttag
aggaaggagaagtcgtaacaaggt------------------------------------
------------------------------------------------------------
------------------------------------------------------------
------------------------------------------------------------
------------------------------------------------------------
------------------------------------------------------------
------------------------------------------------------------
------------------------------------------------------------
------------------------------------------------------------
------------------------------------------------------------
------------------------------------------------------------
------------------------------------------------------------
------------------------------------------------------------
------------------------------------------------------------
------------------------------------------------------------
------------------------------------------------------------
-------------------------------ttccgtaggtgaacctgcggaaggatcat
tcacaaactattgacacaaacacacgcgaaccaaatcgtccgcacagcaggtggcaatga
aagagagactctgcatgtcgcctttcgggg--tagcaagcagcactcttctgccatcgcc
tctgtgcagggcgtaaattcccattgagaggatgtgagggcaccacaggaagcttttcca
caggaagtcgccatccttctccctccacggcggcg----cttctgtgcttggcag-----
-----ttttttttgccttctagggaggatgccgccaatggagccccttat-atatataca
tcatcatatgccc--cttttttttcttaaccgcttgccacagcttcttctatgaggagctt
gtgggaagactgtctccatacaataac-aaagatacaactcttagcggtggatatcttgg

ttctcgcaacgatgaagaacgcagccaactgcgataactaatgtgaattgcaggacttcg
tgaatcattaagtctttgaacgcaagttgcgctcatctccatgaggaggtgagcatgcct
gtttgagtgtcgatgggaaaaacaatac-ataacaacaatcggctctcttatcttcctgc
cttcttgggggcaaaaggtagtgcatgccggagatgggctttgtcgcccctttctcttga
aagaggcgtcgtccgaaacacgtatcgttgcccgctggaggtcatagagcatgagttt--
aaagcaccacatctctcaagtcagagtgag-cgtcttgagtaggtcc-gcattgattcag
tttcaacccaggtggcgttggttacttgccat-aagcctctctggggtttctttgcttgg
acctctcttgtcagtttctctgacgtagagacaggt--gccgtctcatggacgttgccct
ctggaacgtgcta-aaaaaaaaaaaaaga----------aaaaaccgtaccacatgtcaa
aagaggcagatcgtgcttgacggcgggaggtgttgcatttcttgcttgtcctgagcgtct
gccagtctct----ga--cgagtg----tgttccttccataaacccgatctcaaatcagg
taggaccacccgctgaatttaagcatataattaagcggaggaaaagaaactaacaaggat
tccctcagtagcggcgagcgaagcgggaagagctcaacatgaaaaccgttggggggttcgc
gctttccttcgggttgtatagtaaagtggtgtcgcctgcgtcgcgccgggtataaatccc
gtctgaaatggggtgtcgtagagggttatatccccgtctgtgacccggtgtgcgatgcgt
tacggtgaccttacgatgagtcgggttgcttgggcatgcagccctaagtgggtggtaaac
ttcatctaaagctaaatactagcgagagaccgatagcgtacaagtaccgtaagggaaaga
tgcaaagaactttgaaaagagagttaaacagggcttgaaaccgttaagagtgaagcgaag
gaggtcagtgatcatggggccagatccccaagtgcctgtcata-gggcaccgtagcgttc
tggtttcccggggcagcgccggccgggggcgggggttaacagggaggtggttgagggatt
cgttttccgcctgtactcctttatagcttgcaccccggccgagggatccgcgacttgtcg
tgtgagggcgctgtcaaaatggccttctccaccccgtcttgaaacacggaccaaggagtc
taacatactcgcgagtgtttgggtggttaaaacccgaacgcacaatgaaagtaaacggtg
gctctgcgccgctgcttggtgggaaccctgcttttataatggggggcactatcacccgac
cccgatccctttggggtggttggttttgaggtagagcgagtatgatgggacccgaaagat
ggtgaactattcctgagtagggcgaagccagaggaaactctggtggaggctcgtagcgat
gctgacgtgcaaatcgctcgtcgaacttgggtataggggcgaaagactaatcgaaccatc
tagtagctggttccctccgaagtttccctcaggatagcgggagcaaatgcaagtcagtat
catccggtaaagcgaatgattagaggcctcgggggagtaactccttcgacctattctcaa
actttaaataggtgaggagccggcaacgttgcttaaaacaatgaacgccgtgcggacaat
gattgctctcagtgggccgattttggtaagcagaactggcgatgcgggatgaaccgaaag
cgacgctaaggtgccaaactcccagctcattcagacaccagaaaaggtgttgattcatac
agacagcaggacggtggccatggaagtcggaatccgctaaggagtgtgtaacaactcacc
tgccgaatggatcagccctgaaaatggatggcgcttaagcgggggaccgaggcgtcgccg
ttggggtaaatacgaagccccaatgagtaggagggcgtgggggttttgaaacacggtaca
ctcgcatggaagccttgggcgtaagctcgggtggagcgacccctagtgtagatcttggtg

gtagtagcaaatattcaagtgagaaccttgaagactgaagtggggaagggttccacgcga

acagtgcttggacgtgggttagtcggtcctaagagatggggaaactccgtattcaagtgc

gcttttgtagcgcagtccatcgaaagggaagcgggttaacattcccgcaccaggatgtgg

aaactgtggtttaccgcaaactgcggcaacgcaactgaacctggggacgcccgtaggagc

tccgagcagggatctctttcctgtttaacgcgcccaacccccctggaatcggtttgtccgg

agataggggggcgaaaggccggaaaagcaccgcacgtcttgtggtgttcggaacgcttgta

atggcccttgaaaacccaggggaaagcagccgtcttacac--ttgtgtgagacgaggaat

tctcacacctggccgtacccatcaccgcatcaggtccccaaggtgaacagcctctagtca

atagaataatgtagataagggaagtcggcaaaatagatccgtaactttgggaaaaggatt

ggctctaatgttcgggcttgtcgggacgcgcgggcgtagatgccaatgagcggcagacgt

ctagcctctttggaggctggattaacgctgcttgtgggaagaaaccgtgtcgatggaggc

aggaacgcttatcgatcttgtctcttcccgcttgcgatcaacgaacggattagaactggt

acggacaaggggaatccgactgtttaattaaaacaaagcattgcgacggccggaaacggt

gttggcgcaatgtgatttctgcccagtgctctgaatgtcaaagtgaagaaattcaaccaa

gcgcgggtaaacggcgggagtaactatgactct--taaac-aaaccatactacctac---

-------------------gggagtcaaaagtgggggcggcgtctggtgtagaaatacac

gtac--------------------------------------------------------

-------------------gacacgctttttttcctgctagtcgatctgacctttgttct

ttggactgtgg-------------------------------------------------

--------------------------gcggtcagcggcaacataacc-------------

------------------------------------------------------------

------------------------------------------------------------

----------------------------tggtacggg---------------------ga

aggcctcgacgttga---------------------------------------------

--------------------------------gtgctgcatcttggaagtatc-------

-----------------------------ttgacacaataagaga---------------

------------------------------aaagagattgcgcttaatagg---------

----gggttgatctcgt-------------------------------------------

-------------------------------------------ggcgagccgacgc----

---------------------------------cggtgtgaactggt-------------

------------------------------------------------------------

-----------------------ggcaat-------------------------------

----ggccgtcgtagagcacgcaaaggtatgggctgactttttctcctca----aaagtt

ggcttaagggac-gtgccatcccacacgataataaagtgtgcagtctg------------

--------------------taagagctcccacagggcaaagac---------agactgg

gatgcagacggagacggcaaaaacaacaacatccgttctgcttctgggcgcaaatgctta

aggtagccaaatgcctcgtcatctaaactaattgg-at----------------------
-----------------------gagatagtc----------tgagtgaaaagcttatgc
ag-----------------gctagtaccggaa-aaaaaaaaaaaaagaggggctctgcca
ct----ctgac------------------------ggtgcgacactccttgtacagggaa
accctcc--gcatggaaaggcacgtggtaccaccctcgcggggaaactcgcgagcgggat
cccgggtaatgaccggagccgatggtaaaaactcacgtgaccgatggggtgaccctgtgg
cgagacgctaagttgtgggacatcaccctctcacaatacgcatccgtcgcagagactgaa
cgggagtgggctccgttga-----cggccggccggccagccagccgtcttcca-------
---ggggcttaagatacagtccgggcctcagggatgg----aaaaacctggggaacta-c
cagaattagtgacgcgcatgaatggattaacgagattcccactgtccctatctactatct
agcgaaaccacagccaagggaacgggcttggaagaatcagcggggaaagaagaccctgtt
gagcttgactctagtccgactttgtgagatgagttgggaggtgtagtctaagagggaggg
gcaactcgaccgtgaaatacctctactcccaagcttgttttacttactccggcgagcagg
gatgggtggtaacaccctttttctaggattaacacctcttcgggggtggacctgcgcgga
ggacattgtcaggtggggagtttggctggggcggcacatctgttaaaagataacgcaggt
gtcctaagatgagctcagtgagaacagaaatctcacgtagaacaaaagggtaaaagctca
tttgattttgattttcagtacgaatacaaactgtgaaagcatggcctatcgatcctttttg
accttcaaggtttgaagcaagaggtgtcagaaaagttaacacgcgttgtggtagctttta
agagtggactcaaagtgttccgctagtccccccctcggaccgccacttacaaaaggcctgg
ggatgggggcgacactgtcaaattgcggggactccctaaagcctgtctctaccaagccag
cgcggcgaaagcccctggtggccggggtaatgacctagggtatggtaaaagcgagacagg
atg-gaacaatgggcaatccgcagccaagtcctaagg----ctttttgaaaagcggctat
ggatgcagttcaacgactagatggcagtgggctgagagagcccaccgcggcaccctttcg
gcttaagatatagtcttgcccacgtgaaagcgtgcctgcagatgaacgccaccatgtatg
gcaggagctgcaggaggccgctaaacataccagcggctgggaacataacagtaccacagg
gatgaaccatacatagaattagccagtttggtgagacaataattgtgcacgaacgcggta
ttgtccactttgtgcggtgtcctgctcggggccgtccgtgtgcgtcatggctgacagaga
acgccctcctaagcgagtggcgttcgccgttgcgtcggcaaccaacagtgtgcatgtgca
gatacattcaggtggaagctcacaggagtccacggactcttcgtttactcctcacagcga
tcacccggatgcgggaatgagtggtactgaagacagcggcgatgaagacggcgaagagag
tgttgaagaggcccctttggcgcgcaccacggtattcactcaaatcttggaaaacaccga
ggttgccaaacaagaactcttcaatgttactgagaatcgctcacacacggagggctggtg
ctgcatttctgatcttgcgccctctaagaagaatggatacgtccaagtgtctattcaggg
ccataatcacttaatggttatgcagaacctagtcctggctgcgagcggctactcaagcgc
tgtgtacgatgtgaacgttcaggcgagccacctttgcgacaatcgtcgtt---gttgcac
gctaggacacattgttccagagtcggccaaactcaacaatagccgtaaaggctgccgtgg

gtgggtaccttgcccacacacggcgtgcaccactcctatcttcatttgtccgcactctcc
gccttgtataaggtaccactcggcgtacgccgacatgcaggccttgatcgacgcgcacac
agacggtcataagtaaggcacagtagtttcgtggcctatcgctggagtgctgtgtgtttc
atcttctttttctggaactatcgtcccaggcgggtgtccctctctcacccgtctagtccg
gccttggtgcgcaaaga---cgtgcgtggatgcaccttggttaaggaaaatgctcctagt
agttttacaaggacacttcttgtagggcaacacatctggatgcgaagaactcccggaaaa
gacaacggtaccaagtgactggggtaacgccccattcatgaaggcaaataatcaaaaaga
ggtgccgggtaagaacccgttgtatagggacaatccgcagctgacttggctccctg--cc
cat--ccatgggcatagcttagcggttcaacgctcggtaagatgtgggttggcgcgctgt
ccgacagcgcactggcttaaggtacgggctactcccaccggagacggtgtctttgtctac
ctcgggtaacacgcctcgggagaagcaaagagggtgcaatggaaaaaaaactggttgcgg
ggaaatccctaccagagtcacctggtacaaaggaactggcttgtggcagccaagcgttca
tagcgacgttgctttttgatccttcgatgtcggctcttcctatcattgcgaagcagaatt
cgccaagtgttggattgttcacccactaatagggaacgtgagctgggtagccactttta--
-----tgcccaacagctggatagctgggcttctgtctta-ggccgggacttctcacgacc
cgtgccttggtcaccc-aaaaaagggcgaccgcagagctcaggatggagcttctggctta
actagtagaggacagatagagatcgtcaaattgcgggaaaccccttagagcctcgagcta
ccaagtgcgtgtcgaaaggcatagacgcagccagcgatggctgtcgcatggcttga-ttc
catcaagataggtaaaaatgctcgaggattgggcaatccgcagccaagcccctaaggt--
---ccccccgcgggctatggggaaggttcaacgactgtaaggcgtaccgtgcaagcggaa
gatacagtctagccccgctggaaagcagcgccaaagtaagtaggt-tgcagtgggtctgt
cctc
>Form_B_China_SRR9587932
gcatgttttggtatgtgtttcgacagtcatgcatgtcatagccgtaggatggggagtgag
cttcatagcccgaccccttctcaggagcatgttttcatagcatattgagcatactggggt
ggtctcccatgtagctcgctctctcggtgcaggacctcccttttgcgggccacttcgatg
gcctccattatgga-gagcatgttttggcatatgaagcgttggtggtgaggctcgtagcc
tgtatatacataccaacgcagtcgtcatatgtctctctggcgggggtcggtcgtatgacc
ttgtccctacctggagcatgtattttgtatgat---atactggac-ggcagctgcgctgt
tcagtcgtcggccgtacctcatgtgcgcgtttccaggtccgaacgtttgacatgcaaggg
catcataccagaagcctgttttctcgtatgagcatgtaggtaggcaggctcttcagtcgt
ctacccataaatgcacttcgaagga-actttttggtattgaatggctttgtgcccagacg
ttcttgtgagcttgttctcgctctgcgatacgctggaagctgtggtattacctacagcag
acgcattcctagggcggacggcctctctgcagatgggcacatctgtagaaagaggctccc
cttgatgacgcaatgcaatggctgggtaaacggagtgctatacagaagggtcaaacctc-
tgtggaagtacacccataccagcattgtttgttgagaggggtcggctctacgagtgagga

cctatactgccatctcattcagatgcgtagacggtctttactcagactgcctttctggct
gattgtcctcggtccctcgtgcgcacgcgccgctggctgaatgggcttgtccttatggag
tgccgcgttcttc-aaagaatccgtcttcatacagcgacctcatttacatattgccggct
gaatgggcttgtccttatggagcactgcattcttctaggaattgctatctatacagcgac
ctcatttacatacaaggcacct-ttttctacatagacttgccatttggctgattgttctg
acagtcgctgcctcttcggagttctgtgtggacgagtggcc--tcctgagtgggccggtc
cttacggagtgcccacgtctgttatgactgggttctccatacagcgacctcactcagaat
tgtatgg--------------caccgtgccgat-----------tgaatggacaagcagt
ccttagggaatcgcatgaaagcatgacggc-acatggtgtatggacccgttgggccttgc
ctgtgctatcttctcgagtcgaccgg------------tcctcgtctcctcctttcgcga
cgggaagctcttgagtttaccgtgggtcaggacgctgtggtatctgtttgtctgtagtgt
gggctctctatcagaacagcgtatcagcacacgtcgtctgggtctatggtatttttgctt
aacggccctgggtttggtagtatggttccatgtcggtcctctcacctcccatccgtcatg
tctctga-----------------------------------------------------
------------------------------------------------------------
--------------------------------------------tatcatctgcacagta
cgtcttgtcagaatatcattctgtatggcagtcgatagactaggccctatgccgcagctg
tctctgaccactatgcgagcgcatacctacctcaatgtgttcatagctgttatggatctt
gggtgtggtgtgccagaacgatacgccaccagattgtggaatggtgcttgtagtattaga
agaggacatgtatgaagagcacgttcttttctcttatgatggtgccgcgggtgggggccg
ttctcattctgcttgtcgggtgctcaaggtgcatgtaggattgctgaacccac-tgaatg
gtcatgagaatggtcatatggtgaggttcacttctccatcgctttgtgtgtgttggatgt
ttgggtggcttcggccgcatggaaaatctcggtctgtaactcatcagggagggtttccat
gtgagcaatctctccagtcgaacacacacccatgggcgtcccgctaggcaaaatggaact
ctcgatccgacacagcctgtggagtgtctgttccgtgtggcgtgaaacggcttatactgg
atgtcttccc-tttggacaccagtctgttggtatg--tacgcataccatctagagggagc
aagaaaaaggcatgcaagtgccacaaaggaagggaggccttcctgctaccgtgtgtcc-t
cttgacgatcacgggggagtatatagagacatttggacgagatcctttttttcacgaggcg
gtgccctcgcgcaaaggtgttgctctgcacgcatctttatgatgatagttgctcgctcct
atgcgtccactcgtgaaaatacctgctctctctgtctgtttccatgacacccc-catgcg
ttgtcatagagcacgctcaggtacagaaagtgctgtgactttcttggcctgctgtcactg
agtcagcctgatactttggaggaatcttttctgggcactgcccagctgagagttcttatc
aggctctggagtgtgcatatgcctatattcattatagtgcatgtggctacgcagaaaaag
catcgacgtggaatggagaagaacaccattctgaatggaacggttcttactctacttctg
ttgcttgacgctgtatgggaatcgctttgctctgggggtacaccttccattccatgacaa
agcatttgatctatgccccga-gggtcattctctatgaagagcccgaaacttgtctgagt

gagaggcgaaggtagtccttcattt--atgcggggtccgccctcacactgtagtcggcac

agtcccttgtttatggactgcgcagctcaatcttcgactggagagg--aaaaacaaaaca

caagtatgacacgcacatacgtgttttctaggactgtgacctgtcggctgcctcgcggtg

gttggcatgatgccccaagt-ggggggggagaaacaatcagtatgattgataaaaacca

gtgggg-aaaaaggcgcgctgtcgcacacccttggtcacggcctc---aagtcgacggcc

ctcactggccca-ccctgcgactcctgttcgcttttgacaagacacacctgcaata-cgg

aggaaatgcctgagcagctgtgccgcagcaactgcggacgcgtagcgatatgcatccctg

ttgtttgcgacttggtacaaagaacgtgcgccgggagacggattgccctttgacttttcc

acccagcccgcggctctatacccttgacgtacctatttacctggttgatcctgccagtag

tcatatgcttgtctcaaagactaagccatgcatgtctaagtataaacacttttatactcg

tgaaactgcgaatggctcattaaaacagttatagttcctttgggaacaacactacttgga

taaccgtagtaattctagagctaatacatgcctcaacgcccgactcacgaagggtggtat

ttattggataaaaaaccatcgtgctctgtccagagcgctttgagatgattcacaataact

tgtcggatcgcatggcctcgtgctggcgactcctcattcaaatttctgccctatcaactt

tcgatggtagagtattggtctaccatggtgtcgacgggtgacggggaattagggttcgat

tccggagagggagcctgagaaacggctaccacatccaaggaaggcagcaggcgcgcaaat

tacccaatcccgactcggggaggtagtgacaaaaaataacaataggggccctttgggtc

ttctaattggaatgagaacaatttaaatcccttatcgaggatccattggagggcaagtct

ggtgaacgta----------------------------------tgcatta---------

--aagccctccgtaat-aaaaaaaaacagcgccagatgtct-------------------

------------------------------------------------------------

------------------------------------------------------------

------------------------------------------------------------

------------------------------------------------------------

------------------------------------------------------------

-------ctctctggaagcacacca-----------------------------------

---tgcatggggtttgttcctctc-----------------------------tgcatgg

ccgtg-----------------------------------------tgaaaaaggcgagt

aggcaaacgatgggaaacccggcttggaatagtaaccagccacagagcccccatcataca

cttctaaattgccggggagtgcccgcgaggccttgactaccgcggtccgccgagaggtga

gagcgagcaatcgttcaggacgcagcagtgtacgtagtggtacatggatggtaaaaacgt

caaggatagggttgatccgcagggaagcgcgagtgccgccatg---------gtcggtgg

gcaaaggcgaaccttcagagactctaatggagtgggcgcaagcttaagggagagtccaat

ccactgggaaaccaggtccacagcagtaacggggtgatgctcgccccagaggactgtgga

aggcaaccccacc----aaaagggtgtgtgttgtcgggtagc-aatggccagcagccgcg

gtaattccagctccaatagcgtatattaaagttgttgcagttaaaacgctcgtagtcgga

cctcggagcgcccccacgggccgggcttttgccttgttgccagtgggagtttgtctcctt
ttgtcgggacgcggtgtgcagggcctttactgtctctgcgcgtttacctgcgggccgacc
gtttactgtgaagaaagtaaagtgttcaaagcaggccaatgccttgaatatgtgagcatg
gaataatagaataggacttgggctctattttgttggtttccagtgaccaagtaatgatta
atagggacggttgggggcattcgtatttcattgtcagaggtgaaattcttggattgatgg
aagacgtacaactgcgaaagcatctgccatggatgttttcattgatcaagaacgaaagtt
aggggatcgaagacgatcagataccgtcgtagtcttaaccataaacgatgccgactgggg
attggcgggggcacttttatatgactctgtcagcaccctgagggaaaccaaagtctttgg
gttctgggggggagtatggtcgcaaggctgaaacttaaaggaattgacggaagggcaccac
caggagtggagcctgcggcttaatttgactcaacacgggaaaacttaccaggtccggaca
gaagaatgattgacagactgaaagctctttcttgatttttggttggtggtgcatggccg
ttcttagttggtggagtgatttgtctggttaattccgttaacgaacgagacctcgtcctg
ctaaataggtgcgcgcatgccagcactgcgttcttaccttcttagagggactatgcgcgt
ctagcgtatggaagattgaggcaataacaggtctgtgatgcccttagatgttctgggccg
cacgcgcgctacactgatgcattcaacgagtttcacttgatagtcctgggtcggaaggct
cgggtaatcttttgaaagtgcatcgtgctggggatagatcattgcaattattgatcttca
acgaggaattccttgtaggcgcaggtcatcagcctgcgccgaatacgtccctgccctttg
tacacaccgcccgtcgctcctaccgattgaatggtccgatgaaatgtccggatcgcggct
agtgtgatggactcgtccagagcggtagccgtgagaagctcattaaatcttatcatttag
aggaaggagaagtcgtaacaaggt------------------------------------
------------------------------------------------------------
------------------------------------------------------------
------------------------------------------------------------
------------------------------------------------------------
------------------------------------------------------------
------------------------------------------------------------
------------------------------------------------------------
------------------------------------------------------------
------------------------------------------------------------
------------------------------------------------------------
------------------------------------------------------------
------------------------------------------------------------
------------------------------------------------------------
------------------------------------------------------------
------------------------------------------------------------
-------------------------------ttccgtaggtgaacctgcggaaggatcat

tcacaaactattgacacaaacacacgcgaaccaaatcgtccgcacagcaggtggcaatga
aagagagactctgcatgtcgcctttcgggg--tagcaagcagcactcttctgccatcgcc
tctgtgcagggcgtaaattcccattgagaggatgtgagggcaccacaggaagcttttcca
caggaagtcgccatccttctccctccacggcggcg----cttctgtgcttggcag-----
-----ttttttttgccttctagggaggatgccgccaatggagccccttat-atatataca
tcatcatatgccc--cttttttttcttaaccgcttgccacagcttcttctatgaggagctt
gtgggaagactgtctccatacaataac-aaagatacaactcttagcggtggatatcttgg
ttctcgcaacgatgaagaacgcagccaactgcgataactaatgtgaattgcaggacttcg
tgaatcattaagtctttgaacgcaagttgcgctcatctccatgaggaggtgagcatgcct
gtttgagtgtcgatgggaaaaacaatac-ataacaacaatcggctctcttatcttcctgc
cttcttgggggcaaaaggtagtgcatgccggagatgggctttgtcgcccctttctcttga
aagaggcgtcgtccgaaacacgtatcgttgcccgctggaggtcatagagcatgagttt--
aaagcaccacatctctcaagtcagagtgag-cgtcttgagtaggtcc-gcattgattcag
tttcaacccaggtggcgttggttacttgccat-aagcctctctggggtttctttgcttgg
acctctcttgtcagtttctctgacgtagagacaggt--gccgtctcatggacgttgccct
ctggaacgtgcta-aaaaaaaaaaaaaga----------aaaaaccgtaccacatgtcaa
aagaggcagatcgtgcttgacggcgggaggtgttgcatttcttgcttgtcctgagcgtct
gccagtctct----ga--cgagtg----tgttccttccataaacccgatctcaaatcagg
taggaccacccgctgaatttaagcatataattaagcggaggaaaagaaactaacaaggat
tccctcagtagcggcgagcgaagcgggaagagctcaacatgaaaaccgttgggggttcgc
gctttccttcgggttgtatagtaaagtggtgtcgcctgcgtcgcgccgggtataaatccc
gtctgaaatggggtgtcgtagagggttatatccccgtctgtgacccggtgtgcgatgcgt
tacggtgaccttacgatgagtcgggttgcttgggcatgcagccctaagtgggtggtaaac
ttcatctaaagctaaatactagcgagagaccgatagcgtacaagtaccgtaagggaaaga
tgcaaagaactttgaaaagagagttaaacagggcttgaaaccgttaagagtgaagcgaag
gaggtcagtgatcatggggccagatccccaagtgcctgtcata-gggcaccgtagcgttc
tggtttcccggggcagcgccggccgggggcgggggttaacagggaggtggttgagggatt
cgttttccgcctgtactcctttatagcttgcaccccggccgagggatccgcgacttgtcg
tgtgagggcgctgtcaaaatggccttctccaccccgtcttgaaacacggaccaaggagtc
taacatactcgcgagtgtttgggtggttaaaacccgaacgcacaatgaaagtaaacggtg
gctctgcgccgctgcttggtgggaaccctgcttttataatggggggcactatcacccgac
cccgatccctttggggtggttggttttgaggtagagcgagtatgatgggacccgaaagat
ggtgaactattcctgagtagggcgaagccagaggaaactctggtggaggctcgtagcgat
gctgacgtgcaaatcgctcgtcgaacttgggtataggggcgaaagactaatcgaaccatc
tagtagctggttccctccgaagtttccctcaggatagcgggagcaaatgcaagtcagtat
catccggtaaagcgaatgattagaggcctcgggggagtaactccttcgacctattctcaa

actttaaataggtgaggagccggcaacgttgcttaaaacaatgaacgccgtgcggacaat

gattgctctcagtgggccgattttggtaagcagaactggcgatgcgggatgaaccgaaag

cgacgctaaggtgccaaactcccagctcattcagacaccagaaaaggtgttgattcatac

agacagcaggacggtggccatggaagtcggaatccgctaaggagtgtgtaacaactcacc

tgccgaatggatcagccctgaaaatggatggcgcttaagcgggggaccgaggcgtcgccg

ttggggtaaatacgaagccccaatgagtaggagggcgtgggggttttgaaacacggtaca

ctcgcatggaagccttgggcgtaagctcgggtggagcgacccctagtgtagatcttggtg

gtagtagcaaatattcaagtgagaaccttgaagactgaagtggggaagggttccacgcga

acagtgcttggacgtgggttagtcggtcctaagagatggggaaactccgtattcaagtgc

gcttttgtagcgcagtccatcgaaagggaagcgggttaacattcccgcaccaggatgtgg

aaactgtggtttaccgcaaactgcggcaacgcaactgaacctggggacgcccgtaggagc

tccgagcagggatctctttcctgtttaacgcgcccaacccccctggaatcggtttgtccgg

agatagggggcgaaaggccggaaaagcaccgcacgtcttgtggtgttcggaacgcttgta

atggcccttgaaaacccaggggaaagcagccgtcttacac--ttgtgtgagacgaggaat

tctcacacctggccgtacccatcaccgcatcaggtccccaaggtgaacagcctctagtca

atagaataatgtagataagggaagtcggcaaaatagatccgtaactttgggaaaaggatt

ggctctaatgttcgggcttgtcgggacgcgcgggcgtagatgccaatgagcggcagacgt

ctagcctctttggaggctggattaacgctgcttgtgggaagaaaccgtgtcgatggaggc

aggaacgcttatcgatcttgtctcttcccgcttgcgatcaacgaacggattagaactggt

acggacaaggggaatccgactgtttaattaaaacaaagcattgcgacggccggaaacggt

gttggcgcaatgtgatttctgcccagtgctctgaatgtcaaagtgaagaaattcaaccaa

gcgcgggtaaacggcgggagtaactatgactct--taaac-aaaccatactacctac---

-------------------gggagtcaaaagtgggggcggcgtctggtgtagaaatacac

gtac--------------------------------------------------------

-------------------gacacgctttttttcctgctagtcgatctgacctttgttct

ttggactgtgg-------------------------------------------------

--------------------------gcggtcagcggcaacataacc-------------

------------------------------------------------------------

------------------------------------------------------------

----------------------------tggtacggg---------------------ga

aggcctcgacgttga---------------------------------------------

--------------------------------gtgctgcatcttggaagtatc-------

-----------------------------ttgacacaataagaga---------------

------------------------------aaagagattgcgcttaatagg---------

----gggttgatctcgt-------------------------------------------

-------------------------------------------ggcgagccgacgc----

---------------------------------cggtgtgaactggt-------------
------------------------------------------------------------
-----------------------ggcaat-------------------------------
----ggccgtcgtagagcacgcaaaggtatgggctgactttttctcctca----aaagtt
ggcttaagggac-gtgccatcccacacgataataaagtgtgcagtctg------------
--------------------taagagctcccacagggcaaagac---------agactgg
gatgcagacggagacggcaaaaacaacaacatccgttctgcttctgggcgcaaatgctta
aggtagccaaatgcctcgtcatctaaactaattgg-at----------------------
-----------------------gagatagtc----------tgagtgaaaagcttatgc
ag-----------------gctagtaccggaa-aaaaaaaaaaaaagaggggctctgcca
ct----ctgac------------------------ggtgcgacactccttgtacagggaa
accctcc--gcatggaaaggcacgtggtaccaccctcgcggggaaactcgcgagcgggat
cccgggtaatgaccggagccgatggtaaaaactcacgtgaccgatggggtgaccctgtgg
cgagacgctaagttgtgggacatcaccctctcacaatacgcatccgtcgcagagactgaa
cgggagtgggctccgttga-----cggccggccggccagccagccgtcttcca-------
---ggggcttaagatacagtccgggcctcagggatgg----aaaaacctggggaacta-c
cagaattagtgacgcgcatgaatggattaacgagattcccactgtccctatctactatct
agcgaaaccacagccaagggaacgggcttggaagaatcagcggggaaagaagaccctgtt
gagcttgactctagtccgactttgtgagatgagttgggaggtgtagtctaagagggaggg
gcaactcgaccgtgaaatacctctactcccaagcttgttttacttactccggcgagcagg
gatgggtggtaacaccctttttctaggattaacacctcttcgggggtggacctgcgcgga
ggacattgtcaggtggggagtttggctggggcggcacatctgttaaaagataacgcaggt
gtcctaagatgagctcagtgagaacagaaatctcacgtagaacaaaagggtaaaagctca
tttgattttgattttcagtacgaatacaaactgtgaaagcatggcctatcgatccttttg
accttcaaggtttgaagcaagaggtgtcagaaaagttaacacgcgttgtggtagctttta
agagtggactcaaagtgttccgctagtcccccctcggaccgccacttacaaaaggcctgg
ggatgggggcgacactgtcaaattgcggggactccctaaagcctgtctctaccaagccag
cgcggcgaaagcccctggtggccggggtaatgacctagggtatggtaaaagcgagacagg
atg-gaacaatgggcaatccgcagccaagtcctaagg----ctttttgaaaagcggctat
ggatgcagttcaacgactagatggcagtgggctgagagagcccaccgcggcacccctttcg
gcttaagatatagtcttgcccacgtgaaagcgtgcctgcagatgaacgccaccatgtatg
gccggagctgcaggagaccgc--acttgatgtgcagtcgggaat-cgacagtaccacagg
gatgaac-----------------------------------------------------
------------------------------------------------------------
------------------------------------------------------------
------------------------------------------------------------

------------------------------------------------------------

-------------------------------------------------aaaaacac---

------------------------------------------------------------

------------------------------------------------------------

--acaatta---------------------------------------------------

----------------------------------------atttgtggcc---atcatac

----------------------------------------agtcgta-------------

------------------------------------------------------------

------------------------------------------------------------

------------------------------------------------------------

------------tgatattatcgtcccaggcgggtgcgcctctctcacccgtctagtccg

gccttggtgcgcaaaaaatgcttgcgtggatgcgccttggttaaggaaaatgctcctagt

agttttacaaggacacttcttgtagggcaacacatctggatgcgaagaactcccggaaaa

gacaacggtaccaagtgactggggtaacgccccattcatgaaggcaaataatcaaaaaga

ggtgccgggtaagaacccgttgtatagggacaatccgcagctgacttggctccctg--cc

cat--ccatgggcatagcttagcggttcaacgctcggtaagatgtgggttggcgcgctgt

ccgacagcgcactggcttaaggtacgggctactcccaccggagacggtgtctttgtctac

ctcgggtaacacgcctcgggagaagcaaagagggtgcaatgg-aaaaaaactggttgcgg

ggaaatccctaccagagtcacctggtacaaaggaactggcttgtggcagccaagcgttca

tagcgacgttgctttttgatccttcgatgtcggctcttcctatcattgcgaagcagaatt

cgccaagtgttggattgttcacccactaatagggaacgtgagctgggtagccacttta--

-----tgcccaacagctggatagctgggcttctgtctta-ggccgggacttctcacgacc

cgtgccttggtcaccc-aaaaaagggcgaccgcagagctcaggatggagcttctggctta

actagtagaggacagatagagatcgtcaaattgcgggaaaccccttagagcctcgagcta

ccaagtgcgtgtcgaaaggcatagacgcagccagcgatggctgtcgcatggcttga-ttc

catcaagataggtaaaaatgctcgaggattgggcaatccgcagccaagcccctaaggt--

---cccccgcgggctatggggaaggttcaacgactgtaaggcgtaccgtgcaagcggaa

gatacagtctagccccgctggaaagcagcgccaaagtaagtaggt-tgcagtgggtctgt

cctc

>Form_B_China_SRR9587955

gcatgttttggtatgtgtttcgacagtcatgcatgtcatagccgtaggatggggagtgag

cttcatagcccgaccccttctcaggagcatgttttcatagcatattgagcatactggggt

ggtctcccatgtagctcgctctctcggtgcaggacctcccttttgcgggccacttcgatg

gcctccattatgga-gagcatgttttggcatatgaagcgttggtggtgaggctcgtagcc

tgtatatacataccaacgcagtcgtcatatgtctctctggcgggggtcggtcgtatgacc

ttgtccctacctggagcatgtattttgtatgat---atactggac-ggcagctgcgctgt

tcagtcgtcggccgtacctcatgtgcgcgtttccaggtccgaacgtttgacatgcaaggg
catcataccagaagcctgttttcttgtatgagcatgtaggtaggcaggctcttcagtcgt
ctacccataaatgcacttcgaagga-actttttggtattgaatggctttgtgcccagacg
ttcttgtgagcttgttctcgctctgcgatacgctggaagctgtggtattacctacagcag
acgcattcctagggcggacggcctctctgcagatgcgcacatctgtagaaagaggctccc
cttgatgacgcaatgcaatggctgggtaaacggagtgctatacagaagggtcaaacctc-
tgtggaagtacactcataccagcattgtttgttgagaggggtcggctctacgagtgagga
cctatactgccatctcattcagatgcgtagacggtctttactcagactgcctttctggct
gattgtcctcggtccctcgtgcgcacgcgccgctggctgaatgggcttgtccttatggag
tgccgcgttcttc-aaagaatccgtcttcatacagcgacctcatttacatattgccggct
gaatgggcttgtccttatggagcactgcattcttctaggaattgctatctatacagcgac
ctcatttacatacaaggcacct-ttttctacatagacttgccatttggctgattgttctg
acagtcgctgcctcttcggagttctgtgtggacgagtggcc--tcctgagtgggccggtc
cttacggagtgcccacgtctgttatgactgggttctccatacagcgacctcactcagaat
tgtatgg--------------caccgtgccgat-----------tgaatggacaagcagt
ccttagggaatcgcatgaaagcatgacggc-acatggtgtatggacccgttgggccttgc
ctgtgctatcttctcgagtcgaccgg------------tcctcgtctcctcctttcgcga
cgggaagctcttgagtttaccgtgggtcaggacgctgtggtatctgtttgtctgtagtgt
gggctctctatcagaacagcgtatcagcacacgtcgtctgggtctatggtatttttgctt
aacggccctgggtttggtagtatggttccatgtcggtcctctcacctcccaaccgtcatg
tctctga-----------------------------------------------------
------------------------------------------------------------
--------------------------------------------tatcatctgcacagta
cgtcttgtcagaatatcattctgtatggcagtcgatagactaggccctatgccgcagctg
tctctgaccactatgcgagcgcatacctacctcaatgtgttcatagctgttatggatctt
gggtgtggtgtgccagaacgatacgccaccagattgtggaatggtgcttgtagtattaga
agaggacatgtatgaagagcacgttcttttctcttatgatggtgccgcgggtgcgggccg
ttctcattctgcttgtcgggtgctcaaggtgcatgtaggattgctgaacccac-tgaatg
gtcatgagaatggtcatatggtgaggttcacttctccatcgctttgtgtgtgttggatgt
ttgggtggcttcggccgcatggaaaatctcggtctgtaactcatcagggagggtttccat
gtgagcaatctctccagtcgaacacacacccatgggcgtcccgctaggcaaaatggaact
ctcgatccgacacagcctgtggagtgtttgttccgtgtggcgtgaaacggcttatactgg
atgtcttccc-tttggacaccagtctgttggtatg--tacgcataccatctagagggagc
aagaaaaaggcatgcaagtgccacaaaggaagggaggccttcctgctaccgtgtgtcc-t
cttgacgatcacgggggagtatatagagacatttggacgagatccgtttttcacgaggcg
gtgccctcgcgcaaaggtgttgctctgcacgcatctttatgatgatagttgctcgctcct

atgcgtccactcgtgaaaatacctgctctctctgtctgtttccatgacacccc-catgcg
ttgtcatagagcatgctcaggtacagaaagtgctgtgactttcttggcctgctgtcactg
agtcagcctgatactttggaggaatcttttctgggcactgcccagctgagagttcttatc
aggctctggagtgtgcatatgcctatattcattatagtgcatgtggctacgcagaaaaag
catcgacgtggaatggagaagaacaccattctgaatggaacggttcttactctacttctg
ttgcttgacgctgtatgggaatcgctttgctctgggggtacaccttccattccatgacaa
agcatttgatctatgccccga-gggtcattctctatgaagagcccgaaacttgtctgagt
gagaggcgaaggtagtccttcattt--atgcggggtccgccctcacactgtagtcggcac
agtcccttgtttacggactgcgcagctcaatcttcgactggagagg--aaaaacaaaaca
caagtatgacacgcacatacgtgttttctaggactgtgacctgtcggctgcctcgcggtg
gttggcatgatgccccaagt-gggggggggagaaacaatcagtatgattgataaaaacca
gtgggg-aaaaaggcgcgctgtcgcacacccttggtcacggcctc---aagtcgacggcc
ctcactggccca-ccctgcgactcctgttcgcttttgacaagacacacctgcaata-cgg
aggaaatgcctgagcagctgtgccgcagcaactgcggacgcgtagcgatatgcatccctg
ttgtttgcgacttggtacaaagaacgtgcgccgggagacggattgccctttgacttttcc
acccagcccgcggctctatacccttgacgtacctatttacctggttgatcctgccagtag
tcatatgcttgtctcaaagactaagccatgcatgtctaagtataaacacttttatactcg
tgaaactgcgaatggctcattaaaacagttatagttcctttgggaacaacactacttgga
taaccgtagtaattctagagctaatacatgcctcaacgcccgactcacgaagggtggtat
ttattggataaaaaaccatcgtgctctgtccagagcgctttgagatgattcacaataact
tgtcggatcgcatggcctcgtgctggcgactcctcattcaaatttctgccctatcaactt
tcgatggtagagtattggtctaccatggtgtcgacgggtgacggggaattagggttcgat
tccggagagggagcctgagaaacggctaccacatccaaggaaggcagcaggcgcgcaaat
tacccaatcccgactcggggaggtagtgacaaaaaataacaatagggggccctttgggtc
ttctaattggaatgagaacaatttaaatcccttatcgaggatccattggagggcaagtct
ggtgaacgta----------------------------------tgcatta---------
--aagccctccgtaat-aaaaaaaaacagcgccagatgtct-------------------
------------------------------------------------------------
------------------------------------------------------------
------------------------------------------------------------
------------------------------------------------------------
------------------------------------------------------------
-------ctctctggaagcacacca-----------------------------------
---tgcatggggtttgttcctctc-----------------------------tgcatgg
ccgtg-----------------------------------------tgaaaaaggcgagt
aggcaaacgatgggaaacccggcttggaatagtaaccagccacagagcccccatcataca

cttctaaattgccggggagtgcccgcgaggccttgactaccgcggtccgccgagaggtga

gagcgagcaatcgttcaggacgcagcagtgtacgtagtggtacatggatggtaaaaacgt

caagtatagggttgatccgcagggaagcgcgagtgccgccatg---------gtcggtgg

gcaaaggcgaaccttcagagactctaatggagtgggcgcaagcttaagggagagtccaat

ccactgggaaaccaggtccacagcagtaacggggtgatgctcgccccagaggactgtgga

aggcaaccccacc----aaaagggtgtgtgttgtcgggtagc-aatggccagcagccgcg

gtaattccagctccaatagcgtatattaaagttgttgcagttaaaacgctcgtagtcgga

cctcggagcgcccccacgggccgggcttttgccttgttgccagtgggagtttgtctcctt

ttgtcgggacgcggtgtgcagggcctttactgtctctgcgcgtttacctgcgggccgacc

gtttactgtgaagaaagtaaagtgttcaaagcaggccaatgccttgaatatgtgagcatg

gaataatagaataggacttgggctctattttgttggtttccagtgaccaagtaatgatta

atagggacggttgggggcattcgtatttcattgtcagaggtgaaattcttggattgatgg

aagacgaacaactgcgaaagcatctgccatggatgttttcattgatcaagaacgaaagtt

aggggatcgaagacgatcagataccgtcgtagtcttaaccataaacgatgccgactgggg

attggcgggggcacttttatatgactctgtcagcaccctgagggaaaccaaagtctttgg

gttctgggggggagtatggtcgcaaggctgaaacttaaaggaattgacggaagggcaccac

caggagtggagcctgcggcttaatttgactcaacacgggaaaacttaccaggtccggaca

gaagaatgattgacagactgaaagctctttcttgatttttttggttggtggtgcatggccg

ttcttagttggtggagtgatttgtctggttaattccgttaacgaacgagacctcgtcctg

ctaaataggtgcgcgcatgccagcactgcgttcttaccttcttagagggactatgcgcgt

ctagcgtatggaagattgaggcaataacaggtctgtgatgcccttagatgttctgggccg

cacgcgcgctacactgatgcattcaacgagtttcacttgatagtcctgggtcggaaggct

cgggtaatcttttgaaagtgcatcgtgctggggatagatcattgcaattattgatcttca

acgaggaattccttgtaggcgcaggtcatcagcctgcgccgaatacgtccctgccctttg

tacacaccgcccgtcgctcctaccgattgaatggtccgatgaaatgtccggatcgcggct

agtgtgatggactcgtccagagcggtagccgtgagaagctcattaaatcttatcatttag

aggaaggagaagtcgtaacaaggt------------------------------------

------------------------------------------------------------

------------------------------------------------------------

------------------------------------------------------------

------------------------------------------------------------

------------------------------------------------------------

------------------------------------------------------------

------------------------------------------------------------

------------------------------------------------------------

------------------------------------------------------------

------------------------------------------------------------
------------------------------------------------------------
------------------------------------------------------------
------------------------------------------------------------
------------------------------------------------------------
------------------------------------------------------------
-------------------------------ttccgtaggtgaacctgcggaaggatcat
tcacaaactattgacacaaacacacgcgaaccaaatcgtccgcacagcaggtggcaatga
aagagagactctgcatgtcgcctttcgggg--tagcaagcagcactcttctgccatcgcc
tctgtgcagggcgtaaattcccattgagaggatgtgagggcaccacaggaagcttttcca
caggaagtcgccatccttctccctccacggcggcg----cttctgtgcttggcag-----
-----ttttttttgccttctagggaggatgccgccaatggagccccttat-atatataca
tcatcatatgccc--cttttttttcttaaccgcttgccacagcttcttctatgaggagctt
gtgggaagactgtctccatacaataac-aaagatacaactcttagcggtggatatcttgg
ttctcgcaacgatgaagaacgcagccaactgcgataactaatgtgaattgcaggacttcg
tgaatcattcagtctttgaacgcaagttgcgctcatctccatgaggaggtgagcatgcct
gtttgagtgtcgatgggaaaaacaatac-ataacaacaatcggctctcttatcttcctgc
cttcttgggggcaaaaggtagtgcatgccggagatgggctttgtcgcccctttctcttga
aagaggcgtcgtccgaaacacgtatcgttgcccgctggaggtcatagagcatgagttt--
aaagcaccacatctctcaagtcagagtgag-cgtcttgagtaggtcc-gcattgattcag
tttcaacccaggtggcgttggttacttgccat-aagcctctctggggtttctttgcttgg
acctctcttgtcagtttctctgacgtagagacaggt--gccgtctcatggacgttgccct
ctggaacgtgcta-aaaaaaaaaaaaaga----------aaaaaccgtaccacatgtcaa
aagaggcagatcgtgcttgacggcgggaggtgttgcatttcttgcttgtcctgagcgtct
gccagtctct----ga--cgagtg----tgttccttccataaacccgatctcaaatcagg
taggaccacccgctgaatttaagcatataattaagcggaggaaaagaaactaacaaggat
tccctcagtagcggcgagcgaagcgggaagagctcaacatgaaaaccgttgggggttcgc
gctttccttcgggttgtatagtaaagtggtgtcgcctgcgtcgcgccgggtataaatccc
gtctgaaatggggtgtcgtagagggttatatccccgtctgtgacccggtgtgcgatgcgt
tacggtgaccttacgatgagtcgggttgcttgggcatgcagccctaagtgggtggtaaac
ttcatctaaagctaaatactagcgagagaccgatagcgtacaagtaccgtaagggaaaga
tgcaaagaactttgaaaagagagttaaacagggcttgaaaccgttaagagtgaagcgaag
gaggtcagtgatcatggggccagatccccaagtgcctgtcata-gggcaccgtagcgttc
tggtttcccggggcagcgccggccgggggcgggggttaacagggaggtggttgagggatt
cgttttccgcctgtactcctttatagcttgcaccccggccgagggatccgcgacttgtcg
tgtgagggcgctgtcaaaatggccttctccaccccgtcttgaaacacggaccaaggagtc

taacatactcgcgagtgtttgggtggttaaaacccgaacgcacaatgaaagtaaacggtg
gctctgcgccgctgcttggtgggaaccctgcttttataatggggggcactatcacccgac
cccgatccctttggggtggttggttttgaggtagagcgagtatgatgggacccgaaagat
ggtgaactattcctgagtagggcgaagccagaggaaactctggtggaggctcgtagcgat
gctgacgtgcaaatcgctcgtcgaacttgggtatagggggcgaaagactaatcgaaccatc
tagtagctggttccctccgaagtttccctcaggatagcgggagcaaatgcaagtcagtat
catccggtaaagcgaatgattagaggcctcgggggagtaactccttcgacctattctcaa
actttaaataggtgaggagccggcaacgttgcttaaaacaatgaacgccgtgcggacaat
gattgctctcagtgggccgattttggtaagcagaactggcgatgcgggatgaaccgaaag
cgacgctaaggtgccaaactcccagctcattcagacaccagaaaaggtgttgattcatac
agacagcaggacggtggccatggaagtcggaatccgctaaggagtgtgtaacaactcacc
tgccgaatggatcagccctgaaaatggatggcgcttaagcgggggaccgaggcgtcgccg
ttggggtaaatacgaagccccaatgagtaggagggcgtgggggttttgaaacacggtaca
ctcgcatggaagccttgggcgtaagctcgggtggagcgacccctagtgtagatcttggtg
gtagtagcaaatattcaagtgagaaccttgaagactgaagtggggaagggttccacgcga
acagtgcttggacgtgggttagtcggtcctaagagatggggaaactccgtattcaagtgc
gcttttgtagcgcagtccatcgaaagggaagcgggttaacattcccgcaccaggatgtgg
aaactgtggtttaccgcaaactgcggcaacgcaactgaacctggggacgcccgtaggagc
tccgagcagggatctctttcctgtttaacgcgcccaacccccctggaatcggtttgtccgg
agatagggggcgaaaggccggaaaagcaccgcacgtcttgtggtgttcggaacgcttgta
atggcccttgaaaacccaggggaaagcagccgtcttacac--ttgtgtgagacgaggaat
tctcacacctggccgtacccatcaccgcatcaggtccccaaggtgaacagcctctagtca
atagaataatgtagataagggaagtcggcaaaatagatccgtaactttgggaaaaggatt
ggctctaatgttcgggcttgtcgggacgcgcgggcgtagatgccaatgagcggcagacgt
ctagcctctttggaggctggattaacgctgcttgtgggaagaaaccgtgtcgatggaggc
aggaacgcttatcgatcttgtctcttcccgcttgcgatcaacgaacggattagaactggt
acggacaaggggaatccgactgtttaattaaaacaaagcattgcgacggccggaaacggt
gttggcgcaatgtgatttctgcccagtgctctgaatgtcaaagtgaagaaattcaaccaa
gcgcgggtaaacggcgggagtaactatgactct--taaac-aaaccatactacctac---
---------------------gggagtcaaaagtgggggcggcgtctggtgtagaaatacac
gtac---------------------------------------------------------
---------------------gacacgcttttttcctgctagtcgatctgacctttgttct
ttggactgtgg--------------------------------------------------
-----------------------------gcggtcagcggcaacataacc-----------
-------------------------------------------------------------
-------------------------------------------------------------

----------------------------tggtacggg---------------------ga
aggcctcgacgttga---------------------------------------------
--------------------------------gtgctgcatcttggaagtatc-------
-----------------------------ttgacacaataagaga---------------
------------------------------aaagagattgcgcttaatagg---------
----gggttgatctcgt-------------------------------------------
-------------------------------------------ggcgagccgacgc----
---------------------------------cggtgtgaactggt-------------
------------------------------------------------------------
-----------------------ggcaat-------------------------------
----ggccgtcgtagagcacgcaaaggtatgggctgactttttctcctca----aaagtt
ggcttagggac-gtgccatcccacacgataataaagtgtgcagtctg-------------
--------------------taagagctcccacagggcaaagac---------agactgg
gatgcagacggagacggcaaaaacaacaacatccgttctgcttctgggcgcaaatgctta
aggtagccaaatgcctcgtcatctaaactaattgg-at----------------------
-----------------------gagatagtc----------tgagtgaaaagcttatgc
ag-----------------gctagtaccgg---aaaaaaaaaaaaagaggggctctgcca
ct----ctgac------------------------ggtgcgacactccttgtacagggaa
accctcc--gcatggaaaggcacgtggtaccaccctcgcggggaaactcgcgagcgggat
cccgggtaatgaccggagccgatggtaaaaactcacgtgaccgatggggtgaccctgtgg
cgagacgctaagttgtgggacatcaccctctcacaatacgcatccgtcgcagagactgaa
cgggagtgggctccgttga-----cggccggccggccagccagccgtcttcca-------
---ggggcttaagatacagtccgggcctcagggatgg----aaaaacctggggaacta-c
cagaattagtgacgcgcatgaatggattaacgagattcccactgtccctatctactatct
agcgaaaccacagccaagggaacgggcttggaagaatcagcggggaaagaagaccctgtt
gagcttgactctagtccgactttgtgagatgagttgggaggtgtagtctaagagggaggg
gcaactcgaccgtgaaatacctctactcccaagcttgttttacttactccggcgagcagg
gatgggtggtaacacccttttttctaggattaacacctcttcgggggtggacctgcgcgga
ggacattgtcaggtggggagtttggctggggcggcacatctgttaaaagataacgcaggt
gtcctaagatgagctcagtgagaacagaaatctcacgtagaacaaaagggtaaaagctca
tttgattttgattttcagtacgaatacaaactgtgaaagcatggcctatcgatccttttg
accttcaaggtttgaagcaagaggtgtcagaaaagttaacacgcgttgtggtagctttta
agagtggactcaaagtgttccgctagtccccccctcggaccgccacttacaaaaggcctgg
ggatgggggcgacactgtcaaattgcggggactccctaaagcctgtctctaccaagccag
cgcggcgaaagcccctggtggccggggtaatgacctagggtatggtaaaagcgagacagg
atg-gaacaatgggcaatccgcagccaagtcctaagg----ctttttgaaaagcggctat

ggatgcagttcaacgactagatggcagtgggctgagagagcccaccgcggcaccctttcg
gcttaagatatagtcttgcccacgtgaaagcgtgcctgcagatgaacgccaccatgtatg
gccggagctgcaggagaccgc--acttgatgtgcagtcgggaat-cgacagtaccacagg
gatgaac-----------------------------------------------------
------------------------------------------------------------
------------------------------------------------------------
------------------------------------------------------------
------------------------------------------------------------
-------------------------------------------------aaaaacac---
------------------------------------------------------------
------------------------------------------------------------
------------------------------------------------------------
--------------------------------------acaattgtggcc---atcatac
----------------------------------------agtcgta-------------
------------------------------------------------------------
------------------------------------------------------------
------------------------------------------------------------
------------tgatattatcgtcccaggcgggtgcgcctctctcacccgtctagtccg
gccttggtgcgcaaaaaatgcttgcgtggatgcgccttggttaaggaaaatgctcctagt
agttttacaaggacacttcttgtagggcaacacatctggatgcgaagaactcccggaaaa
gacaacggtaccaagtgactggggtaacgccccattcatgaaggcaaataatcaaaaaga
ggtgccgggtaagaacccgttgtatagggacaatccgcagctgacttggctccctg--cc
catccccatgggcatagcttagcggttcaacgctcggtaagatgtgggttggcgcgctgt
ccgacagcgcactggcttaaggtacgggctactcccaccggagacggtgtctttgtctac
ctcgggtaacacgcctcgggagaagcaaagagggtgcaatgg-aaaaaaactggttgcgg
ggaaatccctaccagagtcacctggtacaaaggaactggcttgtggcagccaagcgttca
tagcgacgttgctttttgatccttcgatgtcggctcttcctatcattgcgaagcagaatt
cgccaagtgttggattgttcacccactaatagggaacgtgagctgggtagccacttta--
-----tgcccaacagctggatagctgggcttctgtctta-ggccgggacttctcacgacc
cgtgccttggtcaccc-aaaaaagggcgaccgcagagctcaggatggagcttctggctta
actagtagaggacagatagagatcgtcaaattgcgggaaaccccttagagcctcgagcta
ccaagtgcgtgtcgaaaggcatagacgcagccagcgatggctgtcgcatggcttga-ttc
catcaagataggtaaaaatgctcgaggattgggcaatccgcagccaagcccctaaggt--
---ccccccgcgggctatggggaaggttcaacgactgtaaggcgtaccgtgcaagcggaa
gatacagtctagccccgctggaaagcagcgccaaagtaagtaggt-tgcagtgggtctgt
cctc

>Form_M_China
gcatgttttggtatgtgtttcgacagtcatgcatgtcatagccgtaggatggggagtgag
cttcatagcccgaccccttctcaggagcatgttttcatagcatattgagcatactggggt
ggtctcccatgtagctcgctctctcggtgcaggacctcccttttgcgggccacttcgatg
gcctccattatgga-gagcatgttttggcatatgaagcgttggtggtgaggctcgtagcc
tgtatatacataccaacgcagtcgtcatatgtctctctggcgggggtcggtcgtatgacc
ttgtccctacctggagcatgtattttgtatgat---atactggac-ggcagctgcgctgt
tcagtcgtcagccgtacctcatgtgcgcgtttccaggtccgaacgtttgacatgcaaggg
catcataccagaagcctgttttctcgtatgagcatgtaggtaggcaggctcttcagtcgt
ctacccataaatgcacttcgaagga-actttttggtattgaatggctttgtgcccagacg
ttcttgtgagcttgttctcgctctgcgatacgctggaagctgtggtattacctacagcag
acgcattcctagggcggacggcctctctgcagatgggcacatctgtagaaagaggctccc
cttgatgacgcaatgcaatggctgggtaaacggagtgctatacagaagggtcaaacctc-
tgtggaagtacactcataccagcattgtttgttgagaggggtcggctctacgagtgagga
cctatactgccatctcattcagatgcgtagacggtctttactcagactgcctttctggct
gattgtcctcggtccctcgtgcgcacgcgccgctggctgaatgggcttgtccttatggag
tgccgcgttcttc-aaagaatccgtcttcatacagcgacctcatttacatattgccggct
gaatgggcttgtccttatggagcactgcattcttctaggaattgctatctatacagcgac
ctcatttacatacaaggcacct-ttttctacatagacttgccatttggctgattgttctg
acagtcgctgcctcttcggagttctgtgtggacgagtggcc--tcctgagtgggccggtc
cttacggagtgcccacgtctgttatgactgggttctccatacagcgacctcactcagaat
tgtatgg--------------caccgtgccgat-----------tgaatggacaagcagt
ccttagggaatcgcatgaaagcatgacggc-acatggtgtatggacccgttgggccttgc
ctgtgctatcttctcgagtcgaccgg------------tcctcgtctcctcctttcgcga
cgggaagctcttgagtttaccgtgggtcaggacgctgtggtatctgtttgtctgtagtgt
gggctctctatcagaacagcgtatcagcacacgtcgtctgggtctatggtatttttgctt
aacggccctgggtttggtagtatggttccatgtcggtcctctcacctcccaaccgtcatg
tctctga-----------------------------------------------------
------------------------------------------------------------
--------------------------------------------tatcatctgcacagta
cgtcttgtcagaatatcattctgtatggcagtcgatagactaggccctatgccgcagctg
tctctgaccactatgcgagcgcatacctacctcaatgtgttcatagctgttatggatctt
gggtgtggtgtgccagaacgatacgccaccagattgtggaatggtgcttgtagtattaga
agaggacatgtatgaagagcacgttcttttctcttatgatggtgccgcgggtgcgggccg
ttctcattctgcttgtcgggcgctcaaggtgcatgtaggattgctgaacccac-tgaatg
gtcatgagaatggtcatatggtgaggttcacttctccatcgctttgtgtgtgttggatgt

ttgggtggcttcggccgcatggaaaatctcggtctgtaactcatcagggagggtttccat
gtgagcaatctctccagtcgaacacacacccatgggcgtcccgctaggcaaaatggaact
ctcgatccgacacagcctgtggagtgtttgttccgtgtggcgtgaaacggcttatactgg
atgtcttccc-tttggacaccagtctgttggtatg--tacgcataccatctagagggagc
aagaaaaaggcatgcaagtgccacaaaggaagggaggccttcctgctaccgtgtgtcc-t
cttgacgatcacgggggagtatatagagacatttggacgagatccgtttttcacgaggcg
gtgccctcgcgcaaaggtgttgctctgcacgcatctttatgatgatagttgctcgctcct
atgcgtccactcgtgaaaatacctgctctctctgtctgtttccatgacacccc-catgcg
ttgtcatagagcatgctcaggtacagaaagtgctgtgactttcttggcctgctgtcactg
agtcagcctgatactttggaggaatcttttctgggcactgcccagctgagagttcttatc
aggctctggagtgtgcatatgcctatattcattatagtgcatgtggctacgcagaaaaag
catcgacgtggaatggagaagaacaccattctgaatggaacggttcttactctacttctg
ttgcttgacgctgtatgggaatcgctttgctctgggggtacaccttccattccatgacaa
agcatttgatctatgccccga-gggtcattctctatgaagagcccgaaacttgtctgagt
gagaggcgaaggtagtccttcattt--atgcggggtccgccctcacactgtagtcggcac
agtcccttgtttatggactgcgcagctcaatcttcgactggagagg--aaaaacaaaaca
caagtatgacacgcacatacgtgttttctaggactgtgacctgtcggctgcctcgcggtg
gttggcatgatgccccaagt-gggggggggagaaacaatcagtatgattgataaaaacca
gtgggg-aaaaaggcgcgctgtcgcacacccttggtcacggcctc---aagtcgacggcc
ctcactggccca-ccctgcgactcctgttcgcttttgacaagacacacctgcaata-cgg
aggaaatgcctgagcagctgtgccgcagcaactgcggacgcgtagcgatatgcatccctg
ttgtttgcgacttggtacaaagaacgtgcgccgggagacggattgccctttgacttttcc
acccagcccgcggctctatacccttgacgtacctatttacctggttgatcctgccagtag
tcatatgcttgtctcaaagactaagccatgcatgtctaagtataaacacttttatactcg
tgaaactgcgaatggctcattaaaacagttatagttcctttgggaacaacactacttgga
taaccgtagtaattctagagctaatacatgcctcaacgcccgactcacgaagggtggtat
ttattggataaaaaaccatcgtgctctgtccagagcgctttgagatgattcacaataact
tgtcggatcgcatggcctcgtgctggcgactcctcattcaaatttctgccctatcaactt
tcgatggtagagtattggtctaccatggtgtcgacgggtgacggggaattagggttcgat
tccggagagggagcctgagaaacggctaccacatccaaggaaggcagcaggcgcgcaaat
tacccaatcccgactcggggaggtagtgacaaaaaataacaatagggggccctttgggtc
ttctaattggaatgagaacaatttaaatcccttatcgaggatccattggagggcaagtct
ggtgaacgta----------------------------------tgcatta---------
--aagccctccgtaat-aaaaaaaaacagcgccagatgtct-------------------
------------------------------------------------------------
------------------------------------------------------------

------------------------------------------------------------
------------------------------------------------------------
------------------------------------------------------------
-------ctctctggaagcacacca-----------------------------------
---tgcatggggtttgttcctctc-----------------------------tgcatgg
ccgtg-----------------------------------------tgaaaaaggcgagt
aggcaaacgatgggaaacccggcttggaatagtaaccagccacagagcccccatcataca
cttctaaattgccggggagtgcccgcgaggccttgactaccgcggtccgccgagaggtga
gagcgagcaatcgttcaggacgcagcagtgtacgtagtggtacatggatggtaaaaacgt
caaggatagggttgatccgcagggaagcgcgagtgccgccatg---------gtcggtgg
gcaaaggcgaaccttcagagactctaatggagtgggcgcaagcttaagggagagtccaat
ccactgggaaaccaggtccacagcagtaacggggtgatgctcgccacagaggactgtgga
aggcaaccccacc----aaaagggtgtgtgttgtcgggtagc-aatggccagcagccgcg
gtaattccagctccaatagcgtatattaaagttgttgcagttaaaacgctcgtagtcgga
cctcggagcgcccccacgggccgggcttttgccttgttgccagtgggagtttgtctcctt
ttgtcgggacgcggtgtgcagggcctttactgtctctgcgcgtttacctgcgggccgacc
gtttactgtgaagaaagtaaagtgttcaaagcaggccaatgccttgaatatgtgagcatg
gaataatagaataggacttgggctctattttgttggtttccagtgaccaagtaatgatta
atagggacggttgggggcattcgtatttcattgtcagaggtgaaattcttggattgatgg
aagacgtacaactgcgaaagcatctgccatggatgttttcattgatcaagaacgaaagtt
aggggatcgaagacgatcagataccgtcgtagtcttaaccataaacgatgccgactgggg
attggcgggggcacttttatatgactctgtcagcaccctgagggaaaccaaagtctttgg
gttctgggggagtatggtcgcaaggctgaaacttaaaggaattgacggaagggcaccac
caggagtggagcctgcggcttaatttgactcaacacgggaaaacttaccaggtccggaca
gaagaatgattgacagactgaaagctctttcttgatttttggttggtggtgcatggccg
ttcttagttggtggagtgatttgtctggttaattccgttaacgaacgagacctcgtcctg
ctaaataggtgcgcgcatgccagcactgcgttcttaccttcttagagggactatgcgcgt
ctagcgtatggaagattgaggcaataacaggtctgtgatgcccttagatgttctgggccg
cacgcgcgctacactgatgcattcaacgagtttcacttgatagtcctgggtcggaaggct
cgggtaatcttttgaaagtgcatcgtgctggggatagatcattgcaattattgatcttca
acgaggaattccttgtaggcgcaggtcatcagcctgcgccgaatacgtccctgccctttg
tacacaccgcccgtcgctcctaccgattgaatggtccgatgaaatgtccggatcgcggct
agtgtgatggactcgtccagagcggtagccgtgagaagctcattaaatcttatcatttag
aggaaggagaagtcgtaacaaggtttccgtattttgtgggtggataccgattcaataaca
tctgccgtttttctcaatgtactatatccctagtctgcttgcccagttcttggagtctcg
tgcgatgtgtgctattacaggcacgctgggtcatgaagacagacaaggttaggatcttca

cggaagtacgcgcaataggagcgcgtcttgttcaacgcacctgactcaagagtcatatgc
gagggacggacgcagcgcttgtcgccgcacagatgggaagcttcctctccaggcatagga
tggcgctgcatttgcatcatcgctgctagtaagtggcaataatacttgattccccgccag
cgcacttgtacgtagccgttggcggcgggaacgtagttggtgatgtgacagcctcgagcg
ttcacctctgtgtgtccgccaatctgcacaaagatgaacgccatatcggagacaggggga
ggtgccattg---tgatgatgacgatgttgttgggtggtccttccgtaggaaaacgtaca
cgaagcctttgcggcccgaaagggtgcacagtgcgcgactataaacactgattctgtggt
caacgcaagcagcggttctgtggaagaacagctgcgactttctcaaattgcgggaacacc
ctaaagtcaacaagcaccgcggcttgacgtgaaaagcgtcaggcgcagcagtcggggtag
cgccctggcggatggtaatagccttgttgaatggcacaatgggcaatccgcagccaagcc
cctaaggtcccccgctgggctatggggaaggttcacagactgtaagggagggggtttctt
ccatga--aagaagcttaagagacagtcggtcccctgcgagagcagtgtcgcatgtgagg
aaggcacccatgaaaagtgggtgctggagagccatgtgagccctctatgagggcgggaag
ttttccga---tacaatgagaaacaaacacgttccgtaggtgaacctgcggaaggatcat
tcacaaactattgacacaaacacacgcgaaccaaatcgtccgcacagcaggtggcaatga
aagagagactctgcatgtcgcctttcgggg--tagcaagcagcactcttctgccatcgcc
tctgtgcagggcgtaaattcccattgagaggatgtgagggcaccacaggaagcttttcca
caggaagtcgccatccttctccctccacggcggcg----cttctgtgcttggcag-----
-----ttttttttgccttctagggaggatgccgccaatggagccccttat-atatataca
tcatcatatgccc--ctttttttcttaaccgcttgccacagcttcttctatgaggagctt
gtgggaagactgtctccatacaataac-aaagatacaactcttagcggtggatatcttgg
ttctcgcaacgatgaagaacgcagccaactgcgataactaatgtgaattgcaggacttcg
tgaatcattaagtctttgaacgcaagttgcgctcatctccatgaggaggtgagcatgcct
gtttgagtgtcgatgggaaaaacaatac-ataacaacaatcggctctcttatcttcctgc
cttcttgggggcaaaaggtagtgcatgccggagatgggctttgtcgcccctttctcttga
aagaggcgtcgtccgaaacacgtatcgttgcccgctggaggtcatagagcatgagttt--
aaagcaccacatctctcaagtcagagtgag-cgtcttgagtaggtcc-gcattgattcag
tttcaacccaggtggcgttggttacttgccat-aagcctctctggggtttctttgcttgg
acctctcttgtcagtttctctgacgtagagacaggt--gccgtctcatggacgttgccct
ctggaacgtgcta-aaaaaaaaaaaaaga----------aaaaaccgtaccacatgtcaa
aagaggcagatcgtgcttgacggcgggaggtgttgcatttcttgcttgtcctgagcgtct
gccagtctct----ga--cgagtg----tgttccttccataaacccgatctcaaatcagg
taggaccacccgctgaatttaagcatataattaagcggaggaaaagaaactaacaaggat
tccctcagtagcggcgagcgaagcgggaagagctcaacatgaaaaccgttgggggttcgc
gctttccttcgggttgtatagtaaagtggtgtcgcctgcgtcgcgccgggtataaatccc
gtctgaaatggggtgtcgtagagggttatatccccgtctgtgacccggtgtgcgatgcgt

tacggtgaccttacgatgagtcgggttgcttgggcatgcagccctaagtgggtggtaaac
ttcatctaaagctaaatactagcgagagaccgatagcgtacaagtaccgtaagggaaaga
tgcaaagaactttgaaaagagagttaaacagggcttgaaaccgttaagagtgaagcgaag
gaggtcagtgatcatggggccagatccccaagtgcctgtcata-gggcaccgtagcgttc
tggtttcccggggcagcgccggccgggggcgggggttaacagggaggtggttgagggatt
cgttttccgcctgtactcctttatagcttgcaccccggccgagggatccgcgacttgtcg
tgtgagggcgctgtcaaaatggccttctccaccccgtcttgaaacacggaccaaggagtc
taacatactcgcgagtgtttgggtggttaaaacccgaacgcacaatgaaagtaaacggtg
gctctgcgccgctgcttggtgggaaccctgcttttataatgggggcactatcacccgac
cccgatccctttggggtggttggttttgaggtagagcgagtatgatgggacccgaaagat
ggtgaactattcctgagtagggcgaagccagaggaaactctggtggaggctcgtagcgat
gctgacgtgcaaatcgctcgtcgaacttgggtataggggcgaaagactaatcgaaccatc
tagtagctggttccctccgaagtttccctcaggatagcgggagcaaatgcaagtcagtat
catccggtaaagcgaatgattagaggcctcgggggagtaactccttcgacctattctcaa
actttaaataggtgaggagccggcaacgttgcttaaaacaatgaacgccgtgcggacaat
gattgctctcagtgggccgattttggtaagcagaactggcgatgcgggatgaaccgaaag
cgacgctaaggtgccaaactcccagctcattcagacaccagaaaaggtgttgattcatac
agacagcaggacggtggccatggaagtcggaatccgctaaggagtgtgtaacaactcacc
tgccgaatggatcagccctgaaaatggatggcgcttaagcgggggaccgaggcgtcgccg
ttggggtaaatacgaagccccaatgagtaggagggcgtgggggttttgaaacacggtaca
ctcgcatggaagccttgggcgtaagctcgggtggagcgacccctagtgtagatcttggtg
gtagtagcaaatattcaagtgagaaccttgaagactgaagtggggaagggttccacgcga
acagtgcttggacgtgggttagtcggtcctaagagatggggaaactccgtattcaagtgc
gcttttgtagcgcagtccatcgaaagggaagcgggttaacattcccgcaccaggatgtgg
aaactgtggtttaccgcaaactgcggcaacgcaactgaacctggggacgcccgtaggagc
tccgagcagggatctctttcctgtttaacgcgcccaacccccctggaatcggtttgtccgg
agataggggcgaaaggccggaaaagcaccgcacgtcttgtggtgttcggaacgcttgta
atggcccttgaaaacccaggggaaagcagccgtcttacac--ttgtgtgagacgaggaat
tctcacacctggccgtacccatcaccgcatcaggtccccaaggtgaacagcctctagtca
atagaataatgtagataagggaagtcggcaaaatagatccgtaactttgggaaaaggatt
ggctctaatgttcgggcttgtcgggacgcgcgggcgtagatgccaatgagcggcagacgt
ctagcctctttggaggctggattaacgctgcttgtgggaagaaaccgtgtcgatggaggc
aggaacgcttatcgatcttgtctcttcccgcttgcgatcaacgaacggattagaactggt
acggacaaggggaatccgactgtttaattaaaacaaagcattgcgacggccggaaacggt
gttggcgcaatgtgatttctgcccagtgctctgaatgtcaaagtgaagaaattcaaccaa
gcgcgggtaaacggcgggagtaactatgactct--taaac-aaaccatactacctac---

-------------------gggagtcaaaagtgggggcggcgtctggtgtagaaatacac
gtac--------------------------------------------------------
-------------------gacacgctttttttcctgctagtcgatctgacctttgttct
ttggactgtgg-------------------------------------------------
--------------------------gcggtcagcggcaacataacc-------------
------------------------------------------------------------
------------------------------------------------------------
----------------------------tggtacggg---------------------ga
aggcctcgacgttga---------------------------------------------
--------------------------------gtgctgcatcttggaagtatc-------
-----------------------------ttgacacaataagaga---------------
------------------------------aaagagattgcgcttaatagg---------
----gggttgatctcgt-------------------------------------------
-------------------------------------------ggcgagccgacgc----
---------------------------------cggtgtgaactggt-------------
------------------------------------------------------------
-----------------------ggcaat-------------------------------
----ggccgtcgtagagcacgcaaaggtatgggctgactttttctcctca----aaagtt
ggcttaagggac-gtgccatcccacacgataataaagtgtgcagtctg------------
--------------------taagagctcccacagggcaaagac---------agactgg
gatgcagacggagacggcaaaaacaacaacatccgttctgcttctgggcgcaaatgctta
aggtagccaaatgcctcgtcatctaaactaattgg-at----------------------
-----------------------gagatagtc----------tgagtgaaaagcttatgc
ag-----------------gctagtaccgg---aaaaaaaaaaaaagaggggctctgcca
ct----ctgac------------------------ggtgcgacactccttgtacagggaa
accctcc--gcatggaaaggcacgtggtaccaccctcgcggggaaactcgcgagcgggat
cccgggtaatgaccggagccgatggtaaaaactcacgtgaccgatggggtgaccctgtgg
cgagacgctaagttgtgggacatcaccctctcacaatacgcatccgtcgcagagactgaa
cgggagtgggctccgttga-----cggccggccggccagccagccgtcttcca-------
---ggggcttaagatacagtccgggcctcagggatgg----aaaaacctggggaacta-c
cagaattagtgacgcgcatgaatggattaacgagattcccactgtccctatctactatct
agcgaaaccacagccaagggaacgggcttggaagaatcagcggggaaagaagaccctgtt
gagcttgactctagtccgactttgtgagatgagttgggaggtgtagtctaagagggaggg
gcaactcgaccgtgaaatacctctactcccaagcttgttttacttactccggcgagcagg
gatgggtggtaacacccttttttctaggattaacacctcttcgggggtggacctgcgcgga
ggacattgtcaggtggggagtttggctggggcggcacatctgttaaaagataacgcaggt

gtcctaagatgagctcagtgagaacagaaatctcacgtagaacaaaagggtaaaagctca
tttgattttgattttcagtacgaatacaaactgtgaaagcatggcctatcgatccttttg
accttcaaggtttgaagcaagaggtgtcagaaaagt------------------------
------------------------------------------------------------
------------------------------------------------------------
------------------------------------------------------------
------------------------------------------------------------
------------------------------------------------------------
------------------------------------------------------------
---------------------------------------------------taccacagg
gat---------------------------------------------------------
------------------------------------------------------------
------------------------------------------------------------
------------------------------------------------------------
------------------------------------------------------------
------------------------------------------------------------
------------------------------------------------------------
------------------------------------------------------------
------------------------------------------------------------
------------------------------------------------------------
------------------------------------------------------------
------------------------------------------------------------
------------------------------------------------------------
------------------------------------------------------------
------------------------------------------------------------
------------------------------------------------------------
------------------------------------------------------------
------------------------------------------------------------
------------------------------------------------------------
------------------------------------------------------------
------------------------------------------------------------
------------------------------------------------------------
---------------------------------aactggcttgtggcagccaagcgttca
tagcgacgttgctttttgatccttcgatgtcggctcttcctatcattgcgaagcagaatt
cgccaagtgttggattgttcacccactaatagggaacgtgagctgggtagccacttta--
-----tgcccaacagctggatagctgggcttctgtctta-ggccgggacttctcacgacc

cgtgccttggtcaccc-aaaaaagggcgaccgcagagctcaggatggagcttctggctta
actagtagaggacagatagagatcgtcaaattgcgggaaaccccttagagcctcgagcta
ccaagtgcgtgtcgaaaggcatagacgcagccagcgatggctgtcgcatggcttga-ttc
catcaagataggtaaaaatgctcgaggattgggcaatccgcagccaagcccctaaggt--
---ccccccgcgggctatggggaaggttcaacgactgtaaggcgtaccgtgcaagcggaa
gatacagtctagccccgctggaaagcagcgccaaagtaagtaggt-tgcagtgggtctgt
cctc
>Form_A_China_SRR9587962
gcatgttttggtatgtgtttcgacagtcatgcatgtcatagccgtaggatggggagtgag
cttcatagcccgaccccttctcaggagcatgttttcatagcatattgagcatactggggt
ggtctcccatgtagctcgctctctcggtgcaggacctcccttttgcgggccacttcgatg
gcctccattatgga-gagcatgttttggcatatgaagcgttggtggtgaggctcgtagcc
tgtatatacataccaacgcagtcgtcatatgtctctctggcgggggtcggtcgtatgacc
ttgtccctacctggagcatgtattttgtatgat---atactggac-ggcagctgcgctgt
tcagtcgtcagccgtacctcatgtgcgcgtttccaggtccgaacgtttgacatgcaaggg
catcataccagaagcctgttttctcgtatgagcatgtaggtaggcaggctcttcagtcgt
ctacccataaatgcacttcgaagga-actttttggtattgaatggctttgtgcccagacg
ttcttgtgagcttgttctcgctctgcgatacgctggaagctgtggtattacctacagcag
acgcattcctagggcggacggcctctctgcagatgggcacatctgtagaaagaggctccc
cttgatgacgcaatgcaatggctgggtaaacggagtgctatacagaagggtcaaacctc-
tgtggaagtacactcataccagcattgtttgttgagaggggtcggctctacgagtgagga
cctatactgccatctcattcagatgcgtagacggtctttactcagactgcctttctggct
gattgtcctcggtccctcgtgcgcacgcgccgctggctgaatgggcttgtccttatggag
tgccgcgttcttc-aaagaatccgtcttcatacagcgacctcatttacatattgccggct
gaatgggcttgtccttatggagcactgcattcttctaggaattgctatctatacagcgac
ctcatttacatacaaggcacct-ttttctacatagacttgccatttggctgattgttctg
acagtcgctgcctcttcggagttctgtgtggacgagtggcc--tcctgagtgggccggtc
cttacggagtgcccacgtctgttatgactgggttctccatacagcgacctcactcagaat
tgtatgg--------------caccgtgccgat-----------tgaatggacaagcagt
ccttagggaatcgcatgaaagcatgacggc-acatggtgtatggacccgttgggccttgc
ctgtgctatcttctcgagtcgaccgg------------tcctcgtctcctcctttcgcga
cgggaagctcttgagtttaccgtgggtcaggacgctgtggtatctgtttgtctgtagtgt
gggctctctatcagaacagcgtatcagcacacgtcgtctgggtctatggtatttttgctt
aacggccctgggtttggtagtatggttccatgtcggtcctctcacctcccaaccgtcatg
tctctga-----------------------------------------------------
------------------------------------------------------------

--------------------------------------------tatcatctgcacagta
cgtcttgtcagaatatcattctgtatggcagtcgatagactaggccctatgccgcagctg
tctctgaccactatgcgagcgcatacctacctcaatgtgttcatagctgttatggatctt
gggtgtggtgtgccagaacgatacgccaccagattgtggaatggtgcttgtagtattaga
agaggacatgtatgaagagcacgttcttttctcttatgatggtgccgcgggtgcgggccg
ttctcattctgcttgtcgggcgctcaaggtgcatgtaggattgctgaacccac-tgaatg
gtcatgagaatggtcatatggtgaggttcacttctccatcgctttgtgtgtgttggatgt
ttgggtggcttcggccgcatggaaaatctcggtctgtaactcatcagggagggtttccat
gtgagcaatctctccagtcgaacacacacccatgggcgtcccgctaggcaaaatggaact
ctcgatccgacacagcctgtggagtgtttgttccgtgtggcgtgaaacggcttatactgg
atgtcttccc-tttggacaccagtctgttggtatg--tacgcataccatctagagggagc
aagaaaaaggcatgcaagtgccacaaaggaagggaggccttcctgctaccgtgtgtcc-t
cttgacgatcacgggggagtatatagagacatttggacgagatccgtttttcacgaggcg
gtgccctcgcgcaaaggtgttgctctgcacgcatctttatgatgatagttgctcgctcct
atgcgtccactcgtgaaaatacctgctctctctgtctgtttccatgacacccc-catgcg
ttgtcatagagcatgctcaggtacagaaagtgctgtgactttcttggcctgctgtcactg
agtcagcctgatactttggaggaatcttttctgggcactgcccagctgagagttcttatc
aggctctggagtgtgcatatgcctatattcattatagtgcatgtggctacgcagaaaaag
catcgacgtggaatggagaagaacaccattctgaatggaacggttcttactctacttctg
ttgcttgacgctgtatgggaatcgctttgctctgggggtacaccttccattccatgacaa
agcatttgatctatgccccga-gggtcattctctatgaagagcccgaaacttgtctgagt
gagaggcgaaggtagtccttcattt--atgcggggtccgccctcacactgtagtcggcac
agtcccttgtttatggactgcgcagctcaatcttcgactggagagg--aaaaacaaaaca
caagtatgacacgcacatacgtgttttctaggactgtgacctgtcggctgcctcgcggtg
gttggcatgatgccccaagt-gggggggggagaaacaatcagtatgattgataaaaacca
gtgggg-aaaaaggcgcgctgtcgcacacccttggtcacggcctc---aagtcgacggcc
ctcactggccca-ccctgcgactcctgttcgcttttgacaagacacacctgcaata-cgg
aggaaatgcctgagcagctgtgccgcagcaactgcggacgcgtagcgatatgcatccctg
ttgtttgcgacttggtacaaagaacgtgcgccgggagacggattgccctttgacttttcc
acccagcccgcggctctatacccttgacgtacctatttacctggttgatcctgccagtag
tcatatgcttgtctcaaagactaagccatgcatgtctaagtataaacacttttatactcg
tgaaactgcgaatggctcattaaaacagttatagttcctttgggaacaacactacttgga
taaccgtagtaattctagagctaatacatgcctcaacgcccgactcacgaagggtggtat
ttattggataaaaaaccatcgtgctctgtccagagcgctttgagatgattcacaataact
tgtcggatcgcatggcctcgtgctggcgactcctcattcaaatttctgccctatcaactt
tcgatggtagagtattggtctaccatggtgtcgacgggtgacggggaattagggttcgat

tccggagagggagcctgagaaacggctaccacatccaaggaaggcagcaggcgcgcaaat
tacccaatcccgactcggggaggtagtgacaaaaaataacaatagggggccctttgggtc
ttctaattggaatgagaacaatttaaatcccttatcgaggatccattggagggcaagtct
ggtgaacgta----------------------------------tgcatta---------
--aagccctccgtaat-aaaaaaaaacagcgccagatgtct-------------------
------------------------------------------------------------
------------------------------------------------------------
------------------------------------------------------------
------------------------------------------------------------
------------------------------------------------------------
-------ctctctggaagcacacca-----------------------------------
---tgcatggggtttgttcctctc-----------------------------tgcatgg
ccgtg-----------------------------------------tgaaaaaggcgagt
aggcaaacgatgggaaacccggcttggaatagtaaccagccacagagcccccatcataca
cttctaaattgccggggagtgcccgcgaggccttgactaccgcggtccgccgagaggtga
gagcgagcaatcgttcaggacgcagcagtgtacgtagtggtacatggatggtaaaaacgt
caaggatagggttgatccgcagggaagcgcgagtgccgccatg---------gtcggtgg
gcaaaggcgaaccttcagagactctaatggagtgggcgcaagcttaagggagagtccaat
ccactgggaaaccaggtccacagcagtaacggggtgatgctcgccacagaggactgtgga
aggcaaccccacc----aaaagggtgtgtgttgtcgggtagc-aatggccagcagccgcg
gtaattccagctccaatagcgtatattaaagttgttgcagttaaaacgctcgtagtcgga
cctcggagcgcccccacgggccgggcttttgccttgttgccagtgggagtttgtctcctt
ttgtcgggacgcggtgtgcagggcctttactgtctctgcgcgtttacctgcgggccgacc
gtttactgtgaagaaagtaaagtgttcaaagcaggccaatgccttgaatatgtgagcatg
gaataatagaataggacttgggctctattttgttggtttccagtgaccaagtaatgatta
atagggacggttgggggcattcgtatttcattgtcagaggtgaaattcttggattgatgg
aagacgtacaactgcgaaagcatctgccatggatgttttcattgatcaagaacgaaagtt
aggggatcgaagacgatcagataccgtcgtagtcttaaccataaacgatgccgactgggg
attggcgggggcacttttatatgactctgtcagcaccctgagggaaaccaaagtctttgg
gttctgggggagtatggtcgcaaggctgaaacttaaaggaattgacggaagggcaccac
caggagtggagcctgcggcttaatttgactcaacacgggaaaacttaccaggtccggaca
gaagaatgattgacagactgaaagctctttcttgatttttggttggtggtgcatggccg
ttcttagttggtggagtgatttgtctggttaattccgttaacgaacgagacctcgtcctg
ctaaataggtgcgcgcatgccagcactgcgttcttaccttcttagagggactatgcgcgt
ctagcgtatggaagattgaggcaataacaggtctgtgatgcccttagatgttctgggccg
cacgcgcgctacactgatgcattcaacgagtttcacttgatagtcctgggtcggaaggct

cgggtaatcttttgaaagtgcatcgtgctggggatagatcattgcaattattgatcttca
acgaggaattccttgtaggcgcaggtcatcagcctgcgccgaatacgtccctgccctttg
tacacaccgcccgtcgctcctaccgattgaatggtccgatgaaatgtccggatcgcggct
agtgtgatggactcgtccagagcggtagccgtgagaagctcattaaatcttatcatttag
aggaaggagaagtcgtaacaaggtttccgtattttgtgggtggataccgattcaataaca
tctgccgtttttctcaatgtactatatccctagtctgcttgcccagttcttggagtctcg
tgcgatgtgtgctattacaggcacgctgggtcatgaagacagacaaggttaggatcttca
cggaagtacgcgcaataggagcgcgtcttgttcaacgcacctgactcaagagtcatatgc
gagggacggacgcagcgcttgtcgccgcacagatgggaagcttcctctccaggcatagga
tggcgctgcatttgcatcatcgctgctagtaagtggcaataatacttgattccccgccag
cgcacttgtacgtagccgttggcggcgggaacgtagttggtgatgtgacagcctcgagcg
ttcacctctgtgtgtccgccaatctgcacaaagatgaacgccatatcggagacagggga
ggtgccattg---tgatgatgacgatgttgttgggtggtccttccgtaggaaaacgtaca
cgaagcctttgcggcccgaaagggtgcacagtgcgcgactataaacactgattctgtggt
caacgcaagcagcggttctgtggaagaacagctgcgactttctcaaattgcgggaacacc
ctaaagtcaacaagcaccgcggcttgacgtgaaaagcgtcaggcgcagcagtcggggtag
cgccctggcggatggtaatagccttgttgaatggcacaatgggcaatccgcagccaagcc
cctaaggtcccccgctgggctatggggaaggttcacagactgtaagggagggggtttctt
ccatga--aagaagcttaagagacagtcggtcccctgcgagagcagtgtcgcatgtgagg
aaggcacccatgaaaagtgggtgctggagagccatgtgagccctctatgagggcgggaag
ttttccga---tacaatgagaaacaaacacgttccgtaggtgaacctgcggaaggatcat
tcacaaactattgacacaaacacacgcgaaccaaatcgtccgcacagcaggtggcaatga
aagagagactctgcatgtcgcctttcgggg--tagcaagcagcactcttctgccatcgcc
tctgtgcagggcgtaaattcccattgagaggatgtgagggcaccacaggaagcttttcca
caggaagtcgccatccttctccctccacggcggcg----cttctgtgcttggcag-----
-----ttttttttgccttctagggaggatgccgccaatggagccccttat-atatataca
tcatcatatgccc--cttttttttcttaaccgcttgccacagcttcttctatgaggagctt
gtgggaagactgtctccatacaataac-aaagatacaactcttagcggtggatatcttgg
ttctcgcaacgatgaagaacgcagccaactgcgataactaatgtgaattgcaggacttcg
tgaatcattaagtctttgaacgcaagttgcgctcatctccatgaggaggtgagcatgcct
gtttgagtgtcgatgggaaaaacaatac-ataacaacaatcggctctcttatcttcctgc
cttcttgggggcaaaaggtagtgcatgccggagatgggctttgtcgcccctttctcttga
aagaggcgtcgtccgaaacacgtatcgttgcccgctggaggtcatagagcatgagttt--
aaagcaccacatctctcaagtcagagtgag-cgtcttgagtaggtcc-gcattgattcag
tttcaacccaggtggcgttggttacttgccataaagcctctctggggtttctttgcttgg
acctctcttgtcagtttctctgacgtagagacaggt--gccgtctcatggacgttgccct

ctggaacgtgct---aaaaaaaaaaaaaga----------aaaaaccgtaccacatgtcaa
aagaggcagatcgtgcttgacggcgggaggtgttgcatttcttgcttgtcctgagcgtct
gccagtctct----ga--cgagtg----tgttccttccataaacccgatctcaaatcagg
taggaccacccgctgaatttaagcatataattaagcggaggaaaagaaactaacaaggat
tccctcagtagcggcgagcgaagcgggaagagctcaacatgaaaaccgttgggggttcgc
gctttccttcgggttgtatagtaaagtggtgtcgcctgcgtcgcgccgggtataaatccc
gtctgaaatggggtgtcgtagagggttatatccccgtctgtgacccggtgtgcgatgcgt
tacggtgaccttacgatgagtcgggttgcttgggcatgcagccctaagtgggtggtaaac
ttcatctaaagctaaatactagcgagagaccgatagcgtacaagtaccgtaagggaaaga
tgcaaagaactttgaaaagagagttaaacagggcttgaaaccgttaagagtgaagcgaag
gaggtcagtgatcatggggccagatccccaagtgcctgtcata-gggcaccgtagcgttc
tggtttcccggggcagcgccggccgggggcgggggttaacagggaggtggttgagggatt
cgttttccgcctgtactcctttatagcttgcaccccggccgagggatccgcgacttgtcg
tgtgagggcgctgtcaaaatggccttctccaccccgtcttgaaacacggaccaaggagtc
taacatactcgcgagtgtttgggtggttaaaacccgaacgcacaatgaaagtaaacggtg
gctctgcgccgctgcttggtgggaaccctgcttttataatggggggcactatcacccgac
cccgatccctttggggtggttggttttgaggtagagcgagtatgatgggacccgaaagat
ggtgaactattcctgagtagggcgaagccagaggaaactctggtggaggctcgtagcgat
gctgacgtgcaaatcgctcgtcgaacttgggtataggggcgaaagactaatcgaaccatc
tagtagctggttccctccgaagtttccctcaggatagcgggagcaaatgcaagtcagtat
catccggtaaagcgaatgattagaggcctcgggggagtaactccttcgacctattctcaa
actttaaataggtgaggagccggcaacgttgcttaaaacaatgaacgccgtgcggacaat
gattgctctcagtgggccgattttggtaagcagaactggcgatgcgggatgaaccgaaag
cgacgctaaggtgccaaactcccagctcattcagacaccagaaaaggtgttgattcatac
agacagcaggacggtggccatggaagtcggaatccgctaaggagtgtgtaacaactcacc
tgccgaatggatcagccctgaaaatggatggcgcttaagcgggggaccgaggcgtcgccg
ttggggtaaatacgaagccccaatgagtaggagggcgtgggggttttgaaacacggtaca
ctcgcatggaagccttgggcgtaagctcgggtggagcgacccctagtgtagatcttggtg
gtagtagcaaatattcaagtgagaaccttgaagactgaagtggggaagggttccacgcga
acagtgcttggacgtgggttagtcggtcctaagagatggggaaactccgtattcaagtgc
gcttttgtagcgcagtccatcgaaagggaagcgggttaacattcccgcaccaggatgtgg
aaactgtggtttaccgcaaactgcggcaacgcaactgaacctggggacgcccgtaggagc
tccgagcagggatctctttcctgtttaacgcgcccaacccccctggaatcggtttgtccgg
agatagggggcgaaaggccggaaaagcaccgcacgtcttgtggtgttcggaacgcttgta
atggcccttgaaaacccaggggaaagcagccgtcttacac--ttgtgtgagacgaggaat
tctcacacctggccgtacccatcaccgcatcaggtccccaaggtgaacagcctctagtca

atagaataatgtagataagggaagtcggcaaaatagatccgtaactttgggaaaaggatt
ggctctaatgttcgggcttgtcgggacgcgcgggcgtagatgccaatgagcggcagacgt
ctagcctctttggaggctggattaacgctgcttgtgggaagaaaccgtgtcgatggaggc
aggaacgcttatcgatcttgtctcttcccgcttgcgatcaacgaacggattagaactggt
acggacaaggggaatccgactgtttaattaaaacaaagcattgcgacggccggaaacggt
gttggcgcaatgtgatttctgcccagtgctctgaatgtcaaagtgaagaaattcaaccaa
gcgcgggtaaacggcgggagtaactatgactct--taaac-aaaccatactacctac---
-------------------gggagtcaaaagtgggggcggcgtctggtgtagaaatacac
gtac--------------------------------------------------------
-------------------gacacgctttttttcctgctagtcgatctgacctttgttct
ttggactgtgg-------------------------------------------------
--------------------------gcggtcagcggcaacataacc-------------
------------------------------------------------------------
------------------------------------------------------------
----------------------------tggtacggg---------------------ga
aggcctcgacgttga---------------------------------------------
--------------------------------gtgctgcatcttggaagtatc-------
-----------------------------ttgacacaatgagaga---------------
------------------------------aaagagattgcgcttaatagg---------
----gggttgatctcgt-------------------------------------------
-------------------------------------------ggcgagccgacgc----
---------------------------------cggtgtgaactggt-------------
------------------------------------------------------------
-----------------------ggcaat-------------------------------
----ggccgtcgtagagcacgcaaaggtatgggctgactttttctcctca----aaagtt
ggcttaagggac-gtgccatcccacacgataataaagtgtgcagtctg------------
--------------------taagagctcccacagggcaaagac---------agactgg
gatgcagacggagacggcaaaaacaacaacatccgttctgcttctgggcgcaaatgctta
aggtagccaaatgcctcgtcatctaaactaattgg-at----------------------
-----------------------gagatagtc----------tgagtgaaaagcttatgc
ag-----------------gctagtaccgg----aaaaaaaaaaaagaggggctctgcca
ct----ctgac------------------------ggtgcgacactccttgtacagggaa
accctcc--gcatggaaaggcacgtggtaccaccctcgcggggaaactcgcgagcgggat
cccgggtaatgaccggagccgatggtaaaaactcacgtgaccgatggggtgaccctgtgg
cgagacgctaagttgtgggacatcaccctctcacaatacgcatccgtcgcagagactgaa
cgggagtgggctccgttga-----cggccggccggccagttagccgtcttcca-------

---ggggcttaagatacagtccgggcctcagggatgg----aaaaacctggggaacta-c

cagaattagtgacgcgcatgaatggattaacgagattcccactgtccctatctactatct

agcgaaaccacagccaagggaacgggcttggaagaatcagcggggaaagaagaccctgtt

gagcttgactctagtccgactttgtgagatgagttgggaggtgtagtctaagagggaggg

gcaactcgaccgtgaaatacctctactcccaagcttgttttacttactccggcgagcagg

gatgggtggtaacacccttttctaggattaacacctcttcgggggtggacctgcgcgga

ggacattgtcaggtggggagtttggctggggcggcacatctgttaaaagataacgcaggt

gtcctaagatgagctcagtgagaacagaaatctcacgtagaacaaaagggtaaaagctca

tttgattttgattttcagtacgaatacaaactgtgaaagcatggcctatcgatcctttg

accttcaaggtttgaagcaagaggtgtcagaaaagttaacacgcgttgtggtagcttta

agagtggactcaaagtgttccgctagtccccctcggaccgccacttacaaaaggcctgg

ggatgggggcgacactgtcaaattgcggggactccctaaagcctgtctctaccaagccag

cgcggcgaaagcccctggtggccggggtaatgacctagggtatggtaaaagcgagacagg

atg-gaacaatgggcaatccgcagccaagtcctaagg----ctttttgaaaagcggctat

ggatgcagttcaacgactagatggcagtgggctgagagagcccaccgcggcaccctttcg

gcttaagatatagtcttgcccacgtgaaagcgtgcctgcagatgaacgccaccatgtatg

gccggagctgcaggagaccgc--acttgatgtgcagtcgggaat-cgacagtaccacagg

gatgaac-----------------------------------------------------

------------------------------------------------------------

------------------------------------------------------------

------------------------------------------------------------

------------------------------------------------------------

-------------------------------------------------aaaaacac---

------------------------------------------------------------

------------------------------------------------------------

--acaatta---------------------------------------------------

----------------------------------------atttgtggcc---atcatac

----------------------------------------agtcgta-------------

------------------------------------------------------------

------------------------------------------------------------

------------------------------------------------------------

------------tgatattatcgtcccaggcgggtgcgcctctctcacccgtctagtccg

gccttggtgcgcaaaaaatgcttgcgtggatgcgccttggttaaggaaaatgctcctagt

agttttacaaggacacttcttgtagggcaacacatctggatgcgaagaactcccggaaaa

gacaacggtaccaagtgactggggtaacgccccattcatgaaggcaaataatcaaaaaga

ggtgccgggtaagaacccgttgtatagggacaatccgcagctgacttggctccctg--cc

cat--ccatgggcatagcttagcggttcaacgctcggtaagatgtgggttggcgcgctgt
ccgacagcgcactggcttaaggtacgggctactcccaccggagacggtgtctttgtctac
ctcgggtaacacgcctcgggagaagcaaagagggtgcaatggaaaaaaaactggttgcgg
ggaaatccctaccagagtcacctggtacaaaggaactggcttgtggcagccaagcgttca
tagcgacgttgctttttgatccttcgatgtcggctcttcctatcattgcgaagcagaatt
cgccaagtgttggattgttcacccactaatagggaacgtgagctgggtagccacttta--
-----tgcccaacagctggatagctgggcttctgtctta-ggccgggacttctcacgacc
cgtgccttggtcaccc-aaaaaagggcgaccgcagagctcaggatggagcttctggctta
actagtagaggacagatagagatcgtcaaattgcgggaaaccccttagagcctcgagcta
ccaagtgcgtgtcgaaaggcatagacgcagccagcgatggctgtcgcatggcttga-ttc
catcaagataggtaaaaatgctcgaggattgggcaatccgcagccaagcccctaaggt--
---ccccccgcgggctatggggaaggttcaacgactgtaaggcgtaccgtgcaagcggaa
gatacagtctagccccgctggaaagcagcgccaaagtaagtaggt-tgcagtgggtctgt
cctc
>Form_A_China_SRR9587944
gcatgttttggtatgtgtttcgacagtcatgcatgtcatagccgtaggatggggagtgag
cttcatagcccgaccccttctcaggagcatgttttcatagcatattgagcatactggggt
ggtctcccatgtagctcgctctctcggtgcaggacctcccttttgcgggccacttcgatg
gcctccattatgga-gagcatgttttggcatatgaagcgttggtggtgaggctcgtagcc
tgtatatacataccaacgcagtcgtcatatgtctctctggcgggggtcggtcgtatgacc
ttgtccctacctggagcatgtattttgtatgat---atactggac-ggcagctgcgctgt
tcagtcgtcagccgtacctcatgtgcgcgtttccaggtccgaacgtttgacatgcaaggg
catcataccagaagcctgttttctcgtatgagcatgtaggtaggcaggctcttcagtcgt
ctacccataaatgcacttcgaagga-actttttggtattgaatggctttgtgcccagacg
ttcttgtgagcttgttctcgctctgcgatacgctggaagctgtggtattacctacagcag
acgcattcctagggcggacggcctctctgcagatgggcacatctgtagaaagaggctccc
cttgatgacgcaatgcaatggctgggtaaacggagtgctatacagaagggtcaaacctc-
tgtggaagtacactcataccagcattgtttgttgagaggggtcggctctacgagtgagga
cctatactgccatctcattcagatgcgtagacggtctttactcagactgcctttctggct
gattgtcctcggtccctcgtgcgcacgcgccgctggctgaatgggcttgtccttatggag
tgccgcgttcttc-aaagaatccgtcttcatacagcgacctcatttacatattgctggct
gaatgggcttgtccttatggagcactgcattcttctaggaattgctatctatacagcgac
ctcatttacatacaaggcacct-ttttctacatagacttgccatttggctgattgttctg
acagtcgctgcctcttcggagttctgtgtggacgagtggcc--tcctgagtgggccggtc
cttacggagtgcccacgtctgttatgactgggttctccatacagcgacctcactcagaat
tgtatgg--------------caccgtgccgat-----------tgaatggacaagcagt

ccttagggaatcgcatgaaagcatgacggc-acatggtgtatggacccgttgggccttgc
ctgtgctatcttctcgagtcgaccgg------------tcctcgtctcctcctttcgcga
cgggaagctcttgagtttaccgtgggtcaggacgctgtggtatctgtttgtctgtagtgt
gggctctctatcagaacagcgtatcagcacacgtcgtctgggtctatggtatttttgctt
aacggccctgggtttggtagtatggttccatgtcggtcctctcacctcccaaccgtcatg
tctctga-----------------------------------------------------
------------------------------------------------------------
--------------------------------------------tatcatctgcacagta
cgtcttgtcagaatatcattctgtatggcagtcgatagactaggccctatgccgcagctg
tctctgaccactatgcgagcgcatacctacctcaatgtgttcatagctgttatggatctt
gggtgtggtgtgccagaacgatacgccaccagattgtggaatggtgcttgtagtattaga
agaggacatgtatgaagagcacgttcttttctcttatgatggtgccgcgggtgcgggccg
ttctcattctgcttgtcgggtgctcaaggtgcatgtaggattgctgaacccac-tgaatg
gtcatgagaatggtcatatggtgaggttcacttctccatcgctttgtgtgtgttggatgt
ttgggtggcttcggccgcatggaaaatctcggtctgtaactcatcagggagggtttccat
gtgagcaatctctccagtcgaacacacacccatgggcgtcccgctaggcaaaatggaact
ctcgatccgacacagcctgtggagtgtttgttccgtgtggcgtgaaacggcttatactgg
atgtcttccc-tttggacaccagtctgttggtatg--tacgcataccatctagagggagc
aagaaaaaggcatgcaagtgccacaaaggaagggaggccttcctgctaccgtgtgtcc-t
cttgacgatcacgggggagtatatagagacatttggacgagatccgtttttcacgaggcg
gtgccctcgcgcaaaggtgttgctctgcacgcatctttatgatgatagttgctcgctcct
atgcgtccactcgtgaaaatacctgctctctctgtctgtttccatgacacccc-catgcg
ttgtcatagagcatgctcaggtacagaaagtgctgtgactttcttggcctgctgtcactg
agtcagcctgatactttggaggaatcttttctgggcactgcccagctgagagttcttatc
aggctctggagtgtgcatatgcctatattcattatagtgcatgtggctacgcagaaaaag
catcgacgtggaatggagaagaacaccattctgaatggaacggttcttactctacttctg
ttgcttgacgctgtatgggaatcgctttgctctgggggtacaccttccattccatgacaa
agcatttgatctatgccccga-gggtcattctctatgaagagcccgaaacttgtctgagt
gagaggcgaaggtagtccttcattt--atgcggggtccgccctcacactgtagtcggcac
agtcccttgtttatggactgcgcagctcaatcttcgactggagagg--aaaaacaaaaca
caagtatgacacgcacatacgtgttttctaggactgtgacctgtcggctgcctcgcggtg
gttggcatgatgccccaagt-ggggggggagaaacaatcagtatgattgataaaaacca
gtgggg-aaaaaggcgcgctgtcgcacacccttggtcacggcctc---aagtcgacggcc
ctcactggccca-ccctgcgactcctgttcgcttttgacaagacacacctgcaata-cgg
aggaaatgcctgagcagctgtgccgcagcaactgcggacgcgtagcgatatgcatccctg
ttgtttgcgacttggtacaaagaacgtgcgccgggagacggattgccctttgacttttcc

acccagcccgcggctctatacccttgacgtacctatttacctggttgatcctgccagtag
tcatatgcttgtctcaaagactaagccatgcatgtctaagtataaacacttttatactcg
tgaaactgcgaatggctcattaaaacagttatagttcctttgggaacaacactacttgga
taaccgtagtaattctagagctaatacatgcctcaacgcccgactcacgaagggtggtat
ttattggataaaaaaccatcgtgctctgtccagagcgctttgagatgattcacaataact
tgtcggatcgcatggcctcgtgctggcgactcctcattcaaatttctgccctatcaactt
tcgatggtagagtattggtctaccatggtgtcgacgggtgacggggaattagggttcgat
tccggagagggagcctgagaaacggctaccacatccaaggaaggcagcaggcgcgcaaat
tacccaatcccgactcggggaggtagtgacaaaaataacaatagggggccctttgggtc
ttctaattggaatgagaacaatttaaatcccttatcgaggatccattggagggcaagtct
ggtgaacgta----------------------------------tgcatta---------
--aagccctccgtaat-aaaaaaaaacagcgccagatgtct-------------------
------------------------------------------------------------
------------------------------------------------------------
------------------------------------------------------------
------------------------------------------------------------
------------------------------------------------------------
-------ctctctggaagcacacca-----------------------------------
---tgcatggggtttgttcctctc-----------------------------tgcatgg
ccgtg-----------------------------------------tgaaaaaggcgagt
aggcaaacgatgggaaacccggcttggaatagtaaccagccacagagcccccatcataca
cttctaaattgccggggagtgcccgcgaggccttgactaccgcggtccgccgagaggtga
gagcgagcaatcgttcaggacgcagcagtgtacgtagtggtacatggatggtaaaaacgt
caaggatagggttgatccgcagggaagcgcgagtgccgccatg---------gtcggtgg
gcaaaggcgaaccttcagagactctaatggagtgggcgcaagcttaagggagagtccaat
ccactgggaaaccaggtccacagcagtaacggggtgatgctcgccccagaggactgtgga
aggcaaccccacc----aaaagggtgtgtgttgtcgggtagc-aatggccagcagccgcg
gtaattccagctccaatagcgtatattaaagttgttgcagttaaaacgctcgtagtcgga
cctcggagcgcccccacgggccgggcttttgccttgttgccagtgggagtttgtctcctt
ttgtcgggacgcggtgtgcagggcctttactgtctctgcgcgtttacctgcgggccgacc
gtttactgtgaagaaagtaaagtgttcaaagcaggccaatgccttgaatatgtgagcatg
gaataatagaataggacttgggctctattttgttggtttccagtgaccaagtaatgatta
atagggacggttgggggcattcgtatttcattgtcagaggtgaaattcttggattgatgg
aagacgtacaactgcgaaagcatctgccatggatgttttcattgatcaagaacgaaagtt
aggggatcgaagacgatcagataccgtcgtagtcttaaccataaacgatgccgactgggg
attggcgggggcacttttatatgactctgtcagcaccctgagggaaaccaaagtctttgg

gttctggggggagtatggtcgcaaggctgaaacttaaaggaattgacggaagggcaccac
caggagtggagcctgcggcttaatttgactcaacacgggaaaacttaccaggtccggaca
gaagaatgattgacagactgaaagctctttcttgatttttggttggtggtgcatggccg
ttcttagttggtggagtgatttgtctggttaattccgttaacgaacgagacctcgtcctg
ctaaataggtgcgcgcatgccagcactgcgttcttaccttcttagagggactatgcgcgt
ctagcgtatggaagattgaggcaataacaggtctgtgatgcccttagatgttctgggccg
cacgcgcgctacactgatgcattcaacgagtttcacttgatagtcctgggtcggaaggct
cgggtaatcttttgaaagtgcatcgtgctggggatagatcattgcaattattgatcttca
acgaggaattccttgtaggcgcaggtcatcagcctgcgccgaatacgtccctgccctttg
tacacaccgcccgtcgctcctaccgattgaatggtccgatgaaatgtccggatcgcggct
agtgtgatggactcgtccagagcggtagccgtgagaagctcattaaatcttatcatttag
aggaaggagaagtcgtaacaaggtttccgtattttgtgggtggataccgattcaataaca
tctgccgtttttctcaatgtactatatccctagtctgcttgcccagttcttggagtctcg
tgcgatgtgtgctattacaggcacgctgggtcatgaagacagacaaggttaggatcttca
cggaagtacgcgcaataggagcgcgtcttgttcaacgcacctgactcaagagtcatatgc
gagggacggacgcagcgcttgtcgccgcacagatgggaagcttcctctccaggcatagga
tggcgctgcatttgcatcatcgctgctagtaagtggcaataatacttgattccccgccag
cgcacttgtacgtagccgttggcggcgggaacgtagttggtgatgtgacagcctcgagcg
ttcacctctgtgtgtccgccaatctgcacaaagatgaacgccatatcggagacaggggga
ggtgccattg---tgatgatgacgatgttgttgggtggtccttccgtaggaaaacgtaca
cgaagcctttgcggcccgaaagggtgcacagtgcgcgactataaacactgattctgtggt
caacgcaagcagcggttctgtggaagaacagctgcgactttctcaaattgcgggaacacc
ctaaagtcaacaagcaccgcggcttgacgtgaaaagcgtcaggcgcagcagtcggggtag
cgccctggcggatggtaatagccttgttgaatggcacaatgggcaatccgcagccaagcc
cctaaggtcccccgctgggctatggggaaggttcacagactgtaagggagggggtttctt
ccatga--aagaagcttaagagacagtcggtcccctgcgagagcagtgtcgcatgtgagg
aaggcacccatgaaaagtgggtgctggagagccatgtgagccctctatgagggcgggaag
ttttccga---tacaatgagaaacaaacacgttccgtaggtgaacctgcggaaggatcat
tcacaaactattgacacaaacacacgcgaaccaaatcgtccgcacagcaggtggcaatga
aagagagactctgcatgtcgcctttcgggg--tagcaagcagcactcttctgccatcgcc
tctgtgcagggcgtaaattcccattgagaggatgtgagggcaccacaggaagcttttcca
caggaagtcgccatccttctccctccacggcggcg----cttctgtgcttggcag-----
-----ttttttttgccttctagggaggatgccgccaatggagccccttat-atatataca
tcatcatatgccc--cttttttttcttaaccgcttgccacagcttcttctatgaggagctt
gtgggaagactgtctccatacaataac-aaagatacaactcttagcggtggatatcttgg
ttctcgcaacgatgaagaacgcagccaactgcgataactaatgtgaattgcaggacttcg

tgaatcattaagtctttgaacgcaagttgcgctcatctccatgaggaggtgagcatgcct
gtttgagtgtcgatgggaaaaacaatac-ataacaacaatcggctctcttatcttcctgc
cttcttgggggcaaaaggtagtgcatgccggagatgggctttgtcgcccctttctcttga
aagaggcgtcgtccgaaacacgtatcgttgcccgctggaggtcatagagcatgagttt--
aaagcaccacatctctcaagtcagagtgag-cgtcttgagtaggtcc-gcattgattcag
tttcaacccaggtggcgttggttacttgccat-aagcctctctggggtttctttgcttgg
acctctcttgtcagtttctctgacgtagagacaggt--gccgtctcatggacgttgccct
ctggaacgtgctaaaaaaaaaaaaaaaga----------aaaaaccgtaccacatgtcaa
aagaggcagatcgtgcttgacggcgggaggtgttgcatttcttgcttgtcctgagcgtct
gccagtctct----ga--cgagtg----tgttccttccataaacccgatctcaaatcagg
taggaccacccgctgaatttaagcatataattaagcggaggaaaagaaactaacaaggat
tccctcagtagcggcgagcgaagcgggaagagctcaacatgaaaaccgttgggggttcgc
gctttccttcgggttgtatagtaaagtggtgtcgcctgcgtcgcgccgggtataaatccc
gtctgaaatggggtgtcgtagagggttatatccccgtctgtgacccggtgtgcgatgcgt
tacggtgaccttacgatgagtcgggttgcttgggcatgcagccctaagtgggtggtaaac
ttcatctaaagctaaatactagcgagagaccgatagcgtacaagtaccgtaagggaaaga
tgcaaagaactttgaaaagagagttaaacagggcttgaaaccgttaagagtgaagcgaag
gaggtcagtgatcatggggccagatccccaagtgcctgtcata-gggcaccgtagcgttc
tggtttcccggggcagcgccggccgggggcgggggttaacagggaggtggttgagggatt
cgttttccgcctgtactcctttatagcttgcaccccggccgagggatccgcgacttgtcg
tgtgagggcgctgtcaaaatggccttctccaccccgtcttgaaacacggaccaaggagtc
taacatactcgcgagtgtttgggtggttaaaacccgaacgcacaatgaaagtaaacggtg
gctctgcgccgctgcttggtgggaaccctgcttttataatgggggcactatcacccgac
cccgatccctttggggtggttggttttgaggtagagcgagtatgatgggacccgaaagat
ggtgaactattcctgagtagggcgaagccagaggaaactctggtggaggctcgtagcgat
gctgacgtgcaaatcgctcgtcgaacttgggtataggggcgaaagactaatcgaaccatc
tagtagctggttccctccgaagtttccctcaggatagcgggagcaaatgcaagtcagtat
catccggtaaagcgaatgattagaggcctcgggggagtaactccttcgacctattctcaa
actttaaataggtgaggagccggcaacgttgcttaaaacaatgaacgccgtgcggacaat
gattgctctcagtgggccgattttggtaagcagaactggcgatgcgggatgaaccgaaag
cgacgctaaggtgccaaactcccagctcattcagacaccagaaaaggtgttgattcatac
agacagcaggacggtggccatggaagtcggaatccgctaaggagtgtgtaacaactcacc
tgccgaatggatcagccctgaaaatggatggcgcttaagcgggggaccgaggcgtcgccg
ttggggtaaatacgaagccccaatgagtaggagggcgtgggggttttgaaacacggtaca
ctcgcatggaagccttgggcgtaagctcgggtggagcgacccctagtgtagatcttggtg
gtagtagcaaatattcaagtgagaaccttgaagactgaagtggggaagggttccacgcga

acagtgcttggacgtgggttagtcggtcctaagagatggggaaactccgtattcaagtgc
gcttttgtagcgcagtccatcgaaagggaagcgggttaacattcccgcaccaggatgtgg
aaactgtggtttaccgcaaactgcggcaacgcaactgaacctggggacgcccgtaggagc
tccgagcagggatctctttcctgtttaacgcgcccaacccccctggaatcggtttgtccgg
agatagggggcgaaaggccggaaaagcaccgcacgtcttgtggtgttcggaacgcttgta
atggcccttgaaaacccaggggaaagcagccgtcttacac--ttgtgtgagacgaggaat
tctcacacctggccgtacccatcaccgcatcaggtccccaaggtgaacagcctctagtca
atagaataatgtagataagggaagtcggcaaaatagatccgtaactttgggaaaaggatt
ggctctaatgttcgggcttgtcgggacgcgcgggcgtagatgccaatgagcggcagacgt
ctagcctctttggaggctggattaacgctgcttgtgggaagaaaccgtgtcgatggaggc
aggaacgcttatcgatcttgtctcttcccgcttgcgatcaacgaacggattagaactggt
acggacaaggggaatccgactgtttaattaaaacaaagcattgcgacggccggaaacggt
gttggcgcaatgtgatttctgcccagtgctctgaatgtcaaagtgaagaaattcaaccaa
gcgcgggtaaacggcgggagtaactatgactct--taaac-aaaccatactacctac---
-------------------gggagtcaaaagtgggggcggcgtctggtgtagaaatacac
gtac--------------------------------------------------------
-------------------gacacgctttttttcctgctagtcgatctgacctttgttct
ttggactgtgg-------------------------------------------------
--------------------------gcggtcagcggcaacataacc-------------
------------------------------------------------------------
------------------------------------------------------------
----------------------------tggtacggg---------------------ga
aggcctcgacgttga---------------------------------------------
--------------------------------gtgctgcatcttggaagtatc-------
-----------------------------ttgacacaataagaga---------------
------------------------------aaagagattgcgcttaatagg---------
----gggttgatctcgt-------------------------------------------
-------------------------------------------ggcgagccgacgc----
---------------------------------cggtgtgaactggt-------------
------------------------------------------------------------
-----------------------ggcaat-------------------------------
----ggccgtcgtagagcacgcaaaggtatgggctgactttttctcctca----aaagtt
ggcttaagggac-gtgccatcccacacgataataaagtgtgcagtctg------------
--------------------taagagctcccacagggcaaagac---------agactgg
gatgcagacggagacggcaaaaacaacaacatccgttctgcttctgggcgcaaatgctta
aggtagccaaatgcctcgtcatctaaactaattgg-at----------------------

-----------------------gagatagtc----------tgagtgaaaagcttatgc
ag-----------------gctagtaccgg---aaaaaaaaaaaaagaggggctctgcca
ct----ctgac------------------------ggtgcgacactccttgtacagggaa
accctcc--gcatggaaaggcacgtggtaccaccctcgcggggaaactcgcgagcgggat
cccgggtaatgaccggagccgatggtaaaaactcacgtgaccgatggggtgaccctgtgg
cgagacgctaagttgtgggacatcaccctctcacaatacgcatccgtcgcagagactgaa
cgggagtgggctccgtcga-------------cggctaaccagccttcgtcgt-------
---ggggcttaagatacagtccgggcctcagggatgg----aaaaacctggggaacta-c
cagaattagtgacgcgcatgaatggattaacgagattcccactgtccctatctactatct
agcgaaaccacagccaagggaacgggcttggaagaatcagcggggaaagaagaccctgtt
gagcttgactctagtccgactttgtgagatgagttgggaggtgtagtctaagagggaggg
gcaactcgaccgtgaaatacctctactcccaagcttgttttacttactccggcgagcagg
gatgggtggtaacacccttttctaggattaacacctcttcgggggtggacctgcgcgga
ggacattgtcaggtggggagtttggctggggcggcacatctgttaaaagataacgcaggt
gtcctaagatgagctcagtgagaacagaaatctcacgtagaacaaaagggtaaaagctca
tttgattttgattttcagtacgaatacaaactgtgaaagcatggcctatcgatcctttg
accttcaaggtttgaagcaagaggtgtcagaaaagttaacacgcgttgtggtagctttta
agagtggactcaaagtgttccgctagtccccccctcggaccgccacttacaaaaggcctgg
ggatgggggcgacactgtcaaattgcggggactccctaaagcctgtctctaccaagccag
cgcggcgaaagcccctggtggccggggtaatgacctagggtatggtaaaagcgagacagg
atg-gaacaatgggcaatccgcagccaagtcctaaggc--tctttttgaaaagcggctat
ggatgcagttcaacgactagatggcagtgggctgagagagcccaccgcggcacccttcg
gcttaagatatagtcttgcccacgtgaaagcgtgcctgcagatgaacgccaccatgtatg
gccggagctgcaggagaccgc--acttgatgtgcagtcgggaat-cgacagtaccacagg
gatgaac-----------------------------------------------------
------------------------------------------------------------
------------------------------------------------------------
------------------------------------------------------------
------------------------------------------------------------
-------------------------------------------------aaaaacac---
------------------------------------------------------------
------------------------------------------------------------
------------------------------------------------------------
--------------------------------------acaattgtggcc---atcatac
----------------------------------------agtcgta-------------
------------------------------------------------------------

------------------------------------------------------------
------------------------------------------------------------
------------tgatattatcgtcccaggcgggtgcgcctctctcacccgtctagtccg
gccttggtgcgcaaaaaatgcttgcgtggatgcgccttggttaaggaaaatgctcctagt
agttttacaaggacacttcttgtagggcaacacatctggatgcgaagaactcccggaaaa
gacaacggtaccaagtgactggggtaacgccccattcatgaaggcaaataatcaaaaaga
ggtgccgggtaagaacccgttgtatagggacaatccgcagctgacttggctccctg--cc
catccccatgggcatagcttagcggttcaacgctcggtaagatgtgggttggcgcgctgt
ccgacagcgcactggcttaaggtacgggctactcccaccggagacggtgtctttgtctac
ctcgggtaacacgcctcgggagaagcaaagagggtgcaatgg-aaaaaaactggttgcgg
ggaaatccctaccagagtcacctggtacaaaggaactggcttgtggcagccaagcgttca
tagcgacgttgctttttgatccttcgatgtcggctcttcctatcattgcgaagcagaatt
cgccaagtgttggattgttcacccactaatagggaacgtgagctgggtagccactttа--
-----tgcccaacagctggatagctgggcttctgtctta-ggccgggacttctcacgacc
cgtgccttggtcaccc-aaaaaagggcgaccgcagagctcaggatggagcttctggctta
actagtagaggacagatagagatcgtcaaattgcgggaaaccccttagagcctcgagcta
ccaagtgcgtgtcgaaaggcatagacgcagccagcgatggctgtcgcatggcttga-ttc
catcaagataggtaaaaatgctcgaggattgggcaatccgcagccaagcccctaaggt--
---ccccccgcgggctatggggaaggttcaacgactgtaaggcgtaccgtgcaagcggaa
gatacagtctagccccgctggaaagcagcgccaaagtaagtaggt-tgcagtgggtctgt
cctc
>Form_E_China_SRR9587930
gcatgttttggtatgtgtttcgacagtcatgcatgtcatagccgtaggatggggagtgag
cttcatagcccgaccccttctcaggagcatgttttcatagcatattgagcatactggggt
ggtctcccatgtagctcgctctctcggtgcaggacctcccttttgcgggccacttcgatg
gcctccattatgga-gagcatgttttggcatatgaagcgttggtggtgaggctcgtagcc
tgtatatacataccaacgcagtcgtcatatgtctctctggcgggggtcggtcgtatgacc
ttgtccctacctggagcatgtatttcgtatgat---atactggac-ggcagctgcgctgt
tcagtcgtcggccgtacctcatgtgcgcgtttccaggtccgaacgtttgacatgcaaggg
catcataccagaagcctgttttctcgtatgagcatgtaggtaggcaggctcttcagtcgt
ctacccataaatgcacttcgaagga-actttttggtattgaatggctttgtgcccagacg
ttcttgtgagcttgttctcgctctgcgatacgctggaagctgtggtattacctacagcag
acgcattcctagggcggacggcctctctgcagatgggcacatctgtagaaagaggctccc
cttgatgacgcaatgcaatggctgggtaaacggagtgctatacagaagggtcaaacctc-
tgtggaagtacactcataccagcattgtttgttgagaggggtcggctctacgagtgagga
cctatactgccatctcattcagatgcgtagacggtctttactcagactgcctttctggct

gattgtcctcggtccctcgtgcgcacgcgccgctggctgaatgggcttgtccttatggag
tgccgcgttcttc-aaagaatccgtcttcatacagcgacctcatttacatattgccggct
gaatgggcttgtccttatggagcactgcattcttctaggaattgctatctatacagcgac
ctcatttacatacaaggcacct-ttttctacatagacttgccatttggctgattgttctg
acagtcgctgcctcttcggagttctgtgtggacgagtggcc--tcctgagtgggccggtc
cttacggagtgcccacgtctgttatgactgggttctccatacagcgacctcactcagaat
tgtatgg--------------caccgtgccgat-----------tgaatggacaagcagt
ccttagggaatcgcatgaaagcatgatggc-acatggtgtatggacccgttgggccttgc
ctgtgctatcttctcgagtcgaccgg------------tcctcgtctcctcctttcgcga
cgggaagctcttgagtttaccgtgggtcaggacgctgtggtatctgtttgtctgtagtgt
gggctctctatcagaacagcgtatcagcacacgtcgtctgggtctatggtatttttgctt
aacggccctgggtttggtagtatggttccatgtcggtcctctcacctcccatccgtcatg
tctctga-----------------------------------------------------
------------------------------------------------------------
--------------------------------------------tatcatctgcacagta
cgtcttgtcagaatatcattctgtatggcagtcgatagactaggccctatgccgcagctg
tctctgaccactatgcgagcgcatacctacctcaatgtgttcatagctgttatggatctt
gggtgtggtgtgccagaacgatacgccaccagattgtggaatggtgcttgtagtattaga
agaggacatgtatgaagagcacgttcttttctcttatgatggtgccgcgggtgggggccg
ttctcattctgcttgtcgggtgctcaaggtgcatgtaggattgctgaacccac-tgaatg
gtcatgagaatggtcatatggtgaggttcacttctccatcgctttgtgtgtgttggatgt
ttgggtggcttcggccgcatggaaaatctcggtctgtaactcatcagggagggtttccat
gtgagcaatctctccagtcgaacacacacccatgggcgtcccgctaggcaaaatggaact
ctcgatccgacacagcctgtggagtgtctgttccgtgtggcgtgaaacggcttatactgg
atgtcttccc-tttggacaccagtctgttggtatg--tacgcataccatctagagggagc
aagaaaaaggcatgcaagtgccacaaaggaagggaggcctccctgctaccgtgtgtcc-t
cttgacgatcacgggggagtatatagagacatttggacgagatcctttttttcacgaggcg
gtgccctcgcgcaaaggtgttgctctgcacgcatctttatgatgatagttgctcgctcct
atgcgtccactcgtgaaaatacctgctctctctgtctgtttccatgacacccc-catgcg
ttgtcatagagcacgctcaggtacagaaagtgctgtgactttcttggcctgctgtcactg
agtcagcctgatactttggaggaatcttttctgggcgctgcccagctgagagttcttatc
aggctctggagtgtgcatatggctataatcataatagtgcatgtggccacgcagaaaaag
catcgacgtggaatggagaagaacaccattctgaatggaacggttcttactctgcttccg
ttgcttgacgctgtatgggaatcgctttgccctgggggtacaccttccattccatgacaa
agcatttgatctatgccccga-gggtcattctctatgaagagcccgaaacttgtctgagt
gagaggcgaaggtagcccttcattt--atgcggggtccgccctcacactgtagtcggcac

agtcccttgtttatggactgcgcagctcaatcttcgactggagagg--aaaaacaaaaca

caagtatgacacgcacatacgtgttttctaggactgtgacctgtcggctgcctcgcggtg

gttggcatgatgccccaagtggggggggggagaaacaatcagtatgattgataaaaacca

gtgggg-aaaaaggcgcgctgtcgcacacccttggtcacggcctc---aagtcgacggcc

ctcactggccca-ccctgcgactcctgttcgcttttgacaagacacacctgcaata-cgg

aggaaatgcctgagcagctgtgccgcagcaactgcggacgcgtagcgatatgcatccctg

ttgtttgcgacttggtacaaagaacgtgcgccgggagacggattgccctttgacttttcc

acccagcccgcggctctatacccttgacgtacctatttacctggttgatcctgccagtag

tcatatgcttgtctcaaagactaagccatgcatgtctaagtataaacacttttatactcg

tgaaactgcgaatggctcattaaaacagttatagttcctttgggaacaacactacttgga

taaccgtagtaattctagagctaatacatgcctcaacgcccgactcacgaagggtggtat

ttattggataaaaaaccatcgtgctctgtccagagcgctttgagatgattcacaataact

tgtcggatcgcatggcctcgtgctggcgactcctcattcaaatttctgccctatcaactt

tcgatggtagagtattggtctaccatggtgtcgacgggtgacggggaattagggttcgat

tccggagagggagcctgagaaacggctaccacatccaaggaaggcagcaggcgcgcaaat

tacccaatcccgactcggggaggtagtgacaaaaaataacaatagggggccctttgggtc

ttctaattggaatgagaacaatttaaatcccttatcgaggatccattggagggcaagtct

ggt---------------------------------------------------------

------------------------------------------------------------

------------------------------------------------------------

------------------------------------------------------------

------------------------------------------------------------

------------------------------------------------------------

------------------------------------------------------------

------------------------------------------------------------

------------------------------------------------------------

------------------------------------------------------------

------------------------------------------------------------

------------------------------------------------------------

------------------------------------------------------------

------------------------------------------------------------

------------------------------------------------------------

------------------------------------------------------------

-----------------------------------------------gccagcagccgcg

gtaattccagctccaatagcgtatattaaagttgttgcagttaaaacgctcgtagtcgga

cctcggagcgcccccacgggccgggcttttgccttgttgccagtgggagtttgtctcctt

ttgtcgggacgcggtgtgcggggcctttactgtctctgcgcgtttacctgcgggccgacc
gtttactgtgaagaaagtaaagtgttcaaagcaggccaatgccttgaatatgtgagcatg
gaataatagaataggacttgggctctattttgttggtttccagtgaccaagtaatgatta
atagggacggttgggggcattcgtatttcattgtcagaggtgaaattcttggattgatgg
aagacgtacaactgcgaaagcatctgccatggatgttttcattgatcaagaacgaaagtt
aggggatcgaagacgatcagataccgtcgtagtcttaaccataaacgatgccgactgggg
attggcgggggcacttttatatgactctgtcagcaccctgagggaaaccaaagtctttgg
gttctgggggggagtatggtcgcaaggctgaaacttaaaggaattgacggaagggcaccac
caggagtggagcctgcggcttaatttgactcaacacgggaaaacttaccaggtccggaca
gaagaatgattgacagactgaaagctctttcttgatttttggttggtggtgcatggccg
ttcttagttggtggagtgatttgtctggttaattccgttaacgaacgagacctcgtcctg
ctaaataggtgcgcgcatgccagcactgcgttcttaccttcttagagggactatgcgcgt
ctagcgtatggaagattgaggcaataacaggtctgtgatgcccttagatgttctgggccg
cacgcgcgctacactgatgcattcaacgagtttcacttgatagtcctgggtcggaaggct
cgggtaatcttttgaaagtgcatcgtgctggggatagatcattgcaattattgatcttca
acgaggaattccttgtaggcgcaggtcatcagcctgcgccgaatacgtccctgccctttg
tacacaccgcccgtcgctcctaccgattgaatggtccgatgaaatgtccggatcgcggct
agtgtgatggactcgtccagagcggtagccgtgagaagctcattaaatcttatcatttag
aggaaggagaagtcgtaacaaggt------------------------------------
------------------------------------------------------------
------------------------------------------------------------
------------------------------------------------------------
------------------------------------------------------------
------------------------------------------------------------
------------------------------------------------------------
------------------------------------------------------------
------------------------------------------------------------
------------------------------------------------------------
------------------------------------------------------------
------------------------------------------------------------
------------------------------------------------------------
------------------------------------------------------------
------------------------------------------------------------
------------------------------------------------------------
-------------------------------ttccgtaggtgaacctgcggaaggatcat
tcacaaactattgacacaaacacacgcgaaccaaatcgtccgcacagcaggtggcaatga

aagagagactctgcatgtcgcctttcgggg--tagcaagcagcactcttctgccatcgcc
tctgtgcagggcgtaaattcccattgagaggatgtgagggcaccacaggaagcttttcca
caggaagtcgccatccttctccctccacggcggcg----cttctgtgcttggcag-----
----tttttttttgccttctagggaggatgccgccaatggagccccttat-atatataca
tcatcatatgccc--cttttttttcttaaccgcttgccacagcttcttctatgaggagctt
gtgggaagactgtctccatacaataac-aaagatacaactcttagcggtggatatcttgg
ttctcgcaacgatgaagaacgcagccaactgcgataactaatgtgaattgcaggacttcg
tgaatcattaagtctttgaacgcaagttgcgctcatctccatgaggaggtgagcatgcct
gtttgagtgtcgatgggaaaaacaatac-ataacaacaatcggctctcttatcttcctgc
cttcttgggggcaaaaggtagtgcatgccggagatgggctttgtcgcccctttctcttga
aagaggcgtcgtccgaaacacgtatcgttgcccgctggaggtcatagagcatgagttt--
aaagcaccacatctctcaagtcagagtgag-cgtcttgagtaggtcc-gcattgattcag
tttcaacccaggtggcgttggttacttgccat-aagcctctctggggtttctttgcttgg
acctctcttgtcagtttctctgacgtagagacaggt--gccgtctcatggacgttgccct
ctggaacgtgcta-aaaaaaaaaaaaaga----------aaaaaccgtaccacatgtcaa
aagaggcagatcgtgcttgacggcgggaggtgttgcatttcttgcttgtcctgagcgtct
gccagtctct----ga--cgagtg----tgttccttccataaacccgatctcaaatcagg
taggaccacccgctgaatttaagcatataattaagcggaggaaaagaaactaacaaggat
tccctcagtagcggcgagcgaagcgggaagagctcaacatgaaaaccgttgggggttcgc
gctttccttcgggttgtatagtaaagtggtgtcgcctgcgtcgcgccgggtataaatccc
gtctgaaatggggtgtcgtagagggttatatccccgtctgtgacccggtgtgcgatgcgt
tacggtgaccttacgatgagtcgggttgcttgggcatgcagccctaagtgggtggtaaac
ttcatctaaagctaaatactagcgagagaccgatagcgtacaagtaccgtaagggaaaga
tgcaaagaactttgaaaagagagttaaacagggcttgaaaccgttaagagtgaagcgaag
gaggtcagtgatcatggggccagatccccaagtgcctgtcata-gggcaccgtagcgttc
tggtttcccggggcagcgccggccgggggcgggggttaacagggaggtggttgagggatt
cgttttccgcctgtactcctttatagcttgcaccccggccgagggatccgcgacttgtcg
tgtgagggcgctgtcaaaatggccttctccaccccgtcttgaaacacggaccaaggagtc
taacatactcgcgagtgtttgggtggttaaaacccgaacgcacaatgaaagtaaacggtg
gctctgcgccgctgcttggtgggaaccctgcttttataatgggggcactatcacccgac
cccgatccctttggggtggttggttttgaggtagagcgagtatgatgggacccgaaagat
ggtgaactattcctgagtagggcgaagccagaggaaactctggtggaggctcgtagcgat
gctgacgtgcaaatcgctcgtcgaacttgggtataggggcgaaagactaatcgaaccatc
tagtagctggttccctccgaagtttccctcaggatagcgggagcaaatgcaagtcagtat
catccggtaaagcgaatgattagaggcctcgggggagtaactccttcgacctattctcaa
actttaaataggtgaggagccggcaacgttgcttaaaacaatgaacgccgtgcggacaat

gattgctctcagtgggccgattttggtaagcagaactggcgatgcgggatgaaccgaaag

cgacgctaaggtgccaaactcccagctcattcagacaccagaaaaggtgttgattcatac

agacagcaggacggtggccatggaagtcggaatccgctaaggagtgtgtaacaactcacc

tgccgaatggatcagccctgaaaatggatggcgcttaagcgggggaccgaggcgtcgccg

ttggggtaaatacgaagccccaatgagtaggagggcgtgggggttttgaaacacggtaca

ctcgcatggaagccttgggcgtaagctcgggtggagcgacccctagtgtagatcttggtg

gtagtagcaaatattcaagtgagaaccttgaagactgaagtggggaagggttccacgcga

acagtgcttggacgtgggttagtcggtcctaagagatggggaaactccgtattcaagtgc

gcttttgtagcgcagtccatcgaaagggaagcgggttaacattcccgcaccaggatgtgg

aaactgtggtttaccgcaaactgcggcaacgcaactgaacctggggacgcccgtaggagc

tccgagcagggatctctttcctgtttaacgcgcccaacccccctggaatcggtttgtccgg

agatagggggcgaaaggccggaaaagcaccgcacgtcttgtggtgttcggaacgcttgta

atggcccttgaaaacccaggggaaagcagccgtcttacac--ttgtgtgagacgaggaat

tctcacacctggccgtacccatcaccgcatcaggtccccaaggtgaacagcctctagtca

atagaataatgtagataagggaagtcggcaaaatagatccgtaactttgggaaaaggatt

ggctctaatgttcgggcttgtcgggacgcgcgggcgtagatgccaatgagcggcagacgt

ctagcctctttggaggctggattaacgctgcttgtgggaagaaaccgtgtcgatggaggc

aggaacgcttatcgatcttgtctcttcccgcttgcgatcaacgaacggattagaactggt

acggacaaggggaatccgactgtttaattaaaacaaagcattgcgacggccggaaacggt

gttggcgcaatgtgatttctgcccagtgctctgaatgtcaaagtgaagaaattcaaccaa

gcgcgggtaaacggcgggagtaactatgactct---------------------------

------------------------------------------------------------

------------------------------------------------------------

------------------------------------------------------------

------------------------------------------------------------

------------------------------------------------------------

------------------------------------------------------------

------------------------------------------------------------

------------------------------------------------------------

------------------------------------------------------------

------------------------------------------------------------

------------------------------------------------------------

------------------------------------------------------------

------------------------------------------------------------

------------------------------------------------------------

------------------------------------------------------------

------------------------------------------------------------
------------------------------------------------------------
------------------------------------------------------------
------------------------------------------------------------
------------------------------------------------------------
--------------------------------------------------------ctta
aggtagccaaatgcctcgtcatctaaactaattgg-at----------------------
-----------------------gagatagtc----------tgagtgaaaagcttatgc
ag-----------------gctagtaccggaa-aaaaaaaaaaaaagaggggctctgcca
ct----ctgac------------------------ggtgcgacactccttgtacagggaa
accctcc--gcatggaaaggcacgtggtaccaccctcgcggggaaactcgcgagcgggat
cccgggtaatgaccggagccgatggtaaaaactcacgtgaccgatggggtgaccctgtgg
cgagacgctaagttgtgggacatcaccctctcacaatacgcatccgtcgcagagactgaa
cgggagtgggctccgttga-----cggccggccggccagccagccgtcttcca-------
---ggggcttaagatacagtccgggcctcagggatgg----aaaaacctggggaacta-c
cagaattagtgacgcgcatgaatggattaacgagattcccactgtccctatctactatct
agcgaaaccacagccaagggaacgggcttggaagaatcagcggggaaagaagaccctgtt
gagcttgactctagtccgactttgtgagatgagttgggaggtgtagtctaagagggaggg
gcaactcgaccgtgaaatacctctactcccaagcttgttttacttactccggcgagcagg
gatgggtggtaacacccttttttctaggattaacacctcttcgggggtggacctgcgcgga
ggacattgtcaggtggggagtttggctggggcggcacatctgttaaaagataacgcaggt
gtcctaagatgagctcagtgagaacagaaatctcacgtagaacaaaagggtaaaagctca
tttgattttgattttcagtacgaatacaaactgtgaaagcatggcctatcgatccttttg
accttcaaggtttgaagcaagaggtgtcagaaaagttaacacgcgttgtggtagctttta
agagtggactcaaagtgttccgctagtccccccctcggaccgccacttacaaaaggcctgg
ggatgggggcgacactgtcaaattgcggggactccctaaagcctgtctctaccaagccag
cgcggcgaaagcccctggtggccggggtaatgacctagggtatggtaaaagcgagacagg
atg-gaacaatgggcaatccgcagccaagtcctaagg----ctttttgaaaagcggctat
ggatgcagttcaacgactagatggcagtgggctgagagagcccaccgcggcaccctttcg
gcttaagatatagtcttgcccacgtgaaagcgtgcctgcagatgaacgccaccatgtatg
gccggagctgcaggagaccgc--acttgatgtgcagtcgggaat-cgacagtaccacagg
gatg-aa-----------------------------------------------------
------------------------------------------------------------
------------------------------------------------------------
------------------------------------------------------------
------------------------------------------------------------

------------------------------------------------caaaaacac---
------------------------------------------------------------
------------------------------------------------------------
--acaatta---------------------------------------------------
----------------------------------------atttgtggcc---atcatac
----------------------------------------agtcgta-------------
------------------------------------------------------------
------------------------------------------------------------
------------------------------------------------------------
------------tgatattatcgtcccaggcgggtgcgcctctctcacccgtctagtccg
gccttggtgcgcaaaaaatgcttgcgtggatgcgccttggttaaggaaaatgctcctagt
agttttacaaggacacttcttgtagggcaacacatctggatgcgaagaactcccggaaaa
gacaacggtaccaagtgactggggtaacgccccattcatgaaggcaaataatcaaaaaga
ggtgccgggtaagaacccgttgtatagggacaatccgcagctgacttggctccctg--cc
cat--ccatgggcatagcttagcggttcaacgctcggtaagatgtgggttggcgcgctgt
ccgacagcgcactggcttaaggtacgggctactcccaccggagacggtgtctttgtctac
ctcgggtaacacgcctcgggagaagcaaagagggtgcaatggaaaaaaaactggttgcgg
ggaaatccctaccagagtcacctggtacaaaggaactggcttgtggcagccaagcgttca
tagcgacgttgctttttgatccttcgatgtcggctcttcctatcattgcgaagcagaatt
cgccaagtgttggattgttcacccactaatagggaacgtgagctgggtagccacttta--
-----tgcccaacagctggatagctgggcttctgtctta-ggccgggacttctcacgacc
cgtgccttggtcaccc-aaaaaagggcgaccgcagagctcaggatggagcttctggctta
actagtagaggacagatagagatcgtcaaattgcgggaaaccccttagagcctcgagcta
ccaagtgcgtgtcgaaaggcatagacgcagccagcgatggctgtcgcatggcttga-ttc
catcaagataggtaaaaatgctcgaggattgggcaatccgcagccaagcccctaaggt--
---ccccccgcgggctatggggaaggttcaacgactgtaaggcgtaccgtgcaagcggaa
gatacagtctagccccgctggaaagcagcgccaaagtaagtaggt-tgcagtgggtctgt
cctc
>Form_D_China
gcatgttttggtatgtgtttcgacagtcatgcatgtcatagccgtaggatggggagtgag
cttcatagcccgaccccttctcaggagcatgttttcatagcatattgagcatactggggt
ggtctcccatgtagctcgctctctcggtgcaggacctcccttttgcgggccacttcgatg
gcctccattatgga-gagcatgttttggcatatgaagcgttggtggtgaggctcgtagcc
tgtatatacataccaacgcagtcgtcatatgtctctctggcgggggtcggtcgtatgacc
ttgtccctacctggagcatgtatttcgtatgat---atactggac-ggcagctgcgctgt
tcagtcgtcggccgtacctcatgtgcgcgtttccaggtccgaacgtttgacatgcaaggg

catcataccagaagcctgttttctcgtatgagcatgtaggtaggcaggctcttcagtcgt
ctacccataaatgcacttcgaagga-actttttggtattgaatggctttgtgcccagacg
ttcttgtgagcttgttctcgctctgcgatacgctggaagctgtggtattacctacagcag
acgcattcctagggcggacggcctctctgcagatgggcacatctgtagaaagaggctccc
cttgatgacgcaatgcaatggctgggtaaacggagtgctatacagaagggtcaaacctc-
tgtggaagtacactcataccagcattgtttgttgagaggggtcggctctacgagtgagga
cctatactgccatctcattcagatgcgtagacggtctttactcagactgcctttctggct
gattgtcctcggtccctcgtgcgcacgcgccgctggctgaatgggcttgtccttatggag
tgccgcgttcttc-aaagaatccgtcttcatacagcgacctcatttacatattgccggct
gaatgggcttgtccttatggagcactgcattcttctaggaattgctatctatacagcgac
ctcatttacatacaaggcacct-ttttctacatagacttgccatttggctgattgttctg
acagtcgctgcctcttcggagttctgtgtggacgagtggcc--tcctgagtgggccggtc
cttacggagtgcccacgtctgttatgactgggttctccatacagcgacctcactcagaat
tgtatgg--------------caccgtgccgat-----------tgaatggacaagcagt
ccttagggaatcgcatgaaagcatgatggc-acatggtgtatggacccgttgggccttgc
ctgtgctatcttctcgagtcgaccgg------------tcctcgtctcctcctttcgcga
cgggaagctcttgagtttaccgtgggtcaggacgctgtggtatctgtttgtctgtagtgt
gggctctctatcagaacagcgtatcagcacacgtcgtctgggtctatggtatttttgctt
aacggccctgggtttggtagtatggttccatgtcggtcctctcacctcccatccgtcatg
tctctga-----------------------------------------------------
------------------------------------------------------------
--------------------------------------------tatcatctgcacagta
cgtcttgtcagaatatcattctgtatggcagtcgatagactaggccctatgccgcagctg
tctctgaccactatgcgagcgcatacctacctcaatgtgttcatagctgttatggatctt
gggtgtggtgtgccagaacgatacgccaccagattgtggaatggtgcttgtagtattaga
agaggacatgtatgaagagcacgttcttttctcttatgatggtgccgcgggtgggggccg
ttctcattctgcttgtcgggtgctcaaggtgcatgtaggattgctgaacccac-tgaatg
gtcatgagaatggtcatatggtgaggttcacttctccatcgctttgtgtgtgttggatgt
ttgggtggcttcggccgcatggaaaatctcggtctgtaactcatcagggagggtttccat
gtgagcaatctctccagtcgaacacacacccatgggcgtcccgctaggcaaaatggaact
ctcgatccgacacagcctgtggagtgtctgttccgtgtggcgtgaaacggcttatactgg
atgtcttccc-tttggacaccagtctgttggtatg--tacgcataccatctagagggagc
aagaaaaaggcatgcaagtgccacaaaggaagggaggcctccctgctaccgtgtgtcc-t
cttgacgatcacgggggagtatatagagacatttggacgagatcctttttttcacgaggcg
gtgccctcgcgcaaaggtgttgctctgcacgcatctttatgatgatagttgctcgctcct
atgcgtccactcgtgaaaatacctgctctctctgtctgtttccatgacacccc-catgcg

ttgtcatagagcacgctcaggtacagaaagtgctgtgactttcttggcctgctgtcactg
agtcagcctgatactttggaggaatcttttctgggcgctgcccagctgagagttcttatc
aggctctggagtgtgcatatggctataatcataatagtgcatgtggccacgcagaaaaag
catcgacgtggaatggagaagaacaccattctgaatggaacggttcttactctgcttccg
ttgcttgacgctgtatgggaatcgctttgccctgggggtacaccttccattccatgacaa
agcatttgatctatgccccga-gggtcattctctatgaagagcccgaaacttgtctgagt
gagaggcgaaggtagcccttcattt--atgcggggtccgccctcacactgtagtcggcac
agtcccttgtttatggactgcgcagctcaatcttcgactggagagg--aaaaacaaaaca
caagtatgacacgcacatacgtgttttctaggactgtgacctgtcggctgcctcgcggtg
gttggcatgatgccccaagtgggggggggagaaacaatcagtatgattgataaaaacca
gtgggg-aaaaaggcgcgctgtcgcacacccttggtcacggcctc---aagtcgacggcc
ctcactggccca-ccctgcgactcctgttcgcttttgacaagacacacctgcaata-cgg
aggaaatgcctgagcagctgtgccgcagcaactgcggacgcgtagcgatatgcatccctg
ttgtttgcgacttggtacaaagaacgtgcgccgggagacggattgccctttgacttttcc
acccagcccgcggctctatacccttgacgtacctatttacctggttgatcctgccagtag
tcatatgcttgtctcaaagactaagccatgcatgtctaagtataaacacttttatactcg
tgaaactgcgaatggctcattaaaacagttatagttcctttgggaacaacactacttgga
taaccgtagtaattctagagctaatacatgcctcaacgcccgactcacgaagggtggtat
ttattggataaaaaaccatcgtgctctgtccagagcgctttgagatgattcacaataact
tgtcggatcgcatggcctcgtgctggcgactcctcattcaaatttctgccctatcaactt
tcgatggtagagtattggtctaccatggtgtcgacgggtgacggggaattagggttcgat
tccggagagggagcctgagaaacggctaccacatccaaggaaggcagcaggcgcgcaaat
tacccaatcccgactcggggaggtagtgacaaaaaataacaatagggggccctttgggtc
ttctaattggaatgagaacaatttaaatcccttatcgaggatccattggagggcaagtct
ggt---------------------------------------------------------
------------------------------------------------------------
------------------------------------------------------------
------------------------------------------------------------
------------------------------------------------------------
------------------------------------------------------------
------------------------------------------------------------
------------------------------------------------------------
------------------------------------------------------------
------------------------------------------------------------
------------------------------------------------------------
------------------------------------------------------------

------------------------------------------------------------
------------------------------------------------------------
------------------------------------------------------------
------------------------------------------------------------
-----------------------------------------------gccagcagccgcg
gtaattccagctccaatagcgtatattaaagttgttgcagttaaaacgctcgtagtcgga
cctcggagcgcccccacgggccgggcttttgccttgttgccagtgggagtttgtctcctt
ttgtcgggacgcggtgtgcagggcctttactgtctctgcgcgtttacctgcgggccgacc
gtttactgtgaagaaagtaaagtgttcaaagcaggccaatgccttgaatatgtgagcatg
gaataatagaataggacttgggctctattttgttggtttccagtgaccaagtaatgatta
atagggacggttgggggcattcgtatttcattgtcagaggtgaaattcttggattgatgg
aagacgtacaactgcgaaagcatctgccatggatgttttcattgatcaagaacgaaagtt
aggggatcgaagacgatcagataccgtcgtagtcttaaccataaacgatgccgactgggg
attggcgggggcacttttatatgactctgtcagcaccctgagggaaaccaaagtctttgg
gttctggggggagtatggtcgcaaggctgaaacttaaaggaattgacggaagggcaccac
caggagtggagcctgcggcttaatttgactcaacacgggaaaacttaccaggtccggaca
gaagaatgattgacagactgaaagctctttcttgattttttggttggtggtgcatggccg
ttcttagttggtggagtgatttgtctggttaattccgttaacgaacgagacctcgtcctg
ctaaataggtgcgcgcatgccagcactgcgttcttaccttcttagagggactatgcgcgt
ctagcgtatggaagattgaggcaataacaggtctgtgatgcccttagatgttctgggccg
cacgcgcgctacactgatgcattcaacgagtttcacttgatagtcctgggtcggaaggct
cgggtaatcttttgaaagtgcatcgtgctggggatagatcattgcaattattgatcttca
acgaggaattccttgtaggcgcaggtcatcagcctgcgccgaatacgtccctgccctttg
tacacaccgcccgtcgctcctaccgattgaatggtccgatgaaatgtccggatcgcggct
agtgtgatggactcgtccagagcggtagccgtgagaagctcattaaatcttatcatttag
aggaaggagaagtcgtaacaaggt------------------------------------
------------------------------------------------------------
------------------------------------------------------------
------------------------------------------------------------
------------------------------------------------------------
------------------------------------------------------------
------------------------------------------------------------
------------------------------------------------------------
------------------------------------------------------------
------------------------------------------------------------
------------------------------------------------------------

------------------------------------------------------------

------------------------------------------------------------

------------------------------------------------------------

------------------------------------------------------------

------------------------------------------------------------

-------------------------------ttccgtaggtgaacctgcggaaggatcat

tcacaaactattgacacaaacacacgcgaaccaaatcgtccgcacagcaggtggcaatga

aagagagactctgcatgtcgcctttcgggg--tagcaagcagcactcttctgccatcgcc

tctgtgcagggcgtaaattcccattgagaggatgtgagggcaccacaggaagcttttcca

caggaagtcgccatccttctccctccacggcggcg----cttctgtgcttggcag-----

----tttttttttgccttctagggaggatgccgccaatggagccccttat-atatataca

tcatcatatgccc--cttttttttcttaaccgcttgccacagcttcttctatgaggagctt

gtgggaagactgtctccatacaataac-aaagatacaactcttagcggtggatatcttgg

ttctcgcaacgatgaagaacgcagccaactgcgataactaatgtgaattgcaggacttcg

tgaatcattaagtctttgaacgcaagttgcgctcatctccatgaggaggtgagcatgcct

gtttgagtgtcgatgggaaaaacaatac-ataacaacaatcggctctcttatcttcctgc

cttcttgggggcaaaaggtagtgcatgccggagatgggctttgtcgcccctttctcttga

aagaggcgtcgtccgaaacacgtatcgttgcccgctggaggtcatagagcatgagttt--

aaagcaccacatctctcaagtcagagtgag-cgtcttgagtaggtcc-gcattgattcag

tttcaacccaggtggcgttggttacttgccat-aagcctctctggggtttctttgcttgg

acctctcttgtcagtttctctgacgtagagacaggt--gccgtctcatggacgttgccct

ctggaacgtgcta-aaaaaaaaaaaaaga----------aaaaaccgtaccacatgtcaa

aagaggcagatcgtgcttgacggcgggaggtgttgcatttcttgcttgtcctgagcgtct

gccagtctct----ga--cgagtg----tgttccttccataaacccgatctcaaatcagg

taggaccacccgctgaatttaagcatataattaagcggaggaaaagaaactaacaaggat

tccctcagtagcggcgagcgaagcgggaagagctcaacatgaaaaccgttgggggttcgc

gctttccttcgggttgtatagtaaagtggtgtcgcctgcgtcgcgccgggtataaatccc

gtctgaaatggggtgtcgtagagggttatatccccgtctgtgacccggtgtgcgatgcgt

tacggtgaccttacgatgagtcgggttgcttgggcatgcagccctaagtgggtggtaaac

ttcatctaaagctaaatactagcgagagaccgatagcgtacaagtaccgtaagggaaaga

tgcaaagaactttgaaaagagagttaaacagggcttgaaaccgttaagagtgaagcgaag

gaggtcagtgatcatggggccagatccccaagtgcctgtcata-gggcaccgtagcgttc

tggtttcccggggcagcgccggccgggggcgggggttaacagggaggtggttgagggatt

cgttttccgcctgtactcctttatagcttgcaccccggccgagggatccgcgacttgtcg

tgtgagggcgctgtcaaaatggccttctccaccccgtcttgaaacacggaccaaggagtc

taacatactcgcgagtgtttgggtggttaaaacccgaacgcacaatgaaagtaaacggtg

gctctgcgccgctgcttggtgggaaccctgcttttataatgggggcactatcacccgac
cccgatccctttggggtggttggttttgaggtagagcgagtatgatgggacccgaaagat
ggtgaactattcctgagtagggcgaagccagaggaaactctggtggaggctcgtagcgat
gctgacgtgcaaatcgctcgtcgaacttgggtataggggcgaaagactaatcgaaccatc
tagtagctggttccctccgaagtttccctcaggatagcgggagcaaatgcaagtcagtat
catccggtaaagcgaatgattagaggcctcgggggagtaactccttcgacctattctcaa
actttaaataggtgaggagccggcaacgttgcttaaaacaatgaacgccgtgcggacaat
gattgctctcagtgggccgattttggtaagcagaactggcgatgcgggatgaaccgaaag
cgacgctaaggtgccaaactcccagctcattcagacaccagaaaaggtgttgattcatac
agacagcaggacggtggccatggaagtcggaatccgctaaggagtgtgtaacaactcacc
tgccgaatggatcagccctgaaaatggatggcgcttaagcgggggaccgaggcgtcgccg
ttggggtaaatacgaagccccaatgagtaggagggcgtgggggttttgaaacacggtaca
ctcgcatggaagccttgggcgtaagctcgggtggagcgacccctagtgtagatcttggtg
gtagtagcaaatattcaagtgagaaccttgaagactgaagtggggaagggttccacgcga
acagtgcttggacgtgggttagtcggtcctaagagatggggaaactccgtattcaagtgc
gcttttgtagcgcagtccatcgaaagggaagcgggttaacattcccgcaccaggatgtgg
aaactgtggtttaccgcaaactgcggcaacgcaactgaacctggggacgcccgtaggagc
tccgagcagggatctctttcctgtttaacgcgcccaacccccctggaatcggtttgtccgg
agatagggggcgaaaggccggaaaagcaccgcacgtcttgtggtgttcggaacgcttgta
atggcccttgaaaacccaggggaaagcagccgtcttacac--ttgtgtgagacgaggaat
tctcacacctggccgtacccatcaccgcatcaggtccccaaggtgaacagcctctagtca
atagaataatgtagataagggaagtcggcaaaatagatccgtaactttgggaaaaggatt
ggctctaatgttcgggcttgtcgggacgcgcgggcgtagatgccaatgagcggcagacgt
ctagcctctttggaggctggattaacgctgcttgtgggaagaaaccgtgtcgatggaggc
aggaacgcttatcgatcttgtctcttcccgcttgcgatcaacgaacggattagaactggt
acggacaaggggaatccgactgtttaattaaaacaaagcattgcgacggccggaaacggt
gttggcgcaatgtgatttctgcccagtgctctgaatgtcaaagtgaagaaattcaaccaa
gcgcgggtaaacggcgggagtaactatgactct--taaacaaaaccatactacctac---
-------------------gggagtcaaaagtgggggcggcgtctggtgtagaaatacac
gtac--------------------------------------------------------
-------------------gacacgctttttttcctgctagtcgatctgacctttgttct
ttggactgtgg-------------------------------------------------
--------------------------gcggtcagcggcaacataacc-------------
------------------------------------------------------------
------------------------------------------------------------
----------------------------tggtacggg---------------------ga

aggcctcgacgttga---------------------------------------------
--------------------------------gtgctgcatcttggaagtatc-------
-----------------------------ttgacacaataagaga---------------
------------------------------aaagagattgcgcttaatagg---------
----gggttgatctcgt-------------------------------------------
-------------------------------------------ggcgagccgacgc----
---------------------------------cggtgtgaactggt-------------
------------------------------------------------------------
-----------------------ggcaat-------------------------------
----ggccgtcgtagagcacgcaaaggtatgggctgactttttctcctca----aaagtt
ggcttaaggac-gtgccatcccacacgataataaagtgtgcagtctg-------------
--------------------taagagctcccacagggcaaagac---------agactgg
gatgcagacggagacggcaaaaacaacaacatccgttctgcttctgggcgcaaatgctta
aggtagccaaatgcctcgtcatctaaactaattgg-at----------------------
-----------------------gagatagtc----------tgagtgaaaagcttatgc
ag-----------------gctagtaccggaa-aaaaaaaaaaaaagaggggctctgcca
ct----ctgac------------------------ggtgcgacactccttgtacagggaa
accctcc--gcatggaaaggcacgtggtaccaccctcgcggggaaactcgcgagcgggat
cccgggtaatgaccggagccgatggtaaaaactcacgtgaccgatggggtgaccctgtgg
cgagacgctaagttgtgggacatcaccctctcacaatacgcatccgtcgcagagactgaa
cgggagtgggctccgttgac-ggccggccggccggccagccagccgtcttcca-------
---ggggcttaagatacagtccgggcctcagggatgg----aaaaacctggggaacta-c
cagaattagtgacgcgcatgaatggattaacgagattcccactgtccctatctactatct
agcgaaaccacagccaagggaacgggcttggaagaatcagcggggaaagaagaccctgtt
gagcttgactctagtccgactttgtgagatgagttgggaggtgtagtctaagagggaggg
gcaactcgaccgtgaaatacctctactcccaagcttgttttacttactccggcgagcagg
gatgggtggtaacaccctttttctaggattaacacctcttcgggggtggacctgcgcgga
ggacattgtcaggtggggagtttggctggggcggcacatctgttaaaagataacgcaggt
gtcctaagatgagctcagtgagaacagaaatctcacgtagaacaaaagggtaaaagctca
tttgattttgattttcagtacgaatacaaactgtgaaagcatggcctatcgatcctttttg
accttcaaggtttgaagcaagaggtgtcagaaaagttaacacgcgttgtggtagctttta
agagtggactcaaagtgttccgctagtccccccctcggaccgccacttacaaaaggcctgg
ggatgggggcgacactgtcaaattgcggggactccctaaagcctgtctctaccaagccag
cgcggcgaaagcccctggtggccggggtaatgacctagggtatggtaaaagcgagacagg
atg-gaacaatgggcaatccgcagccaagtcctaagg----cttttgaaaagcggctat
ggatgcagttcaacgactagatggcagtgggctgagagagcccaccgcggcacccctttcg

gcttaagatatagtcttgcccacgtgaaagcgtgcctgcagatgaacgccaccatgtatg
gccggagctgcaggagaccgc--acttgatgtgcagtcgggaat-cgacagtaccacagg
gatgaaa-----------------------------------------------------
------------------------------------------------------------
------------------------------------------------------------
------------------------------------------------------------
------------------------------------------------------------
------------------------------------------------aaaaaacac---
------------------------------------------------------------
------------------------------------------------------------
--acaatta---------------------------------------------------
----------------------------------------atttgtggcc---atcatac
----------------------------------------agtcgta-------------
------------------------------------------------------------
------------------------------------------------------------
------------------------------------------------------------
------------tgatattatcgtcccaggcgggtgcgcctctctcacccgtctagtccg
gccttggtgcgcaaaaaatgcttgcgtggatgcgccttggttaaggaaaatgctcctagt
agttttacaaggacacttcttgtagggcaacacatctggatgcgaagaactcccggaaaa
gacaacggtaccaagtgactggggtaacgccccattcatgaaggcaaataatcaaaaaga
ggtgccgggtaagaacccgttgtatagggacaatccgcagctgacttggctccctg--cc
cat--ccatgggcatagcttagcggttcaacgctcggtaagatgtgggttggcgcgctgt
ccgacagcgcactggcttaaggtacgggctactcccaccggagacggtgtctttgtctac
ctcgggtaacacgcctcgggagaagcaaagagggtgcaatggaaaaaaaactggttgcgg
ggaaatccctaccagagtcacctggtacaaaggaactggcttgtggcagccaagcgttca
tagcgacgttgctttttgatccttcgatgtcggctcttcctatcattgcgaagcagaatt
cgccaagtgttggattgttcacccactaatagggaacgtgagctgggtagccacttta--
-----tgcccaacagctggatagctgggcttctgtctta-ggccgggacttctcacgacc
cgtgccttggtcaccc-aaaaaagggcgaccgcagagctcaggatggagcttctggctta
actagtagaggacagatagagatcgtcaaattgcgggaaaccccttagagcctcgagcta
ccaagtgcgtgtcgaaaggcatagacgcagccagcgatggctgtcgcatggcttga-ttc
catcaagataggtaaaaatgctcgaggattgggcaatccgcagccaagcccctaaggt--
---ccccccgcgggctatggggaaggttcaacgactgtaaggcgtaccgtgcaagcggaa
gatacagtctagccccgctggaaagcagcgccaaagtaagtaggt-tgcagtgggtctgt
cctc
>Form_B_China_SRR9587936

gcatgttttggtatgtgtttcgacagtcatgcatgtcatagccgtaggatggggagtgag
cttcatagcccgaccccttctcaggagcatgttttcatagcatattgagcatactggggt
ggtctcccatgtagctcgctctctcggtgcaggacctccctttttgcgggccacttcgatg
gcctccattatgga-gagcatgttttggcatatgaagcgttggtggtgaggctcgtagcc
tgtatatacataccaacgcagtcgtcatatgtctctctggcgggggtcggtcgtatgacc
ttgtccctacctggagcatgtattttgtatgat---atactggac-ggcagctgcgctgt
tcagtcgtcggccgtacctcatgtgcgcgtttccaggtccgaacgtttgacatgcaaggg
catcataccagaagcctgttttctcgtatgagcatgtaggtaggcaggctcttcagtcgt
ctacccataaatgcacttcgaagga-actttttggtattgaatggctttgtgcccagacg
ttcttgtgagcttgttctcgctctgcgatacgctggaagctgtggtattacctacagcag
acgcattcctagggcggacggcctctctgcagatgggcacatctgtagaaagaggctccc
cttgatgacgcaatgcaatggctgggtaaacggagtgctatacagaagggtcaaacctc-
tgtggaagtacacccataccagcattgtttgttgagaggggtcggctctacgagtgagga
cctatactgccatctcattcagatgcgtagacggtctttactcagactgcctttctggct
gattgtcctcggtccctcgtgcgcacgcgccgctggctgaatgggcttgtccttatggag
tgccgcgttcttc-aaagaatccgtcttcatacagcgacctcatttacatattgccggct
gaatgggcttgtccttatggagcactgcattcttctaggaattgctatctatacagcgac
ctcatttacatacaaggcacct-ttttctacatagacttgccatttggctgattgttctg
acagtcgctgcctcttcggagttctgtgtggacgagtggcc--tcctgagtgggccggtc
cttacggagtgcccacgtctgttatgactgggttctccatacagcgacctcactcagaat
tgtatgg--------------caccgtgccgat-----------tgaatggacaagcagt
ccttagggaatcgcatgaaagcatgacggc-acatggtgtatggacccgttgggccttgc
ctgtgctatcttctcgagtcgaccgg------------tcctcgtctcctcctttcgcga
cgggaagctcttgagtttaccgtgggtcaggacgctgtggtatctgtttgtctgtagtgt
gggctctctatcagaacagcgtatcagcacacgtcgtctgggtctatggtatttttgctt
aacggccctgggtttggtagtatggttccatgtcggtcctctcacctcccaaccgtcatg
tctctga-----------------------------------------------------
------------------------------------------------------------
--------------------------------------------tatcatctgcacagta
cgtcttgtcagaatatcattctgtatggcagtcgatagactaggccctatgccgcagctg
tctctgaccactatgcgagcgcatacctacctcaatgtgttcatagctgttatggatctt
gggtgtggtgtgccagaacgatacgccaccagattgtggaatggtgcttgtagtattaga
agaggacatgtatgaagagcacgttcttttctcttatgatggtgccgcgggtgcgggccg
ttttcattctgcttgtcgggtgctcaaggtgcatgtaggattgctgaacccac-tgaatg
gtcatgagaatggtcatatggtgaggttcacttctccatcgctttgtgtgtgttggatgt
ttgggtggcttcggccgcatggaaaatctcggtctgtaactcatcagggagggtttccat

gtgagcaatctctccagtcgaacacacacccatgggcgtcccgctaggcaaaatggaact
ctcgatccgacacagcctgtggagtgtttgttccgtgtggcgtgaaacggcttatactgg
atgtcttccc-tttggacaccagtctgttggtatg--tacgcataccatctagagggagc
aagaaaaaggcatgcaagtgccacaaaggaagggaggccttcctgctaccgtgtgtcc-t
cttgacgatcacgggggagtatatagagacatttggacgagatccgtttttcacgaggcg
gtgccctcgcgcaaaggtgttgctctgcacgcatctttatgatgatagttgctcgctcct
atgcgtccactcgtgaaaatacctgctctctctgtctgtttccatgacacccc-catgcg
ttgtcatagagcatgctcaggtacagaaagtgctgtgactttcttggcctgctgtcactg
agtcagcctgatactttggaggaatcttttctgggcactgcccagctgagagttcttatc
aggctctggagtgtgcatatgcttatattcattatagtgcatgtggctacgcagaaaaag
catcgacgtggaatggagaagaacaccattctgaatggaacggttcttactctacttctg
ttgcttgacgctgtatgggaatcgctttgctctgggggtacaccttccattccatgacaa
agcatttgatctatgccccga-gggtcattctctatgaagagcccgaaacttgtctgagt
gagaggcgaaggtagtccttcattt--atgcggggtccgccctcacactgtagtcggcac
agtcccttgtttatggactgcgcagctcaatcttcgactggagagg--aaaaacaaaaca
caagtatgacacgcacatacgtgttttctaggactgtgacctgtcggctgcctcgcggtg
gttggcatgatgccccaagt-gggggggggagaaacaatcagtatgattgataaaaacca
gtgggg-aaaaaggcgcgctgtcgcacacccttggtcacggcctc---aagtcgacggcc
ctcactggccca-ccctgcgactcctgttcgcttttgacaagacacacctgcaata-cgg
aggaaatgcctgagcagctgtgccgcagcaactgcggacgcgtagcgatatgcatccctg
ttgtttgcgacttggtacaaagaacgtgcgccgggagacggattgccctttgacttttcc
acccagcccgcggctctatacccttgacgtacctatttacctggttgatcctgccagtag
tcatatgcttgtctcaaagactaagccatgcatgtctaagtataaacacttttatactcg
tgaaactgcgaatggctcattaaaacagttatagttcctttgggaacaacactacttgga
taaccgtagtaattctagagctaatacatgcctcaacgcccgactcacgaagggtggtat
ttattggataaaaaaccatcgtgctctgtccagagcgctttgagatgattcacaataact
tgtcggatcgcatggcctcgtgctggcgactcctcattcaaatttctgccctatcaactt
tcgatggtagagtattggtctaccatggtgtcgacgggtgacggggaattagggttcgat
tccggagagggagcctgagaaacggctaccacatccaaggaaggcagcaggcgcgcaaat
tacccaatcccgactcggggaggtagtgacaaaaaataacaatagggggccctttgggtc
ttctaattggaatgagaacaatttaaatcccttatcgaggatccattggagggcaagtct
ggtgaacgta----------------------------------tgcatca---------
--aagccctccgtaat-aaaaaaaaacagcgccagatgtct-------------------
------------------------------------------------------------
------------------------------------------------------------
------------------------------------------------------------

------------------------------------------------------------
------------------------------------------------------------
-------ctctctggaagcacacca-----------------------------------
---cgcacggggttcgttcctctc-----------------------------tgcatgg
ccgtg-----------------------------------------tgaaaaaggcgagt
aggcaaacgatgggaaacccggcttggaatagtaaccagccacagagcccccatcataca
cttctaaattgccggggagtgcccgcgaggccttgactaccgcggtccgccgagaggtga
gagcgagcaatcgctcaggacgcagcagtgtacgtagtggtacatggatggtaaaaacgt
caaggatagggttgatccgcagggaagcgcgagtgccgccatg---------gtcggtgg
gcaaaggcgaaccttcagagactctaatggagtgggcgcaagcttaagggagagtccaat
ccactgggaaaccaggtccacagcagtaacggggtgatgctcgccccagaggactgtgga
aggcaaccccacc----aaaagggtgtgtgttgtcgggtagc-aatggccagcagccgcg
gtaattccagctccaatagcgtatattaaagttgttgcagttaaaacgctcgtagtcgga
cctcggagcgcccccacgggccgggcttttgccttgttgccagtgggagtttgtctcctt
ttgtcgggacgcggtgtgcagggcctttactgtctctgcgcgtttacctgcgggccgacc
gtttactgtgaagaaagtaaagtgttcaaagcaggccaatgccttgaatatgtgagcatg
gaataatagaataggacttgggctctattttgttggttttcagtgaccaagtaatgatta
atagggacggttgggggcattcgtatttcattgtcagaggtgaaattcttggattgatgg
aagacgtacaactgcgaaagcatctgccatggatgttttcattgatcaagaacgaaagtt
aggggatcgaagacgatcagataccgtcgtagtcttaaccataaacgatgccgactgggg
attggcgggggcacttttatatgactctgtcagcaccctgagggaaaccaaagtctttgg
gttctgggggagtatggtcgcaaggctgaaacttaaaggaattgacggaagggcaccac
caggagtggagcctgcggcttaatttgactcaacacgggaaaacttaccaggtccggaca
gaagaatgattgacagactgaaagctctttcttgatttttggttggtggtgcatggccg
ttcttagttggtggagtgatttgtctggttaattccgttaacgaacgagacctcgtcctg
ctaaataggtgcgcgcatgccagcactgcgttcttaccttcttagagggactatgcgcgt
ctagcgtatggaagattgaggcaataacaggtctgtgatgcccttagatgttctgggccg
cacgcgcgctacactgatgcattcaacgagtttcacttgatagtcctgggtcggaaggct
cgggtaatcttttgaaagtgcatcgtgctggggatagatcattgcaattattgatcttca
acgaggaattccttgtaggcgcaggtcatcagcctgcgccgaatacgtccctgccctttg
tacacaccgcccgtcgctcctaccgattgaatggtccgatgaaatgtccggatcgcggct
agtgtgatggactcgtccagagcggtagccgtgagaagctcattaaatcttatcatttag
aggaaggagaagtcgtaacaaggt------------------------------------
------------------------------------------------------------
------------------------------------------------------------
------------------------------------------------------------

------------------------------------------------------------
------------------------------------------------------------
------------------------------------------------------------
------------------------------------------------------------
------------------------------------------------------------
------------------------------------------------------------
------------------------------------------------------------
------------------------------------------------------------
------------------------------------------------------------
------------------------------------------------------------
------------------------------------------------------------
------------------------------------------------------------
-------------------------------ttccgtaggtgaacctgcggaaggatcat
tcacaaactattgacacaaacacacgcgaaccaaatcgtccgcacagcaggtggcaatga
aagagagactctgcatgtcgcctttcgggg--tagcaagcagcactcttctgccatcgcc
tctgtgcagggcgtaaattcccattgagaggatgtgagggcaccacaggaagcttttcca
caggaagtcgccatccttctccctccacggcggcg----cttctgtgcttggcag-----
----tttttttttgccttctagggaggatgccgccaatggagccccttat-atatataca
tcatcatatgccc--cttttttttcttaaccgcttgccacagcttcttctatgaggagctt
gtgggaagactgtctccatacaataac-aaagatacaactcttagcggtggatatcttgg
ttctcgcaacgatgaagaacgcagccaactgcgataactaatgtgaattgcaggacttcg
tgaatcattaagtctttgaacgcaagttgcgctcatctccatgaggaggtgagcatgcct
gtttgagtgtcgatgggaaaaacaatac-ataacaacaatcggctctcttatcttcctgc
cttcttgggggcaaaaggtagtgcatgccggagatgggctttgtcgcccctttctcttga
aagaggcgtcgtccgaaacacgtatcgttgcccgctggaggtcatagagcatgagttt--
aaagcaccacatctctcaagtcagagtgag-cgtcttgagtaggtcc-gcattgattcag
tttcaacccaggtggcgttggttacttgccat-aagcctctctggggtttctttgcttgg
acctctcttgtcagtttctctgacgtagagacaggt--gccgtctcatggacgttgccct
ctggaacgtgcta-aaaaaaaaaaaaaga----------aaaaaccgtaccacatgtcaa
aagaggcagatcgtgcttgacggcgggaggtgttgcatttcttgcttgtcctgagcgtct
gccagtctct----ga--cgagtg----tgttccttccataaacccgatctcaaatcagg
taggaccacccgctgaatttaagcatataattaagcggaggaaaagaaactaacaaggat
tccctcagtagcggcgagcgaagcgggaagagctcaacatgaaaaccgttgggggttcgc
gctttccttcgggttgtatagtaaagtggtgtcgcctgcgtcgcgccgggtataaatccc
gtctgaaatggggtgtcgtagagggttatatccccgtctgtgacccggtgtgcgatgcgt
tacggtgaccttacgatgagtcgggttgcttgggcatgcagccctaagtgggtggtaaac

ttcatctaaagctaaatactagcgagagaccgatagcgtacaagtaccgtaagggaaaga
tgcaaagaactttgaaaagagagttaaacagggcttgaaaccgttaagagtgaagcgaag
gaggtcagtgatcatggggccagatccccaagtgcctgtcata-gggcaccgtagcgttc
tggtttcccggggcagcgccggccgggggcgggggttaacagggaggtggttgagggatt
cgttttccgcctgtactcctttatagcttgcaccccggccgagggatccgcgacttgtcg
tgtgagggcgctgtcaaaatggccttctccaccccgtcttgaaacacggaccaaggagtc
taacatactcgcgagtgtttgggtggttaaaacccgaacgcacaatgaaagtaaacggtg
gctctgcgccgctgcttggtgggaaccctgcttttataatgggggcactatcacccgac
cccgatccctttggggtggttggttttgaggtagagcgagtatgatgggacccgaaagat
ggtgaactattcctgagtagggcgaagccagaggaaactctggtggaggctcgtagcgat
gctgacgtgcaaatcgctcgtcgaacttgggtataggggcgaaagactaatcgaaccatc
tagtagctggttccctccgaagtttccctcaggatagcgggagcaaatgcaagtcagtat
catccggtaaagcgaatgattagaggcctcgggggagtaactccttcgacctattctcaa
actttaaataggtgaggagccggcaacgttgcttaaaacaatgaacgccgtgcggacaat
gattgctctcagtgggccgattttggtaagcagaactggcgatgcgggatgaaccgaaag
cgacgctaaggtgccaaactcccagctcattcagacaccagaaaaggtgttgattcatac
agacagcaggacggtggccatggaagtcggaatccgctaaggagtgtgtaacaactcacc
tgccgaatggatcagccctgaaaatggatggcgcttaagcgggggaccgaggcgtcgccg
ttggggtaaatacgaagccccaatgagtaggagggcgtgggggttttgaaacacggtaca
ctcgcatggaagccttgggcgtaagctcgggtggagcgacccctagtgtagatcttggtg
gtagtagcaaatattcaagtgagaaccttgaagactgaagtggggaagggttccacgcga
acagtgcttggacgtgggttagtcggtcctaagagatggggaaactccgtattcaagtgc
gcttttgtagcgcagtccatcgaaagggaagcgggttaacattcccgcaccaggatgtgg
aaactgtggtttaccgcaaactgcggcaacgcaactgaacctggggacgcccgtaggagc
tccgagcagggatctctttcctgtttaacgcgcccaacccccctggaatcggtttgtccgg
agataggggggcgaaaggccggaaaagcaccgcacgtcttgtggtgttcggaacgcttgta
atggcccttgaaaacccaggggaaagcagccgtcttacac--ttgtgtgagacgaggaat
tctcacacctggccgtacccatcaccgcatcaggtccccaaggtgaacagcctctagtca
atagaataatgtagataagggaagtcggcaaaatagatccgtaactttgggaaaaggatt
ggctctaatgttcgggcttgtcgggacgcgcgggcgtagatgccaatgagcggcagacgt
ctagcctctttggaggctggattaacgctgcttgtgggaagaaaccgtgtcgatggaggc
aggaacgcttatcgatcttgtctcttcccgcttgcgatcaacgaacggattagaactggt
acggacaaggggaatccgactgtttaattaaaacaaagcattgcgacggccggaaacggt
gttggcgcaatgtgatttctgcccagtgctctgaatgtcaaagtgaagaaattcaaccaa
gcgcgggtaaacggcgggagtaactatgactct--taaacaaaaccatactacctac---
-------------------gggagtcaaaagtgggggcggcgtctggtgtagaaatacac

gtac--------------------------------------------------------
-------------------gacacgctttttttcctgctagtcgatctgacctttgttct
ttggactgtgg-------------------------------------------------
--------------------------gcggtcagcggcaacataacc-------------
------------------------------------------------------------
------------------------------------------------------------
----------------------------tggtacggg---------------------ga
aggcctcgacgttga---------------------------------------------
--------------------------------gtgctgcatcttggaagtatc-------
-----------------------------ttgacacaataagaga---------------
------------------------------aaagagattgcgcttaatagg---------
----gggttgatctcgt-------------------------------------------
-------------------------------------------ggcgagccgacgc----
---------------------------------cggtgtgaactggt-------------
------------------------------------------------------------
-----------------------ggcaat-------------------------------
----ggccgtcgtagagcacgcaaaggtatgggctgactttttctcctca----aaagtt
ggcttaagggac-gtgccatcccacacgataataaagtgtgcagtctg------------
--------------------taagagctcccacagggcaaagac---------agactgg
gatgcagacggagacggcaaaaacaacaacatccgttctgcttctgggcgcaaatgctta
aggtagccaaatgcctcgtcatctaaactaattgg-at----------------------
-----------------------gagatagtc----------tgagtgaaaagcttatgc
ag-----------------gctagtaccggaa-aaaaaaaaaaaaagaggggctctgcca
ct----ctgac------------------------ggtgcgacactccttgtacagggaa
accctcc--gcatggaaaggcacgtggtaccaccctcgcggggaaactcgcgagcgggat
cccgggtaatgaccggagccgatggtaaaaactcacgtgaccgatggggtgaccctgtgg
cgagacgctaagttgtgggacatcaccctctcacaatacgcatccgtcgcagagactgaa
cgggagtgggctccgttgac-ggccggccggccggccagccagccgtcttcca-------
---ggggcttaagatacagtccgggcctcagggatgg----aaaaacctggggaacta-c
cagaattagtgacgcgcatgaatggattaacgagattcccactgtccctatctactatct
agcgaaaccacagccaagggaacgggcttggaagaatcagcggggaaagaagaccctgtt
gagcttgactctagtccgactttgtgagatgagttgggaggtgtagtctaagagggaggg
gcaactcgaccgtgaaatacctctactcccaagcttgttttacttactccggcgagcagg
gatgggtggtaacacccttttttctaggattaacacctcttcgggggtggacctgcgcgga
ggacattgtcaggtggggagtttggctggggcggcacatctgttaaaagataacgcaggt
gtcctaagatgagctcagtgagaacagaaatctcacgtagaacaaaagggtaaaagctca

tttgattttgattttcagtacgaatacaaactgtgaaagcatggcctatcgatcctttttg

accttcaaggtttgaagcaagaggtgtcagaaaagttaacacgcgttgtggtagcttttta

agagtggactcaaagtgttccgctagtcccccctcggaccgccacttacaaaaggcctgg

ggatgggggcgacactgtcaaattgcggggactccctaaagcctgtctctaccaagccag

cgcggcgaaagcccctggtggccggggtaatgacctagggtatggtaaaagcgagacagg

atg-gaacaatgggcaatccgcagccaagtcctaagg----ctttttgaaaagcggctat

ggatgcagttcaacgactagatggcagtgggctgagagagcccaccgcggcaccctttcg

gcttaagatatagtcttgcccacgtgaaagcgtgcctgcagatgaacgccaccatgtatg

gccggagctgcaggagaccgc--acttgatgtgcagtcgggaat-cgacagtaccacagg

gatgaaa-----------------------------------------------------

------------------------------------------------------------

------------------------------------------------------------

------------------------------------------------------------

------------------------------------------------------------

------------------------------------------------aaaaaacac---

------------------------------------------------------------

------------------------------------------------------------

--acaatta---------------------------------------------------

----------------------------------------atttgtggcc---atcatac

----------------------------------------agtcgta-------------

------------------------------------------------------------

------------------------------------------------------------

------------------------------------------------------------

------------tgatattatcgtcccaggcgggtgcgcctctctcacccgtctagtccg

gccttggtgcgcaaaaaatgcttgcgtggatgcgccttggttaaggaaaatgctcctagt

agttttacaaggacacttcttgtagggcaacacatctggatgcgaagaactcccggaaaa

gacaacggtaccaagtgactggggtaacgccccattcatgaaggcaaataatcaaaaaga

ggtgccgggtaagaacccgttgtatagggacaatccgcagctgacttggctccctg--cc

cat--ccatgggcatagcttagcggttcaacgctcggtaagatgtgggttggcgcgctgt

ccgacagcgcactggcttaaggtacgggctactcccaccggagacggtgtctttgtctac

ctcgggtaacacgcctcgggagaagcaaagagggtgcaatggaaaaaaaactggttgcgg

ggaaatccctaccagagtcacctggtacaaaggaactggcttgtggcagccaagcgttca

tagcgacgttgctttttgatccttcgatgtcggctcttcctatcattgcgaagcagaatt

cgccaagtgttggattgttcacccactaatagggaacgtgagctgggtagccacttta--

-----tgcccaacagctggatagctgggcttctgtctta-ggccgggacttctcacgacc

cgtgccttggtcaccc-aaaaaagggcgaccgcagagctcaggatggagcttctggctta

actagtagaggacagatagagatcgtcaaattgcgggaaaccccttagagcctcgagcta
ccaagtgcgtgtcgaaaggcatagacgcagccagcgatggctgtcgcatggcttga-ttc
catcaagataggtaaaaatgctcgaggattgggcaatccgcagccaagcccctaaggt--
---ccccccgcgggctatggggaaggttcaacgactgtaaggcgtaccgtgcaagcggaa
gatacagtctagccccgctggaaagcagcgccaaagtaagtaggt-tgcagtgggtctgt
cctc
>Form_E_China_SRR9587922
gcatgttttggtatgtgtttcgacagtcatgcatgtcatagccgtaggatggggagtgag
cttcatagcccgaccccttctcaggagcatgttttcatagcatattgagcatactggggt
ggtctcccatgtagctcgctctctcggtgcaggacctcccttttgcgggccacttcgatg
gcctccattatgga-gagcatgttttggcatatgaagcgttggtggtgaggctcgtagcc
tgtatatacataccaacgcagtcgtcatatgtctctctggcgggggtcggtcgtatgacc
ttgtccctacctggagcatgtattttgtatgat---atactggac-ggcagctgcgctgt
tcagtcgtcggccgtacctcatgtgcgcgtttccaggtccgaacgtttgacatgcaaggg
catcataccagaagcctgttttcttgtatgagcatgtaggtaggcaggctcttcagtcgt
ctacccataaatgcacttcgaagga-actttttggtattgaatggctttgtgcccagacg
ttcttgtgagcttgttctcgctctgcgatacgctggaagctgtggtattacctacagcag
acgcattcctagggcggacggcctctctgcagatgggcacatctgtagaaagaggctccc
cttgatgacgcaatgcaatggctgggtaaacggagtgctatacagaagggtcaaacctc-
tgtggaagtacactcataccagcattgtttgttgagaggggtcggctctacgagtgaggc
cctatactgccatctcattcagatgcgtagacggtctttactcagactgcctttctggct
gattgtcctcggtccctcgtgcgcacgcgccgctggctgaatgggcttgtccttatggag
tgccgcgttcttc-aaagaatccgtcttcatacagcgacctcatttacatattgccggct
gaatgggcttgtccttatggagcactgcattcttctaggaattgctatctatacagcgac
ctcatttacatacaaggcacct-ttttctacatagacttgccatttggctgattgttctg
acagtcgctgcctcttcggagttctgtgtggacgagtggcc--tcctgagtgggccggtc
cttacggagtgcccaagtctgttatgactgggttctccatacagcgacctcactcagaat
tgtatgg--------------caccgtgccgat-----------tgaatggacaagcagt
ccttagggaatcgcatgaaagcatgacggcaacatggtgtatggacccgttgggccttgc
ctgtgctatcttctcgagtcgaccgg------------tcctcgtctcctcctttcgcga
cgggaagctcttgagtttaccgtgggtcaggacgctgtggtatctgtttgtctgtagtgt
gggctctctatcagaacagcgtatcagcacacgtcgtctgggtctatggtatttttgctt
aacggccctgggtttggtagtatggttccatgtcggtcctctcacctcccaaccgtcatg
tctctga-----------------------------------------------------
------------------------------------------------------------
-------------------------------------------tatcatctgcacagta

cgtcttgtcagaatatcattctgtatggcagtcgatagactaggccctatgccgcagctg
tctctgaccactatgcgagcgcatacctacctcaatgtgttcatagctgttatggatctt
gggtgtggtgtgccagaacgatacgccaccagattgtggaatggtgcttgtagtattaga
agaggacatgtatgaagagcacgttcttttctcttatgatggtgccgcgggtgcgggccg
ttctcattctgcttgtcgggtgctcaaggtgcatgtaggattgctgaacccac-tgaatg
gtcatgagaatggtcatatggtgaggttcacttctccatcgctttgtgtgtgttggatgt
ttgggtggcttcggccgcatggaaaatctcggtctgtaactcatcagggagggtttccat
gtgagcaatctctccagtcgaacacacacccatgggcgtcccgctaggcaaaatggaact
ctcgatccgacacagcctgtggagtgtttgttccgtgtggcgtgaaacggcttatactgg
atgtcttccc-tttggacaccagtctgttggtatg--tacgcataccatctagagggagc
aagaaaaaggcatgcaagtgccacaaaggaagggaggccttcctgctaccgtgtgtcc-t
cttgacgatcacgggggagtatatagagacatttggacgagatccgtttttcacgaggcg
gtgccctcgcgcaaaggtgttgctctgcacgcatctttatgatgatagttgctcgctcct
atgcgtccactcgtgaaaatacctgctctctctgtctgtttccatgacacccc-catgcg
ttgtcatagagcatgctcaggtacagaaagtgctgtgactttcttggcctgctgtcactg
agtcagcctgatactttggaggaatcttttctgggcactgcccagctgagagttcttatc
aggctctggagtgtgcatatggctataatcataatagtgcatgtggccacgcagaaaaag
catcgacgtggaatggagaagaacaccattctgaatggaacggttcttactctgcttccg
ttgcttgacgctgtatgggaatcgctttgctctgggggtacaccttccattccatgacaa
agcatttgatctatgccccga-gggtcattctctatgaagagcccgaaacttgtctgagt
gagaggcgaaggtagtccttcattt--atgcggggtccgccctcacactgtagtcggcac
agtcccttgtttatggactgcgcagctcaatcttcgactggagagg--aaaaacaaaaca
caagtatgacacgcacatacgtgttttctaggactgtgacctgtcggctgcctcgcggtg
gttggcatgatgccccaagt-gggggggggagaaacaatcagtatgattgataaaaacca
gtgggg-aaaaaggcgcgctgtcgcacacccttggtcacggcctc---aagtcgacggcc
ctcactggccca-ccctgcgactcctgttcgcttttgacaagacacacctgcaata-cgg
aggaaatgcctgagcagctgtgccgcagcaactgcggacgcgtagcgatatgcatccctg
ttgtttgcgacttggtacaaagaacgtgcgccgggagacggattgccctttgacttttcc
acccagcccgcggctctatacccttgacgtacctatttacctggttgatcctgccagtag
tcatatgcttgtctcaaagactaagccatgcatgtctaagtataaacacttttatactcg
tgaaactgcgaatggctcattaaaacagttatagttcctttgggaacaacactacttgga
taaccgtagtaattctagagctaatacatgcctcaacgcccgactcacgaagggtggtat
ttattggataaaaaaccatcgtgctctgtccagagcgctttgagatgattcacaataact
tgtcggatcgcatggcctcgtgctggcgactcctcattcaaatttctgccctatcaactt
tcgatggtagagtattggtctaccatggtgtcgacgggtgacggggaattagggttcgat
tccggagagggagcctgagaaacggctaccacatccaaggaaggcagcaggcgcgcaaat

tacccaatcccgactcggggaggtagtgacaaaaaataacaatagggggccctttgggtc
ttctaattggaatgagaacaatttaaatcccttatcgaggatccattggagggcaagtct
ggt---------------------------------------------------------
------------------------------------------------------------
------------------------------------------------------------
------------------------------------------------------------
------------------------------------------------------------
------------------------------------------------------------
------------------------------------------------------------
------------------------------------------------------------
------------------------------------------------------------
------------------------------------------------------------
------------------------------------------------------------
------------------------------------------------------------
------------------------------------------------------------
------------------------------------------------------------
------------------------------------------------------------
------------------------------------------------------------
-----------------------------------------------gccagcagccgcg
gtaattccagctccaatagcgtatattaaagttgttgcagttaaaacgctcgtagtcgga
cctcggagcgcccccacgggccgggcttttgccttgttgccagtgggagtttgtctcctt
ttgtcgggacgcggtgtgcagggcctttactgtctctgcgcgtttacctgcgggccgacc
gtttactgtgaagaaagtaaagtgttcaaagcaggccaatgccttgaatatgtgagcatg
gaataatagaataggacttgggctctattttgttggttttcagtgaccaagtaatgatta
atagggacggttgggggcattcgtatttcattgtcagaggtgaaattcttggattgatgg
aagacgtacaactgcgaaagcatctgccatggatgttttcattgatcaagaacgaaagtt
aggggatcgaagacgatcagataccgtcgtagtcttaaccataaacgatgccgactgggg
attggcgggggcacttttatatgactctgtcagcaccctgagggaaaccaaagtctttgg
gttctgggggggagtatggtcgcaaggctgaaacttaaaggaattgacggaagggcaccac
caggagtggagcctgcggcttaatttgactcaacacgggaaaacttaccaggtccggaca
gaagaatgattgacagactgaaagctctttcttgatttttttggttggtggtgcatggccg
ttcttagttggtggagtgatttgtctggttaattccgttaacgaacgagacctcgtcctg
ctaaataggtgcgcgcatgccagcactgcgttcttaccttcttagagggactatgcgcgt
ctagcgtatggaagattgaggcaataacaggtctgtgatgcccttagatgttctgggccg
cacgcgcgctacactgatgcattcaacgagtttcacttgatagtcctgggtcggaaggct
cgggtaatcttttgaaagtgcatcgtgctggggatagatcattgcaattattgatcttca

acgaggaattccttgtaggcgcaggtcatcagcctgcgccgaatacgtccctgccctttg
tacacaccgcccgtcgctcctaccgattgaatggtccgatgaaatgtccggatcgcggct
agtgtgatggactcgtccagagcggtagccgtgagaagctcattaaatcttatcatttag
aggaaggagaagtcgtaacaaggt------------------------------------
------------------------------------------------------------
------------------------------------------------------------
------------------------------------------------------------
------------------------------------------------------------
------------------------------------------------------------
------------------------------------------------------------
------------------------------------------------------------
------------------------------------------------------------
------------------------------------------------------------
------------------------------------------------------------
------------------------------------------------------------
------------------------------------------------------------
------------------------------------------------------------
------------------------------------------------------------
------------------------------------------------------------
-------------------------------ttccgtaggtgaacctgcggaaggatcat
tcacaaactattgacacaaacacacgcgaaccaaatcgtccgcacagcaggtggcaatga
aagagagactctgcatgtcgcctttcgggg--tagcaagcagcactcttctgccatcgcc
tctgtgcagggcgtaaattcccattgagaggatgtgagggcaccacaggaagcttttcca
caggaagtcgccatccttctccctccacggcggcg----cttctgtgcttggcag-----
----tttttttttgccttctagggaggatgccgccaatggagccccttat-atatataca
tcatcatatgccc--ctttttttcttaaccgcttgccacagcttcttctatgaggagctt
gtgggaagactgtctccatacaataac-aaagatacaactcttagcggtggatatcttgg
ttctcgcaacgatgaagaacgcagccaactgcgataactaatgtgaattgcaggacttcg
tgaatcattaagtctttgaacgcaagttgcgctcatctccatgaggaggtgagcatgcct
gtttgagtgtcgatgggaaaaacaatac-ataacaacaatcggctctcttatcttcctgc
cttcttgggggcaaaaggtagtgcatgccggagatgggctttgtcgcccctttctcttga
aagaggcgtcgtccgaaacacgtatcgttgcccgctggaggtcatagagcatgagttt--
aaagcaccacatctctcaagtcagagtgag-cgtcttgagtaggtcc-gcattgattcag
tttcaacccaggtggcgttggttacttgccat-aagcctctctggggtttctttgcttgg
acctctcttgtcagtttctctgacgtagagacaggt--gccgtctcatggacgttgccct
ctggaacgtgcta-aaaaaaaaaaaaaga----------aaaaaccgtaccacatgtcaa

aagagacagatcgtgcttgacggcgggaggtgttgcatttcttgcttgtcctgagcgtct
gccagtctct----ga--cgagtg----tgttccttccataaacccgatctcaaatcagg
taggaccacccgctgaatttaagcatataattaagcggaggaaaagaaactaacaaggat
tccctcagtagcggcgagcgaagcgggaagagctcaacatgaaaaccgttgggggttcgc
gctttccttcgggttgtatagtaaagtggtgtcgcctgcgtcgcgccgggtataaatccc
gtctgaaatggggtgtcgtagagggttatatccccgtctgtgacccggtgtgcgatgcgt
tacggtgaccttacgatgagtcgggttgcttgggcatgcagccctaagtgggtggtaaac
ttcatctaaagctaaatactagcgagagaccgatagcgtacaagtaccgtaagggaaaga
tgcaaagaactttgaaaagagagttaaacagggcttgaaaccgttaagagtgaagcgaag
gaggtcagtgatcatggggccagatccccaagtgcctgtcata-gggcaccgtagcgttc
tggtttcccggggcagcgccggccgggggcgggggttaacagggaggtggttgagggatt
cgttttccgcctgtactcctttatagcttgcaccccggccgagggatccgcgacttgtcg
tgtgagggcgctgtcaaaatggccttctccaccccgtcttgaaacacggaccaaggagtc
taacatactcgcgagtgtttgggtggttaaaacccgaacgcacaatgaaagtaaacggtg
gctctgcgccgctgcttggtgggaaccctgcttttataatgggggcactatcacccgac
cccgatccctttggggtggttggttttgaggtagagcgagtatgatgggacccgaaagat
ggtgaactattcctgagtagggcgaagccagaggaaactctggtggaggctcgtagcgat
gctgacgtgcaaatcgctcgtcgaacttgggtataggggcgaaagactaatcgaaccatc
tagtagctggttccctccgaagtttccctcaggatagcgggagcaaatgcaagtcagtat
catccggtaaagcgaatgattagaggcctcgggggagtaactccttcgacctattctcaa
actttaaataggtgaggagccggcaacgttgcttaaaacaatgaacgccgtgcggacaat
gattgctctcagtgggccgattttggtaagcagaactggcgatgcgggatgaaccgaaag
cgacgctaaggtgccaaactcccagctcattcagacaccagaaaaggtgttgattcatac
agacagcaggacggtggccatggaagtcggaatccgctaaggagtgtgtaacaactcacc
tgccgaatggatcagccctgaaaatggatggcgcttaagcgggggaccgaggcgtcgccg
ttggggtaaatacgaagccccaatgagtaggagggcgtgggggttttgaaacacggtaca
ctcgcatggaagccttgggcgtaagctcgggtggagcgacccctagtgtagatcttggtg
gtagtagcaaatattcaagtgagaaccttgaagactgaagtggggaagggttccacgcga
acagtgcttggacgtgggttagtcggtcctaagagatggggaaactccgtattcaagtgc
gcttttgtagcgcagtccatcgaaagggaagcgggttaacattcccgcaccaggatgtgg
aaactgtggtttaccgcaaactgcggcaacgcaactgaacctggggacgcccgtaggagc
tccgagcagggatctctttcctgtttaacgcgcccaacccccctggaatcggtttgtccgg
agatagggggcgaaaggccggaaaagcaccgcacgtcttgtggtgttcggaacgcttgta
atggcccttgaaaacccaggggaaagcagccgtcttacac--ttgtgtgagacgaggaat
tctcacacctggccgtacccatcaccgcatcaggtccccaaggtgaacagcctctagtca
atagaataatgtagataagggaagtcggcaaaatagatccgtaactttgggaaaaggatt

ggctctaatgttcgggcttgtcgggacgcgcgggcgtagatgccaatgagcggcagacgt
ctagcctctttggaggctggattaacgctgcttgtgggaagaaaccgtgtcgatggaggc
aggaacgcttatcgatcttgtctcttcccgcttgcgatcaacgaacggattagaactggt
acggacaaggggaatccgactgtttaattaaaacaaagcattgcgacggccggaaacggt
gttggcgcaatgtgatttctgcccagtgctctgaatgtcaaagtgaagaaattcaaccaa
gcgcgggtaaacggcgggagtaactatgactctcttagac-agatcaagacaccttcaat
aaattggttcattatgtcgggtggtccgtcgtccgggaagcgaccaaagcaaaaccccgt
agactgtgatgtgcatctgtcagacgaaactgacgtccgtgttcatcaagccttagctat
ctttcaacgccggcttcgagacgtggcgtctcttaaggtgtttgaagtggctctctctgt
gggaactactgatgcactggtagagatgggcactatcacttgctggagtattgatgctgt
tgggaaaggtttctcaaggaacagcagcggtcgctggcaatttgacctcactggcaagat
taatggcaagcctctaatatccaccacaaaggatcgttggaggccggttgtgacggctgt
gccctactacctttcgtgcgccatcgatggggacaagacgacgctggaggaaaagcagtg
gacttcatcacacatgtgtcaccatgaatggtgtgtgaacgtggagcacattgtttatga
aagcctcgacgtcaac--aagtctcggaactggtg--ccagggaaaaaacgcatgcctcc
atctgcctgtttgcattcgaaccggccgccgtgtgcaggagctcaggaagagcaatgggg
ttcgtgcaatggacctcctcctgcagaaattggctggattggagattgaggttgaggagc
ccgctgcagctgtactccgagagtctgtcgaacgggaggacgcctagtgagtgactcgtg
acacgggttggctgcgcatgatgcgtcaactacatggacatcttttgagagagataggcg
ggctgaaaaggttatagccgtctagtctcgctctctttcattgagcggggcgacacacct
ggttgcgggaaccctgcccacctttctatcaagaagtgggaaccagtccattcctactcc
tgcttgtgaggcatatagt--aacgtagcagcaggaggatctcggcgagaggccgaaccc
gaaaggaataacgtgggctgggaggcgatccgcaggcgagtccgtaactatatgattatc
gacgggccgtctcaacgactggtaaggtgtgggtttg-tcctcctcctcacgtggggaca
ggcttaaggtacagtctaatcccacccgccaagggtgcttggg---tg------------
--------------------taatccctcttccatagagggcactagccca-ggggctgg
gacgctgggaggaa----------------atgccccgtccctatagccggtagaaatga
aggtagccaaatgcctcgtcatctaaactaattgg-at----------------------
----------------------gagatagtc----------tgagtgaaaagcttatgc
ag-----------------gctagtaccgg---aaaaaaaaaaaaaagaggggctctgcca
ct----ctgac------------------------ggtgcgacactccttgtacagggaa
accctcc--gcatggaaaggcacgtggtaccaccctcgcggggaaactcgcgagcgggat
cccgggtaatgaccggagccgatggtaaaaactcacgtgaccgatggggtgaccctgtgg
cgagacgctaagttgtgggacatcaccctctcacaatacgcatccgtcgcagagactgaa
cgggagtgggctccgttga-----cggccggccggccagccagccgtcttcca-------
---ggggcttaagatacagtccgggcctcagggatgg----aaaaacctggggaacta-c

cagaattagtgacgcgcatgaatggattaacgagattcccactgtccctatctactatct
agcgaaaccacagccaagggaacgggcttggaagaatcagcggggaaagaagaccctgtt
gagcttgactctagtccgactttgtgagatgagttgggaggtgtagtctaagagggaggg
gcaactcgaccgtgaaatacctctactcccaagcttgttttacttactccggcgagcagg
gatgggtggtaacacccttttctaggattaacacctcttcgggggtggacctgcgcgga
ggacattgtcaggtggggagtttggctggggcggcacatctgttaaaagataacgcaggt
gtcctaagatgagctcagtgagaacagaaatctcacgtagaacaaaagggtaaaagctca
tttgattttgattttcagtacgaatacaaactgtgaaagcatggcctatcgatcctttg
accttcaaggtttgaagcaagaggtgtcagaaaagttaacacgcgttgtggtagcttta
agagtggactcaaagtgttccgctagtccccctcggaccgccacttacaaaaggcctgg
ggatgggggcgacactgtcaaattgcggggactccctaaagcctgtctctaccaagccag
cgcggcgaaagcccctggtggccggggtaatgacctagggtatggtaaaagcgagacagg
atg-gaacaatgggcaatccgcagccaagtcctaagg----ctttttgaaaagcggctat
ggatgcagttcaacgactagatggcagtgggctgagagagcccaccgcggcaccctttcg
gcttaagatatagtcttgcccacgtgaaagcgtgcctgcagatgaacgccaccatgtatg
gccggagctgcaggagaccgc--acttgatgtgcagtcgggaat-cgacagtaccacagg
gatgaac-----------------------------------------------------
------------------------------------------------------------
------------------------------------------------------------
------------------------------------------------------------
------------------------------------------------------------
-------------------------------------------------aaaaacac---
------------------------------------------------------------
------------------------------------------------------------
--acaatta---------------------------------------------------
----------------------------------------atttgtggcc---atcatac
----------------------------------------agtcgta-------------
------------------------------------------------------------
------------------------------------------------------------
------------------------------------------------------------
------------tgatattatcgtcccaggcgggtgcgcctctctcacccgtctagtccg
gccttggtgcgcaaaaaaatgcttgcgtggatgcgccttggttaaggaaaatgctcctagt
agttttacaaggacacttcttgtagggcaacacatctggatgcgaagaactcccggaaaa
gacaacggtaccaagtgactggggtaacgccccattcatgaaggcaaataatcaaaaaga
ggtgccgggtaagaacccgttgtatagggacaatccgcagctgacttggctccctg--cc
catccccatgggcatagcttagcggttcaacgctcggtaagatgtgggttggcgcgctgt

ccgacagcgcactggcttaaggtacgggctactcccaccggagacggtgtctttgtctac
ctcgggtaacacgcctcgggagaagcaaagagggtgcaatgg-aaaaaaactggttgcgg
ggaaatccctaccagagtcacctggtacaaaggaactggcttgtggcagccaagcgttca
tagcgacgttgctttttgatccttcgatgtcggctcttcctatcattgcgaagcagaatt
cgccaagtgttggattgttcacccactaatagggaacgtgagctgggtagccactttta--
-----tgcccaacagctggatagctgggcttctgtctta-ggccgggacttctcacgacc
cgtgccttggtcaccc-aaaaaagggcgaccgcagagctcaggatggagcttctggctta
actagtagaggacagatagagatcgtcaaattgcgggaaaccccttagagcctcgagcta
ccaagtgcgtgtcgaaaggcatagacgcagccagcgatggctgtcgcatggcttga-ttc
catcaagataggtaaaaatgctcgaggattgggcaatccgcagccaagcccctaaggt--
---ccccccgcgggctatggggaaggttcaacgactgtaaggcgtaccgtgcaagcggaa
gatacagtctagccccgctggaaagcagcgccaaagtaagtaggt-tgcagtgggtctgt
cctc
>Form_E_China_SRR9587923
gcatgttttggtatgtgtttcgacagtcatgcatgtcatagccgtaggatggggagtgag
cttcatagcccgaccccttctcaggagcatgttttcatagcatattgagcatactggggt
ggtctcccatgtagctcgctctctcggtgcaggacctcccttttgcgggccacttcgatg
gcctccattatgga-gagcatgttttggcatatgaagcgttggtggtgaggctcgtagcc
tgtatatacataccaacgcagtcgtcatatgtctctctggcgggggtcggtcgtatgacc
ttgtccctacctggagcatgtattttgtatgat---atactggac-ggcagctgcgctgt
tcagtcgtcggccgtacctcatgtgcgcgtttccaggtccgaacgtttgacatgcaaggg
catcataccagaagcctgttttctcgtatgagcatgtaggtaggcaggctcttcagtcgt
ctacccataaatgcacttcgaagga-actttttggtattgaatggctttgtgcccagacg
ttcttgtgagcttgttctcgctctgcgatacgctggaagctgtggtattacctacagcag
acgcattcctagggcggacggcctctctgcagatgggcacatctgtagaaagaggctccc
cttgatgacgcaatgcaatggctgggtaaacggagtgctatacagaagggtcaaacctc-
tgtggaagtacactcataccagcattgtttgttgagaggggtcggctctacgagtgaggc
cctatactgccatctcattcagatgcgtagacggtctttactcagactgcctttctggct
gattgtcctcggtccctcgtgcgcacgcgccgctggctgaatgggcttgtccttatggag
tgccgcgttcttc-aaagaatccgtcttcatacagcgacctcatttacatattgccggct
gaatgggcttgtccttatggagcactgcattcttctaggaattgctatctatacagcgac
ctcatttacatacaaggcacct-ttttctacatagacttgccatttggctgattgttctg
acagtcgctgcctcttcggagttctgtgtggacgagtggcc--tcctgagtgggccggtc
cttacggagtgcccaagtctgttatgactgggttctccatacagcgacctcactcagaat
tgtatgg--------------caccgtgccgat-----------tgaatggacaagcagt
ccttagggaatcgcatgaaagcatgacggcaacatggtgtatggacccgttgggccttgc

ctgtgctatcttctcgagtcgaccgg------------tcctcgtctcctcctttcgcga
cgggaagctcttgagtttaccgtgggtcaggacgctgtggtatctgtttgtctgtagtgt
gggctctctatcagaacagcgtatcagcacacgtcgtctgggtctatggtatttttgctt
aacggccctgggtttggtagtatggttccatgtcggtcctctcacctcccaaccgtcatg
tctctga-----------------------------------------------------
------------------------------------------------------------
--------------------------------------------tatcatctgcacagta
cgtcttgtcagaatatcattctgtatggcagtcgatagactaggccctatgccgcagctg
tctctgaccactatgcgagcgcatacctacctcaatgtgttcatagctgttatggatctt
gggtgtggtgtgccagaacgatacgccaccagattgtggaatggtgcttgtagtattaga
agaggacatgtatgaagagcacgttcttttctcttatgatggtgccgcgggtgcgggccg
ttctcattctgcttgtcgggtgctcaaggtgcatgtaggattgctgaacccac-tgaatg
gtcatgagaatggtcatatggtgaggttcacttctccatcgctttgtgtgtgttggatgt
ttgggtggcttcggccgcatggaaaatctcggtctgtaactcatcagggagggtttccat
gtgagcaatctctccagtcgaacacacacccatgggcgtcccgctaggcaaaatggaact
ctcgatccgacacagcctgtggagtgtttgttccgtgtggcgtgaaacggcttatactgg
atgtcttccc-tttggacaccagtctgttggtatg--tacgcataccatctagagggagc
aagaaaaaggcatgcaagtgccacaaaggaagggaggccttcctgctaccgtgtgtcc-t
cttgacgatcacgggggagtatatagagacatttggacgagatccgtttttcacgaggcg
gtgccctcgcgcaaaggtgttgctctgcacgcatctttatgatgatagttgctcgctcct
atgcgtccactcgtgaaaatacctgctctctctgtctgtttccatgacacccc-catgcg
ttgtcatagagcatgctcaggtacagaaagtgctgtgactttcttggcctgctgtcactg
agtcagcctgatactttggaggaatcttttctgggcactgcccagctgagagttcttatc
aggctctggagtgtgcatatgcctatattcattatagtgcatgtggctacgcagaaaaag
catcgacgtggaatggagaagaacaccattctgaatggaacggttcttactctacttctg
ttgcttgacgctgtatgggaatcgctttgctctgggggtacaccttccattccatgacaa
agcatttgatctatgccccga-gggtcattctctatgaagagcccgaaacttgtctgagt
gagaggcgaaggtagtccttcattt--atgcggggtccgccctcacactgtagtcggcac
agtcccttgtttatggactgcgcagctcaatcttcgactggagagg--aaaaacaaaaca
caagtatgacacgcacatacgtgttttctaggactgtgacctgtcggctgcctcgcggtg
gttggcatgatgccccaagtgggggggggggagaaacaatcagtatgattgataaaaacca
gtgggg-aaaaaggcgcgctgtcgcacacccttggtcacggcctc---aagtcgacggcc
ctcactggccca-ccctgcgactcctgttcgcttttgacaagacacacctgcaata-cgg
aggaaatgcctgagcagctgtgccgcagcaactgcggacgcgtagcgatatgcatccctg
ttgtttgcgacttggtacaaagaacgtgcgccgggagacggattgccctttgacttttcc
acccagcccgcggctctatacccttgacgtacctatttacctggttgatcctgccagtag

tcatatgcttgtctcaaagactaagccatgcatgtctaagtataaacacttttatactcg
tgaaactgcgaatggctcattaaaacagttatagttcctttgggaacaacactacttgga
taaccgtagtaattctagagctaatacatgcctcaacgcccgactcacgaagggtggtat
ttattggataaaaaaccatcgtgctctgtccagagcgctttgagatgattcacaataact
tgtcggatcgcatggcctcgtgctggcgactcctcattcaaatttctgccctatcaactt
tcgatggtagagtattggtctaccatggtgtcgacgggtgacggggaattagggttcgat
tccggagagggagcctgagaaacggctaccacatccaaggaaggcagcaggcgcgcaaat
tacccaatcccgactcggggaggtagtgacaaaaaataacaatagggggccctttgggtc
ttctaattggaatgagaacaatttaaatcccttatcgaggatccattggagggcaagtct
ggt---------------------------------------------------------
------------------------------------------------------------
------------------------------------------------------------
------------------------------------------------------------
------------------------------------------------------------
------------------------------------------------------------
------------------------------------------------------------
------------------------------------------------------------
------------------------------------------------------------
------------------------------------------------------------
------------------------------------------------------------
------------------------------------------------------------
------------------------------------------------------------
------------------------------------------------------------
------------------------------------------------------------
------------------------------------------------------------
-----------------------------------------------gccagcagccgcg
gtaattccagctccaatagcgtatattaaagttgttgcagttaaaacgctcgtagtcgga
cctcggagcgcccccacgggccgggcttttgccttgttgccagtgggagtttgtctcctt
ttgtcgggacgcggtgtgcagggcctttactgtctctgcgcgtttacctgcgggccgacc
gtttactgtgaagaaagtaaagtgttcaaagcaggccaatgccttgaatatgtgagcatg
gaataatagaataggacttgggctctattttgttggttttcagtgaccaagtaatgatta
atagggacggttgggggcattcgtatttcattgtcagaggtgaaattcttggattgatgg
aagacgtacaactgcgaaagcatctgccatggatgttttcattgatcaagaacgaaagtt
aggggatcgaagacgatcagataccgtcgtagtcttaaccataaacgatgccgactgggg
attggcgggggcacttttatatgactctgtcagcaccctgagggaaaccaaagtctttgg
gttctgggggagtatggtcgcaaggctgaaacttaaaggaattgacggaagggcaccac

caggagtggagcctgcggcttaatttgactcaacacgggaaaacttaccaggtccggaca
gaagaatgattgacagactgaaagctctttcttgatttttggttggtggtgcatggccg
ttcttagttggtggagtgatttgtctggttaattccgttaacgaacgagacctcgtcctg
ctaaataggtgcgcgcatgccagcactgcgttcttaccttcttagagggactatgcgcgt
ctagcgtatggaagattgaggcaataacaggtctgtgatgcccttagatgttctgggccg
cacgcgcgctacactgatgcattcaacgagtttcacttgatagtcctgggtcggaaggct
cgggtaatcttttgaaagtgcatcgtgctggggatagatcattgcaattattgatcttca
acgaggaattccttgtaggcgcaggtcatcagcctgcgccgaatacgtccctgccctttg
tacacaccgcccgtcgctcctaccgattgaatggtccgatgaaatgtccggatcgcggct
agtgtgatggactcgtccagagcggtagccgtgagaagctcattaaatcttatcatttag
aggaaggagaagtcgtaacaaggt------------------------------------
------------------------------------------------------------
------------------------------------------------------------
------------------------------------------------------------
------------------------------------------------------------
------------------------------------------------------------
------------------------------------------------------------
------------------------------------------------------------
------------------------------------------------------------
------------------------------------------------------------
------------------------------------------------------------
------------------------------------------------------------
------------------------------------------------------------
------------------------------------------------------------
------------------------------------------------------------
------------------------------------------------------------
-------------------------------ttccgtaggtgaacctgcggaaggatcat
tcacaaactattgacacaaacacacgcgaaccaaatcgtccgcacagcaggtggcaatga
aagagagactctgcatgtcgcctttcgggg--tagcaagcagcactcttctgccatcgcc
tctgtgcagggcgtaaattcccattgagaggatgtgagggcaccacaggaagcttttcca
caggaagtcgccatccttctccctccacggcggcg----cttctgtgcttggcag-----
----ttttttttttgccttctagggaggatgccgccaatggagccccttat-atatataca
tcatcatatgccc--ctttttttcttaaccgcttgccacagcttcttctatgaggagctt
gtgggaagactgtctccatacaataac-aaagatacaactcttagcggtggatatcttgg
ttctcgcaacgatgaagaacgcagccaactgcgataactaatgtgaattgcaggacttcg
tgaatcattaagtctttgaacgcaagttgcgctcatctccatgaggaggtgagcatgcct

gtttgagtgtcgatgggaaaaacaatac-ataacaacaatcggctctcttatcttcctgc
cttcttgggggcaaaaggtagtgcatgccggagatgggctttgtcgcccctttctcttga
aagaggcgtcgtccgaaacacgtatcgttgcccgctggaggtcatagagcatgagttt--
aaagcaccacatctctcaagtcagagtgag-cgtcttgagtaggtcc-gcattgattcag
tttcaacccaggtggcgttggttacttgccat-aagcctctctggggtttctttgcttgg
acctctcttgtcagtttctctgacgtagagacaggt--gccgtctcatggacgttgccct
ctggaacgtgcta-aaaaaaaaaaaaaga----------aaaaaccgtaccacatgtcaa
aagagacagatcgtgcttgacggcgggaggtgttgcatttcttgcttgtcctgagcgtct
gccagtctct----ga--cgagtg----tgttccttccataaacccgatctcaaatcagg
taggaccacccgctgaatttaagcatataattaagcggaggaaaagaaactaacaaggat
tccctcagtagcggcgagcgaagcgggaagagctcaacatgaaaaccgttgggggttcgc
gctttccttcgggttgtatagtaaagtggtgtcgcctgcgtcgcgccgggtataaatccc
gtctgaaatggggtgtcgtagagggttatatccccgtctgtgacccggtgtgcgatgcgt
tacggtgaccttacgatgagtcgggttgcttgggcatgcagccctaagtgggtggtaaac
ttcatctaaagctaaatactagcgagagaccgatagcgtacaagtaccgtaagggaaaga
tgcaaagaactttgaaaagagagttaaacagggcttgaaaccgttaagagtgaagcgaag
gaggtcagtgatcatggggccagatccccaagtgcctgtcata-gggcaccgtagcgttc
tggtttcccggggcagcgccggccgggggcgggggttaacagggaggtggttgagggatt
cgttttccgcctgtactcctttatagcttgcaccccggccgagggatccgcgacttgtcg
tgtgagggcgctgtcaaaatggccttctccaccccgtcttgaaacacggaccaaggagtc
taacatactcgcgagtgtttgggtggttaaaacccgaacgcacaatgaaagtaaacggtg
gctctgcgccgctgcttggtgggaaccctgcttttataatgggggcactatcacccgac
cccgatccctttggggtggttggttttgaggtagagcgagtatgatgggacccgaaagat
ggtgaactattcctgagtagggcgaagccagaggaaactctggtggaggctcgtagcgat
gctgacgtgcaaatcgctcgtcgaacttgggtataggggcgaaagactaatcgaaccatc
tagtagctggttccctccgaagtttccctcaggatagcgggagcaaatgcaagtcagtat
catccggtaaagcgaatgattagaggcctcgggggagtaactccttcgacctattctcaa
actttaaataggtgaggagccggcaacgttgcttaaaacaatgaacgccgtgcggacaat
gattgctctcagtgggccgattttggtaagcagaactggcgatgcgggatgaaccgaaag
cgacgctaaggtgccaaactcccagctcattcagacaccagaaaaggtgttgattcatac
agacagcaggacggtggccatggaagtcggaatccgctaaggagtgtgtaacaactcacc
tgccgaatggatcagccctgaaaatggatggcgcttaagcgggggaccgaggcgtcgccg
ttggggtaaatacgaagccccaatgagtaggagggcgtgggggttttgaaacacggtaca
ctcgcatggaagccttgggcgtaagctcgggtggagcgacccctagtgtagatcttggtg
gtagtagcaaatattcaagtgagaaccttgaagactgaagtggggaagggttccacgcga
acagtgcttggacgtgggttagtcggtcctaagagatggggaaactccgtattcaagtgc

gcttttgtagcgcagtccatcgaaagggaagcgggttaacattcccgcaccaggatgtgg
aaactgtggtttaccgcaaactgcggcaacgcaactgaacctggggacgcccgtaggagc
tccgagcagggatctctttcctgtttaacgcgcccaaccccctggaatcggtttgtccgg
agatagggggcgaaaggccggaaaagcaccgcacgtcttgtggtgttcggaacgcttgta
atggcccttgaaaacccaggggaaagcagccgtcttacac--ttgtgtgagacgaggaat
tctcacacctggccgtacccatcaccgcatcaggtccccaaggtgaacagcctctagtca
atagaataatgtagataagggaagtcggcaaaatagatccgtaactttgggaaaaggatt
ggctctaatgttcgggcttgtcgggacgcgcgggcgtagatgccaatgagcggcagacgt
ctagcctctttggaggctggattaacgctgcttgtgggaagaaaccgtgtcgatggaggc
aggaacgcttatcgatcttgtctcttcccgcttgcgatcaacgaacggattagaactggt
acggacaaggggaatccgactgtttaattaaaacaaagcattgcgacggccggaaacggt
gttggcgcaatgtgatttctgcccagtgctctgaatgtcaaagtgaagaaattcaaccaa
gcgcgggtaaacggcgggagtaactatgactctctt------------------------
------------------------------------------------------------
------------------------------------------------------------
------------------------------------------------------------
------------------------------------------------------------
------------------------------------------------------------
------------------------------------------------------------
------------------------------------------------------------
------------------------------------------------------------
------------------------------------------------------------
------------------------------------------------------------
------------------------------------------------------------
------------------------------------------------------------
------------------------------------------------------------
------------------------------------------------------------
------------------------------------------------------------
------------------------------------------------------------
------------------------------------------------------------
------------------------------------------------------------
------------------------------------------------------------
------------------------------------------------------------
-----------------------------------------------------------a
aggtagccaaatgcctcgtcatctaaactaattgg-at----------------------
-----------------------gagatagtc----------tgagtgaaaagcttatgc

ag-----------------gctagtaccgg---aaaaaaaaaaaaagaggggctctgcca
ct----ctgac------------------------ggtgcgacactccttgtacagggaa
accctcc--gcatggaaaggcacgtggtaccaccctcgcggggaaactcgcgagcgggat
cccgggtaatgaccggagccgatggtaaaaactcacgtgaccgatggggtgaccctgtgg
cgagacgctaagttgtgggacatcaccctctcacaatacgcatccgtcgcagagactgaa
cgggagtgggctccgttga-----cggccggccggccagccagccgtcttcca-------
---ggggcttaagatacagtccgggcctcagggatgg----aaaaacctggggaacta-c
cagaattagtgacgcgcatgaatggattaacgagattcccactgtccctatctactatct
agcgaaaccacagccaagggaacgggcttggaagaatcagcggggaaagaagaccctgtt
gagcttgactctagtccgactttgtgagatgagttgggaggtgtagtctaagagggaggg
gcaactcgaccgtgaaatacctctactcccaagcttgttttacttactccggcgagcagg
gatgggtggtaacacccttttctaggattaacacctcttcgggggtggacctgcgcgga
ggacattgtcaggtggggagtttggctggggcggcacatctgttaaaagataacgcaggt
gtcctaagatgagctcagtgagaacagaaatctcacgtagaacaaaagggtaaaagctca
tttgattttgattttcagtacgaatacaaactgtgaaagcatggcctatcgatcctttg
accttcaaggtttgaagcaagaggtgtcagaaaagttaacacgcgttgtggtagcttta
agagtggactcaaagtgttccgctagtccccccctcggaccgccacttacaaaaggcctgg
ggatgggggcgacactgtcaaattgcggggactccctaaagcctgtctctaccaagccag
cgcggcgaaagcccctggtggccggggtaatgacctagggtatggtaaaagcgagacagg
atg-gaacaatgggcaatccgcagccaagtcctaagg----ctttttgaaaagcggctat
ggatgcagttcaacgactagatggcagtgggctgagagagcccaccgcggcaccctttcg
gcttaagatatagtcttgcccacgtgaaagcgtgcctgcagatgaacgccaccatgtatg
gccggagctgcaggagaccgc--acttgatgtgcagtcgggaat-cgacagtaccacagg
gatgaac-----------------------------------------------------
------------------------------------------------------------
------------------------------------------------------------
------------------------------------------------------------
------------------------------------------------------------
-------------------------------------------------aaaaacac---
------------------------------------------------------------
------------------------------------------------------------
--acaa------------------------------------------------------
------------------------------------------ttgtggcc---atcatac
----------------------------------------agtcgta-------------
------------------------------------------------------------
------------------------------------------------------------

------------------------------------------------------------
------------tgatattatcgtcccaggcgggtgcgcctctctcacccgtctagtccg
gccttggtgcgcaaaaaatgcttgcgtggatgcgccttggttaaggaaaatgctcctagt
agttttacaaggacacttcttgtagggcaacacatctggatgcgaagaactcccggaaaa
gacaacggtaccaagtgactggggtaacgccccattcatgaaggcaaataatcaaaaaga
ggtgccgggtaagaacccgttgtatagggacaatccgcagctgacttggctccctg--cc
cat--ccatgggcatagcttagcggttcaacgctcggtaagatgtgggttggcgcgctgt
ccgacagcgcactggcttaaggtacgggctactcccaccggagacggtgtctttgtctac
ctcgggtaacacgcctcgggagaagcaaagagggtgcaatgg-aaaaaaactggttgcgg
ggaaatccctaccagagtcacctggtacaaaggaactggcttgtggcagccaagcgttca
tagcgacgttgctttttgatccttcgatgtcggctcttcctatcattgcgaagcagaatt
cgccaagtgttggattgttcacccactaatagggaacgtgagctgggtagccacttta--
-----tgcccaacagctggatagctgggcttctgtctta-ggccgggacttctcacgacc
cgtgccttggtcaccc-aaaaaagggcgaccgcagagctcaggatggagcttctggctta
actagtagaggacagatagagatcgtcaaattgcgggaaaccccttagagcctcgagcta
ccaagtgcgtgtcgaaaggcatagacgcagccagcgatggctgtcgcatggcttga-ttc
catcaagataggtaaaaatgctcgaggattgggcaatccgcagccaagcccctaaggt--
---ccccccgcgggctatggggaaggttcaacgactgtaaggcgtaccgtgcaagcggaa
gatacagtctagccccgctggaaagcagcgccaaagtaagtaggt-tgcagtgggtctgt
cctc
>Form_E_China_SRR9587924_SRR9587965
gcatgttttggtatgtgtttcgacagtcatgcatgtcatagccgtaggatggggagtgag
cttcatagcccgaccccttctcaggagcatgttttcatagcatattgagcatactggggt
ggtctcccatgtagctcgctctctcggtgcaggacctcccttttgcgggccacttcgatg
gcctccattatgga-gagcatgttttggcatatgaagcgttggtggtgaggctcgtagcc
tgtatatacataccaacgcagtcgtcatatgtctctctggcgggggtcggtcgtatgacc
ttgtccctacctggagcatgtattttgtatgat---atactggac-ggcagctgcgctgt
tcagtcgtcggccgtacctcatgtgcgcgtttccaggtccgaacgtttgacatgcaaggg
catcataccagaagcctgttttctcgtatgagcatgtaggtaggcaggctcttcagtcgt
ctacccataaatgcacttcgaagga-actttttggtattgaatggctttgtgcccagacg
ttcttgtgagcttgttctcgctctgcgatacgctggaagctgtggtattacctacagcag
acgcattcctagggcggacggcctctctgcagatgggcacatctgtagaaagaggctccc
cttgatgacgcaatgcaatggctgggtaaacggagtgctatacagaagggtcaaacctc-
tgtggaagtacactcataccagcattgtttgttgagaggggtcggctctacgagtgaggc
cctatactgccatctcattcagatgcgtagacggtctttactcagactgcctttctggct
gattgtcctcggtccctcgtgcgcacgcgccgctggctgaatgggcttgtccttatggag

tgccgcgttcttc-aaagaatccgtcttcatacagcgacctcatttacatattgccggct
gaatgggcttgtccttatggagcactgcattcttctaggaattgctatctatacagcgac
ctcatttacatacaaggcacct-ttttctacatagacttgccatttggctgattgttctg
acagtcgctgcctcttcggagttctgtgtggacgagtggcc--tcctgagtgggccggtc
cttacggagtgcccaagtctgttatgactgggttctccatacagcgacctcactcagaat
tgtatgg--------------caccgtgccgat-----------tgaatggacaagcagt
ccttagggaatcgcatgaaagcatgacggcaacatggtgtatggacccgttgggccttgc
ctgtgctatcttctcgagtcgaccgg------------tcctcgtctcctcctttcgcga
cgggaagctcttgagtttaccgtgggtcaggacgctgtggtatctgtttgtctgtagtgt
gggctctctatcagaacagcgtatcagcacacgtcgtctgggtctatggtatttttgctt
aacggccctgggtttggtagtatggttccatgtcggtcctctcacctcccaaccgtcatg
tctctga-----------------------------------------------------
------------------------------------------------------------
--------------------------------------------tatcatctgcacagta
cgtcttgtcagaatatcattctgtatggcagtcgatagactaggccctatgccgcagctg
tctctgaccactatgcgagcgcatacctacctcaatgtgttcatagctgttatggatctt
gggtgtggtgtgccagaacgatacgccaccagattgtggaatggtgcttgtagtattaga
agaggacatgtatgaagagcacgttcttttctcttatgatggtgccgcgggtgcgggccg
ttctcattctgcttgtcgggtgctcaaggtgcatgtaggattgctgaacccac-tgaatg
gtcatgagaatggtcatatggtgaggttcacttctccatcgctttgtgtgtgttggatgt
ttgggtggcttcggccgcatggaaaatctcggtctgtaactcatcagggagggtttccat
gtgagcaatctctccagtcgaacacacacccatgggcgtcccgctaggcaaaatggaact
ctcgatccgacacagcctgtggagtgtttgttccgtgtggcgtgaaacggcttatactgg
atgtcttccc-tttggacaccagtctgttggtatg--tacgcataccatctagagggagc
aagaaaaaggcatgcaagtgccacaaaggaagggaggccttcctgctaccgtgtgtcc-t
cttgacgatcacgggggagtatatagagacatttggacgagatccgtttttcacgaggcg
gtgccctcgcgcaaaggtgttgctctgcacgcatctttatgatgatagttgctcgctcct
atgcgtccactcgtgaaaatacctgctctctctgtctgtttccatgacacccc-catgcg
ttgtcatagagcatgctcaggtacagaaagtgctgtgactttcttggcctgctgtcactg
agtcagcctgatactttggaggaatcttttctgggcactgcccagctgagagttcttatc
aggctctggagtgtgcatatgcctatattcattatagtgcatgtggctacgcagaaaaag
catcgacgtggaatggagaagaacaccattctgaatggaacggttcttactctacttctg
ttgcttgacgctgtatgggaatcgctttgctctgggggtacaccttccattccatgacaa
agcatttgatctatgccccga-gggtcattctctatgaagagcccgaaacttgtctgagt
gagaggcgaaggtagtccttcattt--atgcggggtccgccctcacactgtagtcggcac
agtcccttgtttatggactgcgcagctcaatcttcgactggagagg--aaaaacaaaaca

caagtatgacacgcacatacgtgttttctaggactgtgacctgtcggctgcctcgcggtg
gttggcatgatgccccaagtggggggggggagaaacaatcagtatgattgataaaaacca
gtgggg-aaaaaggcgcgctgtcgcacacccttggtcacggcctc---aagtcgacggcc
ctcactggccca-ccctgcgactcctgttcgcttttgacaagacacacctgcaata-cgg
aggaaatgcctgagcagctgtgccgcagcaactgcggacgcgtagcgatatgcatccctg
ttgtttgcgacttggtacaaagaacgtgcgccgggagacggattgccctttgacttttcc
acccagcccgcggctctatacccttgacgtacctatttacctggttgatcctgccagtag
tcatatgcttgtctcaaagactaagccatgcatgtctaagtataaacacttttatactcg
tgaaactgcgaatggctcattaaaacagttatagttcctttgggaacaacactacttgga
taaccgtagtaattctagagctaatacatgcctcaacgcccgactcacgaagggtggtat
ttattggataaaaaaccatcgtgctctgtccagagcgctttgagatgattcacaataact
tgtcggatcgcatggcctcgtgctggcgactcctcattcaaatttctgccctatcaactt
tcgatggtagagtattggtctaccatggtgtcgacgggtgacggggaattagggttcgat
tccggagagggagcctgagaaacggctaccacatccaaggaaggcagcaggcgcgcaaat
tacccaatcccgactcggggaggtagtgacaaaaaataacaatagggggccctttgggtc
ttctaattggaatgagaacaatttaaatcccttatcgaggatccattggagggcaagtct
ggt---------------------------------------------------------
------------------------------------------------------------
------------------------------------------------------------
------------------------------------------------------------
------------------------------------------------------------
------------------------------------------------------------
------------------------------------------------------------
------------------------------------------------------------
------------------------------------------------------------
------------------------------------------------------------
------------------------------------------------------------
------------------------------------------------------------
------------------------------------------------------------
------------------------------------------------------------
------------------------------------------------------------
------------------------------------------------------------
-----------------------------------------------gccagcagccgcg
gtaattccagctccaatagcgtatattaaagttgttgcagttaaaacgctcgtagtcgga
cctcggagcgcccccacgggccgggcttttgccttgttgccagtgggagtttgtctcctt
ttgtcgggacgcggtgtgcagggcctttactgtctctgcgcgtttacctgcgggccgacc

gtttactgtgaagaaagtaaagtgttcaaagcaggccaatgccttgaatatgtgagcatg
gaataatagaataggacttgggctctattttgttggttttcagtgaccaagtaatgatta
atagggacggttgggggcattcgtatttcattgtcagaggtgaaattcttggattgatgg
aagacgtacaactgcgaaagcatctgccatggatgttttcattgatcaagaacgaaagtt
aggggatcgaagacgatcagataccgtcgtagtcttaaccataaacgatgccgactgggg
attggcgggggcacttttatatgactctgtcagcaccctgagggaaaccaaagtctttgg
gttctgggggggagtatggtcgcaaggctgaaacttaaaggaattgacggaagggcaccac
caggagtggagcctgcggcttaatttgactcaacacgggaaaacttaccaggtccggaca
gaagaatgattgacagactgaaagctctttcttgatttttttggttggtggtgcatggccg
ttcttagttggtggagtgatttgtctggttaattccgttaacgaacgagacctcgtcctg
ctaaataggtgcgcgcatgccagcactgcgttcttaccttcttagagggactatgcgcgt
ctagcgtatggaagattgaggcaataacaggtctgtgatgcccttagatgttctgggccg
cacgcgcgctacactgatgcattcaacgagtttcacttgatagtcctgggtcggaaggct
cgggtaatcttttgaaagtgcatcgtgctggggatagatcattgcaattattgatcttca
acgaggaattccttgtaggcgcaggtcatcagcctgcgccgaatacgtccctgccctttg
tacacaccgcccgtcgctcctaccgattgaatggtccgatgaaatgtccggatcgcggct
agtgtgatggactcgtccagagcggtagccgtgagaagctcattaaatcttatcatttag
aggaaggagaagtcgtaacaaggt------------------------------------
------------------------------------------------------------
------------------------------------------------------------
------------------------------------------------------------
------------------------------------------------------------
------------------------------------------------------------
------------------------------------------------------------
------------------------------------------------------------
------------------------------------------------------------
------------------------------------------------------------
------------------------------------------------------------
------------------------------------------------------------
------------------------------------------------------------
------------------------------------------------------------
------------------------------------------------------------
------------------------------------------------------------
-------------------------------ttccgtaggtgaacctgcggaaggatcat
tcacaaactattgacacaaacacacgcgaaccaaatcgtccgcacagcaggtggcaatga
aagagagactctgcatgtcgcctttcgggg--tagcaagcagcactcttctgccatcgcc

tctgtgcagggcgtaaattcccattgagaggatgtgagggcaccacaggaagcttttcca
caggaagtcgccatccttctccctccacggcggcg----cttctgtgcttggcag-----
----tttttttttgccttctagggaggatgccgccaatggagccccttat-atatataca
tcatcatatgccc--ctttttttcttaaccgcttgccacagcttcttctatgaggagctt
gtgggaagactgtctccatacaataac-aaagatacaactcttagcggtggatatcttgg
ttctcgcaacgatgaagaacgcagccaactgcgataactaatgtgaattgcaggacttcg
tgaatcattaagtctttgaacgcaagttgcgctcatctccatgaggaggtgagcatgcct
gtttgagtgtcgatgggaaaaacaatac-ataacaacaatcggctctcttatcttcctgc
cttcttgggggcaaaaggtagtgcatgccggagatgggctttgtcgcccctttctcttga
aagaggcgtcgtccgaaacacgtatcgttgcccgctggaggtcatagagcatgagttt--
aaagcaccacatctctcaagtcagagtgag-cgtcttgagtaggtcc-gcattgattcag
tttcaacccaggtggcgttggttacttgccat-aagcctctctggggtttctttgcttgg
acctctcttgtcagtttctctgacgtagagacaggt--gccgtctcatggacgttgccct
ctggaacgtgcta-aaaaaaaaaaaaaga----------aaaaaccgtaccacatgtcaa
aagagacagatcgtgcttgacggcgggaggtgttgcatttcttgcttgtcctgagcgtct
gccagtctct----ga--cgagtg----tgttccttccataaacccgatctcaaatcagg
taggaccacccgctgaatttaagcatataattaagcggaggaaaagaaactaacaaggat
tccctcagtagcggcgagcgaagcgggaagagctcaacatgaaaaccgttgggggttcgc
gctttccttcgggttgtatagtaaagtggtgtcgcctgcgtcgcgccgggtataaatccc
gtctgaaatggggtgtcgtagagggttatatccccgtctgtgacccggtgtgcgatgcgt
tacggtgaccttacgatgagtcgggttgcttgggcatgcagccctaagtgggtggtaaac
ttcatctaaagctaaatactagcgagagaccgatagcgtacaagtaccgtaagggaaaga
tgcaaagaactttgaaaagagagttaaacagggcttgaaaccgttaagagtgaagcgaag
gaggtcagtgatcatggggccagatccccaagtgcctgtcata-gggcaccgtagcgttc
tggtttcccggggcagcgccggccgggggcgggggttaacagggaggtggttgagggatt
cgttttccgcctgtactcctttatagcttgcaccccggccgagggatccgcgacttgtcg
tgtgagggcgctgtcaaaatggccttctccaccccgtcttgaaacacggaccaaggagtc
taacatactcgcgagtgtttgggtggttaaaacccgaacgcacaatgaaagtaaacggtg
gctctgcgccgctgcttggtgggaaccctgcttttataatgggggcactatcacccgac
cccgatccctttggggtggttggttttgaggtagagcgagtatgatgggacccgaaagat
ggtgaactattcctgagtagggcgaagccagaggaaactctggtggaggctcgtagcgat
gctgacgtgcaaatcgctcgtcgaacttgggtataggggcgaaagactaatcgaaccatc
tagtagctggttccctccgaagtttccctcaggatagcgggagcaaatgcaagtcagtat
catccggtaaagcgaatgattagaggcctcgggggagtaactccttcgacctattctcaa
actttaaataggtgaggagccggcaacgttgcttaaaacaatgaacgccgtgcggacaat
gattgctctcagtgggccgattttggtaagcagaactggcgatgcgggatgaaccgaaag

cgacgctaaggtgccaaactcccagctcattcagacaccagaaaaggtgttgattcatac
agacagcaggacggtggccatggaagtcggaatccgctaaggagtgtgtaacaactcacc
tgccgaatggatcagccctgaaaatggatggcgcttaagcgggggaccgaggcgtcgccg
ttggggtaaatacgaagccccaatgagtaggagggcgtgggggttttgaaacacggtaca
ctcgcatggaagccttgggcgtaagctcgggtggagcgacccctagtgtagatcttggtg
gtagtagcaaatattcaagtgagaaccttgaagactgaagtggggaagggttccacgcga
acagtgcttggacgtgggttagtcggtcctaagagatggggaaactccgtattcaagtgc
gcttttgtagcgcagtccatcgaaagggaagcgggttaacattcccgcaccaggatgtgg
aaactgtggtttaccgcaaactgcggcaacgcaactgaacctggggacgcccgtaggagc
tccgagcagggatctctttcctgtttaacgcgcccaacccccctggaatcggtttgtccgg
agatagggggcgaaaggccggaaaagcaccgcacgtcttgtggtgttcggaacgcttgta
atggcccttgaaaacccaggggaaagcagccgtcttacac--ttgtgtgagacgaggaat
tctcacacctggccgtacccatcaccgcatcaggtccccaaggtgaacagcctctagtca
atagaataatgtagataagggaagtcggcaaaatagatccgtaactttgggaaaaggatt
ggctctaatgttcgggcttgtcgggacgcgcgggcgtagatgccaatgagcggcagacgt
ctagcctctttggaggctggattaacgctgcttgtgggaagaaaccgtgtcgatggaggc
aggaacgcttatcgatcttgtctcttcccgcttgcgatcaacgaacggattagaactggt
acggacaaggggaatccgactgtttaattaaaacaaagcattgcgacggccggaaacggt
gttggcgcaatgtgatttctgcccagtgctctgaatgtcaaagtgaagaaattcaaccaa
gcgcgggtaaacggcgggagtaactatgactctcttagac-agatcaagacaccttcaat
aaattggttcattatgtcgggtggtccgtcgtccgggaagcgaccaaagcaaaaccccgt
agactgtgatgtgcatctgtcagacgaaactgacgtccgtgttcatcaagccttagctat
ctttcaacgccggcttcgagacgtggcgtctcttaaggtgtttgaagtggctctctctgt
gggaactactgatgcactggtagagatgggcactatcacttgctggagtattgatgctgt
tgggaaaggtttctcaaggaacagcagcggtcgctggcaatttgacctcactggcaagat
taatggcaagcctctaatatccaccacaaaggatcgttggaggccggttgtgacggctgt
gccctactacctttcgtgcgccatcgatggggacaagacgacgctggaggaaaagcagtg
gacttcatcacacatgtgtcaccatgaatggtgtgtgaacgtggagcacattgtttatga
aagcctcgacgtcaac--aagtctcggaactggtg--ccagggaaaaaacgcatgcctcc
atctgcctgtttgcattcgaaccggccgccgtgtgcaggagctcaggaagagcaatgggg
ttcgtgcaatggacctcctcctgcagaaattggctggattggagattgaggttgaggagc
ccgctgcagctgtactccgagagtctgtcgaacgggaggacgcctagtgagtgactcgtg
acacgggttggctgcgcatgatgcgtcaactacatggacatcttttgagagagataggcg
ggctgaaaaggttatagccgtctagtctcgctctctttcattgagcggggcgacacacct
ggttgcgggaaccctgcccacctttctatcaagaagtgggaaccagtccattcctactcc
tgcttgtgaggcatatagt--aacgtagcagcaggaggatctcggcgagaggccgaaccc

gaaaggaataacgtgggctgggaggcgatccgcaggcgagtccgtaactatatgattatc
gacgggccgtctcaacgactggtaaggtgtgggtttg-tcctcctcctcacgtggggaca
ggcttaaggtacagtctaatcccacccgccaagggtgcttggg---tg------------
--------------------taatccctcttccatagagggcactagccca-ggggctgg
gacgctgggaggaa----------------atgccccgtccctatagccggtagaaatga
aggtagccaaatgcctcgtcatctaaactaattgg-at----------------------
-----------------------gagatagtc----------tgagtgaaaagcttatgc
ag-----------------gctagtaccgg---aaaaaaaaaaaaagaggggctctgcca
ct----ctgac------------------------ggtgcgacactccttgtacagggaa
accctcc--gcatggaaaggcacgtggtaccaccctcgcggggaaactcgcgagcgggat
cccgggtaatgaccggagccgatggtaaaaactcacgtgaccgatggggtgaccctgtgg
cgagacgctaagttgtgggacatcaccctctcacaatacgcatccgtcgcagagactgaa
cgggagtgggctccgttga-----cggccggccggccagccagccgtcttcca-------
---ggggcttaagatacagtccgggcctcagggatgg----aaaaacctggggaacta-c
cagaattagtgacgcgcatgaatggattaacgagattcccactgtccctatctactatct
agcgaaaccacagccaagggaacgggcttggaagaatcagcggggaaagaagaccctgtt
gagcttgactctagtccgactttgtgagatgagttgggaggtgtagtctaagagggaggg
gcaactcgaccgtgaaatacctctactcccaagcttgttttacttactccggcgagcagg
gatgggtggtaacaccctttttctaggattaacacctcttcgggggtggacctgcgcgga
ggacattgtcaggtggggagtttggctggggcggcacatctgttaaaagataacgcaggt
gtcctaagatgagctcagtgagaacagaaatctcacgtagaacaaaagggtaaaagctca
tttgattttgattttcagtacgaatacaaactgtgaaagcatggcctatcgatcctttttg
accttcaaggtttgaagcaagaggtgtcagaaaagttaacacgcgttgtggtagctttta
agagtggactcaaagtgttccgctagtccccccctcggaccgccacttacaaaaggcctgg
ggatgggggcgacactgtcaaattgcggggactccctaaagcctgtctctaccaagccag
cgcggcgaaagcccctggtggccggggtaatgacctagggtatggtaaaagcgagacagg
atg-gaacaatgggcaatccgcagccaagtcctaagg----ctttttgaaaagcggctat
ggatgcagttcaacgactagatggcagtgggctgagagagcccaccgcggcaccctttcg
gcttaagatatagtcttgcccacgtgaaagcgtgcctgcagatgaacgccaccatgtatg
gccggagctgcaggagaccgc--acttgatgtgcagtcgggaat-cgacagtaccacagg
gatgaac-----------------------------------------------------
------------------------------------------------------------
------------------------------------------------------------
------------------------------------------------------------
------------------------------------------------------------
-------------------------------------------------aaaaacac---

------------------------------------------------------------
------------------------------------------------------------
--acaa------------------------------------------------------
------------------------------------------ttgtggcc---atcatac
----------------------------------------agtcgta-------------
------------------------------------------------------------
------------------------------------------------------------
------------------------------------------------------------
------------tgatattatcgtcccaggcgggtgcgcctctctcacccgtctagtccg
gccttggtgcgcaaaaaatgcttgcgtggatgcgccttggttaaggaaaatgctcctagt
agttttacaaggacacttcttgtagggcaacacatctggatgcgaagaactcccggaaaa
gacaacggtaccaagtgactggggtaacgccccattcatgaaggcaaataatcaaaaaga
ggtgccgggtaagaacccgttgtatagggacaatccgcagctgacttggctccctg--cc
cat--ccatgggcatagcttagcggttcaacgctcggtaagatgtgggttggcgcgctgt
ccgacagcgcactggcttaaggtacgggctactcccaccggagacggtgtctttgtctac
ctcgggtaacacgcctcgggagaagcaaagagggtgcaatgg-aaaaaaactggttgcgg
ggaaatccctaccagagtcacctggtacaaaggaactggcttgtggcagccaagcgttca
tagcgacgttgctttttgatccttcgatgtcggctcttcctatcattgcgaagcagaatt
cgccaagtgttggattgttcacccactaatagggaacgtgagctgggtagccacttta--
-----tgcccaacagctggatagctgggcttctgtctta-ggccgggacttctcacgacc
cgtgccttggtcaccc-aaaaaagggcgaccgcagagctcaggatggagcttctggctta
actagtagaggacagatagagatcgtcaaattgcgggaaaccccttagagcctcgagcta
ccaagtgcgtgtcgaaaggcatagacgcagccagcgatggctgtcgcatggcttga-ttc
catcaagataggtaaaaatgctcgaggattgggcaatccgcagccaagcccctaaggt--
---ccccccgcgggctatggggaaggttcaacgactgtaaggcgtaccgtgcaagcggaa
gatacagtctagccccgctggaaagcagcgccaaagtaagtaggt-tgcagtgggtctgt
cctc
>Form_E_China_SRR9587968
gcatgttttggtatgtgtttcgacagtcatgcatgtcatagccgtaggatggggagtgag
cttcatagcccgaccccttctcaggagcatgttttcatagcatattgagcatactggggt
ggtctcccatgtagctcgctctctcggtgcaggacctcccttttgcgggccacttcgatg
gcctccattatgga-gagcatgttttggcatatgaagcgttggtggtgaggctcgtagcc
tgtatatacataccaacgcagtcgtcatatgtctctctggcgggggtcggtcgtatgacc
ttgtccctacctggagcatgtattttgtatgat---atactggac-ggcagctgcgctgt
tcagtcgtcggccgtacctcatgtgcgcgtttccaggtccgaacgtttgacatgcaaggg
catcataccagaagcctgttttctcgtatgagcatgtaggtaggcaggctcttcagtcgt

ctacccataaatgcacttcgaagga-actttttggtattgaatggctttgtgcccagacg
ttcttgtgagcttgttctcgctctgcgatacgctggaagctgtggtattacctacagcag
acgcattcctagggcggacggcctctctgcagatgggcacatctgtagaaagaggctccc
cttgatgacgcaatgcaatggctgggtaaacggagtgctatacagaagggtcaaacctc-
tgtggaagtacactcataccagcattgtttgttgagaggggtcggctctacgagtgaggc
cctatactgccatctcattcagatgcgtagacggtctttactcagactgcctttctggct
gattgtcctcggtccctcgtgcgcacgcgccgctggctgaatgggcttgtccttatggag
tgccgcgttcttc-aaagaatccgtcttcatacagcgacctcatttacatattgccggct
gaatgggcttgtccttatggagcactgcattcttctaggaattgctatctatacagcgac
ctcatttacatacaaggcacct-ttttctacatagacttgccatttggctgattgttctg
acagtcgctgcctcttcggagttctgtgtggacgagtggcc--tcctgagtgggccggtc
cttacggagtgcccaagtctgttatgactgggttctccatacagcgacctcactcagaat
tgtatgg--------------caccgtgccgat-----------tgaatggacaagcagt
ccttagggaatcgcatgaaagcatgacggcaacatggtgtatggacccgttgggccttgc
ctgtgctatcttctcgagtcgaccgg------------tcctcgtctcctcctttcgcga
cgggaagctcttgagtttaccgtgggtcaggacgctgtggtatctgtttgtctgtagtgt
gggctctctatcagaacagcgtatcagcacacgtcgtctgggtctatggtatttttgctt
aacggccctgggtttggtagtatggttccatgtcggtcctctcacctcccaaccgtcatg
tctctga-----------------------------------------------------
------------------------------------------------------------
--------------------------------------------tatcatctgcacagta
cgtcttgtcagaatatcattctgtatggcagtcgatagactaggccctatgccgcagctg
tctctgaccactatgcgagcgcatacctacctcaatgtgttcatagctgttatggatctt
gggtgtggtgtgccagaacgatacgccaccagattgtggaatggtgcttgtagtattaga
agaggacatgtatgaagagcacgttcttttctcttatgatggtgccgcgggtgcgggccg
ttctcattctgcttgtcgggtgctcaaggtgcatgtaggattgctgaacccac-tgaatg
gtcatgagaatggtcatatggtgaggttcacttctccatcgctttgtgtgtgttggatgt
ttgggtggcttcggccgcatggaaaatctcggtctgtaactcatcagggagggtttccat
gtgagcaatctctccagtcgaacacacacccatgggcgtcccgctaggcaaaatggaact
ctcgatccgacacagcctgtggagtgtttgttccgtgtggcgtgaaacggcttatactgg
atgtcttccc-tttggacaccagtctgttggtatg--tacgcataccatctagagggagc
aagaaaaaggcatgcaagtgccacaaaggaagggaggccttcctgctaccgtgtgtcc-t
cttgacgatcacgggggagtatatagagacatttggacgagatccgtttttcacgaggcg
gtgccctcgcgcaaaggtgttgctctgcacgcatctttatgatgatagttgctcgctcct
atgcgtccactcgtgaaaatacctgctctctctgtctgtttccatgacacccc-catgcg
ttgtcatagagcatgctcaggtacagaaagtgctgtgactttcttggcctgctgtcactg

agtcagcctgatactttggaggaatcttttctgggcactgcccagctgagagttcttatc
aggctctggagtgtgcatatgcctatattcattatagtgcatgtggctacgcagaaaaag
catcgacgtggaatggagaagaacaccattctgaatggaacggttcttactctacttctg
ttgcttgacgctgtatgggaatcgctttgctctgggggtacaccttccattccatgacaa
agcatttgatctatgccccga-gggtcattctctatgaagagcccgaaacttgtctgagt
gagaggcgaaggtagtccttcattt--atgcggggtccgccctcacactgtagtcggcac
agtcccttgtttatggactgcgcagctcaatcttcgactggagagg--aaaaacaaaaca
caagtatgacacgcacatacgtgttttctaggactgtgacctgtcggctgcctcgcggtg
gttggcatgatgccccaagtgggggggggagaaacaatcagtatgattgataaaaacca
gtgggg-aaaaaggcgcgctgtcgcacacccttggtcacggcctc---aagtcgacggcc
ctcactggccca-ccctgcgactcctgttcgcttttgacaagacacacctgcaata-cgg
aggaaatgcctgagcagctgtgccgcagcaactgcggacgcgtagcgatatgcatccctg
ttgtttgcgacttggtacaaagaacgtgcgccgggagacggattgccctttgacttttcc
acccagcccgcggctctatacccttgacgtacctatttacctggttgatcctgccagtag
tcatatgcttgtctcaaagactaagccatgcatgtctaagtataaacacttttatactcg
tgaaactgcgaatggctcattaaaacagttatagttcctttgggaacaacactacttgga
taaccgtagtaattctagagctaatacatgcctcaacgcccgactcacgaagggtggtat
ttattggataaaaaaccatcgtgctctgtccagagcgctttgagatgattcacaataact
tgtcggatcgcatggcctcgtgctggcgactcctcattcaaatttctgccctatcaactt
tcgatggtagagtattggtctaccatggtgtcgacgggtgacggggaattagggttcgat
tccggagagggagcctgagaaacggctaccacatccaaggaaggcagcaggcgcgcaaat
tacccaatcccgactcggggaggtagtgacaaaaaataacaatagggggccctttgggtc
ttctaattggaatgagaacaatttaaatcccttatcgaggatccattggagggcaagtct
ggt---------------------------------------------------------
------------------------------------------------------------
------------------------------------------------------------
------------------------------------------------------------
------------------------------------------------------------
------------------------------------------------------------
------------------------------------------------------------
------------------------------------------------------------
------------------------------------------------------------
------------------------------------------------------------
------------------------------------------------------------
------------------------------------------------------------
------------------------------------------------------------

------------------------------------------------------------
------------------------------------------------------------
------------------------------------------------------------
-----------------------------------------------gccagcagccgcg
gtaattccagctccaatagcgtatattaaagttgttgcagttaaaacgctcgtagtcgga
cctcggagcgcccccacgggccgggcttttgccttgttgccagtgggagtttgtctcctt
ttgtcgggacgcggtgtgcagggcctttactgtctctgcgcgtttacctgcgggccgacc
gtttactgtgaagaaagtaaagtgttcaaagcaggccaatgccttgaatatgtgagcatg
gaataatagaataggacttgggctctattttgttggtttccagtgaccaagtaatgatta
atagggacggttgggggcattcgtatttcattgtcagaggtgaaattcttggattgatgg
aagacgtacaactgcgaaagcatctgccatggatgttttcattgatcaagaacgaaagtt
aggggatcgaagacgatcagataccgtcgtagtcttaaccataaacgatgccgactgggg
attggcgggggcacttttatatgactctgtcagcaccctgagggaaaccaaagtctttgg
gttctgggggggagtatggtcgcaaggctgaaacttaaaggaattgacggaagggcaccac
caggagtggagcctgcggcttaatttgactcaacacgggaaaacttaccaggtccggaca
gaagaatgattgacagactgaaagctctttcttgatttttggttggtggtgcatggccg
ttcttagttggtggagtgatttgtctggttaattccgttaacgaacgagacctcgtcctg
ctaaataggtgcgcgcatgccagcactgcgttcttaccttcttagagggactatgcgcgt
ctagcgtatggaagattgaggcaataacaggtctgtgatgcccttagatgttctgggccg
cacgcgcgctacactgatgcattcaacgagtttcacttgatagtcctgggtcggaaggct
cgggtaatcttttgaaagtgcatcgtgctggggatagatcattgcaattattgatcttca
acgaggaattccttgtaggcgcaggtcatcagcctgcgccgaatacgtccctgccctttg
tacacaccgcccgtcgctcctaccgattgaatggtccgatgaaatgtccggatcgcggct
agtgtgatggactcgtccagagcggtagccgtgagaagctcattaaatcttatcatttag
aggaaggagaagtcgtaacaaggt------------------------------------
------------------------------------------------------------
------------------------------------------------------------
------------------------------------------------------------
------------------------------------------------------------
------------------------------------------------------------
------------------------------------------------------------
------------------------------------------------------------
------------------------------------------------------------
------------------------------------------------------------
------------------------------------------------------------
------------------------------------------------------------

------------------------------------------------------------
------------------------------------------------------------
------------------------------------------------------------
------------------------------------------------------------
-------------------------------ttccgtaggtgaacctgcggaaggatcat
tcacaaactattgacacaaacacacgcgaaccaaatcgtccgcacagcaggtggcaatga
aagagagactctgcatgtcgcctttcgggg--tagcaagcagcactcttctgccatcgcc
tctgtgcagggcgtaaattcccattgagaggatgtgagggcaccacaggaagcttttcca
caggaagtcgccatccttctccctccacggcggcg----cttctgtgcttggcag-----
-----ttttttttgccttctagggaggatgccgccaatggagccccttat-atatataca
tcatcatatgccc--ctttttttcttaaccgcttgccacagcttcttctatgaggagctt
gtgggaagactgtctccatacaataac-aaagatacaactcttagcggtggatatcttgg
ttctcgcaacgatgaagaacgcagccaactgcgataactaatgtgaattgcaggacttcg
tgaatcattaagtctttgaacgcaagttgcgctcatctccatgaggaggtgagcatgcct
gtttgagtgtcgatgggaaaaacaatac-ataacaacaatcggctctcttatcttcctgc
cttcttgggggcaaaaggtagtgcatgccggagatgggctttgtcgcccctttctcttga
aagaggcgtcgtccgaaacacgtatcgttgcccgctggaggtcatagagcatgagttt--
aaagcaccacatctctcaagtcagagtgag-cgtcttgagtaggtcc-gcattgattcag
tttcaacccaggtggcgttggttacttgccat-aagcctctctggggtttctttgcttgg
acctctcttgtcagtttctctgacgtagagacaggt--gccgtctcatggacgttgccct
ctggaacgtgcta-aaaaaaaaaaaaaga----------aaaaaccgtaccacatgtcaa
aagagacagatcgtgcttgacggcgggaggtgttgcatttcttgcttgtcctgagcgtct
gccagtctct----ga--cgagtg----tgttccttccataaacccgatctcaaatcagg
taggaccacccgctgaatttaagcatataattaagcggaggaaaagaaactaacaaggat
tccctcagtagcggcgagcgaagcgggaagagctcaacatgaaaaccgttgggggttcgc
gctttccttcgggttgtatagtaaagtggtgtcgcctgcgtcgcgccgggtataaatccc
gtctgaaatggggtgtcgtagagggttatatccccgtctgtgacccggtgtgcgatgcgt
tacggtgaccttacgatgagtcgggttgcttgggcatgcagccctaagtgggtggtaaac
ttcatctaaagctaaatactagcgagagaccgatagcgtacaagtaccgtaagggaaaga
tgcaaagaactttgaaaagagagttaaacagggcttgaaaccgttaagagtgaagcgaag
gaggtcagtgatcatggggccagatccccaagtgcctgtcata-gggcaccgtagcgttc
tggtttcccggggcagcgccggccgggggcgggggttaacagggaggtggttgagggatt
cgttttccgcctgtactcctttatagcttgcaccccggccgagggatccgcgacttgtcg
tgtgagggcgctgtcaaaatggccttctccaccccgtcttgaaacacggaccaaggagtc
taacatactcgcgagtgtttgggtggttaaaacccgaacgcacaatgaaagtaaacggtg
gctctgcgccgctgcttggtgggaaccctgcttttataatgggggggcactatcacccgac

cccgatccctttggggtggttggttttgaggtagagcgagtatgatgggacccgaaagat
ggtgaactattcctgagtagggcgaagccagaggaaactctggtggaggctcgtagcgat
gctgacgtgcaaatcgctcgtcgaacttgggtataggggcgaaagactaatcgaaccatc
tagtagctggttccctccgaagtttccctcaggatagcgggagcaaatgcaagtcagtat
catccggtaaagcgaatgattagaggcctcgggggagtaactccttcgacctattctcaa
actttaaataggtgaggagccggcaacgttgcttaaaacaatgaacgccgtgcggacaat
gattgctctcagtgggccgattttggtaagcagaactggcgatgcgggatgaaccgaaag
cgacgctaaggtgccaaactcccagctcattcagacaccagaaaaggtgttgattcatac
agacagcaggacggtggccatggaagtcggaatccgctaaggagtgtgtaacaactcacc
tgccgaatggatcagccctgaaaatggatggcgcttaagcgggggaccgaggcgtcgccg
ttggggtaaatacgaagccccaatgagtaggagggcgtgggggttttgaaacacggtaca
ctcgcatggaagccttgggcgtaagctcgggtggagcgacccctagtgtagatcttggtg
gtagtagcaaatattcaagtgagaaccttgaagactgaagtggggaagggttccacgcga
acagtgcttggacgtgggttagtcggtcctaagagatggggaaactccgtattcaagtgc
gcttttgtagcgcagtccatcgaaagggaagcgggttaacattcccgcaccaggatgtgg
aaactgtggtttaccgcaaactgcggcaacgcaactgaacctggggacgcccgtaggagc
tccgagcagggatctctttcctgtttaacgcgcccaacccccctggaatcggtttgtccgg
agatagggggcgaaaggccggaaaagcaccgcacgtcttgtggtgttcggaacgcttgta
atggcccttgaaaacccaggggaaagcagccgtcttacac--ttgtgtgagacgaggaat
tctcacacctggccgtacccatcaccgcatcaggtccccaaggtgaacagcctctagtca
atagaataatgtagataagggaagtcggcaaaatagatccgtaactttgggaaaaggatt
ggctctaatgttcgggcttgtcgggacgcgcgggcgtagatgccaatgagcggcagacgt
ctagcctctttggaggctggattaacgctgcttgtgggaagaaaccgtgtcgatggaggc
aggaacgcttatcgatcttgtctcttcccgcttgcgatcaacgaacggattagaactggt
acggacaaggggaatccgactgtttaattaaaacaaagcattgcgacggccggaaacggt
gttggcgcaatgtgatttctgcccagtgctctgaatgtcaaagtgaagaaattcaaccaa
gcgcgggtaaacggcgggagtaactatgactctcttagac-agatcaagacaccttcaat
aaattggttcattatgtcgggtggtccgtcgtccgggaagcgaccaaagcaaaaccccgt
agactgtgatgtgcatctgtcagacgaaactgacgtccgtgttcatcaagccttagctat
ctttcaacgccggcttcgagacgtggcgtctcttaaggtgtttgaagtggctctctctgt
gggaactactgatgcactggtagagatgggcactatcacttgctggagtattgatgctgt
tgggaaaggtttctcaaggaacagcagcggtcgctggcaatttgacctcactggcaagat
taatggcaagcctctaatatccaccacaaaggatcgttggaggccggttgtgacggctgt
gccctactacctttcgtgcgccatcgatggggacaagacgacgctggaggaaaagcagtg
gacttcatcacacatgtgtcaccatgaatggtgtgtgaacgtggagcacattgtttatga
aagcctcgacgtcaac--aagtctcggaactggtg--ccagggaaaaaacgcatgcctcc

atctgcctgtttgcattcgaaccggccgccgtgtgcaggagctcaggaagagcaatgggg
ttcgtgcaatggacctcctcctgcagaaattggctggattggagattgaggttgaggagc
ccgctgcagctgtactccgagagtctgtcgaacgggaggacgcctagtgagtgactcgtg
acacgggttggctgcgcatgatgcgtcaactacatggacatcttttgagagagataggcg
ggctgaaaaggttatagccgtctagtctcgctctctttcattgagcggggcgacacacct
ggttgcgggaaccctgcccacctttctatcaagaagtgggaaccagtccattcctactcc
tgcttgtgaggcatatagt--aacgtagcagcaggaggatctcggcgagaggccgaaccc
gaaaggaataacgtgggctgggaggcgatccgcaggcgagtccgtaactatatgattatc
gacgggccgtctcaacgactggtaaggtgtgggtttg-tcctcctcctcacgtggggaca
ggcttaaggtacagtctaatcccacccgccaagggtgcttggg---tg------------
--------------------taatccctcttccatagagggcactagccca-ggggctgg
gacgctgggaggaa----------------atgccccgtccctatagccggtagaaatga
aggtagccaaatgcctcgtcatctaaactaattgg-at----------------------
-----------------------gagatagtc----------tgagtgaaaagcttatgc
ag-----------------gctagtaccgg---aaaaaaaaaaaaagaggggctctgcca
ct----ctgac------------------------ggtgcgacactccttgtacagggaa
accctcc--gcatggaaaggcacgtggtaccaccctcgcggggaaactcgcgagcgggat
cccgggtaatgaccggagccgatggtaaaaactcacgtgaccgatggggtgaccctgtgg
cgagacgctaagttgtgggacatcaccctctcacaatacgcatccgtcgcagagactgaa
cgggagtgggctccgttga-----cggccggccggccagccagccgtcttcca-------
---ggggcttaagatacagtccgggcctcagggatgg----aaaaacctggggaacta-c
cagaattagtgacgcgcatgaatggattaacgagattcccactgtccctatctactatct
agcgaaaccacagccaagggaacgggcttggaagaatcagcggggaaagaagaccctgtt
gagcttgactctagtccgactttgtgagatgagttgggaggtgtagtctaagagggaggg
gcaactcgaccgtgaaatacctctactcccaagcttgttttacttactccggcgagcagg
gatgggtggtaacacccttttctaggattaacacctcttcgggggtggacctgcgcgga
ggacattgtcaggtggggagtttggctggggcggcacatctgttaaaagataacgcaggt
gtcctaagatgagctcagtgagaacagaaatctcacgtagaacaaaagggtaaaagctca
tttgattttgattttcagtacgaatacaaactgtgaaagcatggcctatcgatcctttg
accttcaaggtttgaagcaagaggtgtcagaaaagttaacacgcgttgtggtagcttta
agagtggactcaaagtgttccgctagtccccccctcggaccgccacttacaaaaggcctgg
ggatgggggcgacactgtcaaattgcggggactccctaaagcctgtctctaccaagccag
cgcggcgaaagcccctggtggccggggtaatgacctagggtatggtaaaagcgagacagg
atg-gaacaatgggcaatccgcagccaagtcctaagg----ctttttgaaaagcggctat
ggatgcagttcaacgactagatggcagtgggctgagagagcccaccgcggcaccctttcg
gcttaagatatagtcttgcccacgtgaaagcgtgcctgcagatgaacgccaccatgtatg

gccggagctgcaggagaccgc--acttgatgtgcagtcgggaat-cgacagtaccacagg
gatgaac-----------------------------------------------------
------------------------------------------------------------
------------------------------------------------------------
------------------------------------------------------------
------------------------------------------------------------
-------------------------------------------------aaaaacac---
------------------------------------------------------------
------------------------------------------------------------
--acaa------------------------------------------------------
------------------------------------------ttgtggcc---atcatac
----------------------------------------agtcgta-------------
------------------------------------------------------------
------------------------------------------------------------
------------------------------------------------------------
------------tgatattatcgtcccaggcgggtgcgcctctctcacccgtctagtccg
gccttggtgcgcaaaaaatgcttgcgtggatgcgccttggttaaggaaaatgctcctagt
agttttacaaggacacttcttgtagggcaacacatctggatgcgaagaactcccggaaaa
gacaacggtaccaagtgactggggtaacgccccattcatgaaggcaaataatcaaaaaga
ggtgccgggtaagaacccgttgtatagggacaatccgcagctgacttggctccctg--cc
cat--ccatgggcatagcttagcggttcaacgctcggtaagatgtgggttggcgcgctgt
ccgacagcgcactggcttaaggtacgggctactcccaccggagacggtgtctttgtctac
ctcgggtaacacgcctcgggagaagcaaagagggtgcaatgg-aaaaaaactggttgcgg
ggaaatccctaccagagtcacctggtacaaaggaactggcttgtggcagccaagcgttca
tagcgacgttgctttttgatccttcgatgtcggctcttcctatcattgcgaagcagaatt
cgccaagtgttggattgttcacccactaatagggaacgtgagctgggtagccacttta--
-----tgcccaacagctggatagctgggcttctgtctta-ggccgggacttctcacgacc
cgtgccttggtcaccc-aaaaaagggcgaccgcagagctcaggatggagcttctggctta
actagtagaggacagatagagatcgtcaaattgcgggaaaccccttagagcctcgagcta
ccaagtgcgtgtcgaaaggcatagacgcagccagcgatggctgtcgcatggcttga-ttc
catcaagataggtaaaaatgctcgaggattgggcaatccgcagccaagcccctaaggt--
---ccccccgcgggctatggggaaggttcaacgactgtaaggcgtaccgtgcaagcggaa
gatacagtctagccccgctggaaagcagcgccaaagtaagtaggt-tgcagtgggtctgt
cctc
>Form_F_China
gcatgttttggtatgtgtttcgacagtcatgcatgtcatagccgtaggatggggagtgag

cttcatagcccgaccccttctcaggagcatgttttcatagcatattgagcatactggggt
ggtctcccatgtagctcgctctctcggtgcaggacctcccttttgcgggccacttcgatg
gcctccattatgga-gagcatgttttggcatatgaagcgttggtggtgaggctcgtagcc
tgtatatacataccaacgcagtcgtcatatgtctctctggcgggggtcggtcgtatgacc
ttgtccctacctggagcatgtatttcgtatgat---atactggac-ggcagctgcgctgt
tcagtcgtcggccgtacctcatgtgcgcgtttccaggtccgaacgtttgacatgcaaggg
catcataccagaagcctgttttctcgtatgagcatgtaggtaggcaggctcttcagtcgt
ctacccataaatgcacttcgaagga-actttttggtattgaatggctttgtgcccagacg
ttcttgtgagcttgttctcgctctgcgatacgctggaagctgtggtattacctacagcag
acgcattcctagggcggacggcctctctgcagatgggcacatctgtagaaagaggctccc
cttgatgacgcaatgcaatggctgggtaaacggagtgctatacagaagggtcaaacctc-
tgtggaagtacactcataccagcattgtttgttgagaggggtcggctctacgagtgagga
cctatactgccatctcattcagatgcgtagacggtctttactcagactgcctttctggct
gattgtcctcggtccctcgtgcgcacgcgccgctggctgaatgggcttgtccttatggag
tgccgcgttcttc-aaagaatccgtcttcatacagcgacctcatttacatattgccggct
gaatgggcttgtccttatggagcactgcattcttctaggaattgctatctatacagcgac
ctcatttacatacaaggcacct-ttttctacatagacttgccatttggctgattgttctg
acagtcgctgcctcttcggagttctgtgtggacgagtggcc--tcctgagtgggccggtc
cttacggagtgcccacgtctgttatgactgggttctccatacagcgacctcactcagaat
tgtatgg--------------caccgtgccgat-----------tgaatggacaagcagt
ccttagggaatcgcatgaaagcatgatggc-acatggtgtatggacccgttgggccttgc
ctgtgctatcttctcgagtcgaccgg------------tcctcgtctcctcctttcgcga
cgggaagctcttgagtttaccgtgggtcaggacgctgtggtatctgtttgtctgtagtgt
gggctctctatcagaacagcgtatcagcacacgtcgtctgggtctatggtatttttgctt
aacggccctgggtttggtagtatggttccatgtcggtcctctcacctcccatccgtcatg
tctctga-----------------------------------------------------
------------------------------------------------------------
--------------------------------------------tatcatctgcacagta
cgtcttgtcagaatatcattctgtatggcagtcgatagactaggccctatgccgcagctg
tctctgaccactatgcgagcgcatacctacctcaatgtgttcatagctgttatggatctt
gggtgtggtgtgccagaacgatacgccaccagattgtggaatggtgcttgtagtattaga
agaggacatgtatgaagagcacgttcttttctcttatgatggtgccgcgggtgggggccg
ttctcattctgcttgtcgggtgctcaaggtgcatgtaggattgctgaacccac-tgaatg
gtcatgagaatggtcatatggtgaggttcacttctccatcgctttgtgtgtgttggatgt
ttgggtggcttcggccgcatggaaaatctcggtctgtaactcatcagggagggtttccat
gtgagcaatctctccagtcgaacacacacccatgggcgtcccgctaggcaaaatggaact

ctcgatccgacacagcctgtggagtgtctgttccgtgtggcgtgaaacggcttatactgg
atgtcttccc-tttggacaccagtctgttggtatg--tacgcataccatctagagggagc
aagaaaaaggcatgcaagtgccacaaaggaagggaggcctccctgctaccgtgtgtcc-t
cttgacgatcacgggggagtatatagagacatttggacgagatcctttttttcacgaggcg
gtgccctcgcgcaaaggtgttgctctgcacgcatctttatgatgatagttgctcgctcct
atgcgtccactcgtgaaaatacctgctctctctgtctgtttccatgacacccc-catgcg
ttgtcatagagcacgctcaggtacagaaagtgctgtgactttcttggcctgctgtcactg
agtcagcctgatactttggaggaatcttttctgggcgctgcccagctgagagttcttatc
aggctctggagtgtgcatatggctataatcataatagtgcatgtggccacgcagaaaaag
catcgacgtggaatggagaagaacaccattctgaatggaacggttcttactctgcttccg
ttgcttgacgctgtatgggaatcgctttgccctgggggtacaccttccattccatgacaa
agcatttgatctatgccccga-gggtcattctctatgaagagcccgaaacttgtctgagt
gagaggcgaaggtagcccttcattt--atgcggggtccgccctcacactgtagtcggcac
agtcccttgtttatggactgcgcagctcaatcttcgactggagagg--aaaaacaaaaca
caagtatgacacgcacatacgtgttttctaggactgtgacctgtcggctgcctcgcggtg
gttggcatgatgccccaagtgggggggggagaaacaatcagtatgattgataaaaacca
gtgggg-aaaaaggcgcgctgtcgcacacccttggtcacggcctc---aagtcgacggcc
ctcactggccca-ccctgcgactcctgttcgcttttgacaagacacacctgcaata-cgg
aggaaatgcctgagcagctgtgccgcagcaactgcggacgcgtagcgatatgcatccctg
ttgtttgcgacttggtacaaagaacgtgcgccgggagacggattgccctttgacttttcc
acccagcccgcggctctatacccttgacgtacctatttacctggttgatcctgccagtag
tcatatgcttgtctcaaagactaagccatgcatgtctaagtataaacacttttatactcg
tgaaactgcgaatggctcattaaaacagttatagttcctttgggaacaacactacttgga
taaccgtagtaattctagagctaatacatgcctcaacgcccgactcacgaagggtggtat
ttattggataaaaaaccatcgtgctctgtccagagcgctttgagatgattcacaataact
tgtcggatcgcatggcctcgtgctggcgactcctcattcaaatttctgccctatcaactt
tcgatggtagagtattggtctaccatggtgtcgacgggtgacggggaattagggttcgat
tccggagagggagcctgagaaacggctaccacatccaaggaaggcagcaggcgcgcaaat
tacccaatcccgactcggggaggtagtgacaaaaaataacaatagggggccctttgggtc
ttctaattggaatgagaacaatttaaatcccttatcgaggatccattggagggcaagtct
ggt---------------------------------------------------------
------------------------------------------------------------
------------------------------------------------------------
------------------------------------------------------------
------------------------------------------------------------
------------------------------------------------------------

------------------------------------------------------------

------------------------------------------------------------

------------------------------------------------------------

------------------------------------------------------------

------------------------------------------------------------

------------------------------------------------------------

------------------------------------------------------------

------------------------------------------------------------

------------------------------------------------------------

------------------------------------------------------------

-----------------------------------------------gccagcagccgcg

gtaattccagctccaatagcgtatattaaagttgttgcagttaaaacgctcgtagtcgga

cctcggagcgcccccacgggccgggcttttgccttgttgccagtgggagtttgtctcctt

ttgtcgggacgcggtgtgcagggcctttactgtctctgcgcgtttacctgcgggccgacc

gtttactgtgaagaaagtaaagtgttcaaagcaggccaatgccttgaatatgtgagcatg

gaataatagaataggacttgggctctattttgttggtttccagtgaccaagtaatgatta

atagggacggttgggggcattcgtatttcattgtcagaggtgaaattcttggattgatgg

aagacgtacaactgcgaaagcatctgccatggatgttttcattgatcaagaacgaaagtt

aggggatcgaagacgatcagataccgtcgtagtcttaaccataaacgatgccgactgggg

attggcgggggcacttttatatgactctgtcagcaccctgagggaaaccaaagtctttgg

gttctgggggggagtatggtcgcaaggctgaaacttaaaggaattgacggaagggcaccac

caggagtggagcctgcggcttaatttgactcaacacgggaaaacttaccaggtccggaca

gaagaatgattgacagactgaaagctctttcttgatttttggttggtggtgcatggccg

ttcttagttggtggagtgatttgtctggttaattccgttaacgaacgagacctcgtcctg

ctaaataggtgcgcgcatgccagcactgcgttcttaccttcttagagggactatgcgcgt

ctagcgtatggaagattgaggcaataacaggtctgtgatgcccttagatgttctgggccg

cacgcgcgctacactgatgcattcaacgagtttcacttgatagtcctgggtcggaaggct

cgggtaatcttttgaaagtgcatcgtgctggggatagatcattgcaattattgatcttca

acgaggaattccttgtaggcgcaggtcatcagcctgcgccgaatacgtccctgccctttg

tacacaccgcccgtcgctcctaccgattgaatggtccgatgaaatgtccggatcgcggct

agtgtgatggactcgtccagagcggtagccgtgagaagctcattaaatcttatcatttag

aggaaggagaagtcgtaacaaggt------------------------------------

------------------------------------------------------------

------------------------------------------------------------

------------------------------------------------------------

------------------------------------------------------------

------------------------------------------------------------
------------------------------------------------------------
------------------------------------------------------------
------------------------------------------------------------
------------------------------------------------------------
------------------------------------------------------------
------------------------------------------------------------
------------------------------------------------------------
------------------------------------------------------------
------------------------------------------------------------
------------------------------------------------------------
-------------------------------ttccgtaggtgaacctgcggaaggatcat
tcacaaactattgacacaaacacacgcgaaccaaatcgtccgcacagcaggtggcaatga
aagagagactctgcatgtcgcctttcgggg--tagcaagcagcactcttctgccatcgcc
tctgtgcagggcgtaaattcccattgagaggatgtgagggcaccacaggaagcttttcca
caggaagtcgccatccttctccctccacggcggcg----cttctgtgcttggcag-----
----tttttttttgccttctagggaggatgccgccaatggagccccttat-atatataca
tcatcatatgccc--ctttttttcttaaccgcttgccacagcttcttctatgaggagctt
gtgggaagactgtctccatacaataac-aaagatacaactcttagcggtggatatcttgg
ttctcgcaacgatgaagaacgcagccaactgcgataactaatgtgaattgcaggacttcg
tgaatcattaagtctttgaacgcaagttgcgctcatctccatgaggaggtgagcatgcct
gtttgagtgtcgatgggaaaaacaatac-ataacaacaatcggctctcttatcttcctgc
cttcttgggggcaaaaggtagtgcatgccggagatgggctttgtcgcccctttctcttga
aagaggcgtcgtccgaaacacgtatcgttgcccgctggaggtcatagagcatgagttt--
aaagcaccacatctctcaagtcagagtgag-cgtcttgagtaggtcc-gcattgattcag
tttcaacccaggtggcgttggttacttgccat-aagcctctctggggtttctttgcttgg
acctctcttgtcagtttctctgacgtagagacaggt--gccgtctcatggacgttgccct
ctggaacgtgcta-aaaaaaaaaaaaaga----------aaaaaccgtaccacatgtcaa
aagaggcagatcgtgcttgacggcgggaggtgttgcatttcttgcttgtcctgagcgtct
gccagtctct----ga--cgagtg----tgttccttccataaacccgatctcaaatcagg
taggaccacccgctgaatttaagcatataattaagcggaggaaaagaaactaacaaggat
tccctcagtagcggcgagcgaagcgggaagagctcaacatgaaaaccgttgggggttcgc
gctttccttcgggttgtatagtaaagtggtgtcgcctgcgtcgcgccgggtataaatccc
gtctgaaatggggtgtcgtagagggttatatccccgtctgtgacccggtgtgcgatgcgt
tacggtgaccttacgatgagtcgggttgcttgggcatgcagccctaagtgggtggtaaac
ttcatctaaagctaaatactagcgagagaccgatagcgtacaagtaccgtaaggaaaga

tgcaaagaactttgaaaagagagttaaacagggcttgaaaccgttaagagtgaagcgaag
gaggtcagtgatcatggggccagatccccaagtgcctgtcata-gggcaccgtagcgttc
tggtttcccggggcagcgccggccgggggcgggggttaacagggaggtggttgagggatt
cgttttccgcctgtactcctttatagcttgcaccccggccgagggatccgcgacttgtcg
tgtgagggcgctgtcaaaatggccttctccaccccgtcttgaaacacggaccaaggagtc
taacatactcgcgagtgtttgggtggttaaaacccgaacgcacaatgaaagtaaacggtg
gctctgcgccgctgcttggtgggaaccctgcttttataatgggggcactatcacccgac
cccgatccctttggggtggttggttttgaggtagagcgagtatgatgggacccgaaagat
ggtgaactattcctgagtagggcgaagccagaggaaactctggtggaggctcgtagcgat
gctgacgtgcaaatcgctcgtcgaacttgggtataggggcgaaagactaatcgaaccatc
tagtagctggttccctccgaagtttccctcaggatagcgggagcaaatgcaagtcagtat
catccggtaaagcgaatgattagaggcctcgggggagtaactccttcgacctattctcaa
actttaaataggtgaggagccggcaacgttgcttaaaacaatgaacgccgtgcggacaat
gattgctctcagtgggccgattttggtaagcagaactggcgatgcgggatgaaccgaaag
cgacgctaaggtgccaaactcccagctcattcagacaccagaaaaggtgttgattcatac
agacagcaggacggtggccatggaagtcggaatccgctaaggagtgtgtaacaactcacc
tgccgaatggatcagccctgaaaatggatggcgcttaagcgggggaccgaggcgtcgccg
ttggggtaaatacgaagccccaatgagtaggagggcgtgggggttttgaaacacggtaca
ctcgcatggaagccttgggcgtaagctcgggtggagcgacccctagtgtagatcttggtg
gtagtagcaaatattcaagtgagaaccttgaagactgaagtggggaagggttccacgcga
acagtgcttggacgtgggttagtcggtcctaagagatggggaaactccgtattcaagtgc
gcttttgtagcgcagtccatcgaaagggaagcgggttaacattcccgcaccaggatgtgg
aaactgtggtttaccgcaaactgcggcaacgcaactgaacctggggacgcccgtaggagc
tccgagcagggatctctttcctgtttaacgcgcccaaccccctggaatcggtttgtccgg
agatagggggcgaaaggccggaaaagcaccgcacgtcttgtggtgttcggaacgcttgta
atggcccttgaaaacccaggggaaagcagccgtcttacac--ttgtgtgagacgaggaat
tctcacacctggccgtacccatcaccgcatcaggtccccaaggtgaacagcctctagtca
atagaataatgtagataagggaagtcggcaaaatagatccgtaactttgggaaaaggatt
ggctctaatgttcgggcttgtcgggacgcgcgggcgtagatgccaatgagcggcagacgt
ctagcctctttggaggctggattaacgctgcttgtgggaagaaaccgtgtcgatggaggc
aggaacgcttatcgatcttgtctcttcccgcttgcgatcaacgaacggattagaactggt
acggacaaggggaatccgactgtttaattaaaacaaagcattgcgacggccggaaacggt
gttggcgcaatgtgatttctgcccagtgctctgaatgtcaaagtgaagaaattcaaccaa
gcgcgggtaaacggcgggagtaactatgactctcttagac-agatcaagacaccttcaat
aaattggttcattatgtcgggtggtccgtcgtccgggaagcgaccaaagcaaaaccccgt
agactgtgatgtgcatctgtcagacgaaactgacgtccgtgttcatcaagccttagctat

ctttcaacgccggcttcgagacgtggcgtctcttaaggtgtttgaagtggctctctctgt

gggaactactgatgcactggtagagatgggcactatcacttgctggagtattgatgctgt

tgggaaaggtttctcaaggaacagcagcggtcgctggcaatttgacctcactggcaagat

taatggcaagcctctaatatccaccacaaaggatcgttggaggccggttgtgacggctgt

gccctactacctttcgtgcgccatcgatggggacaagacgacgctggaggaaaagcagtg

gacttcatcacacatgtgtcaccatgaatggtgtgtgaacgtggagcacattgtttatga

aagcctcgacgtcaac--aagtctcggaactggtg--ccagggaaaaaacgcatgcctcc

atctgcctgtttgcattcgaaccggccgccgtgtgcaggagctcaggaagagcaatgggg

ttcgtgcaatggacctcctcctgcagaaattggctggattggagattgaggttgaggagc

ccgctgcagctgtactccgagagtctgtcgaacgggaggacgcctagtgagtgactcgtg

acacgggttggctgcgcatgatgcgtcaactacatggacatcttttgagagagataggcg

ggctgaaaaggttatagccgtctagtctcgctctctttcattgagcggggcgacacacct

ggttgcgggaaccctgcccacctttctatcaagaagtgggaaccagtccattcctactcc

tgcttgtgaggcatatagt--aacgtagcagcaggaggatctcggcgagaggccgaaccc

gaaaggaataacgtgggctgggaggcgatccgcaggcgagtccgtaactatatgattatc

gacgggccgtctcaacgactggtaaggtgtgggtttg-tcctcctcctcacgtggggaca

ggcttaaggtacagtctaatcccacccgccaagggtgcttggg---tg------------

--------------------taatccctcttccatagagggcactagccca-ggggctgg

gacgctgggaggaa----------------atgccccgtccctatagccggtagaaatga

aggtagccaaatgcctcgtcatct------------------------------------

------------------------------------------------------------

------------------------------------------------------------

------------------------------------------------------------

------------------------------------------------------------

------------------------------------------------------------

------------------------------------------------------------

------------------------------------------------------------

------------------------------------------------------------

---aattagtgacgcgcatgaatggattaacgagattcccactgtccctatctactatct

agcgaaaccacagccaagggaacgggcttggaagaatcagcggggaaagaagaccctgtt

gagcttgactctagtccgactttgtgagatgagttgggaggtgtagtctaagagggaggg

gcaactcgaccgtgaaatacctctactcccaagcttgttttacttactccggcgagcagg

gatgggtggtaacacccttttctaggattaacacctcttcgggggtggacctgcgcgga

ggacattgtcaggtggggagtttggctggggcggcacatctgttaaaagataacgcaggt

gtcctaagatgagctcagtgagaacagaaatctcacgtagaacaaaagggtaaaagctca

tttgattttgattttcagtacgaatacaaactgtgaaagcatggcctatcgatcctttg

accttcaaggtttgaagcaagaggtgtcagaaaagttaacacgcgttgtggtagctttta
agagtggactcaaagtgttccgctagtcccccctcggaccgccacttacaaaaggcctgg
ggatgggggcgacactgtcaaattgcggggactccctaaagcctgtctctaccaagccag
cgcggcgaaagcccctggtggccggggtaatgacctagggtatggtaaaagcgagacagg
atg-gaacaatgggcaatccgcagccaagtcctaagg----ctttttgaaaagcggctat
ggatgcagttcaacgactagatggcagtgggctgagagagcccaccgcggcacccctttcg
gcttaagatatagtcttgcccacgtgaaagcgtgcctgcagatgaacgccaccatgtatg
gccggagctgcaggagaccgc--acttgatgtgcagtcgggaat-cgacagtaccacagg
gatgaac-----------------------------------------------------
------------------------------------------------------------
------------------------------------------------------------
------------------------------------------------------------
------------------------------------------------------------
-------------------------------------------------aaaaacac---
------------------------------------------------------------
------------------------------------------------------------
--acaatta---------------------------------------------------
----------------------------------------atttgtggcc---atcatac
----------------------------------------agtcgta-------------
------------------------------------------------------------
------------------------------------------------------------
------------------------------------------------------------
------------tgatattatcgtcccaggcgggtgcgcctctctcacccgtctagtccg
gccttggtgcgcaaaaaatgcttgcgtggatgcgccttggttaaggaaaatgctcctagt
agttttacaaggacacttcttgtagggcaacacatctggatgcgaagaactcccggaaaa
gacaacggtaccaagtgactggggtaacgccccattcatgaaggcaaataatcaaaaaga
ggtgccgggtaagaacccgttgtatagggacaatccgcagctgacttggctccctg--cc
cat--ccatgggcatagcttagcggttcaacgctcggtaagatgtgggttggcgcgctgt
ccgacagcgcactggcttaaggtacgggctactcccaccggagacggtgtctttgtctac
ctcgggtaacacgcctcgggagaagcaaagagggtgcaatggaaaaaaaactggttgcgg
ggaaatccctaccagagtcacctggtacaaaggaactggcttgtggcagccaagcgttca
tagcgacgttgctttttgatccttcgatgtcggctcttcctatcattgcgaagcagaatt
cgccaagtgttggattgttcacccactaatagggaacgtgagctgggtagccacttta--
-----tgcccaacagctggatagctgggcttctgtctta-ggccgggacttctcacgacc
cgtgccttggtcaccc-aaaaaagggcgaccgcagagctcaggatggagcttctggctta
actagtagaggacagatagagatcgtcaaattgcgggaaacccccttagagcctcgagcta

ccaagtgcgtgtcgaaaggcatagacgcagccagcgatggctgtcgcatggcttga-ttc
catcaagataggtaaaaatgctcgaggattgggcaatccgcagccaagcccctaaggt--
---ccccccgcgggctatggggaaggttcaacgactgtaaggcgtaccgtgcaagcggaa
gatacagtctagccccgctggaaagcagcgccaaagtaagtaggt-tgcagtgggtctgt
cctc
>Form_G_China
gcatgttttggtatgtgtttcgacagtcatgcatgtcatagccgtaggatggggagtgag
cttcatagcccgaccccttctcaggagcatgttttcatagcatattgagcatactggggt
ggtctcccatgtagctcgctctctcggtgcaggacctcccttttgcgggccacttcgatg
gcctccattatgga-gagcatgttttggcatatgaagcgttggtggtgaggctcgtagcc
tgtatatacataccaacgcagtcgtcatatgtctctctggcgggggtcggtcgtatgacc
ttgtccctacctggagcatgtatttcgtatgat---atactggac-ggcagctgcgctgt
tcagtcgtcggccgtacctcatgtgcgcgtttccaggtccgaacgtttgacatgcaaggg
catcataccagaagcctgttttctcgtatgagcatgtaggtaggcaggctcttcagtcgt
ctacccataaatgcacttcgaagga-actttttggtattgaatggctttgtgcccagacg
ttcttgtgagcttgttctcgctctgcgatacgctggaagctgtggtattacctacagcag
acgcattcctagggcggacggcctctctgcagatgggcacatctgtagaaagaggctccc
cttgatgacgcaatgcaatggctgggtaaacggagtgctatacagaagggtcaaacctc-
tgtggaagtacactcataccagcattgtttgttgagaggggtcggctctacgagtgagga
cctatactgccatctcattcagatgcgtagacggtctttactcagactgcctttctggct
gattgtcctcggtccctcgtgcgcacgcgccgctggctgaatgggcttgtccttatggag
tgccgcgttcttc-aaagaatccgtcttcatacagcgacctcatttacatattgccggct
gaatgggcttgtccttatggagcactgcattcttctaggaattgctatctatacagcgac
ctcatttacatacaaggcacct-ttttctacatagacttgccatttggctgattgttctg
acagtcgctgcctcttcggagttctgtgtggacgagtggcc--tcctgagtgggccggtc
cttacggagtgcccacgtctgttatgactgggttctccatacagcgacctcactcagaat
tgtatgg--------------caccgtgccgat-----------tgaatggacaagcagt
ccttagggaatcgcatgaaagcatgatggc-acatggtgtatggacccgttgggccttgc
ctgtgctatcttctcgagtcgaccgg------------tcctcgtctcctcctttcgcga
cgggaagctcttgagtttaccgtgggtcaggacgctgtggtatctgtttgtctgtagtgt
gggctctctatcagaacagcgtatcagcacacgtcgtctgggtctatggtatttttgctt
aacggccctgggtttggtagtatggttccatgtcggtcctctcacctcccatccgtcatg
tctctga-----------------------------------------------------
------------------------------------------------------------
--------------------------------------------tatcatctgcacagta
cgtcttgtcagaatatcattctgtatggcagtcgatagactaggccctatgccgcagctg

tctctgaccactatgcgagcgcatacctacctcaatgtgttcatagctgttatggatctt
gggtgtggtgtgccagaacgatacgccaccagattgtggaatggtgcttgtagtattaga
agaggacatgtatgaagagcacgttcttttctcttatgatggtgccgcgggtgggggccg
ttctcattctgcttgtcgggtgctcaaggtgcatgtaggattgctgaacccac-tgaatg
gtcatgagaatggtcatatggtgaggttcacttctccatcgctttgtgtgtgttggatgt
ttgggtggcttcggccgcatggaaaatctcggtctgtaactcatcagggagggtttccat
gtgagcaatctctccagtcgaacacacacccatgggcgtcccgctaggcaaaatggaact
ctcgatccgacacagcctgtggagtgtctgttccgtgtggcgtgaaacggcttatactgg
atgtcttccc-tttggacaccagtctgttggtatg--tacgcataccatctagagggagc
aagaaaaaggcatgcaagtgccacaaaggaagggaggcctccctgctaccgtgtgtcc-t
cttgacgatcacgggggagtatatagagacatttggacgagatcctttttttcacgaggcg
gtgccctcgcgcaaaggtgttgctctgcacgcatctttatgatgatagttgctcgctcct
atgcgtccactcgtgaaaatacctgctctctctgtctgtttccatgacacccc-catgcg
ttgtcatagagcacgctcaggtacagaaagtgctgtgactttcttggcctgctgtcactg
agtcagcctgatactttggaggaatcttttctgggcgctgcccagctgagagttcttatc
aggctctggagtgtgcatatggctataatcataatagtgcatgtggccacgcagaaaaag
catcgacgtggaatggagaagaacaccattctgaatggaacggttcttactctgcttccg
ttgcttgacgctgtatgggaatcgctttgccctgggggtacaccttccattccatgacaa
agcatttgatctatgccccga-gggtcattctctatgaagagcccgaaacttgtctgagt
gagaggcgaaggtagcccttcattt--atgcggggtccgccctcacactgtagtcggcac
agtcccttgtttatggactgcgcagctcaatcttcgactggagagg--aaaaacaaaaca
caagtatgacacgcacatacgtgttttctaggactgtgacctgtcggctgcctcgcggtg
gttggcatgatgccccaagtgggggggggagaaacaatcagtatgattgataaaaacca
gtgggg-aaaaaggcgcgctgtcgcacacccttggtcacggcctc---aagtcgacggcc
ctcactggccca-ccctgcgactcctgttcgcttttgacaagacacacctgcaata-cgg
aggaaatgcctgagcagctgtgccgcagcaactgcggacgcgtagcgatatgcatccctg
ttgtttgcgacttggtacaaagaacgtgcgccgggagacggattgccctttgacttttcc
acccagcccgcggctctatacccttgacgtacctatttacctggttgatcctgccagtag
tcatatgcttgtctcaaagactaagccatgcatgtctaagtataaacacttttatactcg
tgaaactgcgaatggctcattaaaacagttatagttcctttgggaacaacactacttgga
taaccgtagtaattctagagctaatacatgcctcaacgcccgactcacgaagggtggtat
ttattggataaaaaaccatcgtgctctgtccagagcgctttgagatgattcacaataact
tgtcggatcgcatggcctcgtgctggcgactcctcattcaaatttctgccctatcaactt
tcgatggtagagtattggtctaccatggtgtcgacgggtgacggggaattagggttcgat
tccggagagggagcctgagaaacggctaccacatccaaggaaggcagcaggcgcgcaaat
tacccaatcccgactcggggaggtagtgacaaaaaataacaatagggggccctttgggtc

ttctaattggaatgagaacaatttaaatcccttatcgaggatccattggagggcaagtct
ggtgaacgta----------------------------------tgcatca---------
--aagccctccgtaat-aaaaaaaaacagcgccagatgtct-------------------
------------------------------------------------------------
------------------------------------------------------------
------------------------------------------------------------
------------------------------------------------------------
------------------------------------------------------------
-------ctctctggaagcacacca-----------------------------------
---cgcacggggttcgttcctctc-----------------------------tgcatgg
ccgtg-----------------------------------------tgaaaaaggcgagt
aggcaaacgatgggaaacccggcttggaatagtaaccagccacagagcccccatcataca
cttctaaattgccggggagtgcccgcgaggccttgactaccgcggtccgccgagaggtga
gagcgagcaatcgctcaggacgcagcagtgtacgtagtggtacatggatggtaaaaacgt
caaggatagggttgatccgcagggaagcgcgagtgccgccatg---------gtcggtgg
gcaaaggcgaaccttcagagactctaatggagtgggcgcaagcttaagggagagtccaat
ccactgggaaaccaggtccacagcagtaacggggtgatgctcgccccagaggactgtgga
aggcaaccccacc----aaaagggtgtgtgttgtcgggtagc-aatggccagcagccgcg
gtaattccagctccaatagcgtatattaaagttgttgcagttaaaacgctcgtagtcgga
cctcggagcgcccccacgggccgggcttttgccttgttgccagtgggagtttgtctcctt
ttgtcgggacgcggtgtgcagggcctttactgtctctgcgcgtttacctgcgggccgacc
gtttactgtgaagaaagtaaagtgttcaaagcaggccaatgccttgaatatgtgagcatg
gaataatagaataggacttgggctctattttgttggtttccagtgaccaagtaatgatta
atagggacggttgggggcattcgtatttcattgtcagaggtgaaattcttggattgatgg
aagacgtacaactgcgaaagcatctgccatggatgttttcattgatcaagaacgaaagtt
aggggatcgaagacgatcagataccgtcgtagtcttaaccataaacgatgccgactgggg
attggcgggggcacttttatatgactctgtcagcaccctgagggaaaccaaagtctttgg
gttctgggggggagtatggtcgcaaggctgaaacttaaaggaattgacggaagggcaccac
caggagtggagcctgcggcttaatttgactcaacacgggaaaacttaccaggtccggaca
gaagaatgattgacagactgaaagctctttcttgatttttttggttggtggtgcatggccg
ttcttagttggtggagtgatttgtctggttaattccgttaacgaacgagacctcgtcctg
ctaaataggtgcgcgcatgccagcactgcgttcttaccttcttagagggactatgcgcgt
ctagcgtatggaagattgaggcaataacaggtctgtgatgcccttagatgttctgggccg
cacgcgcgctacactgatgcattcaacgagtttcacttgatagtcctgggtcggaaggct
cgggtaatcttttgaaagtgcatcgtgctggggatagatcattgcaattattgatcttca
acgaggaattccttgtaggcgcaggtcatcagcctgcgccgaatacgtccctgccctttg

tacacaccgcccgtcgctcctaccgattgaatggtccgatgaaatgtccggatcgcggct
agtgtgatggactcgtccagagcggtagccgtgagaagctcattaaatcttatcatttag
aggaaggagaagtcgtaacaaggt------------------------------------
------------------------------------------------------------
------------------------------------------------------------
------------------------------------------------------------
------------------------------------------------------------
------------------------------------------------------------
------------------------------------------------------------
------------------------------------------------------------
------------------------------------------------------------
------------------------------------------------------------
------------------------------------------------------------
------------------------------------------------------------
------------------------------------------------------------
------------------------------------------------------------
------------------------------------------------------------
------------------------------------------------------------
-------------------------------ttccgtaggtgaacctgcggaaggatcat
tcacaaactattgacacaaacacacgcgaaccaaatcgtccgcacagcaggtggcaatga
aagagagactctgcatgtcgcctttcgggg--tagcaagcagcactcttctgccatcgcc
tctgtgcagggcgtaaattcccattgagaggatgtgagggcaccacaggaagcttttcca
caggaagtcgccatccttctccctccacggcggcg----cttctgtgcttggcag-----
-----ttttttttgccttctagggaggatgccgccaatggagccccttat-atatataca
tcatcatatgccc--ctttttttcttaaccgcttgccacagcttcttctatgaggagctt
gtgggaagactgtctccatacaataac-aaagatacaactcttagcggtggatatcttgg
ttctcgcaacgatgaagaacgcagccaactgcgataactaatgtgaattgcaggacttcg
tgaatcattaagtctttgaacgcaagttgcgctcatctccatgaggaggtgagcatgcct
gtttgagtgtcgatgggaaaaacaatac-ataacaacaatcggctctcttatcttcctgc
cttcttgggggcaaaaggtagtgcatgccggagatgggctttgtcgcccctttctcttga
aagaggcgtcgtccgaaacacgtatcgttgcccgctggaggtcatagagcatgagttt--
aaagcaccacatctctcaagtcagagtgag-cgtcttgagtaggtcc-gcattgattcag
tttcaacccaggtggcgttggttacttgccat-aagcctctctggggtttctttgcttgg
acctctcttgtcagtttctctgacgtagagacaggt--gccgtctcatggacgttgccct
ctggaacgtgcta-aaaaaaaaaaaaaga----------aaaaaccgtaccacatgtcaa
aagaggcagatcgtgcttgacggcgggaggtgttgcatttcttgcttgtcctgagcgtct

gccagtctct----ga--cgagtg----tgttccttccataaacccgatctcaaatcagg
taggaccacccgctgaatttaagcatataattaagcggaggaaaagaaactaacaaggat
tccctcagtagcggcgagcgaagcgggaagagctcaacatgaaaaccgttgggggttcgc
gctttccttcgggttgtatagtaaagtggtgtcgcctgcgtcgcgccgggtataaatccc
gtctgaaatggggtgtcgtagagggttatatccccgtctgtgacccggtgtgcgatgcgt
tacggtgaccttacgatgagtcgggttgcttgggcatgcagccctaagtgggtggtaaac
ttcatctaaagctaaatactagcgagagaccgatagcgtacaagtaccgtaagggaaaga
tgcaaagaactttgaaaagagagttaaacagggcttgaaaccgttaagagtgaagcgaag
gaggtcagtgatcatggggccagatccccaagtgcctgtcata-gggcaccgtagcgttc
tggtttcccggggcagcgccggccgggggcgggggttaacagggaggtggttgagggatt
cgttttccgcctgtactcctttatagcttgcaccccggccgagggatccgcgacttgtcg
tgtgagggcgctgtcaaaatggccttctccaccccgtcttgaaacacggaccaaggagtc
taacatactcgcgagtgtttgggtggttaaaacccgaacgcacaatgaaagtaaacggtg
gctctgcgccgctgcttggtgggaaccctgcttttataatggggggcactatcacccgac
cccgatccctttggggtggttggttttgaggtagagcgagtatgatgggacccgaaagat
ggtgaactattcctgagtagggcgaagccagaggaaactctggtggaggctcgtagcgat
gctgacgtgcaaatcgctcgtcgaacttgggtataggggcgaaagactaatcgaaccatc
tagtagctggttccctccgaagtttccctcaggatagcgggagcaaatgcaagtcagtat
catccggtaaagcgaatgattagaggcctcgggggagtaactccttcgacctattctcaa
actttaaataggtgaggagccggcaacgttgcttaaaacaatgaacgccgtgcggacaat
gattgctctcagtgggccgattttggtaagcagaactggcgatgcgggatgaaccgaaag
cgacgctaaggtgccaaactcccagctcattcagacaccagaaaaggtgttgattcatac
agacagcaggacggtggccatggaagtcggaatccgctaaggagtgtgtaacaactcacc
tgccgaatggatcagccctgaaaatggatggcgcttaagcgggggaccgaggcgtcgccg
ttggggtaaatacgaagccccaatgagtaggagggcgtgggggttttgaaacacggtaca
ctcgcatggaagccttgggcgtaagctcgggtggagcgacccctagtgtagatcttggtg
gtagtagcaaatattcaagtgagaaccttgaagactgaagtggggaagggttccacgcga
acagtgcttggacgtgggttagtcggtcctaagagatggggaaactccgtattcaagtgc
gcttttgtagcgcagtccatcgaaagggaagcgggttaacattcccgcaccaggatgtgg
aaactgtggtttaccgcaaactgcggcaacgcaactgaacctggggacgcccgtaggagc
tccgagcagggatctctttcctgtttaacgcgcccaaccccctggaatcggtttgtccgg
agatagggggcgaaaggccggaaaagcaccgcacgtcttgtggtgttcggaacgcttgta
atggcccttgaaaacccaggggaaagcagccgtcttacac--ttgtgtgagacgaggaat
tctcacacctggccgtacccatcaccgcatcaggtccccaaggtgaacagcctctagtca
atagaataatgtagataagggaagtcggcaaaatagatccgtaactttgggaaaaggatt
ggctctaatgttcgggcttgtcgggacgcgcgggcgtagatgccaatgagcggcagacgt

ctagcctctttggaggctggattaacgctgcttgtgggaagaaaccgtgtcgatggaggc
aggaacgcttatcgatcttgtctcttcccgcttgcgatcaacgaacggattagaactggt
acggacaaggggaatccgactgtttaattaaaacaaagcattgcgacggccggaaacggt
gttggcgcaatgtgatttctgcccagtgctctgaatgtcaaagtgaagaaattcaaccaa
gcgcgggtaaacggcgggagtaactatgactctcttagac-agatcaagacaccttcaat
aaattggttcattatgtcgggtggtccgtcgtccgggaagcgaccaaagcaaaaccccgt
agactgtgatgtgcatctgtcagacgaaactgacgtccgtgttcatcaagccttagctat
ctttcaacgccggcttcgagacgtggcgtctcttaaggtgtttgaagtggctctctctgt
gggaactactgatgcactggtagagatgggcactatcacttgctggagtattgatgctgt
tgggaaaggtttctcaaggaacagcagcggtcgctggcaatttgacctcactggcaagat
taatggcaagcctctaatatccaccacaaaggatcgttggaggccggttgtgacggctgt
gccctactacctttcgtgcgccatcgatggggacaagacgacgctggaggaaaagcagtg
gacttcatcacacatgtgtcaccatgaatggtgtgtgaacgtggagcacattgtttatga
aagcctcgacgtcaac--aagtctcggaactggtg--ccagggaaaaaacgcatgcctcc
atctgcctgtttgcattcgaaccggccgccgtgtgcaggagctcaggaagagcaatgggg
ttcgtgcaatggacctcctcctgcagaaattggctggattggagattgaggttgaggagc
ccgctgcagctgtactccgagagtctgtcgaacgggaggacgcctagtgagtgactcgtg
acacgggttggctgcgcatgatgcgtcaactacatggacatcttttgagagagataggcg
ggctgaaaaggttatagccgtctagtctcgctctctttcattgagcggggcgacacacct
ggttgcgggaaccctgcccacctttctatcaagaagtgggaaccagtccattcctactcc
tgcttgtgaggcatatagt--aacgtagcagcaggaggatctcggcgagaggccgaaccc
gaaaggaataacgtgggctgggaggcgatccgcaggcgagtccgtaactatatgattatc
gacgggccgtctcaacgactggtaaggtgtgggtttg-tcctcctcctcacgtggggaca
ggcttaaggtacagtctaatcccacccgccaagggtgcttggg---tg------------
--------------------taatccctcttccatagagggcactagccca-ggggctgg
gacgctgggaggaa----------------atgccccgtccctatagccggtagaaatga
aggtagccaaatgcctcgtcatctaaactaattgg-at----------------------
-----------------------gagatagtc----------tgagtgaaaagcttatgc
ag-----------------gctagtaccgg---aaaaaaaaaaaaagaggggctctgcca
ct----ctgac------------------------ggtgcgacactccttgtacagggaa
accctcc--gcatggaaaggcacgtggtaccaccctcgcggggaaactcgcgagcgggat
cccgggtaatgaccggagccgatggtaaaaactcacgtgaccgatggggtgaccctgtgg
cgagacgctaagttgtgggacatcaccctctcacaatacgcatccgtcgcagagactgaa
cgggagtgggctccgttga-----cggccggccggccagccagccgtcttcca-------
---ggggcttaagatacagtccgggcctcagggatgg----aaaaacctggggaacta-c
cagaattagtgacgcgcatgaatggattaacgagattcccactgtccctatctactatct

agcgaaaccacagccaagggaacgggcttggaagaatcagcggggaaagaagaccctgtt
gagcttgactctagtccgactttgtgagatgagttgggaggtgtagtctaagagggaggg
gcaactcgaccgtgaaatacctctactcccaagcttgttttacttactccggcgagcagg
gatgggtggtaacacccttttctaggattaacacctcttcgggggtggacctgcgcgga
ggacattgtcaggtggggagtttggctggggcggcacatctgttaaaagataacgcaggt
gtcctaagatgagctcagtgagaacagaaatctcacgtagaacaaaagggtaaaagctca
tttgattttgattttcagtacgaatacaaactgtgaaagcatggcctatcgatcctttg
accttcaaggtttgaagcaagaggtgtcagaaaagttaacacgcgttgtggtagctttta
agagtggactcaaagtgttccgctagtccccccctcggaccgccacttacaaaaggcctgg
ggatgggggcgacactgtcaaattgcggggactccctaaagcctgtctctaccaagccag
cgcggcgaaagcccctggtggccggggtaatgacctagggtatggtaaaagcgagacagg
atg-gaacaatgggcaatccgcagccaagtcctaagg----ctttttgaaaagcggctat
ggatgcagttcaacgactagatggcagtgggctgagagagcccaccgcggcaccctttcg
gcttaagatatagtcttgcccacgtgaaagcgtgcctgcagatgaacgccaccatgtatg
gccggagctgcaggagaccgc--acttgatgtgcagtcgggaat-cgacagtaccacagg
gatgaac-----------------------------------------------------
------------------------------------------------------------
------------------------------------------------------------
------------------------------------------------------------
------------------------------------------------------------
-------------------------------------------------aaaaacac---
------------------------------------------------------------
------------------------------------------------------------
--acaatta---------------------------------------------------
----------------------------------------atttgtggcc---atcatac
----------------------------------------agtcgta-------------
------------------------------------------------------------
------------------------------------------------------------
------------------------------------------------------------
------------tgatattatcgtcccaggcgggtgcgcctctctcacccgtctagtccg
gccttggtgcgcaaaaaatgcttgcgtggatgcgccttggttaaggaaaatgctcctagt
agttttacaaggacacttcttgtagggcaacacatctggatgcgaagaactcccggaaaa
gacaacggtaccaagtgactggggtaacgccccattcatgaaggcaaataatcaaaaaga
ggtgccgggtaagaacccgttgtatagggacaatccgcagctgacttggctccctg--cc
cat--ccatgggcatagcttagcggttcaacgctcggtaagatgtgggttggcgcgctgt
ccgacagcgcactggcttaaggtacgggctactcccaccggagacggtgtctttgtctac

ctcgggtaacacgcctcgggagaagcaaagagggtgcaatggaaaaaaaactggttgcgg
ggaaatccctaccagagtcacctggtacaaaggaactggcttgtggcagccaagcgttca
tagcgacgttgctttttgatccttcgatgtcggctcttcctatcattgcgaagcagaatt
cgccaagtgttggattgttcacccactaatagggaacgtgagctgggtagccacttta--
-----tgcccaacagctggatagctgggcttctgtctta-ggccgggacttctcacgacc
cgtgccttggtcaccc-aaaaaagggcgaccgcagagctcaggatggagcttctggctta
actagtagaggacagatagagatcgtcaaattgcgggaaaccccttagagcctcgagcta
ccaagtgcgtgtcgaaaggcatagacgcagccagcgatggctgtcgcatggcttga-ttc
catcaagataggtaaaaatgctcgaggattgggcaatccgcagccaagcccctaaggt--
---ccccccgcgggctatggggaaggttcaacgactgtaaggcgtaccgtgcaagcggaa
gatacagtctagccccgctggaaagcagcgccaaagtaagtaggt-tgcagtgggtctgt
cctc
>Form_K_China
gcatgttttggtatgtgtttcgacagtcatgcatgtcatagccgtaggatggggagtgag
cttcatagcccgaccccttctcaggagcatgttttcatagcatattgagcatactggggt
ggtctcccatgtagctcgctctctcggtgcaggacctcccttttgcgggccacttcgatg
gcctccattatgga-gagcatgttttggcatatgaagcgttggtggtgaggctcgtagcc
tgtatatacataccaacgcagtcgtcatatgtctctctggcgggggtcggtcgtatgacc
ttgtccctacctggagcatgtattttgtatgat---atactggac-ggcagctgcgctgt
tcagtcgtcagccgtacctcatgtgcgcgtttccaggtccgaacgtttgacatgcaaggg
catcataccagaagcctgttttctcgtatgagcatgtaggtaggcaggctcttcagtcgt
ctacccataaatgcacttcgaagga-actttttggtattgaatggctttgtgcccagacg
ttcttgtgagcttgttctcgctctgcgatacgctggaagctgtggtattacctacagcag
acgcattcctagggcggacggcctctctgcagatgggcacatctgtagaaagaggctccc
cttgatgacgcaatgcaatggctgggtaaacggagtgctatacagaagggtcaaacctc-
tgtggaagtacactcataccagcattgtttgttgagaggggtcggctctacgagtgagga
cctatactgccatctcattcagatgcgtagacggtctttactcagactgcctttctggct
gattgtcctcggtccctcgtgcgcacgcgccgctggctgaatgggcttgtccttatggag
tgccgcgttcttc-aaagaatccgtcttcatacagcgacctcatttacatattgccggct
gaatgggcttgtccttatggagcactgcattcttctaggaattgctatctatacagcgac
ctcatttacatacaaggcacct-ttttctacatagacttgccatttggctgattgttctg
acagtcgctgcctcttcggagttctgtgtggacgagtggcc--tcctgagtgggccggtc
cttacggagtgcccacgtctgttatgactgggttctccatacagcgacctcactcagaat
tgtatgg--------------caccgtgccgat-----------tgaatggacaagcagt
ccttagggaatcgcatgaaagcatgacggc-acatggtgtatggacccgttgggccttgc
ctgtgctatcttctcgagtcgaccgg------------tcctcgtctcctcctttcgcga

cgggaagctcttgagtttaccgtgggtcaggacgctgtggtatctgtttgtctgtagtgt
gggctctctatcagaacagcgtatcagcacacgtcgtctgggtctatggtatttttgctt
aacggccctgggtttggtagtatggttccatgtcggtcctctcacctcccaaccgtcatg
tctctga-----------------------------------------------------
------------------------------------------------------------
--------------------------------------------tatcatctgcacagta
cgtcttgtcagaatatcattctgtatggcagtcgatagactaggccctatgccgcagctg
tctctgaccactatgcgagcgcatacctacctcaatgtgttcatagctgttatggatctt
gggtgtggtgtgccagaacgatacgccaccagattgtggaatggtgcttgtagtattaga
agaggacatgtatgaagagcacgttcttttctcttatgatggtgccgcgggtgcgggccg
ttctcattctgcttgtcgggtgctcaaggtgcatgtaggattgctgaacccac-tgaatg
gtcatgagaatggtcatatggtgaggttcacttctccatcgctttgtgtgtgttggatgt
ttgggtggcttcggccgcatggaaaatctcggtctgtaactcatcagggagggtttccat
gtgagcaatctctccagtcgaacacacacccatgggcgtcccgctaggcaaaatggaact
ctcgatccgacacagcctgtggagtgtttgttccgtgtggcgtgaaacggcttatactgg
atgtcttccc-tttggacaccagtctgttggtatg--tacgcataccatctagagggagc
aagaaaaaggcatgcaagtgccacaaaggaagggaggccttcctgctaccgtgtgtcc-t
cttgacgatcacgggggagtatatagagacatttggacgagatccgtttttcacgaggcg
gtgccctcgcgcaaaggtgttgctctgcacgcatctttatgatgatagttgctcgctcct
atgcgtccactcgtgaaaatacctgctctctctgtctgtttccatgacacccc-catgcg
ttgtcatagagcatgctcaggtacagaaagtgctgtgactttcttggcctgctgtcactg
agtcagcctgatactttggaggaatcttttctgggcactgcccagctgagagttcttatc
aggctctggagtgtgcatatgcctatattcattatagtgcatgtggctacgcagaaaaag
catcgacgtggaatggagaagaacaccattctgaatggaacggttcttactctacttctg
ttgcttgacgctgtatgggaatcgctttgctctgggggtacaccttccattccatgacaa
agcatttgatctatgccccga-gggtcattctctatgaagagcccgaaacttgtctgagt
gagaggcgaaggtagtccttcattt--atgcggggtccgccctcacactgtagtcggcac
agtcccttgtttatggactgcgcagctcaatcttcgactggagagg--aaaaacaaaaca
caagtatgacacgcacatacgtgttttctaggactgtgacctgtcggctgcctcgcggtg
gttggcatgatgccccaagtggggggggggagaaacaatcagtatgattgataaaaacca
gtgggg-aaaaaggcgcgctgtcgcacacccttggtcacggcctc---aagtcgacggcc
ctcactggccca-ccctgcgactcctgttcgcttttgacaagacacacctgcaata-cgg
aggaaatgcctgagcagctgtgccgcagcaactgcggacgcgtagcgatatgcatcccta
ttgtttgcgacttggtacaaagaacgtgcgccgggagacggattgccctttgacttttcc
acccagcccgcggctctatacccttgacgtacctatttacctggttgatcctgccagtag
tcatatgcttgtctcaaagactaagccatgcatgtctaagtataaacacttttatactcg

tgaaactgcgaatggctcattaaaacagttatagttcctttgggaacaacactacttgga
taaccgtagtaattctagagctaatacatgcctcaacgcccgactcacgaagggtggtat
ttattggataaaaaaccatcgtgctctgtccagagcgctttgagatgattcacaataact
tgtcggatcgcatggcctcgtgctggcgactcctcattcaaatttctgccctatcaactt
tcgatggtagagtattggtctaccatggtgtcgacgggtgacggggaattagggttcgat
tccggagagggagcctgagaaacggctaccacatccaaggaaggcagcaggcgcgcaaat
tacccaatcccgactcggggaggtagtgacaaaaaataacaatagggggccctttgggtc
ttctaattggaatgagaacaatttaaatcccttatcgaggatccattggagggcaagtct
ggtgaacgta----------------------------------tgcatta---------
--aagccctccgtaat-aaaaaaaaacagcgccagatgtct-------------------
------------------------------------------------------------
------------------------------------------------------------
------------------------------------------------------------
------------------------------------------------------------
------------------------------------------------------------
-------ctctctggaagcacacca-----------------------------------
---tgcatggggtttgttcctctc-----------------------------tgcatgg
ccgtg-----------------------------------------tgaaaaaggcgagt
aggcaaacgatgggaaacccggcttggaatagtaaccagccacagagcccccatcataca
cttctaaattgccggggagtgcccgcgaggccttgactaccgcggtccgccgagaggtga
gagcgagcaatcgttcaggacgcagcagtgtacgtagtggtacatggatggtaaaaacgt
caaggatagggttgatccgcagggaagcgcgagtgccgccatg---------gtcggtgg
gcaaaggcgaaccttcagagactctaatggagtgggcgcaagcttaagggagagtccaat
ccactgggaaaccaggtccacagcagtaacggggtgatgctcgccccagaggactgtgga
aggcaaccccacc----aaaagggtgtgtgttgtcgggtagc-aatggccagcagccgcg
gtaattccagctccaatagcgtatattaaagttgttgcagttaaaacgctcgtagtcgga
cctcggagcgcccccacgggccgggcttttgccttgttgccagtgggagtttgtctcctt
ttgtcgggacgcggtgtgcagggcctttactgtctctgcgcgtttacctgcgggccgacc
gtttactgtgaagaaagtaaagtgttcaaagcaggccaatgccttgaatatgtgagcatg
gaataatagaataggacttgggctctattttgttggtttccagtgaccaagtaatgatta
atagggacggttgggggcattcgtatttcattgtcagaggtgaaattcttggattgatgg
aagacgtacaactgcgaaagcatctgccatggatgttttcattgatcaagaacgaaagtt
aggggatcgaagacgatcagataccgtcgtagtcttaaccataaacgatgccgactgggg
attggcgggggcacttttatatgactctgtcagcaccctgagggaaaccaaagtctttgg
gttctgggggggagtatggtcgcaaggctgaaacttaaaggaattgacggaagggcaccac
caggagtggagcctgcggcttaatttgactcaacacgggaaaacttaccaggtccggaca

gaagaatgattgacagactgaaagctctttcttgatttttttggttggtggtgcatggccg
ttcttagttggtggagtgatttgtctggttaattccgttaacgaacgagacctcgtcctg
ctaaataggtgcgcgcatgccagcactgcgttcttaccttcttagagggactatgcgcgt
ctagcgtatggaagattgaggcaataacaggtctgtgatgcccttagatgttctgggccg
cacgcgcgctacactgatgcattcaacgagtttcacttgatagtcctgggtcggaaggct
cgggtaatcttttgaaagtgcatcgtgctggggatagatcattgcaattattgatcttca
acgaggaattccttgtaggcgcaggtcatcagcctgcgccgaatacgtccctgccctttg
tacacaccgcccgtcgctcctaccgattgaatggtccgatgaaatgtccggatcgcggct
agtgtgatggactcgtccagagcggtagccgtgagaagctcattaaatcttatcatttag
aggaaggagaagtcgtaacaaggtttccgtattttgtggatgggcactcgtc--------
------------------------------------------------------------
------------------------------------------------------------
------------------------------------------------------------
---------gtggcgttctttgcgccacgcggat--------------------------
------------------------------------------------------------
------------------------------------------------------------
------------------------------------------------------------
-----------------------------------gtccccttccgtaggaaaacgtaca
cgaagcctttgcggcccgaaagggtgcacagtgcgcgactataaacactgattctgtggt
caacgcaagcagcggttctgtgacagaacagctgcgactttctcaaattgcgggaacacc
ctaaagtcaacaaacaccgcggccggacgtg-aaagcgtcaggcgcagcagtcggggtag
cgccctggcggatggtaagagctttgttgaatggcacaatgggcaatccgcagccaagcc
cctaaggtccttcgc--ggctatggggaaggttcacagactgtaagggagggggtttctt
ccat---gaagaagcttaagagacagtcggtcccctgcgagagcagtgtctcatgtgagg
aaggtgcccatgaaaagtgggcgctggagagccatgggagccctctatgagggtggggag
ttttcctagataataatgagaaacaaacacgttccgtaggtgaacctgcggaaggatcat
tcacaaactattgacacaaacacacgcgaaccaaatcgtccgcacagcaggtggcaatga
aagagagactctgcatgtcgcctttcgggg--tagcaagcagcactcttctgccatcgcc
tctgtgcagggcgtaaattcccattgagaggatgtgagggcaccacaggaagcttttcca
caggaagtcgccatccttctccctccacggcggcg----cttctgtgcttggcag-----
-----ttttttttgccttctagggaggatgccgccaatggagccccttat-atatataca
tcatcatatgccc--cttttttttcttaaccgcttgccacagcttcttctatgaggagctt
gtgggaagactgtctccatacaataac-aaagatacaactcttagcggtggatatcttgg
ttctcgcaacgatgaagaacgcagccaactgcgataactaatgtgaattgcaggacttcg
tgaatcattaagtctttgaacgcaagttgcgctcatctccatgaggaggtgagcatgcct
gtttgagtgtcgatgggaaaaacaatac-ataacaacaatcggctctcttatcttcctgc

cttcttgggggcaaaaggtagtgcatgccggagatgggctttgtcgcccctttctcttga
aagaggcgtcgtccgaaacacgtatcgttgcccgctggaggtcatagagcatgagttt--
aaagcaccacatctctcaagtcagagtgag-cgtcttgagtaggtcc-gcattgattcag
tttcaacccaggtggcgttggttacttgccat-aagcctctctggggtttctttgcttgg
acctctcttgtcagtttctctgacgtagagacaggt--gccgtctcatggacgttgccct
ctggaacgtgct--aaaaaaaaaaaaaga----------aaaaaccgtaccacatgtcaa
aagaggcagatcgtgcttgacggcgggaggtgttgcatttcttgcttgtcctgagcgtct
gccagtctct----ga--cgagtg----tgttccttccataaacccgatctcaaatcagg
taggaccacccgctgaatttaagcatataattaagcggaggaaaagaaactaacaaggat
tccctcagtagcggcgagcgaagcgggaagagctcaacatgaaaaccgttgggggttcgc
gctttccttcgggttgtatagtaaagtggtgtcgcctgcgtcgcgccgggtataaatccc
gtctgaaatggggtgtcgtagagggttatatccccgtctgtgacccggtgtgcgatgcgt
tacggtgaccttacgatgagtcgggttgcttgggcatgcagccctaagtgggtggtaaac
ttcatctaaagctaaatactagcgagagaccgatagcgtacaagtaccgtaagggaaaga
tgcaaagaactttgaaaagagagttaaacagggcttgaaaccgttaagagtgaagcgaag
gaggtcagtgatcatggggccagatccccaagtgcctgtcata-gggcaccgtagcgttc
tggtttcccggggcagcgccggccgggggcgggggttaacagggaggtggttgagggatt
cgttttccgcctgtactcctttatagcttgcaccccggccgagggatccgcgacttgtcg
tgtgagggcgctgtcaaaatggccttctccaccccgtcttgaaacacggaccaaggagtc
taacatactcgcgagtgtttgggtggttaaacccgaacgcacaatgaaagtaaacggtg
gctctgcgccgctgcttggtgggaaccctgcttttataatgggggcactatcacccgac
cccgatccctttggggtggttggttttgaggtagagcgagtatgatgggacccgaaagat
ggtgaactattcctgagtagggcgaagccagaggaaactctggtggaggctcgtagcgat
gctgacgtgcaaatcgctcgtcgaacttgggtataggggcgaaagactaatcgaaccatc
tagtagctggttccctccgaagtttccctcaggatagcgggagcaaatgcaagtcagtat
catccggtaaagcgaatgattagaggcctcgggggagtaactccttcgacctattctcaa
actttaaataggtgaggagccggcaacgttgcttaaaacaatgaacgccgtgcggacaat
gattgctctcagtgggccgattttggtaagcagaactggcgatgcgggatgaaccgaaag
cgacgctaaggtgccaaactcccagctcattcagacaccagaaaaggtgttgattcatac
agacagcaggacggtggccatggaagtcggaatccgctaaggagtgtgtaacaactcacc
tgccgaatggatcagccctgaaaatggatggcgcttaagcgggggaccgaggcgtcgccg
ttggggtaaatacgaagccccaatgagtaggagggcgtgggggttttgaaacacggtaca
ctcgcatggaagccttgggcgtaagctcgggtggagcgacccctagtgtagatcttggtg
gtagtagcaaatattcaagtgagaaccttgaagactgaagtggggaagggttccacgcga
acagtgcttggacgtgggttagtcggtcctaagagatggggaaactccgtattcaagtgc
gcttttgtagcgcagtccatcgaaagggaagcgggttaacattcccgcaccaggatgtgg

aaactgtggtttaccgcaaactgcggcaacgcaactgaacctggggacgcccgtaggagc
tccgagcagggatctctttcctgtttaacgcgcccaacccccctggaatcggtttgtccgg
agatagggggcgaaaggccggaaaagcaccgcacgtcttgtggtgttcggaacgcttgta
atggcccttgaaaacccaggggaaagcagccgtcttacac--ttgtgtgagacgaggaat
tctcacacctggccgtacccatcaccgcatcaggtccccaaggtgaacagcctctagtca
atagaataatgtagataagggaagtcggcaaaatagatccgtaactttgggaaaaggatt
ggctctaatgttcgggcttgtcgggacgcgcgggcgtagatgccaatgagcggcagacgt
ctagcctctttggaggctggattaacgctgcttgtgggaagaaaccgtgtcgatggaggc
aggaacgcttatcgatcttgtctcttcccgcttgcgatcaacgaacggattagaactggt
acggacaaggggaatccgactgtttaattaaaacaaagcattgcgacggccggaaacggt
gttggcgcaatgtgatttctgcccagtgctctgaatgtcaaagtgaagaaattcaaccaa
gcgcgggtaaacggcgggagtaactatgactctcttaagg--------------------
------------------------------------------------------------
------------------------------------------------------------
------------------------------------------------------------
------------------------------------------------------------
------------------------------------------tagcc-------------
------------------------------------------------------------
------------------------------------------------------------
----------------------------------------------------------aa
atgcctcgtcatct----------------------------------------------
------------------------------------------------------------
------------------------------------------------------------
------------------------------------------------------------
------------------------------------------------------------
------------------------------------------------------------
------------------------------------------------------------
------------------------------------------------------------
------------------------------------------------------------
------------------------------------------------------------
------------------------------------------------------------
------------------------------------------------------------
------------------------------------------------------------
------------------------------------------------------------
------------------------------------------------------------
------------------------------------------------------------

------------------------------------------------------------
------------------------------------------------------------
------------------------------------------------------------
------------------------------------------------------------
------------------------------------------------------------
------------------------------------------------------------
---aattagtgacgcgcatgaatggattaacgagattcccactgtccctatctactatct
agcgaaaccacagccaagggaacgggcttggaagaatcagcggggaaagaagaccctgtt
gagcttgactctagtccgactttgtgagatgagttgggaggtgtagtctaagagggaggg
gcaactcgaccgtgaaatacctctactcccaagcttgttttacttactccggcgagcagg
gatgggtggtaacacccttttctaggattaacacctcttcgggggtggacctgcgcgga
ggacattgtcaggtggggagtttggctggggcggcacatctgttaaaagataacgcaggt
gtcctaagatgagctcagtgagaacagaaatctcacgtagaacaaaagggtaaaagctca
tttgattttgattttcagtacgaatacaaactgtgaaagcatggcctatcgatcctttttg
accttcaaggtttgaagcaagaggtgtcagaaaagt------------------------
------------------------------------------------------------
------------------------------------------------------------
------------------------------------------------------------
------------------------------------------------------------
------------------------------------------------------------
------------------------------------------------------------
---------------------------------------------------taccacagg
gat---------------------------------------------------------
------------------------------------------------------------
------------------------------------------------------------
------------------------------------------------------------
------------------------------------------------------------
------------------------------------------------------------
------------------------------------------------------------
------------------------------------------------------------
------------------------------------------------------------
------------------------------------------------------------
------------------------------------------------------------
------------------------------------------------------------
------------------------------------------------------------
------------------------------------------------------------

------------------------------------------------------------
------------------------------------------------------------
------------------------------------------------------------
------------------------------------------------------------
------------------------------------------------------------
------------------------------------------------------------
------------------------------------------------------------
------------------------------------------------------------
---------------------------------aactggcttgtggcagccaagcgttca
tagcgacgttgctttttgatccttcgatgtcggctcttcctatcattgcgaagcagaatt
cgccaagtgttggattgttcacccactaatagggaacgtgagctgggtagccacttta--
-----tgcccaacagctggatagctgggcttctgtctta-ggccgggacttctcacgacc
cgtgccttggtcaccc-aaaaaagggcgaccgcagagctcaggatggagcttctggctta
actagtagaggacagatagagatcgtcaaattgcgggaaaccccttagagcctcgagcta
ccaagtgcgtgtcgaaaggcatagacgcagccagcgatggctgtcgcatggcttga-ttc
catcaagataggtaaaaatgctcgaggattgggcaatccgcagccaagcccctaaggt--
---ccccccgcgggctatggggaaggttcaacgactgtaaggcgtaccgtgcaagcggaa
gatacagtctagccccgctggaaagcagcgccaaagtaagtaggt-tgcagtgggtctgt
cctc
>Form_J_China
gcatgttttggtatgtgtttcgacagtcatgcatgtcatagccgtaggatggggagtgag
cttcatagcccgaccccttctcaggagcatgttttcatagcatattgagcatactggggt
ggtctcccatgtagctcgctctctcggtgcaggacctcccttttgcgggccacttcgatg
gcctccattatgga-gtgcatgttttggcatatgaagcgttggtggtgaggctcgtagcc
tgtatatacataccaacgcagtcgtcatatgtctctctggcgggggtcggtcgtatgacc
ttgtccctacctggagcatgtattttgtatgat---atactggac-ggcagctgcgctgt
tcagtcgtcagccgtacctcatgtgcgcgtttccaggtccgaacgtttgacatgcaaggg
catcataccagaagcctgttttctcgtatgagcatgtaggtaggcaggctcttcagtcgt
ctacccataaatgcacttcgaagga-actttttggtattgaatggctttgtgcccagacg
ttcttgtgagcttgttctcgctctgcgatacgctggaagctgtggtattacctacagcag
acgcattcctagggcggacggcctctctgcagatgggcacatctgtagaaagaggctccc
cttgatgacgcaatgcaatggctgggtaaacggagtgctatacagaagggtcaaacctc-
tgtggaagtacactcataccagcattgtttgttgagaggggtcggctctacgagtgaggc
cctatactgccatctcattcagatgcgtagacggtctttactcagactgcctttctggct
gattgtcctcggtccctcgtgcgcacgcgccgctggctgaatgggcttgtccttatggag
tgccgcgttcttc-aaagaatccgtcttcatacagcgacctcatttacatattgccggct

gaatgggcttgtccttatggagcactgcattcttctaggaattgctatctatacagcgac
ctcatttacatacaaggcacct-ttttctacatagacttgccatttggctgattgttctg
acagtcgctgcctcttcggagttctgtgtggacgagtggcc--tcctgagtgggccggtc
cttacggagtgcccaagtctgttatgactgggttctccatacagcgacctcactcagaat
tgtatgg--------------caccgtgccgat-----------tgaatggacaagcagt
ccttagggaatcgcatgaaagcatgacggcaacatggtgtatggacccgttgggccttgc
ctgtgctatcttctcgagtcgaccgg------------tcctcgtctcctcctttcgcga
cgggaagctcttgagtttaccgtgggtcaggacgctgtggtatctgtttgtctgtagtgt
gggctctctatcagaacagcgtatcagcacacgtcgtctgggtctatggtatttttgctt
aacggccctgggtttggtagtatggttccatgtcggtcctctcacctcccaaccgtcatg
tctc--------------------------------------------------------
------------------------------------------------------------
--------------------------------------------tatcatctgcacagta
cgtcttgtcagaatatcattctgtatggcagtcgatagactaggccctatgccgcagctg
tctctgaccactatgcgagcgcatacctacctcaatgtgttcatagctgttatggatctt
gggtgtggtgtgccagaacgatacgccaccagattgtggaatggtgcttgtagtattaga
agaggacatgtatgaagagcacgttcttttctcttatgatggtgccgcgggtgcgggccg
ttctcattctgcttgtcgggtgctcaaggtgcatgtaggattgctgaacccac-tgaatg
gtcatgagaatggtcatatggtgaggttcacttctccatcgctttgtgtgtgttggatgt
ttgggtggcttcggccgcatggaaaatctcggtctgtaactcatcagggagggtttccat
gtgagcaatctctccagtcgaacacacacccatgggcgtcccgctaggcaaaatggaatt
ctcgatccgacacagcctgtggagtgtttgttccgtgtggcgtgaaacggcttatactgg
atgtcttccc-tttggacaccagtctgttggtatg--tacgcataccatctagagggagc
aagaaaaaggcatgcaagtgccacaaaggaagggaggccttcctgctaccgtgtgtcc-t
cttgacgatcacgggggagtatatagagacatttggacgagatcctttttttcacgaggcg
gtgccctcgcgcaaaggtgttgctctgcacgcatctttatgatgatagttgctcgctcct
atgcgtccactcgtgaaaatacctgctctctctgtctgtttccatgacacccc-catgcg
ttgtcatagagcatgctcaggtacagaaagtgctgtgactttcttggcctgctgtcactg
agtcagcctgatactttggaggaatcttttctgggcactgcccagctgagagttcttatc
aggctctggagtgtgcatatgcctatattcattatagtgcatgtggctacgcagaaaaag
catcgacgtggaatggagaagaacaccattctgaatggaacggttcttactctacttctg
ttgcttgacgctgtatgggaatcgctttgctctgggggtacaccttccattccatgacaa
agcatttgatctatgccccga-gggtcattctctatgaagagcccgaaacttgtctgagt
gagaggcgaaggtagtccttcattt--atgcggggtccgccctcacactgtagtcggcac
agtcccttgtttatggactgcgcagctcaatcttcgactggagagg--aaaaacaaaaca
caagtatgacacgcacatacgtgttttctaggactgtgacctgtcggctgcctcgcggtg

gttggcatgatgccccaagt-gggggggggagaaacaatcagtatgattgataaaaacca
gtgggg-aaaaaggcgcgctgtcgcacacccttggtcacggcctc---aagtcgacggcc
ctcactggccca-ccctgcgactcctgttcgcttttgacaagacacacctgcaata-cgg
aggaaatgcctgagcagctgtgccgcagcaactgcggacgcgtagcgatatgcatccctg
ttgtttgcgacttggtacaaagaacgtgcgccgggagacggattgccctttgacttttcc
acccagcccgcggctctatacccttgacgtacctatttacctggttgatcctgccagtag
tcatatgcttgtctcaaagactaagccatgcatgtctaagtataaacacttttatactcg
tgaaactgcgaatggctcattaaaacagttatagttcctttgggaacaacactacttgga
taaccgtagtaattctagagctaatacatgcctcaacgcccgactcacgaagggtggtat
ttattggataaaaaaccatcgtgctctgtccagagcgctttgagatgattcacaataact
tgtcggatcgcatggcctcgtgctggcgactcctcattcaaatttctgccctatcaactt
tcgatggtagagtattggtctaccatggtgtcgacgggtgacggggaattagggttcgat
tccggagagggagcctgagaaacggctaccacatccaaggaaggcagcaggcgcgcaaat
tacccaatcccgactcggggaggtagtgacaaaaaataacaatagggggccctttgggtc
ttctaattggaatgagaacaatttaaatcccttatcgaggatccattggagggcaagtct
ggtgaacgta----------------------------------tgcatta---------
--aagccctccgtaataaaaaaaaaacagcgccagatgtct-------------------
------------------------------------------------------------
------------------------------------------------------------
------------------------------------------------------------
------------------------------------------------------------
------------------------------------------------------------
-------ctctctggaagcacacca-----------------------------------
---tgcatggggtttgttcctctc-----------------------------tgcatgg
ccgtg-----------------------------------------tgaaaaaggcgagt
aggcaaacgatgggaaacccggcttggaatagtaaccagccacagagcccccatcataca
cttctaaattgccggggagtgcccgcgaggccttgactaccgcggtccgccgagaggtga
gagcgagcaatcgttcaggacgcagcagtgtacgtagtggtacatggatggtaaaaacgt
caaggatagggttgatccgcagggaagcgcgagtgccgccatg---------gtcggtgg
gcaaaggcgaaccttcagagactctaatggagtgggcgcaagcttaagggagagtccaat
ccactgggaaaccaggtccacagcagtaacggggtgatgctcgccccagaggactgtgga
aggcaaccccacc----aaaagggtgtgtgttgtcgggtagc-aatggccagcagccgcg
gtaattccagctccaatagcgtatattaaagttgttgcagttaaaacgctcgtagtcgga
cctcggagcgcccccacgggccgggcttttgccttgttgccagtgggagtttgtctcctt
ttgtcgggacgcggtgtgcagggcctttactgtctctgcgcgtttacctgcgggccgacc
gtttactgtgaagaaagtaaagtgttcaaagcaggccaatgccttgaatatgtgagcatg

gaataatagaataggacttgggctctattttgttggtttccagtgaccaagtaatgatta
atagggacggttgggggcattcgtatttcattgtcagaggtgaaattcttggattgatgg
aagacgtacaactgcgaaagcatctgccatggatgttttcattgatcaagaacgaaagtt
aggggatcgaagacgatcagataccgtcgtagtcttaaccataaacgatgccgactgggg
attggcgggggcacttttatatgactctgtcagcaccctgagggaaaccaaagtctttgg
gttctgggggggagtatggtcgcaaggctgaaacttaaaggaattgacggaagggcaccac
caggagtggagcctgcggcttaatttgactcaacacgggaaaacttaccaggtccggaca
gaagaatgattgacagactgaaagctctttcttgatttttttggttggtggtgcatggccg
ttcttagttggtggagtgatttgtctggttaattccgttaacgaacgagacctcgtcctg
ctaaataggtgcgcgcatgccagcactgcgttcttaccttcttagagggactatgcgcgt
ctagcgtatggaagattgaggcaataacaggtctgtgatgcccttagatgttctgggccg
cacgcgcgctacactgatgcattcaacgagtttcacttgatagtcctgggtcggaaggct
cgggtaatcttttgaaagtgcatcgtgctggggatagatcattgcaattattgatcttca
acgaggaattccttgtaggcgcaggtcatcagcctgcgccgaatacgtccctgccctttg
tacacaccgcccgtcgctcctaccgattgaatggtccgatgaaatgtccggatcgcggct
agtgtgatggactcgtccagagcggtagccgtgagaagctcattaaatcttatcatttag
aggaaggagaagtcgtaacaaggtttccgtattttgtggatgggcactcgtc--------
------------------------------------------------------------
------------------------------------------------------------
------------------------------------------------------------
---------gtggcgttctttgcgccacgcggat--------------------------
------------------------------------------------------------
------------------------------------------------------------
------------------------------------------------------------
-----------------------------------gtccccttccgtaggaaaacgtaca
cgaagcctttgcggcccgaaagggtgcacagtgcgcgactataaacactgattctgtggt
caacgcaagcagcggttctgtgacagaacagctgcgactttctcaaattgcgggaacacc
ctaaagtcaacaaacaccgcggccggacgtg-aaagcgtcaggcgcagcagtcggggtag
cgccctggcggatggtaagagctttgttgaatggcacaatgggcaatccgcagccaagcc
cctaaggtccttcgc--ggctatggggaaggttcacagactgtaagggaggggggtttctt
ccatgaagaagaagcttaagagacagtcggtcccctgcgagagcagtgtctcatgtgagg
aaggtgcccatgaaaagtgggcgctggagagccatgggagccctctatgagggtggggag
ttttcctagataataatgagaaacaaacacgttccgtaggtgaacctgcggaaggatcat
tcacaaactattgacacaaacacacgcgaaccaaatcgtccgcacagcaggtggcaatga
aagagagactctgcatgtcgcctttcgggg--tagcaagcagcactcttctgccatcgcc
tctgtacagggcgtaaattcccattgagaggatgtgagggcaccacaggaagcttttcca

caggaagtcgccatccttctccctccacggcggcg----cttctgtgcttggcag-----
-----ttttttttgccttctagggaggatgccgccaatggagccccttat-atatataca
tcatcatatgccc--cttttttttcttaaccgcttgccacagcttcttctatgaggagctt
gtgggaagactgtctccatacaataac-aaagatacaactcttagcggtggatatcttgg
ttctcgcaacgatgaagaacgcagccaactgcgataactaatgtgaattgcaggacttcg
tgaatcattaagtctttgaacgcaagttgcgctcatctccatgaggaggtgagcatgcct
gtttgagtgtcgatgggaaaaacaatac-ataacaacaatcggctctcttatcttcctgc
cttcttgggggcaaaaggtagtgcatgccggagatgggctttgtcgcccctttctcttga
aagaggcgtcgtccgaaacacgtatcgttgcccgctggaggtcatagagcatgagttt--
aaagcaccacatctctcaagtcagagtgag-cgtcttgagtaggtcc-gcattgattcag
tttcaacccaggtggcgttggttacttgccataaagcctctctggggtttctttgcttgg
acctctcttgtcagtttctctgacgtagagacaggt--gccgtctcatggacgttgccct
ctggaacgtgct--aaaaaaaaaaaaaga----------aaaaaccgtaccacatgtcaa
aagaggcagatcgtgcttgacggcgggaggtgttgcatttcttgcttgtcctgagcgtct
gccagtctct----ga--cgagtg----tgttccttccataaacccgatctcaaatcagg
taggaccacccgctgaatttaagcatataattaagcggaggaaaagaaactaacaaggat
tccctcagtagcggcgagcgaagcgggaagagctcaacatgaaaaccgttgggggttcgc
gctttccttcgggttgtatagtaaagtggtgtcgcctgcgtcgcgccgggtataaatccc
gtctgaaatggggtgtcgtagagggttatatccccgtctgtgacccggtgtgcgatgcgt
tacggtgaccttacgatgagtcgggttgcttgggcatgcagccctaagtgggtggtaaac
ttcatctaaagctaaatactagcgagagaccgatagcgtacaagtaccgtaagggaaaga
tgcaaagaactttgaaaagagagttaaacagggcttgaaaccgttaagagtgaagcgaag
gaggtcagtgatcatggggccagatccccaagtgcctgtcata-gggcaccgtagcgttc
tggtttcccggggcagcgccggccgggggcgggggttaacagggaggtggttgagggatt
cgttttccgcctgtactcctttatagcttgcaccccggccgagggatccgcgacttgtcg
tgtgagggcgctgtcaaaatggccttctccaccccgtcttgaaacacggaccaaggagtc
taacatactcgcgagtgtttgggtggttaaaacccgaacgcacaatgaaagtaaacggtg
gctctgcgccgctgcttggtgggaaccctgcttttataatgggggcactatcacccgac
cccgatccctttggggtggttggttttgaggtagagcgagtatgatgggacccgaaagat
ggtgaactattcctgagtagggcgaagccagaggaaactctggtggaggctcgtagcgat
gctgacgtgcaaatcgctcgtcgaacttgggtataggggcgaaagactaatcgaaccatc
tagtagctggttccctccgaagtttccctcaggatagcgggagcaaatgcaagtcagtat
catccggtaaagcgaatgattagaggcctcgggggagtaactccttcgacctattctcaa
actttaaataggtgaggagccggcaacgttgcttaaaacaatgaacgccgtgcggacaat
gattgctctcagtgggccgattttggtaagcagaactggcgatgcgggatgaaccgaaag
cgacgctaaggtgccaaactcccagctcattcagacaccagaaaaggtgttgattcatac

agacagcaggacggtggccatggaagtcggaatccgctaaggagtgtgtaacaactcacc
tgccgaatggatcagccctgaaaatggatggcgcttaagcgggggaccgaggcgtcgccg
ttggggtaaatacgaagccccaatgagtaggagggcgtgggggttttgaaacacggtaca
ctcgcatggaagccttgggcgtaagctcgggtggagcgacccctagtgtagatcttggtg
gtagtagcaaatattcaagtgagaaccttgaagactgaagtggggaagggttccacgcga
acagtgcttggacgtgggttagtcggtcctaagagatggggaaactccgtattcaagtgc
gcttttgtagcgcagtccatcgaaagggaagcgggttaacattcccgcaccaggatgtgg
aaactgtggtttaccgcaaactgcggcaacgcaactgaacctggggacgcccgtaggagc
tccgagcagggatctctttcctgtttaacgcgcccaacccccctggaatcggtttgtccgg
agatagggggcgaaaggccggaaaagcaccgcacgtcttgtggtgttcggaacgcttgta
atggcccttgaaaacctaggggaaagcagccgtcttacac--ttgtgtgagacgaggaat
tctcacacctggccgtacccatcaccgcatcaggtccccaaggtgaacagcctctagtca
atagaataatgtagataagggaagtcggcaaaatagatccgtaactttgggaaaaggatt
ggctctaatgttcgggcttgtcgggacgcgcgggcgtagatgccaatgagcggcagacgt
ctagcctctttggaggctggattaacgctgcttgtgggaagaaaccgtgtcgatggaggc
aggaacgcttatcgatcttgtctcttcccgcttgcgatcaacgaacggattagaactggt
acggacaaggggaatccgactgtttaattaaaacaaagcattgcgacggccggaaacggt
gttggcgcaatgtgatttctgcccagtgctctgaatgtcaaagtgaagaaattcaaccaa
gcgcgggtaaacggcgggagtaactatgactctcttaagg--------------------
------------------------------------------------------------
------------------------------------------------------------
------------------------------------------------------------
------------------------------------------------------------
------------------------------------------tagcc-------------
------------------------------------------------------------
------------------------------------------------------------
----------------------------------------------------------aa
atgcctcgtcatctaa--actaattggatgagatagtctgagtgaaaagcttatgcaggc
tagtacctgtct-------aatcgtccacagtgcaattctattcgtcagcgacaggctca
ctcacg-----------------------ttaacaccacgagaagc--------------
------------------------------aatggagcagcgc-----------------
----gggtggtgcttgc--------------acaccaacacccat---------------
-------------------------------------------tgctgcccatcgt----
---------------------------------cgatgaaaacaactccc----------
------------------------------------------------------------
-----------------------ggacatccg----------cgccgcgatatgttgata

aatcgactctcgtagcacagg---------------------------------tgggcc
gggttgcagcaccaggtgttgccgcacaggtgcgatgtgtcggaattgaccggcttaaca
ccgcagcgtacctcggcaactagcacgtgcaccatggcatgtct--gcccataaaactgg
tgcgataatagcca----------gacggctggcgaaatacacctggtccagtcggctgg
cggcacttgaagatattcttgtaaccaccaactgacat----------------------
-----------------------gacacggtcggccaacaagtcgttgacgaacgcatct
agtgtagtcaacgaggtgtactggttgcggtg-gcgagacatggtgtagaaacgtgtgtg
tg----ttgacgaagtgaaatgtattgtgtagatgggtgcgacactccttgtacagggaa
accctcc--gcatggaaaggcacgtggtaccaccctcgcggggaaactcgcgagcgggat
cccgggtaatgaccggagccgatggtaaaaacccacgtgaccgatggggtgaccctgtgg
cgagacgctaagttgtgggacatcaccctctcacaatacgcatccgtcgcagagactgaa
cgggagtgggctccgtcga-------------cggccagccagccgtcttccag------
--gggggcttaagatacagtccgggcctcagggatgg----aaaaacctggggaacta-c
cagaattagtgacgcgcatgaatggattaacgagattcccactgtccctatctactatct
agcgaaaccacagccaagggaacgggcttggaagaatcagcggggaaagaagaccctgtt
gagcttgactctagtccgactttgtgagatgagttgggaggtgtagtctaagagggaggg
gcaactcgaccgtgaaatacctctactcccaagcttgttttacttactccggcgagcagg
gatgggtggtaacaccctttttctaggattaacacctcttcgggggtggacctgcgcgga
ggacattgtcaggtggggagtttggctggggcggcacatctgttaaaagataacgcaggt
gtcctaagatgagctcagtgagaacagaaatctcacgtagaacaaaagggtaaaagctca
tttgattttgattttcagtacgaatacaaactgtgaaagcatggcctatcgatcctttttg
accttcaaggtttgaagcaagaggtgtcagaaaagt------------------------
------------------------------------------------------------
------------------------------------------------------------
------------------------------------------------------------
------------------------------------------------------------
------------------------------------------------------------
------------------------------------------------------------
---------------------------------------------------taccacagg
gat---------------------------------------------------------
------------------------------------------------------------
------------------------------------------------------------
------------------------------------------------------------
------------------------------------------------------------
------------------------------------------------------------
------------------------------------------------------------

------------------------------------------------------------
------------------------------------------------------------
------------------------------------------------------------
------------------------------------------------------------
------------------------------------------------------------
------------------------------------------------------------
------------------------------------------------------------
------------------------------------------------------------
------------------------------------------------------------
------------------------------------------------------------
------------------------------------------------------------
------------------------------------------------------------
------------------------------------------------------------
------------------------------------------------------------
------------------------------------------------------------
---------------------------------aactggcttgtggcagccaagcgttca
tagcgacgttgctttttgatccttcgatgtcggctcttcctatcattgcgaagcagaatt
cgccaagtgttggattgttcacccactaatagggaacgtgagctgggtagccacttta--
-----tgcccaacagctggatagctgggcttctgtctta-ggccgggacttctcacgacc
cgtgccttggtcaccc-aaaaaagggcgaccgcagagctcaggatggagcttctggctta
actagtagaggacagatagagatcgtcaaattgcgggaaaccccttagagcctcgagcta
ccaagtgcgtgtcgaaaggcatagacgcagccagcgatggctgtcgcatggcttga-ttc
catcaagataggtaaaaatgctcgaggattgggcaatccgcagccaagcctctaaggt--
---ccccccgcgggctatggggaaggttcaacgactgtaaggcgtaccgtgcaagcggaa
gatacagtctagccccgctggaaagcagcgccaaagtaagtaggt-tgcagtgggtctgt
cctc
>form_H_China
gcatgttttggtatgtgtttcgacagtcatgcatgtcatagccgtaggatggggagtgag
cttcatagcccgaccccttctcaggagcatgttttcatagcatattgagcatactggggt
ggtctcccatgtagctcgctctctcggtgcaggacctcccttttgcgggccacttcgatg
gcctccattatgga-gagcatgttttggcatatgaagcgttggtggtgaggctcgtagcc
tgtatatacataccaacgcagtcgtcatatgtctctctggcgggggtcggtcgtatgacc
ttgtccctacctggagcatgtattttgtatgat---atactggac-ggcagctgcgctgt
tcagtcgtcggccgtacctcatgtgcgcgtttccaggtccgaacgtttgacatgcaaggg
catcataccagaagcctgttttctcgtatgagcatgtaggtaggcaggctcttcagtcgt
ctacccataaatgcacttcgaaggga-actttttggtattgaatggctttgtgcccagacg

ttcttgtgagcttgttctcgctctgcgatacgctggaagctgtggtattacctacagcag
acgcattcctagggcggacggcctctctgcagatgggcacatctgtagaaagaggctccc
cttgatgacgcaatgcaatggctgggtaaacggagtgctatacagaagggtcaaacctc-
tgtggaagtacactcataccagcattgtttgttgagaggggtcggctctacgagtgaggc
cctatactgccatctcattcagatgcgtagacggtctttactcagactgcctttctggct
gattgtcctcggtccctcgtgcgcacgcgccgctggctgaatgggcttgtccttatggag
tgccgcgttcttc-aaagaatccgtcttcatacagcgacctcatttacatattgccggct
gaatgggcttgtccttatggagcactgcattcttctaggaattgctatctatacagcgac
ctcatttacatacaaggcacct-ttttctacatagacttgccatttggctgattgttctg
acagtcgctgcctcttcggagttctgtgtggacgagtggcc--tcctgagtgggccggtc
cttacggagtgcccaagtctgttatgactgggttctccatacagcgacctcactcagaat
tgtatgg--------------caccgtgccgat-----------tgaatggacaagcagt
ccttagggaatcgcatgaaagcatgacggcaacatggtgtatggacccgttgggccttgc
ctgtgctatcttctcgagtcgaccgg------------tcctcgtctcctcctttcgcga
cgggaagctcttgagtttaccgtgggtcaggacgctgtggtatctgtttgtctgtagtgt
gggctctctatcagaacagcgtatcagcacacgtcgtctgggtctatggtatttttgctt
aacggccctgggtttggtagtatggttccatgtcggtcctctcacctcccaaccgtcatg
tctctga-----------------------------------------------------
------------------------------------------------------------
--------------------------------------------tatcatctgcacagta
cgtcttgtcagaatatcattctgtatggcagtcgatagactaggccctatgccgcagctg
tctctgaccactatgcgagcgcatacctacctcaatgtgttcatagctgttatggatctt
gggtgtggtgtgccagaacgatacgccaccagattgtggaatggtgcttgtagtattaga
agaggacatgtatgaagagcacgttcttttctcttatgatggtgccgcgggtgcgggccg
ttctcattctgcttgtcgggtgctcaaggtgcatgtaggattgctgaacccac-tgaatg
gtcatgagaatggtcatatggtgaggttcacttctccatcgctttgtgtgtgttggatgt
ttgggtggcttcggccgcatggaaaatctcggtctgtaactcatcagggagggtttccat
gtgagcaatctctccagtcgaacacacacccatgggcgtcccgctaggcaaaatggaact
ctcgatccgacacagcctgtggagtgtttgttccgtgtggcgtgaaacggcttatactgg
atgtcttccc-tttggacaccagtctgttggtatg--tacgcataccatctagagggagc
aagaaaaaggcatgcaagtgccacaaaggaagggaggccttcctgctaccgtgtgtcc-t
cttgacgatcacgggggagtatatagagacatttggacgagatccgtttttcacgaggcg
gtgccctcgcgcaaaggtgttgctctgcacgcatctttatgatgatagttgctcgctcct
atgcgtccactcgtgaaaatacctgctctctctgtctgtttccatgacacccc-catgcg
ttgtcatagagcatgctcaggtacagaaagtgctgtgactttcttggcctgctgtcactg
agtcagcctgatactttggaggaatcttttctgggcactgcccagctgagagttcttatc

aggctctggagtgtgcatatgcctatattcattatagtgcatgtggctacgcagaaaaag
catcgacgtggaatggagaagaacaccattctgaatggaacggttcttactctacttctg
ttgcttgacgctgtatgggaatcgctttgctctgggggtacaccttccattccatgacaa
agcatttgatctatgccccga-gggtcattctctatgaagagcccgaaacttgtctgagt
gagaggcgaaggtagtccttcattt--atgcggggtccgccctcacactgtagtcggcac
agtcccttgtttatggactgcgcagctcaatcttcgactggagagg--aaaaacaaaaca
caagtatgacacgcacatacgtgttttctaggactgtgacctgtcggctgcctcgcggtg
gttggcatgatgccccaagt-gggggggggagaaacaatcagtatgattgataaaaacca
gtgggg-aaaaaggcgcgctgtcgcacacccttggtcacggcctc---aagtcgacggcc
ctcactggccca-ccctgcgactcctgttcgcttttgacaagacacacctgcaata-cgg
aggaaatgcctgagcagctgtgccgcagcaactgcggacgcgtagcgatatgcatccctg
ttgtttgcgacttggtacaaagaacgtgcgccgggagacggattgccctttgacttttcc
acccagcccgcggctctatacccttgacgtacctatttacctggttgatcctgccagtag
tcatatgcttgtctcaaagactaagccatgcatgtctaagtataaacacttttatactcg
tgaaactgcgaatggctcattaaaacagttatagttcctttgggaacaacactacttgga
taaccgtagtaattctagagctaatacatgcctcaacgcccgactcacgaagggtggtat
ttattggataaaaaaccatcgtgctctgtccagagcgctttgagatgattcacaataact
tgtcggatcgcatggcctcgtgctggcgactcctcattcaaatttctgccctatcaactt
tcgatggtagagtattggtctaccatggtgtcgacgggtgacggggaattagggttcgat
tccggagagggagcctgagaaacggctaccacatccaaggaaggcagcaggcgcgcaaat
tacccaatcccgactcggggaggtagtgacaaaaaataacaatagggggccctttgggtc
ttctaattggaatgagaacaatttaaatcccttatcgaggatccattggagggcaagtct
ggtgaacgta----------------------------------tgcatta---------
--aagccctccgtaat-aaaaaaaaacagcgccagatgtct-------------------
------------------------------------------------------------
------------------------------------------------------------
------------------------------------------------------------
------------------------------------------------------------
------------------------------------------------------------
-------ctctctggaagcacacca-----------------------------------
---tgcatggggtttgttcctctc-----------------------------tgcatgg
ccgtg------------------------------------------tgaaaaggcgagt
aggcaaacgatgggaaacccggcttggaatagtaaccagccacagagcccccatcataca
cttctaaattgccggggagtgcccgcgaggccttgactaccgcggtccgccgagaggtga
gagcgagcaatcgttcaggacgcagcagtgtacgtagtggtacatggatggtaaaaacgt
caaggatagggttgatccgcagggaagcgcgagtgccgccatg---------gtcggtgg

gcaaaggcgaaccttcagagactctaatggagtgggcgcaagcttaagggagagtccaat
ccactgggaaaccaggtccacagcagtaacggggtgatgctcgccccagaggactgtgga
aggcaaccccacc----aaaagggtgtgtgttgtcgggtagc-aatggccagcagccgcg
gtaattccagctccaatagcgtatattaaagttgttgcagttaaaacgctcgtagtcgga
cctcggagcgcccccacgggccgggcttttgccttgttgccagtgggagtttgtctcctt
ttgtcgggacgcggtgtgcagggcctttactgtctctgcgcgtttacctgcgggccgacc
gtttactgtgaagaaagtaaagtgttcaaagcaggccaatgccttgaatatgtgagcatg
gaataatagaataggacttgggctctattttgttggtttccagtgaccaagtaatgatta
atagggacggttgggggcattcgtatttcattgtcagaggtgaaattcttggattgatgg
aagacgaacaactgcgaaagcatctgccatggatgttttcattgatcaagaacgaaagtt
aggggatcgaagacgatcagataccgtcgtagtcttaaccataaacgatgccgactgggg
attggcgggggcacttttatatgactctgtcagcaccctgagggaaaccaaagtctttgg
gttctgggggggagtatggtcgcaaggctgaaacttaaaggaattgacggaagggcaccac
caggagtggagcctgcggcttaatttgactcaacacgggaaaacttaccaggtccggaca
gaagaatgattgacagactgaaagctctttcttgatttttttggttggtggtgcatggccg
ttcttagttggtggagtgatttgtctggttaattccgttaacgaacgagacctcgtcctg
ctaaataggtgcgcgcatgccagcactgcgttcttaccttcttagagggactatgcgcgt
ctagcgtatggaagattgaggcaataacaggtctgtgatgcccttagatgttctgggccg
cacgcgcgctacactgatgcattcaacgagtttcacttgatagtcctgggtcggaaggct
cgggtaatcttttgaaagtgcatcgtgctggggatagatcattgcaattattgatcttca
acgaggaattccttgtaggcgcaggtcatcagcctgcgccgaatacgtccctgccctttg
tacacaccgcccgtcgctcctaccgattgaatggtccgatgaaatgtccggatcgcggct
agtgtgatggactcgtccagagcggtagccgtgagaagctcattaaatcttatcatttag
aggaaggagaagtcgtaacaaggt------------------------------------
------------------------------------------------------------
------------------------------------------------------------
------------------------------------------------------------
------------------------------------------------------------
------------------------------------------------------------
------------------------------------------------------------
------------------------------------------------------------
------------------------------------------------------------
------------------------------------------------------------
------------------------------------------------------------
------------------------------------------------------------
------------------------------------------------------------

------------------------------------------------------------
------------------------------------------------------------
------------------------------------------------------------
-------------------------------ttccgtaggtgaacctgcggaaggatcat
tcacaaactattgacacaaacacacgcgaaccaaatcgtccgcacagcaggtggcaatga
aagagagactctgcatgtcgcctttcgggg--tagcaagcagcactcttctgccatcgcc
tctgtgcagggcgtaaattcccattgagaggatgtgagggcaccacaggaagcttttcca
caggaagtcgccatccttctccctccacggcggcg----cttctgtgcttggcag-----
-----ttttttttgccttctagggaggatgccgccaatggagccccttat-atatataca
tcatcatatgccc--ctttttttcttaaccgcttgccacagcttcttctatgaggagctt
gtgggaagactgtctccatacaataac-aaagatacaactcttagcggtggatatcttgg
ttctcgcaacgatgaagaacgcagccaactgcgataactaatgtgaattgcaggacttcg
tgaatcattaagtctttgaacgcaagttgcgctcatctccatgaggaggtgagcatgcct
gtttgagtgtcgatgggaaaaacaatac-agaacaacaatcggctctcttatcttcctgc
cttcttgggggcaaaaggtagtgcatgccggagatgggctttgtcgcccctttctcttga
aagaggcgtcgtccgaaacacgtatcgttgcccgctggaggtcatagagcatgagttt--
aaagcaccacatctctcaagtcagagtgag-cgtcttgagtaggtcc-gcattgattcag
tttcaacccaggtggcgttggttacttgccat-aagcctctctggggtttctttgcttgg
acctctcttgtcagtttctctgacgtagagacaggt--gccgtctcatggacgttgccct
ctggaacgtgcta-aaaaaaaaaaaaaga----------aaaaaccgtaccacatgtcaa
aagaggcagatcgtgcttgacggcgggaggtgttgcatttcttgcttgtcctgagcgtct
gccagtctct----ga--cgagtg----tgttccttccataaacccgatctcaaatcagg
taggaccacccgctgaatttaagcatataattaagcggaggaaaagaaactaacaaggat
tccctcagtagcggcgagcgaagcgggaagagctcaacatgaaaaccgttgggggttcgc
gctttccttcgggttgtatagtaaagtggtgtcgcctgcgtcgcgccgggtataaatccc
gtctgaaatggggtgtcgtagagggttatatccccgtctgtgacccggtgtgcgatgcgt
tacggtgaccttacgatgagtcgggttgcttgggcatgcagccctaagtgggtggtaaac
ttcatctaaagctaaatactagcgagagaccgatagcgtacaagtaccgtaagggaaaga
tgcaaagaactttgaaaagagagttaaacagggcttgaaaccgttaagagtgaagcgaag
gaggtcagtgatcatggggccagatccccaagtgcctgtcata-gggcactgtagcgttc
tggtttcccggggcagcgccggccgggggcgggggttaacagggaggtggttgagggatt
cgttttccgcctgtactcctttatagcttgcaccccggccgagggatccgcgacttgtcg
tgtgagggcgctgtcaaaatggccttctccaccccgtcttgaaacacggaccaaggagtc
taacatactcgcgagtgtttgggtggttaaaacccgaacgcacaatgaaagtaaacggtg
gctctgcgccgctgcttggtgggaaccctgcttttataatgggggcactatcacccgac
cccgatccctttggggtggttggttttgaggtagagcgagtatgatgggacccgaaagat

ggtgaactattcctgagtagggcgaagccagaggaaactctggtggaggctcgtagcgat
gctgacgtgcaaatcgctcgtcgaacttgggtataggggcgaaagactaatcgaaccatc
tagtagctggttccctccgaagtttccctcaggatagcgggagcaaatgcaagtcagtat
catccggtaaagcgaatgattagaggcctcgggggagtaactccttcgacctattctcaa
actttaaataggtgaggagccggcaacgttgcttaaaacaatgaacgccgtgcggacaat
gattgctctcagtgggccgattttggtaagcagaactggcgatgcgggatgaaccgaaag
cgacgctaaggtgccaaactcccagctcattcagacaccagaaaaggtgttgattcatac
agacagcaggacggtggccatggaagtcggaatccgctaaggagtgtgtaacaactcacc
tgccgaatggatcagccctgaaaatggatggcgcttaagcgggggaccgaggcgtcgccg
ttggggtaaatacgaagccccaatgagtaggagggcgtgggggttttgaaacacggtaca
ctcgcatggaagccttgggcgtaagctcgggtggagcgacccctagtgtagatcttggtg
gtagtagcaaatattcaagtgagaaccttgaagactgaagtggggaagggttccacgcga
acagtgcttggacgtgggttagtcggtcctaagagatggggaaactccgtattcaagtgc
gcttttgtagcgcagtccatcgaaagggaagcgggttaacattcccgcaccaggatgtgg
aaactgtggtttaccgcaaactgcggcaacgcaactgaacctggggacgcccgtaggagc
tccgagcagggatctctttcctgtttaacgcgcccaacccccctggaatcggtttgtccgg
agatagggggcgaaaggccggaaaagcaccgcacgtcttgtggtgttcggaacgcttgta
atggcccttgaaaacccaggggaaagcagccgtcttacac--ttgtgtgagacgaggaat
tctcacacctggccgtacccatcaccgcatcaggtccccaaggtgaacagcctctagtca
atagaataatgtagataagggaagtcggcaaaatagatccgtaactttgggaaaaggatt
ggctctaatgttcgggcttgtcgggacgcgcgggcgtagatgccaatgagcggcagacgt
ctagcctctttggaggctggattaacgctgcttgtgggaagaaaccgtgtcgatggaggc
aggaacgcttatcgatcttgtctcttcccgcttgcgatcaacgaacggattagaactggt
acggacaaggggaatccgactgtttaattaaaacaaagcattgcgacggccggaaacggt
gttggcgcaatgtgatttctgcccagtgctctgaatgtcaaagtgaagaaattcaaccaa
gcgcgggtaaacggcgggagtaactatgactctcttaagg--------------------
------------------------------------------------------------
------------------------------------------------------------
------------------------------------------------------------
------------------------------------------------------------
------------------------------------------tagcc-------------
------------------------------------------------------------
------------------------------------------------------------
----------------------------------------------------------aa
atgcctcgtcatctagacactacttggatgagatagtctgggttaaaagcttatgcaggc
tagtaccctctgggtaaagaacaa------atatacgactccacaggcgcatc-------

-------------------------acatttcacaccaagcgaaac--------------

------------------------------aatcttgcaatgc-----------------

----ggcgtgtgctggcaca----------------------------------------

-------------------------------------------cgcggacccactg----

-------------ctgctgctcct---------cgatgaagacagttccc----------

------------------------------------------------------------

-----------------------gcacatccg----------cggcgggagatgtttttg

tcgcggctttcgaagcataag---------------------------------tggtcc

gggttgcagcaccacgcgttgccgcagaggtgagaggtgtcat-------cggcatcaca

--------------------------ctcctttgtgacgagcgt---ctcg-tggatgaa

cttgtgcagggtgacctt----------gcgtcctttgtagttcgtg----------cgg

aagtagccgttggcggcttggtcaacgccaattcccatcggcattttgcacttgtagatg

ttcttgacgcctccgatcgagtagacgcggttcgccatgaattcgtcgacgaacgcctcc

ggcgtgacgtccg------gcgtgtattggt-------------------tgctcttccg

cg----acatcgtgttggcgtgcagcg--------ggtgcgacactccttgtacagggga

accctccaaacagaaaaaggcgcgtggtaccacccctgcgaggaaacgcgcgggcgggat

cccgggtaatgaccggagccgaaggtaaaaacccacgcgaccgatggggcaaccctgtgg

cgagaggctaagttgcgag-cgtc-ttcgcgcgcgatatgcctccgtcgcagagactgaa

cgggagtgggctcgagatg-----ctaccactggggacacgtgatgttccttgtgccaag

taccgggcttaagatacagtccaggccccaggtatgg-----aaaacctggggaactacc

ctgaattagtgacgcgcatgaatggattaacgagattcccactgtccctatctactatct

agcgaaaccacagccaagggaacgggcttggaagaatcagcggggaaagaagaccctgtt

gagcttgactctagtccgactttgtgagatgagttgggaggtgtagtctaagagggaggg

gcaactcgaccgtgaaatacctctactcccaagcttgttttacttactccggcgagcagg

gatgggtggtaacacccttttctaggattaacacctcttcgggggtggacctgcgcgga

ggacattgtcaggtggggagtttggctggggcggcacatctgttaaaagataacgcaggt

gtcctaagatgagctcagtgagaacagaaatctcacgtagaacaaaagggtaaaagctca

tttgattttgattttcagtacgaatacaaactgtgaaagcatggcctatcgatcctttg

accttcaaggtttgaagcaagaggtgtcagaaaagttaacacgcgttgtggtagctttta

agagtggactcaaagtgttccgctagtccccccctcggaccgccacttacaaaaggcctgg

ggatgggggcgacactgtcaaattgcggggactccctaaagcctgtctctaccaagccag

cgcggcgaaagcccctggtggccggggtaatgacctagggtatggtaaaagcgagacagg

atg-gaacaatgggcaatccgcagccaagtcctaagg----ctttttgaaaagcggctat

ggatgcagttcaacgactagatggcagtgggctgagagagcccaccgcggcacccctttcg

gcttaagatatagtcttgcccacgtgaaagcgtgcctgcagatgaacgccaccatgtatg

gccggagctgcaggagaccgc--acttgatgtgcagtcgggaat-cgacagtaccacagg

gatgaac-----------------------------------------------------
------------------------------------------------------------
------------------------------------------------------------
------------------------------------------------------------
------------------------------------------------------------
-------------------------------------------------aaaaacac---
------------------------------------------------------------
------------------------------------------------------------
--acaatta---------------------------------------------------
----------------------------------------atttgtggcc---atcatac
----------------------------------------agtcgta-------------
------------------------------------------------------------
------------------------------------------------------------
------------------------------------------------------------
------------tgatattatcgtcccaggcgggtgcgcctctctcacccgtctagtccg
gccttggtgcgcaaaaaatgcttgcgtggatgcgccttggttaaagaaaatgctcctagt
agttttacaaggacacttcttgtagggcaacacatctggatgcgaagaactcccggaaaa
gacaacggtaccaagtgactggggtaacgccccattcatgaaggcaaataatcaaaaaga
ggtgccgggtaagaacccgttgtatagggacaatccgcagctgacttggctccctg--cc
catccccatgggcatagcttagcggttcaacgctcggtaagatgtgggttggcgcgctgt
ccgacagcgcactggcttaaggtacgggctactcccaccggagacggtgtctttgtctac
ctcgggtaacacgcctcgggagaagcaaagagggtgcaatgg-aaaaaaactggttgcgg
ggaaatccctaccagagtcacctggtacaaaggaactggcttgtggcagccaagcgttca
tagcgacgttgctttttgatccttcgatgtcggctcttcctatcattgcgaagcagaatt
cgccaagtgttggattgttcacccactaatagggaacgtgagctgggtagccacttta--
-----tgcccaacagctggatagctgggcttctgtctta-ggccgggacttctcacgacc
cgtgccttggtcaccc-aaaaaagggcgaccgcagagctcaggatggagcttctggctta
actagtagaggacagatagagatcgtcaaattgcgggaaaccccttagagcctcgagcta
ccaagtgcgtgtcgaaaggcatagacgcagccagcgatggctgtcgcatggcttga-ttc
catcaagataggtaaaaatgctcgaggattgggcaatccgcagccaagcccctaaggt--
---ccccccgcgggctatggggaaggttcaacgactgtaaggcgtaccgtgcaagcggaa
gatacagtctagccccgctggaaagcagcgccaaagtaagtaggt-tgcagtgggtctgt
cctc
>Form_P_Japan
gcctgttttgatatgtgttttgacagtcatacgggtcatggccgagggatggtgggtgag
cttcatagcgcaatcccccgtcaggagcatgttttcatagcattttgagcatactagggt

ggtctcccatgtagctcgctctctcggctcaggaccttccgtatggggcactttttgacg
agctcctttggggaggagcatgttctggcatatgaagcgttggtcacgagcttcgtagcc
cg---acacataccaaatcagtcttcatatgtctttctggcgggggtcggtcgtatgacc
acgcccttaccttgagcatgtttttcgtatgataccatacctgacgggcagcgccgctgt
tcagtcgtcggccgtacctcctatgcgcgttttctgattcgaacgtgtgacatgcaagcg
catcatatcagaagcatgtttttttcatatgagcatgttggtaggcaggctcttcagtcgt
ctacccataaatgcactttgaaagacgtttttttggtactgaacggctttgtggtcagacg
ttctcgtgagcctattctcgccctctgatacgctggatgctgcggtattacccacagcag
acat---------ccaggagctccctctgcggataag--catctgtagaagggggtttct
ctcggtgac-----gcactggctgggaaaacggagtgctatccaaaagggtgaaacctct
tggggatgtactcccataccagcataatatgctgagaggaattgactcaatgagtgaggt
cctatactgcaacctcttttggatatgtaggcggtctgtacccagactgtcctactggct
gatttcccttggtcc--------cacgcgtcgctggctgaatgggctagtccttatgaag
tgccgcgttcatcaaaagaaccggtcttcatacagcgacctcattc-tatactgccagct
ggatgggctggtccttatggagcgccgcattcttcgaagaatcggtttctatagagcgac
cccatttacatataaggaaccg-----ttacatagacttgccatttggctgatggctctg
acagtcgctgcctcttcggagttctgtgtaaacgtggcgccatttttgagtgggccagtc
cttatggagtgtccgtgtcttctatgacagggtcccccatacagcgacctcactcagaac
cgtatgggcacgatggtgaaagaccatgcagatagtccgccttccagagggggaagtagt
atatggggaatgctggcgaagcatgaggt--acatggttcatggacccgttgggccatgc
ctgtgccatcttctcgagtcaaaccggtccgtgtcgcttccttatctcttcctttcgcga
cgacaagctcttgagctggtcgtgggtcaggacgctgtggtatctgtttgtctgtagtgt
gggct-tccaccagaacagcgtatcagcacacgtcgtctgggcatatgatgtttttgctt
tgtggtcccgggtttggtagtatggttccatgtcggtccgctcacctcccaaccataatg
tctttcacatcatcttcacagtacgtcttgtcagcataccattctgtatagaacagtgta
tcagcacacgtcgtctgagcatagggtatttttgctttgtggtcccgggtttggtagtat
ggttccatgtcggtccgctcacctcccaaccataatgtcatttctatcatctgcacagta
cgtcttgtcagaataccattctctatggcagtcgatagactaggccttatgccgcagcag
tctctgatcactgtgcgagtgcatatatctctcaatgtgttcatggctagcatgggtctt
gagtgagatgtgccagagagatacgccaccagtctatggaaaggtgctttttgtatgaga
agagtacatgtatgaagagcatgttctctcctcttatgatggtaccgcgggtacgggctg
ctcgcattatgcttggcaggtgtccactgtgcatgcaggattgctgaacccacttgtatg
gccatgagaatggtcattgggtgaggttcacttctccatcgctttgtgtgtgttagatgt
ttgggtggcttcggccgcatggaaattctcggtctgtaactcatcagggagggtttccat
gtgagta--ttctccagtcgaccacacacccttgggcgtcccgctagacaaaatgtaact
ctcgatccgacctagcctgtggagtgtct-ttccgtatggcgtgaaagggcccagactgg

atgttttcccttttggacaccagtctgt---------tatatagatcatctacagggagc

aaaggacaggcatgcaagtgccacaaaggcagggaggccttcctgctactgtgtgtcctt

cttgacgatcacgggggagtatatagaaacatttgaaggagatcc-ttttccacaaggcg

gtgccctcgcgcaaaggtgttgctctgcacgcatctgtatgatgatagttgctcgctcct

atgcgtccacttgtgaaaataccttctctctttgtctgtttctatgacgccccttctgct

ttgtcatagagcatactcaggtacagaaagtgccgtgacttccttggcctgctgtcaaag

aaatagcctggtgctatagaggactc-ttcctgggcgccgcctggtgaagaggtcctatc

aggctctggagtgtgcgtatgcctatgttcatcatagctcatgtgcacgcgc-ggaaaag

catcgacgtggaatggggaagaacaccattctgaatgagccagttcttcctctgctcccc

ttgcttgacgctgtatgggaatcgctttgccttgggggtacaccttccattccatgacaa

agcattcgatctatgtcccgaggggtgattctacatgaagaacccgaaacttgtctgagt

gagaggcgaaggtagtccttcatttt-atgcgggatccgccctcacactgtagtcggcac

agtcccttgtttagggactgcgcagctcaatcttcgactggagaggaaaaaaacaaaaca

caagtatgacacgcacatacgtgttttctaggccattgacctgttggctgcctcgcggtg

gttggcatgatgccccaagtgggggggggggaaatcaatcagtatggttgacaaaatcca

gtggggaaaaaaggcgcgctgtcgcacacccttggtcacggcctcaaaaagtcgacggcc

ctcactggcccacccctgcgactcctgttcgcttttgacaagacacacctgcaata-tgg

aggaaatgcctgagcagctgtgccgcagcagctgtggacgcgtagcgatatgcatcccta

ttgtttgcgacttggtacaaagaacgcgtgccgggagacggattgccctttgacttttcc

acccggcccgcggctctgccctttacgtacccttatttacctggttgatcctgccagtag

tcatatgcttgtctcaaagactaagccatgcatgtctaagtataaacacttttatactcg

tgaaactgcgaatggctcattaaaacagttatagttcctttgggaacaacactacttgga

taaccgtagtaattctagagctaatacatgcctcaacgcccgactcacgaagggtggtat

ttattggataaaaaaccatcgtgctctctccagagcgctttgagatgattcacaataact

tgtcggatcgcatggcctcgtgctggcgactcctcattcaaatttctgccctatcaactt

tcgatggtagagtattggtctaccatggtgtcgacgggtgacggggaattagggttcgat

tccggagagggagcctgagaaacggctaccacatccaaggaaggcagcaggcgcgcaaat

tacccaatcccgactcggggaggtagtgacaaaaaataacaatagggggccctttgggtc

ttctaattggaatgagaacaatttaaatcccttatcgaggatccattggagggcaagtct

ggtgaacgta----------------------------------tgcatca---------

--aagccctcgat----aaaaaaaaacagcgccagatgtct-------------------

------------------------------------------------------------

------------------------------------------------------------

------------------------------------------------------------

------------------------------------------------------------

------------------------------------------------------------

-------ctctctggaagcgcacca-----------------------------------

---tgcat-ggctttgttcctctc-----------------------------tgcatgg

ccg-------------------------------------------tgtggaaggcgagt

aggcaaacgatgggaaaccaggcttggaatagtaaccaaccacagagcccccatcataca

cttctaaattgccggggagtgcccgcgaggccttgactaccgcggtccgccgagaggtga

gagcgagcaatcgttcaggacgcagcagtgtacgtagtggtacatggatggtaaaaacgt

caaggatagggttgatccgcagggaagcgcgagtgccgccatg---------gtcggtgg

gcaaaggcgaaccttcagagactctaatggagtgggcgcaagcttaagggagagtccaat

ccactgggaaaccaggtccacagcagtaacggggtgatgctcgccccagaggactgtgga

aggcaccaccaccccaagagggggtgggtgttgtcgggtagcaaatggccagcagccgcg

gtaattccagctccaatagcgtatattaaagttgttgcagttaaaacgctcgtagtcgga

cctcggagcgcccccacgggccgggcttttgccttgttgccagtgggagtttgtctcctt

ttgtcgggacgcggtgtgcagggcctttactgtctctgcgcgtttacctgcgggccgacc

gtttactgtgaagaaagtaaagtgttcaaagcaggccaatgccttgaatatgtgagcatg

gaataatagaataggacttgggctctattttgttggtttccagtgaccaagtaatgatta

atagggacggttgggggcattcgtatttcattgtcagaggtgaaattcttggattgatgg

aagacgtacaactgcgaaagcatctgccatggatgttttcattgatcaagaacgaaagtt

aggggatcgaagacgatcagataccgtcgtagtcttaaccataaacgatgccgactgggg

attggcgggggcactattatatgactctgtcagcaccctgagggaaaccaaagtctttgg

gttctgggggggagtatggtcgcaaggctgaaacttaaaggaattgacggaagggcaccac

caggagtggagcctgcggcttaatttgactcaacacgggaaaacttaccaggtccggaca

gaagaatgattgacagactgaaagctctttcttgatttttttggttggtggtgcatggccg

ttcttagttggtggagtgatttgtctggttaattccgttaacgaacgagacctcgtcctg

ctaaataggtgcgcgcatgccagcactgcgttcttaccttcttagagggactatgcgcgt

ctagcgtatggaagattgaggcaataacaggtctgtgatgcccttagatgttctgggccg

cacgcgcgctacactgatgcattcaacgagtttcacttgatagtcctgggtcggaaggct

cgggtaatcttttgaaagtgcatcgtgctggggatagatcattgcaattattgatcttca

acgaggaattccttgtaggtgcaggtcatcagcctgcgccgaatacgtccctgccctttg

tacacaccgcccgtcgctcctaccgattgaatggtccgatgaaatgtccggatcgcggct

agtgtgatggactcgtccagagcggtagccgtgagaagctcattaaatcttatcatttag

aggaaggagaagtcgtaacaaggtttccgtattttgtgggtggataccgattcaataaca

tccgccgtttttctcaatgtactatatccctagtctgcttgcctagttcttggagtcttg

tgcgatgtgtgctattacaggcatgccgggtcatgaagacagacaaggttgggatcttca

cggaagtacgcgcaataggagcgcgtcttgttcaacgcacccgactcaagagtcatatgc

gagggacggacgcagcgcttgtcgccgcacagatggga----------------------

------------------------------------------------------------

------------------------------------------------------------
------------------------------------cgcgccatatcggagacaggggga
cgtgccattgtgatgatgatgatgatgttgttgggtggtccttccgtaggaaaacgtaca
cgaagcctttgcggcccgaaagggtgcacagtgcgcgactataaacactgattctgtggt
caacgcaagcagcggttctgtggaagaacagctgcgactttctcaaattgcgggaacacc
ctaaagtcaacaaacaccgcggccggacgtg-aaagcgtcaggcgcagcagtcggggtag
cgccctggtggaaggtaatagctttgttgaatggcacaatgggcaatccgcagccaagcc
cctaaggtcccccgctgggctatggggaaggttcacagactgtaagggagggggtttctt
ccatga--aagaagcttaagagacagtcggtcccctgcgagagcagtgtcgcatgtgagg
aaggcacccatgaaaagtgggtgctggagagccatgtgagccctctatgagggtgggaag
ttttccga---tacaatgagaaacaaacacgttccgtaggtgaacctgcggaaggatcat
tcacaaactattgacacaaacacacgcgaaccaaatcgtccacaca---ggtgccgatga
aagagagaatctgtttgtcgccttttggggtat----acaagcactctttt-ccatcgcc
tctgtgcagggcgtaacttcccattgagaggatgtgagggcaccacaggaagcttttcca
caggaagtcgccatccttctccctccacgacggcggcgcttcacaggcttggcagttttt
ttttttttttattgccttccagggaggatgccgccaatggagccacatag-acatttaca
tcatcatatgcccgtttttctttcttaaccgcttgccacagtttctcctgtgaggagctt
gtgggaagaccgtctccatacaataacaaaagatacaactcttagcggtggatatcttgg
ttctcgcaacgatgaagaacgcagccaactgcgatacctagtgtgaattgcaggactccg
tgaatcattaagtctttgaacgcaagttgcgctcatctccatgaggaggtgagcatgcct
gtttgagtgtcgatgggaaaaacaatac-ataacaacaatcggctctcttcccttcctgc
cttccccggcaaaaaaggtagtgcatgccggagatgggcctggtcgcccct---tcatga
aagaggcgtcgtccgaaacatgtgtcgttgcccgctggaggtcatacagcatgagtttca
aaaacaccatggctctcaaggcagagtaaatcgtcttgagacggtct-gcattgattcac
tttcaacccaggtggcgttggttacttgccat-aagcctctctggggtttctttgcttgg
acctctcttgtcagttcttctgacgtagagacaggtgcgcggtctcatgggcgttgccct
ctggaacgtgctaaaaaaaacaaaaca---aacaacaacaacaaccgtatcacacgtcaa
gagggtcggatcgtacttgacggcgggagatcctttgtttctcgcttgtcctgagcgtct
gccaagccttctttga--cgagtgtgtttgttccttccataaacccgatctcacatcagg
taggaccacccgctgaatttaagcatataattaagcggaggaaaagaaactaacaaggat
tccctcagtagcggcgagcgaagcgggaagagctcaacatgaaaaccgtaggggggttcgc
gctcaccttcgggttgtatagtaaagtggtgtcgcctgcgtcgcgccgggtataaatccc
gtttgaaatggggtgtcgtagagggttataaccccgtctgtgacccggtgtgcgatgcgt
tacggtgaccttacgatgagtcgggttgcttgggcatgcagccctaagtgggtggtaaac
ttcatctaaagctaaatactagcgagagaccgatagcgtacaagtaccgtaagggaaaga
tgcaaagaactttgaaaagagagttaaacagggcttgaaaccgttaagagtgaagcgaag

gaggtcagtgatcatggggccagatccccacgtgcccgtcata-gggcaccgtagcgttc
tgg-ttcccggggcagcgccggccgggggcgggtgtcaacagggaggtggttgagggatt
aactttccgcctgtactcctttctagcttgcaccccggccgagggatccgcgacttgtcg
tgtgagggcgctgtcaaaatggccttctccaccccgtcttgaaacacggaccaaggagtc
taacatactcgcgagtgtttgggtggttaaaacccgaacgcacaatgaaagtaaacggtg
gctctgcgctgctgcttggtgggaaccctgctcttataacggggggcaccatcacccgac
cccgatccctttggggtggttggttttgaggcagagcgagtatgatgggacccgaaagat
ggtgaactattcctgagtagggcgaagccagaggaaactctggtggaggctcgtagcgat
gctgacgtgcaaatcgctcgtcgaacttgggtataggggcgaaagactaatcgaaccatc
tagtagctggttccctccgaagtttccctcaggatagcgggagcaaatgcaagtcagtat
catccggtaaagcgaatgattagaggcctcgggggagtaactccttcgacctattctcaa
actttaaataggtgaggagccggcaacgttgcttaaaacaatgaacgccgtgcggacaat
gattgctctcagtgggccgattttggtaagcagaactggcgatgcgggatgaaccgaaag
cgacgctaaggtgccaaactcccagctcattcagacaccagaaaaggtgttgattcatac
agacagcaggacggtggccatggaagtcggaatccgctaaggagtgtgtaacaactcacc
tgccgaatggatcagccctgaaaatggatggcgcttaagcgggggaccgaggcgtcgccg
ttggggtaaatacgaagcctcaatgagtaggagggcgtgggggttttgaaacacggtaca
cacgcatggaagccttgggcgtaagctcgggtggagcgacccctagtgtagatcttggtg
gtagtagcaaatattcaagtgagaaccttgaagactgaagtggggaagggttccacgcga
acagtgcttggacgtgggttagtcggtcctaagagatggggaaactccgtattcaagtgc
gctcttccagcgcagtccatcgaaagggaagcgggttaacattcccgcaccaggatgtgg
aaactgtgg-ttaccgcaaactgcggcaacgcaactgaacttggggacgcccgtaggagc
tccgagcagggatctctttcctgtttaacgcgcccaaccccctggaatcggtttgtccgg
agatagggggcgaaaggccggaaaagcaccgcacgtcttgtggtgttcggaacgcttgta
atggcccttgaaaacccaggggaaagcagccgtcttacactttttgtgggacgaggaat
tctcacacctggccgtacccatcaccgcatcaggtccccaaggtgaacagcctctagtca
atagaataatgtagataagggaagtcggcaaaatagatccgtaactttgggaaaaggatt
ggctctaatgttcgggcttgctgggacgcgcgggcgtagacgccaatgagcggcagacgt
ctagcctcctcggaggctggattgacgctgcttgtgggaagaaaccgtgtcgatggaggc
aggagcgcttgccgttcttgtctcttcctgcttgcgatcaacgaacggattagaactggt
acggacaaggggaatccgactgtttaattaaaacaaagcattgcgacggccggaaacggt
gttggcgcaatgtgatttctgcccagtgctctgaatgtcaaagtgaagaaattcaaccaa
gcgcgggtaaacggcgggagtaactatgactctcttaact-------------------
------------------------------------------------------------
------------------------------------------------------------
------------------------------------------------------------

------------------------------------------------------------
------------------------------------------------------------
------------------------------------------------------------
------------------------------------------------------------
-----------------------------------------------aca----------
------------------------------------------------------------
------------------------------------------------------------
-----------------------------tggacatcttttgaga---------------
------------------------------gagataggcgggctgaaaaggt------ta
tagccgtctagtctcgc-------------------------------------------
---------------------------------tctttcatg-agcggggcgacacacct
ggttgcgggaaccctgcccacctctctatcag-aagtgggaaccagtccattcctactcc
tgcttgtgaggcatatagtaaaaagtagcagcaggaggatctcggcgagaggccgaaccc
gaaaggaataacgtgggctgggaggcaatccgcaggcgagtccgtaactatatgattatc
gacgggccgtctcaacgactggtaaggtgtggg-------tttgtcctcacgc-gggaca
ggcttaaggtacagtctaatcccacccgccaagggtgcttggg---tg------------
--------------------taatccctcttacatagagggcactagccca-ggggctgg
gacgctgggaggaa----------------atgccccgtccctatagccggtagaaatga
aggtagccaaatgcctcgtcatctaaactaattgg-at----------------------
-----------------------gagatagtc----------tgagtgaaaagcttatgc
ag-----------------gctagtaccggta-------aaaaaacgagaggctctgcca
ct----ctgac------------------------ggtgcgacactccttgtacagggaa
accctcc--gcatggaaaggcacgtggtaccaccctcgcgaggaaactcgtgagcgggat
cccgggtaatgaccggagccgatggtaaaaacccacgtgaccgatggggtgaccctgtgg
cgagaagctacgttgtga-----cttgtgtgcacaatatgcttccgtcgcagagactgaa
cgggagtgggctccgtcga---------cggccg----gccagccgtcttcca-------
---ggagcttaagatacagtccggcccacagggatgg---cttaaacctgtggaacaa-g
cagaattagtgacgcgcatgaatggattaacgagattcccactgtccctatctactatct
agcgaaaccacagccaagggaacgggcttggaagaatcagcggggaaagaagaccctgtt
gagcttgactctagtccgactttgtgagatgagttgggaggtgtagtctaagagggaggg
gcaactcgaccgtgaaatacctctactcccaagcttgttttacttactccggcgagcagg
aatgggtggcaacacccttttctaggattaacacctcttcgggggtggacctgcgcgga
ggacattgtcaggtggggagtttggctggggcggcacatctgttaaaagataacgcaggt
gtcctaagatgagctcagtgagaacagaaatctcacgtagaacaaaagggtaaaagctca
tttgattttgattttcagtacgaatacaaactgtgaaagcatggcctatcgatccttttg
accttcaaggtttgaagcaagaggtgtcagaaaagttagcacgcgttgtggtagctttta

agagtggactcaaagtgttccgctagtccccccctcggaccgccacttacaaaaaggacag
aggagggggcgacactgtcaaattgcggggactccctaaagcctgtctctaccaagccag
-gcggcgaaagctcctggtggccggggtaatgacctagggtatggtaaaagcgagacagg
atgagaacaatgggcaatccgcagccaagtcctacggctcttttcttggagaagagctat
ggatgcagttcaacgactagatggcagtgggctgagggagcctacggcggcacccctttcg
gcttaagatatagtcttgcccacgtgaaagcgtgcctgcagatgaacgccaccatgtatg
gcaggagctgcaggaggccgctaaacataccagcggctgggaacataacagtaccacagg
gatgaac-----------------------------------------------------
------------------------------------------------------------
------------------------------------------------------------
------------------------------------------------------------
------------------------------------------------------------
-------------------------------------------------aaaaacac---
------------------------------------------------------------
------------------------------------------------------------
--acaattg---------------------------------------------------
------------------------------------------ttgtggca---atcatac
----------------------------------------agtcgta-------------
------------------------------------------------------------
------------------------------------------------------------
------------------------------------------------------------
------------tgatattatcgtcccaggcgggtgcgcctctctcacccgtctagtccg
gccttggtgcgcaaaaaatgcttgcgtggatgcgccttggttaaggaaaatgctcctcgt
agttttacaaggacatttcttgtagggcaacacatctggatgcgaagaactcccgg-aaa
gacaacggtaccaagtgactggggtaacgccccattcataaaggcaatgaat---aaaga
ggtgccgggtaataatctgttgtatagggacaatccgcagctgacttggctccctgcccc
cat--cggtgggcatagcttagcggttcaacgctcggtaagatgtgggttggcgcgctgt
ccaacagcgcactggcttaaggtacgggctactcccaccggagacggtgtctttgtctac
ctcgggtaacacgcctcgggagaagcaaagagggtgcaagta--aaaaaactggtagtgg
ggaaatccctaccagagtcacctggtacaaaggaactggcttgtggcagccaagcgttca
tagcgacgttgctttttgatccttcgatgtcggctcttcctatcattgcgaagcagaatt
cgccaagtgttggattgttcacccactaatagggaacgtgagctgggtaggcattttatg
ccctatgcccaacagctggatagctgggcttctgtccta-ggccgggacttctcacgacc
cgtgccttggtcaccc-aaaaaagggcgaccacagagctcaggatggatcttctggctca
actagtagaggacaggtagagatcgtcaaattgcgggaaaacccttagagcctcgagcta
ccaagtgcgtgtcgaaaggcatagacacagccagcaatggctgtcgcatggcttga-ttc

catcaagataggtaaaaatgctcgaggattgggcaatccgcagccaagcccctaaggt--
--ccccccgtggggctatggggaaggttcaacgactgtaaggcgtaccgtgcaagcggaa
gatacagtctagccccgctggaaagcagcgccaaagtaagtaggt-tgcagtggatctgt
cctc
>Form_Q_China
gcctgttttgatatgtgttttgacagtcatacgggtcatggccgagggatggtgggtgag
cttcatagcgcaatcccccgtcaggagcatgttttcatagcattttgagcatactagggt
ggtctcccatgtagctcgctctctcggctcaggaccttccgtatggggcactttttgacg
agctcctttggggaggagcatgttctggcatatgaagcgttggtcacgagcttcgtagcc
cg---acacataccaaatcagtcttcatatgtctttctggcgggggtcggtcgtatgacc
acgcccttaccttgagcatgtttttcgtatgataccatacctgacgggcagcgccgctgt
tcagtcgtcggccgtacctcctatgcgcgttttctgattcgaacgtgtgacatgcaagcg
catcatatcagaagcatgtttttttcatatgagcatgttggtaggcaggctcttcagtcgt
ctacccataaatgcactttgaaagacgtttttttggtactgaacggctttgtggtcagacg
ttctcgtgagcctattctcgccctctgatacgctggatgctgcggtattacccacagcag
acat---------ccaggagctccctctgcggataag--catctgtagaagggggtttct
ctcggtgac-----gcactggctgggaaaacggagtgctatccaaaagggtgaaacctct
tggggatgtactcccataccagcataatatgctgagaggaattgactcaatgagtgaggt
cctatactgcaacctcttttggatatgtaggcggtctgtacccagactgtcctactggct
gatttcccttggtcc--------cacgcgtcgctggctgaatgggctagtccttatgaag
tgccgcgttcatcaaaagaaccggtcttcatacagcgacctcattc-tatactgccagct
ggatgggctggtccttatggagcgccgcattcttcgaagaatcggtttctatagagcgac
cccatttacatataaggaaccg-----ttacatagacttgccatttggctgatggctctg
acagtcgctgcctcttcggagttctgtgtaaacgtggcgccatttttgagtgggccagtc
cttatggagtgtccgtgtcttctatgacagggtcccccatacagcgacctcactcagaac
cgtatgggcacgatggtgaaagaccatgcagatagtccgccttccagaggggggaagtagt
atatggggaatgctggcgaagcatgaggt--acatggttcatggacccgttgggccatgc
ctgtgccatcttctcgagtcaaaccggtccgtgtcgcttccttatctcttcctttcgcga
cgacaagctcttgagctggtcgtgggtcaggacgctgtggtatctgtttgtctgtagtgt
gggct-tccaccagaacagcgtatcagcacacgtcgtctgggcatatgatgtttttgctt
tgtggtcccgggtttggtagtatggttccatgtcggtccgctcacctcccaaccataatg
tctttcacatcatcttcacagtacgtcttgtcagcataccattctgtatagaacagtgta
tcagcacacgtcgtctgagcatagggtatttttgctttgtggtcccgggtttggtagtat
ggttccatgtcggtccgctcacctcccaaccataatgtcatttctatcatctgcacagta
cgtcttgtcagaataccattctctatggcagtcgatagactaggccttatgccgcagcag
tctctgatcactgtgcgagtgcatatatctctcaatgtgttcatggctagcatgggtctt

gagtgagatgtgccagagagatacgccaccagtctatggaaaggtgctttttgtatgaga
agagtacatgtatgaagagcatgttctctcctcttatgatggtaccgcgggtacgggctg
ctcgcattatgcttggcaggtgtccactgtgcatgcaggattgctgaacccacttgtatg
gccatgagaatggtcattgggtgaggttcacttctccatcgctttgtgtgtgttagatgt
ttgggtggcttcggccgcatggaaattctcggtctgtaactcatcagggagggtttccat
gtgagta--ttctccagtcgaccacacacccttgggcgtcccgctagacaaaatgtaact
ctcgatccgacctagcctgtggagtgtct-ttccgtatggcgtgaaagggcccagactgg
atgttttcccttttggacaccagtctgt---------tatatagatcatctacagggagc
aaaggacaggcatgcaagtgccacaaaggcagggaggccttcctgctactgtgtgtcctt
cttgacgatcacgggggagtatatagaaacatttgaaggagatcc-ttttccacaaggcg
gtgccctcgcgcaaaggtgttgctctgcacgcatctgtatgatgatagttgctcgctcct
atgcgtccacttgtgaaaataccttctctctttgtctgtttctatgacgccccttctgct
ttgtcatagagcatactcaggtacagaaagtgccgtgacttccttggcctgctgtcaaag
aaatagcctggtgctatagaggactc-ttcctgggcgccgcctggtgaagaggtcctatc
aggctctggagtgtgcgtatgcctatgttcatcatagctcatgtgcacgcgc-ggaaaag
catcgacgtggaatggggaagaacaccattctgaatgagccagttcttcctctgctcccc
ttgcttgacgctgtatgggaatcgctttgccttgggggtacaccttccattccatgacaa
agcattcgatctatgtcccgaggggtgattctacatgaagaacccgaaacttgtctgagt
gagaggcgaaggtagtccttcatttt-atgcgggatccgccctcacactgtagtcggcac
agtcccttgtttagggactgcgcagctcaatcttcgactggagaggaaaaaaacaaaaca
caagtatgacacgcacatacgtgttttctaggccattgacctgttggctgcctcgcggtg
gttggcatgatgccccaagtgggggggggggaaatcaatcagtatggttgacaaaatcca
gtggggaaaaaaggcgcgctgtcgcacacccttggtcacggcctcaaaaagtcgacggcc
ctcactggcccacccctgcgactcctgttcgcttttgacaagacacacctgcaata-tgg
aggaaatgcctgagcagctgtgccgcagcagctgtggacgcgtagcgatatgcatcccta
ttgtttgcgacttggtacaaagaacgcgtgccgggagacggattgccctttgacttttcc
acccggcccgcggctctgccctttacgtacccttatttacctggttgatcctgccagtag
tcatatgcttgtctcaaagactaagccatgcatgtctaagtataaacacttttatactcg
tgaaactgcgaatggctcattaaaacagttatagttcctttgggaacaacactacttgga
taaccgtagtaattctagagctaatacatgcctcaacgcccgactcacgaagggtggtat
ttattggataaaaaaccatcgtgctctctccagagcgctttgagatgattcacaataact
tgtcggatcgcatggcctcgtgctggcgactcctcattcaaatttctgccctatcaactt
tcgatggtagagtattggtctaccatggtgtcgacgggtgacggggaattagggttcgat
tccggagagggagcctgagaaacggctaccacatccaaggaaggcagcaggcgcgcaaat
tacccaatcccgactcggggaggtagtgacaaaaaataacaatagggggccctttgggtc
ttctaattggaatgagaacaatttaaatcccttatcgaggatccattggagggcaagtct

ggtgaacctagttccacccgctcctcgtgttgatcgcacgcaggcgcaccgtgtgaagca
tcaagccctcgtt----aaaataaaacagcgccagatgtctcgtccggaaggaggtccct
caggtagagcacacaacaaaagcggccagtcaactctgggcgtgtgtgatgctgcggctg
aagtgcacatagatcggtgcttgcatgctgtgacaaaaagtggggtcgaggtgctgaaat
tgtccagaacgatgtgtctcctacaagaaacacctcgcgatggacgctacacgcacggtc
aaagaaagcacttctatgtttatcaggcggtggcttatgccaggtttggtagacaggaaa
tgcttaaagtgaccacctcaaagatcggcacagacctgaccatatcgcatgtgtgtggca
cccatcactgcctagcagcgcatcacttaatgttagaaccaaagcatgtgaatgatgatc
gcgtgcac-tgccatgtttttctcgggcgcgtttctgacgcttcagagaggcatgcgttg
gcagagcgtcattgcccacatttaccgcggtgctattctgtttgactgctaacggacaag
aggcaaaagatgggaatc---gtttataa--------aaccattgagcccccatcataca
cttctaaattgccggggagtgcccgcgaggccttgactaccgcggtccgccgagaggtga
gagcgagcaatcgttcaggacgcagcagtgtacgtagtggtacatggatggtaaaaacgt
caaggatagggttgatccgcagggaagcgcgagtgccgccatg---------gtcggtgg
gcaaaggcgaaccttcagagactctaatggagtgggcgcaagcttaagggagagtccaat
ccactgggaaaccaggtccacagcagtaacggggtgatgctcgccccagaggactgtgga
aggcaccaccaccccaagaggggggtgggtgttgtcgggtagcaaatggccagcagccgcg
gtaattccagctccaatagcgtatattaaagttgttgcagttaaaacgctcgtagtcgga
cctcggagcgcccccacgggccgggcttttgccttgttgccagtgggagtttgtctcctt
ttgtcgggacgcggtgtgcagggcctttactgtctctgcgcgtttacctgcgggccgacc
gtttactgtgaagaaagtaaagtgttcaaagcaggccaatgccttgaatatgtgagcatg
gaataatagaataggacttgggctctattttgttggtttccagtgaccaagtaatgatta
atagggacggttgggggcattcgtatttcattgtcagaggtgaaattcttggattgatgg
aagacgtacaactgcgaaagcatctgccatggatgttttcattgatcaagaacgaaagtt
aggggatcgaagacgatcagataccgtcgtagtcttaaccataaacgatgccgactgggg
attggcgggggcactattatatgactctgtcagcaccctgagggaaaccaaagtctttgg
gttctgggggggagtatggtcgcaaggctgaaacttaaaggaattgacggaagggcaccac
caggagtggagcctgcggcttaatttgactcaacacgggaaaacttaccaggtccggaca
gaagaatgattgacagactgaaagctctttcttgatttttttggttggtggtgcatggccg
ttcttagttggtggagtgatttgtctggttaattccgttaacgaacgagacctcgtcctg
ctaaataggtgcgcgcatgccagcactgcgttcttaccttcttagagggactatgcgcgt
ctagcgtatggaagattgaggcaataacaggtctgtgatgcccttagatgttctgggccg
cacgcgcgctacactgatgcattcaacgagtttcacttgatagtcctgggtcggaaggct
cgggtaatcttttgaaagtgcatcgtgctggggatagatcattgcaattattgatcttca
acgaggaattccttgtaggtgcaggtcatcagcctgcgccgaatacgtccctgccctttg
tacacaccgcccgtcgctcctaccgattgaatggtccgatgaaatgtccggatcgcggct

agtgtgatggactcgtccagagcggtagccgtgagaagctcattaaatcttatcatttag
aggaaggagaagtcgtaacaaggtttccgtattttgtgggtggataccgattcaataaca
tccgccgtttttctcaatgtactatatccctagtctgcttgcctagttcttggagtcttg
tgcgatgtgtgctattacaggcatgccgggtcatgaagacagacaaggttgggatcttca
cggaagtacgcgcaataggagcgcgtcttgttcaacgcacccgactcaagagtcatatgc
gagggacggacgcagcgcttgtcgccgcacagatggga----------------------
------------------------------------------------------------
------------------------------------------------------------
------------------------------------cgcgccatatcggagacaggggga
cgtgccattgtgatgatgatgatgatgttgttgggtggtccttccgtaggaaaacgtaca
cgaagcctttgcggcccgaaagggtgcacagtgcgcgactataaacactgattctgtggt
caacgcaagcagcggttctgtggaagaacagctgcgactttctcaaattgcgggaacacc
ctaaagtcaacaaacaccgcggccggacgtg-aaagcgtcaggcgcagcagtcggggtag
cgccctggtggaaggtaatagctttgttgaatggcacaatgggcaatccgcagccaagcc
cctaaggtcccccgctgggctatggggaaggttcacagactgtaagggagggggtttctt
ccatga--aagaagcttaagagacagtcggtcccctgcgagagcagtgtcgcatgtgagg
aaggcacccatgaaaagtgggtgctggagagccatgtgagccctctatgagggtgggaag
ttttccga---tacaatgagaaacaaacacgttccgtaggtgaacctgcggaaggatcat
tcacaaactattgacacaaacacacgcgaaccaaatcgtccacaca---ggtgccgatga
aagagagaatctgtttgtcgccttttggggtat----acaagcactctttt-ccatcgcc
tctgtgcagggcgtaacttcccattgagaggatgtgagggcaccacaggaagcttttcca
caggaagtcgccatccttctccctccacgacggcggcgcttcacaggcttggcagttttt
ttttttttttattgccttccagggaggatgccgccaatggagccacatag-acatttaca
tcatcatatgcccgtttttctttcttaaccgcttgccacagtttctcctgtgaggagctt
gtgggaagaccgtctccatacaataacaaaagatacaactcttagcggtggatatcttgg
ttctcgcaacgatgaagaacgcagccaactgcgatacctagtgtgaattgcaggactccg
tgaatcattaagtctttgaacgcaagttgcgctcatctccatgaggaggtgagcatgcct
gtttgagtgtcgatgggaaaaacaatac-ataacaacaatcggctctcttcccttcctgc
cttccccggcaaaaaaggtagtgcatgccggagatgggcctggtcgcccct---tcatga
aagaggcgtcgtccgaaacatgtgtcgttgcccgctggaggtcatacagcatgagtttca
aaaacaccatggctctcaaggcagagtaaatcgtcttgagacggtct-gcattgattcac
tttcaacccaggtggcgttggttacttgccat-aagcctctctggggtttctttgcttgg
acctctcttgtcagttcttctgacgtagagacaggtgcgcggtctcatgggcgttgccct
ctggaacgtgctaaaaaaaacaaaaca---aacaacaacaacaaccgtatcacacgtcaa
gagggtcggatcgtacttgacggcgggagatcctttgtttctcgcttgtcctgagcgtct
gccaagccttctttga--cgagtgtgtttgttccttccataaacccgatctcacatcagg

taggaccacccgctgaatttaagcatataattaagcggaggaaaagaaactaacaaggat
tccctcagtagcggcgagcgaagcgggaagagctcaacatgaaaaccgtaggggttcgc
gctcaccttcgggttgtatagtaaagtggtgtcgcctgcgtcgcgccgggtataaatccc
gtttgaaatggggtgtcgtagagggttataaccccgtctgtgacccggtgtgcgatgcgt
tacggtgaccttacgatgagtcgggttgcttgggcatgcagccctaagtgggtggtaaac
ttcatctaaagctaaatactagcgagagaccgatagcgtacaagtaccgtaagggaaaga
tgcaaagaactttgaaaagagagttaaacagggcttgaaaccgttaagagtgaagcgaag
gaggtcagtgatcatggggccagatccccacgtgcccgtcata-gggcaccgtagcgttc
tgg-ttcccggggcagcgccggccgggggcgggtgtcaacagggaggtggttgagggatt
aactttccgcctgtactcctttctagcttgcaccccggccgagggatccgcgacttgtcg
tgtgagggcgctgtcaaaatggccttctccaccccgtcttgaaacacggaccaaggagtc
taacatactcgcgagtgtttgggtggttaaaacccgaacgcacaatgaaagtaaacggtg
gctctgcgctgctgcttggtgggaaccctgctcttataacgggggcaccatcacccgac
cccgatccctttggggtggttggttttgaggcagagcgagtatgatgggacccgaaagat
ggtgaactattcctgagtagggcgaagccagaggaaactctggtggaggctcgtagcgat
gctgacgtgcaaatcgctcgtcgaacttgggtataggggcgaaagactaatcgaaccatc
tagtagctggttccctccgaagtttccctcaggatagcgggagcaaatgcaagtcagtat
catccggtaaagcgaatgattagaggcctcgggggagtaactccttcgacctattctcaa
actttaaataggtgaggagccggcaacgttgcttaaaacaatgaacgccgtgcggacaat
gattgctctcagtgggccgattttggtaagcagaactggcgatgcgggatgaaccgaaag
cgacgctaaggtgccaaactcccagctcattcagacaccagaaaaggtgttgattcatac
agacagcaggacggtggccatggaagtcggaatccgctaaggagtgtgtaacaactcacc
tgccgaatggatcagccctgaaaatggatggcgcttaagcgggggaccgaggcgtcgccg
ttggggtaaatacgaagcctcaatgagtaggagggcgtgggggttttgaaacacggtaca
cacgcatggaagccttgggcgtaagctcgggtggagcgacccctagtgtagatcttggtg
gtagtagcaaatattcaagtgagaaccttgaagactgaagtggggaagggttccacgcga
acagtgcttggacgtgggttagtcggtcctaagagatggggaaactccgtattcaagtgc
gctcttccagcgcagtccatcgaaagggaagcgggttaacattcccgcaccaggatgtgg
aaactgtgg-ttaccgcaaactgcggcaacgcaactgaacttggggacgcccgtaggagc
tccgagcagggatctctttcctgtttaacgcgcccaacccccctggaatcggtttgtccgg
agatagggggcgaaaggccggaaaagcaccgcacgtcttgtggtgttcggaacgcttgta
atggcccttgaaaacccaggggaaagcagccgtcttacactttttttgtgggacgaggaat
tctcacacctggccgtacccatcaccgcatcaggtccccaaggtgaacagcctctagtca
atagaataatgtagataagggaagtcggcaaaatagatccgtaactttgggaaaaggatt
ggctctaatgttcgggcttgctgggacgcgcgggcgtagacgccaatgagcggcagacgt
ctagcctcctcggaggctggattgacgctgcttgtgggaagaaaccgtgtcgatggaggc

aggagcgcttgccgttcttgtctcttcctgcttgcgatcaacgaacggattagaactggt
acggacaaggggaatccgactgtttaattaaaacaaagcattgcgacggccggaaacggt
gttggcgcaatgtgatttctgcccagtgctctgaatgtcaaagtgaagaaattcaaccaa
gcgcgggtaaacggcgggagtaactatgactctcttaact--------------------
------------------------------------------------------------
------------------------------------------------------------
------------------------------------------------------------
------------------------------------------------------------
------------------------------------------------------------
------------------------------------------------------------
------------------------------------------------------------
-----------------------------------------------aca----------
------------------------------------------------------------
------------------------------------------------------------
-----------------------------tggacatcttttgaga---------------
------------------------------gagataggcgggctgaaaaggt------ta
tagccgtctagtctcgc-------------------------------------------
---------------------------------tctttcatg-agcggggcgacacacct
ggttgcgggaaccctgcccacctctctatcag-aagtgggaaccagtccattcctactcc
tgcttgtgaggcatatagtaaaaagtagcagcaggaggatctcggcgagaggccgaaccc
gaaaggaataacgtgggctgggaggcaatccgcaggcgagtccgtaactatatgattatc
gacgggccgtctcaacgactggtaaggtgtggg-------tttgtcctcacgc-gggaca
ggcttaaggtacagtctaatcccacccgccaagggtgcttggg---tg------------
--------------------taatccctcttacatagagggcactagccca-ggggctgg
gacgctgggaggaa----------------atgccccgtccctatagccggtagaaatga
aggtagccaaatgcctcgtcatctaaactaattgg-at----------------------
-----------------------gagatagtc----------tgagtgaaaagcttatgc
ag-----------------gctagtaccggta-------aaaaaacgagaggctctgcca
ct----ctgac------------------------ggtgcgacactccttgtacagggaa
accctcc--gcatggaaaggcacgtggtaccaccctcgcgaggaaactcgtgagcgggat
cccgggtaatgaccggagccgatggtaaaaacccacgtgaccgatggggtgaccctgtgg
cgagaagctacgttgtga-----cttgtgtgcacaatatgcttccgtcgcagagactgaa
cgggagtgggctccgtcga---------cggccg----gccagccgtcttcca-------
---ggagcttaagatacagtccggcccacagggatgg---cttaaacctgtggaacaa-g
cagaattagtgacgcgcatgaatggattaacgagattcccactgtccctatctactatct
agcgaaaccacagccaagggaacgggcttggaagaatcagcggggaaagaagaccctgtt

gagcttgactctagtccgactttgtgagatgagttgggaggtgtagtctaagagggaggg
gcaactcgaccgtgaaatacctctactcccaagcttgttttacttactccggcgagcagg
aatgggtggcaacacccttttctaggattaacacctcttcgggggtggacctgcgcgga
ggacattgtcaggtggggagtttggctggggcggcacatctgttaaaagataacgcaggt
gtcctaagatgagctcagtgagaacagaaatctcacgtagaacaaaagggtaaaagctca
tttgattttgattttcagtacgaatacaaactgtgaaagcatggcctatcgatcctttg
accttcaaggtttgaagcaagaggtgtcagaaaagttagcacgcgttgtggtagctttta
agagtggactcaaagtgttccgctagtccccctcggaccgccacttacaaaaaggacag
aggaggggcgacactgtcaaattgcggggactccctaaagcctgtctctaccaagccag
-gcggcgaaagctcctggtggccggggtaatgacctagggtatggtaaaagcgagacagg
atgagaacaatgggcaatccgcagccaagtcctacggctcttttcttggagaagagctat
ggatgcagttcaacgactagatggcagtgggctgagggagcctacggcggcacccttcg
gcttaagatatagtcttgcccacgtgaaagcgtgcctgcagatgaacgccaccatgtatg
gcaggagctgcaggaggccgctaaacataccagcggctgggaacataacagtaccacagg
gatgaac-----------------------------------------------------
------------------------------------------------------------
------------------------------------------------------------
------------------------------------------------------------
------------------------------------------------------------
-------------------------------------------------aaaaacac---
------------------------------------------------------------
------------------------------------------------------------
--acaattg---------------------------------------------------
------------------------------------------ttgtggca---atcatac
----------------------------------------agtcgta-------------
------------------------------------------------------------
------------------------------------------------------------
------------------------------------------------------------
------------tgatattatcgtcccaggcgggtgcgcctctctcacccgtctagtccg
gccttggtgcgcaaaaaatgcttgcgtggatgcgccttggttaaggaaaatgctcctcgt
agttttacaaggacatttcttgtagggcaacacatctggatgcgaagaactcccgg-aaa
gacaacggtaccaagtgactggggtaacgccccattcataaaggcaatgaat---aaaga
ggtgccgggtaataatctgttgtatagggacaatccgcagctgacttggctccctgcccc
cat--cggtgggcatagcttagcggttcaacgctcggtaagatgtgggttggcgcgctgt
ccaacagcgcactggcttaaggtacgggctactcccaccggagacggtgtctttgtctac
ctcgggtaacacgcctcgggagaagcaaagagggtgcaagta--aaaaaactggtagtgg

ggaaatccctaccagagtcacctggtacaaaggaactggcttgtggcagccaagcgttca
tagcgacgttgctttttgatccttcgatgtcggctcttcctatcattgcgaagcagaatt
cgccaagtgttggattgttcacccactaatagggaacgtgagctgggtaggcattttatg
ccctatgcccaacagctggatagctgggcttctgtccta-ggccgggacttctcacgacc
cgtgccttggtcaccc-aaaaaagggcgaccacagagctcaggatggatcttctggctca
actagtagaggacaggtagagatcgtcaaattgcgggaaaacccttagagcctcgagcta
ccaagtgcgtgtcgaaaggcatagacacagccagcaatggctgtcgcatggcttga-ttc
catcaagataggtaaaaatgctcgaggattgggcaatccgcagccaagcccctaaggt--
--ccccccgtggggctatggggaaggttcaacgactgtaaggcgtaccgtgcaagcggaa
gatacagtctagccccgctggaaagcagcgccaaagtaagtaggt-tgcagtggatctgt
cctc
>Form_O_China
gcctgttttgatatgtgttttgacagtcatacgggtcatggccgagggatggtgggtgag
cttcatagcgcaacccccgtcaggagcatgttttcatagcattttgagcatactagggt
ggtctcccatgtagctcgctctctcggctcaggaccttccgtatggggcactttttgacg
agctcctttggggaggagcatgttctggcatatgaagcgttggtcacgagcttcgtagcc
cg---acacataccaaatcagtcttcatatgtctttctggcgggggtcggtcgtatgacc
acgcccttaccttgagcatgtttttcgtatg-----atacctgacgggcagcgccgctct
tcagtcgtcggccgtacctcgtatgcgcgttttctgattcgaacgtgtgacatgcaagcg
catcatatcagaagcatgtttttttcatatgagcatgttggtaggcaggctcttcagtcgt
ctacccataaatgcactttgaaagacgtttttttggtactgaacggctttgtggtcagacg
ttctcgtgagcttattctcgccctctgatacgctggatgctgcggtattacccacagcag
acat---------ccaggagctccctctgcggataag--catctgtagaagggggtttct
ctcggtgac-----gcaatggctgggaaaacggagtgctatccagaagggtgaaacctct
tggggatgtactcccataccagcataatatgctgagaggaattgactcaatgagtgaggt
cctatactgcaacctcttttggatatgtaggcggtctgtacccagactgtcctactggct
gatttcccttggtcc--------cacgcgtcgctggctgaatgggctagtccttatgaag
tgccgcgttcatcaaaagaaccggtcttcatacagcgacctcattc-tatactgccagct
ggatgggctggtccttatggagcgccgcattcttcgaagaatcggtttctatagagcgac
cccatttacatataaggaaccg-----ttacatagacttgccatttggctgattgctctg
acagtcgctgcctcttcggagttctgtgtaaacgtggcgccatttttgagtgggccagtc
cttatggagtgtccgtgtcttctatgacagggtcccccatacagcgacctcactcagaac
cgtatgggcacgatggtgaaagaccatgcagatagtccgccttccaaagggggaagtagt
atatggggaatgctggcgaagcatgaggt--acatggttcatggacccgttgggccatgc
ctgtgccatcttctcgagtcaaaccggtccgtgtcgcttccttatctcttcctttcgcga
cgacaagctcttgagctggtcgtgggtcaggacgctgtggtatctgtttgtctgtagtgt

gggct-tccaccagaacagcgtatcagcacacgtcgtctgggcatatgatgtttttgctt
tgtggtcccgggtttggtagtatggttccatgtcggtccgctcacctcccaaccataatg
tctttcacatcatcttcacagaacgtcttgtcagcataccattctgtatagaacagtgta
tcagcacacgtcgtctgagcatagggtatttttgctttgtggtcccgggtttggtagtat
ggttccatgtcggtccgctcacctcccaaccataatgtcatttctatcatctgcacagta
cgtcttgtcagaataccattctctatggcagtcgatagactaggccttatgccgcagcag
tctctgatcactgtgcgagtgcatatatctctcaatgtgttcatggctagcatgggtctt
gagtgagatgtgccagagagatgcgccaccagtctatggaaaggtgctttttgtatgaga
agagtacatgtatgaagagcatgttctctcctcttatgatggtaccgcgggtacgggctg
ctcgcattatgcttggcaggtgtccactgtgcatgcaggattgctgaacccacttgtatg
gccatgagaatggtcattgggtgaggttcacttctccatcgctttgtgtgtgttagatgt
ttgggtggcttcggccgcatggaaattctcggtctgtaactcatcagggagggtttccat
gtgagta--ttctccagtcgaccacacacccttgggcgtcccgctagacaaaatgtaact
ctcgatccgacctagcctgtggagtgtct-ttccgtatggcgtgaaagggcccacactgg
atgttttcccttttggacaccagtctgttagtatgtatatatagatcatctacagggagc
aaaggacaggcatgcaagtgccacaaaggcagggaggccttcctgctactgtgtgtcctt
cttgacgatcacgggggagtatatagaaacatttgaaggagatcc-ttttccacaaggcg
gtgccctcgcgcaaaggtgttgctctgcacgcatctgtatgatgatagttgctcgctcct
atgcgtccacttgtgaaaataccttctctctttgtctgtttctatgacgccccttctgct
ttgtcatagagcatactcaggtacagaaagtgccgtgacttccttggcctgctgtcaaag
aaatagcctggtgctatagaggactc-ttcctgggcgccgcctggtgaagaggtcctatc
aggctctggagtgtgcgtatgcctatgttcatcatagctcatgtgcacgtgc-ggaaaag
catcgacgtggaatggggaagaacaccattctgaatgagccagttcttcctctgctcccc
ttgcttgacgctgtatgggaatcgctttgccttgggggtacaccttccattccatgacaa
agcattcgatctatgtcccgaggggtgattctacatgaagaacccgaaacttgtctgagt
gagaggcgaaggtagtcctt----------ctggatccgccctcacactgtagtcggcac
agtcccttgtttagggactgcgcagctcaatcttcgactggagaggaaaaaaacaaaaca
caagtatgacacgcacatacgtgttttctaggccattgacctgttggctgcctcgcggtg
gttggcatgatgccccaagtgggggggggggaaatcaatcagtatggttgacaaaatcca
gtgggg-aaaaaggcgcgctgtcgcacacccttggtcacggcctcaaaaagtcgacggcc
ctcactggcccacccctgcgactcctgttcgcttttgacaagacacacctgcaata-tgg
aggaaatgcctgagcagctgtgccgcagcagctgtggacgcgtagcgatatgcatcccta
ttgtttgcgacttggtacaaagaacgcgtgccgggagacggattgccctttgacttttcc
acccggcccgcggctctgccctttacgtacccttatttacctggttgatcctgccagtag
tcatatgcttgtctcaaagactaagccatgcatgtctaagtataaacacttttatactcg
tgaaactgcgaatggctcattaaaacagttatagttcctttgggaacaacactacttgga

taaccgtagtaattctagagctaatacatgcctcaacgcccgactcacgaagggtggtat
ttattggataaaaaaccatcgtgctctctccagagcgctttgagatgattcacaataact
tgtcggatcgcatggcctcgtgctggcgactcctcattcaaatttctgccctatcaactt
tcgatggtagagtattggtctaccatggtgtcgacgggtgacggggaattagggttcgat
tccggagagggagcctgagaaacggctaccacatccaaggaaggcagcaggcgcgcaaat
tacccaatcccgactcggggaggtagtgacaaaaaataacaataggggggccctttgggtc
ttctaattggaatgagaacaatttaaatcccttatcgaggatccattggagggcaagtct
ggtgaacgta----------------------------------tgcatca---------
--aagccctcgat----aaaaaaaaacagcgccagatgtct-------------------
------------------------------------------------------------
------------------------------------------------------------
------------------------------------------------------------
------------------------------------------------------------
------------------------------------------------------------
-------ctctctggaagcgcacca-----------------------------------
---tgcat-ggctttgttcctctc-----------------------------tgcatgg
ccg-------------------------------------------tgtggaaggcgagt
aggcaaacgatgggaaaccaggcttggaatagtaaccaaccacagagcccccatcataca
cttctaaattgccggggagtgcccgcgaggccttgactaccgcggtccgccgagaggtga
gagcgagcaatcgttcaggacgcagcagtgtacgtagtggtacatggatggtaaaaacgt
caaggatagggttgatccgcagggaagcgcgagtgccgccatg---------gtcggtgg
gcaaaggcgaaccttcagagactctaatggagtgggcgcaagcttaagggagagtccaat
ccactgggaaaccaggtccacagcagtaacggggtgatgctcgccccagaggactgtgga
aggcaccaccaccccaagagggggtgggtgttgtcgggtagcaaatggccagcagccgcg
gtaattccagctccaatagcgtatattaaagttgttgcagttaaaacgctcgtagtcgga
cctcggagcgcccccacgggccgggcttttgccttgttgccagtgggagtttgtctcctt
ttgtcgggacgcggtgtgcagggcctttactgtctctgcgcgtttacctgcgggccgacc
gtttactgtgaagaaagtaaagtgttcaaagcaggccaatgccttgaatatgtgagcatg
gaataatagaataggacttgggctctattttgttggtttccagtgaccaagtaatgatta
atagggacggttgggggcattcgtatttcattgtcagaggtgaaattcttggattgatgg
aagacgtacaactgcgaaagcatctgccatggatgttttcattgatcaagaacgaaagtt
aggggatcgaagacgatcagataccgtcgtagtcttaaccataaacgatgccgactgggg
attggcgggggcactattatatgactctgtcagcaccctgagggaaaccaaagtctttgg
gttctgggggggagtatggtcgcaaggctgaaacttaaaggaattgacggaagggcaccac
caggagtggagcctgcggcttaatttgactcaacacgggaaaacttaccaggtccggaca
gaagaatgattgacagactgaaagctctttcttgatttttttggttggtggtgcatggccg

ttcttagttggtggagtgatttgtctggttaattccgttaacgaacgagacctcgtcctg

ctaaataggtgcgcgcatgccagcactgcgttcttaccttcttagagggactatgcgcgt

ctagcgtatggaagattgaggcaataacaggtctgtgatgcccttagatgttctgggccg

cacgcgcgctacactgatgcattcaacgagtttcacttgatagtcctgggtcggaaggct

cgggtaatcttttgaaagtgcatcgtgctggggatagatcattgcaattattgatcttca

acgaggaattccttgtaggtgcaggtcatcagcctgcgccgaatacgtccctgccctttg

tacacaccgcccgtcgctcctaccgattgaatggtccgatgaaatgtccggatcgcggct

agtgtgatggactcgtccagagcggtagccgtgagaagctcattaaatcttatcatttag

aggaaggagaagtcgtaacaaggtttccgtattttgtgggtggataccgattcaataaca

tccgccgtttttctcaatgtactatatccctagtctgcttgcctagttcttggagtctcg

tgcgatgtgtgc------------------------------------------------

------------------------------------------------------------

-aaagatgga--------------------------------------------------

------------------------------------------------------------

------------------------------------------------------------

--------------------------------------cgccatatcggagacaggggga

ggtgccattg---tgatgatgacgatgttgttgggtggtccttccgtaggaagacgtaca

cgaagcctttgcggcccgaaagggtgcacagtgcgcgactataaacactgattctgtggt

caacgcaagcagcggttctgtggaagaacagctgcgactttctcaaattgcgggaacacc

ctaaagtcaacaagcaccgcggcttgacgtgaaaagcgtcaggcgcagcagtcggggtag

cgccctggcggatggtaatagccttgttgaatggcacaatgggcaatccgcagccaagcc

cctaaggtcccccgctgggctatggggaaggttcacagactgtaagggagggggtttctt

ccatga--aagaagcttaagagacagtcggtcccctgcgagagcagtgtcgcatgtgagg

aaggcacccatgaaaagtgggtgctggagagccatgtgagccctctatgagggtgggaag

ttttccga---tacaatgagaaacaaacacgttccgtaggtgaacctgcggaaggatcat

tcacaaactattgacacaaacacacgcgaaccaaatcgtccacaca---ggtgccgatga

aagagagaatctgattgtcgccttttggggtat----acaagcactctttt-ccatcgcc

tctgtgcagggcgtaacttcccattgagaggatgtgagggcaccacaggaagcttttcca

caggaagtcgccatccttctccctccacgacggcggcgcttcacaggcttggcag-tttt

ttttttttttattgccttccagggaggatgccgccaatggagccacatag-acatttaca

tcatcatatgcccgtttttctttcttaaccgcttgccacagtttctcctgtgaggagctt

gtgggaagaccgtctccatacaataacaaaagatacaactcttagcggtggatatcttgg

ttctcgcaacgatgaagaacgcagccaactgcgatacctagtgtgaattgcaggactccg

tgaatcattaagtctttgaacgcaagttgcgctcatctccatgaggaggtgagcatgcct

gtttgagtgtcgatgggaaaaacaatacaataacaacaatcggctctcttcccttcctgc

cttccccggc-aaaaaggtagtgcatgccggagatgggcctggtcgcccct---tcatga

aagaggcgtcgtccgaaacacgtgtcgttgcccgctggaggtcatacagcatgagtttc-
aaaacaccatggctctcaaggcagagtaaatcgtcttgagacggtct-gcattgattcac
tttcaacccaggtggcgttggttacttgccat-aagcctctctggggtttctttgcttgg
acctctcttgtcagttcttctgacgtagagacaggt--gcggtctcatgggcgttgccct
ctggaacgtgctaaaaaaaacaaaacaaacaacaacaacaacaaccgtatcacacgtcaa
gagggtcggatcgtacttgacggcgggagatcctttgtttctcgcttgtcctgagcgtct
gccaagccttctttgacgcgtgtgtgtttgttccttccataaacccgatctcacatcagg
taggaccacccgctgaatttaagcatataattaagcggaggaaaagaaactaacaaggat
tccctcagtagcggcgagcgaagcgggaagagctcaacatgaaaaccgtaggggggttcgc
gctcaccttcgggttgtatagtaaagtggtgtcgcctgcgtcgcgccgggtataaatccc
gtttgaaatggggtgtcgtagagggttataaccccgtctgtgacccggtgtgcgatgcgt
tacggtgaccttacgatgagtcgggttgcttgggcatgcagccctaagtgggtggtaaac
ttcatctaaagctaaatactagcgagagaccgatagcgtacaagtaccgtaagggaaaga
tgcaaagaactttgaaaagagagttaaacagggcttgaaaccgttaagagtgaagcgaag
gaggtcagtgatcatggggccagatccccacgtgcccgtcataggggcaccgtagcgttc
tgg-ttcccggggcagcgccggccgggggcgggtgtcaatagggaggtggttgagggatt
aactttccgcctgtactcctttctagcttgcaccccggccgagggatccgcgacttgtcg
tgtgagggcgctgtcaaaatggccttctccaccccgtcttgaaacacggaccaaggagtc
taacatactcgcgagtgtttgggtggttaaaacccgaacgcacaatgaaagtaaacggtg
gctctgcgctgctgcttggtgggaaccctgctcttataacgggggcaccatcacccgac
cccgatccctttggggtggttggttttgaggcagagcgagtatgatgggacccgaaagat
ggtgaactattcctgagtagggcgaagccagaggaaactctggtggaggctcgtagcgat
gctgacgtgcaaatcgctcgtcgaacttgggtataggggcgaaagactaatcgaaccatc
tagtagctggttccctccgaagtttccctcaggatagcgggagcaaatgcaagtcagtat
catccggtaaagcgaatgattagaggcctcgggggagtaactccttcgacctattctcaa
actttaaataggtgaggagccggcaacgttgcttaaaacaatgaacgccgtgcggacaat
gattgctctcagtgggccgattttggtaagcagaactggcgatgcgggatgaaccgaaag
cgacgctaaggtgccaaactcccagctcattcagacaccagaaaaggtgttgattcatac
agacagcaggacggtggccatggaagtcggaatccgctaaggagtgtgtaacaactcacc
tgccgaatggatcagccctgaaaatggatggcgcttaagcgggggaccgaggcgtcgccg
ttggggtaaatacgaagcctcaatgagtaggagggcgtgggggttttgaaacacggtaca
cacgcatggaagccttgggcgtaagctcgggtggagcgacccctagtgtagatcttggtg
gtagtagcaaatattcaagtgagaaccttgaagactgaagtggggaagggttccacgcga
acagtgcttggacgtgggttagtcggtcctaagagatggggaaactccgtattcaagtgc
gctcttccagcgcagtccatcgaaagggaagcgggttaacattcccgcaccaggatgtgg
aaactgtgg-ttaccgcaaactgcggcaacgcaactgaacttggggacgcccgtaggagc

tccgagcagggatctctttcctgtttaacgcgcccaaccccctggaatcggtttgtccgg
agatagggggcgaaaggccggaaaagcaccgcacgtcttgtggtgttcggaacgcttgta
atggcccttgaaaacccaggggaaagcagccgtcttacactttttttgtgggacgaggaat
tctcacacctggccgtacccatcaccgcatcaggtccccaaggtgaacagcctctagtca
atagaataatgtagataagggaagtcggcaaaatagatccgtaactttgggaaaaggatt
ggctctaatgttcgggcttgctgggacgcgcgggcgtagacgccaatgagcggcagacgt
ctagcctcctcggaggctggattgacgctgcttgtgggaagaaaccgtgtcgatggaggc
aggagcgcttgccgttcttgtctcttcctgcttgcgatcaacgaacggattagaactggt
acggacaaggggaatccgactgtttaattaaaacaaagcattgcgacggccggaaacggt
gttggcgcaatgtgatttctgcccagtgctctgaatgtcaaagtgaagaaattcaaccaa
gcgcgggtaaacggcgggagtaactatgactctcttag----------------------
------------------------------------------------------------
------------------------------------------------------------
------------------------------------------------------------
------------------------------------------------------------
------------------------------------------------------------
------------------------------------------------------------
------------------------------------------------------------
-----------------------------------------------aca----------
------------------------------------------------------------
------------------------------------------------------------
-----------------------------tggacatcttttgaga---------------
------------------------------gagataggcgggctgaaaaggt------ta
tagccgtctagtctcgc-------------------------------------------
---------------------------------tctttcatg-agcggggcgacacacct
ggttgcgggaaccctgcccacctctctatcag-aagtgggaaccagtccattcctactcc
tgcttgtgaggcatatagt-aaaagtagcagcaggaggatctcggcgagaggccgaaccc
gaaaggaataacgtgggctgggaggcaatccgcaggcgagtccgtaactatatgattatc
gacgggccgtctcaacgactggtaaggtgtggg-------tttgtcctcacgc-gggaca
ggcttaaggtacagtctaatcccacccgccaagggtgcttggg---tg------------
--------------------taatccctcttacatagagggcactagccca-ggggctgg
gacgctgggaggaa----------------atgccccgtccctatagccggtagaaatga
aggtagccaaatgcctcgtcatctaaactaattgg-at----------------------
-----------------------gagatagtc----------tgagtgaaaagcttatgc
ag-----------------gctagtaccggga-------aaaaaaagaggggctctgcca
ctctgactgac------------------------ggtgcgacactccttgtacagggaa

```
accctcc--gcatggaaaggcacgtggtaccaccctcgcggggaaactcgcgagcgggat
cccgggtaatgaccggagccgatggtaaaaacccacgtgaccgatggggtgaccctgtgg
cgaggcgctaagttgtgggacatcaccctctcacaatacgcatccgtcgcagagactgaa
cgggagtgggctccgtcga---------cggccggccagccagccgtcttcca-------
---ggggcttaagatacagtccgggcctcagggatgg----aaaaacctggggaacta-c
cagaattagtgacgcgcatgaatggattaacgagattcccactgtccctatctactatct
agcgaaaccacagccaagggaacgggcttggaagaatcagcggggaaagaagaccctgtt
gagcttgactctagtccgactttgtgagatgagttgggaggtgtagtctaagagggaggg
gcaactcgaccgtgaaatacctctactcccaagcttgttttacttactccggcgagcagg
gatgggtggtaacacccttttttctaggattaacacctctttgggggtggacctgcgcgga
ggacattgtcaggtggggagtttggctggggcggcacatctgttaaaagataacgcaggt
gtcctaagatgagctcagtgagaacagaaatctcacgtagaacaaaagggtaaaagctca
tttgattttgattttcagtacgaatacaaactgtgaaagcatggcctatcgatcctttttg
accttcaaggtttgaagcaagaggtgtcagaaaagttagcacgcgttgtggtagctttta
agagtggactcaaagtgttccgctagtccccccctcggaccgccacttacaaaaaggacag
aggagggggcgacactgtcaaattgcggggactccctaaagcctgtctctaccaagccag
-gcggcgaaagctcctggtggccggggtaatgacctagggtatggtaaaagcgagacagg
atgagaacaatgggcaatccgcagccaagtcctacggctcttttcttggagaagagctat
ggatgcagttcaacgactagatggcagtgggctgagggagcctacggcggcaccctttcg
gcttaagatatagtcttgcccacgtgaaagcgtgcctgcagatgaacgccaccatgtatg
gcaggagctgcaggaggccgctaaacataccagcggctgggaacataacagtaccacagg
gatgaac-----------------------------------------------------
------------------------------------------------------------
------------------------------------------------------------
------------------------------------------------------------
------------------------------------------------------------
-------------------------------------------------aaaaacac---
------------------------------------------------------------
------------------------------------------------------------
--acaattg---------------------------------------------------
------------------------------------------ttgtggcaatcatcatac
----------------------------------------agtcgta-------------
------------------------------------------------------------
------------------------------------------------------------
------------------------------------------------------------
------------tgatagtatcgtcccaggcgggtgcgcctctctcacccgtctagtccg
```

gccttggtgcgcaaaaaatgcttgcgtggatgcgccttggttaaggaaaatgctcctcgt
agttttacaaggacatttcttgtagggcaacacatctggatgcgaagaactcccgg-aaa
gacaacggtaccaagtgcttggggtaacgccccattcatgaaggcaatgaat---aaaga
ggtgccgggtaataatctgttgtatagggacaatccgcagctgacttggctccctg--cc
cat--cggtgggcatagcttagcggttcaacgctcggtaagatgtgggttggcgcgctgt
ctgacagcgcactggcttaaggtacgggctactcccaccggagacggtgtctttgtctac
ctcgggtaacacgcctcgggagaagcaaagagggtgcaaat----aaaaactggtagcgg
ggaaatccctaccagagtcacctggtacaaaggaactggcttgtggcagccaagcgttca
tagcgacgttgctttttgatccttcgatgtcggctcttcctatcattgcgaagcagaatt
cgccaagtgttggattgttcacccactaatagggaacgtgagctgggtaggcattttatg
ccctatgcccaacagctggatagctgggcttctgtccta-ggccgggacttctcacgacc
cgtgccttggtcaccc-aaaaaagggcgaccacagagctcaggatggatcttctggctca
actagtagaggacaggtagagatcgtcaaattgcgggaaaccccttagagcctcgagcta
ccaagtgcgtgtcgaaaggcatagacacagccagcaatggctgtcgcatggcttga-ttc
catcaagataggtaaaaatgctcgaggattgggcaatccgcagccaagcccctaaggt--
--ccccccgtggggctatggggaaggttcaacgactgtaaggcgtaccgtgcaagcggaa
gatacagtctagccccgctggaaagcagcgccaaagtaagtaggt-tgcagtggatctgt
cctc
>Form_N_Japan
gcctgttttgatatgtgttttgacagtcatacgggtcatggccgagggatggtgggtgag
cttcatagcgcaacccccccgtcaggagcatgttttcatagcattttgagcatactagggt
ggtctcccatgtagctcgctctctcggctcaggaccttccgtatggggcactttttgacg
agctcctttggggaggagcatgttctggcatatgaagcgttggtcacgagcttcgtagcc
cg---acacataccaaatcagtcttcatatgtctttctggcgggggtcggtcgtatgacc
acgcccttaccttgagcatgtttttcgtatg-----atacctgacgggcagcgccgctgt
tcagtcgtcggccgtacctcctatgcgcgttttctgattcgaacgtgtgacattcaagcg
catcatatcagaagcatgtttttttcatatgagcatgttggtaggcaggctcttcagtcgt
ctacccataaatgcactttgaaagacgttttttggtactgaacggctttgtggtcagacg
ttctcgtgagcttattctcgccctctgatacgctggatgctgcggtattacccacagcag
acat---------ccaggagctccctctgcggataag--catctgtagaaggggtttct
ctcggtgac-----gcaatggctgggaaaacggagtgctatccagaagggtgaaacctct
tggggatgtactcccataccagcataatatgctgagaggaattgactcaatgagtgaggt
cctatactgcaacctcttttggatatgtaggcggtctgtacccagactgtcctactggct
gatttcccttggtcc--------cacgcgtcgctggctgaatgggctagtccttatgaag
tgccgcgttcatcaaaagaaccggtcttcatacagcgacctcattc-tatactgccagct
ggatgggctggtccttatggagcgccgcattcttcgaagaatcggtttctatagagcgac

cccatttacatataaggaaccg-----ttacatagacttgccatttggctgattgctctg
acagtcgctgcctcttcggagttctgtgtaaacgtggcgccatttttgagtgggccagtc
cttatggagtgtccgtgtcttctatgacagggtcccccatacagcgacctcactcagaac
cgtatgggcacgatggtgaaagaccatgcagatagtccgccttccaaagggggaagcagt
atatggggaatgctggcgaagcatgaggt--acatggttcatggacccgttgggccatgc
ctgtgccatcttctcgagtcaaaccggtccgtgtcgcttccttatctcttcctttcgcga
cgacaagctcttgagctggtcgtgggtcaggacgctgtggtatctgtttgtctgtagtgt
gggct-tccaccagaacagcgtatcagcacacgtcgtctgggcatatgatgtttttgctt
tgtggtcccgggtttggtagtatggttccatgtcggtccgctcacctcccaaccataatg
tctttcacatcatcttcacagtacgtcttgtcagcataccattctgtatagaacagtgta
tcagcacacgtcgtctgagcatagggtatttttgctttgtggtcccgggtttggtagtat
ggttccatgtcggtccgctcacctcccaaccataatgtcatttctatcatctgcacagta
cgtcttgtcagaataccattctctatggcagtcgatagactaggccttatgccgcagcag
tctctgatcactgtgcgagtgcatatatctctcaatgtgttcatggctagcatgggtctt
gagtgagttgtgccagagagatacgccaccagtctatggaaaggtgctttttgtatgaga
agagtacatgtatgaagagcatgttctctcctcttatgatggtaccgcgggtacgggctg
ctcgcattatgcttggcaggtgtccactgtgcatgcaggattgctgaacccacttgtatg
gccatgagaatggtcattgggtgaggttcacttctccatcgctttgtgtgtgttagatgt
ttgggtggcttcggccgcatggaaattctcggtctgtaactcatcagggagggtttccat
gtgagta--ttctccagtcgaccacacacccttgggcgtcccgctaggcaaaatgtaact
ctcgatccgacctagcctgtggagtgtct-ttccgtatggcgtgaaagggcccacactgg
atgttttcccttttggacaccagtctggtagtatgtatatatagatcatctacagggagc
aaaggacaggcatgcaagtgccacaaaggcagggaggccttcctgctactgtgtgtcctt
cttgacgatcacgggggagtatatagaaacatttgaaggagatcc-ttttccacaaggcg
gtgccctcgcgcaaaggtgttgctctgcacgcatctgtatgatgatagttgctcgctcct
atgcgtccacttgtgaaaataccttctctctttgtctgtttctatgacgccccttctgct
ttgtcatagagcatactcaggtacagaaagtgccgtgacttccttggcctgctgtcaaag
aaatagcctggtgctatagaggactc-ttcctgggcgccgcctggtgaagaggtcctatc
aggctctggagtgtgcgtatgcctatgttcatcatagctcatttgcacgcgc-ggaaaag
catcgacgtggaatggggaagaacaccattctgaatgagccagttcttcctctgctcccc
ttgcttgacgctgtatgggaatcgttttgccttgggggtacaccttccattccatgacaa
agcattcgatctatgtcccgaggggtgattctacatgaagaacccgaaacttgtctgagt
gagaggcgaaggtagtccttcattttaatgcgggatccgccctcacactgtagtcggcac
agtcccttgtttagggactgcgcagctcaatcttcgactggagaggaaaaaaacaaaaca
caagtatgacacgcacatacgtgttttctaggccatggacctgttggctgcctcgcggtg
gttggcatgatgccccaagtgggggggggggaaatcaatcagtatggttgacaaaatcca

gtgggg-aaaaaggcgcgctgtcgcacacccttggtcacggcctcaaaaagtcgacggcc
ctcactggcccacccctgcgactcctgttcgcttttgacaagacacacctgcaata-tgg
aggaaatgcctgagcagctgtgccgcagcagctgtggacgcgtagcgatatgcatcccta
ttgtttgcgacttggtacaaagaacgcgtgccgggagacggattgccctttgacttttcc
acccggcccgcggctctgccctttacgtacccttatttacctggttgatcctgccagtag
tcatatgcttgtctcaaagactaagccatgcatgtctaagtataaacacttttatactcg
tgaaactgcgaatggctcattaaaacagttatagttcctttgggaacaacactacttgga
taaccgtagtaattctagagctaatacatgcctcaacgcccgactcacgaagggtggtat
ttattggataaaaaaccatcgtgctctctccagagcgctttgagatgattcacaataact
tgtcggatcgcatggcctcgtgctggcgactcctcattcaaatttctgccctatcaactt
tcgatggtagagtattggtctaccatggtgtcgacgggtgacggggaattagggttcgat
tccggagagggagcctgagaaacggctaccacatccaaggaaggcagcaggcgcgcaaat
tacccaatcccgactcggggaggtagtgacaaaaaataacaatagggggccctttgggtc
ttctaattggaatgagaacaatttaaatcccttatcgaggatccattggagggcaagtct
ggtgaacgta----------------------------------tgcatca---------
--aagccctcgat----aaaaaaaaacagcgccagatgtct-------------------
------------------------------------------------------------
------------------------------------------------------------
------------------------------------------------------------
------------------------------------------------------------
------------------------------------------------------------
-------ctctctggaagcgcacca-----------------------------------
---tgcat-ggctttgttcctctc-----------------------------tgcatgg
ccg-------------------------------------------tgtggaaggcgagt
aggcaaacgatgggaaaccaggcttggaatagtaaccaaccacagagcccccatcataca
cttctaaattgccggggagtgcccgtgaggccttgactaccgcggtccgccgagaggtga
gagcgagtaatcgttcaggacgcagcagtgtacgtagtggtacatggatggtaaaaacgt
caaggatagggttgatccgcagggaagcgcgagtgccgccatc---------gtcggtgg
gcaaaggcgaaccttcagagactctaatggagtgggcgcaagcttaagggagagtccaat
ccactgggaaaccaggtccacagcagtaacggggtgatgctcgccccagaggactgtgga
aggcaccaccaccccaagagggggtgggtgttgtcgggtagcaaatggccagcagccgcg
gtaattccagctccaatagcgtatattaaagttgttgcagttaaaacgctcgtagtcgga
cctcggagcgcccccacgggccgggcttttgccttgttgccagtgggagtttgtctcctt
ttgtcgggacgcggtgtgcagggcctttactgtctctgcgcgtttacctgcgggccgacc
gtttactgtgaagaaagtaaagtgttcaaagcaggccaatgccttgaatatgtgagcatg
gaataatagaataggacttgggctctattttgttggtttccagtgaccaagtaatgatta

atagggacggttgggggcattcgtatttcattgtcagaggtgaaattcttggattgatgg
aagacgtacaactgcgaaagcatctgccatggatgttttcattgatcaagaacgaaagtt
aggggatcgaagacgatcagataccgtcgtagtcttaaccataaacgatgccgactgggg
attggcgggggcactattatatgactctgtcagcaccctgagggaaaccaaagtctttgg
gttctgggggggagtatggtcgcaaggctgaaacttaaaggaattgacggaagggcaccac
caggagtggagcctgcggcttaatttgactcaacacgggaaaacttaccaggtccggaca
gaagaatgattgacagactgaaagctctttcttgatttttttggttggtggtgcatggccg
ttcttagttggtggagtgatttgtctggttaattccgttaacgaacgagacctcgtcctg
ctaaataggtgcgcgcatgccagcactgcgttcttaccttcttagagggactatgcgcgt
ctagcgtatggaagattgaggcaataacaggtctgtgatgcccttagatgttctgggccg
cacgcgcgctacactgatgcattcaacgagtttcacttgatagtcctgggtcggaaggct
cgggtaatcttttgaaagtgcatcgtgctggggatagatcattgcaattattgatcttca
acgaggaattccttgtaggtgcaggtcatcagcctgcgccgaatacgtccctgccctttg
tacacaccgcccgtcgctcctaccgattgaatggtccgatgaaatgtccggatcgcggct
agtgtgatggactcgtccagagcggtagccgtgagaagctcattaaatcttatcatttag
aggaaggagaagtcgtaacaaggtttccgtattttgtgggtggataccgattcaataaca
tccgccgtttttctcaatgtactatatccctagtctgcttgcctagttcttggagtcttg
tgcgatgtgtgctattacaggcatgccgggtcatgaagacagacaaggttgggatcttca
cggaagtacgcgcaataggagcgcgtcttgttcaacgcacctgactcaagagtcatatgc
gagggacggacgcagcgcttgtcgccgcacagatgggaagcttcctctccaggcatagga
tggcgctgcatttgcatcaccgctgctagtaagtggcaataatacttgactccccgccag
cgcacttgtacgtagccgttggcggcgggaacgtagttggtgatgtgacagccttgagcg
ttcacctctgtgtgtccgccaatctgcacaaagatggacgccatatcggagacaggggga
ggtgccattg---tgatgatgacgatgttgttgggtggtccttccgtaggaagacgtaca
cgaagcctttgcggcccgaaagggtgcacagtgcgcgactataaacactgattctgtggt
caacgcaagcagcggttctgtggaagaacagctgcgactttctcaaattgcgggaacacc
ctaaagtcaacaagcaccgcggcttgacgtgaaaagcgtcaggcgcagcagtcggggtag
cgccctggcggatggtaatagccttgttgaatggcacaatgggcaatccgcagccaagcc
cctaaggtcccccgctgggctatggggaaggttcacagactgtaagggaggggggtttctt
ccatga--aagaagcttaagagacagtcggtcccctgcgagagcagtgtcgcatgtgagg
aaggcacccatgaaaagtgggtgctggagagccatgtgagccctctatgagggtgggaag
ttttccga---tacaatgagaaacaaacacgttccgtaggtgaacctgcggaaggatcat
tcacaaactattgacacaaacacacgcgaaccaaatcgtccacaca---ggtgccgatga
aagagagaatctgtttgtcgccttttggggtat----acaagcactctttt-ccatcgcc
tctgtgcagggcgtaacttcccattgagaggatgtgagggcaccacaggaagcttttcca
caggaagtcgccatccttctccctccacgacggcggcgcttcacaggcttggcag-tttt

tttttttttttattgccttccagggaggatgccgccaatggagccacatag-acatttaca
tcaccatatgcccgtttttctttcttaaccgcttgccacagtttctcctgtgaggagctt
gtgggaagaccgtctccatacaataacaaaagatacaactcttagcggtggatatcttgg
ttctcgcaacgatgaagaacgcagccaactgcgatacctagtgtgaattgcaggactccg
tgaatcattaagtctttgaacgcaagttgcgctcatctccatgaggaggtgagcatgcct
gtttgagtgtcgatgggaaaaacaatac-ataacaacaatcggctctcttcccttcctgc
cttccccggcaaaaaaggtagtgcatgccggagatgggcctggtcgcccct---tcatga
aagaggcgtcgtccgaaacacgtgtcgttgcccgctggaggtcatacagcatgagtttca
aaaacaccatggctctcaaggcagcgtaaatcgtcttgagacggtctggcattgattcac
tttcaacccaggtggcgttggttacttgccat-aagcctctctggggtttctttgcttgg
acctctcttgtcagttcttctgacgtagagacaggtgcgcggtctcatgggcgttgccct
ctggaacgtgct-aaaaaaacaaaacaaacaacaacaacaacaaccgtatcacacgtcaa
gagggtcggatcgtacttgacggcgggagatcctttgtttctcgcttgtcctgagcgtct
gccaagccttctttga--cgagtgtgtttgttccttccataaacccgatctcacatcagg
taggaccacccgctgaatttaagcatataattaagcggaggaaaagaaactaacaaggat
tccctcagtagcggcgagcgaagcgggaagagctcaacatgaaaaccgtaggggggttcgc
gctcaccttcgggttgtatagtaaagtggtgtcgcctgcgtcgcgccgggtataaatccc
gtttgaaatggggtgtcgtagagggttataaccccgtctgtgacccggtgtgcgatgcgt
tacggtgaccttacgatgagtcgggttgcttgggcatgcagccctaagtgggtggtaaac
ttcatctaaagctaaatactagcgagagaccgatagcgtacaagtaccgtaagggaaaga
tgcaaagaactttgaaaagagagttaaacagggcttgaaaccgttaagagtgaagcgaag
gaggtcagtgatcatggggccagatccccacgtgcccgtcata-gggcaccgtagcgttc
tgg-ttcccggggcagcgccggccgggggcgggtgtcaatagggaggtggttgagggatt
aactttccgcctgtactcctttctagcttgcaccccggccgagggatccgcgactcgtcg
tgtgagggcgctgtcaaaatggccttctccaccccgtcttgaaacacggaccaaggagtc
taacatactcgcgagtgtttgggtggttaaaacccgaacgcacaatgaaagtaaacggtg
gctctgcgctgctgcttggtgggaaccctgctcttataacgggggcaccatcacccgac
cccgatccctttggggtggttggttttgaggcagagcgagtatgatgggacccgaaagat
ggtgaactattcctgagtagggcgaagccagaggaaactctggtggaggctcgtagcgat
gctgacgtgcaaatcgctcgtcgaacttgggtataggggcgaaagactaatcgaaccatc
tagtagctggttccctccgaagtttccctcaggatagcgggagcaaatgcaagtcagtat
catccggtaaagcgaatgattagaggcctcgggggagtaactccttcgacctattctcaa
actttaaataggtgaggagccggcaacgttgcttaaaacaatgaacgccgtgcggacaat
gattgctctcagtgggccgattttggtaagcagaactggcgatgcgggatgaaccgaaag
cgacgctaaggtgccaaactcccagctcattcagacaccagaaaaggtgttgattcatac
agacagcaggacggtggccatggaagtcggaatccgctaaggagtgtgtaacaactcacc

tgccgaatggatcagccctgaaaatggatggcgcttaagcgggggaccgaggcgtcgccg
ttggggtaaatacgaagcctcaatgagtaggagggcgtgggggttttgaaacacggtaca
cacgcatggaagccttgggcgtaagctcgggtggagcgacccctagtgtagatcttggtg
gtagtagcaaatattcaagtgagaaccttgaagactgaagtggggaagggttccacgcga
acagtgcttggacgtgggttagtcggtcctaagagatggggaaactccgtattcaagtgc
gctcttccagcgcagtccatcgaaagggaagcgggttaacattcccgcaccaggatgtgg
aaactgtgg-ttaccgcaaactgcggcaacgcaactgaacttggggacgcccgtaggagc
tccgagcagggatctctttcctgtttaacgcgcccaacccccctggaatcggtttgtccgg
agatagggggcgaaaggccggaaaagcaccgcacgtcttgtggtgttcggaacgcttgta
atggcccttgaaaacccaggggaaagcagccgtcttacactttttttgtgggacgaggaat
tctcacacctggccgtacccatcaccgcatcaggtccccaaggtgaacagcctctagtca
atagaataatgtagataagggaagtcggcaaaatagatccgtaactttgggaaaaggatt
ggctctaatgttcgggcttgctgggacgcgcgggcgtagacgccaatgagcggcagacgt
ctagcctcctcggaggctggattgacgctgcttgtgggaagaaaccgtgtcgatggaggc
aggagcgcttgccgttcttgtctcttcctgcttgcgatcaacgaacggattagaactggt
acggacaaggggaatccgactgtttaattaaaacaaagcattgcgacggccggaaacggt
gttggcgcaatgtgatttctgcccagtgctctgaatgtcaaagtgaagaaattcaaccaa
gcgcgggtaaacggcgggagtaactatgactctcttaa----------------------
------------------------------------------------------------
------------------------------------------------------------
------------------------------------------------------------
------------------------------------------------------------
------------------------------------------------------------
------------------------------------------------------------
------------------------------------------------------------
------------------------------------------------------------
------------------------------------------------------------
------------------------------------------------------------
------------------------------------------------------------
------------------------------------------------------------
------------------------------------------------------------
------------------------------------------------------------
------------------------------------------------------------
------------------------------------------------------------
------------------------------------------------------------
------------------------------------------------------------

------------------------------------------------------------
------------------------------------------------------------
------------------------------------------------------------
-ggtagccaaatgcctcgtcatct------------------------------------
------------------------------------------------------------
------------------------------------------------------------
------------------------------------------------------------
------------------------------------------------------------
------------------------------------------------------------
------------------------------------------------------------
------------------------------------------------------------
------------------------------------------------------------
---aattagtgacgcgcatgaatggattaacgagattcccactgtccctatctactatct
agcgaaaccacagccaagggaacgggcttggaagaatcagcggggaaagaagaccctgtt
gagcttgactctagtccgactttgtgagatgagttgggaggtgtagtctaagagggaggg
gcaactcgaccgtgaaatacctctactcccaagcttgttttacttactccggcgagcagg
aatgggtggcaacacccttttttctaggattaacacctcttcgggggtggacctgcgcgga
ggacattgtcaggtggggagtttggctggggcggcacatctgttaaaagataacgcaggt
gtcctaagatgagctcagtgagaacagaaatctcacgtagaacaaaagggtaaaagctca
tttgattttgattttcagtacgaatacaaactgtgaaagcatggcctatcgatccttttg
accttcaaggtttgaagcaagaggtgtcagaaaagt------------------------
------------------------------------------------------------
------------------------------------------------------------
------------------------------------------------------------
------------------------------------------------------------
------------------------------------------------------------
------------------------------------------------------------
---------------------------------------------------taccacagg
gat---------------------------------------------------------
------------------------------------------------------------
------------------------------------------------------------
------------------------------------------------------------
------------------------------------------------------------
------------------------------------------------------------
------------------------------------------------------------
------------------------------------------------------------

------------------------------------------------------------

------------------------------------------------------------

------------------------------------------------------------

------------------------------------------------------------

------------------------------------------------------------

------------------------------------------------------------

------------------------------------------------------------

------------------------------------------------------------

------------------------------------------------------------

------------------------------------------------------------

------------------------------------------------------------

------------------------------------------------------------

------------------------------------------------------------

------------------------------------------------------------

---------------------------------aactggcttgtggcagccaagcgttca

tagcgacgttgctttttgatccttcgatgtcggctcttcctatcattgcgaagcagaatt

cgccaagtgttggattgttcacccactaatagggaacgtgagctgggtaggcattttatg

ccctatgcccaacagctggatagctgggcttctgtccta-ggccgggacttctcacgacc

cgtgccttggtcacccaaaaaaagggcgaccacagagctcaggatggatcttctggctca

actagtagaggacaggtagagatcgtcaaattgcgggaaaacccttagagcctcgagcta

ccaagtgcgtgtcgaaaggcatagacacagccagcaatggctgtcgcatggcttga-ttc

catcaagataggtaaaaatgctcgaggattgggcaatccgcagccaagcccctaaggt--

--ccccccgtggggctatggggaaggttcaacgactgtaaggcgtaccgtgcaagcggaa

gatacagtctagccccgctggaaagcagcgccaaagtaagtaggt-tgcagtggatctgt

cctc

**Supplementary Figure 5.** Phylogenetic trees.

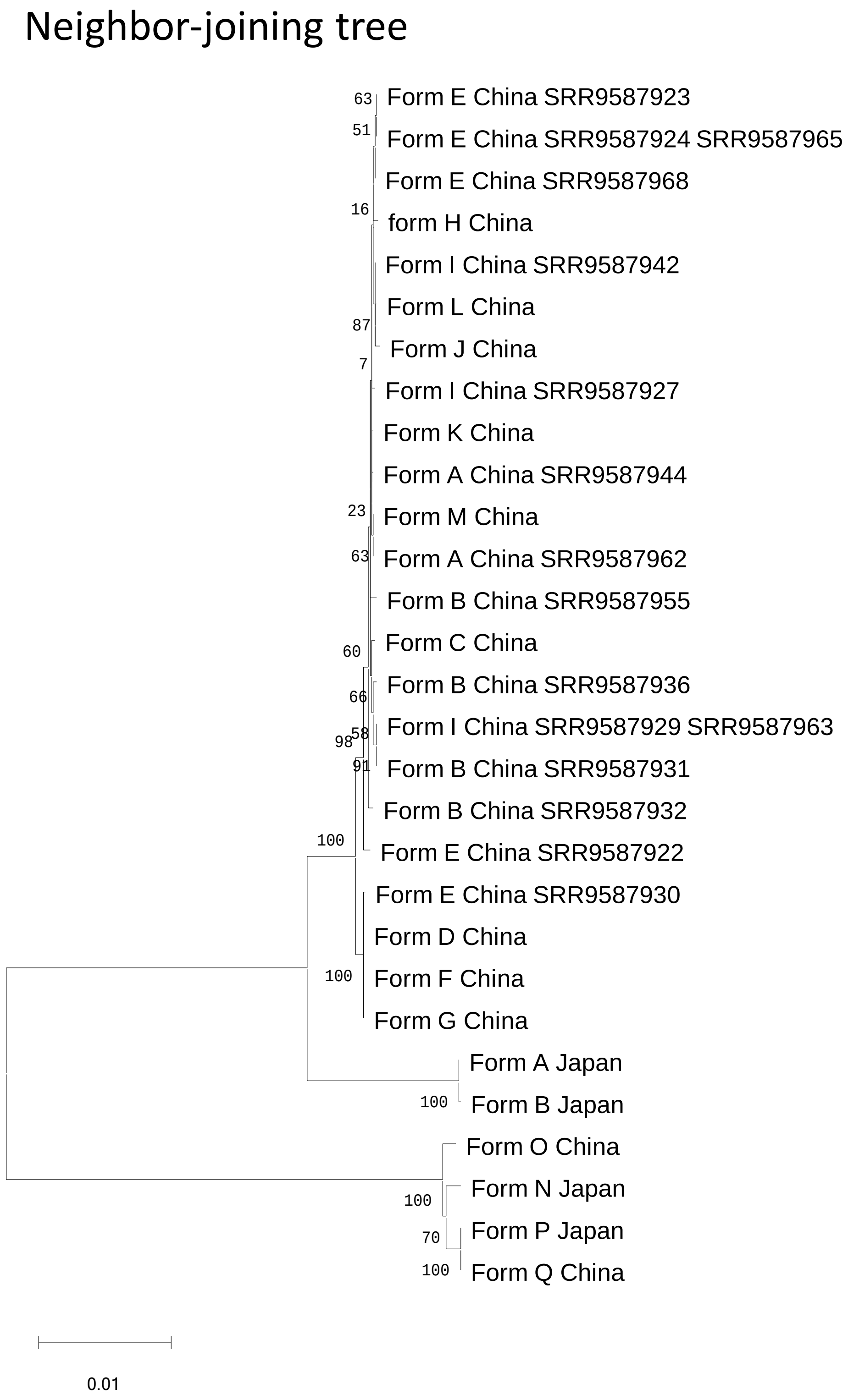

### Maximum likelihood tree

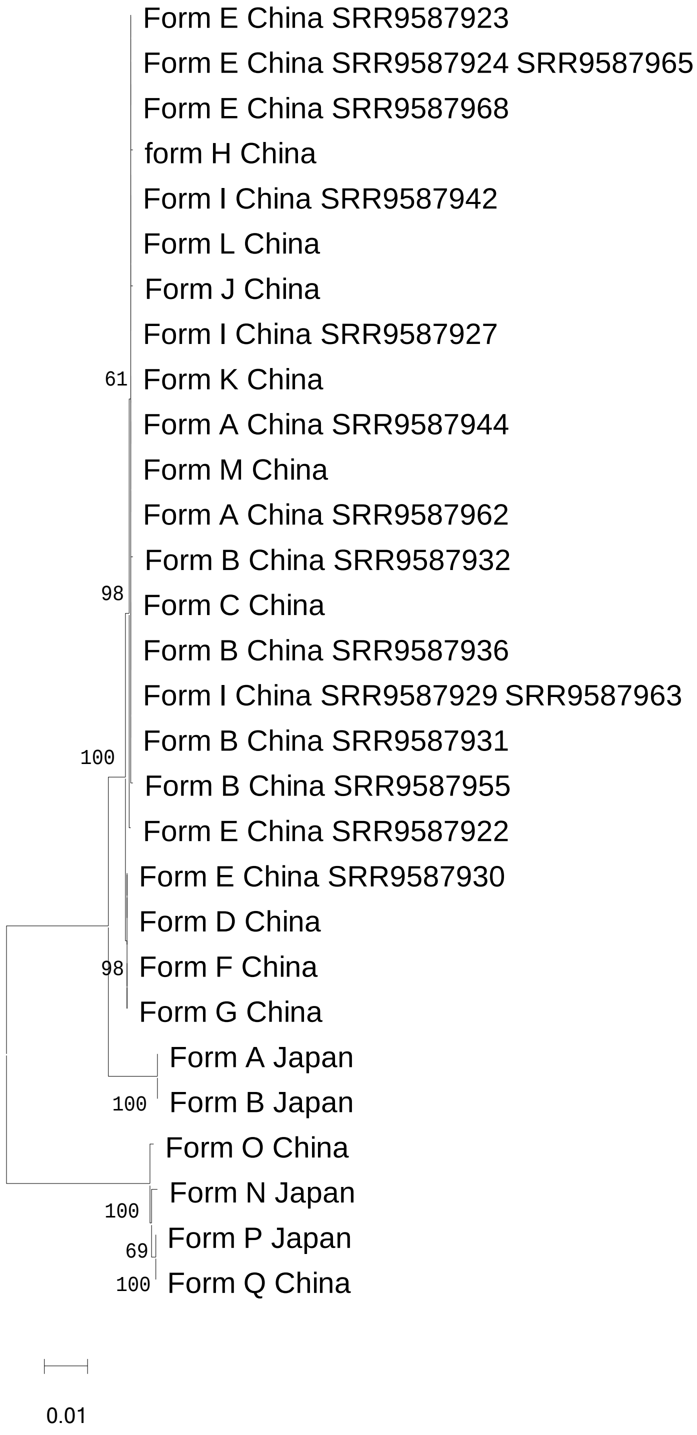

**Supplementary Figure 6.** The results of mapping the reads of Chinese *N. yezoensis* to the sequence of SRR9587944.

SRR9587917

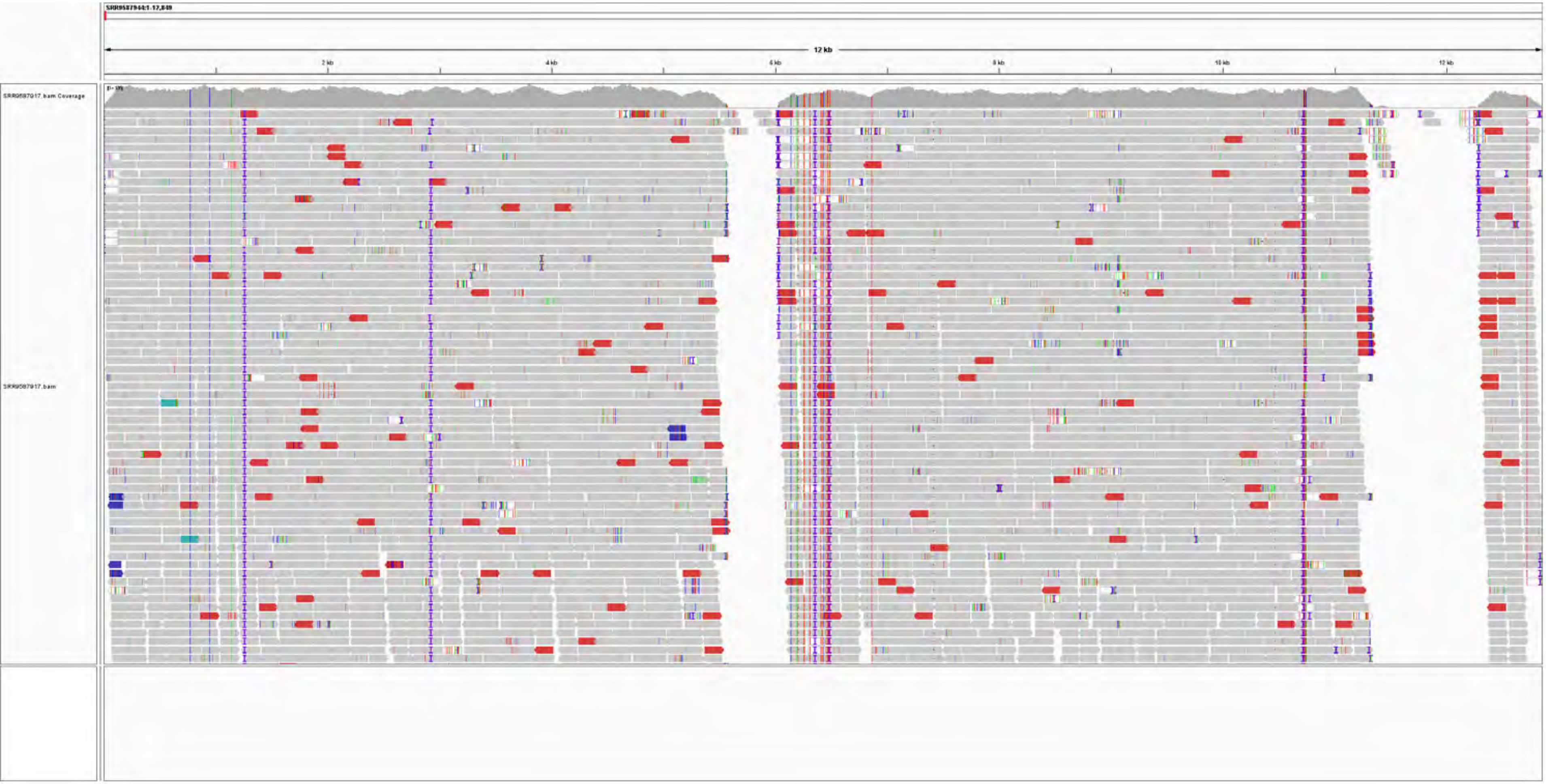

### SRR9587918

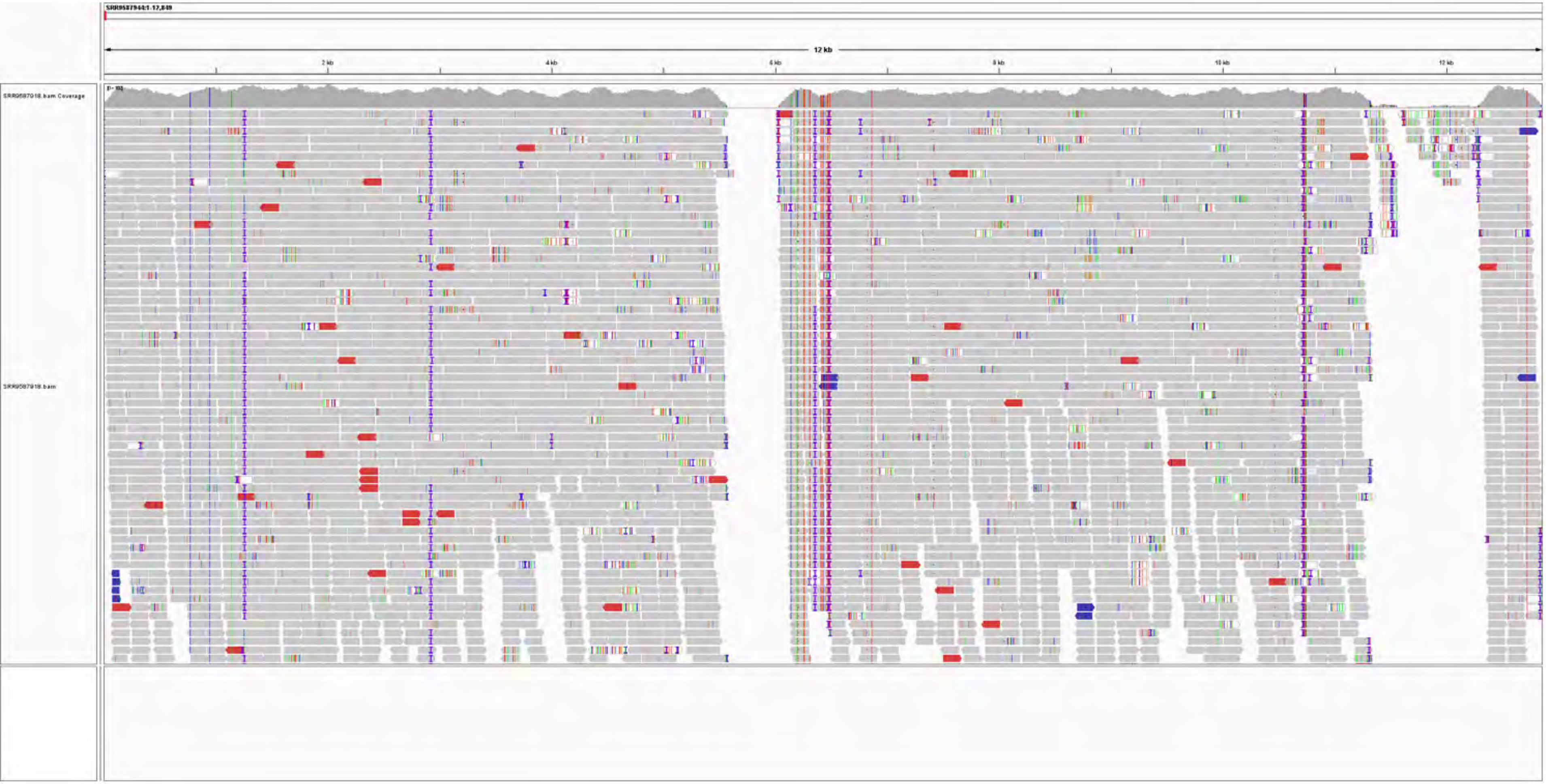

### SRR9587919

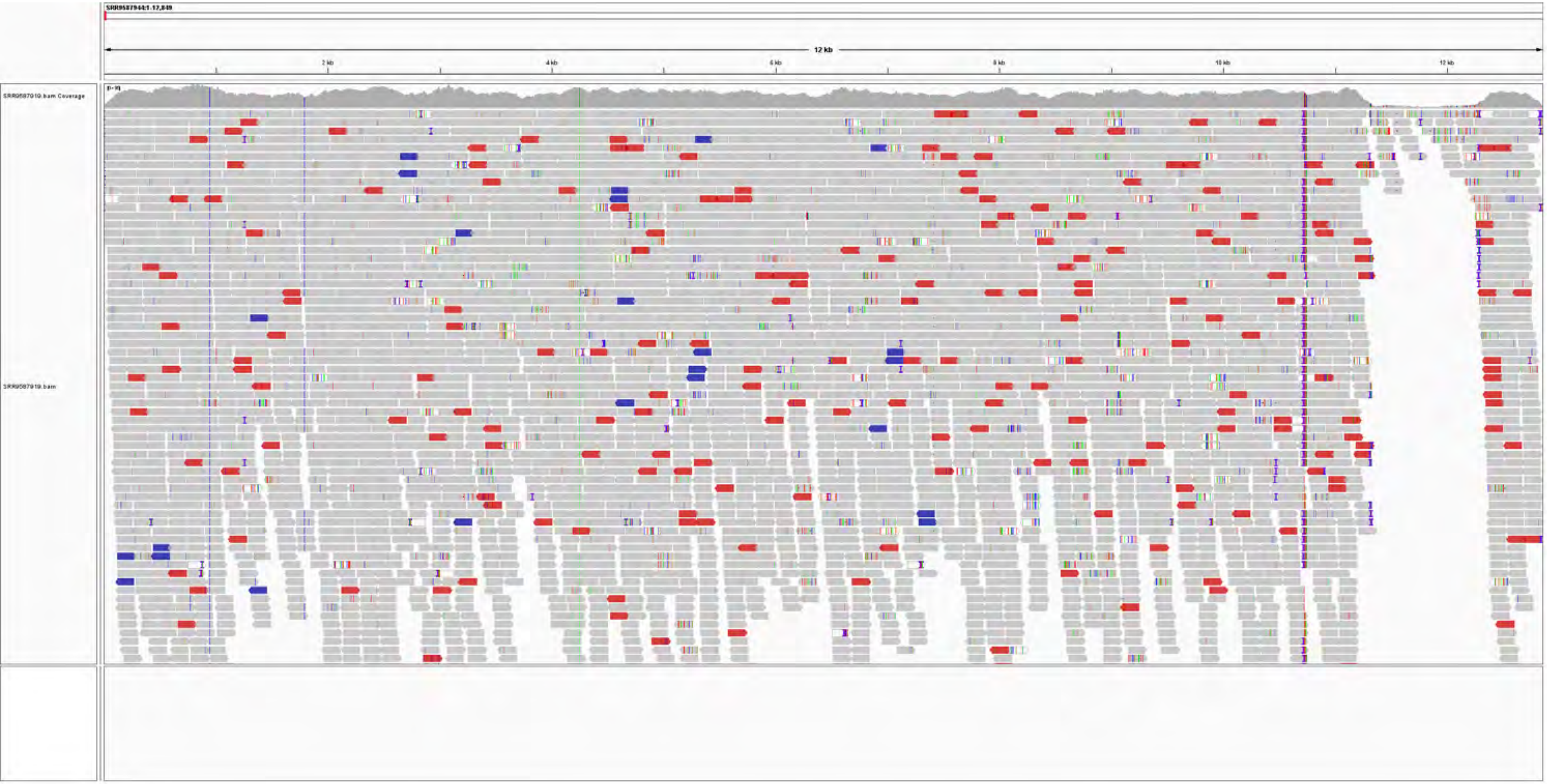

### SRR9587920

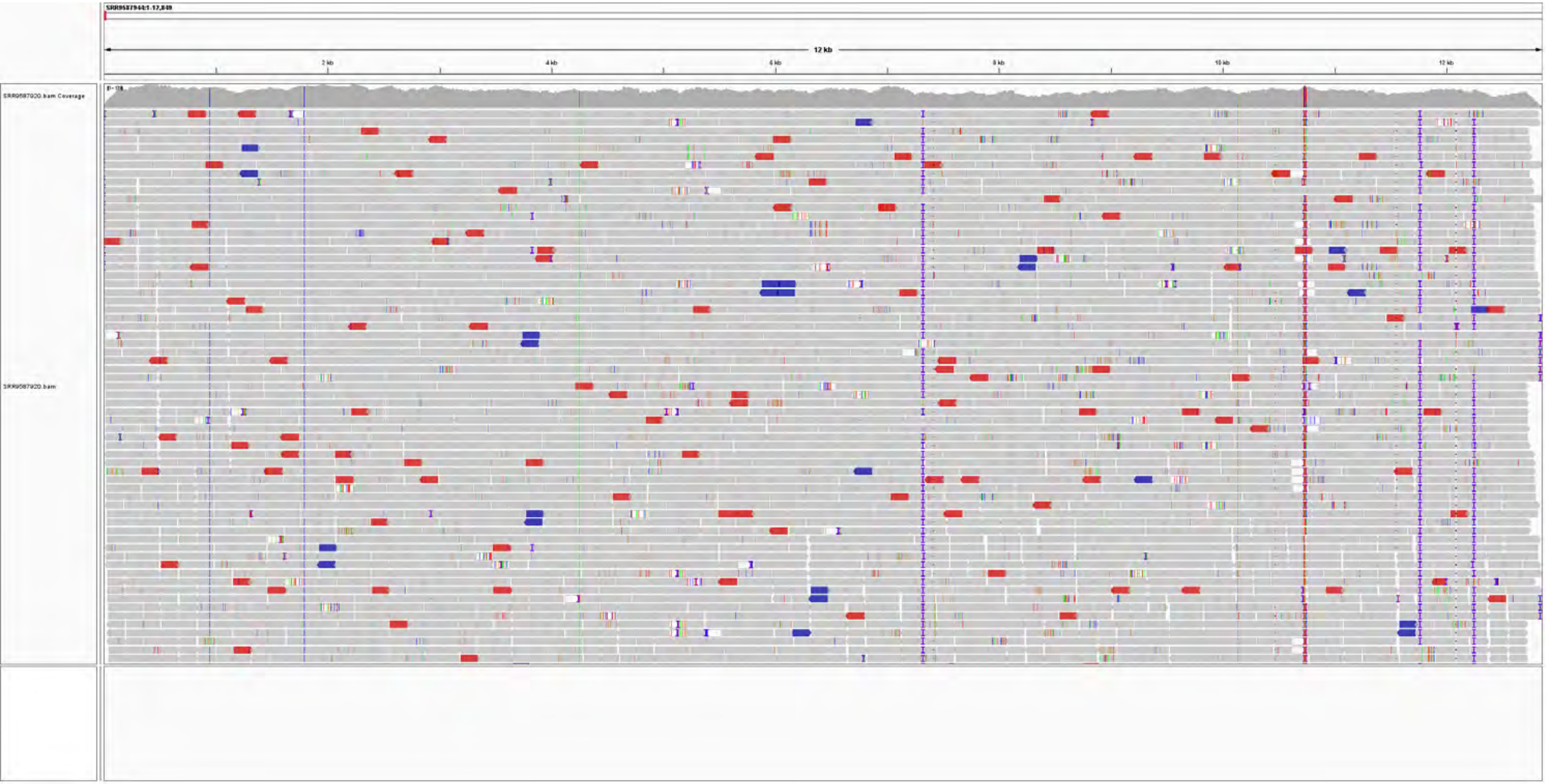

### SRR9587921

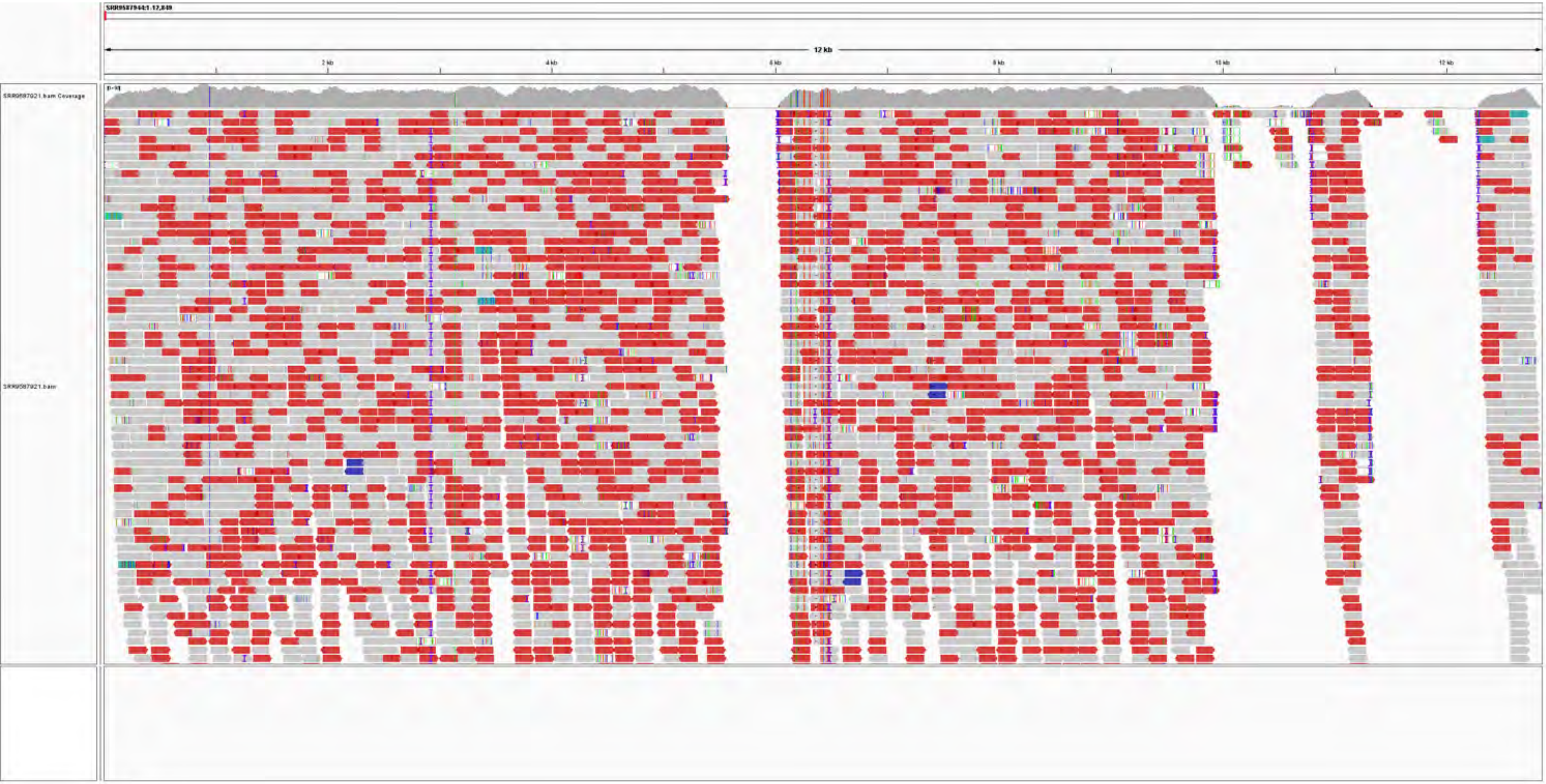

SRR9587944:1-12,849
12 kb
2 kb
4 kb
6 kb
8 kb
10 kb
12 kb
SRR9587921.bam Coverage
SRR9587921.bam

### SRR9587922

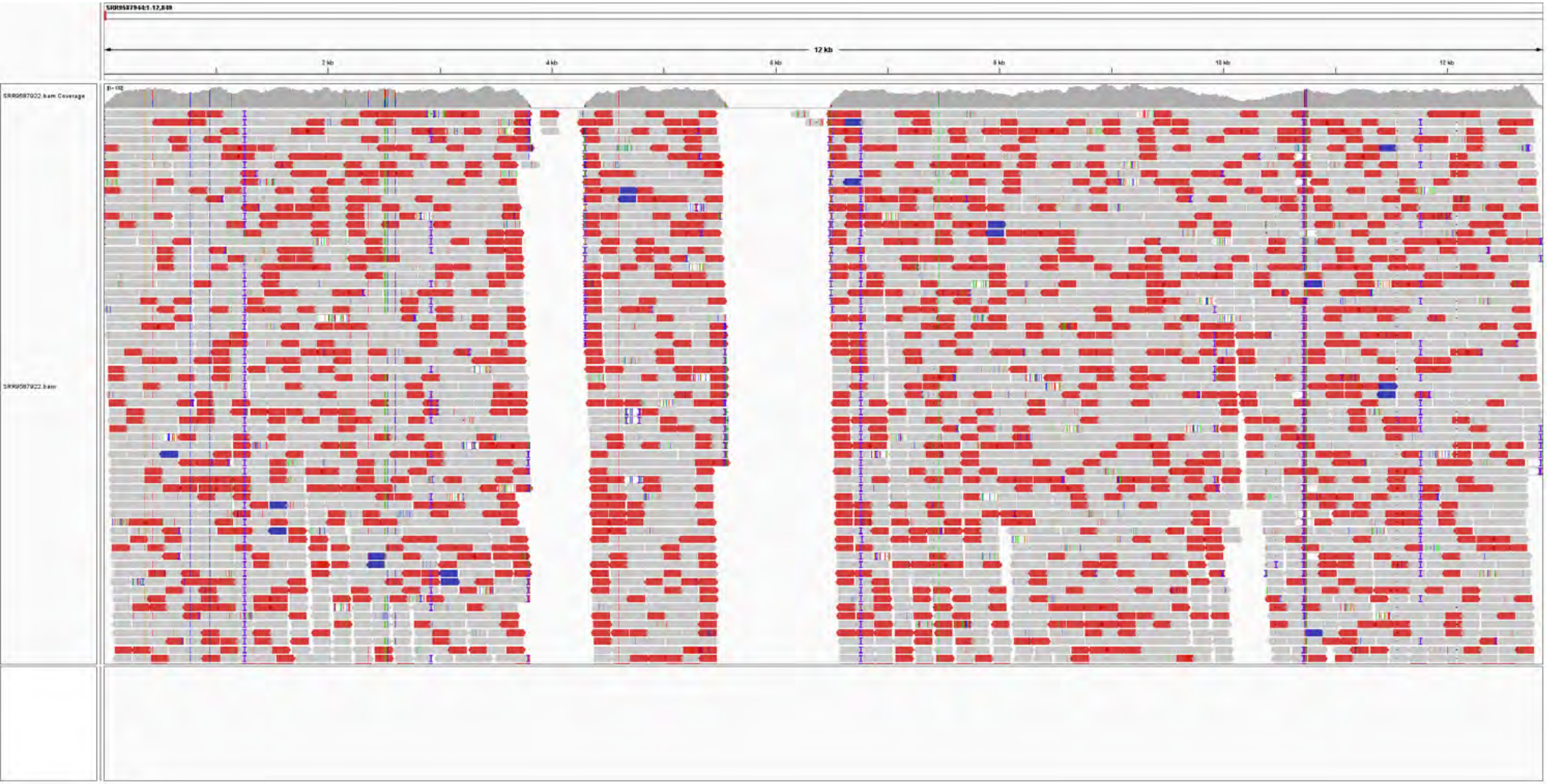
SRR9587944:1-12,849
12 kb
2 kb
4 kb
6 kb
8 kb
10 kb
12 kb
SRR9587922.bam Coverage
SRR9587922.bam

### SRR9587923

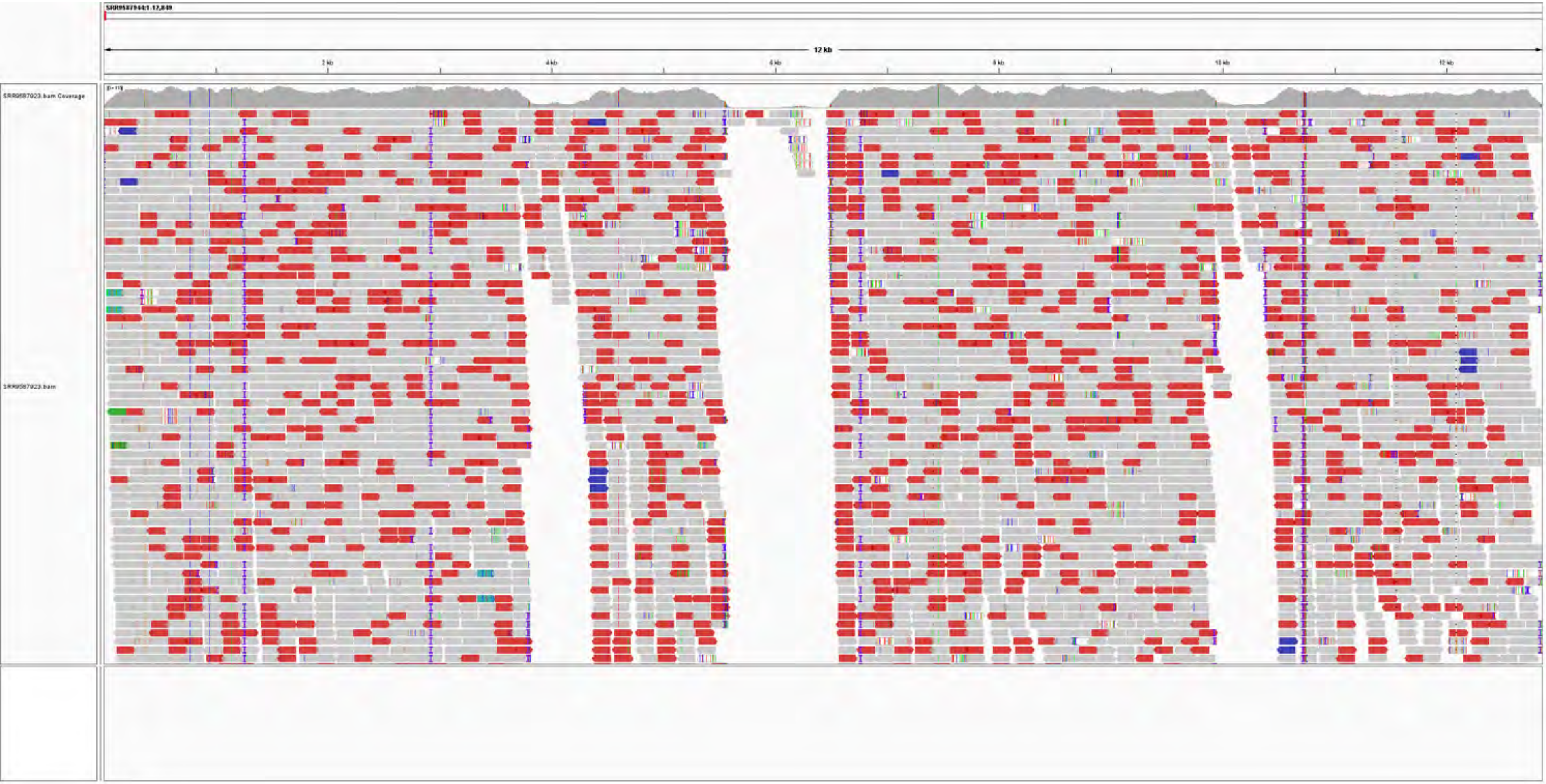

12 kb
2 kb
4 kb
6 kb
8 kb
10 kb
12 kb
SRR9587923.bam Coverage
SRR9587923.bam

### SRR9587924

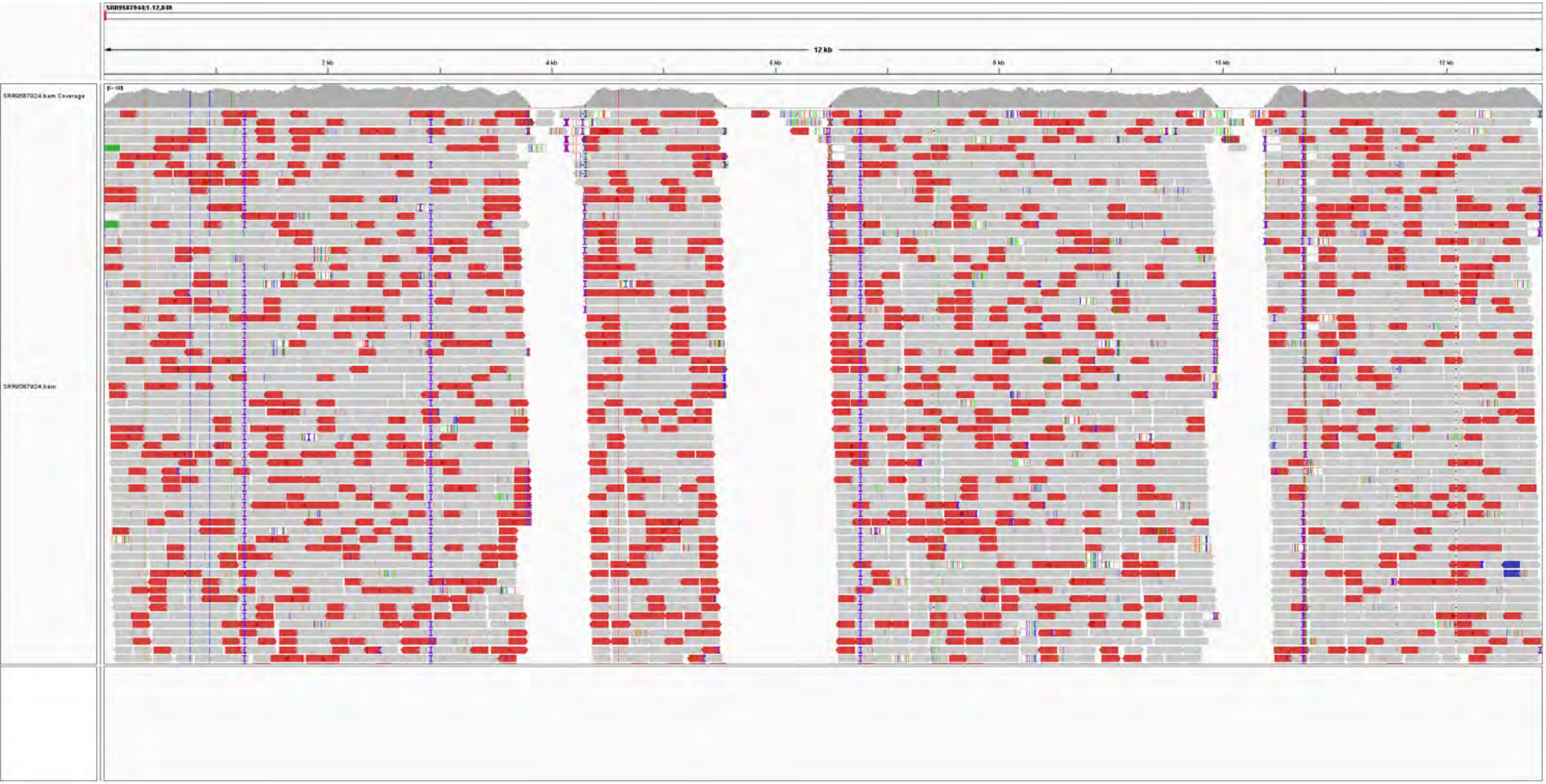

### SRR9587927

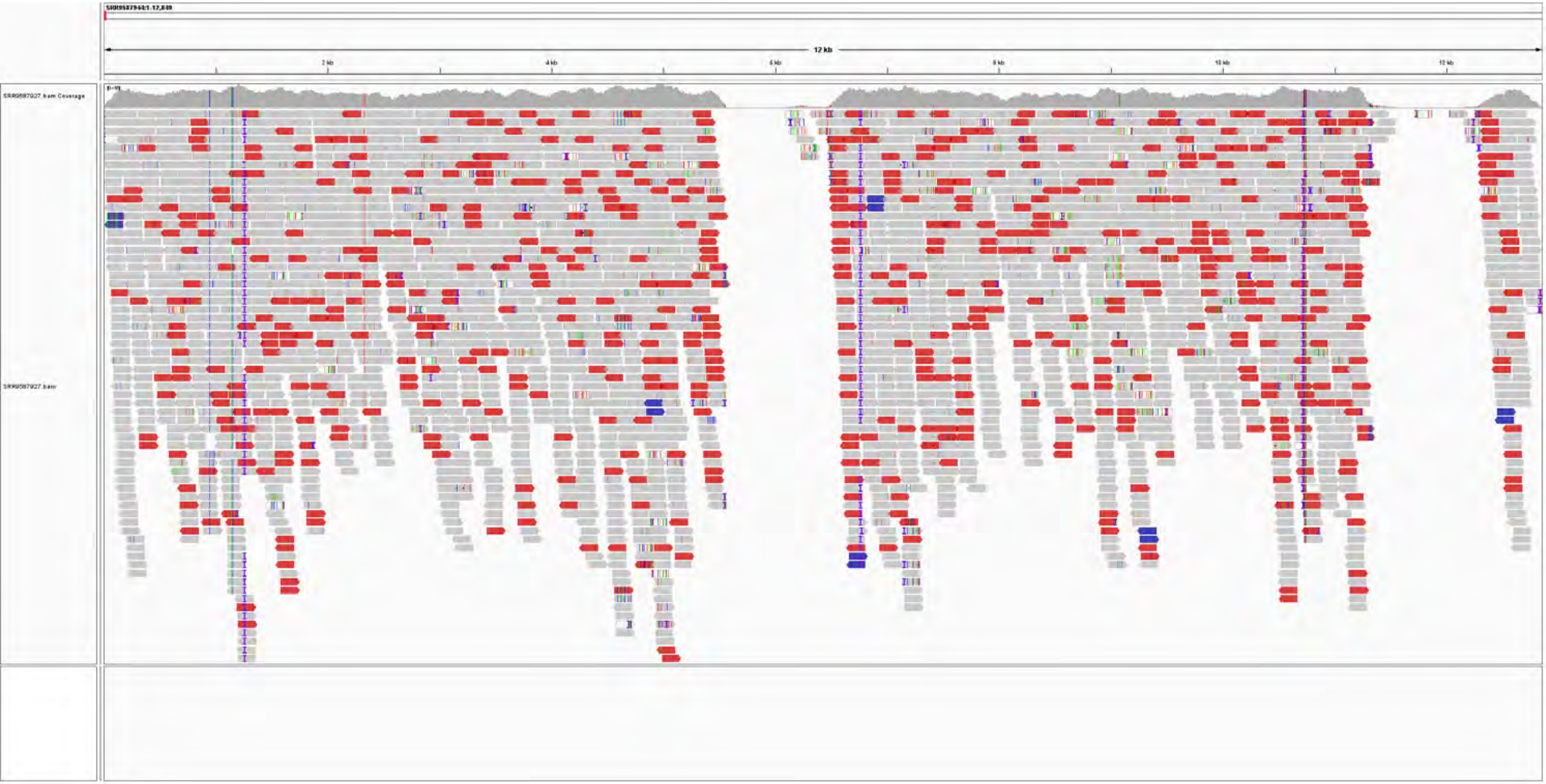

### SRR9587928

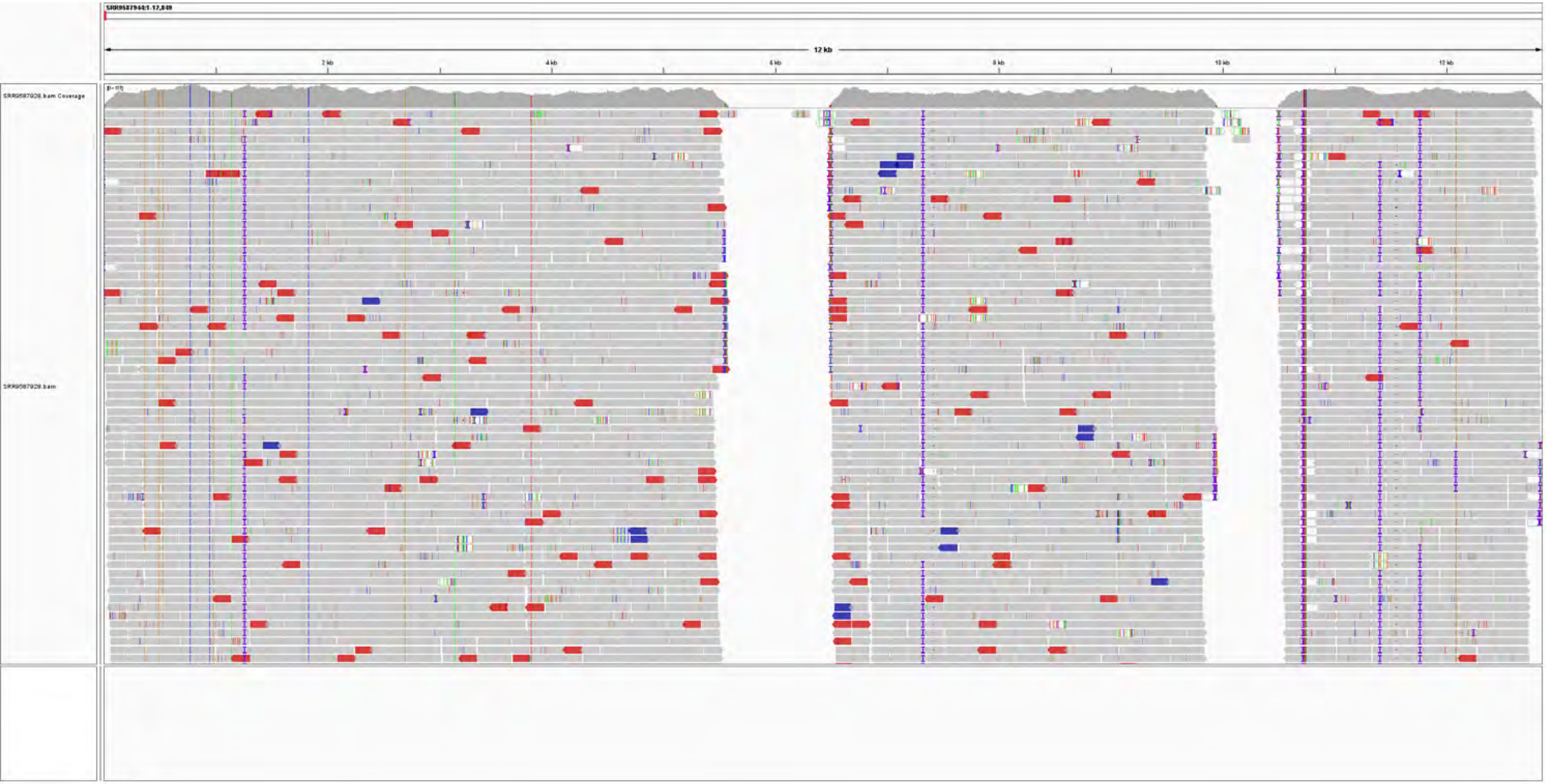

SRR9587944:1-17,849
12 kb
2 kb
4 kb
6 kb
8 kb
10 kb
12 kb
SRR9587928.bam Coverage
SRR9587928.bam

### SRR9587929

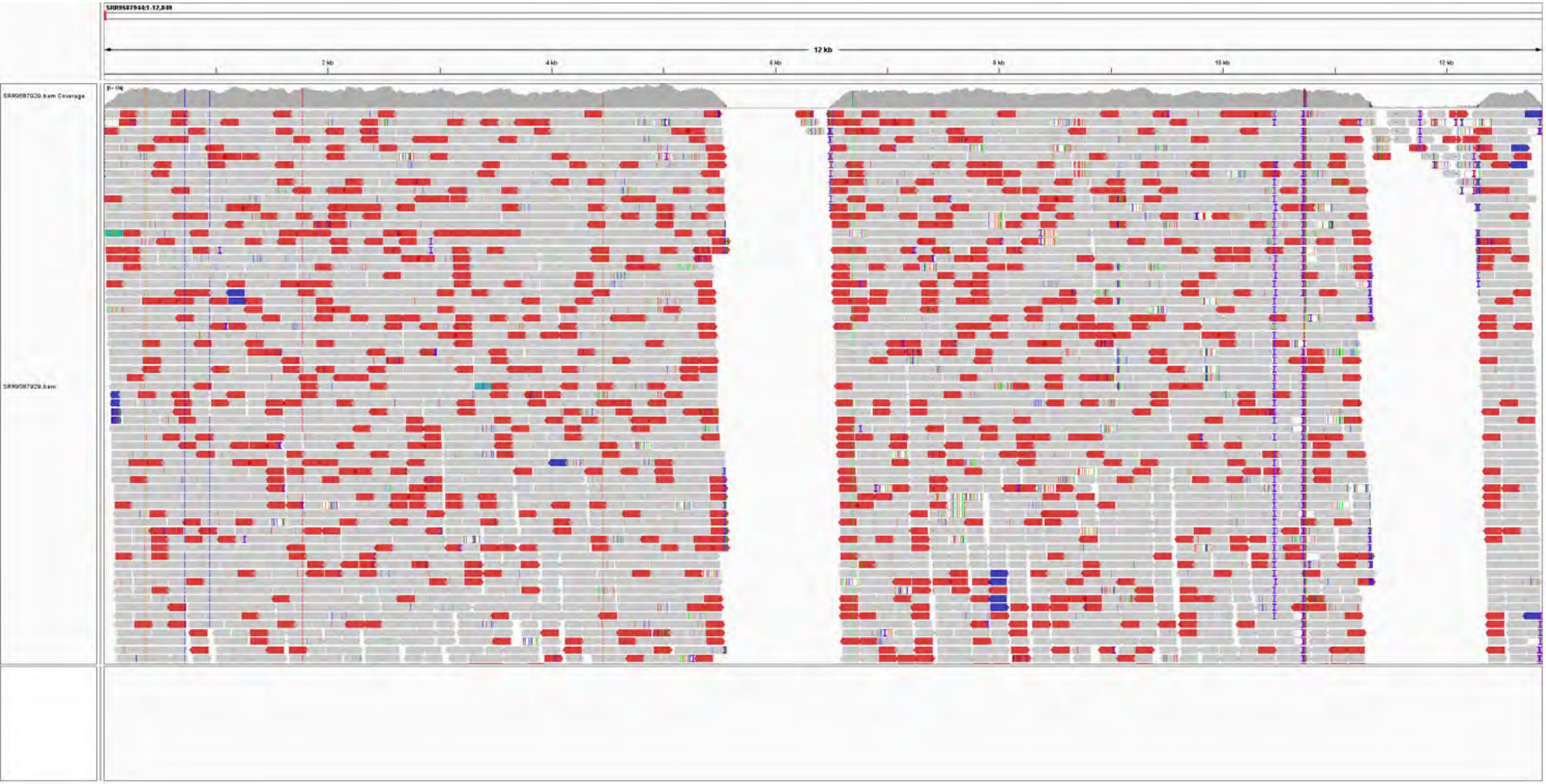

### SRR9587930

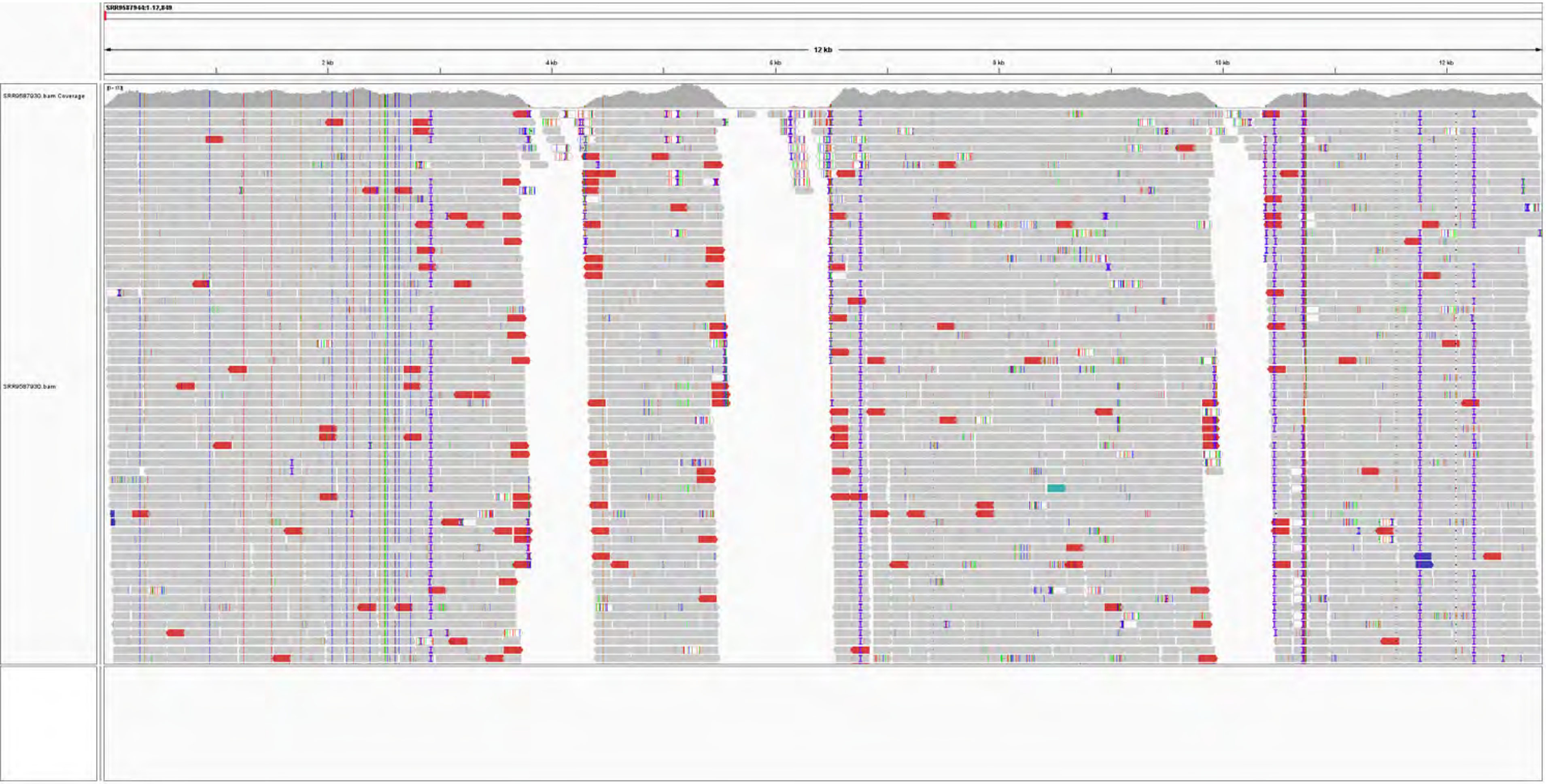

12 kb
2 kb
4 kb
6 kb
8 kb
10 kb
12 kb
SRR9587930.bam Coverage
SRR9587930.bam

### SRR9587931

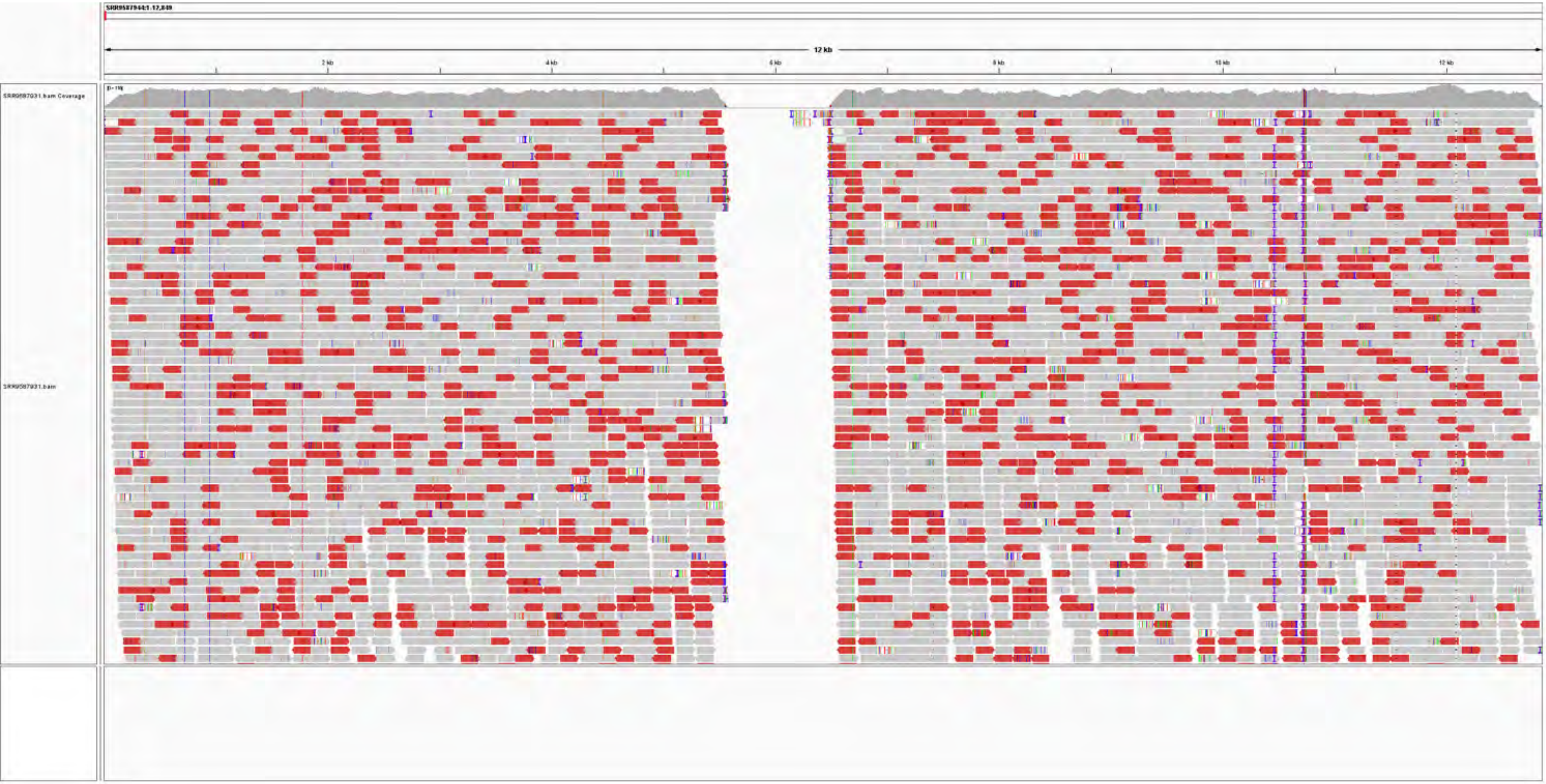

### SRR9587932

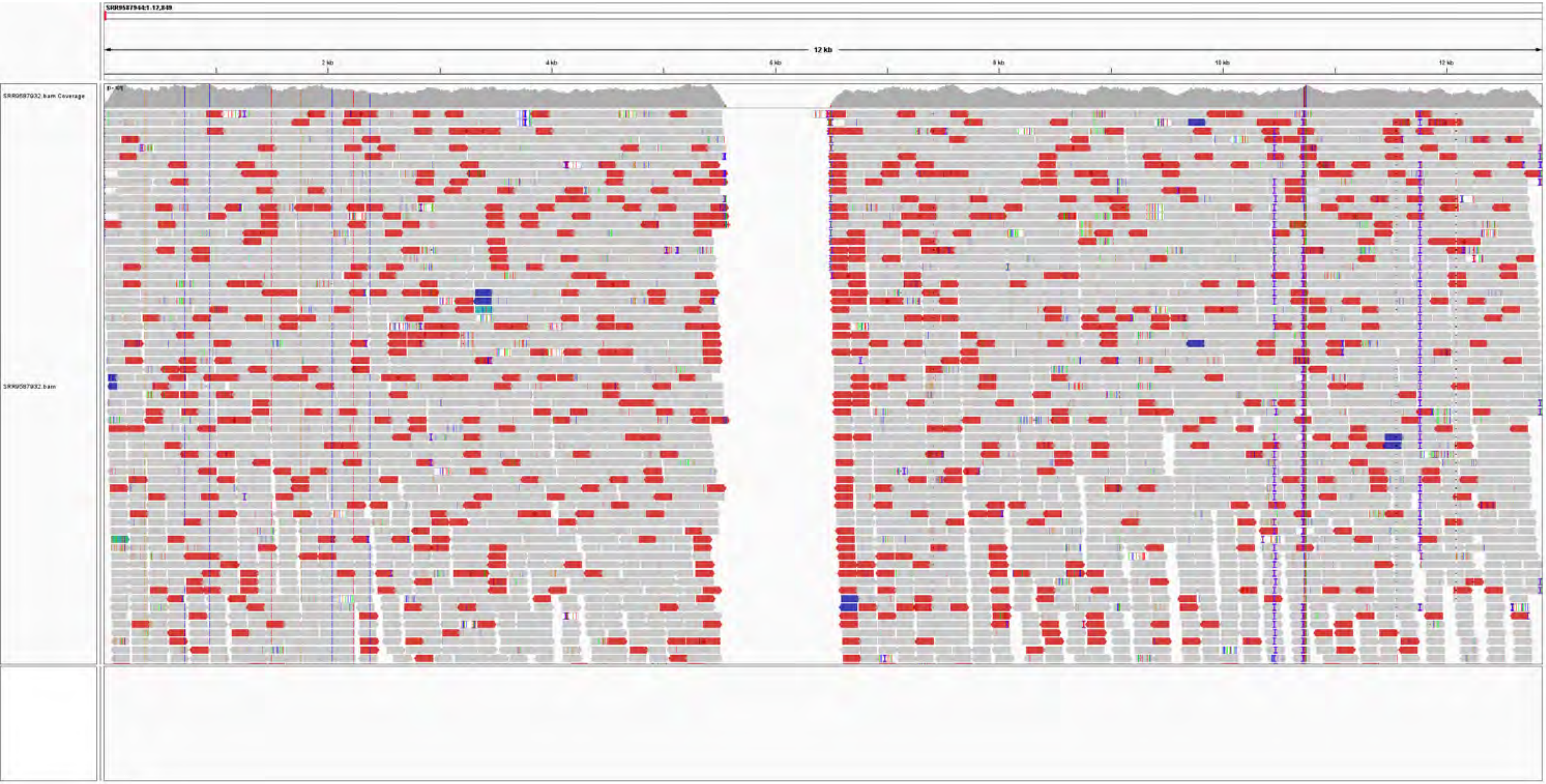

### SRR9587933

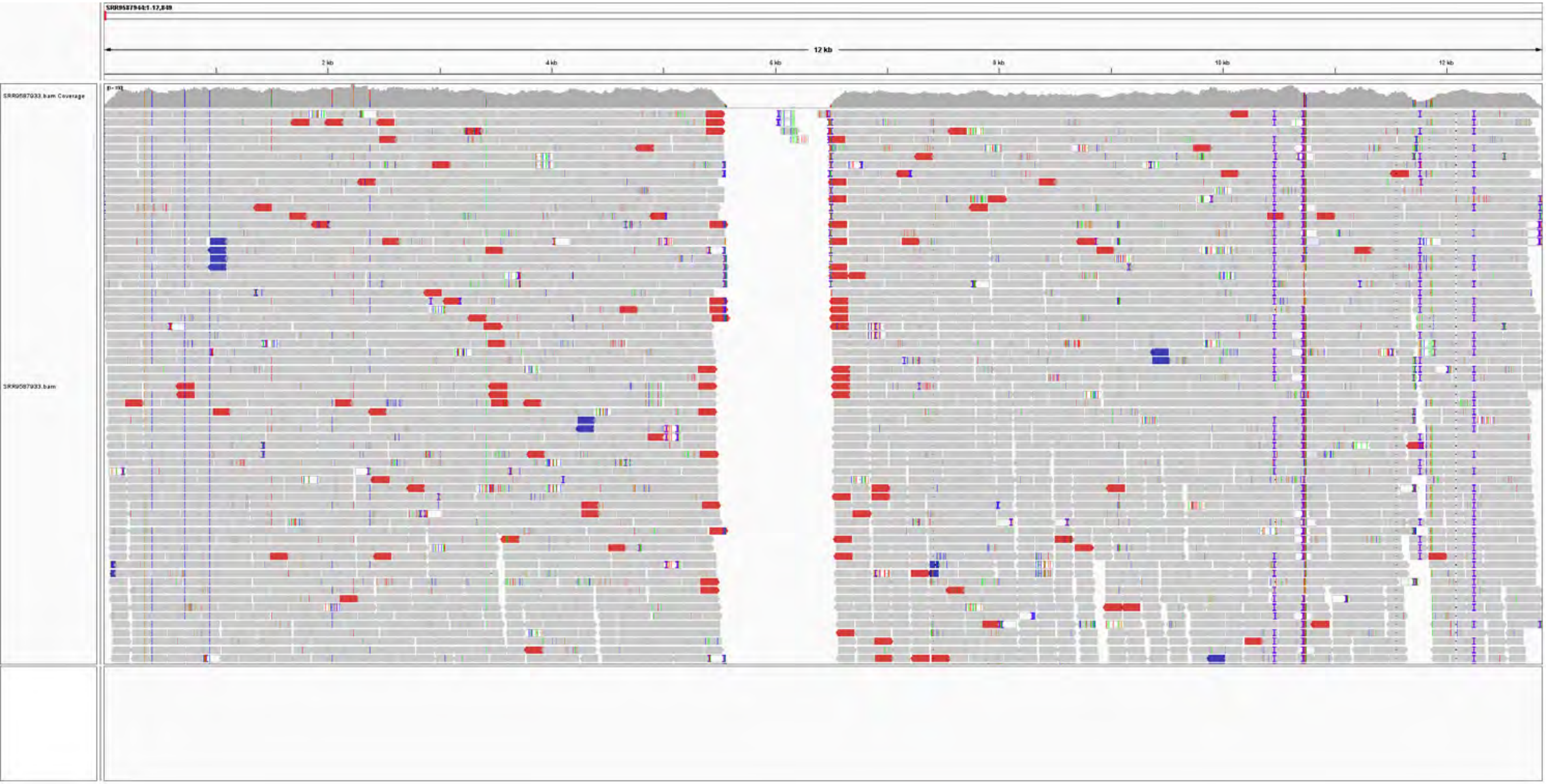

12 kb
2 kb
4 kb
6 kb
8 kb
10 kb
12 kb
SRR9587933.bam Coverage
SRR9587933.bam

### SRR9587934

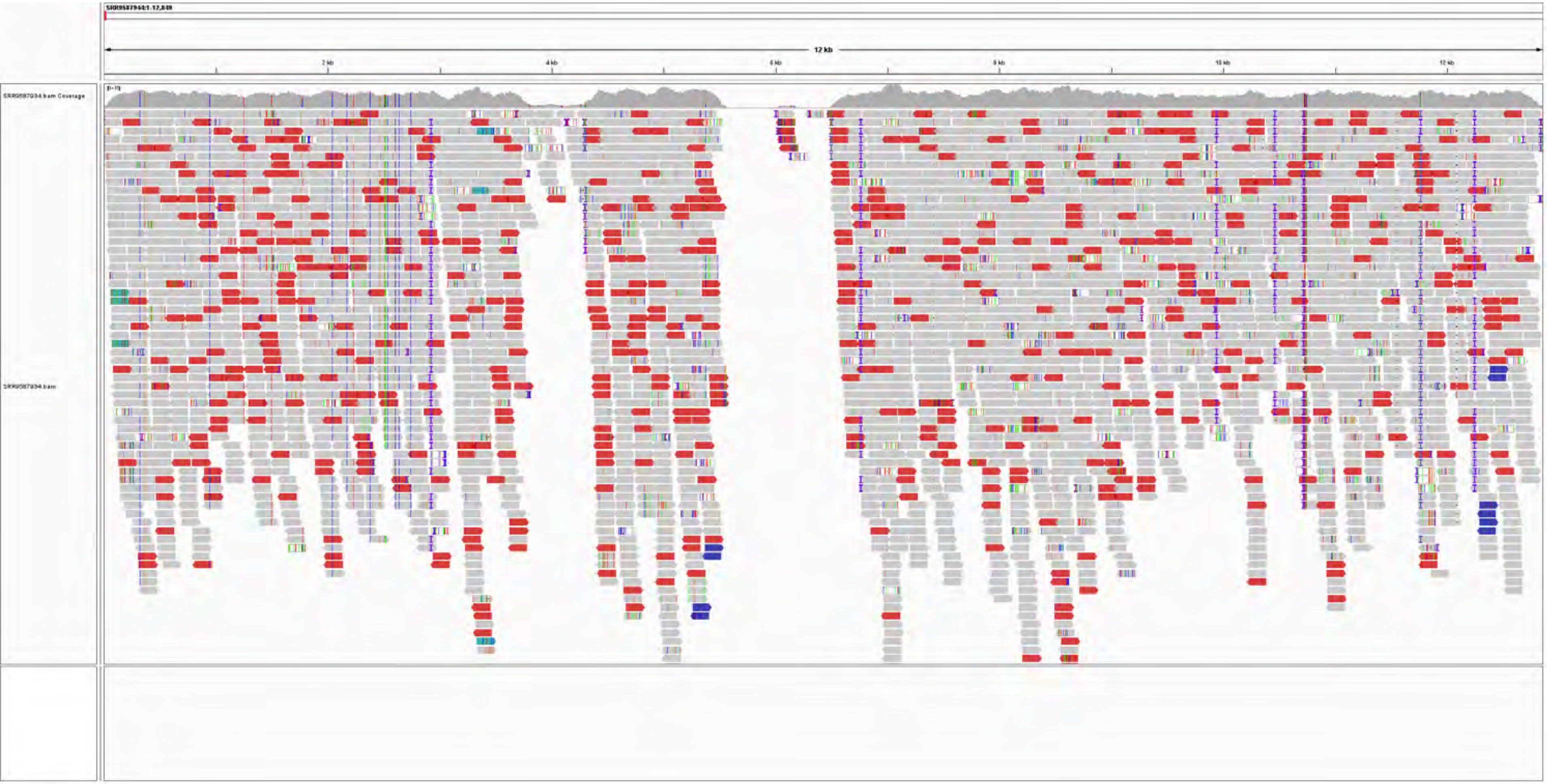

### SRR9587936

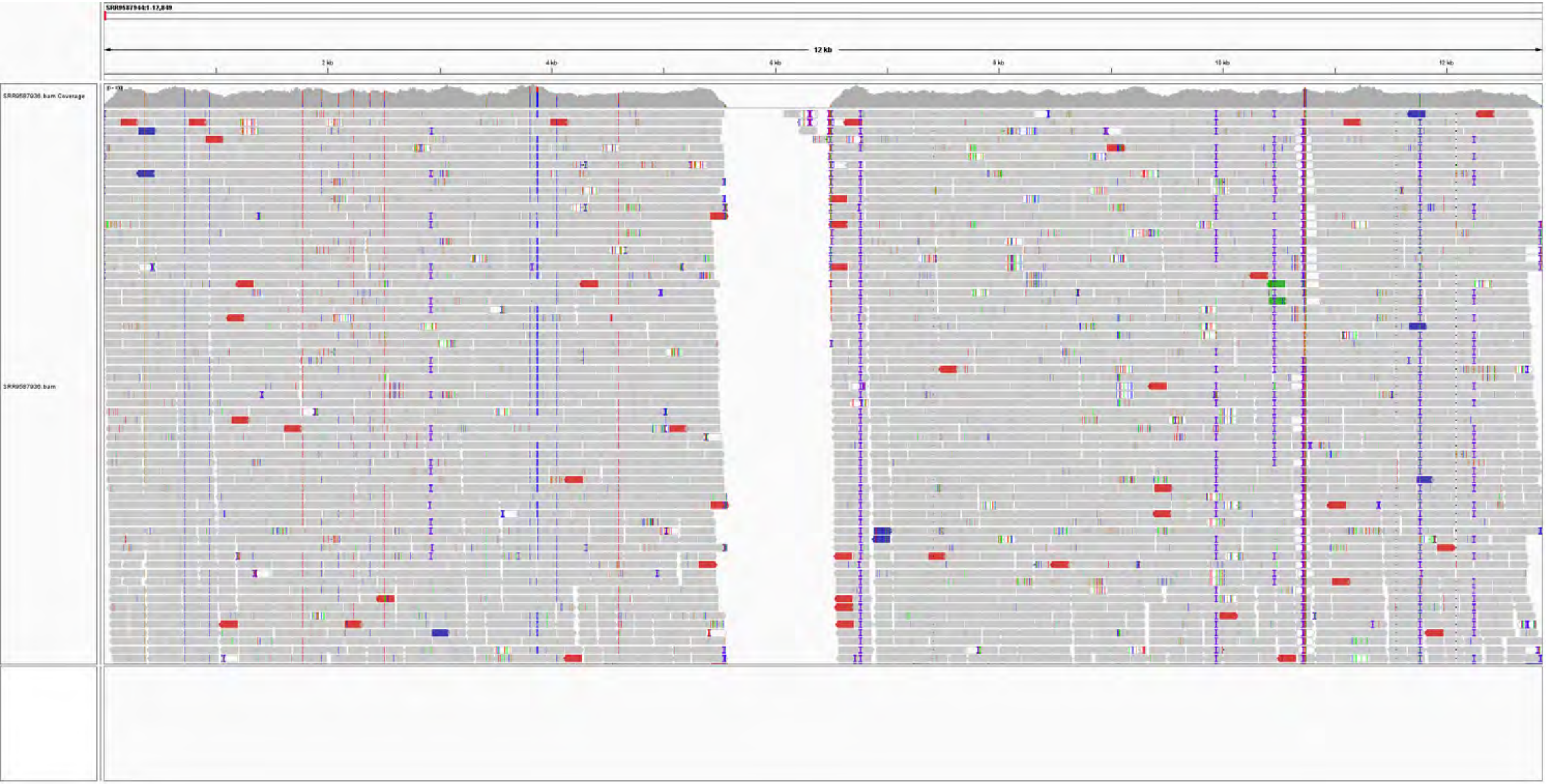

SRR9587944:1-17,849
12 kb
2 kb
4 kb
6 kb
8 kb
10 kb
12 kb
SRR9587936.bam Coverage
SRR9587936.bam

### SRR9587937

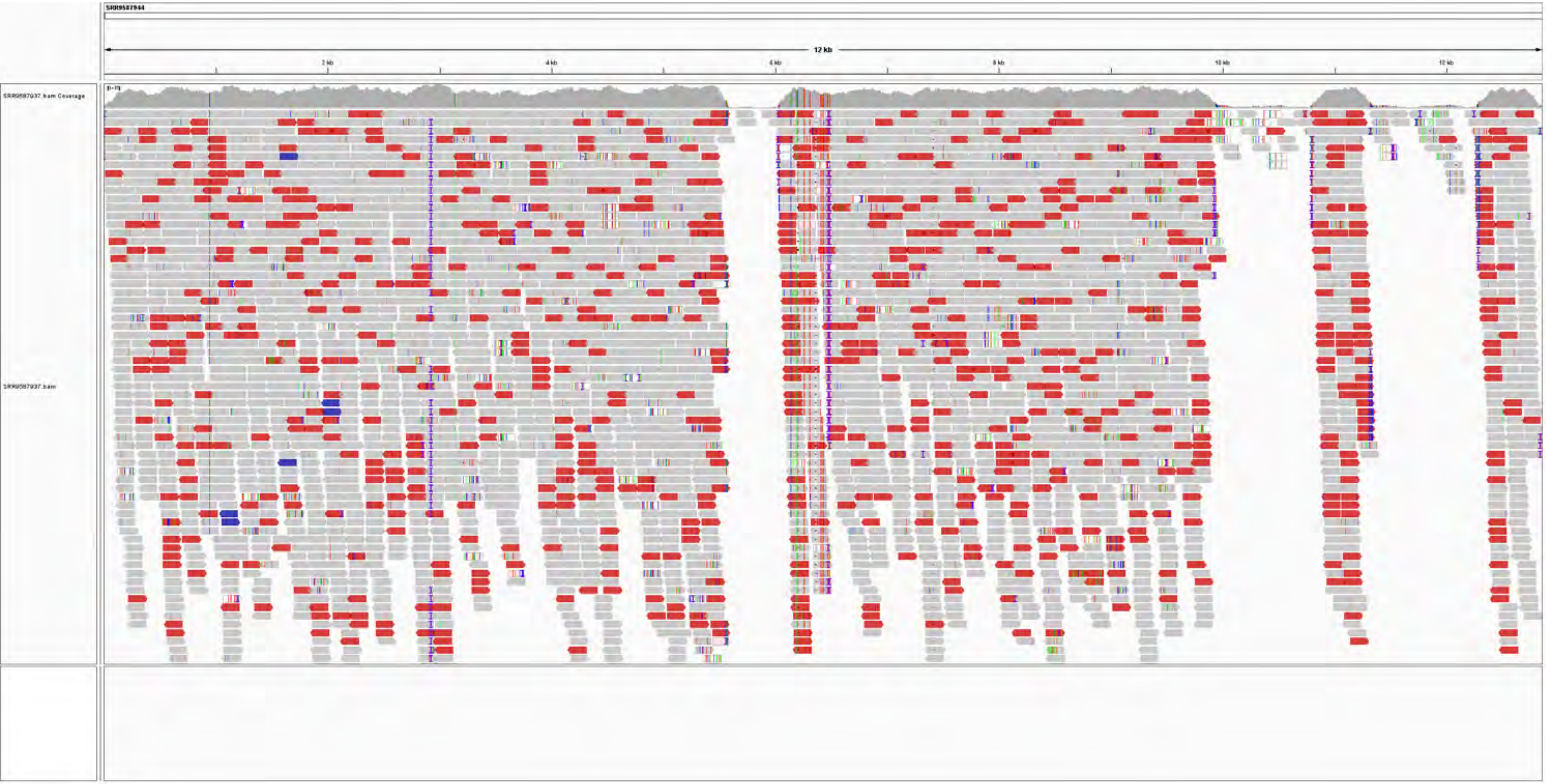

SRR9587944
12 kb
2 kb
4 kb
6 kb
8 kb
10 kb
12 kb
SRR9587937.bam Coverage
SRR9587937.bam

### SRR9587938

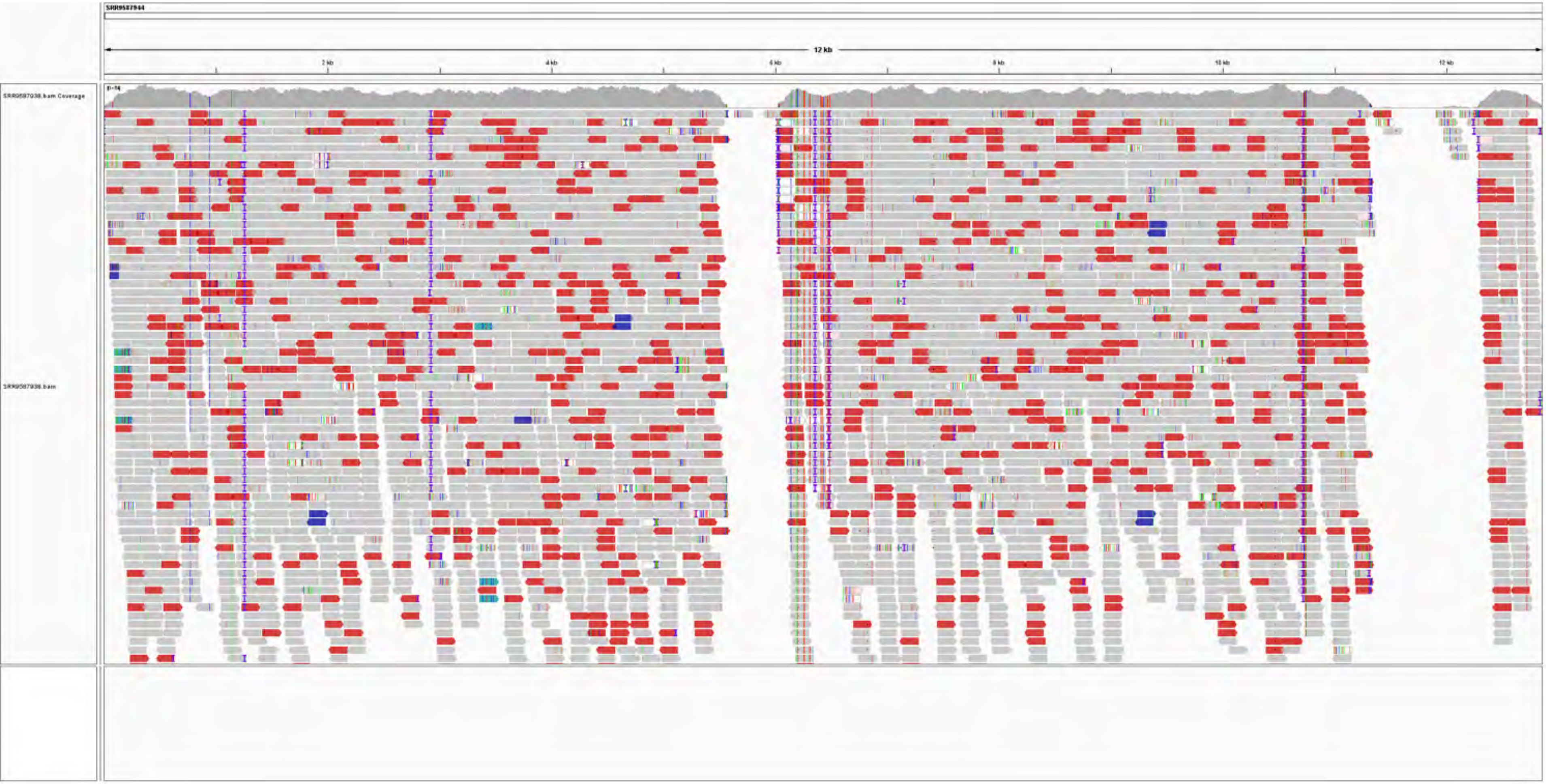

### SRR9587939

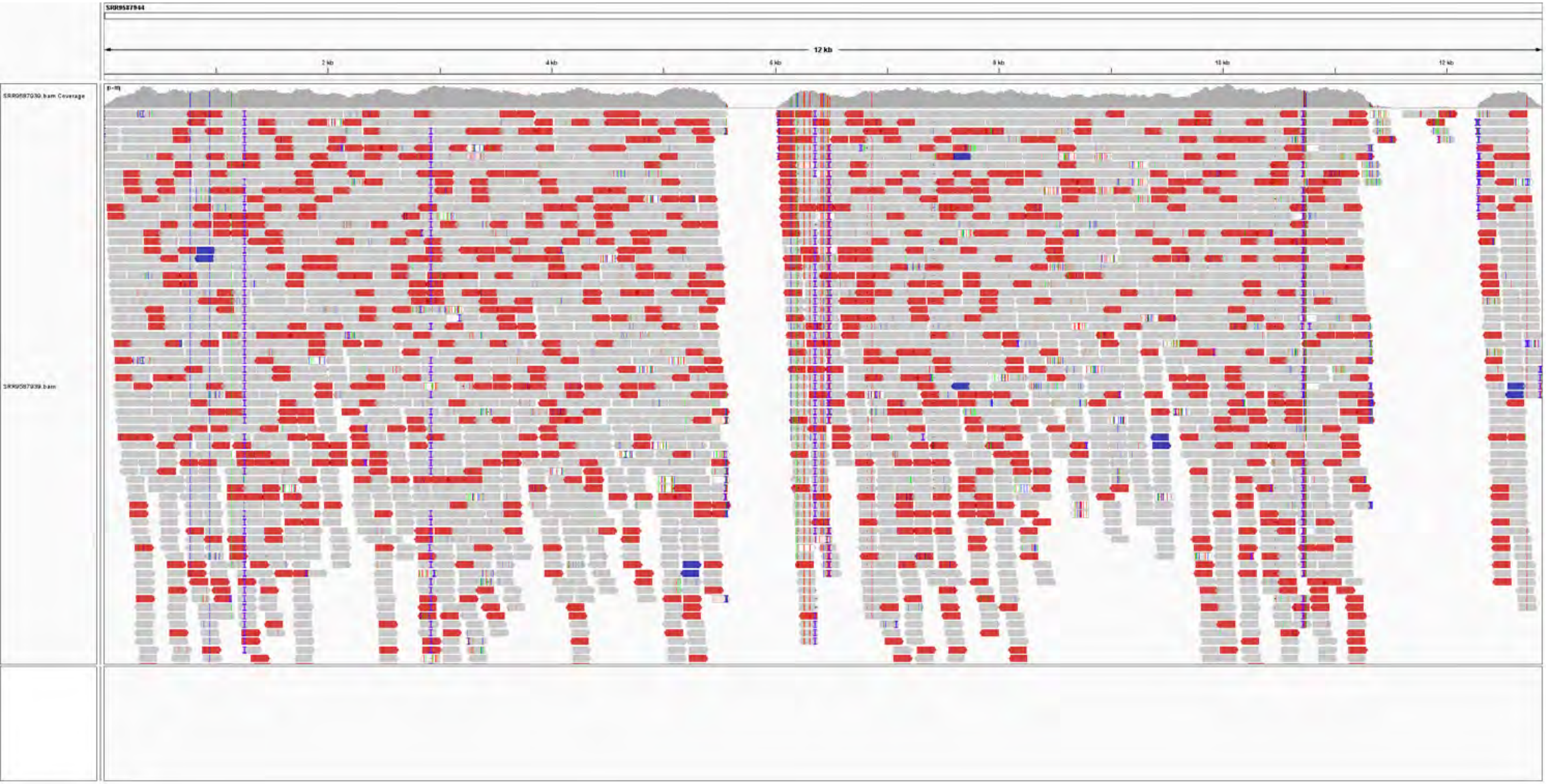

### SRR9587941

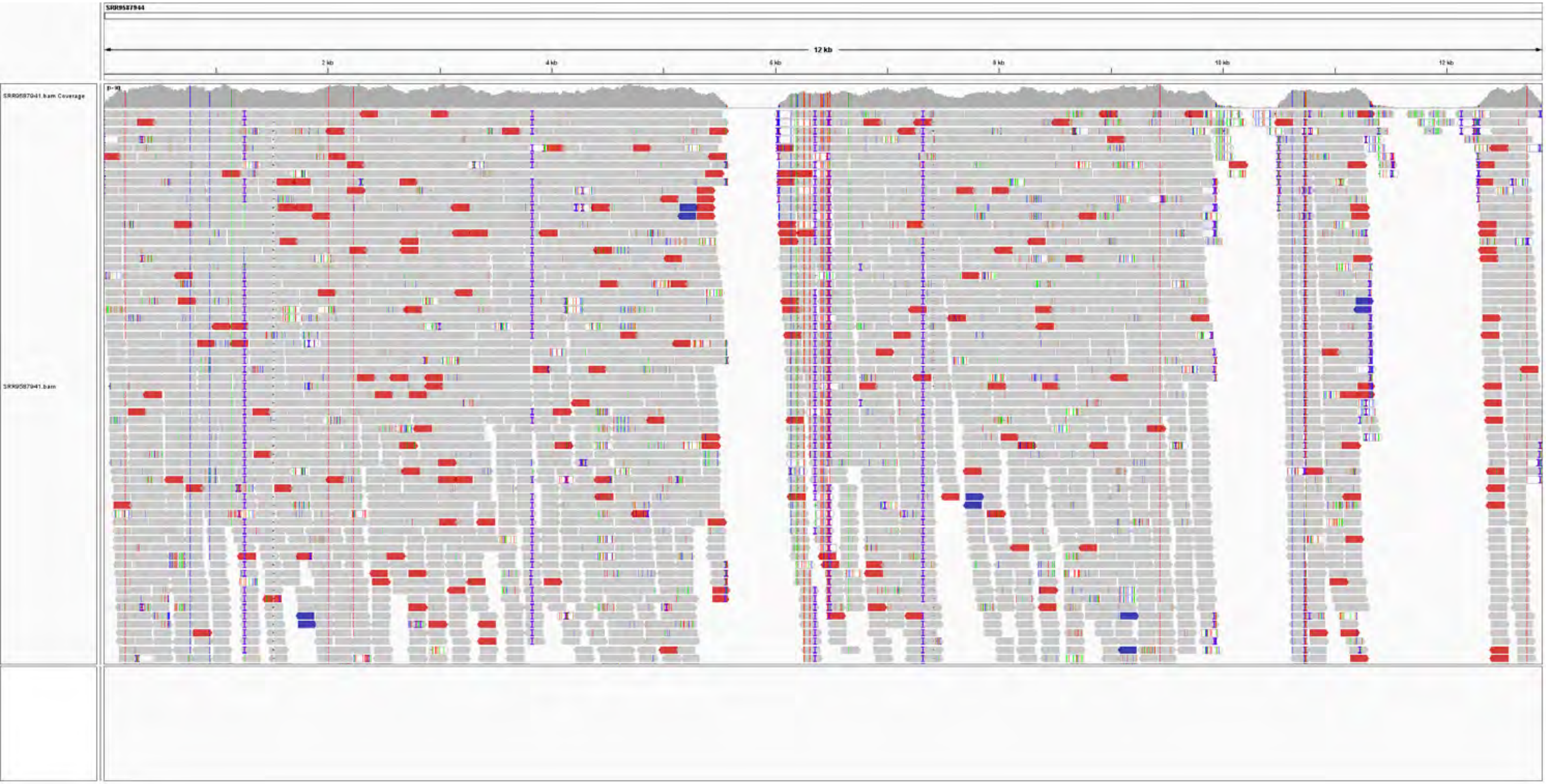

12 kb
2 kb
4 kb
6 kb
8 kb
10 kb
12 kb
[0 - 90]
SRR9587941.bam Coverage
SRR9587941.bam

### SRR9587942

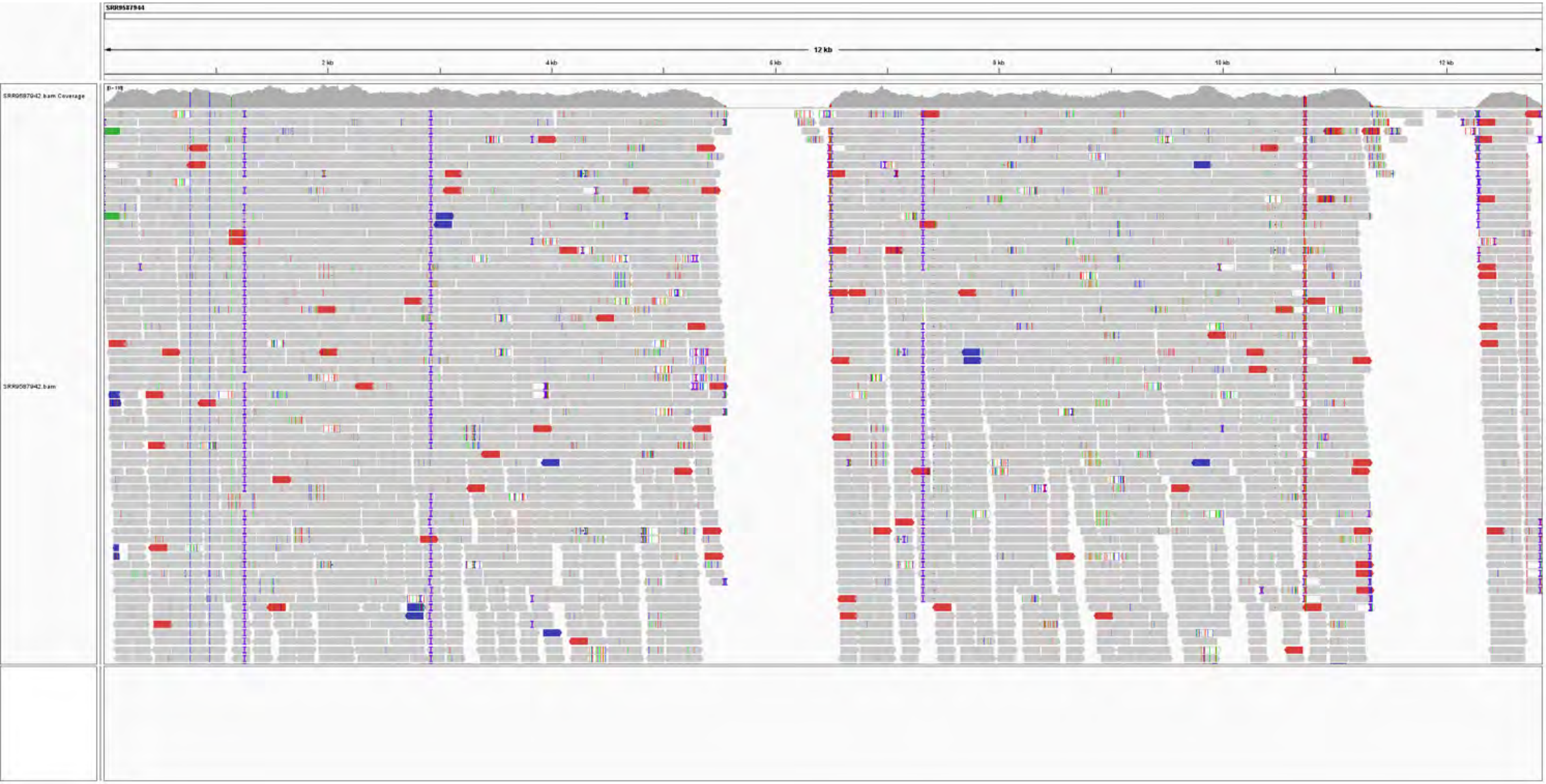
SRR9587944
12 kb
2 kb
4 kb
6 kb
8 kb
10 kb
12 kb
SRR9587942.bam Coverage
SRR9587942.bam

### SRR9587943

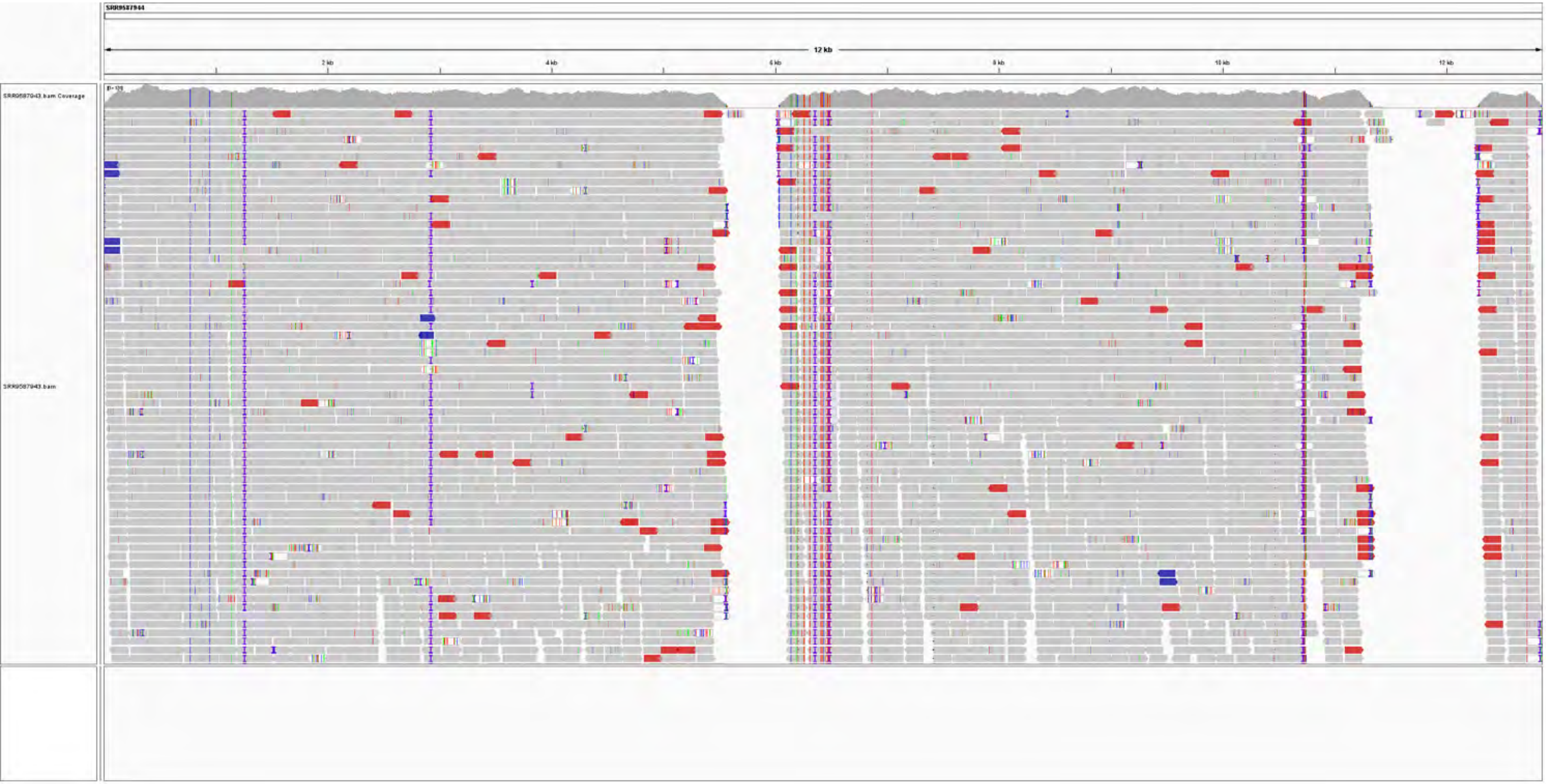

### SRR9587944

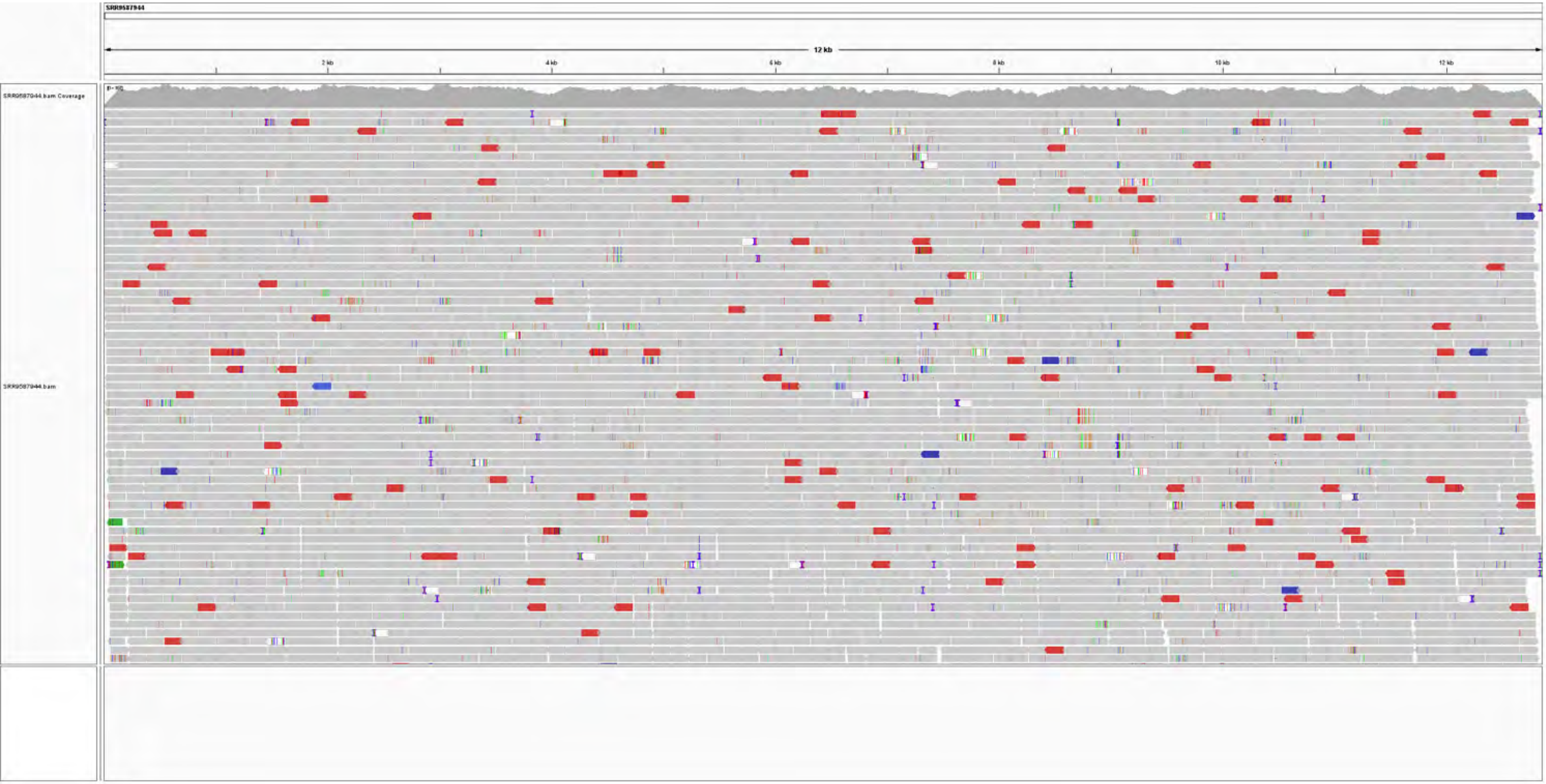

SRR9587944
12 kb
2 kb
4 kb
6 kb
8 kb
10 kb
12 kb
SRR9587944.bam Coverage
SRR9587944.bam

### SRR9587946

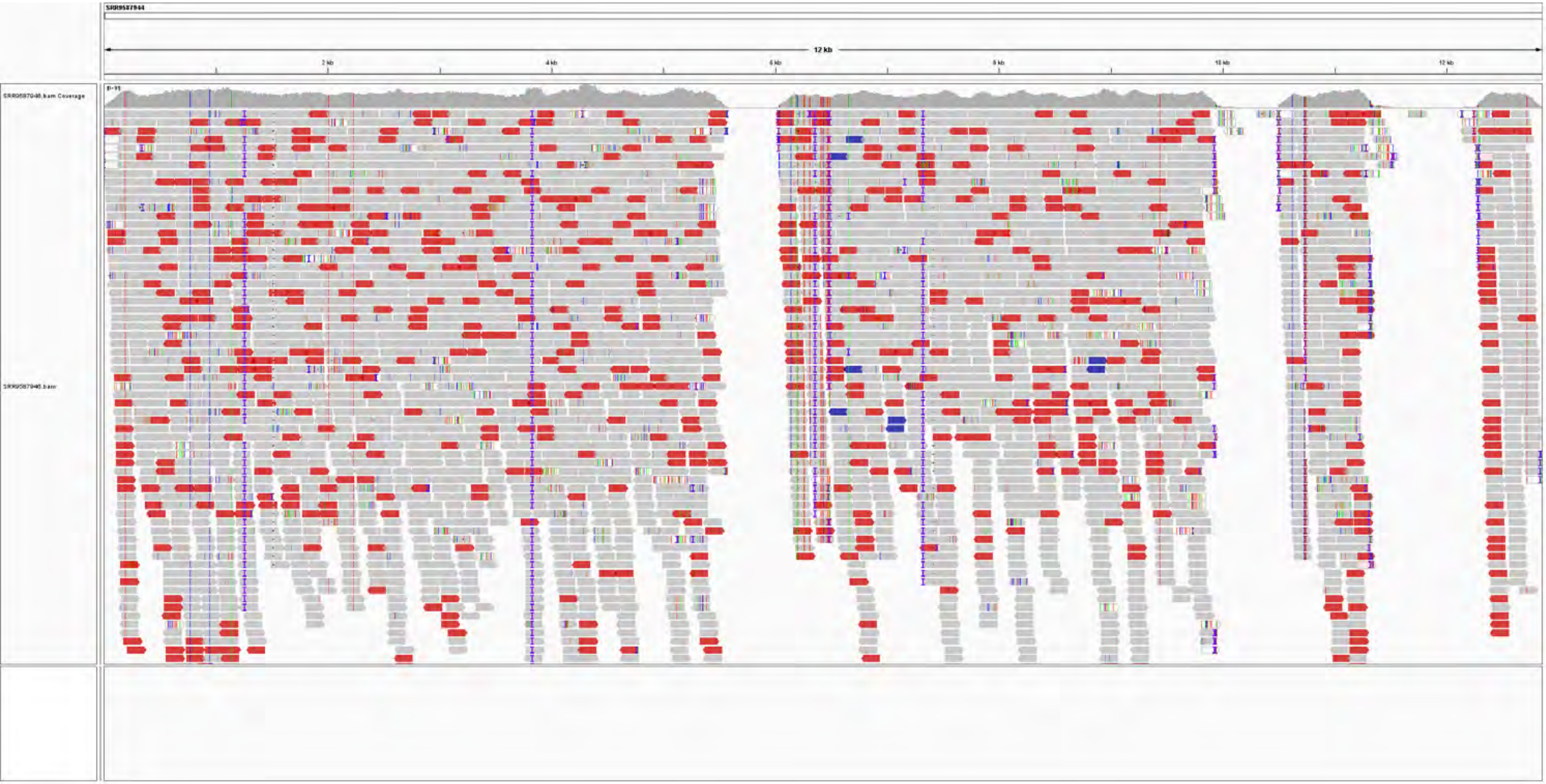

### SRR9587947

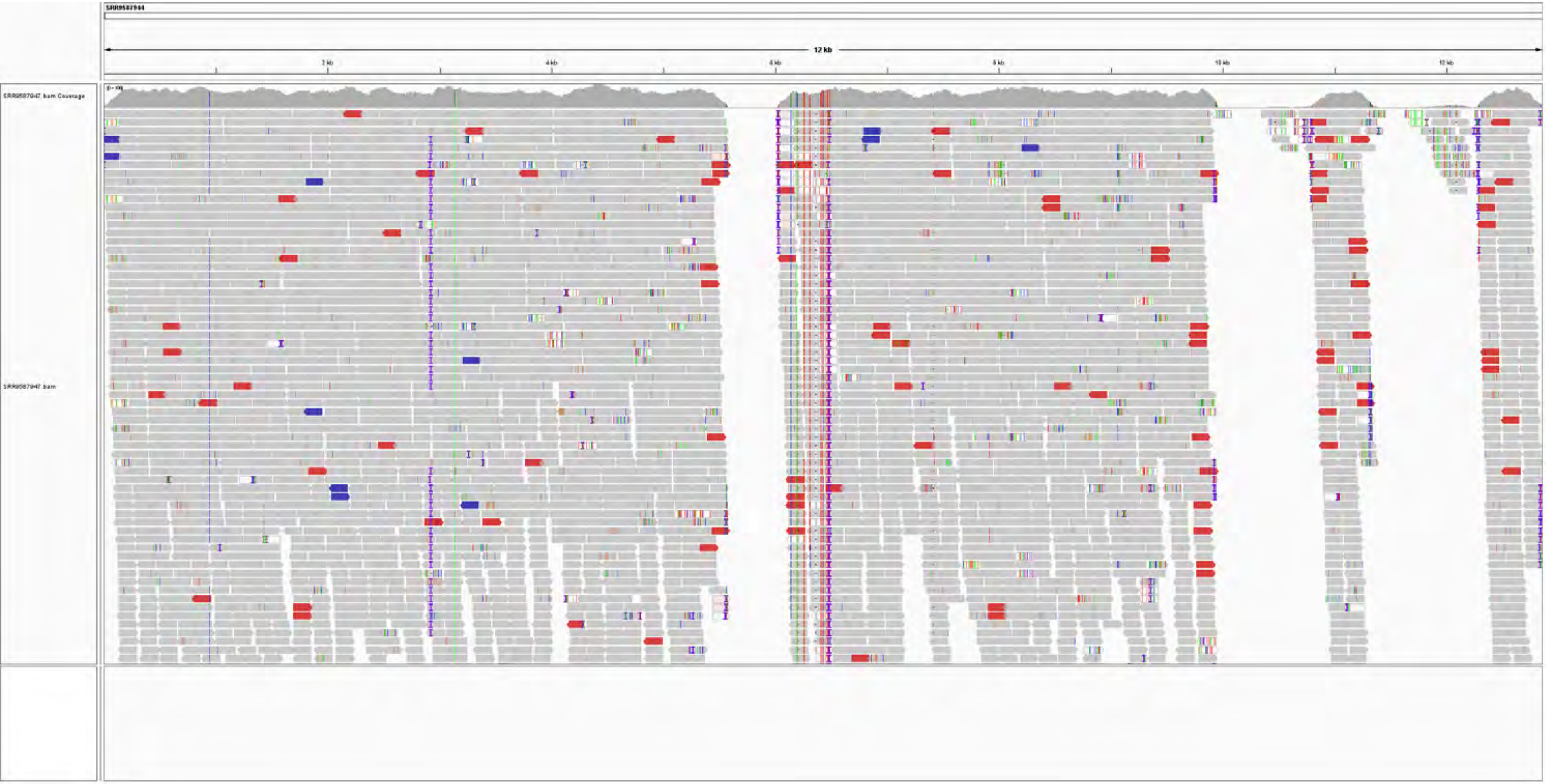

### SRR9587948

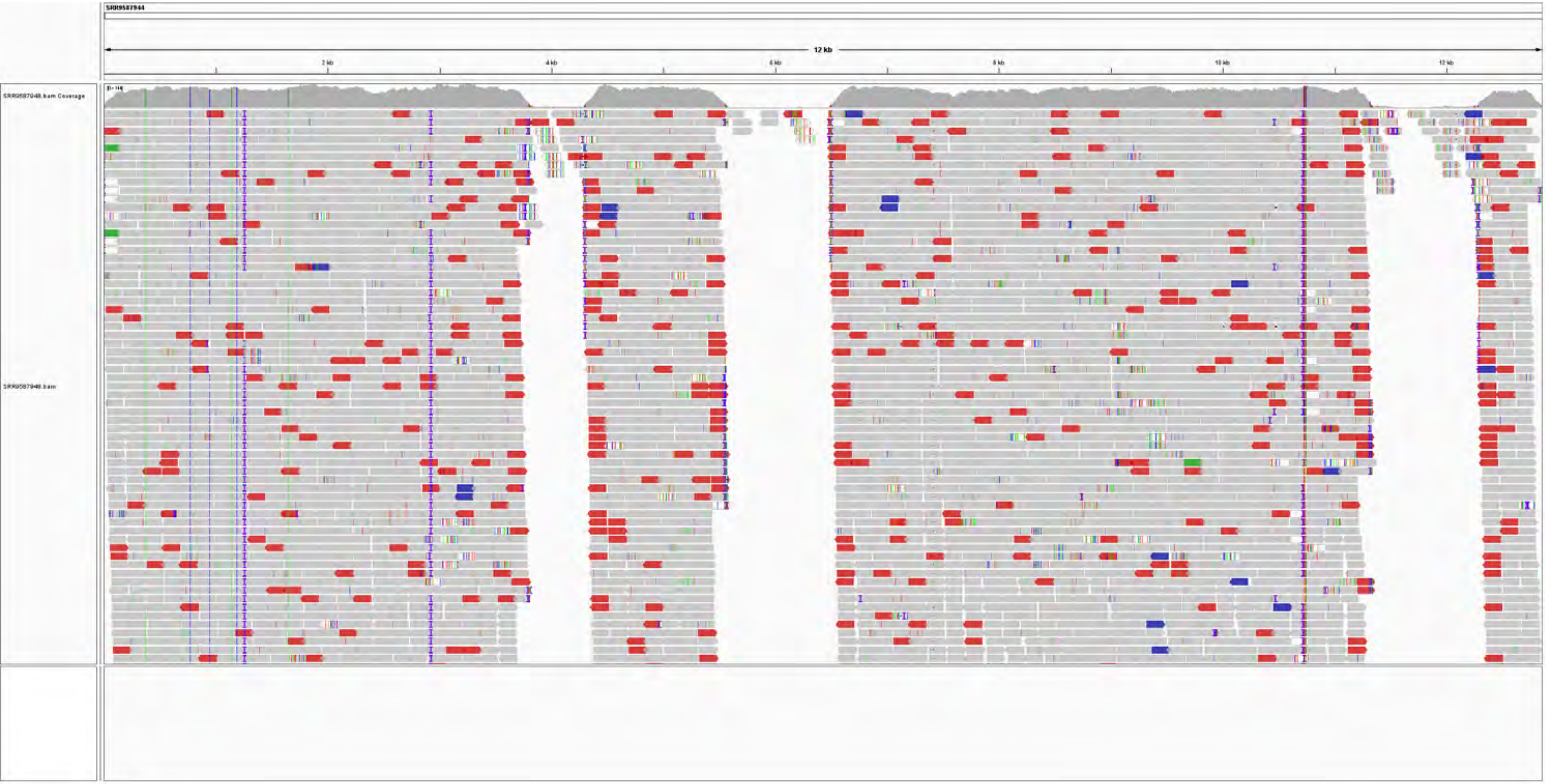

### SRR9587949

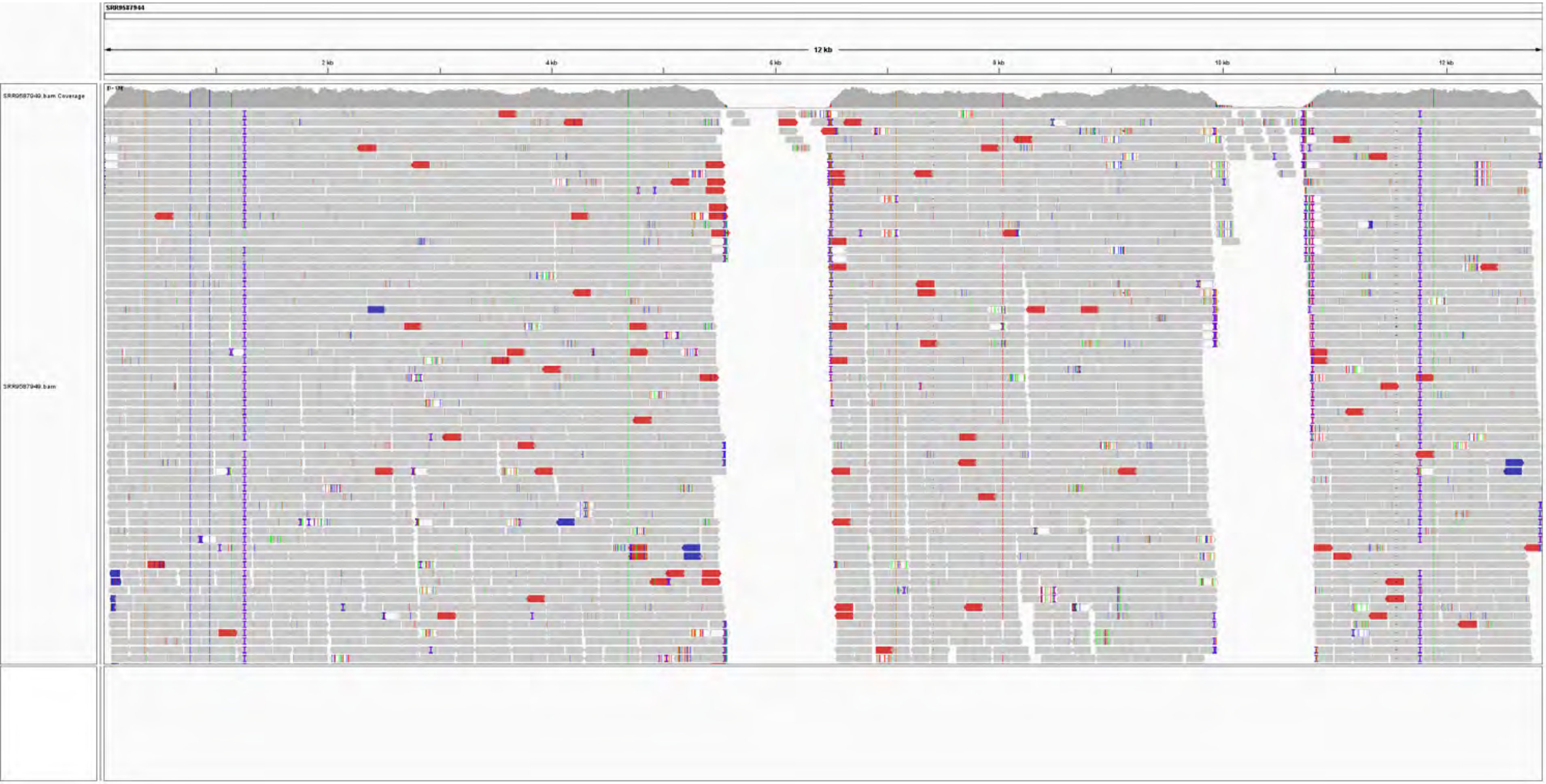

12 kb
2 kb
4 kb
6 kb
8 kb
10 kb
12 kb
SRR9587949.bam Coverage
SRR9587949.bam

### SRR9587950

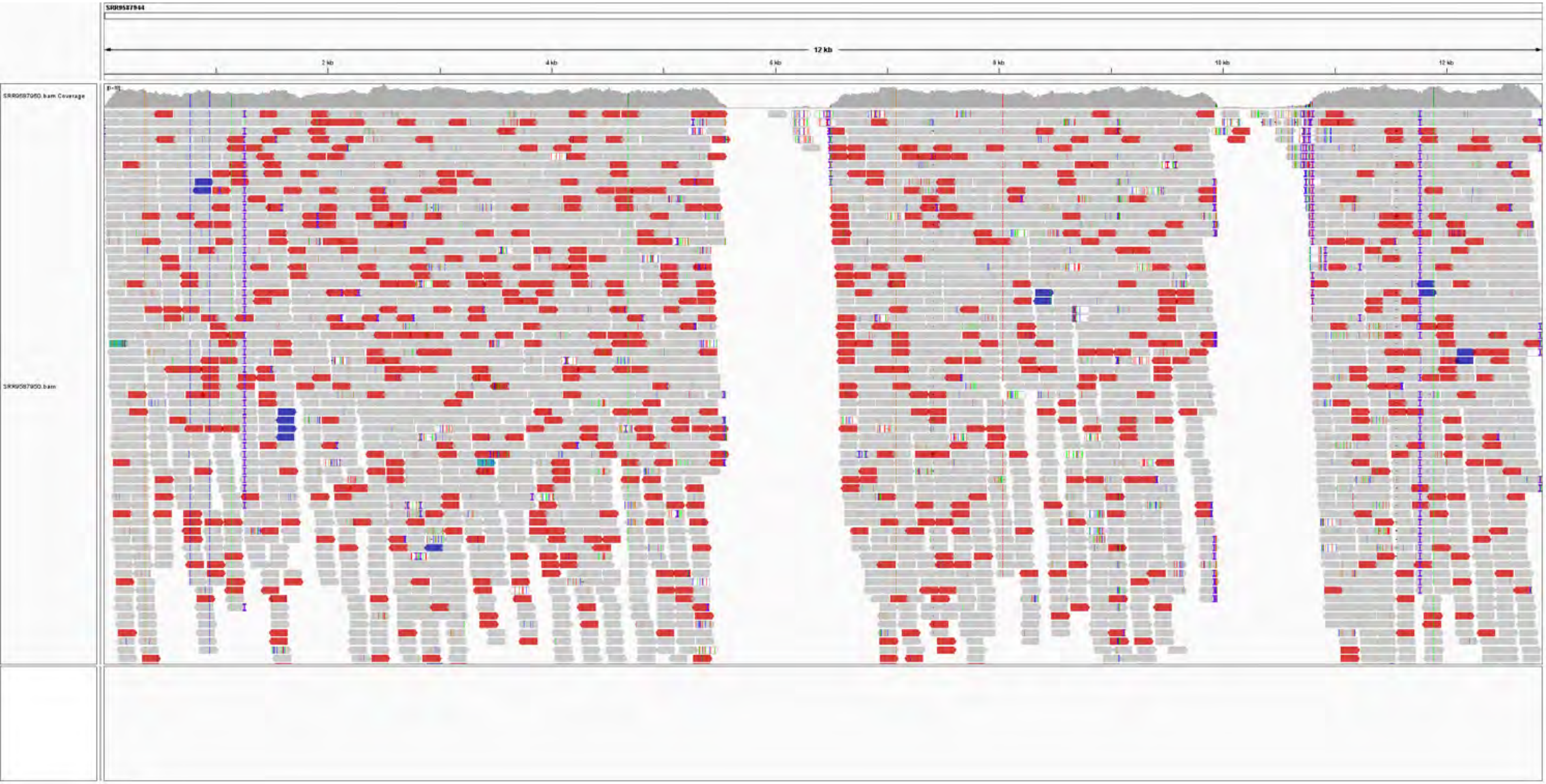

### SRR9587951

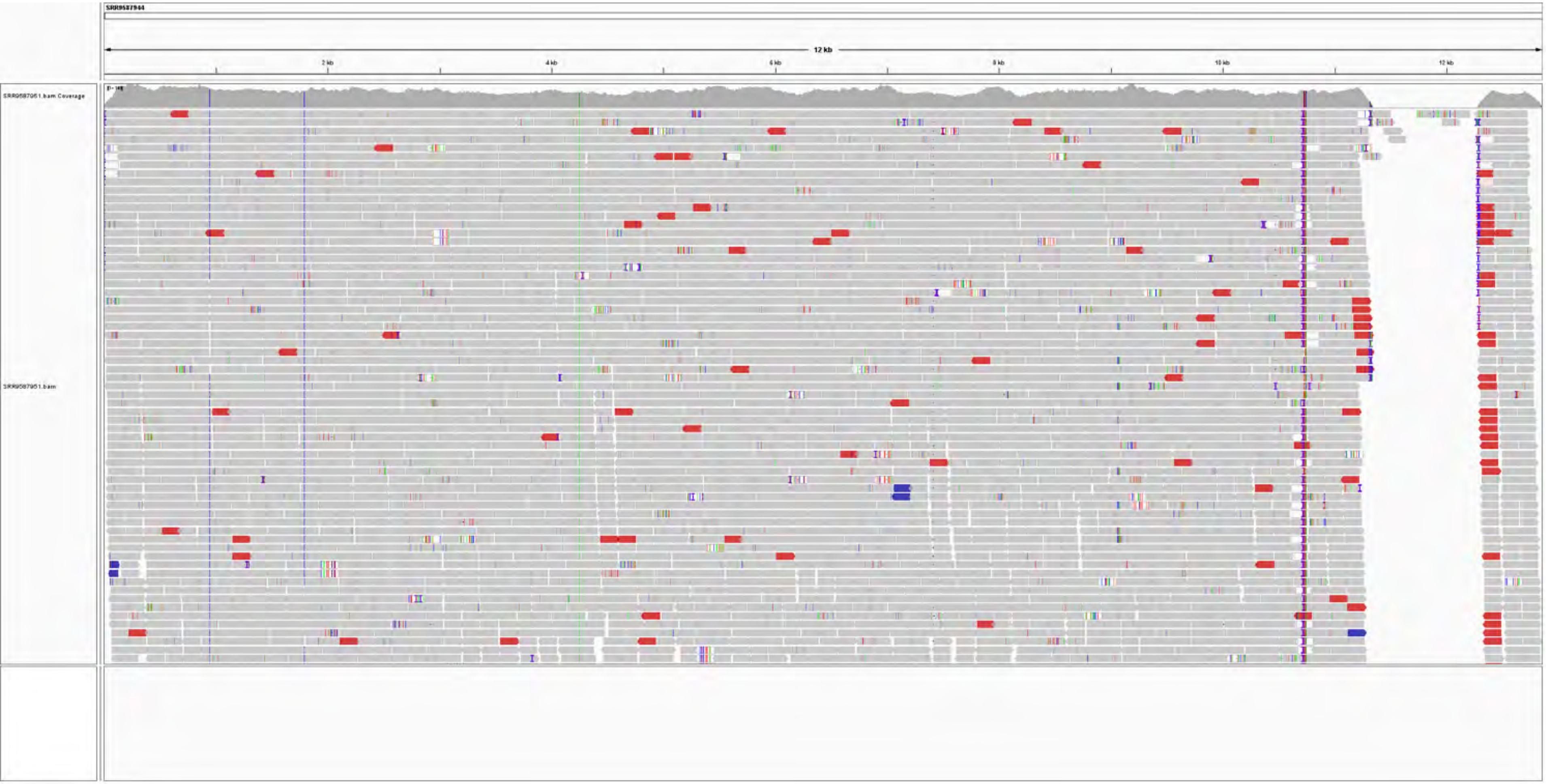

SRR9587944
12 kb
2 kb
4 kb
6 kb
8 kb
10 kb
12 kb
SRR9587951.bam Coverage
SRR9587951.bam

### SRR9587952

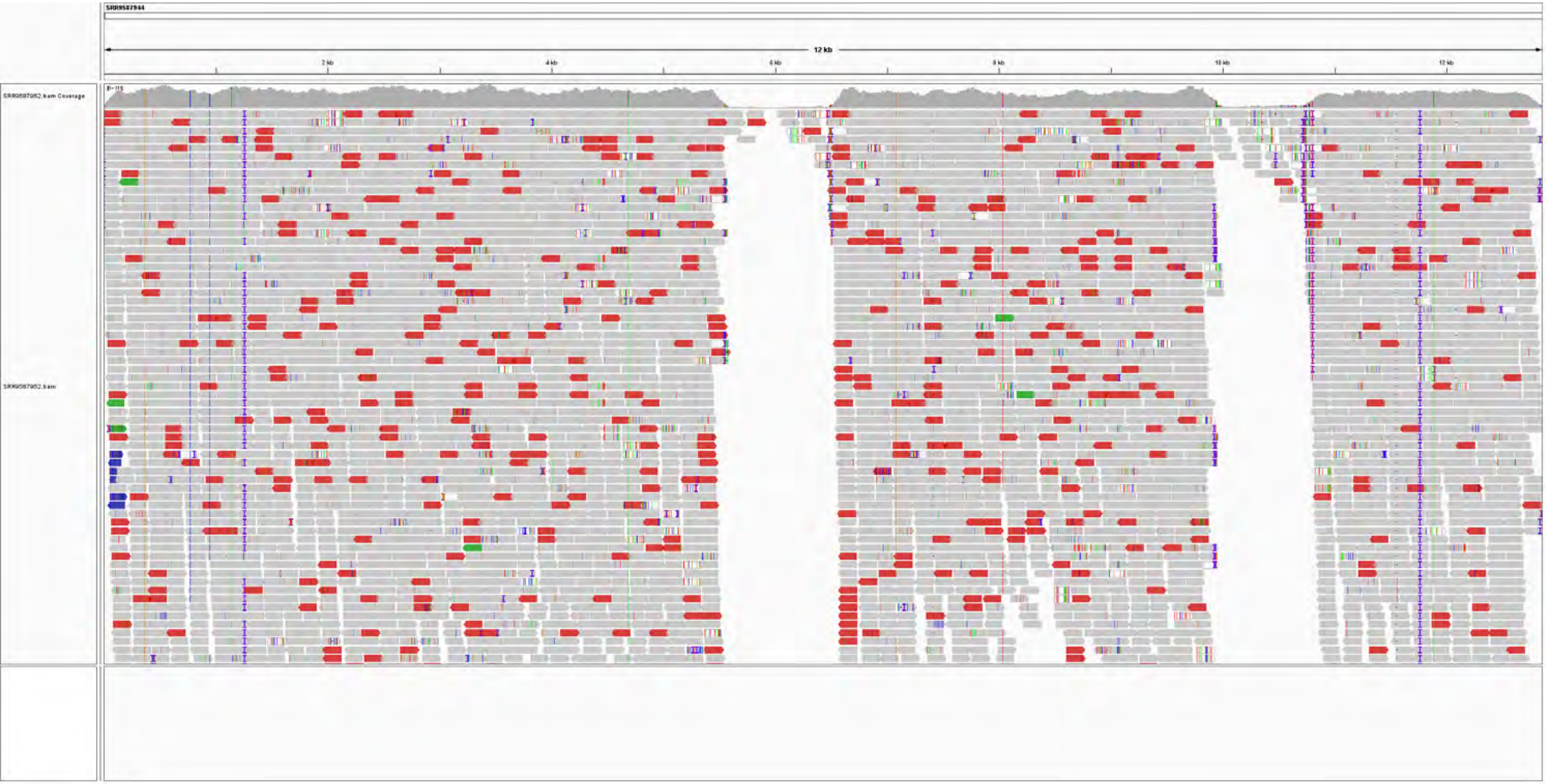

### SRR9587953

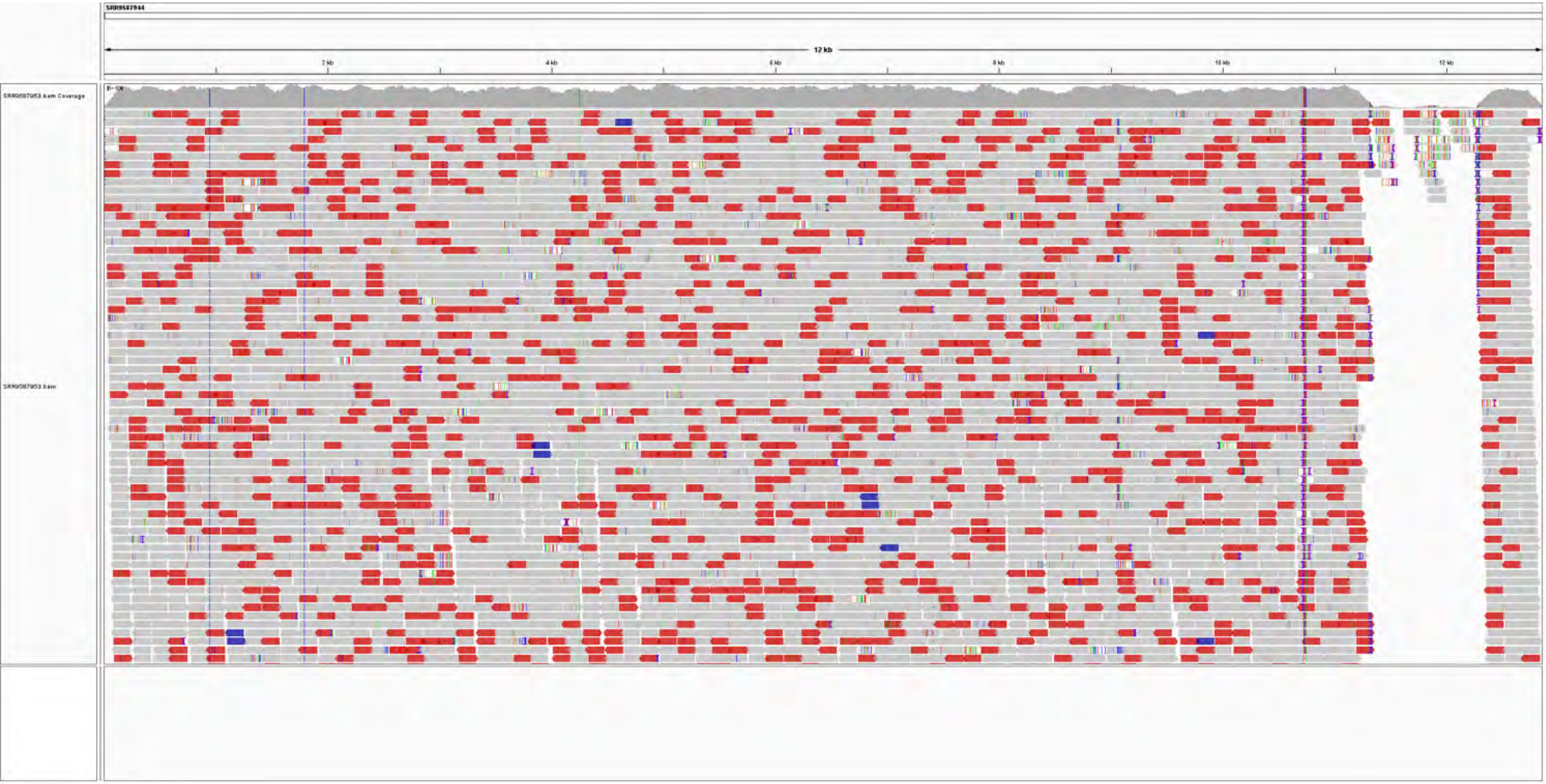

### SRR9587954

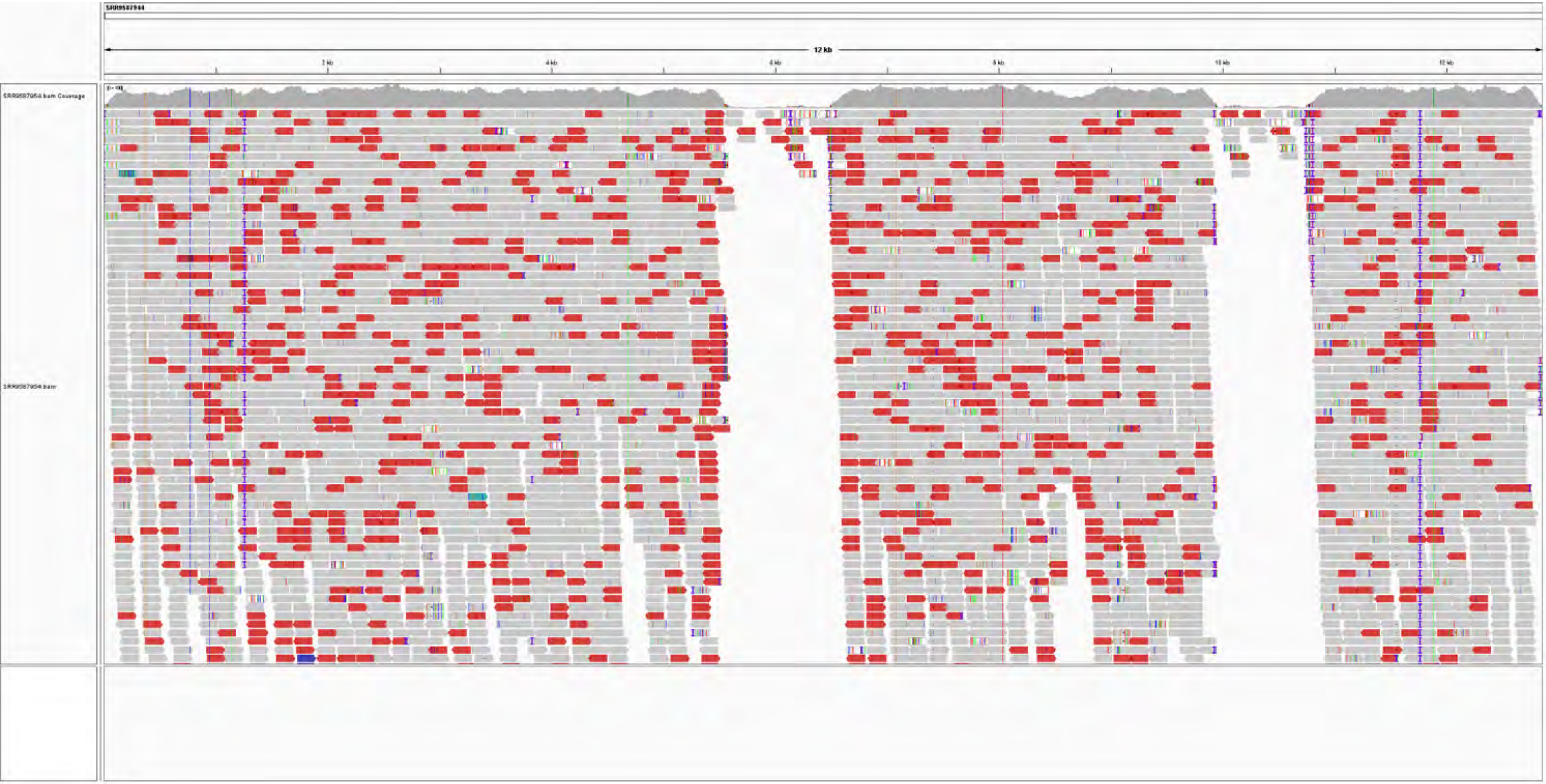
12 kb
2 kb
4 kb
6 kb
8 kb
10 kb
12 kb
SRR9587954.bam Coverage
SRR9587954.bam

### SRR9587955

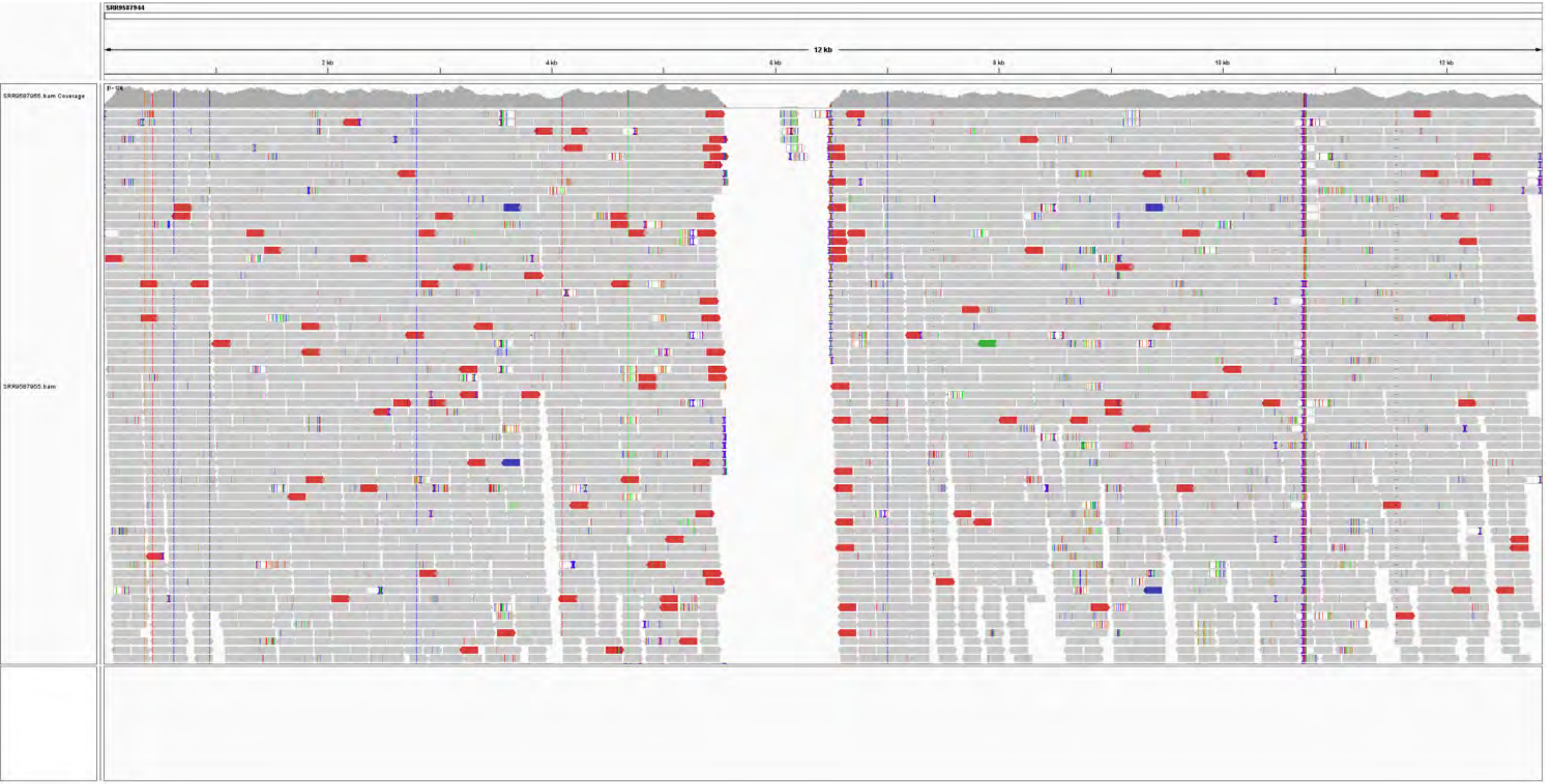

SRR9587944
12 kb
2 kb
4 kb
6 kb
8 kb
10 kb
12 kb
SRR9587955.bam Coverage
SRR9587955.bam

### SRR9587956

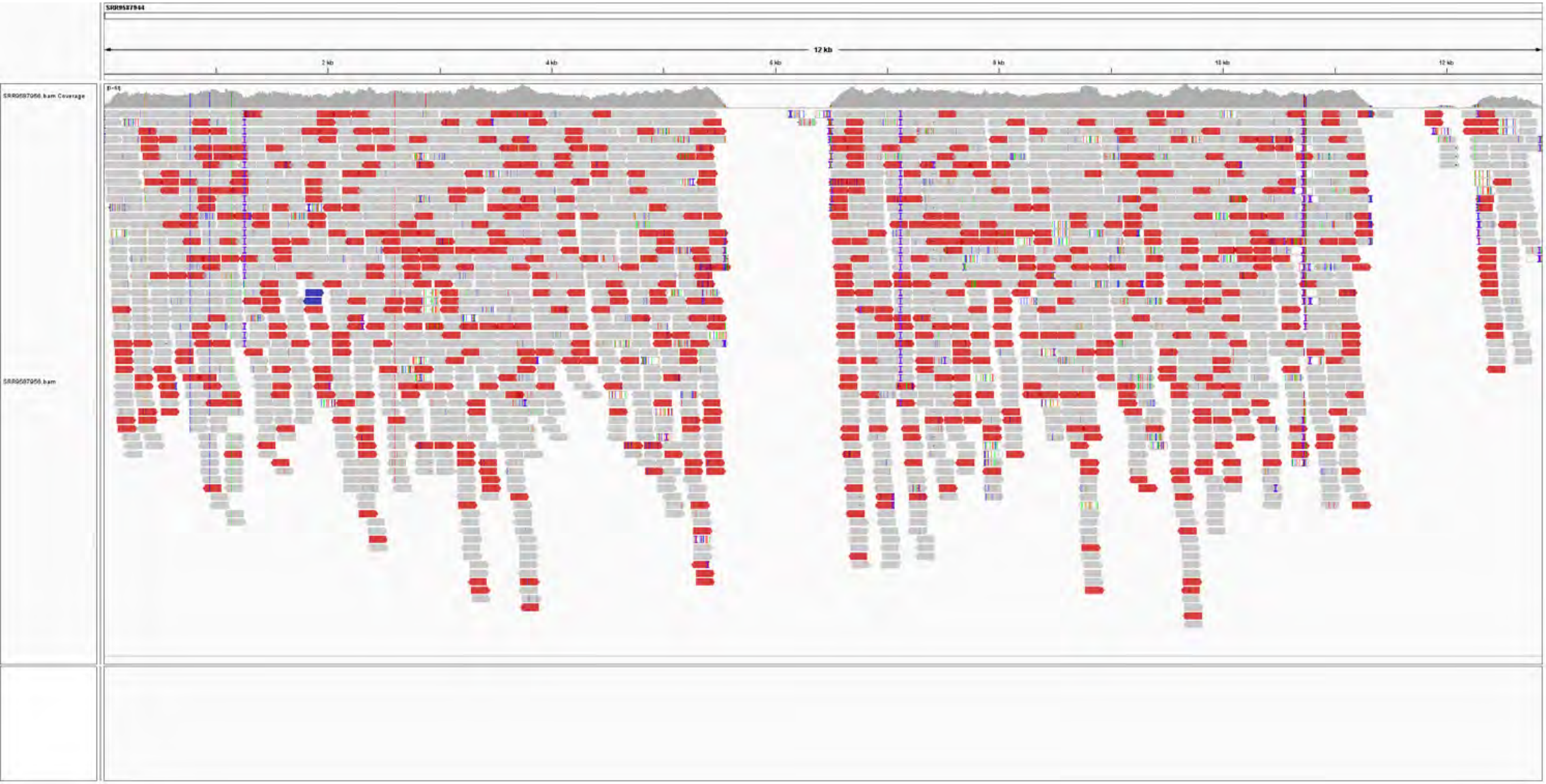

### SRR9587958

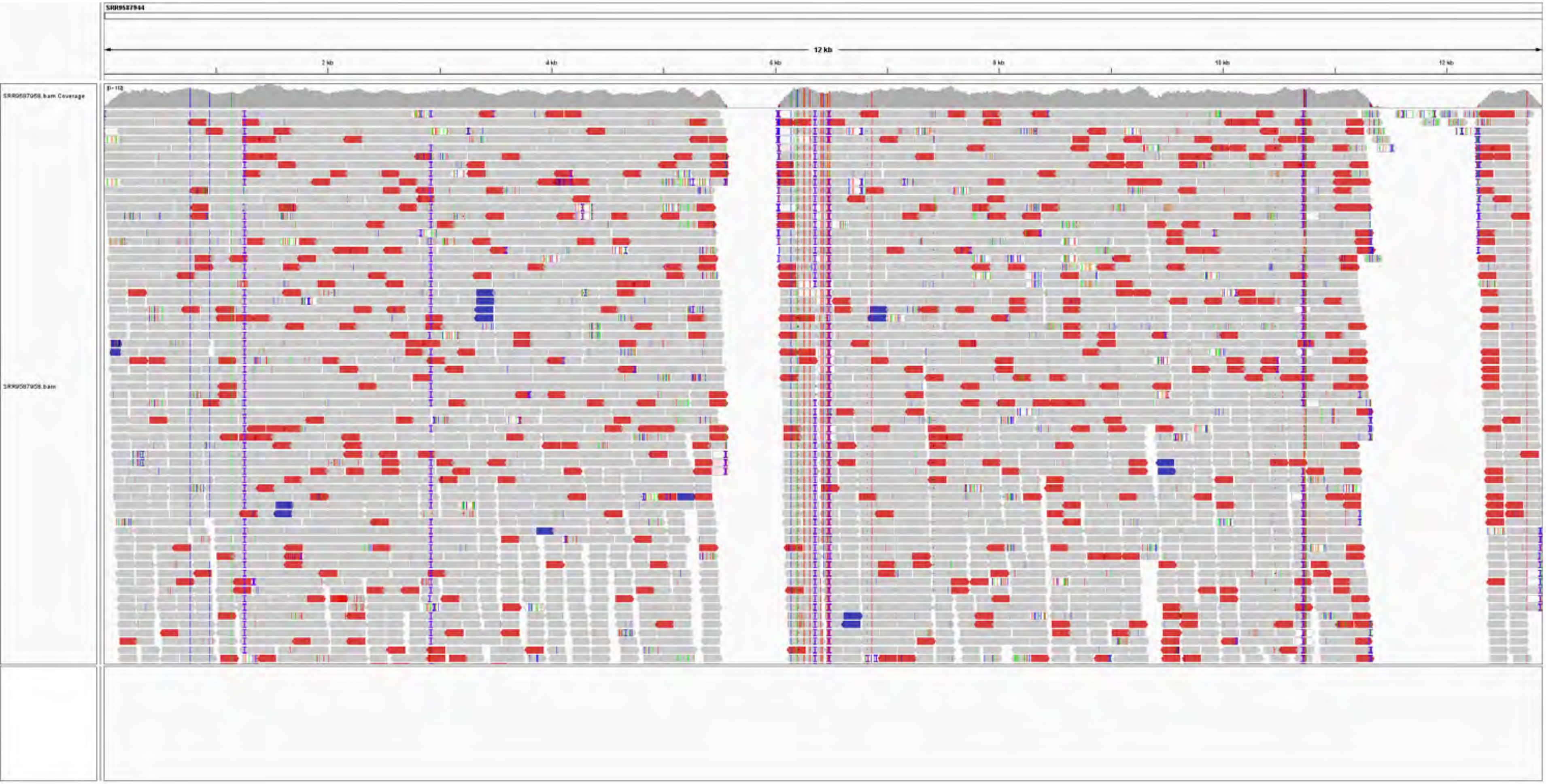

### SRR9587959

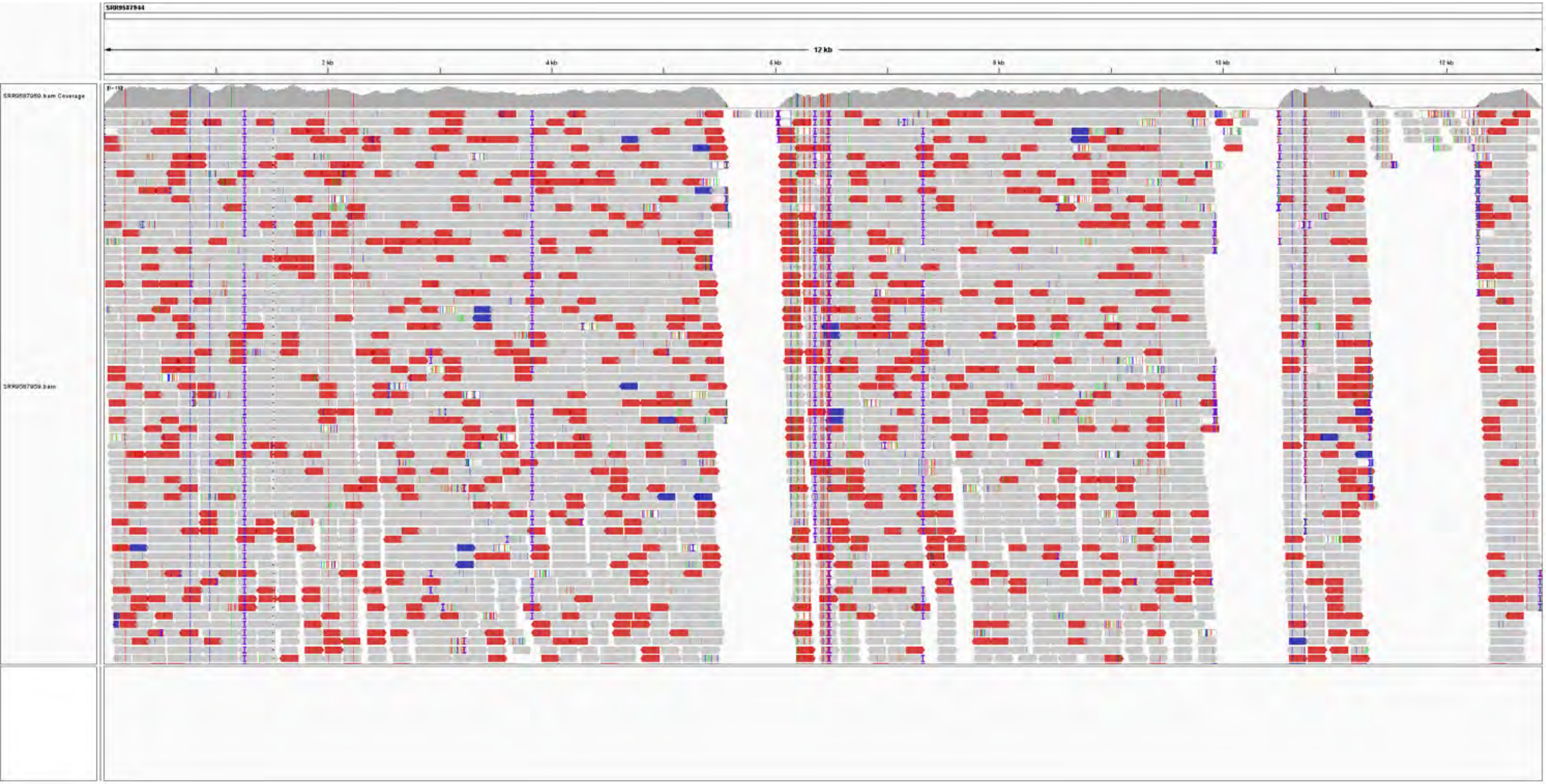

SRR9587944
12 kb
2 kb
4 kb
6 kb
8 kb
10 kb
12 kb
SRR9587959.bam Coverage
SRR9587959.bam

### SRR9587960

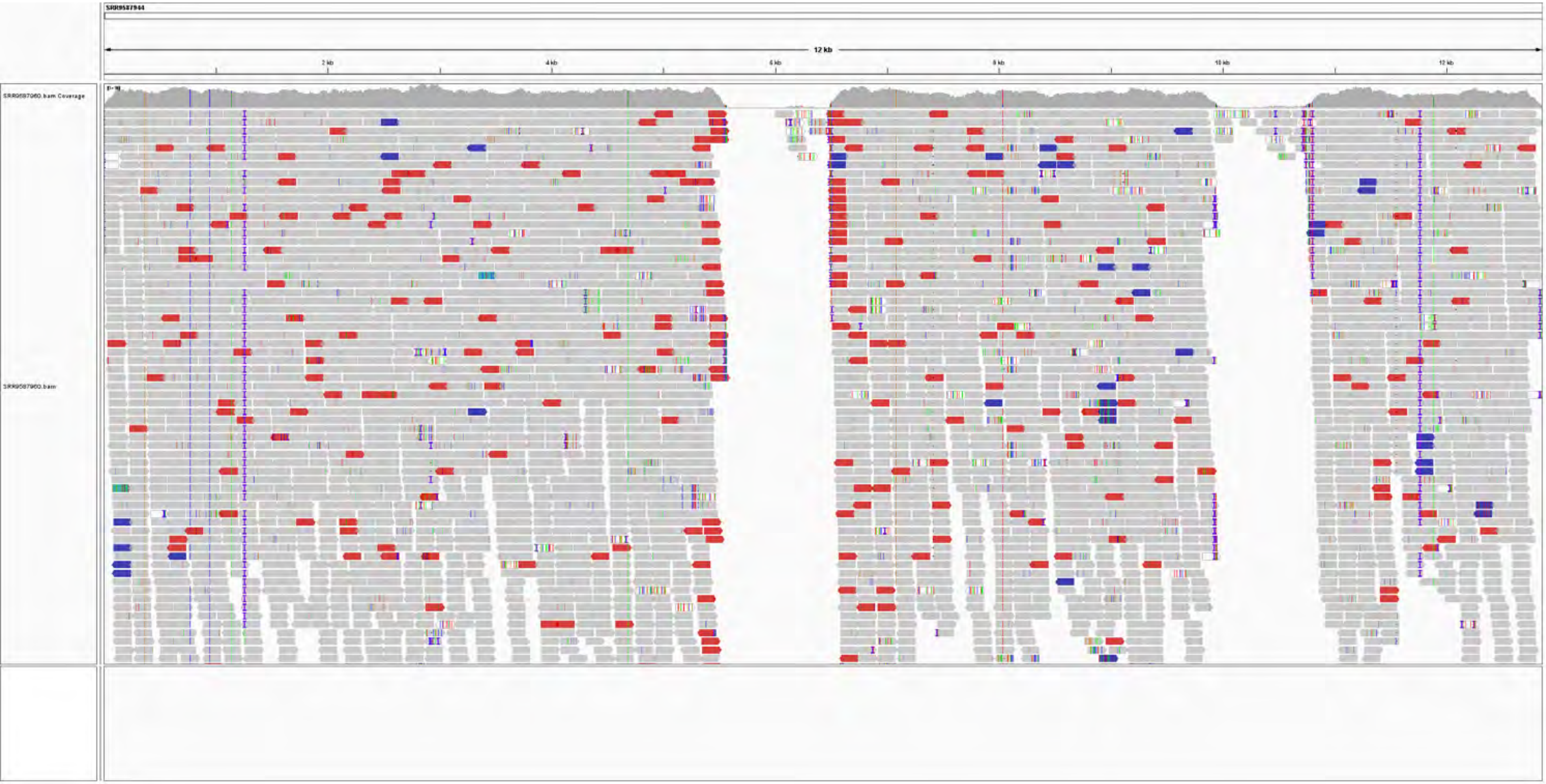

SRR9587944
12 kb
2 kb
4 kb
6 kb
8 kb
10 kb
12 kb
SRR9587960.bam Coverage
SRR9587960.bam

### SRR9587961

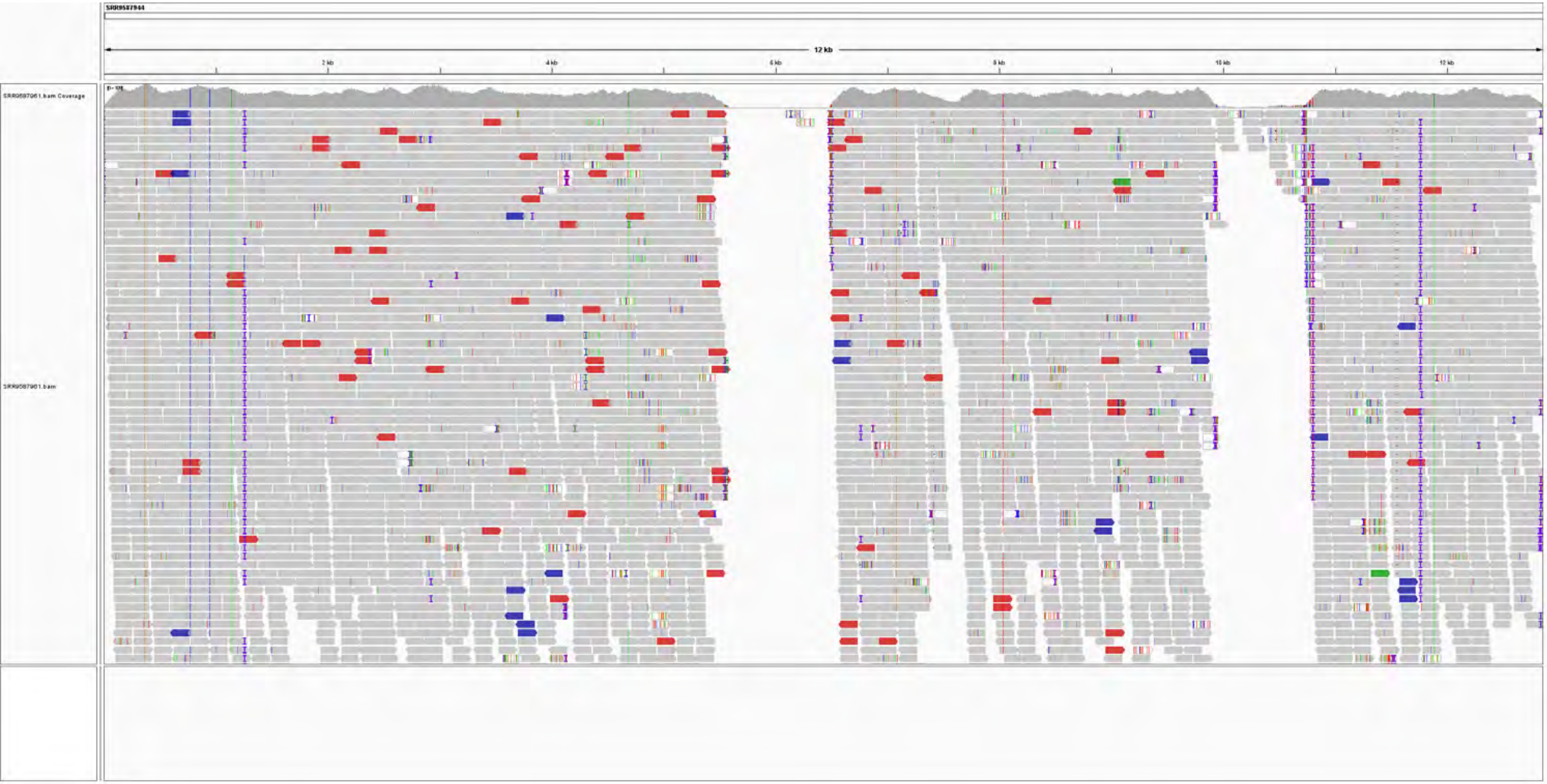

### SRR9587962

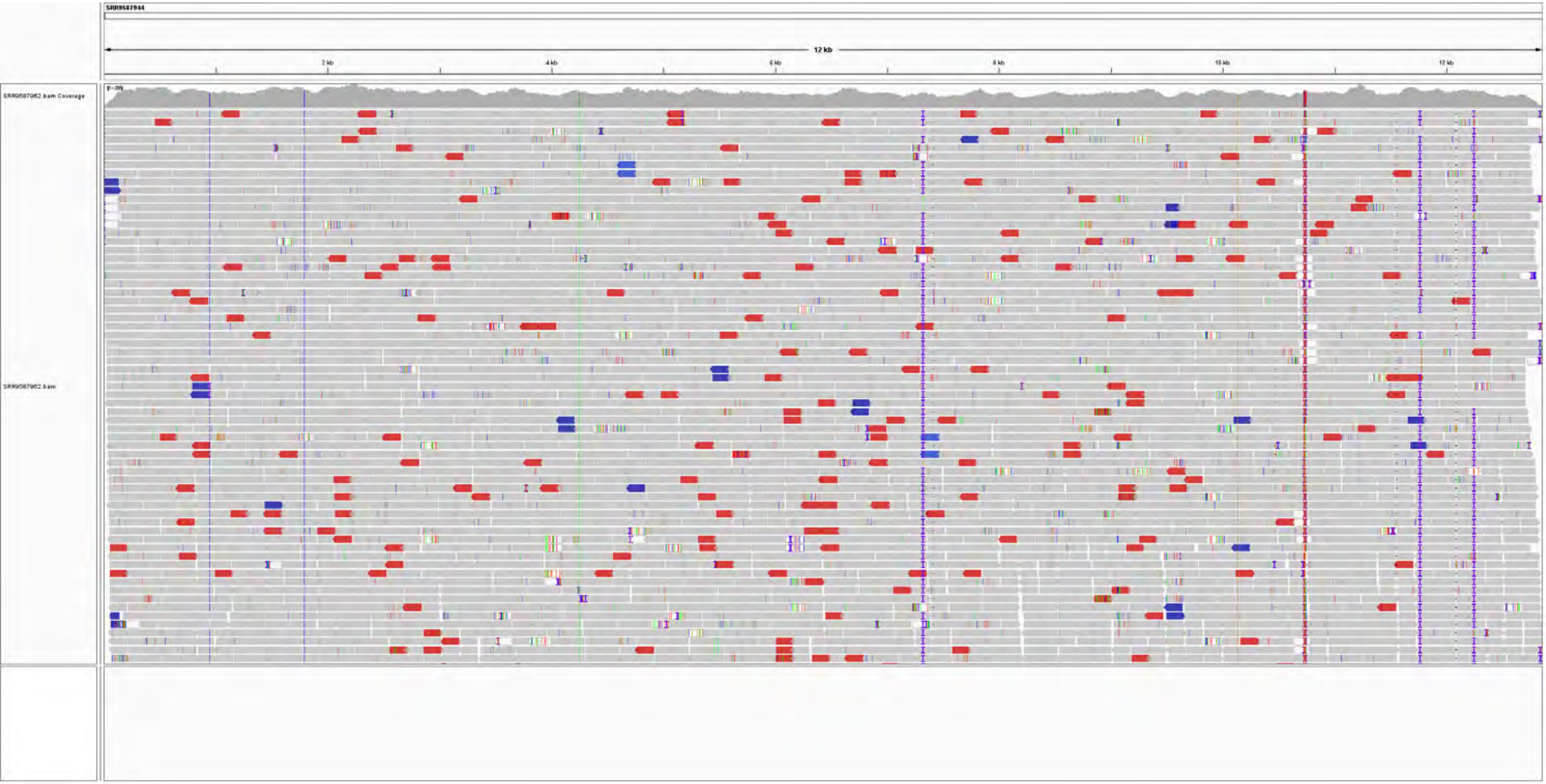

### SRR9587963

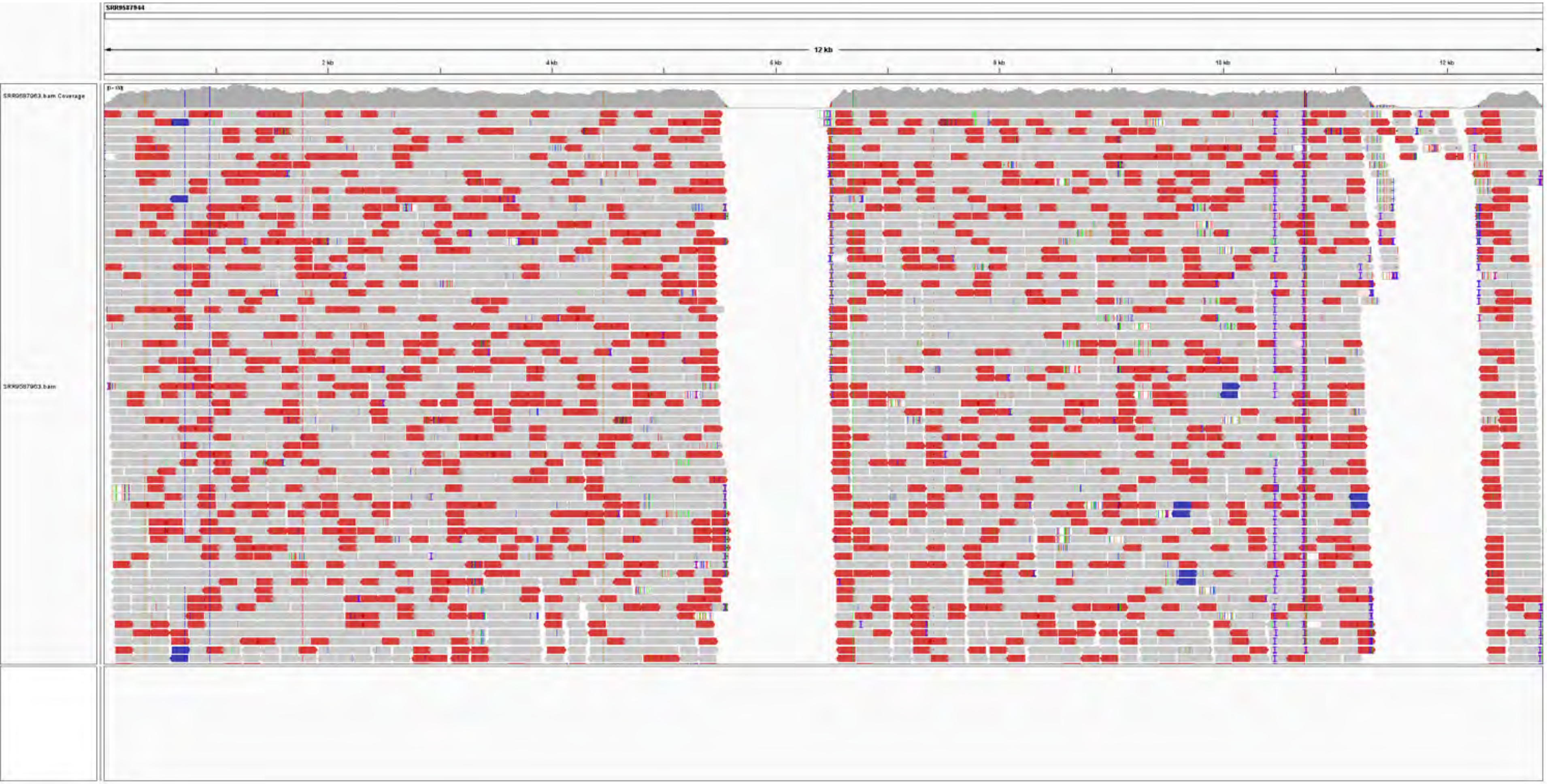

SRR9587944
12 kb
2 kb
4 kb
6 kb
8 kb
10 kb
12 kb
SRR9587963.bam Coverage
SRR9587963.bam

### SRR9587964

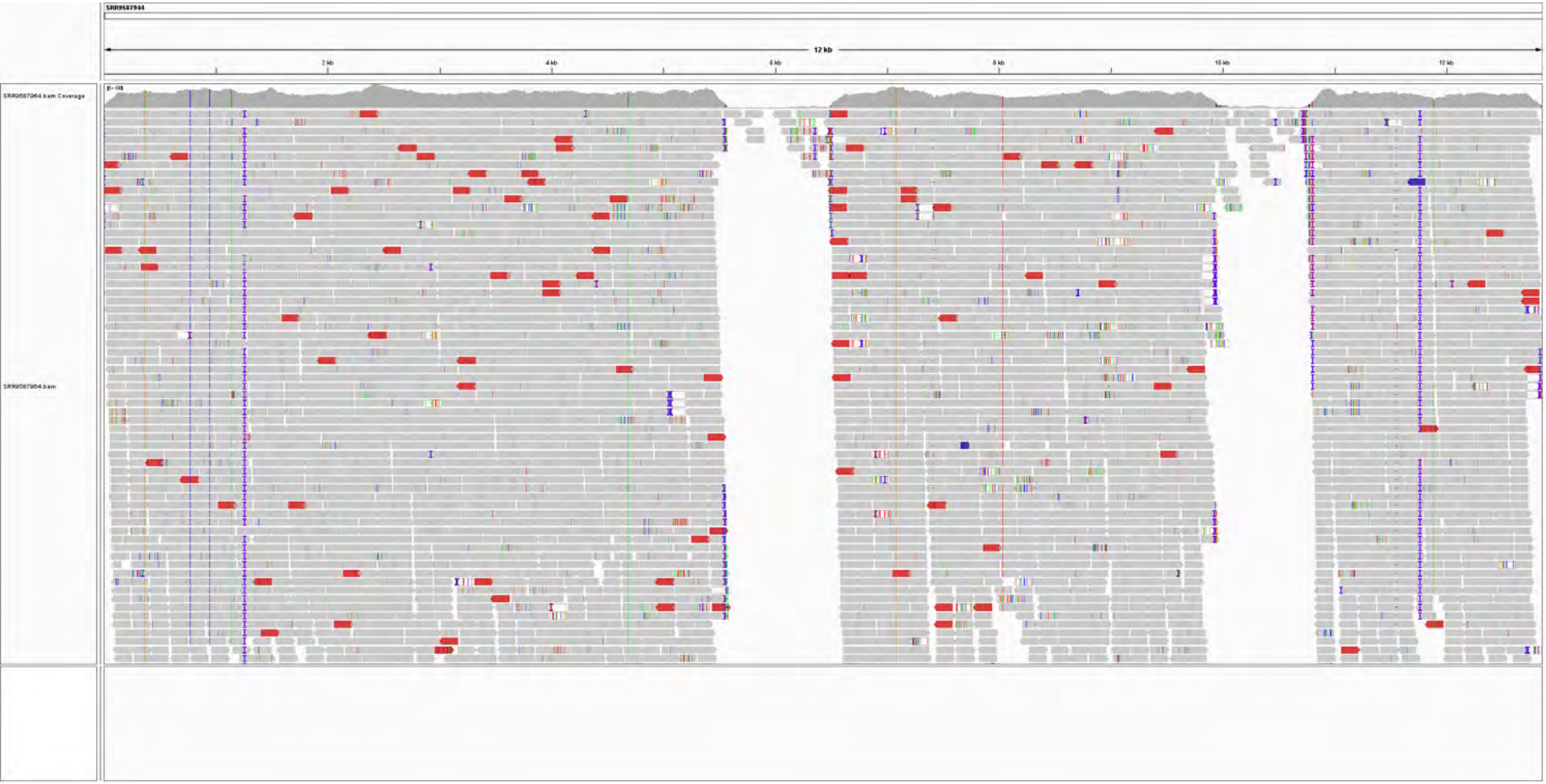

12 kb
2 kb
4 kb
6 kb
8 kb
10 kb
12 kb
SRR9587964.bam Coverage
SRR9587964.bam

### SRR9587965

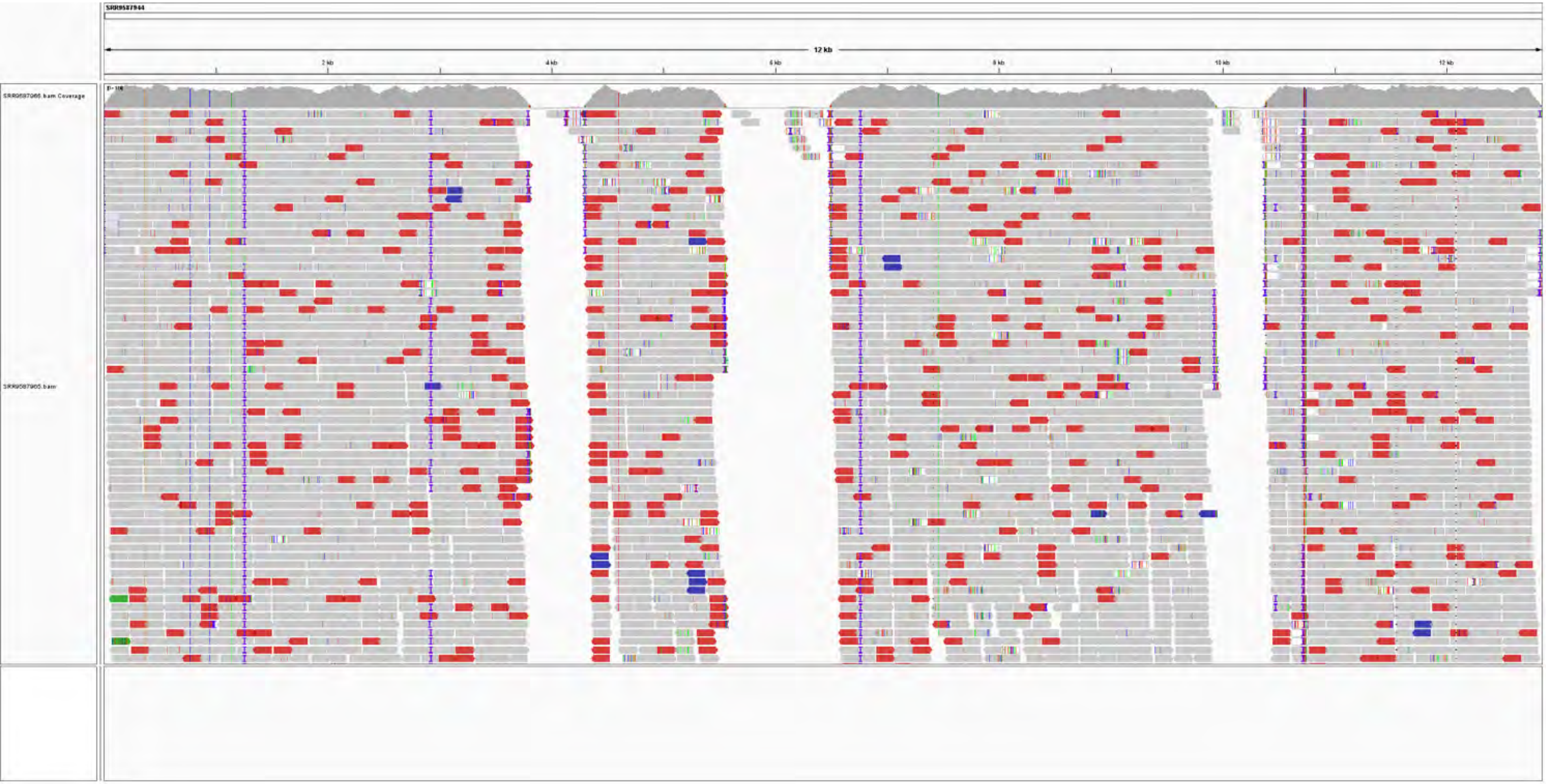

### SRR9587966

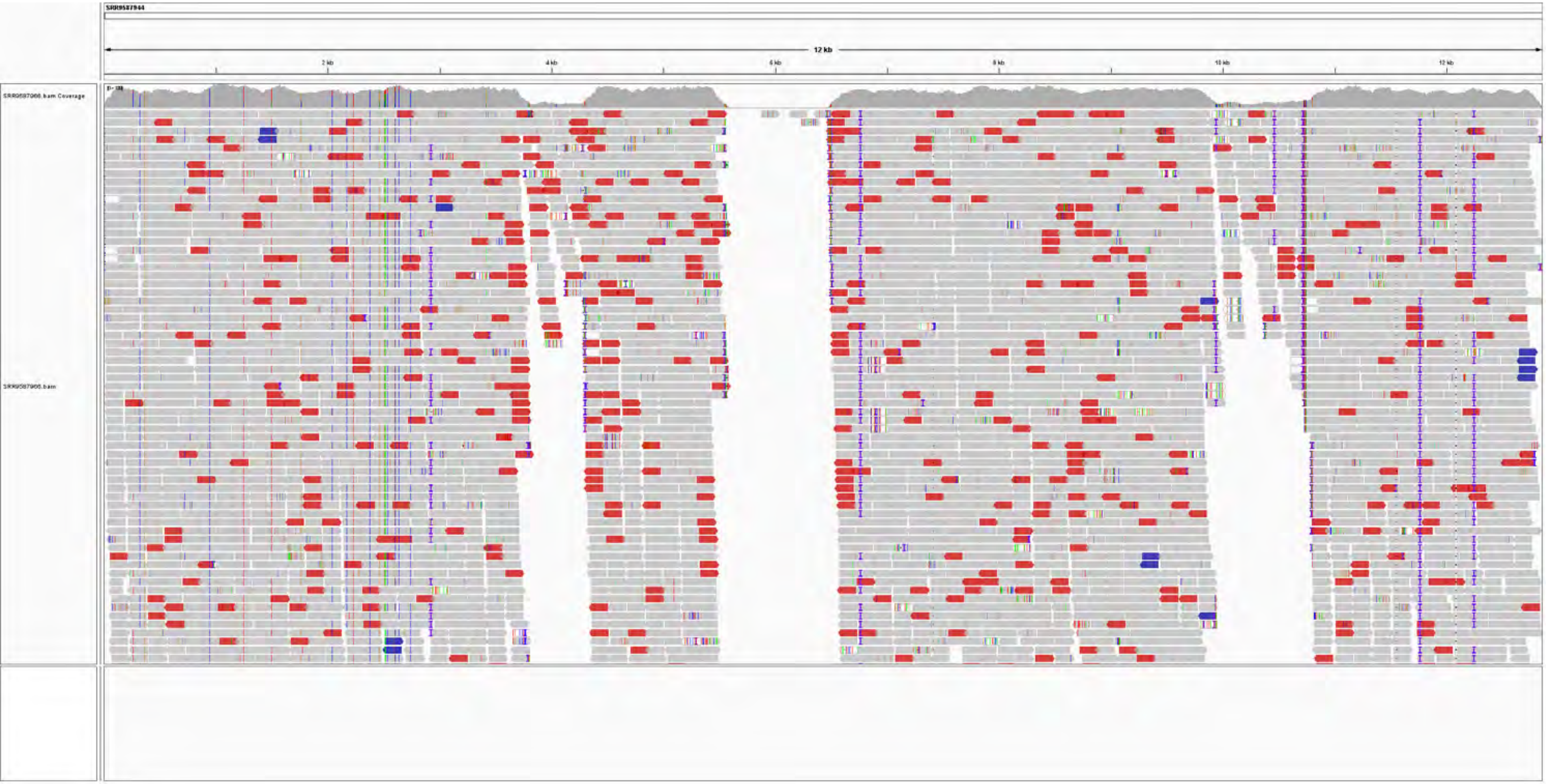

### SRR9587967

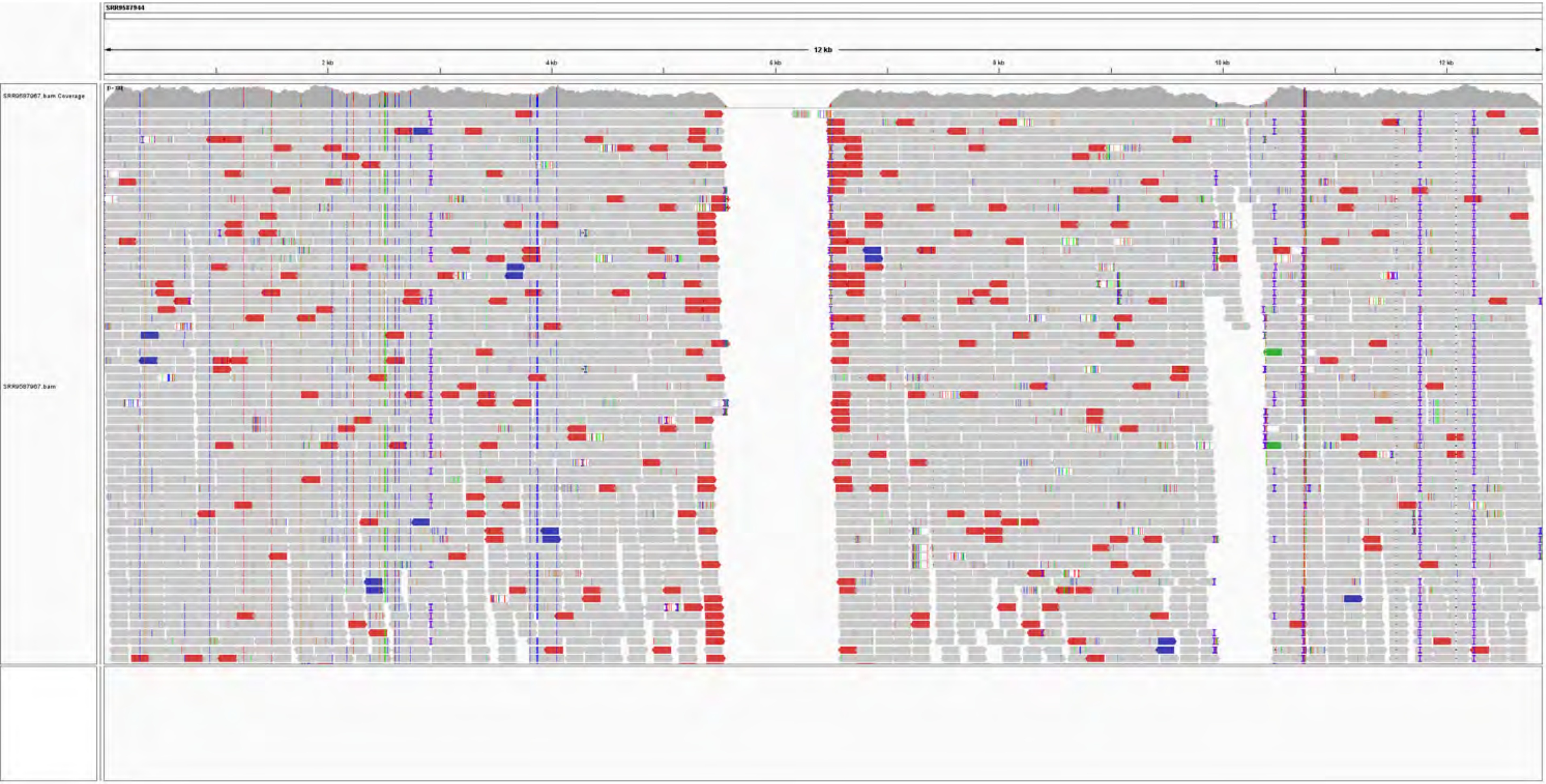
SRR9587944
12 kb
2 kb
4 kb
6 kb
8 kb
10 kb
12 kb
SRR9587967.bam Coverage
SRR9587967.bam

### SRR9587968

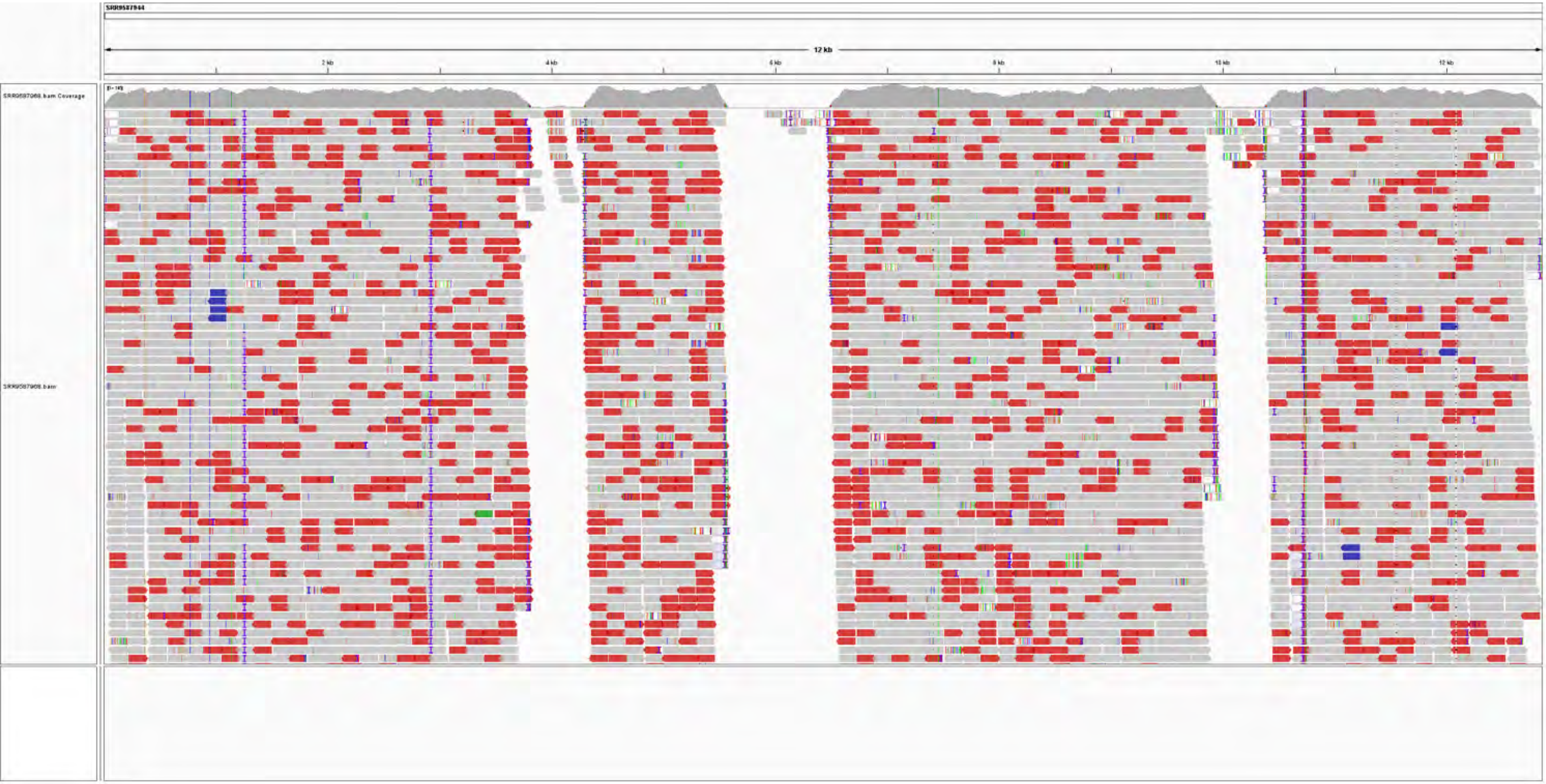

### SRR9587969

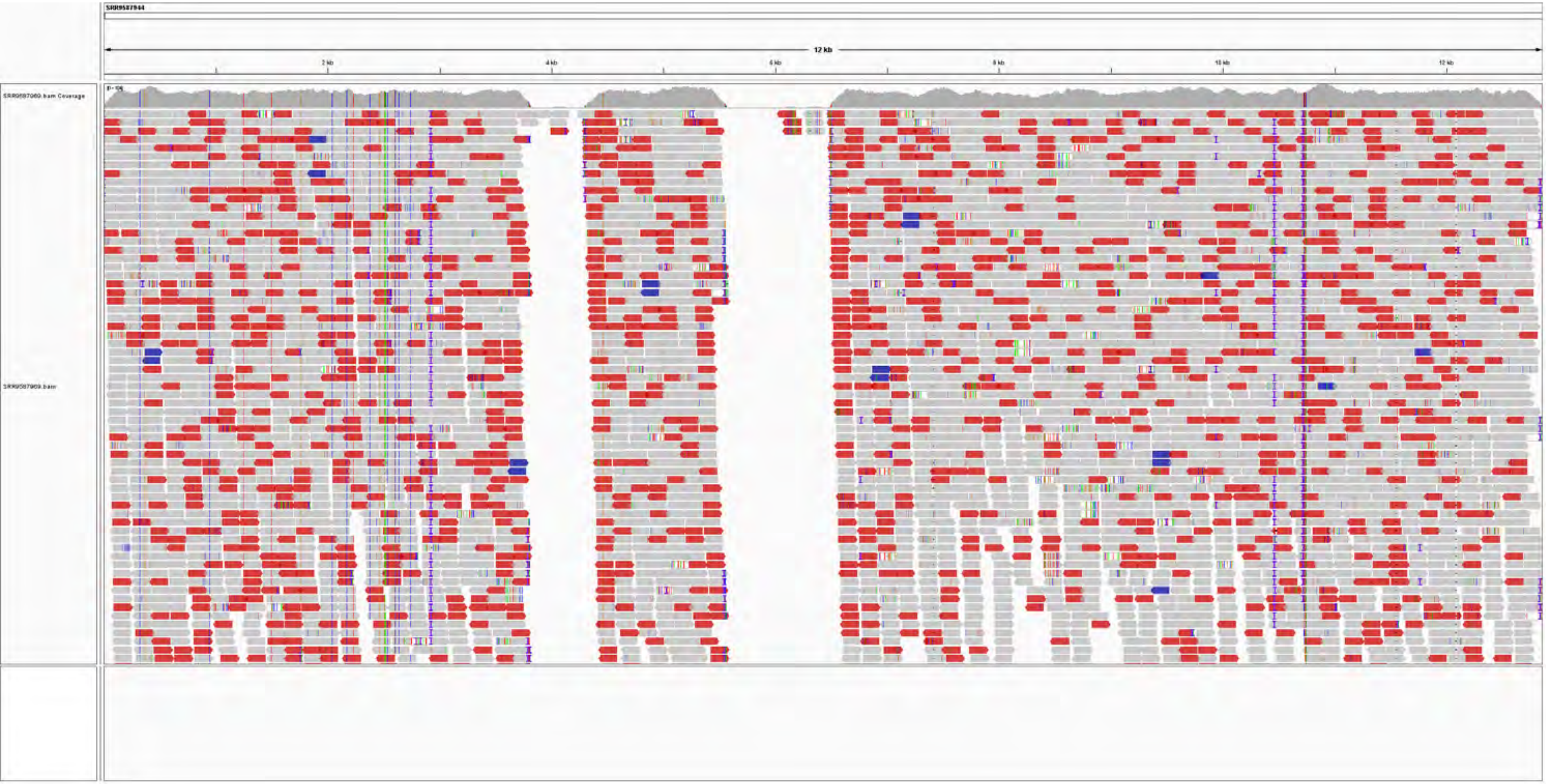
SRR9587944
12 kb
2 kb
4 kb
6 kb
8 kb
10 kb
12 kb
SRR9587969.bam Coverage
SRR9587969.bam

**Supplementary Figure 7.** The results of mapping the reads of Japanese *N. yezoensis* to the sequence of Pyr_16.

### Pyr_1

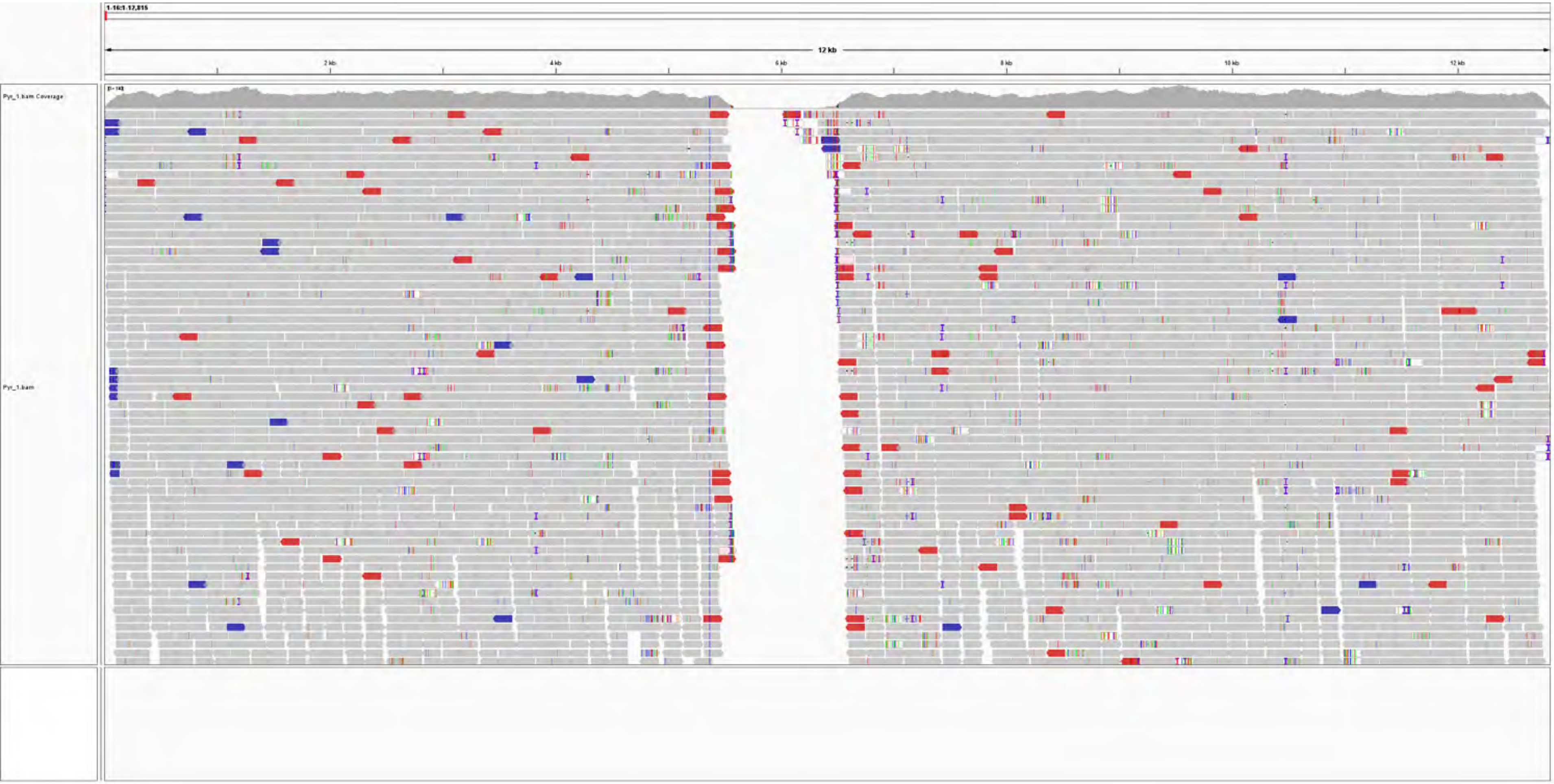

### Pyr_2

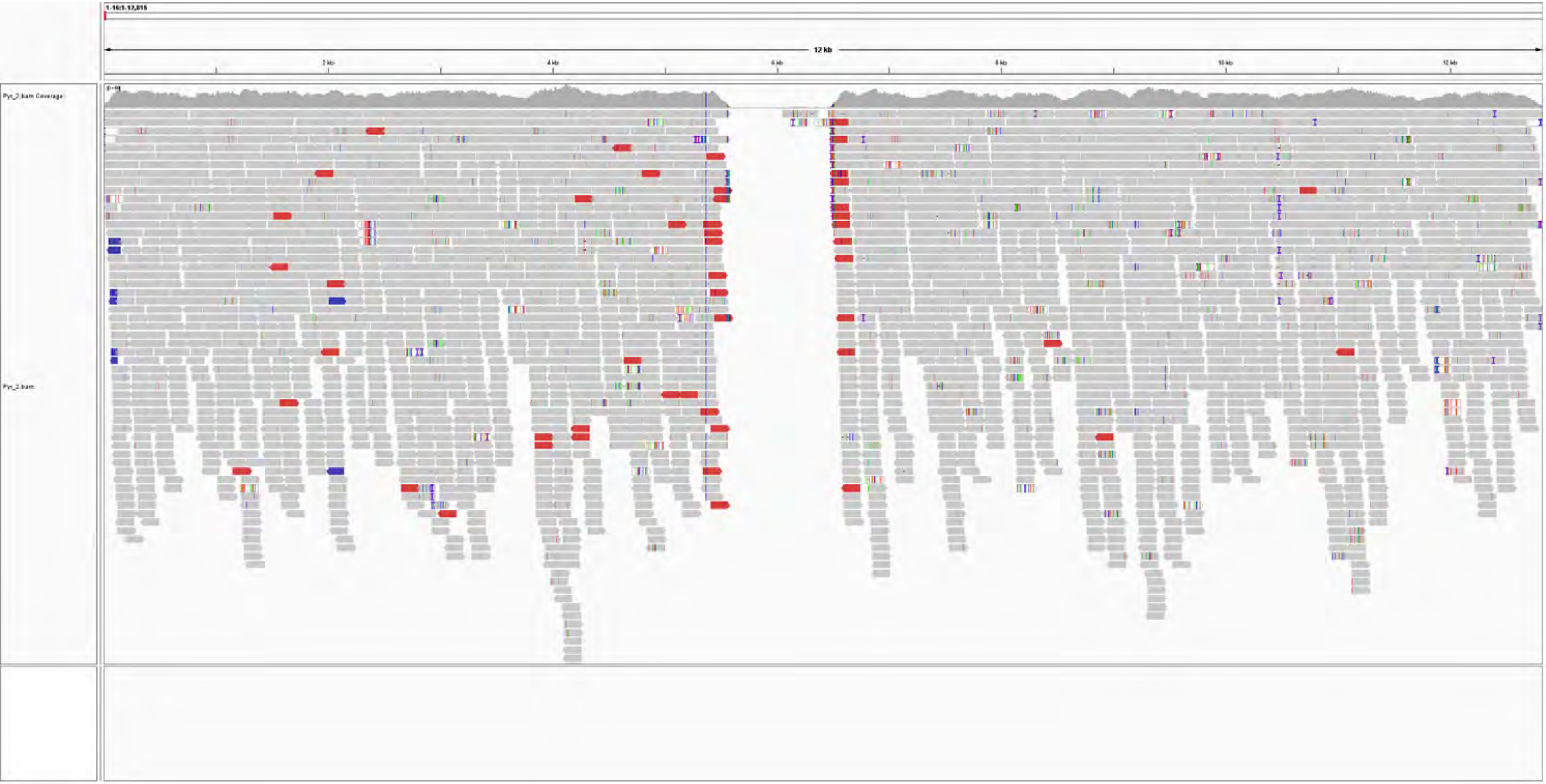

1-16:1-12,815
12 kb
2 kb
4 kb
6 kb
8 kb
10 kb
12 kb
Pyr_2.bam Coverage
Pyr_2.bam

### Pyr_3

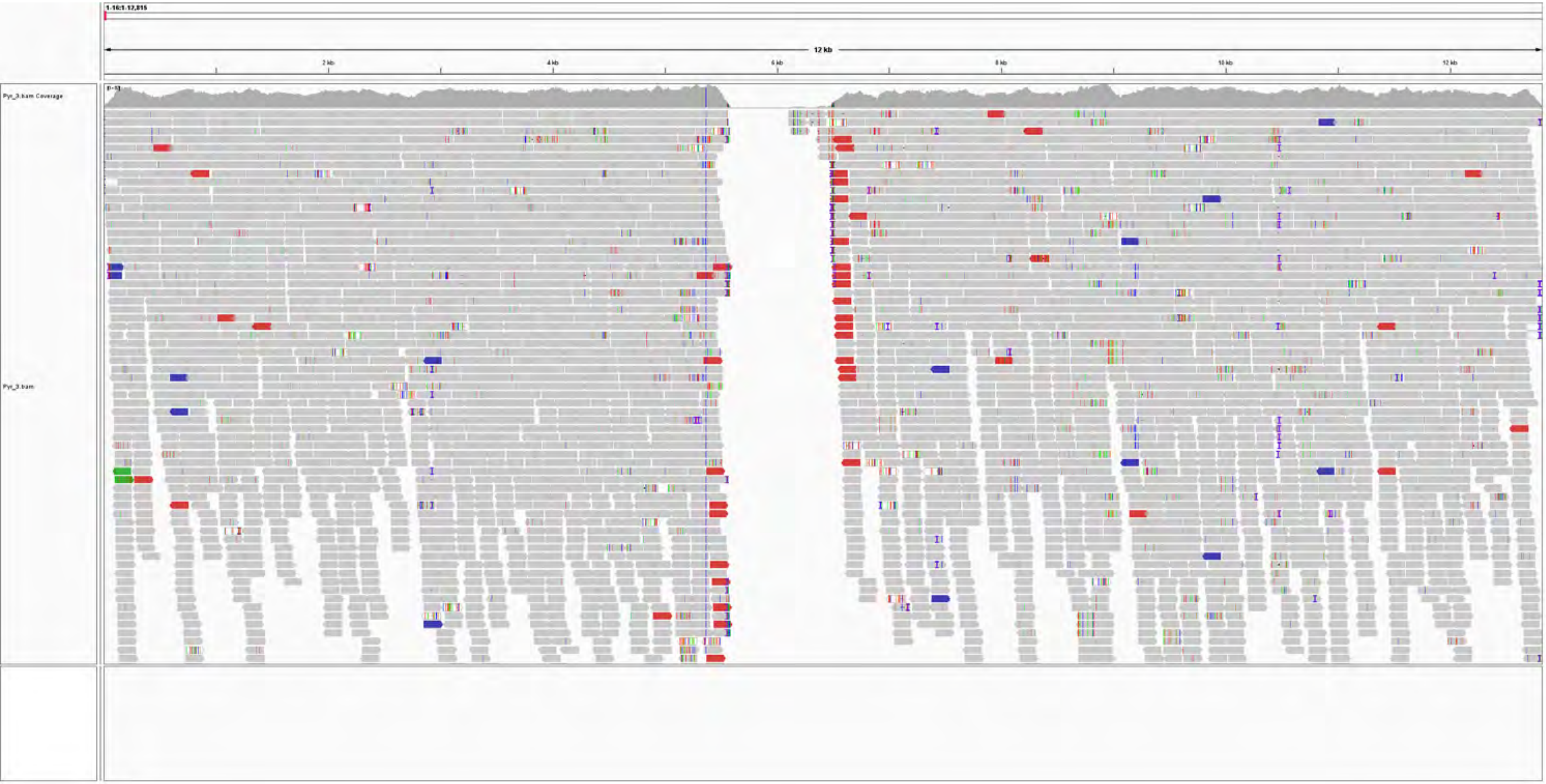

1-16:1-12,815
12 kb
2 kb
4 kb
6 kb
8 kb
10 kb
12 kb
Pyr_3.bam Coverage
Pyr_3.bam

### Pyr_4

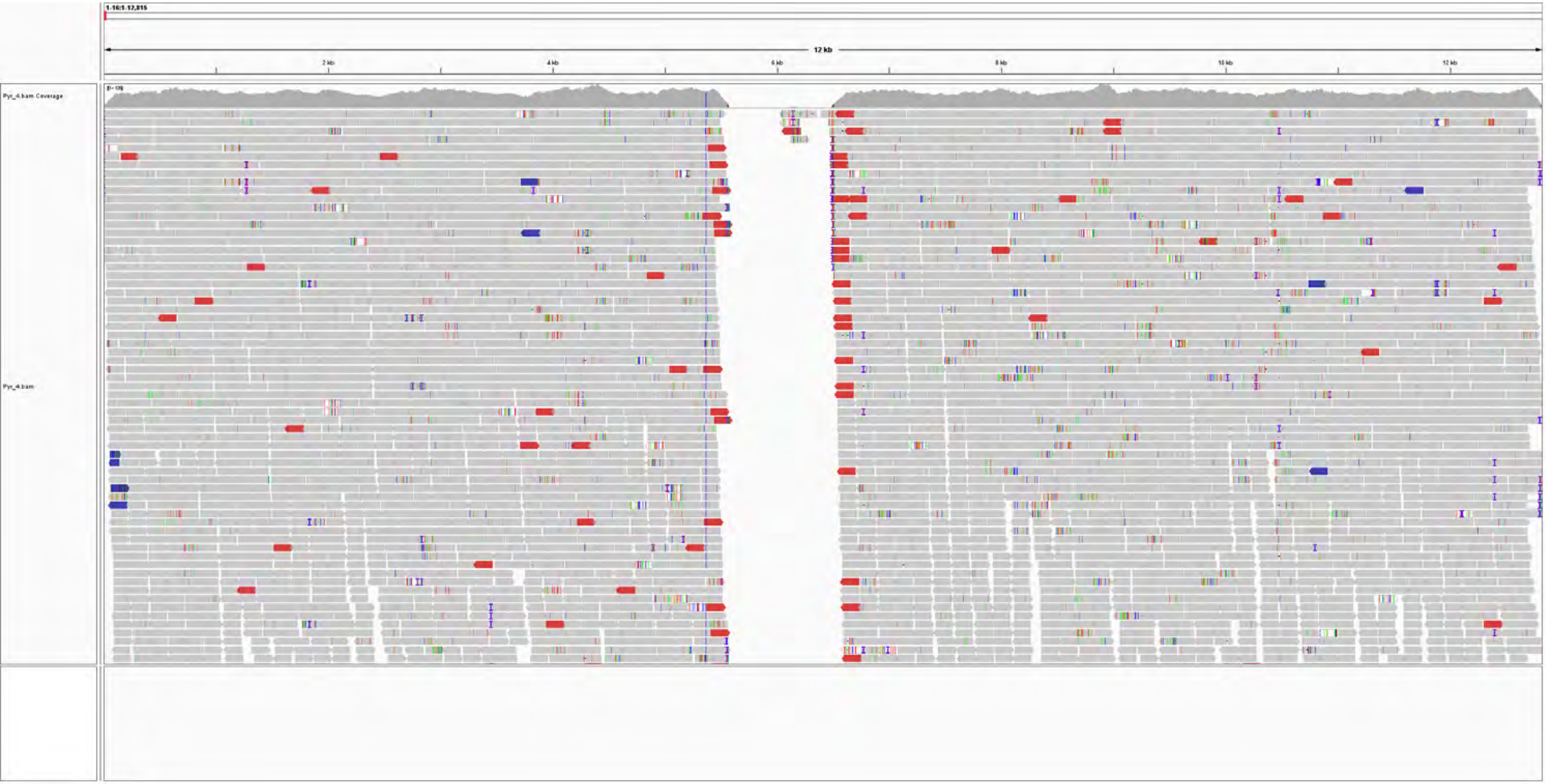

1-16:1-12,815
12 kb
2 kb
4 kb
6 kb
8 kb
10 kb
12 kb
Pyr_4.bam Coverage
Pyr_4.bam

### Pyr_7

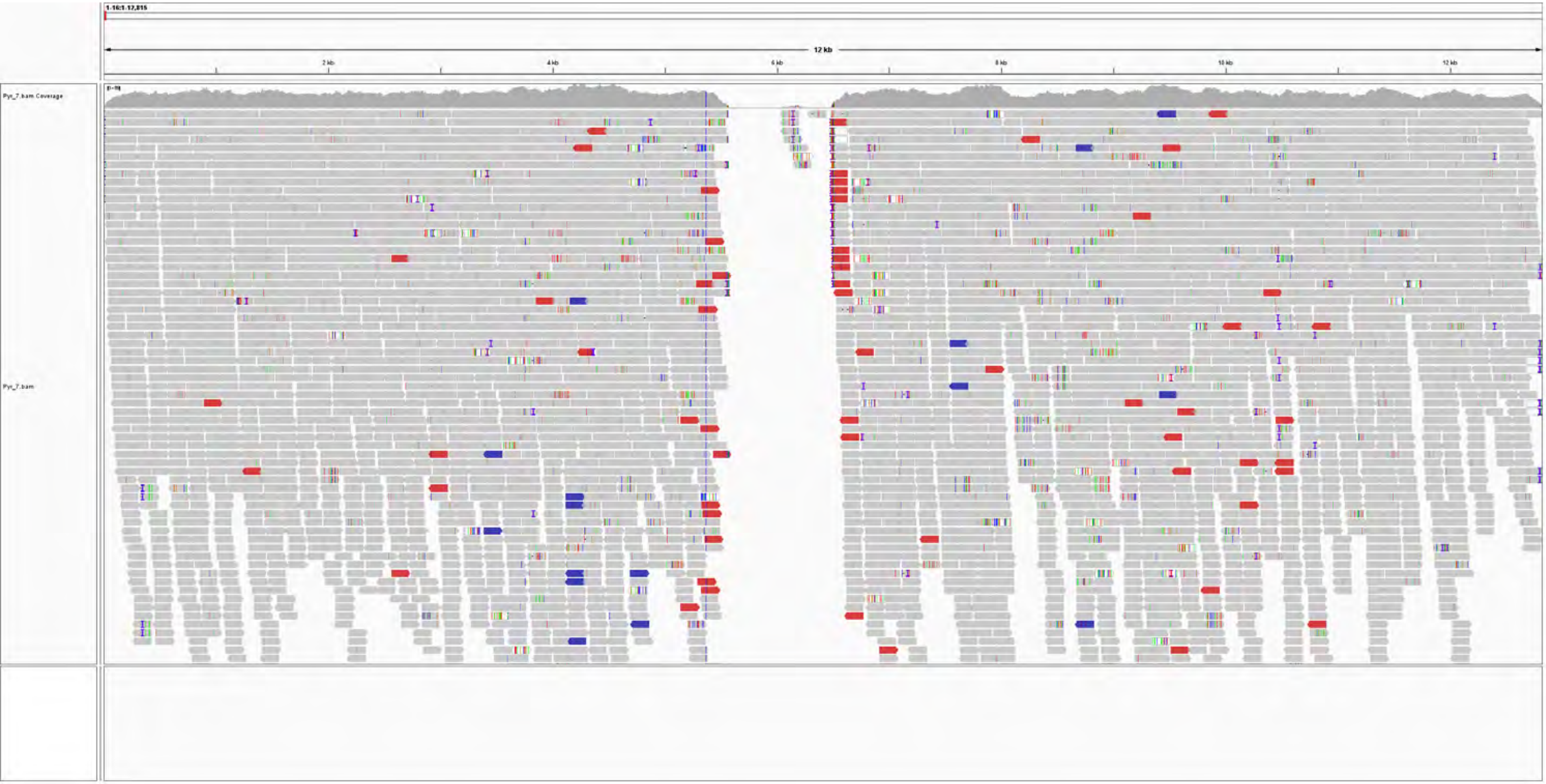

1-16:1-12,815
12 kb
2 kb
4 kb
6 kb
8 kb
10 kb
12 kb
Pyr_7.bam Coverage
Pyr_7.bam

### Pyr_8

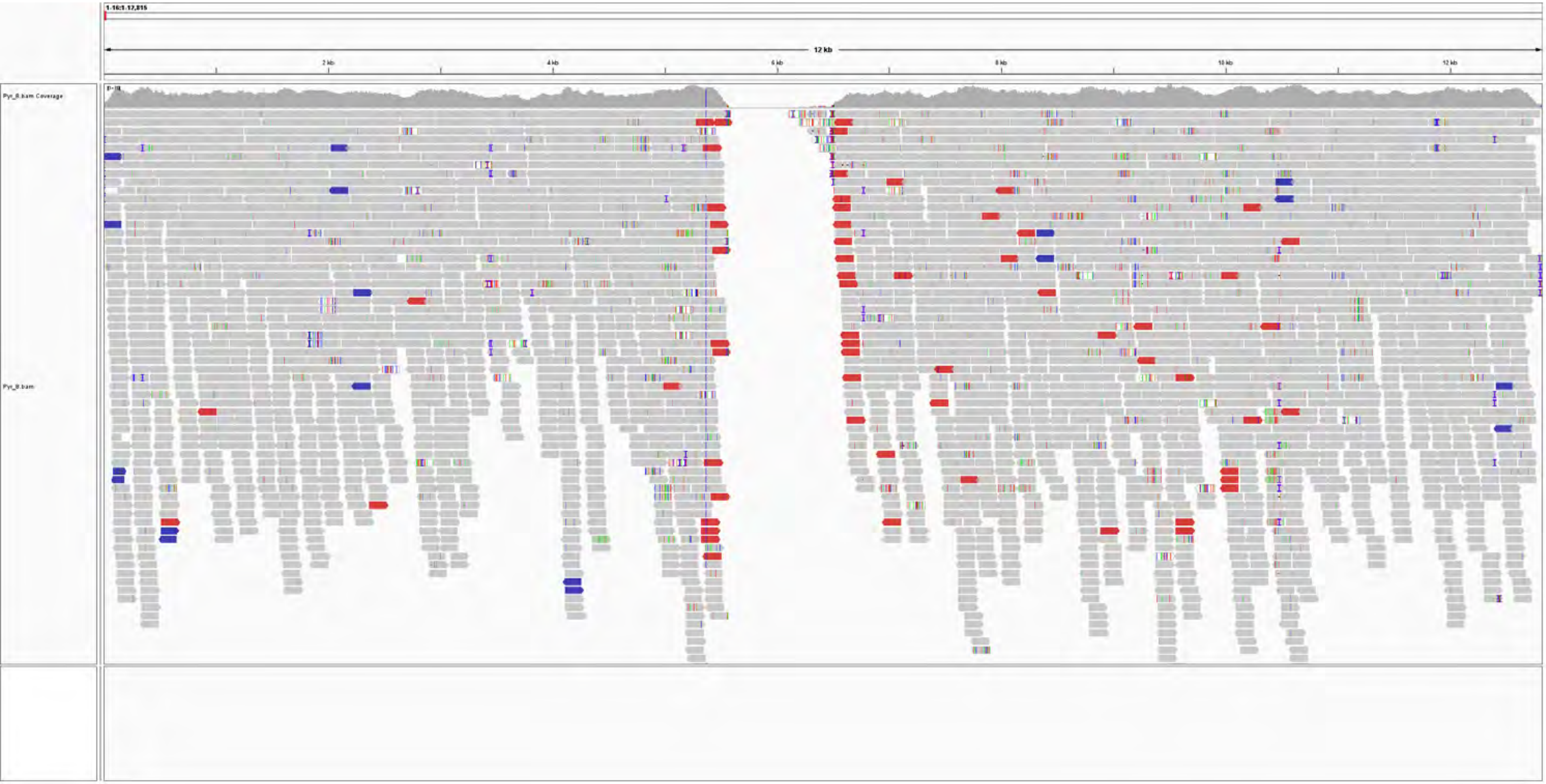

1-16:1-17,815
12 kb
2 kb
4 kb
6 kb
8 kb
10 kb
12 kb
Pyr_8.bam Coverage
Pyr_8.bam

### Pyr_9

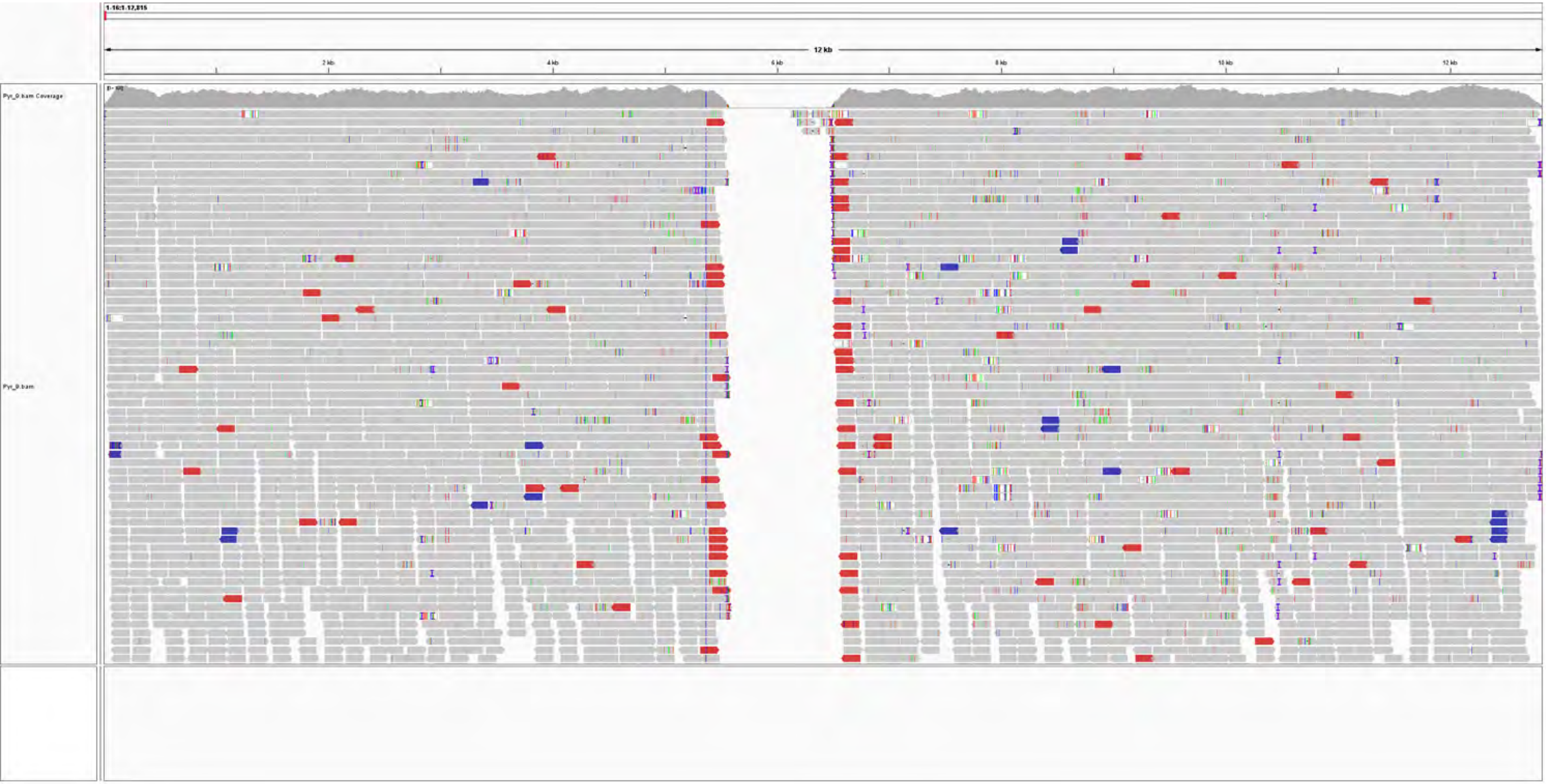

### Pyr_10

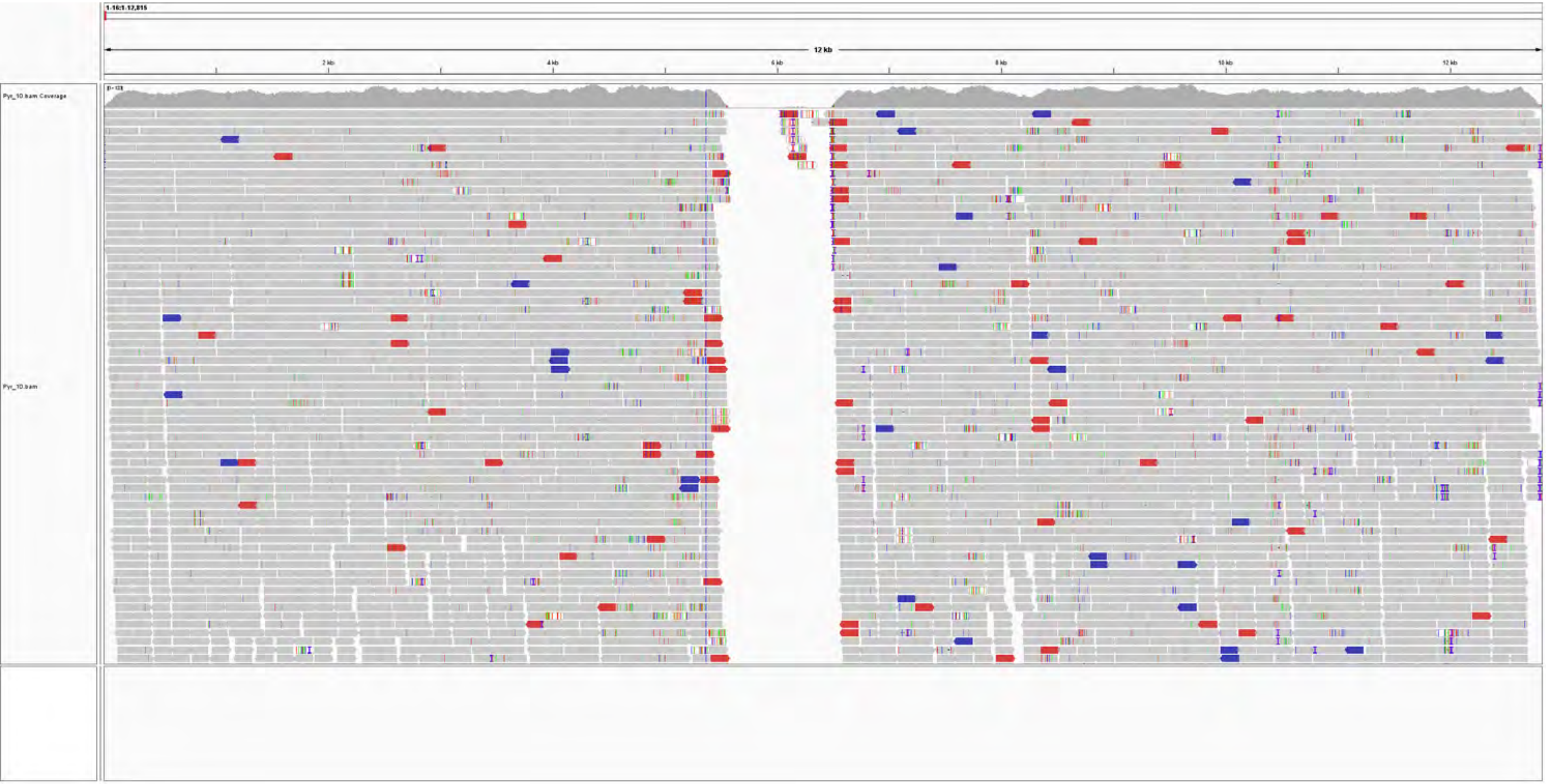

1-16:1-12,815
12 kb
2 kb
4 kb
6 kb
8 kb
10 kb
12 kb
Pyr_10.bam Coverage
Pyr_10.bam

### Pyr_11

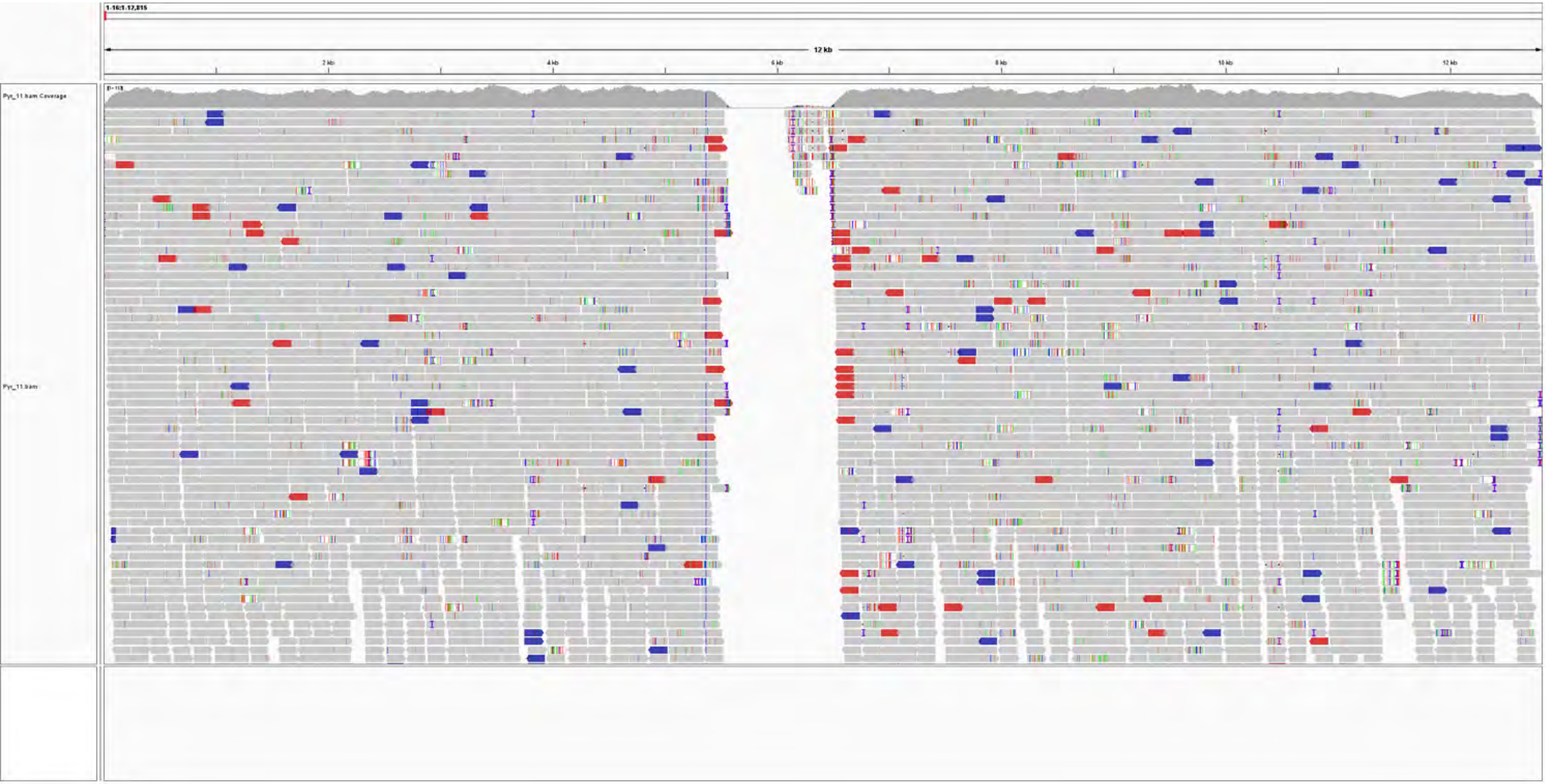

### Pyr_12

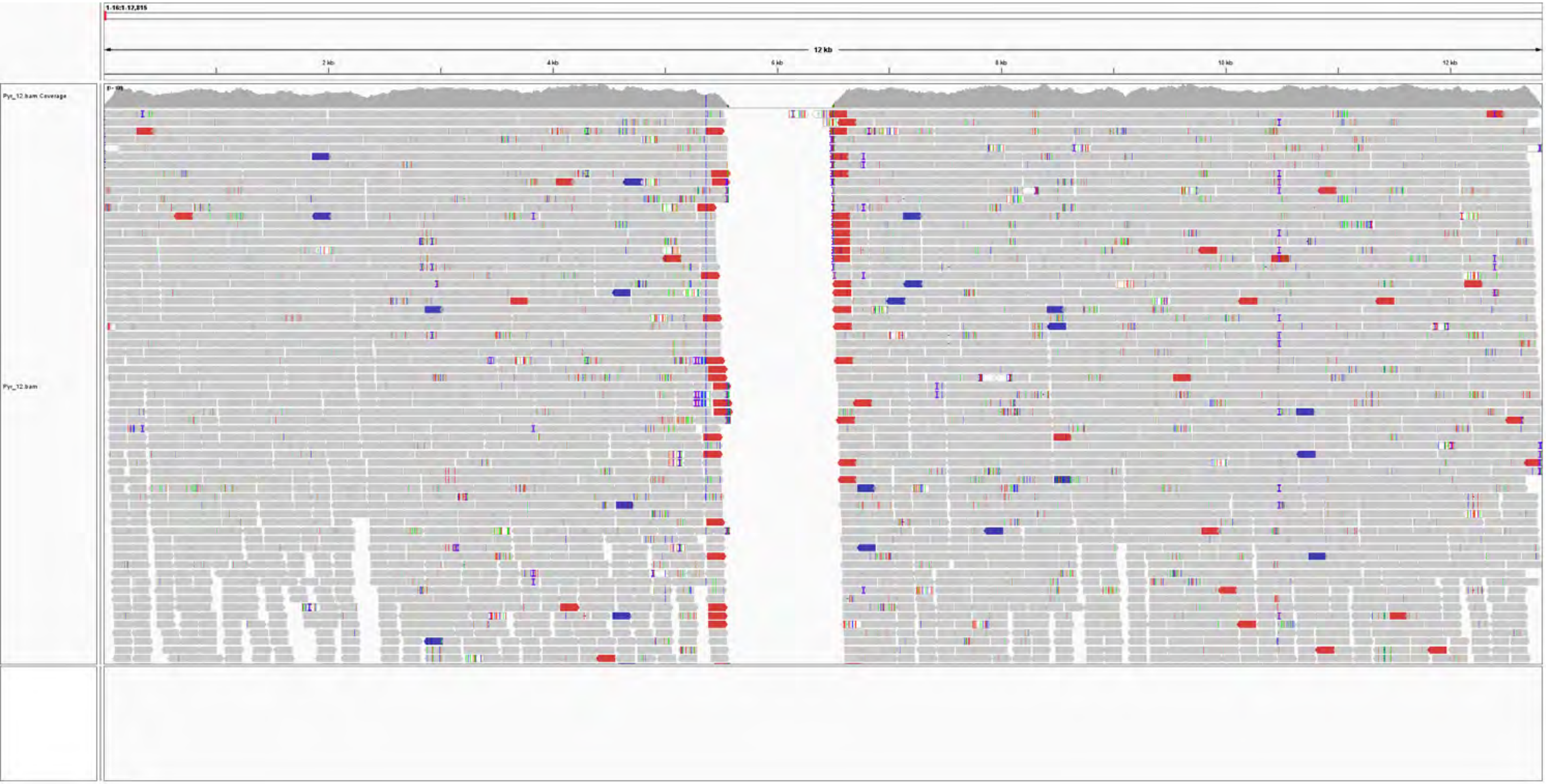

1-16:1-12,815
12 kb
2 kb
4 kb
6 kb
8 kb
10 kb
12 kb
Pyr_12.bam Coverage
Pyr_12.bam

### Pyr_13

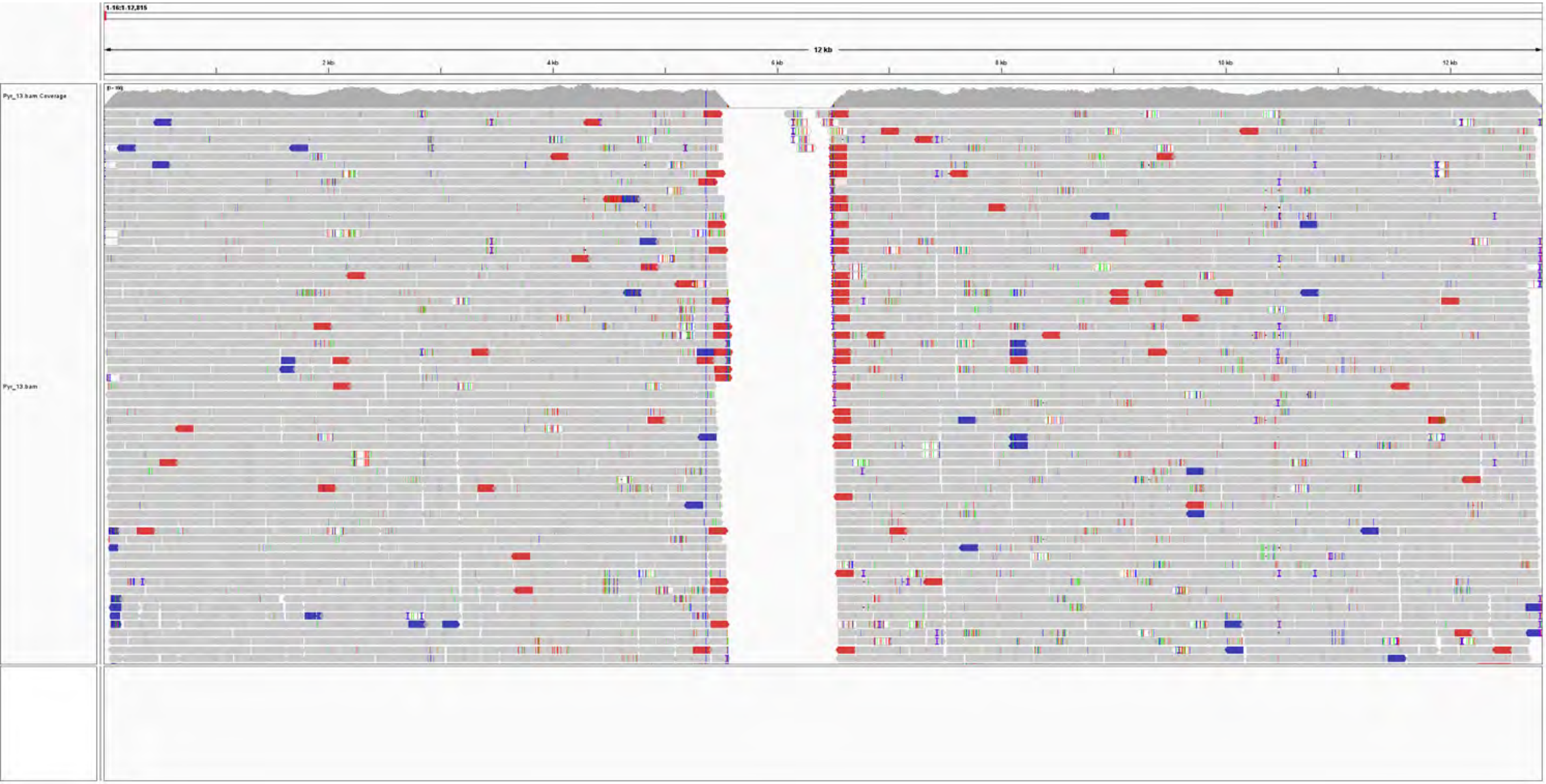

1-16:1-12,815
12 kb
2 kb
4 kb
6 kb
8 kb
10 kb
12 kb
Pyr_13.bam Coverage
Pyr_13.bam

### Pyr_14

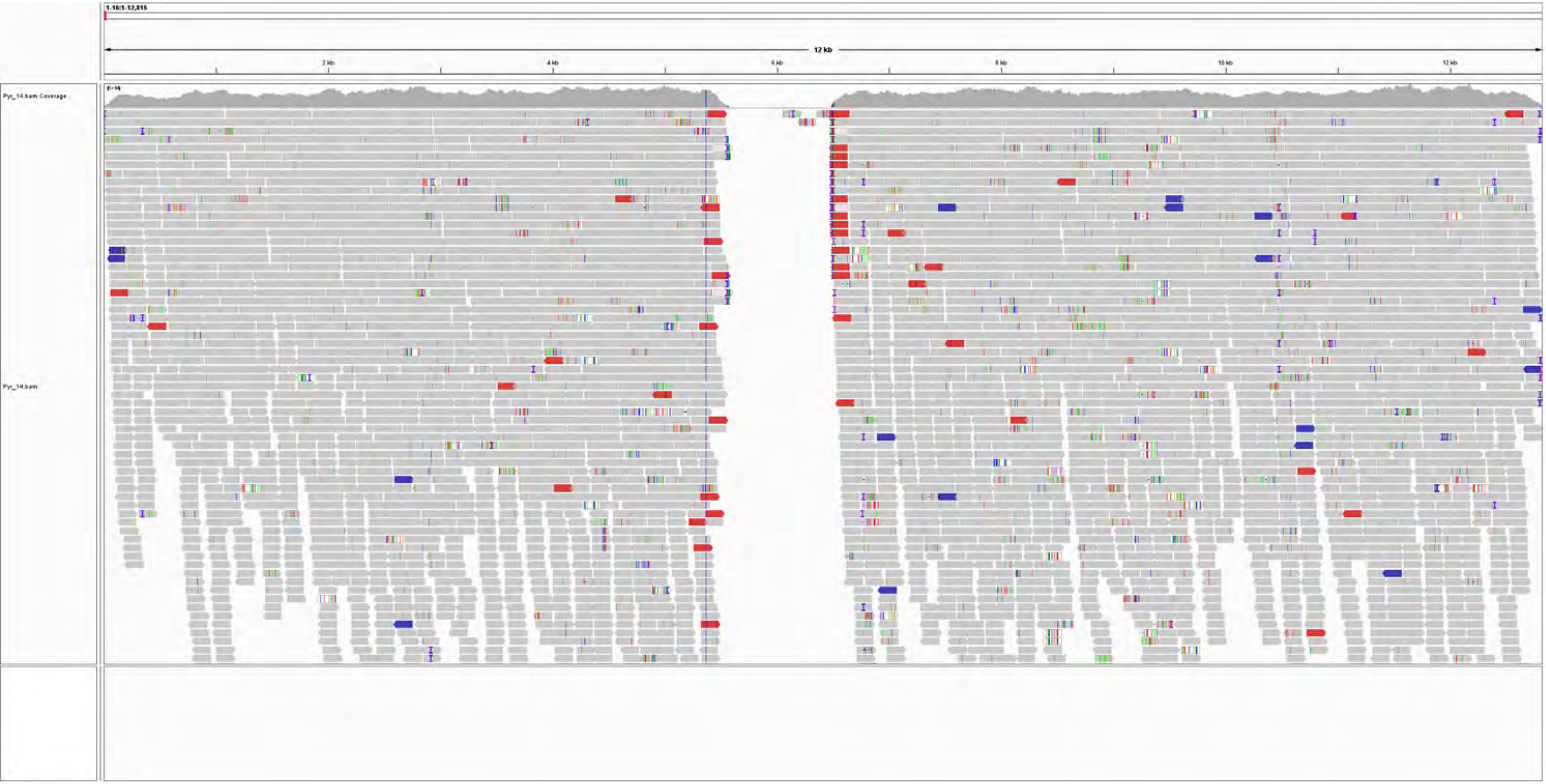

### Pyr_15

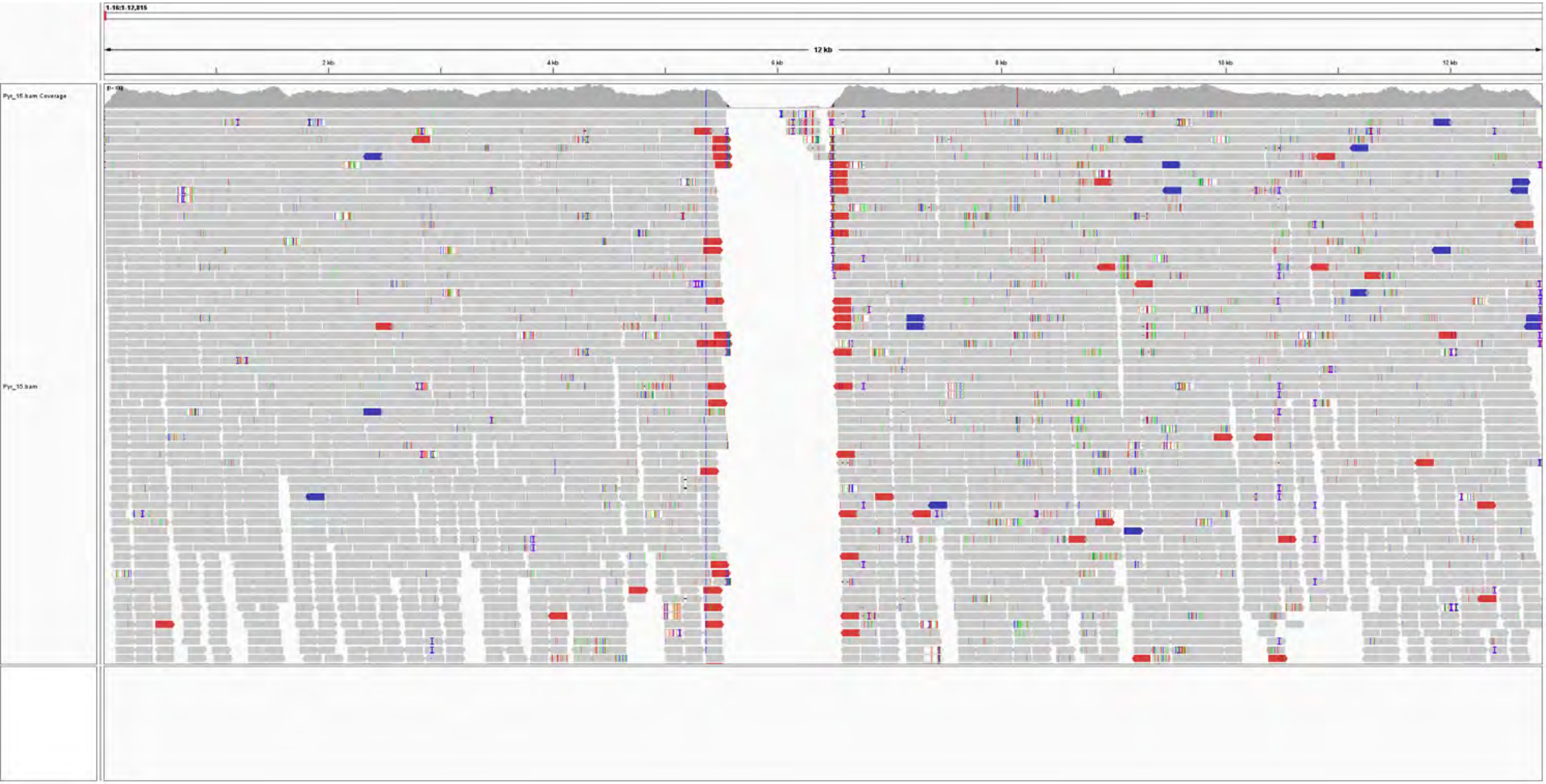

1-16:1-12,815
12 kb
2 kb
4 kb
6 kb
8 kb
10 kb
12 kb
Pyr_15.bam Coverage
Pyr_15.bam

### Pyr_16

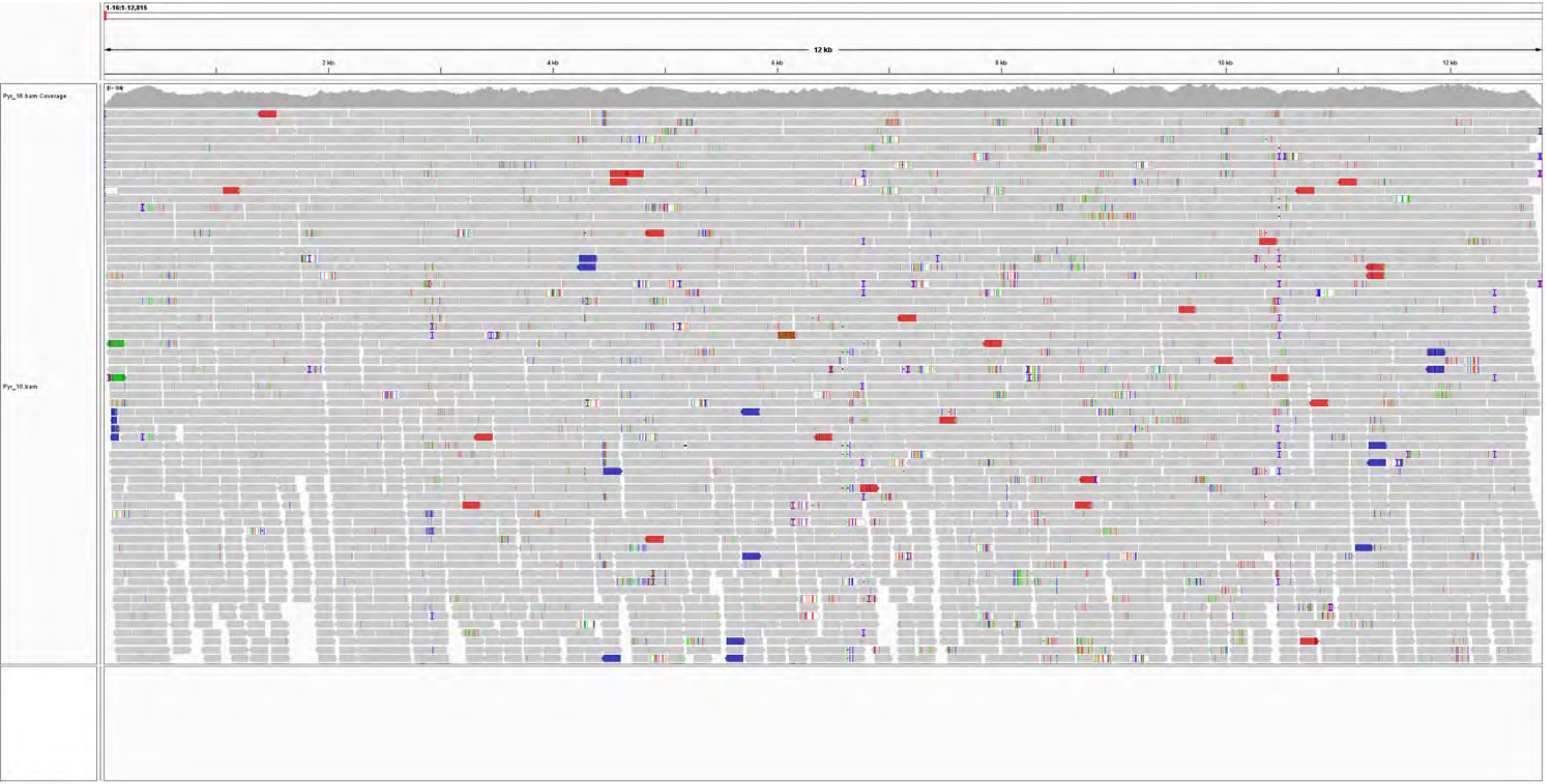

1-16:1-12,815
12 kb
2 kb
4 kb
6 kb
8 kb
10 kb
12 kb
Pyr_16.bam Coverage
Pyr_16.bam

### Pyr_17

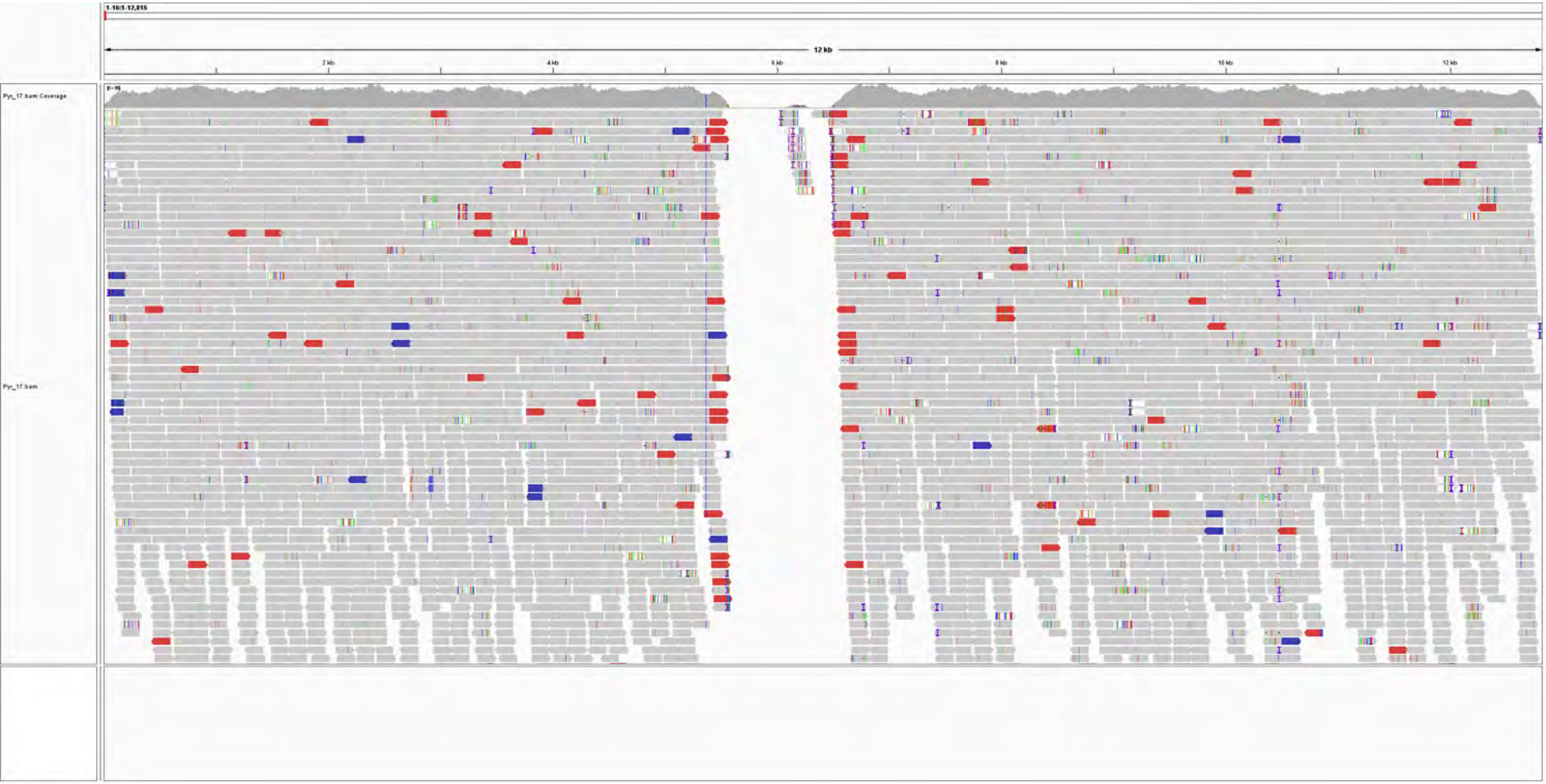

### Pyr_18

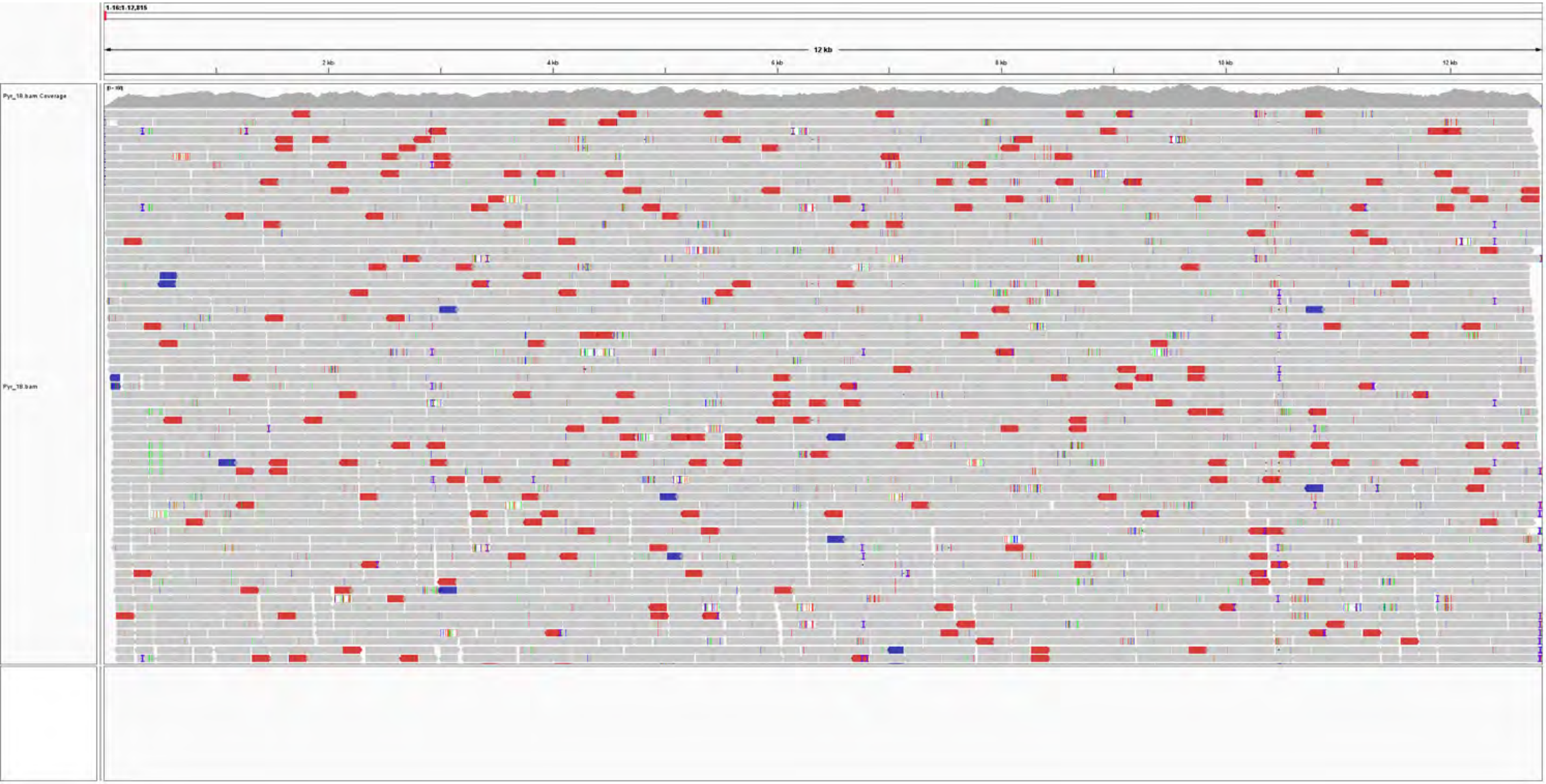

### Pyr_20

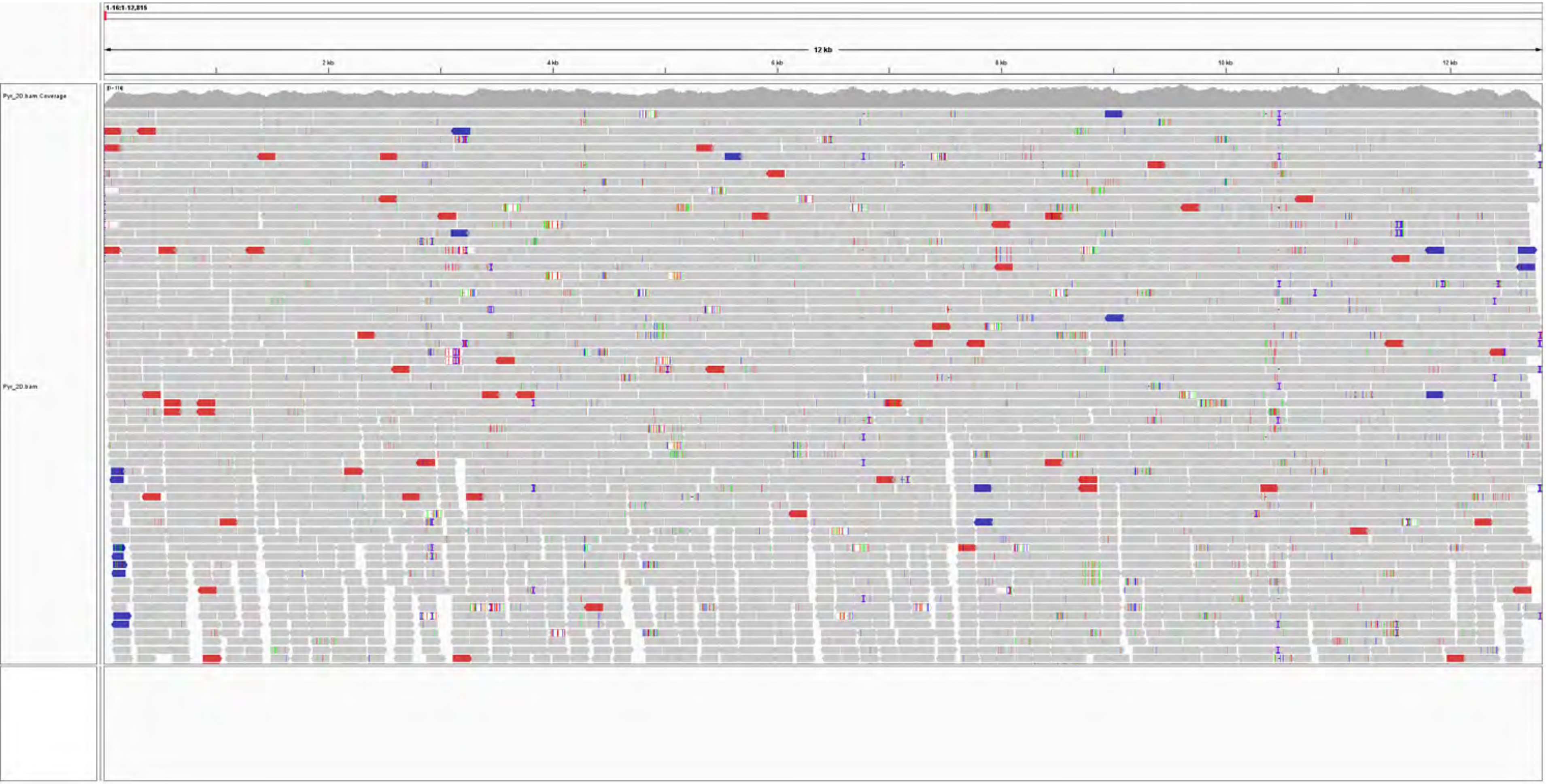
1-16:1-12,815
12 kb
2 kb
4 kb
6 kb
8 kb
10 kb
12 kb
Pyr_20.bam Coverage
[0 - 116]
Pyr_20.bam

### Pyr_21

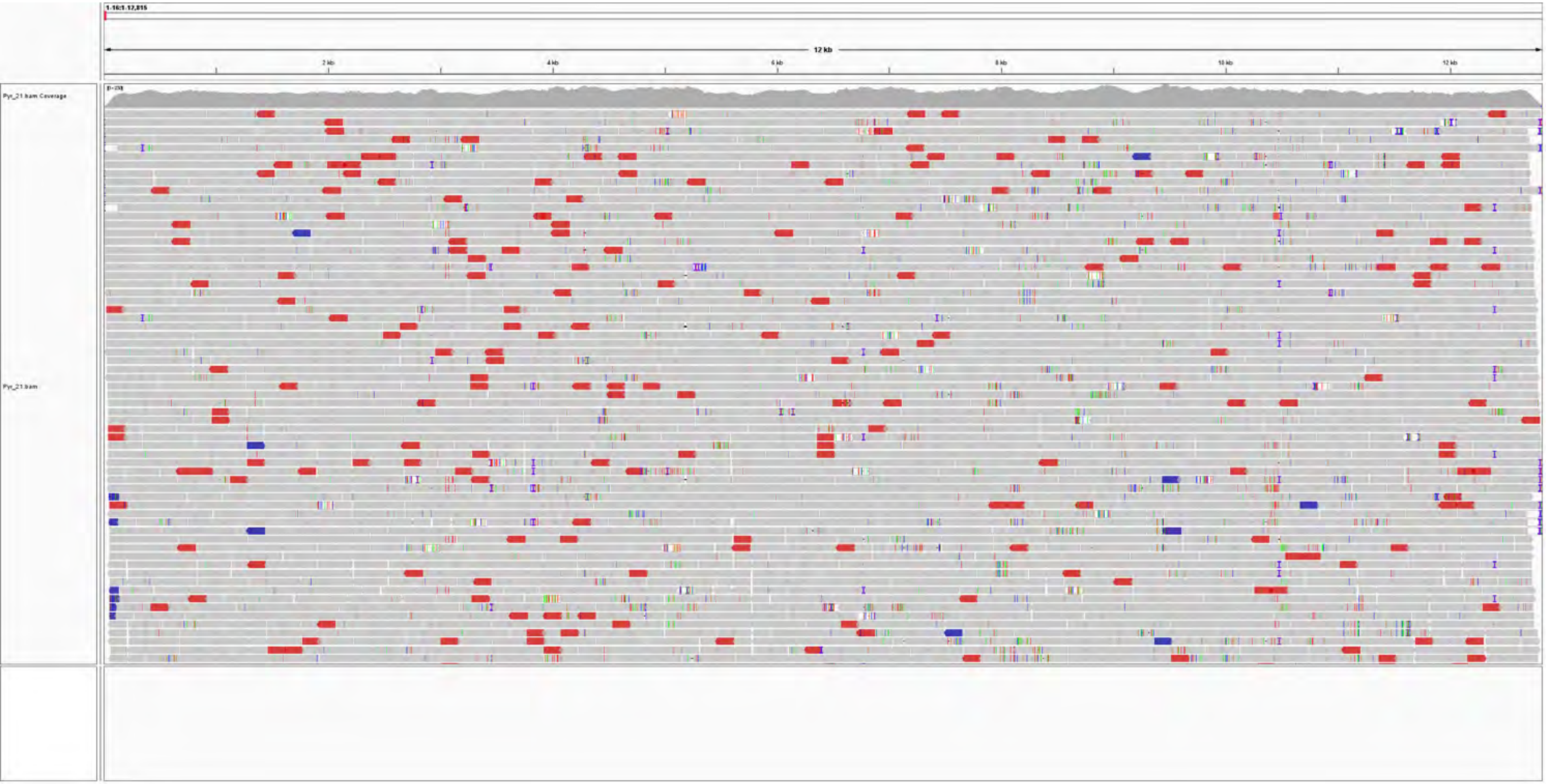

1:16:1-12,815
12 kb
2 kb
4 kb
6 kb
8 kb
10 kb
12 kb
Pyr_21.bam Coverage
Pyr_21.bam

### Pyr_22

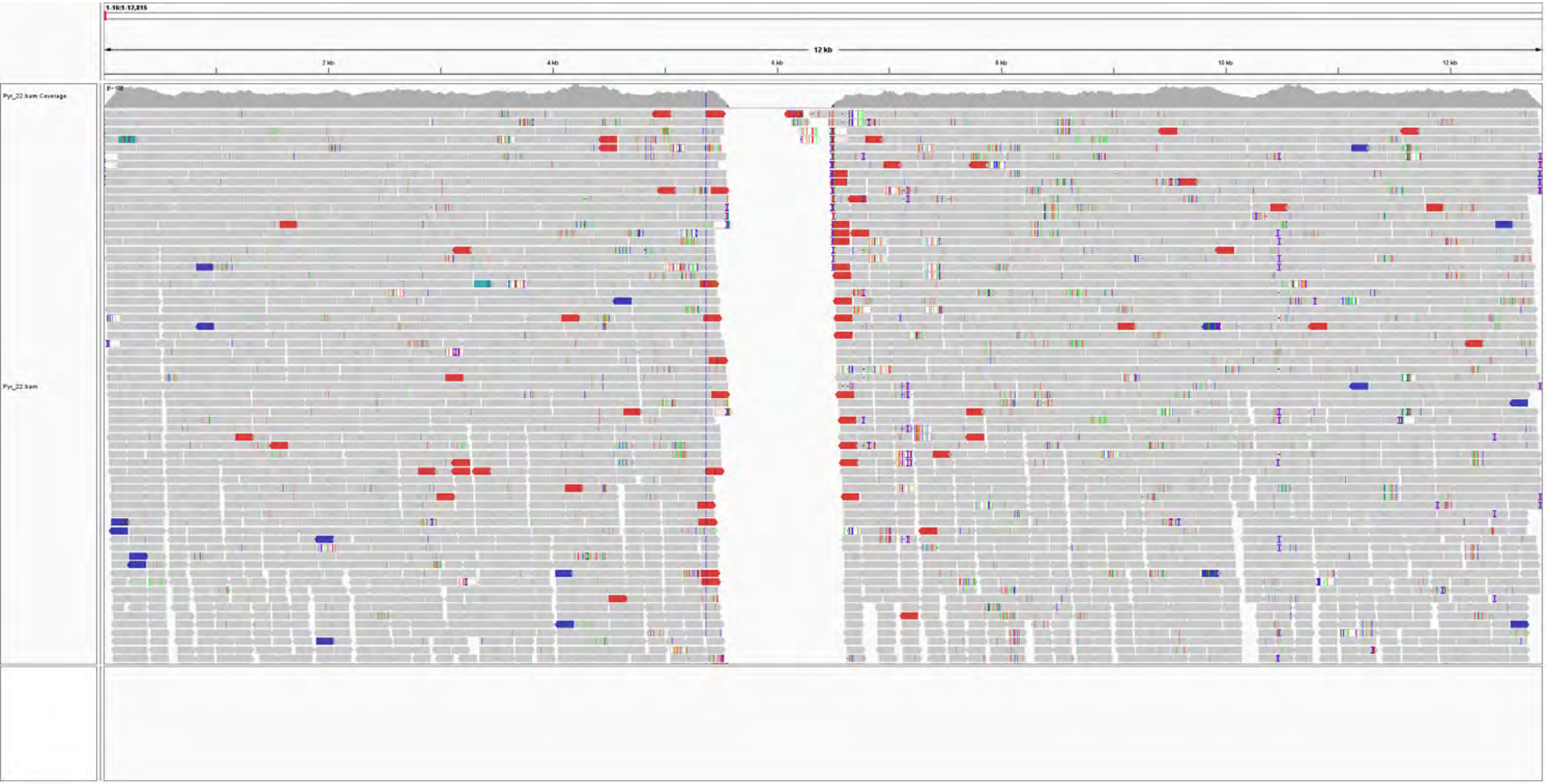
1-16:1-12,815
12 kb
2 kb
4 kb
6 kb
8 kb
10 kb
12 kb
Pyr_22.bam Coverage
Pyr_22.bam

### Pyr_23

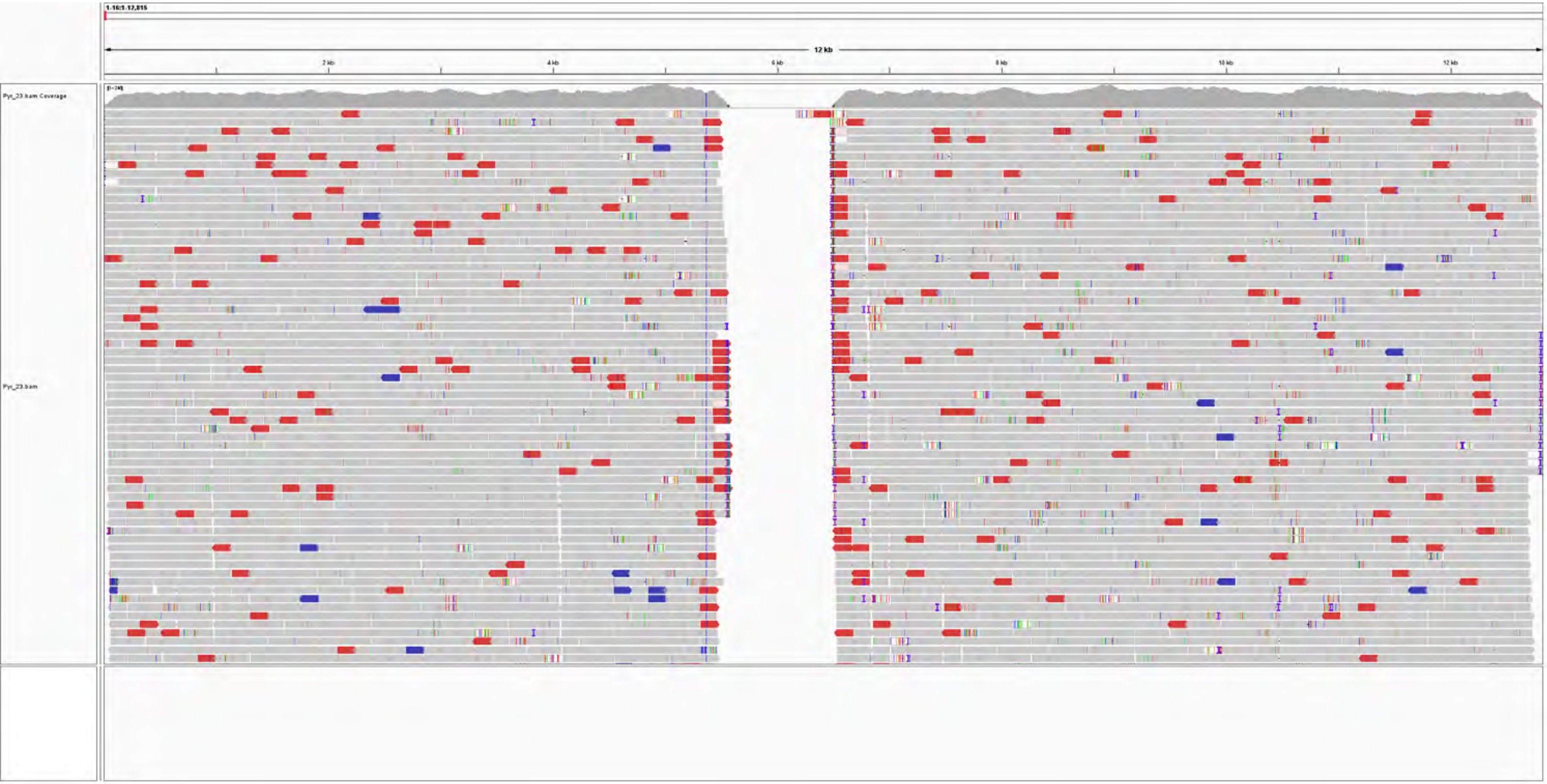

### Pyr_24

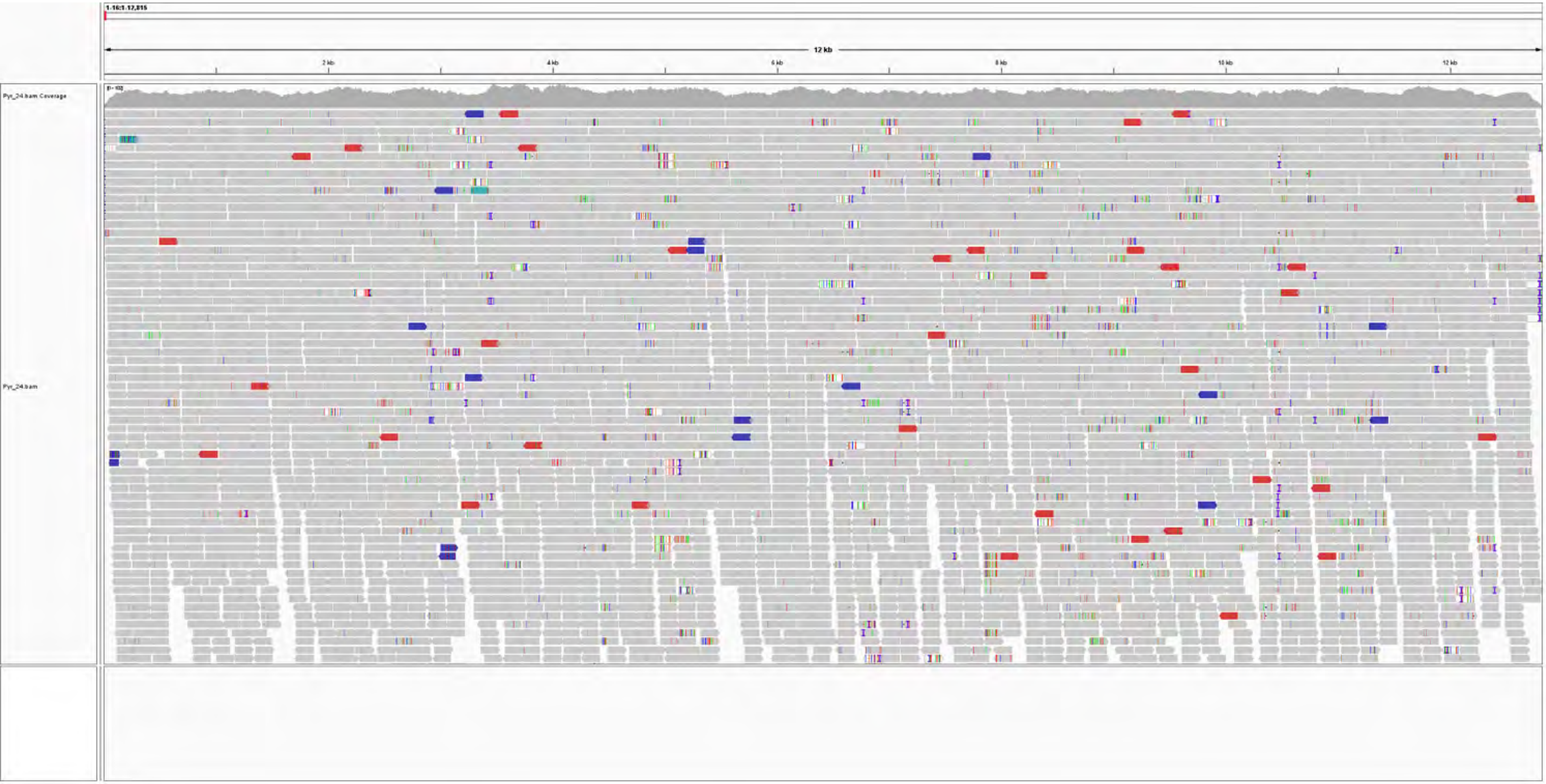
1-16:1-12,815
12 kb
2 kb
4 kb
6 kb
8 kb
10 kb
12 kb
Pyr_24.bam Coverage
Pyr_24.bam

### Pyr_25

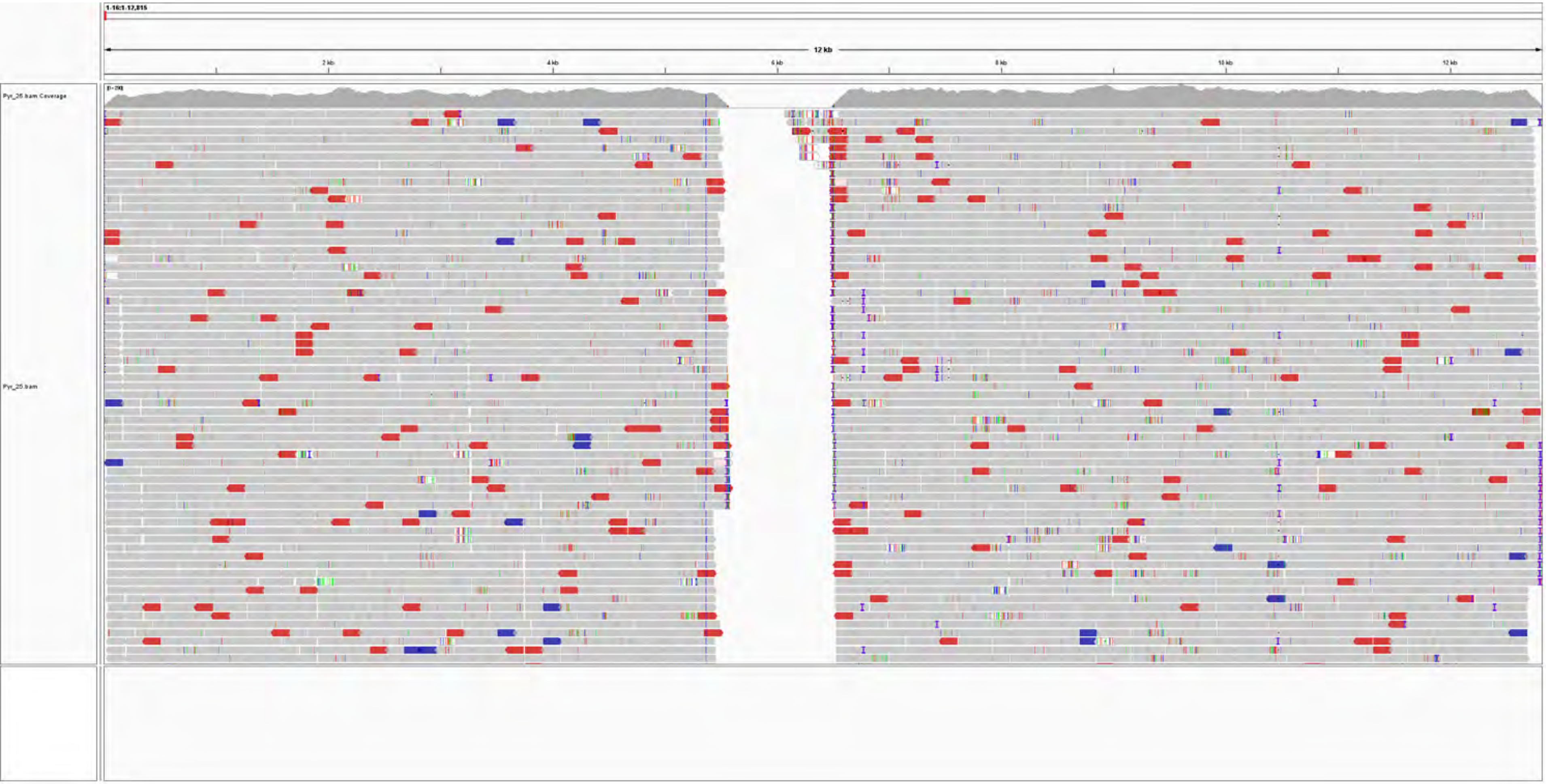

### Pyr_26

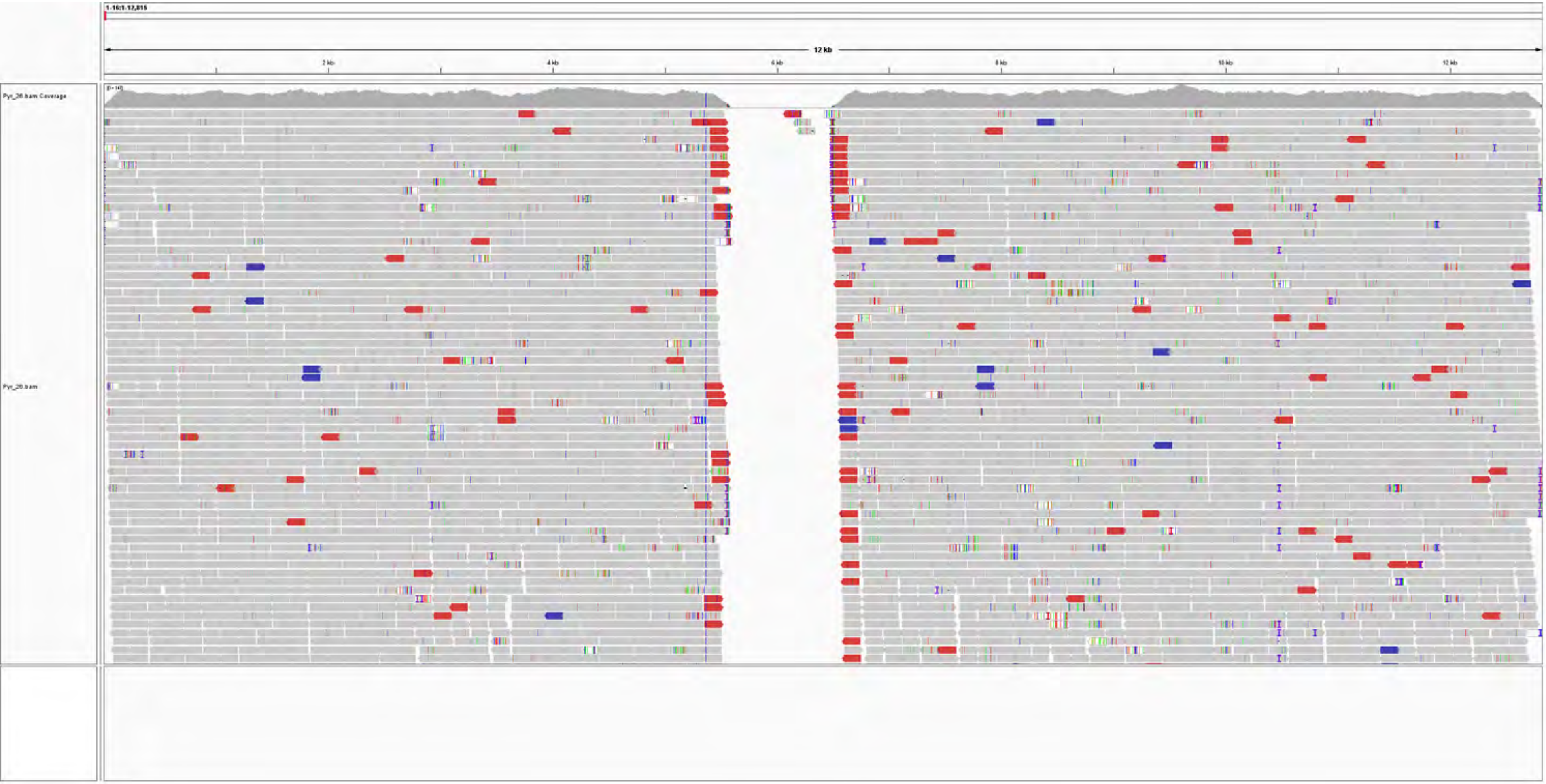

1-16:1-12,815
12 kb
2 kb
4 kb
6 kb
8 kb
10 kb
12 kb
Pyr_26.bam Coverage
Pyr_26.bam

### Pyr_28

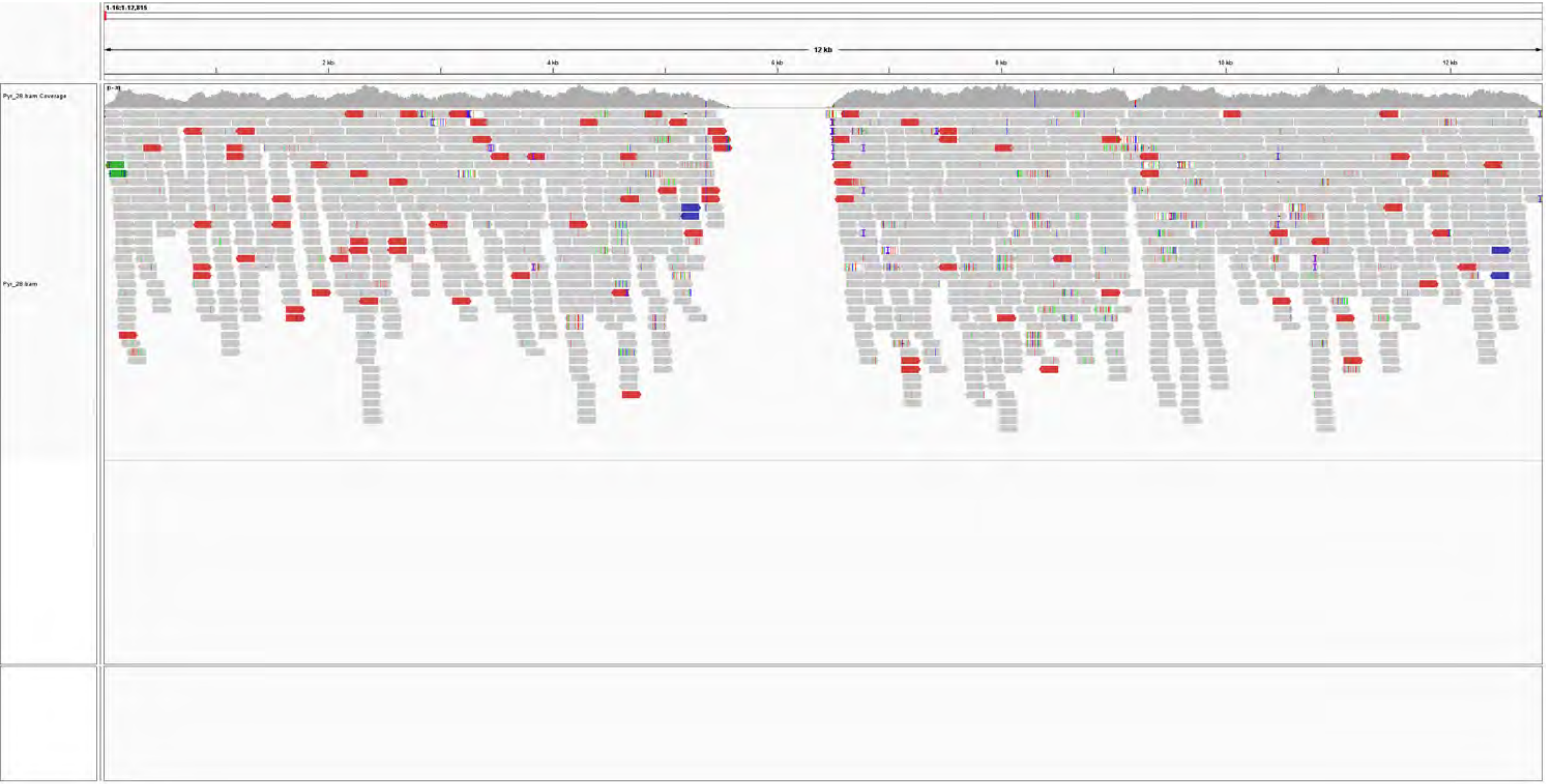

### Pyr_29

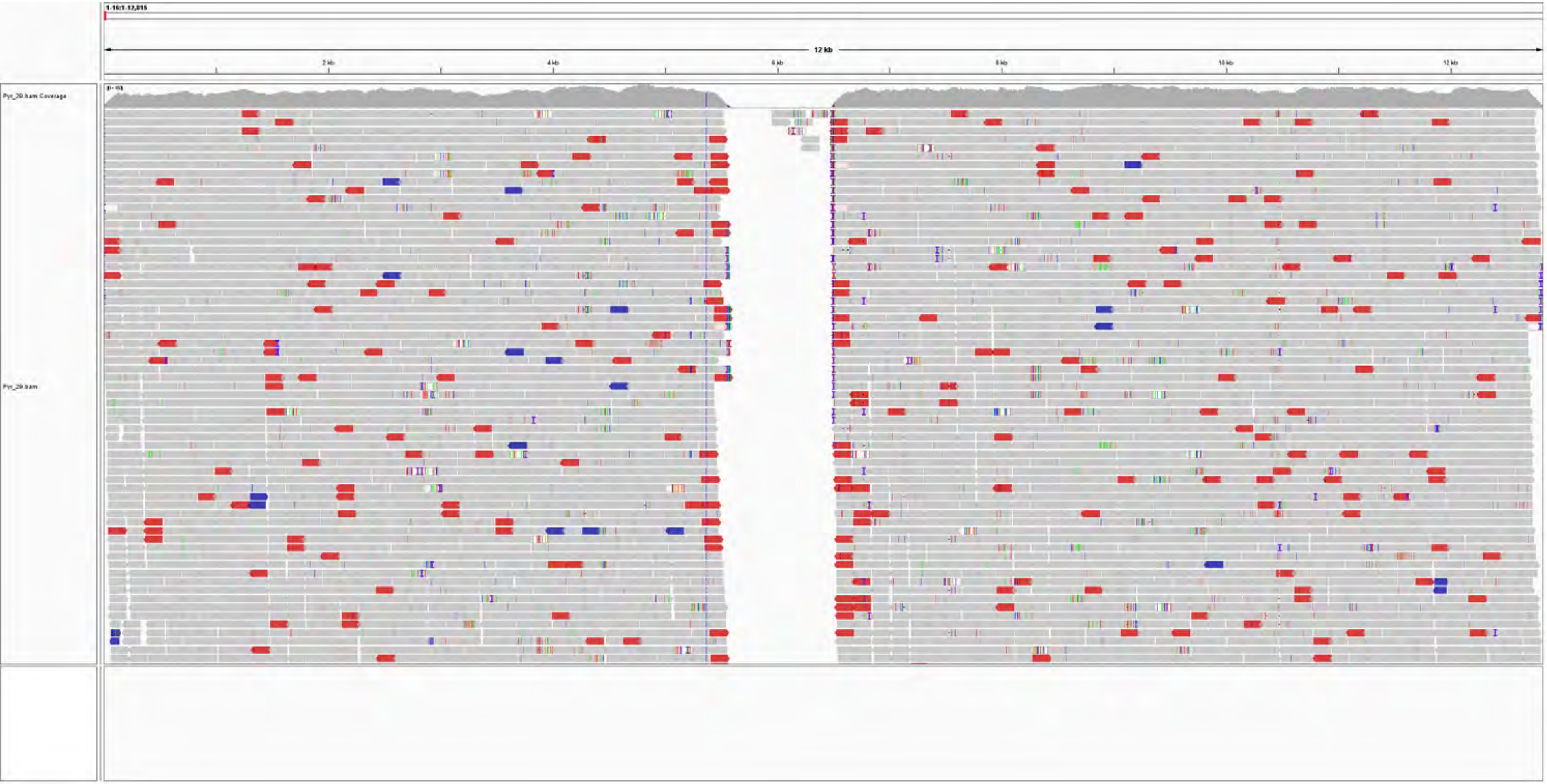

### Pyr_30

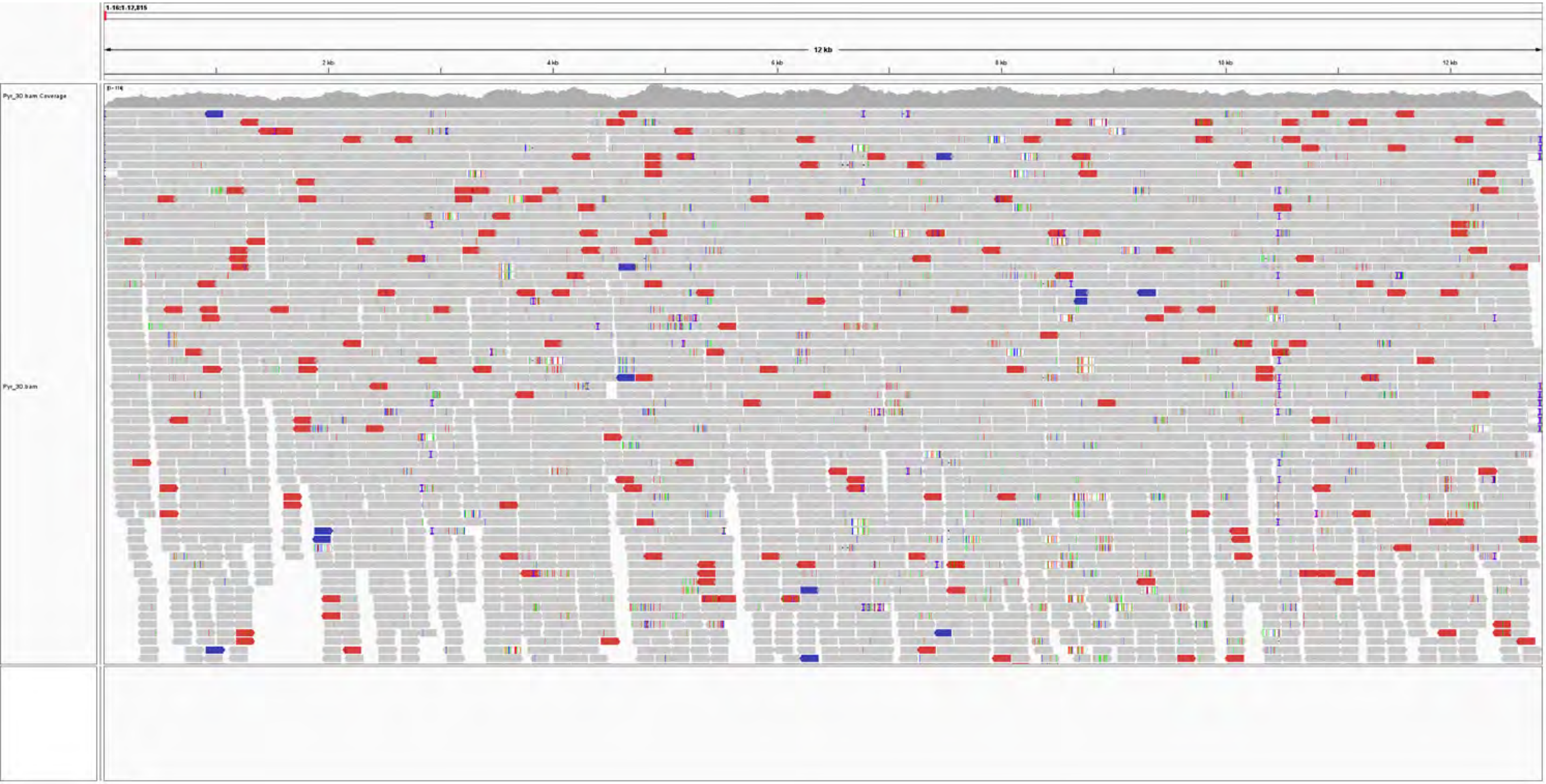

1-16:1-12,815
12 kb
2 kb
4 kb
6 kb
8 kb
10 kb
12 kb
Pyr_30.bam Coverage
[0 - 114]
Pyr_30.bam

### Pyr_33

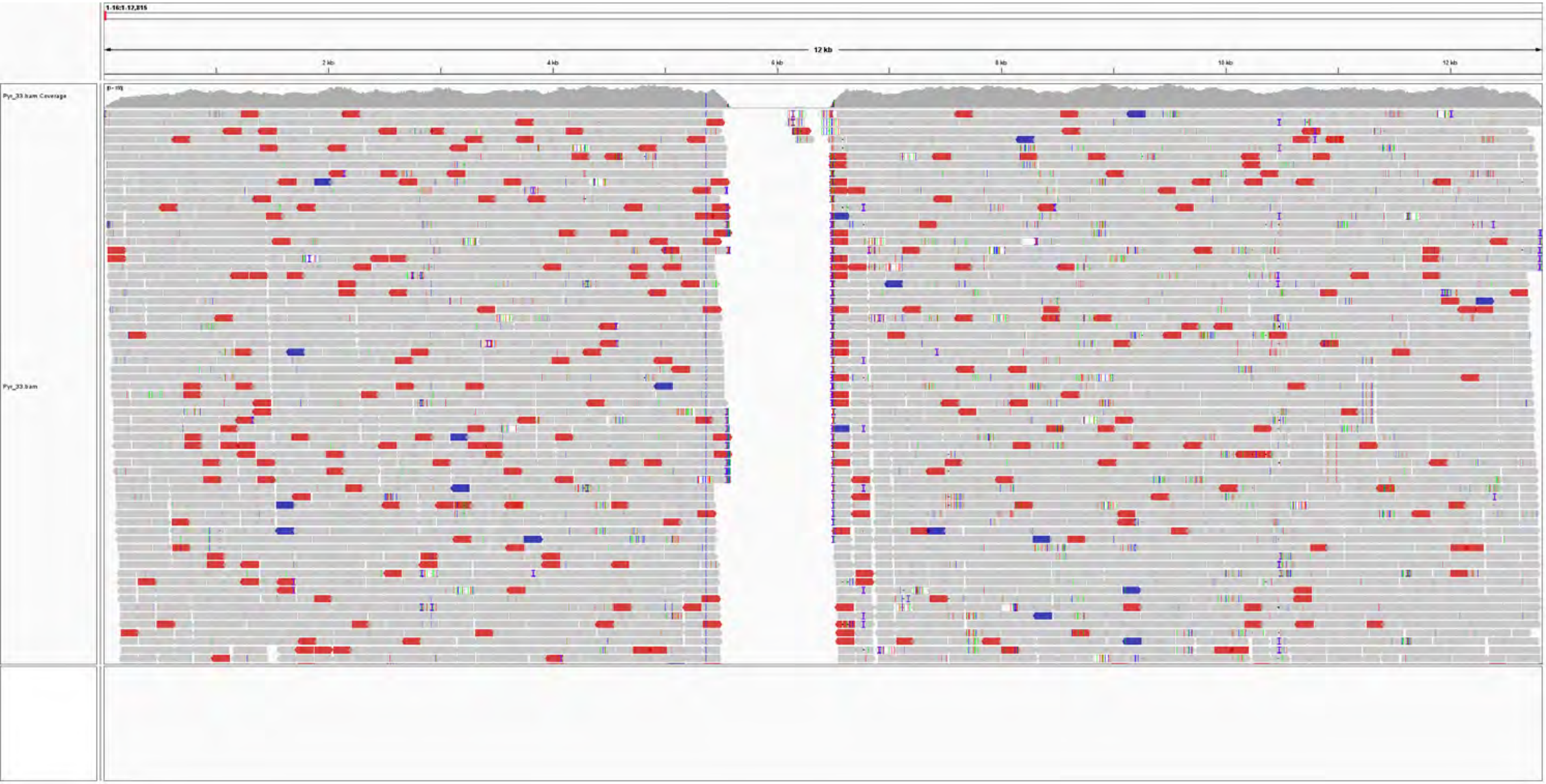

### Pyr_34

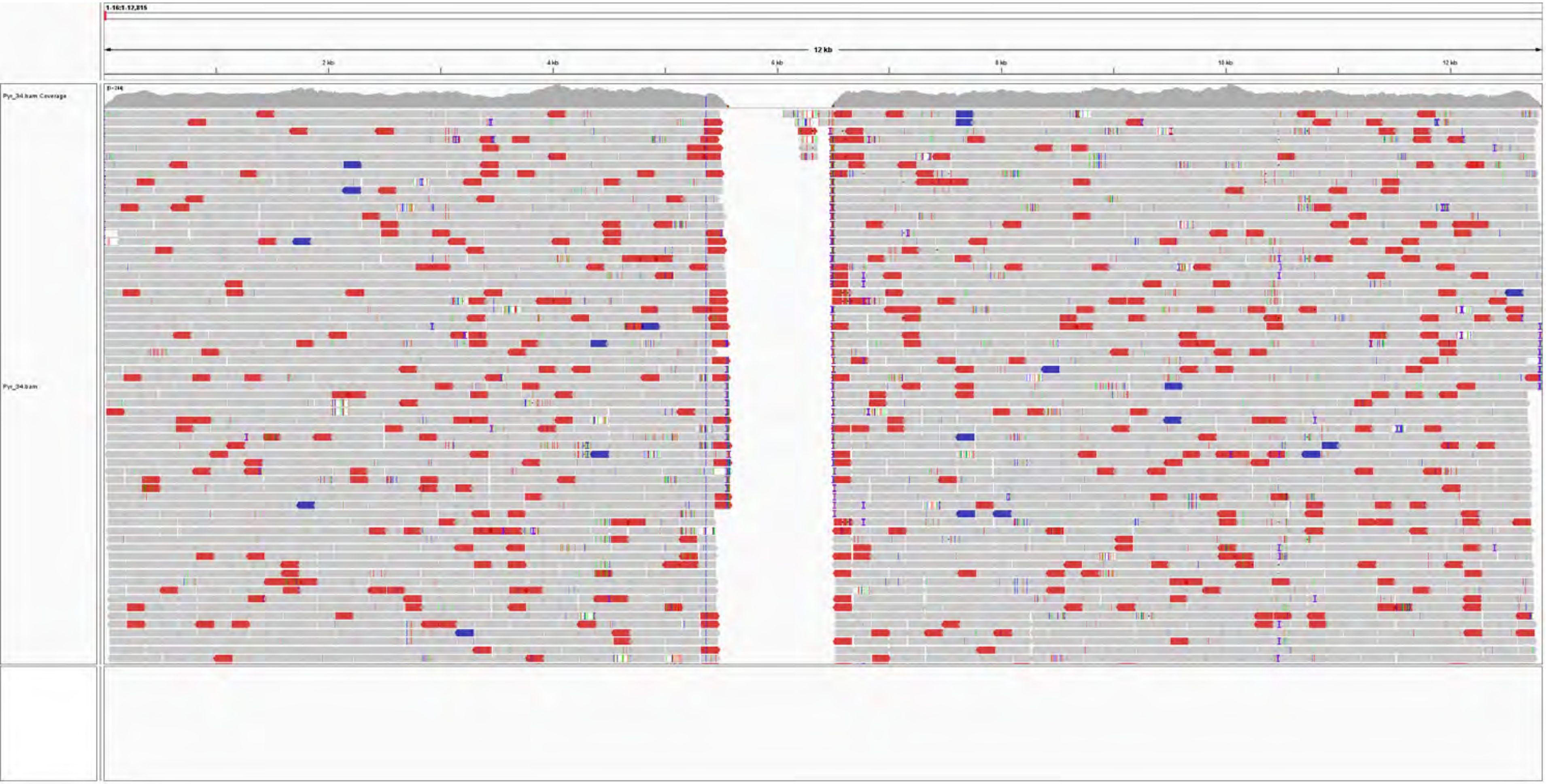

### Pyr_36

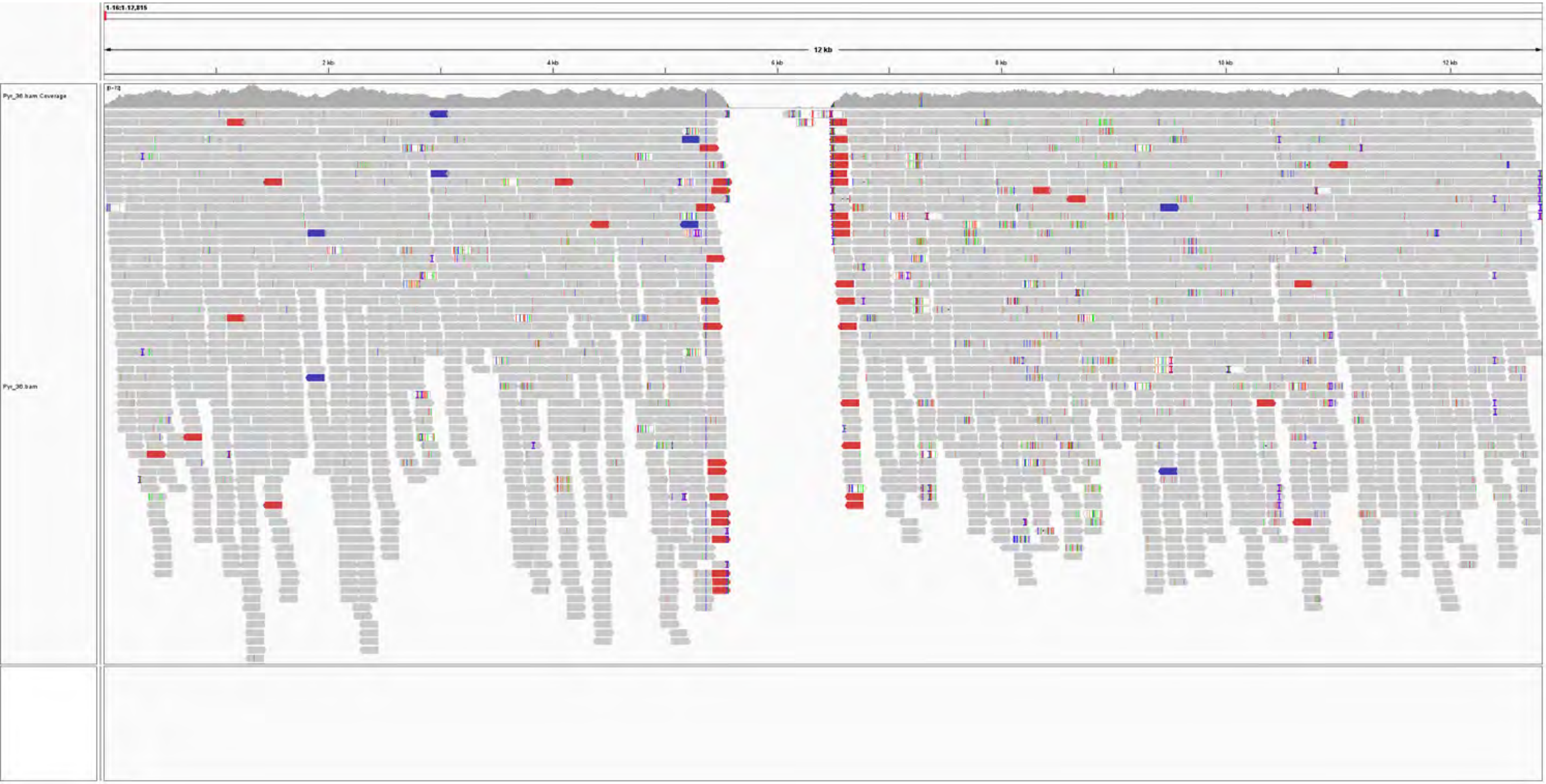

### Pyr_38

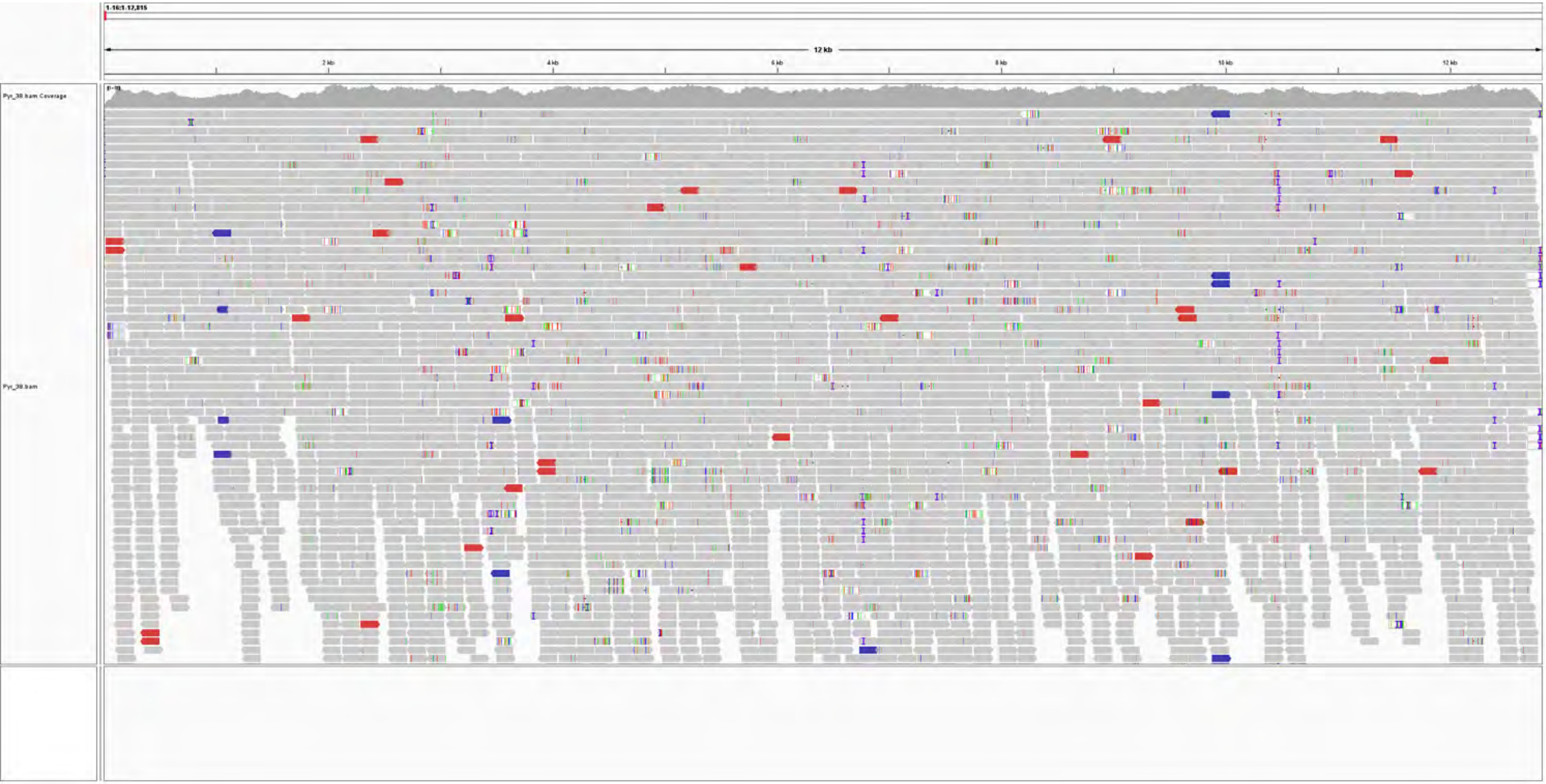

1-16:1-12,815
12 kb
2 kb
4 kb
6 kb
8 kb
10 kb
12 kb
Pyr_38.bam Coverage
Pyr_38.bam

### Pyr_39

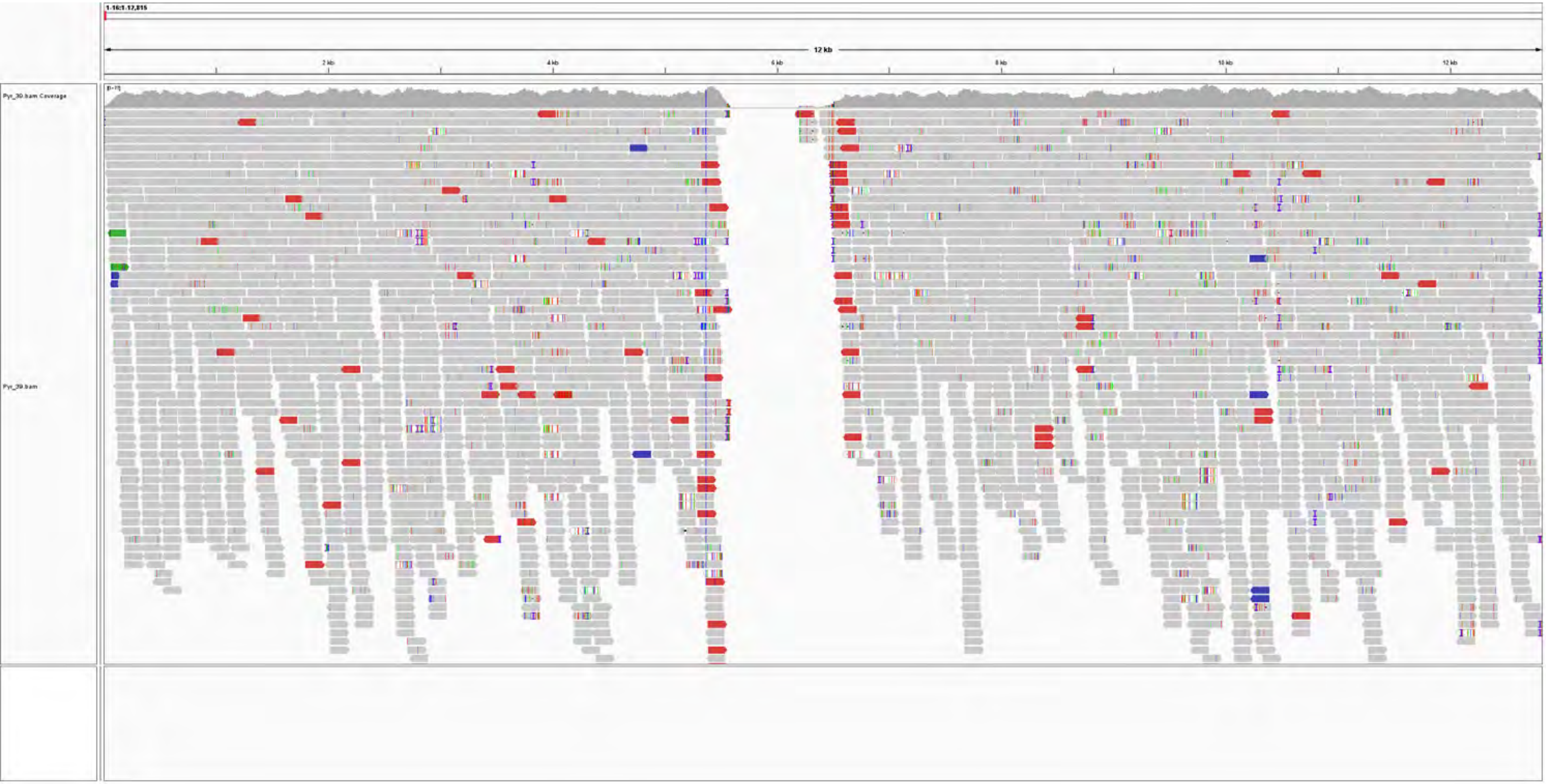

### Pyr_40

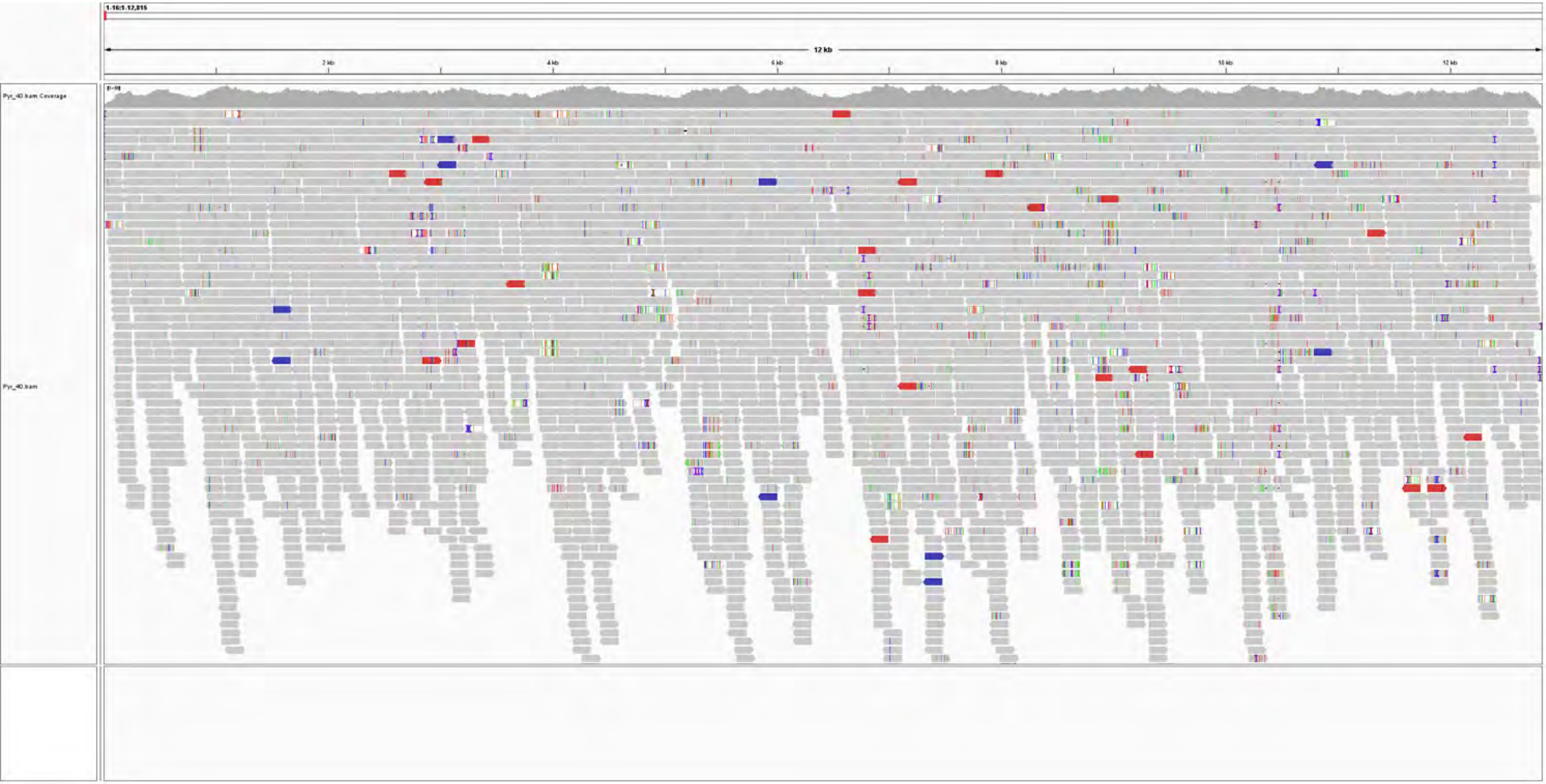

### Pyr_41

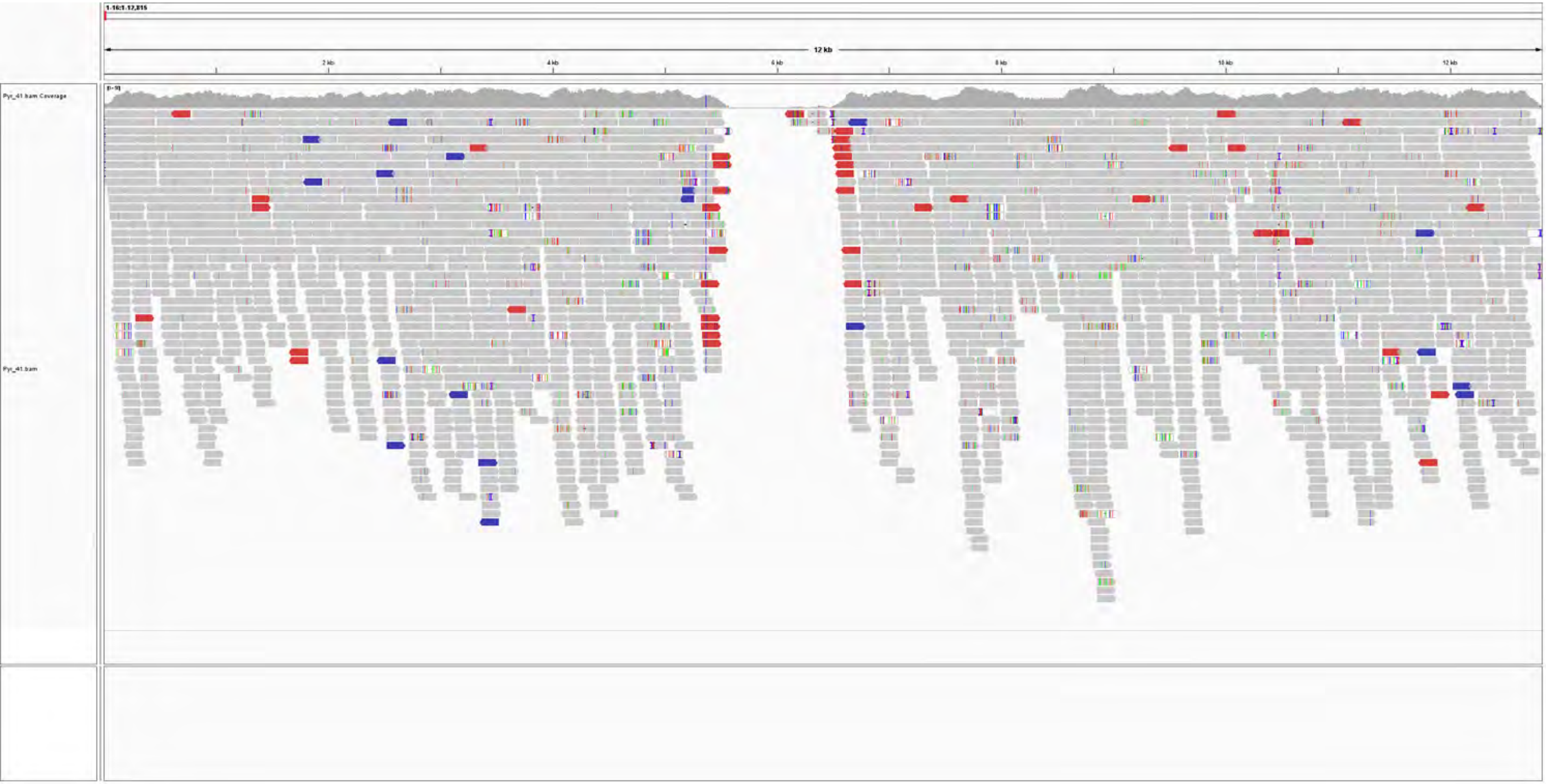
1-16:1-12,815
12 kb
2 kb
4 kb
6 kb
8 kb
10 kb
12 kb
Pyr_41.bam Coverage
Pyr_41.bam

### Pyr_42

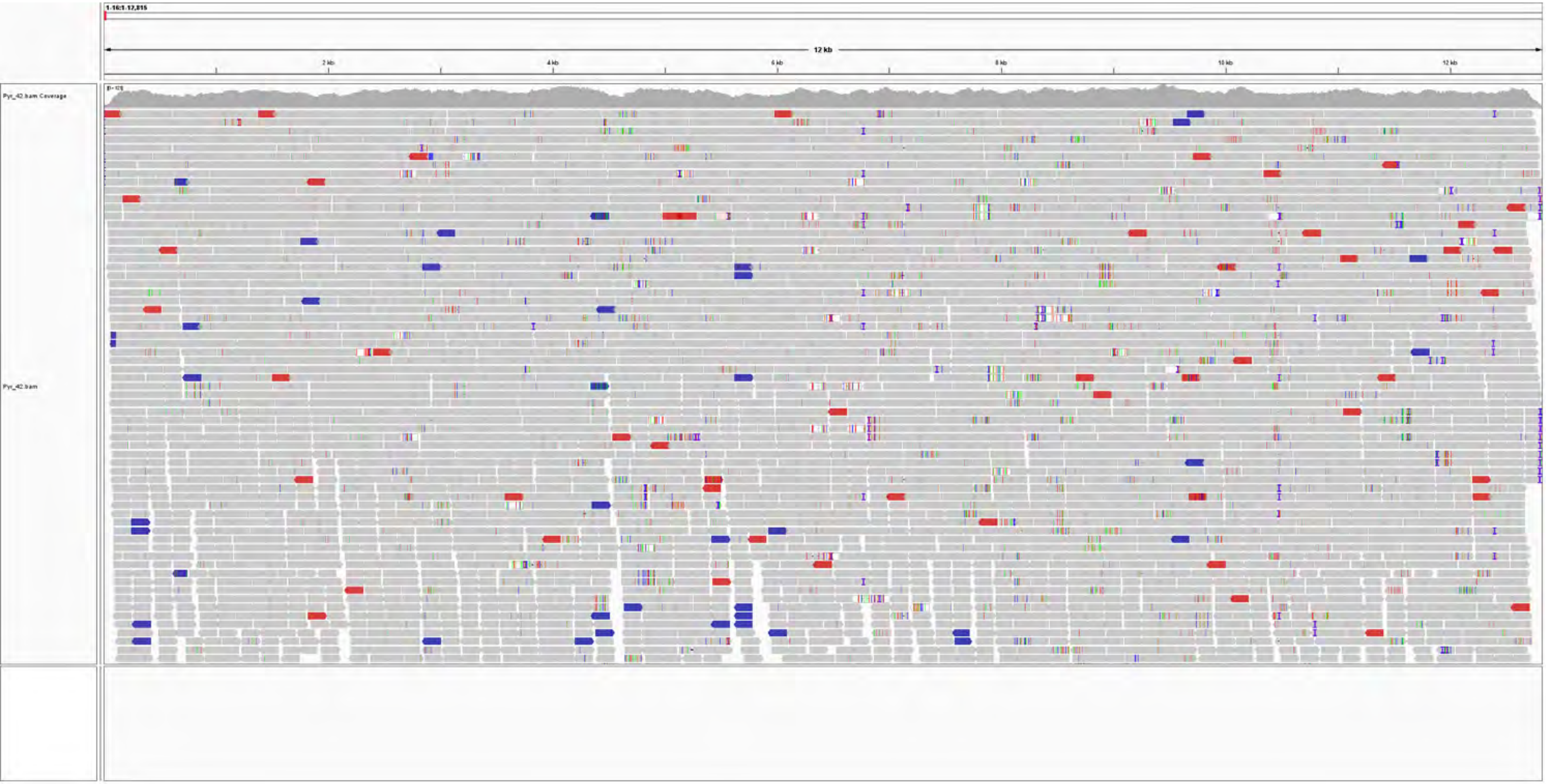

1-16:1-12,815
12 kb
2 kb
4 kb
6 kb
8 kb
10 kb
12 kb
Pyr_42.bam Coverage
Pyr_42.bam

**Supplementary Figure 8.** The result of mapping the reads of each sample to its own mitochondrial sequence.

### SRR9587923

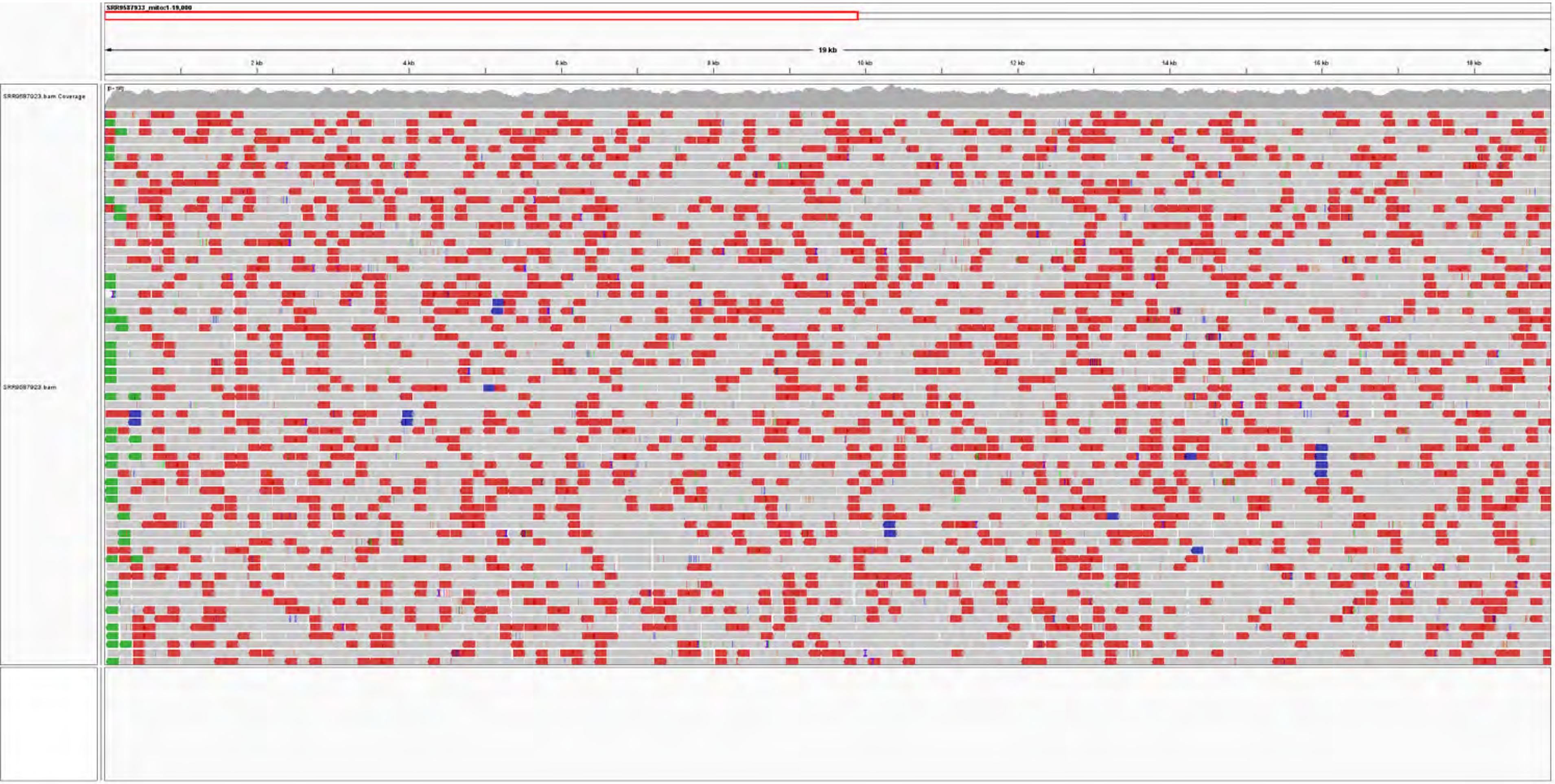

### SRR9587923 (continued)

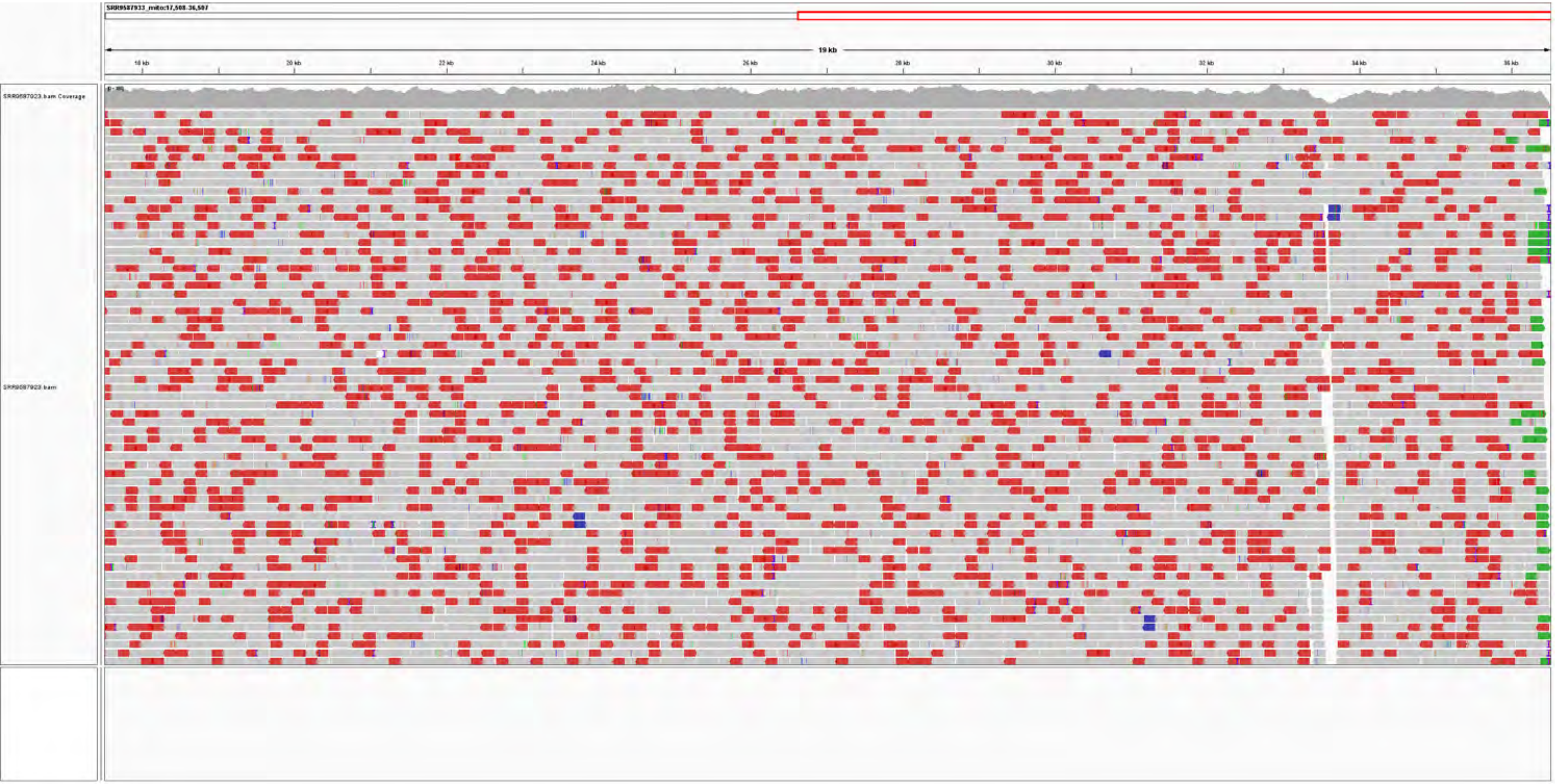

### SRR9587933

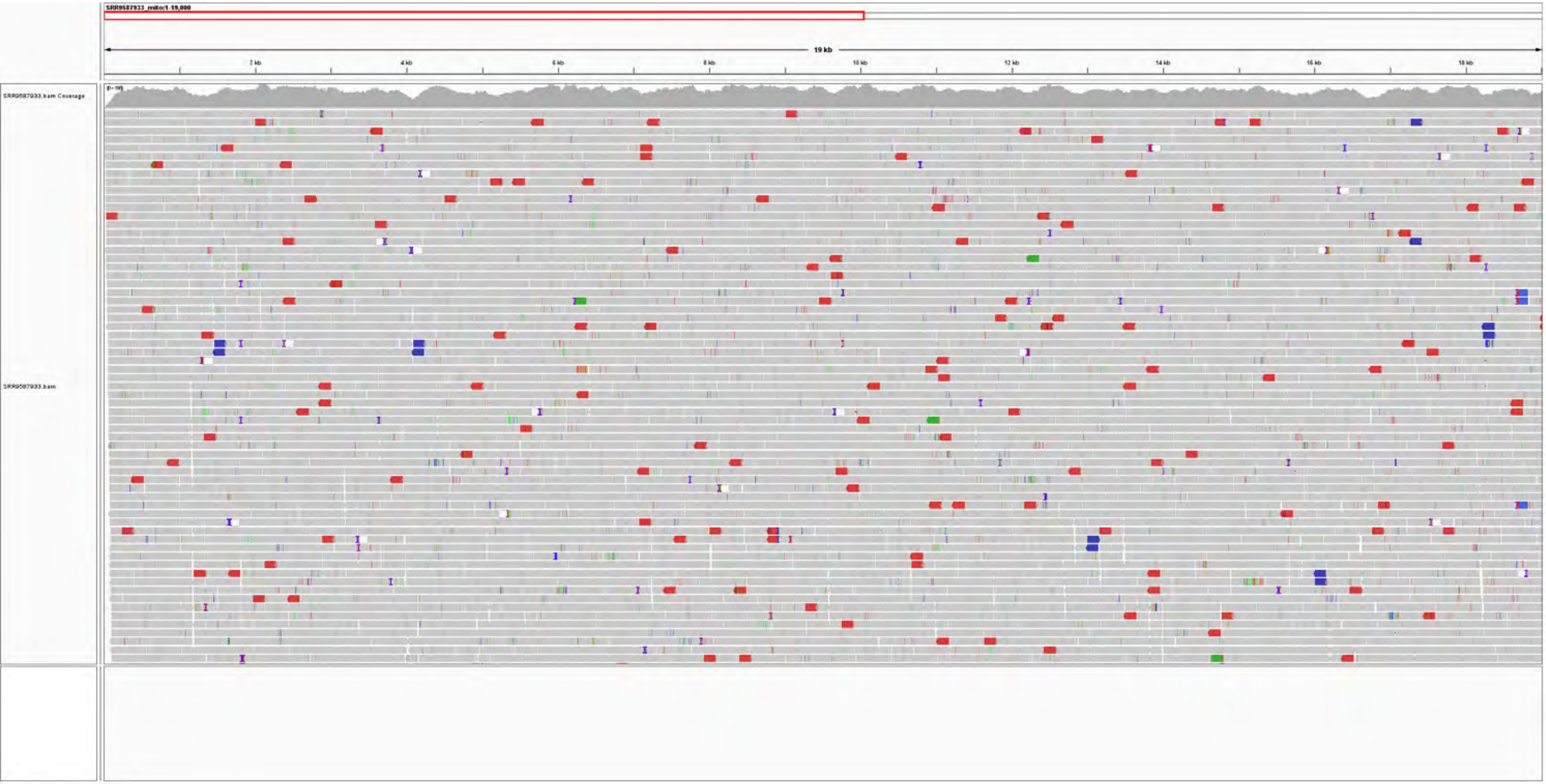

### SRR9587933 (continued)

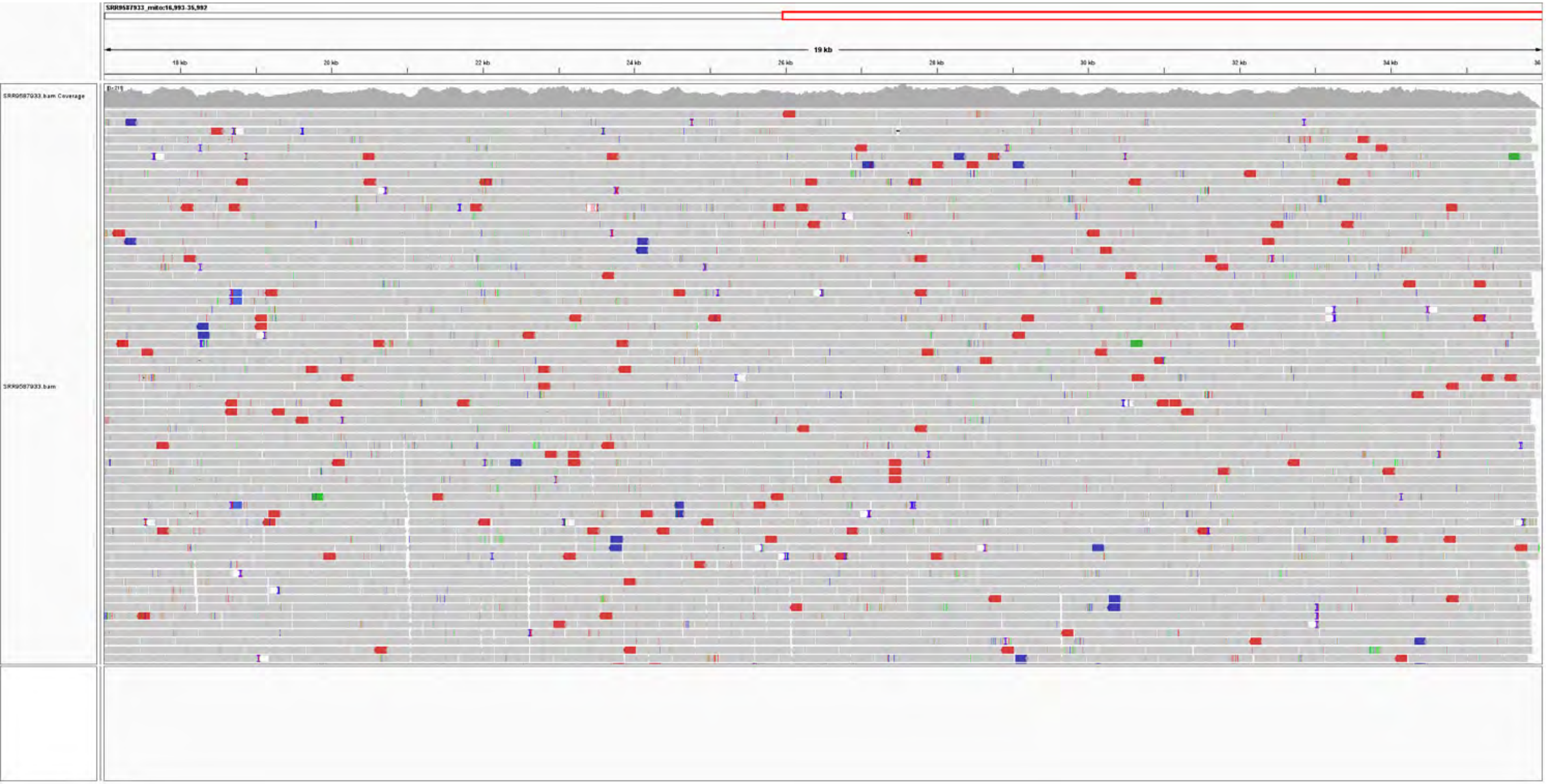

### SRR9587936

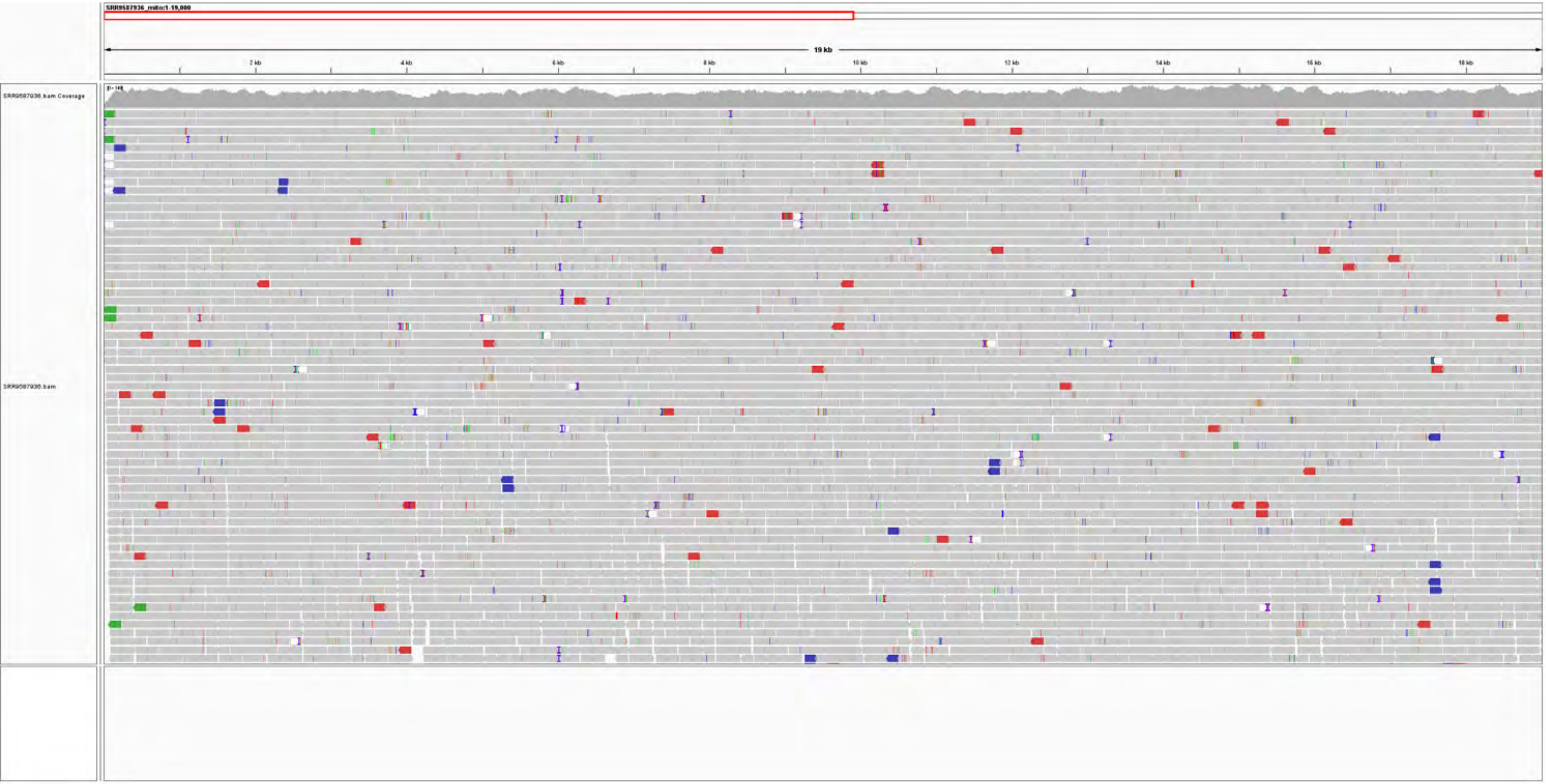

### SRR9587936 (continued)

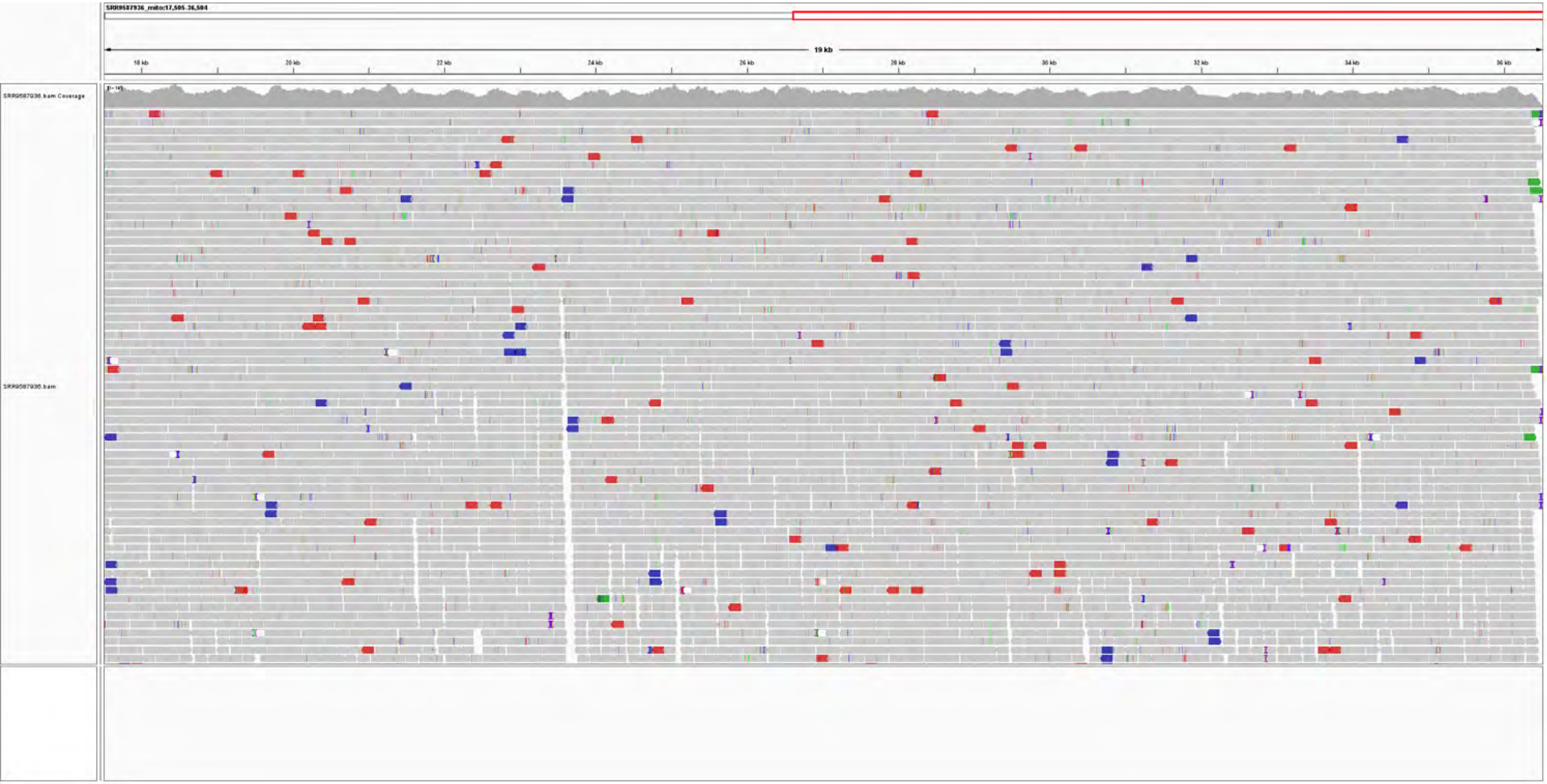

### SRR9587966

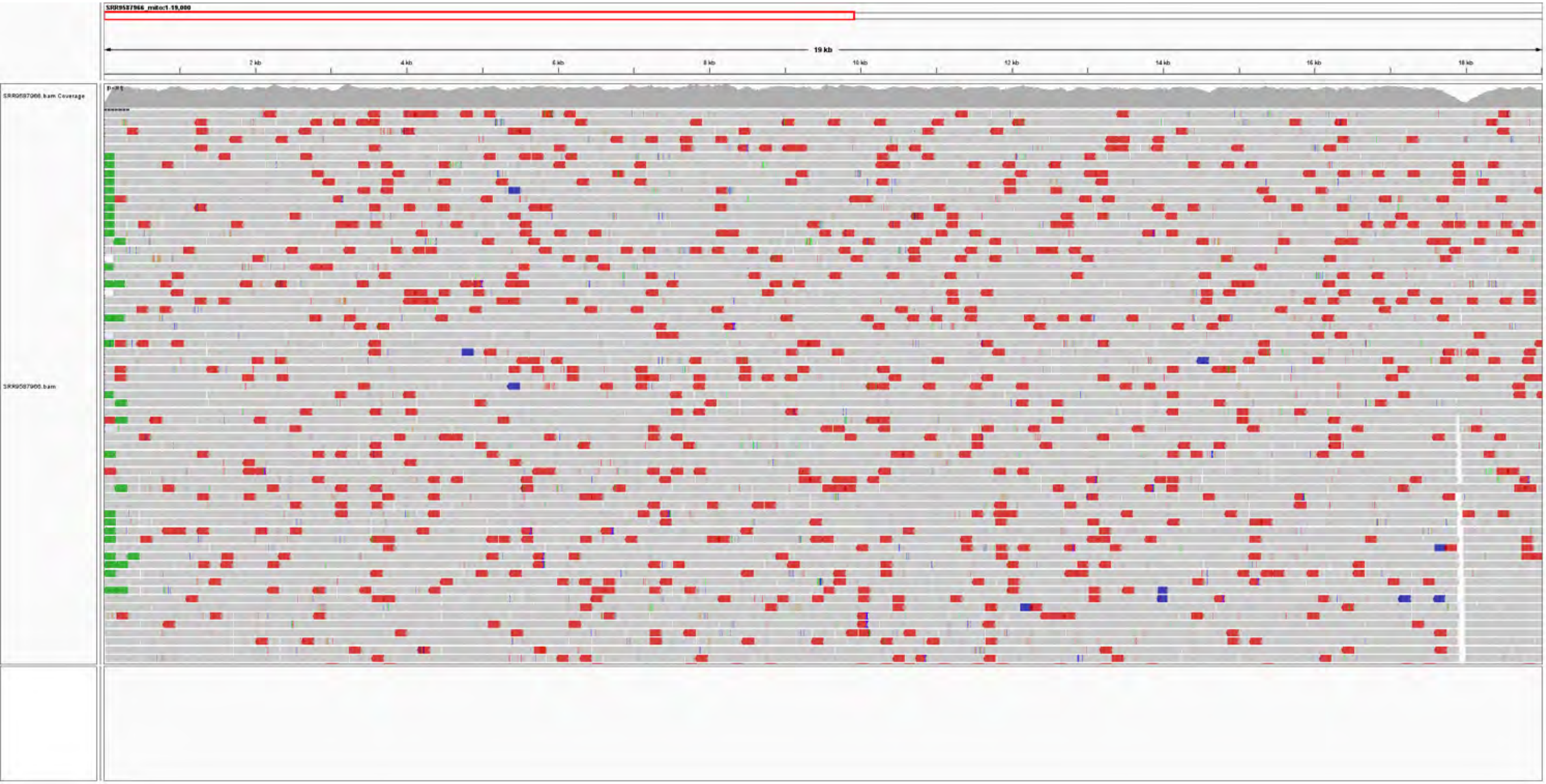

SRR9587966_mito:1-19,000
19 kb
2 kb
4 kb
6 kb
8 kb
10 kb
12 kb
14 kb
16 kb
18 kb
SRR9587966.bam Coverage
SRR9587966.bam

### SRR9587966 (continued)

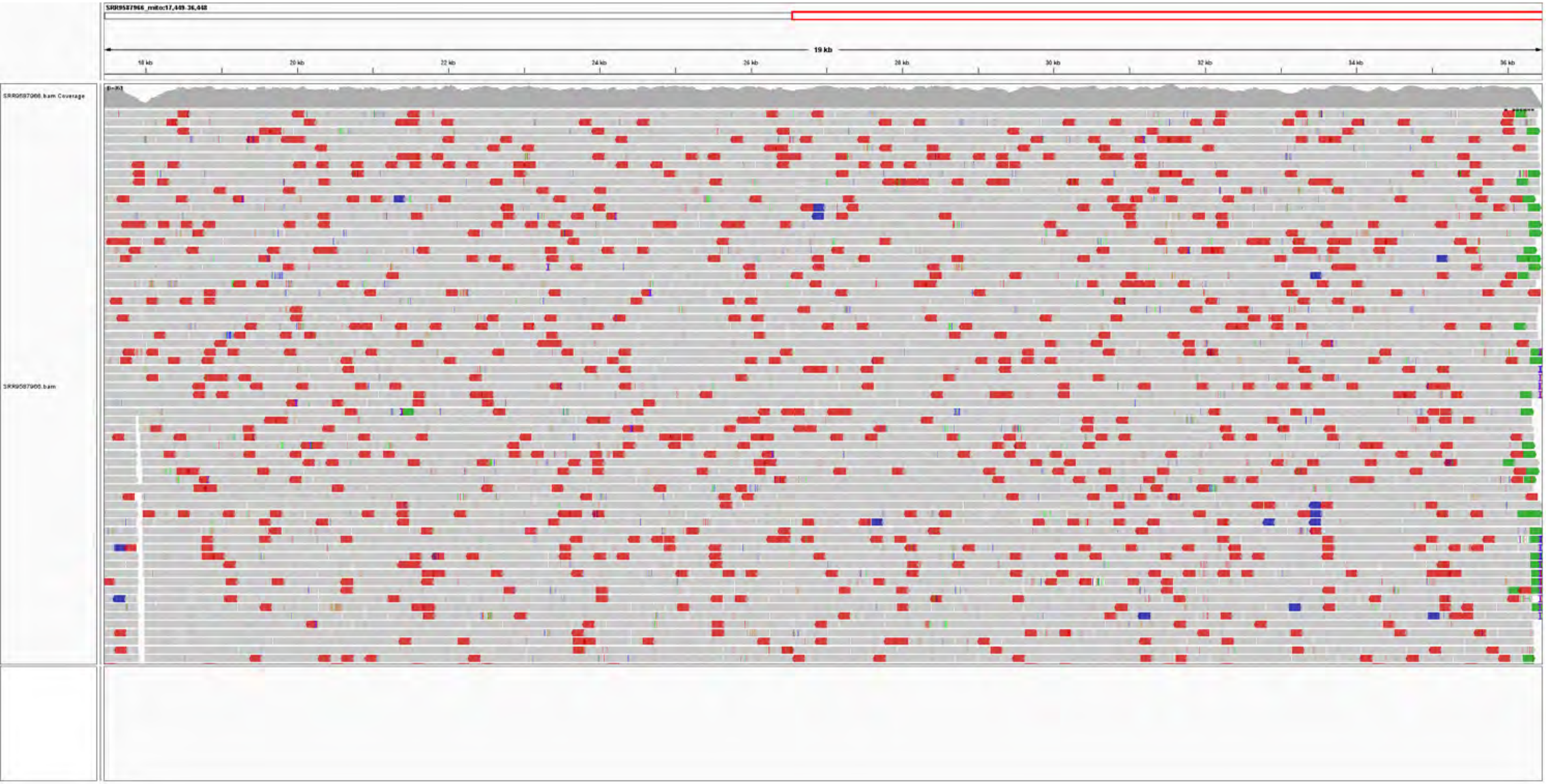

### SRR9587967

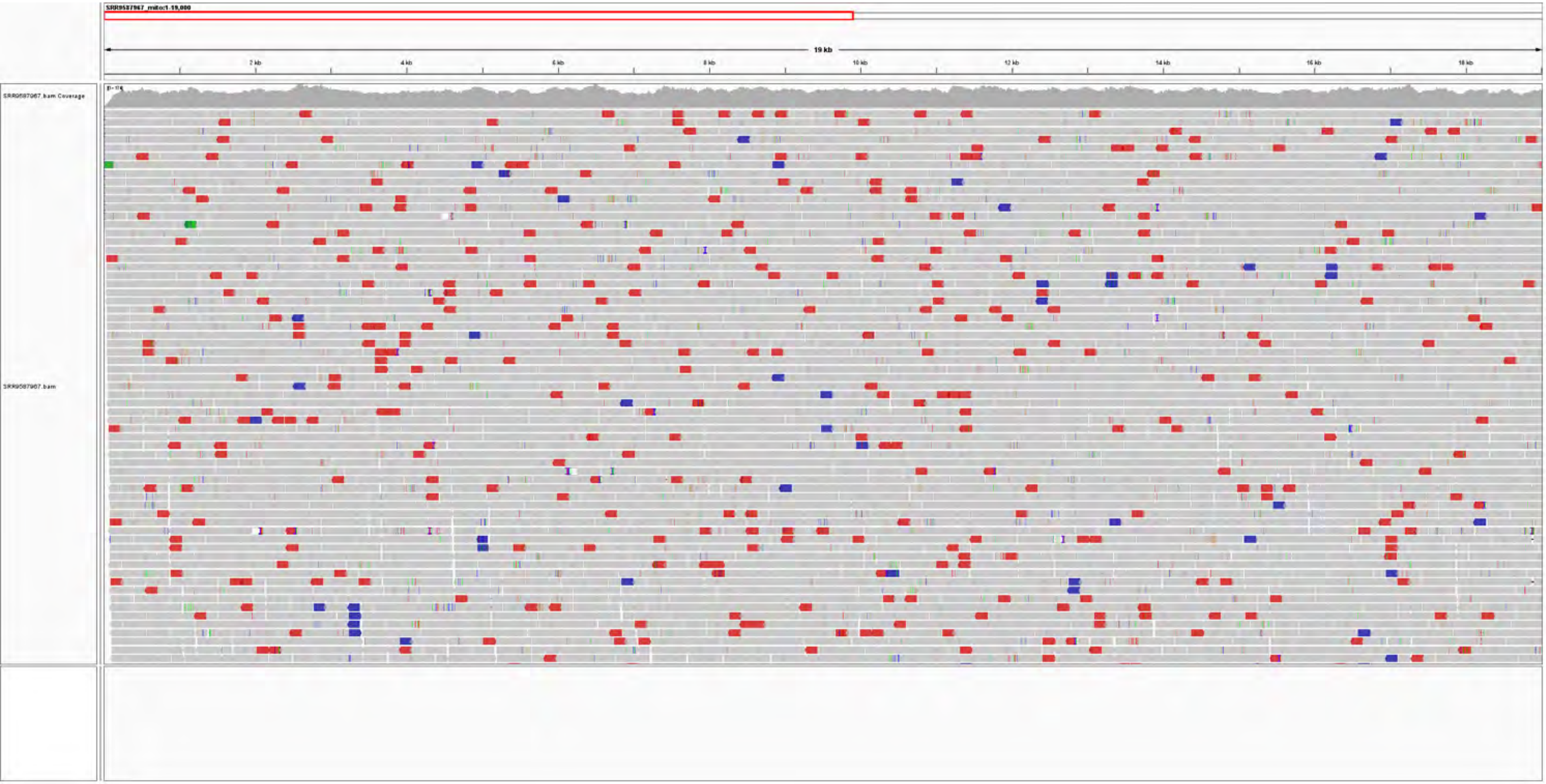

SRR9587967_mito:1-19,000
19 kb
2 kb
4 kb
6 kb
8 kb
10 kb
12 kb
14 kb
16 kb
18 kb
SRR9587967.bam Coverage
SRR9587967.bam

### SRR9587967 (continued)

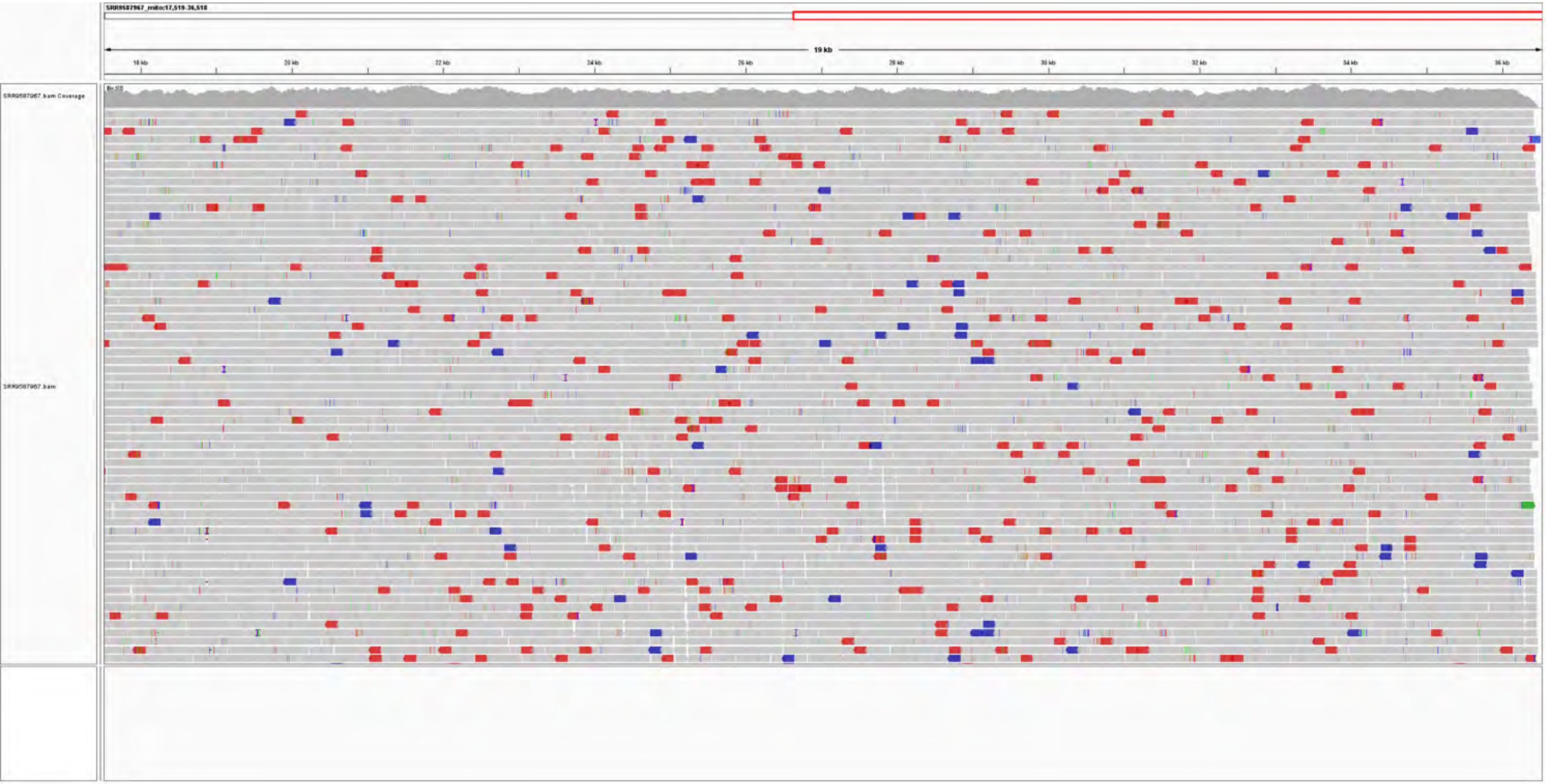
